# Formation of human long intergenic non-coding RNA genes and pseudogenes: ancestral sequences are key players

**DOI:** 10.1101/826784

**Authors:** Nicholas Delihas

## Abstract

Pathways leading to formation of non-coding RNA and protein genes are varied and complex. We report finding a highly conserved repeat sequence present in both human and chimpanzee genomes that appears to have originated from a common primate ancestor. This sequence is repeatedly copied in human chromosome 22 (chr22) low copy repeats (LCR22) or segmental duplications and forms twenty-one different genes, which include human long intergenic non-coding RNA (lincRNA) gene and pseudogene families, as well as the gamma-glutamyltransferase (*GGT*) protein gene family and the RNA pseudogenes that originate from *GGT* sequences. In sharp contrast, only predicted protein genes stem from the homologous repeat sequence present in chr22 of chimpanzee. The data point to an ancestral DNA sequence, highly conserved through evolution and duplicated in humans by chromosomal repeat sequences that serves as a functional genomic element in the development of new and diverse genes in humans and chimpanzee.

## Introduction

Models presented for the pathways in formation of genes are diverse (1). These include formation of long non-coding RNA (lncRNA) genes from protein genes (2-5), with one study based on similarities in open reading frames (5), and the reverse pathway of human protein gene formation from lncRNA genes that are found in rhesus macaque and chimpanzee (6). Here we report new findings on an evolutionarily conserved repeat sequence that is present in multiple and diverse RNA and protein genes and propose that a similar sequence serves as a proto-gene forming unit, a nucleation site for formation of new genes, both non-coding RNA (ncRNA) and protein genes. The repeat sequence is highly prevalent in different segmental duplications or low copy repeats (LCR22) of human chromosome 22 (chr22), specifically in region 22q11.2. Chr22 has the largest number of segmental duplications per unit chromosomal length of any human chromosome (7). These duplications are dynamic (8). Several may have arisen after the separation of human and macaque lineages (9) and they are continuously evolving, as shown by comparisons and differences found in current human populations (10). Segmental duplications have been considered to be important for new gene formation and human evolution (11-13). Additionally, the 22q11.2 region in itself is of special interest as it is prone to genetic deletions formed during fetal development that result in a high rate of genetic abnormalities (14). Segmental duplications have been shown to participate in the deletion process via meiotic nonallelic homologous recombination (9, 11).

In this paper we propose a model for the evolutionarily conserved human/chimpanzee repeat sequence and show that it serves as a starting point for formation of new lncRNA genes with subsequent base pair changes, sequence additions and/or deletions. The core sequence consists of the common sequence shared by the gamma-glutamyltransferase (*GGT*) protein gene family, where *GGT* is linked to three phylogenetically conserved and distinct sequences. In humans, these sequences form families of long intergenic non-coding RNA genes and pseudogenes that are linked to *GGT* sequences present in chromosomal segmental duplications; in chimpanzee, the conserved repeat sequence has not been found to form lncRNA genes but to form predicted protein genes. The presence of *GGT* in the long arm of chr22 was determined several decades ago (15) and its duplication in segmental duplications has also been reported (11). The *GGT* family is well characterized (16, 17).

In addition to the *GGT*-linked gene segments, we describe another protein gene family in LCR22s, the ubiquitin specific peptidase (*USP*) family that is found linked to lincRNA genes in humans. In chimpanzee, the *USP*-linked sequences encode only predicted protein genes, which is similar to the findings with *GGT*–associated sequences. The significance of chromosomal segmental duplications to gene development described here has parallels to the importance of human genome expansion of repeat units in the evolution of regulatory elements (18).

## Methods

### Reference genomes for primate species

Homo sapiens chromosome 22, GRCh38.p12 Primary Assembly NCBI was the source of sequences and properties of RNA and protein genes. Pan troglodytes, isolate Yerkes chimp pedigree #C0471 (Clint) chromosome 22, Clint_PTRv2, NCBI was the source of chimpanzee sequences and protein genes. In addition, a cloned sequence from Pan troglodytes, clone rp43-41g5, complete sequence GenBank: AC099533.36 provided an additional copy of the conserved repeat sequence. Gorilla gorilla (western gorilla) chromosome 22, gorGor4, NCBI Reference Sequence: NC_018446.2, locus: NC_018446 (19) was used to search for the presence of *GGT* genes and the conserved repeat sequence.

### Gene properties and gene searches

NCBI and Ensembl websites: (https://www.ncbi.nlm.nih.gov/gene) (19-21) and (http://useast.ensembl.org/Homo_sapiens/Info/Index) (22, 23) were used as the primary sources for gene properties. However *FAM230A* gene annotations provided a partial gene sequence as there is a 50 kbp unsequenced gap within the gene. For, sequence analysis both the *FAM230A* NMD RNA transcript sequence from NCBI and the *FAM230A* gene sequence provided by Ensembl was used. Additional databases employed for gene properties were: Gene Cards: GeneCards – the human gene database, (www.genecards.org) (24), HGNC: (Genenames.org) (25), RNAcentral: rnacentral.org/ (26). For chimpanzee gene searches, the NCBI Reference sequence (RefSeq) database was used (19). The NCBI annotation of chimpanzee protein genes are with the Gnomon-The NCBI eukaryotic gene prediction tool (https://www.ncbi.nlm.nih.gov/genome/annotation_euk/gnomon/).

### Genomic coordinates

The NCBI and Ensembl gene coordinates differ for a number of genes, especially at the 5’ ends. For uniformity, all coordinates used here were according to NCBI with the expectation of *AC023490.3* that has been annotated only by Ensembl. The description of linked gene segments in Table 1 is in the order of *FAM230*-clincRNA-*GGT* or pseudogenene-*GGT*. However, in the genome, the gene order for several of the repeat segments is in the reverse orientation. For consistency, all linked genes are shown in the same orientation as in Table 1.

**Table 1.** *GGT*-linked genes present in human LCR22*.

| Linked gene segments | LCR22 location | chr22 coordinates** |
| --- | --- | --- |
| *FAM230B--LOC105372935--GGT2* | LCR22D | chr22:21166903 -21283023 |
| *FAM230E--LOC105377182--GGT3P* | LCR22A | chr22:18733914-18791961 |
| *FAM230A--AC023490.3--GGTLC3* | LCR22A | chr22:18487127-18518165 |
| *FAM230J--LOC105372942--GGTLC5P* | LCR22A | chr22:18340163-18386526 |
| *POM121L1P--GGTLC2* | LCR22E | chr22: 22631557-22647898 |
| *POM121L10P--BCRP3--GGT1* | LCR22H | chr22:24583150-24650612 |
| *POM121L9P (BCRP1)--GGTLC4P--GGT5* | LCR22G | chr22:24219654-24265524 |
| *FAM230C--LOC101060145--GGT4P* | (chr13) | chr13:18195297-18271624 |

\*The *FAM230* family consists of ten genes. Genes *FAM230F*, *FAM230H*, and *FAM230I* are not linked to a conserved *lincRNA*-spacer-*GGT* sequence and appear to have been formed separately in LCR22s.
\*\*Coordinates are from the NCBI.

### Blast, BLAT searches and sequence identity determinations

The Blast search engine (https://blast.ncbi.nlm.nih.gov/Blast.cgi?CMD=Web&PAGE_TYPE=BlastHome (27) and Blat search engine (http://useast.ensembl.org/Homo_sapiens/Tools/Blast?db=core) (22) were both used to find similarities is gene sequences and to initially detect gene families.

### Nucleotide and amino acid sequence alignments and identity determinations

**T**he EMBL-EBI Clustal Omega Multiple Sequence Alignment program,website: http://www.ebi.ac.uk/Tools/msa/clustalo/ was used for alignment of two or more nucleotide or amino acid sequences. This program was also used to determine phylogenetic relationships via generation of a phylogram.

The identity between two sequences was determined by the NCBI Basic Local Alignment Search Tools, blastn and blastp, align two or more sequences with the Program Selection: Optimize for Highly similar sequences (megablast) (27). The identities represent only aligned sequences and do not including gaps sequences.

### RNA expression

The expression of RNA from normal tissues were obtained from website: www.ncbi.nlm.nih.gov/gene/, human tissue-specific expression (HPA) RNA-seq normal tissues (28). The expression of circular RNAs: Tissue-specific circular RNA induction during human fetal development was obtained from website: www.ncbi.nlm.nih.gov/gene/ (29) where RNA-seq was performed on 27 different human tissues with samples from 95 individuals.

### Protein properties

UniProtKB (uniprot.org/uniprot/) was the source of human protein amino acid sequences. For chimpanzee proteins, amino acid sequences and regions of the protein sequence that have predicted functional domains were from: www.ncbi.nlm.nih.gov/protein (20).

### RepeatMasker analysis of nt sequences

The RepeatMasker program, www.repeatmasker.org/cgi-bin/WEBRepeatMasker (30) was employed to search for transposable elements. The search engine, “hmmer” for DNA sequences and interspersed repeats was used to detect transposable elements.

## Results

### Background on *GGT*-linked gene repeat sequences

The DNA repeat sequence was detected in human chr22 segmental duplications LCR22A and LCR22D while analyzing the *FAM230* lincRNA genes (31). The repeat represents three gene families, whose sequences are linked (Figure 1a): the *FAM230* lincRNA gene family (highlighted in yellow), a newly found **<u>c</u>**onserved **<u>l</u>**ong **i**ntergenic **<u>n</u>**on-**<u>c</u>**oding RNA (*clincRNA*) gene family (highlighted in green) and the sequence of the *GGT* protein family as well as *GGT*-related pseudogenes (highlighted in red). An uncharacterized spacer sequence that resides between the *clincRNA* and *GGT* sequences (highlighted in gray) is also highly conserved in LCR22A and LCR22D. We refer to *GGT* as the sequence shared by *GGT1* and *GGT2* that comprises ∼20,000 bp. Figure 1b is a representation of the linked gene segment *FAM230B*--*LOC105372935*--*GGT2*, which we use as a guide and model for sequence comparisons. Listed are bp numbers that show the ends of genes present in the linked gene segment, which comprises a total of 116,120 bp. The drawings are representational and not to scale.

**Figure 1.**
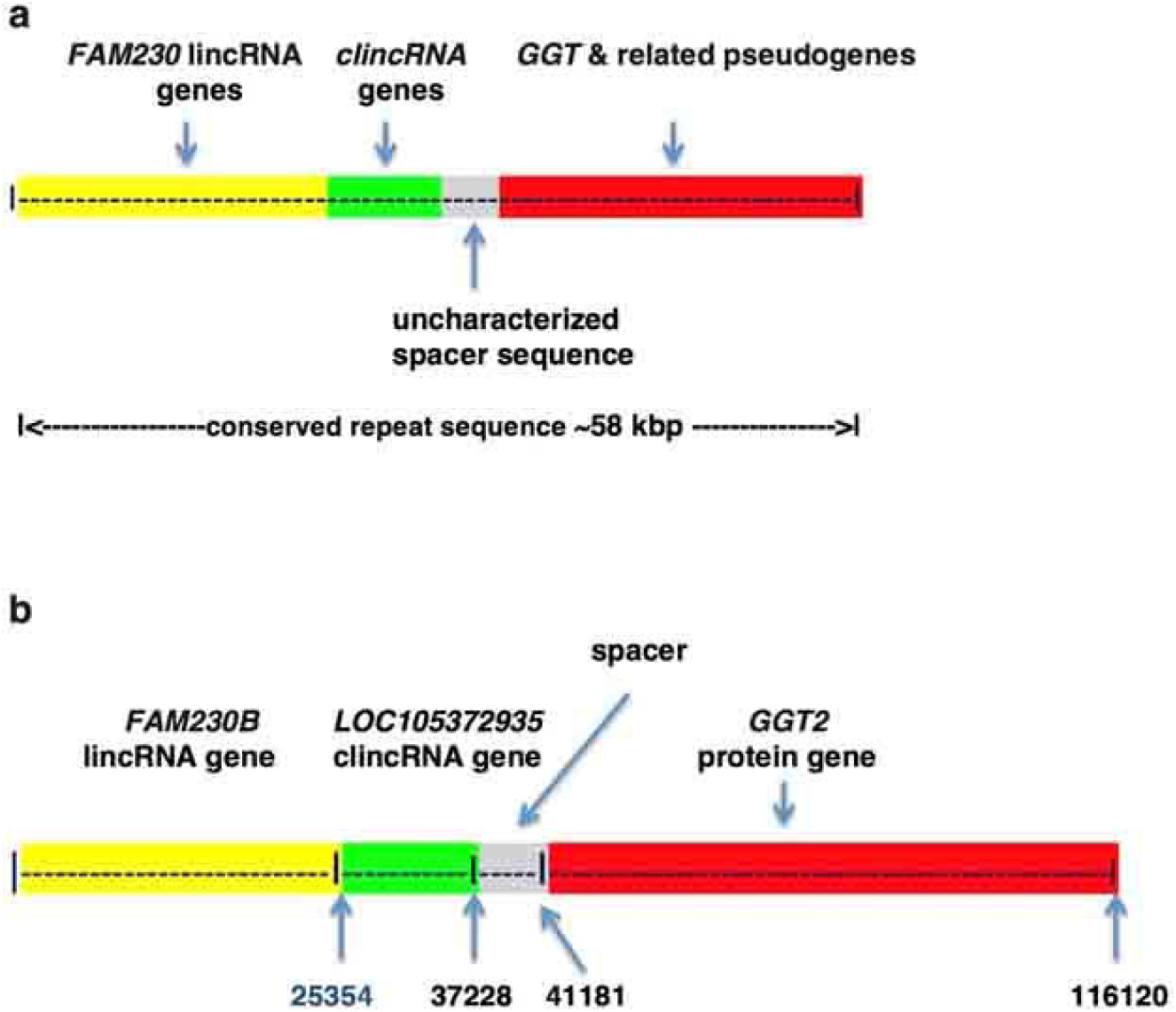
**a**. Diagrammatic representation of the conserved core sequence that comprises linked gene families found in the human chr22 LCR22A and D segmental duplications; these are: *FAM230* lincRNA gene family (highlighted in yellow), the *clincRNA* family genes (highlighted in green), a spacer sequence (highlighted in gray) and the *GGT* gene family (highlighted in red. **b.** Schematic of the linked gene segment *FAM230B*--*LOC105372935*--*GGT2.* Diagrams are approximate and not drawn to scale.

Table 1 lists the linked gene segments, which represent copies of the conserved sequence, and their location in LCR22s. The *clincRNA* genes are those starting with the prefix LOC or AC and are linked to *FAM230* genes in LCR22A and LCR22D. Also grouped together in Table 1 are linked genes in segmental duplications LCR22E and LCR22H; these carry the repeat sequence but do not have the *FAM230* sequence, and some also differ with respect to the uncharacterized spacer sequence, which may be partially or totally missing. In segmental duplications LCR22E and LCR22H, pseudogenes *POM121* transmembrane nucleoporin like 1 pseudogene *POM121L1P* and the BCR activator of RhoGEF family pseudogene *BCRP3* are found linked to *GGT*; these pseudogenes stem from the *clincRNA* sequence. Thus the homologous sequence that forms the *clincRNA* gene family in LCR22A and D is found to generate pseudogenes in chromosomal segmental duplications LCR22E, and H. The *FAM230C* gene and linked genes reside in chr13 and not in chr22 or an LCR22 (Table 1).

A diagrammatic representation of the eight LCR22s in human chr22 shows the location of the *GGT*-linked gene segments in LCR22s (Figure 2). The four linked-gene units that contain the *FAM230* gene family (Figure 1) are present only in LCR22A and LCR22D..

**Figure 2.**
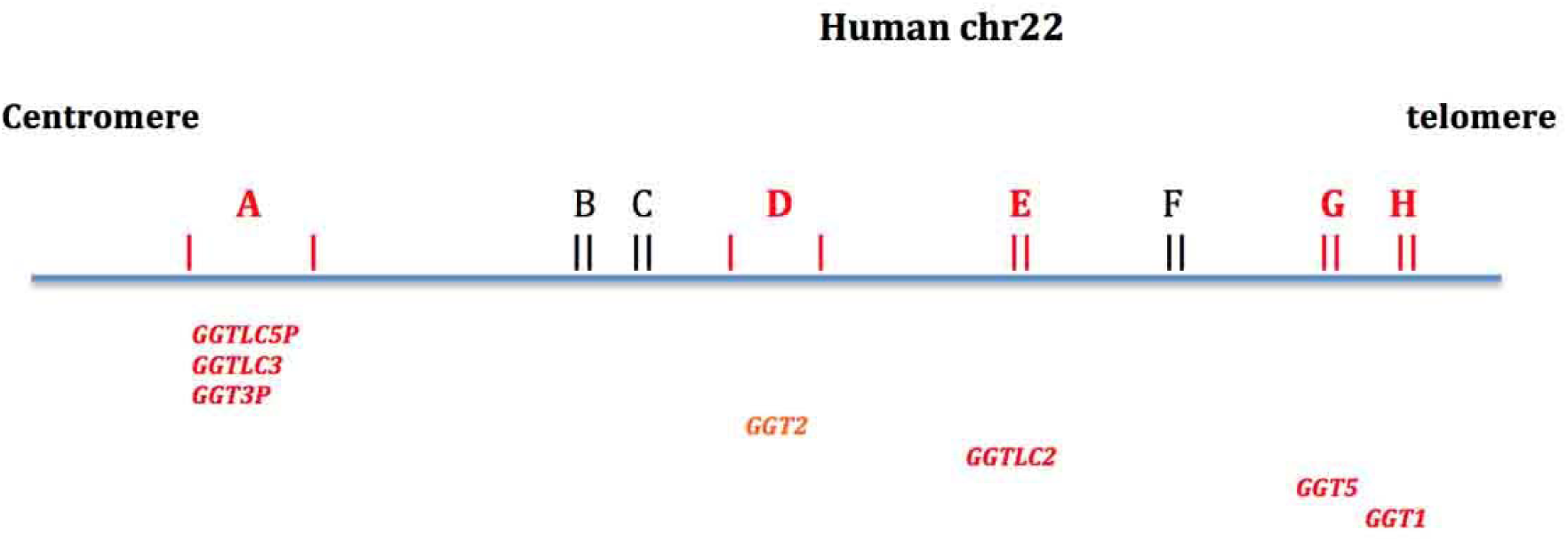
A schematic of segmental duplications found in the 22q.11.2 region of human chr22. A-H (in red) represent the eight LCR22s. The *GGT*-linked gene segments are represented by the *GGT*-related symbols in the drawing.

The human DNA repeat sequence is also found present in the chimpanzee genome with high identity. A nt sequence alignment of four human *GGT*-linked gene sequences together with two homologous sequences from chr22 of the chimpanzee genome reveals the high similarity between most of the human and chimpanzee sequences (Supplementary Figure S1). Figure 3 shows a small segment of the sequences, which is taken from the complete nt sequence alignment of six *GGT*-linked gene segments. It visually displays the near perfect similarity in shared sequences at the *FAM230B* gene/*LOC105372935* (clincRNA) gene junction site (yellow/green highlighted junction). The divergence between the six sequences can be seen in Supplementary Figure S1.

**Figure 3.**
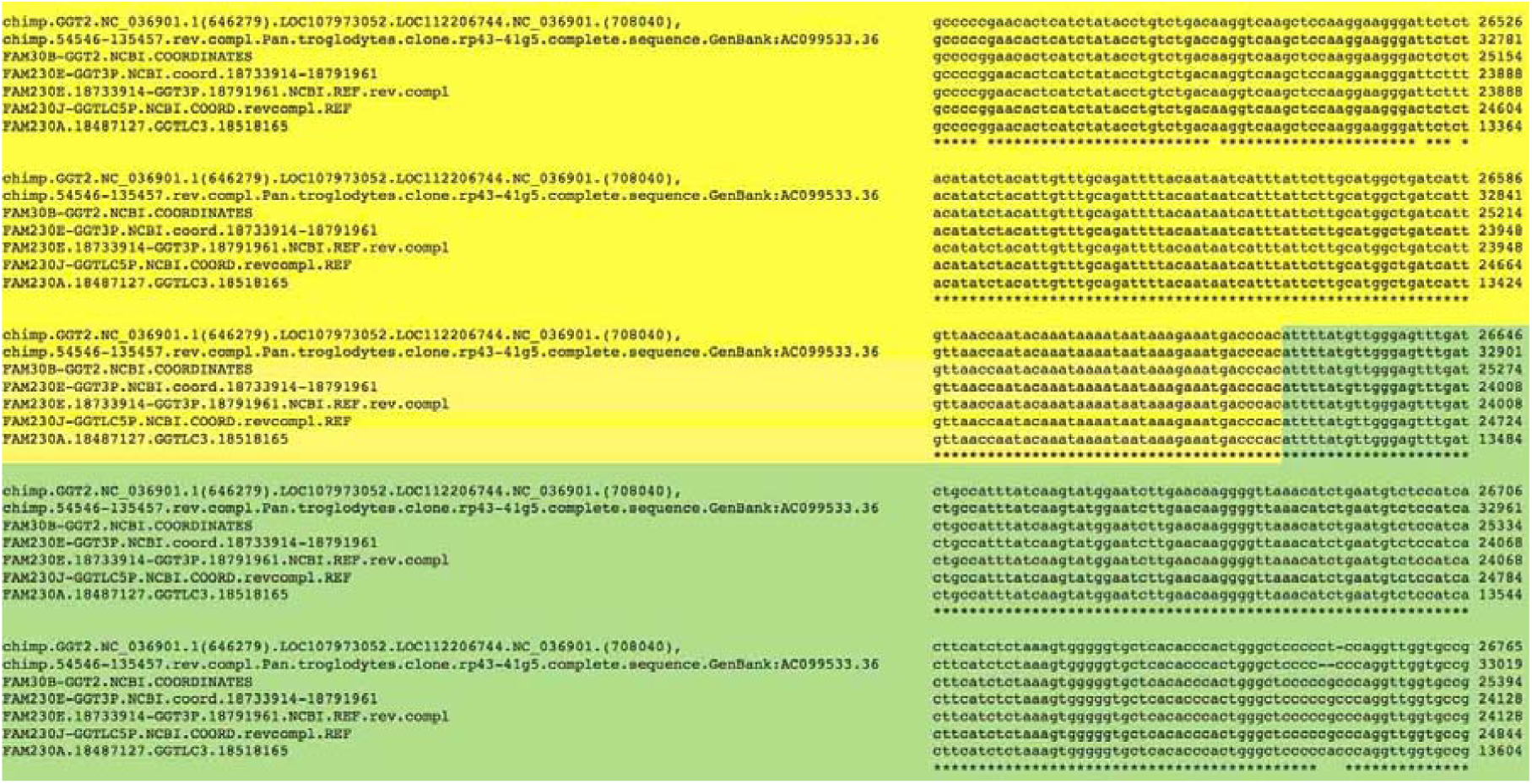
A segment of the alignment of four *FAM230-clincRNA-GGT linked genes* (Table 1) with two chimpanzee sequences. The complete sequence alignment is in Supplementary Figure S1. The yellow highlight denotes sequences of the *FAM230* genes, green highlight denotes the *clincRNA* genes with the *FAM230B--LOC105372935--GGT2* coordinates used for guideposts. The figure displays the 3’ end *FAM230B* gene/*LOC105372935* junction. The two chimpanzee sequences are from: Pan troglodytes isolate Yerkes chimp pedigree #C0471 (Clint) chromosome 22, Clint_PTRv2, NCBI Reference Sequence: NC_036901.1 and chimp.54546-135457.revcompl. from Pan.troglodytes.clone.rp43-41g5.GenBank:AC099533.36. The human sequences are from Homo sapiens chromosome 22, GRCh38.p12 Primary Assembly NCBI Reference Sequence: NC_000022.

As a model for the conserved repeat sequence, the sequence of *FAM230B*-*LOC105372935*-*GGT2* is used here for all comparisons as it contains 96% of the length of the linked gene-*GGT2* counterpart in chimpanzee and displays a very high nt sequence identity (97%-98%) with the chimpanzee sequence. In this manuscript the term *FAM230/clincRNA /GGT* is used to signify the conserved repeat sequence (Fig 1a) and to represent the putative ancestral conserved sequence. The chimpanzee genes are further discussed below.

### Analyses of *GGT*-linked genes in segmental duplications LCR22A and LCR22D

NCBI displays maps of *GGT* genes and surrounding genes (www.ncbi.nlm.nih.gov/gene). These maps are shown in Figure 4, left panel. In the right panel of Figure 4, schematic diagrams represent homologous sequences, with color identification that depict the *GGT* associated gene families found in the LCR22 duplications. Table 2 shows the percent nt sequence identity obtained from sequence alignments of the *GGT*-linked gene segments with the sequence of *FAM230B-LOC105372935*-*GGT2*. Figure 1b serves as a guide for the association of the percent identity relative to each gene family as it shows the positional ends of genes. Table 2 shows the conservation of sequence, which reveals a 98%-99% identity throughout most of the lengths of the four segments. Lower identities largely correspond to changes within *FAM230* lincRNA genes.

**Table 2.** Percent identity of *FAM230-*linked genes relative to *FAM230B-LOC105372935-GGT2*.

| *FAM230*-linked genes | % identity | nt positions from *FAM230B-LOC105372935-GGT2* |
| --- | --- | --- |
| *FAM230E-LOC105377182-GGT3P* | 98% | 393-16584 |
|  | 90% | 16436-18838 |
|  | 99% | 18864-59136 |
| *FAM230A-AC023490.3-GGTLC3* | 99% | 1507-11568 |
|  | 97% | 11517-16855 |
|  | 98% | 16647-40312 |
|  | 99% | 40295-42900 |
| *\*FAM230J-LOC105372942-GGTLC5P* | 98% | 1477-40312 |
|  | 99% | 40312-46976 |

\*Note; small segment of *FAM230J-LOC105372942-GGTLC5P* at nt 16600-17494 has identity of 77%; this reflects changes in the *FAM230J* sequence relative to *FAM230B*.

**Figure 4.**
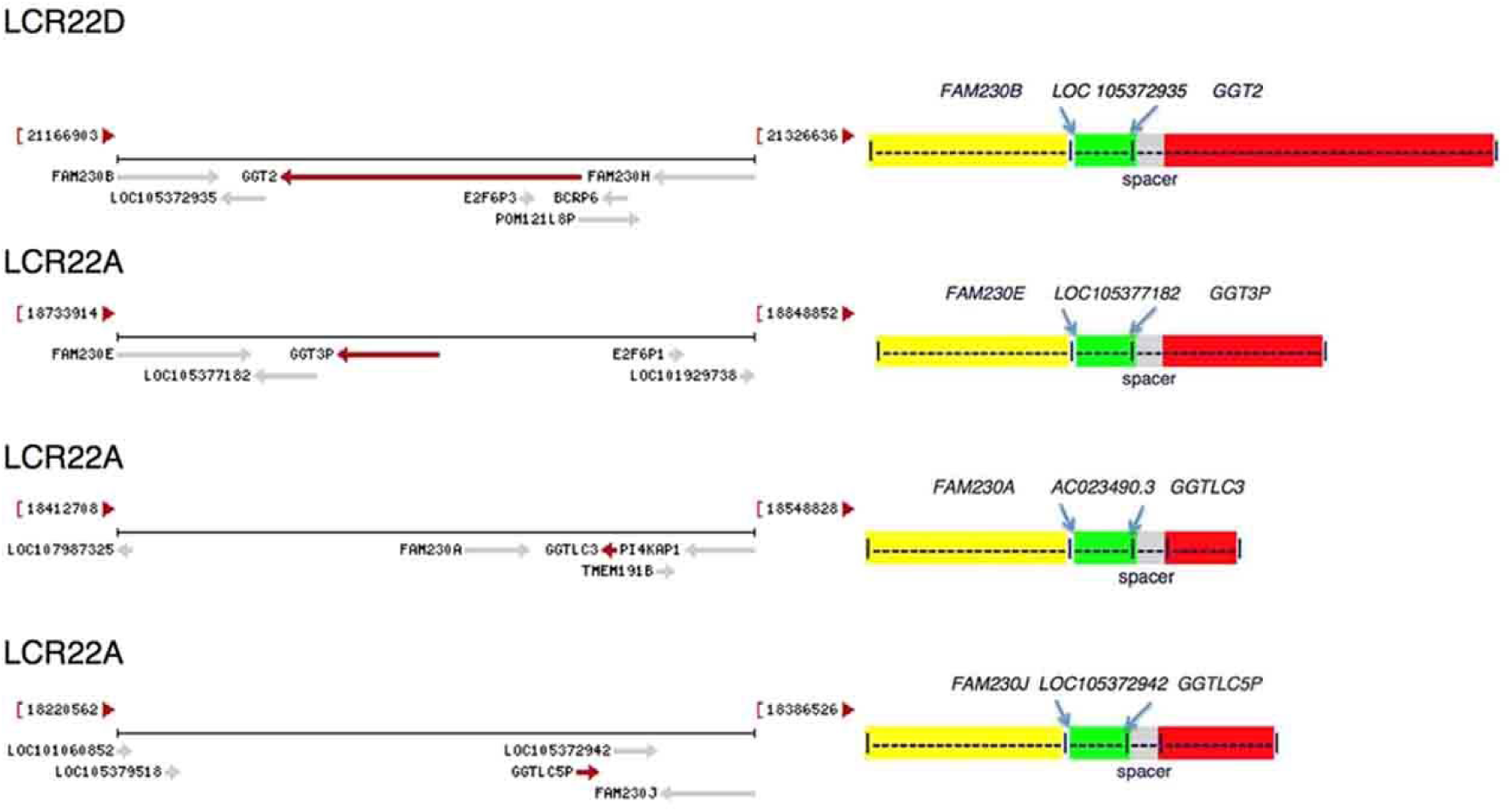
**Left panel**: *GGT* and the associated genes in LCR22s of chr22. The end chromosomal coordinates are shown in parentheses. The gene arrangement diagrams are directly from the NCBI website: https://www.ncbi.nlm.nih.gov/gene (21). **Right panel**: Schematic representation of *GGT*-linked genes (but not drawn to scale). *GGTLC5P* and its associated genes (bottom figure) are presented in the reverse orientation. Note: the *FAM230A* gene has a 50 kbp sequence gap, thus sequences from both Ensembl and NCBI were used for alignments to obtain more complete identity values. In addition, only Ensembl has annotated the clincRNA gene, *AC023490.3.*

A comparison of *FAM230E*-*LOC105377182*-*GGT3P* and *FAM230B--LOC105372935*--*GGT2* sequences indicates that the major sequence changes are between the lincRNA *FAM230B* and *FAM230E* genes. Figure 5 shows significant mutational changes in one region involving a large sequence deletion and several point mutations between the two *FAM230* sequences. This region is followed by over 8 kbp that show no major additions/deletions/point mutation. These differences may show the development of the *FAM230* genes into distinct structures and possibly different functions. For example, lincRNA transcripts from *FAM230B* and *FAM230E* differ in nt sequence, length and exon sequences (19, 21). Although the expression of RNA in normal somatic tissues from these genes is found only in testes (28, 31), the expression of circular RNAs (circRNA)s during fetal development shows differences between certain tissues (29) (Supplementary Figure S2). *FAM230E* circRNA is expressed in fetal heart tissue at 10 weeks and 17 weeks development, whereas *FAM230B* circRNA is not expressed in this tissue. This may be of significance in terms of possible *FAM230E* RNA gene function in the 22q11.2 chromosomal region during fetal development, as a 22q11.2 deletion results in abnormal heart development (14). There are genetic factors that may influence expression of circRNAs, resulting in differences in circRNA expression and the onset of various diseases (32).

**Figure 5.**
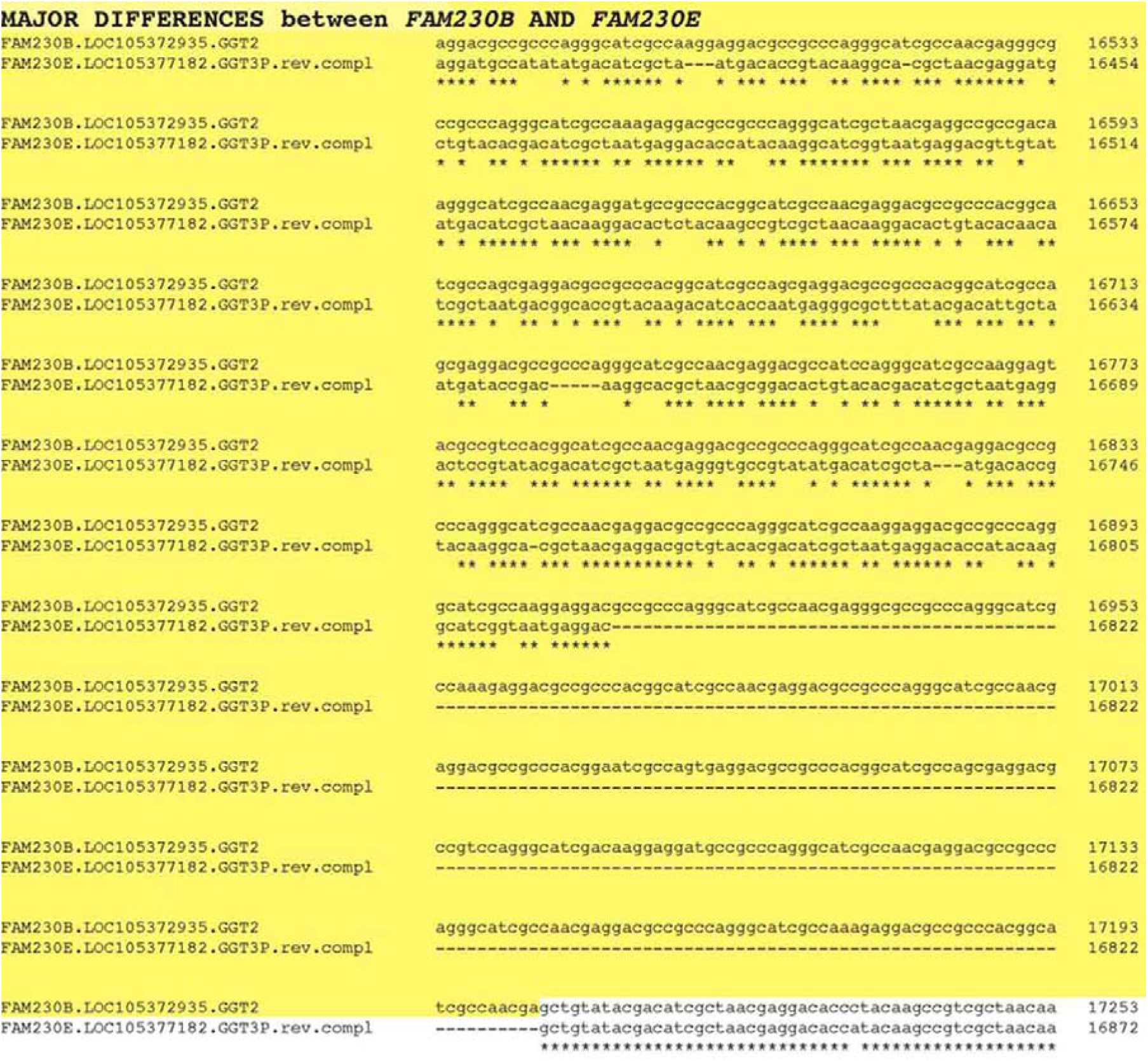
A section of the alignment of *FAM230B-LOC105372935*-*GGT2* and *FAM230E*-*LOC105377182*--*GGT* sequences with yellow highlighted sequences showing differences (point mutations, deletions/insertions) between *FAM230* genes.

The *clincRNA* genes *LOC105372935* and *LOC105377182* (Figure 4, right panel, top two drawings) are nearly identical, both in sequence and length. The expression of RNA from these two genes in somatic tissues, as well as the expression of circular RNA during fetal development, are also nearly identical (21, 29). Formation of the *clincRNA* genes may be recent, as they have not significantly diverged in sequence or in tissue-specific transcript expression.

*GGT3P* (Figure 4, right panel, second drawing from top) is an unprocessed pseudogene and comprises 18,273 bp. Its entire sequence is homologous to the 3’ end nt sequence of *GGT2* and *GGT1* protein genes. RNA transcript expression from *GGT2* and *GGT3P* in normal somatic tissues between the two genes is similar (21). The expression of circular RNAs during fetal development also shows similar patterns (29).

*GGT2* (Figure 4, right panel, top drawing) is a complex gene that encodes thirteen different transcripts. Most transcripts differ in size due to the presence of multiple exons in the *GGT2* 5’ UTR (see:www.ncbi.nlm.nih.gov/gene/728441) (20). The transcript that is used here as a model for gene size is the longest (NCBI GenBank ACCESSION NM_001351304 XM_016999937). *GGT2* produces a protein product [www.uniprot.org/uniprot/P36268], however the protein is inactive in glutathione hydrolase activity and its enzymatic activity has not been fully characterized (17).

Two other *GGT*-linked gene segments found in LCR22A, *FAM230A*-*AC023490.3*-*GGTLC3* and *FAM230J*-*LOC105372942*-*GGTLC5P* also show a high identity with *FAM230B*--*LOC105372935*--*GGT2* (Table 2), but here there are regions of major sequence changes within *FAM230* and differences in *GGT*-related genes in sequence lengths. In these segments, the *GGT* sequence forms the protein gene *GGTLC3*, the gamma-glutamyltransferase light chain family member 3, and the unprocessed pseudogene *GGTLC5P* is the gamma-glutamyltransferase light chain 5 pseudogene. The *GGTLC3* sequence consists only of the 3’ end sequences of *GGT1/GGT2* and displays an identity of 97% with *GGT1*, but closer identity with *GGT2*, 99%. An alignment of the three gene sequences reveals thirty-nine point mutations and three deletion/insertion mutations that are unique to *GGT1* relative to the other two sequences, and only one point mutation that is unique to *GGT2* and there are no deletions/insertions. This highly biased mutational pattern suggests that *GGTLC3* originated from a sequence similar to that of *GGT2*.

### *GGT*-linked genes in segmental duplications LCR22E, H and G

Linked gene sequences in segmental duplications LCR22E, H and G (Table 1) differ from those in LCR22A and D. They do not carry the *FAM230* sequence, and in one case, sections of the clincRNA sequence and the spacer region are missing. Figure 6 shows a schematic drawing depicting the differences relative the *FAM230B-LOC105373935-GGT2* model.

**Fig. 6.**
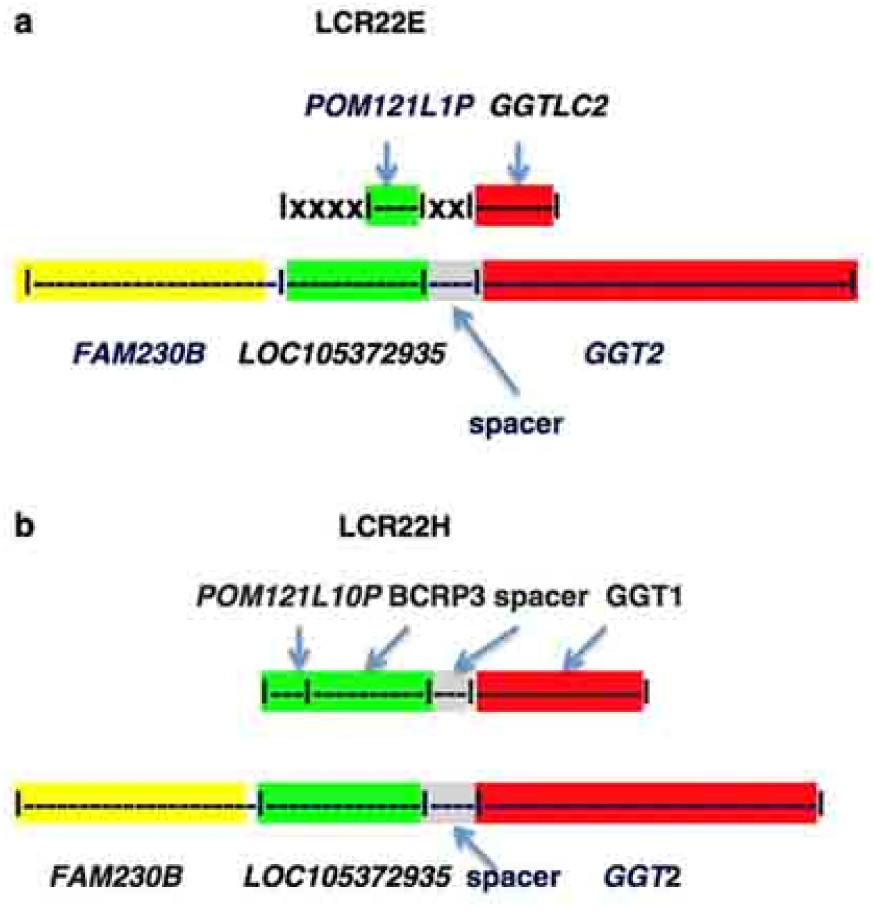
Schematic representation of *GGT*-linked genes in LCR22E, and H and comparisons with *FAM230B-LOC105372935*-*GGT2*. **a**. |xxx| represents the absence of parts of the *LOC105372935* clincRNA and the entire spacer sequences. The percent identity relative to nt positions of *FAM230B-LOC105372935-GGT2* are, *POM121L1P*, nt positions 28645-30907 96%; *GGTLC2*, positions 40838-43793, 96%; **b**. Percent identity of nt postions 28900-40312 of *FAM230B-LOC105372935-GGT2* with *POM121L10P--BCRP3*--spacer, 96%; with *GGT1*, nt positions, 40295 to 57246, 97%. The lengths of genes in the figure are not to scale.

#### POM121L1P-GTTLC2

The *POM121L1P-GTTLC2* linked gene segment is found in LCR22E. It resides in a complex chromosomal region, the immunoglobulin lambda gene locus *IGL*. There are six genes packed into a space of ∼3.2 kbp that also contains the *POM121L1P-GTTLC2* linked gene segment (19). There is evidence that the conserved repeat sequence was duplicated in this chromosomal region but it is significantly different; there is a partial *clincRNA* gene sequence present in *POM121L1P*-*GGTLC2* and *FAM230* and spacer sequences are missing (Figure 6a) (the symbol |XXX| refers to clincRNA and spacer sequences missing).

*POM121L1P* is termed a POM121 transmembrane nucleoporin like 1 unprocessed pseudogene (22). 2279 bp of the *POM121L1P* pseudogene sequence has an identity of 96% with aligned sequences of the *clincRNA* gene *LOC105372935* of *FAM230B*--*LOC105372935*--*GGT2*. Thus, part of the conserved sequence that forms *clincRNA* genes LCR22A and LCR22D (green highlight, Figure 6) forms part of this pseudogene in LCR22E.

*GGTLC2* encodes a gamma-glutamyltransferase light chain 2 protein and displays glutathione hydrolase activity [https://www.ebi.ac.uk/interpro/protein/Q14390]. It shares most of its sequence with *GGT2* and *GGT1* and displays an identity of 96% with *GGT2* but 98% with *GGT1*, however a mutational analysis to determine the closeness of *GGTLC2* with *GGT1* relative to *GGT2* is inconclusive.

#### POM121L10P-BCRP3-GGT1

*POM121L10P-BCRP3*-*GGT1* resides in LCR22H. Based on a sequence alignment with *FAM230B--LOC105372935--GGT2*, the *clincRNA* sequence and the entire uncharacterized spacer sequence are present and are linked to *GGT1* (Figure 6b). Sequences of the *POM121L10P and BCRP3* genes both stem from the *clincRNA* sequence and are highly similar to sections of the *clincRNA LOC105372935* sequence; 2264 bp of the *POM121L10P* sequence are homologous to *LOC105372935* (*clincRNA*) and an adjacent 6498 bp encoding *BCRP3* are also homologous to the clincRNA sequence with 96% identity. *BCRP3* is one of the eight *BCRP* family of pseudogenes that contain sequences from the breakpoint cluster region (*BCR*) gene. The *BCRP* pseudogenes are complex. Part of the *BCR* gene is in LCR22F. Of the eight *BCRP* family genes only one, *BCRP8* resides within the *BCR* gene sequence and thus stems from the *BCR* gene locus. Functions of *BCRP3* are needed to understand the relationship of this pseudogene to *BCR.*

GGT1 is a well-characterized enzyme. Over two decades ago it was pointed out that there are several human genes for *GGT* that produce different mRNAs but that *GGT1* produces an active gamma-glutamyltransferase enzyme (33). This was confirmed by Heisterkamp et al. (17).

#### POM121L9P(BCRP1)-GGTLC4P-GGT5

The *POM121L9P(BCRP1)*-*GTLC4P-GGT5* segment resides in LCR22G (Table 1). The *BCRP1* gene is situated entirely within the *POM121L9P* sequence and is an antisense gene. However, *GGT5* is an anomaly. Although gene positions relative to each other in chr22 are *POM121L9P(BCRP1)*-*GTLC4P-GGT5*, there is no evidence that *GGT5* originates from a *GGT* locus, however data point to the origin from a clincRNA sequence.

An alignment of the *POM121L9P (BCRP1)*-*GGTLC4P-GGT5* sequence with that of *FAM230B*-*LOC105372935*-*GGT2* shows that *POM121L9P (BCRP1)*-*GGTLC4P-GGT5* contains spacer and *GGT* sequences, and most of the *clincRNA* sequence. Of significance, a sequence alignment of the *GGT5* sequence with that of the *clincRNA LOC105372935* shows that *GGT5* contains part of this *clincRNA* sequence but carries no *GGT* sequences (Table 3 and Supplementary Figure S3). *GGT5* is 25489 bp in length and carries 5803 bp.of the *LOC10537293* clincRNA sequence. Thus ∼21% of *GGT5* contains clincRNA sequences and the identity is 90-92% (Table 3). It has been pointed out before that there is little nt sequence homology between the *GGT5* and *GGT1* genes (17).

**Table 3.** Percent Identities of *GGT5* and *GGTLC4P* with *GGT2* and *clincRNA* gene.

| Gene | length | # bp with identity<br>with *GGT2* | % identity | # bp with identity<br>with *clincRNA*<br>*LOC105372935* | % identity |
| --- | --- | --- | --- | --- | --- |
| *GGT5* | 25489 | 0 | 0 | 5803 | 90-92 |
| *GGTLC4P* | 1553 | 1553 | 96% | 0 | 0 |

Table 3 shows the close similarity of pseudogene *GGTLC4P* with *GGT2* sequences, where the entire sequence of *GGTLC4P* consists of *GGT2* sequences. *GGTLC4P* also displays a high identity with *GGT1* (not shown).

Although there is no significant nt sequence homology between *GGT5* and *GGT2*, amino acid sequences of the protein products have similarities where approximately half of the amino acid residues are identical (17). In addition, the GGT5 protein displays gamma-glutamyltransferase activity (17).

The chimpanzee *GGT5* nt sequence is also present in the chimpanzee genome and it is found to be highly similar to the human sequence (with 98% identity over 90% of the human *GGT5* sequence). In addition, both human and chimpanzee genes contain the clincRNA signature sequences. A phylogram tree analysis was performed with *GGT*-related sequences. It shows that human *GGT5*, chimpanzee *GGT5* and human *clincRNA LOC105372935* DNA sequences form a phylogenetic branch that is separate from the branch grouping of *GGT1, GGT2*, and *GGTLC* (Figure 7). This is consistent with the data of Table 3. Because of the close similarities, *GGT5* appears to have originated in a primate ancestor.

**Figure 7.**
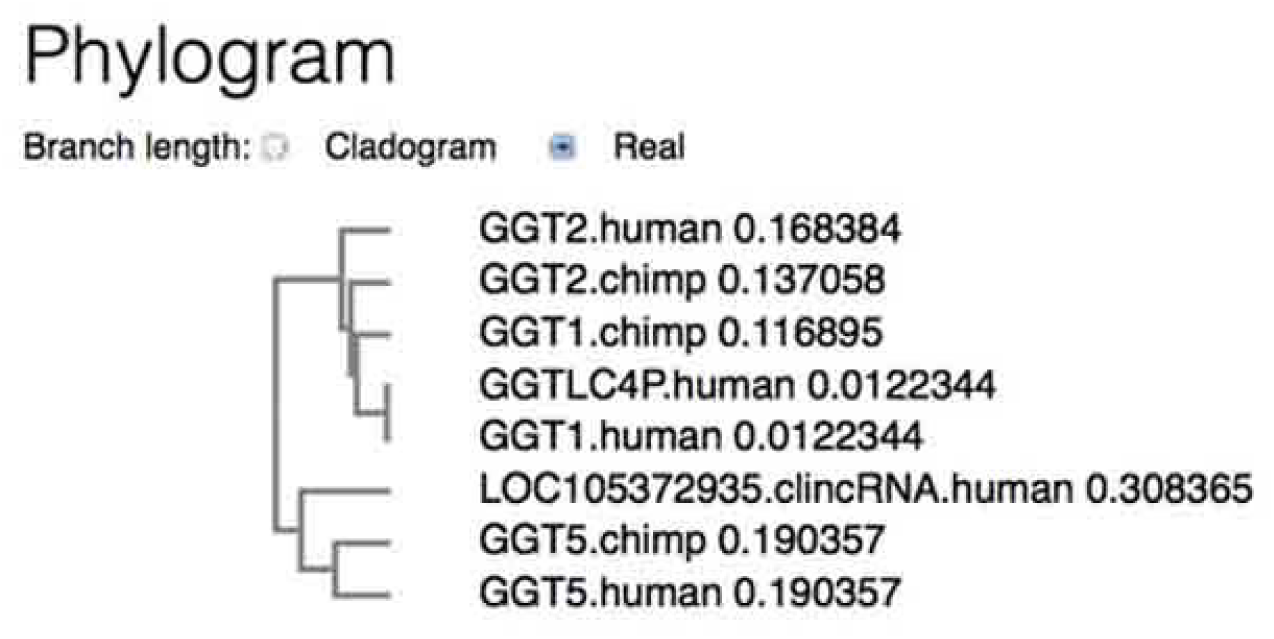
Phylogram shows the phylogenetic branch relationships between *GGT* genes and the *LOC105372935 lincRNA* gene. The data were obtained using the EBI Clustal Omega sequence alignment and phylogeny program.

#### FAM230C-LOC101060145-GGT4P

The *FAM230C-LOC101060145-GGT4P* linked gene locus is in chr13, which distinguishes it from the other *FAM230* family genes that are all in chr 22. What stands out in the *FAM230C-LOC101060145-GGT4P* linked gene segment is that no genes stem from the clincRNA sequence and that two pseudogenes, *LOC101060145* and *GGT4P* originate from the *GGT* sequence (Supplementary Figure S4). *LOC101060145* is annotated as a glutathione hydrolase light chain 1-like pseudogene by NCBI and *GGT4P* is a gamma-glutamyltransferase 4 pseudogene annotated by Ensembl. Supplementary Figure S4 also shows that the *FAM230C*-*LOC101060145*-*GGT4P* linked gene sequence has a high identity with the *FAM230B-LOC105372935-GGT2* sequence. This shows the presence of the *FAM230*-*clincRNA*-spacer-*GGT* repeat sequence outside of chr22.

### The human *FAM230B-LOC105372935-GGT2* sequence is present in the chimpanzee genome

#### Chimpanzee *LOC112206744*-*LOC107973052-GGT2*

Chimpanzee sequences in chr22 that are analogous to human *GGT*-linked sequences show high similarities. An analysis of the chimpanzee sequence adjacent to *GGT2* shows that 94% of the chimpanzee *LOC112206744*-*LOC107973052-GGT2* sequence has an identity of 97%-98% compared with the human *FAM230B-LOC105372935-GGT2* (Table 4). A sequence alignment between the human and chimpanzee sequences is shown in Supplementary Figure S5.

**Table 4.** Percent identity of chimpanzee *GGT*-linked sequences relative to human *FAM230B-LOC105372935-GGT2*.

| Chimpanzee GGT-linked genes | %identity | bp positions from<br>*FAM230B-LOC105372935-GGT2* |
| --- | --- | --- |
| *LOC112206744-LOC107973052-GGT2\** | 98% | 2423-16644 |
|  | 98% | 17177-23523 |
|  | 97% | 23509-40312 |
| ; | 98% | 41007-60800 |
| *LOC112206779-LOC107973089-GGT1* | 94% | 635-1495 |
|  | 98% | 2423-17049 |
|  | 97% | 17936-40312 |
|  | 97% | 40295-61048 |

\*61762 bp is total length of chimpanzee *LOC112206744-LOC107973052-GGT2*

In sharp contrast to the human lincRNA genes *FAM230B* and *LOC105372935*, annotations reveal that there are predicted protein genes, *LOC112206744* and *LOC107973052* encoded in the homologous chimpanzee sequence (Figure 8a). The annotated protein genes were derived by NCBI with the automated computational analysis program, “Gnomon, the gene prediction method” (20). Thus, homologous sequences that form lincRNA genes in humans form predicted protein genes in the chimpanzee. The spacer sequence is also present with a 97% identity to the human spacer.

**Figure 8.**
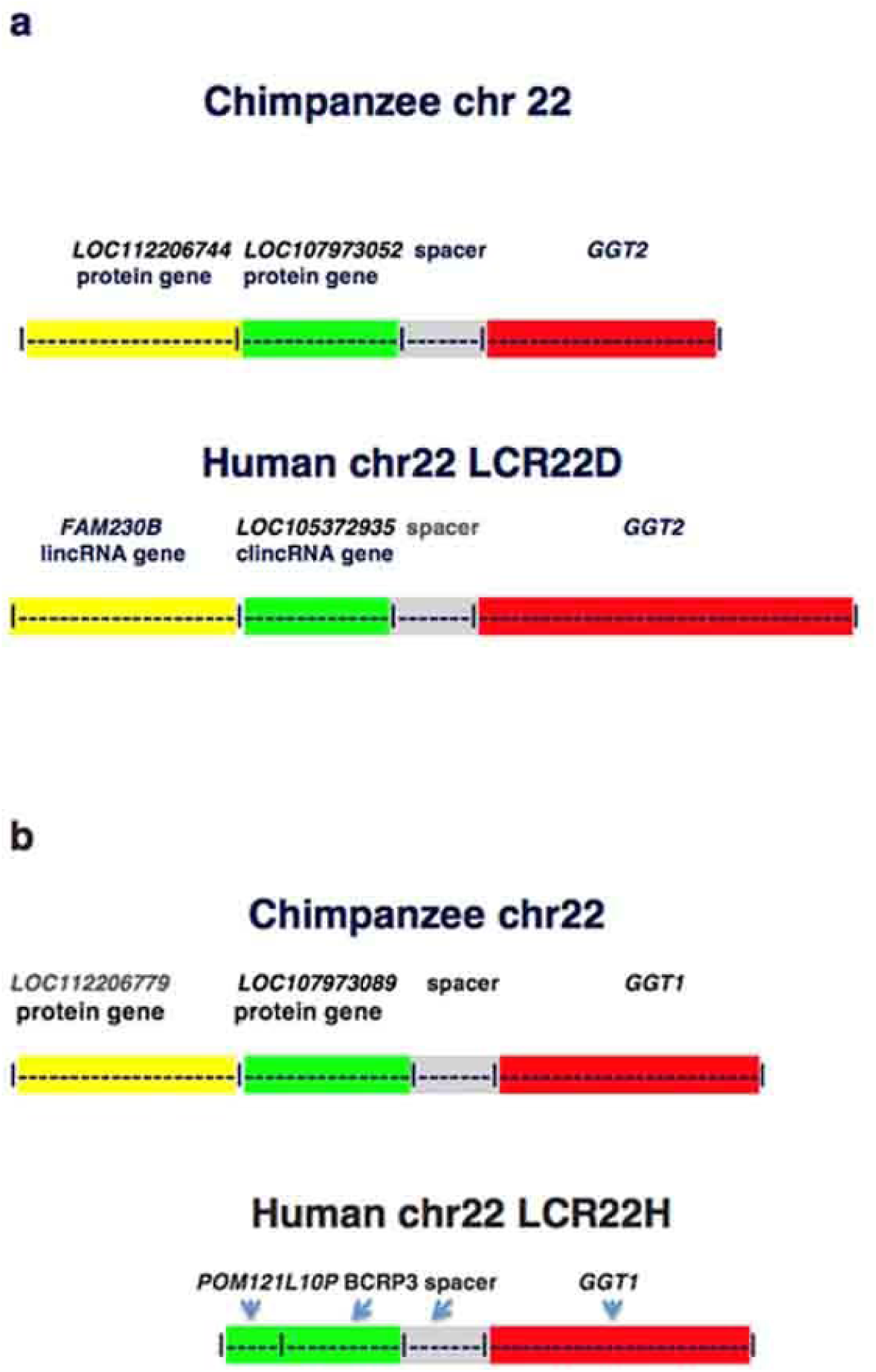
Schematic of nt sequences of *FAM230B-LOC105372935-GGT2* that are present in the chimpanzee genomic sequence: source: Pan troglodytes isolate Yerkes chimp pedigree #C0471 (Clint) chr22. **a.** the chimpanzee *LOC112206744* -*LOC107973052-GGT2* gene sequence identities with nt positions of human *FAM230B--LOC105372935* --*GGT2* are: 2423-16644, 98%identity *^*with *LOC112206744*; 23509-40312, 97%, *LOC107973052* and spacer; 41007-60800, 98%, *GGT2*. The identity between the two sequences starts with position 1 of the chimpanzee *GGT2*-linked gene sequence and position 185 of the human *GGT2*-linked gene sequence and ends at chimpanzee position 61762 and human position 60800. **b.** The chimpanzee *LOC112206779--LOC107973089*--*GGT1* gene sequence identities with nt positions of *FAM230B--LOC105372935* --*GGT2* are: 635 to 1495, 94%, *LOC112206779*; 2423 to 17049 98% *LOC112206779*; 17936-40312, 97%, *LOC107973089*-spacer; 40295-61048, 97%, *GGT1*. Human *POM121L10P--BCRP3--GGT1*, which lacks the *FAM230* sequence is shown for comparison.

#### Chimpanzee LOC112206779-LOC107973089-GGT1

Chimpanzee *GGT1* and its adjacent sequence also have a high identity with the *FAM230B--LOC105372935--GGT2* sequence (Table 4). But unlike the human *GGT2* linked gene locus, the human *GGT1* linked gene segment lacks the *FAM230* sequence (Fig 8b, bottom schematic). The chimpanzee sequence has (and possibly retained from an ancestral sequence) the *FAM230* sequence; it shows a 94%-98% identity with the *FAM230B* gene sequence.

Similar to the chimpanzee *GGT2-*linked gene segment, the sequence that is homologous to human *FAM230B* in chimpanzee encodes a predicted protein gene, *LOC112206779*. The *LOC112206779-LOC107973089*-*GGT1* locus also has the clincRNA-spacer sequence signature, and the homologous clincRNA sequence encodes another predicted protein gene, *LOC107973089* (Fig 8b). Thus taken together with the *GGT2*-linked genes, there are four protein genes annotated in homologous sequences of the chimpanzee genome that encode lincRNA genes in humans.

### Human *FAM230D-USP* linked gene sequences are present in the chimpanzee genome

#### *FAM230D-USP18* linked gene sequences are present in the chimpanzee genome

In addition to the *GGT*-linked genes, there is another example of the human *FAM230*-*lincRNA* sequence that forms protein genes in chimpanzee, sequences that are linked to the *USP* protein genes; *USP* is an ubiquitin specific peptidase.

In the human *USP18* linked sequences, the *FAM230D* sequence and part of the clincRNA sequence are linked to *USP18*, however the uncharacterized spacer is missing. This unit is found in segmental duplication LCR22A (Supplementary Figure S6). Thus only a section of the conserved repeat sequence represented in Fig. 1, the *FAM230*-clincRNA section is present in the *FAM230D*-*USP18* locus.

A sequence annotated as *LOC104003670*-*LOC745914*-*USP18* is present in chimpanzee. Both *LOC104003670* and *LOC745914* are predicted protein genes and are homologous to sections of the lincRNA *FAM230D* gene (with 97% identity); thus both these genes appear to stem from the *FAM230* sequence (Supplementary Figure S6). The clincRNA sequence does not form a part of the chimpanzee protein genes. The chimpanzee *LOC104003670* amino acid sequence is 727 aa in length and its peptide carries a predicted conserved protein domain, the RNA recognition motif superfamily (RRM) with an e-value of 6.45-03 (20). This domain is at aa positions 192-341 of *LOC104003670*. The other chimpanzee gene, *LOC745914* translates to a 287 aa peptide in length. No putative domains are found in this protein sequence.

The human *FAM230* family genes do display open reading frames. Chimpanzee *LOC104003670* has an aa sequence similarity to an 158 aa open reading frame of *FAM230D* with 50% identity and an 85% aa sequence identity with a translated open reading frame of 440 aa from the human nonsense mediated decay transcript from *FAM230A* (Supplementary Figure S7). As all *FAM230* genes share a high degree of similar nt sequences, there are also similarities in open reading frames. The *FAM230* sequences may serve as a foundation for protein gene development in addition to lincRNA gene formation.

#### *FAM230G-USP41* linked gene sequences are present in the chimpanzee genome

Another example of an *FAM230-USP* sequence, *FAM230G-USP41* that forms protein genes in chimpanzee in place of lincRNA genes are the genes linked to *USP41*: *LOC112206746-LOC112206769*-*USP41* (Supplementary Figure S8). USP41 is termed ubiquitin specific peptidase 41. The *FAM230G-USP41* segment is in LCR22B. There are major changes in both the human and chimpanzee *USP41*-linked sequences compared to the *FAM230*-*clincRNA*-spacer segment of the conserved repeat depicted in Figure. 1. In humans, *FAM230G* is linked directly to the *USP41* sequence without clincRNA or spacer sequences and the *FAM230G* gene lacks the 3’ half sequence of *FAM230B.* The chimpanzee sequence that is homologous to *FAM230G*-*USP41, LOC112206746-LOC112206769*-*USP41* also does not carry the *clincRNA* or uncharacterized spacer sequences, but there are two annotated protein genes, *LOC112206746* and *LOC112206769* that have a nt sequence identity (97% and 84%, respectively) with the *FAM230B* sequence. Of note, *LOC112206769* is specified by a section of the *FAM230B* sequence that is not present in *FAM230G*, the 3’ end segment. This is the second example of the *FAM230B* sequence, or part of it, that is missing in the human analog (the other being *POM121L10P*-*BCRP3*-*GGT1*) but is present in the chimpanzee and forms predicted protein genes.

The two chimpanzee proteins, *LOC112206746* and *LOC112206769* show a predicted super domain. *LOC112206746* is termed uncharacterized protein DKFZp434B061-like and is 417 aa in length (20). A protein Blast Search gives a non-specific hit for a ribonuclease E super domain with an e-value of 5.08e-04. *LOC112206769* is also termed a DKFZp434B061-like protein and also gives a hit for a ribonuclease E super domain with an e-value, 1.21e-04. These two proteins, as well as chimpanzee *LOC104003670* (linked to *USP18*), all carry predicted super domains in their amino acid sequences. In humans, there is also an uncharacterized protein termed DKFZp434B061 (UniProtKB/Swiss-Prot: Q9UF83.2) and there is experimental evidence for the protein at the transcript level (21). Ensembl has annotated the human gene as *AL356585.9*. A characterization of this putative protein and determination of its relationship to the chimpanzee DKFZp434B061-like proteins is of interest.

## Discussion

The proposed ancestral proto-gene forming element is based on the findings that *GGT* and the three distinct linked sequences are conserved between humans and chimpanzee to the extent of primate protein genes and that these sequences evolutionarily withstood mutational drift and shift. In humans, the sequence has been repeatedly duplicated in the genome by chromosomal expansion through segmental duplications where the *GGT*-linked sequences is found to primarily form lincRNA gene and pseudogene families. In chimpanzee, these sequences form predicted protein genes; thus the informational content of the DNA element is such that it can lead to development of either lncRNA or protein genes. The *GGT*-related and the *USP*-sequences described here may not be isolated cases as there are other genes in LCR22s, unrelated to *GGT* sequences that appear to have also originated by the same process of sequence duplication where an homologous duplicated sequence can result in the formation of different genes (unpublished data).

The concept of segmental duplications as vehicles for the proliferation of *GGT*- and *USP*-related repeat sequences with the development of new genes parallels the findings of the effects of human chromosomal expansion, which consists primarily of repeat sequences, on the evolution and development of gene regulation (18). In addition, there is a parallel of the proposed ancestral proto-gene forming element described here with that of enhancer regulatory elements that have developed from ancestral sequences or proto-enhancers (34). These studies point to the role of ancestral sequences in the evolution of regulatory elements, and in the current work, that of gene development.

The study here adds another aspect to the work of others that suggests a number of lncRNA genes originated from protein genes (2-5). For example, Talyan *et al*. (5) showed that RNA and protein genes share partial open reading frames and that a number of RNA genes may have originated from protein genes. In the work presented here, some of the *GGT*-related RNA pseudogenes stem from protein *GGT* sequences, however the *BCRP* and *POM121* family pseudogenes <u>and</u> the lincRNA genes originate from the evolutionarily conserved *clincRNA* sequence and *FAM230* RNA sequences, respectively, and not from existing protein genes. On another scale, other lncRNAs have been shown to come from enhancer sequences (35). Thus various studies show that lncRNAs have different origins.

With respect to protein gene formation, open reading frames in *FAM230* and *clincRNA* sequences may provide the foundation for protein gene development. We show that there are translated open reading frames in lincRNA *FAM230* sequences that have similarities to those of predicted chimpanzee protein genes. It is not surprising that the aa sequence of protein gene *LOC104003670*, which is linked to *USP18* in chimpanzee has a high identity to the translated open reading frame of several human *FAM230* lincRNA sequences (e.g., see Supplementary Figure S4). However lincRNA sequences lack protein coding capacity, as was previously pointed out (36).

In the chimpanzee, a total of eight predicted protein genes arise from the nt sequences homologous to the human *FAM230* sequences with several protein structures showing proposed functional super domains, however no lncRNA genes have been assigned to these sequences in the chimpanzee. This suggests preferential formation of lncRNA genes in humans from homologous sequences that form only protein genes in the chimpanzee. Experimental evidence for a protein product would further support this finding.

There are sixteen lincRNA genes and pseudogenes that arise from the *FAM230*-*clincRNA* sequence in humans, and one protein gene, *GGT5. GGT5* belongs to the *GGT* family of protein genes. It is a well-characterized gene whose protein product displays gamma-glutamyltransferase activity but the gene nt sequence displays no significant DNA sequence homology with other members of the *GGT* family, as shown by Heisterkamp et al (17) and the work presented here. Thus *GGT5* is an anomaly in that its DNA sequence does not stem from a *GGT* locus. Although its gene position in chr22 is: *POM121L9P(BCRP1)*--*GTLC4P--GGT5*, there is no evidence that *GGT5* contains *GGT* DNA sequences but data point to the origin from the *clincRNA* locus. This is consistent with the predicted protein genes in chimpanzee and shows that protein genes can be formed from the clincRNA sequence. As the chimpanzee *GGT5* gene is found to be homologous to the human *GGT5* and its sequence also contains clincRNA nt sequences, *GGT5* and the process of its formation from *clincRNA* sequences appear to have originated in a common primate ancestor.

Why is *GGT5* formed from an unrelated DNA sequence and not from the *GGT* sequence itself? The primate cell may have performed its own “genetic and molecular engineering” to form a protein similar in sequence and function to the *GGT* family proteins but from a different genomic sequence. This does not address why the *GGT* nt sequence is not used to form the *GGT5* gene as it is for other *GGT*-related genes. Genes that are descended from an ancestral gene, share nucleotide sequences, have similar translated protein aa sequences and share similar functions are generally classified as a gene family. The *GGT5* gene offers an interesting variation to this definition.

The *clincRNA* gene family found in LCR22 A and D may have formed recently, as there is little or no difference in sequence or in RNA transcript expression in different tissues. Whether some of these genes will develop important functions or eventually disappear is not known. The *FAM230* genes show more development with differences in DNA nt sequence, specific RNA transcript structures and a differential expression of circRNAs in fetal tissues (21, 29).

The uncharacterized spacer sequence is not found to form a part of genes. This sequence is highly conserved in homologous sequences of the linked gene loci found in LCR22A and D, and it is also present in chimpanzee counterparts and conserved to 97%. But it has totally dissipated in linked genes *POM121L1P*-*GGTLC2*, found in LCR22E and in *FAM230D--USP18* in LCR22A. Its evolutionary conservation points to a function, but perhaps a non-essential one as it has disappeared in some segmental duplications.

The formation of the *FAM230* family, which consists of a total of eight lincRNA genes in chr22, is only partially understood. This gene family is formed from the 3’ half sequence of *FAM230C* with a remnant of the 5’ half *FAM230C* sequence found in the upstream region of seven of the genes (31, 37). *FAM230C* resides in chr13. Six *FAM230* genes are part of the conserved repeat units described here, either with *GGT* or *USP* genes and may have have formed while part of these repeats. The remaining three *FAM230* genes have not been found to be part of a repeat element and may have formed by a separate process. The origin of *FAM230C*, which found in chr13 is also uncertain.

The gorilla genome (19) has also been searched for the presence of the conserved repeat sequence by alignment of the gorilla genomic sequence with the *FAM230B*--*LOC105372935*--*GGT2* sequence and with gorilla sequences neighboring *GGT* genes. *GGT1, GGT2 and GGT5*, clincRNA and the uncharacterized spacer sequences were detected in the gorilla chr22 genomic sequence. However, the current gorilla sequence has a large number of unsequenced gaps, which currently precludes a detailed analysis.

After this manuscript was completed, Stewart and Rogers [38] published work with Drosophila showing the importance of chromosomal rearrangements in the formation of new protein genes.

## Availability of data on websites

Gene searches, gene properties, and gene transcript expression data: www.ncbi.nlm.nih.gov/gene/ (http://useast.ensembl.org/Homo_sapiens/Info/Index

Additional databases for gene properties:

GeneCards – the human gene database: (www.genecards.org)

HGNC: (Genenames.org)

RNAcentral: (rnacentral.org/)

Blast and BLAT searches and sequence identity determinations: (https://blast.ncbi.nlm.nih.gov/Blast.cgi?CMD=Web&PAGE_TYPE=BlastHome (http://useast.ensembl.org/Homo_sapiens/Tools/Blast?db=core)

Nucleotide and amino acid sequence alignments:

The EMBL-EBI Clustal Omega Multiple Sequence Alignment program: (http://www.ebi.ac.uk/Tools/msa/clustalo/)

Protein properties:

UniProtKB (uniprot.org/uniprot/)

Predicted functional domains (www.ncbi.nlm.nih.gov/protein)

RepeatMasker analysis of nt sequences:

RepeatMasker program (www.repeatmasker.org/cgi-bin/WEBRepeatMasker)

## Supplementary Data

Supplementary Figure S1. Nt sequence alignment of two chimpanzee and four human sequences that contain the repeat core sequence.

Supplementary Figure S2. Circular RNA expression during human fetal development.

Supplementary Figure S3. Alignment of human *GGT5* with *LOC105372935* clincRNA gene sequence.

Supplementary Figure S4. a. Color highlighted sections represent the *FAM230B-LOC105372935* -*GGT2* sequences that are found in *FAM230C*-*LOC101060145-GGT4P* with the respective percent identities. **b.** Schematic of *FAM230B-LOC105372935-spacer*-*GGT2* for comparisons.

Supplementary Figure S5. Nt alignment of chimpanzee *LOC112206744-LOC107973052-GGT2* with human *FAM230B-LOC105372935GGT2.*

Supplementary Figure S6. Identity of chimpanzee *LOC104003670*-*LOC745914*-*USP18 with FAM230D-USP18*.

Supplementary Figure S7. Alignment of aa sequences from chimpanzee LOC104003670 amino acid sequence (query) with the aa sequence of the open reading frame of FAM230A transcript from UniProteinKB-A0A1W2PPH8 (A0A1W2PPH8_HUMAN)(subject).

Supplementary Figure S8. The *FAM230G-USP41* sequence is analogous to chimpanzee *LOC112206746--LOC112206769--USP41*.

