## Supplementary Figure S1 for "Formation of human long intergenic non-coding RNA genes and pseudogenes: ancestral sequences are key players"

### A conserved primate sequence forms long non-coding RNA genes, pseudogenes and GGT-related protein genes

**Nicholas Delihas**

Supplementary Figure S1. Nt sequence alignment of two chimpanzee and four human sequences that contain the repeat core sequence. The chimp.LOC112206744-LOC107973052-GGT2.revcompl sequence was from Pan troglodytes isolate Yerkes chimp pedigree #C0471 (Clint) chromosome 22, Clint_PTRv2. The source of the other chimpanzee sequence is from a clone and is as shown below. The human sequences were from Homo sapiens chromosome 22, GRCh38.p12 Primary Assembly NCBI Reference Sequence: NC_000022 with the human *FAM230A* sequence from the NCBI NMD transcript. The Clustal Omega, Multiple sequence alignment program was used for sequence alignment.

```
CLUSTAL O(1.2.4) multiple sequence alignment


chimp.LOC112206744-LOC107973052-GGT2.revcompl                            ------------------------------------------------------------ 0
chimp54546-135457.revcompl.Pan.troglodytes.clone.rp43-41g5.GenBank:AC099533.36      AGCAGTTGGACCTCACAGTGTGGATTGTGCCTTCACCCTGGAATGTTTATGCCCTATCGC 60
human.FAM30B-LOC105372935-GGT2.NCBI.                                     agcagttggacctcacagtgtggattgtgccttcaccctggaatgtttatgccctatcac 60
human.FAM230E-LOC105377182-GGT3P.NCBI.                                   ------------------------------------------------------------ 0
human.FAM230J-LOC105372942-GGTLC5P.NCBI.rev.compl                        ------------------------------------------------------------ 0
human.FAM230A.ncRNA.18487127.18500594-GGTLC3.18516335.18518165.rev.compl.REF      ------------------------------------------------------------ 0

chimp.LOC112206744-LOC107973052-GGT2.revcompl                            ------------------------------------------------------------ 0
chimp54546-135457.revcompl.Pan.troglodytes.clone.rp43-41g5.GenBank:AC099533.36      CATGGTGATGGGATTAGGGATCTCCTGCCCTTGGTCCTAAGTGCCACTGTCTGTGCTGAG 120
human.FAM30B-LOC105372935-GGT2.NCBI.                                     catggtgatgggattagggatctcctgcccttggtcctaagtgccactatctgtgctgag 120
human.FAM230E-LOC105377182-GGT3P.NCBI.                                   ------------------------------------------------------------ 0
human.FAM230J-LOC105372942-GGTLC5P.NCBI.rev.compl                        ------------------------------------------------------------ 0
human.FAM230A.ncRNA.18487127.18500594-GGTLC3.18516335.18518165.rev.compl.REF      ------------------------------------------------------------ 0

chimp.LOC112206744-LOC107973052-GGT2.revcompl                            ------------------------------------------------------------ 0
chimp54546-135457.revcompl.Pan.troglodytes.clone.rp43-41g5.GenBank:AC099533.36      TTTTTCAAAGGTCAGAGCAGATTGAACCTCTGTGGTTTCATTTTCCCTGATTTTGATTTT 180
human.FAM30B-LOC105372935-GGT2.NCBI.                                     tttttcaaaggtcagagcagattgaaccattgtggtttcattttccctgattttgatttt 180
human.FAM230E-LOC105377182-GGT3P.NCBI.                                   ------------------------------------------------------------ 0
human.FAM230J-LOC105372942-GGTLC5P.NCBI.rev.compl                        ------------------------------------------------------------ 0
human.FAM230A.ncRNA.18487127.18500594-GGTLC3.18516335.18518165.rev.compl.REF      ------------------------------------------------------------ 0

chimp.LOC112206744-LOC107973052-GGT2.revcompl                            ----atggggaacctgtgtggctgcattcaaggtatgttcatactggcctgtcaaatgcg 56
chimp54546-135457.revcompl.Pan.troglodytes.clone.rp43-41g5.GenBank:AC099533.36      TCTTATGGGGAACCTGTGTGGCTGCATTCAAGGTATGTTCATACTGGCCTGTCAAATGCG 240
human.FAM30B-LOC105372935-GGT2.NCBI.                                     tcttatggggaacctgtgtggctgcattcaaggtatgttcatactggcctgtcaaatgcg 240
human.FAM230E-LOC105377182-GGT3P.NCBI.                                   ------------------------------------------------------------ 0
human.FAM230J-LOC105372942-GGTLC5P.NCBI.rev.compl                        ------------------------------------------------------------ 0
human.FAM230A.ncRNA.18487127.18500594-GGTLC3.18516335.18518165.rev.compl.REF      ------------------------------------------------------------ 0

chimp.LOC112206744-LOC107973052-GGT2.revcompl                            atcttttcaaattactagttaatgctttcaaaatatgttatttaaaaaattatcctctgt 116
chimp54546-135457.revcompl.Pan.troglodytes.clone.rp43-41g5.GenBank:AC099533.36      ATCTTTTCAAATTACTAGTTAATGCTTTCAAAATATGTTATTTAAAAAATTATCCTCTGT 300
human.FAM30B-LOC105372935-GGT2.NCBI.                                     atcttttcaaattactagttaatgctttcaaaatatgttatttaaaaaattagcctctgt 300
human.FAM230E-LOC105377182-GGT3P.NCBI.                                   ------------------------------------------------------------ 0
human.FAM230J-LOC105372942-GGTLC5P.NCBI.rev.compl                        ------------------------------------------------------------ 0
human.FAM230A.ncRNA.18487127.18500594-GGTLC3.18516335.18518165.rev.compl.REF      ------------------------------------------------------------ 0

chimp.LOC112206744-LOC107973052-GGT2.revcompl                            attttccatatggagttataaatatgtttcatggttatgttttattcctcaatttatata 176
chimp54546-135457.revcompl.Pan.troglodytes.clone.rp43-41g5.GenBank:AC099533.36      ATTTTCCATATGCAGTTATAAATATGTTTCATGGTTATGTTTTATTCCTCAATTTATAGA 360
human.FAM30B-LOC105372935-GGT2.NCBI.                                     attttccatatgcagttataaatatgtttcatgattatgttttattcctcaatttatata 360
human.FAM230E-LOC105377182-GGT3P.NCBI.                                   ------------------------------------------------------------ 0
human.FAM230J-LOC105372942-GGTLC5P.NCBI.rev.compl                        ------------------------------------------------------------ 0
human.FAM230A.ncRNA.18487127.18500594-GGTLC3.18516335.18518165.rev.compl.REF      ------------------------------------------------------------ 0

chimp.LOC112206744-LOC107973052-GGT2.revcompl                            tttgattattgtaccaagcagagtacctttgaaatttttcttcatttaaaaaatatgtct 236
chimp54546-135457.revcompl.Pan.troglodytes.clone.rp43-41g5.GenBank:AC099533.36      TTTGATTATTGTACCAAGTAGAGTACCTTTGAAATTTTTCTTCATTTAAAAAATATGTAT 420
human.FAM30B-LOC105372935-GGT2.NCBI.                                     tttgattattgtaccaagcagagtatctttgaaatttttcttcatttaaaaaatatgtat 420
human.FAM230E-LOC105377182-GGT3P.NCBI.                                   ------------------------------------------------------------ 0
human.FAM230J-LOC105372942-GGTLC5P.NCBI.rev.compl                        ------------------------------------------------------------ 0
human.FAM230A.ncRNA.18487127.18500594-GGTLC3.18516335.18518165.rev.compl.REF      ------------------------------------------------------------ 0

chimp.LOC112206744-LOC107973052-GGT2.revcompl                            cttggctcaggcctgtaatcccagcactttgggaggccaatgcaagaggatcacaaggtg 296
chimp54546-135457.revcompl.Pan.troglodytes.clone.rp43-41g5.GenBank:AC099533.36      CTTGGCTCAGGCCTGTAATCCCAGCACTTTGGGAGGCCAAGGCAAGAGGATCACAAGGTG 480
human.FAM30B-LOC105372935-GGT2.NCBI.                                     cttgactcaggcctgtaatcccagcactttgggaggccaaggcaagaggatcacaaggtg 480
human.FAM230E-LOC105377182-GGT3P.NCBI.                                   ------------------------------------------------------------ 0
human.FAM230J-LOC105372942-GGTLC5P.NCBI.rev.compl                        ------------------------------------------------------------ 0
human.FAM230A.ncRNA.18487127.18500594-GGTLC3.18516335.18518165.rev.compl.REF      ------------------------------------------------------------ 0

chimp.LOC112206744-LOC107973052-GGT2.revcompl                            aggagatcaagaccatcctggtcaatacagtgaaaccctgtctgtacgaaaagtacaaaa 356
chimp54546-135457.revcompl.Pan.troglodytes.clone.rp43-41g5.GenBank:AC099533.36      AGGAGATCAAGACCATCCTGCCCAATACAGTAAAACCCTGTCTCTATCAAAAATACAAAA 540
human.FAM30B-LOC105372935-GGT2.NCBI.                                     aggagatcaagaccatcctggccaatacagtgaaaccctgtctctactacaaatacaaaa 540
human.FAM230E-LOC105377182-GGT3P.NCBI.                                   ------------------------------------------------------------ 0
human.FAM230J-LOC105372942-GGTLC5P.NCBI.rev.compl                        ------------------------------------------------------------ 0
human.FAM230A.ncRNA.18487127.18500594-GGTLC3.18516335.18518165.rev.compl.REF      ------------------------------------------------------------ 0

chimp.LOC112206744-LOC107973052-GGT2.revcompl                            aattagccaggaatggtggcagctggtgtagtcccagtgtgaattgg------------- 403
chimp54546-135457.revcompl.Pan.troglodytes.clone.rp43-41g5.GenBank:AC099533.36      AATTAGCCTGGCATGGTGGCAGCTGCTGTAGTCACAGTGTGGTGTAGTCCCAGCTACCTG 600
human.FAM30B-LOC105372935-GGT2.NCBI.                                     atttagccaggcatggtggcagctggtgtagtcccagtgtgaattgg------------- 587
human.FAM230E-LOC105377182-GGT3P.NCBI.                                   ------------------------------------------------------------ 0
human.FAM230J-LOC105372942-GGTLC5P.NCBI.rev.compl                        ------------------------------------------------------------ 0
human.FAM230A.ncRNA.18487127.18500594-GGTLC3.18516335.18518165.rev.compl.REF      ------------------------------------------------------------ 0

chimp.LOC112206744-LOC107973052-GGT2.revcompl                            ------------------------------------------------------------ 403
chimp54546-135457.revcompl.Pan.troglodytes.clone.rp43-41g5.GenBank:AC099533.36      GGAGGCTGAGGCAGGACAATCGCTTGAACCCGTGAGGCAGAGGTTGCAGTGAGCCAAGAT 660
human.FAM30B-LOC105372935-GGT2.NCBI.                                     ------------------------------------------------------------ 587
human.FAM230E-LOC105377182-GGT3P.NCBI.                                   ------------------------------------------------------------ 0
human.FAM230J-LOC105372942-GGTLC5P.NCBI.rev.compl                        ------------------------------------------------------------ 0
human.FAM230A.ncRNA.18487127.18500594-GGTLC3.18516335.18518165.rev.compl.REF      ------------------------------------------------------------ 0

chimp.LOC112206744-LOC107973052-GGT2.revcompl                            -------------------------------gattcagtttattcccaaattcccaaatt 432
chimp54546-135457.revcompl.Pan.troglodytes.clone.rp43-41g5.GenBank:AC099533.36      GGCGCCATTGCACTCCAGCCTGTGCAACAGAACAAGACTCTGTTAAAAAAAAAAAAACAT 720
human.FAM30B-LOC105372935-GGT2.NCBI.                                     ------------------------------------------------------------ 587
human.FAM230E-LOC105377182-GGT3P.NCBI.                                   ------------------------------------------------------------ 0
human.FAM230J-LOC105372942-GGTLC5P.NCBI.rev.compl                        ------------------------------------------------------------ 0
human.FAM230A.ncRNA.18487127.18500594-GGTLC3.18516335.18518165.rev.compl.REF      ------------------------------------------------------------ 0

chimp.LOC112206744-LOC107973052-GGT2.revcompl                            atatatattatatatataatatattatatatacagtatatataatatattatatatacag 492
chimp54546-135457.revcompl.Pan.troglodytes.clone.rp43-41g5.GenBank:AC099533.36      ATATATATAATGTATATTATATATATTATATAT------TATAGAATATATGATATACAA 774
human.FAM30B-LOC105372935-GGT2.NCBI.                                     ------------------------------------------------------------ 587
human.FAM230E-LOC105377182-GGT3P.NCBI.                                   ------------------------------------------------------------ 0
human.FAM230J-LOC105372942-GGTLC5P.NCBI.rev.compl                        ------------------------------------------------------------ 0
human.FAM230A.ncRNA.18487127.18500594-GGTLC3.18516335.18518165.rev.compl.REF      ------------------------------------------------------------ 0

chimp.LOC112206744-LOC107973052-GGT2.revcompl                            tatataatatactatgtatacagtatatattatatactgtattatatac-attatatact 551
chimp54546-135457.revcompl.Pan.troglodytes.clone.rp43-41g5.GenBank:AC099533.36      TATGTAATATATAATATAT--GATACATAATATATAATATATGATATACAATTATATATC 832
human.FAM30B-LOC105372935-GGT2.NCBI.                                     ------------------------------------------------------------ 587
human.FAM230E-LOC105377182-GGT3P.NCBI.                                   ------------------------------------------------------------ 0
human.FAM230J-LOC105372942-GGTLC5P.NCBI.rev.compl                        ------------------------------------------------------------ 0
human.FAM230A.ncRNA.18487127.18500594-GGTLC3.18516335.18518165.rev.compl.REF      ------------------------------------------------------------ 0

chimp.LOC112206744-LOC107973052-GGT2.revcompl                            gtatacattatatactgtattatatattatatactgtatatataatatatattatatata 611
chimp54546-135457.revcompl.Pan.troglodytes.clone.rp43-41g5.GenBank:AC099533.36      ATATATATTAAATGTCACATTATATATATTATATATTATATATAATATACATAATGTATA 892
human.FAM30B-LOC105372935-GGT2.NCBI.                                     ------------------------------------------------------------ 587
human.FAM230E-LOC105377182-GGT3P.NCBI.                                   ------------------------------------------------------------ 0
human.FAM230J-LOC105372942-GGTLC5P.NCBI.rev.compl                        ------------------------------------------------------------ 0
human.FAM230A.ncRNA.18487127.18500594-GGTLC3.18516335.18518165.rev.compl.REF      ------------------------------------------------------------ 0

chimp.LOC112206744-LOC107973052-GGT2.revcompl                            cagtatatagtacatattatatatactatatgtta--tataaaatatataatataaaata 669
chimp54546-135457.revcompl.Pan.troglodytes.clone.rp43-41g5.GenBank:AC099533.36      ACATATATAATATATAATATATAACAAATATATTAGGTATATAATATATCTTATATGACA 952
human.FAM30B-LOC105372935-GGT2.NCBI.                                     ------------------------------------------------------------ 587
human.FAM230E-LOC105377182-GGT3P.NCBI.                                   ------------------------------------------------------------ 0
human.FAM230J-LOC105372942-GGTLC5P.NCBI.rev.compl                        ------------------------------------------------------------ 0
human.FAM230A.ncRNA.18487127.18500594-GGTLC3.18516335.18518165.rev.compl.REF      ------------------------------------------------------------ 0
```

```
chimp.LOC112206744-LOC107973052-GGT2.revcompl                               tataatatataatatataatatataatatataatatgtataatatataatatataatata 729
chimp54546-135457.revcompl.Pan.troglodytes.clone.rp43-41g5.GenBank:AC099533.36   TATAATATATATTATGTTATATGACATATATAATATATCTTACATATTATATATTATA-- 1010
human.FAM30B-LOC105372935-GGT2.NCBI.                                        ------------------------------------------------------------ 587
human.FAM230E-LOC105377182-GGT3P.NCBI.                                      ------------------------------------------------------------ 0
human.FAM230J-LOC105372942-GGTLC5P.NCBI.rev.compl                           ------------------------------------------------------------ 0
human.FAM230A.ncRNA.18487127.18500594-GGTLC3.18516335.18518165.rev.compl.REF ------------------------------------------------------------ 0

chimp.LOC112206744-LOC107973052-GGT2.revcompl                               tagtatatatatcgtataatatatataatatatattatatataatatatattataaatta 789
chimp54546-135457.revcompl.Pan.troglodytes.clone.rp43-41g5.GenBank:AC099533.36   ------ATATATTATATATAATATATTATATATAATATATATAATATATATTATATATTT 1064
human.FAM30B-LOC105372935-GGT2.NCBI.                                        -------------------------------gattcagtttattcccaaattcccaatta 617
human.FAM230E-LOC105377182-GGT3P.NCBI.                                      ------------------------------------------------------------ 0
human.FAM230J-LOC105372942-GGTLC5P.NCBI.rev.compl                           ------------------------------------------------------------ 0
human.FAM230A.ncRNA.18487127.18500594-GGTLC3.18516335.18518165.rev.compl.REF ------------------------------------------------------------ 0

chimp.LOC112206744-LOC107973052-GGT2.revcompl                               taatttataatttatatatattatatatataataataaattataattataaattataatt 849
chimp54546-135457.revcompl.Pan.troglodytes.clone.rp43-41g5.GenBank:AC099533.36   T-ATATATATGTTATATATATAATATATAC-ATTATATATCATATATATTATATGTAATA 1122
human.FAM30B-LOC105372935-GGT2.NCBI.                                        t-atatatata-tacatataaaatacatat-ataatatattaaatatattatatatataa 674
human.FAM230E-LOC105377182-GGT3P.NCBI.                                      ------------------------------------------------------------ 0
human.FAM230J-LOC105372942-GGTLC5P.NCBI.rev.compl                           ------------------------------------------------------------ 0
human.FAM230A.ncRNA.18487127.18500594-GGTLC3.18516335.18518165.rev.compl.REF ------------------------------------------------------------ 0

chimp.LOC112206744-LOC107973052-GGT2.revcompl                               tataatatatattatatataatatatattatatattataatttataatatacattattta 909
chimp54546-135457.revcompl.Pan.troglodytes.clone.rp43-41g5.GenBank:AC099533.36   TATATCATATATTACATATAATATATATCATATATTAT--------------ATATGTAT 1168
human.FAM30B-LOC105372935-GGT2.NCBI.                                        tatatattatattatatataat-------------------------------------- 696
human.FAM230E-LOC105377182-GGT3P.NCBI.                                      ------------------------------------------------------------ 0
human.FAM230J-LOC105372942-GGTLC5P.NCBI.rev.compl                           ------------------------------------------------------------ 0
human.FAM230A.ncRNA.18487127.18500594-GGTLC3.18516335.18518165.rev.compl.REF ------------------------------------------------------------ 0

chimp.LOC112206744-LOC107973052-GGT2.revcompl                               tatataatatataatgtatattatatattatatatttaatatataatatgttatatataa 969
chimp54546-135457.revcompl.Pan.troglodytes.clone.rp43-41g5.GenBank:AC099533.36   TATATATTATATATCATATATATTATATTATATATATCATATATAATATAATATATATAA 1228
human.FAM30B-LOC105372935-GGT2.NCBI.                                        ------------------------------------------------------------ 696
human.FAM230E-LOC105377182-GGT3P.NCBI.                                      ------------------------------------------------------------ 0
human.FAM230J-LOC105372942-GGTLC5P.NCBI.rev.compl                           ------------------------------------------------------------ 0
human.FAM230A.ncRNA.18487127.18500594-GGTLC3.18516335.18518165.rev.compl.REF ------------------------------------------------------------ 0

chimp.LOC112206744-LOC107973052-GGT2.revcompl                               tatataatatat------------------------------------------------ 981
chimp54546-135457.revcompl.Pan.troglodytes.clone.rp43-41g5.GenBank:AC099533.36   TATGTATTATATATAATTATTATACATAAGTATATATAATAATTATATATAATATAAGGA 1288
human.FAM30B-LOC105372935-GGT2.NCBI.                                        ------------------------------------------------------------ 696
human.FAM230E-LOC105377182-GGT3P.NCBI.                                      ------------------------------------------------------------ 0
human.FAM230J-LOC105372942-GGTLC5P.NCBI.rev.compl                           ------------------------------------------------------------ 0
human.FAM230A.ncRNA.18487127.18500594-GGTLC3.18516335.18518165.rev.compl.REF ------------------------------------------------------------ 0

chimp.LOC112206744-LOC107973052-GGT2.revcompl                               ------------------------------------------------------------ 981
chimp54546-135457.revcompl.Pan.troglodytes.clone.rp43-41g5.GenBank:AC099533.36   TGCAGGATGTAAAAGGAAATTATATATATGTTATATATATTATATTATATATATATGTAT 1348
human.FAM30B-LOC105372935-GGT2.NCBI.                                        -------------------------------------------------tatatatatat 707
human.FAM230E-LOC105377182-GGT3P.NCBI.                                      ------------------------------------------------------------ 0
human.FAM230J-LOC105372942-GGTLC5P.NCBI.rev.compl                           ------------------------------------------------------------ 0
human.FAM230A.ncRNA.18487127.18500594-GGTLC3.18516335.18518165.rev.compl.REF ------------------------------------------------------------ 0

chimp.LOC112206744-LOC107973052-GGT2.revcompl                               ------------------------------------------------------------ 981
chimp54546-135457.revcompl.Pan.troglodytes.clone.rp43-41g5.GenBank:AC099533.36   ATATTTGGGGGTGCCCTATTTCCCATCTCATAACTTATTTTAAGAAGCACAGCATAATAA 1408
human.FAM30B-LOC105372935-GGT2.NCBI.                                        atatttgtgggtgccctatttcccgtctcataacttattttaagaagc-cagcataataa 766
human.FAM230E-LOC105377182-GGT3P.NCBI.                                      ------------------------------------------------------------ 0
human.FAM230J-LOC105372942-GGTLC5P.NCBI.rev.compl                           ------------------------------------------------------------ 0
human.FAM230A.ncRNA.18487127.18500594-GGTLC3.18516335.18518165.rev.compl.REF ------------------------------------------------------------ 0

chimp.LOC112206744-LOC107973052-GGT2.revcompl                               ------------------------------------------------------------ 981
chimp54546-135457.revcompl.Pan.troglodytes.clone.rp43-41g5.GenBank:AC099533.36   TGTGTGGGCTTGGGATTCAGTTTTTGAAACAAAACACTGAGCCTTCAATGACCTTCCTGT 1468
human.FAM30B-LOC105372935-GGT2.NCBI.                                        tgtgtgggcttgggattcagtttttgaaacaaaacactgagcctttgatgaccttcctgt 826
human.FAM230E-LOC105377182-GGT3P.NCBI.                                      ------------------------------------------------------------ 0
human.FAM230J-LOC105372942-GGTLC5P.NCBI.rev.compl                           ------------------------------------------------------------ 0
human.FAM230A.ncRNA.18487127.18500594-GGTLC3.18516335.18518165.rev.compl.REF ------------------------------------------------------------ 0

chimp.LOC112206744-LOC107973052-GGT2.revcompl                               ------------------------------------------------------------ 981
chimp54546-135457.revcompl.Pan.troglodytes.clone.rp43-41g5.GenBank:AC099533.36   ACATGTAAAAGCACACCTGTCTGCATGGCAGCAGTTGGACCTCACAGTGTGGATTGTGCC 1528
human.FAM30B-LOC105372935-GGT2.NCBI.                                        acttgtaaaagcccacctgtctgcatggcagcagttggacctcacagtgtggattgtgcc 886
human.FAM230E-LOC105377182-GGT3P.NCBI.                                      ------------------------------------------------------------ 0
human.FAM230J-LOC105372942-GGTLC5P.NCBI.rev.compl                           ------------------------------------------------------------ 0
human.FAM230A.ncRNA.18487127.18500594-GGTLC3.18516335.18518165.rev.compl.REF ------------------------------------------------------------ 0

chimp.LOC112206744-LOC107973052-GGT2.revcompl                               ------------------------------------------------------------ 981
chimp54546-135457.revcompl.Pan.troglodytes.clone.rp43-41g5.GenBank:AC099533.36   TTCACCCTGGAATGTTTATGCCCTATCGCCATGGTGATGGGATTAGGGATCTCCTGCCTT 1588
human.FAM30B-LOC105372935-GGT2.NCBI.                                        ttcaccctggaatgtttatgccctatcgccatggtgatgggattagggatctcctgccct 946
human.FAM230E-LOC105377182-GGT3P.NCBI.                                      ----------------------------------------gattagggatctcctgccct 20
human.FAM230J-LOC105372942-GGTLC5P.NCBI.rev.compl                           ----------------------------------------gattagggatctcctgccct 20
human.FAM230A.ncRNA.18487127.18500594-GGTLC3.18516335.18518165.rev.compl.REF ------------------------------------------------------------ 0

chimp.LOC112206744-LOC107973052-GGT2.revcompl                               ------------------------------------------------------------ 981
chimp54546-135457.revcompl.Pan.troglodytes.clone.rp43-41g5.GenBank:AC099533.36   TGGTCCTAAGTGCCACTATCTGTGCTGAGTTTTTCAAAGGTCAGAGCAGATTGAATCTTT 1648
human.FAM30B-LOC105372935-GGT2.NCBI.                                        tggtcctaagtgccactatctgtgctgagtttttcaaaggtcagagcagattgaaccatt 1006
human.FAM230E-LOC105377182-GGT3P.NCBI.                                      tggtcctaagtgccactatctgtgctgagtttttcaaaggtcagagcagattgaaccatt 80
human.FAM230J-LOC105372942-GGTLC5P.NCBI.rev.compl                           tggtcgtaagtgccactacctgtgctgagtttttcaaaggtcagagcagattgaaccatt 80
human.FAM230A.ncRNA.18487127.18500594-GGTLC3.18516335.18518165.rev.compl.REF ------------------------------------------------------------ 0

chimp.LOC112206744-LOC107973052-GGT2.revcompl                               ------------------------------------------------------------ 981
chimp54546-135457.revcompl.Pan.troglodytes.clone.rp43-41g5.GenBank:AC099533.36   GTGGTTTCATTTTCCCTGATTTTGATTTTTCTCATGGGGAACCTGTGTGGCTGCATTCAA 1708
human.FAM30B-LOC105372935-GGT2.NCBI.                                        gtggtttcattttccctgattttgatttttcttatggggaacctgtgtggctgcattcaa 1066
human.FAM230E-LOC105377182-GGT3P.NCBI.                                      gtggtttcattttccctgattttgatttttcttatggggaacctgtgtggctgcattcaa 140
human.FAM230J-LOC105372942-GGTLC5P.NCBI.rev.compl                           gtggtttcattttccctgattttgatttttcttatggggaacctgtgtggctgcattcaa 140
human.FAM230A.ncRNA.18487127.18500594-GGTLC3.18516335.18518165.rev.compl.REF ------------------------------------------------------------ 0

chimp.LOC112206744-LOC107973052-GGT2.revcompl                               ------------------------------------------------------------ 981
chimp54546-135457.revcompl.Pan.troglodytes.clone.rp43-41g5.GenBank:AC099533.36   GGTATGTTCATACTGGCCTGTCAAGTGTGATCTTTTCAAATTACTAGTTAATGCTTTCAA 1768
human.FAM30B-LOC105372935-GGT2.NCBI.                                        ggtatgttcatactggcctgtcaaatgcgatcttttcaaattactagttaatgctttcaa 1126
human.FAM230E-LOC105377182-GGT3P.NCBI.                                      ggtatgttcatactggcctgtcaaatgcgatcttttcaaattactagttaatgctttcaa 200
human.FAM230J-LOC105372942-GGTLC5P.NCBI.rev.compl                           ggtatgttcatactggcctgtcaaatgcgatcttttcaaattactagttaatgctttcaa 200
human.FAM230A.ncRNA.18487127.18500594-GGTLC3.18516335.18518165.rev.compl.REF ------------------------------------------------------------ 0

chimp.LOC112206744-LOC107973052-GGT2.revcompl                               ------------------------------------------------------------ 981
chimp54546-135457.revcompl.Pan.troglodytes.clone.rp43-41g5.GenBank:AC099533.36   AATATGTTATTTAAAAAATTATCCTCTGTATTTTCCATATGCAGTTATAAATATGTTTCA 1828
human.FAM30B-LOC105372935-GGT2.NCBI.                                        aatatgttatttaaaaaattagcctctgtattttccatatgcagttataaatatgtttca 1186
human.FAM230E-LOC105377182-GGT3P.NCBI.                                      aatatgttatttaaaaaattagcctctgtattttccatatgcagttataaatatgtttca 260
human.FAM230J-LOC105372942-GGTLC5P.NCBI.rev.compl                           aatatgttatttaaaaaattagcctctgtattttccatatgcagttataaatatgtttca 260
human.FAM230A.ncRNA.18487127.18500594-GGTLC3.18516335.18518165.rev.compl.REF ------------------------------------------------------------ 0

chimp.LOC112206744-LOC107973052-GGT2.revcompl                               ------------------------------------------------------------ 981
chimp54546-135457.revcompl.Pan.troglodytes.clone.rp43-41g5.GenBank:AC099533.36   TGGTTATGTTTTATTCCTCAATTTATAGATTTGATTATTGTACCAAGCAGAGTACCTTTG 1888
human.FAM30B-LOC105372935-GGT2.NCBI.                                        tgattatgttttattcctcaatttatatatttgattattgtaccaagcagagtatctttg 1246
human.FAM230E-LOC105377182-GGT3P.NCBI.                                      tgattatgttttattcctcaatttatatatttgattattgtaccaagcagagtatctttg 320
human.FAM230J-LOC105372942-GGTLC5P.NCBI.rev.compl                           tgattatgttttattcctcaatttatatatttgattattgtaccaagcagagtatctttc 320
human.FAM230A.ncRNA.18487127.18500594-GGTLC3.18516335.18518165.rev.compl.REF ------------------------------------------------------------ 0

chimp.LOC112206744-LOC107973052-GGT2.revcompl                               ------------------------------------------------------------ 981
chimp54546-135457.revcompl.Pan.troglodytes.clone.rp43-41g5.GenBank:AC099533.36   AAATTTTTCTTCATTTAAAAAATATGTATCTTAACTCAGGCCTGTAATCCCAGCACATTG 1948
human.FAM30B-LOC105372935-GGT2.NCBI.                                        aaatttttcttcatttaaaaaatatgtatcttgactcaggcctgtaatcccagcactttg 1306
human.FAM230E-LOC105377182-GGT3P.NCBI.                                      aaatttttcttcatttaaaaaatatgtatcttgactcaggcctgtaatcccagcactttg 380
human.FAM230J-LOC105372942-GGTLC5P.NCBI.rev.compl                           aaatttttcttcatttaaaaaatatgtatcttgactcaggcctataatcccagcactttg 380
human.FAM230A.ncRNA.18487127.18500594-GGTLC3.18516335.18518165.rev.compl.REF ------------------------------------------------------------ 0

chimp.LOC112206744-LOC107973052-GGT2.revcompl                               ------------------------------------------------------------ 981
chimp54546-135457.revcompl.Pan.troglodytes.clone.rp43-41g5.GenBank:AC099533.36   GGAGTCCAAGGCAAGAGGATCACAAGGTGAGGAGATCAAGACCATCCTGCCCAATACAGT 2008
human.FAM30B-LOC105372935-GGT2.NCBI.                                        ggaggccaaggcaagaggatcacaaggtgaggagatcaagaccatcctggccaatacagt 1366
human.FAM230E-LOC105377182-GGT3P.NCBI.                                      ggaggccaaggcaagaggatcacaaggtgaggagatcaagaccatcctggccattacagt 440
human.FAM230J-LOC105372942-GGTLC5P.NCBI.rev.compl                           ggaggccaaggcaagaggatcacaaggtgaggagatcaagaccatcctggccaatacagt 440
human.FAM230A.ncRNA.18487127.18500594-GGTLC3.18516335.18518165.rev.compl.REF ------------------------------------------------------------ 0

chimp.LOC112206744-LOC107973052-GGT2.revcompl                               ------------------------------------------------------------ 981
```

```
chimp54546-135457.revcompl.Pan.troglodytes.clone.rp43-41g5.GenBank:AC099533.36      GAAACCCTGTCTCTACGAAAAATACAAAAAATTAGCCAGGCATGGTGGCAGCTGGTGTAG 2068
human.FAM30B-LOC105372935-GGT2.NCBI.                                                gaaaccctgtctctactacaaatacaaaaatttagccaggcatggtggcagctggtgtag 1426
human.FAM230E-LOC105377182-GGT3P.NCBI.                                              gaaaccctgtctctactacaaatacaaaaaattagccaggcatggtggcagctggtgtag 500
human.FAM230J-LOC105372942-GGTLC5P.NCBI.rev.compl                                   gaaaccctgtctctactacaaatacaaaaaattagccaggcatggtggcagctggtgtag 500
human.FAM230A.ncRNA.18487127.18500594-GGTLC3.18516335.18518165.rev.compl.REF        ------------------------------------------------------------ 0

chimp.LOC112206744-LOC107973052-GGT2.revcompl                                       ------------------------------------------------------------ 981
chimp54546-135457.revcompl.Pan.troglodytes.clone.rp43-41g5.GenBank:AC099533.36      TCCCAATGTGAATCGGG---------ATTCAGTTTATTCCCAAATTCCCAAAT------- 2112
human.FAM30B-LOC105372935-GGT2.NCBI.                                                tcccagtgtgaattgggattcagtttattcagtttattcccaaattcccaaat------- 1479
human.FAM230E-LOC105377182-GGT3P.NCBI.                                              tcccagtgtgaattgggattcagtttattcagtttattcccaaattccaaattatatat- 559
human.FAM230J-LOC105372942-GGTLC5P.NCBI.rev.compl                                   tcccagtgtgaattgggattcagtttattc--------ccaaattcccaaattatatata 552
human.FAM230A.ncRNA.18487127.18500594-GGTLC3.18516335.18518165.rev.compl.REF        ------------------------------------------------------------ 0

chimp.LOC112206744-LOC107973052-GGT2.revcompl                                       ------------------------------------------------------------ 981
chimp54546-135457.revcompl.Pan.troglodytes.clone.rp43-41g5.GenBank:AC099533.36      ------------------------------------------------------------ 2112
human.FAM30B-LOC105372935-GGT2.NCBI.                                                ------------------------------------------------------------ 1479
human.FAM230E-LOC105377182-GGT3P.NCBI.                                              ------------------------------------------------------------ 559
human.FAM230J-LOC105372942-GGTLC5P.NCBI.rev.compl                                   tatattatatatatatatataaaaaatatatatatatataaaatatatatataatgtatt 612
human.FAM230A.ncRNA.18487127.18500594-GGTLC3.18516335.18518165.rev.compl.REF        ------------------------------------------------------------ 0

chimp.LOC112206744-LOC107973052-GGT2.revcompl                                       ------------------------------------------------------------ 981
chimp54546-135457.revcompl.Pan.troglodytes.clone.rp43-41g5.GenBank:AC099533.36      ------------------------------------------------------------ 2112
human.FAM30B-LOC105372935-GGT2.NCBI.                                                ------------------------------------------------------------ 1479
human.FAM230E-LOC105377182-GGT3P.NCBI.                                              ------------------------------------------------------------ 559
human.FAM230J-LOC105372942-GGTLC5P.NCBI.rev.compl                                   taatatattatatataatatattatatatattatatataatatattatatatataatata 672
human.FAM230A.ncRNA.18487127.18500594-GGTLC3.18516335.18518165.rev.compl.REF        ------------------------------------------------------------ 0

chimp.LOC112206744-LOC107973052-GGT2.revcompl                                       ------------------------------------------------------------ 981
chimp54546-135457.revcompl.Pan.troglodytes.clone.rp43-41g5.GenBank:AC099533.36      ------------------------------------------------------------ 2112
human.FAM30B-LOC105372935-GGT2.NCBI.                                                ------------------------------------------------------------ 1479
human.FAM230E-LOC105377182-GGT3P.NCBI.                                              ------------------------------------------------------------ 559
human.FAM230J-LOC105372942-GGTLC5P.NCBI.rev.compl                                   tataatatattaaatataatatataatatatataatatattatatatattatagataata 732
human.FAM230A.ncRNA.18487127.18500594-GGTLC3.18516335.18518165.rev.compl.REF        ------------------------------------------------------------ 0

chimp.LOC112206744-LOC107973052-GGT2.revcompl                                       ------------------------------------------------------------ 981
chimp54546-135457.revcompl.Pan.troglodytes.clone.rp43-41g5.GenBank:AC099533.36      ------------------------------------------------------------ 2112
human.FAM30B-LOC105372935-GGT2.NCBI.                                                ------------------------------------------------------------ 1479
human.FAM230E-LOC105377182-GGT3P.NCBI.                                              ------------------------------------------------------------ 559
human.FAM230J-LOC105372942-GGTLC5P.NCBI.rev.compl                                   tatattatattatatataatatatattatacattatataattatatattatgtatataat 792
human.FAM230A.ncRNA.18487127.18500594-GGTLC3.18516335.18518165.rev.compl.REF        ------------------------------------------------------------ 0

chimp.LOC112206744-LOC107973052-GGT2.revcompl                                       ------------------------------------------------------------ 981
chimp54546-135457.revcompl.Pan.troglodytes.clone.rp43-41g5.GenBank:AC099533.36      ------------------------------------------------------------ 2112
human.FAM30B-LOC105372935-GGT2.NCBI.                                                ------------------------------------------------------------ 1479
human.FAM230E-LOC105377182-GGT3P.NCBI.                                              ------------------------------------------------------------ 559
human.FAM230J-LOC105372942-GGTLC5P.NCBI.rev.compl                                   atataatatataatatatattatataattatatattatgtacattatatattatttatat 852
human.FAM230A.ncRNA.18487127.18500594-GGTLC3.18516335.18518165.rev.compl.REF        ------------------------------------------------------------ 0

chimp.LOC112206744-LOC107973052-GGT2.revcompl                                       ------------------------------------------------------------ 981
chimp54546-135457.revcompl.Pan.troglodytes.clone.rp43-41g5.GenBank:AC099533.36      ------------------------------------------------------------ 2112
human.FAM30B-LOC105372935-GGT2.NCBI.                                                ------------------------------------------------------------ 1479
human.FAM230E-LOC105377182-GGT3P.NCBI.                                              ------------------------------------------------------------ 559
human.FAM230J-LOC105372942-GGTLC5P.NCBI.rev.compl                                   atattatatataacatataatatatataataaataatatatatattatatatatcatata 912
human.FAM230A.ncRNA.18487127.18500594-GGTLC3.18516335.18518165.rev.compl.REF        ------------------------------------------------------------ 0

chimp.LOC112206744-LOC107973052-GGT2.revcompl                                       -----------attgtatataatatatattatattatatattataatacataatataata 1030
chimp54546-135457.revcompl.Pan.troglodytes.clone.rp43-41g5.GenBank:AC099533.36      -----------TATATATATAATATATATAATATGTATTATATATACAGTGTATATAATA 2161
human.FAM30B-LOC105372935-GGT2.NCBI.                                                -----------tatat-------------------------------------------- 1484
human.FAM230E-LOC105377182-GGT3P.NCBI.                                              ------------------------------------------------------------ 559
human.FAM230J-LOC105372942-GGTLC5P.NCBI.rev.compl                                   atatataatatatata-------------------------------------------- 928
human.FAM230A.ncRNA.18487127.18500594-GGTLC3.18516335.18518165.rev.compl.REF        ------------------------------------------------------------ 0

chimp.LOC112206744-LOC107973052-GGT2.revcompl                                       tataatat------attatatattatatatattatata------atatattatatattat 1078
chimp54546-135457.revcompl.Pan.troglodytes.clone.rp43-41g5.GenBank:AC099533.36      TATATTATATATACATTATATATAATACATATTATATACACTGTATATATAATATATATT 2221
human.FAM30B-LOC105372935-GGT2.NCBI.                                                ------------------------------------------------------------ 1484
human.FAM230E-LOC105377182-GGT3P.NCBI.                                              ------------------------------------------------------------ 559
human.FAM230J-LOC105372942-GGTLC5P.NCBI.rev.compl                                   ------------------------------------------------------------ 928
human.FAM230A.ncRNA.18487127.18500594-GGTLC3.18516335.18518165.rev.compl.REF        ------------------------------------------------------------ 0

chimp.LOC112206744-LOC107973052-GGT2.revcompl                                       atatatatataatatataatatattatatatatgt-----------tatatattatatat 1127
chimp54546-135457.revcompl.Pan.troglodytes.clone.rp43-41g5.GenBank:AC099533.36      ATATATAAAATATAGATAATATATATTATATATACAGTAGGTAAAATATATATTACATAT 2281
human.FAM30B-LOC105372935-GGT2.NCBI.                                                ------------------------------------------------------------ 1484
human.FAM230E-LOC105377182-GGT3P.NCBI.                                              ------------------------------------------------------------ 559
human.FAM230J-LOC105372942-GGTLC5P.NCBI.rev.compl                                   ------------------------------------------------------------ 928
human.FAM230A.ncRNA.18487127.18500594-GGTLC3.18516335.18518165.rev.compl.REF        ------------------------------------------------------------ 0

chimp.LOC112206744-LOC107973052-GGT2.revcompl                                       atatgttatatattatagaatatgttatatattatatattatagaatatgttatatatta 1187
chimp54546-135457.revcompl.Pan.troglodytes.clone.rp43-41g5.GenBank:AC099533.36      ACTGTATATATAATATATATTATATATATTATATTATATTATATTATATTATATATATTA 2341
human.FAM30B-LOC105372935-GGT2.NCBI.                                                ------------------------------------------------------------ 1484
human.FAM230E-LOC105377182-GGT3P.NCBI.                                              ------------------------------------------------------------ 559
human.FAM230J-LOC105372942-GGTLC5P.NCBI.rev.compl                                   ------------------------------------------------------------ 928
human.FAM230A.ncRNA.18487127.18500594-GGTLC3.18516335.18518165.rev.compl.REF        ------------------------------------------------------------ 0

chimp.LOC112206744-LOC107973052-GGT2.revcompl                                       tatattatata------atatgttatatattatatattataga----------------- 1224
chimp54546-135457.revcompl.Pan.troglodytes.clone.rp43-41g5.GenBank:AC099533.36      TATATTATATATATGATATATGATATATAATATATATTATATATACAGTATATGTAATAT 2401
human.FAM30B-LOC105372935-GGT2.NCBI.                                                ------------------------------------------------------------ 1484
human.FAM230E-LOC105377182-GGT3P.NCBI.                                              ------------------------------------------------------------ 559
human.FAM230J-LOC105372942-GGTLC5P.NCBI.rev.compl                                   ------------------------------------------------------------ 928
human.FAM230A.ncRNA.18487127.18500594-GGTLC3.18516335.18518165.rev.compl.REF        ------------------------------------------------------------ 0

chimp.LOC112206744-LOC107973052-GGT2.revcompl                                       -----------------------------------------------------------a 1225
chimp54546-135457.revcompl.Pan.troglodytes.clone.rp43-41g5.GenBank:AC099533.36      GTATTATATATACAGTATATGTAATATGTATTATATATATTATTTATAATATGTATTATA 2461
human.FAM30B-LOC105372935-GGT2.NCBI.                                                ------------------------------------------------------------ 1484
human.FAM230E-LOC105377182-GGT3P.NCBI.                                              ------------------------------------------------------------ 559
human.FAM230J-LOC105372942-GGTLC5P.NCBI.rev.compl                                   ------------------------------------------------------------ 928
human.FAM230A.ncRNA.18487127.18500594-GGTLC3.18516335.18518165.rev.compl.REF        ------------------------------------------------------------ 0

chimp.LOC112206744-LOC107973052-GGT2.revcompl                                       tatgttatatattatataatatgttatatattatatattatataatatgttatatattat 1285
chimp54546-135457.revcompl.Pan.troglodytes.clone.rp43-41g5.GenBank:AC099533.36      TATATTATTTATAATAT-GTTTGATATATATTATATATAATAT-ATATTATATATATTAT 2519
human.FAM30B-LOC105372935-GGT2.NCBI.                                                ------------------------------------------------------------ 1484
human.FAM230E-LOC105377182-GGT3P.NCBI.                                              ------------------------------------------------------------ 559
human.FAM230J-LOC105372942-GGTLC5P.NCBI.rev.compl                                   ------------------------------------------------------------ 928
human.FAM230A.ncRNA.18487127.18500594-GGTLC3.18516335.18518165.rev.compl.REF        ------------------------------------------------------------ 0

chimp.LOC112206744-LOC107973052-GGT2.revcompl                                       atatattatataatatgttatatattatataatatgt----tatatattatatatattat 1341
chimp54546-135457.revcompl.Pan.troglodytes.clone.rp43-41g5.GenBank:AC099533.36      AAATAATATATAGTATA-TATATTATATATAATATATAGTATATATATTATATATAATAT 2578
human.FAM30B-LOC105372935-GGT2.NCBI.                                                ------------------------------------------------------------ 1484
human.FAM230E-LOC105377182-GGT3P.NCBI.                                              ------------------------------------------------------------ 559
human.FAM230J-LOC105372942-GGTLC5P.NCBI.rev.compl                                   ------------------------------------------------------------ 928
human.FAM230A.ncRNA.18487127.18500594-GGTLC3.18516335.18518165.rev.compl.REF        ------------------------------------------------------------ 0

chimp.LOC112206744-LOC107973052-GGT2.revcompl                                       ataatatgt----tatatattatatatattatataatatgt------tatatattatata 1391
chimp54546-135457.revcompl.Pan.troglodytes.clone.rp43-41g5.GenBank:AC099533.36      ATAATATATAGTATATATATTATATATTATATATAATATATATAGTATATATATTATATA 2638
human.FAM30B-LOC105372935-GGT2.NCBI.                                                ------------------------------------------------------------ 1484
human.FAM230E-LOC105377182-GGT3P.NCBI.                                              ------------------------------------------------------------ 559
human.FAM230J-LOC105372942-GGTLC5P.NCBI.rev.compl                                   ------------------------------------------------------------ 928
human.FAM230A.ncRNA.18487127.18500594-GGTLC3.18516335.18518165.rev.compl.REF        ------------------------------------------------------------ 0

chimp.LOC112206744-LOC107973052-GGT2.revcompl                                       ttatataatatgttatatattatatatattatataatatgttatatattatatattatat 1451
chimp54546-135457.revcompl.Pan.troglodytes.clone.rp43-41g5.GenBank:AC099533.36      TAGTAT-ATAGTATATATATTATTTATAATATATAGTA---TATATATTATTTATAATAT 2694
human.FAM30B-LOC105372935-GGT2.NCBI.                                                ------------------------------------------------------------ 1484
human.FAM230E-LOC105377182-GGT3P.NCBI.                                              ------------------------------------------------------------ 559
human.FAM230J-LOC105372942-GGTLC5P.NCBI.rev.compl                                   ------------------------------------------------------------ 928
human.FAM230A.ncRNA.18487127.18500594-GGTLC3.18516335.18518165.rev.compl.REF        ------------------------------------------------------------ 0

chimp.LOC112206744-LOC107973052-GGT2.revcompl                                       aatatgttatatattatatatattatataatatgttatatattatatattatataatatg 1511
chimp54546-135457.revcompl.Pan.troglodytes.clone.rp43-41g5.GenBank:AC099533.36      -ATAGTATATATATTATTTATATTCCAGGATATATTATATTTTATAATATATATTATTTC 2753
```

```
human.FAM30B-LOC105372935-GGT2.NCBI.                                              ------------------------------------------------------------ 1484
human.FAM230E-LOC105377182-GGT3P.NCBI.                                            ------------------------------------------------------------ 559
human.FAM230J-LOC105372942-GGTLC5P.NCBI.rev.compl                                 ------------------------------------------------------------ 928
human.FAM230A.ncRNA.18487127.18500594-GGTLC3.18516335.18518165.rev.compl.REF      ------------------------------------------------------------ 0

chimp.LOC112206744-LOC107973052-GGT2.revcompl                                     t------tatatattatatatattatataatatatattatatcttatataatataattta 1565
chimp54546-135457.revcompl.Pan.troglodytes.clone.rp43-41g5.GenBank:AC099533.36    CATACTATATATAATGTGTATATTATATATTATATATTTAATA-TATAATGTATATATTG 2812
human.FAM30B-LOC105372935-GGT2.NCBI.                                              ------------------------------------------------------------ 1484
human.FAM230E-LOC105377182-GGT3P.NCBI.                                            ------------------------------------------------------------ 559
human.FAM230J-LOC105372942-GGTLC5P.NCBI.rev.compl                                 ------------------------------------------------------------ 928
human.FAM230A.ncRNA.18487127.18500594-GGTLC3.18516335.18518165.rev.compl.REF      ------------------------------------------------------------ 0

chimp.LOC112206744-LOC107973052-GGT2.revcompl                                     tatattatatattatacaatatatattatatattatattatatattatacaatatatatt 1625
chimp54546-135457.revcompl.Pan.troglodytes.clone.rp43-41g5.GenBank:AC099533.36    TATGTAATATATTATATTATATGTTATATGTAATATATTATATATTATATGTTATATGTA 2872
human.FAM30B-LOC105372935-GGT2.NCBI.                                              ------------------------------------------------------------ 1484
human.FAM230E-LOC105377182-GGT3P.NCBI.                                            ------------------------------------------------------------ 559
human.FAM230J-LOC105372942-GGTLC5P.NCBI.rev.compl                                 ------------------------------------------------------------ 928
human.FAM230A.ncRNA.18487127.18500594-GGTLC3.18516335.18518165.rev.compl.REF      ------------------------------------------------------------ 0

chimp.LOC112206744-LOC107973052-GGT2.revcompl                                     atata-tatattatatattatacaatatatattatataatatataatatatattatat-- 1682
chimp54546-135457.revcompl.Pan.troglodytes.clone.rp43-41g5.GenBank:AC099533.36    ATATATTATATTATATATTATATGTAATATATTATATATTATATGTAATATATTATATAT 2932
human.FAM30B-LOC105372935-GGT2.NCBI.                                              ------------------------------------------------------------ 1484
human.FAM230E-LOC105377182-GGT3P.NCBI.                                            ------------------------------------------------------------ 559
human.FAM230J-LOC105372942-GGTLC5P.NCBI.rev.compl                                 ------------------------------------------------------------ 928
human.FAM230A.ncRNA.18487127.18500594-GGTLC3.18516335.18518165.rev.compl.REF      ------------------------------------------------------------ 0

chimp.LOC112206744-LOC107973052-GGT2.revcompl                                     --tatataatatatatataatttccttttacatgctgcatccttcaacattccatccccc 1740
chimp54546-135457.revcompl.Pan.troglodytes.clone.rp43-41g5.GenBank:AC099533.36    TACATATAATATATATATAATTTCCTTTTACATCCTGCATCCTTCAACGTTCCATCCCCC 2992
human.FAM30B-LOC105372935-GGT2.NCBI.                                              --------atatatatataatttccttttacatcctgcatccttcaacgttacatccccc 1536
human.FAM230E-LOC105377182-GGT3P.NCBI.                                            --------atatatatataatttccttttacatcctgcatccttcaacgttccatccccc 611
human.FAM230J-LOC105372942-GGTLC5P.NCBI.rev.compl                                 --------atatatatataatttccttttacatcctgcatccttcaacgttccattcccc 980
human.FAM230A.ncRNA.18487127.18500594-GGTLC3.18516335.18518165.rev.compl.REF      ------------------------------------------------------------ 0

chimp.LOC112206744-LOC107973052-GGT2.revcompl                                     accccacagattaagttattccccaggggagaatatggcaaactctattttaatgcagtt 1800
chimp54546-135457.revcompl.Pan.troglodytes.clone.rp43-41g5.GenBank:AC099533.36    ACCCCACAGATTAAGTTATTCCCCAAGGGAGAATATGGCAAAGTCTATTTTAATGCAGTA 3052
human.FAM30B-LOC105372935-GGT2.NCBI.                                              accccacagattaagttattccccaggggagaatatggcaaagtctattttaatgcagtt 1596
human.FAM230E-LOC105377182-GGT3P.NCBI.                                            accccacagattaagttattccccaggggagaatatggcaaagtctattttaatgcagtt 671
human.FAM230J-LOC105372942-GGTLC5P.NCBI.rev.compl                                 accccacagattaagttattccccaggggagaatatggcagagtctattttaatgctgtt 1040
human.FAM230A.ncRNA.18487127.18500594-GGTLC3.18516335.18518165.rev.compl.REF      ------------------------------------------------------------ 0

chimp.LOC112206744-LOC107973052-GGT2.revcompl                                     tttagcctaatt------------------------------------------------ 1812
chimp54546-135457.revcompl.Pan.troglodytes.clone.rp43-41g5.GenBank:AC099533.36    GTTACCCTAATTAAGAACTTATGAGGCCGGGCGCGGTGGCTCACGACTGTAATCCCAGCA 3112
human.FAM30B-LOC105372935-GGT2.NCBI.                                              tttaacccaatt------------------------------------------------ 1608
human.FAM230E-LOC105377182-GGT3P.NCBI.                                            tttaacccaatt------------------------------------------------ 683
human.FAM230J-LOC105372942-GGTLC5P.NCBI.rev.compl                                 tttaacccaatt------------------------------------------------ 1052
human.FAM230A.ncRNA.18487127.18500594-GGTLC3.18516335.18518165.rev.compl.REF      ------------------------------------------------------------ 0

chimp.LOC112206744-LOC107973052-GGT2.revcompl                                     ------------------------------------------------------------ 1812
chimp54546-135457.revcompl.Pan.troglodytes.clone.rp43-41g5.GenBank:AC099533.36    CTTTGGGAGGCCGAGGTGGGGGGATCACGAGGTCAGGAGATCGAGACCATCCTGGCTAAC 3172
human.FAM30B-LOC105372935-GGT2.NCBI.                                              ------------------------------------------------------------ 1608
human.FAM230E-LOC105377182-GGT3P.NCBI.                                            ------------------------------------------------------------ 683
human.FAM230J-LOC105372942-GGTLC5P.NCBI.rev.compl                                 ------------------------------------------------------------ 1052
human.FAM230A.ncRNA.18487127.18500594-GGTLC3.18516335.18518165.rev.compl.REF      ------------------------------------------------------------ 0

chimp.LOC112206744-LOC107973052-GGT2.revcompl                                     ------------------------------------------------------------ 1812
chimp54546-135457.revcompl.Pan.troglodytes.clone.rp43-41g5.GenBank:AC099533.36    ATGGTGAAACCCCGTCTCTACTAAAAATACAAAAAATTAGCCGGGCGTGGTGGCGGGCGC 3232
human.FAM30B-LOC105372935-GGT2.NCBI.                                              ------------------------------------------------------------ 1608
human.FAM230E-LOC105377182-GGT3P.NCBI.                                            ------------------------------------------------------------ 683
human.FAM230J-LOC105372942-GGTLC5P.NCBI.rev.compl                                 ------------------------------------------------------------ 1052
human.FAM230A.ncRNA.18487127.18500594-GGTLC3.18516335.18518165.rev.compl.REF      ------------------------------------------------------------ 0

chimp.LOC112206744-LOC107973052-GGT2.revcompl                                     ------------------------------------------------------------ 1812
chimp54546-135457.revcompl.Pan.troglodytes.clone.rp43-41g5.GenBank:AC099533.36    CTGTAGTCCCAGCTACTTGGGAGGCTGAGGCAGGAGAATGGCGTGAACCCGGGAGGCGGA 3292
human.FAM30B-LOC105372935-GGT2.NCBI.                                              ------------------------------------------------------------ 1608
human.FAM230E-LOC105377182-GGT3P.NCBI.                                            ------------------------------------------------------------ 683
human.FAM230J-LOC105372942-GGTLC5P.NCBI.rev.compl                                 ------------------------------------------------------------ 1052
human.FAM230A.ncRNA.18487127.18500594-GGTLC3.18516335.18518165.rev.compl.REF      ------------------------------------------------------------ 0

chimp.LOC112206744-LOC107973052-GGT2.revcompl                                     ------------------------------------------------------------ 1812
chimp54546-135457.revcompl.Pan.troglodytes.clone.rp43-41g5.GenBank:AC099533.36    GCTTGCAGTGAGCCAAGATCGCGCCACTGCACTCCAGCCTGGGCGACAGAGCGAGACTCC 3352
human.FAM30B-LOC105372935-GGT2.NCBI.                                              ------------------------------------------------------------ 1608
human.FAM230E-LOC105377182-GGT3P.NCBI.                                            ------------------------------------------------------------ 683
human.FAM230J-LOC105372942-GGTLC5P.NCBI.rev.compl                                 ------------------------------------------------------------ 1052
human.FAM230A.ncRNA.18487127.18500594-GGTLC3.18516335.18518165.rev.compl.REF      ------------------------------------------------------------ 0

chimp.LOC112206744-LOC107973052-GGT2.revcompl                                     -------------------------aagaa-ttatgaaatcattactttccaaatatttg 1846
chimp54546-135457.revcompl.Pan.troglodytes.clone.rp43-41g5.GenBank:AC099533.36    GTCTCAAAAAAAAAAAAAAAAAAAAAGAACTTATGAAATCATTACTTTCCAAATCTTTG 3412
human.FAM30B-LOC105372935-GGT2.NCBI.                                              -------------------------aagaacctatgaaatcattactttccaaaactttg 1643
human.FAM230E-LOC105377182-GGT3P.NCBI.                                            -------------------------aagaacctatgaaatcattactttccaaaactttg 718
human.FAM230J-LOC105372942-GGTLC5P.NCBI.rev.compl                                 -------------------------aagaacctatgaaatcattactttccaaaactttg 1087
human.FAM230A.ncRNA.18487127.18500594-GGTLC3.18516335.18518165.rev.compl.REF      ------------------------------------------------------------ 0

chimp.LOC112206744-LOC107973052-GGT2.revcompl                                     gaacaaagccacagtagtatgcatgggttggaggcttttcacagaataaaatgtacctat 1906
chimp54546-135457.revcompl.Pan.troglodytes.clone.rp43-41g5.GenBank:AC099533.36    GAACAAAGCCACAGTAGTAGGGATGGGTTGGAGGCTTTTCACACAATAAAATGTACCTCT 3472
human.FAM30B-LOC105372935-GGT2.NCBI.                                              gaacaaagccacagtagtatggatccgttggaggcttttcacacaataaaatgtacctct 1703
human.FAM230E-LOC105377182-GGT3P.NCBI.                                            gaacaaagccacagtagtatggatccgttggaggcttttcacacaataaaatgtacatct 778
human.FAM230J-LOC105372942-GGTLC5P.NCBI.rev.compl                                 gaacaaagccacagtagtaaggatccgttggaggcttttcacacaataaaatgtaactct 1147
human.FAM230A.ncRNA.18487127.18500594-GGTLC3.18516335.18518165.rev.compl.REF      ------------------------------------------------------------ 0

chimp.LOC112206744-LOC107973052-GGT2.revcompl                                     ctttgtttttaacatgtttttcccttcctctcttctttctttgtgaaatgtgtatttact 1966
chimp54546-135457.revcompl.Pan.troglodytes.clone.rp43-41g5.GenBank:AC099533.36    CTTTGTTTTTAACATGTTTTTCCCTTCCTCTCTTCTTTCTTTGTGAAATGTGTATTTACT 3532
human.FAM30B-LOC105372935-GGT2.NCBI.                                              ctttgtttttaacatgtttttcccttcctctcttctttttttgtgaaatgtgtatttact 1763
human.FAM230E-LOC105377182-GGT3P.NCBI.                                            ctttgtttttaacatgtttttcccttcctctcttctttttttgtgaaatgtgtatttact 838
human.FAM230J-LOC105372942-GGTLC5P.NCBI.rev.compl                                 ctttgtttttaacatgtttttcccttcctctcttctttttttgtgaaatgtgtatttact 1207
human.FAM230A.ncRNA.18487127.18500594-GGTLC3.18516335.18518165.rev.compl.REF      ------------------------------------------------------------ 0

chimp.LOC112206744-LOC107973052-GGT2.revcompl                                     ttaataaatttgtagtaagtcacttccattcacatattaatttttaaagtaataagtat 2026
chimp54546-135457.revcompl.Pan.troglodytes.clone.rp43-41g5.GenBank:AC099533.36    TTAATAAGTTTGTAGTAAGTCACTTCTATTCACATATTAATTTTTAAAGTAATAAGTAT 3592
human.FAM30B-LOC105372935-GGT2.NCBI.                                              ttaatatatttgtagtaagtcacttccatgcacatattaatttttaaagtaataagtat 1823
human.FAM230E-LOC105377182-GGT3P.NCBI.                                            ttaatatatttgtagtaagtcacttccatgcacatattaatttttaaagtaataagtat 898
human.FAM230J-LOC105372942-GGTLC5P.NCBI.rev.compl                                 ttaatatatttgtagtaagtcacttccatgcacatattaatttttaaagtaataagcat 1267
human.FAM230A.ncRNA.18487127.18500594-GGTLC3.18516335.18518165.rev.compl.REF      ------------------------------------------------------------ 0

chimp.LOC112206744-LOC107973052-GGT2.revcompl                                     gtgtattgtctactagtgaaataaaacacacatttatttttattctttggaagttatcca 2086
chimp54546-135457.revcompl.Pan.troglodytes.clone.rp43-41g5.GenBank:AC099533.36    GTGTATTGTCTACGTGTGAAATAAAACACACATTTATTTTTATGCTTTGGAAGTTATCCA 3652
human.FAM30B-LOC105372935-GGT2.NCBI.                                              gtgtattgtctacgtgtgaaagaaaacacacatttatttttatgctttggaagttatcca 1883
human.FAM230E-LOC105377182-GGT3P.NCBI.                                            gtgtattgtctacgtgtgaaagaaaacacacatttatttttatgccttggaagttatcca 958
human.FAM230J-LOC105372942-GGTLC5P.NCBI.rev.compl                                 gtgtattgtctacgtgtgaaagaaaacacacatttatttttatgctttggaagttatcca 1327
human.FAM230A.ncRNA.18487127.18500594-GGTLC3.18516335.18518165.rev.compl.REF      ------------------------------------------------------------ 0

chimp.LOC112206744-LOC107973052-GGT2.revcompl                                     gaatcatggcattgtcaatcacagtcaatcacccaacctactcacctttccagtgtaatc 2146
chimp54546-135457.revcompl.Pan.troglodytes.clone.rp43-41g5.GenBank:AC099533.36    GAATCATGGAATTGTCAATCACAGTCAATCACCCAACCTACTCACCTTTCCAGTGTAATC 3712
human.FAM30B-LOC105372935-GGT2.NCBI.                                              gaatcatggaattgtcaatcacagtcaatcacccaacctactcacctttccagtgtaatc 1943
human.FAM230E-LOC105377182-GGT3P.NCBI.                                            gaatcatggaattgtcaatcacagtcaatcacccaacctactcacctttccagtgtaatc 1018
human.FAM230J-LOC105372942-GGTLC5P.NCBI.rev.compl                                 gaatcatggaattgtcaatcacagtcaatcacccaacctactcacctttccagtgtaatc 1387
human.FAM230A.ncRNA.18487127.18500594-GGTLC3.18516335.18518165.rev.compl.REF      ------------------------------------------------------------ 0

chimp.LOC112206744-LOC107973052-GGT2.revcompl                                     ttagtctaa-ttttttttttgttatccaatgagatgcagtatttcaactcagagagataa 2205
chimp54546-135457.revcompl.Pan.troglodytes.clone.rp43-41g5.GenBank:AC099533.36    TTAGTCAAG--TTTTTTTTTGTTATCCAATGAGATGCAGTATTTCAACTCAGAAAGATAA 3770
human.FAM30B-LOC105372935-GGT2.NCBI.                                              ttagtcaaa-ttttttttttgttatccaatgagatgcagtatttcaactcagaaagataa 2002
human.FAM230E-LOC105377182-GGT3P.NCBI.                                            ttagtcaaa-ttttttttttgttatccaatgagatgcagtatttcaactcagaaagataa 1077
human.FAM230J-LOC105372942-GGTLC5P.NCBI.rev.compl                                 ttagtcaaattttttttttttgttatccaatgagatgcagtatttcaactcagaaagataa 1447
human.FAM230A.ncRNA.18487127.18500594-GGTLC3.18516335.18518165.rev.compl.REF      ------------------------------------------------------------ 0

chimp.LOC112206744-LOC107973052-GGT2.revcompl                                     atagaatgaattggtagagactattaactaagaatatacagttttatttatgctcagaag 2265
chimp54546-135457.revcompl.Pan.troglodytes.clone.rp43-41g5.GenBank:AC099533.36    ATAGAATGAATTGATAGAGACTATTAACTAAGAATATACAGTTTTATTTATACTCAGAAG 3830
human.FAM30B-LOC105372935-GGT2.NCBI.                                              atagagtgaatttatagagactattaactaagaacatacagttttatttatactcagaag 2062
```

```
human.FAM230E-LOC105377182-GGT3P.NCBI.                                          atagagtgaatttatagagactattaactaagaacatacagttttatttatactcagaag 1137
human.FAM230J-LOC105372942-GGTLC5P.NCBI.rev.compl                               atagagtgaatttatagagactattaactaagaacatacagtttgatttatactcagaag 1507
human.FAM230A.ncRNA.18487127.18500594-GGTLC3.18516335.18518165.rev.compl.REF    ------------------------------------------------------------ 0

chimp.LOC112206744-LOC107973052-GGT2.revcompl                                   caagtagattatgtatatatgtatgaagataaaaattaaaaggataattgtgtaaatttg 2325
chimp54546-135457.revcompl.Pan.troglodytes.clone.rp43-41g5.GenBank:AC099533.36  CAAGTAGATTATGTATATATGTATGAAGATAAAAATTAAAAGGATAATTGTGTAAATTTG 3890
human.FAM30B-LOC105372935-GGT2.NCBI.                                            caagtagattatgtacatatatatgaagataaaaattaaaaggataattgtgtaaatttg 2122
human.FAM230E-LOC105377182-GGT3P.NCBI.                                          caagtagattatgtacatatatatgaagataaaaattaaaaggataattgtgtaaatttg 1197
human.FAM230J-LOC105372942-GGTLC5P.NCBI.rev.compl                               caagtagattatgtacatatatatgaagataaaaattaaaaggataattgtgtaaatttg 1567
human.FAM230A.ncRNA.18487127.18500594-GGTLC3.18516335.18518165.rev.compl.REF    ------------------------------------------------------------ 0

chimp.LOC112206744-LOC107973052-GGT2.revcompl                                   catgtagagagctttgaaaacctgtttacttgttaatgctgttttgatgtattgtgtgtc 2385
chimp54546-135457.revcompl.Pan.troglodytes.clone.rp43-41g5.GenBank:AC099533.36  CATGTGGAGAGCTTTGAAAACCTGTTTACTTGTTAATGCTGTTTTGATGTATTGTGTGTC 3950
human.FAM30B-LOC105372935-GGT2.NCBI.                                            catgtagagggctttgaaaacctgtttacttgttaatgctgttttgatgtat--tgtgtc 2180
human.FAM230E-LOC105377182-GGT3P.NCBI.                                          catgtagagggctttgaaaacctgtttacttgttaatgctgttttgatgtat--tgtgtc 1255
human.FAM230J-LOC105372942-GGTLC5P.NCBI.rev.compl                               catgtagagggctttgaaaacctgtttacttgtgaatgctgttttgatgtat--tgtgtc 1625
human.FAM230A.ncRNA.18487127.18500594-GGTLC3.18516335.18518165.rev.compl.REF    ------------------------------------------------------------ 0

chimp.LOC112206744-LOC107973052-GGT2.revcompl                                   tttgttctcccgacccagcatccagagctctctgctggagctaagtgctcatcagttcca 2445
chimp54546-135457.revcompl.Pan.troglodytes.clone.rp43-41g5.GenBank:AC099533.36  TTTGTTCTCCCGACCCAGCATCCAGAGCTCTCTGCTGGAGCTAAGTGCTCATCAGTTCCA 4010
human.FAM30B-LOC105372935-GGT2.NCBI.                                            tttgttctcccgacccatcgtccagagctctctgcaggagctaagtgctcatcagttcca 2240
human.FAM230E-LOC105377182-GGT3P.NCBI.                                          tttgttctcccgacccatcgtccagagctctctgcaggagctaagtgctcatcagttcca 1315
human.FAM230J-LOC105372942-GGTLC5P.NCBI.rev.compl                               tttgttctcccgacccatcatccagagctctctgcaggagctaagtgctcatcagttcca 1685
human.FAM230A.ncRNA.18487127.18500594-GGTLC3.18516335.18518165.rev.compl.REF    ------------------------------------------------------------ 0

chimp.LOC112206744-LOC107973052-GGT2.revcompl                                   tgacttggaaactgtctaagtttagaggcacttgtatttgttagtaaataagtcaagatg 2505
chimp54546-135457.revcompl.Pan.troglodytes.clone.rp43-41g5.GenBank:AC099533.36  TGACTTGGAAACTGTCTAAGTTTAGAGGCACTTGTATTTGTTAGTAAATAAGTCAAGATG 4070
human.FAM30B-LOC105372935-GGT2.NCBI.                                            tgacttggaaactgtctaagtttagaggcacttgtatttgttagtaaataaggcaagatg 2300
human.FAM230E-LOC105377182-GGT3P.NCBI.                                          tgacttggaaactgtctaagtttagaggcacttgtatttgttagtaaataaggcaagatg 1375
human.FAM230J-LOC105372942-GGTLC5P.NCBI.rev.compl                               tgacttggaaactgtctaagtttagaggcacttgtatttgttagtaaataaggcaagatg 1745
human.FAM230A.ncRNA.18487127.18500594-GGTLC3.18516335.18518165.rev.compl.REF    ------------------------------------------------------------ 0

chimp.LOC112206744-LOC107973052-GGT2.revcompl                                   atattgtttcacaggctttagtgcagaagactgaatagagaagctgctccacccagtaca 2565
chimp54546-135457.revcompl.Pan.troglodytes.clone.rp43-41g5.GenBank:AC099533.36  ATATTGTTTCACAGGCTTTAGTGCAGAAGACTGAATAGAGAAGCTGCTCCACCCAGTACA 4130
human.FAM30B-LOC105372935-GGT2.NCBI.                                            atattgtttcacaggttttagtgccgaagactgaatagataagctgctccacccagtaca 2360
human.FAM230E-LOC105377182-GGT3P.NCBI.                                          atattgtttcacaggttttagtgccgaagactgaatagataagctgctccacccagtaca 1435
human.FAM230J-LOC105372942-GGTLC5P.NCBI.rev.compl                               atattgtttcacaggttttagtgccgaagactgaatagataagctgctccacccagtaca 1805
human.FAM230A.ncRNA.18487127.18500594-GGTLC3.18516335.18518165.rev.compl.REF    ------------------------------------------------------------ 0

chimp.LOC112206744-LOC107973052-GGT2.revcompl                                   ctggtgttcatttcatggtcatctcatctgttaaccatggatataaaacacttatcttca 2625
chimp54546-135457.revcompl.Pan.troglodytes.clone.rp43-41g5.GenBank:AC099533.36  CTGGTGTTCATTTCATGGTCATCTCATCTGTTAACCATGGATATAAAACACTTATCTTCA 4190
human.FAM30B-LOC105372935-GGT2.NCBI.                                            ctggtgttcatttcatggtcatctcatctgttaaccatggatataaaacatttatcttca 2420
human.FAM230E-LOC105377182-GGT3P.NCBI.                                          ctggtgttcatttcatggtcatctcatctgttaaccatggatataaaacatttatcttca 1495
human.FAM230J-LOC105372942-GGTLC5P.NCBI.rev.compl                               ctggtgttcatttcatggtcatctcatctgttaaccatggatataaaacatttatcttca 1865
human.FAM230A.ncRNA.18487127.18500594-GGTLC3.18516335.18518165.rev.compl.REF    ------------------------------------------------------------ 0

chimp.LOC112206744-LOC107973052-GGT2.revcompl                                   atgatcttcctgtacatgtaaaaacacacctgtctgcatggcagcagttggacctcacag 2685
chimp54546-135457.revcompl.Pan.troglodytes.clone.rp43-41g5.GenBank:AC099533.36  ATGATCTTCCTGTACATATAAAAACACACCTGTCTACATGGCAGCAGTTGGACCTCACAA 4250
human.FAM30B-LOC105372935-GGT2.NCBI.                                            at---------------------------------------------------------- 2422
human.FAM230E-LOC105377182-GGT3P.NCBI.                                          at---------------------------------------------------------- 1497
human.FAM230J-LOC105372942-GGTLC5P.NCBI.rev.compl                               at---------------------------------------------------------- 1867
human.FAM230A.ncRNA.18487127.18500594-GGTLC3.18516335.18518165.rev.compl.REF    ------------------------------------------------------------ 0

chimp.LOC112206744-LOC107973052-GGT2.revcompl                                   tgtggattgtgccttcaccctggaatgtctatgatgccctatcgaccatggtgatgggat 2745
chimp54546-135457.revcompl.Pan.troglodytes.clone.rp43-41g5.GenBank:AC099533.36  TGTGGATTGTGCCTTCACCCTGGAATGTCTATGATGCCCTATCGACCATGGTGATGGGAT 4310
human.FAM30B-LOC105372935-GGT2.NCBI.                                            ------------------------------------------------------------ 2422
human.FAM230E-LOC105377182-GGT3P.NCBI.                                          ------------------------------------------------------------ 1497
human.FAM230J-LOC105372942-GGTLC5P.NCBI.rev.compl                               ------------------------------------------------------------ 1867
human.FAM230A.ncRNA.18487127.18500594-GGTLC3.18516335.18518165.rev.compl.REF    ------------------------------------------------------------ 0

chimp.LOC112206744-LOC107973052-GGT2.revcompl                                   tagggatctcttcccctttgctcggccactgtctgtgccagccaggccactgggccattg 2805
chimp54546-135457.revcompl.Pan.troglodytes.clone.rp43-41g5.GenBank:AC099533.36  TAGGGATCTCTTGCCCTTTGCTCTGCCACTGTCTGTGCCAGCCAGGCCACTGGGCCATTG 4370
human.FAM30B-LOC105372935-GGT2.NCBI.                                            ------------------------------------------------------------ 2422
human.FAM230E-LOC105377182-GGT3P.NCBI.                                          ------------------------------------------------------------ 1497
human.FAM230J-LOC105372942-GGTLC5P.NCBI.rev.compl                               ------------------------------------------------------------ 1867
human.FAM230A.ncRNA.18487127.18500594-GGTLC3.18516335.18518165.rev.compl.REF    ------------------------------------------------------------ 0

chimp.LOC112206744-LOC107973052-GGT2.revcompl                                   tggcggatggtgatgccctttgtgtggagcgcactgagtgtgcgctgaggcagacacagt 2865
chimp54546-135457.revcompl.Pan.troglodytes.clone.rp43-41g5.GenBank:AC099533.36  TGGCAGATGGTGATGCCCTTTGTGTGGAGCGCACTGAGTGTGCGCTGAGGCAGACACAGT 4430
human.FAM30B-LOC105372935-GGT2.NCBI.                                            ------------------------------------------------------------ 2422
human.FAM230E-LOC105377182-GGT3P.NCBI.                                          ------------------------------------------------------------ 1497
human.FAM230J-LOC105372942-GGTLC5P.NCBI.rev.compl                               ------------------------------------------------------------ 1867
human.FAM230A.ncRNA.18487127.18500594-GGTLC3.18516335.18518165.rev.compl.REF    ------------------------------------------------------------ 0

chimp.LOC112206744-LOC107973052-GGT2.revcompl                                   acttttgacaactttatctgcttctttcaaaactgggtctgcattatgataaagtggtcg 2925
chimp54546-135457.revcompl.Pan.troglodytes.clone.rp43-41g5.GenBank:AC099533.36  ACTTTTGACAACTTTATCTGCTTCTTTCAAAACTGGGTCTGCATTATGATAAAGTGGTCG 4490
human.FAM30B-LOC105372935-GGT2.NCBI.                                            ------------------------------------------------------------ 2422
human.FAM230E-LOC105377182-GGT3P.NCBI.                                          ------------------------------------------------------------ 1497
human.FAM230J-LOC105372942-GGTLC5P.NCBI.rev.compl                               ------------------------------------------------------------ 1867
human.FAM230A.ncRNA.18487127.18500594-GGTLC3.18516335.18518165.rev.compl.REF    ------------------------------------------------------------ 0

chimp.LOC112206744-LOC107973052-GGT2.revcompl                                   gttttaggccaagcaggtgtatgtcatagagcaatgacttgggagttcagaatcctgggc 2985
chimp54546-135457.revcompl.Pan.troglodytes.clone.rp43-41g5.GenBank:AC099533.36  GTTTTAGGCCAAGCAGGTGTATGTCATAGAACAATGACTTGGGAGTTCAGAATCCTGGGC 4550
human.FAM30B-LOC105372935-GGT2.NCBI.                                            ------------------------------------------------------------ 2422
human.FAM230E-LOC105377182-GGT3P.NCBI.                                          ------------------------------------------------------------ 1497
human.FAM230J-LOC105372942-GGTLC5P.NCBI.rev.compl                               ------------------------------------------------------------ 1867
human.FAM230A.ncRNA.18487127.18500594-GGTLC3.18516335.18518165.rev.compl.REF    ------------------------------------------------------------ 0

chimp.LOC112206744-LOC107973052-GGT2.revcompl                                   gtgaataagcctctgagccttattaagctgtgaaatctaaagcaagacatttcactgata 3045
chimp54546-135457.revcompl.Pan.troglodytes.clone.rp43-41g5.GenBank:AC099533.36  GTGAATAAGCCTCTGAGCCTTATTAAGCTGTGAAATCTAAAGCAAGACATTTCACTGATA 4610
human.FAM30B-LOC105372935-GGT2.NCBI.                                            ------------------------------------------------------------ 2422
human.FAM230E-LOC105377182-GGT3P.NCBI.                                          ------------------------------------------------------------ 1497
human.FAM230J-LOC105372942-GGTLC5P.NCBI.rev.compl                               ------------------------------------------------------------ 1867
human.FAM230A.ncRNA.18487127.18500594-GGTLC3.18516335.18518165.rev.compl.REF    ------------------------------------------------------------ 0

chimp.LOC112206744-LOC107973052-GGT2.revcompl                                   attaagcctcatgagcacttatactgtacactttacctacatcaaatcattcggccatca 3105
chimp54546-135457.revcompl.Pan.troglodytes.clone.rp43-41g5.GenBank:AC099533.36  ATTAAGCCTCATGAGCACTTATACTGTACACTTTACCTACATGAAATCATTCGGCCATCG 4670
human.FAM30B-LOC105372935-GGT2.NCBI.                                            ------------------------------------------------------------ 2422
human.FAM230E-LOC105377182-GGT3P.NCBI.                                          ------------------------------------------------------------ 1497
human.FAM230J-LOC105372942-GGTLC5P.NCBI.rev.compl                               ------------------------------------------------------------ 1867
human.FAM230A.ncRNA.18487127.18500594-GGTLC3.18516335.18518165.rev.compl.REF    ------------------------------------------------------------ 0

chimp.LOC112206744-LOC107973052-GGT2.revcompl                                   ccacaaacaaatggcatgattgatttactatcacctttcgctgggtagaaaaagacagag 3165
chimp54546-135457.revcompl.Pan.troglodytes.clone.rp43-41g5.GenBank:AC099533.36  CCACAGACAAATGGCATGATTGATTTACTATCACCTTTTGCTGGGCAGAAAAAGACAGAG 4730
human.FAM30B-LOC105372935-GGT2.NCBI.                                            ------------------------------------------------------------ 2422
human.FAM230E-LOC105377182-GGT3P.NCBI.                                          ------------------------------------------------------------ 1497
human.FAM230J-LOC105372942-GGTLC5P.NCBI.rev.compl                               ------------------------------------------------------------ 1867
human.FAM230A.ncRNA.18487127.18500594-GGTLC3.18516335.18518165.rev.compl.REF    ------------------------------------------------------------ 0

chimp.LOC112206744-LOC107973052-GGT2.revcompl                                   atgaatgtgcccaagcttatgagcccctcagcaaaagagccaagatgggaacccaagcat 3225
chimp54546-135457.revcompl.Pan.troglodytes.clone.rp43-41g5.GenBank:AC099533.36  ATGAATGTGCCCAAGCTTATGAGCCCCTCAGCAAAAGAGCCAAGATGGGAACCCAAGCAT 4790
human.FAM30B-LOC105372935-GGT2.NCBI.                                            ------------------------------------------------------------ 2422
human.FAM230E-LOC105377182-GGT3P.NCBI.                                          ------------------------------------------------------------ 1497
human.FAM230J-LOC105372942-GGTLC5P.NCBI.rev.compl                               ------------------------------------------------------------ 1867
human.FAM230A.ncRNA.18487127.18500594-GGTLC3.18516335.18518165.rev.compl.REF    ------------------------------------------------------------ 0

chimp.LOC112206744-LOC107973052-GGT2.revcompl                                   acagccccaatgctgaggctctgaactactgacctgccctcagcactcagccttgggatc 3285
chimp54546-135457.revcompl.Pan.troglodytes.clone.rp43-41g5.GenBank:AC099533.36  ACAGCCCCAATGCTGAGGCTCTGAACTACTGACCTGCCCTCAGCACTCAGCCTTGGGATC 4850
human.FAM30B-LOC105372935-GGT2.NCBI.                                            ------------------------------------------------------------ 2422
human.FAM230E-LOC105377182-GGT3P.NCBI.                                          ------------------------------------------------------------ 1497
human.FAM230J-LOC105372942-GGTLC5P.NCBI.rev.compl                               ------------------------------------------------------------ 1867
human.FAM230A.ncRNA.18487127.18500594-GGTLC3.18516335.18518165.rev.compl.REF    ------------------------------------------------------------ 0

chimp.LOC112206744-LOC107973052-GGT2.revcompl                                   atgagtcactgtgcaagggagttccaacatctgcatgtatgtctggaatgatctgagcct 3345
chimp54546-135457.revcompl.Pan.troglodytes.clone.rp43-41g5.GenBank:AC099533.36  ATGAGTCACTGTGCAAGGGAGTTCCAACATCTGCATGTATGTCTGGAATGATCTGAGCCT 4910
human.FAM30B-LOC105372935-GGT2.NCBI.                                            ------------------------------------------------------------ 2422
human.FAM230E-LOC105377182-GGT3P.NCBI.                                          ------------------------------------------------------------ 1497
```

```
human.FAM230J-LOC105372942-GGTLC5P.NCBI.rev.compl                            ------------------------------------------------------------ 1867
human.FAM230A.ncRNA.18487127.18500594-GGTLC3.18516335.18518165.rev.compl.REF ------------------------------------------------------------ 0

chimp.LOC112206744-LOC107973052-GGT2.revcompl                                gtagagttcctacacactggccacattatagggtggtgtctgtggtcacacagctcaggg 3405
chimp54546-135457.revcompl.Pan.troglodytes.clone.rp43-41g5.GenBank:AC099533.36 GTAGAGTTCCTACACACTGGCCACATTATAGGGTGGTGTCTGTGGTCACACAGCTCAGGG 4970
human.FAM30B-LOC105372935-GGT2.NCBI.                                         ------------------------------------------------------------ 2422
human.FAM230E-LOC105377182-GGT3P.NCBI.                                       ------------------------------------------------------------ 1497
human.FAM230J-LOC105372942-GGTLC5P.NCBI.rev.compl                            ------------------------------------------------------------ 1867
human.FAM230A.ncRNA.18487127.18500594-GGTLC3.18516335.18518165.rev.compl.REF ------------------------------------------------------------ 0

chimp.LOC112206744-LOC107973052-GGT2.revcompl                                caggtattagtacatgaatagcttagatgtgtcatagtctttatgtgaaaggcacataaa 3465
chimp54546-135457.revcompl.Pan.troglodytes.clone.rp43-41g5.GenBank:AC099533.36 CAGGTATTAGTACATGAATAGCTTAGCTGTGTCATAGTCTTTATGTGAAAGGCACATAAA 5030
human.FAM30B-LOC105372935-GGT2.NCBI.                                         ------------------------------------------------------------ 2422
human.FAM230E-LOC105377182-GGT3P.NCBI.                                       ------------------------------------------------------------ 1497
human.FAM230J-LOC105372942-GGTLC5P.NCBI.rev.compl                            ------------------------------------------------------------ 1867
human.FAM230A.ncRNA.18487127.18500594-GGTLC3.18516335.18518165.rev.compl.REF ------------------------------------------------------------ 0

chimp.LOC112206744-LOC107973052-GGT2.revcompl                                agcccaaatgcttttgaactggttgcggattttgagttggaggacaacacttgggacaac 3525
chimp54546-135457.revcompl.Pan.troglodytes.clone.rp43-41g5.GenBank:AC099533.36 AGCCCAAATGCTTTTGAACTGGTTGCGGATTTTGAGTTGGAGGACAACACTTGGGACAAC 5090
human.FAM30B-LOC105372935-GGT2.NCBI.                                         ------------------------------------------------------------ 2422
human.FAM230E-LOC105377182-GGT3P.NCBI.                                       ------------------------------------------------------------ 1497
human.FAM230J-LOC105372942-GGTLC5P.NCBI.rev.compl                            ------------------------------------------------------------ 1867
human.FAM230A.ncRNA.18487127.18500594-GGTLC3.18516335.18518165.rev.compl.REF ------------------------------------------------------------ 0

chimp.LOC112206744-LOC107973052-GGT2.revcompl                                cggtcacattccttcagtgcgcggccactcctcagcacatatcccaaagccggtaaataa 3585
chimp54546-135457.revcompl.Pan.troglodytes.clone.rp43-41g5.GenBank:AC099533.36 CGGTCACATTCCTTCAGTGCGCGGCCACTCCTCAGCACATATCCCAAAGCCGGTAAATAA 5150
human.FAM30B-LOC105372935-GGT2.NCBI.                                         ------------------------------------------------------------ 2422
human.FAM230E-LOC105377182-GGT3P.NCBI.                                       ------------------------------------------------------------ 1497
human.FAM230J-LOC105372942-GGTLC5P.NCBI.rev.compl                            ------------------------------------------------------------ 1867
human.FAM230A.ncRNA.18487127.18500594-GGTLC3.18516335.18518165.rev.compl.REF ------------------------------------------------------------ 0

chimp.LOC112206744-LOC107973052-GGT2.revcompl                                tgggaagaccctgtactgatgtttgatggtgttatgaacttcaaatgacccctatagaaa 3645
chimp54546-135457.revcompl.Pan.troglodytes.clone.rp43-41g5.GenBank:AC099533.36 TGGGAAGACCCTGTACTGATGTTTGATGGTGTTATGAACTTCAAATGACCCCTATAGAAA 5210
human.FAM30B-LOC105372935-GGT2.NCBI.                                         ------------------------------------------------------------ 2422
human.FAM230E-LOC105377182-GGT3P.NCBI.                                       ------------------------------------------------------------ 1497
human.FAM230J-LOC105372942-GGTLC5P.NCBI.rev.compl                            ------------------------------------------------------------ 1867
human.FAM230A.ncRNA.18487127.18500594-GGTLC3.18516335.18518165.rev.compl.REF ------------------------------------------------------------ 0

chimp.LOC112206744-LOC107973052-GGT2.revcompl                                tgttaaaaaaaaaaaaaaaaaaaaagtctgtgagctacacaacctggggacaagacaagt 3705
chimp54546-135457.revcompl.Pan.troglodytes.clone.rp43-41g5.GenBank:AC099533.36 TGTT--------AAAAAAAAAAAAAGTCTGTGAGCTACACAACCTGGGGACAAGACAAGT 5262
human.FAM30B-LOC105372935-GGT2.NCBI.                                         ------------------------------------------------------------ 2422
human.FAM230E-LOC105377182-GGT3P.NCBI.                                       ------------------------------------------------------------ 1497
human.FAM230J-LOC105372942-GGTLC5P.NCBI.rev.compl                            ------------------------------------------------------------ 1867
human.FAM230A.ncRNA.18487127.18500594-GGTLC3.18516335.18518165.rev.compl.REF ------------------------------------------------------------ 0

chimp.LOC112206744-LOC107973052-GGT2.revcompl                                aagaacacaaaataccagtaaataatgagcctcatgtgtcaccctgcttgcaactccctc 3765
chimp54546-135457.revcompl.Pan.troglodytes.clone.rp43-41g5.GenBank:AC099533.36 AAGAACACAAAATACCACTAAATAATGAGCCTCATGTGTCACCCTGCTTGCAACTCCCTC 5322
human.FAM30B-LOC105372935-GGT2.NCBI.                                         ------------------------------------------------------------ 2422
human.FAM230E-LOC105377182-GGT3P.NCBI.                                       ------------------------------------------------------------ 1497
human.FAM230J-LOC105372942-GGTLC5P.NCBI.rev.compl                            ------------------------------------------------------------ 1867
human.FAM230A.ncRNA.18487127.18500594-GGTLC3.18516335.18518165.rev.compl.REF ------------------------------------------------------------ 0

chimp.LOC112206744-LOC107973052-GGT2.revcompl                                ggcagctggtttgaagagcctcagtgctcaccaggggccatctccccgctctgccctctc 3825
chimp54546-135457.revcompl.Pan.troglodytes.clone.rp43-41g5.GenBank:AC099533.36 GGCAGCTGGTTTGAAGAGCCTCAGTGCTCACCAGGGGCCATCTCCCCGCTCTGCCCTCTC 5382
human.FAM30B-LOC105372935-GGT2.NCBI.                                         ------------------------------------------------------------ 2422
human.FAM230E-LOC105377182-GGT3P.NCBI.                                       ------------------------------------------------------------ 1497
human.FAM230J-LOC105372942-GGTLC5P.NCBI.rev.compl                            ------------------------------------------------------------ 1867
human.FAM230A.ncRNA.18487127.18500594-GGTLC3.18516335.18518165.rev.compl.REF ------------------------------------------------------------ 0

chimp.LOC112206744-LOC107973052-GGT2.revcompl                                tcgctgtgggttgcctgggctgaacataggcagcccctcacatgatgactgcaaacacag 3885
chimp54546-135457.revcompl.Pan.troglodytes.clone.rp43-41g5.GenBank:AC099533.36 TCGCTGTGGGTTGCCTGGGCTGAACATAGGCAGCCCCTCACATGATGACTGCAAACACAG 5442
human.FAM30B-LOC105372935-GGT2.NCBI.                                         ------------------------------------------------------------ 2422
human.FAM230E-LOC105377182-GGT3P.NCBI.                                       ------------------------------------------------------------ 1497
human.FAM230J-LOC105372942-GGTLC5P.NCBI.rev.compl                            ------------------------------------------------------------ 1867
human.FAM230A.ncRNA.18487127.18500594-GGTLC3.18516335.18518165.rev.compl.REF ------------------------------------------------------------ 0

chimp.LOC112206744-LOC107973052-GGT2.revcompl                                ggcaaaggaagactgaacggaaaatgtatggttatagtggttattttaaaatataagtat 3945
chimp54546-135457.revcompl.Pan.troglodytes.clone.rp43-41g5.GenBank:AC099533.36 GGCAAAGGAAGACTGAACGGAAAATGTATGGTTATAGTGGTTATTTTAAAATATAAGTAT 5502
human.FAM30B-LOC105372935-GGT2.NCBI.                                         ------------------------------------------------------------ 2422
human.FAM230E-LOC105377182-GGT3P.NCBI.                                       ------------------------------------------------------------ 1497
human.FAM230J-LOC105372942-GGTLC5P.NCBI.rev.compl                            ------------------------------------------------------------ 1867
human.FAM230A.ncRNA.18487127.18500594-GGTLC3.18516335.18518165.rev.compl.REF ------------------------------------------------------------ 0

chimp.LOC112206744-LOC107973052-GGT2.revcompl                                ttcagtattttctaatctttaataattgattatttttaggggaggttttttttttttttt 4005
chimp54546-135457.revcompl.Pan.troglodytes.clone.rp43-41g5.GenBank:AC099533.36 TTCAGTATTTTCTAATCTTTAATAATTGATTATTTTTAGGGGAGG--TTTTTTTTTTTTT 5560
human.FAM30B-LOC105372935-GGT2.NCBI.                                         ------------------------------------------------------------ 2422
human.FAM230E-LOC105377182-GGT3P.NCBI.                                       ------------------------------------------------------------ 1497
human.FAM230J-LOC105372942-GGTLC5P.NCBI.rev.compl                            ------------------------------------------------------------ 1867
human.FAM230A.ncRNA.18487127.18500594-GGTLC3.18516335.18518165.rev.compl.REF ------------------------------------------------------------ 0

chimp.LOC112206744-LOC107973052-GGT2.revcompl                                tttgagacagagtcttgcgctgttgcccaggctggagtgcagtggcatgatctcagctca 4065
chimp54546-135457.revcompl.Pan.troglodytes.clone.rp43-41g5.GenBank:AC099533.36 TTTGAGACAGAGTCTTGCACTGTTGCCCAGGCTGGAGTGCAGTGGCATGATCTCAGCTCA 5620
human.FAM30B-LOC105372935-GGT2.NCBI.                                         ------------------------------------------------------------ 2422
human.FAM230E-LOC105377182-GGT3P.NCBI.                                       ------------------------------------------------------------ 1497
human.FAM230J-LOC105372942-GGTLC5P.NCBI.rev.compl                            ------------------------------------------------------------ 1867
human.FAM230A.ncRNA.18487127.18500594-GGTLC3.18516335.18518165.rev.compl.REF ------------------------------------------------------------ 0

chimp.LOC112206744-LOC107973052-GGT2.revcompl                                ctgcaacctctacctcctgtgttcaagcgattctcctgcctcagcctcccgagtagctgg 4125
chimp54546-135457.revcompl.Pan.troglodytes.clone.rp43-41g5.GenBank:AC099533.36 CTGCAACCTCTGCCTCCTGTGTTCAAGCGATTCTCCTGCCTCAGCCTCCCGAGTAGCTGG 5680
human.FAM30B-LOC105372935-GGT2.NCBI.                                         ------------------------------------------------------------ 2422
human.FAM230E-LOC105377182-GGT3P.NCBI.                                       ------------------------------------------------------------ 1497
human.FAM230J-LOC105372942-GGTLC5P.NCBI.rev.compl                            ------------------------------------------------------------ 1867
human.FAM230A.ncRNA.18487127.18500594-GGTLC3.18516335.18518165.rev.compl.REF ------------------------------------------------------------ 0

chimp.LOC112206744-LOC107973052-GGT2.revcompl                                gactacaggcacgtgccaccacgcctggcaaagtttttgtgtttttagtagagacggggt 4185
chimp54546-135457.revcompl.Pan.troglodytes.clone.rp43-41g5.GenBank:AC099533.36 GACTACAGGCACGTGCCACCACGCCTGGCAAAGTTTTTGTGTTTTTAGTAGAGACGGGGT 5740
human.FAM30B-LOC105372935-GGT2.NCBI.                                         ------------------------------------------------------------ 2422
human.FAM230E-LOC105377182-GGT3P.NCBI.                                       ------------------------------------------------------------ 1497
human.FAM230J-LOC105372942-GGTLC5P.NCBI.rev.compl                            ------------------------------------------------------------ 1867
human.FAM230A.ncRNA.18487127.18500594-GGTLC3.18516335.18518165.rev.compl.REF ------------------------------------------------------------ 0

chimp.LOC112206744-LOC107973052-GGT2.revcompl                                ttcactctgttagccaggatggtctcaatcttctgacctcatgatccacccgcctcggcc 4245
chimp54546-135457.revcompl.Pan.troglodytes.clone.rp43-41g5.GenBank:AC099533.36 TTCACTCTGTTAGCCAGGATGGTCTCAATCTTCTGACCTCATGATCCACCCGCCTCGGCC 5800
human.FAM30B-LOC105372935-GGT2.NCBI.                                         ------------------------------------------------------------ 2422
human.FAM230E-LOC105377182-GGT3P.NCBI.                                       ------------------------------------------------------------ 1497
human.FAM230J-LOC105372942-GGTLC5P.NCBI.rev.compl                            ------------------------------------------------------------ 1867
human.FAM230A.ncRNA.18487127.18500594-GGTLC3.18516335.18518165.rev.compl.REF ------------------------------------------------------------ 0

chimp.LOC112206744-LOC107973052-GGT2.revcompl                                tcccaaagtgctgggattacaggtgtgagccaccacacctggcctgatttttttattttt 4305
chimp54546-135457.revcompl.Pan.troglodytes.clone.rp43-41g5.GenBank:AC099533.36 TCCCAAAGTGCTGGGATTACAGGTGTGAGCCACCACACCTGGCCTGATTTTTTACTTTTT 5860
human.FAM30B-LOC105372935-GGT2.NCBI.                                         ------------------------------------------------------------ 2422
human.FAM230E-LOC105377182-GGT3P.NCBI.                                       ------------------------------------------------------------ 1497
human.FAM230J-LOC105372942-GGTLC5P.NCBI.rev.compl                            ------------------------------------------------------------ 1867
human.FAM230A.ncRNA.18487127.18500594-GGTLC3.18516335.18518165.rev.compl.REF ------------------------------------------------------------ 0

chimp.LOC112206744-LOC107973052-GGT2.revcompl                                tatttttgagatgaagtctctgtcacccaggctggagtgcagtggtgcaatctcagctc 4365
chimp54546-135457.revcompl.Pan.troglodytes.clone.rp43-41g5.GenBank:AC099533.36 TATTTTTGAGATGAAGTCTCTGTCACCCAGGCTGGAGTGCAGTGGTGCAATCTCAGCTC 5920
human.FAM30B-LOC105372935-GGT2.NCBI.                                         ------------------------------------------------------------ 2422
human.FAM230E-LOC105377182-GGT3P.NCBI.                                       ------------------------------------------------------------ 1497
human.FAM230J-LOC105372942-GGTLC5P.NCBI.rev.compl                            ------------------------------------------------------------ 1867
human.FAM230A.ncRNA.18487127.18500594-GGTLC3.18516335.18518165.rev.compl.REF ------------------------------------------------------------ 0

chimp.LOC112206744-LOC107973052-GGT2.revcompl                                actgcaatctctgcctccaggttcaagggattgtcctgcctcagcctcctgagtagctgg 4425
chimp54546-135457.revcompl.Pan.troglodytes.clone.rp43-41g5.GenBank:AC099533.36 ACTGCAATCTCTGCCTCCAGGTTCAAGGATTCTCCTGCCTCAGCCTCCTGAGTAGCTGG 5980
human.FAM30B-LOC105372935-GGT2.NCBI.                                         ------------------------------------------------------------ 2422
human.FAM230E-LOC105377182-GGT3P.NCBI.                                       ------------------------------------------------------------ 1497
human.FAM230J-LOC105372942-GGTLC5P.NCBI.rev.compl                            ------------------------------------------------------------ 1867
```

```
human.FAM230A.ncRNA.18487127.18500594-GGTLC3.18516335.18518165.rev.compl.REF        ------------------------------------------------------------ 0

chimp.LOC112206744-LOC107973052-GGT2.revcompl                                        ----gttttttttttttttagttgagatggggttttaccatgttggtcaggctggtctcg 4481
chimp54546-135457.revcompl.Pan.troglodytes.clone.rp43-41g5.GenBank:AC099533.36       TTTTTTTTTTTTTTTTTTTAGTTGAGATGGGGTTTTACCATGTTGGTCAGGCTGGTCTCG 6040
human.FAM30B-LOC105372935-GGT2.NCBI.                                                 -------------------------gatggggttttaccatgttggtcaggctggtctcg 2457
human.FAM230E-LOC105377182-GGT3P.NCBI.                                               -------------------------gatggggttttaccatgttggtcaggctggtctcg 1532
human.FAM230J-LOC105372942-GGTLC5P.NCBI.rev.compl                                    -------------------------gatggggttttaccatgttggtcaggctggtctcg 1902
human.FAM230A.ncRNA.18487127.18500594-GGTLC3.18516335.18518165.rev.compl.REF        ------------------------------------------------------------ 0

chimp.LOC112206744-LOC107973052-GGT2.revcompl                                        aactcctgacctcaaatgatccgcccacctccaccttccaaactgctgggattacaggtg 4541
chimp54546-135457.revcompl.Pan.troglodytes.clone.rp43-41g5.GenBank:AC099533.36       AACTCCTGACCTCAAATGATCCGCCCACCTCCACCTTCCAAACTGCTGGGATTACAGGTG 6100
human.FAM30B-LOC105372935-GGT2.NCBI.                                                 aactcctgacctcaaatgatccacccacctccaccttccaaactgctgggattacaggtg 2517
human.FAM230E-LOC105377182-GGT3P.NCBI.                                               aactcctgacctcaaatgatccacccacctccaccttccaaactgctgggattacaggtg 1592
human.FAM230J-LOC105372942-GGTLC5P.NCBI.rev.compl                                    aactcctgacctcaaatgatccacccacctccaccttccaaactgctgggattacaggtg 1962
human.FAM230A.ncRNA.18487127.18500594-GGTLC3.18516335.18518165.rev.compl.REF        ------------------------------------------------------------ 0

chimp.LOC112206744-LOC107973052-GGT2.revcompl                                        tgagccactatgcctggctgattattttcataaccaagaaaagaaataaatataattaat 4601
chimp54546-135457.revcompl.Pan.troglodytes.clone.rp43-41g5.GenBank:AC099533.36       TGAGCCACTATGCCTGGCTGATTATTTTCATAACCAAGAAAAGAAATAAATATAATTAAT 6160
human.FAM30B-LOC105372935-GGT2.NCBI.                                                 tgagccactatgcctgactgattattttcataaccaagaaaagaaataaatacaattaat 2577
human.FAM230E-LOC105377182-GGT3P.NCBI.                                               tgagccactatgcctgactgattattttcataaccaagaaaagaaataaatacaattaat 1652
human.FAM230J-LOC105372942-GGTLC5P.NCBI.rev.compl                                    tgagccactatgcctgactgattattttcataaccaagaaaagaaataaatacaattaat 2022
human.FAM230A.ncRNA.18487127.18500594-GGTLC3.18516335.18518165.rev.compl.REF        ------------------------------------------------------------ 0

chimp.LOC112206744-LOC107973052-GGT2.revcompl                                        gctggtgcatggtattaaatctagtttttaaaaaattcacacataaaccaggcagaaccc 4661
chimp54546-135457.revcompl.Pan.troglodytes.clone.rp43-41g5.GenBank:AC099533.36       GCTGGTGCATGGTATTAAATCTAGTTTTTAAAAAAATCACACATAAACCAGGCAGAACCC 6220
human.FAM30B-LOC105372935-GGT2.NCBI.                                                 gctggtgcatggtattaaatctagtttttaaaaaattcacacataaacaaggcagaaccc 2637
human.FAM230E-LOC105377182-GGT3P.NCBI.                                               gctggtgcatggtattaaatctagtttttaaaaaattcacacataaacaaggcagaaccc 1712
human.FAM230J-LOC105372942-GGTLC5P.NCBI.rev.compl                                    gctggtgcatggtattaaatctagtttttaaaaaattcacacataaacagggcagaaccc 2082
human.FAM230A.ncRNA.18487127.18500594-GGTLC3.18516335.18518165.rev.compl.REF        ------------------------------------------------------------ 0

chimp.LOC112206744-LOC107973052-GGT2.revcompl                                        tataccctccatgataaatgcagtagcagtgtatgtgggtctgtggaggttgaaagggac 4721
chimp54546-135457.revcompl.Pan.troglodytes.clone.rp43-41g5.GenBank:AC099533.36       TATACCCTCCATGATAAATGCAGTAGCAGTGTATGTGGGTCTGTGGAGGTTGAAAGGGAC 6280
human.FAM30B-LOC105372935-GGT2.NCBI.                                                 tataccctccatgataaatgcagtagcagtgtatgtgggtctgtggaggttgaaagggac 2697
human.FAM230E-LOC105377182-GGT3P.NCBI.                                               tataccctccatgataaatgcagtagcagtgtatgtgggtctgtggaggttgaaagggac 1772
human.FAM230J-LOC105372942-GGTLC5P.NCBI.rev.compl                                    tataccctccatgataaatgcagtagcagtgtatgtgggtctgtggaggttgaaagggac 2142
human.FAM230A.ncRNA.18487127.18500594-GGTLC3.18516335.18518165.rev.compl.REF        ------------------------------------------------------------ 0

chimp.LOC112206744-LOC107973052-GGT2.revcompl                                        ttggtagatgtcaagaaggtagtggcagtcctgctgggcttttaaagggtctgaagaagt 4781
chimp54546-135457.revcompl.Pan.troglodytes.clone.rp43-41g5.GenBank:AC099533.36       TTGGTAGATGTCAAGAAGGTAGTGGCAGTCCTGCTGGGCTTTTAAAGGGTCTGAAGAAGT 6340
human.FAM30B-LOC105372935-GGT2.NCBI.                                                 ttggtagatgtcaagaagctagtggcagtcctgctgggcttttaaagggtctgaagaagt 2757
human.FAM230E-LOC105377182-GGT3P.NCBI.                                               ttggtagatgtcaagaaggtagtggcagtcctgctgggcttttaaagggtctgaagaagt 1832
human.FAM230J-LOC105372942-GGTLC5P.NCBI.rev.compl                                    ttggtcgatgtcaagaaggtagtggcagtcctgctgggcttttaaagggtctgaagaagt 2202
human.FAM230A.ncRNA.18487127.18500594-GGTLC3.18516335.18518165.rev.compl.REF        ------------------------------------------------------------ 0

chimp.LOC112206744-LOC107973052-GGT2.revcompl                                        gacaggatgctgtggttgaatcctagcatgtattttagcatttgttcatttggagtttga 4841
chimp54546-135457.revcompl.Pan.troglodytes.clone.rp43-41g5.GenBank:AC099533.36       GACAGGATGCTGTGGTTGAATCCTAGCATGTATTTTAGCATTTGTTCATTTGGAGTTTGA 6400
human.FAM30B-LOC105372935-GGT2.NCBI.                                                 gacaggatgctgtggttgaatcctagcatgtattttagcatttgttcatttggag-ttga 2816
human.FAM230E-LOC105377182-GGT3P.NCBI.                                               gacaggatgctgtggttgaatcctagcatgtattttagcatttgttcatttggag-ttga 1891
human.FAM230J-LOC105372942-GGTLC5P.NCBI.rev.compl                                    gacaggatgctgtggttgaatcgtagcatgtgttttagcatttgttcatttggag-ttga 2261
human.FAM230A.ncRNA.18487127.18500594-GGTLC3.18516335.18518165.rev.compl.REF        ------------------------------------------------------------ 0

chimp.LOC112206744-LOC107973052-GGT2.revcompl                                        ttatttcacgttgctttcatttgccattacctggaaagccaagggctctactctcatttc 4901
chimp54546-135457.revcompl.Pan.troglodytes.clone.rp43-41g5.GenBank:AC099533.36       TTATTTCACGTTGCTTTCATTTGCCATTACCTGGAAAGCCAAGGGCTGTACTCTTATTTC 6460
human.FAM30B-LOC105372935-GGT2.NCBI.                                                 ttatttcacgttgctttcatttgccattacctggaaagccaagggctctactctcatttc 2876
human.FAM230E-LOC105377182-GGT3P.NCBI.                                               ttatttcacgttgctttcatttgccattacctggaaagccaagggctctactctcatttc 1951
human.FAM230J-LOC105372942-GGTLC5P.NCBI.rev.compl                                    ttatttcacgttgctttcatttgccattacctggaaagccaagggctctactctcatttc 2321
human.FAM230A.ncRNA.18487127.18500594-GGTLC3.18516335.18518165.rev.compl.REF        ------------------------------------------------------------ 0

chimp.LOC112206744-LOC107973052-GGT2.revcompl                                        cttgctgctctttctttgccttccttggtccatgaagaagatggtccaggagaagctcat 4961
chimp54546-135457.revcompl.Pan.troglodytes.clone.rp43-41g5.GenBank:AC099533.36       CTTGCTACTCTTTCTTTGCCTTCCTTGGTCCATGAAGAAGATGGTCCAGGAGAAGCTCAT 6520
human.FAM30B-LOC105372935-GGT2.NCBI.                                                 cttgctgctctttctttgccttccttggtccgtgaagaagatggtccaggagaagctcat 2936
human.FAM230E-LOC105377182-GGT3P.NCBI.                                               cttgctgctctttctttgccttccttggtccgtgaagaagatggtccaggagaagctcat 2011
human.FAM230J-LOC105372942-GGTLC5P.NCBI.rev.compl                                    cttgctgctctttctttgccttccttggtccgtgaagaagatggtccaggagaagctcat 2381
human.FAM230A.ncRNA.18487127.18500594-GGTLC3.18516335.18518165.rev.compl.REF        ------------------------------------------------------------ 0

chimp.LOC112206744-LOC107973052-GGT2.revcompl                                        tccatgcttgttaaccaggcacacccctaagttccagtccctgagtcattcatgagtagc 5021
chimp54546-135457.revcompl.Pan.troglodytes.clone.rp43-41g5.GenBank:AC099533.36       TCCATGCTTGTTAACCAGGCACACCCCTAAGTTCCAGTCCCTGAGTCATTCATGAGTAGC 6580
human.FAM30B-LOC105372935-GGT2.NCBI.                                                 tccatgcttgttaaccaggcacgcccctaagttccagtccctgagtcattcatgagtagc 2996
human.FAM230E-LOC105377182-GGT3P.NCBI.                                               tccatgcttgttaaccaggcacgcccctaagttccagtccctgagtcattcatgagtagc 2071
human.FAM230J-LOC105372942-GGTLC5P.NCBI.rev.compl                                    tccatgcttgttaaccaggcacgcccctaagttccagtccctgagtcattcatgagtagc 2441
human.FAM230A.ncRNA.18487127.18500594-GGTLC3.18516335.18518165.rev.compl.REF        ------------------------------------------------------------ 0

chimp.LOC112206744-LOC107973052-GGT2.revcompl                                        actgccaatgaactgacagccatgctgtgtccctccacatcccctaggtgactcgaagaa 5081
chimp54546-135457.revcompl.Pan.troglodytes.clone.rp43-41g5.GenBank:AC099533.36       ACTGCCAATGAACTGACAGCCGTGCTGTGTCCCTCCACATCCCCTAGGTGACTCGAAGAA 6640
human.FAM30B-LOC105372935-GGT2.NCBI.                                                 actgccaatgaactgacagccatgctgtgtccctccacatcccctaggtgactcgaagaa 3056
human.FAM230E-LOC105377182-GGT3P.NCBI.                                               actgccaatgaactgacagccatgctgtgtccctccacatcccctaggtgactcgaagaa 2131
human.FAM230J-LOC105372942-GGTLC5P.NCBI.rev.compl                                    actgccaatgaactgacagccatgctgtgtccctccacatcccctaggtgactcgaagaa 2501
human.FAM230A.ncRNA.18487127.18500594-GGTLC3.18516335.18518165.rev.compl.REF        ------------------------------------------------------------ 0

chimp.LOC112206744-LOC107973052-GGT2.revcompl                                        gccttccaaaaagcatgtgaaaagggagccctcctctactaccaaggtaaagtagcctgt 5141
chimp54546-135457.revcompl.Pan.troglodytes.clone.rp43-41g5.GenBank:AC099533.36       GCCTTCCAAAAAGCATGTGAAAAGGGAGCCCTCCTCTACTACCAAGGTAAAGTAGCCTGT 6700
human.FAM30B-LOC105372935-GGT2.NCBI.                                                 gccttccaaaaagcatgtgaaaaggaagccctactctactaccaaggtaaagtagcctgt 3116
human.FAM230E-LOC105377182-GGT3P.NCBI.                                               gccttccaaaaagcgtgtgaaaaggaagccctactctactaccaaggtaaagtagcctgt 2191
human.FAM230J-LOC105372942-GGTLC5P.NCBI.rev.compl                                    gccttccaaaaagcgtgtgaaaaggaagccctactctactaccaaggtaaagtagcctgt 2561
human.FAM230A.ncRNA.18487127.18500594-GGTLC3.18516335.18518165.rev.compl.REF        ------------------------------------------------------------ 0

chimp.LOC112206744-LOC107973052-GGT2.revcompl                                        ctttgcctaagatgtaaatgttgttttcttggatcctttatttttcagttgatatctgct 5201
chimp54546-135457.revcompl.Pan.troglodytes.clone.rp43-41g5.GenBank:AC099533.36       CTTTGCCTAAGATGTAAATGTTGTTTTCTTGGATCCTTTATTTTTCAGTTGATATCAGCT 6760
human.FAM30B-LOC105372935-GGT2.NCBI.                                                 ctttgcctaagatgtaaatgttgttttcttggatcctttatttttcagttgatatcagct 3176
human.FAM230E-LOC105377182-GGT3P.NCBI.                                               ctttgcctaagatgtaaatgttgttttcttggatcctttatttttcagttgatatcagct 2251
human.FAM230J-LOC105372942-GGTLC5P.NCBI.rev.compl                                    ctttgcctaagatgtaaatgttgttttcttggatcctttatttttcagttgatatcagct 2621
human.FAM230A.ncRNA.18487127.18500594-GGTLC3.18516335.18518165.rev.compl.REF        ------------------------------------------------------------ 0

chimp.LOC112206744-LOC107973052-GGT2.revcompl                                        atgggaaaattatccactacattatagatgttagataatatttccttggggatggaggag 5261
chimp54546-135457.revcompl.Pan.troglodytes.clone.rp43-41g5.GenBank:AC099533.36       ATGGGAAAATTATCCACTACATTATAGATGTTAGATAATATTTCCTTGGGAATGGAGGAG 6820
human.FAM30B-LOC105372935-GGT2.NCBI.                                                 atgggaaaattctccactacattataggtgttagataatatttccttggggatggaggag 3236
human.FAM230E-LOC105377182-GGT3P.NCBI.                                               atgggaaaattctccactacattataggtgttagataatatttccttggggatggaggag 2311
human.FAM230J-LOC105372942-GGTLC5P.NCBI.rev.compl                                    atgggaaaattctccactacattataggtgttagataatatttccttggggatggaggag 2681
human.FAM230A.ncRNA.18487127.18500594-GGTLC3.18516335.18518165.rev.compl.REF        ------------------------------------------------------------ 0

chimp.LOC112206744-LOC107973052-GGT2.revcompl                                        gtgtattttaccaactgacacctgattccagaggatgtgcaaaattggcagtgtcagata 5321
chimp54546-135457.revcompl.Pan.troglodytes.clone.rp43-41g5.GenBank:AC099533.36       GTGTATTTTACCAACTGACACCTGATTCCAGAGGACGTGCAAAATTGGCAGTGTCAGATA 6880
human.FAM30B-LOC105372935-GGT2.NCBI.                                                 gtgtattttaccaactgacacctgattccagaggacgtgcaaaattggcagtgtcagata 3296
human.FAM230E-LOC105377182-GGT3P.NCBI.                                               gtgtattttaccaactgacacctgattccagaggacgtgcaaaattggcagtgtcagata 2371
human.FAM230J-LOC105372942-GGTLC5P.NCBI.rev.compl                                    gtgtattttaccaactgacacctgattccagaggacgtgcaaaattggcagtgtcagata 2741
human.FAM230A.ncRNA.18487127.18500594-GGTLC3.18516335.18518165.rev.compl.REF        ------------------------------------------------------------ 0

chimp.LOC112206744-LOC107973052-GGT2.revcompl                                        gtacactgcgtgttaagggatgttttcttcaggaacaagctttccactttagataagaat 5381
chimp54546-135457.revcompl.Pan.troglodytes.clone.rp43-41g5.GenBank:AC099533.36       GTACACTGCGTGTTAAGGGATGTTTTCTTCAGGAACAAGCTTTCCACTTTAGATAAGAAT 6940
human.FAM30B-LOC105372935-GGT2.NCBI.                                                 gtacactgggtgttaagggatgttttcttcaggaacaagctttccactttagataagaat 3356
human.FAM230E-LOC105377182-GGT3P.NCBI.                                               gtacactgggtgttaagggatgttttcttcaggaacaagctttccactttagataagaat 2431
human.FAM230J-LOC105372942-GGTLC5P.NCBI.rev.compl                                    gtacactgggtgttaagggatgttttcttcaggaacaagctttccactttagataagaat 2801
human.FAM230A.ncRNA.18487127.18500594-GGTLC3.18516335.18518165.rev.compl.REF        ------------------------------------------------------------ 0

chimp.LOC112206744-LOC107973052-GGT2.revcompl                                        tctgcaattgctactcaaaaattacctagacagaaacattcttcaagaaaagctcctgtg 5441
chimp54546-135457.revcompl.Pan.troglodytes.clone.rp43-41g5.GenBank:AC099533.36       TCTGCAATTGCTACTCAAAAATTACCTAGACAGAAACATTCTTCAAGAAAAGCTCCTGTG 7000
human.FAM30B-LOC105372935-GGT2.NCBI.                                                 tctgcaattgctactcaaaaattacctagacagaaacattcttcaagaaaagctcctgtg 3416
human.FAM230E-LOC105377182-GGT3P.NCBI.                                               tctgcaattgctactcaaaaattacctagacagaaacattcttcaagaaaagctcctgtg 2491
human.FAM230J-LOC105372942-GGTLC5P.NCBI.rev.compl                                    tctgcaattgctactcaaaaattacctagacagaaacattcttcaagaaaagctcctgtg 2861
human.FAM230A.ncRNA.18487127.18500594-GGTLC3.18516335.18518165.rev.compl.REF        ------------------------------------------------------------ 0

chimp.LOC112206744-LOC107973052-GGT2.revcompl                                        ctttcctaagggaactctactctagagttggggcttttgacttgaaccttatttccaatc 5501
chimp54546-135457.revcompl.Pan.troglodytes.clone.rp43-41g5.GenBank:AC099533.36       CTTTCCTAAGGGAACTCTACTCTAGAGTTGGGGCTTTTGACTTGAACCTTATTTCCAATC 7060
human.FAM30B-LOC105372935-GGT2.NCBI.                                                 ctttcctaagggaactctactctagagttggggcttttgacttgaaccttatttccagtc 3476
human.FAM230E-LOC105377182-GGT3P.NCBI.                                               ctttcctaagggaactctactctagagttggggcttttgacttgaaccttatttccagtc 2551
human.FAM230J-LOC105372942-GGTLC5P.NCBI.rev.compl                                    ctttcctaagggaactctactctagagttggggcttttgacttgaaccttatttccaatc 2921
human.FAM230A.ncRNA.18487127.18500594-GGTLC3.18516335.18518165.rev.compl.REF        ------------------------------------------------------------ 0
```

```
chimp.LOC112206744-LOC107973052-GGT2.revcompl                                       ttggttacccagagtttccaagtgaacaaaagacctgtgtgagccatccatagcatagcc 5561
chimp54546-135457.revcompl.Pan.troglodytes.clone.rp43-41g5.GenBank:AC099533.36      TTGGTTACCCAGAGTTTCCAAGTGAACAAAAGACCTGTGTGAGCCATCCATAGCATAGCC 7120
human.FAM30B-LOC105372935-GGT2.NCBI.                                                ttggttacccagagtttccaagtgaacaaaagacctgtgtgagccatccatagcatagcc 3536
human.FAM230E-LOC105377182-GGT3P.NCBI.                                              ttggttacccagagtttccaagtgaacaaaagacctgtgtgagccatccatagcatagcc 2611
human.FAM230J-LOC105372942-GGTLC5P.NCBI.rev.compl                                   ttggttacccagagtttccaagtgaacaaaagacctgtgtgagccatccatagcatagcc 2981
human.FAM230A.ncRNA.18487127.18500594-GGTLC3.18516335.18518165.rev.compl.REF        ------------------------------------------------------------ 0

chimp.LOC112206744-LOC107973052-GGT2.revcompl                                       tgattctcagagtgttttccttctctaattacaggtgacttcagggagcacattcaatgg 5621
chimp54546-135457.revcompl.Pan.troglodytes.clone.rp43-41g5.GenBank:AC099533.36      TGATTCTCAGAGTGTTTTCCTTCTCTAATTACAGGTGACTTCAGGGAGCACATTCAATGG 7180
human.FAM30B-LOC105372935-GGT2.NCBI.                                                tgattctcagagtgttttccttctctaattacaggtgacttcagggagcacattcaatgg 3596
human.FAM230E-LOC105377182-GGT3P.NCBI.                                              tgattctcagagtgttttccttctctaattacaggtgacttcagggagcacattcaatgg 2671
human.FAM230J-LOC105372942-GGTLC5P.NCBI.rev.compl                                   tgattctcagagtgttttccttctctaattacaggtgacttcagggagcacattcaatgg 3041
human.FAM230A.ncRNA.18487127.18500594-GGTLC3.18516335.18518165.rev.compl.REF        ------------------------------------------------------------ 0

chimp.LOC112206744-LOC107973052-GGT2.revcompl                                       tacgtattctggaatcactcactggttgttagaaaaggattctataggaaatctggagct 5681
chimp54546-135457.revcompl.Pan.troglodytes.clone.rp43-41g5.GenBank:AC099533.36      TACGTATTCTGGAATCACTCACTGGTTGTTAGAAAAGGATTCTATAGGAAATCTGGAGCT 7240
human.FAM30B-LOC105372935-GGT2.NCBI.                                                tacgtattctggaatcactcactggttgttagaaaaggattctacaggaaatctggagct 3656
human.FAM230E-LOC105377182-GGT3P.NCBI.                                              tacgtattctggaatcactcactggttgttagaaaaggattctacaggaaatctggagct 2731
human.FAM230J-LOC105372942-GGTLC5P.NCBI.rev.compl                                   tacgtattctggaatcactcactggttgttagaaaaggattctacaggaaatctggagct 3101
human.FAM230A.ncRNA.18487127.18500594-GGTLC3.18516335.18518165.rev.compl.REF        ------------------------------------------------------------ 0

chimp.LOC112206744-LOC107973052-GGT2.revcompl                                       taactgctggcttttgtctagagaggctccatgatccaagacatctggtgggaatgagga 5741
chimp54546-135457.revcompl.Pan.troglodytes.clone.rp43-41g5.GenBank:AC099533.36      TAACTGCTGGCTTTTGTCTGGAAAGCCTCCATGATCCAAGACATCTGGTGGGAATGAGGA 7300
human.FAM30B-LOC105372935-GGT2.NCBI.                                                taactgctggcttttgtctggagagcctccatgatccaagacatctggtgggaatgagga 3716
human.FAM230E-LOC105377182-GGT3P.NCBI.                                              taactgctggcttttgtctggagagcctccatgatccaagacatctggtgggaatgagga 2791
human.FAM230J-LOC105372942-GGTLC5P.NCBI.rev.compl                                   taactgctggcttttgtccggagagcctccatgatccaagacatctggtgggaatgagga 3161
human.FAM230A.ncRNA.18487127.18500594-GGTLC3.18516335.18518165.rev.compl.REF        ------------------------------------------------------------ 0

chimp.LOC112206744-LOC107973052-GGT2.revcompl                                       tgtgggatatagtaaaggaaactggttttccaggtgacatactctttttatctatgtata 5801
chimp54546-135457.revcompl.Pan.troglodytes.clone.rp43-41g5.GenBank:AC099533.36      TGTGGGGTATAGTAAAGGAAACTGGTTTTCCAGGTGACATACTCTTTTTATCTACGTATA 7360
human.FAM30B-LOC105372935-GGT2.NCBI.                                                tgtagggtatagtaaaagaaactggttttcctggtgacatactctttttatctatgtata 3776
human.FAM230E-LOC105377182-GGT3P.NCBI.                                              tgtagggtatagtaaaagaaactggttttcctggtgacatactctttttatctatgtata 2851
human.FAM230J-LOC105372942-GGTLC5P.NCBI.rev.compl                                   tgtagggtatagtaaaagaaactggttttcctggtgacatactctttttatctatgtata 3221
human.FAM230A.ncRNA.18487127.18500594-GGTLC3.18516335.18518165.rev.compl.REF        ------------------------------------------------------------ 0

chimp.LOC112206744-LOC107973052-GGT2.revcompl                                       gtttctgggaatgtgttcacattaggttgtgtgtgggtatgtgtgtattagggcgggggt 5861
chimp54546-135457.revcompl.Pan.troglodytes.clone.rp43-41g5.GenBank:AC099533.36      GTTTCTGGGAATGTGTTCACATTAGGTTGTGTGTGGGTATGTGTGTATTAGGGCGGGGGT 7420
human.FAM30B-LOC105372935-GGT2.NCBI.                                                gtttctgggaacatgttcacattaggttgtgtgtgggtatgtgtgtattagggcgggggt 3836
human.FAM230E-LOC105377182-GGT3P.NCBI.                                              gtttctgggaacatgttcacattaggttgtgtgtgggtatgtgtgtattagggcgggggt 2911
human.FAM230J-LOC105372942-GGTLC5P.NCBI.rev.compl                                   gtttctgggaacatgttcacattaggttgtgtgtgggtatgtgtgtattagggcgggggt 3281
human.FAM230A.ncRNA.18487127.18500594-GGTLC3.18516335.18518165.rev.compl.REF        ------------------------------------------------------------ 0

chimp.LOC112206744-LOC107973052-GGT2.revcompl                                       gggatgaggtggtctgtgtgcacatctgcatgatttgcttgtgaatgtgtgtctatgtgt 5921
chimp54546-135457.revcompl.Pan.troglodytes.clone.rp43-41g5.GenBank:AC099533.36      GGGGTGAGGTGGTCTGTGTGCACGTCTGCATGATTTGCTTGTGAATGTGTGTCTATGTGT 7480
human.FAM30B-LOC105372935-GGT2.NCBI.                                                ggggtgaggtggtctgtgtgcaagtctgcatgatttgcttgtgaatgtgtgtctatgtgt 3896
human.FAM230E-LOC105377182-GGT3P.NCBI.                                              ggggtgaggtggtctgtgtgcaagtctgcatgatttgcttgtgaatgtgtgtctatgtgt 2971
human.FAM230J-LOC105372942-GGTLC5P.NCBI.rev.compl                                   ggggtgaggtggtctgtgtgcaagtctgcatgatttgcttgtgaatgtgtgtctatctgt 3341
human.FAM230A.ncRNA.18487127.18500594-GGTLC3.18516335.18518165.rev.compl.REF        ------------------------------------------------------------ 0

chimp.LOC112206744-LOC107973052-GGT2.revcompl                                       gtttcccccaggaaaaaaatgttgtgtttacccagcacaactctcagtgccatttttctt 5981
chimp54546-135457.revcompl.Pan.troglodytes.clone.rp43-41g5.GenBank:AC099533.36      GTTTCCCCCAGGAAAAAAATGTTGTGTTTACCCAGCACAACTCTCAGTGCCATTTTTCTT 7540
human.FAM30B-LOC105372935-GGT2.NCBI.                                                gtttcccctaggaaaaaaatgttgtgtttacccagcacaactctcagtgccatttttctt 3956
human.FAM230E-LOC105377182-GGT3P.NCBI.                                              gtttcccctaggaaaaaaatgttgtgtttacccagcacaactctcagtgccatgtttctt 3031
human.FAM230J-LOC105372942-GGTLC5P.NCBI.rev.compl                                   gtttcccccaggaaaaaaatgttgtgtttacccagcacaactctcagtgccatttttctt 3401
human.FAM230A.ncRNA.18487127.18500594-GGTLC3.18516335.18518165.rev.compl.REF        ------------------------------------------------------------ 0

chimp.LOC112206744-LOC107973052-GGT2.revcompl                                       aatttaacaaatcagaccacatactttacttacattagttcacacctcatcatcatcatg 6041
chimp54546-135457.revcompl.Pan.troglodytes.clone.rp43-41g5.GenBank:AC099533.36      AATTTAACAAATCAGACCACATACTTTACTTACATTAGTTCACACCTCATCATCATCATG 7600
human.FAM30B-LOC105372935-GGT2.NCBI.                                                aatttaacaaatcagaccacatactttacttacattagttcacacctcatcatcatcatg 4016
human.FAM230E-LOC105377182-GGT3P.NCBI.                                              aatttaacaaatcagaccacatactttacttacattagttcacacctcatcatcatcatg 3091
human.FAM230J-LOC105372942-GGTLC5P.NCBI.rev.compl                                   aatttaacaaatcagaccacatactttacttacattagttcacacctcatcatcatcatg 3461
human.FAM230A.ncRNA.18487127.18500594-GGTLC3.18516335.18518165.rev.compl.REF        ------------------------------------------------------------ 0

chimp.LOC112206744-LOC107973052-GGT2.revcompl                                       cccatatgttgtgagcttgtttattgagcccacatgccagatggagaaactaagccacat 6101
chimp54546-135457.revcompl.Pan.troglodytes.clone.rp43-41g5.GenBank:AC099533.36      CCCATATGTCGTGAGCTTGTTTATTGAGCCCACATGCCAGATGGAGAAACTAAGCCACAT 7660
human.FAM30B-LOC105372935-GGT2.NCBI.                                                cccatatgttgtgagcttgtttattgagcccacatgccagatggagaaactcagccacat 4076
human.FAM230E-LOC105377182-GGT3P.NCBI.                                              cccatatgttgtgagcttgtttattgagcccacatgccagatggagaaactaagccacat 3151
human.FAM230J-LOC105372942-GGTLC5P.NCBI.rev.compl                                   cccatatgttgtgagcttgtttattgagcccacatgccagatggagaaactaagccacat 3521
human.FAM230A.ncRNA.18487127.18500594-GGTLC3.18516335.18518165.rev.compl.REF        ------------------------------------------------------------ 0

chimp.LOC112206744-LOC107973052-GGT2.revcompl                                       aaataaatgtgccctggttcacttgctgcatagtgaagagtcaaaatgtttactcatacg 6161
chimp54546-135457.revcompl.Pan.troglodytes.clone.rp43-41g5.GenBank:AC099533.36      AAATAAATGTGCCCTGGTTCACTTGCTGCATAGTGAAGAGTCAAAATGTTTACTCATACG 7720
human.FAM30B-LOC105372935-GGT2.NCBI.                                                aaataaatgtgccctggttcacttgctgcatagtgaagagtcaaaatgtttactcatacg 4136
human.FAM230E-LOC105377182-GGT3P.NCBI.                                              aaataaatgtgccctggttcacttgctgcatagtgaagagtcaaaatgtttactcaaacg 3211
human.FAM230J-LOC105372942-GGTLC5P.NCBI.rev.compl                                   aaataaatgtgctctggttcacttgctgcatagtgaagagtcaaaatgtttcctcatacg 3581
human.FAM230A.ncRNA.18487127.18500594-GGTLC3.18516335.18518165.rev.compl.REF        ------------------------------------------------------------ 0

chimp.LOC112206744-LOC107973052-GGT2.revcompl                                       gtgctaatgttgaaggcctgaactacaacctctatttatcagccagtgaagagatcacta 6221
chimp54546-135457.revcompl.Pan.troglodytes.clone.rp43-41g5.GenBank:AC099533.36      GTGCTAATGTTGAAGGCCTGAACTACAACCTCTATTTATCAGCCAGTGAAGAGATCACTA 7780
human.FAM30B-LOC105372935-GGT2.NCBI.                                                gtgctaatgttgaaggcctgaactacaacctctatttatcagccagtgaagagatcacta 4196
human.FAM230E-LOC105377182-GGT3P.NCBI.                                              gtgctaatgttgaaggcctgaactacaacctctatttatcagccagtgaagagatcacta 3271
human.FAM230J-LOC105372942-GGTLC5P.NCBI.rev.compl                                   gtgctaatgttgaaggcctgaactacaacctctatttatcagccagtgaagagatcacta 3641
human.FAM230A.ncRNA.18487127.18500594-GGTLC3.18516335.18518165.rev.compl.REF        ------------------------------------------------------------ 0

chimp.LOC112206744-LOC107973052-GGT2.revcompl                                       ttcaccatgcaagggagttccagcaccctctatgcctggaattacccacgcctgcagaga 6281
chimp54546-135457.revcompl.Pan.troglodytes.clone.rp43-41g5.GenBank:AC099533.36      TTCACCATGCAAGGGAGTTCCAGCACCCTCTATGCCTGGAATTACCCATGCCTGCAGAGA 7840
human.FAM30B-LOC105372935-GGT2.NCBI.                                                ttccccatgcaagggagttccagcaccctctatgcctggaattacccacacctgcagaga 4256
human.FAM230E-LOC105377182-GGT3P.NCBI.                                              ttcaccatgcaagggagttccagcaccctctatgcctggaattacccacgcctgcagaga 3331
human.FAM230J-LOC105372942-GGTLC5P.NCBI.rev.compl                                   ttcaccatgcaagggagttccagcaccctctatgcctggaattacccacgcctgcagaga 3701
human.FAM230A.ncRNA.18487127.18500594-GGTLC3.18516335.18518165.rev.compl.REF        ------------------------------------------------------------ 0

chimp.LOC112206744-LOC107973052-GGT2.revcompl                                       tcccaaacgccatccctcacataagagagcctcatgatctcataatccaggtagctatgt 6341
chimp54546-135457.revcompl.Pan.troglodytes.clone.rp43-41g5.GenBank:AC099533.36      TCCCAAACGCCATCCCTCACATAAGAGAGCCTCATGATCTCATAATCCAGGTAGCTATGT 7900
human.FAM30B-LOC105372935-GGT2.NCBI.                                                tcccaaacgccatccctcacataagacagcctcatgatctcataatccaggtagctatgt 4316
human.FAM230E-LOC105377182-GGT3P.NCBI.                                              tcccaaacgccatccctcacataagacagcctcatgatctcataatccaggtagctatgt 3391
human.FAM230J-LOC105372942-GGTLC5P.NCBI.rev.compl                                   tcccaaacgccatccctcacataagacagcctcatgatctcataatccaggtagctatgt 3761
human.FAM230A.ncRNA.18487127.18500594-GGTLC3.18516335.18518165.rev.compl.REF        ------------------------------------------------------------ 0

chimp.LOC112206744-LOC107973052-GGT2.revcompl                                       agacatcttcctgcaggtgtcacatagtccttagtgggaaaccaacatagaaagcccatg 6401
chimp54546-135457.revcompl.Pan.troglodytes.clone.rp43-41g5.GenBank:AC099533.36      AGACATCTTCCTGCAGGTGTCACATAGTCCTTAGTGTGAAACCAACATAGAAAGCCCATG 7960
human.FAM30B-LOC105372935-GGT2.NCBI.                                                agacatcttcctgcaggtgtcacatagtccttagtgtgaaaccaacatagaaagcccatg 4376
human.FAM230E-LOC105377182-GGT3P.NCBI.                                              agacatcttcctgcaggtgtcacatagtccttagtgtgaaaccaacatagaaagcccatg 3451
human.FAM230J-LOC105372942-GGTLC5P.NCBI.rev.compl                                   agacatcttcctgcaggtgtcacatagtccttagtgtgaaaccaacatagaaagcccatg 3821
human.FAM230A.ncRNA.18487127.18500594-GGTLC3.18516335.18518165.rev.compl.REF        ------------------------------------------------------------ 0

chimp.LOC112206744-LOC107973052-GGT2.revcompl                                       tttctgatcaaatcacaggttctgaaacactaagggaagcactaagtaggacaatgtggt 6461
chimp54546-135457.revcompl.Pan.troglodytes.clone.rp43-41g5.GenBank:AC099533.36      TTTCTGATCAAATCACAGGTTCTGAAACACTAAGTGAGGCACTAAGTAGGACAATGTGGT 8020
human.FAM30B-LOC105372935-GGT2.NCBI.                                                tttctgatcaaatcacaggttctgaaacactaagggaggcactaagtaggacaacgtggt 4436
human.FAM230E-LOC105377182-GGT3P.NCBI.                                              tttctgatcaaatcacaggttctgaaacactaagggaggcactaagtaggacaacgtggt 3511
human.FAM230J-LOC105372942-GGTLC5P.NCBI.rev.compl                                   tttctgatcaaatcacaggttctgaaacactaagggaggcactaagtaggacaacgtggt 3881
human.FAM230A.ncRNA.18487127.18500594-GGTLC3.18516335.18518165.rev.compl.REF        ------------------------------------------------------------ 0

chimp.LOC112206744-LOC107973052-GGT2.revcompl                                       gcctgtgtgtcatagctgggtctcctcaagacatggatcaagtccaataagaattgggga 6521
chimp54546-135457.revcompl.Pan.troglodytes.clone.rp43-41g5.GenBank:AC099533.36      GCCTGCATGTCATAGCTGGGTCTCCTCAAGACATGGATCAAGTCCAATAAGAATTGGAGA 8080
human.FAM30B-LOC105372935-GGT2.NCBI.                                                gcctgcgtgtcatagctgggtctcctcaagacatggatcaagtccagtaagaattgggga 4496
human.FAM230E-LOC105377182-GGT3P.NCBI.                                              gcctgcgtgtcatagctgggtctcctcaagacatggatcaagtccagtaagaattgggga 3571
human.FAM230J-LOC105372942-GGTLC5P.NCBI.rev.compl                                   gcctgcgtgtcatagctgggtctcctcaagacatggatcaagtccagtaagaattgggga 3941
human.FAM230A.ncRNA.18487127.18500594-GGTLC3.18516335.18518165.rev.compl.REF        ------------------------------------------------------------ 0

chimp.LOC112206744-LOC107973052-GGT2.revcompl                                       gatgctttagagtcttgatggagttatcaccacaagccctctgagctacacactttaggg 6581
chimp54546-135457.revcompl.Pan.troglodytes.clone.rp43-41g5.GenBank:AC099533.36      GATGCTTTAGAGTCTTGATGGAGTTATCACCACAAGCCCTCTGAGCTACACACTTTAGGG 8140
human.FAM30B-LOC105372935-GGT2.NCBI.                                                gatgctttagagtcttgatggagttatcaccacaagccctctgagctacacactttaggg 4556
human.FAM230E-LOC105377182-GGT3P.NCBI.                                              gatgctttagagtcttgatggagttatcaccacaagccctctgagctacacactttaggg 3631
human.FAM230J-LOC105372942-GGTLC5P.NCBI.rev.compl                                   gatgctttagagtcttgatggagttatcaccacaagccctctgagctacacactttaggg 4001
human.FAM230A.ncRNA.18487127.18500594-GGTLC3.18516335.18518165.rev.compl.REF        ------------------------------------------------------------ 0
```

```
chimp.LOC112206744-LOC107973052-GGT2.revcompl                                   atcatgaccattaagtactcaaattaccacttggttgttatccgggtatccgtcgtcctt 6641
chimp54546-135457.revcompl.Pan.troglodytes.clone.rp43-41g5.GenBank:AC099533.36  ATCATGACCATTAAGTACTCAAATTACCATTTGGTTGTTATCTGGGTATCCGTCGTCCTT 8200
human.FAM30B-LOC105372935-GGT2.NCBI.                                            atcatgaccattaagtactcaaattaccatttggttgctatccgggtatccgtcgtcctt 4616
human.FAM230E-LOC105377182-GGT3P.NCBI.                                          atcatgaccattaagtactcaaattaccatttggttgttatccgggtatccgtcgtcctt 3691
human.FAM230J-LOC105372942-GGTLC5P.NCBI.rev.compl                               atcatgaccattaagtactcaaattaccatttggttgttatccgggtatccgtcgtcctt 4061
human.FAM230A.ncRNA.18487127.18500594-GGTLC3.18516335.18518165.rev.compl.REF    ------------------------------------------------------------ 0

chimp.LOC112206744-LOC107973052-GGT2.revcompl                                   gtggcaactctcttgtgaagctggtgtggacagcctcagtgctggagctgtgcctgcctt 6701
chimp54546-135457.revcompl.Pan.troglodytes.clone.rp43-41g5.GenBank:AC099533.36  GTGGCAACTCTCTTGTGAAGCTGGTGTGGACAGTCTCAGTGCTGGAGCTGTGCCTGCCTT 8260
human.FAM30B-LOC105372935-GGT2.NCBI.                                            gtggcaaccctcttgtgaagctggtgtggacagcctcagtgctggagctgtgcctgcctt 4676
human.FAM230E-LOC105377182-GGT3P.NCBI.                                          gtggcaaccctcttgtgaagctggtgtggacagcctcagtgctggagctgtgcctgcctt 3751
human.FAM230J-LOC105372942-GGTLC5P.NCBI.rev.compl                               gtggcaaccctcttgtgaagctggtgtggacagcctcagtgctggagctgtgcctccctt 4121
human.FAM230A.ncRNA.18487127.18500594-GGTLC3.18516335.18518165.rev.compl.REF    ------------------------------------------------------------ 0

chimp.LOC112206744-LOC107973052-GGT2.revcompl                                   ctgaatggaccctttctgtgttagcaggtgggtacaagcatgggggtcagcacactcagt 6761
chimp54546-135457.revcompl.Pan.troglodytes.clone.rp43-41g5.GenBank:AC099533.36  CTGAATGGACCCTTTCTGTGTTAGCAGGTGGGTACAAGCATGGGGGTCAGCACACTCAGT 8320
human.FAM30B-LOC105372935-GGT2.NCBI.                                            ctgagtggaccctttctgtgttagcaggtgggtacaagcgtgggggtcagcacactcagt 4736
human.FAM230E-LOC105377182-GGT3P.NCBI.                                          ctgagtggaccctttctgtgttagcaggtgggtacaagcgtgggggtcagcacactcagt 3811
human.FAM230J-LOC105372942-GGTLC5P.NCBI.rev.compl                               ctgagtggaccctttctgtgttagcaggtgggtacaagcgtgggggtcagcacactcagt 4181
human.FAM230A.ncRNA.18487127.18500594-GGTLC3.18516335.18518165.rev.compl.REF    ------------------------------------------------------------ 0

chimp.LOC112206744-LOC107973052-GGT2.revcompl                                   ggatttacacacacagcattgaagagtaaggctgggcttcattatttatacattttcaat 6821
chimp54546-135457.revcompl.Pan.troglodytes.clone.rp43-41g5.GenBank:AC099533.36  GGATTTACACACACAGCATTGAAGAGTAAGGCTGGGCTTCATTATTTATACATTTTCAAT 8380
human.FAM30B-LOC105372935-GGT2.NCBI.                                            ggatttacacacacagcgttgaagagtaaggctgggcttcattatttatacattttcaat 4796
human.FAM230E-LOC105377182-GGT3P.NCBI.                                          ggatttacacacacagcgttgaagagtaaggctgggcttcgttatttatacattttcaat 3871
human.FAM230J-LOC105372942-GGTLC5P.NCBI.rev.compl                               ggatttacacacacagcgttgaagagtaaggctgggcttcattatttatacattttcaat 4241
human.FAM230A.ncRNA.18487127.18500594-GGTLC3.18516335.18518165.rev.compl.REF    ------------------------------------------------------------ 0

chimp.LOC112206744-LOC107973052-GGT2.revcompl                                   aaatgatgatcttcataacataaaatcaatgacgtagtacactagaatactgtccctagt 6881
chimp54546-135457.revcompl.Pan.troglodytes.clone.rp43-41g5.GenBank:AC099533.36  AAATGATGATCTTCATAACATAAAATCAATGATGTAGTACACTAGAATACTGTCCCTAGT 8440
human.FAM30B-LOC105372935-GGT2.NCBI.                                            aaatgatgatcttcataacataaaatcaatgatgtagtacactagaatactgtccctagt 4856
human.FAM230E-LOC105377182-GGT3P.NCBI.                                          aaatgatgatcttcataacataaaatcaatgatgtagtacactagaatactgtccctagt 3931
human.FAM230J-LOC105372942-GGTLC5P.NCBI.rev.compl                               aaatgatgatcttcataacataaaatcaatgatgtagtacactagaatactgtccctagt 4301
human.FAM230A.ncRNA.18487127.18500594-GGTLC3.18516335.18518165.rev.compl.REF    ------------------------------------------------------------ 0

chimp.LOC112206744-LOC107973052-GGT2.revcompl                                   attgaatcttgtctctcaacaaagggttgcttaaagtcacatgacagattccattcaact 6941
chimp54546-135457.revcompl.Pan.troglodytes.clone.rp43-41g5.GenBank:AC099533.36  ATTGAATCTTGTCTCTCAACAAAGGGTTGCTTAAAGTCACATGACAGATTCCATTCAACT 8500
human.FAM30B-LOC105372935-GGT2.NCBI.                                            attgaatcttgtctctcagcaaagggttgcttaaagtcacgtgacagattccattcaact 4916
human.FAM230E-LOC105377182-GGT3P.NCBI.                                          attgaatcttgtctctcagcaaagggttgcttaaagtcacgtgacagattccattcaact 3991
human.FAM230J-LOC105372942-GGTLC5P.NCBI.rev.compl                               attgaatcttgtctctcagcaaagggttgcttaaagtcacgtgacagattccattcaact 4361
human.FAM230A.ncRNA.18487127.18500594-GGTLC3.18516335.18518165.rev.compl.REF    ------------------------------------------------------------ 0

chimp.LOC112206744-LOC107973052-GGT2.revcompl                                   gatgacacatgctgtagcagcagttaaagcagtcatttgaaaaggcttttactataaact 7001
chimp54546-135457.revcompl.Pan.troglodytes.clone.rp43-41g5.GenBank:AC099533.36  GACGACACATGCTGTAGCAGCAGTTAAAGCAGTCATTTGAAAAGGCTTTTACTATAAACT 8560
human.FAM30B-LOC105372935-GGT2.NCBI.                                            gatgacacatgctgtagcagcagttaaagcagtcatttgaaaaggcttttactataaact 4976
human.FAM230E-LOC105377182-GGT3P.NCBI.                                          gatgacacatgctgtagcagcagttaaagcagtcatttgaaaaggcttttactataaact 4051
human.FAM230J-LOC105372942-GGTLC5P.NCBI.rev.compl                               gatgacacatgctgtagcagcagttaaagcagtcatttgaaaaggcttttactataaact 4421
human.FAM230A.ncRNA.18487127.18500594-GGTLC3.18516335.18518165.rev.compl.REF    ------------------------------------------------------------ 0

chimp.LOC112206744-LOC107973052-GGT2.revcompl                                   tatgtgtgagcctgaagtggggggataaaagaggtgattagctcccctgtgccatgtttct 7061
chimp54546-135457.revcompl.Pan.troglodytes.clone.rp43-41g5.GenBank:AC099533.36  TATGTGTGAGCCTGAAGTGGGGGATAAAAGAGGTGATTAGCTCCCCTGTGCCATGTTTCT 8620
human.FAM30B-LOC105372935-GGT2.NCBI.                                            tacgtgtgagcctgaagtgggggataaaagaggcgattagctcccctgtgccatgtttct 5036
human.FAM230E-LOC105377182-GGT3P.NCBI.                                          tatgtgtgagcctgaagtgggggataaaagaggtgattagctcccctgtgccatgtttct 4111
human.FAM230J-LOC105372942-GGTLC5P.NCBI.rev.compl                               tatgtgtgagcctgaagtgggggataaaagaggcgattagctcccctgtgccatgtttct 4481
human.FAM230A.ncRNA.18487127.18500594-GGTLC3.18516335.18518165.rev.compl.REF    ------------------------------------------------------------ 0

chimp.LOC112206744-LOC107973052-GGT2.revcompl                                   attatgtgcatggtggaggaaaattacacaggaaggtgatggagagaacagagcaaagga 7121
chimp54546-135457.revcompl.Pan.troglodytes.clone.rp43-41g5.GenBank:AC099533.36  ATTATGTGCGTGGTGGAGGAAAATTACACAGGAAGGTGATGGAGAGAACAGAGCAAAGGT 8680
human.FAM30B-LOC105372935-GGT2.NCBI.                                            attatgtgcgtggtggaggaaaattacacaggaaggtgatggagagaacagagcaaagga 5096
human.FAM230E-LOC105377182-GGT3P.NCBI.                                          gttatgtgcgtggtggaggaaaattacacaggaaggtgatggagagaacagagcaaagga 4171
human.FAM230J-LOC105372942-GGTLC5P.NCBI.rev.compl                               attatgtgcgtggtggaggaaaattacacaggaaggtgatggagagaacagagcaaagga 4541
human.FAM230A.ncRNA.18487127.18500594-GGTLC3.18516335.18518165.rev.compl.REF    ------------------------------------------------------------ 0

chimp.LOC112206744-LOC107973052-GGT2.revcompl                                   ttggacaggtccattgaacccataagactatggtgaggttagtgaatgagattggtcatt 7181
chimp54546-135457.revcompl.Pan.troglodytes.clone.rp43-41g5.GenBank:AC099533.36  TTGGACGGGTCCATTGAACCCATAAGACTATGGTGAGGTTAGTGAATGAGATTGGTCATT 8740
human.FAM30B-LOC105372935-GGT2.NCBI.                                            ttggacaggtccattgaacccataagactatggtgaggttagtgaatgagattggtcatt 5156
human.FAM230E-LOC105377182-GGT3P.NCBI.                                          ttggacaggtccattgaacccataagactatggtgaggttagtgaatgagactggtcatt 4231
human.FAM230J-LOC105372942-GGTLC5P.NCBI.rev.compl                               ttggacaggtccattgaacccataagactatggtgaggttagtgaatgagattggtcatt 4601
human.FAM230A.ncRNA.18487127.18500594-GGTLC3.18516335.18518165.rev.compl.REF    ------------------------------------------------------------ 0

chimp.LOC112206744-LOC107973052-GGT2.revcompl                                   ttaggtcaaattctacccagagctggtgcagccactgcccattcttagccagaccttatt 7241
chimp54546-135457.revcompl.Pan.troglodytes.clone.rp43-41g5.GenBank:AC099533.36  TTAGGTCAAATTTTACCCAGAGCTGGTGCAGCCACTGCCCATTCTTAGCCAGACCTTATT 8800
human.FAM30B-LOC105372935-GGT2.NCBI.                                            ttaggtcaaattttacccagagctggtgcagccactgcccattcttagccagaccttatt 5216
human.FAM230E-LOC105377182-GGT3P.NCBI.                                          ttaggtcaaattttacccagagctggtacagccactgccgattcttagccagaccttatt 4291
human.FAM230J-LOC105372942-GGTLC5P.NCBI.rev.compl                               ttaggtcaaattttacccagagctggtgcagccactgcccattcttagccagaccttatt 4661
human.FAM230A.ncRNA.18487127.18500594-GGTLC3.18516335.18518165.rev.compl.REF    ------------------------------------------------------------ 0

chimp.LOC112206744-LOC107973052-GGT2.revcompl                                   gcaggcagctctgatcaatagtcaaggaggcagtggggcttgcagacttaattcattaaa 7301
chimp54546-135457.revcompl.Pan.troglodytes.clone.rp43-41g5.GenBank:AC099533.36  GCAGGCAGCTCTGATCAATAGTCAAGGAGGCAGTGGGGGTTGCAGACTTAATTCATTAAA 8860
human.FAM30B-LOC105372935-GGT2.NCBI.                                            gcaggcagctctgatcaatagtcaaggaggcagtggggcttgcagacttaattcattaaa 5276
human.FAM230E-LOC105377182-GGT3P.NCBI.                                          gcaggcagctctgatcaatagtcaaggaggcagtgggggttgcagactttattcattaaa 4351
human.FAM230J-LOC105372942-GGTLC5P.NCBI.rev.compl                               gcaggcagctctgatcaatagtcaaggaggcagtggggcttgcagacttaattcattaaa 4721
human.FAM230A.ncRNA.18487127.18500594-GGTLC3.18516335.18518165.rev.compl.REF    ------------------------------------------------------------ 0

chimp.LOC112206744-LOC107973052-GGT2.revcompl                                   tcaccaaagcaccagcccacacggccacttttccagttaattcacagtagcttgcatatt 7361
chimp54546-135457.revcompl.Pan.troglodytes.clone.rp43-41g5.GenBank:AC099533.36  TCACCAAAGCACCAGCCCACACGGCCACTTTTCCAGTTAATTCACAGTAGCTTGCATATT 8920
human.FAM30B-LOC105372935-GGT2.NCBI.                                            tcaccaaagcaccagcccaaacggccacttttccagttaattgacagtagcttgcatatt 5336
human.FAM230E-LOC105377182-GGT3P.NCBI.                                          tcaccaaagcaccagcccacacggccacttttccagttaattcacagtagcttgcatatt 4411
human.FAM230J-LOC105372942-GGTLC5P.NCBI.rev.compl                               tcaccaaagcaccagcccacacggccacttttccagttaattgacagtagcttgcatatt 4781
human.FAM230A.ncRNA.18487127.18500594-GGTLC3.18516335.18518165.rev.compl.REF    ------------------------------------------------------------ 0

chimp.LOC112206744-LOC107973052-GGT2.revcompl                                   caggtttgatcagtggaagggaagttactctttgcagacccatcttttgacaatcatttt 7421
chimp54546-135457.revcompl.Pan.troglodytes.clone.rp43-41g5.GenBank:AC099533.36  CAGGTTTGATCAGTGGAAGGGAAGTTACTCTTTGCAGACCCATCTTTTGACAATCATTTT 8980
human.FAM30B-LOC105372935-GGT2.NCBI.                                            caggtttgatcagtggaagggaagttactctttgcagacccatcttttgacaatcatttt 5396
human.FAM230E-LOC105377182-GGT3P.NCBI.                                          caggtttgatcagtggaagggaagttactctttgcagactcat-ttttgacaatcatttt 4470
human.FAM230J-LOC105372942-GGTLC5P.NCBI.rev.compl                               caggtttgatcagtggaagggaagttactctttgcagacccat-ttttgacaatcatttt 4840
human.FAM230A.ncRNA.18487127.18500594-GGTLC3.18516335.18518165.rev.compl.REF    ------------------------------------------------------------ 0

chimp.LOC112206744-LOC107973052-GGT2.revcompl                                   gcagtgtcagaaggtctgagcagcctcgggaggcaagcagtccctggtccctcagtgtag 7481
chimp54546-135457.revcompl.Pan.troglodytes.clone.rp43-41g5.GenBank:AC099533.36  GCAGTGTCAGAAGGTCTGAGCAGCCTCGGGAGGCAAGCAGTCCCTGGTCCCTCAGTGTAG 9040
human.FAM30B-LOC105372935-GGT2.NCBI.                                            gtggtgtcggaaggtctgagcagcctcgggaggcaagcagtccctggtccctcagtgtag 5456
human.FAM230E-LOC105377182-GGT3P.NCBI.                                          gcggtgtcagaaggtctgagcagcctcgggaggcaagcagtccctggtccctcagtgtag 4530
human.FAM230J-LOC105372942-GGTLC5P.NCBI.rev.compl                               gcggtgtcagaaggtgtgagcagcctcgggaggcaagcagtccctggtccctcagtgtag 4900
human.FAM230A.ncRNA.18487127.18500594-GGTLC3.18516335.18518165.rev.compl.REF    ------------------------------------------------------------ 0

chimp.LOC112206744-LOC107973052-GGT2.revcompl                                   tcactggaggagacagtcactgagaggcagctggcagggtgaagggaaaggggaggcagg 7541
chimp54546-135457.revcompl.Pan.troglodytes.clone.rp43-41g5.GenBank:AC099533.36  TCACTGGAGGAGACAGTCACTGAGAGGCAGCTGGCAGGGTGAAGGGAAAGGGGAGGCAGG 9100
human.FAM30B-LOC105372935-GGT2.NCBI.                                            tcactggaggagacagtcactgagaggcagctggcagggtgaagggaaaggggaggcagg 5516
human.FAM230E-LOC105377182-GGT3P.NCBI.                                          tcactggaggagacagtcactgagaggcagctggcagggtgaagggaaaggggaggcagg 4590
human.FAM230J-LOC105372942-GGTLC5P.NCBI.rev.compl                               tcactggaggagacagtcactgagaggcagctggcagggtgaagggcaaggggaggcagg 4960
human.FAM230A.ncRNA.18487127.18500594-GGTLC3.18516335.18518165.rev.compl.REF    ------------------------------------------------------------ 0

chimp.LOC112206744-LOC107973052-GGT2.revcompl                                   ccacagagatgacagcctttaagctgtcatactgg-gaggtcaaggatctgaaagaggaa 7600
chimp54546-135457.revcompl.Pan.troglodytes.clone.rp43-41g5.GenBank:AC099533.36  CCACAGAGATGACAGCCTTTAAGCTGTCATACTGG-GAGGTCAAGGATCTGAAAGAGGAA 9159
human.FAM30B-LOC105372935-GGT2.NCBI.                                            ccacagagatgacagcctttaagctgtcatactgg-gaggtcaaggatctgaaagaggaa 5575
human.FAM230E-LOC105377182-GGT3P.NCBI.                                          ccacagagatgacagtctttaagctgtcatactggtgaggtcaaggatctgaaagaggaa 4650
human.FAM230J-LOC105372942-GGTLC5P.NCBI.rev.compl                               ccacagagatgacagcctttaagctgtcatactgg-gaggtcaaggatctgaaagaggaa 5019
human.FAM230A.ncRNA.18487127.18500594-GGTLC3.18516335.18518165.rev.compl.REF    ------------------------------------------------------------ 0

chimp.LOC112206744-LOC107973052-GGT2.revcompl                                   ggagaattctttatcattaaggacctgtccttatctcaggcatttcctccagagcatcac 7660
chimp54546-135457.revcompl.Pan.troglodytes.clone.rp43-41g5.GenBank:AC099533.36  GGAGAATTCTTTATCATTAAGGACCTGTCCTTATCTCAGGCATTTCCTCCAGAGCATCAC 9219
human.FAM30B-LOC105372935-GGT2.NCBI.                                            ggagaattctttatcattaaggacctgtccttatctcaggcatttcctccagagcatcac 5635
human.FAM230E-LOC105377182-GGT3P.NCBI.                                          ggagaattctttatcattaaggacctgtccttatctcaggcatttcctccagagcatcac 4710
human.FAM230J-LOC105372942-GGTLC5P.NCBI.rev.compl                               ggagaattctttatcattaaggacctgtccttatctcaggcatttcctccagagcaccac 5079
human.FAM230A.ncRNA.18487127.18500594-GGTLC3.18516335.18518165.rev.compl.REF    ------------------------------------------------------------ 0
```

```
chimp.LOC112206744-LOC107973052-GGT2.revcompl                                   ctttgtccacccacacaccttgggctaggaggactggagaaagacagtgaggggtctctt 7720
chimp54546-135457.revcompl.Pan.troglodytes.clone.rp43-41g5.GenBank:AC099533.36  TTTTGTCCACCCACACACCTTGGGCTAGGAGGACTGGAGAAAGACAGTGAGGGGTCTCTT 9279
human.FAM30B-LOC105372935-GGT2.NCBI.                                            ctttgtccacccacacaccttgggctaggaggactggagaaagacagtgaggggtctctt 5695
human.FAM230E-LOC105377182-GGT3P.NCBI.                                          ctttgtccacccacacaccttgggctaggaggactggagaaagacagtgaggggtctctt 4770
human.FAM230J-LOC105372942-GGTLC5P.NCBI.rev.compl                               ctttgtccacccacacaccttgggctaggaggactggagaaagacagtgaggggtctctt 5139
human.FAM230A.ncRNA.18487127.18500594-GGTLC3.18516335.18518165.rev.compl.REF    ------------------------------------------------------------ 0

chimp.LOC112206744-LOC107973052-GGT2.revcompl                                   gggtctctggcacagggcgtgatgaagaggtggcagtttttcaggaatctctctctctag 7780
chimp54546-135457.revcompl.Pan.troglodytes.clone.rp43-41g5.GenBank:AC099533.36  GGGTCTCTGGCACAGGGCGTGATGAAGAGGTGGCAGTTTTTCAGGAATCTCTCTCTCTAG 9339
human.FAM30B-LOC105372935-GGT2.NCBI.                                            gggtctctggcacagggcgtgatgaagaggtggcagtttttcaggaatctctctctctag 5755
human.FAM230E-LOC105377182-GGT3P.NCBI.                                          gggtctctggcacagggcgtgatgaagaggtggcagtttttcaggaatctctctctctag 4830
human.FAM230J-LOC105372942-GGTLC5P.NCBI.rev.compl                               gggtctgtggcacagggcgtgatgaagaggtggcagtttttcaggaatctctctctctag 5199
human.FAM230A.ncRNA.18487127.18500594-GGTLC3.18516335.18518165.rev.compl.REF    ------------------------------------------------------------ 0

chimp.LOC112206744-LOC107973052-GGT2.revcompl                                   ggaaccaaatacatttcccatctcagctccttcactcagcggggttgaggttctgctcgt 7840
chimp54546-135457.revcompl.Pan.troglodytes.clone.rp43-41g5.GenBank:AC099533.36  GGAACCAAATACATTTCCCATCTCAGGTCCTTCACTCAGCGGGGTTGAGGTTCTGCTCGT 9399
human.FAM30B-LOC105372935-GGT2.NCBI.                                            ggaaccaaatacatttcccatctcaggtccttcactcagcggggttgaggttctgctcgt 5815
human.FAM230E-LOC105377182-GGT3P.NCBI.                                          ggaaccaaatacatttcccatctcaggtccttcactcagcggggttgaggttctgctcgt 4890
human.FAM230J-LOC105372942-GGTLC5P.NCBI.rev.compl                               ggaaccaaatacatttcccatctcaggtccttcactcagcggggttgaggttctgctcgt 5259
human.FAM230A.ncRNA.18487127.18500594-GGTLC3.18516335.18518165.rev.compl.REF    ------------------------------------------------------------ 0

chimp.LOC112206744-LOC107973052-GGT2.revcompl                                   cacttatcatctctgaatgtcagcaccctcaagtgtaaaatctcagccacagccccttct 7900
chimp54546-135457.revcompl.Pan.troglodytes.clone.rp43-41g5.GenBank:AC099533.36  CACTTATCATCTCTGAATGTCAGCACCCTCAAGTGTAAAATCTCAGCCACAGCCCCTCCT 9459
human.FAM30B-LOC105372935-GGT2.NCBI.                                            cacttatcatctctgaatgtcagcaccctcaagtgtaaaatctcagccacagcccctcct 5875
human.FAM230E-LOC105377182-GGT3P.NCBI.                                          cacttatcatctctgaatgtcagcaccctcaagtgtaaaatctcagccacagcccctcct 4950
human.FAM230J-LOC105372942-GGTLC5P.NCBI.rev.compl                               cacttatcatctctgaatgtcagcaccctcaagtgtaaaatctcagccacagcccctcct 5319
human.FAM230A.ncRNA.18487127.18500594-GGTLC3.18516335.18518165.rev.compl.REF    ------------------------------------------------------------ 0

chimp.LOC112206744-LOC107973052-GGT2.revcompl                                   ctgcaccccctgcagggctgatgttctccataaaccataaggcatcatgcccacggaaaa 7960
chimp54546-135457.revcompl.Pan.troglodytes.clone.rp43-41g5.GenBank:AC099533.36  CTGCACCCCCTGCAGGGCTGATGTTCTCCATAAACCATAAGGCATCATGCCCACGGAAAA 9519
human.FAM30B-LOC105372935-GGT2.NCBI.                                            ctgcaccccctgcagggctgatgttctccataaaccataaggcgtcatgcccacggaaaa 5935
human.FAM230E-LOC105377182-GGT3P.NCBI.                                          ctgcaccccctgcagggctgatgttctccataaaccataaggcgtcatgcccacggaaaa 5010
human.FAM230J-LOC105372942-GGTLC5P.NCBI.rev.compl                               ctgcaccccctgcagggctgatgttctccataaaccataaggcgtcatgcccacggaaaa 5379
human.FAM230A.ncRNA.18487127.18500594-GGTLC3.18516335.18518165.rev.compl.REF    ------------------------------------------------------------ 0

chimp.LOC112206744-LOC107973052-GGT2.revcompl                                   gccgaacaggaaagcatgctccactgccccagagccatccaagttcccccctccatattcc 8020
chimp54546-135457.revcompl.Pan.troglodytes.clone.rp43-41g5.GenBank:AC099533.36  GCCGAACAGGAAAGCATGCTCCACTGCCCCGGAGCCATCCAAGTTCCCCCTCCATATTCC 9579
human.FAM30B-LOC105372935-GGT2.NCBI.                                            gccgaacaggaaagcatgctccactgccccggagtcatgcaagttcccccctccatattcc 5995
human.FAM230E-LOC105377182-GGT3P.NCBI.                                          gccgaacaggaaagcatgctccactgccccggagccatccaagttcccccctccatattcc 5070
human.FAM230J-LOC105372942-GGTLC5P.NCBI.rev.compl                               gccgaacaggaaagcatgctccactgccccggagccatccaagttcccccctccatattcc 5439
human.FAM230A.ncRNA.18487127.18500594-GGTLC3.18516335.18518165.rev.compl.REF    ------------------------------------------------------------ 0

chimp.LOC112206744-LOC107973052-GGT2.revcompl                                   gccactgctaagtgtccagcctattcctcctggcatgtagtaaacacttagagaacatta 8080
chimp54546-135457.revcompl.Pan.troglodytes.clone.rp43-41g5.GenBank:AC099533.36  GCCACTGCTAAGTGTCCAGCCTATTCCTCCTGGCATGTAGTAAACACTTAGAGAACATTA 9639
human.FAM30B-LOC105372935-GGT2.NCBI.                                            gccactgctaagtgtccagcttattcctcctggcatgtagtaaacacttagagaacatta 6055
human.FAM230E-LOC105377182-GGT3P.NCBI.                                          gccactgctaagtgtccagcttattcctcctggcctgtagtaaacacttagagaacatta 5130
human.FAM230J-LOC105372942-GGTLC5P.NCBI.rev.compl                               gccactgctaagtgtccagcttattcctcctggcatgtagtaaacacttagagaacatta 5499
human.FAM230A.ncRNA.18487127.18500594-GGTLC3.18516335.18518165.rev.compl.REF    ------------------------------------------------------------ 0

chimp.LOC112206744-LOC107973052-GGT2.revcompl                                   ctgaagtaccagtcctctccaaggttttcctgtatttagtgatttttttagccctgtactg 8140
chimp54546-135457.revcompl.Pan.troglodytes.clone.rp43-41g5.GenBank:AC099533.36  CTGAAGTACCAGTCCTCTGTAAGGTTTTCCTGTATTTAGTGATTTTTTAGCCCTGTACTG 9699
human.FAM30B-LOC105372935-GGT2.NCBI.                                            ctgaagtaccagtcctctctaaggttttcctgtatttagtgatttttttagccctgcactg 6115
human.FAM230E-LOC105377182-GGT3P.NCBI.                                          ctgaagtaccagtcctctctaaggttttcctgtatttagtgatttttttagccctgtactg 5190
human.FAM230J-LOC105372942-GGTLC5P.NCBI.rev.compl                               ctgaagtaccagtcctctctaaggttttcctgtatttagtgatttttttagccctgtactg 5559
human.FAM230A.ncRNA.18487127.18500594-GGTLC3.18516335.18518165.rev.compl.REF    ------------------------------------------------------------ 0

chimp.LOC112206744-LOC107973052-GGT2.revcompl                                   tgatactaagaagtagggcctaaatagggcctaaaaagtattgctaaaattacattatga 8200
chimp54546-135457.revcompl.Pan.troglodytes.clone.rp43-41g5.GenBank:AC099533.36  TGATACTAAGAAGTAGGGCCTAAATAGGGCCTAAAAAGTATTGCTAAAATTACATTATGA 9759
human.FAM30B-LOC105372935-GGT2.NCBI.                                            tgatactaagaagtagggcttaaatagggcctaaaaagtattgctaaaattacattatga 6175
human.FAM230E-LOC105377182-GGT3P.NCBI.                                          tgatactaagaagtagggcctaaatagggcctaaaaagtattgctaaaattacattatga 5250
human.FAM230J-LOC105372942-GGTLC5P.NCBI.rev.compl                               tgatactaagaagtagggcttaaatagggcctaaaaagtattgctaaaattacattatga 5619
human.FAM230A.ncRNA.18487127.18500594-GGTLC3.18516335.18518165.rev.compl.REF    ------------------------------------------------------------ 0

chimp.LOC112206744-LOC107973052-GGT2.revcompl                                   cagtgcagagaactgagggcagagggaggacatgagcttgccaggtccacatggcttagt 8260
chimp54546-135457.revcompl.Pan.troglodytes.clone.rp43-41g5.GenBank:AC099533.36  CAGTGCAGAGAACTGAGGGCAGAGGGAGGACATGAGCTTGCCAGGTCTACATGGCTTAGT 9819
human.FAM30B-LOC105372935-GGT2.NCBI.                                            cagtgcagagaactgagggcagagggaggacatgagcttgccaggtccacatggcttagt 6235
human.FAM230E-LOC105377182-GGT3P.NCBI.                                          cagtgcagagaactgagggcagagggaagacatgagcttgccaggtccacatggcttagt 5310
human.FAM230J-LOC105372942-GGTLC5P.NCBI.rev.compl                               cagtgcagagaactgagggcagagggaggacatgagcttgccaggtccacatggcttagt 5679
human.FAM230A.ncRNA.18487127.18500594-GGTLC3.18516335.18518165.rev.compl.REF    ------------------------------------------------------------ 0

chimp.LOC112206744-LOC107973052-GGT2.revcompl                                   ggaatttgaatccgggcccccactctgcaccagccctgcactcacagtcatcctgctgtg 8320
chimp54546-135457.revcompl.Pan.troglodytes.clone.rp43-41g5.GenBank:AC099533.36  GGAATTTGAATCCGGGCCCCCACTCTGCACCAGCCCCGCACTCACAGTCATCCTGCTGTA 9879
human.FAM30B-LOC105372935-GGT2.NCBI.                                            ggaatttgaatccgggcccccactctgcaccagccctgcactcacagtcatccggctgtg 6295
human.FAM230E-LOC105377182-GGT3P.NCBI.                                          ggaatttgaatccgggcccccactctgcaccagccctgcactcacagtcatccggctgtg 5370
human.FAM230J-LOC105372942-GGTLC5P.NCBI.rev.compl                               ggaatttgaatccgggcccccactctgcaccagccctgcactcacagtcatccggctgtg 5739
human.FAM230A.ncRNA.18487127.18500594-GGTLC3.18516335.18518165.rev.compl.REF    ------------------------------------------------------------ 0

chimp.LOC112206744-LOC107973052-GGT2.revcompl                                   ttctcctctccaggaaggcactgcccacgcagtctgtctgatagaggtgttgagtgctca 8380
chimp54546-135457.revcompl.Pan.troglodytes.clone.rp43-41g5.GenBank:AC099533.36  TTCTCCTCTCCAGGAAGGCACTGCCCATGCAGTCTGTCTGATAGAGGTGTTGAGTGCTCA 9939
human.FAM30B-LOC105372935-GGT2.NCBI.                                            ttctcctctccaggaaggcactgcccacgcagtctgtctgatagaggtgttgagtgctca 6355
human.FAM230E-LOC105377182-GGT3P.NCBI.                                          ttcccctctccaggaaggcactgcccacgcagtctgtctgatagaggtgttgagtgctca 5430
human.FAM230J-LOC105372942-GGTLC5P.NCBI.rev.compl                               ttcccctctccaggaaggcactgcccacgcagtctgtctgatagaggtgttgagtgctca 5799
human.FAM230A.ncRNA.18487127.18500594-GGTLC3.18516335.18518165.rev.compl.REF    ------------------------------------------------------------ 0

chimp.LOC112206744-LOC107973052-GGT2.revcompl                                   ctgaactccgtgatcttcctgaaacccaactttgattcagtgggttctgcttggaagcct 8440
chimp54546-135457.revcompl.Pan.troglodytes.clone.rp43-41g5.GenBank:AC099533.36  CTGAACTCCGTGATCTTCCTGAAACCCAACTTTGATTCAGTGGGCTCTGCTTGGAAGCCT 9999
human.FAM30B-LOC105372935-GGT2.NCBI.                                            ctgaactccgtgatcttcctgaaacccaactttgattcagtgggctctgcttggaagcct 6415
human.FAM230E-LOC105377182-GGT3P.NCBI.                                          ctgaactccgtgatcttcctgaaacccaactttgattcagtgggctctgcttggaagcct 5490
human.FAM230J-LOC105372942-GGTLC5P.NCBI.rev.compl                               ctgaactccgtgatcttcctgaaacccaactttgattcagtgggctctgcttggaagcct 5859
human.FAM230A.ncRNA.18487127.18500594-GGTLC3.18516335.18518165.rev.compl.REF    ------------------------------------------------------------ 0

chimp.LOC112206744-LOC107973052-GGT2.revcompl                                   gtaaagaaaaggatcataagtttaaacttagaacagattatcactattttccctctggtc 8500
chimp54546-135457.revcompl.Pan.troglodytes.clone.rp43-41g5.GenBank:AC099533.36  GTAAAGAAAAGGATCATAAGTTTAAACTTAGAACAGATTATCACTATTTTCCCTCTGGTC 10059
human.FAM30B-LOC105372935-GGT2.NCBI.                                            gtaaagaaaaggatcataagtttaaacttagaacagattatcactattttccctctggtc 6475
human.FAM230E-LOC105377182-GGT3P.NCBI.                                          gtaaagaaaaggatcataagtttaaacttagaacagattatcactattttccctctggcc 5550
human.FAM230J-LOC105372942-GGTLC5P.NCBI.rev.compl                               gtaaa-aaaaggatcataagtttaaacttagaacagattatcactattttccctctggtc 5918
human.FAM230A.ncRNA.18487127.18500594-GGTLC3.18516335.18518165.rev.compl.REF    ------------------------------------------------------------ 0

chimp.LOC112206744-LOC107973052-GGT2.revcompl                                   ttctgtcagcaagatgtcaacagccctatctattgtaaatgcattaaccagcatcttctc 8560
chimp54546-135457.revcompl.Pan.troglodytes.clone.rp43-41g5.GenBank:AC099533.36  TTCTGTCAGCAAGATGTCAACAGCCCTATCTATTGTAAATGCATTAACCAGCATCTTCTC 10119
human.FAM30B-LOC105372935-GGT2.NCBI.                                            ttctgtcagcaagatgtcaacagccctatctattgtcaatgcattaaccagcatcttctc 6535
human.FAM230E-LOC105377182-GGT3P.NCBI.                                          ttctgtcagcaagatgtcaacagccctatctattgtcaatgcattaaccagcgtcttctc 5610
human.FAM230J-LOC105372942-GGTLC5P.NCBI.rev.compl                               ttctgtcagcaagatgtcaacagccctatctattgtcaatgcattaaccagcatcttctc 5978
human.FAM230A.ncRNA.18487127.18500594-GGTLC3.18516335.18518165.rev.compl.REF    ------------------------------------------------------------ 0

chimp.LOC112206744-LOC107973052-GGT2.revcompl                                   tgatagagaatataagaagatatgctgtgcacaccaaccagtgttggagacctcatggct 8620
chimp54546-135457.revcompl.Pan.troglodytes.clone.rp43-41g5.GenBank:AC099533.36  TGATAGAGAATACAAGAAGATATGCTGTGCACACCAACCAGTGTCGGAGACCTCATGGCT 10179
human.FAM30B-LOC105372935-GGT2.NCBI.                                            tgatagagaatacaagaagatatgctgtgcacaccaaccagtgtaggagacctcatggct 6595
human.FAM230E-LOC105377182-GGT3P.NCBI.                                          tgatagagaatacaagaagatatgctgtgcacaccaaccagtgtaggagacctcatggct 5670
human.FAM230J-LOC105372942-GGTLC5P.NCBI.rev.compl                               tgatagagaatacaagaagatatgctgtgcacaccaaccagtgtaggagacctcatggct 6038
human.FAM230A.ncRNA.18487127.18500594-GGTLC3.18516335.18518165.rev.compl.REF    ------------------------------------------------------------ 0

chimp.LOC112206744-LOC107973052-GGT2.revcompl                                   cccgggtaaagaagaagaggtacccacaagaaggtactgtggaagttcattaattaagtt 8680
chimp54546-135457.revcompl.Pan.troglodytes.clone.rp43-41g5.GenBank:AC099533.36  CCCGGGTAAAGAAGAAGAGGTACCCACAAGAAGGTACTGTGGAAGTTCATTAATTAAGTT 10239
human.FAM30B-LOC105372935-GGT2.NCBI.                                            cccgggtaaagaagaagaggtacccacaagaaggtactgtggaagttcattaattaagtt 6655
human.FAM230E-LOC105377182-GGT3P.NCBI.                                          cccgggtaaagaagaagaggtacccacaagaaggtactgtggaagttcattaattaagtt 5730
human.FAM230J-LOC105372942-GGTLC5P.NCBI.rev.compl                               cccgggtaaagaagaagaggtacccacaagaaggtactgtggaagttcattaattaagtt 6098
human.FAM230A.ncRNA.18487127.18500594-GGTLC3.18516335.18518165.rev.compl.REF    ------------------------------------------------------------ 0

chimp.LOC112206744-LOC107973052-GGT2.revcompl                                   gattcaagaattgcaattgcggggagtattcagtgtcccatatgtaagaggaaactatga 8740
chimp54546-135457.revcompl.Pan.troglodytes.clone.rp43-41g5.GenBank:AC099533.36  GATTCAAGAATTGCAATTGCGGGGAGTATTCAGTGTCCCATATGTAAGAGGAAACTATGA 10299
human.FAM30B-LOC105372935-GGT2.NCBI.                                            gattcaagaattgcagttgcggggagtattcagtgtcccatatgtaagaggaaactatga 6715
human.FAM230E-LOC105377182-GGT3P.NCBI.                                          gattcaagaattgcagttgcggggagtattcagtgtcccatatgtaagaggaaactatga 5790
human.FAM230J-LOC105372942-GGTLC5P.NCBI.rev.compl                               gattcaagaattgcagttgcggggagtattcagtgtcccatatgtaagaggaaactatga 6158
human.FAM230A.ncRNA.18487127.18500594-GGTLC3.18516335.18518165.rev.compl.REF    ------------------------------------------------------------ 0

chimp.LOC112206744-LOC107973052-GGT2.revcompl                                   agagactaagccatatttttttaatgtgtcaggattctaatttgcctggtcagtaaatatt 8800
```

```
chimp54546-135457.revcompl.Pan.troglodytes.clone.rp43-41g5.GenBank:AC099533.36      AGAGACTAAGCCATATTTTTTAATGTGTCAGGATTCAAATTTGCCTGGTCAGTAAATATT 10359
human.FAM30B-LOC105372935-GGT2.NCBI.                                                agagactaagccatattttttaatgtgtcaggattctaatttgcctggtcagtaaatatt 6775
human.FAM230E-LOC105377182-GGT3P.NCBI.                                              agagactaagccatattttttaatgtgtcaggattctaatttgcctggtcagtaaatatt 5850
human.FAM230J-LOC105372942-GGTLC5P.NCBI.rev.compl                                   agagactaagccatattttttaatgtgtcaggattctaatttgcctggtcagtaaatatt 6218
human.FAM230A.ncRNA.18487127.18500594-GGTLC3.18516335.18518165.rev.compl.REF        ------------------------------------------------------------ 0

chimp.LOC112206744-LOC107973052-GGT2.revcompl                                       gctaccaccacaaaagtaaatatctacttaaaagtcaattttggttcatgttgaatgata 8860
chimp54546-135457.revcompl.Pan.troglodytes.clone.rp43-41g5.GenBank:AC099533.36      GCTACCACCACAAAAGTAAATATCTACTTAAAAGACAATTTTGGTTCATGTTTAATGATA 10419
human.FAM30B-LOC105372935-GGT2.NCBI.                                                gctaccaccacaaaagtaaatatctacttaaaagtcaattttggttcatgtttaatgata 6835
human.FAM230E-LOC105377182-GGT3P.NCBI.                                              gctaccaccacaaaagtaaatatctacttaaaagtcaattttggttcatgtttaatgata 5910
human.FAM230J-LOC105372942-GGTLC5P.NCBI.rev.compl                                   gctaccaccacaaaagtaaatatctacttaaaagtcaattttggttcatgtttaatgata 6278
human.FAM230A.ncRNA.18487127.18500594-GGTLC3.18516335.18518165.rev.compl.REF        ------------------------------------------------------------ 0

chimp.LOC112206744-LOC107973052-GGT2.revcompl                                       gacaatgtttcaagctaatgtctagaacttacctggttgttaaacataagcatagatctc 8920
chimp54546-135457.revcompl.Pan.troglodytes.clone.rp43-41g5.GenBank:AC099533.36      GACAATGTTTCAAGCTAATGTCTAGAACTTACCTGGTTGTTAAACATAAGCATAGATCTC 10479
human.FAM30B-LOC105372935-GGT2.NCBI.                                                gacaatgtttcaagctaatgtctagaacttacctggttgttaaacataagcatagatctc 6895
human.FAM230E-LOC105377182-GGT3P.NCBI.                                              gacaatgtttcaagctaatgtctagaacttacctggttgttaaacaaaagcatagatctc 5970
human.FAM230J-LOC105372942-GGTLC5P.NCBI.rev.compl                                   gacaatgcttcaagctaatgtctagaacttacctggttgttaaacataagcatagatctc 6338
human.FAM230A.ncRNA.18487127.18500594-GGTLC3.18516335.18518165.rev.compl.REF        ------------------------------------------------------------ 0

chimp.LOC112206744-LOC107973052-GGT2.revcompl                                       cctgaaagagtggtgctatattattatttttcaattaatatatttctttagagagtttta 8980
chimp54546-135457.revcompl.Pan.troglodytes.clone.rp43-41g5.GenBank:AC099533.36      CCTGAAAGAGTGGTGCTATATTATTATTTTTCAATTAATATATTTCTTTAGAGAGTTTTA 10539
human.FAM30B-LOC105372935-GGT2.NCBI.                                                cctgaaagagtggtgctatattattatttttcaattaatatatttctttagagagtttta 6955
human.FAM230E-LOC105377182-GGT3P.NCBI.                                              cctgaaagagtggtgctatattattatttttcaattaatatatttctttagagaatttta 6030
human.FAM230J-LOC105372942-GGTLC5P.NCBI.rev.compl                                   cctgaaagagtggtgctatattattatttttcaattaatatatttctttagagagtttta 6398
human.FAM230A.ncRNA.18487127.18500594-GGTLC3.18516335.18518165.rev.compl.REF        ------------------------------------------------------------ 0

chimp.LOC112206744-LOC107973052-GGT2.revcompl                                       aattgacataaaaactgagcatatggccgggcgtggtggctcacacttataatcccagca 9040
chimp54546-135457.revcompl.Pan.troglodytes.clone.rp43-41g5.GenBank:AC099533.36      AATTGACATAAAAACTGAGCATATGGCCGGGCGTGGTGGCTCACACTTATAATCCCAGCA 10599
human.FAM30B-LOC105372935-GGT2.NCBI.                                                aattgacataaaaactgagcatatggccgggcgtggtggctcacacttataatcccagca 7015
human.FAM230E-LOC105377182-GGT3P.NCBI.                                              aattgacataaaaactgagcatatggccgggtgtggtggctcacacttataatcccagca 6090
human.FAM230J-LOC105372942-GGTLC5P.NCBI.rev.compl                                   aattgacataaaaactgagcatatggccgtgtgtggtggctcacacttataatcccagca 6458
human.FAM230A.ncRNA.18487127.18500594-GGTLC3.18516335.18518165.rev.compl.REF        ------------------------------------------------------------ 0

chimp.LOC112206744-LOC107973052-GGT2.revcompl                                       ctttaggaggccaaggcaggcggatcatctgaggtcaagagttggagaccagcctggcca 9100
chimp54546-135457.revcompl.Pan.troglodytes.clone.rp43-41g5.GenBank:AC099533.36      CTTTAGGAGGCCAAGGCAGGCGGATCATCTGAGGTCAGGAGTTGGAGACCAGCCTGGCCA 10659
human.FAM30B-LOC105372935-GGT2.NCBI.                                                ctttaggaggccaaggcaggcggatcatctgaggtcaggagttggagaccagcctggcca 7075
human.FAM230E-LOC105377182-GGT3P.NCBI.                                              ctttaggaggccaaggcaggcggatcatctgaggtcaggagttggagaccagcctggcca 6150
human.FAM230J-LOC105372942-GGTLC5P.NCBI.rev.compl                                   ctttaggaggccaaggcaggcggatcatctgaggtcaggagttggagaccagcctggcca 6518
human.FAM230A.ncRNA.18487127.18500594-GGTLC3.18516335.18518165.rev.compl.REF        ------------------------------------------------------------ 0

chimp.LOC112206744-LOC107973052-GGT2.revcompl                                       acatggtgaaaccccatctctactaaaaatacaaaaaaattagccgggtgtggtggcagg 9160
chimp54546-135457.revcompl.Pan.troglodytes.clone.rp43-41g5.GenBank:AC099533.36      ACATGGTGAAACCCCATCTCTACTAAAAATACAAAAAAATTAGCCGGGTGTGGTGGCAGG 10719
human.FAM30B-LOC105372935-GGT2.NCBI.                                                acatggtgaaaccccatctctactaaaaatccaaaaaaattagccgggtgtggtggcagg 7135
human.FAM230E-LOC105377182-GGT3P.NCBI.                                              acatggtgaaaccccatctctactaaaaatacaaaaaaattagccgggtgtggtggcagg 6210
human.FAM230J-LOC105372942-GGTLC5P.NCBI.rev.compl                                   acatggtgaaaccccatctctactaaaaatccaaaaaaattagccgggtgtggtggcagg 6578
human.FAM230A.ncRNA.18487127.18500594-GGTLC3.18516335.18518165.rev.compl.REF        ------------------------------------------------------------ 0

chimp.LOC112206744-LOC107973052-GGT2.revcompl                                       tgcctgtaatcccagctactcaggaggctgaggcaggagaatcgcttgaacccaggaggc 9220
chimp54546-135457.revcompl.Pan.troglodytes.clone.rp43-41g5.GenBank:AC099533.36      TGCCTGTAATCCCAGCTACTCAGGAGGCTGAGGCAGGAGAATCGCTTGAACCCAGGAGGC 10779
human.FAM30B-LOC105372935-GGT2.NCBI.                                                cgcctgtaatcccagctactcaggaggctgaggcaggagaatcgcttgaacccaggaggc 7195
human.FAM230E-LOC105377182-GGT3P.NCBI.                                              cgcctgtaatcccagctactcaggaggctgaggcaggagaatcgcttgaacccaggaggc 6270
human.FAM230J-LOC105372942-GGTLC5P.NCBI.rev.compl                                   cgcctgtaatcccagctactcagaaggctgaggcaggagaatcgcttgaacccgggaggc 6638
human.FAM230A.ncRNA.18487127.18500594-GGTLC3.18516335.18518165.rev.compl.REF        ------------------------------------------------------------ 0

chimp.LOC112206744-LOC107973052-GGT2.revcompl                                       agaggttgcagtgagccaagatcatgccattgcactccagcctgggtgacaagagtgaaa 9280
chimp54546-135457.revcompl.Pan.troglodytes.clone.rp43-41g5.GenBank:AC099533.36      AGAGGTTGCAGTGAGCCAAGATCATGCCATTGCACTCCAGCCTGGGTGACAAGAGTGAAA 10839
human.FAM30B-LOC105372935-GGT2.NCBI.                                                agaggttgcagtgagccaagatcatgccattgcactccagcctgggtgacaagagtgaaa 7255
human.FAM230E-LOC105377182-GGT3P.NCBI.                                              agaggttgcagtgagccaagatcatgccattgcactccagcctgggtgac--gagtgaaa 6328
human.FAM230J-LOC105372942-GGTLC5P.NCBI.rev.compl                                   agaggttgcagtgagccaagatcatgccattgcactccagcctgggtgacaagagtgaaa 6698
human.FAM230A.ncRNA.18487127.18500594-GGTLC3.18516335.18518165.rev.compl.REF        ------------------------------------------------------------ 0

chimp.LOC112206744-LOC107973052-GGT2.revcompl                                       ctccatctca----aaataaataaataaataaataaataaataaataaaaattgagtata 9336
chimp54546-135457.revcompl.Pan.troglodytes.clone.rp43-41g5.GenBank:AC099533.36      CTCCATCTCA--------------------AAATAAATAAATAAATAAAAATTGAGTATA 10879
human.FAM30B-LOC105372935-GGT2.NCBI.                                                ctccatctca------------aaataaataaataaataaataaataaaaattgagtata 7303
human.FAM230E-LOC105377182-GGT3P.NCBI.                                              ctccatctca------------aaataaataaataaataaataaataaaaattgactata 6376
human.FAM230J-LOC105372942-GGTLC5P.NCBI.rev.compl                                   ctccatctcaaaataaataaataaataaataaataaataaataaataaaaattgagtata 6758
human.FAM230A.ncRNA.18487127.18500594-GGTLC3.18516335.18518165.rev.compl.REF        ------------------------------------------------------------ 0

chimp.LOC112206744-LOC107973052-GGT2.revcompl                                       taatatacgaagttcccatattattctgtctcctcaccctcacttcctaatttcacctat 9396
chimp54546-135457.revcompl.Pan.troglodytes.clone.rp43-41g5.GenBank:AC099533.36      TAATACACGAAGTTCCCATATTATTCTGTCTCCTCACCCTCACTTCCTAATTTCACCTAT 10939
human.FAM30B-LOC105372935-GGT2.NCBI.                                                taatacacgaagttcccatattattctgtctcctcaccctcacttcctaatttcacctat 7363
human.FAM230E-LOC105377182-GGT3P.NCBI.                                              taatacacgaagttcccatattattctgactcctcaccctcacttcctaatttcacctat 6436
human.FAM230J-LOC105372942-GGTLC5P.NCBI.rev.compl                                   taatacacgaagttcccatattattctgtctcctcaccctcacttcctaatttcacctat 6818
human.FAM230A.ncRNA.18487127.18500594-GGTLC3.18516335.18518165.rev.compl.REF        ------------------------------------------------------------ 0

chimp.LOC112206744-LOC107973052-GGT2.revcompl                                       tagtaacatcttacattactgtggtacatttgctagaataatgagaaaatattgacacat 9456
chimp54546-135457.revcompl.Pan.troglodytes.clone.rp43-41g5.GenBank:AC099533.36      TAGTAACATCTTACATTACTGTGGTACATTTGCTAGAATAATGAGAAAATATTGACACAT 10999
human.FAM30B-LOC105372935-GGT2.NCBI.                                                tagtaacatcttacattactgtggtacatttgctagaataatgagaaaatattgacacat 7423
human.FAM230E-LOC105377182-GGT3P.NCBI.                                              tagtaacatcttacattactgtggtacatttgctagaataatgagaacatattgatacat 6496
human.FAM230J-LOC105372942-GGTLC5P.NCBI.rev.compl                                   tagtaacatcttacattactgtggtacatttgctagaataatgagaaaatattgacacat 6878
human.FAM230A.ncRNA.18487127.18500594-GGTLC3.18516335.18518165.rev.compl.REF        ------------------------------------------------------------ 0

chimp.LOC112206744-LOC107973052-GGT2.revcompl                                       tattatctgaagtctacatttgcataatgttcattctttctgttatacatatatatgaat 9516
chimp54546-135457.revcompl.Pan.troglodytes.clone.rp43-41g5.GenBank:AC099533.36      TATTATCTGAAGTCTACATTTGCATAATGTTCATTCTTTCTGTTATACATATATATGAAT 11059
human.FAM30B-LOC105372935-GGT2.NCBI.                                                tattatctgaagtctacatttgcataatgttcattctttctgttatacatatatatgaat 7483
human.FAM230E-LOC105377182-GGT3P.NCBI.                                              tattatctgaagtctacatttgcataatgttcattctttctgttatacatatatatgaat 6556
human.FAM230J-LOC105372942-GGTLC5P.NCBI.rev.compl                                   tattatctgaagtctgcatttgcataatgttcattctttctgttatacatatatatgaat 6938
human.FAM230A.ncRNA.18487127.18500594-GGTLC3.18516335.18518165.rev.compl.REF        ------------------------------------------------------------ 0

chimp.LOC112206744-LOC107973052-GGT2.revcompl                                       tttgaaatatttaaaacattatgttcacccttatggtctcataaagaaaatgttcacttc 9576
chimp54546-135457.revcompl.Pan.troglodytes.clone.rp43-41g5.GenBank:AC099533.36      TTTGAAATATTTAAAACATTATGTTCACCCTTATGGTCTCATAAAGAAAATGTTCACTTC 11119
human.FAM30B-LOC105372935-GGT2.NCBI.                                                tttgaaatatttaaaacattatgttcacccttatggtctcataaagaaaatgttcacttc 7543
human.FAM230E-LOC105377182-GGT3P.NCBI.                                              tttgaaatatttaaaacattatgttcacccttatggtctcataaagaaaatgttcacttc 6616
human.FAM230J-LOC105372942-GGTLC5P.NCBI.rev.compl                                   tttgaaatatttaaaacattatgttcacccttatggtctcataaagaaaatgttcacttc 6998
human.FAM230A.ncRNA.18487127.18500594-GGTLC3.18516335.18518165.rev.compl.REF        ------------------------------------------------------------ 0

chimp.LOC112206744-LOC107973052-GGT2.revcompl                                       cctaaaaatcctctcttctcattaatctctgtcctctttctccagaaaccttggcaacta 9636
chimp54546-135457.revcompl.Pan.troglodytes.clone.rp43-41g5.GenBank:AC099533.36      CCTAAAAATCCTCTCTTCTCATTAATCTCTGTCCTCTTTCTCCAGAAACCTTGGCAACTA 11179
human.FAM30B-LOC105372935-GGT2.NCBI.                                                cctaaaaatcctctcttctcattaatctctgtcctctttctccagaaaccttggcaacta 7603
human.FAM230E-LOC105377182-GGT3P.NCBI.                                              cctaaaaatcctctcttctcattaatctctgtcctctttctccagaaaccttggcaacta 6676
human.FAM230J-LOC105372942-GGTLC5P.NCBI.rev.compl                                   cctaaaaatcctctcttctcattaatctctgtcctctttctccagaaaccttggcaacta 7058
human.FAM230A.ncRNA.18487127.18500594-GGTLC3.18516335.18518165.rev.compl.REF        ------------------------------------------------------------ 0

chimp.LOC112206744-LOC107973052-GGT2.revcompl                                       ttaatatttttactatcacttcagctttgccttttccagaatgtcatatagttggaatca 9696
chimp54546-135457.revcompl.Pan.troglodytes.clone.rp43-41g5.GenBank:AC099533.36      TTAATATTTTTACTACCGCTTCAGCTTTGCCTTTTCCAGAATGTCATATAGTTGGAATCA 11239
human.FAM30B-LOC105372935-GGT2.NCBI.                                                ttaatatttttactatcgcttcagctttgccttttccagaatgtcatatagttggaatca 7663
human.FAM230E-LOC105377182-GGT3P.NCBI.                                              ttaatatttttactatcgcttcagctttgccttttccagaatgtcatatagttggaatca 6736
human.FAM230J-LOC105372942-GGTLC5P.NCBI.rev.compl                                   ttaatatttttactatcgcttcagctttgccttttccagaatgtcatatagttggaatca 7118
human.FAM230A.ncRNA.18487127.18500594-GGTLC3.18516335.18518165.rev.compl.REF        ------------------------------------------------------------ 0

chimp.LOC112206744-LOC107973052-GGT2.revcompl                                       tatattatgtagtttttcagatgaatttattgcactaaattgatgtacgctttagctgc 9756
chimp54546-135457.revcompl.Pan.troglodytes.clone.rp43-41g5.GenBank:AC099533.36      TATATTATGTAGTTTTTCAGATGAATTTATTGCACTAAATTGATGTACGCTTTAGCTGC 11299
human.FAM30B-LOC105372935-GGT2.NCBI.                                                tatattatgtagtttttcagatgaatttattgcactaaattgatgtacgctttagctgc 7723
human.FAM230E-LOC105377182-GGT3P.NCBI.                                              tatattatgtagtttttcagatgaatttattgcactaaattgatgtatgctttagctgc 6796
human.FAM230J-LOC105372942-GGTLC5P.NCBI.rev.compl                                   tatattatgtagtttttcagatgaatttattgcactaaattgatgtacgctttagctgc 7178
human.FAM230A.ncRNA.18487127.18500594-GGTLC3.18516335.18518165.rev.compl.REF        ------------------------------------------------------------ 0

chimp.LOC112206744-LOC107973052-GGT2.revcompl                                       tttcatgtctttttttatgccttaatggcaaaaaatggcacattaaatcaccaaatatat 9816
chimp54546-135457.revcompl.Pan.troglodytes.clone.rp43-41g5.GenBank:AC099533.36      TTTCATGTCTTTTTTATGCCTTAATGGCAAAAAATGGCACATTAAATCACCAAATATAT 11359
human.FAM30B-LOC105372935-GGT2.NCBI.                                                tttcatgtctttttttatgccttaatggcaaaaaatggcacattaaatcaccaaatatat 7783
human.FAM230E-LOC105377182-GGT3P.NCBI.                                              tttcatgtctttttttatgccttaatggcaaaaaatggcacattaaatcaccaaatatat 6856
human.FAM230J-LOC105372942-GGTLC5P.NCBI.rev.compl                                   tttcatgtctttttttatgccttaatggcaaaaaatggcacattaaatcaccaaatatat 7238
human.FAM230A.ncRNA.18487127.18500594-GGTLC3.18516335.18518165.rev.compl.REF        ------------------------------------------------------------ 0

chimp.LOC112206744-LOC107973052-GGT2.revcompl                                       tgcattaaatgaatttttgtcttttattcacctgttgaagaattcggtagatttcatga 9876
chimp54546-135457.revcompl.Pan.troglodytes.clone.rp43-41g5.GenBank:AC099533.36      TGCATTAAAAGAATTTTTGTCTTTTTATTCACCTGTTGAAGAATTCGGTAGATTTCATGA 11419
```

```
human.FAM30B-LOC105372935-GGT2.NCBI.                                              tgcattcaatgaatttttgtctttttattcacctgttgaagaattcggtagatttcatga 7843
human.FAM230E-LOC105377182-GGT3P.NCBI.                                            tgcattaaatgaattttgtctttttattcacctgttgaagaattcggtagatttcatga 6916
human.FAM230J-LOC105372942-GGTLC5P.NCBI.rev.compl                                 tgcattaaatgaattttgtctttttattcacctgttgaagaattcggtagatttcatga 7298
human.FAM230A.ncRNA.18487127.18500594-GGTLC3.18516335.18518165.rev.compl.REF      ------------------------------------------------------------ 0

chimp.LOC112206744-LOC107973052-GGT2.revcompl                                     gagaaaccatctgggcctggtgctttctttttcagaatgctcttaatgtgaattcaactt 9936
chimp54546-135457.revcompl.Pan.troglodytes.clone.rp43-41g5.GenBank:AC099533.36    GAGAAACCATCTGGGCCTGGTGCTTTCTTTTTCGGAATGCTCTTAATGTGAATTCAACTT 11479
human.FAM30B-LOC105372935-GGT2.NCBI.                                              gagaaaccatctgggcctggtgctttctttttcggaatgctcttaatgtgaattcaactt 7903
human.FAM230E-LOC105377182-GGT3P.NCBI.                                            gagaaaccatctgggcctggtgctttctttttcggaatgctcttaatgtgaattcaactt 6976
human.FAM230J-LOC105372942-GGTLC5P.NCBI.rev.compl                                 gagaaaccatctgggcctggtgctttctttttcggaatgctcttaatgtgaattcaactt 7358
human.FAM230A.ncRNA.18487127.18500594-GGTLC3.18516335.18518165.rev.compl.REF      ------------------------------------------------------------ 0

chimp.LOC112206744-LOC107973052-GGT2.revcompl                                     atttaatagacataagtttattcaaattaggatcctagcatgaccttgggaagattgcct 9996
chimp54546-135457.revcompl.Pan.troglodytes.clone.rp43-41g5.GenBank:AC099533.36    ATTTAATAGACATAAGTTTATTCAAATTAGGATCCTAGCATGACCTTGGGAAGATTGCCT 11539
human.FAM30B-LOC105372935-GGT2.NCBI.                                              atttaatagacataagtttattcaaattaggatcctagcatgaccttgggaagattgcct 7963
human.FAM230E-LOC105377182-GGT3P.NCBI.                                            atttaatagacataagtttattcaaattaggattctagcgtgaccttgggaagattgcct 7036
human.FAM230J-LOC105372942-GGTLC5P.NCBI.rev.compl                                 atttaatagacataagtttattcaaattaggatcctagcatgaccttgggaagattgcct 7418
human.FAM230A.ncRNA.18487127.18500594-GGTLC3.18516335.18518165.rev.compl.REF      ------------------------------------------------------------ 0

chimp.LOC112206744-LOC107973052-GGT2.revcompl                                     ttcaaggaattgatacatttcactgaggttatcaaactgcggtcatagaactgttcatga 10056
chimp54546-135457.revcompl.Pan.troglodytes.clone.rp43-41g5.GenBank:AC099533.36    TTCAAGGAATTGATACATTTCACTGAGGTTATCAAACTGCGGTCATAGAACTGTTCATGA 11599
human.FAM30B-LOC105372935-GGT2.NCBI.                                              ttcaaggaattgatacatttcactgaggttatcaaactgcggtcatagaactgttcatga 8023
human.FAM230E-LOC105377182-GGT3P.NCBI.                                            ttcaaggaattgatacatttcactgaggttatcagactgcggtcatagaactgttcataa 7096
human.FAM230J-LOC105372942-GGTLC5P.NCBI.rev.compl                                 ttcaaggaattgatacatttcactgaggttatcaaactgcgctcatagaactgttcatga 7478
human.FAM230A.ncRNA.18487127.18500594-GGTLC3.18516335.18518165.rev.compl.REF      ------------------------------------------------------------ 0

chimp.LOC112206744-LOC107973052-GGT2.revcompl                                     tattccttttaatgcctaacagttcagtagagatggctcctcttttatttctgaaattgg 10116
chimp54546-135457.revcompl.Pan.troglodytes.clone.rp43-41g5.GenBank:AC099533.36    TATTCCTTTTAATGCCTAACAGTTCAGTAGAGATGGCTCCTCTTTTATTTCTGAAATTGG 11659
human.FAM30B-LOC105372935-GGT2.NCBI.                                              tattccttttaatgcctaacagttcagtagagatggctcctcttttatttctgaaattgg 8083
human.FAM230E-LOC105377182-GGT3P.NCBI.                                            tattccttttaatgcctaacagttcagtagagatggctcctcttttatttctgaaattgg 7156
human.FAM230J-LOC105372942-GGTLC5P.NCBI.rev.compl                                 tattccttttaatgcctaacagttcagtagagatggctcctcttttatttctgaaattgg 7538
human.FAM230A.ncRNA.18487127.18500594-GGTLC3.18516335.18518165.rev.compl.REF      ------------------------------------------------------------ 0

chimp.LOC112206744-LOC107973052-GGT2.revcompl                                     tcatttgtgttatcttctttttcttggttagcctgcatatcaattcattcattgtaatga 10176
chimp54546-135457.revcompl.Pan.troglodytes.clone.rp43-41g5.GenBank:AC099533.36    TCATTTGTGTTATCTTCTTTTTCTTGGTTAGCCTGCATATCAATTCATTCATTGTAATGA 11719
human.FAM30B-LOC105372935-GGT2.NCBI.                                              tcatttgtgttatcttctttttcttggttagcctgcatatcaattcattcattgtaatga 8143
human.FAM230E-LOC105377182-GGT3P.NCBI.                                            tcatttgtgttatcttctttttcttggttagcctgcatatcaattcattcattgtaatga 7216
human.FAM230J-LOC105372942-GGTLC5P.NCBI.rev.compl                                 tcatttgtgttatcttctttttcttggttagcctgcatatcaattcattcattgtaatga 7598
human.FAM230A.ncRNA.18487127.18500594-GGTLC3.18516335.18518165.rev.compl.REF      ------------------------------------------------------------ 0

chimp.LOC112206744-LOC107973052-GGT2.revcompl                                     gcatatcaaagaaccagcttttggttttattgatttctgatggtttcagtgttttaatt 10236
chimp54546-135457.revcompl.Pan.troglodytes.clone.rp43-41g5.GenBank:AC099533.36    GCATATCAAAGAACCAGCTTTTGGTTTTATTGATTTCTGATGATTTCAGTGTTTTAATT 11779
human.FAM30B-LOC105372935-GGT2.NCBI.                                              gcatatcaaagaaccagcttttggttttattgatttctgatgatttcagtgttttaatt 8203
human.FAM230E-LOC105377182-GGT3P.NCBI.                                            gcatatcaaagaaccagcttttggttttattgatttctgatgatttcagtgttttaatt 7276
human.FAM230J-LOC105372942-GGTLC5P.NCBI.rev.compl                                 gcatatcaaagaaccagcttttggttttattgatttctgatgatttcagtgttttaatt 7658
human.FAM230A.ncRNA.18487127.18500594-GGTLC3.18516335.18518165.rev.compl.REF      ------------------------------------------------------------ 0

chimp.LOC112206744-LOC107973052-GGT2.revcompl                                     ttattgatttctgtgatgttgtttattacttttacttgctttccattgcattcctctatt 10296
chimp54546-135457.revcompl.Pan.troglodytes.clone.rp43-41g5.GenBank:AC099533.36    TTATTGATTTCTGTGATGTTGTTTATTACTTTTACTTGCTTTCCATTGCATTCCTCTATT 11839
human.FAM30B-LOC105372935-GGT2.NCBI.                                              ttattgatttctgtgatgttgtttattacttttacttgctttccattgcattcctctatt 8263
human.FAM230E-LOC105377182-GGT3P.NCBI.                                            ttattgatttctgtgatgttgtttattacttttacttgctttccattgcattcctctatt 7336
human.FAM230J-LOC105372942-GGTLC5P.NCBI.rev.compl                                 ttattgatttctgtgatgttgtttattacttttacttgctttccattgcattcctctatt 7718
human.FAM230A.ncRNA.18487127.18500594-GGTLC3.18516335.18518165.rev.compl.REF      ------------------------------------------------------------ 0

chimp.LOC112206744-LOC107973052-GGT2.revcompl                                     ttctatagttccctaattgaaacatgatattactgattttaggtcttgtgatttttagta 10356
chimp54546-135457.revcompl.Pan.troglodytes.clone.rp43-41g5.GenBank:AC099533.36    TTCTATAGTTCCCTAATTGAAACATGATATTACTGATTTTAGGTCTTGTGATTTTTAGTA 11899
human.FAM30B-LOC105372935-GGT2.NCBI.                                              ttctatagttccctaattgaaacatgatattactgattttaggtcttgtgatttttagta 8323
human.FAM230E-LOC105377182-GGT3P.NCBI.                                            ttctatagttccctaattgaaacatgatattactgattttaggtcttgtgatttttagta 7396
human.FAM230J-LOC105372942-GGTLC5P.NCBI.rev.compl                                 ttctatagttccctaattgaaacatgatattactgattttaggtcttgtgatttttagta 7778
human.FAM230A.ncRNA.18487127.18500594-GGTLC3.18516335.18518165.rev.compl.REF      ------------------------------------------------------------ 0

chimp.LOC112206744-LOC107973052-GGT2.revcompl                                     tattgcatccaatgctatagatttccctctaaggactgcttttgctacatccagaaatct 10416
chimp54546-135457.revcompl.Pan.troglodytes.clone.rp43-41g5.GenBank:AC099533.36    TATTGCATCCAATGCTATAGATTTCCCTCTAAGGACTGCTTTTGCTACATCCAGAAATCT 11959
human.FAM30B-LOC105372935-GGT2.NCBI.                                              tattgcatccaatgctatagatttccctctaaggactgcttttgctacatccagaaatct 8383
human.FAM230E-LOC105377182-GGT3P.NCBI.                                            tattgcatccaatgctgtagatttccctctaaggactgcttttgctacatccagaaatct 7456
human.FAM230J-LOC105372942-GGTLC5P.NCBI.rev.compl                                 tattgcatccaatgctatagatttccctctaaggactgcttttgctacatccagaaatct 7838
human.FAM230A.ncRNA.18487127.18500594-GGTLC3.18516335.18518165.rev.compl.REF      ------------------------------------------------------------ 0

chimp.LOC112206744-LOC107973052-GGT2.revcompl                                     tgccaagtcacatttcttttaatgtagttaaaagtatttttaattttctattgagactt 10476
chimp54546-135457.revcompl.Pan.troglodytes.clone.rp43-41g5.GenBank:AC099533.36    TGCCAAGTCACATTTCTTTTAATGTAGTTAAAAGTATTTTTAATTTTCTATTGAGACTT 12019
human.FAM30B-LOC105372935-GGT2.NCBI.                                              tgccaagtcacatttcttttaatgtagttaaaagtatttttaattttctattgagactt 8443
human.FAM230E-LOC105377182-GGT3P.NCBI.                                            tgccaagtgacatttcttttaatgtagttaaaagtatttttaattttctattgagactt 7516
human.FAM230J-LOC105372942-GGTLC5P.NCBI.rev.compl                                 tgccaagtcacatttcttttaatgtagttaaaagtacttttaattttctattgagactt 7898
human.FAM230A.ncRNA.18487127.18500594-GGTLC3.18516335.18518165.rev.compl.REF      ------------------------------------------------------------ 0

chimp.LOC112206744-LOC107973052-GGT2.revcompl                                     cttctttaacccatgagttatttaaaagtgcattgctaatttgcaaatatttggggattt 10536
chimp54546-135457.revcompl.Pan.troglodytes.clone.rp43-41g5.GenBank:AC099533.36    CTTCTTTAACCCATGAGTTATTTAAAAGTGCATTGCTAATTTGCAAATATTTGGGGATTT 12079
human.FAM30B-LOC105372935-GGT2.NCBI.                                              cttctttaacccatgagttatttaaaagtgcattgctaatttgcaaatatttggggattt 8503
human.FAM230E-LOC105377182-GGT3P.NCBI.                                            cttcttttacccatgagttatttaaaagtgcattgctaatttgcaaatatttggggattt 7576
human.FAM230J-LOC105372942-GGTLC5P.NCBI.rev.compl                                 cttctttaacccaacagttatttaaaagtgcattgctaatttgcaaatatttggggattt 7958
human.FAM230A.ncRNA.18487127.18500594-GGTLC3.18516335.18518165.rev.compl.REF      ------------------------------------------------------------ 0

chimp.LOC112206744-LOC107973052-GGT2.revcompl                                     tgtggctcttttacagttgttgatttttttgttgtcaggtatgtgttgcaaaagcagtcgt 10596
chimp54546-135457.revcompl.Pan.troglodytes.clone.rp43-41g5.GenBank:AC099533.36    TCTGGCTCTTTTACAGTTGTTGATTTTTTGTTGTCAGGTATGTGTTGCAAAAGCAGTCGT 12139
human.FAM30B-LOC105372935-GGT2.NCBI.                                              tgtggctcttttacagttgttgaattttttgttgtcaggtatgtgttgcaaaagcagtcgt 8563
human.FAM230E-LOC105377182-GGT3P.NCBI.                                            tgtggctcttttacagttgttgatttttttgttgtcaggtgtgtgttgcaaaagcagtcgt 7636
human.FAM230J-LOC105372942-GGTLC5P.NCBI.rev.compl                                 tgtggctcttttacagttgttgatttttttgttgtcaggtgtgtgttgcaaaagcagtcgt 8018
human.FAM230A.ncRNA.18487127.18500594-GGTLC3.18516335.18518165.rev.compl.REF      ------------------------------------------------------------ 0

chimp.LOC112206744-LOC107973052-GGT2.revcompl                                     ctacctcatcttgccaccagccaagatggcccaggatgtgggctctccctgagtgaatct 10656
chimp54546-135457.revcompl.Pan.troglodytes.clone.rp43-41g5.GenBank:AC099533.36    CTACCTCATCTTGCCACCAGCCAAGATGGCCCAGGATGTGGGCTCTCCCTGAGTGAATCT 12199
human.FAM30B-LOC105372935-GGT2.NCBI.                                              ctacctcatcttgccaccacccaagatggcccaggatgtgggctctccctgagtgaatct 8623
human.FAM230E-LOC105377182-GGT3P.NCBI.                                            ctacctcatcttgccaccacccaagatggcccaggatgtgggctctccctgagtgaatct 7696
human.FAM230J-LOC105372942-GGTLC5P.NCBI.rev.compl                                 ctacctcatcttgccaccacccaagatggcccaggatgtgggctctccctgagtgaatct 8078
human.FAM230A.ncRNA.18487127.18500594-GGTLC3.18516335.18518165.rev.compl.REF      ------------------------------------------------------------ 0

chimp.LOC112206744-LOC107973052-GGT2.revcompl                                     ttggcaatctgccaacctgatgtgttcggcctccttctttagtctgagtttcccttctgc 10716
chimp54546-135457.revcompl.Pan.troglodytes.clone.rp43-41g5.GenBank:AC099533.36    TTGGCAATCTGCCAACCTGATGTGTTCGGCCTCCTTCTTTAGTCTGAGTTTGCCTTCTGC 12259
human.FAM30B-LOC105372935-GGT2.NCBI.                                              ttggcaatctgccaacctgatgtgttcggcctccttctttagtctgagcttgccttctgc 8683
human.FAM230E-LOC105377182-GGT3P.NCBI.                                            ttggcaatctgccaacctgatgtgttcggcctccttctttagtctgagcttgccttctgc 7756
human.FAM230J-LOC105372942-GGTLC5P.NCBI.rev.compl                                 ttggcaatctgccaacctgatgtgttcggcctccttctttagtctgagcttgccttctgc 8138
human.FAM230A.ncRNA.18487127.18500594-GGTLC3.18516335.18518165.rev.compl.REF      ------------------------------------------------------------ 0

chimp.LOC112206744-LOC107973052-GGT2.revcompl                                     ttagaaagggccattctcagttctggcagggagttttcccaacattgagaaggtggcatt 10776
chimp54546-135457.revcompl.Pan.troglodytes.clone.rp43-41g5.GenBank:AC099533.36    TTAGAAAGGGCCATTCTCAGTTCTGGCAGGGAGTTTTCCCAACATTGAGAAGGTGGCATT 12319
human.FAM30B-LOC105372935-GGT2.NCBI.                                              ttagaaagggccattctcagttctggcagggagttttcccaacattgagaaggtggcatt 8743
human.FAM230E-LOC105377182-GGT3P.NCBI.                                            ttagaaagggccattctcagttctggcagggagttttcccaacattgagaaggtggcatt 7816
human.FAM230J-LOC105372942-GGTLC5P.NCBI.rev.compl                                 ttagaaagggccattctcagttctggcagggagttttcccaacattgagaaggtggcatt 8198
human.FAM230A.ncRNA.18487127.18500594-GGTLC3.18516335.18518165.rev.compl.REF      ------------------------------------------------------------ 0

chimp.LOC112206744-LOC107973052-GGT2.revcompl                                     cttactccccactgcagcctgcacctctgaccggtggtcagcagacaggacagaggtcct 10836
chimp54546-135457.revcompl.Pan.troglodytes.clone.rp43-41g5.GenBank:AC099533.36    CTTACTCCCCACTACAGCCTGCACCTCTGACCGGTGGTCAGCAGACAGGACAGAGGTCCT 12379
human.FAM30B-LOC105372935-GGT2.NCBI.                                              cttactccccactgcagcctgcacctctgaccggtggtcagcagacaggacagaggtcct 8803
human.FAM230E-LOC105377182-GGT3P.NCBI.                                            cttactccccactgcagcctgcacctctgaccggtggtcagcagacaggacagaggtcct 7876
human.FAM230J-LOC105372942-GGTLC5P.NCBI.rev.compl                                 cttactccccactgcagcctgcacctctgaccggtggtcagcagacaggacagaggtcct 8258
human.FAM230A.ncRNA.18487127.18500594-GGTLC3.18516335.18518165.rev.compl.REF      ------------------------------------------------------------ 0

chimp.LOC112206744-LOC107973052-GGT2.revcompl                                     cagtagacagagttcagcggggtctctgaccaaagtgcatcttcagagtctgcacctacc 10896
chimp54546-135457.revcompl.Pan.troglodytes.clone.rp43-41g5.GenBank:AC099533.36    CATTAGACAGAGTTCAGCGGGGTCTCTGACCAAAGTGCATCTTCAGAGTCTACACCTACC 12439
human.FAM30B-LOC105372935-GGT2.NCBI.                                              catcagacagagttcagcagggtctctgaccaaagtgcatcttcagagtctgcacctacc 8863
human.FAM230E-LOC105377182-GGT3P.NCBI.                                            cattagacagagttcagcagggtctctgaccaaagtgcatcttcagagtctgcacctacc 7936
human.FAM230J-LOC105372942-GGTLC5P.NCBI.rev.compl                                 cattagacagagttcagcagggtctctgaccaaagtgcatcttcagagtctgcacctacc 8318
human.FAM230A.ncRNA.18487127.18500594-GGTLC3.18516335.18518165.rev.compl.REF      ------------------------------------------------------------ 0

chimp.LOC112206744-LOC107973052-GGT2.revcompl                                     cactgtgaccacgggcaggctctgagtcctaaagcaggaggaactgtgcgaccatcctga 10956
chimp54546-135457.revcompl.Pan.troglodytes.clone.rp43-41g5.GenBank:AC099533.36    CACTGTGACCACGGGCAGGCTCTGAGTCCTAAAGCAGGAGGAGCTGTGCGACCATCCTGA 12499
human.FAM30B-LOC105372935-GGT2.NCBI.                                              cactgtgaccacgggcaggctctgagtcctaaagcaggaggaactgtgcgaccatcctga 8923
```

```
human.FAM230E-LOC105377182-GGT3P.NCBI.                                      cactgtgaccacgggcaggctctgagtcctaaagcaggaggaactgtgcgaccatcctga 7996
human.FAM230J-LOC105372942-GGTLC5P.NCBI.rev.compl                           cactgtgaccacgggcaggctctgagtcctaaagcaggaggaactgtgcgaccatcctga 8378
human.FAM230A.ncRNA.18487127.18500594-GGTLC3.18516335.18518165.rev.compl.REF ------------------------------------------------------------ 0

chimp.LOC112206744-LOC107973052-GGT2.revcompl                               ttggaaatttgtgaggatcaccgtgttactcaagtaaggtctttggaaagtgtcgtatta 11016
chimp54546-135457.revcompl.Pan.troglodytes.clone.rp43-41g5.GenBank:AC099533.36 TTGGAAATTTGTGAGGATCACCGTGTTACTCAAGTAAGGTCTTTGGAAAGTGTCGTATTA 12559
human.FAM30B-LOC105372935-GGT2.NCBI.                                        ttggaaatttgtgaggatcaccgtgttactcaagtaaggtctttggaaagtgtcgtatta 8983
human.FAM230E-LOC105377182-GGT3P.NCBI.                                      ttggaaatttgtgaggatcaccgtgttactcaagtaaggtctttggaaagtgtcgtatta 8056
human.FAM230J-LOC105372942-GGTLC5P.NCBI.rev.compl                           ttggaaatttgtgaggatcaccgtgttactcaagtaaggtctttggaaagtgtcgtatta 8438
human.FAM230A.ncRNA.18487127.18500594-GGTLC3.18516335.18518165.rev.compl.REF ------------------------------------------------------------ 0

chimp.LOC112206744-LOC107973052-GGT2.revcompl                               ctactgtttgtgaactgcttgttggtggcctggctgagccacacactttatgaaaaccag 11076
chimp54546-135457.revcompl.Pan.troglodytes.clone.rp43-41g5.GenBank:AC099533.36 CTACTGTTTGTGAACTGCTTGTTGGTGGCCTGGCTGAGCCACACACTTTATGAAAACCAG 12619
human.FAM30B-LOC105372935-GGT2.NCBI.                                        ctactgtttgtgaactgcttgttggtggcctggctgagccacacactttatgaaaaccag 9043
human.FAM230E-LOC105377182-GGT3P.NCBI.                                      ctactgtttgtgaactgcttgttggtggcctggctgagccacacactttatgaaaaccag 8116
human.FAM230J-LOC105372942-GGTLC5P.NCBI.rev.compl                           ctactgtttgtgaactgcttgttggtggcctggctgagccacacactttatgaaaaccag 8498
human.FAM230A.ncRNA.18487127.18500594-GGTLC3.18516335.18518165.rev.compl.REF ------------------------------------------------------------ 0

chimp.LOC112206744-LOC107973052-GGT2.revcompl                               gacccctcagctggtgtgggtgtctatgcagactgagaccctcatgtgaacagccttgtg 11136
chimp54546-135457.revcompl.Pan.troglodytes.clone.rp43-41g5.GenBank:AC099533.36 GACCCCTCAGCTGGTGTGGGTGTCTATGCAGCCTGAGACCCTCATGTGAACAGCCTCGTG 12679
human.FAM30B-LOC105372935-GGT2.NCBI.                                        gacccctcagctggtgtgggtgtctatgcagcctgagaccctcatgtgaacagcctcgtg 9103
human.FAM230E-LOC105377182-GGT3P.NCBI.                                      gacccctcagctggtgtgggtgtctatgcagcctgagaccctcatgtgaacagcctcgtg 8176
human.FAM230J-LOC105372942-GGTLC5P.NCBI.rev.compl                           gacccctcagctggtgtgggtgtctatgcagcctgagaccctcatgtgaacagcctcgtg 8558
human.FAM230A.ncRNA.18487127.18500594-GGTLC3.18516335.18518165.rev.compl.REF ------------------------------------------------------------ 0

chimp.LOC112206744-LOC107973052-GGT2.revcompl                               gcagctgtctttgccccttgccaccatcagtgcctccttgttcctgggcactgctttctc 11196
chimp54546-135457.revcompl.Pan.troglodytes.clone.rp43-41g5.GenBank:AC099533.36 GCAGCTGTCTTTGCCCCTTGCCACCATCAGTGCCTCCTTGTTCCTGGGCACTGCTTTCTC 12739
human.FAM30B-LOC105372935-GGT2.NCBI.                                        gcagctgtctttgccccttgccaccatcagtgcctccttgttcctgggcactgctttctc 9163
human.FAM230E-LOC105377182-GGT3P.NCBI.                                      gcagctgtctttgccccttgccaccatcagtgcctccttgttcctgggcactgctttctc 8236
human.FAM230J-LOC105372942-GGTLC5P.NCBI.rev.compl                           gcagctgtctttgccccttgccaccatcagtgcctccttgttcctgggcactgctttctc 8618
human.FAM230A.ncRNA.18487127.18500594-GGTLC3.18516335.18518165.rev.compl.REF ------------------------------------------------------------ 0

chimp.LOC112206744-LOC107973052-GGT2.revcompl                               tgatggtgctccattgttttcctgcacctcagtgtctacagctggatgtctcttctgcaa 11256
chimp54546-135457.revcompl.Pan.troglodytes.clone.rp43-41g5.GenBank:AC099533.36 TGATGGTGCTCCATTGTTTTCCTGCACCTCAGTGTCTACAGCTGGATGTCTCTTCTGCAA 12799
human.FAM30B-LOC105372935-GGT2.NCBI.                                        tgatggtgctccattgttttcctgcacctcagtgtctacagctggatgtctcttccgcaa 9223
human.FAM230E-LOC105377182-GGT3P.NCBI.                                      tgatggtgctccattgttttcctgcacctcagtgtctacagctggatgtctcttccacag 8296
human.FAM230J-LOC105372942-GGTLC5P.NCBI.rev.compl                           tgatggtgctccattgttttcctgcacctcagtgtctacagctggatgtctcttccgcaa 8678
human.FAM230A.ncRNA.18487127.18500594-GGTLC3.18516335.18518165.rev.compl.REF ------------------------------------------------------------ 0

chimp.LOC112206744-LOC107973052-GGT2.revcompl                               tctaggcgagggggcatcaatggcagttctgctgtggcactgccctccttcttagcttgt 11316
chimp54546-135457.revcompl.Pan.troglodytes.clone.rp43-41g5.GenBank:AC099533.36 TCTAGGCGAGGGGGCATCAATGGCAGTTCTGCTGTGGCACTGCCCTCCTTCTTAGCTTGT 12859
human.FAM30B-LOC105372935-GGT2.NCBI.                                        tctaggcgagggggcatcaacggcagttctgctgtggcactgccctccttcttagcttgt 9283
human.FAM230E-LOC105377182-GGT3P.NCBI.                                      tctaggcgagggggcatcaatggcagttctgctgtggcactgccctccttcttagcttgt 8356
human.FAM230J-LOC105372942-GGTLC5P.NCBI.rev.compl                           tctaggcgagggggcatcaatggcagttctgctgtggcactgccctccttcttagcttgt 8738
human.FAM230A.ncRNA.18487127.18500594-GGTLC3.18516335.18518165.rev.compl.REF ------------------------------------------------------------ 0

chimp.LOC112206744-LOC107973052-GGT2.revcompl                               cttgctctgtcttaggctccctcaaagatcccacccttcaggttcttccacaagtttctt 11376
chimp54546-135457.revcompl.Pan.troglodytes.clone.rp43-41g5.GenBank:AC099533.36 CTTGCTCTGTCTTAGGCTCCCTGAAAGATCCCACCCTTCGGGTTCTTCCACAAGTTTCTT 12919
human.FAM30B-LOC105372935-GGT2.NCBI.                                        cttgctctgtcttaggctccctcaaagatcccacccttcaggttcttccacaagtttctt 9343
human.FAM230E-LOC105377182-GGT3P.NCBI.                                      cttgctctgtcttaggctccctcaaagatcccacccttcgggttcttccacaagtttctt 8416
human.FAM230J-LOC105372942-GGTLC5P.NCBI.rev.compl                           cttgctctgtcttaggctccctcaaagatcccacccttcaggttcttccacaagtttctt 8798
human.FAM230A.ncRNA.18487127.18500594-GGTLC3.18516335.18518165.rev.compl.REF ------------------------------------------------------------ 0

chimp.LOC112206744-LOC107973052-GGT2.revcompl                               actgaaatccgagcagaaaactatgaccaatatgaccaatcctatacccactgaagacat 11436
chimp54546-135457.revcompl.Pan.troglodytes.clone.rp43-41g5.GenBank:AC099533.36 ATTGAAATCCGAGCAGAAAACTATGACCAATATGACCAATCCTATACCCACTGAAGACAT 12979
human.FAM30B-LOC105372935-GGT2.NCBI.                                        attgaaatccgagcagaaaactatgaccaatatgaccaatcctatacccactgaagacat 9403
human.FAM230E-LOC105377182-GGT3P.NCBI.                                      attgaaatccgagcagaaaactatgaccaatatgaccaatcctatacccactgaagacat 8476
human.FAM230J-LOC105372942-GGTLC5P.NCBI.rev.compl                           attgaaatccgagcagaaaactatgaccaatatgaccaatcctatacccactgaagacat 8858
human.FAM230A.ncRNA.18487127.18500594-GGTLC3.18516335.18518165.rev.compl.REF ------------------------------------------------------------ 0

chimp.LOC112206744-LOC107973052-GGT2.revcompl                               gaatgaagaattaaaacaattctccatggactctaccatatagatccctagaagtaattt 11496
chimp54546-135457.revcompl.Pan.troglodytes.clone.rp43-41g5.GenBank:AC099533.36 GAATGAAGAATTAAAACAATTCTCCATGGACTCTACCATATAGATCCCTAGAAGTAATTT 13039
human.FAM30B-LOC105372935-GGT2.NCBI.                                        gaatgaagaattaaaacaattctccatggactctaccatatagatccctagaagtaattt 9463
human.FAM230E-LOC105377182-GGT3P.NCBI.                                      gaatgaagaattaaaacaattctccatggactctaccatatagatccctagaagtaattt 8536
human.FAM230J-LOC105372942-GGTLC5P.NCBI.rev.compl                           gaatgaagaattaaaacaattctccatggactctaccatatagatccctagaagtaattt 8918
human.FAM230A.ncRNA.18487127.18500594-GGTLC3.18516335.18518165.rev.compl.REF ------------------------------------------------------------ 0

chimp.LOC112206744-LOC107973052-GGT2.revcompl                               ctaaaaaaaaaaaaaaaa----caaggaagatgtaatagttttccataaattagaatacc 11552
chimp54546-135457.revcompl.Pan.troglodytes.clone.rp43-41g5.GenBank:AC099533.36 CT------AAAAAAAAAAAAATCAAGGAAGATGTAATAGTTTTCCATAAATTAGAATACC 13093
human.FAM30B-LOC105372935-GGT2.NCBI.                                        ctaaaaaaaaaaaaaaaaaaatcaaggaagatgtaatagttttccataaattagaatacc 9523
human.FAM230E-LOC105377182-GGT3P.NCBI.                                      ct-----aaaaaaaaaaaaaatcaaggaagatgtaatagttttccataaattagaatacc 8591
human.FAM230J-LOC105372942-GGTLC5P.NCBI.rev.compl                           ct------aaaaaaaaaaaaatcaaggaaaatgtaatagttttccataaattagaatacc 8972
human.FAM230A.ncRNA.18487127.18500594-GGTLC3.18516335.18518165.rev.compl.REF ------------------------------------------------------------ 0

chimp.LOC112206744-LOC107973052-GGT2.revcompl                               ctacatgtacaattaaatgaaatggctagtatagtcttgaaaccaaaaccagataaggta 11612
chimp54546-135457.revcompl.Pan.troglodytes.clone.rp43-41g5.GenBank:AC099533.36 CTACATGTACAATTAAATGAAATGGCTAGTATAGTCTTGAAACCAAAACCAGATAAGGTA 13153
human.FAM30B-LOC105372935-GGT2.NCBI.                                        ctacatgtacaattaaatgaaatggctagtatagtcttgaaaccaaaaccagataaggta 9583
human.FAM230E-LOC105377182-GGT3P.NCBI.                                      ctacatgtacaattaaatgaaatggctagtatagtcttgaaaccaaatccagataaggta 8651
human.FAM230J-LOC105372942-GGTLC5P.NCBI.rev.compl                           ctacatgtacaattaaatgaaatggctactatagtcttgaaaccaaaaccagataaggta 9032
human.FAM230A.ncRNA.18487127.18500594-GGTLC3.18516335.18518165.rev.compl.REF ------------------------------------------------------------ 0

chimp.LOC112206744-LOC107973052-GGT2.revcompl                               aattaaattctgtgatattttaaaatactgtaaattctgactaatgtgagttaatctcaa 11672
chimp54546-135457.revcompl.Pan.troglodytes.clone.rp43-41g5.GenBank:AC099533.36 AATTAAATTCTGTGATATTTTAAAATACTGTAAATTCTGACTAATGTGAGTT------AA 13207
human.FAM30B-LOC105372935-GGT2.NCBI.                                        aattaaattctgtgatattttaaaatactgtaaattctgactaatgtgagttaatctcaa 9643
human.FAM230E-LOC105377182-GGT3P.NCBI.                                      aattaaattctgtgatattttaaaatactttaaattctgactaatgtgagttaatctcaa 8711
human.FAM230J-LOC105372942-GGTLC5P.NCBI.rev.compl                           aattaaattctgtgatattttaaaatactttaaattctgactaatgtgagttaatctcaa 9092
human.FAM230A.ncRNA.18487127.18500594-GGTLC3.18516335.18518165.rev.compl.REF ------------------------------------------------------------ 0

chimp.LOC112206744-LOC107973052-GGT2.revcompl                               tatatgaaatagtagattaacattgaaaatgcaataaataaaattagctacctcaagagt 11732
chimp54546-135457.revcompl.Pan.troglodytes.clone.rp43-41g5.GenBank:AC099533.36 TATATGAAATAGTAGATTAACATTGAAAATGCAATAAATAAAATTAGCTACCTCAAGAGT 13267
human.FAM30B-LOC105372935-GGT2.NCBI.                                        tatatgaaatagtagattaacattgaaaatgcaataaataaaattagctacctcaagagt 9703
human.FAM230E-LOC105377182-GGT3P.NCBI.                                      catatgaaatagtagattaacattgaaaatgcaataaataaaattagctacctcaagagt 8771
human.FAM230J-LOC105372942-GGTLC5P.NCBI.rev.compl                           catatgaaatagtagattaacattgaaaatgcaataaataaaattagctacctcaagagt 9152
human.FAM230A.ncRNA.18487127.18500594-GGTLC3.18516335.18518165.rev.compl.REF ------------------------------------------------------------ 0

chimp.LOC112206744-LOC107973052-GGT2.revcompl                               ttaatggaaaaaaatgtgattattgcaatagattcaggaaattcatgaataacattcacc 11792
chimp54546-135457.revcompl.Pan.troglodytes.clone.rp43-41g5.GenBank:AC099533.36 TTAATGGAAAAAAATGTGATTATTGCAATAGATTCAGGAAATTCATGAATAACATTCACC 13327
human.FAM30B-LOC105372935-GGT2.NCBI.                                        ttaatggaaaaaaatgtgattattgcaatagattcaggaaattcatgaataacattcacc 9763
human.FAM230E-LOC105377182-GGT3P.NCBI.                                      ttaatggaaaaaaatgtgattattgcaatagattcaggaaattcatgaataacattcacc 8831
human.FAM230J-LOC105372942-GGTLC5P.NCBI.rev.compl                           ttaatggaaaaaaatgtgattattgcaatagattcaggaaattcatgaataacattcacc 9212
human.FAM230A.ncRNA.18487127.18500594-GGTLC3.18516335.18518165.rev.compl.REF ------------------------------------------------------------ 0

chimp.LOC112206744-LOC107973052-GGT2.revcompl                               ctatatttgtaggacaactatctgattaactttaagtgacctgtgacaaccatttgaatt 11852
chimp54546-135457.revcompl.Pan.troglodytes.clone.rp43-41g5.GenBank:AC099533.36 CTATATTTGTAGGACAACTATCTGATTAACTTTAAGTGACCTGTGACAACCATTTGAATT 13387
human.FAM30B-LOC105372935-GGT2.NCBI.                                        ctatatttgtaggacagctatc----taactttaagtgacctgtgacaaccatttgaatt 9819
human.FAM230E-LOC105377182-GGT3P.NCBI.                                      ctatatttgtaggacaactatctgattaactttaagtgacctgtgacaaccacttgaatt 8891
human.FAM230J-LOC105372942-GGTLC5P.NCBI.rev.compl                           ctatatttgtaggacaactatctgattaactttaagtgacctgtgacaaccacttgaatt 9272
human.FAM230A.ncRNA.18487127.18500594-GGTLC3.18516335.18518165.rev.compl.REF ------------------------------------------------------------ 0

chimp.LOC112206744-LOC107973052-GGT2.revcompl                               aatgctgctttcacagcatatctcttggcttgttaaaaacccgacaagaatttccgtaac 11912
chimp54546-135457.revcompl.Pan.troglodytes.clone.rp43-41g5.GenBank:AC099533.36 AATGCTGCTTTCACAGCATATCTCTTGGCTTGTTAAAAATCCGACAAGAATTTCCGTAAC 13447
human.FAM30B-LOC105372935-GGT2.NCBI.                                        aatgctgctttcacagcatatctcttggcttgttaaaaacccgacaagaatttccgtaac 9879
human.FAM230E-LOC105377182-GGT3P.NCBI.                                      aatgctgctttcacagcatatctcttggcttgttaaaaacccgacaagaatttccgtaac 8951
human.FAM230J-LOC105372942-GGTLC5P.NCBI.rev.compl                           aatgctgctttcacagcatatctcttggcttgttaaaaacccgacaagaatttccgtaac 9332
human.FAM230A.ncRNA.18487127.18500594-GGTLC3.18516335.18518165.rev.compl.REF ------------------------------------------------------------ 0

chimp.LOC112206744-LOC107973052-GGT2.revcompl                               attaatttattttaacacctatattgggtgtgaacccaccataaagtttgcccactgaa 11972
chimp54546-135457.revcompl.Pan.troglodytes.clone.rp43-41g5.GenBank:AC099533.36 ATTAATTTATTTTTAACACCTATATTGGGTGTGAACCCACCATAAAGTTTGCCCACTGAA 13507
human.FAM30B-LOC105372935-GGT2.NCBI.                                        attaatttatttttaacacctatattgggtgtgaacccaccataaagtttgcccactgaa 9939
human.FAM230E-LOC105377182-GGT3P.NCBI.                                      a-----ttatttttaacacctatattgggtgtgaacccaccataaagtttgcccactgaa 9006
human.FAM230J-LOC105372942-GGTLC5P.NCBI.rev.compl                           a-----ttatttttaacacctatattgggtgtgaacccaccataaagtttgcccactgaa 9387
human.FAM230A.ncRNA.18487127.18500594-GGTLC3.18516335.18518165.rev.compl.REF ------------------------------------------------------------ 0

chimp.LOC112206744-LOC107973052-GGT2.revcompl                               aaggtctacaatttgatgctttattaaattgatactgtgtgcacccatcaccaccatcta 12032
chimp54546-135457.revcompl.Pan.troglodytes.clone.rp43-41g5.GenBank:AC099533.36 AAGGTCTACAATTTGATGCTTTATTAAATTGATACTGTGTGCACCCATCACCACGATCTA 13567
human.FAM30B-LOC105372935-GGT2.NCBI.                                        aaggtctacaatttgatgctttattaaattgatactgtgtgcacccatcaccacgatcta 9999
human.FAM230E-LOC105377182-GGT3P.NCBI.                                      aaggtctagaatttgatgctttattaaattgatactgtgtgcacccaacaccacgatcta 9066
```

```
human.FAM230J-LOC105372942-GGTLC5P.NCBI.rev.compl                              aaggtctagaatttgatgctttattaaattgatactgtgtgcacccaacaccacgatcta 9447
human.FAM230A.ncRNA.18487127.18500594-GGTLC3.18516335.18518165.rev.compl.REF   ------------------------------------------------------------ 0

chimp.LOC112206744-LOC107973052-GGT2.revcompl                                  atttaaatatgtttccctcacccaaattctctcttgcgcactggcagttaatccccaccc 12092
chimp54546-135457.revcompl.Pan.troglodytes.clone.rp43-41g5.GenBank:AC099533.36 ATTTAAATATGTTTCCCTCACCCAAATTCTCTCTTGCACACTGGCAGTTAATCCCCACCC 13627
human.FAM30B-LOC105372935-GGT2.NCBI.                                           atttaaatatgtttccctcacccaaattctctcttgcgcactggcagttaatccccactc 10059
human.FAM230E-LOC105377182-GGT3P.NCBI.                                         atttaaatatgtttccctcacccaagttctctcttgcgcactggcagttaatccccactc 9126
human.FAM230J-LOC105372942-GGTLC5P.NCBI.rev.compl                              atttaaatatgtttccctcacccaagttctctcttgcgcactggcagttaatccccactc 9507
human.FAM230A.ncRNA.18487127.18500594-GGTLC3.18516335.18518165.rev.compl.REF   ------------------------------------------------------------ 0

chimp.LOC112206744-LOC107973052-GGT2.revcompl                                  ccatctccagccctaagcaatactgctgtgacattccatctccataaatttcccacttgc 12152
chimp54546-135457.revcompl.Pan.troglodytes.clone.rp43-41g5.GenBank:AC099533.36 CCATCTCCAGCCCTAAGCAATACTGCTGTGACATTCCATCTCCATAAATTTCCCACTTGC 13687
human.FAM30B-LOC105372935-GGT2.NCBI.                                           ccatctccagccctaagcaatactgctgtgacattccatctccataaatttcccatttgc 10119
human.FAM230E-LOC105377182-GGT3P.NCBI.                                         ccatctccagccctaagcaatactgctgtgacattccatctccataaatttcccacttgc 9186
human.FAM230J-LOC105372942-GGTLC5P.NCBI.rev.compl                              ccatctccagccctaagcaatactgctgtgacattccatctccataaatttcccacttgc 9567
human.FAM230A.ncRNA.18487127.18500594-GGTLC3.18516335.18518165.rev.compl.REF   ------------------------------------------------------------ 0

chimp.LOC112206744-LOC107973052-GGT2.revcompl                                  ttaatagaaatggacatatatatatatttggaatctgacttccttcatttagcatactat 12212
chimp54546-135457.revcompl.Pan.troglodytes.clone.rp43-41g5.GenBank:AC099533.36 TTAATAGAATTGGACATATATATATATTTGGAATCTGACTTCCTTCATTTAGCATACTAT 13747
human.FAM30B-LOC105372935-GGT2.NCBI.                                           ttaatagaaatggacatatatatatatttggaatctgacttccttcatttagcatactat 10179
human.FAM230E-LOC105377182-GGT3P.NCBI.                                         tttatagaaatggacatatatatatatttggaatctgacttccctcatttagcatactat 9246
human.FAM230J-LOC105372942-GGTLC5P.NCBI.rev.compl                              tttatagaaatggacatatatatatatttggaatctgacttccctcatttagcatactat 9627
human.FAM230A.ncRNA.18487127.18500594-GGTLC3.18516335.18518165.rev.compl.REF   ------------------------------------------------------------ 0

chimp.LOC112206744-LOC107973052-GGT2.revcompl                                  gtttgaagttaattgacgtgttagcacgtgctggtcatgtgttttccttcatagtctgct 12272
chimp54546-135457.revcompl.Pan.troglodytes.clone.rp43-41g5.GenBank:AC099533.36 GTTTGAAGTTAATTGACGTGTTAGCACGTGCTGGTCATGTGTTTTCCTTCATAGTCTGCT 13807
human.FAM30B-LOC105372935-GGT2.NCBI.                                           gtttgaagttaattgacgtgttagcacgtgctggtcatgtgttttccttcatagtctgct 10239
human.FAM230E-LOC105377182-GGT3P.NCBI.                                         gtttgaagttaattgacgtgttagcacgtgctggtcatgtgttttccttcatagtctgct 9306
human.FAM230J-LOC105372942-GGTLC5P.NCBI.rev.compl                              gtttgaagttaattgacgtgttagcacgtgctggtcatgtgttttccttcatagtctgct 9687
human.FAM230A.ncRNA.18487127.18500594-GGTLC3.18516335.18518165.rev.compl.REF   ------------------------------------------------------------ 0

chimp.LOC112206744-LOC107973052-GGT2.revcompl                                  gtgtttactcatacagatagtgtttattcgtttatcagttaatggacatttaattgtttt 12332
chimp54546-135457.revcompl.Pan.troglodytes.clone.rp43-41g5.GenBank:AC099533.36 GTGTTTACTCATACAGATAGTGTTTATTCATTTATCAGTTAATGGACATTTAATTGTTTT 13867
human.FAM30B-LOC105372935-GGT2.NCBI.                                           gtgtttactcatacagatagtgtttattcatttatcagttaatggacatttaattgtttt 10299
human.FAM230E-LOC105377182-GGT3P.NCBI.                                         gtgtttattcatacagatagtgtttattcatttatcagttaatggacatttaattgtttt 9366
human.FAM230J-LOC105372942-GGTLC5P.NCBI.rev.compl                              gtgtttattcatacagatagtgtttattcatttatcagttaatggacatttaattgtttt 9747
human.FAM230A.ncRNA.18487127.18500594-GGTLC3.18516335.18518165.rev.compl.REF   ------------------------------------------------------------ 0

chimp.LOC112206744-LOC107973052-GGT2.revcompl                                  gttattttctttgatgagtaacgtagcttcgagcattcatatacagtcaggtaatgcata 12392
chimp54546-135457.revcompl.Pan.troglodytes.clone.rp43-41g5.GenBank:AC099533.36 GTTATTTTCTTTGATGAGTAATATAGCTTTGAGCATTCATATACAGTCACGTAATGCATA 13927
human.FAM30B-LOC105372935-GGT2.NCBI.                                           gttattttctttgatgagtaatgtagctttgagcattcatatacagtcatgtaatgcata 10359
human.FAM230E-LOC105377182-GGT3P.NCBI.                                         gttattttctttgatgagtaatgtagctttgtgcattcatatacagtcatgtaatgcata 9426
human.FAM230J-LOC105372942-GGTLC5P.NCBI.rev.compl                              gttattttctttgatgagtaatgtagctttgagcattcatatacagtcatgtaatgcata 9807
human.FAM230A.ncRNA.18487127.18500594-GGTLC3.18516335.18518165.rev.compl.REF   ------------------------------------------------------------ 0

chimp.LOC112206744-LOC107973052-GGT2.revcompl                                  atgacattttggtcaaaaaaaa-tttttttttctgagacccaggctggagtgcagtggc 12451
chimp54546-135457.revcompl.Pan.troglodytes.clone.rp43-41g5.GenBank:AC099533.36 ATGACGTTTTCGTCAAAAAAAA-TTTTTTTTTCTGAGACCCAGGCTGGAGTGCAGTGGC 13986
human.FAM30B-LOC105372935-GGT2.NCBI.                                           atgacattttggtc-aaaaaaatttttttttttctgagacccaggctggagtgcagtggc 10418
human.FAM230E-LOC105377182-GGT3P.NCBI.                                         atgacattttggtcaaaaaaaatttttttttttctgagacccaggctggagtgcagtggc 9486
human.FAM230J-LOC105372942-GGTLC5P.NCBI.rev.compl                              atgacattttggtcaaaaaaaatttttttttttctgagacccaggctggagtgcagtggc 9867
human.FAM230A.ncRNA.18487127.18500594-GGTLC3.18516335.18518165.rev.compl.REF   ------------------------------------------------------------ 0

chimp.LOC112206744-LOC107973052-GGT2.revcompl                                  acaatctcggctcactggaacctccacctcccaggtttaagcaattcttgtgcctcaacc 12511
chimp54546-135457.revcompl.Pan.troglodytes.clone.rp43-41g5.GenBank:AC099533.36 ACAATCTCGGCTCACTGGAACCTCCACCTCCCAGGTTTAAGCAATTCTCCTGCCTCAACT 14046
human.FAM30B-LOC105372935-GGT2.NCBI.                                           acaatctcggctcactggaacctccacctcccaggtttaagcaattctcatgcctcaacc 10478
human.FAM230E-LOC105377182-GGT3P.NCBI.                                         acaatctcggctcactggaacctccatctcccaggtttaagcaattctcatgcctcaacc 9546
human.FAM230J-LOC105372942-GGTLC5P.NCBI.rev.compl                              acaatctcggctcactggaacctccatctcccaggtttaagcaattctcgtgcctcaacc 9927
human.FAM230A.ncRNA.18487127.18500594-GGTLC3.18516335.18518165.rev.compl.REF   ------------------------------------------------------------ 0

chimp.LOC112206744-LOC107973052-GGT2.revcompl                                  tcccgagtagctgggacaactggcacacgccaccatgcctggataattttttgtattttca 12571
chimp54546-135457.revcompl.Pan.troglodytes.clone.rp43-41g5.GenBank:AC099533.36 TCCCGAGTAGCTGGGACAACTGTCACACGCCACCATGCCTGGATAATTTTTGTATTTTCA 14106
human.FAM30B-LOC105372935-GGT2.NCBI.                                           tcccgagtagctgggacaactggcacacgccaccatgcctggataatttttgtattttca 10538
human.FAM230E-LOC105377182-GGT3P.NCBI.                                         tcccgagtagctgggacaactggcacacaccaccacgcctggataatttttgtattttca 9606
human.FAM230J-LOC105372942-GGTLC5P.NCBI.rev.compl                              tcccgagtagctgggacaactggcacacaccaccacacctggataatttttgtattttca 9987
human.FAM230A.ncRNA.18487127.18500594-GGTLC3.18516335.18518165.rev.compl.REF   ------------------------------------------------------------ 0

chimp.LOC112206744-LOC107973052-GGT2.revcompl                                  gtagagacaggattttgctgtgttggtcaggctagtctcaaactcctagcctcaggtgat 12631
chimp54546-135457.revcompl.Pan.troglodytes.clone.rp43-41g5.GenBank:AC099533.36 GTAGAGACAGGATTTTGCTGTGTTGGTCAGGCTAGTCTCAAACTCCTAGCCTCAGGTGAT 14166
human.FAM30B-LOC105372935-GGT2.NCBI.                                           gtagagacaggattttgctgtgttggtcaggctagtctcaaactcccagcctcaggtgat 10598
human.FAM230E-LOC105377182-GGT3P.NCBI.                                         gtagagacaggattttgctgtgttggtcaggctagtctcaaactcctagcctcaggtgat 9666
human.FAM230J-LOC105372942-GGTLC5P.NCBI.rev.compl                              gtagagacaggattttgctgtgttggtcaggctagtctcaaactcctagcctcaggtgat 10047
human.FAM230A.ncRNA.18487127.18500594-GGTLC3.18516335.18518165.rev.compl.REF   ------------------------------------------------------------ 0

chimp.LOC112206744-LOC107973052-GGT2.revcompl                                  ccacccatctctgcctcccaaagtgctgggattataggcatgagccaccacacccagcct 12691
chimp54546-135457.revcompl.Pan.troglodytes.clone.rp43-41g5.GenBank:AC099533.36 CCACCCATCTCTGCCTCCCAAAGTGCTGGGATTATAGGCATGAGCCACCACACCCAGCCT 14226
human.FAM30B-LOC105372935-GGT2.NCBI.                                           ccacctatctctgcctcccaaagtgctgggattataggcatgagccaccacacccagcct 10658
human.FAM230E-LOC105377182-GGT3P.NCBI.                                         ccacccacctctgcctcccaaagtgctgggattataggcatgagccaccacacccagcct 9726
human.FAM230J-LOC105372942-GGTLC5P.NCBI.rev.compl                              ccacccacctctgcctcccaaagtgctgggattataggcatgagccaccacacccagcct 10107
human.FAM230A.ncRNA.18487127.18500594-GGTLC3.18516335.18518165.rev.compl.REF   ------------------------------------------------------------ 0

chimp.LOC112206744-LOC107973052-GGT2.revcompl                                  aatttttttaagaaacaagggaactattttctaaattacttttgccaatttatatttcta 12751
chimp54546-135457.revcompl.Pan.troglodytes.clone.rp43-41g5.GenBank:AC099533.36 AATTTTTTTAAGAAACAAGGGAACTATTTTCTAAATTACTTTTGCCAATTGATATTTCTA 14286
human.FAM30B-LOC105372935-GGT2.NCBI.                                           aatttttttaagaaacaagggaactattttctaaattacttttgccaatttatatttcta 10718
human.FAM230E-LOC105377182-GGT3P.NCBI.                                         aatttttttaagaaagaagggaactattttctaaattacttttgccaatttatattccta 9786
human.FAM230J-LOC105372942-GGTLC5P.NCBI.rev.compl                              aatttttttaagaaagaagggaactattttctaaattacttttgccaatttatattccta 10167
human.FAM230A.ncRNA.18487127.18500594-GGTLC3.18516335.18518165.rev.compl.REF   ------------------------------------------------------------ 0

chimp.LOC112206744-LOC107973052-GGT2.revcompl                                  ccatgatgcatagcactaatttcaccgtacaatgtatggtaggccccaatatgtaagaaa 12811
chimp54546-135457.revcompl.Pan.troglodytes.clone.rp43-41g5.GenBank:AC099533.36 CCATGATGCATAGCACTAATTTCACTGTACAATGTATGGTAGGCCCCAGTATGTAAGAAA 14346
human.FAM30B-LOC105372935-GGT2.NCBI.                                           ccatgatgcatagcactaatttcaccgtacaatgtatggtaggccccaatatgtaagaaa 10778
human.FAM230E-LOC105377182-GGT3P.NCBI.                                         ccatgatgcatagcactaatttcaccgtacaatgtatggtaggccccaatatgtaagaaa 9846
human.FAM230J-LOC105372942-GGTLC5P.NCBI.rev.compl                              ccatgatgcatagcactaatttcaccgtacaatgtatggtaggccccaatatgtaagaaa 10227
human.FAM230A.ncRNA.18487127.18500594-GGTLC3.18516335.18518165.rev.compl.REF   ------------------------------------------------------------ 0

chimp.LOC112206744-LOC107973052-GGT2.revcompl                                  tgatgaaagtaacacataaagattagtataaaacaaataaaattatcattgttgctatca 12871
chimp54546-135457.revcompl.Pan.troglodytes.clone.rp43-41g5.GenBank:AC099533.36 TGATGAAAGTAACACATAAAGATTAGTATAAAACAAATAAGATTATCATTGTTGCTATCA 14406
human.FAM30B-LOC105372935-GGT2.NCBI.                                           tgatgaaaataacacataaagattggtataaaacaaataagattatcattgttgctatca 10838
human.FAM230E-LOC105377182-GGT3P.NCBI.                                         tgatgaaagtaacacataaagattagtataagacaaatgagattatcgttgttgctatca 9906
human.FAM230J-LOC105372942-GGTLC5P.NCBI.rev.compl                              tgatgaaagtaacacataaagattagtataagacaaatgagattatcgttgttgctatca 10287
human.FAM230A.ncRNA.18487127.18500594-GGTLC3.18516335.18518165.rev.compl.REF   ------------------------------------------------------------ 0

chimp.LOC112206744-LOC107973052-GGT2.revcompl                                  tctttgtcaaattctgaaaacaatctgagtatattttttatataaatatgcttggcaacat 12931
chimp54546-135457.revcompl.Pan.troglodytes.clone.rp43-41g5.GenBank:AC099533.36 TCTTTGTCAAATTCTGAAAACAATCTGAGTATATTTTTATATAAATATGCTTGGCAACAT 14466
human.FAM30B-LOC105372935-GGT2.NCBI.                                           tctttgtcaaattctgaaaacaatctgagtatatttttatataaatatgcttggcaacat 10898
human.FAM230E-LOC105377182-GGT3P.NCBI.                                         tctttgtcaaattctgaaaacaatctgagtatatttttatataaatatgcttggcaacat 9966
human.FAM230J-LOC105372942-GGTLC5P.NCBI.rev.compl                              tctttgtcaaattctgaaaacaatctgagtatatttttatataaatatgcttggcaacat 10347
human.FAM230A.ncRNA.18487127.18500594-GGTLC3.18516335.18518165.rev.compl.REF   ------------------------------------------------------------ 0

chimp.LOC112206744-LOC107973052-GGT2.revcompl                                  agctgaaaaacgcattatcagttacatttatcagtaacaaagacataaatttgaaggggg 12991
chimp54546-135457.revcompl.Pan.troglodytes.clone.rp43-41g5.GenBank:AC099533.36 AGCTGAAAAACGCATTATCAGTTACATTTATCAGTAACAAAGACATAAATTTGAAGGGGG 14526
human.FAM30B-LOC105372935-GGT2.NCBI.                                           agctgaaaaacgcattatcagttacatttatcagtaacaaagacataaatttgaaggggg 10958
human.FAM230E-LOC105377182-GGT3P.NCBI.                                         agctgaaaaaagcattatcagttacatttatcagtaacaaagacataaatttgaaggggg 10026
human.FAM230J-LOC105372942-GGTLC5P.NCBI.rev.compl                              agctgaaaaaagcattatcagttacatttatcagtaacaaagacataaatttgaaggggg 10407
human.FAM230A.ncRNA.18487127.18500594-GGTLC3.18516335.18518165.rev.compl.REF   ------------------------------------------------------------ 0

chimp.LOC112206744-LOC107973052-GGT2.revcompl                                  aaaaacacttgtactaacaacgcaatgtcagaattcacataaaaattctgctggtcactt 13051
chimp54546-135457.revcompl.Pan.troglodytes.clone.rp43-41g5.GenBank:AC099533.36 AAAAACACTTGTACTAACAACGCAATGTCAGAATTAACATAAAAATTCTGCTGGTCACTT 14586
human.FAM30B-LOC105372935-GGT2.NCBI.                                           aacaacacttgtactaacaacgcaatgtcagaattaacataaaaattctgctggtcactt 11018
human.FAM230E-LOC105377182-GGT3P.NCBI.                                         aaaaacacttgtactaacaatgcaatgtcagaattaacataaaaattctgctggtcactt 10086
human.FAM230J-LOC105372942-GGTLC5P.NCBI.rev.compl                              aaaaacacttgtactaacaatgcaatgtcagaattaacataaaaattctgctggtcactt 10467
human.FAM230A.ncRNA.18487127.18500594-GGTLC3.18516335.18518165.rev.compl.REF   ------------------------------------------------------------ 0

chimp.LOC112206744-LOC107973052-GGT2.revcompl                                  tggaatatttaattgcctggggcagtgtttagtagacaaatgagcatctatggagcaccc 13111
chimp54546-135457.revcompl.Pan.troglodytes.clone.rp43-41g5.GenBank:AC099533.36 TGGAATATTTAATTGCCTGGGGCAGTGTTTAGTAGACAAATGAGCATCTATGGAGCACCC 14646
human.FAM30B-LOC105372935-GGT2.NCBI.                                           tggaatatttaattgcctggggcagtgtttagtagacaaatgagcatctatggagcactc 11078
human.FAM230E-LOC105377182-GGT3P.NCBI.                                         tggaatatttaattgcctggggcagtgtttagtagacacatgagcatctatggagcaccc 10146
human.FAM230J-LOC105372942-GGTLC5P.NCBI.rev.compl                              tggaatatttaattgcctggggcagtgtttagtagacaaatgagcatctatggagcaccc 10527
```

```
human.FAM230A.ncRNA.18487127.18500594-GGTLC3.18516335.18518165.rev.compl.REF        ------------------------------------------------------------ 0

chimp.LOC112206744-LOC107973052-GGT2.revcompl                                       aaagtaggggaatcaacagaacttcggtttgaaaagttatctgggtttagagcgtgaaac 13171
chimp54546-135457.revcompl.Pan.troglodytes.clone.rp43-41g5.GenBank:AC099533.36      AAAGTAGGGGAATCAACAGAACTTGGATTTGAAAAGTTATCTGGGTTTAGAGCGTGAAAC 14706
human.FAM30B-LOC105372935-GGT2.NCBI.                                                aaagtaggggaatcaacagaacttgggtttgaaaagttatctgggtttagagcgtgaaac 11138
human.FAM230E-LOC105377182-GGT3P.NCBI.                                              aaagtaggggaatcaacagaacttgggtttcaaaagttatctgggtttagagcgtgaaac 10206
human.FAM230J-LOC105372942-GGTLC5P.NCBI.rev.compl                                   aaagtaggggaatcaacagaacttgggtttcaaaagttatctgggtttagagcgtgaaac 10587
human.FAM230A.ncRNA.18487127.18500594-GGTLC3.18516335.18518165.rev.compl.REF        ------------------------------------------------------------ 0

chimp.LOC112206744-LOC107973052-GGT2.revcompl                                       tttgttagaggacacacgccttgcatgagcgaggtgccttggtgtgtgtggacgtaccat 13231
chimp54546-135457.revcompl.Pan.troglodytes.clone.rp43-41g5.GenBank:AC099533.36      TTTGTTAGAGGACACACACCTTGCATGAGCGAGGTGCCTTGGTGTGTGTGGATGTACCAT 14766
human.FAM30B-LOC105372935-GGT2.NCBI.                                                tttgttagaggacacgcaccttgcatgagcgaggtgccctggtgtgtgtggacgtaccat 11198
human.FAM230E-LOC105377182-GGT3P.NCBI.                                              tttgttagaggacacacaccttgcatgagtgaggtgccttggtgtgtgtggacgtatcat 10266
human.FAM230J-LOC105372942-GGTLC5P.NCBI.rev.compl                                   tttgttagaggacacacaccttgcatgagtgaggtgccttggtgtgtgtggacgtatcat 10647
human.FAM230A.ncRNA.18487127.18500594-GGTLC3.18516335.18518165.rev.compl.REF        ------------------------------------------------------------ 0

chimp.LOC112206744-LOC107973052-GGT2.revcompl                                       tatgcttggaggtacagcataatggtggcttcctccagaaagggacatttggggtggatt 13291
chimp54546-135457.revcompl.Pan.troglodytes.clone.rp43-41g5.GenBank:AC099533.36      TATGCTTGGAGGTACAGCATAATGGTGGCTTCCTCCAGAAAGGGACATTTGGGGTAGATT 14826
human.FAM30B-LOC105372935-GGT2.NCBI.                                                tatgcttggaggtacagcataatggtggcttcctccagaaagggacatttggggtggatt 11258
human.FAM230E-LOC105377182-GGT3P.NCBI.                                              tatgcttggaggtacagcataatggtggcttcctccagaaagggacatttggggtagatt 10326
human.FAM230J-LOC105372942-GGTLC5P.NCBI.rev.compl                                   tatgcttggaggtacagcataatggtggcttcctccagaaagggacatttggggtagatt 10707
human.FAM230A.ncRNA.18487127.18500594-GGTLC3.18516335.18518165.rev.compl.REF        ------------------------------------------------------------ 0

chimp.LOC112206744-LOC107973052-GGT2.revcompl                                       cattccatctagacaacacagcctgatgtggcatggacatgaatggaggtgaaatggtca 13351
chimp54546-135457.revcompl.Pan.troglodytes.clone.rp43-41g5.GenBank:AC099533.36      CATTCCATCTAGACAACACAGCCTGATGTGGCATGGACATGAATGGAGGTGAAATGGTCA 14886
human.FAM30B-LOC105372935-GGT2.NCBI.                                                cattccatctagacaacacagcctgatgtggcatggacatgaatggaggtgaaatggtca 11318
human.FAM230E-LOC105377182-GGT3P.NCBI.                                              catgccatctagacaacacagcctgatgtggcatggacatgaatggaggtgaaatggtca 10386
human.FAM230J-LOC105372942-GGTLC5P.NCBI.rev.compl                                   catgccatctagacaacacagcctgatgtggcatggacatgaatggaggtgaaatggtca 10767
human.FAM230A.ncRNA.18487127.18500594-GGTLC3.18516335.18518165.rev.compl.REF        ------------------------------------------------------------ 0

chimp.LOC112206744-LOC107973052-GGT2.revcompl                                       gtagttgagaggatcagtcctgacaagggccaaggtgaaaaacctgggaaccccttcagg 13411
chimp54546-135457.revcompl.Pan.troglodytes.clone.rp43-41g5.GenBank:AC099533.36      GTAGTTGAGAGGATCAGTCCTGACAAGGGCCGAGGTGAAAAACCTGGGAACCCCTTCAGG 14946
human.FAM30B-LOC105372935-GGT2.NCBI.                                                gtagttgagaggatcagtcctgacaagggccgaggtgaaaaacctgggaaccccttcagg 11378
human.FAM230E-LOC105377182-GGT3P.NCBI.                                              gtagttgagaggatcagtcctgacaagggccgaggtgaaaaacctgggaaccccttcagg 10446
human.FAM230J-LOC105372942-GGTLC5P.NCBI.rev.compl                                   gtagttgagaggatcagtcctgacaagggccgaggtgaaaaacctgggaaccccttcagg 10827
human.FAM230A.ncRNA.18487127.18500594-GGTLC3.18516335.18518165.rev.compl.REF        ------------------------------------------------------------ 0

chimp.LOC112206744-LOC107973052-GGT2.revcompl                                       tacaaagtcttcagttgaaaaaggaggtggtcacaggaaatactgagacaggacagcaat 13471
chimp54546-135457.revcompl.Pan.troglodytes.clone.rp43-41g5.GenBank:AC099533.36      TACAAAGTCTTCAGTTGAAAAAGGAGGTGGTCACAGGAAATACTGAGACAGGACAGCAAT 15006
human.FAM30B-LOC105372935-GGT2.NCBI.                                                tacaaagtcttcagttgaaaaaggaggtggtcacaggaaatactgagacaggacagcaaa 11438
human.FAM230E-LOC105377182-GGT3P.NCBI.                                              tgcaaagtcttcagttgaaaaaggaggtggtcacaggaaatactgagatgggtcagcaat 10506
human.FAM230J-LOC105372942-GGTLC5P.NCBI.rev.compl                                   tgcaaagtcttcagttgaaaaaggaggtggtcacaggaaatactgagatgggtcagcaat 10887
human.FAM230A.ncRNA.18487127.18500594-GGTLC3.18516335.18518165.rev.compl.REF        ------------------------------------------------------------ 0

chimp.LOC112206744-LOC107973052-GGT2.revcompl                                       gcatgggagacagagttcttggccctgcagggtgagtagtgtggattctcaagttttctc 13531
chimp54546-135457.revcompl.Pan.troglodytes.clone.rp43-41g5.GenBank:AC099533.36      GCATGGGAGACAGAGTTCTTGGCCCTGCAAGGTGAGTAGTGTGGATTCTCAAGTTTTCTC 15066
human.FAM30B-LOC105372935-GGT2.NCBI.                                                gcatgggagacagagttcttggccctgcagggtgagtagtgtggattctcaagttttctc 11498
human.FAM230E-LOC105377182-GGT3P.NCBI.                                              gcatgggagacagagttcttggccctgcagggtgagtagtgtggattctcaagttttctc 10566
human.FAM230J-LOC105372942-GGTLC5P.NCBI.rev.compl                                   gcatgggagacagagttcttggccctgcagggtgagtagtgtggattctcaagttttctc 10947
human.FAM230A.ncRNA.18487127.18500594-GGTLC3.18516335.18518165.rev.compl.REF        ------------------------------------------------------------ 0

chimp.LOC112206744-LOC107973052-GGT2.revcompl                                       ctctctccattaatttctttcccaatgcagatgacttccatcatacagtcttcagcaacc 13591
chimp54546-135457.revcompl.Pan.troglodytes.clone.rp43-41g5.GenBank:AC099533.36      CTCTCTCCATTAATTTCTTTCCCAATGCAGATGACTTCCATCATACAGTCTTCAGCAACC 15126
human.FAM30B-LOC105372935-GGT2.NCBI.                                                ctctctccattaatttctttcccaatgcagatgacttccatcatacagtcttcagcaacc 11558
human.FAM230E-LOC105377182-GGT3P.NCBI.                                              ctctctccattaatttctttcccaatgcagatgacttccatcatacagtcttcagcaacc 10626
human.FAM230J-LOC105372942-GGTLC5P.NCBI.rev.compl                                   ctctctccattaatttctttcccaatgcagatgacttccatcatacagtcttcagcaacc 11007
human.FAM230A.ncRNA.18487127.18500594-GGTLC3.18516335.18518165.rev.compl.REF        ------------------ttcccaatgcagatgacttccatcatacagtcttcagcaacc 42
                                                                                                      ******************************************

chimp.LOC112206744-LOC107973052-GGT2.revcompl                                       ttgaaagattggacaagcttcagcccactcttgaaggtaaaggaaggcagctaacaagac 13651
chimp54546-135457.revcompl.Pan.troglodytes.clone.rp43-41g5.GenBank:AC099533.36      TTGAAAGATTGGACAAGCTTCAGCCCACTCTTGAAGGTAAAGGAAGGCAGCTAACAAGAC 15186
human.FAM30B-LOC105372935-GGT2.NCBI.                                                ttgaaagattggacaagcttcagcccactcttgaaggtaaaggaaggcagctaacaagac 11618
human.FAM230E-LOC105377182-GGT3P.NCBI.                                              ttgaaagattggacaagcttcagcccactcttgaaggtaaaggaaggcagctaacaagac 10686
human.FAM230J-LOC105372942-GGTLC5P.NCBI.rev.compl                                   ttgaaagattggacaagcttcagcccactcttgaaggtaaaggaaggcagctaacaagac 11067
human.FAM230A.ncRNA.18487127.18500594-GGTLC3.18516335.18518165.rev.compl.REF        ttgaaagattggacaagcttcagcccactcttgaaggtaaaggaaggcagctaacaagac 102
                                                                                    ************************************************************

chimp.LOC112206744-LOC107973052-GGT2.revcompl                                       tggcatctgggcttggctgtgcgtgttttctattgtggggaaatatatataacacaatat 13711
chimp54546-135457.revcompl.Pan.troglodytes.clone.rp43-41g5.GenBank:AC099533.36      TGGCATCTGGGCTTGGCTGTGCGTGTTTTCTATCGTGGGGAAATATATATAACACATTAT 15246
human.FAM30B-LOC105372935-GGT2.NCBI.                                                tggcatctgggcttggctgtgcgtgttttctatcgtggggaaatatatataacacaatat 11678
human.FAM230E-LOC105377182-GGT3P.NCBI.                                              tggcatctgggcttggctgtgcatgttttctatcgtggggaaatatatataacacattat 10746
human.FAM230J-LOC105372942-GGTLC5P.NCBI.rev.compl                                   tggcatctgggcttggctgtgcatgttttctatcgtggggaaatatatataacacattat 11127
human.FAM230A.ncRNA.18487127.18500594-GGTLC3.18516335.18518165.rev.compl.REF        tggcatctgggcttggctgtgcatgttttctatcgtggggaaatatatgtaacacattat 162
                                                                                    ********************** ********** ************** ******* ***

chimp.LOC112206744-LOC107973052-GGT2.revcompl                                       ttatcatttgaaccttttaaccaaagtgtgcactccatggcattcaatatattcacaggg 13771
chimp54546-135457.revcompl.Pan.troglodytes.clone.rp43-41g5.GenBank:AC099533.36      TTATCATTTGAACCTTTTAACCAAAGTGTGCACTCCATGGCATTCAATATATTCACAGGG 15306
human.FAM30B-LOC105372935-GGT2.NCBI.                                                ttatcatttgaaccttttaaccaaagtgtgcactccatggcattcaatatattcacaggg 11738
human.FAM230E-LOC105377182-GGT3P.NCBI.                                              ttatcatttgaaccttttaaccaaagtgtgcactccgtggcattcaatatattcacaggg 10806
human.FAM230J-LOC105372942-GGTLC5P.NCBI.rev.compl                                   ttatcatttgaaccttttaaccaaagtgtgcactccgtggcattcaatatattcacaggg 11187
human.FAM230A.ncRNA.18487127.18500594-GGTLC3.18516335.18518165.rev.compl.REF        ttatcatttgaaccttttaaccaaagtgtgcactccgtggcattcaatatattcacaggg 222
                                                                                    ************************************ ***********************

chimp.LOC112206744-LOC107973052-GGT2.revcompl                                       ttgcataaccaacaccactatctacacccacaattttgatgatttcttacaaaaccttgt 13831
chimp54546-135457.revcompl.Pan.troglodytes.clone.rp43-41g5.GenBank:AC099533.36      TTGCATAACCAACACCACTATCTACACCCACAATTTTGATGATTTCTTACAAAACCTTGT 15366
human.FAM30B-LOC105372935-GGT2.NCBI.                                                ttgcataaccaacaccactatctacacccacaattttgatgatttcttacgaaaccttgt 11798
human.FAM230E-LOC105377182-GGT3P.NCBI.                                              ttgcataaccaacaccactatctacacccacaattttgatgatttcttacaaaaccttgt 10866
human.FAM230J-LOC105372942-GGTLC5P.NCBI.rev.compl                                   ttgcataaccaacaccactatctacacccacaattttgatgatttcttacaaaaccttgt 11247
human.FAM230A.ncRNA.18487127.18500594-GGTLC3.18516335.18518165.rev.compl.REF        ttgcataaccaacaccactatctacacccacaattttgatgatttcttacaaaaccttgt 282
                                                                                    ************************************************** *********

chimp.LOC112206744-LOC107973052-GGT2.revcompl                                       ccactataagcaatatagcaccttcccctatttccagctcatggtgattcctatcccac 13891
chimp54546-135457.revcompl.Pan.troglodytes.clone.rp43-41g5.GenBank:AC099533.36      CCACAATAAGCAATATAGCACCTTCCCCCTATTTCCAGCCCATGGTGATTCCTATCCCAC 15426
human.FAM30B-LOC105372935-GGT2.NCBI.                                                ccacaataagcaatatagcaccttccccctatttccagcccatggtgattcctatcccac 11858
human.FAM230E-LOC105377182-GGT3P.NCBI.                                              ccacaataagcaatatagcgccttccccctatttccagcccatggtgattcctatcccac 10926
human.FAM230J-LOC105372942-GGTLC5P.NCBI.rev.compl                                   ccacaataagcaatatagcgccttccccctatttccagcccatggtgattcctatccgac 11307
human.FAM230A.ncRNA.18487127.18500594-GGTLC3.18516335.18518165.rev.compl.REF        ccacaataagcaatatagcaccttccccctatttccagcccatggtgattcctatccgac 342
                                                                                    **** ************** ******************* ***************** **

chimp.LOC112206744-LOC107973052-GGT2.revcompl                                       tttctcttgtatgaatttgactattctaggcacttcatgtaattacaattatacaatata 13951
chimp54546-135457.revcompl.Pan.troglodytes.clone.rp43-41g5.GenBank:AC099533.36      TTTCTCTTGTATGAATTTGACTATTCTAGGCACTTCATGTCATTACAATTATACAATATA 15486
human.FAM30B-LOC105372935-GGT2.NCBI.                                                tttctcttgtatgaatttgactattctaggcacttcatgtaattacaattatacaatata 11918
human.FAM230E-LOC105377182-GGT3P.NCBI.                                              --tctctcgtatgaatttgactattctaggcacttcatgtaattacaattatacaatata 10984
human.FAM230J-LOC105372942-GGTLC5P.NCBI.rev.compl                                   tttctcttgtatgaatttgactattctaggcacttcatgtaattacaattatacaatata 11367
human.FAM230A.ncRNA.18487127.18500594-GGTLC3.18516335.18518165.rev.compl.REF        tttctcttgtatgaatttgactattctaggcacttcatgtaattacaattatacaatata 402
                                                                                      ***** ******************************** *******************

chimp.LOC112206744-LOC107973052-GGT2.revcompl                                       ttccttttgtgtctggcttatttcactaagcataatgttctcaatgtccacccatgttgt 14011
chimp54546-135457.revcompl.Pan.troglodytes.clone.rp43-41g5.GenBank:AC099533.36      TTCCTTTTGTGTCTGGCTTATTTCACTAAGCATAATGTTCTCAATGTCCACCCATGTTGT 15546
human.FAM30B-LOC105372935-GGT2.NCBI.                                                ttccttttgtgtctggcttatttcactaagcataatgttctcaatgtccacccatgttgt 11978
human.FAM230E-LOC105377182-GGT3P.NCBI.                                              ttccttttgtgtctggcttatttcactaagcataatgttctcaatgtccacccatgttgt 11044
human.FAM230J-LOC105372942-GGTLC5P.NCBI.rev.compl                                   ttccttttgtgtctggcttatttcactaagcataatgttctcaatgtccacccatgttgt 11427
human.FAM230A.ncRNA.18487127.18500594-GGTLC3.18516335.18518165.rev.compl.REF        ttccttttgtgtctggcttatttcactaagcataatgttctcaatgtccacccatgttgt 462
                                                                                    ************************************************************

chimp.LOC112206744-LOC107973052-GGT2.revcompl                                       atcatctatcaaaattatgttcgtttttacagatggatgatgtagcattgcatgcagac 14071
chimp54546-135457.revcompl.Pan.troglodytes.clone.rp43-41g5.GenBank:AC099533.36      ATCATCTATCAAAATTATGTTCGTTTTTTACAGATGGATGATGTAGCATTGCATGCAGAC 15606
human.FAM30B-LOC105372935-GGT2.NCBI.                                                atcatctttcaaaatgatgttcgtttttacagatggatgatgtagcattgcatgcagac 12038
human.FAM230E-LOC105377182-GGT3P.NCBI.                                              atcatctatcaaaatgatgttcatttttacagatggatgatgtagcattgcatgcagac 11104
human.FAM230J-LOC105372942-GGTLC5P.NCBI.rev.compl                                   atcatctatcaaaatgatgttcgtttttacagatggatgatgtagcattgcatgcagac 11487
human.FAM230A.ncRNA.18487127.18500594-GGTLC3.18516335.18518165.rev.compl.REF        atcatctatcaaaatgatgttcgtttttacagatggatgatgtagcattgcatgcagac 522
                                                                                    ******* ******* ****** *************************************

chimp.LOC112206744-LOC107973052-GGT2.revcompl                                       caccttgcttttattacattcatttgttcactgatggttggattatttccaccttttggc 14131
chimp54546-135457.revcompl.Pan.troglodytes.clone.rp43-41g5.GenBank:AC099533.36      CACCTTGCTTTTATTACATTCATTTGTTCACTGATGGTTGGATTATTTCCACCTTTTGGC 15666
human.FAM30B-LOC105372935-GGT2.NCBI.                                                caccttgcttttattacattcatttgttcactgatggttggattatttccaccttttggc 12098
human.FAM230E-LOC105377182-GGT3P.NCBI.                                              caccttgcttttattacattcatttgttcactgatggttggattatttccaccttttggc 11164
human.FAM230J-LOC105372942-GGTLC5P.NCBI.rev.compl                                   caccttgcttttattacattcatttgttcactgatggttggattatttccaccttttggc 11547
human.FAM230A.ncRNA.18487127.18500594-GGTLC3.18516335.18518165.rev.compl.REF        caccttgcttttattacattcatttgttcactgatggttggattatttccaccttttggc 582
                                                                                    ************************************************************

chimp.LOC112206744-LOC107973052-GGT2.revcompl                                       tcctgtgaaaagtgatgctacaaacattagtatacaaacatctgtttgatttctgttctc 14191
chimp54546-135457.revcompl.Pan.troglodytes.clone.rp43-41g5.GenBank:AC099533.36      TCCTGTGAAAAGTGATGCTACAAACATTAGTATACAAACATCTGTTTGATTTCTGTTCTC 15726
human.FAM30B-LOC105372935-GGT2.NCBI.                                                tcctgtgaaaagtgatgctacaaacattagtatacaaacatctgtttgatttccgttctc 12158
human.FAM230E-LOC105377182-GGT3P.NCBI.                                              tcctgtgaaaagtgatgctacaaacattagtatacaaacatctgtttgatttctgttctc 11224
human.FAM230J-LOC105372942-GGTLC5P.NCBI.rev.compl                                   tcctgtgaaaagtgatgctacaaacattagtatacaaacatctgtttgatttctgttctc 11607
human.FAM230A.ncRNA.18487127.18500594-GGTLC3.18516335.18518165.rev.compl.REF        tcctgtgaaaagtgatgctacaaacattagtatacaaacatctgtttgatttctgttctc 642
```

```
                                                                                ***************************************************** ******

chimp.LOC112206744-LOC107973052-GGT2.revcompl                                   tattctttggggtgcctaagagtagagttcctgggtccaacaggggttctatatttaaac 14251
chimp54546-135457.revcompl.Pan.troglodytes.clone.rp43-41g5.GenBank:AC099533.36  TATTCTTTGGGGTGCCTAAGAGTAGAGTTCCTGGGTCCAACAGGGGTTCTATATTTAACC 15786
human.FAM30B-LOC105372935-GGT2.NCBI.                                            tattctttggggtgcctaagagtagagttcctgggtccaacaggggttctatatttaacc 12218
human.FAM230E-LOC105377182-GGT3P.NCBI.                                          tattctttggggtgcctaggagtagagttcctgggtccaacaggggttctatatttaacc 11284
human.FAM230J-LOC105372942-GGTLC5P.NCBI.rev.compl                               tattctttggggtgcctaagagtagagttcctgggtccaacaggggttctatatttaacc 11667
human.FAM230A.ncRNA.18487127.18500594-GGTLC3.18516335.18518165.rev.compl.REF    tattctttggggtgcctaagagtagagttcctgggtccaacaggggttctatatttaacc 702
                                                                                ****************** **************************************** *

chimp.LOC112206744-LOC107973052-GGT2.revcompl                                   ttctgagccactgcagactgtttttcagagtggctgcaactttatccatttctaccatca 14311
chimp54546-135457.revcompl.Pan.troglodytes.clone.rp43-41g5.GenBank:AC099533.36  TTCTGAGCCACTGCAGACTGTTTTTCACAGCGGCTGCAACTTTATCCATTTCTACCATCA 15846
human.FAM30B-LOC105372935-GGT2.NCBI.                                            ttctgagccactgcagactgtttttcacagtggctgcaactttatccatttctaccatca 12278
human.FAM230E-LOC105377182-GGT3P.NCBI.                                          ttctgagccactgcagactgtttttcacagtggctgcaactttatccatttctaccatca 11344
human.FAM230J-LOC105372942-GGTLC5P.NCBI.rev.compl                               ttctgagccactgcagactgtttttcacagtggctgcaactttatccatttctaccatca 11727
human.FAM230A.ncRNA.18487127.18500594-GGTLC3.18516335.18518165.rev.compl.REF    ttctgagccactgcagactgtttttcacagtggctgcaactttatccatttctaccatca 762
                                                                                *************************** ** *****************************

chimp.LOC112206744-LOC107973052-GGT2.revcompl                                   atgtatcagggttacaatttctttacgcccttcttcacacttatttcctttaaatcatc 14371
chimp54546-135457.revcompl.Pan.troglodytes.clone.rp43-41g5.GenBank:AC099533.36  ATGTATCAGGGTTACAATTTCTTTACATCCTTGTTCACACTTATTTCCTTTAAATCATC 15906
human.FAM30B-LOC105372935-GGT2.NCBI.                                            atgtatcagggttacaatttctttacgtccttcttcacacttatttcctttaaatcatc 12338
human.FAM230E-LOC105377182-GGT3P.NCBI.                                          atgtatcagggttacaatttctttacatccttgttcacacttatttcctttaaatcatc 11404
human.FAM230J-LOC105372942-GGTLC5P.NCBI.rev.compl                               atgtatcagggttacaatttctttacatccttgttcacacttatttcctttaaatcatc 11787
human.FAM230A.ncRNA.18487127.18500594-GGTLC3.18516335.18518165.rev.compl.REF    atgtatcagggttacaatttctttacgtccttgttcacacttatttcctttaaatcatc 822
                                                                                **************************  **** ***************************

chimp.LOC112206744-LOC107973052-GGT2.revcompl                                   ctagtaggtgtatattggtggctgcttgtgtttctcatttgcatttccctaatgactaat 14431
chimp54546-135457.revcompl.Pan.troglodytes.clone.rp43-41g5.GenBank:AC099533.36  CTAGTAGGTGTATATTGGTGGCTGCTTGTGTTTTTCATTTGCATTTCCCTAATGACTAAT 15966
human.FAM30B-LOC105372935-GGT2.NCBI.                                            ctagtaggtgtatattggtggctgcttgtgtttttcatttgcatttccctaatgactaat 12398
human.FAM230E-LOC105377182-GGT3P.NCBI.                                          ctagtaggtgtatattggtggctgcttgtgtttttcatttgcatttccctaatgactaat 11464
human.FAM230J-LOC105372942-GGTLC5P.NCBI.rev.compl                               ctagtaggtgtatattggtggctgcttgtgtttttcatttgcatttccctaatgactaat 11847
human.FAM230A.ncRNA.18487127.18500594-GGTLC3.18516335.18518165.rev.compl.REF    ctagtaggtgtatattggtggctgcttgtgtttttcatttgcatttccctaatgactaat 882
                                                                                ********************************* **************************

chimp.LOC112206744-LOC107973052-GGT2.revcompl                                   gatcctgagcagcttttcctgtgctactatctgtggctgtatcttctttagggaaatata 14491
chimp54546-135457.revcompl.Pan.troglodytes.clone.rp43-41g5.GenBank:AC099533.36  GATCCTGAGCAGCTTTTCCTGTGCTACTATCTGTGGCTGTATCTTCTTTAGGGAAATATA 16026
human.FAM30B-LOC105372935-GGT2.NCBI.                                            gatcctgagcagcttttcctgtgctactatctgtggctgtatcttctttagggaaatata 12458
human.FAM230E-LOC105377182-GGT3P.NCBI.                                          gatcctgagcagcttttcctgtgctactatttgtggctatatcttctttagggaaatata 11524
human.FAM230J-LOC105372942-GGTLC5P.NCBI.rev.compl                               gatcctgagcagcttttcctgtgctactatttgtggctatatcttctttagggaaatata 11907
human.FAM230A.ncRNA.18487127.18500594-GGTLC3.18516335.18518165.rev.compl.REF    gatcctgagcagcttttcctgtgctactatctgtggctgtatcttctttagggaaatata 942
                                                                                ****************************** ******* *********************

chimp.LOC112206744-LOC107973052-GGT2.revcompl                                   tgttgaagtcttttgcccatttttaaagagttgtctgattttatttagttagtttgttg 14551
chimp54546-135457.revcompl.Pan.troglodytes.clone.rp43-41g5.GenBank:AC099533.36  TGTTGAAGTCTTTTGCCCATTTTTAAAGAGTTGTCTGATTTTGTTTAGTTAGTTTGTTG 16086
human.FAM30B-LOC105372935-GGT2.NCBI.                                            tgttgaagtcttttgcccatttttaaagagttgtctgattttatttagttagtttgttg 12518
human.FAM230E-LOC105377182-GGT3P.NCBI.                                          tgttgaagtcttttgcccatttttaaagagttgtctgattttatttagttagtttgttg 11584
human.FAM230J-LOC105372942-GGTLC5P.NCBI.rev.compl                               tgttgaagtcttttgcccatttttaaagagttgtctgattttatttagttagtttgttg 11967
human.FAM230A.ncRNA.18487127.18500594-GGTLC3.18516335.18518165.rev.compl.REF    tgttgaagtcttttgcccatttttaaagagttgtctgattttatttagttagtttgttg 1002
                                                                                ***************************************** ******************

chimp.LOC112206744-LOC107973052-GGT2.revcompl                                   ttgtagattttgaatatatcttaaatatatttaaaatagattctaaattttagtctctt 14611
chimp54546-135457.revcompl.Pan.troglodytes.clone.rp43-41g5.GenBank:AC099533.36  TTGTAGATTTTGAATATATCTTAAATATATTTAAAATAGATTCTAAATTTTAGTCTCTT 16146
human.FAM30B-LOC105372935-GGT2.NCBI.                                            ttgtagattttgaatatatcttaaatatatttaaaatatattctaaattttagtctctt 12578
human.FAM230E-LOC105377182-GGT3P.NCBI.                                          ctgtagattttgaatatatcttaaatatatttaaaa-atattctaaattttagtctctt 11643
human.FAM230J-LOC105372942-GGTLC5P.NCBI.rev.compl                               ctgtagattttgaatatatcttaaatatatttaaaa-atattctaaattttagtctctt 12026
human.FAM230A.ncRNA.18487127.18500594-GGTLC3.18516335.18518165.rev.compl.REF    ctgtagattttgaatatatcttaaatatatttaaaa-atattctaaattttagtctctt 1061
                                                                                 ********************************* * ***********************

chimp.LOC112206744-LOC107973052-GGT2.revcompl                                   acaagataaatgatttgcaaatatttccacctttgtgtagaactttagattcacaaactt 14671
chimp54546-135457.revcompl.Pan.troglodytes.clone.rp43-41g5.GenBank:AC099533.36  ACAAGATAAATGATTTGCAAATATTTCCCCCTTTGTGTAGAACTTTAGATTCACAAACTT 16206
human.FAM30B-LOC105372935-GGT2.NCBI.                                            acaagagaaatgatttgcaaatatttccacctttgtgtagaactttagattcacaaactt 12638
human.FAM230E-LOC105377182-GGT3P.NCBI.                                          acaagataaatgatttgcaaatatttccccctttgtgtagaactttagattcacaaactt 11703
human.FAM230J-LOC105372942-GGTLC5P.NCBI.rev.compl                               acaagataaatgatttgcaaatatttccccctttgtgtagaactttagattcacaaactt 12086
human.FAM230A.ncRNA.18487127.18500594-GGTLC3.18516335.18518165.rev.compl.REF    acaagataaatgatttgcaaatatttccccctttgtgtagaactttagattcacaaactt 1121
                                                                                ****** ********************* *******************************

chimp.LOC112206744-LOC107973052-GGT2.revcompl                                   cattaatttgtgtgaaatcctcagcagttgaccccaaacagataagactgaagcagtatt 14731
chimp54546-135457.revcompl.Pan.troglodytes.clone.rp43-41g5.GenBank:AC099533.36  CATTAATTTGTATGAAATCCTCAGCAGTTGACCCCAAAAGGATAAGACTGAAGCAGTATT 16266
human.FAM30B-LOC105372935-GGT2.NCBI.                                            cgttaatttgtatgaaatcctcagcagttgaccccaaacagataagactgaagcagtatt 12698
human.FAM230E-LOC105377182-GGT3P.NCBI.                                          cattaatttgtatgaaatcctcagcagttgaccccaaacagataagactgaagcagtatt 11763
human.FAM230J-LOC105372942-GGTLC5P.NCBI.rev.compl                               cattaatttgtatgaaatcctcagcagttgaccccaaacagataagactgaagcagtatt 12146
human.FAM230A.ncRNA.18487127.18500594-GGTLC3.18516335.18518165.rev.compl.REF    cattaatttgtatgaaatcctcagcagttgaccccaaacagataagactgaagcagtatt 1181
                                                                                * ********* **************************  ********************

chimp.LOC112206744-LOC107973052-GGT2.revcompl                                   ttaggaatagttgaaagtatgatcaccacaaaacataagcgtaatcaaatcctgcaagct 14791
chimp54546-135457.revcompl.Pan.troglodytes.clone.rp43-41g5.GenBank:AC099533.36  TTAGGAATAGTTGAAAGTATGATCACCACAAAACATAAGCGTAATCAAATCCTGCAAGCT 16326
human.FAM30B-LOC105372935-GGT2.NCBI.                                            ttaggaatagttgaaagtatgatcaccacaaaacataagcgtaatcaaatcctgcaagct 12758
human.FAM230E-LOC105377182-GGT3P.NCBI.                                          ttaggaatagttgaaagtatgatcaccacaaaacataagcgtaatcaaatcctgcaagct 11823
human.FAM230J-LOC105372942-GGTLC5P.NCBI.rev.compl                               ttaggaatagttgaaagtatgatcaccacaaaacataagcgtaatcaaatcctgcaagct 12206
human.FAM230A.ncRNA.18487127.18500594-GGTLC3.18516335.18518165.rev.compl.REF    ttaggaatagttgaaagtatgatcaccacaaaacataagcgtaatcaaatcctgcaagct 1241
                                                                                ************************************************************

chimp.LOC112206744-LOC107973052-GGT2.revcompl                                   acatgtaaagcacaatgacaaataaggcagcaaagggccatctggtgattagttcaccac 14851
chimp54546-135457.revcompl.Pan.troglodytes.clone.rp43-41g5.GenBank:AC099533.36  ACATGTAAGGCACAATGACAAATAAGGCAGCAAAGGGCCATCTGGTGATTAGTTCACCAC 16386
human.FAM30B-LOC105372935-GGT2.NCBI.                                            acatgtaaggcacaatgacaaataaggcagcaaagggccatctggtgattagttcaccac 12818
human.FAM230E-LOC105377182-GGT3P.NCBI.                                          acatgtaaggcacaatgacaaataaggcagcaaagggccatctggtgattagttcaccac 11883
human.FAM230J-LOC105372942-GGTLC5P.NCBI.rev.compl                               acatgtaaggcacaatgacaaataaggcagcaaagggccatctggtgattagttcaccac 12266
human.FAM230A.ncRNA.18487127.18500594-GGTLC3.18516335.18518165.rev.compl.REF    acatgtaaggcacaatgacaaataaggcagcaaagggccatctggtgattagttcaccac 1301
                                                                                ******** ***************************************************

chimp.LOC112206744-LOC107973052-GGT2.revcompl                                   acttgttgcaactgtttgcgctgcagagttaaaacacaccagcattcaacccatgtctcc 14911
chimp54546-135457.revcompl.Pan.troglodytes.clone.rp43-41g5.GenBank:AC099533.36  ACTTGTTGCAACTGTTTGTGCTGCAGAGTTAAAACACACCAGCATTCAACCCATGTCTCC 16446
human.FAM30B-LOC105372935-GGT2.NCBI.                                            acttgttgcaactgtttgcgctgcagagttaaaacacaccagcattcaacccgtgtctcc 12878
human.FAM230E-LOC105377182-GGT3P.NCBI.                                          acttgttgcaactgtttgcgctgcagagttaaaacataccagcattcaacccatgtctcc 11943
human.FAM230J-LOC105372942-GGTLC5P.NCBI.rev.compl                               acttgttgcaactgtttgcgctgcagagttaaaacataccagcattcaacccatgtctcc 12326
human.FAM230A.ncRNA.18487127.18500594-GGTLC3.18516335.18518165.rev.compl.REF    acttgttgcaactgtttgcactgcagagttaaaacataccagcattcaacccatgtctcc 1361
                                                                                ******************  **************** *************** *******

chimp.LOC112206744-LOC107973052-GGT2.revcompl                                   tctcttgaggtaaactgtcgtatgttggctggcctgaacaagcgtagatattctccatcc 14971
chimp54546-135457.revcompl.Pan.troglodytes.clone.rp43-41g5.GenBank:AC099533.36  TCTCTTGAAGTAAACTGTCGTATGTTGGCTGGCCTGAACAAGCGTAGATATTCTCCATCC 16506
human.FAM30B-LOC105372935-GGT2.NCBI.                                            tctcttgaagtaaactgtcgtatgttggctggcctgaacaagcgtagatattctccatcc 12938
human.FAM230E-LOC105377182-GGT3P.NCBI.                                          tgtcttgaagtaaactgtcgtatgttggctggcctgaacaagcgtagatattctccatcc 12003
human.FAM230J-LOC105372942-GGTLC5P.NCBI.rev.compl                               tgtcttgaagtaaactgtcgtatgttggctggcctgaacaagcgtagatattctccatcc 12386
human.FAM230A.ncRNA.18487127.18500594-GGTLC3.18516335.18518165.rev.compl.REF    tgtcttgaagtaaactgtcgtatgttggctggcctgaacaagcgtagatattctccatcc 1421
                                                                                * ****** ***************************************************

chimp.LOC112206744-LOC107973052-GGT2.revcompl                                   tcaattaatatgcatgcatgacaaagaaaaggaggcctggatgaaaaaatattgtgtgat 15031
chimp54546-135457.revcompl.Pan.troglodytes.clone.rp43-41g5.GenBank:AC099533.36  TCAATTAATATGCATGCATGACAAAGAAAAGGAGGCCTGGATGAAAAAATATTGTGTGAT 16566
human.FAM30B-LOC105372935-GGT2.NCBI.                                            tcaattaatatgcatgcatgacaaagaaaaggaggcctggatgaaaaaatattgtgtgat 12998
human.FAM230E-LOC105377182-GGT3P.NCBI.                                          tcagttaatatgcatgcatgacaaagaaaaggaggcctggatgaaaaaatattgtgtgat 12063
human.FAM230J-LOC105372942-GGTLC5P.NCBI.rev.compl                               tcaattaatatgcatgcatgacaaagaaaaggaggcctggatgaaaaaatattgtgtgat 12446
human.FAM230A.ncRNA.18487127.18500594-GGTLC3.18516335.18518165.rev.compl.REF    tcaattaatatgcatgcatgacaaagaaaaggaggcctggatgaaaaaatactgtgtgat 1481
                                                                                *** *********************************************** ********

chimp.LOC112206744-LOC107973052-GGT2.revcompl                                   taataattatgctttaattaattttaaaggatataatttcagtacttctaattctcccat 15091
chimp54546-135457.revcompl.Pan.troglodytes.clone.rp43-41g5.GenBank:AC099533.36  T-ATAATTATGCTTTAATTAATTTTAAAGGATATAATTTCAGTACTTCTAATTCTCCCAT 16625
human.FAM30B-LOC105372935-GGT2.NCBI.                                            taataattatgctttaattaattttaaaggatataatttcagtacttctaattctcccat 13058
human.FAM230E-LOC105377182-GGT3P.NCBI.                                          taataattatgctttaattaattttaaaggatataatttcagtacttctaattctcccat 12123
human.FAM230J-LOC105372942-GGTLC5P.NCBI.rev.compl                               taataattatgctttaattaattttaaaggatataatttcagtacttctaattctcccat 12506
human.FAM230A.ncRNA.18487127.18500594-GGTLC3.18516335.18518165.rev.compl.REF    taataattatgctttaattaattttaaaggatataatttcagtacttctaattctcccat 1541
                                                                                * **********************************************************

chimp.LOC112206744-LOC107973052-GGT2.revcompl                                   cagcagttataacaaaggattagtgaataaataccatagactgttttgcctggaattgaa 15151
chimp54546-135457.revcompl.Pan.troglodytes.clone.rp43-41g5.GenBank:AC099533.36  CAGCAGTTATAACAAAGGATTAGTGAATAAATACCATAGACTGTTTTGCCTAGAATTGAA 16685
human.FAM30B-LOC105372935-GGT2.NCBI.                                            cagcagttataacaaaggattagtgaataaataccatagactgttttgcctagaattgaa 13118
human.FAM230E-LOC105377182-GGT3P.NCBI.                                          cagcagttataacaaaggattagtgaataaataccatagactgttttgcctagaattgaa 12183
human.FAM230J-LOC105372942-GGTLC5P.NCBI.rev.compl                               cagcagttataacaaaggattagtgaataaataccatagactgttttgcctagaattgaa 12566
human.FAM230A.ncRNA.18487127.18500594-GGTLC3.18516335.18518165.rev.compl.REF    cagcagttataacaaaggattagtgaataaataccatagactgttttgcctagaattgaa 1601
                                                                                *************************************************** ********

chimp.LOC112206744-LOC107973052-GGT2.revcompl                                   tccaatctatctattaaactttgcttttattcaagtgcaaaatgctaaaacac---ataa 15208
chimp54546-135457.revcompl.Pan.troglodytes.clone.rp43-41g5.GenBank:AC099533.36  TCCAATCTATCTATTAAACTTTGCTTTTATTCAAGTGCAAAATGCTAAAACAC---ATAA 16742
human.FAM30B-LOC105372935-GGT2.NCBI.                                            tccaatctatctattaaactttgcttgtattcaagtgcaaaatgctaaaacac---ataa 13175
human.FAM230E-LOC105377182-GGT3P.NCBI.                                          tccaatctgtctattaaactttgcttttattcaagtgcaaaatgctaaaacacataataa 12243
human.FAM230J-LOC105372942-GGTLC5P.NCBI.rev.compl                               tccaatctgtctattaaactttgcttttattcaagtgcaaaatgctaaaacacataataa 12626
human.FAM230A.ncRNA.18487127.18500594-GGTLC3.18516335.18518165.rev.compl.REF    tccaacctgtctattaaactttgcttttattcaagtgcaaaatgctaaaacacataataa 1661
                                                                                ***** ** ***************** **************************   ****

chimp.LOC112206744-LOC107973052-GGT2.revcompl                                   ctgcagtgacagccactgtggatcctcagaggtaaaagtagtcttgggacataaatcctg 15268
chimp54546-135457.revcompl.Pan.troglodytes.clone.rp43-41g5.GenBank:AC099533.36  CTGCAGTGACAGCCACTGTGGATCCTCAGAGGTAAAAGTAGTCTTGGGACATAAATCCTG 16802
human.FAM30B-LOC105372935-GGT2.NCBI.                                            ctgcagtgacagccactgtggatcctcagaggtaaaagtagtcttgggacataaatcctg 13235
human.FAM230E-LOC105377182-GGT3P.NCBI.                                          ctgcagtgacagccactgtggatcctcagaggtaaaagtagtcttgggacataaatcctg 12303
human.FAM230J-LOC105372942-GGTLC5P.NCBI.rev.compl                               ctgcagtgacagccactgtggatcctcagaggtaaaagtagtcttgggacataaatcctg 12686
human.FAM230A.ncRNA.18487127.18500594-GGTLC3.18516335.18518165.rev.compl.REF    ctgcagtgacagccactgtggatcctcagaggtaaaagtagtcttgggacataaatcctg 1721
                                                                                ************************************************************
```

```
chimp.LOC112206744-LOC107973052-GGT2.revcompl                               caagctaatattgtttttacaggtttagaaaaccatttagctgggtttcaaacctcacag 15328
chimp54546-135457.Pan.troglodytes.clone.rp43-41g5.GenBank:AC099533.36       CAAGCTAATATTGTTTTTACAGGTTTAGAAAACCATTTAGCTGGGTTTCAAACCTCACAG 16862
human.FAM30B-LOC105372935-GGT2.NCBI.                                        caagctaatattgtttttacaggtttagaaaaccatttagctgggtttcaaacctcacag 13295
human.FAM230E-LOC105377182-GGT3P.NCBI.                                      caag-taatattgtttttacaggtttagaaaaccatttagctgggtttcaaacctcacag 12362
human.FAM230J-LOC105372942-GGTLC5P.NCBI.rev.compl                           caag-taatattgtttttacaggtttagaaaaccatttagctgggtttcaaacctcacag 12745
human.FAM230A.ncRNA.18487127.18500594-GGTLC3.18516335.18518165.rev.compl.REF caag-taatattgtttttacaggtttagaaaaccatttagctgggtttcaaacctcacag 1780
                                                                            **** *******************************************************

chimp.LOC112206744-LOC107973052-GGT2.revcompl                               tgtgagcagtgggactctcatcaaactatagcatgtgcttcagtaccatttgtagactga 15388
chimp54546-135457.revcompl.Pan.troglodytes.clone.rp43-41g5.GenBank:AC099533.36 TGTGAGCAGTGGGACTCTCATCAAACTATAGCATGTGCTTCAGTACCATTTGTAGACTGA 16922
human.FAM30B-LOC105372935-GGT2.NCBI.                                        tgtgagcagtgggactctcatcaaactatagcatgtgcttcagtaccatttgtagactga 13355
human.FAM230E-LOC105377182-GGT3P.NCBI.                                      tgtgagcagtgggactctcatcaaactatagcatgtgcttcagtaccatttgtagactga 12422
human.FAM230J-LOC105372942-GGTLC5P.NCBI.rev.compl                           tgtgagcagtgggactctcatcaaactatagcatgtgcttcagtaccatttgtagactga 12805
human.FAM230A.ncRNA.18487127.18500594-GGTLC3.18516335.18518165.rev.compl.REF tgtgagcagtgggactctcatcaaactatagcatgtgcttcagtaccatttgtagactga 1840
                                                                            ************************************************************

chimp.LOC112206744-LOC107973052-GGT2.revcompl                               ttcattcccattttcttaagttgccatcagcaaaatgccagggactctatttcttgctcc 15448
chimp54546-135457.revcompl.Pan.troglodytes.clone.rp43-41g5.GenBank:AC099533.36 CTCATTCCCATTTTCTTAAGTTGCCATCAGCAAAATGCCAGGGACTCTATTTCTTGCTCC 16982
human.FAM30B-LOC105372935-GGT2.NCBI.                                        ctcattcccattttcttcagttgccatcagcaaaatgccagggactctatttcttgctcc 13415
human.FAM230E-LOC105377182-GGT3P.NCBI.                                      ctcattcccattgccttaagttgccatcagcaaaatgccagggactctatttcttgctcc 12482
human.FAM230J-LOC105372942-GGTLC5P.NCBI.rev.compl                           ctcattcccattgccttaagttgccatcagcaaaatgccagggactctatttcttgctcc 12865
human.FAM230A.ncRNA.18487127.18500594-GGTLC3.18516335.18518165.rev.compl.REF ctcattcccattgccttaagttgccatcagcaaaatgccagggactctatttcttgctcc 1900
                                                                             ***********  *** ******************************************

chimp.LOC112206744-LOC107973052-GGT2.revcompl                               ttagctcctcgttcttgcctgtctttccacgagggaagatttctagcaggagctcaagc 15508
chimp54546-135457.revcompl.Pan.troglodytes.clone.rp43-41g5.GenBank:AC099533.36 TTAGCTCCTCGTTCTTGCCTGTCTTTCCACGAGGGAAGATTTCTAGCAGGAGCTCAAGC 17042
human.FAM30B-LOC105372935-GGT2.NCBI.                                        ttagctcctcgttcttgcctgtctttccacgagggaagatttctagcaggagctcaagc 13475
human.FAM230E-LOC105377182-GGT3P.NCBI.                                      ttagctcctcgttcttgcctgtctttccacgagggaggatttctagcaggagctcaagc 12542
human.FAM230J-LOC105372942-GGTLC5P.NCBI.rev.compl                           ttagctcctcgttcttgcctgtctttccacgagggaggatttctagcaggagctcaagc 12925
human.FAM230A.ncRNA.18487127.18500594-GGTLC3.18516335.18518165.rev.compl.REF ttagctcctcgttcttgcctgtctttccacgagggaggatttctagcaggagctcaagc 1960
                                                                            ************************************ ***********************

chimp.LOC112206744-LOC107973052-GGT2.revcompl                               tgtgcttttaatgaaacacatccacacacactgtcctgttgtccacattaagcagagctc 15568
chimp54546-135457.revcompl.Pan.troglodytes.clone.rp43-41g5.GenBank:AC099533.36 TGTGCTTTTAATGAAACACATCCACACACACTGTCCTGTTGTCCACATTAAGCAGAGCTC 17102
human.FAM30B-LOC105372935-GGT2.NCBI.                                        tgtgcttttaatgaaacacatccacacacactgtcctgttgtccacattaagcagagctc 13535
human.FAM230E-LOC105377182-GGT3P.NCBI.                                      tgtgcttttaatgaaacacatccacacacactgtcctgttgtccacattaagcagagctc 12602
human.FAM230J-LOC105372942-GGTLC5P.NCBI.rev.compl                           tgtgcttttaatgaaacacatccacacacactgtcctgttgtccacattaagcagagctc 12985
human.FAM230A.ncRNA.18487127.18500594-GGTLC3.18516335.18518165.rev.compl.REF tgtgcttttaatgaaacacatccacacacactgtcctgttgtccacattaagcagagctc 2020
                                                                            ************************************************************

chimp.LOC112206744-LOC107973052-GGT2.revcompl                               cctgaataactcatgaacaaaagcatctatgactaactgttgctctgtgtcctcctagcc 15628
chimp54546-135457.revcompl.Pan.troglodytes.clone.rp43-41g5.GenBank:AC099533.36 CCTGAATAACTCATGAACAAAAGCATCTATGACTAACTGTTGCTCTGTGTCCTCCTAGCC 17162
human.FAM30B-LOC105372935-GGT2.NCBI.                                        cctgaataactcatgaacaaaagcatctatgactaactgttgctctgtgtcctcctagcc 13595
human.FAM230E-LOC105377182-GGT3P.NCBI.                                      cctgaatagctcatgaacaaaagcatctatgactaactgttgctctgtgtcctcctagcc 12662
human.FAM230J-LOC105372942-GGTLC5P.NCBI.rev.compl                           cctgaatagctcatgaacaaaagcatctatgactaactgttgctctgtgtcctcctagcc 13045
human.FAM230A.ncRNA.18487127.18500594-GGTLC3.18516335.18518165.rev.compl.REF cctgaataactcatgaacaaaagcatctatgactaactgttgctctgtgtcctcctagcc 2080
                                                                            ******** ***************************************************

chimp.LOC112206744-LOC107973052-GGT2.revcompl                               tctgaggagtctctagttcacaaggacagaggagatggagagaggccagtcaacgcgagg 15688
chimp54546-135457.revcompl.Pan.troglodytes.clone.rp43-41g5.GenBank:AC099533.36 TCTGAGGAGTCTCTAGTTCACAAGGACAGAGGAGATGGAGAGAGGCCAGTCAACGCGAGG 17222
human.FAM30B-LOC105372935-GGT2.NCBI.                                        tctgaggagtctctagttcacaaggacagaggagatggagagaggccagtcaacgcgagg 13655
human.FAM230E-LOC105377182-GGT3P.NCBI.                                      tctgaggagtctctagttcacaaggacagaggagatggagagaggccagtcaacgtgagg 12722
human.FAM230J-LOC105372942-GGTLC5P.NCBI.rev.compl                           tctgaggagtctctagttcacaaggacagaggagatggagagaggccagtcaacgtgaag 13105
human.FAM230A.ncRNA.18487127.18500594-GGTLC3.18516335.18518165.rev.compl.REF tctgaggagtctctagttcacaaggacagaggagatggagagaggccagtcaacgtgagg 2140
                                                                            ****************************************************** ** *

chimp.LOC112206744-LOC107973052-GGT2.revcompl                               gtaaggttgccttgctttctctgaaatagaaatgttcctttcttggtgtctttctttttc 15748
chimp54546-135457.revcompl.Pan.troglodytes.clone.rp43-41g5.GenBank:AC099533.36 GTAAGGTTGCCTTGCTTTCTCTGAAATAGAAATGTTCCTTTCTTGGTGTCTTTCTTTTTC 17282
human.FAM30B-LOC105372935-GGT2.NCBI.                                        gtaaggttgccttgctttctctgaaatagaaatgttcctttcttggtgtctttctttttc 13715
human.FAM230E-LOC105377182-GGT3P.NCBI.                                      gtaaggttgccttgctttctctgaaatagaaatgttcctttcttgttgtctttctttttc 12782
human.FAM230J-LOC105372942-GGTLC5P.NCBI.rev.compl                           gtaaggttgccttgctttctctgaaatagaaatgttcctttcttgttgtctttctttttc 13165
human.FAM230A.ncRNA.18487127.18500594-GGTLC3.18516335.18518165.rev.compl.REF gtaaggttgccttgctttctctgaaatagaaatgttcctttcttgttgtctttctttttc 2200
                                                                            ********************************************* **************

chimp.LOC112206744-LOC107973052-GGT2.revcompl                               aactgactttacatgtgaaaaaatgacaatgtccatgacaggtattaaatgcagttttct 15808
chimp54546-135457.revcompl.Pan.troglodytes.clone.rp43-41g5.GenBank:AC099533.36 AACTGACTTTACATGTGAAAAAATGACAATGTCCATGACAGGTATTAAATGCAGTTTTCT 17342
human.FAM30B-LOC105372935-GGT2.NCBI.                                        aactgactttacatgtgaaaaaatgacaatgtccatgacaggtattaaatgcagttttct 13775
human.FAM230E-LOC105377182-GGT3P.NCBI.                                      aactgactttacatgtgaaaagatgacaatgtccatgacaggtattaaatgcagttttct 12842
human.FAM230J-LOC105372942-GGTLC5P.NCBI.rev.compl                           aactgactttacatgtgaaaagatgacaatgtccatgacaggtattaaatgcagttttct 13225
human.FAM230A.ncRNA.18487127.18500594-GGTLC3.18516335.18518165.rev.compl.REF aactgactttacatgtgaaaagatgacaatgtccatgacaggtattaaatgcagttttct 2260
                                                                            ********************* **************************************

chimp.LOC112206744-LOC107973052-GGT2.revcompl                               gaggggaggaagaagtgactcttagcaactgatatgtaatccaaaatgacatttagcta 15868
chimp54546-135457.revcompl.Pan.troglodytes.clone.rp43-41g5.GenBank:AC099533.36 GAGGGGAGGAAGAAGTGACTCTTAGCAACTGATATGTAATCCAAAATGGCATTTAGCTA 17402
human.FAM30B-LOC105372935-GGT2.NCBI.                                        gaggggaggaagaagtgactcttagcaactgatatgtaatccaaaatggcatttagcta 13835
human.FAM230E-LOC105377182-GGT3P.NCBI.                                      gaggggaggaagaagtgactcttagcaactgatatgtaatccaaaatggcatttagcta 12902
human.FAM230J-LOC105372942-GGTLC5P.NCBI.rev.compl                           gaggggaggaagaagtgactcttagcaactgatatgtaatccaaaatggcatttagcta 13285
human.FAM230A.ncRNA.18487127.18500594-GGTLC3.18516335.18518165.rev.compl.REF gaggggaggaagaagtgactcttagcaactgatatgtaatacaaaatggcatttagcta 2320
                                                                            **************************************** ******* **********

chimp.LOC112206744-LOC107973052-GGT2.revcompl                               tgacggcttcaggttgtagactgtatccttggggtccttgtccttggaagcaatgtcttc 15928
chimp54546-135457.revcompl.Pan.troglodytes.clone.rp43-41g5.GenBank:AC099533.36 TGACGGCTTCAGGTTGTAGACTGTATCCTTGGGGTCCTTGTCCTTGGAAGCAATGTCTTC 17462
human.FAM30B-LOC105372935-GGT2.NCBI.                                        tgacggcttcaggttgtggactgtatccttggggtccttgtccttggaagcaatgtcttc 13895
human.FAM230E-LOC105377182-GGT3P.NCBI.                                      tgatggcttcaggttgtagactgtatccttggggtccttgtccttggaagcaatgtcttc 12962
human.FAM230J-LOC105372942-GGTLC5P.NCBI.rev.compl                           tgatggcttcaggttgtagactgtatccttggggtccttgtccttggaagcaatgtcttc 13345
human.FAM230A.ncRNA.18487127.18500594-GGTLC3.18516335.18518165.rev.compl.REF tgatggcttcaggttgtagactgtatccttggggtccttgtccttggaagcaatgtcttc 2380
                                                                            *** ************ *******************************************

chimp.LOC112206744-LOC107973052-GGT2.revcompl                               tccttggattcagtattttgcacttgccaacctacgtggacctgagagatccaccatcca 15988
chimp54546-135457.revcompl.Pan.troglodytes.clone.rp43-41g5.GenBank:AC099533.36 TCCTTGGATTCAGTATTTTGCACTTGCCAACCTACGTGGACCTGAGAGATCCACCATCCA 17522
human.FAM30B-LOC105372935-GGT2.NCBI.                                        tccttggattcaggattttgcacttgccaacctacgtggacctgagagatccaccatcca 13955
human.FAM230E-LOC105377182-GGT3P.NCBI.                                      tccttggattcagtattttgcacttgccaacctacgtggacctgagagatccgccatcca 13022
human.FAM230J-LOC105372942-GGTLC5P.NCBI.rev.compl                           tccttggattcagtattttgcacttgccaacctacgtggacctgagagatccgccatcca 13405
human.FAM230A.ncRNA.18487127.18500594-GGTLC3.18516335.18518165.rev.compl.REF tccttggattcagtattttgcacttgccaacctacgtggacctgagagatccaccatcca 2440
                                                                            ************* ************************************** *******

chimp.LOC112206744-LOC107973052-GGT2.revcompl                               gaagctgatgtcttttccagtgtgtagcctacccttgttttggaggccttgaagttgact 16048
chimp54546-135457.revcompl.Pan.troglodytes.clone.rp43-41g5.GenBank:AC099533.36 GAAGCTGATGTCTTTTCCAGTGTGTAGCCTACCCTTGTTTTGGAGGCCTTGAAGTTGACT 17582
human.FAM30B-LOC105372935-GGT2.NCBI.                                        gaagctgatgtcttttccagtgtgtatcctacccttgttttggaggccttgaagttgact 14015
human.FAM230E-LOC105377182-GGT3P.NCBI.                                      gaagctgatgtcttttccagtgtgtatcctacccttgttttggaggccttgaagttgact 13082
human.FAM230J-LOC105372942-GGTLC5P.NCBI.rev.compl                           gaagctgatgtcttttccagtgtgtatcctacccttgttttggaggccttgaagttgact 13465
human.FAM230A.ncRNA.18487127.18500594-GGTLC3.18516335.18518165.rev.compl.REF gaagctgatgtcttttccagtgtgtatcctacccttgttttggaggccttgaagttgact 2500
                                                                            ************************* **********************************

chimp.LOC112206744-LOC107973052-GGT2.revcompl                               acactttctgatcaagttttcagtattcattgagagaaacagagccttgtgcaaacaatc 16108
chimp54546-135457.revcompl.Pan.troglodytes.clone.rp43-41g5.GenBank:AC099533.36 ACACTTTCTGATCAAGTTTTCAGTATTCATTGAGAGAAACAGAGCCTTGTGCAAACAATC 17642
human.FAM30B-LOC105372935-GGT2.NCBI.                                        acactttctgatcaagttttcagtattcattgagagaaacagagccttgtgcaaacaatc 14075
human.FAM230E-LOC105377182-GGT3P.NCBI.                                      acactttctgatcaagttttcaatattcattgagagaaacacagccttgtgcaaacaatc 13142
human.FAM230J-LOC105372942-GGTLC5P.NCBI.rev.compl                           acactttctgatcaagttttcaatattcattgagagaaacacagccttgtgcaaacaatc 13525
human.FAM230A.ncRNA.18487127.18500594-GGTLC3.18516335.18518165.rev.compl.REF acactttctgatcaagttttcaatattcattgagagaaacacagccttgtgcaaacaatc 2560
                                                                            ********************** ****************** ******************

chimp.LOC112206744-LOC107973052-GGT2.revcompl                               cacaacatgacatacccctcaaaaagctttgtttctgtattgcaggtggtgcaggtggcc 16168
chimp54546-135457.revcompl.Pan.troglodytes.clone.rp43-41g5.GenBank:AC099533.36 CACAACATGACATACCCCTCAAAAAGCTTTGTTTCTGTATTGCAGGTGGTGCAGATGGCC 17702
human.FAM30B-LOC105372935-GGT2.NCBI.                                        cacaacatgagatacccctgaaaaagctttgtttctgtattgcaggtggtgcaggtggcc 14135
human.FAM230E-LOC105377182-GGT3P.NCBI.                                      cacaacatgacatacccctcaaaaagctttgtttctgtattgcaggtggtgcaggtggcc 13202
human.FAM230J-LOC105372942-GGTLC5P.NCBI.rev.compl                           cacaacatgacatacccctcaaaaagctttgtttctgtattgcaggtggtgcaggtggcc 13585
human.FAM230A.ncRNA.18487127.18500594-GGTLC3.18516335.18518165.rev.compl.REF cacaacatgacatacccctcaaaaagctttgtttctgtattgcaggtggtgcaggtggcc 2620
                                                                            ********** ******** ********************************** *****

chimp.LOC112206744-LOC107973052-GGT2.revcompl                               cctctgaggtgtgaatctagtaagtattctggaatcacttgccaagaaaacaatctggat 16228
chimp54546-135457.revcompl.Pan.troglodytes.clone.rp43-41g5.GenBank:AC099533.36 CCTCTGAGGTGTGACTCTAGTAAGTATTCTGGAATCACTTGCCAAGAAAACAATCTGGAT 17762
human.FAM30B-LOC105372935-GGT2.NCBI.                                        cctctgaggcttgaatctagtaagtattctggaatcacttgccaagaaaacaatctggat 14195
human.FAM230E-LOC105377182-GGT3P.NCBI.                                      cctctgaggcgtgaatctagtaagtattctggaatcacttaccaagaaaacaatctggat 13262
human.FAM230J-LOC105372942-GGTLC5P.NCBI.rev.compl                           cctctgaggcgtgaatctagtaagtattctggaatcacttaccaagaaaacaatctggat 13645
human.FAM230A.ncRNA.18487127.18500594-GGTLC3.18516335.18518165.rev.compl.REF cctctgaggcatgaatctagtaagtattctggaatcacttaccaagaaaacaatctggat 2680
                                                                            *********  *** ************************* *******************

chimp.LOC112206744-LOC107973052-GGT2.revcompl                               gccaagaaaggtgtggcatccttgcctggtttcaatgtgaagagcctccctgatcctggg 16288
chimp54546-135457.revcompl.Pan.troglodytes.clone.rp43-41g5.GenBank:AC099533.36 GCCAAGAAAGGTGTGGCATCCTTGCCTGGTTTCAATGTGAAGAGCCTCCCTGATCCTGGG 17822
human.FAM30B-LOC105372935-GGT2.NCBI.                                        gccaagaaaggtgtggcatccttgcctggtttcaatgtgaagagcctccctgatcctggg 14255
human.FAM230E-LOC105377182-GGT3P.NCBI.                                      gccaagaaaggtgtggcatccttgcctggtttcaatgtgaagagccaccctgatcctggg 13322
human.FAM230J-LOC105372942-GGTLC5P.NCBI.rev.compl                           gccaagaaaggtgtggcatccttgcctggtttcaatgtgaagagccaccctgatcctggg 13705
human.FAM230A.ncRNA.18487127.18500594-GGTLC3.18516335.18518165.rev.compl.REF gccaagaaaggtgtggcatccttgcctggtttcaatgtgaagagccaccctgatcctggg 2740
                                                                            ********************************************** *************

chimp.LOC112206744-LOC107973052-GGT2.revcompl                               attgtgataggaataagtataggggaagtgtttttttaaaacctgaattccccagggaaa 16348
chimp54546-135457.revcompl.Pan.troglodytes.clone.rp43-41g5.GenBank:AC099533.36 ATTGTGATAGGAATAAGTATAGGGGAAGTGTTTTTTTAAAACCTGAATTCCCCAGGGAAA 17882
human.FAM30B-LOC105372935-GGT2.NCBI.                                        attgtgataggaataagtataggggaagtgtttttttaaaacctgaattccccagggaaa 14315
human.FAM230E-LOC105377182-GGT3P.NCBI.                                      attgtgataggaataagtataggggaagtgtttttttaaaacctgaattccccagggaaa 13382
human.FAM230J-LOC105372942-GGTLC5P.NCBI.rev.compl                           attgtgataggaataagtataggggaagtgtttttttaaaacctgaattccccagggaaa 13765
human.FAM230A.ncRNA.18487127.18500594-GGTLC3.18516335.18518165.rev.compl.REF attgtgataggaataagtataggggaagtgtttttttaaaacctgaattccccagggaaa 2800
                                                                            ************************************************************
```

```
chimp.LOC112206744-LOC107973052-GGT2.revcompl aattatggccaaattttgaggaagcagttgtgctcccttttgggtggtgccgagttgggt 16408
chimp54546-135457.revcompl.Pan.troglodytes.clone.rp43-41g5.GenBank:AC099533.36 AATTATGGCCAAATTTTGAGGAAGCAGTTGTGCTCCCTTTTGGGTGGTGCTGAGTTGGGT 17942
human.FAM30B-LOC105372935-GGT2.NCBI. aattatggccaaattttgaggaagcagctgtgctcccttttgggtggtgctgagttgggt 14375
human.FAM230E-LOC105377182-GGT3P.NCBI. aattatggccaaattttgaggaagcagctgtgctcccttttgggtggtgctgagttgggt 13442
human.FAM230J-LOC105372942-GGTLC5P.NCBI.rev.compl aattatggccaaattttgaggaagcagctgtgctcccttttgggtggtgctgagttgggt 13825
human.FAM230A.ncRNA.18487127.18500594-GGTLC3.18516335.18518165.rev.compl.REF aattatggccaaattttgaggaagcagctgtgctcccttttgggtggtgctgagttgggt 2860
*************************** ********************** *********

chimp.LOC112206744-LOC107973052-GGT2.revcompl gcttgaggattggtggtgtcttgtgtgaggctgcatcgtgtggtgtgaatatgtgtgttt 16468
chimp54546-135457.revcompl.Pan.troglodytes.clone.rp43-41g5.GenBank:AC099533.36 GCTTGAGGATTGGTGGTGTCTTGTGTGAGGCTGCATCATGTGGTGTGAATGTGTGTGTTT 18002
human.FAM30B-LOC105372935-GGT2.NCBI. gcttgaggattggtggtgtcttgtgtgaggctgcatcgtgtggtgtgaatgtgtgtgttt 14435
human.FAM230E-LOC105377182-GGT3P.NCBI. gcttgaggattggtggtgtcttgtgtgaggctgcatcgtgtggtgtgaatgtgtgtgttt 13502
human.FAM230J-LOC105372942-GGTLC5P.NCBI.rev.compl gcttgaggattggtggtgtcttgtgtgaggctgcatcgtgtggtgtgaatgtgtgtgttt 13885
human.FAM230A.ncRNA.18487127.18500594-GGTLC3.18516335.18518165.rev.compl.REF gcttgaggattggtggtgtcttgtgtgaggctgcatcgtgtggtgtgaatgtgtgtgttt 2920
************************************* ************ *********

chimp.LOC112206744-LOC107973052-GGT2.revcompl ctgtacaggtgaggctgtgtgttttctcaggagagatttcccacttatacaacccaatca 16528
chimp54546-135457.revcompl.Pan.troglodytes.clone.rp43-41g5.GenBank:AC099533.36 CTGTACAGGTGAGGCTGTGTGTTTTCTCAGGAGAGATTTCCCACTTATACAACCCAATCA 18062
human.FAM30B-LOC105372935-GGT2.NCBI. ctgtacaggtgaggctgtgtgttttctcaggagagatttcccacttatacaacccaatca 14495
human.FAM230E-LOC105377182-GGT3P.NCBI. ctgtacaggtgaggctgtgtgttttctcaggagagatttcccacttatacaacccaatca 13562
human.FAM230J-LOC105372942-GGTLC5P.NCBI.rev.compl ctgtacaggtgaggctgtgtgttttctcaggagagatttcccatttatacaacccaatca 13945
human.FAM230A.ncRNA.18487127.18500594-GGTLC3.18516335.18518165.rev.compl.REF ctgtacaggtgaggctgtgtgttttctcaggagagatttcccacttatacaacccaatca 2980
******************************************* ****************

chimp.LOC112206744-LOC107973052-GGT2.revcompl ccagtgtccacttctaacaataaaatccacccccgctctactctctctgtacagtgactc 16588
chimp54546-135457.revcompl.Pan.troglodytes.clone.rp43-41g5.GenBank:AC099533.36 CCAGTGTCCACTTCTAACAATAAAATCCACCCCCGCTCTACTCTCTCTGTACAGTGACTC 18122
human.FAM30B-LOC105372935-GGT2.NCBI. ccagtgtccacttctaacaataaaatccacccccgctctactctctctgtacagtgactc 14555
human.FAM230E-LOC105377182-GGT3P.NCBI. ccagtgtccacttctaacaataaaatccacccccgctctactctctctgtacagtgactc 13622
human.FAM230J-LOC105372942-GGTLC5P.NCBI.rev.compl ccagtgtccacttctaacaataaaatccacccccgctctactctctctgtacagtgactc 14005
human.FAM230A.ncRNA.18487127.18500594-GGTLC3.18516335.18518165.rev.compl.REF ccagtgtccacttctaacaataaaatccacccccgctctactctctctgtacagtgactc 3040
************************************************************

chimp.LOC112206744-LOC107973052-GGT2.revcompl ctccaccctcactagagccatccccgggtctgccttattatccccactgctcaggtggag 16648
chimp54546-135457.revcompl.Pan.troglodytes.clone.rp43-41g5.GenBank:AC099533.36 CTCCACCCTCACCAGAGCCATCCCCAGATCTGCCTTATTATCCCCACTGCTCAGGTGGAG 18182
human.FAM30B-LOC105372935-GGT2.NCBI. ctccaccctcaccagagccatccccgggtctgccttattatccccactgctcaggtggag 14615
human.FAM230E-LOC105377182-GGT3P.NCBI. ctccaccctcaccagagccatccccgggtctgccttattatccccactgctcaggtggag 13682
human.FAM230J-LOC105372942-GGTLC5P.NCBI.rev.compl ctccaccctcaccagagccatccccgggtctgccttattatccccactgctcaggtggag 14065
human.FAM230A.ncRNA.18487127.18500594-GGTLC3.18516335.18518165.rev.compl.REF ctccaccctcaccagagccatccccgggtctgccttattatccccactgctcaggtggag 3100
************ ************ * ********************************

chimp.LOC112206744-LOC107973052-GGT2.revcompl aacctgaagggccaagggagtggccccagctcccgagttcctgaatgaaaaagtgaaaac 16708
chimp54546-135457.revcompl.Pan.troglodytes.clone.rp43-41g5.GenBank:AC099533.36 AACCTGAAGGGCCAAGGGAGTGGCCCCAGCTCCCGAGTTCCTGAATGAAAAAGTGAAAAC 18242
human.FAM30B-LOC105372935-GGT2.NCBI. aacctgaagggccaagggagtggccccagctcccgagttcctgaatgaaacagtgaaaac 14675
human.FAM230E-LOC105377182-GGT3P.NCBI. aacctgaagggccaagggagtggccccagctcccgagttcctgaatgaaacagtgaaaac 13742
human.FAM230J-LOC105372942-GGTLC5P.NCBI.rev.compl aacctgaagggccaagggagtggccccagctcccgagttcctgaatgaaaaagtgaaaac 14125
human.FAM230A.ncRNA.18487127.18500594-GGTLC3.18516335.18518165.rev.compl.REF aacctgaagggccaagggagtggccccagctcccgagttcctgaatgaaaaagtgaaaac 3160
************************************************** *********

chimp.LOC112206744-LOC107973052-GGT2.revcompl acgaacccagaagtgtgggccagtgctgatgctgacatgcacttagtcatggggtgttca 16768
chimp54546-135457.revcompl.Pan.troglodytes.clone.rp43-41g5.GenBank:AC099533.36 ACGAACCCAGGAGTGTGGGCCAGTGCTGACGCTGACATGCACTTAGTCATGGGGTGTTCA 18302
human.FAM30B-LOC105372935-GGT2.NCBI. acgaacccaggagtgtggaccagtgctgacgctgacatgcacttagtcatggggtgttca 14735
human.FAM230E-LOC105377182-GGT3P.NCBI. acgaacccaggagtgtgggccagtgctgacgctgacatgcacttagtcatggggtgttca 13802
human.FAM230J-LOC105372942-GGTLC5P.NCBI.rev.compl acgaacccaggagtgtgggccagtgctgacgctgacatgcacttagtcatggggtgttca 14185
human.FAM230A.ncRNA.18487127.18500594-GGTLC3.18516335.18518165.rev.compl.REF acgaacccaggagtgtgggccagtgctgacgctgacatgcacttagtcatggggtgttca 3220
********** ******* ********** ******************************

chimp.LOC112206744-LOC107973052-GGT2.revcompl ccaccacacagggagtccagcattcatgtataagccctaaagcaacgagcccaaaaggcc 16828
chimp54546-135457.revcompl.Pan.troglodytes.clone.rp43-41g5.GenBank:AC099533.36 CCACCACACAGGGAGTCCAGCATTCATGTATAAGCCCTAAAGCAACGAGCCCAAAAGGCC 18362
human.FAM30B-LOC105372935-GGT2.NCBI. ccaccacacacggagtccagcattcatgtataagccctaaagcaacgagcccaaaaggcc 14795
human.FAM230E-LOC105377182-GGT3P.NCBI. ccaccacacagggagtccagcattcatgtataaaccctaaggcaccgagcccaaaaggcc 13862
human.FAM230J-LOC105372942-GGTLC5P.NCBI.rev.compl ccaccacacagggagtccagcattcatgtataaaccctaaggcaccgagcccaaaaggcc 14245
human.FAM230A.ncRNA.18487127.18500594-GGTLC3.18516335.18518165.rev.compl.REF ccaccacacagggagtccagcattcatgtataaaccctaaggcaccgagcccaaaaggcc 3280
********** ********************** ****** *** ***************

chimp.LOC112206744-LOC107973052-GGT2.revcompl ccagacactgcccatcatcataaagtggcctccgtggtcacacaacccagggcagttata 16888
chimp54546-135457.revcompl.Pan.troglodytes.clone.rp43-41g5.GenBank:AC099533.36 CCAGACACTGCCCATCATCATAAAGTGGCCTCCGTGGTCACACAACCCAGGGCAGTTATA 18422
human.FAM30B-LOC105372935-GGT2.NCBI. ccagacactgcccatcatcataaagtggcctccgtggtcacacaacccagggcagttata 14855
human.FAM230E-LOC105377182-GGT3P.NCBI. ccaggcactgcccatcatcataaagtggcctccgtggtcacacaacccagggcagttata 13922
human.FAM230J-LOC105372942-GGTLC5P.NCBI.rev.compl ccagacactgcccatcatcataaagtggcctccgtggtcacacaacccagggcagttatg 14305
human.FAM230A.ncRNA.18487127.18500594-GGTLC3.18516335.18518165.rev.compl.REF ccaggcactgcccatcatcataaagtggtctccgtggtcacacaacccagggcagttata 3340
**** *********************** ******************************

chimp.LOC112206744-LOC107973052-GGT2.revcompl ggctcatctccccacggacaggcatagtcatcagtgtgtcaaaagcacaaagatccccag 16948
chimp54546-135457.revcompl.Pan.troglodytes.clone.rp43-41g5.GenBank:AC099533.36 GGCTCATCTCCCCACGGACCGGCATAGTCATCAGTGTGTCAAAAGCACAAAGATCCCCAG 18482
human.FAM30B-LOC105372935-GGT2.NCBI. gcctcatctccccacggaccgacatagtcatcagtgtgtcaaaagcacaaagatccccag 14915
human.FAM230E-LOC105377182-GGT3P.NCBI. ggttcatctccccacggacaggcatagtcatcagtgtgtcaaaagcacaaagatccccag 13982
human.FAM230J-LOC105372942-GGTLC5P.NCBI.rev.compl ggttcatctccccacggacaggcatagtcatcagtgtgtcaaaagcacaaagatccccag 14365
human.FAM230A.ncRNA.18487127.18500594-GGTLC3.18516335.18518165.rev.compl.REF ggttcatctccccacggacaggcatagtcatcagtgtgtcaaaagcacaaagatccccag 3400
*  **************** * **************************************

chimp.LOC112206744-LOC107973052-GGT2.revcompl gtgtttggctcagctcacagatcccttttttttt----tttaacttttaagttcaggggt 17004
chimp54546-135457.revcompl.Pan.troglodytes.clone.rp43-41g5.GenBank:AC099533.36 GTGTTTGGCTCAGCTCACAGATCCCCCTTTTTTT----TTTAACTTTTAAGTTCAGGGGT 18538
human.FAM30B-LOC105372935-GGT2.NCBI. gtgtttggctcagctcacagatccctttttttt-----atttaacttttcagttcaggggt 14970
human.FAM230E-LOC105377182-GGT3P.NCBI. gtgtttggctcagctcaccgatccttttttttttt---tttaacttttcagttcaggggt 14039
human.FAM230J-LOC105372942-GGTLC5P.NCBI.rev.compl gtgtttggctcagctcaccgatccttttttttttttatttaacttttcagttcaggggt 14425
human.FAM230A.ncRNA.18487127.18500594-GGTLC3.18516335.18518165.rev.compl.REF gtgtttggctcagctcaccgatccttttttttttttttttaacttttaagttcaggggt 3460
****************** *****   *****      ********** ***********

chimp.LOC112206744-LOC107973052-GGT2.revcompl acatgtgcaggatgtgcaggtttgctacatatataaatgtgtgtcatgagagtttgttgt 17064
chimp54546-135457.revcompl.Pan.troglodytes.clone.rp43-41g5.GenBank:AC099533.36 ACATGTGCAGGATGTGCAGGTTTGCTACATATATGAATGTGTGTTATGAGAGTTTGTTGT 18598
human.FAM30B-LOC105372935-GGT2.NCBI. acatgtgcaggatgtgcaggtttgctacatatataaatgtgtgttatgagagttttttgt 15030
human.FAM230E-LOC105377182-GGT3P.NCBI. acatgtgcaggatgtgcaggtttgctacatatataaatgtgtgtcatgagagtttgttgt 14099
human.FAM230J-LOC105372942-GGTLC5P.NCBI.rev.compl acatgtgcaggatgtgcaggtttgctacatatataaatgtgtgtcatgagagtttgttgt 14485
human.FAM230A.ncRNA.18487127.18500594-GGTLC3.18516335.18518165.rev.compl.REF acatgtgcaggatgtgcaggtttgctacatatataaatgtgtgtcatgagagtttgttgt 3520
********************************** ********* ********** ****

chimp.LOC112206744-LOC107973052-GGT2.revcompl acagattattgcatcacccatatattgagcctaatatcagttatttttcctgatcctccc 17124
chimp54546-135457.revcompl.Pan.troglodytes.clone.rp43-41g5.GenBank:AC099533.36 ACAGATTATTGCATCACCCATATATTGAGGCTAATATCAGTTATTTTTCCTGATCCACCC 18658
human.FAM30B-LOC105372935-GGT2.NCBI. acagattattgcatcacccatatattgagcctaatatcagttatttttcctgatccaccc 15090
human.FAM230E-LOC105377182-GGT3P.NCBI. acagattattgcatcacccatatattgagcctaatatcagttatttttcctgatccaccc 14159
human.FAM230J-LOC105372942-GGTLC5P.NCBI.rev.compl acagattattgcatcacccatatattgagcctaatatcagttatttttcctgatccaccc 14545
human.FAM230A.ncRNA.18487127.18500594-GGTLC3.18516335.18518165.rev.compl.REF acagattattgcatcacccatatattgagcctaatatcagttatttttcctgatccaccc 3580
***************************** ************************** ***

chimp.LOC112206744-LOC107973052-GGT2.revcompl cctcctcccacctcccac---cctccagtaggccccacgctcacaaattctaagaggagt 17181
chimp54546-135457.revcompl.Pan.troglodytes.clone.rp43-41g5.GenBank:AC099533.36 CCTCCTCCCACCTCCCACCCTCCTCCAGTAGGCCCCACGCTCACAAATTCTAAGAGGAGT 18718
human.FAM30B-LOC105372935-GGT2.NCBI. cctcctcccacctcccac---cctccagtaggccccacgctcacaaattctaagaggagt 15147
human.FAM230E-LOC105377182-GGT3P.NCBI. tctcctcccacctcccac---cctccagtaggccccacgctcacaaattctaagaggagt 14216
human.FAM230J-LOC105372942-GGTLC5P.NCBI.rev.compl cctcctcccacctcccac---cctccagtaggccccacgctcacaaattctaagaggagt 14602
human.FAM230A.ncRNA.18487127.18500594-GGTLC3.18516335.18518165.rev.compl.REF cctcctcccacctcccac---cctccagtaggccccacgctcacaaattctaagaggagt 3637
 *****************   ***************************************

chimp.LOC112206744-LOC107973052-GGT2.revcompl gggggaccacaaaggccagtgtggcccacttcagttgtgaagttaatttgctcagcaact 17241
chimp54546-135457.revcompl.Pan.troglodytes.clone.rp43-41g5.GenBank:AC099533.36 GGGGGACCACAAAGGCCAGTGTGGCCCACTTCAGTTGTGAAGTTAATCTGCTCAGCAACT 18778
human.FAM30B-LOC105372935-GGT2.NCBI. gggggaccacaaaggccagtgtggcccacttcagttgtgaagttaatttgctcagcaact 15207
human.FAM230E-LOC105377182-GGT3P.NCBI. gggggaccacaaaggccagtgtggcccacttcagttgtgaagttaatttgctcagcaact 14276
human.FAM230J-LOC105372942-GGTLC5P.NCBI.rev.compl gggggaccacaaaggccagtgtggcccacttcagttgtgaagttaatttgctcagcaact 14662
human.FAM230A.ncRNA.18487127.18500594-GGTLC3.18516335.18518165.rev.compl.REF gggggaccacaaaggccagtgtggcccacttcagttgtgaagttaatttgctcagcaact 3697
*********************************************** ************

chimp.LOC112206744-LOC107973052-GGT2.revcompl ggccaaagtctataaggatgggtgatgtattttagtagatttagtaatactatcttccca 17301
chimp54546-135457.revcompl.Pan.troglodytes.clone.rp43-41g5.GenBank:AC099533.36 GGCCAAAGTCTATAAGGATGGGTGATGTATTTTAGTAGATTTAGTAATACTATCTTCCCA 18838
human.FAM30B-LOC105372935-GGT2.NCBI. ggccaaagtctataaggatgggtgatgtattttagtagatttagtaatactatcttccca 15267
human.FAM230E-LOC105377182-GGT3P.NCBI. ggccaaagtctataaggatgggtgatgtattttagtagatttagtaatactatcttccca 14336
human.FAM230J-LOC105372942-GGTLC5P.NCBI.rev.compl ggccaaagtctataaggatgggtgatgtattttagtagatttagtaatactatcttccca 14722
human.FAM230A.ncRNA.18487127.18500594-GGTLC3.18516335.18518165.rev.compl.REF ggccaaagtctataaggatgggtgatgtattttagtagatttagtaatactatcttccca 3757
************************************************************

chimp.LOC112206744-LOC107973052-GGT2.revcompl agccctaaaatgctcaaatcctgccagccaaaaatggtgaggagggacagataggaactc 17361
chimp54546-135457.revcompl.Pan.troglodytes.clone.rp43-41g5.GenBank:AC099533.36 AGCCCTAAAATGCTCAAATCCTGCCAGCCAAAAATGGTGAGGAGGGACAGATAGGAACTC 18898
human.FAM30B-LOC105372935-GGT2.NCBI. agccctaaaatgctcaaatcctgccagccaaaaatggtgaggagggacagataggaactc 15327
human.FAM230E-LOC105377182-GGT3P.NCBI. agccctaaaatgctcaaatcctgccagccaaaaatggtgaggagggacagataggaactc 14396
human.FAM230J-LOC105372942-GGTLC5P.NCBI.rev.compl agccctaaaatgctcaaatcctgccagccaaaaatggtgaggagggacagataggaactc 14782
human.FAM230A.ncRNA.18487127.18500594-GGTLC3.18516335.18518165.rev.compl.REF agccctaaaatgctcaaatcctgccagccaaaaatggtgaggagggacagataggaactc 3817
************************************************************

chimp.LOC112206744-LOC107973052-GGT2.revcompl tgtgtggcacttggttattagcctggcttccatcccttagtggcaagtctcttgtatatg 17421
chimp54546-135457.revcompl.Pan.troglodytes.clone.rp43-41g5.GenBank:AC099533.36 TGTGTGGCACTTGGTTATTAGCCTGGCTTCCATCCCTTAGTGGCAACTCTCTTGTATATG 18958
human.FAM30B-LOC105372935-GGT2.NCBI. tgtgtggcacttggttattagcctggcttccatcccttagtggcaactctcttgtatatg 15387
human.FAM230E-LOC105377182-GGT3P.NCBI. tgtgtggcacttggttattagcctggcttccatcccttagtggcaactctcttgtatatg 14456
human.FAM230J-LOC105372942-GGTLC5P.NCBI.rev.compl tgtgtggcacttggttattagcctggcttccatcccttagtggcaactctcttgtatatg 14842
human.FAM230A.ncRNA.18487127.18500594-GGTLC3.18516335.18518165.rev.compl.REF tgtgtggcacttggttattagcctggcttccatcccttagtggcaactctcttgtatatg 3877
********************************************** *************

chimp.LOC112206744-LOC107973052-GGT2.revcompl tgggttaaagaccctcagcctcaagccaagcctcctccatgaggagccatctcactattg 17481
```

```
chimp54546-135457.revcompl.Pan.troglodytes.clone.rp43-41g5.GenBank:AC099533.36     TGGGTTAAAGACCCTCAGCCTCAAGCCAAGCCTCCTCCATGAGGAGCCATCTCACTATTG 19018
human.FAM30B-LOC105372935-GGT2.NCBI.                                               tgggttaaagaccctcagcctcaagccaagcctcctccatgaggagccatctcactattg 15447
human.FAM230E-LOC105377182-GGT3P.NCBI.                                             tgggttaaagaccctcagcctcaagccaagcctcctccatgaggagccatctcactattg 14516
human.FAM230J-LOC105372942-GGTLC5P.NCBI.rev.compl                                  tgggttaaagaccctcagcctcaagccaagcctcctccatgaggagccatctcactattg 14902
human.FAM230A.ncRNA.18487127.18500594-GGTLC3.18516335.18518165.rev.compl.REF       tgggttaaagaccctcagcctcaagccaagcctcctccatgaggagccatctcactattg 3937
                                                                                   ************************************************************

chimp.LOC112206744-LOC107973052-GGT2.revcompl                                      actggctagtgccgggtatggccaccagcccaactgaaacaaaatgttgctttaaaacaa 17541
chimp54546-135457.revcompl.Pan.troglodytes.clone.rp43-41g5.GenBank:AC099533.36     ACTGGCTAGTGCCGGGTATGGCCACCAGCCCAACTGAAACAAAATGTTGCTTTAAAACAA 19078
human.FAM30B-LOC105372935-GGT2.NCBI.                                               accggctagtgccgggtatggccaccagcccaactgaaacaaaatgttgctttaaaacaa 15507
human.FAM230E-LOC105377182-GGT3P.NCBI.                                             accggctagtgccgggtatggccaccagcccaactgaaacaaaatgttgctttaaaacaa 14576
human.FAM230J-LOC105372942-GGTLC5P.NCBI.rev.compl                                  actggctagtgccgggtatggccaccagcccaactgaaacaaaatgttgctttaaaacaa 14962
human.FAM230A.ncRNA.18487127.18500594-GGTLC3.18516335.18518165.rev.compl.REF       actggctagtgccgggtatggccaccagcccaactgaaacaaaatgttgctttaaaacaa 3997
                                                                                   ** *********************************************************

chimp.LOC112206744-LOC107973052-GGT2.revcompl                                      gtgtaaatctcatacatacaacaggcaaatgcagaagcagtgtggtctcgcaagttgtaa 17601
chimp54546-135457.revcompl.Pan.troglodytes.clone.rp43-41g5.GenBank:AC099533.36     GTGTAAATCTCATACATACAACAGGCAAATGCAGAAGCAGTGTGGTCTCGCAAGTTGTAA 19138
human.FAM30B-LOC105372935-GGT2.NCBI.                                               gtgtaaatctcatacatacaacaggcaaatgcagaagcagtgtggtctcgcaagttgtaa 15567
human.FAM230E-LOC105377182-GGT3P.NCBI.                                             gtgtaaatctcgtacatacaacaggcaaatgcagaagcagtgtggtctcgcaagttgtaa 14636
human.FAM230J-LOC105372942-GGTLC5P.NCBI.rev.compl                                  gtgtaaatctcatacatacaacaggcaaatgcagaagcagtgtggtctcgcaagttgtaa 15022
human.FAM230A.ncRNA.18487127.18500594-GGTLC3.18516335.18518165.rev.compl.REF       gtgtaaatctcatacatacaacaggcaaatgcagaagcagtgtggtctcgcaagttgtaa 4057
                                                                                   *********** ************************************************

chimp.LOC112206744-LOC107973052-GGT2.revcompl                                      agaggacagtcgcaattttgctggacttcaacctgggtagaagacatgagggaactctgt 17661
chimp54546-135457.revcompl.Pan.troglodytes.clone.rp43-41g5.GenBank:AC099533.36     AGAGGACAGTCGCAATTTTGCTGGACTTCAACCTGGGTAGAAGACATGAGGGAACTCTGT 19198
human.FAM30B-LOC105372935-GGT2.NCBI.                                               agaggacagtcgcaattttgctggacttcaacctgggtagaagacatgagggaac-ctgt 15626
human.FAM230E-LOC105377182-GGT3P.NCBI.                                             agaggacagtcgcaattttgctggacttcaacctgggtagaagacacgagggaactctgt 14696
human.FAM230J-LOC105372942-GGTLC5P.NCBI.rev.compl                                  agaggacagtcgcaattttgctggacttcaacctgggtagaagacacgagggaactctgt 15082
human.FAM230A.ncRNA.18487127.18500594-GGTLC3.18516335.18518165.rev.compl.REF       agaggacagtcgcaattttgctggacttcaacctgggtagaagacacgagggaactctgt 4117
                                                                                   ********************************************* ********* ****

chimp.LOC112206744-LOC107973052-GGT2.revcompl                                      cactaaatcacggcagagttcaaggccacttgcagactatttcatgttacagaaggtggc 17721
chimp54546-135457.revcompl.Pan.troglodytes.clone.rp43-41g5.GenBank:AC099533.36     CACTGAATCACGGCAGAGTTCAAGGCCACTTGCAGACTATTTCATGTTATAGAAGGTGGC 19258
human.FAM30B-LOC105372935-GGT2.NCBI.                                               cactaaatcacggcagagttcaaggccacttgtagactatttcatgttacagaaggtggc 15686
human.FAM230E-LOC105377182-GGT3P.NCBI.                                             cactaaatcacggcagagttcaaggccgcttgtagactatttcgtgttatagaaggtggc 14756
human.FAM230J-LOC105372942-GGTLC5P.NCBI.rev.compl                                  cactaaatcacggcagagttcaagg-----------ctatttcgtgttatagaaggtggc 15131
human.FAM230A.ncRNA.18487127.18500594-GGTLC3.18516335.18518165.rev.compl.REF       cactaaatcacggcagagttcaaggccgcttgtagactatttcgtgttatagaaggtggc 4177
                                                                                   **** ********************           ******* ***** **********

chimp.LOC112206744-LOC107973052-GGT2.revcompl                                      ctttagctactaagcaaaggcctctgtttctcatttctttcctgttcatcttcttggtca 17781
chimp54546-135457.revcompl.Pan.troglodytes.clone.rp43-41g5.GenBank:AC099533.36     CTTTAGCTACTAAGCAAAGGCCTCTGTTTCTCATTTCTTTCCTGTTCATCTTCTTGGTCA 19318
human.FAM30B-LOC105372935-GGT2.NCBI.                                               ctttagctactaagcaaaggcctctgtttctcatttctttcctgttcatcttcttggtca 15746
human.FAM230E-LOC105377182-GGT3P.NCBI.                                             ctttagctactaggcaaaggcctctgtttctcatttctttcctgttcatctccttggtca 14816
human.FAM230J-LOC105372942-GGTLC5P.NCBI.rev.compl                                  ctttagctactaggcaaaggcctctgtttctcatttctttcctgttcatctccttggtca 15191
human.FAM230A.ncRNA.18487127.18500594-GGTLC3.18516335.18518165.rev.compl.REF       ctttagctactaagcaaaggcctctgtttctcatttctttcctgttcatctccttggtca 4237
                                                                                   ************ ************************************** ********

chimp.LOC112206744-LOC107973052-GGT2.revcompl                                      tccttcttccgcaagggaaacgagcccaagcaaaaggcagtttcaatattaatttgaccg 17841
chimp54546-135457.revcompl.Pan.troglodytes.clone.rp43-41g5.GenBank:AC099533.36     TCCTTCTTCCGCAAGGGAAATGAGCCCAAGCAAAAGGCAGTTTCAATATTAATTTGACCG 19378
human.FAM30B-LOC105372935-GGT2.NCBI.                                               tccttcttccgcaagggaaacgagcccaagcaaaaggcagtttcaatattaatttgaccg 15806
human.FAM230E-LOC105377182-GGT3P.NCBI.                                             tccttcttccgcaagggaaacgagcccaagcaaaaggcagtttcaatattaatttgaccg 14876
human.FAM230J-LOC105372942-GGTLC5P.NCBI.rev.compl                                  tccttcttccgcaagggaaacgagcccaagcaaaaggcagtttcaatattaatttgaccg 15251
human.FAM230A.ncRNA.18487127.18500594-GGTLC3.18516335.18518165.rev.compl.REF       tccttcttccacaagggaaacgagcccaagcaaaaggcagtttcaatattaatttgaccg 4297
                                                                                   ********** ********* ***************************************

chimp.LOC112206744-LOC107973052-GGT2.revcompl                                      aggttttgtgcagtttattatcatccaggtaatcaagtgcaacccagtctgcctagcagc 17901
chimp54546-135457.revcompl.Pan.troglodytes.clone.rp43-41g5.GenBank:AC099533.36     AGGTTTTGTGCAGTTTATTATCATCCAGGTAATCAAGTGCAACCCAGTCTGCCTAGCAGC 19438
human.FAM30B-LOC105372935-GGT2.NCBI.                                               aggttttgtgcagtttattatcatccaggtaatcaggtgcaacccagtctgcctagccgc 15866
human.FAM230E-LOC105377182-GGT3P.NCBI.                                             aggttttgtgcagttaattatcatccaggtaatcaggtgcaacccagtctgcctagcagc 14936
human.FAM230J-LOC105372942-GGTLC5P.NCBI.rev.compl                                  aggttttgtgcagttaattatcatccaggtaatcaggtgcaacccagtctgcctagcagc 15311
human.FAM230A.ncRNA.18487127.18500594-GGTLC3.18516335.18518165.rev.compl.REF       aggttttgtgcagttaattatcatccaggtaatcaggtgcaacccagtctgcctagcagc 4357
                                                                                   *************** ****************** ********************** **

chimp.LOC112206744-LOC107973052-GGT2.revcompl                                      cccccctatctctgctctgtgttttcatttaataaacattttggtctacttactatgtgct 17961
chimp54546-135457.revcompl.Pan.troglodytes.clone.rp43-41g5.GenBank:AC099533.36     CCCCCTATCTCTGCTCTGTGTTTTCATTTAATAAACATTTTGGTCTACTTACTATGTGCT 19498
human.FAM30B-LOC105372935-GGT2.NCBI.                                               ccccctatctctgctctgtgttttcatttaataaacattttggtctacttactatgtgct 15926
human.FAM230E-LOC105377182-GGT3P.NCBI.                                             ccccctatctctgctctgtgttttcatttaataaacattttggtctacttactatgtgct 14996
human.FAM230J-LOC105372942-GGTLC5P.NCBI.rev.compl                                  ccccctatctctgctctgtgttttcatttaatgaacattttggtctacttactatgtgct 15371
human.FAM230A.ncRNA.18487127.18500594-GGTLC3.18516335.18518165.rev.compl.REF       ccccctatctctgctctgtgttttcatttaataaacattttggtctacttactatgtgct 4417
                                                                                   ******************************** ***************************

chimp.LOC112206744-LOC107973052-GGT2.revcompl                                      agattttctcgagaccaagtaaatgagatagaatctttatgttggcagctaagttagatt 18021
chimp54546-135457.revcompl.Pan.troglodytes.clone.rp43-41g5.GenBank:AC099533.36     AGATTTTCTCGAGACCAAGTAAATGAGATAGAATCTTTATGTTGGCAGCTAAGTTAGATT 19558
human.FAM30B-LOC105372935-GGT2.NCBI.                                               agattttctcgagaccaagtaaatgagatagaatctttatgctggcagctaagttagatt 15986
human.FAM230E-LOC105377182-GGT3P.NCBI.                                             agattttctcgagaccaagtaaatgagatagaatctttatgttggcagctaagttagatt 15056
human.FAM230J-LOC105372942-GGTLC5P.NCBI.rev.compl                                  agattttctcgagaccaagtaaatgagatagaatctttatgttggcagctaagttagatt 15431
human.FAM230A.ncRNA.18487127.18500594-GGTLC3.18516335.18518165.rev.compl.REF       agattttctcgagaccaagtaaatgagatagaatcctttatgttggcagctaagttagatt 4477
                                                                                   *********************************** ***** ******************

chimp.LOC112206744-LOC107973052-GGT2.revcompl                                      taacataactgacaaaaattaaaatttctgatttcttgtaaaaatattttgtatgtgtga 18081
chimp54546-135457.revcompl.Pan.troglodytes.clone.rp43-41g5.GenBank:AC099533.36     TAACATAACTGACAAAAATTAAAATTTCTGATTTCTTGTAAAAATATTTTGTATGTGTGA 19618
human.FAM30B-LOC105372935-GGT2.NCBI.                                               tcacataactgacaaaaattaaaatttctgatttcttgtaaaaatattttgtatgtgtga 16046
human.FAM230E-LOC105377182-GGT3P.NCBI.                                             tcacataactgacaaaaattaaaatttctgatttcttgtaaaaatattttgtatgtgtga 15116
human.FAM230J-LOC105372942-GGTLC5P.NCBI.rev.compl                                  tcgcataactgacaaaaattaaaatttctgatttcttgtaaaaatattttgtatgtgtga 15491
human.FAM230A.ncRNA.18487127.18500594-GGTLC3.18516335.18518165.rev.compl.REF       tcacataactgacaaaaattaaaatttctgatttcttgtaaaaatattttgtatgtgtga 4537
                                                                                   *  *********************************************************

chimp.LOC112206744-LOC107973052-GGT2.revcompl                                      atgcatactgaatgtaaagtggataaaaaactcacacttgcactcatggaaggcttttca 18141
chimp54546-135457.revcompl.Pan.troglodytes.clone.rp43-41g5.GenBank:AC099533.36     ATGCATACTGAATGTAAAGTGGATAAAAAACTCACACTTGCACTCATGGAAGGCTTTTCA 19678
human.FAM30B-LOC105372935-GGT2.NCBI.                                               atgcatactgaatgtaaagtggataaaaaactcacacttgcactcatggaaggcttttca 16106
human.FAM230E-LOC105377182-GGT3P.NCBI.                                             atgcatactgaatgtaaagtggataaaaaactcacacttgcactcatggaaggcttttca 15176
human.FAM230J-LOC105372942-GGTLC5P.NCBI.rev.compl                                  atgcatactgaatgtaaagtggataaaaaactcacacttgcactcatggaaggcttttca 15551
human.FAM230A.ncRNA.18487127.18500594-GGTLC3.18516335.18518165.rev.compl.REF       atgcatactgaatgtaaagtggataaaaaactcacacttgcactcatggaaggcttttca 4597
                                                                                   ************************************************************

chimp.LOC112206744-LOC107973052-GGT2.revcompl                                      tgaatttgtcaatttttattttttatatttccccacttcaccggataatgcatacctgaa 18201
chimp54546-135457.revcompl.Pan.troglodytes.clone.rp43-41g5.GenBank:AC099533.36     TGAATTTGTCAATTTTTATTTTTTATATTTCCCCACTTCACCGGATAATGCATACCTGAA 19738
human.FAM30B-LOC105372935-GGT2.NCBI.                                               tgaatttgtcaatttttattttttatatttccccacttcaccagataatgcatacctgaa 16166
human.FAM230E-LOC105377182-GGT3P.NCBI.                                             tgaatttgtcaatttttattttttatatttccccacttcactggataatgcatacctgaa 15236
human.FAM230J-LOC105372942-GGTLC5P.NCBI.rev.compl                                  tgaatttgtcaatttttattttttatatttccccacttcactggataatgcatacctgaa 15611
human.FAM230A.ncRNA.18487127.18500594-GGTLC3.18516335.18518165.rev.compl.REF       tgaatttgtcaatttttattttttatatttccccacttcactggataatgcatacctgaa 4657
                                                                                   *****************************************  *****************

chimp.LOC112206744-LOC107973052-GGT2.revcompl                                      cctggaaattgattcccactgcagaaagtgttctgagccacatcccttagcttcactagt 18261
chimp54546-135457.revcompl.Pan.troglodytes.clone.rp43-41g5.GenBank:AC099533.36     CCTGGAAACTGATTCCCACTGCAGAAAGTGTTCTGAGCCACATCCCTTAGCTTCACTAGT 19798
human.FAM30B-LOC105372935-GGT2.NCBI.                                               cctggaaactgattcccactgcagaaagtgttctgagccacatcccttagcttcactagt 16226
human.FAM230E-LOC105377182-GGT3P.NCBI.                                             cctggaaactgatgcccactgcagaaagtgttctgagccacatcccttagcttcactagt 15296
human.FAM230J-LOC105372942-GGTLC5P.NCBI.rev.compl                                  cctggaaactgatgcccactgcagaaagtgttctgagccacatcccttagcttcactagt 15671
human.FAM230A.ncRNA.18487127.18500594-GGTLC3.18516335.18518165.rev.compl.REF       cctggaaactgatgcccactgcagaaagtgttctgagccacatcccttagcttcactagt 4717
                                                                                   ******** **** **********************************************

chimp.LOC112206744-LOC107973052-GGT2.revcompl                                      gcaggtccacctgggaggatgtcccagcatcagcttggcccatgctgtgatcagccacct 18321
chimp54546-135457.revcompl.Pan.troglodytes.clone.rp43-41g5.GenBank:AC099533.36     GCAGGTCCACCTGGGAGGATGTCCCAGCATCAGCTTGGCCCATGCTGTGATCAGCCACCT 19858
human.FAM30B-LOC105372935-GGT2.NCBI.                                               gcaggtccacctgggaggatgtcccagcatcagcttgacccatgctgtgatcagccacct 16286
human.FAM230E-LOC105377182-GGT3P.NCBI.                                             gcaggtccacctgggaggatgtcccagcatcagcttggcccatgctgtgatcagccacct 15356
human.FAM230J-LOC105372942-GGTLC5P.NCBI.rev.compl                                  gcaggtccacctgggaggatgtcccagcatcagcttggcccatgctgtgatcagccacct 15731
human.FAM230A.ncRNA.18487127.18500594-GGTLC3.18516335.18518165.rev.compl.REF       gcaggtccacctgggaggatgtcccagcatcagcttggcccatgctgtgatcagccacct 4777
                                                                                   ************************************* **********************

chimp.LOC112206744-LOC107973052-GGT2.revcompl                                      ccatgcaccacaccaagcaagcccctgggtgattcacagtctccaccaccagggcactga 18381
chimp54546-135457.revcompl.Pan.troglodytes.clone.rp43-41g5.GenBank:AC099533.36     CCATGCACCACACCAAGCAAGCCCCTGGGTGATTCACAGTCTCCACCACCAGGGCACTGA 19918
human.FAM30B-LOC105372935-GGT2.NCBI.                                               ccatgcatcacaccaagcaagcccctgggtgattcacagtctccaccaccagggcactga 16346
human.FAM230E-LOC105377182-GGT3P.NCBI.                                             ccatgcaccacaccaagcaagcccctgggtgattcacagtctccactaccagggcactga 15416
human.FAM230J-LOC105372942-GGTLC5P.NCBI.rev.compl                                  ccatgcaccacaccaagcaagcccctgggtgattcacagtctccactaccagggcactga 15791
human.FAM230A.ncRNA.18487127.18500594-GGTLC3.18516335.18518165.rev.compl.REF       ccatgcaccacaccaagcaagcccctgggtgattcacagtctccactaccagggcactga 4837
                                                                                   ******* ************************************** *************

chimp.LOC112206744-LOC107973052-GGT2.revcompl                                      ccttaactgtgtgttcttctagctccccatgaagacaccgtacacgacatcactaacgag 18441
chimp54546-135457.revcompl.Pan.troglodytes.clone.rp43-41g5.GenBank:AC099533.36     CCTTAACTCTGTGTTCTTCTAGCTCCCCATGAGGACACCGTACACGACATCACTAACGAG 19978
human.FAM30B-LOC105372935-GGT2.NCBI.                                               ccttaactctgtgtttttctagctccccatgaggacaccgtacacgacatcactaacgag 16406
human.FAM230E-LOC105377182-GGT3P.NCBI.                                             ccttaactctgtgttcttctagctccccatgaggacaccgtacacaacatcactaacgag 15476
human.FAM230J-LOC105372942-GGTLC5P.NCBI.rev.compl                                  ccttaactctgtgttcttctagctccccatgaggacaccgtacacaacatcactaacgag 15851
human.FAM230A.ncRNA.18487127.18500594-GGTLC3.18516335.18518165.rev.compl.REF       ccttaactctgtgttcttctagctccccatgaggacaccgtacacaacatcactaacgag 4897
                                                                                   ******** ****** **************** ************ **************

chimp.LOC112206744-LOC107973052-GGT2.revcompl                                      gacgccacacacgacatcgccaacgaggacactgtacatgacaccgctaacgaggccgcc 18501
chimp54546-135457.revcompl.Pan.troglodytes.clone.rp43-41g5.GenBank:AC099533.36     GACGCCACACAGGACATCGCTAACAAGGACACTGTATACGACATCGCTAACGAGGCCGCC 20038
human.FAM30B-LOC105372935-GGT2.NCBI.                                               gacgccacacacgacatcgctaacgaggatactgtacacgacatcgctaacgaggccgcc 16466
human.FAM230E-LOC105377182-GGT3P.NCBI.                                             gatgcctcacacgatatcactaacgaggacgctgtccacggcatcgctaacgaggccgcc 15536
human.FAM230J-LOC105372942-GGTLC5P.NCBI.rev.compl                                  gatgcctcacacgatatcactaacgaggacgctgtccacggcatcgctaacgaggccgcc 15911
human.FAM230A.ncRNA.18487127.18500594-GGTLC3.18516335.18518165.rev.compl.REF       gatgcctcacacgatatcactaacgaggacgctgtccacggcatcgctaacgaggccgcc 4957
                                                                                   ** *** **** ** *** * *** ****  ****  * * ** ****************

chimp.LOC112206744-LOC107973052-GGT2.revcompl                                      gacaagggcatcgccagcgaggacgccgcccagggcatcgccaacgaggccgccgcccag 18561
chimp54546-135457.revcompl.Pan.troglodytes.clone.rp43-41g5.GenBank:AC099533.36     GACAAGGGCATCGCCAACGAGGACGCCGCCCAGGGCATCGCCAACGAGGACGCCTCCCAC 20098
```

```
human.FAM30B-LOC105372935-GGT2.NCBI.                                                 gacaagggcatcgccaacgaggatgccgcccacggcatcgccaacgagtacgccgcccac 16526
human.FAM230E-LOC105377182-GGT3P.NCBI.                                               gcccagggcatcgccaacgaggacgccgcccagggcatcgccaacgaggacgccgcccac 15596
human.FAM230J-LOC105372942-GGTLC5P.NCBI.rev.compl                                    gacaagggcatcgccaacgaggacgccgcccagggcatcgccaacgaggacgccgcccac 15971
human.FAM230A.ncRNA.18487127.18500594-GGTLC3.18516335.18518165.rev.compl.REF         gacaagggcaacgccaacgaggacgccgcccagggcatcgccaacgaggacgccgcccac 5017
                                                                                     * * ****** ***** ****** ******** **************  **** ****

chimp.LOC112206744-LOC107973052-GGT2.revcompl                                        ggcatcgtcagcgaggacgccgcccacggaatcgccaacgaggacgccgtccagggcatc 18621
chimp54546-135457.revcompl.Pan.troglodytes.clone.rp43-41g5.GenBank:AC099533.36       GGCATCGCCAACGAGGACACCGCCCACGGCATCGCCAGCGAGGACGCCGCCCACGGAATC 20158
human.FAM30B-LOC105372935-GGT2.NCBI.                                                 ggcatcgccagcgaggacgccgcccacggcatcgccagcgaggacgccgcccagggcatc 16586
human.FAM230E-LOC105377182-GGT3P.NCBI.                                               ggaatcgccagcgaggacgccgcccagggcatcgccaacgaggtcgccgcccagggcatc 15656
human.FAM230J-LOC105372942-GGTLC5P.NCBI.rev.compl                                    ggaatcgccagcgaggacgccgcccagggcatcgccaacgaggtcgccgcccagggcatc 16031
human.FAM230A.ncRNA.18487127.18500594-GGTLC3.18516335.18518165.rev.compl.REF         ggaatcgccagcgaggacgccgcccagggcatcgccaacgaggtcgccgcccagggcatc 5077
                                                                                     ** **** ** ******* ******* ** ******* ***** ***** *** ** ***

chimp.LOC112206744-LOC107973052-GGT2.revcompl                                        gccaacgtggtcgccgcccacggaatcgccaacgaggacgccgcccagggcatcgc---- 18677
chimp54546-135457.revcompl.Pan.troglodytes.clone.rp43-41g5.GenBank:AC099533.36       GCCAACGAGGATGCCGCCCAGGGCATCGCCAA---------------------------- 20190
human.FAM30B-LOC105372935-GGT2.NCBI.                                                 gccagcgaggacgccgcccagggcatcgccaacgaggacaccatccagggcatcgccaag 16646
human.FAM230E-LOC105377182-GGT3P.NCBI.                                               gccaacgaggacgccgcccagggcatcgccaacgaggacgccgtccatggcatcgccaac 15716
human.FAM230J-LOC105372942-GGTLC5P.NCBI.rev.compl                                    gccaacgaggacgccgcccagggcatcgccaaggaggacgccgcccacggcatcgccaac 16091
human.FAM230A.ncRNA.18487127.18500594-GGTLC3.18516335.18518165.rev.compl.REF         gccaacgaggacgccgcccagggcatcgccaacgaggacgccgtccagggcatcgccaac 5137
                                                                                     **** ** **  ******** ** ********

chimp.LOC112206744-LOC107973052-GGT2.revcompl                                        ------------------------------------------------------------ 18677
chimp54546-135457.revcompl.Pan.troglodytes.clone.rp43-41g5.GenBank:AC099533.36       ------------------------------------------------------------ 20190
human.FAM30B-LOC105372935-GGT2.NCBI.                                                 gagtacgccgtccacggcatcgccaacgaggacgccgcccagggcatcgccaacgaggac 16706
human.FAM230E-LOC105377182-GGT3P.NCBI.                                               gaggacgctgcccacggcattgctaacgaggatgccgtgcacggcatcgctaatgaggac 15776
human.FAM230J-LOC105372942-GGTLC5P.NCBI.rev.compl                                    gaggatgctgcccacggcattgctaacgaggatgccgtgcacggcatcgctaatgaggac 16151
human.FAM230A.ncRNA.18487127.18500594-GGTLC3.18516335.18518165.rev.compl.REF         gaggacgccgcccacggcatcgccaacgaggacgctgcccacggcattgctaacgaggat 5197

chimp.LOC112206744-LOC107973052-GGT2.revcompl                                        ------------------------------------------------------------ 18677
chimp54546-135457.revcompl.Pan.troglodytes.clone.rp43-41g5.GenBank:AC099533.36       ------------------------------------------------------------ 20190
human.FAM30B-LOC105372935-GGT2.NCBI.                                                 gccgcccagggcatcgccaacgaggacgccgcccagggcatcgccaaggaggacgccgcc 16766
human.FAM230E-LOC105377182-GGT3P.NCBI.                                               tccgtatacgacatcgctaatgaggatgccatatatgacatcgcta---atgacaccgta 15833
human.FAM230J-LOC105372942-GGTLC5P.NCBI.rev.compl                                    tccgtatacgacatcgctaatgaggatgccatatatgacatcgcta---atgacaccgta 16208
human.FAM230A.ncRNA.18487127.18500594-GGTLC3.18516335.18518165.rev.compl.REF         gcc--------------------------------------------------------- 5200

chimp.LOC112206744-LOC107973052-GGT2.revcompl                                        ------------------------------------------------------------ 18677
chimp54546-135457.revcompl.Pan.troglodytes.clone.rp43-41g5.GenBank:AC099533.36       ------------------------------------------------------------ 20190
human.FAM30B-LOC105372935-GGT2.NCBI.                                                 cagggcatcgccaacgagggcgccgcccagggcatcgccaaagaggacgccgcccagggc 16826
human.FAM230E-LOC105377182-GGT3P.NCBI.                                               caaggca-cgctaacgaggatgctgtacacgacatcgctaatgaggacaccatacaaggc 15892
human.FAM230J-LOC105372942-GGTLC5P.NCBI.rev.compl                                    caaggca-cactaacgaggacgctgtacacgacatcgctaatgaggacaccatacaaggc 16267
human.FAM230A.ncRNA.18487127.18500594-GGTLC3.18516335.18518165.rev.compl.REF         ------------------------------------------------------------ 5200

chimp.LOC112206744-LOC107973052-GGT2.revcompl                                        ------------------------------------------------------------ 18677
chimp54546-135457.revcompl.Pan.troglodytes.clone.rp43-41g5.GenBank:AC099533.36       ------------------------------------------------------------ 20190
human.FAM30B-LOC105372935-GGT2.NCBI.                                                 atcgctaacgaggccgccgacaagggcatcgccaacgaggatgccgcccacggcatcgcc 16886
human.FAM230E-LOC105377182-GGT3P.NCBI.                                               atcggtaatgaggacgttgtatatgacatcgctaacaaggacactctacaagccgtcgct 15952
human.FAM230J-LOC105372942-GGTLC5P.NCBI.rev.compl                                    attggtaatgaggacgttgtatatgacatcgctaacaaggacactctacaagccgtcgct 16327
human.FAM230A.ncRNA.18487127.18500594-GGTLC3.18516335.18518165.rev.compl.REF         ------------------------------------------------------------ 5200

chimp.LOC112206744-LOC107973052-GGT2.revcompl                                        ------------------------------------------------------------ 18677
chimp54546-135457.revcompl.Pan.troglodytes.clone.rp43-41g5.GenBank:AC099533.36       ------------------------------------------------------------ 20190
human.FAM30B-LOC105372935-GGT2.NCBI.                                                 aacgaggacgccgcccacggcatcgccagcga---------------------------- 16918
human.FAM230E-LOC105377182-GGT3P.NCBI.                                               aacaaggacactgtacacaacatcgctaatgac--------------------------- 15985
human.FAM230J-LOC105372942-GGTLC5P.NCBI.rev.compl                                    aacaaggacactgtacacaacatcgctaatgacggcaccatacaagacatcaccaatgag 16387
human.FAM230A.ncRNA.18487127.18500594-GGTLC3.18516335.18518165.rev.compl.REF         ------------------------------------------------------------ 5200

chimp.LOC112206744-LOC107973052-GGT2.revcompl                                        ------------------------------------------------------------ 18677
chimp54546-135457.revcompl.Pan.troglodytes.clone.rp43-41g5.GenBank:AC099533.36       ------------------------------------------------------------ 20190
human.FAM30B-LOC105372935-GGT2.NCBI.                                                 ------------------------------------------------------------ 16918
human.FAM230E-LOC105377182-GGT3P.NCBI.                                               ------------------------------------------------------------ 15985
human.FAM230J-LOC105372942-GGTLC5P.NCBI.rev.compl                                    ggcgctttatacgacattgctaatggcaccgacaaggcacgctaacgtggacactgtaca 16447
human.FAM230A.ncRNA.18487127.18500594-GGTLC3.18516335.18518165.rev.compl.REF         -------------------------------------------------------gtgca 5205

chimp.LOC112206744-LOC107973052-GGT2.revcompl                                        ------------------------------------------------------------ 18677
chimp54546-135457.revcompl.Pan.troglodytes.clone.rp43-41g5.GenBank:AC099533.36       ------------------------------------------------------------ 20190
human.FAM30B-LOC105372935-GGT2.NCBI.                                                 ------------------------------------------------------------ 16918
human.FAM230E-LOC105377182-GGT3P.NCBI.                                               ------------------------------------------------------------ 15985
human.FAM230J-LOC105372942-GGTLC5P.NCBI.rev.compl                                    cgacattgctaatgaggactccgtatacgacatcgctaatgagggtgccgtatatgacat 16507
human.FAM230A.ncRNA.18487127.18500594-GGTLC3.18516335.18518165.rev.compl.REF         cggcatcgctaatgaggactccgtatacgacatcgctaatgaggatgccatatatgacat 5265

chimp.LOC112206744-LOC107973052-GGT2.revcompl                                        ------------------------------------------------------------ 18677
chimp54546-135457.revcompl.Pan.troglodytes.clone.rp43-41g5.GenBank:AC099533.36       ------------------------------------------------------------ 20190
human.FAM30B-LOC105372935-GGT2.NCBI.                                                 ------------------------------------------------------------ 16918
human.FAM230E-LOC105377182-GGT3P.NCBI.                                               ------------------------------------------------------------ 15985
human.FAM230J-LOC105372942-GGTLC5P.NCBI.rev.compl                                    cgctaatgacaccgtacaaggcacgctaacgaggacgctgtacacgacatcgctaatgag 16567
human.FAM230A.ncRNA.18487127.18500594-GGTLC3.18516335.18518165.rev.compl.REF         cgctaatgacaccgtacaaggcacgctaaagaggacgctgtacacgacatcgctaatgag 5325

chimp.LOC112206744-LOC107973052-GGT2.revcompl                                        ------------------------------------------------------------ 18677
chimp54546-135457.revcompl.Pan.troglodytes.clone.rp43-41g5.GenBank:AC099533.36       ------------------------------------------------------------ 20190
human.FAM30B-LOC105372935-GGT2.NCBI.                                                 ------------------------------------------------------------ 16918
human.FAM230E-LOC105377182-GGT3P.NCBI.                                               ------------------------------------------------------------ 15985
human.FAM230J-LOC105372942-GGTLC5P.NCBI.rev.compl                                    gacaccatacaaggcatcggtaatgaggacgctgtatacgacatcgctaacgaggacacc 16627
human.FAM230A.ncRNA.18487127.18500594-GGTLC3.18516335.18518165.rev.compl.REF         gacaccatacaaggcattggtaatgaggacgttgtatatgacatcgctaacaaggacact 5385

chimp.LOC112206744-LOC107973052-GGT2.revcompl                                        ------------------------------------------------------------ 18677
chimp54546-135457.revcompl.Pan.troglodytes.clone.rp43-41g5.GenBank:AC099533.36       ------------------------------------------------------------ 20190
human.FAM30B-LOC105372935-GGT2.NCBI.                                                 -----------------------------------------------ggacgccgcccac 16931
human.FAM230E-LOC105377182-GGT3P.NCBI.                                               ------------------------------------------------ggcaccgtacaa 15997
human.FAM230J-LOC105372942-GGTLC5P.NCBI.rev.compl                                    atacaagccgtcgctaacaaggacactgtacacaacatcgctaatgatggcaccgtacaa 16687
human.FAM230A.ncRNA.18487127.18500594-GGTLC3.18516335.18518165.rev.compl.REF         ctacaagccgtcgctaacaaggacactgtacacaacatcgctaatgacggcaccgtacaa 5445

chimp.LOC112206744-LOC107973052-GGT2.revcompl                                        ------------------------------------------------------------ 18677
chimp54546-135457.revcompl.Pan.troglodytes.clone.rp43-41g5.GenBank:AC099533.36       -----------CGTGATCGCCGTCCACGGCATCGCCAATGTGGTCGCCGTCCAGGGCATC 20239
human.FAM30B-LOC105372935-GGT2.NCBI.                                                 ggcatcgccagcgaggacgccgcccacggcatcgccagcgaggacgccgcccagggcatc 16991
human.FAM230E-LOC105377182-GGT3P.NCBI.                                               gacatcaccaatgagggcgctttatacgacattgctaatgataccgac-----aaggcac 16052
human.FAM230J-LOC105372942-GGTLC5P.NCBI.rev.compl                                    gacatcaccaatgagggcgctttatacgacattgctaatgataccgac-----aaggcac 16742
human.FAM230A.ncRNA.18487127.18500594-GGTLC3.18516335.18518165.rev.compl.REF         gacatcaccaatgagggcgctttatacgacattgctaatgataccgac-----aaggcac 5500

chimp.LOC112206744-LOC107973052-GGT2.revcompl                                        ------------------------------------------------------------ 18677
chimp54546-135457.revcompl.Pan.troglodytes.clone.rp43-41g5.GenBank:AC099533.36       GCCAACGAGGACGCCACCCAGGGCATCGCCAAGGAGGACGCCGTCCACGGCATCGCCAAC 20299
human.FAM30B-LOC105372935-GGT2.NCBI.                                                 gccaacgaggacgccatccagggcatcgccaaggagtacgccgtccacggcatcgccaac 17051
human.FAM230E-LOC105377182-GGT3P.NCBI.                                               gctaacgcggacactgtacacgacatcgctaatgaggactccgtatacgacatcgctaat 16112
human.FAM230J-LOC105372942-GGTLC5P.NCBI.rev.compl                                    gctatcgtggacgctgtacacgacatcgctaatgaggactccgtatacgacatcgctaat 16802
human.FAM230A.ncRNA.18487127.18500594-GGTLC3.18516335.18518165.rev.compl.REF         gctaacgcggacactgtacacgacatcgctaatgaggactccgtataccacatcgctaat 5560

chimp.LOC112206744-LOC107973052-GGT2.revcompl                                        ------------------------------------------------------------ 18677
chimp54546-135457.revcompl.Pan.troglodytes.clone.rp43-41g5.GenBank:AC099533.36       GTGGTCGCCGTCCAGGGCATCGCCAACGTGGTCGCCGTCCAGGGCATCACCAAGGAGGAC 20359
human.FAM30B-LOC105372935-GGT2.NCBI.                                                 gaggacgccgcccagggcatcgccaacgaggacgccgcccagggcatcgccaacgaggac 17111
human.FAM230E-LOC105377182-GGT3P.NCBI.                                               gagggtgccgtatatgacatcgcta---atgacaccgtacaaggca-cgctaacgaggac 16168
human.FAM230J-LOC105372942-GGTLC5P.NCBI.rev.compl                                    gagggtgccgtatatgacatcgcta---atgacaccgtacaaggca-cgctaacgaggac 16858
human.FAM230A.ncRNA.18487127.18500594-GGTLC3.18516335.18518165.rev.compl.REF         gagggtgccgtatatgacatcgcta---atgacaccgtacaaggca-cgctaacgaggac 5616

chimp.LOC112206744-LOC107973052-GGT2.revcompl                                        ------------------------------------------------------------ 18677
chimp54546-135457.revcompl.Pan.troglodytes.clone.rp43-41g5.GenBank:AC099533.36       GCCGTCCATGGCATCGCCAAGGAGGACGCCGCCCAGGGCATCGCCAACGTGGTCGCCGTC 20419
human.FAM30B-LOC105372935-GGT2.NCBI.                                                 gccgcccagggcatcgccaaggaggacgccgcccagggcatcgccaaggaggacgccgcc 17171
human.FAM230E-LOC105377182-GGT3P.NCBI.                                               gctgtacacgacatcgctaatgaggacaccatacaaggcatcggtaatgaggac------ 16222
human.FAM230J-LOC105372942-GGTLC5P.NCBI.rev.compl                                    gctgtacacgacatcgctaatgaggacaccatacaaggcatcggtaatgaggac------ 16912
human.FAM230A.ncRNA.18487127.18500594-GGTLC3.18516335.18518165.rev.compl.REF         gctgtacacgacatcgctaatgaggacaccatacaaggcatcggtaatgaggac------ 5670

chimp.LOC112206744-LOC107973052-GGT2.revcompl                                        ------------------------------------------------------------ 18677
chimp54546-135457.revcompl.Pan.troglodytes.clone.rp43-41g5.GenBank:AC099533.36       CAGGGCATCGCCAACGAGGACGCCGCCCAGGGCATCGCCAAGGAGGACGCCGTCCACGGC 20479
human.FAM30B-LOC105372935-GGT2.NCBI.                                                 cagggcatcgccaacgagggcgccgcccagggcatcgccaaagaggacgccgcccacggc 17231
human.FAM230E-LOC105377182-GGT3P.NCBI.                                               ------------------------------------------------------------ 16222
human.FAM230J-LOC105372942-GGTLC5P.NCBI.rev.compl                                    ------------------------------------------------------------ 16912
human.FAM230A.ncRNA.18487127.18500594-GGTLC3.18516335.18518165.rev.compl.REF         ------------------------------------------------------------ 5670

chimp.LOC112206744-LOC107973052-GGT2.revcompl                                        ------------------------------------------------------------ 18677
chimp54546-135457.revcompl.Pan.troglodytes.clone.rp43-41g5.GenBank:AC099533.36       ATCGCCAACGAGGACGCCGCCCAGGGCATCGCCAACGAGGACGCCGCCCAGGGCATCGC- 20538
human.FAM30B-LOC105372935-GGT2.NCBI.                                                 atcgccaacgaggacgccgcccagggcatcgccaacgaggacgccgcccacggaatcgcc 17291
```

```
human.FAM230E-LOC105377182-GGT3P.NCBI.                                          ------------------------------------------------------------ 16222
human.FAM230J-LOC105372942-GGTLC5P.NCBI.rev.compl                               ------------------------------------------------------------ 16912
human.FAM230A.ncRNA.18487127.18500594-GGTLC3.18516335.18518165.rev.compl.REF    ------------------------------------------------------------ 5670

chimp.LOC112206744-LOC107973052-GGT2.revcompl                                   ------------------------------------------------------------ 18677
chimp54546-135457.revcompl.Pan.troglodytes.clone.rp43-41g5.GenBank:AC099533.36  ------------------------------------------------------------ 20538
human.FAM30B-LOC105372935-GGT2.NCBI.                                            agtgaggacgccgcccacggcatcgccagcgaggacgccgtccagggcatcgacaaggag 17351
human.FAM230E-LOC105377182-GGT3P.NCBI.                                          ------------------------------------------------------------ 16222
human.FAM230J-LOC105372942-GGTLC5P.NCBI.rev.compl                               ------------------------------------------------------------ 16912
human.FAM230A.ncRNA.18487127.18500594-GGTLC3.18516335.18518165.rev.compl.REF    ------------------------------------------------------------ 5670

chimp.LOC112206744-LOC107973052-GGT2.revcompl                                   ------------------------------------------------------------ 18677
chimp54546-135457.revcompl.Pan.troglodytes.clone.rp43-41g5.GenBank:AC099533.36  ------------------------------------------------------------ 20538
human.FAM30B-LOC105372935-GGT2.NCBI.                                            gatgccgcccagggcatcgccaacgaggacgccgcccagggcatcgccaacgaggacgcc 17411
human.FAM230E-LOC105377182-GGT3P.NCBI.                                          ------------------------------------------------------------ 16222
human.FAM230J-LOC105372942-GGTLC5P.NCBI.rev.compl                               ------------------------------------------------------------ 16912
human.FAM230A.ncRNA.18487127.18500594-GGTLC3.18516335.18518165.rev.compl.REF    ------------------------------------------------------------ 5670

chimp.LOC112206744-LOC107973052-GGT2.revcompl                                   --------catcgctaatgaggatgccatatatgacatcgctaatgacaccgtacaaggc 18729
chimp54546-135457.revcompl.Pan.troglodytes.clone.rp43-41g5.GenBank:AC099533.36  --------CATCGCTAATGAGGATGCCATATATGACATCGCTAATGACACCGTACAAGGC 20590
human.FAM30B-LOC105372935-GGT2.NCBI.                                            gcccagggcatcgccaaagaggacgccgcccacggcatcgccaacgagctgtatacgaca 17471
human.FAM230E-LOC105377182-GGT3P.NCBI.                                          -----------------------------------------------gctgtatacgaca 16235
human.FAM230J-LOC105372942-GGTLC5P.NCBI.rev.compl                               -----------------------------------------------gctgtatacgaca 16925
human.FAM230A.ncRNA.18487127.18500594-GGTLC3.18516335.18518165.rev.compl.REF    -----------------------------------------------gctgtatacgaca 5683

chimp.LOC112206744-LOC107973052-GGT2.revcompl                                   atgctaacgaagacatcgtacaagccgtcgctaacaagtacactgtacacaacatcgcta 18789
chimp54546-135457.revcompl.Pan.troglodytes.clone.rp43-41g5.GenBank:AC099533.36  ATGCTAACGAGGACATCGTACAAGCCGTCGCTAACAAGTACACTGTACACAACATCGCTA 20650
human.FAM30B-LOC105372935-GGT2.NCBI.                                            tcgctaacgaggacaccctacaagccgtcgctaacaagtacactgtacacaacatcgcta 17531
human.FAM230E-LOC105377182-GGT3P.NCBI.                                          tcgctaacgaggacaccatacaagccgtcgctaacaaggacactgtacacaacatcgcta 16295
human.FAM230J-LOC105372942-GGTLC5P.NCBI.rev.compl                               tcgctaacgaggacaccatacaagccgtcactaacaaggacactgtacacaacatcgcta 16985
human.FAM230A.ncRNA.18487127.18500594-GGTLC3.18516335.18518165.rev.compl.REF    tcgctaacgaggacaccatacaagccgtcgctaacaaggacactgtacacaacatcgcta 5743
                                                                                  ******** **** * *********** ******** ********************

chimp.LOC112206744-LOC107973052-GGT2.revcompl                                   atgagggcactgtacaagacatcaccaatgaggacgctttatacgacattgctaatggca 18849
chimp54546-135457.revcompl.Pan.troglodytes.clone.rp43-41g5.GenBank:AC099533.36  ATGAGGGCACTGTACAAGACATCACCAATGAGGACGCTTTATACGACATTGCTAATGGCA 20710
human.FAM30B-LOC105372935-GGT2.NCBI.                                            atgagggcactgtacaagacatcaccaatgagggcgctttatacgacattgctaatggca 17591
human.FAM230E-LOC105377182-GGT3P.NCBI.                                          atgacggcaccgtacaagacatcaccaatgagggcgctttatacgacattgctaatgata 16355
human.FAM230J-LOC105372942-GGTLC5P.NCBI.rev.compl                               atgacggcaccgtacaagacatcaccaatgagggcgctttatacgacattgctaatgata 17045
human.FAM230A.ncRNA.18487127.18500594-GGTLC3.18516335.18518165.rev.compl.REF    atgacggcaccgtacaagacatcaccaatgagggcgctttatacgacattgctaatgata 5803
                                                                                **** ***** ********************* **********************  *

chimp.LOC112206744-LOC107973052-GGT2.revcompl                                   ccgacaaggcacgctaacgtggatgctgtacacgacattgctaatgaggacaccgtataa 18909
chimp54546-135457.revcompl.Pan.troglodytes.clone.rp43-41g5.GenBank:AC099533.36  CCGACAAGGCACGCTAACGTGGATGCTGTACACGACATTGCTAATGAGGACACCGTATAA 20770
human.FAM30B-LOC105372935-GGT2.NCBI.                                            ccgacaaggcacgctaacgtggacgctgtacacgacattgctaatgaggacagcgtataa 17651
human.FAM230E-LOC105377182-GGT3P.NCBI.                                          ccgacaaggcacgctaacgtggacgctgtacacgacattgctaatgaggacaccgtataa 16415
human.FAM230J-LOC105372942-GGTLC5P.NCBI.rev.compl                               ccgacaaggcacgctaacgtggacgctgtacacgacattgctaatgaggacaccgtataa 17105
human.FAM230A.ncRNA.18487127.18500594-GGTLC3.18516335.18518165.rev.compl.REF    ccgacaaggcacgctaacgtggacgctgtacacgacattgctaatgaggacaccgtataa 5863
                                                                                *********************** ***************************** *******

chimp.LOC112206744-LOC107973052-GGT2.revcompl                                   gacatcgctagtaactatcgcaagaacaaaaaaccaaacaccgcatattctcactcatag 18969
chimp54546-135457.revcompl.Pan.troglodytes.clone.rp43-41g5.GenBank:AC099533.36  GACATCGCTAGTAACTATCGCAAGAACAAAAAACCAAACACCGCATATTCTCACTCATAG 20830
human.FAM30B-LOC105372935-GGT2.NCBI.                                            gacatcgctagtaactatcgcaagaacaaaaaaccaaacaccgcatattctcactcatag 17711
human.FAM230E-LOC105377182-GGT3P.NCBI.                                          gacatcgctagtaactatcgcaagaacaaaaaaccaaacaccgcatattctcactcatag 16475
human.FAM230J-LOC105372942-GGTLC5P.NCBI.rev.compl                               gacatcgctagtaactatcgcaagaacaaaaaaccaaacaccgcatattctcactcatag 17165
human.FAM230A.ncRNA.18487127.18500594-GGTLC3.18516335.18518165.rev.compl.REF    gacatcgctagtaactatcgcaagaacaaaaaaccaaacaccgcatattctcactcatag 5923
                                                                                ************************************************************

chimp.LOC112206744-LOC107973052-GGT2.revcompl                                   gtgggaattgaacaatgagatcacatggacacaggaaggggaatatcacactctggggac 19029
chimp54546-135457.revcompl.Pan.troglodytes.clone.rp43-41g5.GenBank:AC099533.36  GTGGGAATTGAACAATGAGATCACATGGACACAGGAAGGGGAATATCACACTCTGGGGAC 20890
human.FAM30B-LOC105372935-GGT2.NCBI.                                            atgggaattgaacaatgagatcacatggacacaggaaggggaatatcacactctggggac 17771
human.FAM230E-LOC105377182-GGT3P.NCBI.                                          gtgggaattgaacaatgagatcacatggacacaggaaggggaatatcacattctggggac 16535
human.FAM230J-LOC105372942-GGTLC5P.NCBI.rev.compl                               gtgggaattgaacaatgagatcacatggacacaggaaggggaatatcacactctggggac 17225
human.FAM230A.ncRNA.18487127.18500594-GGTLC3.18516335.18518165.rev.compl.REF    gtgggaattgaacaatgagatcacatggacacaggaaggggaatatcacattctggggac 5983
                                                                                 ************************************************** *********

chimp.LOC112206744-LOC107973052-GGT2.revcompl                                   tgttgtggggtggggggagggggagggatagcatcgggagatatacctaatgctagatg 19089
chimp54546-135457.revcompl.Pan.troglodytes.clone.rp43-41g5.GenBank:AC099533.36  TGTTGTGG---TGGCGGGGGGGGAAGGGATAGCATCGGGAGATATACCTAATGCTAGATG 20947
human.FAM30B-LOC105372935-GGT2.NCBI.                                            tgttgtggggttgg----ggggggagggatagcatcgggagatatacctaatgctagatg 17827
human.FAM230E-LOC105377182-GGT3P.NCBI.                                          tgttgtggggtggggggagggggagggatagcatcgggagatatacctaatgctagatg 16595
human.FAM230J-LOC105372942-GGTLC5P.NCBI.rev.compl                               tcttgtggggtggg----ggggggagggatagcatcgggagatatacctaatgctagatg 17281
human.FAM230A.ncRNA.18487127.18500594-GGTLC3.18516335.18518165.rev.compl.REF    tgttgtggggtggggggagggggagggatagcatcgggagatatacctaatgctagatg 6043
                                                                                * ******    **    ***** ************************************

chimp.LOC112206744-LOC107973052-GGT2.revcompl                                   atgagttagtgggtgcagcgcaccagcatggcacatgtatacatatgtaactaacctgca 19149
chimp54546-135457.revcompl.Pan.troglodytes.clone.rp43-41g5.GenBank:AC099533.36  ATGAGTTAGTGGGTGCAGCGCACCAGCATGGCACATGTATACATATGTAACTAACCTGCA 21007
human.FAM30B-LOC105372935-GGT2.NCBI.                                            atgagttagtgggtgcagcgcaccagcacggcacatgtatacatatgtaactaacctgca 17887
human.FAM230E-LOC105377182-GGT3P.NCBI.                                          atgagttagtgggtgcagcgcaccagcgtggcacatgtatacatatgtaactaacctgca 16655
human.FAM230J-LOC105372942-GGTLC5P.NCBI.rev.compl                               atgagttagtgggtgcagcgcaccagcgtggcacatgtatacatatgtaactaacctgca 17341
human.FAM230A.ncRNA.18487127.18500594-GGTLC3.18516335.18518165.rev.compl.REF    atgagttagtgggtgcagcgcaccagcgtggcacatgtatacatatgtaactaacctgca 6103
                                                                                ***************************  *******************************

chimp.LOC112206744-LOC107973052-GGT2.revcompl                                   caatgtgcacatgtaccctaaaacttaaagtatatatataaaaaaaaagacatcgctagt 19209
chimp54546-135457.revcompl.Pan.troglodytes.clone.rp43-41g5.GenBank:AC099533.36  CAATGTGCACATGTACCCTAAAACTTAAAG--TATATATAAAAAAAAAGACATCGCTAGT 21065
human.FAM30B-LOC105372935-GGT2.NCBI.                                            caatgtgcacatgtaccctaaaacttaaagtatatatataaaaaaaaaagacatcgctagt 17947
human.FAM230E-LOC105377182-GGT3P.NCBI.                                          caatgtgcacatgtaccctaaaacttaaagtatatat--aaaaaaaaagacatcgctagt 16713
human.FAM230J-LOC105372942-GGTLC5P.NCBI.rev.compl                               caatgtgcacatgtaccctaaaacttaaagtatatataaaaaaaaaaagacatcgctagt 17401
human.FAM230A.ncRNA.18487127.18500594-GGTLC3.18516335.18518165.rev.compl.REF    caatgtgcacatgtaccctaaaacttaaagtatatat-aaaaaaaaaagacatcgctagt 6162
                                                                                ******************************  *****  *********************

chimp.LOC112206744-LOC107973052-GGT2.revcompl                                   gagcacgctgtatacgacatcactaatgaggatgctatatatgacatcgctgatgaggac 19269
chimp54546-135457.revcompl.Pan.troglodytes.clone.rp43-41g5.GenBank:AC099533.36  GAGCACGCTGTATACGACATCGCTAATGAGGACGCTATATATGACATCGCTGATGAGGAC 21125
human.FAM30B-LOC105372935-GGT2.NCBI.                                            gagcacgctgtatacgacatcg-------------------------------------- 17969
human.FAM230E-LOC105377182-GGT3P.NCBI.                                          gagcacgctgtatacgacatcg-------------------------------------- 16735
human.FAM230J-LOC105372942-GGTLC5P.NCBI.rev.compl                               gagcacgctgtatacgacatcg-------------------------------------- 17423
human.FAM230A.ncRNA.18487127.18500594-GGTLC3.18516335.18518165.rev.compl.REF    gagcacgctgtatacgacatcg-------------------------------------- 6184
                                                                                ********************

chimp.LOC112206744-LOC107973052-GGT2.revcompl                                   attgtatacgacatcactaatgaggacaccatacaaggcatcgctaacaatgatgctgta 19329
chimp54546-135457.revcompl.Pan.troglodytes.clone.rp43-41g5.GenBank:AC099533.36  ATTGTATACGACATCACTAATGAGGACACCATACAAGGCATCGCTAACGATGACGCTGTA 21185
human.FAM30B-LOC105372935-GGT2.NCBI.                                            ----------------ctaatgaggacaccatacaaggcatcgctaacgatgacgctgta 18013
human.FAM230E-LOC105377182-GGT3P.NCBI.                                          ----------------ctaatgaggacaccatacaaggcatcgctaacgatgacgctgta 16779
human.FAM230J-LOC105372942-GGTLC5P.NCBI.rev.compl                               ----------------ctaatgaggacaccatacaaggcatcgctaacgatgacgctgta 17467
human.FAM230A.ncRNA.18487127.18500594-GGTLC3.18516335.18518165.rev.compl.REF    ----------------ctaatgaggacaccatacaaggcatcgctaacgatgacgctgta 6228
                                                                                                ******************************** **** ******

chimp.LOC112206744-LOC107973052-GGT2.revcompl                                   cacaacatcactaatgatgacaccgtataacacatcgctaattatgacgctgtatacgac 19389
chimp54546-135457.revcompl.Pan.troglodytes.clone.rp43-41g5.GenBank:AC099533.36  CACAACATCGCTAATGATGACACCGTATAAGACATCGCTAATTATGACGCTGTATACGAC 21245
human.FAM30B-LOC105372935-GGT2.NCBI.                                            cacaacatcactaatgatgacaccgtataagacatcgctaattatgacgctgtatacgac 18073
human.FAM230E-LOC105377182-GGT3P.NCBI.                                          cacaacatcactaatgatgacaccgtataagacatcgctaattatgacgctgtatacgac 16839
human.FAM230J-LOC105372942-GGTLC5P.NCBI.rev.compl                               cacaacatcactaatgatgacaccgtataagacatcgctaattatgacgctgtatacgac 17527
human.FAM230A.ncRNA.18487127.18500594-GGTLC3.18516335.18518165.rev.compl.REF    cacaacatcactaatgatgacaccgtataagacatcgctaattatgacgctgtatacgac 6288
                                                                                ********* ******************** *****************************

chimp.LOC112206744-LOC107973052-GGT2.revcompl                                   atcgctaatgacaccgtacaaggcacactaatgaggatgctgtacacgacatcactaatg 19449
chimp54546-135457.revcompl.Pan.troglodytes.clone.rp43-41g5.GenBank:AC099533.36  ATCGCTAATGACACCGTACAAGGCACGCTAACGAGGATGCTGTACACGACATCACTAATG 21305
human.FAM30B-LOC105372935-GGT2.NCBI.                                            atcgctaatgacaccgtacaaggcacgctaacgaggatgctgtacacgacatcactaatg 18133
human.FAM230E-LOC105377182-GGT3P.NCBI.                                          atcgctaatgacaccgtacgaggcacgctaacaaggatgctgtacacaacgtcgctaatg 16899
human.FAM230J-LOC105372942-GGTLC5P.NCBI.rev.compl                               atcgctaatgacaccgtacgaggcacactaacaaggatgctgtacacaacatcgctaatg 17587
human.FAM230A.ncRNA.18487127.18500594-GGTLC3.18516335.18518165.rev.compl.REF    atcgctaatgacaccgtacgaggcacgctaacaaggatgctgtacacaacatcgctaatg 6348
                                                                                ******************* ****** ****  ************** ** ** ******

chimp.LOC112206744-LOC107973052-GGT2.revcompl                                   aggacagtgtacaagccatcgctaatgaggacactgtatatgacatcgctaacgaggaca 19509
chimp54546-135457.revcompl.Pan.troglodytes.clone.rp43-41g5.GenBank:AC099533.36  AGGACAGTGTACAAGCCATCGCTAATGAGGACACTGTATATGACATCGCTAACGAGGACA 21365
human.FAM30B-LOC105372935-GGT2.NCBI.                                            aggacagtgtacaagccatcactaatgaggacactgtatatggcatcgctaacgaggaca 18193
human.FAM230E-LOC105377182-GGT3P.NCBI.                                          aggacagtgtacaagccatcgctaatgaggacactgtatatgacattgctaacgaggaca 16959
human.FAM230J-LOC105372942-GGTLC5P.NCBI.rev.compl                               aggacagtgtataagccatcgctaatgaggacactgtatatgacattgctaacgaggaca 17647
human.FAM230A.ncRNA.18487127.18500594-GGTLC3.18516335.18518165.rev.compl.REF    aggacagtgtacaagccatcgctaatgaggacactgtatatgacattgctaacgaggaca 6408
                                                                                *********** ******** ********************* *** *************

chimp.LOC112206744-LOC107973052-GGT2.revcompl                                   ctgtacaaggcattgctaacgaggacgctgtac---acatcgctaatgaggacaccatat 19566
chimp54546-135457.revcompl.Pan.troglodytes.clone.rp43-41g5.GenBank:AC099533.36  CTGTACAAGGCATTGCTAACGAGGACGCTGTACACAACACCGCTAATGAGGACACCATAT 21425
human.FAM30B-LOC105372935-GGT2.NCBI.                                            ctgtacaagccattgctaacgaggacgctgtacacaacatcgctaatgaggacaccatat 18253
human.FAM230E-LOC105377182-GGT3P.NCBI.                                          ctgtacaaggcattgctaacgaggacgctgtacacaacatcgctaatgaggacaccatat 17019
human.FAM230J-LOC105372942-GGTLC5P.NCBI.rev.compl                               ctgtacaaggcattgctaacgaggacgctgtacacaacatcgctaat---gacaccatat 17704
human.FAM230A.ncRNA.18487127.18500594-GGTLC3.18516335.18518165.rev.compl.REF    ctgtacaaggcattgctaacgaggacgctgtacacaacatcgctaatgaggacaccatat 6468
                                                                                ********* ***********************   *** *******   **********

chimp.LOC112206744-LOC107973052-GGT2.revcompl                                   aagacatcaccaatgaggatgctgtatatgacatcactaatgacacccacaaggcatgct 19626
chimp54546-135457.revcompl.Pan.troglodytes.clone.rp43-41g5.GenBank:AC099533.36  AAGACATCACCAATGAGGATGCTGTATATGACATCGCTAATGACACCCACAAGGCATGCT 21485
human.FAM30B-LOC105372935-GGT2.NCBI.                                            aagacatcaccaatgaggatgctgtatatgacatcgctaatggcacccacaaggcatgct 18313
human.FAM230E-LOC105377182-GGT3P.NCBI.                                          aagacatcaccaatgaggatgctgtatatgacatcgctaatgacacccacaaggcatgct 17079
```

```
human.FAM230J-LOC105372942-GGTLC5P.NCBI.rev.compl                               aagacatcaccaatgaggatgctgtatatgacatcgctaatgacacccacaaggcatgct 17764
human.FAM230A.ncRNA.18487127.18500594-GGTLC3.18516335.18518165.rev.compl.REF    aagacatcaccaatgaggatgctctatatgacatcgctaatgacacccacaaggcatgct 6528
                                                                                *********************** *********** ****** *****************

chimp.LOC112206744-LOC107973052-GGT2.revcompl                                   aacgaggacgctgtagacgacattgcttataaggacaccgtacaagacatcgctaacgag 19686
chimp54546-135457.revcompl.Pan.troglodytes.clone.rp43-41g5.GenBank:AC099533.36  AACGAGGACGCTGTAGACGACATTGCTTATAAGGACACCGTACAAGACATCGCTAACCAG 21545
human.FAM30B-LOC105372935-GGT2.NCBI.                                            aacgaggacgctgtagacgacattgcttataaggacaccgtacaagacatcgctaacgag 18373
human.FAM230E-LOC105377182-GGT3P.NCBI.                                          aacgaggacgctgtagacgacattgcttataaggacaccgtacaagacatcactaacgag 17139
human.FAM230J-LOC105372942-GGTLC5P.NCBI.rev.compl                               aacgaggacgctgtagacgacattgcttataaggacaccgtacaagacatcactaacgag 17824
human.FAM230A.ncRNA.18487127.18500594-GGTLC3.18516335.18518165.rev.compl.REF    aacgaggacgctgtagacgacattgcttataaggacaccgtacaagacatcgctaacgag 6588
                                                                                *************************************************** ***** **

chimp.LOC112206744-LOC107973052-GGT2.revcompl                                   gacgctgtatacgacatcgctaatgaggacgttgtatatgacatcgctaatgaggatgct 19746
chimp54546-135457.revcompl.Pan.troglodytes.clone.rp43-41g5.GenBank:AC099533.36  GACGCTGTATAC------------------------------------------------ 21557
human.FAM30B-LOC105372935-GGT2.NCBI.                                            gacgctgtatacgacatcgctaatgaggacgttgtatatgacatcgctaatgaggatgct 18433
human.FAM230E-LOC105377182-GGT3P.NCBI.                                          gacgctgtatacgacatcgctaatgaggacgttgtatatgacatcgctaatgaggatgct 17199
human.FAM230J-LOC105372942-GGTLC5P.NCBI.rev.compl                               gacgctgtatacgacatcgctaatgaggacgttgtatatgacatcgctaatgaggatgct 17884
human.FAM230A.ncRNA.18487127.18500594-GGTLC3.18516335.18518165.rev.compl.REF    gacgctgtatacgacatcgctaatgaggacgttgtatatgacatcgctaatgaggatgct 6648
                                                                                ************

chimp.LOC112206744-LOC107973052-GGT2.revcompl                                   ttacaagacatagctaatgaggttgctgtatatgatatcgctaatgaggacattgtatat 19806
chimp54546-135457.revcompl.Pan.troglodytes.clone.rp43-41g5.GenBank:AC099533.36  ---------------------------------GACATCGCTAATGAGGACACTGTATAT 21584
human.FAM30B-LOC105372935-GGT2.NCBI.                                            ttacaagacatagctaatgaggttgctgtatatgacatcgctaatgaggacattgtatat 18493
human.FAM230E-LOC105377182-GGT3P.NCBI.                                          ttacaagacatagcgaatgaggttgctgtatatgacatcgctaatgaggacattgtatat 17259
human.FAM230J-LOC105372942-GGTLC5P.NCBI.rev.compl                               ttacaagacatagcgaatgaggttgctgtatatgacatcgctaatgaggacattgtatat 17944
human.FAM230A.ncRNA.18487127.18500594-GGTLC3.18516335.18518165.rev.compl.REF    ttacaagacatagcgaatgaggttgctgtatatgacatcgctaatgaggacattgtatat 6708
                                                                                                                 ** **************** *******

chimp.LOC112206744-LOC107973052-GGT2.revcompl                                   gacatcgctaatgaggacgctctatacgacatcactaatgaggatgctgtatacaacatc 19866
chimp54546-135457.revcompl.Pan.troglodytes.clone.rp43-41g5.GenBank:AC099533.36  GACATCGCTAATGAGGACGCTCTATACGACATCACTAACGAGGACGCTGTATACAACATT 21644
human.FAM30B-LOC105372935-GGT2.NCBI.                                            gacatcgctaatgaggacgctctatacgacatcactaatgaggacgctgtatacaacatc 18553
human.FAM230E-LOC105377182-GGT3P.NCBI.                                          gacatcgctaatgaggacgctctatacgacatcactaatgaggacgctgtatacaacatc 17319
human.FAM230J-LOC105372942-GGTLC5P.NCBI.rev.compl                               gacatcgctaatgaggacgctctatacgacatcactaatgaggacgctgtatacaacatc 18004
human.FAM230A.ncRNA.18487127.18500594-GGTLC3.18516335.18518165.rev.compl.REF    gacatcgctaatgaggacgctctatacgacatcactaatgaggacgctgtatacaacatc 6768
                                                                                ************************************** ***** **************

chimp.LOC112206744-LOC107973052-GGT2.revcompl                                   gctaatgaggacgctgtatatggcatcgctaatgaggatgctgtatacgaattcgctaat 19926
chimp54546-135457.revcompl.Pan.troglodytes.clone.rp43-41g5.GenBank:AC099533.36  GCTAATGAGGACGCTGTATATGGCATCGCTAATGAGGATGCTGTATACAAATTCGCTAAT 21704
human.FAM30B-LOC105372935-GGT2.NCBI.                                            gctaatgaggacgctgtatatggcatcgctaatgaggatgctgtatacgaattcgctaat 18613
human.FAM230E-LOC105377182-GGT3P.NCBI.                                          gctaatgaggacgctgtatatggcatcgctaatgaggatgctgtatacgaattcgctaat 17379
human.FAM230J-LOC105372942-GGTLC5P.NCBI.rev.compl                               gctaatgaggacgctgtatatggcatcgctaatgaggatgctgtatacgaattcgctaat 18064
human.FAM230A.ncRNA.18487127.18500594-GGTLC3.18516335.18518165.rev.compl.REF    gctaatgaggacgctgtatatggcatcgctaatgaggatgctgtatacgaattcgctaat 6828
                                                                                ************************************************ ***********

chimp.LOC112206744-LOC107973052-GGT2.revcompl                                   aaggacgctgtatatgacattgctaatgaggacactatacaagacatctgtaaaaaagaa 19986
chimp54546-135457.revcompl.Pan.troglodytes.clone.rp43-41g5.GenBank:AC099533.36  AAGGACGCTGTATATGACATTGCTAATGAGGACACTGTACAAGACATCTGTAAAAAAGAA 21764
human.FAM30B-LOC105372935-GGT2.NCBI.                                            aaggacgctgtatatgacattgctaatgaggacactgtacaagacatctgtaaaaaagaa 18673
human.FAM230E-LOC105377182-GGT3P.NCBI.                                          aaggacgctgtatatgacattgctaatgaggacactgtacaagacatctgtaaaaaagaa 17439
human.FAM230J-LOC105372942-GGTLC5P.NCBI.rev.compl                               aaggacgctgtatatgacattgctaatgaggacactgtacaagacatctgtaaaaaagaa 18124
human.FAM230A.ncRNA.18487127.18500594-GGTLC3.18516335.18518165.rev.compl.REF    aaggacgctgtatatgacattgctaatgaggacactgtacaagacatctgtaaaaaagaa 6888
                                                                                ************************************* **********************

chimp.LOC112206744-LOC107973052-GGT2.revcompl                                   gatgctgccaatgtaagacacttttctttgtcttgaacagaaatgttactttcctggctt 20046
chimp54546-135457.revcompl.Pan.troglodytes.clone.rp43-41g5.GenBank:AC099533.36  GATGTTGCCAATGTAAGACACTTTTCTTTGTCTTGAACAGAAATGTTACTTTCCTGGCTT 21824
human.FAM30B-LOC105372935-GGT2.NCBI.                                            gatgctgccaatgtaagacacttttctttgtcttgaacagaaatgttactttcctggctt 18733
human.FAM230E-LOC105377182-GGT3P.NCBI.                                          gatgctgccaatgtaagacacttttctttgtcttgaacagaaatgttactttcctggctt 17499
human.FAM230J-LOC105372942-GGTLC5P.NCBI.rev.compl                               gatgctgccaatgtaagacacttttctttgtcttgaacagaaatgttactttcctggctt 18184
human.FAM230A.ncRNA.18487127.18500594-GGTLC3.18516335.18518165.rev.compl.REF    gatgctgccaatgtaagacacttttctttgtcttgaacagaaatgttactttcctggctt 6948
                                                                                **** *******************************************************

chimp.LOC112206744-LOC107973052-GGT2.revcompl                                   ctttccaatcagatgtagacatgaacatctgccagtgtgcattattgatgtcatctgcag 20106
chimp54546-135457.revcompl.Pan.troglodytes.clone.rp43-41g5.GenBank:AC099533.36  CTTTCCAATCAGACGTAGACATGAACATCTGCCAGTGTGAATTATCGATGTCATCTGCAG 21884
human.FAM30B-LOC105372935-GGT2.NCBI.                                            ctttccaatcagatgtagacatgaacatctgccagtgtgcattatcgatgtcatctgcag 18793
human.FAM230E-LOC105377182-GGT3P.NCBI.                                          ctttccaatcagatgtagacatgaacatctgccagtgtgcattatcgatgtcatctgcag 17559
human.FAM230J-LOC105372942-GGTLC5P.NCBI.rev.compl                               ctttccaatcagatgtagacatgaacatctgccagtgtgcattatggatgtcatctgcag 18244
human.FAM230A.ncRNA.18487127.18500594-GGTLC3.18516335.18518165.rev.compl.REF    ctttccaatcagatgtagacatgaacatctgccagtgtgcattatcgatgtcatctgcag 7008
                                                                                ************* ************************* ***** **************

chimp.LOC112206744-LOC107973052-GGT2.revcompl                                   tttaatcaaatgtagacatgaacatctgccaatgtggactatttatgacatctgcaattc 20166
chimp54546-135457.revcompl.Pan.troglodytes.clone.rp43-41g5.GenBank:AC099533.36  TTTAATCAAATGTAGACATGAACATCTGCCAATGTGGACTATTTATGACATCTGCAATTC 21944
human.FAM30B-LOC105372935-GGT2.NCBI.                                            tttaatcaaatgtagacatgaacatctgccaatgtggactatttatgacatctgcaattc 18853
human.FAM230E-LOC105377182-GGT3P.NCBI.                                          tttaatcaaatgtagacatgaacatctgccaatgtggactatttatgacatctgcaattc 17619
human.FAM230J-LOC105372942-GGTLC5P.NCBI.rev.compl                               tttaatcaaatgtagacatgaacatctgccaatgtggactatttatgacatctgcaattc 18304
human.FAM230A.ncRNA.18487127.18500594-GGTLC3.18516335.18518165.rev.compl.REF    tttaatcaaatgtagacatgaacatctgccaatgtggactatttatgacatctgcaattc 7068
                                                                                ************************************************************

chimp.LOC112206744-LOC107973052-GGT2.revcompl                                   ccttggtgtggtgctattgattggcagcctctcaccaacccatgccaggcacactggggt 20226
chimp54546-135457.revcompl.Pan.troglodytes.clone.rp43-41g5.GenBank:AC099533.36  CCTTGGTGTGGTGCTATTGATTGGCAGCCTCTCACCAACCCATGCCAGGCACACTGGGGT 22004
human.FAM30B-LOC105372935-GGT2.NCBI.                                            ccttggtgtggtgctattgattggcagcctctcaccaacccatgccaggcacactggggt 18913
human.FAM230E-LOC105377182-GGT3P.NCBI.                                          ccttggtgtggtgctattgattggcggcctctcaccaacccatgccaggcacactggggt 17679
human.FAM230J-LOC105372942-GGTLC5P.NCBI.rev.compl                               ccttggtgtggtgctattgattggcagcctctcaccaacccatgccaggcacactggggt 18364
human.FAM230A.ncRNA.18487127.18500594-GGTLC3.18516335.18518165.rev.compl.REF    ccttggtgtggtgctattgattggcggcctctcaccaacccatgccaggcacactggggt 7128
                                                                                ************************* **********************************

chimp.LOC112206744-LOC107973052-GGT2.revcompl                                   gtggtagatggcagcatccacgatccactgcaatgcagaggtgtttccctccacagcagt 20286
chimp54546-135457.revcompl.Pan.troglodytes.clone.rp43-41g5.GenBank:AC099533.36  GTGGTAGATGGCAGCATCCACGATCCACTGCAATGCAGAGGTGTTTCCCTCCACAGCAGT 22064
human.FAM30B-LOC105372935-GGT2.NCBI.                                            gtggtagatggcagcatccacgatccactgcaatgcagaggtgtttccctccacagcagt 18973
human.FAM230E-LOC105377182-GGT3P.NCBI.                                          gtggtagatggcagcatccacgatccactgcaatgcagaggtgtttccctccacagcagt 17739
human.FAM230J-LOC105372942-GGTLC5P.NCBI.rev.compl                               gtggtagatggcagcatccacgatccactgcaatgcagaggtgtttccctccacagcagt 18424
human.FAM230A.ncRNA.18487127.18500594-GGTLC3.18516335.18518165.rev.compl.REF    gtggtagatggcagcatccacgatccactgcaatgcagaggtgtttccctccacagcagt 7188
                                                                                ************************************************************

chimp.LOC112206744-LOC107973052-GGT2.revcompl                                   tttccccatggattaagagttgtgaaactgccaatctagatacactttaaagataaatt 20346
chimp54546-135457.revcompl.Pan.troglodytes.clone.rp43-41g5.GenBank:AC099533.36  TTTCCCCCATGGATTAAGAGTTGTGAAACTGCCAATCTAGATACACTTTAAAGATAAATT 22124
human.FAM30B-LOC105372935-GGT2.NCBI.                                            tttcccccatggattaagagttgtgaaactgccaatctagatacactttaaagataaatt 19033
human.FAM230E-LOC105377182-GGT3P.NCBI.                                          tttcccccatggattaagagttgtgaaactgccaatctagatacactttaaagataaatt 17799
human.FAM230J-LOC105372942-GGTLC5P.NCBI.rev.compl                               tttcccccatggattaagagttgtgaaactgccaatctagatacactttaaagataaatt 18484
human.FAM230A.ncRNA.18487127.18500594-GGTLC3.18516335.18518165.rev.compl.REF    tttcccccatggattaagagttgtgaaactgccaatctagatacactttaaagataaatt 7248
                                                                                ************************************************************

chimp.LOC112206744-LOC107973052-GGT2.revcompl                                   ctgtgggaaaaggtcttgtcttttccacaggtgtcttccgtgccagttttggggggacttc 20406
chimp54546-135457.revcompl.Pan.troglodytes.clone.rp43-41g5.GenBank:AC099533.36  CTGTGGGAAAAGGTCTTGTCTTTTCCACAGGTGTCTTCCGTGCCAGTTTTGGGGGACTTC 22184
human.FAM30B-LOC105372935-GGT2.NCBI.                                            ctgtgggaaaaggtcttgtcttttccacaggtgtcttccgtgccagttttgggggacttc 19093
human.FAM230E-LOC105377182-GGT3P.NCBI.                                          ctgtgggaaaaggtcttgtcttttccacaggtgtcttccgtgccagttttggggga---- 17855
human.FAM230J-LOC105372942-GGTLC5P.NCBI.rev.compl                               ctgtgggaaaaggtcttgtcttttccacaggtgtcttccgtgccagttttgggggacttt 18544
human.FAM230A.ncRNA.18487127.18500594-GGTLC3.18516335.18518165.rev.compl.REF    ctgtgggaaaaggtcttgtcttttccacaggtgtcttccgtgccagttttgggggacttt 7308
                                                                                ********************************************************

chimp.LOC112206744-LOC107973052-GGT2.revcompl                                   gacctttgactcaatcactataccccttcttatttctctctcaagttgtcgagagacta 20466
chimp54546-135457.revcompl.Pan.troglodytes.clone.rp43-41g5.GenBank:AC099533.36  GACCTTTGACTCAATCACTATACCCCTTCTTATTTCTCTCTCAAGTTATCGAGAGACTA 22244
human.FAM30B-LOC105372935-GGT2.NCBI.                                            gacctttgactcaatcactataccccttcctatttctctctcaagttgtcgagagacta 19153
human.FAM230E-LOC105377182-GGT3P.NCBI.                                          --------------------------cttcatatttctctctcaagttgtcgagagacta 17890
human.FAM230J-LOC105372942-GGTLC5P.NCBI.rev.compl                               gacctttgactcaatcactataccccttcttatttctctctcaagttgtcgagagacta 18604
human.FAM230A.ncRNA.18487127.18500594-GGTLC3.18516335.18518165.rev.compl.REF    gacctttgactcaatcactataccccttcttatttctctctcaagttgtcgagagacta 7368
                                                                                                         **** ****************** ***********

chimp.LOC112206744-LOC107973052-GGT2.revcompl                                   tcagatctgtgtgacgtgtatggcatcatttcaccctcctaatgttttcttttctataat 20526
chimp54546-135457.revcompl.Pan.troglodytes.clone.rp43-41g5.GenBank:AC099533.36  TCAGATCTGTGTGACATGTATGGCATAATTTCACCCTCCTAATGTTTTCTTTTCTATAAT 22304
human.FAM30B-LOC105372935-GGT2.NCBI.                                            tcagatctgtgtgacgtgtatggcatcatttcaccctcctaatgttttcttttctataat 19213
human.FAM230E-LOC105377182-GGT3P.NCBI.                                          tcagatctgtgtgacgtgtatggcatcatttcaccctcctaatgttttcttttctataat 17950
human.FAM230J-LOC105372942-GGTLC5P.NCBI.rev.compl                               tcagatctgtgtgacgtgtatggcatcatttcaccctcctaatgttttcttttctataat 18664
human.FAM230A.ncRNA.18487127.18500594-GGTLC3.18516335.18518165.rev.compl.REF    tcagatctgtgtgacgtgtatggcatcatttcaccctcctaatgttttcttttctataat 7428
                                                                                *************** ********** *********************************

chimp.LOC112206744-LOC107973052-GGT2.revcompl                                   tgcaggagccattgacattggagaatgatacgtaccctgaaataactcacttcctgagga 20586
chimp54546-135457.revcompl.Pan.troglodytes.clone.rp43-41g5.GenBank:AC099533.36  TGCAGGAGCCATTGACACTGGAGAATGATACGTACCCTGAAATAACTCACTTCCTGAGGA 22364
human.FAM30B-LOC105372935-GGT2.NCBI.                                            tgcaggagccattgacactggagaatgatacgtaccctgaaataactcacttcctgagga 19273
human.FAM230E-LOC105377182-GGT3P.NCBI.                                          tgcaggagccattgacactggagaatgatacctaccctgaaataactcacttcctgagaa 18010
human.FAM230J-LOC105372942-GGTLC5P.NCBI.rev.compl                               tgcaggagccattgacactggagaatgatacctaccctgaaataactcacttcctgagga 18724
human.FAM230A.ncRNA.18487127.18500594-GGTLC3.18516335.18518165.rev.compl.REF    tgcaggagccattgacactggagaatgatacctaccctgaaataactcacttcctgagga 7488
                                                                                ***************** ************* ************************** *

chimp.LOC112206744-LOC107973052-GGT2.revcompl                                   aaaagcgccatctctagggtacagaaacctgattctgggctcctttttgggaaggaggatt 20646
chimp54546-135457.revcompl.Pan.troglodytes.clone.rp43-41g5.GenBank:AC099533.36  AAAAGTGCCATCTCTAGGGTACAGAAACCTGATTCTGGGCTCCTTTTGGGAAGGAGGATT 22424
human.FAM30B-LOC105372935-GGT2.NCBI.                                            aaaagcgccatctctagggtacagaaacctgattctgggctcctttttgggaaggaggatt 19333
human.FAM230E-LOC105377182-GGT3P.NCBI.                                          aaaagcgccatctctagggtacagaaacctgattctgggctcctttttgggaaggaggatt 18070
human.FAM230J-LOC105372942-GGTLC5P.NCBI.rev.compl                               aaaaacgccatctctagggtacagaaacctgattctgggctcctttttgggaaggaggatt 18784
human.FAM230A.ncRNA.18487127.18500594-GGTLC3.18516335.18518165.rev.compl.REF    aaaaacgccatctctagggtacagaaacctgattctgggctcctttttgggaaggaggatt 7548
                                                                                ****  ******************************************************

chimp.LOC112206744-LOC107973052-GGT2.revcompl                                   tggggtctggtgagagcaaatgattttgcaagtataaaacaatgtccagagaggctgtag 20706
chimp54546-135457.revcompl.Pan.troglodytes.clone.rp43-41g5.GenBank:AC099533.36  TGGGGTCTGGTGAGAGCAAATGATTTTGCAAGTATAAAACAATGTCCAGAGAGGCTGTAG 22484
human.FAM30B-LOC105372935-GGT2.NCBI.                                            tggggtctggtgagagcaaatgattttgcaagtataaaaccatgtccagagaggctgtag 19393
human.FAM230E-LOC105377182-GGT3P.NCBI.                                          tggggtctggtgagagcaaatgattttgcaagtataaaaccatgtccagagaggctgtag 18130
human.FAM230J-LOC105372942-GGTLC5P.NCBI.rev.compl                               tggagtctggtgagagcaaatgattttgcaagtataaaacaatgtccagagaggctgtag 18844
```

```
human.FAM230A.ncRNA.18487127.18500594-GGTLC3.18516335.18518165.rev.compl.REF    tggagtctggtgagagcaaatgattttgcaagtataaaacaatgtccagagaggctgtag 7608
                                                                                *** ************************************ ******************

chimp.LOC112206744-LOC107973052-GGT2.revcompl                                   ggatatctgtgagcccagaggaaacaccaggggatcctgtgcgaagcaccatggcttcag 20766
chimp54546-135457.revcompl.Pan.troglodytes.clone.rp43-41g5.GenBank:AC099533.36  GGATATCTGTGAGCCCAGAGGAAACACCAGGGGATCCTGTGCGAAGCACCATGGCTTCAG 22544
human.FAM30B-LOC105372935-GGT2.NCBI.                                            ggatatctgtgagcccagaggaaacaccaggggatcctgtgcgaagcaccatggcttcag 19453
human.FAM230E-LOC105377182-GGT3P.NCBI.                                          ggatatctgtgagcccagaggaaacaccaggggatcctgtgcgaagcaccatggcttcag 18190
human.FAM230J-LOC105372942-GGTLC5P.NCBI.rev.compl                               ggatatctgtgagcccagaggaaacaccaggggatcctgtgcgaagcaccatggcttcag 18904
human.FAM230A.ncRNA.18487127.18500594-GGTLC3.18516335.18518165.rev.compl.REF    ggatatctgtgagcccagaggaaacaccaggggatcctgtgcgaagcaccatggcttcag 7668
                                                                                ************************************************************

chimp.LOC112206744-LOC107973052-GGT2.revcompl                                   ctagggtgggaggagtgggtgggcctctctctaatgacttatcctggtgtttgtgtttct 20826
chimp54546-135457.revcompl.Pan.troglodytes.clone.rp43-41g5.GenBank:AC099533.36  CTAGGGTGGGAGGAGTGGGTGGGCCTCTCTCTAATGACTTATCCTGGTGTTTGTGTTTCT 22604
human.FAM30B-LOC105372935-GGT2.NCBI.                                            ctagggtgggaggagtgggtgggcctctctctaatgacttatcctggtgtttgtgtttct 19513
human.FAM230E-LOC105377182-GGT3P.NCBI.                                          ctagggtgggaggagtgggtgggcctctctctaatgacttatcctggtgtttgtgtttct 18250
human.FAM230J-LOC105372942-GGTLC5P.NCBI.rev.compl                               ctagggtgggaggagtgggtgggcctctctctaatgacttatcctggtgtttgtgtttct 18964
human.FAM230A.ncRNA.18487127.18500594-GGTLC3.18516335.18518165.rev.compl.REF    ctagggtgggaggagtgggtgggcctctctctaatgacttatcctggtgtttgtgtttct 7728
                                                                                ************************************************************

chimp.LOC112206744-LOC107973052-GGT2.revcompl                                   aaagatttgattgtggagagcatatctgatgatggggatttgtaggtaggtaactacttt 20886
chimp54546-135457.revcompl.Pan.troglodytes.clone.rp43-41g5.GenBank:AC099533.36  AAAGATTTGATTGTGGAGAGCATATCTGATGATGGGGATTTGTAGGTAGGTAACTACTTT 22664
human.FAM30B-LOC105372935-GGT2.NCBI.                                            aaagatttgattgtggagagcatatctgatgatggggatttgtaggtaggtaactgcttt 19573
human.FAM230E-LOC105377182-GGT3P.NCBI.                                          aaagatttgattgtggagagcatatctgatgatggggatttgtaggtaggtaactacttt 18310
human.FAM230J-LOC105372942-GGTLC5P.NCBI.rev.compl                               aaagatttgattgtggagagcatatctgatgatggggatttgtaggtaggtaactacttt 19024
human.FAM230A.ncRNA.18487127.18500594-GGTLC3.18516335.18518165.rev.compl.REF    aaagatttgattgtggagagcatatctgatgatggggatttgtaggtaggtaactacttt 7788
                                                                                ******************************************************* ****

chimp.LOC112206744-LOC107973052-GGT2.revcompl                                   ccacgtaagatccaattggagagagttcccaggggccttcggggtatccatgctgcttgg 20946
chimp54546-135457.revcompl.Pan.troglodytes.clone.rp43-41g5.GenBank:AC099533.36  CCACGTAAGATCCAATTGGAGAGAGTTCCCAGGGGCCTTCGGGGTATCCATGCTGCTTGG 22724
human.FAM30B-LOC105372935-GGT2.NCBI.                                            ccacgtaagatccaattggagagagttcccaggggccttcggggtatccatgctgcttgg 19633
human.FAM230E-LOC105377182-GGT3P.NCBI.                                          ccacgtaagatccaattggagagagttcccaggggccttcggggtatccatgctgcttgg 18370
human.FAM230J-LOC105372942-GGTLC5P.NCBI.rev.compl                               ccacgtaagatccaattggagagagttcccaggggccttcggggtatccatgctgcttgg 19084
human.FAM230A.ncRNA.18487127.18500594-GGTLC3.18516335.18518165.rev.compl.REF    ccacgtaagatccaattggagagagttcccaggggccttcggggtatccatgctgcttgg 7848
                                                                                ************************************************************

chimp.LOC112206744-LOC107973052-GGT2.revcompl                                   gaggttaagggagggggcatgaaatcaaaacgaaacaggaaatatgtgtcatattggatt 21006
chimp54546-135457.revcompl.Pan.troglodytes.clone.rp43-41g5.GenBank:AC099533.36  GAGGTTAAGGGAGGGGGCATGAAATCAAAACGAAACAGGAAATATGTGTCATATTGGATT 22784
human.FAM30B-LOC105372935-GGT2.NCBI.                                            gaggttaagggagggggcatgaaatcaaaacgaaacaggaaatatgtgtcatatcggatt 19693
human.FAM230E-LOC105377182-GGT3P.NCBI.                                          gaggttaagggagggggcatgaaatcaaaacaaaacaggaaatatgtgtcatattggact 18430
human.FAM230J-LOC105372942-GGTLC5P.NCBI.rev.compl                               gaggttaagggagggggcatgaaatcaaaaggaaacaggaaatatgtgtcatattggatt 19144
human.FAM230A.ncRNA.18487127.18500594-GGTLC3.18516335.18518165.rev.compl.REF    gaggttaagggagggggcatgaaatcaaaaggaaacaggaaatatgtgtcatattggatt 7908
                                                                                ******************************  ********************** *** *

chimp.LOC112206744-LOC107973052-GGT2.revcompl                                   tggtcttttccgggtttattggcataatagtaagaactgtctctctgggctatgagggtg 21066
chimp54546-135457.revcompl.Pan.troglodytes.clone.rp43-41g5.GenBank:AC099533.36  TGGTCTTTTCCGGGTTTATTGGCATAATAGTTAGAACTGTCTCTCTGGGCTATGAGGGTG 22844
human.FAM30B-LOC105372935-GGT2.NCBI.                                            tggtcttttccgggtttattggcataatagttagaactgtctctctgggctatgagggtg 19753
human.FAM230E-LOC105377182-GGT3P.NCBI.                                          tggtcttttccgggtttattggcataatagttagaactgtctctctgggctatgagggtg 18490
human.FAM230J-LOC105372942-GGTLC5P.NCBI.rev.compl                               tggtcttttccgggtttattggcataatagttagaactgtctctctgggctatgagggtg 19204
human.FAM230A.ncRNA.18487127.18500594-GGTLC3.18516335.18518165.rev.compl.REF    tggtcttttccgggtttattggcataatagttagaactgtctctctgggctatgagggtg 7968
                                                                                ******************************* ****************************

chimp.LOC112206744-LOC107973052-GGT2.revcompl                                   ctgtgttatttaaaggtggtctttcccagaacacctggccttttcttttccgcctctgcc 21126
chimp54546-135457.revcompl.Pan.troglodytes.clone.rp43-41g5.GenBank:AC099533.36  CTGTGTTATTTAAAGGTGGTCTTTCCCAGAGCACCTGGCCTTTTCTTTTCTGCCTCTGCC 22904
human.FAM30B-LOC105372935-GGT2.NCBI.                                            ctgtgttatttaaaggtggtctttcccagaacacctggccttttcttttctgcctctgcc 19813
human.FAM230E-LOC105377182-GGT3P.NCBI.                                          ctgtgttatttaaaggtggtctttcccagaacacctggccttttcttttctgcctctgcc 18550
human.FAM230J-LOC105372942-GGTLC5P.NCBI.rev.compl                               ctgtgatatttaaaggtggtctttcccagaacacctggccttttcttttctgcctctgcc 19264
human.FAM230A.ncRNA.18487127.18500594-GGTLC3.18516335.18518165.rev.compl.REF    ctgtgatatttaaaggtggtctttcccagaacacctggccttttcttttctgcctctgcc 8028
                                                                                ***** ************************ ******************* *********

chimp.LOC112206744-LOC107973052-GGT2.revcompl                                   aaacatcacagactttgggttggattagtcagcaccccttgggattgtgcagaagaggtt 21186
chimp54546-135457.revcompl.Pan.troglodytes.clone.rp43-41g5.GenBank:AC099533.36  AAACATCACAGCCTTTGGGTTGGATTAGTCAGCACCCCTTGGGATTGTGCAGAAGAGGTT 22964
human.FAM30B-LOC105372935-GGT2.NCBI.                                            aaacatcacagcctttgggttggattagtcagcaccccttgggattgtgcagaagaggtt 19873
human.FAM230E-LOC105377182-GGT3P.NCBI.                                          aaacatcacagcctttgggttggattagtcagcaccccttgggattgtgcagaagaggtt 18610
human.FAM230J-LOC105372942-GGTLC5P.NCBI.rev.compl                               aaacatcacagcctttgggttggattagtcagcaccccttgggattgtgcagaagaggtt 19324
human.FAM230A.ncRNA.18487127.18500594-GGTLC3.18516335.18518165.rev.compl.REF    aaacatcacagcctttgggttggattagtcagcaccccttgggattgtgcagaagaggtt 8088
                                                                                ********** *************************************************

chimp.LOC112206744-LOC107973052-GGT2.revcompl                                   tggggttgcatcgagtgtcacctgtggtgaacagaatctgagggacacaactctctcaca 21246
chimp54546-135457.revcompl.Pan.troglodytes.clone.rp43-41g5.GenBank:AC099533.36  TGGGGTTGCACCGAGTGTCACCTGTGGTGAACAGAATCTGAGGGACACAACTCTCTCACA 23024
human.FAM30B-LOC105372935-GGT2.NCBI.                                            tggggttgcattgagtgtcacctgtggtgaacagaatctgagggacacaactctctcaca 19933
human.FAM230E-LOC105377182-GGT3P.NCBI.                                          tggggttgcatcgagtgtcacctgtggtgaacagaatctgagggacacaactctctcaca 18670
human.FAM230J-LOC105372942-GGTLC5P.NCBI.rev.compl                               tggggttgcatcgagtgtcacctgtggtgaacagaatctgagggacacaactctctcaca 19384
human.FAM230A.ncRNA.18487127.18500594-GGTLC3.18516335.18518165.rev.compl.REF    tggggttgcatcgagtgtcacctgtggtgaacagaatctgagggacacaactctctcaca 8148
                                                                                **********  ************************************************

chimp.LOC112206744-LOC107973052-GGT2.revcompl                                   ggcacttccttcaacctggagacagagttctcctggtgtgtgcccaggggtgcaggagaa 21306
chimp54546-135457.revcompl.Pan.troglodytes.clone.rp43-41g5.GenBank:AC099533.36  GGCACTTCCTTCAACCTGGAGACAGAGTTCTCCTGGTGTGTGCCCAGGGGTGGAGGAGAA 23084
human.FAM30B-LOC105372935-GGT2.NCBI.                                            ggcacttcctccaacctggagacagagttctcctggtgtgtgcccaggggtggaggagaa 19993
human.FAM230E-LOC105377182-GGT3P.NCBI.                                          ggcacttcctccaacctggagacagaattctcctggtgtgtgcccaggggtggaggagaa 18730
human.FAM230J-LOC105372942-GGTLC5P.NCBI.rev.compl                               ggcacttccttcaacctggagacagagttctcctggtgtgtgcccaggggtggaggagaa 19444
human.FAM230A.ncRNA.18487127.18500594-GGTLC3.18516335.18518165.rev.compl.REF    ggcacttccttcaacctggagacagagttctcctggtgtgtgcccaggggtggaggagaa 8208
                                                                                ********** *************** ************************* *******

chimp.LOC112206744-LOC107973052-GGT2.revcompl                                   attgacagtctgcctctgaactttcaggactttaaaaagcactcatgtttccatcctcgc 21366
chimp54546-135457.revcompl.Pan.troglodytes.clone.rp43-41g5.GenBank:AC099533.36  ATTGACAGTCTGCCTCTGAACTTTCAGGACTTTAAAAAGCACTCATGTTTCCATCCTCAC 23144
human.FAM30B-LOC105372935-GGT2.NCBI.                                            attgacagtctgcctctgaactttcaggactttaaaaagcactcatgtttccatcctcac 20053
human.FAM230E-LOC105377182-GGT3P.NCBI.                                          attgacagtctgcctctgaactttcaggactttaaaaagcactcatgtttccatcctcac 18790
human.FAM230J-LOC105372942-GGTLC5P.NCBI.rev.compl                               attgacagtctgcctctgaactttcaggactttaaaaagcactcatgtttccatcctcac 19504
human.FAM230A.ncRNA.18487127.18500594-GGTLC3.18516335.18518165.rev.compl.REF    attgacagtctgcctctgaactttcaggactttaaaaagcactcatgtttccatcctcac 8268
                                                                                ********************************************************** *

chimp.LOC112206744-LOC107973052-GGT2.revcompl                                   tgttgactcctggcttaaagggatctccaggggtgagtgaggaggagggatcggaccctg 21426
chimp54546-135457.revcompl.Pan.troglodytes.clone.rp43-41g5.GenBank:AC099533.36  TGTTGACTCCTGGCTTAAAGGGATCTCCCGGGGTGAGTGAGGAGGCGGGATCGGACCCTG 23204
human.FAM30B-LOC105372935-GGT2.NCBI.                                            tgttgactcctggcttaaagggatctcccggggtgagtgaggaggcgggatcggaccctg 20113
human.FAM230E-LOC105377182-GGT3P.NCBI.                                          tgttgactcctggcttaaagggatctcccggggtgagtgaggaggcgggatcggacccta 18850
human.FAM230J-LOC105372942-GGTLC5P.NCBI.rev.compl                               tgttgactcctggcttaaagggatctcccggggtgagtgaggaggcgggattggaccctg 19564
human.FAM230A.ncRNA.18487127.18500594-GGTLC3.18516335.18518165.rev.compl.REF    tgttgactcctggcttaaagggatctcccggggtgagtgaggaggcgggatcggaccctg 8328
                                                                                **************************** **************** ***** *******

chimp.LOC112206744-LOC107973052-GGT2.revcompl                                   gcagtctgacggcagcacctgtgttcctctgcactgggccgtggatgacattacacacct 21486
chimp54546-135457.revcompl.Pan.troglodytes.clone.rp43-41g5.GenBank:AC099533.36  GCAGTCTGACGGCAGCACCTGTGTTCCTCTGCACTGGGCCGTGGATGACATTACACACCT 23264
human.FAM30B-LOC105372935-GGT2.NCBI.                                            gcagtctgacggcagcacctgtgttcctctgcactgggccgtggatgacattacatacct 20173
human.FAM230E-LOC105377182-GGT3P.NCBI.                                          gcagtctgacggcagcacctgtgttcctctgcactgggccgtggatgacattacacacct 18910
human.FAM230J-LOC105372942-GGTLC5P.NCBI.rev.compl                               gcagtctgacggcagcacctgtgttcctctgcactgggccgtggatgacattacacacct 19624
human.FAM230A.ncRNA.18487127.18500594-GGTLC3.18516335.18518165.rev.compl.REF    gcagtctgacggcagcacctgtgttcctctgcactgggccgtggatgacattacacacct 8388
                                                                                ******************************************************* ****

chimp.LOC112206744-LOC107973052-GGT2.revcompl                                   tggtgagaatcaggaattgaggctaaccacatctgaaattgagatgggccttgagtcata 21546
chimp54546-135457.revcompl.Pan.troglodytes.clone.rp43-41g5.GenBank:AC099533.36  TGGTGAGAATCAGGAATTGAGGCTAACCACATCTGAAATTGAGATGGGCCTTGAGTCATA 23324
human.FAM30B-LOC105372935-GGT2.NCBI.                                            tggtgagaatcaggaattgaggctaaccacatctgaaattgagatgggccttgagtcata 20233
human.FAM230E-LOC105377182-GGT3P.NCBI.                                          tggtgagaatcaggaattgaggctaaccacatctgaaattgagatgggccttgagtcata 18970
human.FAM230J-LOC105372942-GGTLC5P.NCBI.rev.compl                               tggtgagaatcaggaattgaggctaaccacatctgaaattgagatgggccttgagtcata 19684
human.FAM230A.ncRNA.18487127.18500594-GGTLC3.18516335.18518165.rev.compl.REF    tggtgagaatcaggaattgaggctaaccacatctgaaattgagatgggccttgagtcata 8448
                                                                                ************************************************************

chimp.LOC112206744-LOC107973052-GGT2.revcompl                                   taaatagtttggaaaagatgcattttactacgctattgaaagaaaccatttatttctcac 21606
chimp54546-135457.revcompl.Pan.troglodytes.clone.rp43-41g5.GenBank:AC099533.36  TAAATAGTTTGGAAAAGATGCATTTTACTACGCTATTGAAAGAAACCATTTATTTCTCAC 23384
human.FAM30B-LOC105372935-GGT2.NCBI.                                            taaatagtttggaaaagatgcattttactacgctattgaaagaaaccatttatttctcac 20293
human.FAM230E-LOC105377182-GGT3P.NCBI.                                          taaatagtttggaaaagatgcattttactacgctattgaaagaaaccatttatttctcac 19030
human.FAM230J-LOC105372942-GGTLC5P.NCBI.rev.compl                               taaatagtttggaaaagatgcattttactacgctattgaaagaaaccatttatttctcac 19744
human.FAM230A.ncRNA.18487127.18500594-GGTLC3.18516335.18518165.rev.compl.REF    taaatagtttggaaaagatgcattttactacgctattgaaagaaaccatttatttctcac 8508
                                                                                ************************************************************

chimp.LOC112206744-LOC107973052-GGT2.revcompl                                   tccagcaggataaatggttttcagtatccatttaactgctcattgactcttactgtagat 21666
chimp54546-135457.revcompl.Pan.troglodytes.clone.rp43-41g5.GenBank:AC099533.36  TCCAGCAGGATAAATGGTTTTCAGTATCCATTTAACTGCTCATTGACTCTTACTGTGGAT 23444
human.FAM30B-LOC105372935-GGT2.NCBI.                                            tccagcaggataaatggttttcagtatccatttaactgctcattgactcttactgtagat 20353
human.FAM230E-LOC105377182-GGT3P.NCBI.                                          tccagcaggataaatggttttcagtatccatttaactgctcattgactcttactgtagat 19090
human.FAM230J-LOC105372942-GGTLC5P.NCBI.rev.compl                               tccagcaggataaatggttttcagtatccacttaactgctcattgactcttactgtagat 19804
human.FAM230A.ncRNA.18487127.18500594-GGTLC3.18516335.18518165.rev.compl.REF    tccagcaggataaatggttttcagtatccacttaactgctcattgactcttactgtagat 8568
                                                                                ****************************** ************************* ***

chimp.LOC112206744-LOC107973052-GGT2.revcompl                                   gaggaggtggccagcagcccctgccctcccccagtt-gtaggcccaaggtaaccagcaat 21725
chimp54546-135457.revcompl.Pan.troglodytes.clone.rp43-41g5.GenBank:AC099533.36  GAGGAGGTAGCCAGCAGCCCCTGCCCTCCCCCAGTT-GTAGGCCCAAGGTAACCAGCAAT 23503
human.FAM30B-LOC105372935-GGT2.NCBI.                                            gaggaggtggccagcagcccctgccctcccccagttggtaggcccaaggtaaccagcaat 20413
human.FAM230E-LOC105377182-GGT3P.NCBI.                                          gaggaggtggccagcagcccctgccctcccccagttggtaggcccaaggtaaccagcaat 19150
human.FAM230J-LOC105372942-GGTLC5P.NCBI.rev.compl                               gaggaggtggccagcagcccctgccctcccccagtt-gtaggcccaaggtaaccagcaat 19863
human.FAM230A.ncRNA.18487127.18500594-GGTLC3.18516335.18518165.rev.compl.REF    gaggaggtggccagcagcccctgccctcccccagtt-gtaggcccaaggtaaccagcaat 8627
                                                                                ******** *************************** ***********************

chimp.LOC112206744-LOC107973052-GGT2.revcompl                                   ggactggatataatggaagagtggtgcattcggaggtatctgtattaatgggacccacat 21785
chimp54546-135457.revcompl.Pan.troglodytes.clone.rp43-41g5.GenBank:AC099533.36  GGACTGGATATAATGGAAGGGTGGTGCATTCGGAGGTATCTGTATTAATGGGACCCACAT 23563
human.FAM30B-LOC105372935-GGT2.NCBI.                                            tgactggatataatggaagagtggtgcattcgggggtatctgtattaatgggacccacat 20473
human.FAM230E-LOC105377182-GGT3P.NCBI.                                          tgactggatataatggaagagtggtgcattcggaggtatctgtattaatgggacccacat 19210
human.FAM230J-LOC105372942-GGTLC5P.NCBI.rev.compl                               tgactggatataatggaagagtggtgcattcggaggtatctgtattaatgggacccacat 19923
human.FAM230A.ncRNA.18487127.18500594-GGTLC3.18516335.18518165.rev.compl.REF    tgactggatataatggaagagtggtgcattcggaggtatctgtattaatgggacccacat 8687
```

```
                                                                                   ****************** ************* **************************

chimp.LOC112206744-LOC107973052-GGT2.revcompl                                     gatatggatgagagctattagggtgagaaaaagcctgggagcacaatgaaatatttaaat 21845
chimp54546-135457.revcompl.Pan.troglodytes.clone.rp43-41g5.GenBank:AC099533.36    GATATGGATGAGAGCTATTAGGGTGAGAAAAAGCCTGGGAGCACAATGAAATATTTAAAT 23623
human.FAM30B-LOC105372935-GGT2.NCBI.                                              gatatggatgagagctattagggtgagaaaaagcctgggagcacaatgaaatatttaaat 20533
human.FAM230E-LOC105377182-GGT3P.NCBI.                                            gatatggatgagagctattagggtgagaaaaagcctgggagcacaatgaaatatttaaat 19270
human.FAM230J-LOC105372942-GGTLC5P.NCBI.rev.compl                                 gatatggatgagagctattagggtgagaaaaagcctgggagcacaatgaaatatttaaat 19983
human.FAM230A.ncRNA.18487127.18500594-GGTLC3.18516335.18518165.rev.compl.REF      gatatggatgagagctattagggtgagaaaaagcctgggagcacaatgaaatatttaaat 8747
                                                                                  ************************************************************

chimp.LOC112206744-LOC107973052-GGT2.revcompl                                     attaaacaaaacattgttgaaatctccattgtactttagtagttgaagtcattcttgtgg 21905
chimp54546-135457.revcompl.Pan.troglodytes.clone.rp43-41g5.GenBank:AC099533.36    ATTAAACAAAACATTGTTGAAATCTCCATTGTACTTTAGTAGTTGAAGTCATTCTTGTGG 23683
human.FAM30B-LOC105372935-GGT2.NCBI.                                              attaaacaaaacattgttgaaatctccattgtactttagtagttgaagtcattcttgtgg 20593
human.FAM230E-LOC105377182-GGT3P.NCBI.                                            attaaacaaaacattgttgaaatctccattgtactttagtagttgaagtcattcttgtgg 19330
human.FAM230J-LOC105372942-GGTLC5P.NCBI.rev.compl                                 attaaacaaaacattgttgaaatctccattgtactttagtagttgaagtcattcttgtgg 20043
human.FAM230A.ncRNA.18487127.18500594-GGTLC3.18516335.18518165.rev.compl.REF      attaaacaaaacattgttgaaatctccattgtactttagtagttgaagtcattcttgtgg 8807
                                                                                  ************************************************************

chimp.LOC112206744-LOC107973052-GGT2.revcompl                                     tcatcactgcctttcccaagcataacaagctacttaatatcacatagacccgtgccatga 21965
chimp54546-135457.revcompl.Pan.troglodytes.clone.rp43-41g5.GenBank:AC099533.36    TCATCACTGCCTTTCCCAAGCATAACAAGCTACTTAATATCACATAGACCCATGCCATGA 23743
human.FAM30B-LOC105372935-GGT2.NCBI.                                              tcatcactgcctttcccaagcataacaagctacttaatatcacatggacccgtgccatga 20653
human.FAM230E-LOC105377182-GGT3P.NCBI.                                            tcatcactgcctttcccaagcataacaagctacttaatatcacatggacccgtgccatga 19390
human.FAM230J-LOC105372942-GGTLC5P.NCBI.rev.compl                                 tcatcactgcctttcccaagcataacaagctacttaatatcacatggacccgtgccatga 20103
human.FAM230A.ncRNA.18487127.18500594-GGTLC3.18516335.18518165.rev.compl.REF      tcatcactgcctttcccaagcataacaagctacttaatatcacatggacccgtgccatga 8867
                                                                                  ********************************************* ***** ********

chimp.LOC112206744-LOC107973052-GGT2.revcompl                                     ggaatgatgatcagtttgtaaaatgccaataaaacaattgcctatagaagccacaatgtt 22025
chimp54546-135457.revcompl.Pan.troglodytes.clone.rp43-41g5.GenBank:AC099533.36    GGAATGATGATCAGTTTGTAAAATGCCAATAAAACAATTGCCTATATAAGCCACAATGTT 23803
human.FAM30B-LOC105372935-GGT2.NCBI.                                              ggaatgatgatcagtttgtaaaatgccaataaaacaattgcctatataagccacaatgtt 20713
human.FAM230E-LOC105377182-GGT3P.NCBI.                                            ggaatgatgatcagtttgtaaaatgccaataaaacaattgcctatataagccacaatgtt 19450
human.FAM230J-LOC105372942-GGTLC5P.NCBI.rev.compl                                 ggaatgatgatcagtttgtaaaatgccaataaaacaattgcctatataagccacaatgtt 20163
human.FAM230A.ncRNA.18487127.18500594-GGTLC3.18516335.18518165.rev.compl.REF      ggaatgatgatcagtttgtcaaatgccaataaaacaattgcctctataagccacaatgtt 8927
                                                                                  ******************* *********************** ** *************

chimp.LOC112206744-LOC107973052-GGT2.revcompl                                     tcatccatatatttcaatttccatgtgtaagtatagttcaaatttcagaaatttattatt 22085
chimp54546-135457.revcompl.Pan.troglodytes.clone.rp43-41g5.GenBank:AC099533.36    TCATCCATATATTTCAATTTCCATGTGTAAGTATAGTTCAAATTTCAGAAAGTTATTATT 23863
human.FAM30B-LOC105372935-GGT2.NCBI.                                              tcatccatatatttcaatttccatgtgtaagtatagttcaaatttcagaaatttattatt 20773
human.FAM230E-LOC105377182-GGT3P.NCBI.                                            tcatccatatatttcaatttccatgtataagtatagttcaaatttcagaaatttattatt 19510
human.FAM230J-LOC105372942-GGTLC5P.NCBI.rev.compl                                 tcatccatatatttcaatttccatgtgtaagtatagttcaaatttcagaaatttattatt 20223
human.FAM230A.ncRNA.18487127.18500594-GGTLC3.18516335.18518165.rev.compl.REF      tcatccatatatttcaatttccatgtgtaagtatagttcaaatttcagaaatttattatt 8987
                                                                                  ************************** ************************ ********

chimp.LOC112206744-LOC107973052-GGT2.revcompl                                     atctaatagaatatgcatggtatatcaatgagcaattatcatactgtttctattaacaat 22145
chimp54546-135457.revcompl.Pan.troglodytes.clone.rp43-41g5.GenBank:AC099533.36    ATCTAATAGAATATGCATGGTATATCAATGAGAAATTATCATACTGTTTCTATTAACAAT 23923
human.FAM30B-LOC105372935-GGT2.NCBI.                                              atctaatagaatatgcatggtatatcaatgagcaattatcatactgtttctattaacaat 20833
human.FAM230E-LOC105377182-GGT3P.NCBI.                                            atctaatagaatatgcatggtatatcaatgagcaattatcatactgtttctattaacaat 19570
human.FAM230J-LOC105372942-GGTLC5P.NCBI.rev.compl                                 atctaatagaatatgcatggtatatcaatgagcaattatcatactgtttctattaacaat 20283
human.FAM230A.ncRNA.18487127.18500594-GGTLC3.18516335.18518165.rev.compl.REF      atctaatagaatatgcatggtatatcaatgagcaattatcatactgtttctattaacaat 9047
                                                                                  ******************************** ***************************

chimp.LOC112206744-LOC107973052-GGT2.revcompl                                     tatttgtatgatgaaaaaagcagactcccattcttggatttttctcagtttgcacacatt 22205
chimp54546-135457.revcompl.Pan.troglodytes.clone.rp43-41g5.GenBank:AC099533.36    TATTTGTATGATGAAAAAAGCAGACTCCCATTCTTGGATTTTTCTCAGTTTGCACACATT 23983
human.FAM30B-LOC105372935-GGT2.NCBI.                                              tatttgtatgatgaaaaaagcagactcccattcttggatttttctcagtttgcacacatt 20893
human.FAM230E-LOC105377182-GGT3P.NCBI.                                            tatttgtatgatgaaaaaagcagactcccattcttggatttttctcagtttgcacacatt 19630
human.FAM230J-LOC105372942-GGTLC5P.NCBI.rev.compl                                 tatttgtatgatgaaaaaagcagactcccattcttggatttttctcagtttgcacacatt 20343
human.FAM230A.ncRNA.18487127.18500594-GGTLC3.18516335.18518165.rev.compl.REF      tatttgtatgatgaaaaaagcagactcccattcttggatttttctcagtttgcacacatt 9107
                                                                                  ************************************************************

chimp.LOC112206744-LOC107973052-GGT2.revcompl                                     agcatgacagccccatttccacctgacatgtgccagcaagaggccaggaacagaggcttt 22265
chimp54546-135457.revcompl.Pan.troglodytes.clone.rp43-41g5.GenBank:AC099533.36    AGCATGACAGCCCCATTTCCACCTGACATGTGCCAGCAAGAGGCCAGGAACAGAGGCTTT 24043
human.FAM30B-LOC105372935-GGT2.NCBI.                                              agcatgacagccccatttccacctgacatgtgccagcaagaggccaggaacagaggcttt 20953
human.FAM230E-LOC105377182-GGT3P.NCBI.                                            agcatgacagccccatttccacctgacatgtgccagcaagaggccaggaacagaggcttt 19690
human.FAM230J-LOC105372942-GGTLC5P.NCBI.rev.compl                                 agcatgacagccccatttccacctgacatgtgccagcaagaggccaggaacagaggcttt 20403
human.FAM230A.ncRNA.18487127.18500594-GGTLC3.18516335.18518165.rev.compl.REF      agcatgacagccccatttccacctgacatgtgccagcaagaggccaggaacagaggcttt 9167
                                                                                  ************************************************************

chimp.LOC112206744-LOC107973052-GGT2.revcompl                                     tcttat----taagatttctaaatgtattatgtattcacatttagaaactctaaatgtca 22321
chimp54546-135457.revcompl.Pan.troglodytes.clone.rp43-41g5.GenBank:AC099533.36    TCTTATTAACTAAGATTTCTAAATGTATTATGTATTCACATTTAGAAACTCTAAATATCA 24103
human.FAM30B-LOC105372935-GGT2.NCBI.                                              tcttattaactaagatttctaaatgtattacgtattcacatttagaaactctaaatatca 21013
human.FAM230E-LOC105377182-GGT3P.NCBI.                                            tcttattaactaagatttctaaatgtattacgtattcacatttagaaactctaaatatca 19750
human.FAM230J-LOC105372942-GGTLC5P.NCBI.rev.compl                                 tcttattaactaagatttctaaatgtattacgtattcacatttagaaactctaaatatca 20463
human.FAM230A.ncRNA.18487127.18500594-GGTLC3.18516335.18518165.rev.compl.REF      tcttattaactaagatttctaaatgtattacgtattcacatttagaaactctaaatatca 9227
                                                                                  ******    ******************** ************************* ***

chimp.LOC112206744-LOC107973052-GGT2.revcompl                                     taaaaggttagcaaggaagtttcccttccactctgaacttccaaacaccaagtcaacatt 22381
chimp54546-135457.revcompl.Pan.troglodytes.clone.rp43-41g5.GenBank:AC099533.36    TAAAAGGTTAGCAAGGAAGTTTCCCTTCCACTCTGAACTTCCAAACACCAAGTCAACATT 24163
human.FAM30B-LOC105372935-GGT2.NCBI.                                              taaaaggttagcaaggaagtttcccttccactctgaacttccaaacaccaagtcaacatt 21073
human.FAM230E-LOC105377182-GGT3P.NCBI.                                            taaaaggttagcaaggaagtttcccttccactctgaacttccaaacaccaagtcaacatt 19810
human.FAM230J-LOC105372942-GGTLC5P.NCBI.rev.compl                                 taaaaggttagcaaggaagtttcccttccactctgaacttccaaacaccaagtcaacatt 20523
human.FAM230A.ncRNA.18487127.18500594-GGTLC3.18516335.18518165.rev.compl.REF      taaaaggttagcaaggaagtttcccttccactctgaacttccaaacaccaagtcaacgtt 9287
                                                                                  ********************************************************* **

chimp.LOC112206744-LOC107973052-GGT2.revcompl                                     tttgtttgcatatcatccctgcaatctatgtgcaaatagaagcatgcacctggaatgcag 22441
chimp54546-135457.revcompl.Pan.troglodytes.clone.rp43-41g5.GenBank:AC099533.36    TTTGTTTACATATCATCCCTGCAATCTATGTGCAAATAGAAGCATGCACCTGGAATGCAG 24223
human.FAM30B-LOC105372935-GGT2.NCBI.                                              tttgtttgcatatcatccctgcaatctatgtgcaaatagaagcatgcacctggaatgcag 21133
human.FAM230E-LOC105377182-GGT3P.NCBI.                                            tttgtttgcatatcatccctgcaatctatgtgcaaatagaagcatgcacctggaatgcag 19870
human.FAM230J-LOC105372942-GGTLC5P.NCBI.rev.compl                                 tttgtttgcatatcatccctgcaatctatgtgcaaatagaagcatgcacctggaatgcag 20583
human.FAM230A.ncRNA.18487127.18500594-GGTLC3.18516335.18518165.rev.compl.REF      tttgtttgcatatcatccctgcaatctatgtgcaaatagaagcatgcacctggaatgcag 9347
                                                                                  ******* ****************************************************

chimp.LOC112206744-LOC107973052-GGT2.revcompl                                     gctgatgtgtgatcgtgtttacacaaagtcctctgcacctctgcatatatcactgggcaa 22501
chimp54546-135457.revcompl.Pan.troglodytes.clone.rp43-41g5.GenBank:AC099533.36    GCTGATGTGTGATCGTGTTTACACAAAGTCCTCTGCACCTCTGCATGTATCACTGGGCAA 24283
human.FAM30B-LOC105372935-GGT2.NCBI.                                              gctgatgtgtgatcatgtttacacaaagtcctctgcacctctgcatatatcactgggcaa 21193
human.FAM230E-LOC105377182-GGT3P.NCBI.                                            gctgatgtgtgatcgtgtttacacaaagtcctctgcacctctgcatatatcactgggcaa 19930
human.FAM230J-LOC105372942-GGTLC5P.NCBI.rev.compl                                 gctgatgtgtgatcgtgtttacacaaagtcctctgcacctctgcatatatcactgggcaa 20643
human.FAM230A.ncRNA.18487127.18500594-GGTLC3.18516335.18518165.rev.compl.REF      gctgatgtgtgatcgtgtttacacaaagtcctctgcacctctgcatatatcactgggcaa 9407
                                                                                  ************** ****************************** **************

chimp.LOC112206744-LOC107973052-GGT2.revcompl                                     tgcaccttagttatcattccacatttcaaatgtaaatccattgtattgtttcagagctat 22561
chimp54546-135457.revcompl.Pan.troglodytes.clone.rp43-41g5.GenBank:AC099533.36    TGCACCTTAGTTATCATTCCACATTTCAAATGTAAATCCATTGTATTGTTTCAGAGCTAT 24343
human.FAM30B-LOC105372935-GGT2.NCBI.                                              tgcaccttagttatcattccacatttcaaatgtaaatccattgtattgtttcagagctat 21253
human.FAM230E-LOC105377182-GGT3P.NCBI.                                            tgcaccttagttatcattccacatttcaaatgtaaatccattgtattgtttcagagctat 19990
human.FAM230J-LOC105372942-GGTLC5P.NCBI.rev.compl                                 tgcaccttagttatcattccacatttcaaatgtaaatccattgtattgtttcagagctat 20703
human.FAM230A.ncRNA.18487127.18500594-GGTLC3.18516335.18518165.rev.compl.REF      tgcaccttagttatcattccacatttcaaatgtaaatccattgtattgtttcagagctat 9467
                                                                                  ************************************************************

chimp.LOC112206744-LOC107973052-GGT2.revcompl                                     gaagtactgcaccccatgactattcccaaaattacttaagcaccccgctatgggtatcca 22621
chimp54546-135457.revcompl.Pan.troglodytes.clone.rp43-41g5.GenBank:AC099533.36    AAAGTACTGCACCCCATGACTATTCCCAAAATTACTTAAGCACCCCGCTATGGGTATCCA 24403
human.FAM30B-LOC105372935-GGT2.NCBI.                                              aaagtactgcaccccatgactattcccaaaattacttaagcaccccgctatgggtatcca 21313
human.FAM230E-LOC105377182-GGT3P.NCBI.                                            aaagtactgcaccccatgactattcccaaaattacttaagcaccccgctatgggtatcca 20050
human.FAM230J-LOC105372942-GGTLC5P.NCBI.rev.compl                                 aaagtactgcaccccatgactattcccaaaattacttaagcaccccgctatgggtatcca 20763
human.FAM230A.ncRNA.18487127.18500594-GGTLC3.18516335.18518165.rev.compl.REF      aaagtactgcaccccatgactattcccaaaattacttaagcaccccgctatgggtatccg 9527
                                                                                   ********************************************************** 

chimp.LOC112206744-LOC107973052-GGT2.revcompl                                     tttgttctgtttccagtcttgctcttataaccaatgctgtagtgaacagcactgtgttga 22681
chimp54546-135457.revcompl.Pan.troglodytes.clone.rp43-41g5.GenBank:AC099533.36    TTTGTTCTGTTTCCAGTCTTGCTCTTATAACCAATGCTGTAGTGAACAGCACTGTGTTGA 24463
human.FAM30B-LOC105372935-GGT2.NCBI.                                              tttgttctgtttccagtcttgctcttataaccaatgctgtagtgaacagcactgtgttga 21373
human.FAM230E-LOC105377182-GGT3P.NCBI.                                            tttgttctgtttccagtcttgctcttataaccaatgctgtagtgaacagcactgtgttga 20110
human.FAM230J-LOC105372942-GGTLC5P.NCBI.rev.compl                                 tttgttctgtttccagtcttgctcttataaccaatgctgtagtgaacagcactgtgttga 20823
human.FAM230A.ncRNA.18487127.18500594-GGTLC3.18516335.18518165.rev.compl.REF      tttgttctgtttccagtcttgctcttataaccaatgctgtagtgaacagcactgtgttga 9587
                                                                                  ************************************************************

chimp.LOC112206744-LOC107973052-GGT2.revcompl                                     gaagtggtgaacgtgggcatctttgtcttgttcccgtcctcaggggggaatgctttcaact 22741
chimp54546-135457.revcompl.Pan.troglodytes.clone.rp43-41g5.GenBank:AC099533.36    GAAGTGGTGAACGTGGGCATCTTTGTCTTGTTCCCTTCCTCAGGGGGAATGCTTTCAACT 24523
human.FAM30B-LOC105372935-GGT2.NCBI.                                              gaagcggtgaacgtgggcatctttgtcttgttcccttcctcaggcggaatgctttcaact 21433
human.FAM230E-LOC105377182-GGT3P.NCBI.                                            gaagtggtgaacgtgggcatctttgtcttgttcccttcctcagggggaatgctttcaact 20170
human.FAM230J-LOC105372942-GGTLC5P.NCBI.rev.compl                                 gaagcggtgaacgtgggcatctttgtcttgttcccttcctcagggggaatgctttcaact 20883
human.FAM230A.ncRNA.18487127.18500594-GGTLC3.18516335.18518165.rev.compl.REF      gaagcggtgaacgtgggcatctttgtcttgttcccttcctcagggggaatgctttcaact 9647
                                                                                  **** ****************************** ******** ***************

chimp.LOC112206744-LOC107973052-GGT2.revcompl                                     ttcccccattcaggaaactgttggctgtgggtttgtcatagatagcttttattaccttaa 22801
chimp54546-135457.revcompl.Pan.troglodytes.clone.rp43-41g5.GenBank:AC099533.36    CTCCCCCATTCAGGAAAATGTTGGCTGTGGGTTTGTCATAGATAGCTTTTATTACCTTAA 24583
human.FAM30B-LOC105372935-GGT2.NCBI.                                              ctcccccattcaggaaaatgttggctgtgggtttgtcatagatagcttttattaccttaa 21493
human.FAM230E-LOC105377182-GGT3P.NCBI.                                            ctcccccattcaggaaaatgttggctgtgggtttgtcatagatagcttttattaccttaa 20230
human.FAM230J-LOC105372942-GGTLC5P.NCBI.rev.compl                                 ctcccccattcaggaaaatgttggctgtgggtttgtcatagatagcttttattaccttaa 20943
human.FAM230A.ncRNA.18487127.18500594-GGTLC3.18516335.18518165.rev.compl.REF      ctcccccattcaggaaaatgttggctgtgggtttgtcatagatagcttttattaccttaa 9707
                                                                                   *************** *******************************************

chimp.LOC112206744-LOC107973052-GGT2.revcompl                                     ggtatgtccattctatgctgattttgatgaacggttttaatcataaagaaatgctggatt 22861
chimp54546-135457.revcompl.Pan.troglodytes.clone.rp43-41g5.GenBank:AC099533.36    GGTATGTCCGTTCTATGCTGATTTTGATGAACGGTTTTAATCGTAAAGAAATGCTGGATT 24643
human.FAM30B-LOC105372935-GGT2.NCBI.                                              ggtatgtccgttctacgctgattttgatgaacggttttaatcataaagaaatgctggatt 21553
human.FAM230E-LOC105377182-GGT3P.NCBI.                                            ggtatgtccgttctatgctgattttgacgaacggttttaatcataaagaaatgctggatt 20290
human.FAM230J-LOC105372942-GGTLC5P.NCBI.rev.compl                                 ggtatgtccgttctatgctgattgtgacgaacggttttaatcataaagaaatgctggatt 21003
human.FAM230A.ncRNA.18487127.18500594-GGTLC3.18516335.18518165.rev.compl.REF      ggtatgtccgttctatgctgattttgacgaacggttttaatcataaagaaatgctggatt 9767
                                                                                  ********* ***** ******* *** ************** *****************
```

```
chimp.LOC112206744-LOC107973052-GGT2.revcompl                                      ttgtcaaaggctttttctgcatctattcagattatcatgtgattttgtttttagtttta 22921
chimp54546-135457.revcompl.Pan.troglodytes.clone.rp43-41g5.GenBank:AC099533.36     TTGTCAAAGGCTTTTTCTGCATCTATTCAGATTATCATGTGATTTTGTTTTTAGTTTTA 24703
human.FAM30B-LOC105372935-GGT2.NCBI.                                               ttgtcaaaggctttttctgcatctattcagattatcatgtgattttgtttttagtttta 21613
human.FAM230E-LOC105377182-GGT3P.NCBI.                                             ttgtcaaaggctttttctgcatctattcagattatcatgtgattttgtttttagtttta 20350
human.FAM230J-LOC105372942-GGTLC5P.NCBI.rev.compl                                  ttgtcaaaggctttttctgcatctattcagattatcatgtgattttgtttttagtttta 21063
human.FAM230A.ncRNA.18487127.18500594-GGTLC3.18516335.18518165.rev.compl.REF       ttgtcaaaggctttttctgcatctattcagattatcatgtgattttgtttttagtttta 9827
                                                                                   ************************************************************

chimp.LOC112206744-LOC107973052-GGT2.revcompl                                      tttatgtgatgtatcacatttattgacttgcgtatgttaaaccatccctgcatccctagt 22981
chimp54546-135457.revcompl.Pan.troglodytes.clone.rp43-41g5.GenBank:AC099533.36     TTTATGTGATGTATCACATTTATTGACTTGCGTATGTTAAACCATCCCTGCATCCCTAGT 24763
human.FAM30B-LOC105372935-GGT2.NCBI.                                               ttgatgtgatgtatcacatttattgacttgcgtatgttaaaccatccctgcatccctagt 21673
human.FAM230E-LOC105377182-GGT3P.NCBI.                                             ttgatgtgatgtatcacatttattgacttgcgtatgttaaaccatccctgcatccctagt 20410
human.FAM230J-LOC105372942-GGTLC5P.NCBI.rev.compl                                  ttgatgtgatgtatcacatttattgacttgcgtatgttaaaccatccctgcatccctagt 21123
human.FAM230A.ncRNA.18487127.18500594-GGTLC3.18516335.18518165.rev.compl.REF       ttgatgtgatgtatcacatttattgacttgcgtatgttaaaccatccctgcatccctagt 9887
                                                                                   ** *********************************************************

chimp.LOC112206744-LOC107973052-GGT2.revcompl                                      atgaaacccacttgaatcatggtggattatctttttgatatgctgttggattcagttagc 23041
chimp54546-135457.revcompl.Pan.troglodytes.clone.rp43-41g5.GenBank:AC099533.36     ATGAAACCCACTTGAATCATGGTGGATTATCTTTTTGATATGCTGTTGGATTCAGTTAGC 24823
human.FAM30B-LOC105372935-GGT2.NCBI.                                               atgaaacccacttgaatcatggtggattatctttttgatatgctgttggattcagttagc 21733
human.FAM230E-LOC105377182-GGT3P.NCBI.                                             atgaaacccacttgaatcatggtggattatctttttgatatgctgttggattcagttagc 20470
human.FAM230J-LOC105372942-GGTLC5P.NCBI.rev.compl                                  atgaaacccacttgaatcatggtggattatctttttgatatgctgttggattcagttagc 21183
human.FAM230A.ncRNA.18487127.18500594-GGTLC3.18516335.18518165.rev.compl.REF       atgaaacccacttgaatcatggtggattatctttttgatatgctgttggattcagttagc 9947
                                                                                   ************************************************************

chimp.LOC112206744-LOC107973052-GGT2.revcompl                                      ttggttgtagcatttcttttattccatctgtggaatgtattggtttaaataatgaaaac 23101
chimp54546-135457.revcompl.Pan.troglodytes.clone.rp43-41g5.GenBank:AC099533.36     TTGGTTGTAGCATTTCTTATTATTCCATCTGTGGAATGTATTGGTTTAAATAATGAAAAC 24883
human.FAM30B-LOC105372935-GGT2.NCBI.                                               ttggttgtagcatttcttattattccatctgtggaatgtattggttgaaataatgaaaac 21793
human.FAM230E-LOC105377182-GGT3P.NCBI.                                             ttggttgtagcatttcttattattccatctgtggaatgtattggttgaaataatgaaaac 20530
human.FAM230J-LOC105372942-GGTLC5P.NCBI.rev.compl                                  ttggttgtagcatttcttattattccatctgtggaatgtattggttgaaataatgaaaac 21243
human.FAM230A.ncRNA.18487127.18500594-GGTLC3.18516335.18518165.rev.compl.REF       ttggttgtagcatttcttattattccatctgtggaatgtattggttgaaataatgaaaac 10007
                                                                                   ****************** *************************** *************

chimp.LOC112206744-LOC107973052-GGT2.revcompl                                      atgttctatcctcactgcttagcactttgtgtttctttaatagccttcccaacagggcaa 23161
chimp54546-135457.revcompl.Pan.troglodytes.clone.rp43-41g5.GenBank:AC099533.36     ATGTTCTATCCTCACTGCTTAGCACTTTGTGTTTCTTTAATAGCCTTCCCAACAGGGCAA 24943
human.FAM30B-LOC105372935-GGT2.NCBI.                                               atgttctatcctcactgcttagcactttgtgtttctttaatagccttcccaacagggcaa 21853
human.FAM230E-LOC105377182-GGT3P.NCBI.                                             atgttctatcctcactgcttagcactttgtgtttctttaatagccttcccaacagggcaa 20590
human.FAM230J-LOC105372942-GGTLC5P.NCBI.rev.compl                                  atgttctatcctcactgcttagcactttgtgtttctttaatagccttcccaacagggcaa 21303
human.FAM230A.ncRNA.18487127.18500594-GGTLC3.18516335.18518165.rev.compl.REF       atgttctatcctcactgcttagcactttgtgtttctttaatagccttcccaacagggcaa 10067
                                                                                   ************************************************************

chimp.LOC112206744-LOC107973052-GGT2.revcompl                                      cataaaagcaggagccctgctagtcaccccttaacccggaatcccccttctccacagct 23221
chimp54546-135457.revcompl.Pan.troglodytes.clone.rp43-41g5.GenBank:AC099533.36     CATAAAAGCAGGAGCCCTGCTAGTCACCCCTTAACCCGGAATCCCCCTTCTCCACAGCT 25003
human.FAM30B-LOC105372935-GGT2.NCBI.                                               cataaaagcaggagccctgctagtctccccttaacccggaatcccccttctccacagct 21913
human.FAM230E-LOC105377182-GGT3P.NCBI.                                             cataaaagcaggagccctgctagtctccccttaacccggaatcccccttctccacagct 20650
human.FAM230J-LOC105372942-GGTLC5P.NCBI.rev.compl                                  cataaaagcaggagccctgctagtctccccttaacccggaatcccccttctccacagct 21363
human.FAM230A.ncRNA.18487127.18500594-GGTLC3.18516335.18518165.rev.compl.REF       cataaaagcaggagccctgctagtctccccttaacccggaatcccccttctccacagct 10127
                                                                                   ************************* **********************************

chimp.LOC112206744-LOC107973052-GGT2.revcompl                                      cgcttattggacaggatagactgggagcccaggcctcaaggtaaggacgtgctctgtcac 23281
chimp54546-135457.revcompl.Pan.troglodytes.clone.rp43-41g5.GenBank:AC099533.36     CGCTCATTGGACAGGATAGACTGGGCGCCCAGGCTTCAAGGTAAGGACGTGCTCTGTCAC 25063
human.FAM30B-LOC105372935-GGT2.NCBI.                                               cgctcattggacaggatagactgggcgcccaggcttcaaggtaaggacgtgctctgtcac 21973
human.FAM230E-LOC105377182-GGT3P.NCBI.                                             cgctcattggacaggatagactgggcgcccaggcttcaaggtaaggacatgctctgtcac 20710
human.FAM230J-LOC105372942-GGTLC5P.NCBI.rev.compl                                  cgctcattggacaggatagactgggcgcccaggcttcaaggtaaggacgtgctctgtcac 21423
human.FAM230A.ncRNA.18487127.18500594-GGTLC3.18516335.18518165.rev.compl.REF       cgctcattggacaggatagactgggcgcccaggcttcaaggtaaggacgtgctctgtcac 10187
                                                                                   **** ******************** ******** ************* ***********

chimp.LOC112206744-LOC107973052-GGT2.revcompl                                      ctagaggtgcagtgcttgggaaggccaaccttggagggttgcctgccagctttacagtga 23341
chimp54546-135457.revcompl.Pan.troglodytes.clone.rp43-41g5.GenBank:AC099533.36     CTAGAGGTGCAGTGCTTGGGAAGGCCAACCTTGGAGGGTTGCCTGCCAGCTTTACAGTGA 25123
human.FAM30B-LOC105372935-GGT2.NCBI.                                               ctagaggtgcagtgcttgggaaggccaaccttggagggttgcctgccagctttacagtga 22033
human.FAM230E-LOC105377182-GGT3P.NCBI.                                             ctagaggtgcagtgcttgggaaggccaaccttggagggttgcctgccagctttacagtga 20770
human.FAM230J-LOC105372942-GGTLC5P.NCBI.rev.compl                                  ctagaggtgcagtgcttgggaaggccaaccttggagggttgcctgccagctttacagtga 21483
human.FAM230A.ncRNA.18487127.18500594-GGTLC3.18516335.18518165.rev.compl.REF       ctagaggtgcagtgcttgggaaggccaaccttggagggttgcctgccagctttacagtga 10247
                                                                                   ************************************************************

chimp.LOC112206744-LOC107973052-GGT2.revcompl                                      cagaggtgttgggagggactgaccaccagtgcataaggctgtgctttgttggtgacataa 23401
chimp54546-135457.revcompl.Pan.troglodytes.clone.rp43-41g5.GenBank:AC099533.36     CAGAGGTGTTGGGAGGGACTGACCACCAGTGCATAAGGCTGTGCTTTGTTGGTGACATAA 25183
human.FAM30B-LOC105372935-GGT2.NCBI.                                               cagaggtgttgagagggactgaccaccagtgcataaggctgtgctttgttggtgacataa 22093
human.FAM230E-LOC105377182-GGT3P.NCBI.                                             cagaggtgttgagagggactgaccaccagtgcagaaggctgtgctttgttggtgacataa 20830
human.FAM230J-LOC105372942-GGTLC5P.NCBI.rev.compl                                  cagaggtgttgagagggactgaccaccagtgcagaaggctgtgctttgttggtgacataa 21543
human.FAM230A.ncRNA.18487127.18500594-GGTLC3.18516335.18518165.rev.compl.REF       cagaggtgttgagagggactgaccaccagtgcataaggctgtgctttgttggtgacataa 10307
                                                                                   *********** ******************** ***************************

chimp.LOC112206744-LOC107973052-GGT2.revcompl                                      aggattgtttcacaggttgttggggagggacaatcccaaggcctcccctggccctggtgc 23461
chimp54546-135457.revcompl.Pan.troglodytes.clone.rp43-41g5.GenBank:AC099533.36     AGGATTGTTTCACAGGTTGTTGGGGAGGGACAATCCCAAGGCCTCCCCTGGCCCTGGTGC 25243
human.FAM30B-LOC105372935-GGT2.NCBI.                                               aggattgtttcacagattgttggggagggacaatcccaaggcctcccctggccctggtgc 22153
human.FAM230E-LOC105377182-GGT3P.NCBI.                                             aggattgtttcacagattgttggggagggacaatcccaaggcctcccctggccctggtgc 20890
human.FAM230J-LOC105372942-GGTLC5P.NCBI.rev.compl                                  aggattgtttcacagattgttggggagggacaatcccaaggcctcccctggccctggtgc 21603
human.FAM230A.ncRNA.18487127.18500594-GGTLC3.18516335.18518165.rev.compl.REF       aggattgtttcacagattgttggggagggacaatcccaaggcctcccctggccctggtgc 10367
                                                                                   *************** ********************************************

chimp.LOC112206744-LOC107973052-GGT2.revcompl                                      tggctctgcacaaaggcaataagagagggatgctggtaagggctgacctgttgctgtgct 23521
chimp54546-135457.revcompl.Pan.troglodytes.clone.rp43-41g5.GenBank:AC099533.36     TGGCTCTGCACAAAGGCAATAAGAGAGGGATGCTGGTAAGGGCTGACCTGTTGCTGTGCT 25303
human.FAM30B-LOC105372935-GGT2.NCBI.                                               tggctctgcacaaaggcaataagagagggatgctggtaagggctgacctgttgctgtgct 22213
human.FAM230E-LOC105377182-GGT3P.NCBI.                                             tggctctgcacaaaggcaataagagagggatgctggtaagggctgacctgttgctgtgct 20950
human.FAM230J-LOC105372942-GGTLC5P.NCBI.rev.compl                                  tggctctgcacaaaggcaataagagagggatgctggtaagggctgacctgttgctgtgct 21663
human.FAM230A.ncRNA.18487127.18500594-GGTLC3.18516335.18518165.rev.compl.REF       tggctctgcacaaaggcaataagagagggatgctggtaagggctgacctgttgctgtgct 10427
                                                                                   ************************************************************

chimp.LOC112206744-LOC107973052-GGT2.revcompl                                      ggggaggaaggtgctgggctgaaattcaggaggctgaggatgcagcagtcccataggagg 23581
chimp54546-135457.revcompl.Pan.troglodytes.clone.rp43-41g5.GenBank:AC099533.36     GGGGAGGAAGGTGCTGGGCTGAAATTCAGGAGGCTGAGGATGCAGCAATCCCATAGGAGG 25363
human.FAM30B-LOC105372935-GGT2.NCBI.                                               ggggaggaaggtgctgggctgaaattcaggaggctgaggatgcagcagtcccataggagg 22273
human.FAM230E-LOC105377182-GGT3P.NCBI.                                             ggggaggaaggtgctgggctgaaattcaggaggctgaggatgcagcagtcccataggagg 21010
human.FAM230J-LOC105372942-GGTLC5P.NCBI.rev.compl                                  ggggaggaaggtgctgggctgaaattcaggaggctgaggatgcagcagtcccataggagg 21723
human.FAM230A.ncRNA.18487127.18500594-GGTLC3.18516335.18518165.rev.compl.REF       ggggaggaaggtgctgggctgaaattcaggaggctgaggatgcagcagtcccataggagg 10487
                                                                                   ********************************************** *************

chimp.LOC112206744-LOC107973052-GGT2.revcompl                                      tacgtgaccttcaggatacattttcttcattgatgatcaatggaaatgagaaatcactga 23641
chimp54546-135457.revcompl.Pan.troglodytes.clone.rp43-41g5.GenBank:AC099533.36     TACGTGACCTTCAGGGTACATTTTCTTCATTGATGATCAATGGAAATGAGAAATCACTGA 25423
human.FAM30B-LOC105372935-GGT2.NCBI.                                               tacatgaccttcaggatacattttcttcattgatgatcaatggaaatgagaaatcactga 22333
human.FAM230E-LOC105377182-GGT3P.NCBI.                                             tacatgaccttcaggatacattttcttcattgatgatcaatggaaatgagaaatcactga 21070
human.FAM230J-LOC105372942-GGTLC5P.NCBI.rev.compl                                  tacatgaccttcaggatacattttcttcattgatgatcaatggaaatgagaaatcactga 21783
human.FAM230A.ncRNA.18487127.18500594-GGTLC3.18516335.18518165.rev.compl.REF       tacatgaccttcaggatacattttcttcattgatgatcaatggaaatgagaaatcactga 10547
                                                                                   *** *********** ********************************************

chimp.LOC112206744-LOC107973052-GGT2.revcompl                                      ctatttttctatcattggaatctactctctactgctcgtgctgttcctgtcttttgggg 23701
chimp54546-135457.revcompl.Pan.troglodytes.clone.rp43-41g5.GenBank:AC099533.36     CTATTTTTCTATCATTGGAATCTACTCTCTACTGCTCATGCTGTTCCTGTCTTTTGGGG 25483
human.FAM30B-LOC105372935-GGT2.NCBI.                                               ctatttttctatcattggaatctactctccactgctcatgctgttcctgtcttttgggg 22393
human.FAM230E-LOC105377182-GGT3P.NCBI.                                             ctatttttctatcattggaatctactctccactgctcatgctgttcctgtcttttgggg 21130
human.FAM230J-LOC105372942-GGTLC5P.NCBI.rev.compl                                  ctatttttctatcattggaatctactctccactgctcatgctgttcctgtcttttgggg 21843
human.FAM230A.ncRNA.18487127.18500594-GGTLC3.18516335.18518165.rev.compl.REF       ctatttttctatcattggaatctactctccactgctcatgctgttcctgtcttttgggg 10607
                                                                                   ***************************** ******* **********************

chimp.LOC112206744-LOC107973052-GGT2.revcompl                                      aagatggaggatcaatcagtgtgcgctgcactgagtggaaggaaggagaactgcgacaaa 23761
chimp54546-135457.revcompl.Pan.troglodytes.clone.rp43-41g5.GenBank:AC099533.36     AAGATGGAGGATCAATCAGTGTGCGCTGCACTGAGTGGAAGGAAGGAGAACTGTGACAAA 25543
human.FAM30B-LOC105372935-GGT2.NCBI.                                               aagatggaggatcaatcagtgtgcactgcactgagtggaaggaaggagaactgtgacaaa 22453
human.FAM230E-LOC105377182-GGT3P.NCBI.                                             aagatggaggatcaatcagtgtgcgctgccctgagtggaaggaaggagaactgtgacaaa 21190
human.FAM230J-LOC105372942-GGTLC5P.NCBI.rev.compl                                  aagatggaggatcaatcagtgtgcgctgcactgagtggaaggaaggagaactgtgacaaa 21903
human.FAM230A.ncRNA.18487127.18500594-GGTLC3.18516335.18518165.rev.compl.REF       aagatggaggatcaatcagtgtgcgctgcactgagtggaaggaaggagaactgtgacaaa 10667
                                                                                   ************************ **** *********************** ******

chimp.LOC112206744-LOC107973052-GGT2.revcompl                                      aattaaggaaggatgagagacgggagggcccttcatccagctgcttgcagagtcctcctg 23821
chimp54546-135457.revcompl.Pan.troglodytes.clone.rp43-41g5.GenBank:AC099533.36     AATTAAGGAAGGATGAGAGACGGGAGGGCCCTTCATCCAGCTGCTTGCAGAGTCCTCCTG 25603
human.FAM30B-LOC105372935-GGT2.NCBI.                                               aattaaggaaggatgagagacgggagggcccttcatccagctgcttgcagagtcctcctg 22513
human.FAM230E-LOC105377182-GGT3P.NCBI.                                             aattaaggaaggatgagagacgggagggcccttcatccagctgcttgcagagtcctcctg 21250
human.FAM230J-LOC105372942-GGTLC5P.NCBI.rev.compl                                  aattaaggaaggatgagagacgggagggcccttcatccagctgcttgcagagtcctcctg 21963
human.FAM230A.ncRNA.18487127.18500594-GGTLC3.18516335.18518165.rev.compl.REF       aattaaggaaggatgagagacgggagggcccttcatccagctgcttgcagagtcctcctg 10727
                                                                                   ************************************************************

chimp.LOC112206744-LOC107973052-GGT2.revcompl                                      aggaggaaagccccgtggctccctggcgaaggagcagtgagggctgcgtgactcccacag 23881
chimp54546-135457.revcompl.Pan.troglodytes.clone.rp43-41g5.GenBank:AC099533.36     AGGAGGAAAGCCCCGTGGCTCCCTGGCGAAGGAGCAGTGAGGGCTGCGTGACTCCCACAG 25663
human.FAM30B-LOC105372935-GGT2.NCBI.                                               aggaggaaagccccgtggctccctggagaaggagcattgagggctgcgtgactcccacag 22573
human.FAM230E-LOC105377182-GGT3P.NCBI.                                             aggaggaaagccccgtggctccctggcgaaggagcagtgagggctgcgtgactcccacag 21310
human.FAM230J-LOC105372942-GGTLC5P.NCBI.rev.compl                                  aggaggaaagccccgtggctccctggcgaaggagcagtgagggctgcgtgactcccacag 22023
human.FAM230A.ncRNA.18487127.18500594-GGTLC3.18516335.18518165.rev.compl.REF       aggaggaaagccccgtggctccctggcgaaggagcagtgagggctgcgtgactcccacag 10787
                                                                                   ************************** ********* ***********************

chimp.LOC112206744-LOC107973052-GGT2.revcompl                                      tgaagtgtgtggtatgtctgaggacacccaggctggtggtccatgaggagccagtggcag 23941
chimp54546-135457.revcompl.Pan.troglodytes.clone.rp43-41g5.GenBank:AC099533.36     TGAAGTGTGTGGTATGTCTGAGGACACCCAGGCTGGTGGTCCATGAGGAGCCAGTGGCAG 25723
human.FAM30B-LOC105372935-GGT2.NCBI.                                               tgaagtgtgtggtatgtctgaggacacccaggctggtggtccatgaggagccagtggcag 22633
human.FAM230E-LOC105377182-GGT3P.NCBI.                                             tgaagtgtgtggtatgtctgaggacacccaggctggtggtccatgaggagccagtggcag 21370
human.FAM230J-LOC105372942-GGTLC5P.NCBI.rev.compl                                  tgaagtgtgtggtatgtctgaggacacccaggctggtggtccatgaggagccagtggcag 22083
human.FAM230A.ncRNA.18487127.18500594-GGTLC3.18516335.18518165.rev.compl.REF       tgaagtgtgtggtatgtctgaggacacccaggctggtggtccatgaggagccagtggcag 10847
                                                                                   ************************************************************
```

```
chimp.LOC112206744-LOC107973052-GGT2.revcompl                                   agtgagaagcagaaaggccaggagggtggctggaggccaggctctgagtcattctccatg 24001
chimp54546-135457.revcompl.Pan.troglodytes.clone.rp43-41g5.GenBank:AC099533.36  AGTGAGAAGCAGAAAGGCCAGGAGGGTGGCTGGAGGCCAGGCTCTGAGTCATTCTCCATG 25783
human.FAM30B-LOC105372935-GGT2.NCBI.                                            agtgagaagaagaaaggccaggagggtggctggaggccaggctctgagtcattctccatg 22693
human.FAM230E-LOC105377182-GGT3P.NCBI.                                          agtgagaagcagaaaggccaggagggtggctggaggccaggctctgagtcattctccatg 21430
human.FAM230J-LOC105372942-GGTLC5P.NCBI.rev.compl                               agtgagaagcagaaaggccaggagggtggctggaggccaggctctgagtcattctccatg 22143
human.FAM230A.ncRNA.18487127.18500594-GGTLC3.18516335.18518165.rev.compl.REF    agtgagaagaagaaaggccaggagggtggctggaggccaggctctgagtcattctccatg 10907
                                                                                ********* **************************************************

chimp.LOC112206744-LOC107973052-GGT2.revcompl                                   tgatggaaacagccggagcccagtgggcttggaggtacaggatgcggtggctgatgacag 24061
chimp54546-135457.revcompl.Pan.troglodytes.clone.rp43-41g5.GenBank:AC099533.36  TGATGGAAACAGCCGGAGCCCAGTGGGCTTGGAGGTACAGGATGCAGTGGCTGATGACAG 25843
human.FAM30B-LOC105372935-GGT2.NCBI.                                            tgatggaaacagccggagcccagtgggcttggaggtacaggatgcggtggctgatgacag 22753
human.FAM230E-LOC105377182-GGT3P.NCBI.                                          tgatggaaacagccggagcccagtgggcttggaggtacaggatgcggtggctgatgacag 21490
human.FAM230J-LOC105372942-GGTLC5P.NCBI.rev.compl                               tgatggaaacagccggagcccagtggacttggaggtacaggatgcggtggctgatgacag 22203
human.FAM230A.ncRNA.18487127.18500594-GGTLC3.18516335.18518165.rev.compl.REF    tgatggaaacagccggagcccagtgggcttggaggtacaggatgcggtggctgatgacag 10967
                                                                                ************************** ****************** **************

chimp.LOC112206744-LOC107973052-GGT2.revcompl                                   aacaatgtggagagaggcgtcatttgtcaaatccttactttgttctgggcattgtgctaa 24121
chimp54546-135457.revcompl.Pan.troglodytes.clone.rp43-41g5.GenBank:AC099533.36  AACAATGTGGAGAGAGGCGTCATTTGTCAAATCCTTACTTTGTTCTGGGCATTGTGCTAA 25903
human.FAM30B-LOC105372935-GGT2.NCBI.                                            aacaatgtggagagaggcgtcatttgtcaaatccttactttgttctgggcattgtgctaa 22813
human.FAM230E-LOC105377182-GGT3P.NCBI.                                          aacaatgtggagagaggtgtcatttgtcaaatccttactttgttctgggcattgtgctaa 21550
human.FAM230J-LOC105372942-GGTLC5P.NCBI.rev.compl                               aacaatgtggagagaggcgtcatttgtcaaatccttactttgttctgggcattgtgctaa 22263
human.FAM230A.ncRNA.18487127.18500594-GGTLC3.18516335.18518165.rev.compl.REF    aacaatgtggagagaggcgtcatttgtcaaatccttactttgttctgggcattgtactaa 11027
                                                                                ***************** ************************************** ****

chimp.LOC112206744-LOC107973052-GGT2.revcompl                                   aaattctgatggctcatcccatttaggggctgaaagttgcagaggtttaggaagctcacc 24181
chimp54546-135457.revcompl.Pan.troglodytes.clone.rp43-41g5.GenBank:AC099533.36  A------CGTGGCTCATCCCATTTAGGGGCTGAAAGTTGCAGAGGTTTAGGAAGCTCACC 25957
human.FAM30B-LOC105372935-GGT2.NCBI.                                            aaattctgatggctcatcccatttaggggctgaaagttgcagaggtttaggaagctcacc 22873
human.FAM230E-LOC105377182-GGT3P.NCBI.                                          aaattctgatggctcatcccatttaggggctgaaagttgcagaggtttaggaagctcacc 21610
human.FAM230J-LOC105372942-GGTLC5P.NCBI.rev.compl                               aaattctgatggctcatcccatttaggggctgaaagttgcagaggtttaggaagctcacc 22323
human.FAM230A.ncRNA.18487127.18500594-GGTLC3.18516335.18518165.rev.compl.REF    aaattctaatggctcatcccatttaggggctgaaagttgcagaggtttaggaagctcacc 11087
                                                                                *        ***************************************************

chimp.LOC112206744-LOC107973052-GGT2.revcompl                                   catgatactggagcccccatctcctgccctagtgccgtccaccttctcacccagccacca 24241
chimp54546-135457.revcompl.Pan.troglodytes.clone.rp43-41g5.GenBank:AC099533.36  CACGATACTGGAGCCCCCATCTCCTGCCCTAGTGCCGTCCACCTTCTCACCCAGCCACCA 26017
human.FAM30B-LOC105372935-GGT2.NCBI.                                            cacgatactggagcccccatctcctgccctagtgctgtccaccttctcacccagccacca 22933
human.FAM230E-LOC105377182-GGT3P.NCBI.                                          cacgatactggagcccccatctcctgccctagtgctgtccaccttctcacccagccacca 21670
human.FAM230J-LOC105372942-GGTLC5P.NCBI.rev.compl                               cacgatactggagcccccatctcctgccctagtgctgtccaccttctcacccagccacca 22383
human.FAM230A.ncRNA.18487127.18500594-GGTLC3.18516335.18518165.rev.compl.REF    cacgatactggagcccccatctcttgccctagtgctgtccaccttctcacccagccacca 11147
                                                                                ** ******************** *********** ************************

chimp.LOC112206744-LOC107973052-GGT2.revcompl                                   cctgtttcaggggaacacacagaagtggtaacctcttatggagaggcaagtaaattctgc 24301
chimp54546-135457.revcompl.Pan.troglodytes.clone.rp43-41g5.GenBank:AC099533.36  CCTATTTCAGGGGAACACACAGAAGTGGTAACCTCTTATGGAGAGGCAAGTAAATTCTGC 26077
human.FAM30B-LOC105372935-GGT2.NCBI.                                            cctgtttcggggaacacacagaagtggtaacctcttatggagaggcaagtaaattctgc 22993
human.FAM230E-LOC105377182-GGT3P.NCBI.                                          cctgtttcggggaacacacagaagtggtaacctcttatggagaggcaagtaaattctgc 21730
human.FAM230J-LOC105372942-GGTLC5P.NCBI.rev.compl                               cctgtttcggggaacacacagaagtggtaacctcttatggtgaggcaagtaaattctgc 22443
human.FAM230A.ncRNA.18487127.18500594-GGTLC3.18516335.18518165.rev.compl.REF    cctgtttcggggaacacacagaagtggtaacctcttatggagaggcaagtaaattctgc 11207
                                                                                *** **** ******************************** ******************

chimp.LOC112206744-LOC107973052-GGT2.revcompl                                   tgtttttgttattcacagaaaaacactggctcatgtgggttgggaaggtgaaataccaga 24361
chimp54546-135457.revcompl.Pan.troglodytes.clone.rp43-41g5.GenBank:AC099533.36  TGTTTTTGTTATTCACAGAAAAACACTGGCTCTTGTGGGTTGGGAAGGTGAAATACCAGA 26137
human.FAM30B-LOC105372935-GGT2.NCBI.                                            tgtttttgttattcacagaaaaacactggctcgtgtgggttgggaaggtgaaataccaga 23053
human.FAM230E-LOC105377182-GGT3P.NCBI.                                          tgtttttgttattcacagaaaaacactggctcgtgtgggttgggaaggtgaaataccaga 21790
human.FAM230J-LOC105372942-GGTLC5P.NCBI.rev.compl                               tgtttttgttattcacagaaaaacactggctcgtgtgggttgggaaggtgaaataccaga 22503
human.FAM230A.ncRNA.18487127.18500594-GGTLC3.18516335.18518165.rev.compl.REF    tgtttttgttattcacagaaaaacactggctcgtgtgggttgggaaggtgaaataccaga 11267
                                                                                ******************************** ***************************

chimp.LOC112206744-LOC107973052-GGT2.revcompl                                   agtatttcatctagttatttctacccatgcaactcctatagtattgaaatgcataggtta 24421
chimp54546-135457.revcompl.Pan.troglodytes.clone.rp43-41g5.GenBank:AC099533.36  AGTATTTCATCTAGTTATTTCTACCCATGCAACTCCTATAGTATTGAAATGCATAGGTTA 26197
human.FAM30B-LOC105372935-GGT2.NCBI.                                            agtatttcatctggttatttctacccatgcgactcctatagtattgaaatgcataggtta 23113
human.FAM230E-LOC105377182-GGT3P.NCBI.                                          agtatttcatctggttatttctacccatgcgactcctatagtattgaaatgcataggtta 21850
human.FAM230J-LOC105372942-GGTLC5P.NCBI.rev.compl                               agtatttcatctggttatttctacccatgcgactcctatagtattgaaatgcataggtta 22563
human.FAM230A.ncRNA.18487127.18500594-GGTLC3.18516335.18518165.rev.compl.REF    agtatttcatctggttatttctacccatgcaactcctatagtattgaaatgcataggtta 11327
                                                                                ************ ***************** *****************************

chimp.LOC112206744-LOC107973052-GGT2.revcompl                                   gcattttggccaatttactcagcattctgggttaaaggct-------------tttatt 24468
chimp54546-135457.revcompl.Pan.troglodytes.clone.rp43-41g5.GenBank:AC099533.36  GCATTTTGGCCAATTTACTCAGCATTCTGGGTTAAAGGCT-------------TTTATT 26244
human.FAM30B-LOC105372935-GGT2.NCBI.                                            gcattttggccaatttactcagcattctgggttaaaggcttttatttattttatttatt 23173
human.FAM230E-LOC105377182-GGT3P.NCBI.                                          gcattttggccaatttactcagcattctgggttaaaggcttttatttat----tttatt 21906
human.FAM230J-LOC105372942-GGTLC5P.NCBI.rev.compl                               gcattttggccaatttactcagcattctgggttaaaggcttttatttattttatttatt 22623
human.FAM230A.ncRNA.18487127.18500594-GGTLC3.18516335.18518165.rev.compl.REF    gcattttggccaatttactcagcattctgggttaaaggcttttatttat----tttatt 11383
                                                                                ****************************************             ******

chimp.LOC112206744-LOC107973052-GGT2.revcompl                                   tatttatttatttatttcgagatggagtttcgctcttgttgcccaagctggagtgcaat 24528
chimp54546-135457.revcompl.Pan.troglodytes.clone.rp43-41g5.GenBank:AC099533.36  TATTTATTTATTTATTTTGAGATGGAGTTTCGCTCTTGTTGCCCAAGCTGGAGTGCAAT 26304
human.FAM30B-LOC105372935-GGT2.NCBI.                                            tatttatttatttattttgagatggagtttcactcttgttgcccaagctggagtgcaat 23233
human.FAM230E-LOC105377182-GGT3P.NCBI.                                          tatttatttatttactttgagatggagtttcactcttgttgcccaagctggagggcaat 21966
human.FAM230J-LOC105372942-GGTLC5P.NCBI.rev.compl                               tatttatttatttattttgagatggagtttcactcttgttgcccaagctggagtgcaat 22683
human.FAM230A.ncRNA.18487127.18500594-GGTLC3.18516335.18518165.rev.compl.REF    tatttatttatttattttgagatggagtttcactcttgttgcccaagctggagtgcaat 11443
                                                                                ************** *** ************* ********************* *****

chimp.LOC112206744-LOC107973052-GGT2.revcompl                                   ggtgcgatcctggttcactgcaacctccgcctcccaggttcaaactattctcctgtctca 24588
chimp54546-135457.revcompl.Pan.troglodytes.clone.rp43-41g5.GenBank:AC099533.36  GGTGCGATCCTGGTTCACTGCAACCTCCGCCTCCCAGATTCAAACTATTCTCCTTTCTCA 26364
human.FAM30B-LOC105372935-GGT2.NCBI.                                            ggtgcgatcctggttcattgcaacctccgcctcccaggttcaaactattctcctgtctca 23293
human.FAM230E-LOC105377182-GGT3P.NCBI.                                          ggtgcgatcctggttcattgcaacctccgcctcccaggttcaaactattctcctgtctca 22026
human.FAM230J-LOC105372942-GGTLC5P.NCBI.rev.compl                               ggtgcgatcctggttcattgcaacctccgcctcccaggttcaaactattctcctgtctca 22743
human.FAM230A.ncRNA.18487127.18500594-GGTLC3.18516335.18518165.rev.compl.REF    ggtgcgatcctggttcattgcaacctccgcctcccaggttcaaactattctcctgtctca 11503
                                                                                ***************** ******************* **************** *****

chimp.LOC112206744-LOC107973052-GGT2.revcompl                                   gcctcccaagtagctgagattataggcacgtgccaccacactgggctaatttttttgtat 24648
chimp54546-135457.revcompl.Pan.troglodytes.clone.rp43-41g5.GenBank:AC099533.36  GCCTCCCAAGTAGCTGAGATTATAGGCATGTGCCACCACCCTGGGCTAATTTTTTTGTAT 26424
human.FAM30B-LOC105372935-GGT2.NCBI.                                            gcctcccaagtagctgagattataggcacatgccaccacactgggctaatttttttgtat 23353
human.FAM230E-LOC105377182-GGT3P.NCBI.                                          gcctcccaagtagctgagattataggcacatgccaccacactgggctaatttttttgtat 22086
human.FAM230J-LOC105372942-GGTLC5P.NCBI.rev.compl                               gcctcccaagtagctgagattataggcacatgccaccacactgggctaatttttttgtat 22803
human.FAM230A.ncRNA.18487127.18500594-GGTLC3.18516335.18518165.rev.compl.REF    gcctcccaagtagctgagattataggcacatgccaccacactgggctaatttttttgtat 11563
                                                                                ****************************  ********* ********************

chimp.LOC112206744-LOC107973052-GGT2.revcompl                                   ttttagtagaaatgagatttcaccatgttggtcaggctggtttcaaaatcctgacctcag 24708
chimp54546-135457.revcompl.Pan.troglodytes.clone.rp43-41g5.GenBank:AC099533.36  TTTTAGTAGAAATGAGATTTCACCATGTTGGTCAGGCTGGTTTCAAACTCCTGACCTCAG 26484
human.FAM30B-LOC105372935-GGT2.NCBI.                                            ttttagtagaaatgagatttcaccatgttggtcaggctggtttcgaactcctgacctcag 23413
human.FAM230E-LOC105377182-GGT3P.NCBI.                                          ttttagtagaaatgagatttcaccatgttggtcaggctggtttcgaactcctgacctcag 22146
human.FAM230J-LOC105372942-GGTLC5P.NCBI.rev.compl                               ttttagtagaaatgagatttcaccatgttggtcaggctggtttcgaactcctgacctcag 22863
human.FAM230A.ncRNA.18487127.18500594-GGTLC3.18516335.18518165.rev.compl.REF    ttttagtagaaatgagatttcaccatgttggtcaggctggtttcgaactcctgacctcag 11623
                                                                                ******************************************** ** ************

chimp.LOC112206744-LOC107973052-GGT2.revcompl                                   gtgatctgccctcctcggcctcttaaagtgctgggattacaggcgtgagccaccatgccc 24768
chimp54546-135457.revcompl.Pan.troglodytes.clone.rp43-41g5.GenBank:AC099533.36  GTGATCTGCCCTCCTCGGCCTCTTAAAGTGCTGGAATTACAGGCGTGAGCCACCATGCCC 26544
human.FAM30B-LOC105372935-GGT2.NCBI.                                            gtgatctgccccctcggcctcttaaagtgctgggattacaggcgtgagccaccacgcct 23473
human.FAM230E-LOC105377182-GGT3P.NCBI.                                          gtgatctgccctcctcggcctcttaaagtgctgggattacaggcgtgagccaccacgcct 22206
human.FAM230J-LOC105372942-GGTLC5P.NCBI.rev.compl                               gtgatctgccctcctcggcctcttaaagtgctgggattacaggtgtgagccaccacgcct 22923
human.FAM230A.ncRNA.18487127.18500594-GGTLC3.18516335.18518165.rev.compl.REF    gtgatctgccctcctcggcctcttaaagtgctgggattacaggcgtgagccaccacgcct 11683
                                                                                *********** ********************** ******** *********** ***

chimp.LOC112206744-LOC107973052-GGT2.revcompl                                   accctaaagtcttttaaaattcacttgtataagttgacttagtttctttttttttttctt 24828
chimp54546-135457.revcompl.Pan.troglodytes.clone.rp43-41g5.GenBank:AC099533.36  GCCCTAAAGTCTTTTAAAATTCACTTGTATAAGTT------------------------- 26579
human.FAM30B-LOC105372935-GGT2.NCBI.                                            gccctaaagtcttttaaaattcacttgtataagtt------------------------- 23508
human.FAM230E-LOC105377182-GGT3P.NCBI.                                          gccctaaagtcttttaaaattcacttgtataagtt------------------------- 22241
human.FAM230J-LOC105372942-GGTLC5P.NCBI.rev.compl                               gccctaaagtcttttaaaattcacttgtataagtt------------------------- 22958
human.FAM230A.ncRNA.18487127.18500594-GGTLC3.18516335.18518165.rev.compl.REF    gccctaaagtcttttaaaattcacttgtataagtt------------------------- 11718
                                                                                 **********************************

chimp.LOC112206744-LOC107973052-GGT2.revcompl                                   tttctttttctttttttttttttattgatcattcttggacttagttttctttaaccttgt 24888
chimp54546-135457.revcompl.Pan.troglodytes.clone.rp43-41g5.GenBank:AC099533.36  -------------------------------------GACTTAGTTTTCTTTAACCTTGT 26602
human.FAM30B-LOC105372935-GGT2.NCBI.                                            -------------------------------------gacttagttttctttaaccttgt 23531
human.FAM230E-LOC105377182-GGT3P.NCBI.                                          -------------------------------------gacttagttttctttaaccttgt 22264
human.FAM230J-LOC105372942-GGTLC5P.NCBI.rev.compl                               -------------------------------------gacttagttttctttaaccttgt 22981
human.FAM230A.ncRNA.18487127.18500594-GGTLC3.18516335.18518165.rev.compl.REF    -------------------------------------gacttagttttctttaaccttgt 11741
                                                                                                                     ***********************

chimp.LOC112206744-LOC107973052-GGT2.revcompl                                   agaaaaat-caaaaatggcaatctgttttatcacacaaataatgtctttttaatggagtg 24947
chimp54546-135457.revcompl.Pan.troglodytes.clone.rp43-41g5.GenBank:AC099533.36  AGAAAAATACAAAAATGGCAATCTCTTTTATCACACAAATAATGTCTTTTTAATGGAGTG 26662
human.FAM30B-LOC105372935-GGT2.NCBI.                                            agaaaaatacaaaaatggcaatctcttttatcacacaaataatgtctttttaatggagtg 23591
human.FAM230E-LOC105377182-GGT3P.NCBI.                                          agaaaaatacaaaaatggcaatctcttttatcacacaaataatgtctttttaatggagtg 22324
human.FAM230J-LOC105372942-GGTLC5P.NCBI.rev.compl                               agaaaaatacaaaaatagcaatctcttttatcacacaaataatgtctttttaatggagtg 23041
human.FAM230A.ncRNA.18487127.18500594-GGTLC3.18516335.18518165.rev.compl.REF    agaaaaatacaaaaatggcaatctcttttatcacacaaataatgtctttttaatggagtg 11801
                                                                                ******** ******* ******* ***********************************

chimp.LOC112206744-LOC107973052-GGT2.revcompl                                   a-tttttttctaattgaggtattatgtacttttcatttactaattattgtttacatttga 25006
chimp54546-135457.revcompl.Pan.troglodytes.clone.rp43-41g5.GenBank:AC099533.36  A-TTTTTTTCTAATTGAGGTATTATGTACTTTTCATTTACTAATTATTGTTTACATTTGA 26721
human.FAM30B-LOC105372935-GGT2.NCBI.                                            a-tttttttctaattgaggtattatgtacttttcatttactaattattgtttacatttga 23650
human.FAM230E-LOC105377182-GGT3P.NCBI.                                          attttttttctaattgaggtattatgtacttttcatttactaattattgtttacatttga 22384
human.FAM230J-LOC105372942-GGTLC5P.NCBI.rev.compl                               a-tttttttctaattgaggtattatgtacttttcatttactaattattgtttacatttga 23100
human.FAM230A.ncRNA.18487127.18500594-GGTLC3.18516335.18518165.rev.compl.REF    a-tttttttctaattgaggtattatgtacttttcatttacaaattattgtttacatttga 11860
                                                                                * ************************************** *******************

chimp.LOC112206744-LOC107973052-GGT2.revcompl                                   agtgttttatgaattaatatttaattgcatagatgaagattactagttataggcatttta 25066
```

```
chimp54546-135457.revcompl.Pan.troglodytes.clone.rp43-41g5.GenBank:AC099533.36      AGTGTTTTATGAATTAATATTTAATTGCATAGATGAAGATTACTAGTTATAGGCATTTTA 26781
human.FAM30B-LOC105372935-GGT2.NCBI.                                                agtgttttatgaattaatatttaattgcatagatgaagattactagttataggcatttta 23710
human.FAM230E-LOC105377182-GGT3P.NCBI.                                              agtgttttatgaattaatatttaattgcatagatgaagattactagttataggcatttta 22444
human.FAM230J-LOC105372942-GGTLC5P.NCBI.rev.compl                                   agtgttttatgaattaatatttaattgcatagatgaagattactagttataggcatttta 23160
human.FAM230A.ncRNA.18487127.18500594-GGTLC3.18516335.18518165.rev.compl.REF        agtgttttatgaattaatatttaattgcatagatgaagattactagttataggcatttta 11920
                                                                                    ************************************************************

chimp.LOC112206744-LOC107973052-GGT2.revcompl                                       gtaaccaatactcattaagcatagcgtggattcatatgacatcaaggagctattttattt 25126
chimp54546-135457.revcompl.Pan.troglodytes.clone.rp43-41g5.GenBank:AC099533.36      CTAACCAATACTCATTAAGCATAGCGTGGATTCATATGACATCAAGGAGCTATTTTATTT 26841
human.FAM30B-LOC105372935-GGT2.NCBI.                                                ctaaccaatactcattaagcatagcgtggattcctatgacatcaaggagctattttattt 23770
human.FAM230E-LOC105377182-GGT3P.NCBI.                                              ctaaccaatactcattaagcatagcgtggattcatatgacatcaaggagctattttattt 22504
human.FAM230J-LOC105372942-GGTLC5P.NCBI.rev.compl                                   ctaaccaatactcattaagcatagcgtggattcctatgacatcaaggagctattttattt 23220
human.FAM230A.ncRNA.18487127.18500594-GGTLC3.18516335.18518165.rev.compl.REF        ctaaccaatactcattaagcatagcatggattcatatgacatcaaggagctattttattt 11980
                                                                                     ************************ ******* **************************

chimp.LOC112206744-LOC107973052-GGT2.revcompl                                       ggtaaaacgaaaaagcacaagaatgaacgaacgcaagaactgaaacggtggagacaccta 25186
chimp54546-135457.revcompl.Pan.troglodytes.clone.rp43-41g5.GenBank:AC099533.36      GGTAGAACGAAAAAGCACAAGAATGAACGAACGCAAGAACTGAAACAGTGGAGACACCTA 26901
human.FAM30B-LOC105372935-GGT2.NCBI.                                                ggtaaaacgaaaaagcacaagaatgaacgaacgcaagaactgaaacagtggagacaccta 23830
human.FAM230E-LOC105377182-GGT3P.NCBI.                                              ggtaaaacgaaaaagcacaagaatgaacgaacgcaagaactgaaacagtggagacaccta 22564
human.FAM230J-LOC105372942-GGTLC5P.NCBI.rev.compl                                   ggtaaaacgaaaaagcacaagaatgaacgaacgcaagaactgaaacagtggagacaccta 23280
human.FAM230A.ncRNA.18487127.18500594-GGTLC3.18516335.18518165.rev.compl.REF        ggtaaaacgaaaaagcacaagaatgaacgaatgcaagaactgaaacagtggagacaccta 12040
                                                                                    **** ************************** ************** *************

chimp.LOC112206744-LOC107973052-GGT2.revcompl                                       gaatgacttgtctaagatctaaatcattttgttgtcttcccagcgtacttattatcctga 25246
chimp54546-135457.revcompl.Pan.troglodytes.clone.rp43-41g5.GenBank:AC099533.36      GAATGACTTATCTAAGATCTAAATCATTTTGTTGTCTTCCCAGCGTACTTATTATCCTGA 26961
human.FAM30B-LOC105372935-GGT2.NCBI.                                                gaatgacttgtctaagatctaaatcattttgttgtcttcccagcgtacttattatcctga 23890
human.FAM230E-LOC105377182-GGT3P.NCBI.                                              gaatgacttgtctaagatctaaatcattttgttgtcttcccagcgtacttattatcctga 22624
human.FAM230J-LOC105372942-GGTLC5P.NCBI.rev.compl                                   gaatgacttgtctaagatctaaatcattttgttgtcttcccagcgtacttattatcctga 23340
human.FAM230A.ncRNA.18487127.18500594-GGTLC3.18516335.18518165.rev.compl.REF        gaatgacttgtctaagatctaaatcattttgttgtcttcccagcatacttattatcctga 12100
                                                                                    ********* ********************************** ***************

chimp.LOC112206744-LOC107973052-GGT2.revcompl                                       tcattgtcatcagcattgtttgggtccttttagcacagatttctcaaaatgggtaactcc 25306
chimp54546-135457.revcompl.Pan.troglodytes.clone.rp43-41g5.GenBank:AC099533.36      TCATTGTCATCAGCATTGTTTGGGTCCTTTTAGCACAGATTTCTCAAAATGGGTAACTCC 27021
human.FAM30B-LOC105372935-GGT2.NCBI.                                                tcattgtcatcagcattgtttgggtccttttagcacagatttctcaaaatgggtaactcc 23950
human.FAM230E-LOC105377182-GGT3P.NCBI.                                              tcattgtcatcagcattgtttgggtccttttagcacagatttctcaaaatgggtaactcc 22684
human.FAM230J-LOC105372942-GGTLC5P.NCBI.rev.compl                                   tcattgtcatcagcattgtttgggtccttttagcacagatttctcaaaatgggtaactcc 23400
human.FAM230A.ncRNA.18487127.18500594-GGTLC3.18516335.18518165.rev.compl.REF        tcattgtcatcagcattgtttgggtccttttagcacagatttctcaaaatgggtaactcc 12160
                                                                                    ************************************************************

chimp.LOC112206744-LOC107973052-GGT2.revcompl                                       ataacagttggaagcttacgaattcatataatttgtaagaggtcaatttggaagtaccta 25366
chimp54546-135457.revcompl.Pan.troglodytes.clone.rp43-41g5.GenBank:AC099533.36      ATAACAGTTGGAAGCTTATGAATTCATATAATTTGTAAGAGGTCAATTTGGAAGTACCTA 27081
human.FAM30B-LOC105372935-GGT2.NCBI.                                                ataacagttggaagcttacgaattcatataatttgtaagaggtcaatttgggagtaccta 24010
human.FAM230E-LOC105377182-GGT3P.NCBI.                                              ataacagttggaagcttacgaattcatataatttgtaagaggtcaatttggaagtaccta 22744
human.FAM230J-LOC105372942-GGTLC5P.NCBI.rev.compl                                   ataacagttggaagcttacgaattcatataatttgtaagaggtcaatttggaagtaccta 23460
human.FAM230A.ncRNA.18487127.18500594-GGTLC3.18516335.18518165.rev.compl.REF        ataacagttggaagcttacgaattcatataatttgtaagaggtcaatttggaagtaccta 12220
                                                                                    ****************** ******************************** ********

chimp.LOC112206744-LOC107973052-GGT2.revcompl                                       tctattttaaaattccaataacctgggaatttcatcccatgtctagagtcttccatgtaa 25426
chimp54546-135457.revcompl.Pan.troglodytes.clone.rp43-41g5.GenBank:AC099533.36      TGTATTTTAAAATTCCAATAACCTGGGAATTTCATCCCATGTCTAGAGTCTTCTATGTAA 27141
human.FAM30B-LOC105372935-GGT2.NCBI.                                                tctattttaaaattccaataacctgggaatttcatcccatgtctagagtcttttatgtaa 24070
human.FAM230E-LOC105377182-GGT3P.NCBI.                                              tctattttaaaattccaataacctgggaatttcatcccatgtctagagtcttttatgtaa 22804
human.FAM230J-LOC105372942-GGTLC5P.NCBI.rev.compl                                   tctattttaaaattccaataacctgggaatttcatcccatgtctagagtcttttatgtaa 23520
human.FAM230A.ncRNA.18487127.18500594-GGTLC3.18516335.18518165.rev.compl.REF        tctattttaaaattccaataacctgggaatttcatcccatgtctagagtcttttatgtaa 12280
                                                                                    * **************************************************  ******

chimp.LOC112206744-LOC107973052-GGT2.revcompl                                       aatatttccacaattaggagaaatatgtgcatggggattttctatgtagcggtgttttga 25486
chimp54546-135457.revcompl.Pan.troglodytes.clone.rp43-41g5.GenBank:AC099533.36      AATATTTCCACAATTAGGAGAAATATGTGCATGGGGATTTTCTATGTAGCGGTGTTTTGA 27201
human.FAM30B-LOC105372935-GGT2.NCBI.                                                aatatttccacaattaggagaaatatgtgcatggggattttctatgtagcggtgttttga 24130
human.FAM230E-LOC105377182-GGT3P.NCBI.                                              aatatttccacaattaggagaaatatgtgcatggggattttctatgtagcggtgttttga 22864
human.FAM230J-LOC105372942-GGTLC5P.NCBI.rev.compl                                   aatatttccacaattaggagaaatatgtgcatggggattttctatgtagcggtgttttga 23580
human.FAM230A.ncRNA.18487127.18500594-GGTLC3.18516335.18518165.rev.compl.REF        aatatttccacaattaggagaaatatgtgcatggggattttctatgtagcggtgttttga 12340
                                                                                    ************************************************************

chimp.LOC112206744-LOC107973052-GGT2.revcompl                                       tagaatagaaaaatgggataaaccaaatttccatcacgaaggaaatagtaatatgctgaa 25546
chimp54546-135457.revcompl.Pan.troglodytes.clone.rp43-41g5.GenBank:AC099533.36      TAGAATAGAAAAATGGGATAAACCAAATTTCCATCACGAAGGAAATAGTAATATGCTGAA 27261
human.FAM30B-LOC105372935-GGT2.NCBI.                                                tagaatagaaaattgggataaaccaaatttccatcacgaaggaaatagtaatatgctgaa 24190
human.FAM230E-LOC105377182-GGT3P.NCBI.                                              tagaatagaaaattgggataaaccaaatttccatcacgaaggaaatagtaatatgctgaa 22924
human.FAM230J-LOC105372942-GGTLC5P.NCBI.rev.compl                                   tggaatagaaaattgggataaaccaaatttccatcacgaaggaaatagtaatatgctgaa 23640
human.FAM230A.ncRNA.18487127.18500594-GGTLC3.18516335.18518165.rev.compl.REF        tagaatagaaaattgggataaatcaaatttccatcacgaaggaaatagtaatatgctgaa 12400
                                                                                    * ********** ********* *************************************

chimp.LOC112206744-LOC107973052-GGT2.revcompl                                       taataatacagcgaatattatgcaggctttaaacatcaaaaaagagttcaacttctgact 25606
chimp54546-135457.revcompl.Pan.troglodytes.clone.rp43-41g5.GenBank:AC099533.36      TAATAATACAGCGAATATTATGCAGGCTTTAAACATCAAAAAAGAGTTCAACTTCTGACT 27321
human.FAM30B-LOC105372935-GGT2.NCBI.                                                taataatacagcgaatattatgcaggctttaaacatcaaaaaagagttcaacttctgact 24250
human.FAM230E-LOC105377182-GGT3P.NCBI.                                              taataatacagcgaatattatgcaggctttaaacatcaaaaaagagttcaacttctgact 22984
human.FAM230J-LOC105372942-GGTLC5P.NCBI.rev.compl                                   taataatacagcgaatattatgcaggctttaaacatcaaaaaagagttcaacttctgact 23700
human.FAM230A.ncRNA.18487127.18500594-GGTLC3.18516335.18518165.rev.compl.REF        taataatacagcgaatattatgcaggctttaaacatcaaaaaagagttcaacttctgact 12460
                                                                                    ************************************************************

chimp.LOC112206744-LOC107973052-GGT2.revcompl                                       tccgatgatggtgttgaagcaggtcactgctggtttacatttgattttcatgtgggaact 25666
chimp54546-135457.revcompl.Pan.troglodytes.clone.rp43-41g5.GenBank:AC099533.36      TCTGATGATGGTGTTGAAGCAGGTCACTGCTGGTTTACATTTGATTTTCATGTGGGAACT 27381
human.FAM30B-LOC105372935-GGT2.NCBI.                                                tccgatgatggtgttgaagcaggtcactgctggtttacatttgattttcatgtgggaact 24310
human.FAM230E-LOC105377182-GGT3P.NCBI.                                              tccgatgatggtgttgaagcaggtcactgctggtttacatttgattttcatgtgggaact 23044
human.FAM230J-LOC105372942-GGTLC5P.NCBI.rev.compl                                   tccgatgatggtgttgaagcaggtcactgctggtttacatttgattttcatgtgggaact 23760
human.FAM230A.ncRNA.18487127.18500594-GGTLC3.18516335.18518165.rev.compl.REF        tccgatgatggtgttgaagcaggtcactgctggtttacatttgattttcatgtgggaact 12520
                                                                                    ** *********************************************************

chimp.LOC112206744-LOC107973052-GGT2.revcompl                                       ctggaagtctgccttagtgattttacatgtggctaaattgagctaatgacaagctgttcg 25726
chimp54546-135457.revcompl.Pan.troglodytes.clone.rp43-41g5.GenBank:AC099533.36      CTGGAAGTCTGCCTTAGTGATTTTACATGTGGCTAAATTGAGCTAATGACAAGCTGTTCG 27441
human.FAM30B-LOC105372935-GGT2.NCBI.                                                ctggaagtccgccttagtgattttacatgtggctaaattgagctaatgacaagctgtttg 24370
human.FAM230E-LOC105377182-GGT3P.NCBI.                                              ctggaagtccgccttagtgattttacatgtggctaaattgagctaatgacaagctgttcg 23104
human.FAM230J-LOC105372942-GGTLC5P.NCBI.rev.compl                                   ctggaagtccgccttagtgattttacatgtggctaaattgagctaatgacaagctgttcg 23820
human.FAM230A.ncRNA.18487127.18500594-GGTLC3.18516335.18518165.rev.compl.REF        ctggaagtccgccttagtgattttacatgtggctaaattgagctaatgacaagccgttcg 12580
                                                                                    ********* ******************************************** *** *

chimp.LOC112206744-LOC107973052-GGT2.revcompl                                       aagtatggcaaaatggaactttaaaacagtatcttgtcagcaaccaagtggacctgtttc 25786
chimp54546-135457.revcompl.Pan.troglodytes.clone.rp43-41g5.GenBank:AC099533.36      AAGTATGGCAAAATGGAACTTTAAAACAGTATCTTGTCAGCAACCAAGTGGACCTGTTTC 27501
human.FAM30B-LOC105372935-GGT2.NCBI.                                                cagtatggcaaaatggaactttaaaacagtatcttgtcaacaaccaagaggacctgtttc 24430
human.FAM230E-LOC105377182-GGT3P.NCBI.                                              aagtatggcaaaatggaactttaaaacagtatcttgtcaacaaccaagaggacctgtttc 23164
human.FAM230J-LOC105372942-GGTLC5P.NCBI.rev.compl                                   aagtatggcaaaatggaactttaaaacagtatcttgtcaacaaccaagaggacctgtttc 23880
human.FAM230A.ncRNA.18487127.18500594-GGTLC3.18516335.18518165.rev.compl.REF        aagtatggcaaaatggaactttaaaacagtatcttgtcaacaaccaagaggacctgtttc 12640
                                                                                     ************************************** ******** ***********

chimp.LOC112206744-LOC107973052-GGT2.revcompl                                       acgtaaagcccatgcattcatctgcctgcccatcattctgtctgtccatcattctgtctg 25846
chimp54546-135457.revcompl.Pan.troglodytes.clone.rp43-41g5.GenBank:AC099533.36      ACGTAAAGCCCATGCATTCATCTGCCTGCCCATCATTCTGTCTGTCCATCATTCTGTCTG 27561
human.FAM30B-LOC105372935-GGT2.NCBI.                                                acataaagcccacgcattcatctgc----------------ctgtccatcattctgtctg 24474
human.FAM230E-LOC105377182-GGT3P.NCBI.                                              acataaagcccacgcattcatctgc----------------ctgtccatcattctgtctg 23208
human.FAM230J-LOC105372942-GGTLC5P.NCBI.rev.compl                                   acataaagcccacgcattcatctgc----------------ctgtccatcattctgtctg 23924
human.FAM230A.ncRNA.18487127.18500594-GGTLC3.18516335.18518165.rev.compl.REF        acataaagcccatgcattcatctgc----------------ctgtccatcattctgtctg 12684
                                                                                    ** ********* ************                *******************

chimp.LOC112206744-LOC107973052-GGT2.revcompl                                       tccacacgggcatcatttgttagtggaactgagtgcccactgtcgagctgacaagcccat 25906
chimp54546-135457.revcompl.Pan.troglodytes.clone.rp43-41g5.GenBank:AC099533.36      CCCACACGGGCATCATTTGTTAGTGGAACTGAGTGCCCACTGTCGAGCTGACAAGCCCAT 27621
human.FAM30B-LOC105372935-GGT2.NCBI.                                                tccacacgggcatcattcgttagtggaactgagtgcccgctgtcgagctgacaagcccat 24534
human.FAM230E-LOC105377182-GGT3P.NCBI.                                              tccacacgggcatcattcgttagtggaactgagtgcccgctgtcgagctgacaagcccat 23268
human.FAM230J-LOC105372942-GGTLC5P.NCBI.rev.compl                                   tccacacgggcatcattcgttagtggaactgagtgcccgctgtcgagctgacaagcccat 23984
human.FAM230A.ncRNA.18487127.18500594-GGTLC3.18516335.18518165.rev.compl.REF        tccacacgggcatcattcgttagtggaactgagtgcccactgtcgagctgacaagcccat 12744
                                                                                     *************** ******************** **********************

chimp.LOC112206744-LOC107973052-GGT2.revcompl                                       aacctccctgttcctagtcacacattaattcttcaacaagttccttttgatagattgtga 25966
chimp54546-135457.revcompl.Pan.troglodytes.clone.rp43-41g5.GenBank:AC099533.36      AACCTCCCTGTTCCTAGTCACACATTAATTCTTCAACAAGTCCCTTTTGATAGATTGTGA 27681
human.FAM30B-LOC105372935-GGT2.NCBI.                                                aacctccctgttcctagtcacacattaattcttcaacaagtcccttttgatagattgtga 24594
human.FAM230E-LOC105377182-GGT3P.NCBI.                                              aacctccctgttcctagtcacacattaattcttcaacaagtcccttttgatagattgtga 23328
human.FAM230J-LOC105372942-GGTLC5P.NCBI.rev.compl                                   aacctccctgttcctagtcacacattaattcttcaacaagtcccttttgatagattgtga 24044
human.FAM230A.ncRNA.18487127.18500594-GGTLC3.18516335.18518165.rev.compl.REF        aacctccctgttcctagtcacacattaattcttcaacaagtcccttttgatagattgtga 12804
                                                                                    ***************************************** ******************

chimp.LOC112206744-LOC107973052-GGT2.revcompl                                       ttaagcttagctgctatttccaattgcttccccaaacgtacttctcactgttctcccatc 26026
chimp54546-135457.revcompl.Pan.troglodytes.clone.rp43-41g5.GenBank:AC099533.36      TTAAGCTTAGCTACTATTTCCAATTGCTTCCCCAAACGTACTTCTCACTGTTCTCCCATC 27741
human.FAM30B-LOC105372935-GGT2.NCBI.                                                ttaagcttagctactatttccaattgcttccccaaacgtacttctcactgttctcccatc 24654
human.FAM230E-LOC105377182-GGT3P.NCBI.                                              ttaagcttagctactatttccaattgcttccccaaacgtacttctcactgttctcccatc 23388
human.FAM230J-LOC105372942-GGTLC5P.NCBI.rev.compl                                   ttaagcttagctactatttccaattgcttccccaaacgtacttctcactgttctcccatc 24104
human.FAM230A.ncRNA.18487127.18500594-GGTLC3.18516335.18518165.rev.compl.REF        ttaagcttagctactatttccaattgcttccccaaacgtacttctcactgttctcccatc 12864
                                                                                    ************ ***********************************************

chimp.LOC112206744-LOC107973052-GGT2.revcompl                                       acacccttcagcccatccatgcggggttcctttgcttttcccaccttacaccaaactccc 26086
chimp54546-135457.revcompl.Pan.troglodytes.clone.rp43-41g5.GenBank:AC099533.36      ACACCCTTCAGCCCATCCATGCGGGGTTCCTTTGCTTTTCCCACCTTACACCAAACTCCC 27801
human.FAM30B-LOC105372935-GGT2.NCBI.                                                acacccttcagcccatccatgcggggttcctttgcttttcccaccttacaccaaactccc 24714
human.FAM230E-LOC105377182-GGT3P.NCBI.                                              acacccttcagcccatccatgcggggttcctttgcttttcccaccttacaccaaactccc 23448
human.FAM230J-LOC105372942-GGTLC5P.NCBI.rev.compl                                   acacccttcagcccatccatgcggggttcctttgcttttcccaccttacaccaaactccc 24164
human.FAM230A.ncRNA.18487127.18500594-GGTLC3.18516335.18518165.rev.compl.REF        acacccttcagcccatccatgcggggttcctttgcttttcccaccttacaccaaactccc 12924
                                                                                    ************************************************************

chimp.LOC112206744-LOC107973052-GGT2.revcompl                                       tatttttactcccactttttacctcctctccaagacaaaacaaacaaaactagcatttaa 26146
chimp54546-135457.revcompl.Pan.troglodytes.clone.rp43-41g5.GenBank:AC099533.36      TATTTTTACTCCCACTTTTACCTCCTCTTCAAGACAAAACAAACAAAACTAGCATTTTAA 27861
```

```
human.FAM30B-LOC105372935-GGT2.NCBI.                                            tatttttactcccacttttacctcctctccaagacacaacaaacaaaactagcattttaa 24774
human.FAM230E-LOC105377182-GGT3P.NCBI.                                          tatttttactcccacttttacctcctctccaagacaaaacaaacaaaactagcattttaa 23508
human.FAM230J-LOC105372942-GGTLC5P.NCBI.rev.compl                               tatttttactcccacttttacctcctctccaagacaaaacaaacaaaactagcattttaa 24224
human.FAM230A.ncRNA.18487127.18500594-GGTLC3.18516335.18518165.rev.compl.REF    tatttttactcccacttttacctcctctccaagacaaaacaaacaaaactagcattttaa 12984
                                                                                **************************** ******* ***********************

chimp.LOC112206744-LOC107973052-GGT2.revcompl                                   aacttagttgtaatctttcttccttcatgaaaatttctccaacagccactcccacggtcc 26206
chimp54546-135457.revcompl.Pan.troglodytes.clone.rp43-41g5.GenBank:AC099533.36  AACTTAGTTGTAATCTTTCTTCCTTCATGAAAATTTCTCCAACAGCCACTCCCACGGTCC 27921
human.FAM30B-LOC105372935-GGT2.NCBI.                                            aacttagttgtaatctttcttccttcatgaaaatttctccaacagccactcccacggtcc 24834
human.FAM230E-LOC105377182-GGT3P.NCBI.                                          aacttagttgtaatctttcttccttcatgaaaatttctccaacagccactcccacggtcc 23568
human.FAM230J-LOC105372942-GGTLC5P.NCBI.rev.compl                               aacttagttgtaatctttcttccttcatgaaaatttctccaacagccactcccacggtcc 24284
human.FAM230A.ncRNA.18487127.18500594-GGTLC3.18516335.18518165.rev.compl.REF    aacttagttgtaatctttcttccttcatgaaaatttctccaacagccactcccacggtcc 13044
                                                                                ************************************************************

chimp.LOC112206744-LOC107973052-GGT2.revcompl                                   tgtgtgttccggatattttaaaataatggctataaggttgagcacttcaggatatgctgt 26266
chimp54546-135457.revcompl.Pan.troglodytes.clone.rp43-41g5.GenBank:AC099533.36  TGTGTCTTCCGGATATTTTAAAATAATGGCTATAAGGTTGAGCACTTCAGGATATGCTGT 27981
human.FAM30B-LOC105372935-GGT2.NCBI.                                            tgtgtgttccggatattttaaaataatggctataaggttgagcacttcaggatacgctgt 24894
human.FAM230E-LOC105377182-GGT3P.NCBI.                                          tgtgtgttccggatattttaaaataatggctataaggctgagcacttcaggatatgctgt 23628
human.FAM230J-LOC105372942-GGTLC5P.NCBI.rev.compl                               tgtgtgttccggatattttaaaataatggctataaggttgagcacttcaggatacgctgt 24344
human.FAM230A.ncRNA.18487127.18500594-GGTLC3.18516335.18518165.rev.compl.REF    tgtgtgttccagatattttaaaataatggctataaggctgagcacttcaggatatgctgt 13104
                                                                                ***** **** ************************** **************** *****

chimp.LOC112206744-LOC107973052-GGT2.revcompl                                   tttgctgtgtgcagatggaggcagtggctggagtgaatgaacggcaacacttgctggcaa 26326
chimp54546-135457.revcompl.Pan.troglodytes.clone.rp43-41g5.GenBank:AC099533.36  TTTGCTGTGTGCAGATGGAGGCAGTGGCTGGAGTGAATGAACGGCAACACTTGCTGGCAA 28041
human.FAM30B-LOC105372935-GGT2.NCBI.                                            tttgctgtgtgcagatggaggcagtggctggagtgaatgaacggcaacacttgctggcaa 24954
human.FAM230E-LOC105377182-GGT3P.NCBI.                                          tttgctgtgtgcagatggaggcagtggctggagtgaatgaacggcaacacttgctggcaa 23688
human.FAM230J-LOC105372942-GGTLC5P.NCBI.rev.compl                               tttgctgtgtgcagatggaggcagtggctggagtgaatgaacggcaacacttgctggcaa 24404
human.FAM230A.ncRNA.18487127.18500594-GGTLC3.18516335.18518165.rev.compl.REF    tttgctgtgtgcagatggaggcagtggctggagtgaatgaacggcaacacttgctggcaa 13164
                                                                                ************************************************************

chimp.LOC112206744-LOC107973052-GGT2.revcompl                                   ccggcagaagctgagagacagggaacaggctctcctccagagcctccaggagccaggcct 26386
chimp54546-135457.revcompl.Pan.troglodytes.clone.rp43-41g5.GenBank:AC099533.36  CCGGCAGAAGCTGAGAGACAGGGAACAGGCTCTCCTCCAGAGCCTCCAGGAGCCAGGCCT 28101
human.FAM30B-LOC105372935-GGT2.NCBI.                                            ccggcagaagctgagagacagggaacaggctctcctccagagcctccaggagccaggcct 25014
human.FAM230E-LOC105377182-GGT3P.NCBI.                                          ccggcagaagctgagagacagggaacaggctctcctccagagcctccaggagccaggcct 23748
human.FAM230J-LOC105372942-GGTLC5P.NCBI.rev.compl                               ccggcagaagctgagagacagggaacaggctctcctccagagcctccaggagccaggcct 24464
human.FAM230A.ncRNA.18487127.18500594-GGTLC3.18516335.18518165.rev.compl.REF    ccggcagaagctgagagacagggaacaggctctcctccagagcctccaggagccaggcct 13224
                                                                                ************************************************************

chimp.LOC112206744-LOC107973052-GGT2.revcompl                                   ttggacaccttgaatgtgggcttctgggagaccatgcgtttctgttataagcagcccagt 26446
chimp54546-135457.revcompl.Pan.troglodytes.clone.rp43-41g5.GenBank:AC099533.36  TTGGACACCTTGAATGTGGGCTTCTGGGAGACCATGCGTTTCTGTTATAAGCAGCCCAGT 28161
human.FAM30B-LOC105372935-GGT2.NCBI.                                            ttggacaacttgaatgtgggcttctgggagaccatgcgtttctgttataagcagcccagt 25074
human.FAM230E-LOC105377182-GGT3P.NCBI.                                          ttggacaccttgaatgtgggcttctgggagaccatgcgtttctgttataagcagcccagt 23808
human.FAM230J-LOC105372942-GGTLC5P.NCBI.rev.compl                               ttggacaccttgaatgtgggcttctgggagaccatgcgtttctgttataagcagcccagt 24524
human.FAM230A.ncRNA.18487127.18500594-GGTLC3.18516335.18518165.rev.compl.REF    ttggacaccttgaatgtgggcttctgggagaccatgcgtttctgttataagcagcccagt 13284
                                                                                ******* ****************************************************

chimp.LOC112206744-LOC107973052-GGT2.revcompl                                   cgctggcagtttttacggctgcccccgaacactcatctatacctgtctgacaaggtcaag 26506
chimp54546-135457.revcompl.Pan.troglodytes.clone.rp43-41g5.GenBank:AC099533.36  CTCTGGCAGTTTTTACGGCTGCCCCCGAACACTCATCTATACCTGTCTGACCAGGTCAAG 28221
human.FAM30B-LOC105372935-GGT2.NCBI.                                            ctctggcagtttttacggctgccccggaacactcatctatacctgtctgacaaggtcaag 25134
human.FAM230E-LOC105377182-GGT3P.NCBI.                                          ctctggcagtttttacggctgccccggaacactcatctatacctgtctgacaaggtcaag 23868
human.FAM230J-LOC105372942-GGTLC5P.NCBI.rev.compl                               ctctggcagtttttacggctgccccggaacactcatctatacctgtctgacaaggtcaag 24584
human.FAM230A.ncRNA.18487127.18500594-GGTLC3.18516335.18518165.rev.compl.REF    ctctggcagtttttacggctgccccggaacactcatctatacctgtctgacaaggtcaag 13344
                                                                                * *********************** ************************* ********

chimp.LOC112206744-LOC107973052-GGT2.revcompl                                   ctccaaggaagggattctctacatatctacattgtttgcagattttacaataatcattta 26566
chimp54546-135457.revcompl.Pan.troglodytes.clone.rp43-41g5.GenBank:AC099533.36  CTCCAAGGAAGGGATTCTCTACATATCTACATTGTTTGCAGATTTTACAATAATCATTTA 28281
human.FAM30B-LOC105372935-GGT2.NCBI.                                            ctccaaggaagggactctctacatatctacattgtttgcagattttacaataatcattta 25194
human.FAM230E-LOC105377182-GGT3P.NCBI.                                          ctccaaggaagggattctttacatatctacattgtttgcagattttacaataatcattta 23928
human.FAM230J-LOC105372942-GGTLC5P.NCBI.rev.compl                               ctccaaggaagggactctctacatatctacattgtttgcagattttacaataatcattta 24644
human.FAM230A.ncRNA.18487127.18500594-GGTLC3.18516335.18518165.rev.compl.REF    ctccaaggaagggattctctacatatctacattgtttgcagattttacaataatcattta 13404
                                                                                ************* *** ******************************************

chimp.LOC112206744-LOC107973052-GGT2.revcompl                                   ttcttgcatggctgatcattgttaaccaatacaaataaaataataaagaaatgacccaca 26626
chimp54546-135457.revcompl.Pan.troglodytes.clone.rp43-41g5.GenBank:AC099533.36  TTCTTGCATGGCTGATCATTGTTAACCAATACAAATAAAATAATAAAGAAATGACCCACA 28341
human.FAM30B-LOC105372935-GGT2.NCBI.                                            ttcttgcatggctgatcattgttaaccaatacaaataaaataataaagaaatgacccaca 25254
human.FAM230E-LOC105377182-GGT3P.NCBI.                                          ttcttgcatggctgatcattgttaaccaatacaaataaaataataaagaaatgacccaca 23988
human.FAM230J-LOC105372942-GGTLC5P.NCBI.rev.compl                               ttcttgcatggctgatcattgttaaccaatacaaataaaataataaagaaatgacccaca 24704
human.FAM230A.ncRNA.18487127.18500594-GGTLC3.18516335.18518165.rev.compl.REF    ttcttgcatggctgatcattgttaaccaatacaaataaaataataaagaaatgacccaca 13464
                                                                                ************************************************************

chimp.LOC112206744-LOC107973052-GGT2.revcompl                                   ttttatgttgggagtttgatctgccatttatcaagtatggaatcttgaacaaggggttaa 26686
chimp54546-135457.revcompl.Pan.troglodytes.clone.rp43-41g5.GenBank:AC099533.36  TTTTATGTTGGGAGTTTGATCTGCCATTTATCAAGTATGGAATCTTGAACAAGGGGTTAA 28401
human.FAM30B-LOC105372935-GGT2.NCBI.                                            ttttatgttgggagtttgatctgccatttatcaagtatggaatcttgaacaaggggttaa 25314
human.FAM230E-LOC105377182-GGT3P.NCBI.                                          ttttatgttgggagtttgatctgccatttatcaagtatggaatcttgaacaaggggttaa 24048
human.FAM230J-LOC105372942-GGTLC5P.NCBI.rev.compl                               ttttatgttgggagtttgatctgccatttatcaagtatggaatcttgaacaaggggttaa 24764
human.FAM230A.ncRNA.18487127.18500594-GGTLC3.18516335.18518165.rev.compl.REF    ttttatgttgggagtttgatctgccatttatcaagtatggaatcttgaacaaggggttaa 13524
                                                                                ************************************************************

chimp.LOC112206744-LOC107973052-GGT2.revcompl                                   acatctgaatgtctccatcacttcatctctaaagtgggggtgctcacacccactgggctc 26746
chimp54546-135457.revcompl.Pan.troglodytes.clone.rp43-41g5.GenBank:AC099533.36  ACATCTGAATGTCTCCATCACTTCATCTCTAAAGTGGGGGTGCTCACACCCACTGGGCTC 28461
human.FAM30B-LOC105372935-GGT2.NCBI.                                            acatctgaatgtctccatcacttcatctctaaagtgggggtgctcacacccactgggctc 25374
human.FAM230E-LOC105377182-GGT3P.NCBI.                                          acatctgaatgtctccatcacttcatctctaaagtgggggtgctcacacccactgggctc 24108
human.FAM230J-LOC105372942-GGTLC5P.NCBI.rev.compl                               acatctgaatgtctccatcacttcatctctaaagtgggggtgctcacacccactgggctc 24824
human.FAM230A.ncRNA.18487127.18500594-GGTLC3.18516335.18518165.rev.compl.REF    acatctgaatgtctccatcacttcatctctaaagtgggggtgctcacacccactgggctc 13584
                                                                                ************************************************************

chimp.LOC112206744-LOC107973052-GGT2.revcompl                                   cccct-ccaggttggtgccggactctccctgggccccccctgttctctcaccagccacatc 26805
chimp54546-135457.revcompl.Pan.troglodytes.clone.rp43-41g5.GenBank:AC099533.36  CCC--CCCAGGTTGGTGCCGGACTCTCCCTGGGCCCCCCTGTTCTCTCACCAGCCACATC 28519
human.FAM30B-LOC105372935-GGT2.NCBI.                                            ccccgcccaggttggtgccggactctccctgggcccccctgttctctcaccagccacatc 25434
human.FAM230E-LOC105377182-GGT3P.NCBI.                                          ccccgcccaggttggtgccggactctccctgggcccccctgttctctcaccagccacatc 24168
human.FAM230J-LOC105372942-GGTLC5P.NCBI.rev.compl                               ccccgcccaggttggtgccggactctccctgggcccccctgttctctcaccagccacatc 24884
human.FAM230A.ncRNA.18487127.18500594-GGTLC3.18516335.18518165.rev.compl.REF    ccccacccaggttggtgccggactctccctgggcccccctgttctctcaccagccacatc 13644
                                                                                ***   ******************************************************

chimp.LOC112206744-LOC107973052-GGT2.revcompl                                   cattctcccccagaggcgctagtgactgtgcgtggctttccattcccaccacgtttgtc 26865
chimp54546-135457.revcompl.Pan.troglodytes.clone.rp43-41g5.GenBank:AC099533.36  CATTCTCCCCCAGAGGTGCTAGTGACTGTGCGTGGCTTTCCATTCCCACCACGTTTGTC 28579
human.FAM30B-LOC105372935-GGT2.NCBI.                                            cattctcccccagaggcgctagtgactgtgcgtggctttccattcccaccacgtttgtc 25494
human.FAM230E-LOC105377182-GGT3P.NCBI.                                          cattctcccccagaggcgctagtgactgtgcgtggctttccattcccaccacgtttgtc 24228
human.FAM230J-LOC105372942-GGTLC5P.NCBI.rev.compl                               cattctcccccagaggcgctagtgactgtgcgtggctttccattcccaccacgtttgtc 24944
human.FAM230A.ncRNA.18487127.18500594-GGTLC3.18516335.18518165.rev.compl.REF    cattctcccccagaggcgctagtgactgtgcgtggctttccattcccaccatgtttgtc 13704
                                                                                ***************** ********************************** *******

chimp.LOC112206744-LOC107973052-GGT2.revcompl                                   tctaaccccagtggcagatcagtgtaagaacacagctgagtgctcctcgcctccttgccc 26925
chimp54546-135457.revcompl.Pan.troglodytes.clone.rp43-41g5.GenBank:AC099533.36  TCTAACCCCAGTGGCAGATCAGTGTAAGAACACAGCTGAGTGCTCCTCGCCTCCTTGCCC 28639
human.FAM30B-LOC105372935-GGT2.NCBI.                                            tctaaccccagtggcagatcagtgtaagaacacagctgagtgctcctcgcctccttgccc 25554
human.FAM230E-LOC105377182-GGT3P.NCBI.                                          tctaaccccagtggcagatcagtgtaagaacacagctgagtgctcctcgcctccttgccc 24288
human.FAM230J-LOC105372942-GGTLC5P.NCBI.rev.compl                               tctaaccccagtggcagatcagtgtaagaacacagctgagtgctcctcgcctccttgccc 25004
human.FAM230A.ncRNA.18487127.18500594-GGTLC3.18516335.18518165.rev.compl.REF    tctaaccccagtggcagatcagtgtaagaacacagctgagtgctcctcgcctccttgccc 13764
                                                                                ************************************************************

chimp.LOC112206744-LOC107973052-GGT2.revcompl                                   cttcaagggctcctcaccacccaccagatcaggtgcaaacttccaagccttactggatcc 26985
chimp54546-135457.revcompl.Pan.troglodytes.clone.rp43-41g5.GenBank:AC099533.36  CTTCAAGGGCTCCTCACCACCCACCAGATCAGGTGCAAACTTCCAAGCCTTACTGGATCC 28699
human.FAM30B-LOC105372935-GGT2.NCBI.                                            cttcaagggcccctcaccacccaccagatcaggtgcaaacttccaagccttactggatcc 25614
human.FAM230E-LOC105377182-GGT3P.NCBI.                                          cttcaagggctcctcaccacccaccagatcaggtgcaaacttccaagccttactggatcc 24348
human.FAM230J-LOC105372942-GGTLC5P.NCBI.rev.compl                               cttcaagggctcctcaccacccaccagatcaggtgcaaacttccaagccttactggatcc 25064
human.FAM230A.ncRNA.18487127.18500594-GGTLC3.18516335.18518165.rev.compl.REF    cttcaagggctcctcaccacccaccagatcaggtgcaaacttccaagccttactggatcc 13824
                                                                                ********** *************************************************

chimp.LOC112206744-LOC107973052-GGT2.revcompl                                   ccttccacattctgagctccgcctgccttcccatcgctatccttccccacctgcctccct 27045
chimp54546-135457.revcompl.Pan.troglodytes.clone.rp43-41g5.GenBank:AC099533.36  CCTTCCACATTCTGAACTCCGCCTGCCTTCCCATCGCTATCCTTCCCCACCTGCCTCCCT 28759
human.FAM30B-LOC105372935-GGT2.NCBI.                                            ccttccacattctgagctccgcctgccttcccatcgctatccttccccacctgcctccct 25674
human.FAM230E-LOC105377182-GGT3P.NCBI.                                          ctttccacattctgagctccgcctgccttcccatcgctatccttccccacctgcctccct 24408
human.FAM230J-LOC105372942-GGTLC5P.NCBI.rev.compl                               ccttccacattctgagctccgcctgccttcccatcgctatccttccccacctgcctccct 25124
human.FAM230A.ncRNA.18487127.18500594-GGTLC3.18516335.18518165.rev.compl.REF    ccttccacattctgagctccgcctgccttcccatcgctatccttccccacctgcctccct 13884
                                                                                * ************* ********************************************

chimp.LOC112206744-LOC107973052-GGT2.revcompl                                   ggtagagaaaagcagagtgtgtgatgctgtctgaatgctgagcacggccttttgcagcca 27105
chimp54546-135457.revcompl.Pan.troglodytes.clone.rp43-41g5.GenBank:AC099533.36  GGTAGAGAAAAGCAGAGTGTGTGATGCTGTCTGAATGCTGAGCACGGCCTTTTGCAGCCA 28819
human.FAM30B-LOC105372935-GGT2.NCBI.                                            ggtagagaaaagcggagtgtgtgatgctgtctgaatgctgagcacggccttttgcagcca 25734
human.FAM230E-LOC105377182-GGT3P.NCBI.                                          ggtagagaaaagcagagtgtgtgatgctgtctgaatgctgagcacggccttttgcagcca 24468
human.FAM230J-LOC105372942-GGTLC5P.NCBI.rev.compl                               ggtagagaaaagcggagtgtgtgatgctgtctgaatgctgagcacggccttttgcagcca 25184
human.FAM230A.ncRNA.18487127.18500594-GGTLC3.18516335.18518165.rev.compl.REF    ggtagagaaaagcagagtgtgtgatgctgtctgaatgctgagcacggccttttgcagcca 13944
                                                                                ************* **********************************************

chimp.LOC112206744-LOC107973052-GGT2.revcompl                                   gtccactgtgtacgctgcccctatcggagacctccaccttaaccctttccagcctggagg 27165
chimp54546-135457.revcompl.Pan.troglodytes.clone.rp43-41g5.GenBank:AC099533.36  GTCCACTGTGGACGCTGCCCCTATCGGAGACCTCCACCTTAACCCTTTCCAGCCTGGAGG 28879
human.FAM30B-LOC105372935-GGT2.NCBI.                                            gtccactgtgtacgctgcccctatcggagacctccaccttaaccctttccagcctggagg 25794
human.FAM230E-LOC105377182-GGT3P.NCBI.                                          gtccactgtgtacgctgcccctatcggagacctccaccttaaccctttccagcctgtagg 24528
human.FAM230J-LOC105372942-GGTLC5P.NCBI.rev.compl                               gtccactgtgtacgctgcccctatcggagacctccaccttaagcctttccagcctggagg 25244
human.FAM230A.ncRNA.18487127.18500594-GGTLC3.18516335.18518165.rev.compl.REF    gtccactgtgtatgctgcccctatcggagacctccaccttaaccctttccagcctggagg 14004
                                                                                ********** * ***************************** ************* ***

chimp.LOC112206744-LOC107973052-GGT2.revcompl                                   ctcctcccagggccccacaacagaagtgacttcccttgcctttgaatttctatagcacaa 27225
chimp54546-135457.revcompl.Pan.troglodytes.clone.rp43-41g5.GenBank:AC099533.36  CTCCTCCCAGGGCCCCACAACAGAAGTGACTTCCCTTGCCTTTGAATTTCTATAGCACAA 28939
human.FAM30B-LOC105372935-GGT2.NCBI.                                            ctcctcccagggccccacaacagaagtgacttcccttgcctttgaatttctatagcacaa 25854
```

```
human.FAM230E-LOC105377182-GGT3P.NCBI.                                      ctcctcccagggccccacaacagaagtgacttcccttgcctttgaatttctatagcacaa 24588
human.FAM230J-LOC105372942-GGTLC5P.NCBI.rev.compl                           ctcctcccagggccccacaacagaagtgacttcccttgcctttgaatttctatagcacaa 25304
human.FAM230A.ncRNA.18487127.18500594-GGTLC3.18516335.18518165.rev.compl.REF ctcctcccagggccccacaacagaagtgacttcccttgcctttgaatttctatagcacaa 14064
                                                                            ************************************************************

chimp.LOC112206744-LOC107973052-GGT2.revcompl                               tccctactgcccccgttaaaactgcaaagtccttttgtggaaaataactttattcatga 27285
chimp54546-135457.revcompl.Pan.troglodytes.clone.rp43-41g5.GenBank:AC099533.36 TCCCTACTGCCCCCGTTAAAACTGCAAAGTCCTTTTGTGGAAAATAACTTTATTCATGA 28999
human.FAM30B-LOC105372935-GGT2.NCBI.                                        tccctactgcccccgttaaaactgcaaagtccttttgtggaaaataactttattcatga 25914
human.FAM230E-LOC105377182-GGT3P.NCBI.                                      gccctactgcccccgttaaaactgcaaagtccttttgtggaaaataactttattcatga 24648
human.FAM230J-LOC105372942-GGTLC5P.NCBI.rev.compl                           gccctactgcccccgttaaaactgcaaagtccttttgtggaaaataactttattcatga 25364
human.FAM230A.ncRNA.18487127.18500594-GGTLC3.18516335.18518165.rev.compl.REF gccctactgcccccgttaaaactgcaaagtccttttgtggaaaataactttattcatga 14124
                                                                             ***********************************************************

chimp.LOC112206744-LOC107973052-GGT2.revcompl                               ctgtgtttatcacactatcttatggagaagagatgatcaataaatatttgcggaataaat 27345
chimp54546-135457.revcompl.Pan.troglodytes.clone.rp43-41g5.GenBank:AC099533.36 CTGTGTTTATCACACTATCTTATGGAGAAGAGATGATCAATAAATATTTGCTGAATAAAT 29059
human.FAM30B-LOC105372935-GGT2.NCBI.                                        ctgtgtttatcacactatcttatggagaagagatgatcaataaatatttgctgaataaat 25974
human.FAM230E-LOC105377182-GGT3P.NCBI.                                      ctgtgtttatcacactatcttatggagaagagatgatcaataaatatttgctgaataaat 24708
human.FAM230J-LOC105372942-GGTLC5P.NCBI.rev.compl                           ctgtgtttatcacactatcttatggagaagagatgatcaataaatatttgctgaataaat 25424
human.FAM230A.ncRNA.18487127.18500594-GGTLC3.18516335.18518165.rev.compl.REF ctgtgtttatcacactatcttatggagaagagatgatcaataaatatttgttgaataaat 14184
                                                                            **************************************************  ********

chimp.LOC112206744-LOC107973052-GGT2.revcompl                               gaatagcagttacaaaacacttgattcatatggaattaatgttggttctcaaagtgaaaa 27405
chimp54546-135457.revcompl.Pan.troglodytes.clone.rp43-41g5.GenBank:AC099533.36 GAATAGCAGTTACAAAACACTTGATTCATATGGAATTAATGTTGGTTCTCAAAGTGAAAA 29119
human.FAM30B-LOC105372935-GGT2.NCBI.                                        gaatagcagttacaaaacacttgattcatatggaattaatgttggttctcaaagtgaaaa 26034
human.FAM230E-LOC105377182-GGT3P.NCBI.                                      gaatagcagttacaaaacactagattcatatggaattaatgttggttctcaaagtgaaaa 24768
human.FAM230J-LOC105372942-GGTLC5P.NCBI.rev.compl                           gaatagcagttacaaaacacttgattcatatggaattaatgttggttctcaaagtgaaaa 25484
human.FAM230A.ncRNA.18487127.18500594-GGTLC3.18516335.18518165.rev.compl.REF gaatagcagttacaaaacacttgattcatatggaattaatgttggttctcaaagtgaaaa 14244
                                                                            ******************** ***************************************

chimp.LOC112206744-LOC107973052-GGT2.revcompl                               attacaaacagcactgatattcagccagtatacaagtctggtcacagcagttgtataata 27465
chimp54546-135457.revcompl.Pan.troglodytes.clone.rp43-41g5.GenBank:AC099533.36 ATTACAAACAGCACTGATATTCAGCCAGTATACAAGTCTGGTAACAGCAGTTGTATAATA 29179
human.FAM30B-LOC105372935-GGT2.NCBI.                                        attacaaacagcactgatattcagccagtatacaagtctggtcacagcagttgtataata 26094
human.FAM230E-LOC105377182-GGT3P.NCBI.                                      attacaaacagcactgatattcagccagtatacaagtctggtcacagcagttgtataata 24828
human.FAM230J-LOC105372942-GGTLC5P.NCBI.rev.compl                           attacaaacagcactgatattcagccagtatacaagtctggtcacagcagttgtataata 25544
human.FAM230A.ncRNA.18487127.18500594-GGTLC3.18516335.18518165.rev.compl.REF attacaaacagcactgatattcagccagtatacaagtctggtcacagcagttgtataata 14304
                                                                            ****************************************** *****************

chimp.LOC112206744-LOC107973052-GGT2.revcompl                               ctgaatacccccctgccactgacctttggcccccagatgcctcccactgccactgctctc 27525
chimp54546-135457.revcompl.Pan.troglodytes.clone.rp43-41g5.GenBank:AC099533.36 CTGAATACCCCCCTGCCACTGACCTTTGGCCCCCAGATGCCTCCCACTGCCACTGCTCTC 29239
human.FAM30B-LOC105372935-GGT2.NCBI.                                        ctgaatacccccctgccactgacctttggcccccagatgcctcccactgccactgctctc 26154
human.FAM230E-LOC105377182-GGT3P.NCBI.                                      ctgaatacccccctgccactgacctttggcccccagatgcctcccactgccactgctctc 24888
human.FAM230J-LOC105372942-GGTLC5P.NCBI.rev.compl                           ctgaataccccccgccactgacctttggcccccagatgcctcccactgccactgctctc 25604
human.FAM230A.ncRNA.18487127.18500594-GGTLC3.18516335.18518165.rev.compl.REF ctgaatacccccctgccactgacctttggcccccagatgcctcccactgccactgctctc 14364
                                                                            ************* **********************************************

chimp.LOC112206744-LOC107973052-GGT2.revcompl                               cccactgggaacccctgaagttcccacaggctcataactaaagggctaatgtcctgcaca 27585
chimp54546-135457.revcompl.Pan.troglodytes.clone.rp43-41g5.GenBank:AC099533.36 CCCACTGGGAACCCCTGAAGTTCCCACAGGCTCATAACTAAAGGGCTAATGTCTTGCACA 29299
human.FAM30B-LOC105372935-GGT2.NCBI.                                        cccactgggaacccctgaagttcccacaggctcataactaaagggctaatgtcttgcaca 26214
human.FAM230E-LOC105377182-GGT3P.NCBI.                                      cccactgggaacccctgaagttcccacaggctcataactaaagggctaatgtctcgcaca 24948
human.FAM230J-LOC105372942-GGTLC5P.NCBI.rev.compl                           cccactgggaacccctgaagttcccacaggctcataactaaagggctaatgtctcgcaca 25664
human.FAM230A.ncRNA.18487127.18500594-GGTLC3.18516335.18518165.rev.compl.REF cccactgggaacccctgaagttcccacaggctcataactaaagggctaatgtctcgcaca 14424
                                                                            *****************************************************  *****

chimp.LOC112206744-LOC107973052-GGT2.revcompl                               gcagcgagcacccaggaccgagcagccacatggccgggtctgctggtgaaagcatccatt 27645
chimp54546-135457.revcompl.Pan.troglodytes.clone.rp43-41g5.GenBank:AC099533.36 GCAGCGAGCACCCAGGACCGAGCAGCCACATGGCTGGGTCTGCTGGTGAAAGCATCCATT 29359
human.FAM30B-LOC105372935-GGT2.NCBI.                                        gcagcgagcacccaggaccgagcagccacatggccgggtctgctggtgaaagcatccatt 26274
human.FAM230E-LOC105377182-GGT3P.NCBI.                                      gcagcgagcacccaggaccgagcagccacatggccgggtctgctggtgaaagcatccatt 25008
human.FAM230J-LOC105372942-GGTLC5P.NCBI.rev.compl                           gcagcgagcacccaggaccgagcagccacatggccgggtctgctggtgaaagcatccatt 25724
human.FAM230A.ncRNA.18487127.18500594-GGTLC3.18516335.18518165.rev.compl.REF gcagcgagcacccaggaccgagcagccacatggccgggtctgctggtgaaagcatccatt 14484
                                                                            ********************************** *************************

chimp.LOC112206744-LOC107973052-GGT2.revcompl                               ctgactgatcaggacctgaggggcctcatggttacatattttgataatatccctaattat 27705
chimp54546-135457.revcompl.Pan.troglodytes.clone.rp43-41g5.GenBank:AC099533.36 CTGACTGATCAGGACCTGAGGGGCCTCATGGTTACATATTTTGATAATATCCCTAATTAT 29419
human.FAM30B-LOC105372935-GGT2.NCBI.                                        ctgactgatcaggacctgaggggcctcatggttacatattttgataatatccctaattat 26334
human.FAM230E-LOC105377182-GGT3P.NCBI.                                      ctgactgatcaggacctgaggggcctcatggttacatattttgataatatccctaattat 25068
human.FAM230J-LOC105372942-GGTLC5P.NCBI.rev.compl                           ctgactgatcaggacctgaggggcctcatggttacatattttgataatatccctaattat 25784
human.FAM230A.ncRNA.18487127.18500594-GGTLC3.18516335.18518165.rev.compl.REF ctgactgatcaggacctgaggggcctcatggttacatattttgataatatccctaattat 14544
                                                                            ************************************************************

chimp.LOC112206744-LOC107973052-GGT2.revcompl                               aaataaggctcagttatatagtttgaaaacaatgcttctcctcattgcaaaatctcttag 27765
chimp54546-135457.revcompl.Pan.troglodytes.clone.rp43-41g5.GenBank:AC099533.36 AAATAAGGCTCAGTTATATAGTTTGAAAACAATGCTTCTCCTCATTGCAAAATCTCTTAG 29479
human.FAM30B-LOC105372935-GGT2.NCBI.                                        aaataaggctcagttatatagtttgaaaacagtgcttctcctcattgcaaaatctcttag 26394
human.FAM230E-LOC105377182-GGT3P.NCBI.                                      aaataaggctcagttatatagtttgaaaacaatgcttctcctcattgcaaaatctcttag 25128
human.FAM230J-LOC105372942-GGTLC5P.NCBI.rev.compl                           aaataaggctcagttatatagtttgaaaacaatgcttctcctcattgcaaaatctcttag 25844
human.FAM230A.ncRNA.18487127.18500594-GGTLC3.18516335.18518165.rev.compl.REF aaataaggctcagttatatagtttgaaaacaatgcttctcctcatcgcaaaatctcttag 14604
                                                                            ******************************* ************* **************

chimp.LOC112206744-LOC107973052-GGT2.revcompl                               aagactccgtagatccaggaacggaaatggaaaatgacagcatgtcaatctctgaaggtt 27825
chimp54546-135457.revcompl.Pan.troglodytes.clone.rp43-41g5.GenBank:AC099533.36 AAGACTCCGTAGATCCAGGAACGGAAATGGAAAATGACAGCGTGTCAATCTCTGAAGGTT 29539
human.FAM30B-LOC105372935-GGT2.NCBI.                                        aagactccgtagatccaggaacggaaatggaaaatgacagcgtgtcaatctctgaaggtt 26454
human.FAM230E-LOC105377182-GGT3P.NCBI.                                      aagactccgtagatccaggaacggaaatggaaaatgacagcgtgtcaatctctgaaggtt 25188
human.FAM230J-LOC105372942-GGTLC5P.NCBI.rev.compl                           aagactccgtagatccaggaacggaaatggaaaatgacagcgtgtcaatctctgaaggtt 25904
human.FAM230A.ncRNA.18487127.18500594-GGTLC3.18516335.18518165.rev.compl.REF aagactccgtagatccaggaacggaaatggaaaatgacagcgtgtcaatctctgaaggtt 14664
                                                                            ***************************************** ******************

chimp.LOC112206744-LOC107973052-GGT2.revcompl                               ttgggcatttccattagcactccatcttcatgtaaaccagaagatatgcagtttcctgcc 27885
chimp54546-135457.revcompl.Pan.troglodytes.clone.rp43-41g5.GenBank:AC099533.36 TTGAGCATTTCCATTAGCACTCCATCTTCATGTAAACCAGAAGATACGCAGTTTCCTGCC 29599
human.FAM30B-LOC105372935-GGT2.NCBI.                                        ttgggcatttccattagcactccatcttcatgtaaaccagaagatatgcagtttcctgcc 26514
human.FAM230E-LOC105377182-GGT3P.NCBI.                                      ttgggcatttccattagcactccatcttcatgtaaaccagaagatatgcagtttcctgcc 25248
human.FAM230J-LOC105372942-GGTLC5P.NCBI.rev.compl                           ttgggcatttccattagcactccatcttcatgtaaaccagaagatatgcagtttcctgcc 25964
human.FAM230A.ncRNA.18487127.18500594-GGTLC3.18516335.18518165.rev.compl.REF ttgggcatttccattagcactccatcttcatgtaaaccagaagatatgcagtttcctgcc 14724
                                                                            *** ****************************************** *************

chimp.LOC112206744-LOC107973052-GGT2.revcompl                               tagagagaagagaagacacatcagcacagcggcatgaaaccttcatcagaaaacaatgct 27945
chimp54546-135457.revcompl.Pan.troglodytes.clone.rp43-41g5.GenBank:AC099533.36 TAGAGAGAAGAGAAGACACATCAGCACAGTGGCATGAAATCTTCATCAGAAAACAATGCT 29659
human.FAM30B-LOC105372935-GGT2.NCBI.                                        tagagagaagagaagacacatcagcacagcggcatgaaaccttcatcagaaaacaatgct 26574
human.FAM230E-LOC105377182-GGT3P.NCBI.                                      tagagagaagagaagacacatcagcacagcggcatgaaaccttcatcagaaaacaatgct 25308
human.FAM230J-LOC105372942-GGTLC5P.NCBI.rev.compl                           tagagagaagagaagacacatcagcacagcggcatgaaaccttcatcagaaaacaatgct 26024
human.FAM230A.ncRNA.18487127.18500594-GGTLC3.18516335.18518165.rev.compl.REF tagagagaagagaagacacatcagcacagcggcatgaaagcttcatcagaaaacaatgct 14784
                                                                            ***************************** ********* ********************

chimp.LOC112206744-LOC107973052-GGT2.revcompl                               tcattaatccttgacaggacaagcgtcagcaaacttccaggccgctggattaagccttca 28005
chimp54546-135457.revcompl.Pan.troglodytes.clone.rp43-41g5.GenBank:AC099533.36 TCATTAATCCGTGACAGGACAAGTGTCAGCAAACTTCCAGGCCGCTGGATTAAGCCTTCA 29719
human.FAM30B-LOC105372935-GGT2.NCBI.                                        tcattaatccgtgacaggacaagcgtcagcaaacttccaggcggctggattaggccttca 26634
human.FAM230E-LOC105377182-GGT3P.NCBI.                                      tcattaatccgtgacaggacaagcgtcagcaaacttccaggcggctggattaggccttca 25368
human.FAM230J-LOC105372942-GGTLC5P.NCBI.rev.compl                           tcattaatccgtgacaggacaagcgtcagcaaacttccaggcggctggataaggccttca 26084
human.FAM230A.ncRNA.18487127.18500594-GGTLC3.18516335.18518165.rev.compl.REF tcattaatccgtgacaggacaagcgtcagcaaacttccaggcggctggattaggccttca 14844
                                                                            ********** ************ ****************** ******* * *******

chimp.LOC112206744-LOC107973052-GGT2.revcompl                               tctatccatcaccttggagaggaacaaaataggtggcctgggaagataagcactatgttt 28065
chimp54546-135457.revcompl.Pan.troglodytes.clone.rp43-41g5.GenBank:AC099533.36 TCTATCCATCACCTTGGAGAGGAACAAAATAGGTGGCCTGGGAAGATAAGCACTATGTTT 29779
human.FAM30B-LOC105372935-GGT2.NCBI.                                        tctatccatcaccttggagaggaacaaaataggtggcctgggaagataagcactatgttt 26694
human.FAM230E-LOC105377182-GGT3P.NCBI.                                      tctatccatcaccttggagaggaacaaaataggtggcctgggaagttaagcactatgttt 25428
human.FAM230J-LOC105372942-GGTLC5P.NCBI.rev.compl                           tctatccatcaccttggagaggaacaaaataggtggcctgggaagttaagcactatgttt 26144
human.FAM230A.ncRNA.18487127.18500594-GGTLC3.18516335.18518165.rev.compl.REF tctatccatcaccttggagaggaacaaaataggtggcctgggaagataagcactatgttt 14904
                                                                            ********************************************* **************

chimp.LOC112206744-LOC107973052-GGT2.revcompl                               ctattagttaatatctaaagcggaggttaacaagctatggacacacaagccaaacccagc 28125
chimp54546-135457.revcompl.Pan.troglodytes.clone.rp43-41g5.GenBank:AC099533.36 CTATTAGTTAATATCTAAAGCGGAGGTTAACAAGCTATGGACACACAAGCCAAACCCAGC 29839
human.FAM30B-LOC105372935-GGT2.NCBI.                                        ctattagttaatatctaaagcggaggttaacaagctatggacacacaagccaaacccagc 26754
human.FAM230E-LOC105377182-GGT3P.NCBI.                                      ctattagttaatatctaaagcggaggttaacaagctatggacacacaagccaaacccagc 25488
human.FAM230J-LOC105372942-GGTLC5P.NCBI.rev.compl                           ctattcgttaatatctaaagcggaggttaacaagctatggacacacaagccaaacccagc 26204
human.FAM230A.ncRNA.18487127.18500594-GGTLC3.18516335.18518165.rev.compl.REF ctattccttaatatctaaagcggaggttaacaagctatggacacacaagccaaacccagc 14964
                                                                            *****  *****************************************************

chimp.LOC112206744-LOC107973052-GGT2.revcompl                               cctcttggggtttttaaatctactttcaacttttattttagattcagcgggcacatgtg 28185
chimp54546-135457.revcompl.Pan.troglodytes.clone.rp43-41g5.GenBank:AC099533.36 CCTCTTGGGGTTTTTAAATCTACTTTCAACTTTTATTTTAGATTCAGCGGGCACATGTG 29899
human.FAM30B-LOC105372935-GGT2.NCBI.                                        cctcttggggtttttaaatctactttcaacttttattttagattcagcgggcacatgtg 26814
human.FAM230E-LOC105377182-GGT3P.NCBI.                                      cctcttggggtttttaaatctactttcaacttttattttagattcagcgggcacatgtg 25548
human.FAM230J-LOC105372942-GGTLC5P.NCBI.rev.compl                           cctcttggggtttttaaatctactttcaacttttattttagattcagcgggcacatgtg 26264
human.FAM230A.ncRNA.18487127.18500594-GGTLC3.18516335.18518165.rev.compl.REF cctcttggggtttttaaatctactttcaacttttattttagattcagcgggcacatgtg 15024
                                                                            ************************************************************

chimp.LOC112206744-LOC107973052-GGT2.revcompl                               caggtttgtcacgtggatatgagcatactccccaacagttggcctttcacccctcccctc 28245
chimp54546-135457.revcompl.Pan.troglodytes.clone.rp43-41g5.GenBank:AC099533.36 CAGGTTTGTCACGTGGATATGAGCATACTCCCCAACAGTTGGCCTTTTGCCCCTCCCCTC 29959
human.FAM30B-LOC105372935-GGT2.NCBI.                                        caggtttgtcacatggatatgagcatactccccaacagttggcctttcacccctcccctc 26874
human.FAM230E-LOC105377182-GGT3P.NCBI.                                      caggtttgtcacgtggatatgagcatactccccaacagttggcctttcacccctcccctc 25608
human.FAM230J-LOC105372942-GGTLC5P.NCBI.rev.compl                           caggtttgtcacgtggatatgagcatactccccaacagttggcctttcacccctcccctc 26324
human.FAM230A.ncRNA.18487127.18500594-GGTLC3.18516335.18518165.rev.compl.REF caggtttgtcacgtggatatgagcatactccccaacagttggcctttcacccctcccctc 15084
                                                                            *********** ***********************************  ***********

chimp.LOC112206744-LOC107973052-GGT2.revcompl                               cctccccatccagcagttcccagttgttgccatctttaagtcaatgactgcccatgttta 28305
chimp54546-135457.revcompl.Pan.troglodytes.clone.rp43-41g5.GenBank:AC099533.36 CCTCCCCATCCAGCAGTTCCCAGTTGTTGCCATCTTTAAGTCAATGAGTCCCCATGTTTA 30019
human.FAM30B-LOC105372935-GGT2.NCBI.                                        cctccccatccagcagttcccagttgttgccatctttaagtcaatgagtccccatgttta 26934
human.FAM230E-LOC105377182-GGT3P.NCBI.                                      cctccccatccagcagttcccagttgttgccatctttaagtcaatgagtccccatgttta 25668
```

```
human.FAM230J-LOC105372942-GGTLC5P.NCBI.rev.compl                              cctccccatccagcagttcccagttgttgccatctttaagtcaatgagtccccatgttta 26384
human.FAM230A.ncRNA.18487127.18500594-GGTLC3.18516335.18518165.rev.compl.REF   cctccccatccagcagttcccagttgttgccatctttaagtcaatgagtccccatgttta 15144
                                                                               ********************************************* * **********

chimp.LOC112206744-LOC107973052-GGT2.revcompl                                  gctcccatttataagagagaatatgcattatgttttgtttggtttttgctg------gtt 28359
chimp54546-135457.revcompl.Pan.troglodytes.clone.rp43-41g5.GenBank:AC099533.36 GCTCCCATTTATAAGAGAGAACATGCATTATGTTTTGTTTGGTTTTTGCTG------TTT 30073
human.FAM30B-LOC105372935-GGT2.NCBI.                                           gctcccatttataagagagaacatgcattatgttttgtttggtttttgctgg-ttttttt 26993
human.FAM230E-LOC105377182-GGT3P.NCBI.                                         gctcccatttataagagagaacatgcattatgttttgtttggtttttgctgg-ttttttt 25727
human.FAM230J-LOC105372942-GGTLC5P.NCBI.rev.compl                              gctcccatttataagagagaacatgcattatgttttgtttggtttttgctggtttttttt 26444
human.FAM230A.ncRNA.18487127.18500594-GGTLC3.18516335.18518165.rev.compl.REF   gctcccatttataagagagaacatgcattatgttttgtttggtttttgctgg-ttttttt 15203
                                                                               ******************** ****************************       **

chimp.LOC112206744-LOC107973052-GGT2.revcompl                                  ttttttttttaatggagtcttgccctgtagcccaggctagagtgcagtggcacaatcttg 28419
chimp54546-135457.revcompl.Pan.troglodytes.clone.rp43-41g5.GenBank:AC099533.36 TTTTTTTTTTAATGGAGTCTTGCCCTGTAGCCCAGGCTAGAGTGCAGTGGCACAATCTTG 30133
human.FAM30B-LOC105372935-GGT2.NCBI.                                           ttttttttttaatggagtcttgccctgtagcccaggctagagtgcagtggcacaaccttg 27053
human.FAM230E-LOC105377182-GGT3P.NCBI.                                         ttttttttttaatggagtcttgccctgtagcccaggctagagtgcagtggcacaatcttg 25787
human.FAM230J-LOC105372942-GGTLC5P.NCBI.rev.compl                              ttttttttttaatggagtcttgccctgtagcccaggctagagtgcagtggcacaatcttg 26504
human.FAM230A.ncRNA.18487127.18500594-GGTLC3.18516335.18518165.rev.compl.REF   ttttttttttaatggagtcttgccctgtagcccaggctagagtgcagtggcacaatcttg 15263
                                                                               ******************************************************** ****

chimp.LOC112206744-LOC107973052-GGT2.revcompl                                  gctcactgcaacctccgcctcccaggttcaaacaattctccctcctcagcctcccgagta 28479
chimp54546-135457.revcompl.Pan.troglodytes.clone.rp43-41g5.GenBank:AC099533.36 GCTCACTGCAACCTCCGCCTCCCAGGTTCAAACAATTCTCCCTCCTCAGCCTCCCGAGTA 30193
human.FAM30B-LOC105372935-GGT2.NCBI.                                           gctcactgcaacctccgcctcccaggttcaaacgattctccctcctcagcctcccgagtg 27113
human.FAM230E-LOC105377182-GGT3P.NCBI.                                         gctcactgcaacctccgcctcccaggttcaaacgattctccctcctcagcctcccgagtg 25847
human.FAM230J-LOC105372942-GGTLC5P.NCBI.rev.compl                              gctcactgcaacctccgcctcccaggttcaaacgattctccctcctcagcctcccgagtg 26564
human.FAM230A.ncRNA.18487127.18500594-GGTLC3.18516335.18518165.rev.compl.REF   gctcactgcaacctccgcctcccaggttcaaacgattctccctcctcagcctcccgagtg 15323
                                                                               ********************************* ************************* 

chimp.LOC112206744-LOC107973052-GGT2.revcompl                                  gctgggactacaggtgcctgccaccacgcccagctaactttttgtatttttagtagagac 28539
chimp54546-135457.revcompl.Pan.troglodytes.clone.rp43-41g5.GenBank:AC099533.36 GCTGGGACTACAGGTGCCCGCCACCATGCCCAGCTAACTTTTTGTATTTTTAGTAGAGAC 30253
human.FAM30B-LOC105372935-GGT2.NCBI.                                           gctgggactacaggcgcccgccaccacgcccggctaactttttgtatttttagtagagac 27173
human.FAM230E-LOC105377182-GGT3P.NCBI.                                         gctgggactacaggcgcccgccaccacgcccggctaactttttgtatttttagtagagac 25907
human.FAM230J-LOC105372942-GGTLC5P.NCBI.rev.compl                              gctgggactacaggcgcccgccaccacgcccggctaactttttgtatttttagtagagac 26624
human.FAM230A.ncRNA.18487127.18500594-GGTLC3.18516335.18518165.rev.compl.REF   gctgggactacaggcgcccgccaccacgcccggctaactttttgtatttttagtagagac 15383
                                                                               ************** ***  ****** **** ****************************

chimp.LOC112206744-LOC107973052-GGT2.revcompl                                  agggtttcactgtgttagccaggatggtctcaatctcctgacctcatgatctgcccacct 28599
chimp54546-135457.revcompl.Pan.troglodytes.clone.rp43-41g5.GenBank:AC099533.36 AGGGTTTCACCGTGTTAGCCAGGATGGTCTCAATCTCCTGACCTCATGATCTGCCCACCT 30313
human.FAM30B-LOC105372935-GGT2.NCBI.                                           agggtttcaccgtgttagccaggatggtctcaatctcctgacctcatgatctgcccacct 27233
human.FAM230E-LOC105377182-GGT3P.NCBI.                                         agggtttcaccgtgttagccaggatggtctcaatctcctgacctcatgatctgcccacct 25967
human.FAM230J-LOC105372942-GGTLC5P.NCBI.rev.compl                              agggtttcaccgtgttagccaggatggtctcaatctcctgacctcatgatctgcccacct 26684
human.FAM230A.ncRNA.18487127.18500594-GGTLC3.18516335.18518165.rev.compl.REF   agggtttcaccgtgttagccaggatggtctcaatctcctgacctcatgatctgcccacct 15443
                                                                               ********** *************************************************

chimp.LOC112206744-LOC107973052-GGT2.revcompl                                  cagtctcccaaagtgcagggattacaggcatgagccaccgtgcccagtcttttgtttatt 28659
chimp54546-135457.revcompl.Pan.troglodytes.clone.rp43-41g5.GenBank:AC099533.36 CACTCTCCCAAAGTGCAGGGATTACAGGCGTGAGCCACCGTGCCCAGCCTTTTGTTTATT 30373
human.FAM30B-LOC105372935-GGT2.NCBI.                                           tagtctcccaaagtgcagggattacaggcgtgagccaccgtgcccagccttttgtttatt 27293
human.FAM230E-LOC105377182-GGT3P.NCBI.                                         cagtctcccaaagtgcagggattacaggcgtgagccaccgtgcccagccttttgtttatt 26027
human.FAM230J-LOC105372942-GGTLC5P.NCBI.rev.compl                              cagtctcccaaagtgcagggattacaggcgggagccaccgtgcccagccttttgtttatt 26744
human.FAM230A.ncRNA.18487127.18500594-GGTLC3.18516335.18518165.rev.compl.REF   cagtctcccaaagtgcagggattacaggcgtgagccaccgtgcccagccttttgtttatt 15503
                                                                                * **************************  **************** ************

chimp.LOC112206744-LOC107973052-GGT2.revcompl                                  ttttgacgagaccgttcttgctctgtcaccaggctggagtgcactggcacagtaatagct 28719
chimp54546-135457.revcompl.Pan.troglodytes.clone.rp43-41g5.GenBank:AC099533.36 TTTTGACGAGACCGTTCTTGCTCTGTCACCAGGCTGGAGTGCACTGGCACAGTAATAGCT 30433
human.FAM30B-LOC105372935-GGT2.NCBI.                                           ttttgatgagacggttcttgctctgtcaccaggctggagtgcactggcacaataatagct 27353
human.FAM230E-LOC105377182-GGT3P.NCBI.                                         ttttgacgagacggttcttgctctgtcaccaggctggagtgcactggcacaataatagct 26087
human.FAM230J-LOC105372942-GGTLC5P.NCBI.rev.compl                              ttttgacgagaccgttcttgctctgtcaccaggctggagtgcactggcacaataatagct 26804
human.FAM230A.ncRNA.18487127.18500594-GGTLC3.18516335.18518165.rev.compl.REF   ttttgacgagacggttcttgctctgtcaccaggctggagtgcactggcacaataatagct 15563
                                                                               ****** ***** **************************************** *******

chimp.LOC112206744-LOC107973052-GGT2.revcompl                                  caccacagccttgtgctcctgggctcaactgaccctcctgcctcagtttagcttcctga 28779
chimp54546-135457.revcompl.Pan.troglodytes.clone.rp43-41g5.GenBank:AC099533.36 CACCACAGCCTCGTGCTCCTGGGCTCAACTGACCCTCCTGCCTCAGTTTAGCTTCCTGA 30493
human.FAM30B-LOC105372935-GGT2.NCBI.                                           caccacagcctcgtgctcctgggctcaactgaccctcctgcctcagtttagcttcctga 27413
human.FAM230E-LOC105377182-GGT3P.NCBI.                                         caccacagcctcgtgctcctgggctcaactgaccctcctgcctcagtttagcttcctga 26147
human.FAM230J-LOC105372942-GGTLC5P.NCBI.rev.compl                              caccacagcctcgtgctcctgggctcaactgaccctcctgcctcagtttagcttcctga 26864
human.FAM230A.ncRNA.18487127.18500594-GGTLC3.18516335.18518165.rev.compl.REF   caccacagcctcgtgctcctgggctcaactgaccctcctgcctcagtttagcttcctga 15623
                                                                               *********** ************************************************

chimp.LOC112206744-LOC107973052-GGT2.revcompl                                  gtagctaggactacgggtgtgtaccaccatgcctagctataataatttttatttttttgt 28839
chimp54546-135457.revcompl.Pan.troglodytes.clone.rp43-41g5.GenBank:AC099533.36 GTAGCTAGGACTACGGGTGTGTACCACCATGCCTAGCTATAATAATTTTTATTTTTTTGT 30553
human.FAM30B-LOC105372935-GGT2.NCBI.                                           gtagctaggactacgggtgtgtaccaccatgcctagctataataatttttatttttttgt 27473
human.FAM230E-LOC105377182-GGT3P.NCBI.                                         gtagctaggactacaggtgtgtaccaccatgcctagctataataatttttatttttttgt 26207
human.FAM230J-LOC105372942-GGTLC5P.NCBI.rev.compl                              gtagctaggactacgggtgtgtaccaccatgcctagctataataatttttatttttttgt 26924
human.FAM230A.ncRNA.18487127.18500594-GGTLC3.18516335.18518165.rev.compl.REF   gtagctaggactacaggtgtgtaccaccatgcctagctataataatttttatttttttgt 15683
                                                                               ************** *********************************************

chimp.LOC112206744-LOC107973052-GGT2.revcompl                                  agagatggagtcttgctttgttgcccaggctggtcttgaactcctggcttaaagtgatcc 28899
chimp54546-135457.revcompl.Pan.troglodytes.clone.rp43-41g5.GenBank:AC099533.36 AGAGATGGAGACTTGCTTTGTTGCCCAGGCTGGTCTTGAACTCCTGGCTTAAAGTGATCC 30613
human.FAM30B-LOC105372935-GGT2.NCBI.                                           agagatggagtcttgctttgttgcccaggctggtcttgaactcctggcttgaagtgatcc 27533
human.FAM230E-LOC105377182-GGT3P.NCBI.                                         agagatggagtcttgctttgttgcccaggctggtcttgaactcctggcttgaagtgatcc 26267
human.FAM230J-LOC105372942-GGTLC5P.NCBI.rev.compl                              agagatggagtcttgctttgttgcccaggctggtcttgaactcctggcttgaagtgatcc 26984
human.FAM230A.ncRNA.18487127.18500594-GGTLC3.18516335.18518165.rev.compl.REF   agagatggagtcttgctttgttgcccaggctggtcttgaagtcctggcttgaagtgatcc 15743
                                                                               ********** ***************************** ********* *********

chimp.LOC112206744-LOC107973052-GGT2.revcompl                                  tcctgcctcggcctcccaaagtgctgggattaaaggtgtgagatcacacccagtctccaa 28959
chimp54546-135457.revcompl.Pan.troglodytes.clone.rp43-41g5.GenBank:AC099533.36 TCCTGCCTCGGCCTCCCAAAGTGCTGGGATTAAAGGTATGAGATCACACCCAGTCTCCAA 30673
human.FAM30B-LOC105372935-GGT2.NCBI.                                           tcctgcctcggcctcccaaagtgccgggattaaaggtgggagatcgcacccagtctccaa 27593
human.FAM230E-LOC105377182-GGT3P.NCBI.                                         tcctgcctcggcctcccaaattgccgggattaaaggtgggagatcgcacccagtctccaa 26327
human.FAM230J-LOC105372942-GGTLC5P.NCBI.rev.compl                              tcctgcctcggcctcccaaagtgccgggattaaaggtgggagatcgcacccagtctccaa 27044
human.FAM230A.ncRNA.18487127.18500594-GGTLC3.18516335.18518165.rev.compl.REF   tcctgcctcggcctcccaaattgccgggattaaaggtgggagatcgcacccagtctccaa 15803
                                                                               ******************** *** ************  ****** **************

chimp.LOC112206744-LOC107973052-GGT2.revcompl                                  ccctctttttgcaagtaaatgtaactggaccccagccatgctcatctgcccatgtactgt 29019
chimp54546-135457.revcompl.Pan.troglodytes.clone.rp43-41g5.GenBank:AC099533.36 CCCTCTTTTTGCAAGTAAATGTAACTGGACCCCAGCCATGCTCATCTGCCCATGTACTGT 30733
human.FAM30B-LOC105372935-GGT2.NCBI.                                           ccctctttttgcaagtaaatgtaactggaccccagccatgctcatctgcccatgtactgt 27653
human.FAM230E-LOC105377182-GGT3P.NCBI.                                         ccctctttttgcaagtaaatgtaactggaccccagccatgctcatctgcccatgtactgt 26387
human.FAM230J-LOC105372942-GGTLC5P.NCBI.rev.compl                              ccctctttttgcaagtaaatgtaactggaccccagccatgctcatctgcccatgtactgt 27104
human.FAM230A.ncRNA.18487127.18500594-GGTLC3.18516335.18518165.rev.compl.REF   ccctctttttgcaagtaaatgtaactggaccccagccatgctcatctgcccatgtactgt 15863
                                                                               ************************************************************

chimp.LOC112206744-LOC107973052-GGT2.revcompl                                  ctacggctgcttttgctctatagggcagagttaagtggttgcaacagacactgcacagac 29079
chimp54546-135457.revcompl.Pan.troglodytes.clone.rp43-41g5.GenBank:AC099533.36 CTACGGCTGCTTTTGCTCTATAGGGCAGAGTTAAGTGGTTGCAACAGACACTGCACAGAC 30793
human.FAM30B-LOC105372935-GGT2.NCBI.                                           ctacggctgcttttgctctacagggcagagttaagtggttgcaacagacactgcacagac 27713
human.FAM230E-LOC105377182-GGT3P.NCBI.                                         ctgcggctgcttttgctctacagggcagagttaagtggttgcaacagacaccgcacagac 26447
human.FAM230J-LOC105372942-GGTLC5P.NCBI.rev.compl                              ctgcggctgcttttgctctacagggcagagttaagtggttgcaacagacaccgcacagac 27164
human.FAM230A.ncRNA.18487127.18500594-GGTLC3.18516335.18518165.rev.compl.REF   ctacggctgcttttgctctacagggcagagttaagtggttgcaacagacaccgcacagac 15923
                                                                               ** ***************** ****************************** ********

chimp.LOC112206744-LOC107973052-GGT2.revcompl                                  cacaaagtctggaagtactttctctccagccctttacagagaaagtctgccaacctctaa 29139
chimp54546-135457.revcompl.Pan.troglodytes.clone.rp43-41g5.GenBank:AC099533.36 CACAAAGTCT-GAAGTACTTTCTCTCCAGCCCTTTACAGAGAAAGTCTGCCAACCTCTAA 30852
human.FAM30B-LOC105372935-GGT2.NCBI.                                           cacaaagtct-gaagtactttctctccagccctttacagagaaagtctgccaacctctaa 27772
human.FAM230E-LOC105377182-GGT3P.NCBI.                                         cacaaagtct-gaagtactttctctccagccctttacagagaaagtctgccaacctctaa 26506
human.FAM230J-LOC105372942-GGTLC5P.NCBI.rev.compl                              cacaaagtct-gaagtactttctctccagccctttacagagaaagtctgccaacctctaa 27223
human.FAM230A.ncRNA.18487127.18500594-GGTLC3.18516335.18518165.rev.compl.REF   cacaaagtct-gaagtactttctctccagccctttacagagaaagtctgccaacctctaa 15982
                                                                               ********** *************************************************

chimp.LOC112206744-LOC107973052-GGT2.revcompl                                  tctcaataacagggaaatcaatgacaaccacaaagtgacaaagattgggtgtctaagatg 29199
chimp54546-135457.revcompl.Pan.troglodytes.clone.rp43-41g5.GenBank:AC099533.36 TCTCAATAACAGGGAAATCAATGACAACCACAAAGTGACAAAGATTGGGTGTCTAAGATG 30912
human.FAM30B-LOC105372935-GGT2.NCBI.                                           tctcaataacagggaaatcaatgacaaccacaaagtgacaaagattgggtgtctaagatg 27832
human.FAM230E-LOC105377182-GGT3P.NCBI.                                         tctcaataacagggaaatcaatgacaaccacaaagtgacaaagattgggtgtctaagatg 26566
human.FAM230J-LOC105372942-GGTLC5P.NCBI.rev.compl                              tctcaataacagggaaatcaatgacaaccacaaagtgacaaagattgggtgtctaagatg 27283
human.FAM230A.ncRNA.18487127.18500594-GGTLC3.18516335.18518165.rev.compl.REF   tctcaataacagggaaatcaatgacaaccacaaagtgacaaagactgggtgtctaagatg 16042
                                                                               ********************************************* **************

chimp.LOC112206744-LOC107973052-GGT2.revcompl                                  gatggtcagaataaacaagagagaaagatgaaaagtagaaggaggatttcaagcgcaagc 29259
chimp54546-135457.revcompl.Pan.troglodytes.clone.rp43-41g5.GenBank:AC099533.36 GATGGTCAGAATAAACAAGAGAGAAAGATGAAAAGTAGAAGGAGGATTTCAAACGCAAGC 30972
human.FAM30B-LOC105372935-GGT2.NCBI.                                           gatgttcagaataaacaagagagaaagatgagaagtagaaggaggatttcaaacgcaagc 27892
human.FAM230E-LOC105377182-GGT3P.NCBI.                                         gatgctcagaataaacaagagagaaagatgaaaagtagaaggaggatttcaaacgcaagc 26626
human.FAM230J-LOC105372942-GGTLC5P.NCBI.rev.compl                              gatgctcagaataaacaagagagaaagatgaaaagtagaaggaggatttcaaacgcaagc 27343
human.FAM230A.ncRNA.18487127.18500594-GGTLC3.18516335.18518165.rev.compl.REF   gatgctcagaataaacaagagagaaagatgaaaagtagaaggaggatttcaaacgcaagc 16102
                                                                               **** ************************** ******************** *******

chimp.LOC112206744-LOC107973052-GGT2.revcompl                                  ttcacctaatccgttatttttcaaatgaccaggcctatctctgtagctgaaaatcacctc 29319
chimp54546-135457.revcompl.Pan.troglodytes.clone.rp43-41g5.GenBank:AC099533.36 TTCACCTAATCCGTTATTTTTCAAATGACCAGGCCTATCTCTGTAGCTGAAAATCACCTC 31032
human.FAM30B-LOC105372935-GGT2.NCBI.                                           ttcacctaatccgttatttttcaaatgaccaggcctatctctgtagccgaaaatcacctc 27952
human.FAM230E-LOC105377182-GGT3P.NCBI.                                         ttcacctaatccgttatttttcaaatgaccaggcctatctctgtagctgaaaatcacctc 26686
human.FAM230J-LOC105372942-GGTLC5P.NCBI.rev.compl                              ttcacctaatccgttatttttcaaatgaccaggcctatctctgtagctgaaaatcacctc 27403
human.FAM230A.ncRNA.18487127.18500594-GGTLC3.18516335.18518165.rev.compl.REF   ttcacctaatccgttatttttcaaatgaccaggcctatctctgtagctgaaaatcacctc 16162
                                                                               *********************************************** ************

chimp.LOC112206744-LOC107973052-GGT2.revcompl                                  aaataggatctctgatatacagtctccaaaagctcagctaagaaacttac--agtctctc 29377
chimp54546-135457.revcompl.Pan.troglodytes.clone.rp43-41g5.GenBank:AC099533.36 AAATAGGATCTCTGATATACAGTCTCCAAAAGCTCAGCCAAGAAACTTACAAAGTCTCTC 31092
human.FAM30B-LOC105372935-GGT2.NCBI.                                           aaataggatctctgatatacagtctccaaaagctcagctaagaaacttacaaagtctctc 28012
human.FAM230E-LOC105377182-GGT3P.NCBI.                                         aaataagatctctgatatacagtctccaaaagctcagctaagaaacttacaaagtctctc 26746
human.FAM230J-LOC105372942-GGTLC5P.NCBI.rev.compl                              aaataggatctctgatatacagtctccaaaagctcagccaagaaacttacaaagtctctc 27463
```

```
human.FAM230A.ncRNA.18487127.18500594-GGTLC3.18516335.18518165.rev.compl.REF        aaataagatctctgatatacagtctccaaaagctcagccaagaaacttacaaagtctctc 16222
                                                                                    ***** ******************************** ***********  ********

chimp.LOC112206744-LOC107973052-GGT2.revcompl                                       tgccttaacttcatccaccttttttctctccagcttctcctcggtagttaatgattataa 29437
chimp54546-135457.revcompl.Pan.troglodytes.clone.rp43-41g5.GenBank:AC099533.36      TGCCTTAACTTCATCCACCTTTTTTCTCTCCAGCTTCTCCTCGGTAGTTAATGATTATAA 31152
human.FAM30B-LOC105372935-GGT2.NCBI.                                                tgccttaacttcatccaccttttttctctccagcttctcctcggtagttaatgattataa 28072
human.FAM230E-LOC105377182-GGT3P.NCBI.                                              tgccttaacttcatccaccttttttctctccagcttttcctcggtagttaatgattataa 26806
human.FAM230J-LOC105372942-GGTLC5P.NCBI.rev.compl                                   tgccttaacttcatccaccttttttctctccagcttctcctcggtagttaatgattataa 27523
human.FAM230A.ncRNA.18487127.18500594-GGTLC3.18516335.18518165.rev.compl.REF        tgccttaacttcatccaccttttttctctccagcttctcctcggtagttaatgattataa 16282
                                                                                    ************************************ ***********************

chimp.LOC112206744-LOC107973052-GGT2.revcompl                                       aaatatttattggctcatgcctgtaatcccagcactttgagaggccaaggcgggcagatc 29497
chimp54546-135457.revcompl.Pan.troglodytes.clone.rp43-41g5.GenBank:AC099533.36      AAATATTTATTGGCTCATGCCTGTAATCCCAGCACTTTGAGAGGCCAAGGCGGGCAGATC 31212
human.FAM30B-LOC105372935-GGT2.NCBI.                                                aaatatttattggctcatgcctgtaatcccagcactttgagaggccgaggcgggcagatc 28132
human.FAM230E-LOC105377182-GGT3P.NCBI.                                              aaatatttattggctcatgcctgtaatcccagcactttgagaggccgaggcgggcagatc 26866
human.FAM230J-LOC105372942-GGTLC5P.NCBI.rev.compl                                   aaatatttattggctcatgcctgtaatcccagcactttgagaggccgaggcgggcagatc 27583
human.FAM230A.ncRNA.18487127.18500594-GGTLC3.18516335.18518165.rev.compl.REF        aaatatttattggctcatgcctgtaatcccagcactttgagaggccgaggcgggcagatc 16342
                                                                                    ********************************************** *************

chimp.LOC112206744-LOC107973052-GGT2.revcompl                                       acgaggtcaggagagcgagaccatcctggctaacacggtgaaatcccgtctctactaaaa 29557
chimp54546-135457.revcompl.Pan.troglodytes.clone.rp43-41g5.GenBank:AC099533.36      ACGAGGTCAGGAGATCGAGACCATCCTGGCTAACACGGTGAAATCCCATCTCTACTAAAA 31272
human.FAM30B-LOC105372935-GGT2.NCBI.                                                ac--ggtcaggagatcgagaccatcctggctaacacggtgaaatcccgtctctactaaaa 28190
human.FAM230E-LOC105377182-GGT3P.NCBI.                                              acgaggtcaggagattgagaccatcctggctaacacggtgaaatcccgtctctactaaaa 26926
human.FAM230J-LOC105372942-GGTLC5P.NCBI.rev.compl                                   acgaggtcaggagatcgagaccatcctggctaacacggtgaaatcccgtctctactaaaa 27643
human.FAM230A.ncRNA.18487127.18500594-GGTLC3.18516335.18518165.rev.compl.REF        acgaggtcaggagatcgagaccatcctggctaacacggtgaaatcccgtctctactaaaa 16402
                                                                                    **  **********  ******************************* ************

chimp.LOC112206744-LOC107973052-GGT2.revcompl                                       atacaaaaaattagccaggcgtggtggcgggcacctgtaatcccagctactcgggaggct 29617
chimp54546-135457.revcompl.Pan.troglodytes.clone.rp43-41g5.GenBank:AC099533.36      ATACAAAAAATTAGCCAGGCGTGGTGGCGGGCGCCTGTAATCCCAGCTACTCGGGAGGCT 31332
human.FAM30B-LOC105372935-GGT2.NCBI.                                                atacaaaaaattagccaggcgtggtggcgggcgcctgtaatcccagctactcaggaggct 28250
human.FAM230E-LOC105377182-GGT3P.NCBI.                                              atacaaaaaattagccaggcgtggtggcgggcgcctgtaatcccagctactcaggaggct 26986
human.FAM230J-LOC105372942-GGTLC5P.NCBI.rev.compl                                   atacaaaaaattagccaggcgtggtggcgggcgcctgtaatcccagctactcaggaggct 27703
human.FAM230A.ncRNA.18487127.18500594-GGTLC3.18516335.18518165.rev.compl.REF        atacaaaaaattagccaggcgtggtggcgggcgcctgtaatcccagctactcaggaggct 16462
                                                                                    ******************************** ******************* *******

chimp.LOC112206744-LOC107973052-GGT2.revcompl                                       gaggcaggagaatggcgtgaacccacaaggcggagcttgcaatgaggtgagatcccacta 29677
chimp54546-135457.revcompl.Pan.troglodytes.clone.rp43-41g5.GenBank:AC099533.36      GAGGCAGGAGAATGGCGTGAACCCACAAGGCGGAGCTTGCAATGAGGTGAGATCCCACTA 31392
human.FAM30B-LOC105372935-GGT2.NCBI.                                                gaggcaggagaatggcgtgaacccacaaggcggagcttgcaatgaggtgagatcccacta 28310
human.FAM230E-LOC105377182-GGT3P.NCBI.                                              gaggcaggagaatggcatgtacccacaaggcggagcttacaatgaggtgagatcccacta 27046
human.FAM230J-LOC105372942-GGTLC5P.NCBI.rev.compl                                   gaggcaggagaatggcgtgaacccacaaggcggagcttgcaatgaggtgagatcccacta 27763
human.FAM230A.ncRNA.18487127.18500594-GGTLC3.18516335.18518165.rev.compl.REF        gaggcaggagaatggcatgtacccacaaggcggagcttgcaacgaggtgagatcccacta 16522
                                                                                    **************** ** ****************** *** *****************

chimp.LOC112206744-LOC107973052-GGT2.revcompl                                       ctgcactccagcctgggcgacagagcaagactccatctca----------------aaac 29721
chimp54546-135457.revcompl.Pan.troglodytes.clone.rp43-41g5.GenBank:AC099533.36      CTGCACTCCAGCCTGGGCGACAGAGCAAGACTCCATCTCAAAACAAACAAACAAACAAAC 31452
human.FAM30B-LOC105372935-GGT2.NCBI.                                                ctgcactccagcctgggcgacagagcaagactccatctca--------aaacaaacaaac 28362
human.FAM230E-LOC105377182-GGT3P.NCBI.                                              ctgcactccagcctgggcgacagagcaagactccatctca--------aaacaaacaaac 27098
human.FAM230J-LOC105372942-GGTLC5P.NCBI.rev.compl                                   ctgcactccagcctgagcgacagagcaagactccatctca--------aaacaaacaaac 27815
human.FAM230A.ncRNA.18487127.18500594-GGTLC3.18516335.18518165.rev.compl.REF        ctgcactccagcctgggcgacagagcaagactccatctca--------aaacaaacaaac 16574
                                                                                    *************** ************************                ****

chimp.LOC112206744-LOC107973052-GGT2.revcompl                                       aaacaaacaaacaaaaaaacagtgtgatggccaggcgcggtgctcaagcctataatccaa 29781
chimp54546-135457.revcompl.Pan.troglodytes.clone.rp43-41g5.GenBank:AC099533.36      AAACAAACAAACAAAAAAACAGTGTGATGGCCAGGCGCGGTGCTCATGCCTATAATCCAA 31512
human.FAM30B-LOC105372935-GGT2.NCBI.                                                aaacaaa----caaaaaaacagtgtgatggccaggcgcagtgctcatgcctataatccaa 28418
human.FAM230E-LOC105377182-GGT3P.NCBI.                                              aaacaaa----caaaaaaacagtgtgatggccaggcgcagtgctcatgcctataatccaa 27154
human.FAM230J-LOC105372942-GGTLC5P.NCBI.rev.compl                                   aaacaaacaaccaaaaaaacagtgtgatggccaggcgcagtgctcatgcctataatccaa 27875
human.FAM230A.ncRNA.18487127.18500594-GGTLC3.18516335.18518165.rev.compl.REF        aaacaaa----caaaaaaacagtgtgatggccaggcgcagtgctcatgcctataatccaa 16630
                                                                                    *******    *************************** ******* *************

chimp.LOC112206744-LOC107973052-GGT2.revcompl                                       gcactttgggaggctgaaatggatcgatggcttgagcccagtagtttgagacaagcctgg 29841
chimp54546-135457.revcompl.Pan.troglodytes.clone.rp43-41g5.GenBank:AC099533.36      GCACTTTGGGAGGCTGAAATGGATGGATGGCTTGAGCCCAGTAGTTTGAGACAAGCCTGG 31572
human.FAM30B-LOC105372935-GGT2.NCBI.                                                gcactttgggaggctgaaatggatggatggcttgagcccagtagtttgagacaagcctgg 28478
human.FAM230E-LOC105377182-GGT3P.NCBI.                                              gcactttgggaggctgaaatggatggatggcttgagcccagtagtttgagacaagcctgg 27214
human.FAM230J-LOC105372942-GGTLC5P.NCBI.rev.compl                                   gcactttgggaggctgaaatggatggatggcttgagcccagtagtttgagacaagcctgg 27935
human.FAM230A.ncRNA.18487127.18500594-GGTLC3.18516335.18518165.rev.compl.REF        gcactttgggaggctgaaatggatggatggcttgagcccagtagtttgagacaagcctgg 16690
                                                                                    ************************ ***********************************

chimp.LOC112206744-LOC107973052-GGT2.revcompl                                       caacatagcgagacctcatctctacaaacatttttaaaatatgccaggcatggtgtgcag 29901
chimp54546-135457.revcompl.Pan.troglodytes.clone.rp43-41g5.GenBank:AC099533.36      CAACATAGCGAGACCTCATCTCTACAAACATTTTTAAAATATGCCAGGCATGGTG----- 31627
human.FAM30B-LOC105372935-GGT2.NCBI.                                                caacatagcgagacctcatctctacaaacatctttaaaatatgccaggcatggtg----- 28533
human.FAM230E-LOC105377182-GGT3P.NCBI.                                              caacatagcgagacctcatctctacaaacatctttaaaatatgccaggcatggtg----- 27269
human.FAM230J-LOC105372942-GGTLC5P.NCBI.rev.compl                                   caacatagcgagacctcatgtctacaaacatctttaaaatatgccaggcatggtg----- 27990
human.FAM230A.ncRNA.18487127.18500594-GGTLC3.18516335.18518165.rev.compl.REF        caacatagcgagacctcatctctacaaacatctttaaaatatgccaggcatggtg----- 16745
                                                                                    ******************* *********** ***********************

chimp.LOC112206744-LOC107973052-GGT2.revcompl                                       actgcatgcctgtagtcccagctattcgggaggctgaggtgggaggatcacctgtgcccg 29961
chimp54546-135457.revcompl.Pan.troglodytes.clone.rp43-41g5.GenBank:AC099533.36      -GTGCATGCCTGTAGTCCCAGCTATTCAGGAGGCTGAGGTGGGAGGATCACCTGTGCCTG 31686
human.FAM30B-LOC105372935-GGT2.NCBI.                                                -gtgcatgcctgtagtcccagctattcaggaggctgaggtgggaggatcacctgtgcccg 28592
human.FAM230E-LOC105377182-GGT3P.NCBI.                                              -gtgcatgcctgtagtcccagctattcaggaggctgaggtgggaggatcacctgtgcccg 27328
human.FAM230J-LOC105372942-GGTLC5P.NCBI.rev.compl                                   -gtgcatgcctgtagtcccagctattcaggaggctgaggtgggaggatcacctgtgcccg 28049
human.FAM230A.ncRNA.18487127.18500594-GGTLC3.18516335.18518165.rev.compl.REF        -gtgcatgcctgtagtcccagctattcaggaagctgaggtgggaggatcacctgtgcccg 16804
                                                                                     ************************** *** ************************** *

chimp.LOC112206744-LOC107973052-GGT2.revcompl                                       ggagttcaaggctgcagtgagctatgatcacaccacagtgctccagcctgggcaacaaag 30021
chimp54546-135457.revcompl.Pan.troglodytes.clone.rp43-41g5.GenBank:AC099533.36      GGAGTTCAAGGCTGCAGTGAGCTATGATCACACCACAGTGCTCCAGCCTGGGCAACAAAG 31746
human.FAM30B-LOC105372935-GGT2.NCBI.                                                ggagttcaaggctgcagtgagctatgatcacaccacagtgctccagcctgggcaacaaag 28652
human.FAM230E-LOC105377182-GGT3P.NCBI.                                              ggagttcaaggctgcagtgagctatgatcacaccacagtgctccagcctgggcaacaaag 27388
human.FAM230J-LOC105372942-GGTLC5P.NCBI.rev.compl                                   ggagttcaaggctgcagtgagctatgatcacaccacagtgctccagcctgggcaacaaag 28109
human.FAM230A.ncRNA.18487127.18500594-GGTLC3.18516335.18518165.rev.compl.REF        ggagttcaaggctgcagtgagctatgatcacaccacagtgctccagcctgggcaacaaag 16864
                                                                                    ************************************************************

chimp.LOC112206744-LOC107973052-GGT2.revcompl                                       caagactccatctctaaaaataaaataaaatt--aaagaaaagatctttgctgtaaaaga 30079
chimp54546-135457.revcompl.Pan.troglodytes.clone.rp43-41g5.GenBank:AC099533.36      CAAGACTCCATCTCTAAAAATAAAATAAAATT--AAAAAAAAGATCTTCGCTGTAAAAGA 31804
human.FAM30B-LOC105372935-GGT2.NCBI.                                                caagactccatctctaaaaataaaataaaattt-aaaaaaaagatcttcgctgtaaaaga 28711
human.FAM230E-LOC105377182-GGT3P.NCBI.                                              caagactccatctctaaaaataaaataaaatt-aaaaaaaaagatcttcgctgtaaaaga 27447
human.FAM230J-LOC105372942-GGTLC5P.NCBI.rev.compl                                   caagactccatctctaaaaataaaataaaatt-aaaaaaaaagatcttcgctgtaaaaga 28168
human.FAM230A.ncRNA.18487127.18500594-GGTLC3.18516335.18518165.rev.compl.REF        caagactccatctctaaaaataaaataaaattaaaaaaaaaagatcttcgctgtaaaaga 16924
                                                                                    ********************************  *** ********** ***********

chimp.LOC112206744-LOC107973052-GGT2.revcompl                                       ggtacgctcaaatgcaataaaagcatataagaaggccgggtgtggtggctcatgcctgta 30139
chimp54546-135457.revcompl.Pan.troglodytes.clone.rp43-41g5.GenBank:AC099533.36      GGTACGCTCAAATGCAATAAAAGCATATAAGAAGGCCGGGTGTAGTGGCTCATGCCTGTA 31864
human.FAM30B-LOC105372935-GGT2.NCBI.                                                ggtacgctcaaatgcaataaaagcatataagaaggccgggtgtggtggctcatgcctgta 28771
human.FAM230E-LOC105377182-GGT3P.NCBI.                                              ggtacgctcaaatgcaataaaagcatataagaaggccgggtgtggtggctcatgcctgta 27507
human.FAM230J-LOC105372942-GGTLC5P.NCBI.rev.compl                                   ggtacgctcaaatgcaataaaagcatataagaaggccgggtgtggtggctcatgcctgta 28228
human.FAM230A.ncRNA.18487127.18500594-GGTLC3.18516335.18518165.rev.compl.REF        ggtacactcaaatgcaataaaagcatataagaaggccgggtgtggtggctcatgcctgta 16984
                                                                                    ***** ************************************* ****************

chimp.LOC112206744-LOC107973052-GGT2.revcompl                                       atctcagcactttgggaggccaagacgggcggatcacgaggtcaggagatcgagactatc 30199
chimp54546-135457.revcompl.Pan.troglodytes.clone.rp43-41g5.GenBank:AC099533.36      ATCTCAGCACTTTGGGAGGCCGAGACGGGCGGATCACGAGGTCAGGAGATCGAGACTATC 31924
human.FAM30B-LOC105372935-GGT2.NCBI.                                                atcccagcactttgggaggccgagacgggcggatcacgaggtcaggagattgagactatc 28831
human.FAM230E-LOC105377182-GGT3P.NCBI.                                              atcccagcactttgggaggccgagacgggcggatcacgaggtcaggagattgagactatc 27567
human.FAM230J-LOC105372942-GGTLC5P.NCBI.rev.compl                                   atcccagcactttgggaggccgagacgggcggatcacgaggtcaggagattgagactatc 28288
human.FAM230A.ncRNA.18487127.18500594-GGTLC3.18516335.18518165.rev.compl.REF        atcccagcactttgggaggccgagacgggcggatcacgaggtcaggagattgagactatc 17044
                                                                                    *** ***************** **************************** *********

chimp.LOC112206744-LOC107973052-GGT2.revcompl                                       ctggctaacgcggagaaaccccatctcctctaaaagtacaaaaaaattagttgggctagg 30259
chimp54546-135457.revcompl.Pan.troglodytes.clone.rp43-41g5.GenBank:AC099533.36      CTGGCTAACACGGTGAAACCCCATCTCCTCTAAAAGTACAAAAAAATTAGTTGGGCTAGG 31984
human.FAM30B-LOC105372935-GGT2.NCBI.                                                ctggctaacgcggtgaaaccccatctcctctaaaagtacaaaaaaattagctgggctagg 28891
human.FAM230E-LOC105377182-GGT3P.NCBI.                                              ctggctaacgcggtgaaaccccatctcctctaaaagtacaaaaaaattagctgggctagg 27627
human.FAM230J-LOC105372942-GGTLC5P.NCBI.rev.compl                                   ctggctaacgcggtgaaaccccatctcctctaaaagtacaaaaaaattagctgggctagg 28348
human.FAM230A.ncRNA.18487127.18500594-GGTLC3.18516335.18518165.rev.compl.REF        ctggctaacgcggtgaaaccccatctcctctaaaagtacaaaaaaattagctgggctagg 17104
                                                                                    ********* *** ************************************ *********

chimp.LOC112206744-LOC107973052-GGT2.revcompl                                       tggcaggcgcctgtagtcccagctactcaggaggctgaggcaggagaatggcataaaccc 30319
chimp54546-135457.revcompl.Pan.troglodytes.clone.rp43-41g5.GenBank:AC099533.36      TGGCAGGCGCCTGTAGTCCCAGCTACTCAGGAGGCTGAGGCAGGAGAATGGCATAAACCC 32044
human.FAM30B-LOC105372935-GGT2.NCBI.                                                tggcaggcgcctgtagtcccagctactgaggaggctgaggcaggagaatggcataaacca 28951
human.FAM230E-LOC105377182-GGT3P.NCBI.                                              tggcaggcgcctgtagtcccagctactcaggaggctgaggcaggagaatggcataaaccc 27687
human.FAM230J-LOC105372942-GGTLC5P.NCBI.rev.compl                                   tggcaggcgcctgtagtcccagctactcaggaggctgaggcaggagaatggcataaaccc 28408
human.FAM230A.ncRNA.18487127.18500594-GGTLC3.18516335.18518165.rev.compl.REF        tggcaggcgcctgtagtcccagctactcaggaggctgaggcaggagaatggcataaaccc 17164
                                                                                    *************************** ******************************* 

chimp.LOC112206744-LOC107973052-GGT2.revcompl                                       gggaggcagagcttgcagtgagccaagatcgcaccactgcactccagcctgggtgacaga 30379
chimp54546-135457.revcompl.Pan.troglodytes.clone.rp43-41g5.GenBank:AC099533.36      GGGAGGCAGAGCTTGCAGTGAGCCAAGATCGCACCACTGCACTCCACCCTGGGTGACAGA 32104
human.FAM30B-LOC105372935-GGT2.NCBI.                                                gggaggcagagcttgcagtgagcctagatcacaccactgcactccagcctgggtgacaga 29011
human.FAM230E-LOC105377182-GGT3P.NCBI.                                              gggaggcagagcttgcaatgagcctagatcgcaccactgcactccagcctgggtgacaga 27747
human.FAM230J-LOC105372942-GGTLC5P.NCBI.rev.compl                                   gggaggcagagcttgcagtgagcctagatcgcaccactgcactccagcctgggtgacaga 28468
human.FAM230A.ncRNA.18487127.18500594-GGTLC3.18516335.18518165.rev.compl.REF        gggaggcagagcttgcagtgagcctagatagcaccactgccctccagcctgggtgacaga 17224
                                                                                    ***************** ****** ****  ********* ***** *************

chimp.LOC112206744-LOC107973052-GGT2.revcompl                                       gcgagactccgtctcaaaaaaaaaaaaa--aaaagaaaaagaaaagaaaagttcttgtga 30437
chimp54546-135457.revcompl.Pan.troglodytes.clone.rp43-41g5.GenBank:AC099533.36      GCGAGACTCCGTCTCAAAAAAA--------AAAGAAAAAGAAAAGAAAAGTTCTTGTGA 32156
human.FAM30B-LOC105372935-GGT2.NCBI.                                                gcgagactccgtctcaaaaaaaaaaaaaagaaaagaaaaagaaaagaaaagttcttctga 29071
human.FAM230E-LOC105377182-GGT3P.NCBI.                                              gtgagactccgtctc-aaaaaaaaaaaaa-aaaagaaaaagaaaagaaaagttcttctga 27805
human.FAM230J-LOC105372942-GGTLC5P.NCBI.rev.compl                                   gtgagactccgtctc---aaaaaaaaaaa-aaaagaaaaagaaaagaaaagttcttgtga 28524
human.FAM230A.ncRNA.18487127.18500594-GGTLC3.18516335.18518165.rev.compl.REF        gcgagactccgtctcaaaaaaaaaaaaaa-aaaagaaaaagaaaagaaaagttcttgtga 17283
```

```
                                                                                  * *************   ****        ************************** ***

chimp.LOC112206744-LOC107973052-GGT2.revcompl                                     catttgtgtatgaaatcagccttcactacatggataggaccagcacgcttccgcggcacg 30497
chimp54546-135457.revcompl.Pan.troglodytes.clone.rp43-41g5.GenBank:AC099533.36    CATTTGTGTATGAAATCAGCCTTCACTACATGGATAGGACCAGCACGCTTCCATGGCACG 32216
human.FAM30B-LOC105372935-GGT2.NCBI.                                              catttgtgtatgaaatcagccttcactacatggataggaccagcacgcttctgcggcacg 29131
human.FAM230E-LOC105377182-GGT3P.NCBI.                                            catttgtgtatgaaatcagccttcactacatggataggaccagcacgcttctgcggcacg 27865
human.FAM230J-LOC105372942-GGTLC5P.NCBI.rev.compl                                 catttgtgtatgaaatcagccttcactacatggataggaccagcacgcttctgcggcacg 28584
human.FAM230A.ncRNA.18487127.18500594-GGTLC3.18516335.18518165.rev.compl.REF      catttgtgtatgaaatcagccttcactacatggataggaccagcacgcttctgcggcacg 17343
                                                                                  ***************************************************   ******

chimp.LOC112206744-LOC107973052-GGT2.revcompl                                     actctgcaatcttactacatttttttttactttgtattttatttattccttttgagacag 30557
chimp54546-135457.revcompl.Pan.troglodytes.clone.rp43-41g5.GenBank:AC099533.36    ACTCTGCAATCTTACTACATTTTTTTTTACTTTGTATTTTATTTATTCCTTTTGAGACAG 32276
human.FAM30B-LOC105372935-GGT2.NCBI.                                              actctgcaatcttactacatttttttttactttgtattttatttattccttttgagacag 29191
human.FAM230E-LOC105377182-GGT3P.NCBI.                                            actctgcaatcttactacatttttttttactttgtattttatttattccttttgagacag 27925
human.FAM230J-LOC105372942-GGTLC5P.NCBI.rev.compl                                 actctgcaatcttactacatttttttttactttgtattttatttattccttttgagacag 28644
human.FAM230A.ncRNA.18487127.18500594-GGTLC3.18516335.18518165.rev.compl.REF      actctgcaatcttactacatttttttttactttgtattttatttattccttttgagacag 17403
                                                                                  ************************************************************

chimp.LOC112206744-LOC107973052-GGT2.revcompl                                     agtctcactctgtcacccaggctgaagtgcagccgagatctcagctcactgcaacctcca 30617
chimp54546-135457.revcompl.Pan.troglodytes.clone.rp43-41g5.GenBank:AC099533.36    AGTCTCACTCTGTCACCCAGGCTGAAGTGCAGCCAAGATCTCGGCTCACTGCAACCTCCA 32336
human.FAM30B-LOC105372935-GGT2.NCBI.                                              agtctcactctgtcacccaggctgaagtgcagccgagatctcggctcactgcaacctcca 29251
human.FAM230E-LOC105377182-GGT3P.NCBI.                                            agtctcactctgtcacccaggctgaagtgcagccgagatctcggctcactgcaacctcca 27985
human.FAM230J-LOC105372942-GGTLC5P.NCBI.rev.compl                                 agtctcactctgtcacccaggctgaagtgcagccgagatctcggctcactgcaacctcca 28704
human.FAM230A.ncRNA.18487127.18500594-GGTLC3.18516335.18518165.rev.compl.REF      agtctcactctgtcacccaggctgaagtgcagccgagatctcggctcactgcaacctcca 17463
                                                                                  ********************************** ******* *****************

chimp.LOC112206744-LOC107973052-GGT2.revcompl                                     cctcctgggttcaagcaattctcttgtctcagcctcccaagtagctgggactacaggcac 30677
chimp54546-135457.revcompl.Pan.troglodytes.clone.rp43-41g5.GenBank:AC099533.36    CCTCCTGGGTTCAAGCAATTCTCTTGTCTCAGCCTCCCAAGTAGCTGGGACTACAGGCAC 32396
human.FAM30B-LOC105372935-GGT2.NCBI.                                              cctcttgggttcaagcaattctcttgtctcagcctcccaagtagctgggactacaggcac 29311
human.FAM230E-LOC105377182-GGT3P.NCBI.                                            cctcttgggttcaagcaattctcttgtctcagcctcccaagtagctgggactacaggcac 28045
human.FAM230J-LOC105372942-GGTLC5P.NCBI.rev.compl                                 cctcttgggttcaagcaattctcttgtctcagcctcccaagtagctgggactacaggcac 28764
human.FAM230A.ncRNA.18487127.18500594-GGTLC3.18516335.18518165.rev.compl.REF      cctcttgggttcaagcaattctcttgtctcagcctcccaagtagctgggactacaggcac 17523
                                                                                  **** *******************************************************

chimp.LOC112206744-LOC107973052-GGT2.revcompl                                     acgtcaaaacgcccggctaatttttgtatttttagtagagatggagtttagccatattgg 30737
chimp54546-135457.revcompl.Pan.troglodytes.clone.rp43-41g5.GenBank:AC099533.36    ACGTCAAAACGCCCGGCTAATTTTTGTATTTTTAGTAGAGATGGAGTTTTGCCATATTGG 32456
human.FAM30B-LOC105372935-GGT2.NCBI.                                              acgtcaaaacgcccggctaatttttgtatttttagtagagatggagttttgccatattgg 29371
human.FAM230E-LOC105377182-GGT3P.NCBI.                                            acgtcaaaacgcccggctaatttttgtatttttagtagagatggagttttgccctattgg 28105
human.FAM230J-LOC105372942-GGTLC5P.NCBI.rev.compl                                 acgtcaaaacgcccagctaatttttgtatttttagtagagatggagttttgccatattgg 28824
human.FAM230A.ncRNA.18487127.18500594-GGTLC3.18516335.18518165.rev.compl.REF      acgtcaaaacgcccggctaatttttgtatttttagtagagatggagttttgccatattgg 17583
                                                                                  ************** ********************************** *** ******

chimp.LOC112206744-LOC107973052-GGT2.revcompl                                     tcaggctggtctcgaactcctgacctcaggtgatcgacctgtcttagcctcccaaagtgc 30797
chimp54546-135457.revcompl.Pan.troglodytes.clone.rp43-41g5.GenBank:AC099533.36    TCAGGCTGGTCTCGAATTCCTGACCTCAGGTGATTGACCTGTCTTAGCCTCCCAAAGTGC 32516
human.FAM30B-LOC105372935-GGT2.NCBI.                                              tcaggctggtctcgaactcctgacctcaggtgatcgacctgtcttagcctcccaaagtgc 29431
human.FAM230E-LOC105377182-GGT3P.NCBI.                                            tcaggctggtctcaaactcctgacctcaggtgatcgacctgtcttagcctcccaaagtgc 28165
human.FAM230J-LOC105372942-GGTLC5P.NCBI.rev.compl                                 tcaggctggtctcgaactcctgacgtcaggtgatcgacctgtcttagcctcccaaagtgc 28884
human.FAM230A.ncRNA.18487127.18500594-GGTLC3.18516335.18518165.rev.compl.REF      tcaagctggtctcgaactcctgacgtcaggtgatcgacctgtcttagcctcccaaagtgc 17643
                                                                                  *** ********* ** ******* ********* *************************

chimp.LOC112206744-LOC107973052-GGT2.revcompl                                     taggattacaggtgtacatttatttatttatttgagatggaatcctgctctgtatttatt 30857
chimp54546-135457.revcompl.Pan.troglodytes.clone.rp43-41g5.GenBank:AC099533.36    TAGGATTACAGGTGTACATTTATTTATTTATTTGAGATAGAATCTTGCTCTGTATTTATT 32576
human.FAM30B-LOC105372935-GGT2.NCBI.                                              taggattacaggtgtacatttatttatttatttgagatggaatcttgctctgtatttatt 29491
human.FAM230E-LOC105377182-GGT3P.NCBI.                                            taggattacaggtgtacatttatttatttatttgagatggaatcttgctctgtatttatt 28225
human.FAM230J-LOC105372942-GGTLC5P.NCBI.rev.compl                                 taggattacaggtgtacatttatttatttatttgagatggaatcttgctctgtatttatt 28944
human.FAM230A.ncRNA.18487127.18500594-GGTLC3.18516335.18518165.rev.compl.REF      taggattacaggtgtacatttatttatttatttgagatggaatcttgctctgtatttatt 17703
                                                                                  ************************************** ***** ***************

chimp.LOC112206744-LOC107973052-GGT2.revcompl                                     aatttatttatttgagatggagtcttgctccattgcccaggctagagtgcagtggtgcaa 30917
chimp54546-135457.revcompl.Pan.troglodytes.clone.rp43-41g5.GenBank:AC099533.36    AATTTATTTATTTGAGATGGAGTCTTGCTCCATCGCCCAGGCTAGAGTGCAGTGGTGCAA 32636
human.FAM30B-LOC105372935-GGT2.NCBI.                                              aatttatttatttgagatggagtcttgctccatcgcccaggctagagtgcagcggtgcaa 29551
human.FAM230E-LOC105377182-GGT3P.NCBI.                                            aatttatttatttgagatggagtcttgctccatcgcccaggctagagtgcagcagtgcaa 28285
human.FAM230J-LOC105372942-GGTLC5P.NCBI.rev.compl                                 aatttatttagttgagatggagtcttgctccatcgcccaggctagagtgcagcggtgcaa 29004
human.FAM230A.ncRNA.18487127.18500594-GGTLC3.18516335.18518165.rev.compl.REF      aatttatttatttgagatggagtcttgctccatcgcccaggctagagtgcagcggtgcaa 17763
                                                                                  ********** ********************** ******************  ******

chimp.LOC112206744-LOC107973052-GGT2.revcompl                                     tctcggctcattgcaacctctgacttccaggttcaagcgattctcctgcctcagtgtccc 30977
chimp54546-135457.revcompl.Pan.troglodytes.clone.rp43-41g5.GenBank:AC099533.36    TCTCAGCTCATTGCAACCTCTGCCTTCCAGGTTCAAGCGATTCTCCTGCCTCAGTGTCCC 32696
human.FAM30B-LOC105372935-GGT2.NCBI.                                              tctcggctcattgcaacctctgccttccaggttcaagcgattctcctgcctcagtgtccc 29611
human.FAM230E-LOC105377182-GGT3P.NCBI.                                            tctcggctcattgcaacctctgccttccaggttcaagcgattctcctgcctcagtgtccc 28345
human.FAM230J-LOC105372942-GGTLC5P.NCBI.rev.compl                                 tctcggctcattgcaacctctgccttccaggttcaagcgattctcctgcctcagtgtccc 29064
human.FAM230A.ncRNA.18487127.18500594-GGTLC3.18516335.18518165.rev.compl.REF      tctcggctcattgcaacctctgccttccaggttcaagcgattctcctgcctcagtgtccc 17823
                                                                                  **** ***************** *************************************

chimp.LOC112206744-LOC107973052-GGT2.revcompl                                     aagtagctgggattacaggtgcctgccaccacagctggttaatttttgtatttttagtag 31037
chimp54546-135457.revcompl.Pan.troglodytes.clone.rp43-41g5.GenBank:AC099533.36    AAGTAGCTGGGATTACAGGTGCCTGCCACCACAGCTGGCTAATTTTTGTATTTTTAGTAG 32756
human.FAM30B-LOC105372935-GGT2.NCBI.                                              aagtagctgggattacaggtgcctgccaccacagctggctaatttttgtatttttagtag 29671
human.FAM230E-LOC105377182-GGT3P.NCBI.                                            aagtagctgggattacaggtgcctgccaccacagctggctaatttttgtatttttagtag 28405
human.FAM230J-LOC105372942-GGTLC5P.NCBI.rev.compl                                 aagtagctgggattacaggtgcctgccaccacagctggctaatttttgtatttttagtag 29124
human.FAM230A.ncRNA.18487127.18500594-GGTLC3.18516335.18518165.rev.compl.REF      aagtagctgggattacaggtgcctgccaccacagctggctaatttttgtatttttagtag 17883
                                                                                  ************************************** *********************

chimp.LOC112206744-LOC107973052-GGT2.revcompl                                     agacagtgtttcaccatcttggccaggctggtctcgggctcctgacctcatgaaccacct 31097
chimp54546-135457.revcompl.Pan.troglodytes.clone.rp43-41g5.GenBank:AC099533.36    AGACAGTGTTTCACCATCTTGGCCAGGCTGGTCTCAGGCTCCTGACCTCATGAACCACCT 32816
human.FAM30B-LOC105372935-GGT2.NCBI.                                              agacagtgtttcaccatcttggccaggctggtctcgggctcctgacctcatgaaccacct 29731
human.FAM230E-LOC105377182-GGT3P.NCBI.                                            agacagtgtttcaccatcttggccaggctggtctcgggctcctgacctcatgaaccacct 28465
human.FAM230J-LOC105372942-GGTLC5P.NCBI.rev.compl                                 agacagtgtttcaccatcttggccaggctggtctcgggctcctgacctcatgaaccacct 29184
human.FAM230A.ncRNA.18487127.18500594-GGTLC3.18516335.18518165.rev.compl.REF      agacagtgtttcaccatcttggccaggctggtctcgggctcctgacctcatgaaccacct 17943
                                                                                  *********************************** ************************

chimp.LOC112206744-LOC107973052-GGT2.revcompl                                     gcctcagcctcccaaagtgttgggattacaggcctaaggcaccatgttcggccatattta 31157
chimp54546-135457.revcompl.Pan.troglodytes.clone.rp43-41g5.GenBank:AC099533.36    GCCTCAGCCTCCCAAAGTGTTGGGATTACAGGCCTAAGGCACCATGCTCGGCCATATTTA 32876
human.FAM30B-LOC105372935-GGT2.NCBI.                                              gcctcagcctcccaaagtgttgggattacaggcctaaggcaccatgctcggccatattta 29791
human.FAM230E-LOC105377182-GGT3P.NCBI.                                            gcctcagcctcccaaagtgttgggattacaggcctaaggcaccatgctcggccatattta 28525
human.FAM230J-LOC105372942-GGTLC5P.NCBI.rev.compl                                 gcctcagcctcccaaaatgttgggattacaggcctaaggcaccatgctcggccatattta 29244
human.FAM230A.ncRNA.18487127.18500594-GGTLC3.18516335.18518165.rev.compl.REF      gcctcagcctcccaaagtgttgggattacaggcctaaggcaccatgctcggccatattta 18003
                                                                                  *************** ****************************** *************

chimp.LOC112206744-LOC107973052-GGT2.revcompl                                     tttatttaattatttagagacaaagtcttgctctgtcacccaggctggagtgcagtggcg 31217
chimp54546-135457.revcompl.Pan.troglodytes.clone.rp43-41g5.GenBank:AC099533.36    TTTATTTAATTATTTAGAGACAAAGTCTTGCTCTGTCACCCAGGCTGGAGTGCAGTGGCG 32936
human.FAM30B-LOC105372935-GGT2.NCBI.                                              tttatttaattatttagagacaaagtcttgctctgtcacccaggctggagtgcagtggcg 29851
human.FAM230E-LOC105377182-GGT3P.NCBI.                                            tttatttaattatttagagacaaagtcttgctctgtcacccaggctggagtgcagtggcg 28585
human.FAM230J-LOC105372942-GGTLC5P.NCBI.rev.compl                                 tttatttaattatttagagacaaagtcttgctctgtcacccaggctggagtgcagtggcg 29304
human.FAM230A.ncRNA.18487127.18500594-GGTLC3.18516335.18518165.rev.compl.REF      tttatttaattatttagagacaaagtcttgctctgtcacccaggctggagtgcagtggcg 18063
                                                                                  ************************************************************

chimp.LOC112206744-LOC107973052-GGT2.revcompl                                     ccatctcagctcactgcagcctccgcctctgaggtttaagcgattctcatgcctcagcct 31277
chimp54546-135457.revcompl.Pan.troglodytes.clone.rp43-41g5.GenBank:AC099533.36    CCATCTCAGCTCACTGCAGCCTCCGCCTCTGAGGTTTAAGCGATTCTCATGCCTCAGCCT 32996
human.FAM30B-LOC105372935-GGT2.NCBI.                                              ccatctcagctcactgcagcctccgcctccgaggtttaagcaattctcatgcctcagcct 29911
human.FAM230E-LOC105377182-GGT3P.NCBI.                                            ccatctcagctcactgcagcctccgcctccaaggtttaagcaattctcatgcctcagact 28645
human.FAM230J-LOC105372942-GGTLC5P.NCBI.rev.compl                                 ccatctcagctcactgcagcctccgcctccgaggtttaagcaattctcatgcctcagact 29364
human.FAM230A.ncRNA.18487127.18500594-GGTLC3.18516335.18518165.rev.compl.REF      ccatctcagctcactgcagcctccgcctccgaggtttaagcaattctcatgcctcagcct 18123
                                                                                  *****************************  ********** *************** **

chimp.LOC112206744-LOC107973052-GGT2.revcompl                                     cctgaataactaggactacagatactcgccaccacgcagggatttttttttttctatttt 31337
chimp54546-135457.revcompl.Pan.troglodytes.clone.rp43-41g5.GenBank:AC099533.36    CCTGAGTAACTAGGACTACAGATACTCGCCACCACACAGGGA-TTTTTTTTTTCTATTTT 33055
human.FAM30B-LOC105372935-GGT2.NCBI.                                              cctgagtaactgggactacagatacttgccaccacgcaggga--tttttttttctatttt 29969
human.FAM230E-LOC105377182-GGT3P.NCBI.                                            cctgagtaactgggactacagaaacttgccaccacgcaggga--tttttttttctatttt 28703
human.FAM230J-LOC105372942-GGTLC5P.NCBI.rev.compl                                 cctgagtaactgggactacagatacttgccaccacgcaggga-ttttttttttctatttt 29423
human.FAM230A.ncRNA.18487127.18500594-GGTLC3.18516335.18518165.rev.compl.REF      cctgagtaactgggactacagatacttgccaccacgcaggga-tttttttttctattttt 18182
                                                                                  ***** ***** ********** *** ******** ******  ********    ****

chimp.LOC112206744-LOC107973052-GGT2.revcompl                                     tttgtagagacacagtttca-ccatgttggccaggctggtctcgaactcctgaccttagg 31396
chimp54546-135457.revcompl.Pan.troglodytes.clone.rp43-41g5.GenBank:AC099533.36    TTTGTAGAGACACAGTTTCACCCATGTTGGCCAGGCTGGTCTCGAACTCCTGACCTTAGG 33115
human.FAM30B-LOC105372935-GGT2.NCBI.                                              tttgtagagacacagtttca-ccatgttggccaggctggtctcgaactcctgaccttagg 30028
human.FAM230E-LOC105377182-GGT3P.NCBI.                                            tttgtagagacacagtttca-ccatgttggccaggctggtctcgaactcctgactttagg 28762
human.FAM230J-LOC105372942-GGTLC5P.NCBI.rev.compl                                 tttgtagagacacagtttca-ccatgttggccaggctggtctcgaactcctgaccttagg 29482
human.FAM230A.ncRNA.18487127.18500594-GGTLC3.18516335.18518165.rev.compl.REF      tttgtagagacacagtttca-ccatgttggccaggctggtctcgaactcctgaccttagg 18241
                                                                                  ******************** ********************************* *****

chimp.LOC112206744-LOC107973052-GGT2.revcompl                                     tgatctgacagcctcgtcctctctaagtactgggattacaggcatgagccccttgcccgg 31456
chimp54546-135457.revcompl.Pan.troglodytes.clone.rp43-41g5.GenBank:AC099533.36    TGATCTGACAGCCTCGTCCTCTCTAAGTACTGGGATTACAGGCATGAGCCCCTTGCCCGG 33175
human.FAM30B-LOC105372935-GGT2.NCBI.                                              tgatctgacagcctcgtcctctcaaagcactgggattacaggcatgagccccttgcccgg 30088
human.FAM230E-LOC105377182-GGT3P.NCBI.                                            tgatctgacagcctcgtcctctcaaagcactgggattacaggcatgagccccttgcccgg 28822
human.FAM230J-LOC105372942-GGTLC5P.NCBI.rev.compl                                 tgatctgacagcctcgtcctctcaaagcactgggattacaggcatgagccccttgcccgg 29542
human.FAM230A.ncRNA.18487127.18500594-GGTLC3.18516335.18518165.rev.compl.REF      tgatctgacagcctcgtcctctcaaagcactgggattacaggcatgagccccttgcccgg 18301
                                                                                  *********************** *** ********************************

chimp.LOC112206744-LOC107973052-GGT2.revcompl                                     cctctcactacatttaagtgacgccatggctcatgcctgtaatcctagcactttgggagg 31516
chimp54546-135457.revcompl.Pan.troglodytes.clone.rp43-41g5.GenBank:AC099533.36    CCTCTCACTACATTTAAGTGACGCCATGGCTCATGCCTGTAATCCTAGCACTTTGGGAGG 33235
human.FAM30B-LOC105372935-GGT2.NCBI.                                              cctctcactacatttaagtgacgccatggctcatgcctgtaatcctagcactttgggagg 30148
human.FAM230E-LOC105377182-GGT3P.NCBI.                                            cctctcactacatttaagtgacgccatggctcatgcctgtaatcctagcactttgggagg 28882
human.FAM230J-LOC105372942-GGTLC5P.NCBI.rev.compl                                 cctctcactacgtttaagtgatgccatggctcatgcctgtaatcctagcactttgggagg 29602
human.FAM230A.ncRNA.18487127.18500594-GGTLC3.18516335.18518165.rev.compl.REF      cctctcactacatttaagtgacgccatggctcatgcctgtaatcctagcactttgggagg 18361
                                                                                  *********** ********* **************************************
```

```
chimp.LOC112206744-LOC107973052-GGT2.revcompl                                   ccaaggcaggtggatcacctgaggtcaggagttcgacacgagcctggccaacatggggaa 31576
chimp54546-135457.revcompl.Pan.troglodytes.clone.rp43-41g5.GenBank:AC099533.36  CCAAGGCAGGTGGATCACCTGAGGTCAGGAGTTCGACACAAGCCTGGCCAACATGGGGAA 33295
human.FAM30B-LOC105372935-GGT2.NCBI.                                            ccaaggcaggtggatcacctgaggtcaggagttcgacacgagcctggccaacatggggaa 30208
human.FAM230E-LOC105377182-GGT3P.NCBI.                                          ccaaggcaggtggatcacctgaggtcaggagttcgacatgagcctggccaacatggggaa 28942
human.FAM230J-LOC105372942-GGTLC5P.NCBI.rev.compl                               ccaaggcaggtggatcacctgaggtcaggagttcgacacgagcctggccaacatggggaa 29662
human.FAM230A.ncRNA.18487127.18500594-GGTLC3.18516335.18518165.rev.compl.REF    ccaaggcaggtggatcacctgaggtcaggagttcgacatgagcctggccaacatggggaa 18421
                                                                                **************************************  ********************

chimp.LOC112206744-LOC107973052-GGT2.revcompl                                   accccgtctctagtaaaaatacaaaaattagtcacgcgtggtggtacaagcctgtaggcc 31636
chimp54546-135457.revcompl.Pan.troglodytes.clone.rp43-41g5.GenBank:AC099533.36  ACCCCGTCTCTAGTAAAAATACAAAAATTAGTCACGCGTGGTGGTACAAGCCTGTAGGCC 33355
human.FAM30B-LOC105372935-GGT2.NCBI.                                            accccgtctctagtaaaaatacaaaaattagtcaggtgtggtggtacaagcctgtaggcc 30268
human.FAM230E-LOC105377182-GGT3P.NCBI.                                          accccgtctctagtaaaaatacaaaaattagtcaggtgtggtggtacaagcctgtaggcc 29002
human.FAM230J-LOC105372942-GGTLC5P.NCBI.rev.compl                               accctgtctctagtaaaaatacaaaaattagtcaggcgtggtggtacaagcctgtaggcc 29722
human.FAM230A.ncRNA.18487127.18500594-GGTLC3.18516335.18518165.rev.compl.REF    accccgtctctagtaaaaatacaaaaattagtcaggtgtggtggtacaagcctgtaggcc 18481
                                                                                **** ***************************** * ***********************

chimp.LOC112206744-LOC107973052-GGT2.revcompl                                   cagctacttggaagactgaggcaggagaatcactttaagcgggaggcagaggttgcagtg 31696
chimp54546-135457.revcompl.Pan.troglodytes.clone.rp43-41g5.GenBank:AC099533.36  CAGCTACTTGGAAGACTGAGGCAGGAGAATCACTTTAACCGGAGGCAGAGGTTGCAGTG 33415
human.FAM30B-LOC105372935-GGT2.NCBI.                                            cagctacttggaagactgaggcaggagaatcactttaagcgggaggcagaggttgcagtg 30328
human.FAM230E-LOC105377182-GGT3P.NCBI.                                          cagctacttggaagactgaggcaggagaatcactttaagcgggaggcagaggttgcagtg 29062
human.FAM230J-LOC105372942-GGTLC5P.NCBI.rev.compl                               cagctacttggaagactgaggcaggagaatcactttaagcgggaggcagaggttgcagtg 29782
human.FAM230A.ncRNA.18487127.18500594-GGTLC3.18516335.18518165.rev.compl.REF    cagctacttggaagactgaggcaggagaatcactttaagcgggaggcagaggttgcagtg 18541
                                                                                ************************************** *********************

chimp.LOC112206744-LOC107973052-GGT2.revcompl                                   agccaatatcatgccactgcactccagcttgggtgacggagtgagacactgtctc----- 31751
chimp54546-135457.revcompl.Pan.troglodytes.clone.rp43-41g5.GenBank:AC099533.36  AGCCAATATCATGCCACTGCACTCCAGCTTGGGTGACAGAGTGAGACACTGTCTCAAAAA 33475
human.FAM30B-LOC105372935-GGT2.NCBI.                                            agccaatatcatgccactgcactccagcttgggtgacagagtgagatactgtctc----a 30384
human.FAM230E-LOC105377182-GGT3P.NCBI.                                          agccaatatcatgccactgcactccagcttgggtgacagagtgagatactgtctc----a 29118
human.FAM230J-LOC105372942-GGTLC5P.NCBI.rev.compl                               agccaatatcatgccactgcactccagcttgggtgacagagtgagatactgtctc----a 29838
human.FAM230A.ncRNA.18487127.18500594-GGTLC3.18516335.18518165.rev.compl.REF    agccaatatcatgccactgcactccagcttgggtgacagagtgagatactgtctc----a 18597
                                                                                ************************************* ******** ********

chimp.LOC112206744-LOC107973052-GGT2.revcompl                                   ------aaaaaaaaaaaagagaaaaaaatatgatgccggggcatctcggcctcaatacct 31805
chimp54546-135457.revcompl.Pan.troglodytes.clone.rp43-41g5.GenBank:AC099533.36  AAAAAAAAGAAAAAAAAAGAGAAAAAAATATGATGCCGGGGCATCACGGCCTCAATACCT 33535
human.FAM30B-LOC105372935-GGT2.NCBI.                                            aaaaaaaagaaaaaaagagagaaaaacatatgatgccggggcatctcggcctcaatacct 30444
human.FAM230E-LOC105377182-GGT3P.NCBI.                                          aaaaaaaagaaaaaaagagagaaaaacatatgatgccggggcatctcggcctcaatacct 29178
human.FAM230J-LOC105372942-GGTLC5P.NCBI.rev.compl                               aaaaaaaagaaaaaataagagaaaaacatatgatgccggggcatctcggcctcaatacct 29898
human.FAM230A.ncRNA.18487127.18500594-GGTLC3.18516335.18518165.rev.compl.REF    aaaaaaaagaaaaaaagagagaaaaacatatgatgccagggcatctcggcctcaatacct 18657
                                                                                      ** ******  ********* ********** ******* **************

chimp.LOC112206744-LOC107973052-GGT2.revcompl                                   gcgtgagcacagtcatgtccaggccagggctgctggtcgaggtccggtcccatctcttcc 31865
chimp54546-135457.revcompl.Pan.troglodytes.clone.rp43-41g5.GenBank:AC099533.36  GCGTGAGCACAGTCATGTCCAGGCCAGGGCTGCTGGTCGAGGTCCGGCCCCATCTCTTCC 33595
human.FAM30B-LOC105372935-GGT2.NCBI.                                            gggtgagcacagtcatgtccaggccagggctgctggtcgaggtccggccccatctcttcc 30504
human.FAM230E-LOC105377182-GGT3P.NCBI.                                          gggtgagcacagtcatgtccaggccagggctgctggtcgaggtccggccccatctcttcc 29238
human.FAM230J-LOC105372942-GGTLC5P.NCBI.rev.compl                               gggtgagcacagtcatgtccaggccagggctgctggttgaggtccggccccatctcttcc 29958
human.FAM230A.ncRNA.18487127.18500594-GGTLC3.18516335.18518165.rev.compl.REF    gggtgagcacagtcatgtccaggccagggctgctggtcgaggtccggccccatctcttcc 18717
                                                                                * *********************************** ********* ************

chimp.LOC112206744-LOC107973052-GGT2.revcompl                                   agcagaaagggagtaagcttgcagggaggctggggacaagatcccaggatctcggcctc 31925
chimp54546-135457.revcompl.Pan.troglodytes.clone.rp43-41g5.GenBank:AC099533.36  AGCAGAAAGGGAGTAAGCTTGCGGGCGGCTGGGGGACAAGATCCCAGGATCTCGGCCTC 33655
human.FAM30B-LOC105372935-GGT2.NCBI.                                            agcagaaagggagtaagcttgcagggcggctggggacaagatcccaggatctcagcctc 30564
human.FAM230E-LOC105377182-GGT3P.NCBI.                                          agcagaaagggagtaagcttgcagggcggctggggacaagatcccaggatctcagcctc 29298
human.FAM230J-LOC105372942-GGTLC5P.NCBI.rev.compl                               agcagaaagggagtaagcttgcagggcggctggggacaagatcccaggatctcagcctc 30018
human.FAM230A.ncRNA.18487127.18500594-GGTLC3.18516335.18518165.rev.compl.REF    agcagaaagggagtaagcttgcagggaggctggggacaagatcccaggatctcagcctc 18777
                                                                                ********************** *** *************************** *****

chimp.LOC112206744-LOC107973052-GGT2.revcompl                                   cgctcatggatcagctctgataccccgagtgagctgggggtgctctgtgcgcattggttt 31985
chimp54546-135457.revcompl.Pan.troglodytes.clone.rp43-41g5.GenBank:AC099533.36  TGCTCATGGATCAGCTCTGAGACCCCGAGTGAGCTGGGGGTGCTCTGTGCGCATCGGTTT 33715
human.FAM30B-LOC105372935-GGT2.NCBI.                                            tgctcatggatcagctctgagaccccgagtgagctgggggtgctctgtgcgcattggttt 30624
human.FAM230E-LOC105377182-GGT3P.NCBI.                                          tgctcatggatcagctctgagaccccgagtgagctgggggtgctctgtgcgcattggttt 29358
human.FAM230J-LOC105372942-GGTLC5P.NCBI.rev.compl                               tgctcatggatcagctctgagaccccgagtgagctgggggtgctctgtgcgcgttggttt 30078
human.FAM230A.ncRNA.18487127.18500594-GGTLC3.18516335.18518165.rev.compl.REF    tgctcatggatcagctctgagaccccgagtgagctgggggtgctctgtgcgcattggttt 18837
                                                                                 ******************* ******************************* * *****

chimp.LOC112206744-LOC107973052-GGT2.revcompl                                   ccccagctgtcaagtaaagggattgtatgaggaagtcttgtcaaggtggaatgatctcag 32045
chimp54546-135457.revcompl.Pan.troglodytes.clone.rp43-41g5.GenBank:AC099533.36  CCCCAGCTGTCAAGTAAAGGGATTGGATGAGGAAGTCTTGTCAAGGTGGAATGATCTCAG 33775
human.FAM30B-LOC105372935-GGT2.NCBI.                                            ccccagctgtcaagtaaagggattggatgaggaagtcttgtcaaggtggaatgatctcag 30684
human.FAM230E-LOC105377182-GGT3P.NCBI.                                          ccccagctgtcaagtaaagggattggatgaggaagtcttgtcaaggtggaatgatctcag 29418
human.FAM230J-LOC105372942-GGTLC5P.NCBI.rev.compl                               ccccagctgtcaagtaaagggattggatgaggaagtcttgtcaaggtggaatgatgtcag 30138
human.FAM230A.ncRNA.18487127.18500594-GGTLC3.18516335.18518165.rev.compl.REF    ccccagctgtcaagtaaagagattggatgaggaagtcttgtcaaggtggaatgatctcag 18897
                                                                                ******************* ***** ***************************** ****

chimp.LOC112206744-LOC107973052-GGT2.revcompl                                   atttggggcagcagtgaatgatcccgctccctgggccatgccagtggcccggcctcggct 32105
chimp54546-135457.revcompl.Pan.troglodytes.clone.rp43-41g5.GenBank:AC099533.36  CTTTGGGGCAGCAGTGAATGATCCCGCTTCCTGGGCCATGCCAGTGGCCCGGCCTCGGCT 33835
human.FAM30B-LOC105372935-GGT2.NCBI.                                            atttggggcagcagtgaatgatcccgctccctgggccatgccagtggcccggcctcggct 30744
human.FAM230E-LOC105377182-GGT3P.NCBI.                                          atttggggcagcagtgaatgatcccgctccctgggccatgccagtggcccggcctcggct 29478
human.FAM230J-LOC105372942-GGTLC5P.NCBI.rev.compl                               atttggggcagcagtgaatgatcccgttccttgggccatgccagtggcctggcctcggct 30198
human.FAM230A.ncRNA.18487127.18500594-GGTLC3.18516335.18518165.rev.compl.REF    atttggggcagcagtgaatgatcccgctccctgggccatgccagtggcccggcctcggct 18957
                                                                                 ************************* * * ****************** **********

chimp.LOC112206744-LOC107973052-GGT2.revcompl                                   caacacagccccaacactctggaatggggatgagggggcagtcagctcttgctcctagta 32165
chimp54546-135457.revcompl.Pan.troglodytes.clone.rp43-41g5.GenBank:AC099533.36  CAACACAGCCCCAACACTCTGGAATGGGGATGAGGGGGCAGTCAGCTCTTGCTCCTAGTA 33895
human.FAM30B-LOC105372935-GGT2.NCBI.                                            gaacacagccccaacactctggaatggggatgagggggcagtcagctcttgctcctagta 30804
human.FAM230E-LOC105377182-GGT3P.NCBI.                                          gaacacagccccaacactctggaatggggatgagggggcagtcagctcttgctcctagta 29538
human.FAM230J-LOC105372942-GGTLC5P.NCBI.rev.compl                               caacacagccccaacactctggaatggggatgagggggcagtcagctcttgctcctagta 30258
human.FAM230A.ncRNA.18487127.18500594-GGTLC3.18516335.18518165.rev.compl.REF    gaacacagccccaacactctggaatggggatgagggggcagtcagctcttgctcctagta 19017
                                                                                 ***********************************************************

chimp.LOC112206744-LOC107973052-GGT2.revcompl                                   agagagatgcaacggggctctgtggctgagctgggtgccttgcctcacacctgtaatccc 32225
chimp54546-135457.revcompl.Pan.troglodytes.clone.rp43-41g5.GenBank:AC099533.36  AGAGAGATGCAACAGGGCTCTGTGGCTGAGCTGGGTGCCTTGCCTCACACCTGTAATCCC 33955
human.FAM30B-LOC105372935-GGT2.NCBI.                                            agagagatgcaacagggctctgtggctgagctgggtgccttgcctcacacctgtaatccc 30864
human.FAM230E-LOC105377182-GGT3P.NCBI.                                          agagagatgcaacagggctctgtggctgagctgggtgccttgcctcacacctgtaatccc 29598
human.FAM230J-LOC105372942-GGTLC5P.NCBI.rev.compl                               agagagatgcaacagggctctgtggctgagctgggtgccttgcctcacacctgtaatccc 30318
human.FAM230A.ncRNA.18487127.18500594-GGTLC3.18516335.18518165.rev.compl.REF    agagagatgcaacagggctctgtggctgagctgggtgccttgcctcacacctgtaatccc 19077
                                                                                ************* **********************************************

chimp.LOC112206744-LOC107973052-GGT2.revcompl                                   aacctttgagaggctgaggcaggaggattgctcgaggccgggaattttgagaatagcctg 32285
chimp54546-135457.revcompl.Pan.troglodytes.clone.rp43-41g5.GenBank:AC099533.36  AACCTTTGAGAGGCCGAGGCAGGAGGATTGCTCGAGGCCGGGAATTTTGAGAATAGCCTG 34015
human.FAM30B-LOC105372935-GGT2.NCBI.                                            aacctttgagaggccgaggcaggaggattgctcgaggccgggaattttgagaatagcctg 30924
human.FAM230E-LOC105377182-GGT3P.NCBI.                                          aacctttgagaggccgaggcaggaggattgctcgagaccgggaattttgagaatagcctg 29658
human.FAM230J-LOC105372942-GGTLC5P.NCBI.rev.compl                               aacctttgagaggccgaggcaggaggattgctcgaggccgggaattttgagaatagcctg 30378
human.FAM230A.ncRNA.18487127.18500594-GGTLC3.18516335.18518165.rev.compl.REF    aacctttgagaggccgaggcaggaggattgctcgaggccgggaattttgagaatagcctg 19137
                                                                                ************** ********************* ***********************

chimp.LOC112206744-LOC107973052-GGT2.revcompl                                   gacaacatagccagaccccatgtctacaaaataataataaaacatacagctatagtccag 32345
chimp54546-135457.revcompl.Pan.troglodytes.clone.rp43-41g5.GenBank:AC099533.36  GACAACATAGCCAGACCCCATGTCTACAAAATAATAATAAAACATACAGCTATAGTCCAA 34075
human.FAM30B-LOC105372935-GGT2.NCBI.                                            gacaacatagccagaccccatgtctacaaaataataagaaaacacacagctatagtccaa 30984
human.FAM230E-LOC105377182-GGT3P.NCBI.                                          gacaacatagccagaccccatgtctacaaaataataataaaacacacagctatagtccaa 29718
human.FAM230J-LOC105372942-GGTLC5P.NCBI.rev.compl                               gacaacatagccagaccccatgtctacaaaataataataaaacatacagctatagtccaa 30438
human.FAM230A.ncRNA.18487127.18500594-GGTLC3.18516335.18518165.rev.compl.REF    gacaacatagccagaccccatgtctacaaaataataataaaacacacagctatagtccaa 19197
                                                                                ************************************* ****** **************

chimp.LOC112206744-LOC107973052-GGT2.revcompl                                   gctacttggcaggctgaggcaggagggtcccttgagtccaggaattggaggctgcattga 32405
chimp54546-135457.revcompl.Pan.troglodytes.clone.rp43-41g5.GenBank:AC099533.36  GCTACTTGGCAGGCTGAGGCAGGAGGGTCCCTTGAGTCCAGGAATTGGAGGCTGCATTGA 34135
human.FAM30B-LOC105372935-GGT2.NCBI.                                            gctacttggcaggctgaggcaggagggtcccttgagtccaggaattggaggctgcattga 31044
human.FAM230E-LOC105377182-GGT3P.NCBI.                                          gctacttggcaggctgaggcaggagggtcccttgagtccaggaattggaggctgcattga 29778
human.FAM230J-LOC105372942-GGTLC5P.NCBI.rev.compl                               gctacttggcaggctgaggcaggagggtcccttgagtccaggaattggaggctgcattga 30498
human.FAM230A.ncRNA.18487127.18500594-GGTLC3.18516335.18518165.rev.compl.REF    gctacttggcaggctgaggcaggagggtcccttgagtccaggaattggaggctgcattga 19257
                                                                                ************************************************************

chimp.LOC112206744-LOC107973052-GGT2.revcompl                                   gctataatcgcaccactgcactccagcttgggtgacaaagtgagaccctgtctctaaaag 32465
chimp54546-135457.revcompl.Pan.troglodytes.clone.rp43-41g5.GenBank:AC099533.36  GCTATAATCGTACCACTGCACTCCAGCTTGGGTGACAAAGTGAGACCCTGTCTCTAAAAG 34195
human.FAM30B-LOC105372935-GGT2.NCBI.                                            gctataatcgcaccactgcactccagcttgggtgacaaagtgagaccctgtctctaaaag 31104
human.FAM230E-LOC105377182-GGT3P.NCBI.                                          gctataatcgcaccactgcactccagcttgggtgacaaagtgagaccctgtctctaaaag 29838
human.FAM230J-LOC105372942-GGTLC5P.NCBI.rev.compl                               gctataatcgcaccactgcactccagcttgggtgacaaagtgagaccctgtctctaaaag 30558
human.FAM230A.ncRNA.18487127.18500594-GGTLC3.18516335.18518165.rev.compl.REF    gctataatcgcaccactgcactccagcttgggtgacaaagtgagaccctgtctctaaaag 19317
                                                                                ********** *************************************************

chimp.LOC112206744-LOC107973052-GGT2.revcompl                                   -aaaaaacattggcctgtgagcatgggtttgagtttcaaacaggagccggagggtagggt 32524
chimp54546-135457.revcompl.Pan.troglodytes.clone.rp43-41g5.GenBank:AC099533.36  AAAAAAAAATTGGCCTGTGAGCATGGGTTTGATTTTTAAACAGGACCCAGAGGGTAGGGT 34255
human.FAM30B-LOC105372935-GGT2.NCBI.                                            -aaaaaaaattggcctgtgagcatgggcttgattttcaaacaggacccggagggtagggt 31163
human.FAM230E-LOC105377182-GGT3P.NCBI.                                          aaaaaaaaattggcctgtgagcatgggcttgattttcaaacaggacccggagggtagggt 29898
human.FAM230J-LOC105372942-GGTLC5P.NCBI.rev.compl                               aaaaaaaaattggcctgtgagcatgggcttgattttcaaacaggacccggagggtagggt 30618
human.FAM230A.ncRNA.18487127.18500594-GGTLC3.18516335.18518165.rev.compl.REF    aaaaaaaaattggcctgtgagcatgggcttgattttcaaacaggacccggagggtagggt 19377
                                                                                 ****** ******************* **** *** ******** ** ***********

chimp.LOC112206744-LOC107973052-GGT2.revcompl                                   aaacttgtcggtaaatctaaatgaatgttattggtataaaattacaatagtatagaaaat 32584
chimp54546-135457.revcompl.Pan.troglodytes.clone.rp43-41g5.GenBank:AC099533.36  AAACGTGTGGGTAAATCTAAATGAATGCTATTGGTATAAAATTACAGTAGTATAGAAAAT 34315
human.FAM30B-LOC105372935-GGT2.NCBI.                                            aaacgtgtgggtaaatctaaatgaatgttattggtataaaattacagtagtatagaaaat 31223
human.FAM230E-LOC105377182-GGT3P.NCBI.                                          aaacgtgtgggtaaatctaaatgaatgttattggtataaaattacagtagtatagaaaat 29958
human.FAM230J-LOC105372942-GGTLC5P.NCBI.rev.compl                               aaacgtgtgggtaaatctaaatgaatgttattggtataaaattacagtagtatagaaaat 30678
human.FAM230A.ncRNA.18487127.18500594-GGTLC3.18516335.18518165.rev.compl.REF    aaacgtgtgggtaaatctaaatgaatgttattggtataaaattacagtagtatagaaaat 19437
                                                                                **** *** ****************** ****************** *************
```

```
chimp.LOC112206744-LOC107973052-GGT2.revcompl                                      gatatcttgtggggtttaaaataaataaaacattctgaaatatgtatgggtacagttata 32644
chimp54546-135457.revcompl.Pan.troglodytes.clone.rp43-41g5.GenBank:AC099533.36     GATATCTTGTGGGGTTTAAAATAAATGAAACATACTGAAATATGTATGGGTACAGTTATA 34375
human.FAM30B-LOC105372935-GGT2.NCBI.                                               gatatcttgtggggtttaaaataaataaaacatactgaaatatgtattggtacagttata 31283
human.FAM230E-LOC105377182-GGT3P.NCBI.                                             gatatcttgtggggtttaaaagaaataaaacatactgaaatatgtatgggtacagttata 30018
human.FAM230J-LOC105372942-GGTLC5P.NCBI.rev.compl                                  gatatcttgtggggtttaaaagaaataaaacatactgaaatatgtatgggtacagttata 30738
human.FAM230A.ncRNA.18487127.18500594-GGTLC3.18516335.18518165.rev.compl.REF       gatatcttgtggggtttaaaataaataaaacatactgaaatatgtatgggtacagttata 19497
                                                                                   ********************* **** ****** ************* ************

chimp.LOC112206744-LOC107973052-GGT2.revcompl                                      tatctgggatttgcactgaaataatgtggggtagagggaagcaggaaagaggatacatga 32704
chimp54546-135457.revcompl.Pan.troglodytes.clone.rp43-41g5.GenBank:AC099533.36     TATCTGGGATTTGCACTGAAATAATGTGGGGTAGAGGGAAGCAGGAAAGAGGATACATGA 34435
human.FAM30B-LOC105372935-GGT2.NCBI.                                               tatctgggatttgcactgaaataatgtggggtagagggaagcaggaaagagtatacatga 31343
human.FAM230E-LOC105377182-GGT3P.NCBI.                                             tatctgggatttgcactgaaataatgtggggtagagggaagcaggaaagagtatacatga 30078
human.FAM230J-LOC105372942-GGTLC5P.NCBI.rev.compl                                  tatctgggatttgcacggaaataatgtggggtagagggaagcaggaaagagtatacatga 30798
human.FAM230A.ncRNA.18487127.18500594-GGTLC3.18516335.18518165.rev.compl.REF       tatctgggatttgcactgaaataatgtggggtagagggaagcaggaaagagtatacatga 19557
                                                                                   **************** *********************************** ********

chimp.LOC112206744-LOC107973052-GGT2.revcompl                                      aattagcttggccataagattgttgttgaaattgaatggatactcggggcttcattacac 32764
chimp54546-135457.revcompl.Pan.troglodytes.clone.rp43-41g5.GenBank:AC099533.36     AATTAGCTTGGCCATAAGATTGTTGTTGAAATTGAATGGATACTCGGGGCTTCATTACAC 34495
human.FAM30B-LOC105372935-GGT2.NCBI.                                               aatgagcttggccataagattgttgttgaaattgaatggatactcggggcttcattacac 31403
human.FAM230E-LOC105377182-GGT3P.NCBI.                                             aatgagcttggccataagattgttgttgaaattgaatggatactcggggcttcattacac 30138
human.FAM230J-LOC105372942-GGTLC5P.NCBI.rev.compl                                  aatgagcttggccataagattgttgttgaaattgaatggatactcggggcttcattacac 30858
human.FAM230A.ncRNA.18487127.18500594-GGTLC3.18516335.18518165.rev.compl.REF       aatgagcttggccataagattgttgttgaaattgaatggatactctgggcttcattacac 19617
                                                                                   *** ***************************************** **************

chimp.LOC112206744-LOC107973052-GGT2.revcompl                                      aattcttttactcttacatagctctacactctcaacataaataagaataaaaacacaaa 32824
chimp54546-135457.revcompl.Pan.troglodytes.clone.rp43-41g5.GenBank:AC099533.36     AATTCTCTTTACTCTTACATAGCTCTACACTCTCAACATAAATAAGAATAAAAACACAAA 34555
human.FAM30B-LOC105372935-GGT2.NCBI.                                               aattctctttactcttacatagctctacactctcaacataaataagaataaaaacacaaa 31463
human.FAM230E-LOC105377182-GGT3P.NCBI.                                             aattctctttactcttacatagctctacactctcaacataaataagaataaaaacacaaa 30198
human.FAM230J-LOC105372942-GGTLC5P.NCBI.rev.compl                                  aattctctttactcttacatagctctacactctcaacataaataagaataaaaacacaaa 30918
human.FAM230A.ncRNA.18487127.18500594-GGTLC3.18516335.18518165.rev.compl.REF       aattctctttactcttacatagctctacactctcaacataaataagaataaaaacacaaa 19677
                                                                                   ****** *****************************************************

chimp.LOC112206744-LOC107973052-GGT2.revcompl                                      aaacacacagatacatctatgcacacacatgtattttaaatacacaaaaatattagcata 32884
chimp54546-135457.revcompl.Pan.troglodytes.clone.rp43-41g5.GenBank:AC099533.36     AAACACACAGATACATCTATGCACACACATATATTTAAAATACACAAAAATATTAGCATA 34615
human.FAM30B-LOC105372935-GGT2.NCBI.                                               aaacacacagatacatctatgcacacacacatatttaaaatacacaaaaatattagcata 31523
human.FAM230E-LOC105377182-GGT3P.NCBI.                                             aaacacacagatacatctatgcacacacacatatttaaaatacacaaaaatattagcata 30258
human.FAM230J-LOC105372942-GGTLC5P.NCBI.rev.compl                                  aaacacacagatacatctatgcacacacacatatttaaaatacacaaaaatattagcata 30978
human.FAM230A.ncRNA.18487127.18500594-GGTLC3.18516335.18518165.rev.compl.REF       aaacacacagatacatctatgcacacacacatatttaaaatacacaaaaatattagcata 19737
                                                                                   *****************************  ***** ***********************

chimp.LOC112206744-LOC107973052-GGT2.revcompl                                      taagtcactggggggtaaatttagttcctgttccaaggttcttgtactgactaggaagagg 32944
chimp54546-135457.revcompl.Pan.troglodytes.clone.rp43-41g5.GenBank:AC099533.36     TAAGTCACTGGGGGTAAATTTAGTTCCTGTTCCAAGGTTCTTGTACTGACTAGGAAGAGG 34675
human.FAM30B-LOC105372935-GGT2.NCBI.                                               taagtcactgggggtaaatttagttcctgttccaaggttcttgtactgactaggaagagg 31583
human.FAM230E-LOC105377182-GGT3P.NCBI.                                             taagtcactgggggtaaatttagttcctgttccaaggttcttgtactgactaggaagagg 30318
human.FAM230J-LOC105372942-GGTLC5P.NCBI.rev.compl                                  taagtcactgggggtaaatttagttcctgttccaaggttcttgtactgactaggaagagg 31038
human.FAM230A.ncRNA.18487127.18500594-GGTLC3.18516335.18518165.rev.compl.REF       taagtcactgggggtaaatttagttcctgttccaaggttcttgtactgactaggaagagg 19797
                                                                                   ************************************************************

chimp.LOC112206744-LOC107973052-GGT2.revcompl                                      atagaagtactaactcataggctgggcgtggtggctcacgcctgtaatcccaacacttta 33004
chimp54546-135457.revcompl.Pan.troglodytes.clone.rp43-41g5.GenBank:AC099533.36     ATAGAAGTACTAACTCATAGGCTGGGCGTGGTGGCTCACGCCTGTAATCCCAACACTTTA 34735
human.FAM30B-LOC105372935-GGT2.NCBI.                                               atagaagtactaactcataggctgggcgcggtggctcacgcctgtaatcccaacacttta 31643
human.FAM230E-LOC105377182-GGT3P.NCBI.                                             atagaagtactaactcataggctgggtgcggtggctcacgcctgtaatcccaacacttta 30378
human.FAM230J-LOC105372942-GGTLC5P.NCBI.rev.compl                                  atagaagtactaactcataggctgggtgcggtggctcacgcctgtaatcccaacacttta 31098
human.FAM230A.ncRNA.18487127.18500594-GGTLC3.18516335.18518165.rev.compl.REF       atagaagtactaactcataggctgggcgcggtggctcacgcctgtaatcccaacacttta 19857
                                                                                   *************************** * ******************************

chimp.LOC112206744-LOC107973052-GGT2.revcompl                                      ggacgccgaggtaagcagatctcttaaggtcaggagttcaagaccagcctggccaacatg 33064
chimp54546-135457.revcompl.Pan.troglodytes.clone.rp43-41g5.GenBank:AC099533.36     GGACGCCGAGGTAAGCAGATCTCTTAAGGTCAGGAGTTCAAGACCAGCCTGGCCAACATG 34795
human.FAM30B-LOC105372935-GGT2.NCBI.                                               ggatgccgaggtaggcagatctcttaaggtccggagttcaagaccagcctggccaacatg 31703
human.FAM230E-LOC105377182-GGT3P.NCBI.                                             ggatgccgaggtaggcagatctcttaaggtccggagttcaagaccagcctggccaacatg 30438
human.FAM230J-LOC105372942-GGTLC5P.NCBI.rev.compl                                  ggatgccgaggtaggcagatctcttaaggtccggagttcaagaccagcctggccaacatg 31158
human.FAM230A.ncRNA.18487127.18500594-GGTLC3.18516335.18518165.rev.compl.REF       ggatgccgaggtaggcagatctcttaaggtccggagttcaagaccagcctggccaacatg 19917
                                                                                   *** ********* ***************** ****************************

chimp.LOC112206744-LOC107973052-GGT2.revcompl                                      gtgaaccctgtctctactaaaaaagaatacaaaaattggccgggcatagtggtgcacac 33124
chimp54546-135457.revcompl.Pan.troglodytes.clone.rp43-41g5.GenBank:AC099533.36     GTGAAACCCTGTCTCTACTAAAAAAGAATACAAAAATTGGCCGGGCATAGTGGTGCACAC 34855
human.FAM30B-LOC105372935-GGT2.NCBI.                                               gtgaaaccctgtctctactgaaaaagaatacaaaaattggccgggcatagtggtgcacac 31763
human.FAM230E-LOC105377182-GGT3P.NCBI.                                             gtgaaaccctgtctctactgaaaaagaatacaaaaattggccgggcatagtggtgcacac 30498
human.FAM230J-LOC105372942-GGTLC5P.NCBI.rev.compl                                  gtgaaaccctgtctctactgaaaaagaatacaaaaattggccgggcatagtggtgcacac 31218
human.FAM230A.ncRNA.18487127.18500594-GGTLC3.18516335.18518165.rev.compl.REF       gtgaaaccctgtctctactgaaaaagaatacaaaaattggccgggcatagtggtgcacac 19977
                                                                                   ****************** *****************************************

chimp.LOC112206744-LOC107973052-GGT2.revcompl                                      ctgtggtcccagctactcaggtgactgaggcaggagaattgcttgaacccaagaagtgga 33184
chimp54546-135457.revcompl.Pan.troglodytes.clone.rp43-41g5.GenBank:AC099533.36     CTGTGGTCCCAGCTACTCAGGTGACTGAGGCAGGAGAATTGCTTGAACCCAAGAAGTGGA 34915
human.FAM30B-LOC105372935-GGT2.NCBI.                                               ctgtggtcccagctactcaggtgactgaggcaggagaattgcttgaacccaagaagtgga 31823
human.FAM230E-LOC105377182-GGT3P.NCBI.                                             ctgtggtcccagctactcaggtgactgaggcaggagaattgcttgaacccaagaagtgga 30558
human.FAM230J-LOC105372942-GGTLC5P.NCBI.rev.compl                                  ctgtggtcccagctactcaggtgactgaggcaggagaattgcttgaacccaagaagtgga 31278
human.FAM230A.ncRNA.18487127.18500594-GGTLC3.18516335.18518165.rev.compl.REF       ctgtggtcccagctactcaggtgactgaggcaggagaattgcttgaacccaagaagtgga 20037
                                                                                   ************************************************************

chimp.LOC112206744-LOC107973052-GGT2.revcompl                                      ggttgcagtgaaccaagattgctccactgcactccagcctgggcagcagaggaagactct 33244
chimp54546-135457.revcompl.Pan.troglodytes.clone.rp43-41g5.GenBank:AC099533.36     GGTTGCAGTGAACCAAGATTGCTCCACTGCACTCCAGCCTGGGCAGCAGAGGAAGACTCT 34975
human.FAM30B-LOC105372935-GGT2.NCBI.                                               ggttgcagtgaaccaagattgctccactgcactccagcctgggcagcagaggaagactct 31883
human.FAM230E-LOC105377182-GGT3P.NCBI.                                             ggttgcagtgaaccaagattgctccactgcactccagcctgggcagcagaggaagactct 30618
human.FAM230J-LOC105372942-GGTLC5P.NCBI.rev.compl                                  ggttgcagtgaaccaagattgctccactgcactccagcctgggcagcagaggaagactct 31338
human.FAM230A.ncRNA.18487127.18500594-GGTLC3.18516335.18518165.rev.compl.REF       ggttgcagtgaaccaagattgctccactgcactccagcctgggcagcagaggaagactct 20097
                                                                                   ************************************************************

chimp.LOC112206744-LOC107973052-GGT2.revcompl                                      ctctgtctcaacaacaacaaaaagtactagctcatgttagactttgataagtgaaggatg 33304
chimp54546-135457.revcompl.Pan.troglodytes.clone.rp43-41g5.GenBank:AC099533.36     CTCTATCTCAACAACAACAAAAAGTACTAGCTCATGTTAGACTTTGATAAGTGAAGGATG 35035
human.FAM30B-LOC105372935-GGT2.NCBI.                                               ctctatctcaaccacaacaaaaagtactagctcatgttagactttgataagggaaggatg 31943
human.FAM230E-LOC105377182-GGT3P.NCBI.                                             ctctatctcaaccacaacaaaaagtactagctcatgttagactttgataagtgaaggatg 30678
human.FAM230J-LOC105372942-GGTLC5P.NCBI.rev.compl                                  ctctatctcaaccacaacaaaaagtactagctcatgttagactttgataagggaaggatg 31398
human.FAM230A.ncRNA.18487127.18500594-GGTLC3.18516335.18518165.rev.compl.REF       ctctatctcaaccacaacaaaaagtactagctcatgttagactttgataagggaaggatg 20157
                                                                                   **** ******* ************************************** ********

chimp.LOC112206744-LOC107973052-GGT2.revcompl                                      catgttgtaagctctaaaataatccactgtcatcttttaaaataactctaagactgcaca 33364
chimp54546-135457.revcompl.Pan.troglodytes.clone.rp43-41g5.GenBank:AC099533.36     CATGTTGTAAGCTCTAAAATAATCCA--GTCATCTTTTAAAATAACTCTAAGACTGCACA 35093
human.FAM30B-LOC105372935-GGT2.NCBI.                                               catgttgtaagctctaaaataatcca--gtcatcttttaaaataactctaagactgcaca 32001
human.FAM230E-LOC105377182-GGT3P.NCBI.                                             catgttgtaagctctaaaataatcca--gtcatcttttaaaataactctaagactgcaca 30736
human.FAM230J-LOC105372942-GGTLC5P.NCBI.rev.compl                                  catgttgtaagctctaaaataatcca--gtcatcttttaaaagaactctaagactgcaca 31456
human.FAM230A.ncRNA.18487127.18500594-GGTLC3.18516335.18518165.rev.compl.REF       catgttgtaagctctaaaataatcca--gtcatcttttaaaataactctaagactgcaca 20215
                                                                                   **************************  ************** *****************

chimp.LOC112206744-LOC107973052-GGT2.revcompl                                      gttatgaaactaatag---aggaggaaattaaataataaaaataataaatccaaaacaag 33421
chimp54546-135457.revcompl.Pan.troglodytes.clone.rp43-41g5.GenBank:AC099533.36     GTTATGAAACTAATAGAGAAGGAGGAAATTAAATCATAAAAATAATAAATCCAAAACAAG 35153
human.FAM30B-LOC105372935-GGT2.NCBI.                                               gttatgaaactaatagagaaggaggaaattaaataataaaaataataaatccaaaacaag 32061
human.FAM230E-LOC105377182-GGT3P.NCBI.                                             gttatgaaactaatagagaaggaggaaattaaataataaaaataataaatccaaaacaag 30796
human.FAM230J-LOC105372942-GGTLC5P.NCBI.rev.compl                                  gttatgaaactaatagagaaggaggaaattaaataataaaaataataaatccaaaacaag 31516
human.FAM230A.ncRNA.18487127.18500594-GGTLC3.18516335.18518165.rev.compl.REF       gttatgaaactaatagagaaggaggaaattaaataataaaactaataaatccaaaacaag 20275
                                                                                   ****************   *************** ****** ******************

chimp.LOC112206744-LOC107973052-GGT2.revcompl                                      atgtgagaggagataagaagaaatagaataggcatggaaaacaaattggtggtgggtttc 33481
chimp54546-135457.revcompl.Pan.troglodytes.clone.rp43-41g5.GenBank:AC099533.36     ATGTGAGAGAAGATAAGAAGAAATAGAATAGGCATGGAAAACAAATTGGTGGTGGGTTTC 35213
human.FAM30B-LOC105372935-GGT2.NCBI.                                               atgtgagaggagataagaagaaatagaataggcatggaaaacaaattggtggtgggtttc 32121
human.FAM230E-LOC105377182-GGT3P.NCBI.                                             atgtgagaggagataagaagaaatagaataggcatggaaaacaaattggtggtgggtttc 30856
human.FAM230J-LOC105372942-GGTLC5P.NCBI.rev.compl                                  atgtgagaggagataagaagaaatagaataggcatggaaaacaaattggtggtgggtttc 31576
human.FAM230A.ncRNA.18487127.18500594-GGTLC3.18516335.18518165.rev.compl.REF       atgtgagaggagataagaagaaatagaataggcatggaaaacaaattggtggtgggtttc 20335
                                                                                   ********* **************************************************

chimp.LOC112206744-LOC107973052-GGT2.revcompl                                      aacccaaataaatcattagttacatttaaaaggacaataaaaattaaaataatcgaaaat 33541
chimp54546-135457.revcompl.Pan.troglodytes.clone.rp43-41g5.GenBank:AC099533.36     AACGTAAATAAATCATTAGTTACATTTAAAAGGACAATAAAAATTAAAATAATTGAAAAT 35273
human.FAM30B-LOC105372935-GGT2.NCBI.                                               aacccaaataaatcattagttacatttaaaaggacaataaaaattaaaataattgaaaat 32181
human.FAM230E-LOC105377182-GGT3P.NCBI.                                             aacccaaataaatcattagttacatttaaaaggacaataaaaattaaaataattgaaaat 30916
human.FAM230J-LOC105372942-GGTLC5P.NCBI.rev.compl                                  aacccaaataaatcattagttacatttaaaaggacaataaaaattaaaataattgaaaat 31636
human.FAM230A.ncRNA.18487127.18500594-GGTLC3.18516335.18518165.rev.compl.REF       aacccaaataaatcattagttacatttaaaaggacaataaaaattaaaataattgaaaat 20395
                                                                                   ***  ************************************************ ******

chimp.LOC112206744-LOC107973052-GGT2.revcompl                                      aaagtaaaacccaactaatgccttttatataagaatacagagaggtggaagataatgaaa 33601
chimp54546-135457.revcompl.Pan.troglodytes.clone.rp43-41g5.GenBank:AC099533.36     AAAATAAAACCCAACTAATGCCTTTTATATAAGAATACAGAGAGGTGGAAGATAATGAAA 35333
human.FAM30B-LOC105372935-GGT2.NCBI.                                               aaagtaaaacccaactaatgccttttatataagggtacagagaggtggaagatcatgaaa 32241
human.FAM230E-LOC105377182-GGT3P.NCBI.                                             aaagtaaaacccaactaatgccttttatataaggatacagagaggtggaagatcatgaaa 30976
human.FAM230J-LOC105372942-GGTLC5P.NCBI.rev.compl                                  aaagtaaaacccaactaatgccttttatataagggtacagagaggtggaagatcatgaaa 31696
human.FAM230A.ncRNA.18487127.18500594-GGTLC3.18516335.18518165.rev.compl.REF       aaagtaaaacccaactaatgccttttatataagggtacagagaggtggaagatcatgaaa 20455
                                                                                   *** *****************************  ****************** ******

chimp.LOC112206744-LOC107973052-GGT2.revcompl                                      aatatgtcatgcatgcactaaccaagaaagctgtataacttttttttttttttt----- 33656
chimp54546-135457.revcompl.Pan.troglodytes.clone.rp43-41g5.GenBank:AC099533.36     AATATGTCATGCATGCACTAACCAAGAAAGCTGTATCGCTTTTTTTTTTTTTTTTT---- 35389
human.FAM30B-LOC105372935-GGT2.NCBI.                                               aatatgtcatgcatgtactaaccaagaaagctgtataacttttttttttttttttttttt 32301
human.FAM230E-LOC105377182-GGT3P.NCBI.                                             aatatgtcatgcatgtactaaccaagaaagctgtataacttttttttttttttttttttt 31036
human.FAM230J-LOC105372942-GGTLC5P.NCBI.rev.compl                                  aatatgtcatgcatgtactaaccaagaaagctgtataacttttttttttttttttttttt 31756
human.FAM230A.ncRNA.18487127.18500594-GGTLC3.18516335.18518165.rev.compl.REF       aatatgtcatgcatgtactaaccaagaaagctgtataac--ttttttttttttttttttt 20513
                                                                                   *************** ********************  *  **************

chimp.LOC112206744-LOC107973052-GGT2.revcompl                                      --------ggagatagagcctcactctgtctcccaggctggagtgcagtgatgtgatctt 33708
```

```
chimp54546-135457.revcompl.Pan.troglodytes.clone.rp43-41g5.GenBank:AC099533.36  --------GGAGATAGAGCCTCACTCTGTCTCCCAGGCTGGAGTGCAGTGATGTGATCTT 35441
human.FAM30B-LOC105372935-GGT2.NCBI.                                            tttttt--ggagatagagcctcactctgtctcccaggctggagtgcagtgatgtgatctt 32359
human.FAM230E-LOC105377182-GGT3P.NCBI.                                          tttttttggagatagagcctcactctgtctcccaggctggagtgcagtgatgtgatctt 31096
human.FAM230J-LOC105372942-GGTLC5P.NCBI.rev.compl                               ttttttttggagatagagcctcactctgtctcccaggctggagtgcagtgatgtgatctt 31816
human.FAM230A.ncRNA.18487127.18500594-GGTLC3.18516335.18518165.rev.compl.REF    ttttttttggagatagagcctcactctgtctcccaggctggagtgcagtgatgtgatctt 20573
                                                                                        ****************************************************

chimp.LOC112206744-LOC107973052-GGT2.revcompl                                   ggcttacagcaatctctcccttctaggctcaagcgattctcccacctcagcatcccaagt 33768
chimp54546-135457.revcompl.Pan.troglodytes.clone.rp43-41g5.GenBank:AC099533.36  GGGTTACAGCAATCTCTCCCTTCTAGGCTCAAGCGATTCTCCCACCTCAGCATCCCAAGT 35501
human.FAM30B-LOC105372935-GGT2.NCBI.                                            ggcttacagcaatctctcccttctaggctcaagcgattctcccacctcagcatcccaagt 32419
human.FAM230E-LOC105377182-GGT3P.NCBI.                                          ggcttacagcaatctctcccttctaggctcaagcgattctcccacctcagcatcccaagt 31156
human.FAM230J-LOC105372942-GGTLC5P.NCBI.rev.compl                               ggcttacagcaatctctcccttctaggctcaagcgattctcccacctcagcatcccaagt 31876
human.FAM230A.ncRNA.18487127.18500594-GGTLC3.18516335.18518165.rev.compl.REF    ggcttacagcaatctctcccttctaggctcaagcgattctcccacctcagcatcccaagt 20633
                                                                                ** *********************************************************

chimp.LOC112206744-LOC107973052-GGT2.revcompl                                   agctgggactacaagtgtgccaacttagaattatattagccacacccagctaatttttgt 33828
chimp54546-135457.revcompl.Pan.troglodytes.clone.rp43-41g5.GenBank:AC099533.36  AGCTGGGACTACAAGTGTGCCAACTTAGAATTATATTAGCCACACCCAGCTAATTTTTGT 35561
human.FAM30B-LOC105372935-GGT2.NCBI.                                            agctgggactacaagtgtgccaacttagaattatattagccacacccagctaatttttgt 32479
human.FAM230E-LOC105377182-GGT3P.NCBI.                                          agctgggactacaagtgtgccaacttagaattatattagccacacccagctaatttttgt 31216
human.FAM230J-LOC105372942-GGTLC5P.NCBI.rev.compl                               agctgggactacaagtgtgccaacttagaattatactagccacacccagctaatttttgt 31936
human.FAM230A.ncRNA.18487127.18500594-GGTLC3.18516335.18518165.rev.compl.REF    agctgggactacaagtgtgccaacttagaattatattagccacacccagctaatttttgt 20693
                                                                                *********************************** ************************

chimp.LOC112206744-LOC107973052-GGT2.revcompl                                   atttttgtagaggcagggtctcgccatgttgcccaggctggtcttgaactcctgggctt 33888
chimp54546-135457.revcompl.Pan.troglodytes.clone.rp43-41g5.GenBank:AC099533.36  ATTTTTGTAGAGGCAGGGTCTCGCCATGTTGCCCAGGTTGGTCTTGAACTCCTGGGCTT 35621
human.FAM30B-LOC105372935-GGT2.NCBI.                                            atttttgtagaggcagggtctcgccatgttgcccaggttggtcttgaactcctgggctt 32539
human.FAM230E-LOC105377182-GGT3P.NCBI.                                          atttttgtagaggcagggtctcgccatgttgcccaggttggtcttgaactcctgggctt 31276
human.FAM230J-LOC105372942-GGTLC5P.NCBI.rev.compl                               atttttgtagaggcagggtctcgccatgttgcccaggttggtcttgaactcctgggctt 31996
human.FAM230A.ncRNA.18487127.18500594-GGTLC3.18516335.18518165.rev.compl.REF    atttttgtagaggcagggtctcgccatgttgcccaggttggtcttgaactcctgggctt 20753
                                                                                ************************************* **********************

chimp.LOC112206744-LOC107973052-GGT2.revcompl                                   cagtgatccacccacctcgacatccagcaaagtgccaagattacagccatgagccaccat 33948
chimp54546-135457.revcompl.Pan.troglodytes.clone.rp43-41g5.GenBank:AC099533.36  CAGTGATCCACCCACCTCGACCTCCAACAAAGTGCCGAGATTACAGCCATGAGCCACCAT 35681
human.FAM30B-LOC105372935-GGT2.NCBI.                                            cagtgatccacccacctcgacctccagcaaagtgccaagattacagccatgagccaccat 32599
human.FAM230E-LOC105377182-GGT3P.NCBI.                                          cagtgatccacccacctcgacctccagcaaagtgccaagattacagccatgagccaccat 31336
human.FAM230J-LOC105372942-GGTLC5P.NCBI.rev.compl                               cagtgatccacccacctcgacctccagcaaagtgccaagattacagccatgagccaccat 32056
human.FAM230A.ncRNA.18487127.18500594-GGTLC3.18516335.18518165.rev.compl.REF    cagtgatccacccacctcgacctccagcaaagtgccaagattacagccatgagccaccat 20813
                                                                                ********************* **** ********* ***********************

chimp.LOC112206744-LOC107973052-GGT2.revcompl                                   gcccagcataaccatttttaatgaagtagactttaagaagaaaagtattattagaggtaa 34008
chimp54546-135457.revcompl.Pan.troglodytes.clone.rp43-41g5.GenBank:AC099533.36  GCCCAGCATAACTATTTTTAATGAAGTAGACTTTAAGAAGAAAAGTATTATTAGAAGTAA 35741
human.FAM30B-LOC105372935-GGT2.NCBI.                                            gcccagcataactatttttaatgaagtagactttaagaagaaaagtattattagaggtaa 32659
human.FAM230E-LOC105377182-GGT3P.NCBI.                                          gcccagcataactatttttaatgaagtagactttaagaagaaaagtattattagaggtaa 31396
human.FAM230J-LOC105372942-GGTLC5P.NCBI.rev.compl                               gcccagcataactatttttaatgaagtagactttaagaagaaaagtattattagaggtaa 32116
human.FAM230A.ncRNA.18487127.18500594-GGTLC3.18516335.18518165.rev.compl.REF    gcccagcataactatttttaatgaagtagactttaagaagaaaagtattattagaggtaa 20873
                                                                                ************ ****************************************** ****

chimp.LOC112206744-LOC107973052-GGT2.revcompl                                   gagacacatcacggaaaagaagaatttactaggagccaggcgcagtggctcatgcctgta 34068
chimp54546-135457.revcompl.Pan.troglodytes.clone.rp43-41g5.GenBank:AC099533.36  GAGACACATCACGGAAAAGAAGAATTTACTAGGAGCCAGGCGCAGTGGCTCGTGCCTGTA 35801
human.FAM30B-LOC105372935-GGT2.NCBI.                                            gagacacatcacagaaaagaagaatttactaggagccaggcgcaatggctcgtgcctgta 32719
human.FAM230E-LOC105377182-GGT3P.NCBI.                                          gagacacatcacagaaaagaagaatttactaggagccaggcgcaatggctcgtgcctgta 31456
human.FAM230J-LOC105372942-GGTLC5P.NCBI.rev.compl                               gggacacatcacagaaaagaagaatttactaggagccaggcgcaatggctcgtgcctgta 32176
human.FAM230A.ncRNA.18487127.18500594-GGTLC3.18516335.18518165.rev.compl.REF    gggacacatcacagaaaagaagaatttactaggagccaggcgcaatggctcgtgcctgta 20933
                                                                                * ********** ******************************* ****** ********

chimp.LOC112206744-LOC107973052-GGT2.revcompl                                   attccagcactttgtgaggccaaggcggcggatcacctgaggttgggagttcaagaccag 34128
chimp54546-135457.revcompl.Pan.troglodytes.clone.rp43-41g5.GenBank:AC099533.36  ATTCCAGCACTTTGTGAGGCCAAGGCGGCGGATCACCTGAGGTTGGGAGTTCAAGACCAG 35861
human.FAM30B-LOC105372935-GGT2.NCBI.                                            attccagcactttgtgaggccaaggcggcggatcacctgaggttgggagttcaagaccag 32779
human.FAM230E-LOC105377182-GGT3P.NCBI.                                          attccagcactttgtgaggccaaggcggcggatcacctgaggttgggagttcaagaccag 31516
human.FAM230J-LOC105372942-GGTLC5P.NCBI.rev.compl                               attccagcactttgtgaggccaaggcggcggatcacctgaggttgggagttcaagaccag 32236
human.FAM230A.ncRNA.18487127.18500594-GGTLC3.18516335.18518165.rev.compl.REF    attccagcactttgtgaggccaaggcggcggatcacctgaggttgggagttcaagaccag 20993
                                                                                ************************************************************

chimp.LOC112206744-LOC107973052-GGT2.revcompl                                   cctgaccaacatggagaagccctgtctctactaaaaatacaaaaattagccaagcatggt 34188
chimp54546-135457.revcompl.Pan.troglodytes.clone.rp43-41g5.GenBank:AC099533.36  CCTGACCAACATGGAGAAGCCCTGTCTCTACTAAAAATACAAAAATTAGCCAAGCATGGT 35921
human.FAM30B-LOC105372935-GGT2.NCBI.                                            cctgaccaacatggagaagccctgtctctactaaaaatacaaaaattagccaagcatggt 32839
human.FAM230E-LOC105377182-GGT3P.NCBI.                                          cctgaccaacatggagaagccctgtctctactaaaaatacaaaaattagccaagcatggt 31576
human.FAM230J-LOC105372942-GGTLC5P.NCBI.rev.compl                               cctgaccaacatggagaagccctgtctctactaaaaatacaaaaattagccaagcatggt 32296
human.FAM230A.ncRNA.18487127.18500594-GGTLC3.18516335.18518165.rev.compl.REF    cctgaccaacatggagaagccctgtctctactaaaaatacaaaaattagccaagcatggt 21053
                                                                                ************************************************************

chimp.LOC112206744-LOC107973052-GGT2.revcompl                                   ggcacatgcctgtaatcccagctactcaggaggctgagggaggagaattgcttggaccca 34248
chimp54546-135457.revcompl.Pan.troglodytes.clone.rp43-41g5.GenBank:AC099533.36  GGCACATGCCTGTAATCCCAGCTACTCAGGAGGCTGAGGGAGGAGAATTGCTTGGACCCA 35981
human.FAM30B-LOC105372935-GGT2.NCBI.                                            ggcacatgcctgtaatcccagctactcaggaggctgagggaggagaattgcttggaccca 32899
human.FAM230E-LOC105377182-GGT3P.NCBI.                                          ggcacatgcctgtaatcccagctactcaggaggctgagggaggagaattgcttggaccca 31636
human.FAM230J-LOC105372942-GGTLC5P.NCBI.rev.compl                               ggcacatgcctgtaatcccagctactcaggaggctgagggaggagaattgcttggaccca 32356
human.FAM230A.ncRNA.18487127.18500594-GGTLC3.18516335.18518165.rev.compl.REF    ggcacatgcctgtaatcccagctactcaggaggctgagggaggagaattgcttggaccca 21113
                                                                                ************************************************************

chimp.LOC112206744-LOC107973052-GGT2.revcompl                                   ggaagtggaggttgcggtgagctgagattgtgccattgcattccagcctgggcaacaaga 34308
chimp54546-135457.revcompl.Pan.troglodytes.clone.rp43-41g5.GenBank:AC099533.36  GGAAGTGGAGGTTGCGGTGAGCTGAGATTGTGCCATTGCATTCCAGCCTGGGCAACAAGA 36041
human.FAM30B-LOC105372935-GGT2.NCBI.                                            ggaagtggaggttgcggtgagctgagattgtgccattgcattccagcctgggcaacaaga 32959
human.FAM230E-LOC105377182-GGT3P.NCBI.                                          ggaagtggaggttgcggtgagctgagactgtgccattgcattccagcctgggcaacaaga 31696
human.FAM230J-LOC105372942-GGTLC5P.NCBI.rev.compl                               ggaagtggaggttgcggtgagctgagattgtgccattgcattccagcctgggcaacaaga 32416
human.FAM230A.ncRNA.18487127.18500594-GGTLC3.18516335.18518165.rev.compl.REF    ggaagtggaggttgcggtgagctgagattgtgccattgcattccagcctgggcaacaaga 21173
                                                                                ************************** *********************************

chimp.LOC112206744-LOC107973052-GGT2.revcompl                                   gcaaaactctgtctc-------------aaaaaaaaagaagttactagctagttacggta 34355
chimp54546-135457.revcompl.Pan.troglodytes.clone.rp43-41g5.GenBank:AC099533.36  GCAAAACTCTGTCTC---------AAAAAAAAAAAAAGAAGTTACTAGCTAGTTTTGGTA 36092
human.FAM30B-LOC105372935-GGT2.NCBI.                                            gcaaaactctgtctc---aaaaaaaaaaaaaaaaaaagaagttactagctagtttcggta 33016
human.FAM230E-LOC105377182-GGT3P.NCBI.                                          gcaaaactctgtctcaaaaaaaaaaaaaaaaaaaaaagaagttactagctagtttcggta 31756
human.FAM230J-LOC105372942-GGTLC5P.NCBI.rev.compl                               gcaaaactctgtctc-----aaaaaaaaaaaaaaaaagaagttactagctagtttcggta 32471
human.FAM230A.ncRNA.18487127.18500594-GGTLC3.18516335.18518165.rev.compl.REF    gcaaaactctgtctc----------aaaaaaaaaaaagaagttactagctagtttcggta 21223
                                                                                ***************             **************************  ****

chimp.LOC112206744-LOC107973052-GGT2.revcompl                                   attcttaacatccaggaaactgg-ttgtgaaagtttttcagagaaactaaaccaatagat 34414
chimp54546-135457.revcompl.Pan.troglodytes.clone.rp43-41g5.GenBank:AC099533.36  ATTCTTAACATCCAGGAAACTGG-ATGTGAAAGTTTTTCAGAGAAACTAAACCAATAGAT 36151
human.FAM30B-LOC105372935-GGT2.NCBI.                                            attcttaacaaccaggaaactgg-atgtgaaagtttttcagagaaactaaaccaatagat 33075
human.FAM230E-LOC105377182-GGT3P.NCBI.                                          attcttaacaaccaggaaactggaatgtgaaagtttttcagagaaactaaaccaatagat 31816
human.FAM230J-LOC105372942-GGTLC5P.NCBI.rev.compl                               attcttaacatccaggaaactggaatgtgaaagtttttcagagaaactaaaccaatagat 32531
human.FAM230A.ncRNA.18487127.18500594-GGTLC3.18516335.18518165.rev.compl.REF    attcttaacatccaggaaactgg-atgtgaaagtttttcagagaaactaaaccaatagat 21282
                                                                                ********** ************  ***********************************

chimp.LOC112206744-LOC107973052-GGT2.revcompl                                   tatacatagagagagatttattgaggaattggctcacatgattgtggggactagcaag-t 34473
chimp54546-135457.revcompl.Pan.troglodytes.clone.rp43-41g5.GenBank:AC099533.36  TATACATAGAGAGAGATTTATTGAGGAATTGGCTCACATGATTGTGGGGACTAGCAAG-T 36210
human.FAM30B-LOC105372935-GGT2.NCBI.                                            tatacatagagagagatttatttaggaattggctcacatgattgtggggactagcaagtt 33135
human.FAM230E-LOC105377182-GGT3P.NCBI.                                          tatacatagagagagatttatttaggaattggctcacatgattgtggggactagcaagtt 31876
human.FAM230J-LOC105372942-GGTLC5P.NCBI.rev.compl                               tatacatagagagagacttatttaggaattggctcacatgattgtggggactagcaagtt 32591
human.FAM230A.ncRNA.18487127.18500594-GGTLC3.18516335.18518165.rev.compl.REF    tatacatagagagagacttatttaggaattggctcacatgattgtggggactagcaagtt 21342
                                                                                **************** ***** *********************************** *

chimp.LOC112206744-LOC107973052-GGT2.revcompl                                   ttaaaatctgtagggcaagccagcaggctataaattcaggtaagagttgatctcgaagtc 34533
chimp54546-135457.revcompl.Pan.troglodytes.clone.rp43-41g5.GenBank:AC099533.36  TTAAAATCTGTAGGGCAAGCCAGCAGGCTATAGATTCAGGTAAGAGTTGATCTCGAAGTC 36270
human.FAM30B-LOC105372935-GGT2.NCBI.                                            ttaaaatctgtagggcaagccagcaggctataaattcaggtaagagttgatctcgaagtc 33195
human.FAM230E-LOC105377182-GGT3P.NCBI.                                          ttaaaatctgtagggcaagccagcaggctataaattcaggtaagagttgatctcgaagtc 31936
human.FAM230J-LOC105372942-GGTLC5P.NCBI.rev.compl                               ttaaaatctgtagggcaagccagcaggctataaattcaggtaagagttgatctcgaagtc 32651
human.FAM230A.ncRNA.18487127.18500594-GGTLC3.18516335.18518165.rev.compl.REF    ttaaaatctgtagggcaagccagcaggctataaattcaggtaagagttgatctcgaagtc 21402
                                                                                ********************************* **************************

chimp.LOC112206744-LOC107973052-GGT2.revcompl                                   tggaacctaaaatctgtagagcagtcagcaggccagaaactcaggcagggtttgtgtgtt 34593
chimp54546-135457.revcompl.Pan.troglodytes.clone.rp43-41g5.GenBank:AC099533.36  TTGAACCTAAAATCTGTAGAGCAGTCAGCAGGCCAGAAACTCAGGCAGGGTTTGTGTGTT 36330
human.FAM30B-LOC105372935-GGT2.NCBI.                                            tggaacctaaaatctgtagagcagtcagcaggccagaaactcaggcagggtttgtgtgtt 33255
human.FAM230E-LOC105377182-GGT3P.NCBI.                                          tggaacctaaaatctgtagagcagtcagcaggccagaaactcaggcagggtttgtgtgtt 31996
human.FAM230J-LOC105372942-GGTLC5P.NCBI.rev.compl                               tggaacctaaaatctgtagagcagtcagcaggccagaaactcaggtagggtttgtgtgtt 32711
human.FAM230A.ncRNA.18487127.18500594-GGTLC3.18516335.18518165.rev.compl.REF    tggaacctaaaatctgtagagcagtcagcaggccagaaactcaggcagggtttgtgtgtt 21462
                                                                                * ******************************************* **************

chimp.LOC112206744-LOC107973052-GGT2.revcompl                                   acagtcttgaagcagaattcctgcttctttgggaaacct--gtttttgttcttaaggcct 34651
chimp54546-135457.revcompl.Pan.troglodytes.clone.rp43-41g5.GenBank:AC099533.36  ATAGTCTTGAAGCAGAATTCCTGCTTCTCTGGGAAACCTCAGTTTTTGTTCTTAAGGCCT 36390
human.FAM30B-LOC105372935-GGT2.NCBI.                                            acagtcttgaagcagaattcctgcttctctgggaaacctcagtttttgttcttaaggcct 33315
human.FAM230E-LOC105377182-GGT3P.NCBI.                                          acagtcttgaagcagaattcctgcttctctgggaaacctcagtttttgttcttaaggcct 32056
human.FAM230J-LOC105372942-GGTLC5P.NCBI.rev.compl                               acagtcttgaagcagaattcctgcttctctgggaaacctcagtttttgttcttaaggcct 32771
human.FAM230A.ncRNA.18487127.18500594-GGTLC3.18516335.18518165.rev.compl.REF    acagtcttgaagcagaattcctgcttctctgggaaacctcagtttttgttcttaaggcct 21522
                                                                                * ************************** **********  *******************

chimp.LOC112206744-LOC107973052-GGT2.revcompl                                   tcaagtgattggaggtggcccacccatattatggtgggtaatctgttttacttaaagtca 34711
chimp54546-135457.revcompl.Pan.troglodytes.clone.rp43-41g5.GenBank:AC099533.36  TCAACTGATTGGAGGTGGCCCACCCATATTATGGTGGGTAATCTGTTTTACTTAAAGTCA 36450
human.FAM30B-LOC105372935-GGT2.NCBI.                                            tcaactgattggaggtggcccacccatattatggtgggtaatctgttttacttaaagtca 33375
human.FAM230E-LOC105377182-GGT3P.NCBI.                                          tcaactgattggaggtggcccacccatattatggtgggtaatctgttttacttaaagtca 32116
human.FAM230J-LOC105372942-GGTLC5P.NCBI.rev.compl                               tcaactgattggaggtggcccacccatattatggtgggtaatctgttttacttaaagtca 32831
human.FAM230A.ncRNA.18487127.18500594-GGTLC3.18516335.18518165.rev.compl.REF    tcaactgattggaggtggcccacccatattatggtgggtaatctgttttacttaaagtca 21582
                                                                                **** *******************************************************

chimp.LOC112206744-LOC107973052-GGT2.revcompl                                   attgactgtcagtgttaaccacatctatgaaataacctcccagcaagatattgacaagta 34771
chimp54546-135457.revcompl.Pan.troglodytes.clone.rp43-41g5.GenBank:AC099533.36  ACTGACTGTCAGTGTTAACCACATCTATGAAATAACCTCCCAGCAAGATATTGACAAGTA 36510
```

```
human.FAM30B-LOC105372935-GGT2.NCBI.                                        attgactgtcagtgttaatcacatctatgaaataacctcccagcaagatattgacaagta 33435
human.FAM230E-LOC105377182-GGT3P.NCBI.                                      attgactgtcagtgttaatcacatctatgaaataacctcccagcaagatattgacaagta 32176
human.FAM230J-LOC105372942-GGTLC5P.NCBI.rev.compl                           attgactgtcagtgttaatcacatctatgaaataacctcccagcaagatattgacaagta 32891
human.FAM230A.ncRNA.18487127.18500594-GGTLC3.18516335.18518165.rev.compl.REF attgactgtcagtgttaatcacatctatgaaataacctcccagcaagatattgacaagta 21642
                                                                            * ***************** *****************************************

chimp.LOC112206744-LOC107973052-GGT2.revcompl                               tttgaccaaacaacggggcaccatagcttagccaagttgacacataaattaaccatcagg 34831
chimp54546-135457.revcompl.Pan.troglodytes.clone.rp43-41g5.GenBank:AC099533.36 TTTGACCAAACAACGGGGCACCATAGCTTAGCCAAGTTGACACATAAATTAACCATCAGG 36570
human.FAM30B-LOC105372935-GGT2.NCBI.                                        tttgaccaaacaacaggacaccatagcttagccaagttgacacataaattaaccatcagg 33495
human.FAM230E-LOC105377182-GGT3P.NCBI.                                      ttcgaccaaacaacaggacaccatagcttagccaagttgacacataaattaaccatcagg 32236
human.FAM230J-LOC105372942-GGTLC5P.NCBI.rev.compl                           tttgaccaaacaacaggacaccatagcttagccaagttgacacataaattaaccatcagg 32951
human.FAM230A.ncRNA.18487127.18500594-GGTLC3.18516335.18518165.rev.compl.REF tttgaccaaacaacaggacaccatagcttagccaagttgacacataaattaaccatcagg 21702
                                                                            ** *********** ** ******************************************

chimp.LOC112206744-LOC107973052-GGT2.revcompl                               agcgagtagaatatccaaaaaacaacatactaggggtattatatcttatatagc-----t 34886
chimp54546-135457.revcompl.Pan.troglodytes.clone.rp43-41g5.GenBank:AC099533.36 AGCGAGTAGAATATCCAAAAAACAACATACTAGGGGTATTATATCTTATATAGC-----T 36625
human.FAM30B-LOC105372935-GGT2.NCBI.                                        agcaagtagaatatccaaaaaacaacatactaggggtattatatcttatatagctattat 33555
human.FAM230E-LOC105377182-GGT3P.NCBI.                                      agcaagtagaatatccaaaaaacaacatactaggggtattatatcttatatagctattat 32296
human.FAM230J-LOC105372942-GGTLC5P.NCBI.rev.compl                           agcaagtagaatatccaaaaaacaacatactaggggtattatatcttatatagctattat 33011
human.FAM230A.ncRNA.18487127.18500594-GGTLC3.18516335.18518165.rev.compl.REF agcaagtagaatatccaaaaaacaacatactaggggtattatatcttatatagctattat 21762
                                                                            *** **************************************************     *

chimp.LOC112206744-LOC107973052-GGT2.revcompl                               aattatataaaacatataattatagaatgaagatattaagataaccattagaacaaaaat 34946
chimp54546-135457.revcompl.Pan.troglodytes.clone.rp43-41g5.GenBank:AC099533.36 AATTATATAAAACATATAATTATAGAATGAAGATATTAAGATAACCATTAGAACAAAAAT 36685
human.FAM30B-LOC105372935-GGT2.NCBI.                                        aattatataaaacatataattatagaatgacgatattaagataaccattagaacaaaaat 33615
human.FAM230E-LOC105377182-GGT3P.NCBI.                                      aattatataaaacatataattatagaatgacgatattaagataaccattagaacaaaaat 32356
human.FAM230J-LOC105372942-GGTLC5P.NCBI.rev.compl                           aattatataaaacatataattatagaatgacgatattaagataagcattagaacaaaaat 33071
human.FAM230A.ncRNA.18487127.18500594-GGTLC3.18516335.18518165.rev.compl.REF aattatataaaacatataattatagaatgacgatattaagataaccattagaacaaaaat 21822
                                                                            ****************************** ************* ***************

chimp.LOC112206744-LOC107973052-GGT2.revcompl                               ataaacttttctt-------ttcttttttttttttgagaccaagtcttgctctgtcaccca 34999
chimp54546-135457.revcompl.Pan.troglodytes.clone.rp43-41g5.GenBank:AC099533.36 ATAAACTTTTCTT-------TTTTTTTTTTTTTTGAGACCAAGTCTTGCTCTGTCACCCA 36738
human.FAM30B-LOC105372935-GGT2.NCBI.                                        ataaacttttcttt-ctttcttttttttttttttgagaccaagtcttgctctgtcaccca 33674
human.FAM230E-LOC105377182-GGT3P.NCBI.                                      ataaacttttctttcc----ttttttttttttttgagaccaagtcttgctctgtcaccca 32412
human.FAM230J-LOC105372942-GGTLC5P.NCBI.rev.compl                           ataaacttttctttc-ttttttttttttttttttgagaccaagtcttgctctgtcaccca 33130
human.FAM230A.ncRNA.18487127.18500594-GGTLC3.18516335.18518165.rev.compl.REF ataaacttttctttcttttttttttttttttttttgagaccaagtcttgctctgtcaccca 21882
                                                                            *************       ** *************************************

chimp.LOC112206744-LOC107973052-GGT2.revcompl                               ggctggagtgcagtggtgcaatcttggcttactgcaacctttgcctcctgagttcaagtg 35059
chimp54546-135457.revcompl.Pan.troglodytes.clone.rp43-41g5.GenBank:AC099533.36 GGCTGGAGTGCAGTGGTGCAATCTTGGCTTACTGCAACCTTTGCCTCCTGGGTTCAAGTG 36798
human.FAM30B-LOC105372935-GGT2.NCBI.                                        ggctggagtgcagtggtgcaatcttggcttactgaaacctttgcctcctgggttcaagtg 33734
human.FAM230E-LOC105377182-GGT3P.NCBI.                                      ggctggagtgcagtggtgcaatcttggcttactgcaacctttgcctcctgcgttcaagtg 32472
human.FAM230J-LOC105372942-GGTLC5P.NCBI.rev.compl                           ggctggagtgcagtggtgcaatcttggcttactgcaacctttgcctcctgcgttcaagtg 33190
human.FAM230A.ncRNA.18487127.18500594-GGTLC3.18516335.18518165.rev.compl.REF ggctggagtgcagtggtgcaatcttggcttactgcaacctttgcctcctgcgttcaagtg 21942
                                                                            ********************************** *************** *********

chimp.LOC112206744-LOC107973052-GGT2.revcompl                               atcctcctgtctcagcctcccaagtagctgggattacaggcacctgctaccatgcccagc 35119
chimp54546-135457.revcompl.Pan.troglodytes.clone.rp43-41g5.GenBank:AC099533.36 ATTCTCCTGTCTCAGCCTCCCAAGTATCTGGGATTACAGGCATCCGCTACCATGCCCAGC 36858
human.FAM30B-LOC105372935-GGT2.NCBI.                                        attctcctgtctcagcctcccaagtatctgggattacaggcacctgctaccatgcccagc 33794
human.FAM230E-LOC105377182-GGT3P.NCBI.                                      attctcctgtctcagcctcccaagtatctgggattacaggcacctgctaccatgcccagc 32532
human.FAM230J-LOC105372942-GGTLC5P.NCBI.rev.compl                           attctcctgtctcagcctcccaagtatctgggattacaggcacctgctaccatgcccagc 33250
human.FAM230A.ncRNA.18487127.18500594-GGTLC3.18516335.18518165.rev.compl.REF attctcctgtctcagcctcccaagtatctgggattacaggcacctgctaccatgcccagc 22002
                                                                            ** ********************** **************** ** **************

chimp.LOC112206744-LOC107973052-GGT2.revcompl                               taatttttgtatttttagtagagacatggtttcaccatgttgcccaggctggtctccaac 35179
chimp54546-135457.revcompl.Pan.troglodytes.clone.rp43-41g5.GenBank:AC099533.36 TAATTTTTGTATTTTTAGTAGAGACATGGTTTCACCATGTTGCCCAGGCTGGTCTCCAAC 36918
human.FAM30B-LOC105372935-GGT2.NCBI.                                        taatttttgtatttttagtagagacatggtttcaccatgttgcccaggctggtctccaac 33854
human.FAM230E-LOC105377182-GGT3P.NCBI.                                      taatttttgtatttttagtagagacatggtttcaccatgttgcccaggctggtctccaac 32592
human.FAM230J-LOC105372942-GGTLC5P.NCBI.rev.compl                           taatttttgtatttttagtagagacatggtttcaccatgttgcccaggctggtctccaac 33310
human.FAM230A.ncRNA.18487127.18500594-GGTLC3.18516335.18518165.rev.compl.REF taatttttgtatttttagtagagacatggtttcaccatgttgcccaggctggtctccaac 22062
                                                                            ************************************************************

chimp.LOC112206744-LOC107973052-GGT2.revcompl                               tcctgacctcaagtgagccagccccctcggcctcccaaagtgctgggattacaggtgtga 35239
chimp54546-135457.revcompl.Pan.troglodytes.clone.rp43-41g5.GenBank:AC099533.36 TCCTGACCTCAAGTGAGCCACCCCCCTCGGCCTCCCAAAGTGCTGGAATTACAGGTGTGA 36978
human.FAM30B-LOC105372935-GGT2.NCBI.                                        tcctgacctcaagtgagccacccccctcggcctcccaaagtgctgggattacaggtgtga 33914
human.FAM230E-LOC105377182-GGT3P.NCBI.                                      tcctgacctcaagtgagccacccccctcggcctcccaaagtgctgggattacaggtgtga 32652
human.FAM230J-LOC105372942-GGTLC5P.NCBI.rev.compl                           tcctgacctcaagtgagccacccccctcggcctcccaaagtgctgggattacaggtgtga 33370
human.FAM230A.ncRNA.18487127.18500594-GGTLC3.18516335.18518165.rev.compl.REF tcctgacctcaagtgagccaccccccttggcctcccaaagtgctgggattacaggtgtga 22122
                                                                            ******************** ****** ****************** *************

chimp.LOC112206744-LOC107973052-GGT2.revcompl                               gccaccacacccagccaaaaatcacctttttcacaaggatcaaaacagtcattacgctgg 35299
chimp54546-135457.revcompl.Pan.troglodytes.clone.rp43-41g5.GenBank:AC099533.36 GCCACCACACCCAGCCAAAAATCACCTTTTTCACAAGGATCAAAACAGTCATTACGCTGG 37038
human.FAM30B-LOC105372935-GGT2.NCBI.                                        gccaccacacccagccaaaaatcaccttttttacaaggatcaaaacagtcattatgctgg 33974
human.FAM230E-LOC105377182-GGT3P.NCBI.                                      gccaccacacccagccaaaaatcaccttttttacaaggatcaaaacagtcattatgctgg 32712
human.FAM230J-LOC105372942-GGTLC5P.NCBI.rev.compl                           gccaccacacccagccaaaaatcaccttttttacaaggatcaaaacagtcattatgctgg 33430
human.FAM230A.ncRNA.18487127.18500594-GGTLC3.18516335.18518165.rev.compl.REF gccaccacacccagccaaaaatcaccttttttacaaggatcaaaacagtcattatgctgg 22182
                                                                            ******************************* ********************** *****

chimp.LOC112206744-LOC107973052-GGT2.revcompl                               agatgacagacctcactgtcaccatgctccttttgtatgtctactaggcatggtgctggg 35359
chimp54546-135457.revcompl.Pan.troglodytes.clone.rp43-41g5.GenBank:AC099533.36 AGATGACAGACCTCACTGTCACCATGCTCCTTTTGTATGTCTACTAGGCACGGTGCTGGG 37098
human.FAM30B-LOC105372935-GGT2.NCBI.                                        agatgacagacctcactgtcaccatgctccttttgtatgtctactaggcacggtgctggg 34034
human.FAM230E-LOC105377182-GGT3P.NCBI.                                      agatgacagacctcactgtcaccatgctccttttgtatgtctactaggcacggtgctggg 32772
human.FAM230J-LOC105372942-GGTLC5P.NCBI.rev.compl                           agatgacagacctcactgtcaccatgctccttttgtatgtctactaggcacggtgctggg 33490
human.FAM230A.ncRNA.18487127.18500594-GGTLC3.18516335.18518165.rev.compl.REF agatgacagacctcactgtcaccatgctccttttgtatgtctactaggcacggtgctggg 22242
                                                                            ************************************************** *********

chimp.LOC112206744-LOC107973052-GGT2.revcompl                               tccacactcacagaaagcttaggagctcgcacccaggggctccggctgtagcagaatccc 35419
chimp54546-135457.revcompl.Pan.troglodytes.clone.rp43-41g5.GenBank:AC099533.36 TCCACACTCACAGAAAGCTTAGGAGCTCGCACCCAGGGGCTCCGGCTGTAGCAGAATCCC 37158
human.FAM30B-LOC105372935-GGT2.NCBI.                                        tccacactcacagaaaccttaggaactcgcacccaggggctccggctgtagcagaatccc 34094
human.FAM230E-LOC105377182-GGT3P.NCBI.                                      tccacactcacagaaaccttaggaactcgcacccaggggctccggctgtagcagaatccc 32832
human.FAM230J-LOC105372942-GGTLC5P.NCBI.rev.compl                           tccacactcacagaaaccttaggaactcgcacccaggggctccggctgtagcagaatccc 33550
human.FAM230A.ncRNA.18487127.18500594-GGTLC3.18516335.18518165.rev.compl.REF tccacactcacagaaaccttaggaactcgcacccaggggctccggctgtagcagaatccc 22302
                                                                            **************** ******* ***********************************

chimp.LOC112206744-LOC107973052-GGT2.revcompl                               aagaataaaacctggtgctgaaagagtaggagatgaggccggatgccatgactcactcct 35479
chimp54546-135457.revcompl.Pan.troglodytes.clone.rp43-41g5.GenBank:AC099533.36 AAGAATAAAACCTGGTGCTGAAAGAGTAGGAGATGAGGCCGGATGCCATGACTCACTCCT 37218
human.FAM30B-LOC105372935-GGT2.NCBI.                                        aagaataaaacctggtgctgaaagagtaggagatgaggccaggtgccatgactcactcct 34154
human.FAM230E-LOC105377182-GGT3P.NCBI.                                      aagaataaaacctggtgctgaaagagtaggagatgaggccgggtgccatgactcactcct 32892
human.FAM230J-LOC105372942-GGTLC5P.NCBI.rev.compl                           aagaataaaacctggtgctgaaagagtaggagatgaggccgggtgccatgactcactcct 33610
human.FAM230A.ncRNA.18487127.18500594-GGTLC3.18516335.18518165.rev.compl.REF aagaataaaacctggtgctgaaagagtaggagatgaggccgggtgccatgactcactcct 22362
                                                                            **************************************** * *****************

chimp.LOC112206744-LOC107973052-GGT2.revcompl                               gtaatgccagcactttgggtggccaaggcgggtgaatcaagagatagagaccatcct-gg 35538
chimp54546-135457.revcompl.Pan.troglodytes.clone.rp43-41g5.GenBank:AC099533.36 GTAATGCCAGCACTTTGGGTGGCCAAGGCGGGTGAATCAAGAGATAGAGACCATCCT-GG 37277
human.FAM30B-LOC105372935-GGT2.NCBI.                                        gtaatgccagcactttgggtggccaaggcgggtgaatcaagagatagagaccatcct-gg 34213
human.FAM230E-LOC105377182-GGT3P.NCBI.                                      gtaatgccagcactttgggtggccaaggcgggtgaatcaagagatagagaccatcct-gg 32951
human.FAM230J-LOC105372942-GGTLC5P.NCBI.rev.compl                           gtaatgccagcactttgggtggccaaggcgggtgaatcaagagatagagaccatcctggg 33670
human.FAM230A.ncRNA.18487127.18500594-GGTLC3.18516335.18518165.rev.compl.REF gtaatgccagcactttgggtggccaaggcgggtgaatcaagagatagagaccatcct-gg 22421
                                                                            ********************************************************* **

chimp.LOC112206744-LOC107973052-GGT2.revcompl                               ccaacatggtgaaaccccgtctctactaaaaatacaaaaattagctgggcgtggtggctg 35598
chimp54546-135457.revcompl.Pan.troglodytes.clone.rp43-41g5.GenBank:AC099533.36 CCAACATGGTGAAACCCCGTCTCTACTAAAAATACAAAAATTAGCTGGGCGTGGTGGCTG 37337
human.FAM30B-LOC105372935-GGT2.NCBI.                                        ccaacacggtgaaaccccgtctctactaaaaatacagaaattagcggggcgtggtggctg 34273
human.FAM230E-LOC105377182-GGT3P.NCBI.                                      ccaacacggtgaaaccccgtctctactaaaaatacagaaattagcggggcgtggtggctg 33011
human.FAM230J-LOC105372942-GGTLC5P.NCBI.rev.compl                           ccaacatggtgaaaccctgtctctactaaaaatacagaaattagcggggcgtggtggctg 33730
human.FAM230A.ncRNA.18487127.18500594-GGTLC3.18516335.18518165.rev.compl.REF ccaacacggtgaaaccccgtctctactaaaaatacagaaattagcggggcgtggtggctg 22481
                                                                            ****** ********** ****************** ******** **************

chimp.LOC112206744-LOC107973052-GGT2.revcompl                               gcacctgtagtcccagctactcaggaggctgaggcaggagaatcatttgaaccgaggaag 35658
chimp54546-135457.revcompl.Pan.troglodytes.clone.rp43-41g5.GenBank:AC099533.36 GCACCTGTAGTCCCAGCTACTCAGGAGGCTGAGGCAGGAGAATCATTTGAACCGAGGAAG 37397
human.FAM30B-LOC105372935-GGT2.NCBI.                                        gcacctgtagtcccagctactcaggaggctgaggcaggagaatcatttgaaccgaggaag 34333
human.FAM230E-LOC105377182-GGT3P.NCBI.                                      gcacctgtagtcccagctactcaggaggctgaggcaggagaatcatttgaaccgaggaag 33071
human.FAM230J-LOC105372942-GGTLC5P.NCBI.rev.compl                           gcacctgtagtcccagctactcaggaggctgaggcaggagaatcatttgaaccgaggaag 33790
human.FAM230A.ncRNA.18487127.18500594-GGTLC3.18516335.18518165.rev.compl.REF gcacctgtagtcccagctactcaggaggctgaggcaggagaatcatttgaaccgaggaag 22541
                                                                            ************************************************************

chimp.LOC112206744-LOC107973052-GGT2.revcompl                               cagaggttgcagtaagctgagatcgtgccactgcactccagcctggtgacagagtgagac 35718
chimp54546-135457.revcompl.Pan.troglodytes.clone.rp43-41g5.GenBank:AC099533.36 CAGAGGTTGCAGTAAGCTGAGATCACGCCACTGCACTCCAGCCTGGTGACAGAGTGAGAC 37457
human.FAM30B-LOC105372935-GGT2.NCBI.                                        cagaggttgcagtaagctgagatcgcgccactgcactccagcctggtgacagagtgagac 34393
human.FAM230E-LOC105377182-GGT3P.NCBI.                                      cagaggttgcagtaagctgagatcgcgccactgcactccagcctggtgacagagtgagac 33131
human.FAM230J-LOC105372942-GGTLC5P.NCBI.rev.compl                           cagaggttgcagtaagctgagatcgcgccactgcactccagcctggtgacagagtgagac 33850
human.FAM230A.ncRNA.18487127.18500594-GGTLC3.18516335.18518165.rev.compl.REF cagaggttgcagtaagctgagatcgcgccactgcactccagcctggtgacagagtgagac 22601
                                                                            ************************  **********************************

chimp.LOC112206744-LOC107973052-GGT2.revcompl                               accgtctc--aaaaaaaaaaaaaaaaaaaaaaaagcaggacactgaactctgggagggcc 35776
chimp54546-135457.revcompl.Pan.troglodytes.clone.rp43-41g5.GenBank:AC099533.36 ACCGTCTC---AAAAGAAAAAAAAAAAAAAAAAAGCAGGACACTGAACTCTGGGAGGGCC 37514
human.FAM30B-LOC105372935-GGT2.NCBI.                                        accgtcac---------aaaaaaaaaaaaaaaaagcaggacactgaactctgggagggcc 34444
human.FAM230E-LOC105377182-GGT3P.NCBI.                                      accgtcacaaaaaaaaaaaaaaaaaaaaaaaaaagcaggacactgaactctgggagggcc 33191
human.FAM230J-LOC105372942-GGTLC5P.NCBI.rev.compl                           accgtcacaaaaaaaaaaaaaaaaaaaaaaaaaagcaggacactgaactctgggagggcc 33910
human.FAM230A.ncRNA.18487127.18500594-GGTLC3.18516335.18518165.rev.compl.REF accgtccc---aaaaaaaaaaaaaaaaaaaaaaagcaggacactgaactctgggagggcc 22658
                                                                            ****** *         *******************************************

chimp.LOC112206744-LOC107973052-GGT2.revcompl                               tcctggtgagaggtgagcacagaggggagagatggaggcaggagcatgggcttctggtgg 35836
chimp54546-135457.revcompl.Pan.troglodytes.clone.rp43-41g5.GenBank:AC099533.36 TCCTGGTGAGAGGTGAGCACAGAGGGGAGAGATGGAGGCAGGAGCATGGGCTTCTGGTGG 37574
human.FAM30B-LOC105372935-GGT2.NCBI.                                        tcctggtgagaggtgagcacagaggggagagatggaggcaggagcatgggcttctggtgg 34504
```

```
human.FAM230E-LOC105377182-GGT3P.NCBI.                                            tcctggtgagaggtgagcacagaggggagagatggaggcaggagcatgggcttctggtgg 33251
human.FAM230J-LOC105372942-GGTLC5P.NCBI.rev.compl                                 tcctggtgagaggtgagcacagaggggagagatggaggcaggagcatgggcttctggtgg 33970
human.FAM230A.ncRNA.18487127.18500594-GGTLC3.18516335.18518165.rev.compl.REF      tcctggtgagaggtgagcacagaggggagagatggaggcaggagcatgggcttctggtgg 22718
                                                                                  ************************************************************

chimp.LOC112206744-LOC107973052-GGT2.revcompl                                     ccccagcagaccctgtggcagcgtggccagggtcctctgcagggaggaatcttggccagg 35896
chimp54546-135457.revcompl.Pan.troglodytes.clone.rp43-41g5.GenBank:AC099533.36    CCCCAGCAGACCCTGTGGCAATGTGGCCAGGGGCCTCTGCAGGGAGGAATCTTGGCCAGG 37634
human.FAM30B-LOC105372935-GGT2.NCBI.                                              ccccagcagaccctgtggcagcgtggccagggtcctctgcagggaggaatcttggccagg 34564
human.FAM230E-LOC105377182-GGT3P.NCBI.                                            ccccagcagaccctgtggcagcgtggccagggtcctctgcagggaggaatcttggccagg 33311
human.FAM230J-LOC105372942-GGTLC5P.NCBI.rev.compl                                 ccccagcagaccctgtggcagcgtggccagggtcctctgcagggaggaatcttggccagg 34030
human.FAM230A.ncRNA.18487127.18500594-GGTLC3.18516335.18518165.rev.compl.REF      ccccagcagaccctgtggcagcgtggccagggtcctctgcagggaggaatcttggccagg 22778
                                                                                  ********************  ********** ***************************

chimp.LOC112206744-LOC107973052-GGT2.revcompl                                     atgacgctgtagcaggcctcttcctgaggcctccagccagcccggccagggacccagcgt 35956
chimp54546-135457.revcompl.Pan.troglodytes.clone.rp43-41g5.GenBank:AC099533.36    ATGACGCTGTAGCAGGCCTCTTCCTGAGGCCTCCAGCCAGCCCGGCCAGGGTCCCAGCGT 37694
human.FAM30B-LOC105372935-GGT2.NCBI.                                              atgacgctgtagcaggcctcttcctgaggcctccagccagcccggccagggtcccagcgt 34624
human.FAM230E-LOC105377182-GGT3P.NCBI.                                            atgacgctgtagcaggcctcttcctgaggcctccagccagcccggccagggtcccagcgt 33371
human.FAM230J-LOC105372942-GGTLC5P.NCBI.rev.compl                                 atgacgctgtagcaggcctcttcctgaggcctccagccagcccggccagggtcccagcgt 34090
human.FAM230A.ncRNA.18487127.18500594-GGTLC3.18516335.18518165.rev.compl.REF      atgacgctgtagcaggcctcttcctgaggcctccagccagcccggccagggtcccagcgt 22838
                                                                                  *************************************************** ********

chimp.LOC112206744-LOC107973052-GGT2.revcompl                                     ccagtgacccctgtttcacagcagcagctggggccagccccaggctctcttccactccca 36016
chimp54546-135457.revcompl.Pan.troglodytes.clone.rp43-41g5.GenBank:AC099533.36    CCAGTGACCCCTGTTTCGCAGCAGCAGCTGGCGCCAGCCCCAGGCTCTCTTCCACTCCCA 37754
human.FAM30B-LOC105372935-GGT2.NCBI.                                              ccagtgacccctgtttcacagcagcagctggggccagccccaggctctcttccactccca 34684
human.FAM230E-LOC105377182-GGT3P.NCBI.                                            ccagtgacccctgtttcacagcagcagctggggccagccccaggctctcttccactccca 33431
human.FAM230J-LOC105372942-GGTLC5P.NCBI.rev.compl                                 ccagtgacccctgtttcacagcagcagctggggccagccccaggctctcttccactccca 34150
human.FAM230A.ncRNA.18487127.18500594-GGTLC3.18516335.18518165.rev.compl.REF      ccagtgacccctgtttcacagcagcagctggggccagccccaggctctcttccactccca 22898
                                                                                  ***************** ************* ****************************

chimp.LOC112206744-LOC107973052-GGT2.revcompl                                     gcttcttaaaactggaagtggagagagttgtttgataaaacactggggcaaaccacatcc 36076
chimp54546-135457.revcompl.Pan.troglodytes.clone.rp43-41g5.GenBank:AC099533.36    GCTTCTTAAAACTGGAAGTGGAGAGAGTAGTTTGATAAAACAGTGGGGCAAACCACATCC 37814
human.FAM30B-LOC105372935-GGT2.NCBI.                                              gcttcttaaaactggaagtggagagagttgtttgataaaacactggggcaaaccacatcc 34744
human.FAM230E-LOC105377182-GGT3P.NCBI.                                            gcttcttaaaactggaagtggagagagttgtttgataaaacactggggcaaaccacatcc 33491
human.FAM230J-LOC105372942-GGTLC5P.NCBI.rev.compl                                 gcttcttaaaactggaagtggagagagttgtttgataaaacactggggcaaaccacatcc 34210
human.FAM230A.ncRNA.18487127.18500594-GGTLC3.18516335.18518165.rev.compl.REF      gcttcttaaaactggaagtggagagagttgtttgataaaacactggggcaaaccacatcc 22958
                                                                                  **************************** ************* *****************

chimp.LOC112206744-LOC107973052-GGT2.revcompl                                     tctcttcaccaagggagagttcgaggggatgccagcagagggcgctttagagtagagacc 36136
chimp54546-135457.revcompl.Pan.troglodytes.clone.rp43-41g5.GenBank:AC099533.36    TCTCTTCACCAAGGGAGAGTTCGAGGGGATGCCGGCAGAGGGAGCTTTAGAGTAGAGACC 37874
human.FAM30B-LOC105372935-GGT2.NCBI.                                              tctcttcaccaagggagagttggaggggatgccggcagagggagctttagagtagagacc 34804
human.FAM230E-LOC105377182-GGT3P.NCBI.                                            tctcttcaccaagggagagttcgaggggatgccggcagagggagctttagagtagagacc 33551
human.FAM230J-LOC105372942-GGTLC5P.NCBI.rev.compl                                 tctcttcaccaagggagagttcgaggggatgccggcagagggagctttagagtagagacc 34270
human.FAM230A.ncRNA.18487127.18500594-GGTLC3.18516335.18518165.rev.compl.REF      tctcttcaccaagggagagttcgaggggatgccggcagagggagctttagagtagagacc 23018
                                                                                  ********************* *********** ******** *****************

chimp.LOC112206744-LOC107973052-GGT2.revcompl                                     cctacccaaccagtgaccgtcacgcacacagcagggcatgctatggagacccccagacag 36196
chimp54546-135457.revcompl.Pan.troglodytes.clone.rp43-41g5.GenBank:AC099533.36    CCTACCCAACCAGTGACCGTCACGCACACAGCAGGGTATGCTATGGAGACCCCCAGACAG 37934
human.FAM30B-LOC105372935-GGT2.NCBI.                                              cctacccaaccagtgaccgtcacgcacacagcagggcatgctatggagacccccagacag 34864
human.FAM230E-LOC105377182-GGT3P.NCBI.                                            cctacccaaccagtgaccgtcacgcacacagcagggcatgctatggagacccccagacag 33611
human.FAM230J-LOC105372942-GGTLC5P.NCBI.rev.compl                                 cctacccaaccagtgaccgtcacgcacacagcagggcatgctatggagacccccagacag 34330
human.FAM230A.ncRNA.18487127.18500594-GGTLC3.18516335.18518165.rev.compl.REF      cctacccaaccagtgaccgtcacgcacacagcagggcatgctatggagacccccagacag 23078
                                                                                  ************************************ ***********************

chimp.LOC112206744-LOC107973052-GGT2.revcompl                                     tcactcggggagacccagcaggtccagactcttcagagatctgtggcagcaggtccccac 36256
chimp54546-135457.revcompl.Pan.troglodytes.clone.rp43-41g5.GenBank:AC099533.36    TCACTCTGGGAGACCCAGCAGGTCCAGACTCTTCAGAGATCTGTGGCAGCAGGTCCCCAC 37994
human.FAM30B-LOC105372935-GGT2.NCBI.                                              tcactcggggagacccagcaggtccagactcttcagagatctgtggcagcaggtccccac 34924
human.FAM230E-LOC105377182-GGT3P.NCBI.                                            tcactcggggagacccagcaggtccagactcttcagagatctgtggcagcaggtccccac 33671
human.FAM230J-LOC105372942-GGTLC5P.NCBI.rev.compl                                 tcactcggggagacccagcaggtccagactcttcagagatctgtggcagcaggtccccac 34390
human.FAM230A.ncRNA.18487127.18500594-GGTLC3.18516335.18518165.rev.compl.REF      tcactcggggagacccagcaggtccagactcttcagagatctgtggcagcaggtccccac 23138
                                                                                  ****** *****************************************************

chimp.LOC112206744-LOC107973052-GGT2.revcompl                                     tcccaaaagccacgtgcccacgggtggtctctggtgcctgagaccccagtctcatttgca 36316
chimp54546-135457.revcompl.Pan.troglodytes.clone.rp43-41g5.GenBank:AC099533.36    TCCCAAAAGCCACGTGCTCACGGGTGGTCTCTGGTGCCTGAGACCCCAGTCTCATTTGCA 38054
human.FAM30B-LOC105372935-GGT2.NCBI.                                              tcccaaaagccacgtgcccacgggtggtctctggtgcctgagaccccagtctcatttgca 34984
human.FAM230E-LOC105377182-GGT3P.NCBI.                                            tcccaaaagccacgtgcccacgggtggtctctggtgcctgagaccccagtctcatttgca 33731
human.FAM230J-LOC105372942-GGTLC5P.NCBI.rev.compl                                 tcccaaaagccacgtgcccacgggtggtctctggtgcctgagaccccagtctcatttgca 34450
human.FAM230A.ncRNA.18487127.18500594-GGTLC3.18516335.18518165.rev.compl.REF      tcccaaaagccacgtgcccacgggtggtctctggtgcctgagaccccagtctcatttgca 23198
                                                                                  ***************** ******************************************

chimp.LOC112206744-LOC107973052-GGT2.revcompl                                     tctttgcaacttcgagtttaagtgggtgtcgcatctctgtatgtcctcctgagcagagga 36376
chimp54546-135457.revcompl.Pan.troglodytes.clone.rp43-41g5.GenBank:AC099533.36    TCTTTGCAACTTTGAGTTTAAGTGGGTGTCGCATCTCTGTATGTCCCCCTGAGCAGAGGA 38114
human.FAM30B-LOC105372935-GGT2.NCBI.                                              tctttgcaacttcgagtttaagtgggtgtcgcatctctgtatgtcctcccgagcagagga 35044
human.FAM230E-LOC105377182-GGT3P.NCBI.                                            tctttgcaacttcgagtttaagtgggtgtcgcatctctgtatgtcctcccgagcagagga 33791
human.FAM230J-LOC105372942-GGTLC5P.NCBI.rev.compl                                 tctttgcaacttcgagtttaagtgggtgtcgcatctctgtatgtcctcccgagcagagga 34510
human.FAM230A.ncRNA.18487127.18500594-GGTLC3.18516335.18518165.rev.compl.REF      tctttgcaacttcgagtttaagtgggtgtcgcatctctgtatgtcctcccgagcagagga 23258
                                                                                  ************ ********************************* ** **********

chimp.LOC112206744-LOC107973052-GGT2.revcompl                                     ggggcacagcctggggtggcagctggcgtcaaaccctcaaatcccctgagagccactggg 36436
chimp54546-135457.revcompl.Pan.troglodytes.clone.rp43-41g5.GenBank:AC099533.36    GGGGCACAGCCTGGGGTGGCGGCTGGCGTCAAACCCTCAAATCCCCTGAGAGCCACTGGG 38174
human.FAM30B-LOC105372935-GGT2.NCBI.                                              ggggcacagcctggggtggcagctggcgtcaaaccctcaaatcccctgagagccactggg 35104
human.FAM230E-LOC105377182-GGT3P.NCBI.                                            ggggcacagcctggggtggcagctgacgtcaaaccctcaaatcccctgagagccactggg 33851
human.FAM230J-LOC105372942-GGTLC5P.NCBI.rev.compl                                 ggggcacagcctggggtggcagctgacgtcaaaccctcaaatcccctgagagccactggg 34570
human.FAM230A.ncRNA.18487127.18500594-GGTLC3.18516335.18518165.rev.compl.REF      ggggcacagcctggggtggcagctggcgtcaaaccctcaaatcccctgagagccactggg 23318
                                                                                  ******************** **** **********************************

chimp.LOC112206744-LOC107973052-GGT2.revcompl                                     gagactaagcagtccccagcccccacttgtccctgagctgccattctcagccctgtggga 36496
chimp54546-135457.revcompl.Pan.troglodytes.clone.rp43-41g5.GenBank:AC099533.36    GAGACTAAGCAGTCTCCAGCCCCCACTTGTCCCTGAGCTGCCATTCTCAGCCCTGTGGGA 38234
human.FAM30B-LOC105372935-GGT2.NCBI.                                              gagactaagcagtccccagcccctacttgtccctgagctgccattctcagccctgtggga 35164
human.FAM230E-LOC105377182-GGT3P.NCBI.                                            gagactaagcagtccccagcccccacttgtccctgagctgccattctcagccctgtggga 33911
human.FAM230J-LOC105372942-GGTLC5P.NCBI.rev.compl                                 gagactaagcagtccccagcccccacttgtccctgagctgccattctcagccctgtggga 34630
human.FAM230A.ncRNA.18487127.18500594-GGTLC3.18516335.18518165.rev.compl.REF      gagactaagcagtccccagcccccacttgtccctgagctgccattctcagccctgtggga 23378
                                                                                  ************** ******** ************************************

chimp.LOC112206744-LOC107973052-GGT2.revcompl                                     ggagacagaaagccctgaagagaaaccaaaggaccaggtcaggaggggct--gggggggt 36554
chimp54546-135457.revcompl.Pan.troglodytes.clone.rp43-41g5.GenBank:AC099533.36    GGAGACAGAAAGCCCTGAAGAGAAACCAAAGGACCAGGTCAGGAGGGGCT--GGGGGGGT 38292
human.FAM30B-LOC105372935-GGT2.NCBI.                                              ggagacagaaagccctgaagagaaaccaaaggaccaggtcaggaggggctgggggggggt 35224
human.FAM230E-LOC105377182-GGT3P.NCBI.                                            ggagacagaaagccctgaagagaaaccaaaggaccaggtcaggaggggct--gggggggt 33969
human.FAM230J-LOC105372942-GGTLC5P.NCBI.rev.compl                                 ggagacagaaagccctgaagagaaaccaaaggaccaggtcaggaggggct--gggggggt 34688
human.FAM230A.ncRNA.18487127.18500594-GGTLC3.18516335.18518165.rev.compl.REF      ggagacagaaagccctgaagagaaaccaaaggaccaggtcaggaggggct--gggggggt 23436
                                                                                  **************************************************  ********

chimp.LOC112206744-LOC107973052-GGT2.revcompl                                     ggcatgagcaatcagggcagggaaggatggacagatggggaatggagggaagaaggaat 36614
chimp54546-135457.revcompl.Pan.troglodytes.clone.rp43-41g5.GenBank:AC099533.36    GGCATGAGCAATCAGGGCAGGGAAGGATGGACAGATGGGGAATGGAGGGAAGAAGGAAT 38352
human.FAM30B-LOC105372935-GGT2.NCBI.                                              ggcatgagcaatcagggcagggaaggatgcacagatggggaatggagggaagaaggaat 35284
human.FAM230E-LOC105377182-GGT3P.NCBI.                                            ggcatgagcaatcagggcagggaaggatggacagatggggaatggagggaagaaggaat 34029
human.FAM230J-LOC105372942-GGTLC5P.NCBI.rev.compl                                 ggcatgagcaatcagggcagggaaggatggacagatggggaatggagggaagaaggaat 34748
human.FAM230A.ncRNA.18487127.18500594-GGTLC3.18516335.18518165.rev.compl.REF      ggcatgagcaatcagggcagggaaggatggaccgatggggaatggagggaagaaggaat 23496
                                                                                  ***************************** ** ***************************

chimp.LOC112206744-LOC107973052-GGT2.revcompl                                     gaatgaaacggtgaatgaatgaacaaagagagagaacggccattcctcccttgctttagt 36674
chimp54546-135457.revcompl.Pan.troglodytes.clone.rp43-41g5.GenBank:AC099533.36    GAATGAAAAGTTGAATGAATGAACAAAGAGAGAGAATGGCCATTCCTCCCTTGCTTTAGT 38412
human.FAM30B-LOC105372935-GGT2.NCBI.                                              gaatgaaaaggtgaatgaatgaacaaagagagagaacggccactcctcccttgctttagt 35344
human.FAM230E-LOC105377182-GGT3P.NCBI.                                            gaatgaaaaggtgaatgaatgaacaaagagagagaacggccactcctcccttgctttagt 34089
human.FAM230J-LOC105372942-GGTLC5P.NCBI.rev.compl                                 gaatgaaaaggtgaatgaatgaacaaagagagagaacggccactcctcccttgctttagt 34808
human.FAM230A.ncRNA.18487127.18500594-GGTLC3.18516335.18518165.rev.compl.REF      gaatgaaaaggtgaatgaatgaacaaagagagagaacggccactcctcccttgctttagt 23556
                                                                                  ******** * ************************* ***** *****************

chimp.LOC112206744-LOC107973052-GGT2.revcompl                                     ttacaaagtactgggatcctcccaacagcctgcaagacagaatttctgggaagcagacca 36734
chimp54546-135457.revcompl.Pan.troglodytes.clone.rp43-41g5.GenBank:AC099533.36    TTACAAAGTACTGGGATCCTCCCAACAGCCTGCAAGACAGAATTTCTGGGAAGCAGACCA 38472
human.FAM30B-LOC105372935-GGT2.NCBI.                                              ttacaaagtactgggatcctcccaacagcctgcaagacagaatttctgggaagcagacca 35404
human.FAM230E-LOC105377182-GGT3P.NCBI.                                            ttacaaagtactgggatcctcccaacagcctgcaagacagaatttctgggaagcagacca 34149
human.FAM230J-LOC105372942-GGTLC5P.NCBI.rev.compl                                 ttacaaagtactgggatcctcccaacagcctgcaagacagaatttctgggaagcagacca 34868
human.FAM230A.ncRNA.18487127.18500594-GGTLC3.18516335.18518165.rev.compl.REF      ttacaaagtactgggatcctcccaacagcctgcaagacagaatttctgggaagcagacca 23616
                                                                                  ************************************************************

chimp.LOC112206744-LOC107973052-GGT2.revcompl                                     ggtggctggcagggaggggaggcttgccctggcttttgtgggcccaatgggaggcagggg 36794
chimp54546-135457.revcompl.Pan.troglodytes.clone.rp43-41g5.GenBank:AC099533.36    GGTGGCTGGCAGGGAGGGGAGGCTTGCCCTGGCTTTTGTGGGCCCAATGGGAGGCAGGGG 38532
human.FAM30B-LOC105372935-GGT2.NCBI.                                              ggtggctggcagggaggggaggcttgccctggcttttgtgggcccaatgggaggcagggg 35464
human.FAM230E-LOC105377182-GGT3P.NCBI.                                            ggtggctggcagggaggggaggcttgccctggcttttgtgggcccaatgggaggcagggg 34209
human.FAM230J-LOC105372942-GGTLC5P.NCBI.rev.compl                                 ggtggctggcagggaggggaggcttgccctggcttttgtgggcccaatgggaggcagggg 34928
human.FAM230A.ncRNA.18487127.18500594-GGTLC3.18516335.18518165.rev.compl.REF      ggtggctggcagggaggggaggcttgccctggcttttgtgggcccaatgggaggcagggg 23676
                                                                                  ************************************************************

chimp.LOC112206744-LOC107973052-GGT2.revcompl                                     gcaagaaggggcatcctgtgtgtgtcctccctgcagcggcagcagcaccttcctggaaga 36854
chimp54546-135457.revcompl.Pan.troglodytes.clone.rp43-41g5.GenBank:AC099533.36    GCAAGAAGGGGCATCCTGTGTGTGTCCTCCCTGCAGCGGCAGCAGCACCTTCCTGGAAGA 38592
human.FAM30B-LOC105372935-GGT2.NCBI.                                              gcaagaaggggcatcctgtgtgtgtcctccctgcagcggcagcagcaccttcctggaaga 35524
human.FAM230E-LOC105377182-GGT3P.NCBI.                                            gcaagaaggggcatcctgtgtgtgtcctccctgcagcggcagcagcaccttcctggaaga 34269
human.FAM230J-LOC105372942-GGTLC5P.NCBI.rev.compl                                 gcaagaaggggcatcctgtgtgtgtcctccctgcagcggcagcagcaccttcctggaaga 34988
human.FAM230A.ncRNA.18487127.18500594-GGTLC3.18516335.18518165.rev.compl.REF      gcaagaaggggcatcctgtgtgtgtcctccctgcagtggcagcagcaccttcctggaaga 23736
                                                                                  ************************************ ***********************

chimp.LOC112206744-LOC107973052-GGT2.revcompl                                     gggtcaggaaacactcactgtggcccctctccatcacgacctcatccaggacaccaaata 36914
chimp54546-135457.revcompl.Pan.troglodytes.clone.rp43-41g5.GenBank:AC099533.36    GGGTCAGGAAACACCCACTGTGGCCCCTCTCCATCACGCCCTCATCCAGGACACCAAGTA 38652
human.FAM30B-LOC105372935-GGT2.NCBI.                                              gggtcaggaaacacccgctgtggcccctctccaccacgccctcatccaggacaccaagta 35584
human.FAM230E-LOC105377182-GGT3P.NCBI.                                            gggtcaggaaacacccgctgtggcccctctccaccacgccctcatccaggacaccaagta 34329
```

```
human.FAM230J-LOC105372942-GGTLC5P.NCBI.rev.compl                                 gggtcaggaaacacccgctgtggcccctctccaccacgccctcatccaggacaccaagta 35048
human.FAM230A.ncRNA.18487127.18500594-GGTLC3.18516335.18518165.rev.compl.REF      gggtcaggaaacacccgctgtggcccctctccaccacgccctcatccaggacaccaagta 23796
                                                                                  ************** * **************** **** ****************** **

chimp.LOC112206744-LOC107973052-GGT2.revcompl                                     tcaagtcactcagctcacgagacccaggccctgactcagggagagaggatgtgaggggtg 36974
chimp54546-135457.revcompl.Pan.troglodytes.clone.rp43-41g5.GenBank:AC099533.36    TC-AGTCACTCGGCTCACGAGATCCAGGCCCTGACTCAGGGAGAGAGGATGTGAGGGGTG 38711
human.FAM30B-LOC105372935-GGT2.NCBI.                                              tc-agtcactcagctcacgagacccaggccctgactcagggagagaggatgtgaggggtg 35643
human.FAM230E-LOC105377182-GGT3P.NCBI.                                            tc-agtcactcagctcacgagacccaggccctgactcagggagagaggatgtgaggggtg 34388
human.FAM230J-LOC105372942-GGTLC5P.NCBI.rev.compl                                 tc-agtcactcagctcacgagacccaggccctgactcagggagagaggatgtgaggggtg 35107
human.FAM230A.ncRNA.18487127.18500594-GGTLC3.18516335.18518165.rev.compl.REF      tc-agtcactcagctcacgagacccaggccctgactcagggagagaggatgtgaggggtg 23855
                                                                                  ** ******** ********* **************************************

chimp.LOC112206744-LOC107973052-GGT2.revcompl                                     gggcaccgggctcctcaggactgagagacctgagatgtggccccgggctgggtgttgggg 37034
chimp54546-135457.revcompl.Pan.troglodytes.clone.rp43-41g5.GenBank:AC099533.36    GGGCACCGGGCTCCTCAGGACTGAGAGACCTGAGATGTGGCCCCGGGCTGGGTGTTGGGG 38771
human.FAM30B-LOC105372935-GGT2.NCBI.                                              gggcaccgggctcctcaggactgagagacctgagatgtggccccgggctgggtgtagggg 35703
human.FAM230E-LOC105377182-GGT3P.NCBI.                                            gggcaccgggctcctcaggactgagagacctgagatgtggccccgggctgggtgtagggg 34448
human.FAM230J-LOC105372942-GGTLC5P.NCBI.rev.compl                                 gggcaccgggctcctcaggactgagagacctgagatgtggccccgggctgggtgtagggg 35167
human.FAM230A.ncRNA.18487127.18500594-GGTLC3.18516335.18518165.rev.compl.REF      gggcaccgggctcctcaggactgagagacctgggatgtggccccgggctgggtgtagggg 23915
                                                                                  ******************************** ********************** ****

chimp.LOC112206744-LOC107973052-GGT2.revcompl                                     cagactggctatggcagcattgtgtgtaccccagcaggccagtacccatgcagggagcct 37094
chimp54546-135457.revcompl.Pan.troglodytes.clone.rp43-41g5.GenBank:AC099533.36    CAGACTGGCTATGGCAGCATTGTGTGTACCCCAGCAGGCCAGTACCCACGCAGGGAGCCT 38831
human.FAM30B-LOC105372935-GGT2.NCBI.                                              cagactggctatggcagcattgtgtgtaccccagcaggccagtacccacgcagggagcct 35763
human.FAM230E-LOC105377182-GGT3P.NCBI.                                            cagactggctatggcagcattgtgtgtaccccagcaggccagtacccacgcagggagcct 34508
human.FAM230J-LOC105372942-GGTLC5P.NCBI.rev.compl                                 cagactggctatggcagcattgtgtgtaccccagcaggccagtacccacgcagggagcct 35227
human.FAM230A.ncRNA.18487127.18500594-GGTLC3.18516335.18518165.rev.compl.REF      cagactggctatggcagcattgtgtgtaccccagcaggccagtacccacgcagggagcct 23975
                                                                                  ************************************************ ***********

chimp.LOC112206744-LOC107973052-GGT2.revcompl                                     ccaaaccccttcacccatgaccctgggagaagaccgcaggcttggagaattggcctcact 37154
chimp54546-135457.revcompl.Pan.troglodytes.clone.rp43-41g5.GenBank:AC099533.36    CCAAACCCCTTCACCCATGACCCTGGGAGAAGACCCCAGCCTTGGAGAATT-GCCTCACT 38890
human.FAM30B-LOC105372935-GGT2.NCBI.                                              ccaaaccccttcacccatgaccctgggagaagaccgcagccttggagaattggcctcact 35823
human.FAM230E-LOC105377182-GGT3P.NCBI.                                            ccaaaccccttcacccatgaccctgggagaagaccgcagccttggagaattggcctcact 34568
human.FAM230J-LOC105372942-GGTLC5P.NCBI.rev.compl                                 ccaaaccccttcacccatgaccctgggagaagaccgcagccttggagaattggcctcact 35287
human.FAM230A.ncRNA.18487127.18500594-GGTLC3.18516335.18518165.rev.compl.REF      ccaaaccccttcacccatgaccctgggagaagaccgcagccttggagaattggcctcact 24035
                                                                                  *********************************** *** *********** ********

chimp.LOC112206744-LOC107973052-GGT2.revcompl                                     gaagggccctgcaccggccagcagggtcaggcggggccagacaggttcccacctgggata 37214
chimp54546-135457.revcompl.Pan.troglodytes.clone.rp43-41g5.GenBank:AC099533.36    GAAGGGACCTGCACTGGCCAGCAGGGTCAGGCGGGGCCAGACAGGTTCCCACCTGGGATA 38950
human.FAM30B-LOC105372935-GGT2.NCBI.                                              gaaggggcctgcgccggccagcagggtcaggcggggccagacaggttcccacctgggata 35883
human.FAM230E-LOC105377182-GGT3P.NCBI.                                            gaaggggcctgcaccggccagcagggtcaggcggggccagacaggttcccacctgggata 34628
human.FAM230J-LOC105372942-GGTLC5P.NCBI.rev.compl                                 gaaggggcctgcaccggccagcagggtcaggcggggccagacaggttcccacctgggata 35347
human.FAM230A.ncRNA.18487127.18500594-GGTLC3.18516335.18518165.rev.compl.REF      gaaggggcctgcaccggccagcagggtcaggcggggccagacaggttcccacctgggata 24095
                                                                                  ****** ***** * *********************************************

chimp.LOC112206744-LOC107973052-GGT2.revcompl                                     tgcaaatgggcctcctgaatcctggagccaggtatggactcacacaccaccattgtcccc 37274
chimp54546-135457.revcompl.Pan.troglodytes.clone.rp43-41g5.GenBank:AC099533.36    TGCAAATGGGCCTCCTGAATCCTGGAGCCAGGTATGGACTCACACACCACCATTGTCCCC 39010
human.FAM30B-LOC105372935-GGT2.NCBI.                                              tgcaaatgggcctcctgaatcctggagccaggtatggactcacacaccaccattgtcccc 35943
human.FAM230E-LOC105377182-GGT3P.NCBI.                                            tgcaaatgggcctcctgaatcctggagccaggtatggactcacacaccaccattgtcccc 34688
human.FAM230J-LOC105372942-GGTLC5P.NCBI.rev.compl                                 tgcaaatgggcctcctgaatcctggagccaggtatggactcacacaccaccattgtcccc 35407
human.FAM230A.ncRNA.18487127.18500594-GGTLC3.18516335.18518165.rev.compl.REF      tgcaaatgggcctcctgaatcctggagccaggtatggactcacacaccaccattgtcccc 24155
                                                                                  ************************************************************

chimp.LOC112206744-LOC107973052-GGT2.revcompl                                     aagtccccatctgccccacgggcacaccctgccacctgttctgtgcaagagccctgaggc 37334
chimp54546-135457.revcompl.Pan.troglodytes.clone.rp43-41g5.GenBank:AC099533.36    AAGTCCCCATCTGCCCCACGGGCACACCCTGCCACCTGTTCTGTGCAAAGGCCCTGAGGC 39070
human.FAM30B-LOC105372935-GGT2.NCBI.                                              aagtccccatctgccccacgggcacaccctgccgcctgttctgtgcaagggccctgaggc 36003
human.FAM230E-LOC105377182-GGT3P.NCBI.                                            aagtccccatctgccccacgggcacaccctgccgcctgttctgtgcaagggccctgaggc 34748
human.FAM230J-LOC105372942-GGTLC5P.NCBI.rev.compl                                 aagtccccatctgccccacgggcacaccctgccgcctgttctgtgcaagggccctgaggc 35467
human.FAM230A.ncRNA.18487127.18500594-GGTLC3.18516335.18518165.rev.compl.REF      aagtccccatctgccccacgggcacaccctgccgcctgttctgtgcaagggccctgaggc 24215
                                                                                  ********************************* **************  **********

chimp.LOC112206744-LOC107973052-GGT2.revcompl                                     tgtctccttgcgctcaagccctgcaggtgctgaagcccacacacacagctcctgcttcct 37394
chimp54546-135457.revcompl.Pan.troglodytes.clone.rp43-41g5.GenBank:AC099533.36    TGTCTCCTTGCGCTCAAGCCCTGCAGGTGCTGAAGCCCACACACACGGCTCCTGCTTCCT 39130
human.FAM30B-LOC105372935-GGT2.NCBI.                                              tgtctccttgcgctcaagccctgcaggtgctgaagcccacacacacggctcctgcttcct 36063
human.FAM230E-LOC105377182-GGT3P.NCBI.                                            tgtctccttgcgctcaagccctgcaggtgctgaagcccacacacacggctcctgcttact 34808
human.FAM230J-LOC105372942-GGTLC5P.NCBI.rev.compl                                 tgtctccttgcgctcaagccctgcaggtgctgaagcccacacacacggctcctgcttcct 35527
human.FAM230A.ncRNA.18487127.18500594-GGTLC3.18516335.18518165.rev.compl.REF      tgtctccttgcgcccaagacctgcaggtgctgaagcccacacacacggctcctgcttcct 24275
                                                                                  ************* **** *************************** ********** **

chimp.LOC112206744-LOC107973052-GGT2.revcompl                                     gggccagtgcacgtgcacacacacacacgtgcacacacacaccc------acaaatatac 37448
chimp54546-135457.revcompl.Pan.troglodytes.clone.rp43-41g5.GenBank:AC099533.36    GGGCCAGTGCACGTGCACACACACACACATGCACACACACACCCACACATACAC--ATAC 39188
human.FAM30B-LOC105372935-GGT2.NCBI.                                              gggccagtgcacgtgcacacacacacacgtgcacacacacacacacccccacaaatatac 36123
human.FAM230E-LOC105377182-GGT3P.NCBI.                                            gggccagtgcacgtgcacacacacacacgtg--cacacacacacacacccacaaatatac 34866
human.FAM230J-LOC105372942-GGTLC5P.NCBI.rev.compl                                 gggccagtgcacgtgcacacacacacacgtg--cacacacacacacacccacaaatatac 35585
human.FAM230A.ncRNA.18487127.18500594-GGTLC3.18516335.18518165.rev.compl.REF      gggccagtgcacgtgcacacacacacacgtgcacacacacacacacacccacaaatatac 24335
                                                                                  **************************** **  ********* *      ***   ****

chimp.LOC112206744-LOC107973052-GGT2.revcompl                                     ccacacacaatcacacacattcacacatacccaccccca-------tacactcacacatt 37501
chimp54546-135457.revcompl.Pan.troglodytes.clone.rp43-41g5.GenBank:AC099533.36    CCACACACAATCACACACATTCACACATACCCACACCCCCCATACTCACACTCACACACT 39248
human.FAM30B-LOC105372935-GGT2.NCBI.                                              ccacacacaatcacacacattcacacatacccac-ccccccatactcacactcacacatt 36182
human.FAM230E-LOC105377182-GGT3P.NCBI.                                            ccacacacaatcacacacattcacacataccca--ccccccatactcacactcacacatt 34924
human.FAM230J-LOC105372942-GGTLC5P.NCBI.rev.compl                                 ccacacacaatcacacacattcacacataccca--ccccccatactcacactcacacatt 35643
human.FAM230A.ncRNA.18487127.18500594-GGTLC3.18516335.18518165.rev.compl.REF      ccacacacaatcacacacattcacacataccca--ccccccatactcaca--cacacatt 24391
                                                                                  *********************************  ***         ***  ****** *

chimp.LOC112206744-LOC107973052-GGT2.revcompl                                     cacacccac------acccacacatacacacacacaatcacacacatttacacacaccca 37555
chimp54546-135457.revcompl.Pan.troglodytes.clone.rp43-41g5.GenBank:AC099533.36    CACACACACCCACAAACACCCACACATACACACACAATCACACACATTTACACACACCCA 39308
human.FAM30B-LOC105372935-GGT2.NCBI.                                              tacacacacc----------------------------------------cacacaccca 36202
human.FAM230E-LOC105377182-GGT3P.NCBI.                                            tacacacacc----------------------------------------cacacaccca 34944
human.FAM230J-LOC105372942-GGTLC5P.NCBI.rev.compl                                 tacacacacc----------------------------------------cacacaccca 35663
human.FAM230A.ncRNA.18487127.18500594-GGTLC3.18516335.18518165.rev.compl.REF      tacacacacc----------------------------------------cacacaccca 24411
                                                                                   **** ***                                         **********

chimp.LOC112206744-LOC107973052-GGT2.revcompl                                     cacacccacactcacacactcagtcacacacaccctcacacagcgaaacacaat--caca 37613
chimp54546-135457.revcompl.Pan.troglodytes.clone.rp43-41g5.GenBank:AC099533.36    CACTCACACACTCA------CAGTCACACA-ACCCTCACACAGCGAAACGCAATCACACA 39361
human.FAM30B-LOC105372935-GGT2.NCBI.                                              cactcacacactca------cagtcacacacaccctcacacagcgaaacacaatcacaca 36256
human.FAM230E-LOC105377182-GGT3P.NCBI.                                            cactcacacactca------cagtcacacacaccctcacacagcgaaacacaatcacaca 34998
human.FAM230J-LOC105372942-GGTLC5P.NCBI.rev.compl                                 cactcacacactca------cagtcacacacaccctcacacagcgaaacacaatcacaca 35717
human.FAM230A.ncRNA.18487127.18500594-GGTLC3.18516335.18518165.rev.compl.REF      cactcacacactca------cagtcacacacaccctcacacagcgaaacacaatcacaca 24465
                                                                                  *** * ********      ********** ****************** ****  ****

chimp.LOC112206744-LOC107973052-GGT2.revcompl                                     cattcacacccacccacacccccacacactcacactcacatatact--------cacaca 37665
chimp54546-135457.revcompl.Pan.troglodytes.clone.rp43-41g5.GenBank:AC099533.36    CATTCACACCCACCCACACCCCCACACACTCACACTCACATATACTCACACACCCACACA 39421
human.FAM30B-LOC105372935-GGT2.NCBI.                                              cattcacacccacccacacccccacacactcacactcacatatact--------cacaca 36308
human.FAM230E-LOC105377182-GGT3P.NCBI.                                            cattcacacccacccacacccccacacactcacactcacatatact--------cacaca 35050
human.FAM230J-LOC105372942-GGTLC5P.NCBI.rev.compl                                 cattcacacccacccacacccccacacactcacactcacatatact--------cacaca 35769
human.FAM230A.ncRNA.18487127.18500594-GGTLC3.18516335.18518165.rev.compl.REF      cattcacacccacccaca-ccccacacactcacactcacatatact--------cacaca 24516
                                                                                  ****************** ***************************        ******

chimp.LOC112206744-LOC107973052-GGT2.revcompl                                     cacccacacacacatacacaaacacaatcacacacattcacacaca-------------- 37711
chimp54546-135457.revcompl.Pan.troglodytes.clone.rp43-41g5.GenBank:AC099533.36    CACCCACACACACATACACAAACACAATCACACACATTCACACACACCCATACTCACACA 39481
human.FAM30B-LOC105372935-GGT2.NCBI.                                              cacccacacacacatacacaaacacaatcacacacattcacacacacccacactcacaca 36368
human.FAM230E-LOC105377182-GGT3P.NCBI.                                            cacccacacacacatacacaaacacaatcacacacattcacacacacccacactcacaca 35110
human.FAM230J-LOC105372942-GGTLC5P.NCBI.rev.compl                                 cacccacacacacatacacaaacacaatcacacacattcacacacacccacactcacaca 35829
human.FAM230A.ncRNA.18487127.18500594-GGTLC3.18516335.18518165.rev.compl.REF      cacccacacacacatacacaaacacaatcacacacattcacacacacccacactcacaca 24576
                                                                                  **********************************************

chimp.LOC112206744-LOC107973052-GGT2.revcompl                                     ----------------------------------------caccccccacaaacacactc 37731
chimp54546-135457.revcompl.Pan.troglodytes.clone.rp43-41g5.GenBank:AC099533.36    TACACGC------AAACGATCACACACATTCACACACCCACACACCCCACAAACACACTC 39535
human.FAM30B-LOC105372935-GGT2.NCBI.                                              tacacacacccaaacacgatcacacacattcacacacccacacaccccacaaacacactc 36428
human.FAM230E-LOC105377182-GGT3P.NCBI.                                            tacacacacccaaacacgatcacacacattcacacacccacacaccccacaaacacactc 35170
human.FAM230J-LOC105372942-GGTLC5P.NCBI.rev.compl                                 tacacacacccaaacacgatcacacacattcacacacccacacaccccacaaacacactc 35889
human.FAM230A.ncRNA.18487127.18500594-GGTLC3.18516335.18518165.rev.compl.REF      tacacacacccaaacacgatcacacacattcacacacccacacaccccacaaacacactc 24636
                                                                                                                          *** ****************

chimp.LOC112206744-LOC107973052-GGT2.revcompl                                     acacatatacccacacacactcactcacacacataatctcacacacacacacacatgctc 37791
chimp54546-135457.revcompl.Pan.troglodytes.clone.rp43-41g5.GenBank:AC099533.36    ACACATATACCCACACACACTCACTCATACACATAATCTCACACACACACACATATGCTC 39595
human.FAM30B-LOC105372935-GGT2.NCBI.                                              acacatatacccacacacactcac------acataatctctcacacacacac--atgctc 36480
human.FAM230E-LOC105377182-GGT3P.NCBI.                                            acacatatacccacacacactcac------acataatctctcacacacacac--atgctc 35222
human.FAM230J-LOC105372942-GGTLC5P.NCBI.rev.compl                                 acacatatacccacacacactcac------acataatctctcacacacacac--atgctc 35941
human.FAM230A.ncRNA.18487127.18500594-GGTLC3.18516335.18518165.rev.compl.REF      acacatatacccacacacactgac------acataatctctcacacacacac--atgctc 24688
                                                                                  ********************* **      ********** ***********  ******

chimp.LOC112206744-LOC107973052-GGT2.revcompl                                     acacacacacgccttctccaggaggggctggctgccaaaggccacccagcttcctcccac 37851
chimp54546-135457.revcompl.Pan.troglodytes.clone.rp43-41g5.GenBank:AC099533.36    ACACACACACGCCTTCTCCAGGAGGGGCTGGCTGCCAAAGGCCCCCCAGCTTCCTCCCAC 39655
human.FAM30B-LOC105372935-GGT2.NCBI.                                              acacacacacgccttctccaggaggggctggctgccaaaggccacccagcttcctcccac 36540
human.FAM230E-LOC105377182-GGT3P.NCBI.                                            acacacacacgccttctccaggaggggctggctgccaaaggccacccagcttcctcccac 35282
human.FAM230J-LOC105372942-GGTLC5P.NCBI.rev.compl                                 acacacacacgccttctccaggaggggctggctgccaaaggccacccagcttcctcccac 36001
human.FAM230A.ncRNA.18487127.18500594-GGTLC3.18516335.18518165.rev.compl.REF      acacacacacgccttctccaggaggggctggctgccaaaggccacccagcttcctcccac 24748
                                                                                  ******************************************* ****************

chimp.LOC112206744-LOC107973052-GGT2.revcompl                                     gtctcactcaccgtacaatatttgagcagaccttggagtcagcagcaacagcagcgtggg 37911
chimp54546-135457.revcompl.Pan.troglodytes.clone.rp43-41g5.GenBank:AC099533.36    GTCTGACTCACCGTACAATATTTGAGCAGACCTTGGAGTCAGCAGCAACAGCAGCGTGGG 39715
human.FAM30B-LOC105372935-GGT2.NCBI.                                              gtctcactcaccgtacaatatttgagaagaccttggagtcagcagcaacagcggcgtggg 36600
human.FAM230E-LOC105377182-GGT3P.NCBI.                                            gtctcactcaccgtacaatatttgagaagaccttggagtcagcagcaacagcggtgtggg 35342
human.FAM230J-LOC105372942-GGTLC5P.NCBI.rev.compl                                 gtctcactcaccgtacaatatttgagaagaccttggagtcagcagcaacagcggtgtggg 36061
```

```
human.FAM230A.ncRNA.18487127.18500594-GGTLC3.18516335.18518165.rev.compl.REF        gtctcactcaccgtacaatatttgagaagaccttggagtcagcagcaacagcggcgtggg 24808
                                                                                    **** ********************* ************************ * *****

chimp.LOC112206744-LOC107973052-GGT2.revcompl                                       caaaggcctgggagccacatggggcctggtgtcgagagaggaccacagccagcacaatga 37971
chimp54546-135457.revcompl.Pan.troglodytes.clone.rp43-41g5.GenBank:AC099533.36      CAAAGGCCTGGGAGCCACATGGGGCCTGGTGTCGAGAGAGGACCACAGCCAGCACAATGA 39775
human.FAM30B-LOC105372935-GGT2.NCBI.                                                caaaggcc-gggggtcaaatggggcctggtgtcgagagaggaccacagccagcacaatga 36659
human.FAM230E-LOC105377182-GGT3P.NCBI.                                              caaaggcc-gggggtcaaatggggcctggtgtcgagagaggaccacagccagcacaatga 35401
human.FAM230J-LOC105372942-GGTLC5P.NCBI.rev.compl                                   caaaggcc-gggggtcaaatggggcctggtgtcgagagaggaccacagccagcacaatga 36120
human.FAM230A.ncRNA.18487127.18500594-GGTLC3.18516335.18518165.rev.compl.REF        caaaggcc-gggggtcaaatggggcctggtgtcgagagaggaccacagccagcacaatga 24867
                                                                                    ******** *** * ** ****************************************

chimp.LOC112206744-LOC107973052-GGT2.revcompl                                       cagccagcgccagccccagccccagccccagcaggaccaggtcacccatggctctgcagt 38031
chimp54546-135457.revcompl.Pan.troglodytes.clone.rp43-41g5.GenBank:AC099533.36      CAGCCAGCGCCAGCCCCAGCCCCAGCCCCAGCAGGACCAGGTCACCCATGGCTCCGCAGT 39835
human.FAM30B-LOC105372935-GGT2.NCBI.                                                cagccagcgccagccccagccccagccccagcaggaccaggtcacccatggctccgtagt 36719
human.FAM230E-LOC105377182-GGT3P.NCBI.                                              cagccagcgccagccccagccccagccccagcaggaccaggtcacccatggctccgtagt 35461
human.FAM230J-LOC105372942-GGTLC5P.NCBI.rev.compl                                   cagccagcgccagccccagccccagccccagcaggaccaggtcacccatggctccgtagt 36180
human.FAM230A.ncRNA.18487127.18500594-GGTLC3.18516335.18518165.rev.compl.REF        cagccagcgccagccccagccccagccccagcaggaccaggtcacccatggctccgtagt 24927
                                                                                    ****************************************************** * ***

chimp.LOC112206744-LOC107973052-GGT2.revcompl                                       cctgggccatggctctgcggcccagaaggagaggggaggccggtgggcagacggagggac 38091
chimp54546-135457.revcompl.Pan.troglodytes.clone.rp43-41g5.GenBank:AC099533.36      CCTGGGCCATGGCTCTGCGGCCCAGAAGGAGAGGGGAGGCCGGTGGGCAGACGGAGGGAC 39895
human.FAM30B-LOC105372935-GGT2.NCBI.                                                cctgggccatggctctgcggcccagaaggagaggggaggccggtgggcagatggagggac 36779
human.FAM230E-LOC105377182-GGT3P.NCBI.                                              cctgggccatggctctgcggcccagaaggagaggggaggccggtgggcagatggagggac 35521
human.FAM230J-LOC105372942-GGTLC5P.NCBI.rev.compl                                   cctgggccatggctctgcggcccagaaggagaggggaggccggtgggcagatggagggac 36240
human.FAM230A.ncRNA.18487127.18500594-GGTLC3.18516335.18518165.rev.compl.REF        cctgggccatggctctgcggcccagaaggagaggggaggccggtgggcagatggagggac 24987
                                                                                    *************************************************** ********

chimp.LOC112206744-LOC107973052-GGT2.revcompl                                       agatgggtgggcagatgaatggacaagaagatgcatagatagactcacaggtaattggac 38151
chimp54546-135457.revcompl.Pan.troglodytes.clone.rp43-41g5.GenBank:AC099533.36      AGATGGGTGGGCAGATGAATGGACAAGAAGATGCATAGATAGACTCACAGGTAATTGGAC 39955
human.FAM30B-LOC105372935-GGT2.NCBI.                                                agatgggtgggcagatgaatggacaagaagatgcatagatagactcacaggtaattggac 36839
human.FAM230E-LOC105377182-GGT3P.NCBI.                                              agatgggtgggcagatgaatggacaagaagatgcatagatagactcacaggtaattggac 35581
human.FAM230J-LOC105372942-GGTLC5P.NCBI.rev.compl                                   agatgggtgggcagatgaatggacaagaagatgcatagatagactcacaggtaattggac 36300
human.FAM230A.ncRNA.18487127.18500594-GGTLC3.18516335.18518165.rev.compl.REF        agatgggtgggcagatgaatggacaagaagatgcatagatagactcacaggtaattggac 25047
                                                                                    ************************************************************

chimp.LOC112206744-LOC107973052-GGT2.revcompl                                       agatggacaaacaggtgggggctgaagacagacacgaagatggatcgacagacaggccag 38211
chimp54546-135457.revcompl.Pan.troglodytes.clone.rp43-41g5.GenBank:AC099533.36      AGATGGACAAACAGGTGGGGGCTGAAGACAGACACGAAGATGGATCGACAGATAGGCCAG 40015
human.FAM30B-LOC105372935-GGT2.NCBI.                                                agatggacaaacaggtgggggctgaagacagacacgaagatggatcgacagacaggccag 36899
human.FAM230E-LOC105377182-GGT3P.NCBI.                                              agatggacaaacaggtgggggctgaagacagacacgaagatggatcgacagacaggccag 35641
human.FAM230J-LOC105372942-GGTLC5P.NCBI.rev.compl                                   agatggacaaacaggtgggggctgaagacagacacgaagatggatcgacagacaggccag 36360
human.FAM230A.ncRNA.18487127.18500594-GGTLC3.18516335.18518165.rev.compl.REF        agatggacaaacaggtgggggctgaagacagacacgaagatggatcgacagacaggccag 25107
                                                                                    **************************************************** *******

chimp.LOC112206744-LOC107973052-GGT2.revcompl                                       atagctagacaaagaggacagtaagagaaagatggtcagatagacaatggaacagagatg 38271
chimp54546-135457.revcompl.Pan.troglodytes.clone.rp43-41g5.GenBank:AC099533.36      ATAGCTAGACAAAGAGGACAGTAAGAGAAAGATGGTCAGATAGACAATGGGACAGAGATG 40075
human.FAM30B-LOC105372935-GGT2.NCBI.                                                atagctagacaaagaggacagtaagagaaagatggtcagatagacaatgggacagagatg 36959
human.FAM230E-LOC105377182-GGT3P.NCBI.                                              atagctagacaaagaggacagtaagagaaagatggtcagatagacaatgggacagagatg 35701
human.FAM230J-LOC105372942-GGTLC5P.NCBI.rev.compl                                   atagctagacaaagaggacagtaagagaaagatggtcagatagacaatgggacagagatg 36420
human.FAM230A.ncRNA.18487127.18500594-GGTLC3.18516335.18518165.rev.compl.REF        atagctagacaaagaggacagtaagagaaagatggtcagatagacaatgggacagagatg 25167
                                                                                    ************************************************** *********

chimp.LOC112206744-LOC107973052-GGT2.revcompl                                       ggcttacagatgggcggacagacagacaggtctgaacagcgggctgccagatggacagat 38331
chimp54546-135457.revcompl.Pan.troglodytes.clone.rp43-41g5.GenBank:AC099533.36      GGCTTACAGATGGGCGGACAGACAGACAGGTCTGAACAGCGGGCTGCCAGATGGACAGAT 40135
human.FAM30B-LOC105372935-GGT2.NCBI.                                                ggcttacagatgggcggacagacagacaggtctgaacagcgggctgccagatggacagat 37019
human.FAM230E-LOC105377182-GGT3P.NCBI.                                              ggcttacagatgggcggacagacagacaggtctgaacagcgggctgccagatggacagat 35761
human.FAM230J-LOC105372942-GGTLC5P.NCBI.rev.compl                                   ggcttacagatgggcggacagacagacaggtctgaacagcgggctgccagatggacagat 36480
human.FAM230A.ncRNA.18487127.18500594-GGTLC3.18516335.18518165.rev.compl.REF        ggcttacagatgggcggacagacagacaggtctgaacagcgggctgccagatggacagat 25227
                                                                                    ************************************************************

chimp.LOC112206744-LOC107973052-GGT2.revcompl                                       gggtgaatggacagatggctggcagctgtggcgagctgctgccctcaccaagtgcacact 38391
chimp54546-135457.revcompl.Pan.troglodytes.clone.rp43-41g5.GenBank:AC099533.36      GGGTGAATGGACAGATGGCTGGCAGCTGTGGCGAGCTGCTGCCCTCACCAAGTGCACACT 40195
human.FAM30B-LOC105372935-GGT2.NCBI.                                                gggtgaatggacagatggctggcagctgtggcgagctgctgccctcaccaagtgcacact 37079
human.FAM230E-LOC105377182-GGT3P.NCBI.                                              gggtgaatggacagatggctggcagctgtggcgagctgctgccctcaccaagtgcacact 35821
human.FAM230J-LOC105372942-GGTLC5P.NCBI.rev.compl                                   gggtgaatggacagatggctggcagctgtggcgagctgctgccctcaccaagtgcacact 36540
human.FAM230A.ncRNA.18487127.18500594-GGTLC3.18516335.18518165.rev.compl.REF        gggtgaatggacagatggctggcagctgtggcgagctgctgccctcaccaagtgcacact 25287
                                                                                    ************************************************************

chimp.LOC112206744-LOC107973052-GGT2.revcompl                                       acggagtggccaaactcatgcctcaacttctagttttcagctcctgcttgttcctggcag 38451
chimp54546-135457.revcompl.Pan.troglodytes.clone.rp43-41g5.GenBank:AC099533.36      ACGGAGTGGCCAAACTCACGCCTCAACTTCTAGTTTTCAGCTCCTGCTTGTTCCTGGCAG 40255
human.FAM30B-LOC105372935-GGT2.NCBI.                                                acggagtggccaaactcatgcctcaacttctagttttcagctcctgcttgttcctggcag 37139
human.FAM230E-LOC105377182-GGT3P.NCBI.                                              acggagtggccaaactcatgcctcaacttctagttttcagctcctgcttgttcctggcag 35881
human.FAM230J-LOC105372942-GGTLC5P.NCBI.rev.compl                                   acggagtggccaaactcatgcctcaacttctagttttcagctcctgcttgttcctggcag 36600
human.FAM230A.ncRNA.18487127.18500594-GGTLC3.18516335.18518165.rev.compl.REF        acggagtggccaaactcatgcctcaacttctagttttcagctcctgcttgttcctggcag 25347
                                                                                    ****************** *****************************************

chimp.LOC112206744-LOC107973052-GGT2.revcompl                                       gaggccaggcagcaaagcgtctgaggggagttttctttgcctagagaagtcagctgctgt 38511
chimp54546-135457.revcompl.Pan.troglodytes.clone.rp43-41g5.GenBank:AC099533.36      GAGGCCAGGCAGCAAAGCATCTGAGGGGAGTTTTCTTTGCCTAG---AGTCAGCTGCTGT 40312
human.FAM30B-LOC105372935-GGT2.NCBI.                                                gaggccaggcagcaaagcgtctgaggggagttttctttgcctagagaagtcagctgctgt 37199
human.FAM230E-LOC105377182-GGT3P.NCBI.                                              gaggccaggcagcaaagcgtctgaggggagttttctttgcctagagaagtcagctgctgt 35941
human.FAM230J-LOC105372942-GGTLC5P.NCBI.rev.compl                                   gaggccaggcagcaaagcgtctgaggggagttttctttgcctagagaagtcagctgctgt 36660
human.FAM230A.ncRNA.18487127.18500594-GGTLC3.18516335.18518165.rev.compl.REF        gaggccaggcagcaaagcgtctgaggggagttttctttgcctagagaagtcagctgctgt 25407
                                                                                    ****************** *************************   *************

chimp.LOC112206744-LOC107973052-GGT2.revcompl                                       gttaactccctcactgctggtaggtccaaaggccccacctaccgcccgccagagcccatg 38571
chimp54546-135457.revcompl.Pan.troglodytes.clone.rp43-41g5.GenBank:AC099533.36      GTTAACTCCCTCACTGCTGGTAGGTCCAAAGGCCCCACCTACCGCCCGCCAGAGCCCATG 40372
human.FAM30B-LOC105372935-GGT2.NCBI.                                                gttaactccctcactgctggtaggtccaaaggccccacctactgcccgccagagcccatg 37259
human.FAM230E-LOC105377182-GGT3P.NCBI.                                              gttaactccctcactgctggtaggtccaaaggcccgacctaccgcccaccagagcccatg 36001
human.FAM230J-LOC105372942-GGTLC5P.NCBI.rev.compl                                   gttaactccctcactgctggtaggtccaaaggcccgacctaccgcccaccagagcccatg 36720
human.FAM230A.ncRNA.18487127.18500594-GGTLC3.18516335.18518165.rev.compl.REF        gttaactccctcactgctggtaggtccaaaggccccacctaccgcccgccagagcccatg 25467
                                                                                    *********************************** ****** **** ************

chimp.LOC112206744-LOC107973052-GGT2.revcompl                                       gtcacactgtcgcaatgtgcaggagaacttggtgcctgctgcactgcgattgccaggtag 38631
chimp54546-135457.revcompl.Pan.troglodytes.clone.rp43-41g5.GenBank:AC099533.36      GTCACACTGTCGCAATGTGCAGGAGAACTTGGTGCCTGCTGCACTGCGGTTGCCAGGTAG 40432
human.FAM30B-LOC105372935-GGT2.NCBI.                                                gtcacactgtcgcaatgtgcaggagaacttggtgcgtgctgcactgcggttgccaggtag 37319
human.FAM230E-LOC105377182-GGT3P.NCBI.                                              gtcacactgtcgcaatgtgcaggagaacttggtgcgtgctgcactgcggttgccaggtag 36061
human.FAM230J-LOC105372942-GGTLC5P.NCBI.rev.compl                                   gtcacactgtcgcaatgtgcaggagaacttggtgcgtgctgcactgcggttgccaggtag 36780
human.FAM230A.ncRNA.18487127.18500594-GGTLC3.18516335.18518165.rev.compl.REF        gtcacactgtcgcaatgtgcaggagaacttggtgcgtgctgcactgcggttgccaggtag 25527
                                                                                    *********************************** ************ ***********

chimp.LOC112206744-LOC107973052-GGT2.revcompl                                       gggcagggctccctggaacctccacaccattccccacgttctcagcagctctgggaaagc 38691
chimp54546-135457.revcompl.Pan.troglodytes.clone.rp43-41g5.GenBank:AC099533.36      GGGCAGGGCTCCCTGGAACCTCCACACCATTCCCCAGGTTCTCAGCAGCTCTGGGAAAGC 40492
human.FAM30B-LOC105372935-GGT2.NCBI.                                                gggcagggctccctggaacctccacaccattccccaggttctcagcagctctgggaaagc 37379
human.FAM230E-LOC105377182-GGT3P.NCBI.                                              gggcagggctccctggaacctccacaccattccccaggttctcagcagctctgggaaagc 36121
human.FAM230J-LOC105372942-GGTLC5P.NCBI.rev.compl                                   gggcagggctccctggaacctccacaccattccccaggttctcagcagctctgggaaagc 36840
human.FAM230A.ncRNA.18487127.18500594-GGTLC3.18516335.18518165.rev.compl.REF        gggcagggctccctggaacctccacaccattccccaggttctcagcagctctgggaaagc 25587
                                                                                    ************************************ ***********************

chimp.LOC112206744-LOC107973052-GGT2.revcompl                                       agagctggggccgcttaactctgtcctggatccggcaaggctgccccactcccagagtgg 38751
chimp54546-135457.revcompl.Pan.troglodytes.clone.rp43-41g5.GenBank:AC099533.36      AGAGCTGGGGCCGCTTAACTCTGTCCTGGATCCGGCAAGGCTGCCCCCCTCCCAGAGTGG 40552
human.FAM30B-LOC105372935-GGT2.NCBI.                                                agagctggggccgcttaactctgccctggatccggcaaggctgcccacctcccagagtgg 37439
human.FAM230E-LOC105377182-GGT3P.NCBI.                                              agagctggggccgcttaactctgccctggatccggcaaggctgcccccctcccagagtgg 36181
human.FAM230J-LOC105372942-GGTLC5P.NCBI.rev.compl                                   agagctggggccgcttaactctgccctggatccggcaaggctgcccccctcccagagtgg 36900
human.FAM230A.ncRNA.18487127.18500594-GGTLC3.18516335.18518165.rev.compl.REF        agagctggggccgcttaactctgccctggatccggcaaggctgcccccctcccagagtgg 25647
                                                                                    *********************** **********************  ************

chimp.LOC112206744-LOC107973052-GGT2.revcompl                                       agccctgctccccagctcccatctctatcccctaaccctctccgcatggcccagcctagt 38811
chimp54546-135457.revcompl.Pan.troglodytes.clone.rp43-41g5.GenBank:AC099533.36      AGCCCTGCTCCCCAGCTCCCATCTCTATCCCCTAACCCTCTCCGCATGGCCCAGCCTAGT 40612
human.FAM30B-LOC105372935-GGT2.NCBI.                                                agccctgctccccagctcccatctctatcccctaaccctctccgcatggcccagcctagt 37499
human.FAM230E-LOC105377182-GGT3P.NCBI.                                              agccctgctccccagctcccatctctatcccctaaccctctccgcatggcccagcctagt 36241
human.FAM230J-LOC105372942-GGTLC5P.NCBI.rev.compl                                   agccctgctccccagctcccatctctatcccctaaccctctccgcatggcccagcctagt 36960
human.FAM230A.ncRNA.18487127.18500594-GGTLC3.18516335.18518165.rev.compl.REF        agccctgctccccagctcccatctctatcccctaaccctctccgcatggcccagcctagt 25707
                                                                                    ************************************************************

chimp.LOC112206744-LOC107973052-GGT2.revcompl                                       cagcatcaaggtggagctgaacagaggcagaaggaggaggacccaaggtggtgtcactca 38871
chimp54546-135457.revcompl.Pan.troglodytes.clone.rp43-41g5.GenBank:AC099533.36      CAGCATCAAGGTGGAGCTGAACAGAGGCAGAAGGAGGAGGACCCAAGGTGGTGTCACTCA 40672
human.FAM30B-LOC105372935-GGT2.NCBI.                                                cagcatcaaggtggagctgaacagaggcagaaggaggaggacccaaggtggtgtcactca 37559
human.FAM230E-LOC105377182-GGT3P.NCBI.                                              cagcatcaaggtggagctgaacagaggcagaaggaggaggacccaaggtggtgtcactca 36301
human.FAM230J-LOC105372942-GGTLC5P.NCBI.rev.compl                                   cagcatcaaggtggagctgaacagaggcagaaggaggaggacccaaggtggtgtcactca 37020
human.FAM230A.ncRNA.18487127.18500594-GGTLC3.18516335.18518165.rev.compl.REF        cagcatcaaggtggagctgaacagaggcagaaggaggaggacccaaggtggtgtcactca 25767
                                                                                    ************************************************************

chimp.LOC112206744-LOC107973052-GGT2.revcompl                                       ggacccgggttcaagtccttatgcttctgcagcctggcctgggtcccccaatcccccag 38931
chimp54546-135457.revcompl.Pan.troglodytes.clone.rp43-41g5.GenBank:AC099533.36      GGACCCGGGTTCAAGTCCTTATGCTTCTGCAGCCTGGCCTGGGTCCCCCAACCCCCCCAG 40732
human.FAM30B-LOC105372935-GGT2.NCBI.                                                ggacccgggttcaagtccttatgtttctgcagcctggcctgggtcccccaacccccccag 37619
human.FAM230E-LOC105377182-GGT3P.NCBI.                                              ggacccgggttcaagtccttatgcttctgcagcctggcctgggtcccccaactcccccag 36361
human.FAM230J-LOC105372942-GGTLC5P.NCBI.rev.compl                                   ggacccgggttcaagtccttatgcttctgcagcctggcctgggtcccccaactcccccag 37080
human.FAM230A.ncRNA.18487127.18500594-GGTLC3.18516335.18518165.rev.compl.REF        ggacccgggttcaagtccttatgcttctgcagcctggcctgggtcccccaactcccccag 25827
                                                                                    *********************** **************************  *******

chimp.LOC112206744-LOC107973052-GGT2.revcompl                                       ggtgaccaagggcttcccagtctgcacagaggacagggggtcttgacagcatcaaatgct 38991
chimp54546-135457.revcompl.Pan.troglodytes.clone.rp43-41g5.GenBank:AC099533.36      GGTGACCAAGGGCTTCCCAGTCTGCACAGAGGACAGGGGGACTTGACAGCATCAAATGCT 40792
human.FAM30B-LOC105372935-GGT2.NCBI.                                                ggtgaccaagggcttcccagtctgcacagaggacagggggtcttgacagcatcaaatgct 37679
human.FAM230E-LOC105377182-GGT3P.NCBI.                                              ggtgaccaagggcttcccagtctgcacagaggacagggggtcttgacagcatcaaatgct 36421
human.FAM230J-LOC105372942-GGTLC5P.NCBI.rev.compl                                   ggtgaccaagggcttcccagtctgcacagaggacagggggtcttgacagcatcaaatgct 37140
human.FAM230A.ncRNA.18487127.18500594-GGTLC3.18516335.18518165.rev.compl.REF        ggtgaccaagggctttccagtctgcacagaggacagggggtcttgacagcatcaaattct 25887
```

```
                                                                           *************** ************************ **************** **

chimp.LOC112206744-LOC107973052-GGT2.revcompl                              ggtgactatgagacacttacatgggaaatgcagacagaccatgcctctagcccttggtac 39051
chimp54546-135457.revcompl.Pan.troglodytes.clone.rp43-41g5.GenBank:AC099533.36 GGTGACTATGAGACACTTACGTGGGAAATGCAGACAGACCATGCCTCTAGCCCTTGGTAC 40852
human.FAM30B-LOC105372935-GGT2.NCBI.                                       ggtgactatgagacacttacatgggaaatgcagacagaccatgcctctagcccttggcac 37739
human.FAM230E-LOC105377182-GGT3P.NCBI.                                     agtgactacgagacacttacgtgggaaatgcagacagaccatgcctctagcccttggcac 36481
human.FAM230J-LOC105372942-GGTLC5P.NCBI.rev.compl                          agtgactacgagacacttacgtgggaaatgcagacagaccatgcctctagcccttggcac 37200
human.FAM230A.ncRNA.18487127.18500594-GGTLC3.18516335.18518165.rev.compl.REF ggtgactacgagacacttacgtgggaaatgcagacagaccatgcctctagcccttggcac 25947
                                                                            ******* *********** ************************************ **

chimp.LOC112206744-LOC107973052-GGT2.revcompl                              ccggcaccatccatccctgggacttgctgtcctggaaatgcagcatggacctccagggag 39111
chimp54546-135457.revcompl.Pan.troglodytes.clone.rp43-41g5.GenBank:AC099533.36 CCGGCACCATCCATCCCTGGGACTTGCTGTCCTGGAAATGCAGCATGGACCTCCAGGGAG 40912
human.FAM30B-LOC105372935-GGT2.NCBI.                                       caggcaccatccatccctgggacttgctgtcctggaaatgcagcatggacctccagggag 37799
human.FAM230E-LOC105377182-GGT3P.NCBI.                                     caggcaccatccatccctgggacttgctgtcctggaaatgcagcatggtcctccagggag 36541
human.FAM230J-LOC105372942-GGTLC5P.NCBI.rev.compl                          caggcaccatccatccctgggacttgctgtcctggaaatgcagcatggtcctccagggag 37260
human.FAM230A.ncRNA.18487127.18500594-GGTLC3.18516335.18518165.rev.compl.REF caggcaccatccatccctgggacttgctgtcctggaaatgcagcatggtcctccagggag 26007
                                                                           * ********************************************** ***********

chimp.LOC112206744-LOC107973052-GGT2.revcompl                              gggggctgtgccatgtgggggcctcaccccacctgcagctctttcccaccctggctgcag 39171
chimp54546-135457.revcompl.Pan.troglodytes.clone.rp43-41g5.GenBank:AC099533.36 GGGGGCTGTGCCATGTTGGGGCCCCACCCCACCTGCAGCTCTTTCCCACCCTGGCTGCAG 40972
human.FAM30B-LOC105372935-GGT2.NCBI.                                       gggggctgtgccatgtgggggccccaccccacctgcagctctttcccaccctggctgcag 37859
human.FAM230E-LOC105377182-GGT3P.NCBI.                                     gggggctgtgccatgtgggggccccaccccacctgcagctctttcccaccctggctgcag 36601
human.FAM230J-LOC105372942-GGTLC5P.NCBI.rev.compl                          gggggctgtgccatgtgggggccccaccccacctgcagctctttcccaccctggctgcag 37320
human.FAM230A.ncRNA.18487127.18500594-GGTLC3.18516335.18518165.rev.compl.REF gggggctgtgccatgtgggggccccaccccacctgcagctctttcccaccctggctgcag 26067
                                                                           **************** ****** ************************************

chimp.LOC112206744-LOC107973052-GGT2.revcompl                              gtctgcttccctgaatccaaatccgctactactgtgctggcagcgcagcctctctgggga 39231
chimp54546-135457.revcompl.Pan.troglodytes.clone.rp43-41g5.GenBank:AC099533.36 GTCTGCTTCCCTGAATCCAAATCCGCTACTACTGTGCTGGCAGCGCAGCCTCTCTGGGGA 41032
human.FAM30B-LOC105372935-GGT2.NCBI.                                       gtctgcttctctgaatccaaatccgctactactgtgctggcagcgcagcctctctgggga 37919
human.FAM230E-LOC105377182-GGT3P.NCBI.                                     gtctgcttccctgaatccaaatccgctactactgtgctggcagcgcagcctctctgggga 36661
human.FAM230J-LOC105372942-GGTLC5P.NCBI.rev.compl                          gtctgcttccctgaatccaaatccgctactactgtgctggcagcgcagcctctctgggga 37380
human.FAM230A.ncRNA.18487127.18500594-GGTLC3.18516335.18518165.rev.compl.REF gtctgcttccctgaatccaaatccgctactactgtgctggcagcgcagcctctctgggga 26127
                                                                           ********* **************************************************

chimp.LOC112206744-LOC107973052-GGT2.revcompl                              cactggcctggctctgttctccccaggcctcagggtgcctaaatgggaggcaaccagggg 39291
chimp54546-135457.revcompl.Pan.troglodytes.clone.rp43-41g5.GenBank:AC099533.36 CACTGGCCTGGCTCTGTTCTCCCCAGGCCTCAGGGTGCCTAAATGGGAGGCAGCCAGGGG 41092
human.FAM30B-LOC105372935-GGT2.NCBI.                                       cactggcctggctctgttctccccaggcctcagggtgcctaaatgggaggcaaccagggg 37979
human.FAM230E-LOC105377182-GGT3P.NCBI.                                     cactggcctggctctgttctccccaggcctcagggtgcctaaatgggaggcaaccagggg 36721
human.FAM230J-LOC105372942-GGTLC5P.NCBI.rev.compl                          cactggcctggctctgttctccccaggcctcagggtgcctaaatgggaggcaaccagggg 37440
human.FAM230A.ncRNA.18487127.18500594-GGTLC3.18516335.18518165.rev.compl.REF cactggcctggctctgttctccccaggcctcagggtgcctaaatgggaggcaaccagggg 26187
                                                                           **************************************************** *******

chimp.LOC112206744-LOC107973052-GGT2.revcompl                              agtgaggacccactgaggggctccgttgaccaggctcagcaggggtgcaggtgatgtggg 39351
chimp54546-135457.revcompl.Pan.troglodytes.clone.rp43-41g5.GenBank:AC099533.36 AGTGAGGACCCACTGAGGGGCTCCGTTGACCAGGCTCAGCAGGGGTGCAGGTGATGTGGG 41152
human.FAM30B-LOC105372935-GGT2.NCBI.                                       agtgaggacccactgaggggctccgttgaccaggctcagcaggggtgcaggtgatgtggg 38039
human.FAM230E-LOC105377182-GGT3P.NCBI.                                     agtgaggacccactgaggggctccgttgaccaggctcagcaggggtgcaggtgatgtggg 36781
human.FAM230J-LOC105372942-GGTLC5P.NCBI.rev.compl                          agtgaggacccactgaggggctccgttgaccaggctcagcaggggtgcaggtgatgtggg 37500
human.FAM230A.ncRNA.18487127.18500594-GGTLC3.18516335.18518165.rev.compl.REF agtgaggacccactgaggggctccgttgaccaggctcagcaggggtgcaggtgatgtggg 26247
                                                                           ************************************************************

chimp.LOC112206744-LOC107973052-GGT2.revcompl                              ctggacttcttcccacgtggcccccacagtcctccccgcttcctccccagctgaacactg 39411
chimp54546-135457.revcompl.Pan.troglodytes.clone.rp43-41g5.GenBank:AC099533.36 GTGGAATCCTTCCCACATGGCCCCCACAGTCCTCCCCGCTTCCTCCCCAGCTGAACACTG 41212
human.FAM30B-LOC105372935-GGT2.NCBI.                                       gtggaatccttcccacatggcccccacagtcctccccgcttcctccccaactgaacactg 38099
human.FAM230E-LOC105377182-GGT3P.NCBI.                                     gtggaatccttcccacatggcccccacagtcctccccgcttcctccccaactgaacactg 36841
human.FAM230J-LOC105372942-GGTLC5P.NCBI.rev.compl                          gtggaatccttcccacatggcccccacagtcctccccgcttcctccccaactgaacactg 37560
human.FAM230A.ncRNA.18487127.18500594-GGTLC3.18516335.18518165.rev.compl.REF gtggaatccttcccacatggcccccacagtcctccccgcttcctccccaactgaacactg 26307
                                                                            **** * ******** ******************************** **********

chimp.LOC112206744-LOC107973052-GGT2.revcompl                              cctgctccagatgtctacacctggagtccgggcccctccatctgggcagcagagaaactg 39471
chimp54546-135457.revcompl.Pan.troglodytes.clone.rp43-41g5.GenBank:AC099533.36 CCTGCTCCAGATGTCTACACCTGGAGTCTGGGCCCCTCCATCTGGGCAGCAGAGAAACTG 41272
human.FAM30B-LOC105372935-GGT2.NCBI.                                       cctgctccagatgtgtacacctggagtctgggcccctccatctgggcagcagagaaactg 38159
human.FAM230E-LOC105377182-GGT3P.NCBI.                                     cctgctccagatgtctacacctggagtctgggcccctccatctgggcagcagagaaactg 36901
human.FAM230J-LOC105372942-GGTLC5P.NCBI.rev.compl                          cctgctccagatgtctacacctggagtctgggcccctccatctgggcagcagagaaactg 37620
human.FAM230A.ncRNA.18487127.18500594-GGTLC3.18516335.18518165.rev.compl.REF cctgctccagatgtgtacacctggagtctgggcccctccatctgggcagcagagaaactg 26367
                                                                           ************** ************* *******************************

chimp.LOC112206744-LOC107973052-GGT2.revcompl                              aggcacagagacagactgtgtccttacaggccacacagcctgccaggcccctatgtccgg 39531
chimp54546-135457.revcompl.Pan.troglodytes.clone.rp43-41g5.GenBank:AC099533.36 AGGCACAGAGACAGACTGTGTCCTTACAGGCCACACAGCCTGCCAGGCCCCTATGTCCGG 41332
human.FAM30B-LOC105372935-GGT2.NCBI.                                       aggcacagagacagactgtgtccttacagggcacacagcctgccaggcccctatgtccgg 38219
human.FAM230E-LOC105377182-GGT3P.NCBI.                                     aggcacagagacagactgtgtccttacagggcacacagcctgccaggcccctatgtccgg 36961
human.FAM230J-LOC105372942-GGTLC5P.NCBI.rev.compl                          aggcacagagacagactgtgtccttacagggcacacagcctgccaggcccctatgtccgg 37680
human.FAM230A.ncRNA.18487127.18500594-GGTLC3.18516335.18518165.rev.compl.REF aggcacagagacagactgtgtccttacagggcacacagcctgccaggcccctatgtccgg 26427
                                                                           ****************************** *****************************

chimp.LOC112206744-LOC107973052-GGT2.revcompl                              ccagagcccctggtcagcctgggctgcagtgattgtttagaggtaggctgttcccacggc 39591
chimp54546-135457.revcompl.Pan.troglodytes.clone.rp43-41g5.GenBank:AC099533.36 CCAGAGCCCCTGGTCAGCCTGGGCTGCAGTGATTGTTTAGAGGTAGGCTGTTCCCACGGC 41392
human.FAM30B-LOC105372935-GGT2.NCBI.                                       ccagagcccctggtcagcctgggctgcagtgattgtttagaggtaggctgttcccacggc 38279
human.FAM230E-LOC105377182-GGT3P.NCBI.                                     ccagagcccctggccagcctgggctgcagtgattgtttagaggtaggctgttcccacggc 37021
human.FAM230J-LOC105372942-GGTLC5P.NCBI.rev.compl                          ccagagcccctggccagcctgggctgcagtgattgtttagaggtaggctgttcccacggc 37740
human.FAM230A.ncRNA.18487127.18500594-GGTLC3.18516335.18518165.rev.compl.REF ccagagcccctggtcagcctgggctgcagtgattgtttagaggtaggctgttcccacggc 26487
                                                                           ************* **********************************************

chimp.LOC112206744-LOC107973052-GGT2.revcompl                              tgcctctcacggtagggggggcctgcggacgcctcctcccgtccccacccgactcccaagc 39651
chimp54546-135457.revcompl.Pan.troglodytes.clone.rp43-41g5.GenBank:AC099533.36 TGCCTCTCACGGTAGGGGGGCCTGTGGACGCCTCCTCCCACCCCCACCCGACTCCCAAGC 41452
human.FAM30B-LOC105372935-GGT2.NCBI.                                       tgcctctcacggtaggggggcctgcggacgcctcctcccgcccccacccgactcccaagc 38339
human.FAM230E-LOC105377182-GGT3P.NCBI.                                     tgcctctcacggtaggggggcctgcggacgcctcctcccgcccccacccgactcccaagc 37081
human.FAM230J-LOC105372942-GGTLC5P.NCBI.rev.compl                          tgcctctcacggtaggggggcctgcggacgcctcctcccgcccccacccgactcccaagc 37800
human.FAM230A.ncRNA.18487127.18500594-GGTLC3.18516335.18518165.rev.compl.REF tgcctctcacggtaggggggcctgtggacgcctcctcccgcccccacccgactcccaagc 26547
                                                                           ************************ **************  *******************

chimp.LOC112206744-LOC107973052-GGT2.revcompl                              ctcagtgacattgctcaaccaggagctgaagtgcattcctgggctcaggccagcccaccc 39711
chimp54546-135457.revcompl.Pan.troglodytes.clone.rp43-41g5.GenBank:AC099533.36 CTCAGTGACATTGCTCAACCAGGAGCTGAAGTGCATTCCTGGGCTCAGGCCAGCCCACCC 41512
human.FAM30B-LOC105372935-GGT2.NCBI.                                       ctcagtgacattgctcaaccaggaactgaagtgcattcctgggctcaggccagcccaccc 38399
human.FAM230E-LOC105377182-GGT3P.NCBI.                                     ctcagtgacattgctcaaccaggagctgaagtgcattcctgggctcaggccagcccaccc 37141
human.FAM230J-LOC105372942-GGTLC5P.NCBI.rev.compl                          ctcagtgacattgctcaaccaggagctgaagtgcattcctgggctcaggccagcccaccc 37860
human.FAM230A.ncRNA.18487127.18500594-GGTLC3.18516335.18518165.rev.compl.REF ctcagtgacattgctcaaccaggagctgaagtgcattcctgggctccggccagcccaccc 26607
                                                                           ************************ ********************* *************

chimp.LOC112206744-LOC107973052-GGT2.revcompl                              acccacccgctgcagtcctggaagcccagaggcctgggcagcaggaacagtggagacagc 39771
chimp54546-135457.revcompl.Pan.troglodytes.clone.rp43-41g5.GenBank:AC099533.36 ACCCACCCGCTGCAGTCCTGGAAGCTCAGAGGCCTGGGCAGCAGGAACAGTGGAGACAGC 41572
human.FAM30B-LOC105372935-GGT2.NCBI.                                       acccacccgctgcagtcctggaagcccagaggcctgggcagcaggaacagtggagacagc 38459
human.FAM230E-LOC105377182-GGT3P.NCBI.                                     accgacccgctgcagtcctggaagcccagaggcctgggcagcaggaacagtggagacagc 37201
human.FAM230J-LOC105372942-GGTLC5P.NCBI.rev.compl                          accgacccgctgcagtcctggaagcccagaggcctgggcagcaggaacagtggagacagc 37920
human.FAM230A.ncRNA.18487127.18500594-GGTLC3.18516335.18518165.rev.compl.REF acccacccgctgcagtcctggaagcccagaggcctgggcagcaggaacagtggagacagc 26667
                                                                           *** ********************* **********************************

chimp.LOC112206744-LOC107973052-GGT2.revcompl                              agtgtgggggacgtccccccctcctctcccaccatcctcgtcaggcagaggccagggtgc 39831
chimp54546-135457.revcompl.Pan.troglodytes.clone.rp43-41g5.GenBank:AC099533.36 AGTGTGGGGGACATCCCCCCCTCCTCTCCCACCATCCTCGTCAGGCAGAGGCCAGGGTGC 41632
human.FAM30B-LOC105372935-GGT2.NCBI.                                       agtgtgggggacgtccccccctcctctcccaccatcctcgtcaggcagaggccagggtgc 38519
human.FAM230E-LOC105377182-GGT3P.NCBI.                                     agtgtgggggacgt-cccccctcctctcccaccatcctcgtcaggcagaggccagggtgc 37260
human.FAM230J-LOC105372942-GGTLC5P.NCBI.rev.compl                          agtgtgggggacgt-cccccctcctctcccaccatcctcgtcaggcagaggccagggtgc 37979
human.FAM230A.ncRNA.18487127.18500594-GGTLC3.18516335.18518165.rev.compl.REF agtgtgggggacgt-cccccctcctctcccaccatcctcgtcaggcagaggccagggtgc 26726
                                                                           ************ * *********************************************

chimp.LOC112206744-LOC107973052-GGT2.revcompl                              agggaccacccgagcaaaggcccagggaaatgaatgggtgtcattttggtcctgacccga 39891
chimp54546-135457.revcompl.Pan.troglodytes.clone.rp43-41g5.GenBank:AC099533.36 AGGGACCACCCGAGCAAAGGCCCAGGGAAATGAATGGGTGTCATTCTGGTCCTGACCCGA 41692
human.FAM30B-LOC105372935-GGT2.NCBI.                                       agggaccaccggagcaaaggcccagggaaatgaatgggtgtcattctggtcctgacccga 38579
human.FAM230E-LOC105377182-GGT3P.NCBI.                                     agggaccgcccgagcaaaggcccagggaaatgaatgggtgtcattctggtcctgacccga 37320
human.FAM230J-LOC105372942-GGTLC5P.NCBI.rev.compl                          agggaccgcccgagcaaaggcccagggaaatgaatgggtgtcattctggtcctgacccga 38039
human.FAM230A.ncRNA.18487127.18500594-GGTLC3.18516335.18518165.rev.compl.REF agggaccaccggagcaaaggcccagggaaatgaatgggtgtcattctggtcctgacccga 26786
                                                                           ******* ** ********************************* ***************

chimp.LOC112206744-LOC107973052-GGT2.revcompl                              ggcacagccaggaaggtccctgtggggaaaagaaacagatatcagactgttactgtgtct 39951
chimp54546-135457.revcompl.Pan.troglodytes.clone.rp43-41g5.GenBank:AC099533.36 GGCACAGCCAGGAAGGTCCCTGTGGGGAAAAGAAAGAGATATCAGACTGTTACTGTGTCT 41752
human.FAM30B-LOC105372935-GGT2.NCBI.                                       ggcacagccaggaaggtccctgtggggaaaagaaagagatatcagactgttactgtgtct 38639
human.FAM230E-LOC105377182-GGT3P.NCBI.                                     ggcacagccaggaaggtccctgtggggaaaagaaagagatatcagactgttactgtgtct 37380
human.FAM230J-LOC105372942-GGTLC5P.NCBI.rev.compl                          ggcacagccaggaaggtccctgtggggaaaagaaagagatatcagactgttactgtgtct 38099
human.FAM230A.ncRNA.18487127.18500594-GGTLC3.18516335.18518165.rev.compl.REF ggcacagccaggaaggtccctgtggggaaaagaaagagatatcagactgttactgtgtct 26846
                                                                           ************************************ ***********************

chimp.LOC112206744-LOC107973052-GGT2.revcompl                              atgtagaaagaagtagaggtaagaggctccattttgttgtgtagtaagaaaaattctttt 40011
chimp54546-135457.revcompl.Pan.troglodytes.clone.rp43-41g5.GenBank:AC099533.36 ATGTAGAAAGAAGTAGACGTAAGAGGCTCCATTTTGTTCTGTACTAAGAAAAATTCTTTT 41812
human.FAM30B-LOC105372935-GGT2.NCBI.                                       atgtagaaagaagtagacgtaagaggctccattttgttgtgtagtaagaaaaattctttt 38699
human.FAM230E-LOC105377182-GGT3P.NCBI.                                     atgtagaaagaagtagacgtaagaggctccattttgttgtgtagtaagaaaaattctttt 37440
human.FAM230J-LOC105372942-GGTLC5P.NCBI.rev.compl                          atgtagaaagaagtagacgtaagaggctccattttgttgtgtagtaagaaaaattctttt 38159
human.FAM230A.ncRNA.18487127.18500594-GGTLC3.18516335.18518165.rev.compl.REF atgtagaaagaagtagacgtaagaggctccattttgctgtgtagtaagaaaaattctttt 26906
                                                                           ***************** ****************** * **** ****************

chimp.LOC112206744-LOC107973052-GGT2.revcompl                              gccttgagatactgttaatctgtaaccctagccccaaccctgtgctcacagaaacatgta 40071
chimp54546-135457.revcompl.Pan.troglodytes.clone.rp43-41g5.GenBank:AC099533.36 GCCTTGAGATGCTGTTAATCTGTAACCCTAGCCCCAACCCTGTGCTCACAGAAACATGTA 41872
human.FAM30B-LOC105372935-GGT2.NCBI.                                       gccttgagatgccgttaatctgtaaccctagccccaaccctgtgctcacagaaacatgta 38759
human.FAM230E-LOC105377182-GGT3P.NCBI.                                     gccttgagatgccgttaatctgtaaccctagccccaaccctgtgctcacagaaacatgta 37500
human.FAM230J-LOC105372942-GGTLC5P.NCBI.rev.compl                          gccttgagatgccgttaatctgtaaccctagccccaaccctgtgctcacagaaacatgta 38219
human.FAM230A.ncRNA.18487127.18500594-GGTLC3.18516335.18518165.rev.compl.REF gccttgagatgccgttaatctgtaaccctagccccaaccctgtgctcacagaaacatgta 26966
                                                                           ********** * ***********************************************
```

```
chimp.LOC112206744-LOC107973052-GGT2.revcompl                                      ctgtgtcgactcaaggtttaatggatttagggctgtgcaggatgtgctttgttaaacaaa 40131
chimp54546-135457.revcompl.Pan.troglodytes.clone.rp43-41g5.GenBank:AC099533.36     CTGTGTCGACTCAAGGTTTAATGGATTTAGGGCTGTGCAGGATGTGCTTTGTTAAACAAA 41932
human.FAM30B-LOC105372935-GGT2.NCBI.                                               ctgtgtcgactcaaggtttaatggattcagggctgtgcaggatgtgctttgttaaacaaa 38819
human.FAM230E-LOC105377182-GGT3P.NCBI.                                             ctgtgtcgactcaaggtttaatggattcagggctgtgcaggatgtgctttgttaaacaaa 37560
human.FAM230J-LOC105372942-GGTLC5P.NCBI.rev.compl                                  ctgtgtcgactcaaggtttaatggattcagggctgtgcaggatgtgctttgttaaacaaa 38279
human.FAM230A.ncRNA.18487127.18500594-GGTLC3.18516335.18518165.rev.compl.REF       ctgtgtcgactcaaggtttaatggattcagggctgtgcaggatgtgctttgttaaacaaa 27026
                                                                                   *************************** ********************************

chimp.LOC112206744-LOC107973052-GGT2.revcompl                                      tgcttgaaggcagcatgcttgttaaaagtcatcaccactccctaatctcaagtaagcagg 40191
chimp54546-135457.revcompl.Pan.troglodytes.clone.rp43-41g5.GenBank:AC099533.36     TGCTTGAAGGCAGCATGCTTGTTAAAAGTCATCACCACTCCCTAATCTCAAGTAAGCAGG 41992
human.FAM30B-LOC105372935-GGT2.NCBI.                                               tgcttgaaggcagcatgcttgttaagagtcatcaccactccctaatctcaagtaagcagg 38879
human.FAM230E-LOC105377182-GGT3P.NCBI.                                             tgcttgaaggcagcatgcttgttaagagtcatcaccactccctaatctcaagtaagcagg 37620
human.FAM230J-LOC105372942-GGTLC5P.NCBI.rev.compl                                  tgcttgaaggcagcatgcttgttaagagtcatcaccactccctaatctcaagtaagcagg 38339
human.FAM230A.ncRNA.18487127.18500594-GGTLC3.18516335.18518165.rev.compl.REF       tgcttgaaggcagcatgcttgttaagagtcatcaccactccctaatctcaagtaagcagg 27086
                                                                                   ************************* **********************************

chimp.LOC112206744-LOC107973052-GGT2.revcompl                                      gacacaaaacactgcagaaggccgcagggacctctgcctaggaaagccaggtattgtcca 40251
chimp54546-135457.revcompl.Pan.troglodytes.clone.rp43-41g5.GenBank:AC099533.36     GACACAAAACACTGCAGAAGGCCGCAGGGACCTCTGCCTAGGAAAGCCAGGTATTGTCCA 42052
human.FAM30B-LOC105372935-GGT2.NCBI.                                               gacacaaaacactgcagaaggccgcagggacctctgcctaggaaagccaggtatcgtcca 38939
human.FAM230E-LOC105377182-GGT3P.NCBI.                                             gacacaaaacactgcagaaggccgcagggacctctgcctaggaaagccaggtattgtcca 37680
human.FAM230J-LOC105372942-GGTLC5P.NCBI.rev.compl                                  gacacaaaacactgcagaaggccgcagggacctctgcctaggaaagccaggtattgtcca 38399
human.FAM230A.ncRNA.18487127.18500594-GGTLC3.18516335.18518165.rev.compl.REF       gacacaaaacactgcagaaggccgcagggacctctgcctaggaaagccaggtattgtcca 27146
                                                                                   ****************************************************** *****

chimp.LOC112206744-LOC107973052-GGT2.revcompl                                      aggtttctccccatgtgacagtctgaaatatggcctcgtgggaagggaaagacctgaccg 40311
chimp54546-135457.revcompl.Pan.troglodytes.clone.rp43-41g5.GenBank:AC099533.36     AGGTTTCTCCCCATGTGACAGTCTGAAATATGGCCTCGTGGGAAGGGAAAGACCTGACCG 42112
human.FAM30B-LOC105372935-GGT2.NCBI.                                               aggtttctccccaggtgacagtctgaaatatggcctcgtgggaagggaaagacctgaccg 38999
human.FAM230E-LOC105377182-GGT3P.NCBI.                                             aggtttctccccaggtgacagtctgaaatatggcctcgtgggaagggaaagacctgaccg 37740
human.FAM230J-LOC105372942-GGTLC5P.NCBI.rev.compl                                  aggtttctccccaggtgacagtctgaaatatggcctcgtgggaagggaaagacctgaccg 38459
human.FAM230A.ncRNA.18487127.18500594-GGTLC3.18516335.18518165.rev.compl.REF       aggtttctccccaggtgacagtctgaaatatggcctcgtgggaagggaaagacctgaccg 27206
                                                                                   ************* **********************************************

chimp.LOC112206744-LOC107973052-GGT2.revcompl                                      tcccccagcccgacacctgtaaagggtctgtgctgaggaggattagtaaaag----ggga 40367
chimp54546-135457.revcompl.Pan.troglodytes.clone.rp43-41g5.GenBank:AC099533.36     TCCCCCAGCCCGACACCCGTAAAGGGTCTGTGCTGAGGAGGATTAGTATAAGAGGAAGGA 42172
human.FAM30B-LOC105372935-GGT2.NCBI.                                               tcccccagcccgacacctgtaaagggtctgtgctgaggaggattagtaaaag----agga 39055
human.FAM230E-LOC105377182-GGT3P.NCBI.                                             tcccccagcccgacacctgtaaagggtctgtgctgaggaggattagtaaaag----agga 37796
human.FAM230J-LOC105372942-GGTLC5P.NCBI.rev.compl                                  tcccccagcccgacacctgtaaagggtctgtgctgaggaggattagtaaaag----agga 38515
human.FAM230A.ncRNA.18487127.18500594-GGTLC3.18516335.18518165.rev.compl.REF       tcccccagcccgacacctgtaaagggtctgtgctgaggaggattagtaaaag----agga 27262
                                                                                   ***************** ****************************** ***     ***

chimp.LOC112206744-LOC107973052-GGT2.revcompl                                      aggcctctttgcagttgagataagaggaaggcatctctctcctgatcgtccctgggcaaa 40427
chimp54546-135457.revcompl.Pan.troglodytes.clone.rp43-41g5.GenBank:AC099533.36     AGGCCTCTTTGCAGTTGAGATAAGAGGAAGGCATCTGTCTCCTGCCCGTCCCTGGGCAAA 42232
human.FAM30B-LOC105372935-GGT2.NCBI.                                               aggcctctttgcagttgagataagaggaaggcatctctctcctgatcgtccctgggcaaa 39115
human.FAM230E-LOC105377182-GGT3P.NCBI.                                             aggcctctttgcagttgagataagaggaaggcatctctctcctgatcgtccctgggcaaa 37856
human.FAM230J-LOC105372942-GGTLC5P.NCBI.rev.compl                                  aggcctctttgcagttgagataagaggaaggcatctctctcctgatcgtccctgggcaaa 38575
human.FAM230A.ncRNA.18487127.18500594-GGTLC3.18516335.18518165.rev.compl.REF       aggcctctttgcagttgagataagaggaaggcatctctctcctgatcgtccctgggcaaa 27322
                                                                                   ************************************ *******  **************

chimp.LOC112206744-LOC107973052-GGT2.revcompl                                      ggaatgtctcagtgttgatggtatattccatctactgagataggagaaaactgccttagg 40487
chimp54546-135457.revcompl.Pan.troglodytes.clone.rp43-41g5.GenBank:AC099533.36     GGAATGTGTAAAACCCAATTGTATATTCCATCTACTGAGATAGGAGAAAACTGCCTTAGG 42292
human.FAM30B-LOC105372935-GGT2.NCBI.                                               ggaatgtctcagtgttgattgtatattccatctgctgagataggagaaaactgccttagg 39175
human.FAM230E-LOC105377182-GGT3P.NCBI.                                             ggaatgtctcagtgttgattgtatattccatctgctgagataggagaaaactgccttagg 37916
human.FAM230J-LOC105372942-GGTLC5P.NCBI.rev.compl                                  ggaatgtctcagtgttgattgtatattccatctgctgagataggagaaaactgccttagg 38635
human.FAM230A.ncRNA.18487127.18500594-GGTLC3.18516335.18518165.rev.compl.REF       ggaatgtctcagtgttgattgtatattccatctgctgagataggagaaaactgccttagg 27382
                                                                                   ******* * *      ** ************* **************************

chimp.LOC112206744-LOC107973052-GGT2.revcompl                                      gctggaggtgggacatgctggtggcaatactgctatttaatgcattgagatgtttatgta 40547
chimp54546-135457.revcompl.Pan.troglodytes.clone.rp43-41g5.GenBank:AC099533.36     GCTGGAGGTGGGACATGCTGGTGACAATACTGCTCTTTAATGCATTGAGATGTTTATGTA 42352
human.FAM30B-LOC105372935-GGT2.NCBI.                                               gctggagatgggacatgctggtggcaatactgctctttaatgcattgagatgtttatgta 39235
human.FAM230E-LOC105377182-GGT3P.NCBI.                                             gctggaggtgggacatgctggtggcaatactgctctttaatgcattgagatgtttatgta 37976
human.FAM230J-LOC105372942-GGTLC5P.NCBI.rev.compl                                  gctggaggtgggacatgctggtggcaatactgctctttaatgcattgagatgtttatgta 38695
human.FAM230A.ncRNA.18487127.18500594-GGTLC3.18516335.18518165.rev.compl.REF       gctggaggtgggacatgctggtggcaatactgctctttaatgcattgagatgtttatgta 27442
                                                                                   ******* *************** ********** *************************

chimp.LOC112206744-LOC107973052-GGT2.revcompl                                      tatgcacatcaaagcacagcacctttttcttaaccttgtttatgacacagagacatttgt 40607
chimp54546-135457.revcompl.Pan.troglodytes.clone.rp43-41g5.GenBank:AC099533.36     TATGCACATCAAAGCACAGCACCTTTTTCTTAACCTTGTTTATGACACAGAGACATTTGT 42412
human.FAM30B-LOC105372935-GGT2.NCBI.                                               tatgcacatcaaagcatagcacctttttcttaaccttgtttatgacacagagacatttgt 39295
human.FAM230E-LOC105377182-GGT3P.NCBI.                                             tatgcacatcaaagcacagcacctttttcttaaccttgtttatgacacagagacatttgt 38036
human.FAM230J-LOC105372942-GGTLC5P.NCBI.rev.compl                                  tatgcacatcaaagcacagcacctttttcttaaccttgtttatgacacagagacatttgt 38755
human.FAM230A.ncRNA.18487127.18500594-GGTLC3.18516335.18518165.rev.compl.REF       tatgcacatcaaagcacagcacctttttcttaaccttgtttatgacacagagacatttgt 27502
                                                                                   **************** *******************************************

chimp.LOC112206744-LOC107973052-GGT2.revcompl                                      tcacgtgttttcctgctgaccctctccccactattaccctattgtcctgccacatccccc 40667
chimp54546-135457.revcompl.Pan.troglodytes.clone.rp43-41g5.GenBank:AC099533.36     TCACGTGTTTTCCTGCTGACCCTCACCCCACTATTACCATATCGTCCTGCCACATCCCCC 42472
human.FAM30B-LOC105372935-GGT2.NCBI.                                               tcacgtgttttcctgctgaccctctccccactattaccctattgtcctgccacatccccc 39355
human.FAM230E-LOC105377182-GGT3P.NCBI.                                             tcacgtgttttcctgctgaccctctccccactattaccctattgtcctgccacatccccc 38096
human.FAM230J-LOC105372942-GGTLC5P.NCBI.rev.compl                                  tcacgtgttttcctgctgaccctctccccactattaccctattgtcctgccacatccccc 38815
human.FAM230A.ncRNA.18487127.18500594-GGTLC3.18516335.18518165.rev.compl.REF       tcacgtgttttcctgctgaccctctccccactattaccctattgtcctgccacatccccc 27562
                                                                                   ************************ ************* *** *****************

chimp.LOC112206744-LOC107973052-GGT2.revcompl                                      tctctgagatggtagagataatgatcaataaatactagggaactcagagactggtgccag 40727
chimp54546-135457.revcompl.Pan.troglodytes.clone.rp43-41g5.GenBank:AC099533.36     TCTCTGAGATGGTAGAGATAATGATCAATAAATACTAGGGAACTCAGAGACTGGTGCCAG 42532
human.FAM30B-LOC105372935-GGT2.NCBI.                                               tctctgagatggtagagataatgatcaataaatactagggaactcagagactggtgccag 39415
human.FAM230E-LOC105377182-GGT3P.NCBI.                                             tctctgagatggtagagataatgatcaataaatactagggaactcagagactggtgccag 38156
human.FAM230J-LOC105372942-GGTLC5P.NCBI.rev.compl                                  tctctgagatggtagagataatgatcaataaatactagggaactcagagactggtgccag 38875
human.FAM230A.ncRNA.18487127.18500594-GGTLC3.18516335.18518165.rev.compl.REF       tctctgagatggtagagataatgatcaataaatactagggaactcagagactggtgccag 27622
                                                                                   ************************************************************

chimp.LOC112206744-LOC107973052-GGT2.revcompl                                      cgtgggtcctccatatgctgagtgcaggtcccctgggcccactttctttctctatactt 40787
chimp54546-135457.revcompl.Pan.troglodytes.clone.rp43-41g5.GenBank:AC099533.36     CGTGGGTCCTCCATATGCTGAGTGCAGGTCCCCTGGGCCCACTTTTCTTTCTCTATACTT 42592
human.FAM30B-LOC105372935-GGT2.NCBI.                                               cgtggggcctccgtatgctgagcgcaggtcccctgggcccacttttctttctctatactt 39475
human.FAM230E-LOC105377182-GGT3P.NCBI.                                             cgtggggcctccatatgctgagtgcaggtcccctgggcccacttttctttctctatactt 38216
human.FAM230J-LOC105372942-GGTLC5P.NCBI.rev.compl                                  cgtggggcctccgtatgctgagtgcaggtcccctgggcccacttttctttctctatactt 38935
human.FAM230A.ncRNA.18487127.18500594-GGTLC3.18516335.18518165.rev.compl.REF       cgtggggcctccatatgctgagtgcaggtcccctgggcccacttttctttctctatactt 27682
                                                                                   ****** ***** ********* *************************************

chimp.LOC112206744-LOC107973052-GGT2.revcompl                                      tgtctctgtgtctctttcttttctcagtctcttgtcccacctgatgagaaacacccacag 40847
chimp54546-135457.revcompl.Pan.troglodytes.clone.rp43-41g5.GenBank:AC099533.36     TGTCTCTGTGTCTCTTTCTTTTCTCAGTCTCTTGTCCCACCTGATGAGAAACACCCACAG 42652
human.FAM30B-LOC105372935-GGT2.NCBI.                                               tgtctctgtgtctctttcttttctcagtctcttgtcccagctgatgggaaacacccacag 39535
human.FAM230E-LOC105377182-GGT3P.NCBI.                                             tggctctgtggctctttcttttctcagtctcttgtcccagctgatgagaaacacccacag 38276
human.FAM230J-LOC105372942-GGTLC5P.NCBI.rev.compl                                  tgtctctgtgtctctttcttttctcagtctcttgtcccagctgatgagaaacacccacag 38995
human.FAM230A.ncRNA.18487127.18500594-GGTLC3.18516335.18518165.rev.compl.REF       tgtctctgtgtctctttcttttctcagtctcttgtcccagctgatgagaaacacccacag 27742
                                                                                   ** ******* **************************** ****** *************

chimp.LOC112206744-LOC107973052-GGT2.revcompl                                      atgtggaggggcaggccaccccttcaggtccctgaatgtccttcctcaggaaatgatggg 40907
chimp54546-135457.revcompl.Pan.troglodytes.clone.rp43-41g5.GenBank:AC099533.36     ATGTGGAGGGGCAGGCCGCTCCTTCAGGTCCCTGAATGTCCTTCCTCAGGAAATGATGGG 42712
human.FAM30B-LOC105372935-GGT2.NCBI.                                               gtgtggaggggcaggccaccccttcaggtccctgaatgtccttcctcaggaaatgatggg 39595
human.FAM230E-LOC105377182-GGT3P.NCBI.                                             gtgtggaggggcaggccaccccttcaggtccctgaatgtccttcctcaggaaatgatggg 38336
human.FAM230J-LOC105372942-GGTLC5P.NCBI.rev.compl                                  gtggggaggggcaggccaccccttcaggtccctgaatgtccttcctcaggaaatgatggg 39055
human.FAM230A.ncRNA.18487127.18500594-GGTLC3.18516335.18518165.rev.compl.REF       gtgtggaggggcaggccaccccttcaggtccctgaatgtccttcctcaggaaatgatggg 27802
                                                                                    ** ************* * ****************************************

chimp.LOC112206744-LOC107973052-GGT2.revcompl                                      ggaaggggtgatgagaatgaaggagaggatttaagtccctcacccccgaggtagtcctg 40967
chimp54546-135457.revcompl.Pan.troglodytes.clone.rp43-41g5.GenBank:AC099533.36     GGAAGGGGTGATGAGAATGAAGGAGAGGATTTAAGTCCCTCACCCCCCGAGGTAGTCCTG 42772
human.FAM30B-LOC105372935-GGT2.NCBI.                                               ggaaggggtgatgagaatgaaggagacgatttaagtccctcaccccccgaggtagtcctg 39655
human.FAM230E-LOC105377182-GGT3P.NCBI.                                             ggaaggggcgatgagaatgaaggagacgatttaagtccctcaccccccgaggtagtcctg 38396
human.FAM230J-LOC105372942-GGTLC5P.NCBI.rev.compl                                  ggaaggggcgatgagaatgaaggagacgatttaagtccctcaccccccgaggtagtcctg 39115
human.FAM230A.ncRNA.18487127.18500594-GGTLC3.18516335.18518165.rev.compl.REF       ggaaggggcgatgagaatgaaggagacgatttaagtccctcaccccccgaggtagtcctg 27862
                                                                                   ******** ***************** *********************************

chimp.LOC112206744-LOC107973052-GGT2.revcompl                                      ggctgagccccatgggacctggagaaccagggtgtaccccaccagcgtgtcgggtccagg 41027
chimp54546-135457.revcompl.Pan.troglodytes.clone.rp43-41g5.GenBank:AC099533.36     GGCTGAGCCCCATGGGACCTGGAGAACCAGGGTGTACCCCACCAGTGTGTCGAGTCGAGG 42832
human.FAM30B-LOC105372935-GGT2.NCBI.                                               ggctgagccccatgggacctagagaaccagggtgtaccccaccagcgtgtcgggtccagg 39715
human.FAM230E-LOC105377182-GGT3P.NCBI.                                             ggctgagccccatgggacctagagaaccagggtgtaccccaccagcatgtcgggtccagg 38456
human.FAM230J-LOC105372942-GGTLC5P.NCBI.rev.compl                                  ggctgagccccatgggacctagagaaccagggtgtaccccaccagcatttcgggtccagg 39175
human.FAM230A.ncRNA.18487127.18500594-GGTLC3.18516335.18518165.rev.compl.REF       ggctgagccccatgggacctagagaaccagggtgtaccccaccagcatgtcgggtccagg 27922
                                                                                   ******************** ************************  * *** *** ***

chimp.LOC112206744-LOC107973052-GGT2.revcompl                                      aagcctcgtggccagctcccacttctcttgctgctgtgcaacccagagcaaggcctgccc 41087
chimp54546-135457.revcompl.Pan.troglodytes.clone.rp43-41g5.GenBank:AC099533.36     AAGCCTCGTGGCCAGCTCCCACTTCTCTTGCTGCTGTGCAACCCAGAGCAAGGCCTGCCC 42892
human.FAM30B-LOC105372935-GGT2.NCBI.                                               aagcctcgtggccagctcccacttctcttcctgctgtgcaacccagagcaaggcctgccc 39775
human.FAM230E-LOC105377182-GGT3P.NCBI.                                             aagcctcgtggccagctcccacttctcttcctgctgtgcaacccagagcaaggcctgccc 38516
human.FAM230J-LOC105372942-GGTLC5P.NCBI.rev.compl                                  aagcctcgtggccagctcccacttctcttcctgctgtgcaacccagagcaaggcctgccc 39235
human.FAM230A.ncRNA.18487127.18500594-GGTLC3.18516335.18518165.rev.compl.REF       aagcctcgtggccagctcccacttctcttcctgctgtgcaacccagagcaaggcctgccc 27982
                                                                                   ***************************** ******************************

chimp.LOC112206744-LOC107973052-GGT2.revcompl                                      ctccagcttcagtcttctcccctgcaaatggggccacggcctttcctctcaggccaaaat 41147
chimp54546-135457.revcompl.Pan.troglodytes.clone.rp43-41g5.GenBank:AC099533.36     CTCCAGCTTCAGTCTTCTCCCCTGCAAATGGGGCCACGGCCTTTCCTCTCAGGCCAAAAT 42952
human.FAM30B-LOC105372935-GGT2.NCBI.                                               ctccagcttcagtcttctcccctgcaaatggggccacggcctttcctctcaggccaatat 39835
human.FAM230E-LOC105377182-GGT3P.NCBI.                                             ctccagcttcagtcttctcccctgcaaatggggccacggcctttcctctcaggccaatat 38576
human.FAM230J-LOC105372942-GGTLC5P.NCBI.rev.compl                                  ctccagcttcagtcttctcccctgcaaatggggccacggcctttcctctcaggccaatat 39295
human.FAM230A.ncRNA.18487127.18500594-GGTLC3.18516335.18518165.rev.compl.REF       ctccagcttcagtcttctcccctgcaaatggggccacggcctttcctctcaggccaatat 28042
                                                                                   ********************************************************* **
```

```
chimp.LOC112206744-LOC107973052-GGT2.revcompl                                    aaggattgaggccgggtgcagtggctcacccctgtaatcctagcactttgggagactgag 41207
chimp54546-135457.revcompl.Pan.troglodytes.clone.rp43-41g5.GenBank:AC099533.36   AAGGATTGAGGCCGGGTGCAGTGGCTCACCCCTGTAATTTTAGCACTTTGGGAGACTGAG 43012
human.FAM30B-LOC105372935-GGT2.NCBI.                                             aaggattgaggccgggtgcagtggct-acccctgtaatcctagcactttgggagactgag 39894
human.FAM230E-LOC105377182-GGT3P.NCBI.                                           aaggattgaggccgggtgcagtggct-acccctgtaatcctagcactttgggagactgag 38635
human.FAM230J-LOC105372942-GGTLC5P.NCBI.rev.compl                                aaggattgaggccgggtgcagtggct-acccctgtaatcctagcactttgggagactgag 39354
human.FAM230A.ncRNA.18487127.18500594-GGTLC3.18516335.18518165.rev.compl.REF     aaggattgaggccgggtgcagtggct-acccctgtaatcctagcactttgggagactgag 28101
                                                                                 ************************* ***********  *********************

chimp.LOC112206744-LOC107973052-GGT2.revcompl                                    atgggggactgcttgaagtcaggagttaagaccagcctggtcaacatagtgagacccca 41267
chimp54546-135457.revcompl.Pan.troglodytes.clone.rp43-41g5.GenBank:AC099533.36   ATGGGGGACTGCTTGAAGTCAGGAGTTAAGACCAGCCTGGTCAACATAGTGAGACCCCA 43072
human.FAM30B-LOC105372935-GGT2.NCBI.                                             atgggggactgcttgaagtcaggagttaagaccagcctggtcaacatagtgagacccca 39954
human.FAM230E-LOC105377182-GGT3P.NCBI.                                           atgggggactgcttgaagtcaggagttaagaccagcctggtcaacatagtgagacccca 38695
human.FAM230J-LOC105372942-GGTLC5P.NCBI.rev.compl                                atgggggactgcttgaagtcaggagttaagaccagcctggtcaacatagtgagacccca 39414
human.FAM230A.ncRNA.18487127.18500594-GGTLC3.18516335.18518165.rev.compl.REF     atgggggactgcttgaagtcaggagttaagaccagcctggtcaacatagtgagacccca 28161
                                                                                 ************************************************************

chimp.LOC112206744-LOC107973052-GGT2.revcompl                                    tctctattggtttaaattttgttt--tttaattaaataaataaaataaggattgaagagt 41325
chimp54546-135457.revcompl.Pan.troglodytes.clone.rp43-41g5.GenBank:AC099533.36   TCTCTATTGGTTTAAATTTTTTTT---TTAATTAAATAAATAAAATAAGGATTGAAGAGT 43129
human.FAM30B-LOC105372935-GGT2.NCBI.                                             tctctattggtttaaattttttttaaaaaaattaaataaataaaataaggattgaagagt 40014
human.FAM230E-LOC105377182-GGT3P.NCBI.                                           tctctattggtttaaattttttttaaaaaaattaaataaataaaataaggattgaagagt 38755
human.FAM230J-LOC105372942-GGTLC5P.NCBI.rev.compl                                tctctattggtttaaattttttttaaaaaaattaaataaataaaataaggattgaagagt 39474
human.FAM230A.ncRNA.18487127.18500594-GGTLC3.18516335.18518165.rev.compl.REF     tctctattggtttaaattttttttaaaaaaattaaataaataaaataaggattgaagagt 28221
                                                                                 ******************** **      *******************************

chimp.LOC112206744-LOC107973052-GGT2.revcompl                                    gacttgtacaccagttgagcccacctccatctcacccttgcagagccccagagacacagc 41385
chimp54546-135457.revcompl.Pan.troglodytes.clone.rp43-41g5.GenBank:AC099533.36   GACTTGTACACCATTTGAGCCCACCTCCATCTCACCCTTGCAGAGCCCCAGAGACACAGC 43189
human.FAM30B-LOC105372935-GGT2.NCBI.                                             gacttgtacaccagttgagcccacctccatctcacccttgcagagccccagagacacagc 40074
human.FAM230E-LOC105377182-GGT3P.NCBI.                                           gacttgtacaccagttgagcccacctccatctcacccttgcagagccccagagacacagc 38815
human.FAM230J-LOC105372942-GGTLC5P.NCBI.rev.compl                                gacttgtacaccagttgagcccacctccatctcacccttgcagagccccagagacacagc 39534
human.FAM230A.ncRNA.18487127.18500594-GGTLC3.18516335.18518165.rev.compl.REF     gacttgtacaccagttgagcccacctccatctcacccttgcagagccccagagacacagc 28281
                                                                                 ************* **********************************************

chimp.LOC112206744-LOC107973052-GGT2.revcompl                                    cctccagggctcagacccagtgggacttgactccacaggcataaaaccctgtttgtctat 41445
chimp54546-135457.revcompl.Pan.troglodytes.clone.rp43-41g5.GenBank:AC099533.36   CCTCCAGAGCTCAGACCCAGTGGGACTTGACTCCACAGGCATAAAACCCTGTTTGTCTAT 43249
human.FAM30B-LOC105372935-GGT2.NCBI.                                             cctccagagctcagacccagtgggacttgactccacaggcataaaaccctgtttgtctat 40134
human.FAM230E-LOC105377182-GGT3P.NCBI.                                           cctccagagctcagacccagtgggacttgactccacaggcataaaaccctgtttgtctat 38875
human.FAM230J-LOC105372942-GGTLC5P.NCBI.rev.compl                                cctccagagctcagacccagtgggacttgactccacaggcataaaaccctgtttgtctat 39594
human.FAM230A.ncRNA.18487127.18500594-GGTLC3.18516335.18518165.rev.compl.REF     cctccagagctcagacccagtgggacttgactccacaggcataaaaccctgtttgtctat 28341
                                                                                 ******* ****************************************************

chimp.LOC112206744-LOC107973052-GGT2.revcompl                                    gggccctttggaatcaccaggttttcggggatcctgaaggatagccccgacctggcctca 41505
chimp54546-135457.revcompl.Pan.troglodytes.clone.rp43-41g5.GenBank:AC099533.36   GGGCCCTTTGGAATCACCAGGTTTTCGGGGCTCCTGAAGGATAGCCCCGACCTGGCCTCA 43309
human.FAM30B-LOC105372935-GGT2.NCBI.                                             gggccctttggaatcaccaggttttcggggctcctgaaggatagccccgacctggcctca 40194
human.FAM230E-LOC105377182-GGT3P.NCBI.                                           gggccctttggaatcaccaggttttcggggctcctgaaggatagccccgacctggcctca 38935
human.FAM230J-LOC105372942-GGTLC5P.NCBI.rev.compl                                gggccctttggaatcaccaggttttcggggctcctgaaggatagccccgacctggcctca 39654
human.FAM230A.ncRNA.18487127.18500594-GGTLC3.18516335.18518165.rev.compl.REF     gggccctttggaatcaccaggttttcggggctcctgaaggatagccccgacctggcctca 28401
                                                                                 ****************************** *****************************

chimp.LOC112206744-LOC107973052-GGT2.revcompl                                    cctggcccctggccccagtgcccctggtgatatccaggtgc------------------- 41546
chimp54546-135457.revcompl.Pan.troglodytes.clone.rp43-41g5.GenBank:AC099533.36   CCTGGCCCCTGGCCCCAGTGCCCCTGGTGATATCCAGGTGC------------------- 43350
human.FAM30B-LOC105372935-GGT2.NCBI.                                             cctggcccctggccccagtgcccctggtgatatccaggtgctgggctgtgatcaccgcct 40254
human.FAM230E-LOC105377182-GGT3P.NCBI.                                           cctggcccctggccccagtgcccctggtgatatccaggtgc------------------- 38976
human.FAM230J-LOC105372942-GGTLC5P.NCBI.rev.compl                                cctggcccctggccccagtgcccctggtgatatccaggtgc------------------- 39695
human.FAM230A.ncRNA.18487127.18500594-GGTLC3.18516335.18518165.rev.compl.REF     cctggcccctggccccagtgcccctggtgatatccaggtgc------------------- 28442
                                                                                 *****************************************

chimp.LOC112206744-LOC107973052-GGT2.revcompl                                    -------------------------------------------------tgggctgtgat 41557
chimp54546-135457.revcompl.Pan.troglodytes.clone.rp43-41g5.GenBank:AC099533.36   -------------------------------------------------TGGGCTGTGAT 43361
human.FAM30B-LOC105372935-GGT2.NCBI.                                             cccaccagcccacctccaccagcccttcccagaaccctgctccaggtgttgggctgtgat 40314
human.FAM230E-LOC105377182-GGT3P.NCBI.                                           -------------------------------------------------tgggctgtgat 38987
human.FAM230J-LOC105372942-GGTLC5P.NCBI.rev.compl                                -------------------------------------------------tgggctgtgat 39706
human.FAM230A.ncRNA.18487127.18500594-GGTLC3.18516335.18518165.rev.compl.REF     -------------------------------------------------tgggctgtgat 28453
                                                                                                                                  ***********

chimp.LOC112206744-LOC107973052-GGT2.revcompl                                    ctccgcctcccaccagcccacctccaccagcccttcccagaaccctgccccaggtgttgg 41617
chimp54546-135457.revcompl.Pan.troglodytes.clone.rp43-41g5.GenBank:AC099533.36   CACCGCCTCCCACCAGCCCACCTCCACCAGCCCTTCCCAGAACCCTGCTCCAGGTGTTGG 43421
human.FAM30B-LOC105372935-GGT2.NCBI.                                             caccgcctcccaccagcccacctccaccagcccttcccagaaccctgctccaggtgttgg 40374
human.FAM230E-LOC105377182-GGT3P.NCBI.                                           caccgcctcccaccagcccacctccaccagcccttcccagaaccctgctccaggtgttgg 39047
human.FAM230J-LOC105372942-GGTLC5P.NCBI.rev.compl                                caccgcctcccaccagcccacctccaccagcccttcccagaaccctgctccaggtgttgg 39766
human.FAM230A.ncRNA.18487127.18500594-GGTLC3.18516335.18518165.rev.compl.REF     caccgcctcccaccagcccacctccaccagcccttcccagaaccctgctccaggtgttgg 28513
                                                                                 * ********************************************** ***********

chimp.LOC112206744-LOC107973052-GGT2.revcompl                                    aactgtgcacagaggagggagcaggccctgagggaggcctggaggggctgcggatggtga 41677
chimp54546-135457.revcompl.Pan.troglodytes.clone.rp43-41g5.GenBank:AC099533.36   AACTGTGCACAGAGGAGGGAGCAGGCCCCGAGGGAGGCCTGGAGGGACTGCGGATGGTGA 43481
human.FAM30B-LOC105372935-GGT2.NCBI.                                             aactgtgcacagaggagggagcaggccccgagggaggcctggaggggctgccaatggtga 40434
human.FAM230E-LOC105377182-GGT3P.NCBI.                                           aactgtgcacagaggagggagcaggccccgagggaggcctggaggggctgccaatggtga 39107
human.FAM230J-LOC105372942-GGTLC5P.NCBI.rev.compl                                aactgtgcacagaggagggagcaggccccgagggaggcctggaggggctgccgatggtga 39826
human.FAM230A.ncRNA.18487127.18500594-GGTLC3.18516335.18518165.rev.compl.REF     aactgtgcacagaggagggagcaggccccgagggaggcctggaggggctgccaatggtga 28573
                                                                                 **************************** **************** ****   *******

chimp.LOC112206744-LOC107973052-GGT2.revcompl                                    aggctgctgtgtctagctgtttccttctggacccactccctctgggctgcgtccccggct 41737
chimp54546-135457.revcompl.Pan.troglodytes.clone.rp43-41g5.GenBank:AC099533.36   AGGCTGCTGTGTCTAGCTGTTTCCTTCTGGACCCACTCCCTCTGGGCTGCGTCCCCGGCT 43541
human.FAM30B-LOC105372935-GGT2.NCBI.                                             aggctgctgtgtcgagctgtttccttccggacccactccctctgggctgcgtccccggct 40494
human.FAM230E-LOC105377182-GGT3P.NCBI.                                           aggctgctgtgtcgagctgtttccttccggacccactccctctgggctgcgtccccggct 39167
human.FAM230J-LOC105372942-GGTLC5P.NCBI.rev.compl                                aggctgctgtgtcgagctgtttccttccggacccactccctctgggctgcgtccccggct 39886
human.FAM230A.ncRNA.18487127.18500594-GGTLC3.18516335.18518165.rev.compl.REF     aggctgctgtgtcgagctgtttccttccggacccactccctctgggctgcgtccccggct 28633
                                                                                 ************* ************* ********************************

chimp.LOC112206744-LOC107973052-GGT2.revcompl                                    ggtccaagccctgatccctgggatctggggacatcttcccgtttgttgttccctgagaac 41797
chimp54546-135457.revcompl.Pan.troglodytes.clone.rp43-41g5.GenBank:AC099533.36   GGTCCAAGCCCTGATCCCTGGGATCTGGGGACATCTTCCCGTTTGTTGTTCCCTGAGAAC 43601
human.FAM30B-LOC105372935-GGT2.NCBI.                                             ggtccaagccctgatccctgggatctggggacatcttcccgtttgctgttccctgagaac 40554
human.FAM230E-LOC105377182-GGT3P.NCBI.                                           ggtccaagccctgatccctgggatctggggacatcttcccgtttgctgttccctgagaac 39227
human.FAM230J-LOC105372942-GGTLC5P.NCBI.rev.compl                                ggtccaagccctgatccctgggatctggggacatcttcccgtttgctgttccctgagaac 39946
human.FAM230A.ncRNA.18487127.18500594-GGTLC3.18516335.18518165.rev.compl.REF     ggtccaagccctgatccctgggatctggggacatcttcccgtttgctgttccctgagaac 28693
                                                                                 ******************************************** ***************

chimp.LOC112206744-LOC107973052-GGT2.revcompl                                    caggcctccctctggagaggatcacaagcttgggtttcactctgggcttgctcttgggaa 41857
chimp54546-135457.revcompl.Pan.troglodytes.clone.rp43-41g5.GenBank:AC099533.36   CAGGCCTCCCTCTGGAGAGGATCACAAGCTTGGGTTTCACTCTGGGCTTGCTCTTGGGAA 43661
human.FAM30B-LOC105372935-GGT2.NCBI.                                             caggcctccctctggagaggatcacaagcttgggtttcactctgggcttgctcttgggaa 40614
human.FAM230E-LOC105377182-GGT3P.NCBI.                                           caggcctccctctggagaggatcacaagcttgggtttcactctgggcttgctcttgggaa 39287
human.FAM230J-LOC105372942-GGTLC5P.NCBI.rev.compl                                caggcctccctctggagaggatcacaagcttgggtttcactctgggcttgctcttgggaa 40006
human.FAM230A.ncRNA.18487127.18500594-GGTLC3.18516335.18518165.rev.compl.REF     caggcctccctgtggagaggatcacaagcttgggtttcactctgggcttgctcttgggaa 28753
                                                                                 *********** ************************************************

chimp.LOC112206744-LOC107973052-GGT2.revcompl                                    cccccaggggcgtggctctgaccgagatgttttcctccagcctgttgcctagtccccat 41917
chimp54546-135457.revcompl.Pan.troglodytes.clone.rp43-41g5.GenBank:AC099533.36   CCCCCAGGGGCGTGGCTCTGACCGAGATGTTTTCCTCCAGCCTGTTGCCCAGTCCCCAT 43721
human.FAM30B-LOC105372935-GGT2.NCBI.                                             cccccaggggcatggctctgaccgagatgttttcctccagcctgttgcccagtccccat 40674
human.FAM230E-LOC105377182-GGT3P.NCBI.                                           cccccaggggcatggctctgaccgagatgttttcctccagcctgttgcccagtccccat 39347
human.FAM230J-LOC105372942-GGTLC5P.NCBI.rev.compl                                cccccaggggcatggctctgaccgagatgttttcctccagcctgttgcccagtccccat 40066
human.FAM230A.ncRNA.18487127.18500594-GGTLC3.18516335.18518165.rev.compl.REF     cccccaggggcatggctctgaccgagatgttttcctccagcctgttgcccagtccccat 28813
                                                                                 ************ ************************************* *********

chimp.LOC112206744-LOC107973052-GGT2.revcompl                                    tcctcgga-cctcagcttcacctccagtgtcatcggcagggtgagctggacgcctacggg 41976
chimp54546-135457.revcompl.Pan.troglodytes.clone.rp43-41g5.GenBank:AC099533.36   TCCTCGGA-CCTCAGCTTCACCTCCAGTGTCATCGGCAGGGTGAGCTGGACGCCTACGGG 43780
human.FAM30B-LOC105372935-GGT2.NCBI.                                             tcctcggaccctcagcttcacctccagtgtcatcggcagggtgagctggacgcctacggg 40734
human.FAM230E-LOC105377182-GGT3P.NCBI.                                           tcctcggaccctcagcttcacctccagtgtcatcggcagggtgagctggacgcctacggg 39407
human.FAM230J-LOC105372942-GGTLC5P.NCBI.rev.compl                                tcctcggaccctcagcttcacctccagtgtcatcggcagggtgagctggacgcctacggg 40126
human.FAM230A.ncRNA.18487127.18500594-GGTLC3.18516335.18518165.rev.compl.REF     tcctcggaccctcagcttcacctccagtgtcatcggcagggtgagctggacgcctacggg 28873
                                                                                 ******** ***************************************************

chimp.LOC112206744-LOC107973052-GGT2.revcompl                                    tctgagaaggcacccgggttcccagcatcagctggccacc-------------------- 42016
chimp54546-135457.revcompl.Pan.troglodytes.clone.rp43-41g5.GenBank:AC099533.36   TCTGAGAAGGCGCCCGGGTTCCCAGCATCAGCTGGCCACCCTCTGCCTAAGAAAGTGCCA 43840
human.FAM30B-LOC105372935-GGT2.NCBI.                                             tctgagaaggcgcccgggttcccagcatcggctggccaccctctgcctaagaaagcgcca 40794
human.FAM230E-LOC105377182-GGT3P.NCBI.                                           tctgagaaggcgcccgggttcccagcatcggctggccaccctctgcctaagaaagcgcca 39467
human.FAM230J-LOC105372942-GGTLC5P.NCBI.rev.compl                                tctgagaaggcgcccgggttcccagcatcggctggccaccctctgcctaagaaagcgcca 40186
human.FAM230A.ncRNA.18487127.18500594-GGTLC3.18516335.18518165.rev.compl.REF     tctgagaaggcgcccgggttcccagcatcggctggccaccctctgcctaagaaagcgcca 28933
                                                                                 *********** ***************** **********

chimp.LOC112206744-LOC107973052-GGT2.revcompl                                    ------------------------------------------------------------ 42016
chimp54546-135457.revcompl.Pan.troglodytes.clone.rp43-41g5.GenBank:AC099533.36   GGGTCGTGGCACCCCCTGGTGGCTGATCCTAGGTAGTGTCACTGCCCAGCCCCAGTAAGG 43900
human.FAM30B-LOC105372935-GGT2.NCBI.                                             gggtcgtgacacccccctggtggctgatcctaggtagtgtcactgcccagccccagtaagg 40854
human.FAM230E-LOC105377182-GGT3P.NCBI.                                           gggtcgtgacacccccctggtggctgatcctaggtagtgtcactgcccagccccagtaagg 39527
human.FAM230J-LOC105372942-GGTLC5P.NCBI.rev.compl                                gggtcgtgacacccccctggtggctgatcctaggtagtgtcactgcccagccccagtaagg 40246
human.FAM230A.ncRNA.18487127.18500594-GGTLC3.18516335.18518165.rev.compl.REF     gggtcgtgacacccccctggtggctgatcctaggtagtgtcactgcccagccccagtaagg 28993

chimp.LOC112206744-LOC107973052-GGT2.revcompl                                    ------------------------------------------------------------ 42016
chimp54546-135457.revcompl.Pan.troglodytes.clone.rp43-41g5.GenBank:AC099533.36   GAGGGCCTGGCCCCAAAGTCCGAGGGATCAGGGTGGAAAGGGGCAGGGCTTGGTGTGAAT 43960
human.FAM30B-LOC105372935-GGT2.NCBI.                                             gagggcctggccccaaagtctgagggatcagggtgggaaggggcagggtttggtgtgaac 40914
human.FAM230E-LOC105377182-GGT3P.NCBI.                                           gagggcctggccccaaagtctgagggatcagggtgggaaggggcagggtttggtgtgaac 39587
human.FAM230J-LOC105372942-GGTLC5P.NCBI.rev.compl                                gagggcctggccccaaagtctgagggatcagggtgggaaggggcagggtttggtgtgaac 40306
human.FAM230A.ncRNA.18487127.18500594-GGTLC3.18516335.18518165.rev.compl.REF     gagggcctggccccaaagtctgagggatcagggtgggaaggggcagggtttggtgtgaac 29053

chimp.LOC112206744-LOC107973052-GGT2.revcompl                                    ------------------------------------------------------------ 42016
```

```
chimp54546-135457.revcompl.Pan.troglodytes.clone.rp43-41g5.GenBank:AC099533.36      CTTCCCCTGGCCCCCAGCCATGTGCCTGGCTCTCCCCATGCTGAAGATGCTGAGGCTAGT 44020
human.FAM30B-LOC105372935-GGT2.NCBI.                                                 cttcccctggcccccagccatgtgccttgctctccccatgctgaagatgctgaggctagt 40974
human.FAM230E-LOC105377182-GGT3P.NCBI.                                               cttcccctggcccccagccatgtgccttgctctccccatgctgaagatgctgaggctagt 39647
human.FAM230J-LOC105372942-GGTLC5P.NCBI.rev.compl                                    cttcccctggcccccagccatgtgcctggctctccccatgctgaagatgctgaggctagt 40366
human.FAM230A.ncRNA.18487127.18500594-GGTLC3.18516335.18518165.rev.compl.REF         cttcccctggcccccagccatgtgccttgctctccccatgctgaagatgctgaggctagt 29113


chimp.LOC112206744-LOC107973052-GGT2.revcompl                                        --------------------------------------ctctgttcctccccagccagat 42038
chimp54546-135457.revcompl.Pan.troglodytes.clone.rp43-41g5.GenBank:AC099533.36      TCCAGTGTCTGCATTGTGAAGATCTCCGAATCCCACCTCTCTGTTCCTCCCCAGCCAGAT 44080
human.FAM30B-LOC105372935-GGT2.NCBI.                                                 tccagtgtccgcattgtgaagatctccgaatcccacctctctgttcctccccagccagat 41034
human.FAM230E-LOC105377182-GGT3P.NCBI.                                               tccagtgcccgcattgtgaagatctccgaatcccacctctctgttcctccccagccagat 39707
human.FAM230J-LOC105372942-GGTLC5P.NCBI.rev.compl                                    tccagtgtccgcattgtgaagatctccgaatcccacctctctgttcctccccagccagat 40426
human.FAM230A.ncRNA.18487127.18500594-GGTLC3.18516335.18518165.rev.compl.REF         tccagtgcccgcattgtgaagatctccgaatcccacctctctgttcctccccagccagat 29173
                                                                                                                           **********************

chimp.LOC112206744-LOC107973052-GGT2.revcompl                                        ggctccatttcacacacaatacactgaggcccagagagtggggagacaggccagggaggc 42098
chimp54546-135457.revcompl.Pan.troglodytes.clone.rp43-41g5.GenBank:AC099533.36      GGCTCCATTTCACACACAATACACTGAGGCCCAGAGAGTGGGGAGACAGGCCAGGGAGGC 44140
human.FAM30B-LOC105372935-GGT2.NCBI.                                                 ggctccatttcacacacaatacactgaggcccagagagtggggagacaggccagggaggc 41094
human.FAM230E-LOC105377182-GGT3P.NCBI.                                               ggctccatttcacacacaatacactgaggcccagagagtggggagacaggccagggaggc 39767
human.FAM230J-LOC105372942-GGTLC5P.NCBI.rev.compl                                    ggctccatttcacacacaaaacactgaggcccagagagtggggagacaggccagggaggc 40486
human.FAM230A.ncRNA.18487127.18500594-GGTLC3.18516335.18518165.rev.compl.REF         ggctccatttcacacacaatacactgaggcccagagagtggggagacaggccagggaggc 29233
                                                                                     ******************* ****************************************

chimp.LOC112206744-LOC107973052-GGT2.revcompl                                        cacctggagcctggcacagtggcctcatttattatgctgctctgctgctcacaggggaag 42158
chimp54546-135457.revcompl.Pan.troglodytes.clone.rp43-41g5.GenBank:AC099533.36      CACCTGGAGCCTGGCACAGTGGCCTCATTTATTGTGCTGCTCTGCTGCTCACAGGGGAAG 44200
human.FAM30B-LOC105372935-GGT2.NCBI.                                                 cacctggagcctggcacagtggcctcatttattatgctgctctgctgctcacaggggaag 41154
human.FAM230E-LOC105377182-GGT3P.NCBI.                                               cacctggagcctggcacagtggcctcatttattatgctgctctgctgctcacaggggaag 39827
human.FAM230J-LOC105372942-GGTLC5P.NCBI.rev.compl                                    cacctggagcctggcacagtggcctcatttattatgctgctctgctgctcacaggggaag 40546
human.FAM230A.ncRNA.18487127.18500594-GGTLC3.18516335.18518165.rev.compl.REF         cacctggagcctggcacagtggcctcatttattatgctgctctgctgctcacaggggaag 29293
                                                                                     ********************************* **************************

chimp.LOC112206744-LOC107973052-GGT2.revcompl                                        cccgtcccccaaagtcctcttcctcatcctggtgagtatcttgtccctggattgcttgtc 42218
chimp54546-135457.revcompl.Pan.troglodytes.clone.rp43-41g5.GenBank:AC099533.36      CTGGTCCCCCGGAGTCCCCTTCCTGGTCCTGGTGAGTATCTTGTCCCTGGATTGCTTGTC 44260
human.FAM30B-LOC105372935-GGT2.NCBI.                                                 cccgtcccccaaagtcctcttcctcatcctggtgagtatcttgtccctggattgcttgtc 41214
human.FAM230E-LOC105377182-GGT3P.NCBI.                                               cccgtcccccaaagtcctcttcctcatcctggtgagtatcttgtccctggattgcttgtc 39887
human.FAM230J-LOC105372942-GGTLC5P.NCBI.rev.compl                                    cccgtcccccaaagtcctcttcctcatcctggtgagtatcttgtccctggattgcttgtc 40606
human.FAM230A.ncRNA.18487127.18500594-GGTLC3.18516335.18518165.rev.compl.REF         cccgtcccccaaagtcctcttcctcatcctggtgagtatcttgtccctggattgcttgtc 29353
                                                                                     *  *******  ***** ******  **********************************

chimp.LOC112206744-LOC107973052-GGT2.revcompl                                        agccttgtctgcctggagcaatcagtagccagcaggttccccgcctttcctggagtccga 42278
chimp54546-135457.revcompl.Pan.troglodytes.clone.rp43-41g5.GenBank:AC099533.36      AGCCTTGTCCGCCTGGAGCACTCAGTAGCCGGCAGGTTCCCCGCCTTTCCTGGAGTCCGA 44320
human.FAM30B-LOC105372935-GGT2.NCBI.                                                 agccttgtctgcctggagcaatcagtagccagcaggttccccgcctttcctggagtccga 41274
human.FAM230E-LOC105377182-GGT3P.NCBI.                                               agccttgtctgcctggagcactcagtagccagcaggttccccgcctttcctggagtccga 39947
human.FAM230J-LOC105372942-GGTLC5P.NCBI.rev.compl                                    agccttgtctgcctggagcactcagtagccagcaggttccccgcctttcctggagtccga 40666
human.FAM230A.ncRNA.18487127.18500594-GGTLC3.18516335.18518165.rev.compl.REF         agccttgtctgcctggagcaatcagtagccagcaggttccccgcctttcctggagtccga 29413
                                                                                     ********* ********** ********* *****************************

chimp.LOC112206744-LOC107973052-GGT2.revcompl                                        ggcagctgcccagccaccagccatgcggacgatggcttgcaccacagcgatgaaggtgga 42338
chimp54546-135457.revcompl.Pan.troglodytes.clone.rp43-41g5.GenBank:AC099533.36      GGCAGCTGCCCAGCCACCAGCCATGCGGACGATGGCTTGCACCACAGCGATGAAGGTGGA 44380
human.FAM30B-LOC105372935-GGT2.NCBI.                                                 ggcagctgcccagccaccagccgtgcggacgatggcttgcaccacagcgatgaaggtgga 41334
human.FAM230E-LOC105377182-GGT3P.NCBI.                                               ggcagctgcccagccaccagccgtgcggacgatggcttgcaccacagcgatgaaggtgga 40007
human.FAM230J-LOC105372942-GGTLC5P.NCBI.rev.compl                                    ggcagctgcccagccaccagccgtgcggacgatggcttgcaccacagcgatgaaggtgga 40726
human.FAM230A.ncRNA.18487127.18500594-GGTLC3.18516335.18518165.rev.compl.REF         ggcagctgcccagccaccagccgtgcggacgatggcttgcaccacagcgatgaaggtgga 29473
                                                                                     ********************** *************************************

chimp.LOC112206744-LOC107973052-GGT2.revcompl                                        cgcgatctgggtgtgatggtgccgggtctccagggctgcagtcactgcctgggggtggga 42398
chimp54546-135457.revcompl.Pan.troglodytes.clone.rp43-41g5.GenBank:AC099533.36      CGCGATCTGGGTGTGATGGTGCCGGGTCTCCAGGGCTGCAGTCACTGCCTGGGGGTGGGA 44440
human.FAM30B-LOC105372935-GGT2.NCBI.                                                 cgcgatctgggtgtgatggtgccgggtctccagggctgcagtcactgcctgggggtggga 41394
human.FAM230E-LOC105377182-GGT3P.NCBI.                                               cgcgatctgggtgtgatggtgccgggtctccagggctgcagtcactgcctgggggtggga 40067
human.FAM230J-LOC105372942-GGTLC5P.NCBI.rev.compl                                    cgcgatctgggtgtgatggtgccgggtctccagggctgcagtcactgcctgggggtggga 40786
human.FAM230A.ncRNA.18487127.18500594-GGTLC3.18516335.18518165.rev.compl.REF         tgcgatctgggtgtgatggtgccgggtctccagggctgcagtcactgcctgggggtggga 29533
                                                                                      ***********************************************************

chimp.LOC112206744-LOC107973052-GGT2.revcompl                                        ggagaggggaagcctgagcagggctccagatgccacctgaaccacgcctgtgtggtcaca 42458
chimp54546-135457.revcompl.Pan.troglodytes.clone.rp43-41g5.GenBank:AC099533.36      GGAGAGGGGAAACCTGAGCAAGGCTCCAGATGCCACCTGAACCACACCTGTGTGGTCACA 44500
human.FAM30B-LOC105372935-GGT2.NCBI.                                                 ggagaggggaagcctgagcagggctccagatgccacctgaaccacacctgtgtggtcaca 41454
human.FAM230E-LOC105377182-GGT3P.NCBI.                                               ggagaggggaagcctgagcagggctccagatgccacctgaaccacacctgtgtggtcaca 40127
human.FAM230J-LOC105372942-GGTLC5P.NCBI.rev.compl                                    ggagaggggaagcctgagcagggctccagatgccacctgaaccacacctgtgtggtcaca 40846
human.FAM230A.ncRNA.18487127.18500594-GGTLC3.18516335.18518165.rev.compl.REF         ggagaggggaagcctgagcagggctccagatgccacctgaaccacacctgtgtggtcaca 29593
                                                                                     *********** ******** ************************ **************

chimp.LOC112206744-LOC107973052-GGT2.revcompl                                        ggcctcagcccaggtggtgccatttcaggccaggtcatcaggaagagcaggttggggcct 42518
chimp54546-135457.revcompl.Pan.troglodytes.clone.rp43-41g5.GenBank:AC099533.36      GGCCTCAGCCCAGGTGGTGCCATTTCGGGCCAGGTCATCAGGAAGAGCAGGTTGGGGCCT 44560
human.FAM30B-LOC105372935-GGT2.NCBI.                                                 ggcctcagcccaggtggtgccatttcaggccaggtcatcaggaagagcaggttggggcct 41514
human.FAM230E-LOC105377182-GGT3P.NCBI.                                               ggcctcagcccaggtggtgccatttcaggccaggtcatcaggaagagcaggttggggcct 40187
human.FAM230J-LOC105372942-GGTLC5P.NCBI.rev.compl                                    ggcctcagcccaggtggtgccatttcaggccaggtcatcaggaagagcaggttggggcct 40906
human.FAM230A.ncRNA.18487127.18500594-GGTLC3.18516335.18518165.rev.compl.REF         ggcctcagcccaggtggtgccatttcaggccaggtcatcaggaagagcaggttggggcct 29653
                                                                                     ************************** *********************************

chimp.LOC112206744-LOC107973052-GGT2.revcompl                                        gctgggtctcactggagcaggggcttggccctcatggcacaggggctccagatggccca 42578
chimp54546-135457.revcompl.Pan.troglodytes.clone.rp43-41g5.GenBank:AC099533.36      GCTGGGTCTCACTGGAGCAGGGGGCTTGGCCCTCATGGCACAGGGGCTCCAGATGGCCCA 44620
human.FAM30B-LOC105372935-GGT2.NCBI.                                                 gctgggtctcactggagcagggggcttggctctcatgtcacaggggctccagatggccca 41574
human.FAM230E-LOC105377182-GGT3P.NCBI.                                               gctgggtctcactggagcagggggcttggccctcatgtcacaggggctccagatggccca 40247
human.FAM230J-LOC105372942-GGTLC5P.NCBI.rev.compl                                    gctgggtctcactggagcagggggcttggccctcatgtcacaggggctccagatggccca 40966
human.FAM230A.ncRNA.18487127.18500594-GGTLC3.18516335.18518165.rev.compl.REF         gctgggtctcactggagcagggggcttggctctcatgtcacaggggctccagatggccca 29713
                                                                                     ***************************** ****** ***********************

chimp.LOC112206744-LOC107973052-GGT2.revcompl                                        ggcactagagagaggacaccaaccattgtccactctgtgatgatccaggcctccagccca 42638
chimp54546-135457.revcompl.Pan.troglodytes.clone.rp43-41g5.GenBank:AC099533.36      GGCACTAGAGAGAGGACACCAACCATTGTCCACTCTGTGATGATCCAGGCCTCCAGCCCA 44680
human.FAM30B-LOC105372935-GGT2.NCBI.                                                 ggcactagagagaggacaccaaccattgtccactctgtgatgatccaggcctccagccca 41634
human.FAM230E-LOC105377182-GGT3P.NCBI.                                               ggcactagagagaggacaccaaccattgtccactctgtgatgatccaggcctccagccca 40307
human.FAM230J-LOC105372942-GGTLC5P.NCBI.rev.compl                                    ggcactagagagaggacaccaaccattgtccactctgtgatgatccaggcctccagccca 41026
human.FAM230A.ncRNA.18487127.18500594-GGTLC3.18516335.18518165.rev.compl.REF         ggcactagagagaggacaccaaccattgtccactctgtgatgatccaggcctccagccca 29773
                                                                                     ************************************************************

chimp.LOC112206744-LOC107973052-GGT2.revcompl                                        ggatgccctggggccccacaccgtgactcagtttctccaaccccggcccacctggtcaa 42698
chimp54546-135457.revcompl.Pan.troglodytes.clone.rp43-41g5.GenBank:AC099533.36      GGATGCCCTGGGGCCCCACACCGTGACTCAGTTTCTCCAACCCCCGGCCCACCTGGTCAA 44740
human.FAM30B-LOC105372935-GGT2.NCBI.                                                 ggatgccctgggaccccacaccgtgactcagtttctccaacccctggcccacctggtcaa 41694
human.FAM230E-LOC105377182-GGT3P.NCBI.                                               ggatgccctgggaccccacaccgtgactcagtttctccaaccccggcccacctggtcaa 40367
human.FAM230J-LOC105372942-GGTLC5P.NCBI.rev.compl                                    ggatgccctgggaccccacaccgtgactcagtttctccaaccccggcccacctggtcaa 41086
human.FAM230A.ncRNA.18487127.18500594-GGTLC3.18516335.18518165.rev.compl.REF         ggatgccctgggaccccacaccgtgactcagtttctccaacccctggcccacctggtcaa 29833
                                                                                     ************ ******************************* ***************

chimp.LOC112206744-LOC107973052-GGT2.revcompl                                        tgtttctctccactgtcgtgacgttgggcagaagcttgttgtgcagccggggctcctcca 42758
chimp54546-135457.revcompl.Pan.troglodytes.clone.rp43-41g5.GenBank:AC099533.36      TGTTTCTCTCCACTGTCGTGACGTTGGGCAGAAGCTGGTTGTGCAGCCGGGGCTCCTCCA 44800
human.FAM30B-LOC105372935-GGT2.NCBI.                                                 tgtttctctccactgtcgtgacgttgggcagaagcttgttgtgcagccggggctcctcca 41754
human.FAM230E-LOC105377182-GGT3P.NCBI.                                               tgtttctctccactgtcgtgacgttgggcagaagcttgttgtgcagccggggctcctcca 40427
human.FAM230J-LOC105372942-GGTLC5P.NCBI.rev.compl                                    tgtttctctccactgtcgtgacgttgggcagaagcttgttgtgcagccggggctcctcca 41146
human.FAM230A.ncRNA.18487127.18500594-GGTLC3.18516335.18518165.rev.compl.REF         tgtttctctccactgtcgtgacgttgggcagaagcttgttgtgcagccggggctcctcca 29893
                                                                                     ************************************ ***********************

chimp.LOC112206744-LOC107973052-GGT2.revcompl                                        cggccttcttcacgtcatagccgaaccagaggttgtagatgatggcctggggcatgggag 42818
chimp54546-135457.revcompl.Pan.troglodytes.clone.rp43-41g5.GenBank:AC099533.36      CGGCCCGCTTCACGTCATAGCCGAACCAGAGGTTGTAGATGATGGCCTGGGGCATGGGAG 44860
human.FAM30B-LOC105372935-GGT2.NCBI.                                                 cggccctcttcacgtcatagccgaaccagaggttgtagatgatggcctggggcatgggag 41814
human.FAM230E-LOC105377182-GGT3P.NCBI.                                               cggccctcttcacgtcatagccgaaccagaggttgtagatgatggcctggggcatgggag 40487
human.FAM230J-LOC105372942-GGTLC5P.NCBI.rev.compl                                    cggccctcttcacgtcatagccgaaccagaggttgtagatgatggcctggggcatgggag 41206
human.FAM230A.ncRNA.18487127.18500594-GGTLC3.18516335.18518165.rev.compl.REF         cggccctcttcacgtcatagccgaaccagaggttgtagatgatggcctggggcatgggag 29953
                                                                                     *****  *****************************************************

chimp.LOC112206744-LOC107973052-GGT2.revcompl                                        tgtgatcagcgtggcttgggggctgtgcagagtgggcagggccagggagaaaaggggtga 42878
chimp54546-135457.revcompl.Pan.troglodytes.clone.rp43-41g5.GenBank:AC099533.36      TGTGATCAGCGTGGCTTGGGGGCTGTGGAGGGTGGGCACGGCCAGGGAGAAAAGGTGTGA 44920
human.FAM30B-LOC105372935-GGT2.NCBI.                                                 tgtgatcagggtggcttgggggctgtgcagagtgggcagggccagggagaaaaggggtga 41874
human.FAM230E-LOC105377182-GGT3P.NCBI.                                               tgtgatcagcgtggcttgggggctgtgcagagtgggcagggccagggagaaaaggggtga 40547
human.FAM230J-LOC105372942-GGTLC5P.NCBI.rev.compl                                    tgtgatcagcgtggcttgggggctgtgcagagtgggcagggccagggagaaaaggggtga 41266
human.FAM230A.ncRNA.18487127.18500594-GGTLC3.18516335.18518165.rev.compl.REF         tgtgatcagcgtggcttgggggctgtgcagagtgggcagggccagggagaaaaggggtga 30013
                                                                                     ********* ***************** ** ******* **************** ****

chimp.LOC112206744-LOC107973052-GGT2.revcompl                                        cacataccagtgcagtgtctgtggtgatctgcgtgcccccagcagctcccaccaccaacc 42938
chimp54546-135457.revcompl.Pan.troglodytes.clone.rp43-41g5.GenBank:AC099533.36      CACATACCAGTGCAGTGGCTGTGGTGATCTGTGTGCCCCCAGCAGCTCCCACAACCATCT 44980
human.FAM30B-LOC105372935-GGT2.NCBI.                                                 cacataccagtgcagtgtctgtggtgatctgcgtgcccccagcagctcccaccaccatcc 41934
human.FAM230E-LOC105377182-GGT3P.NCBI.                                               cacataccagtgcagtgtctgtggtgatctgcgtgcccccagcagctcccaccaccatcc 40607
human.FAM230J-LOC105372942-GGTLC5P.NCBI.rev.compl                                    cacataccagtgcagtgtctgtggtgatctgcgtgcccccagcagctcccaccaccatcc 41326
human.FAM230A.ncRNA.18487127.18500594-GGTLC3.18516335.18518165.rev.compl.REF         cacataccagtgcagtgtctgtggtgatctgcgtgcccccagcagctcccaccaccatcc 30073
                                                                                     ***************** ************* ******************** **** *

chimp.LOC112206744-LOC107973052-GGT2.revcompl                                        ggacctggccgtcctggcccaccatgatcgtcgggcacatggacgagagcggctgcttcc 42998
chimp54546-135457.revcompl.Pan.troglodytes.clone.rp43-41g5.GenBank:AC099533.36      GGACCTGGCCGTCCTGGCCCACCATGATCGTCGGGCACATGGACGAGAGCGGCTGCTTCC 45040
human.FAM30B-LOC105372935-GGT2.NCBI.                                                 ggacctggccgtcctggcccaccatgatcgtcaggcacatggacaagagcggctgcttcc 41994
human.FAM230E-LOC105377182-GGT3P.NCBI.                                               ggacctggccgtcctggcccaccatgatcgtcgggcacatggacaagagcggctgcttcc 40667
human.FAM230J-LOC105372942-GGTLC5P.NCBI.rev.compl                                    ggacctggccgtcctggcccaccatgatcgtcgggcacatggacaagagcggctgcttcc 41386
human.FAM230A.ncRNA.18487127.18500594-GGTLC3.18516335.18518165.rev.compl.REF         ggacctggccgtcctggcccaccatgatcgtcgggcacatggacaagagcggctgcttcc 30133
                                                                                     ******************************** *********** ***************

chimp.LOC112206744-LOC107973052-GGT2.revcompl                                        ctgcggctgatgggagaagacagggatgcccgtcagctgcctgcccaggacacccgcccc 43058
chimp54546-135457.revcompl.Pan.troglodytes.clone.rp43-41g5.GenBank:AC099533.36      CTGTGGCCGATGGGAGAAGACAGGGATGCCCGTCAGCTGCCTGCCCAGGACACCCGCCCC 45100
```

```
human.FAM30B-LOC105372935-GGT2.NCBI.                                              ctgcggccgatgggagaagacagggatgcccgtcagctgcctgcccaggacacccgcccc 42054
human.FAM230E-LOC105377182-GGT3P.NCBI.                                            ctgcggccgatgggagaagacagggatgcccgtcagctgcctgcccaggacacccgcccc 40727
human.FAM230J-LOC105372942-GGTLC5P.NCBI.rev.compl                                 ctgcggccgatgggagaagacagggatgcccgtcagctgcctgcccaggacacccgcccc 41446
human.FAM230A.ncRNA.18487127.18500594-GGTLC3.18516335.18518165.rev.compl.REF      ctgcggccgatgggagaagacagggatgcccgtcagctgcctgcccaggacacccgcccc 30193
                                                                                  *** *** ****************************************************

chimp.LOC112206744-LOC107973052-GGT2.revcompl                                     tctccaccccagtccccca-cccccagacctccaccccatacctggctggatgaaattgg 43117
chimp54546-135457.revcompl.Pan.troglodytes.clone.rp43-41g5.GenBank:AC099533.36    TCTCCACCCCAGTCCCCCA-TCCCCGGACCCCCACCCCATACCTGGCTGGATGAAATTGG 45159
human.FAM30B-LOC105372935-GGT2.NCBI.                                              tctccaccccagtcccccacccccccggacctccaccccatacctggctggatgaaattgg 42114
human.FAM230E-LOC105377182-GGT3P.NCBI.                                            tctccaccccagtcccccacccccccggacctccaccccatacctggctggatgaaattgg 40787
human.FAM230J-LOC105372942-GGTLC5P.NCBI.rev.compl                                 tctccaccccagtcccccacccccccggacctccaccccatacctggctggatgaaattgg 41506
human.FAM230A.ncRNA.18487127.18500594-GGTLC3.18516335.18518165.rev.compl.REF      tctccaccccagtcccccacccccccagacctccaccccatacctggctggatgaaattgg 30253
                                                                                  *******************  **** **** *****************************

chimp.LOC112206744-LOC107973052-GGT2.revcompl                                     caggtgagggggcaccccaaactcattggtg-atgctgggagagctgaagtcgtcca-t 43175
chimp54546-135457.revcompl.Pan.troglodytes.clone.rp43-41g5.GenBank:AC099533.36    CAGGTGAGGGGGCACCCCAAACTCGTTGGTG-ATGTTGGGAGAGCTGAAGTCGTCCATT 45218
human.FAM30B-LOC105372935-GGT2.NCBI.                                              caggtgaggggggtgccccaaactcattggtgaatgctgggagagctgaagtcgtcca-t 42173
human.FAM230E-LOC105377182-GGT3P.NCBI.                                            caggtgaggggggtgccccaaactcattggtg-atgctgggagagctgaagtcgtcca-t 40845
human.FAM230J-LOC105372942-GGTLC5P.NCBI.rev.compl                                 caggtgaggggggtgccccaaactcattggtg-atgctgggagagctgaagtcgtcca-t 41564
human.FAM230A.ncRNA.18487127.18500594-GGTLC3.18516335.18518165.rev.compl.REF      caggtgaggggggtgccccaaactcattggtgaatgctgggagagctgaagtcgtcca-t 30312
                                                                                  *************  ********** ****** *** ********************* *

chimp.LOC112206744-LOC107973052-GGT2.revcompl                                     tcattattgaacaggatcccactgaccggggagcagaccttggagccaaagctaccgccc 43235
chimp54546-135457.revcompl.Pan.troglodytes.clone.rp43-41g5.GenBank:AC099533.36    TCATCATTGAACAGGATCCCGCTGACCGGGCAGCGGACCTTGGAGCCAAAGCTACCGCCC 45278
human.FAM30B-LOC105372935-GGT2.NCBI.                                              tcattattgaacaggatcccactgaccggggagcagaccttggagccaaagctaccgccc 42233
human.FAM230E-LOC105377182-GGT3P.NCBI.                                            tcattattgaacaggatcccactgaccggggagcagaccttggagccaaagctaccgccc 40905
human.FAM230J-LOC105372942-GGTLC5P.NCBI.rev.compl                                 tcattattgaacaggatcccactgaccggggagcagaccttggagccaaagctaccgccc 41624
human.FAM230A.ncRNA.18487127.18500594-GGTLC3.18516335.18518165.rev.compl.REF      tcattattgaacaggatcccactgaccggggagcagaccttggagccaaagctaccgccc 30372
                                                                                  **** *************** ********* *** *************************

chimp.LOC112206744-LOC107973052-GGT2.revcompl                                     agccaggtcagacagcacctgaccttgcctggcccagcctggtccctatccacccactga 43295
chimp54546-135457.revcompl.Pan.troglodytes.clone.rp43-41g5.GenBank:AC099533.36    AGCCAGGTTAGACAGTGCCCGACCTTGCCTGGCCCAGCCTGGTCCCTATCCACCCACTGA 45338
human.FAM30B-LOC105372935-GGT2.NCBI.                                              agccaggtcagacagcacccgaccttgcctggcccagcctggtccctatccacccactga 42293
human.FAM230E-LOC105377182-GGT3P.NCBI.                                            agccaggtcagacagcacccgaccttgcctggcccagcctggtccctatccacccactga 40965
human.FAM230J-LOC105372942-GGTLC5P.NCBI.rev.compl                                 agccaggtcagacagcacccgaccttgcctggcccagcctggtccctatccacccactga 41684
human.FAM230A.ncRNA.18487127.18500594-GGTLC3.18516335.18518165.rev.compl.REF      agccaggtcagacagcacccgaccttgcctggcccagcctggtccctatccacccactga 30432
                                                                                  ******** ******  ** ****************************************

chimp.LOC112206744-LOC107973052-GGT2.revcompl                                     ggctcaaacatactcactgagaggcccaggataagctaccaaggttgggcctcagtttcc 43355
chimp54546-135457.revcompl.Pan.troglodytes.clone.rp43-41g5.GenBank:AC099533.36    GGCTCAAACATACTCACTGAGAGGCCCAGGATAAGCTACCAAGTTTGGGCCTCAGTTTCC 45398
human.FAM30B-LOC105372935-GGT2.NCBI.                                              ggctcaaacatactcactgagaggcccaggataagctaccaaggttgggcctcagtttcc 42353
human.FAM230E-LOC105377182-GGT3P.NCBI.                                            ggctcaaacatactcactgagaggcccaggataagctaccaaggttgggcctcagtttcc 41025
human.FAM230J-LOC105372942-GGTLC5P.NCBI.rev.compl                                 ggctcaaacatactcactgagaggcccaggataagctaccaaggttgggcctcagtttcc 41744
human.FAM230A.ncRNA.18487127.18500594-GGTLC3.18516335.18518165.rev.compl.REF      ggctcaaacatactcactgagaggcccaggataagctaccaaggttgggcctcagtttcc 30492
                                                                                  ******************************************* ****************

chimp.LOC112206744-LOC107973052-GGT2.revcompl                                     caccaggaaaagaggtgatggagccaccttactggataagtgggcagtccctgggccacc 43415
chimp54546-135457.revcompl.Pan.troglodytes.clone.rp43-41g5.GenBank:AC099533.36    CACCAGGAAAAGAGGTGATGGAGCCACCTTACTGGATAAGTGGGCAGTCCCTGGGCCACC 45458
human.FAM30B-LOC105372935-GGT2.NCBI.                                              caccaggaaaagaggtgatggagccaccttactggataagtgggcagtccctgggccacc 42413
human.FAM230E-LOC105377182-GGT3P.NCBI.                                            caccaggaaaagaggtgatggagccaccttactggataagtgggcagtccctgggccacc 41085
human.FAM230J-LOC105372942-GGTLC5P.NCBI.rev.compl                                 caccaggaaaagaggtgatggagccaccttactggataagtgggcagtccctgggccacc 41804
human.FAM230A.ncRNA.18487127.18500594-GGTLC3.18516335.18518165.rev.compl.REF      caccaggaaaagaggtgatggagccaccttactggataagtgggcagtccctgggccacc 30552
                                                                                  ************************************************************

chimp.LOC112206744-LOC107973052-GGT2.revcompl                                     cgcccctggccctttcccacccaggcggcccagcagcccctactagaggttgatggtgct 43475
chimp54546-135457.revcompl.Pan.troglodytes.clone.rp43-41g5.GenBank:AC099533.36    CGCCCCTGGCCCTTTCCCACCCAGGCAGCCCAGCAGCCCCTACTAGAGGTTCATGGTGCT 45518
human.FAM30B-LOC105372935-GGT2.NCBI.                                              cgcccctggccctttcccacccaggcggcccagcagcccctactagaggttgatggtgct 42473
human.FAM230E-LOC105377182-GGT3P.NCBI.                                            cgcccctggccctttcccacccaggcggcccagcagcccctactagaggttgatggtgct 41145
human.FAM230J-LOC105372942-GGTLC5P.NCBI.rev.compl                                 cgcccctggccctttcccacccaggcggcccagcagcccctactagaggttgatggtgct 41864
human.FAM230A.ncRNA.18487127.18500594-GGTLC3.18516335.18518165.rev.compl.REF      cgcccctggccctttcccacccaggcggcccagcagcccctactagaggttgatggtgct 30612
                                                                                  ************************** ************************ ********

chimp.LOC112206744-LOC107973052-GGT2.revcompl                                     ggtggccgacacagcactgccgtcctctgcgacgacagacaggtgagcagtgcccccgtc 43535
chimp54546-135457.revcompl.Pan.troglodytes.clone.rp43-41g5.GenBank:AC099533.36    GGTGGCAGACACAGCACTGCTGTCCTCTGCGACGACAGACAGGTGAGCAGTGCCCCCATC 45578
human.FAM30B-LOC105372935-GGT2.NCBI.                                              ggtggcggacacagcactgccgtcctctgcgacgacagacaggtgagcagtgcccccgtc 42533
human.FAM230E-LOC105377182-GGT3P.NCBI.                                            ggtggcggacacagcactgccgtcctctgcgacgacagacaggtgagcagtgcccccgtc 41205
human.FAM230J-LOC105372942-GGTLC5P.NCBI.rev.compl                                 ggtggcggacacagcactgccgtcctctgcgacgacagacaggtgagcagtgcccccgtc 41924
human.FAM230A.ncRNA.18487127.18500594-GGTLC3.18516335.18518165.rev.compl.REF      ggtggcggacacagcactgccgtcctctgcgacgacagacaggtgagcagtgcccccgtc 30672
                                                                                  ****** ************* ************************************ **

chimp.LOC112206744-LOC107973052-GGT2.revcompl                                     atccggcgtgtagaactcgggcttgtagtaggagatcgggtgagtggtgtggtcagagat 43595
chimp54546-135457.revcompl.Pan.troglodytes.clone.rp43-41g5.GenBank:AC099533.36    AACCGGCGTGTAGAACTCGGGCTTGTAGTAGGAGATCGGGTGAGTGGTGTCGTCAGAGAT 45638
human.FAM30B-LOC105372935-GGT2.NCBI.                                              atccggcgtgtagaactcgggcttgtagtaggagatcgggtgagtggtgtggtcagagat 42593
human.FAM230E-LOC105377182-GGT3P.NCBI.                                            atccggcgtgtagaactcgggcttgtagtaggagatcgggtgagtggtgtggtcagagat 41265
human.FAM230J-LOC105372942-GGTLC5P.NCBI.rev.compl                                 atccggcgtgtagaactcgggcttgtcgtaggagatcgggtgagtggtgtggtcagagat 41984
human.FAM230A.ncRNA.18487127.18500594-GGTLC3.18516335.18518165.rev.compl.REF      atccggcgtgtagaactcgggcttgtagtaggagatcgggtgagtggtgtggtcagagat 30732
                                                                                  * ************************ *********************** *********

chimp.LOC112206744-LOC107973052-GGT2.revcompl                                     ctgggcctggagctgggcagcgaagaactcggaggtcatgttgcggaccacctgccgaga 43655
chimp54546-135457.revcompl.Pan.troglodytes.clone.rp43-41g5.GenBank:AC099533.36    CTGGGCCCGGAGCTGGGCAGCGAAGAACTCAGAGGTCATGTTGCGGACCACCTGCCGAGA 45698
human.FAM30B-LOC105372935-GGT2.NCBI.                                              ctgggaccggagctgggcagcgaagaactcagaggtcatgttgcggaccacctgctgaga 42653
human.FAM230E-LOC105377182-GGT3P.NCBI.                                            ctgggaccggagctgggcagcgaagaactcagaggtcatgttgcggaccacctgctgaga 41325
human.FAM230J-LOC105372942-GGTLC5P.NCBI.rev.compl                                 ctgggaccggagctgggcagcgaagaactcagaggtcatgttgcggaccacctgctgaga 42044
human.FAM230A.ncRNA.18487127.18500594-GGTLC3.18516335.18518165.rev.compl.REF      ctgggaccggagctgggcagcgaagaactcagaggtcatgttgcggaccacctgctgaga 30792
                                                                                  ***** * ********************** ************************ ****

chimp.LOC112206744-LOC107973052-GGT2.revcompl                                     ccccagagctggcctgaggaggtggggagggagggtggggagggggcacaggtctcagaa 43715
chimp54546-135457.revcompl.Pan.troglodytes.clone.rp43-41g5.GenBank:AC099533.36    CCCCAGAGCTGGCCTGAGGAGGTGGGGAGGGAGGGTGGGGAGGGGGCACAGGTCTCAGAA 45758
human.FAM30B-LOC105372935-GGT2.NCBI.                                              ccccagagctggcctgaggaggtggggagggagggtggggagggggcacaggtctcagaa 42713
human.FAM230E-LOC105377182-GGT3P.NCBI.                                            ccccagagctggcctgagga-------------ggtggggagggggcacaggtctcagaa 41372
human.FAM230J-LOC105372942-GGTLC5P.NCBI.rev.compl                                 ccccagagctggcctgagga-------------ggtggggagggggcacaggtctcagaa 42091
human.FAM230A.ncRNA.18487127.18500594-GGTLC3.18516335.18518165.rev.compl.REF      ccccagagctggcctgaggaggtggggagggagggtggggagggggcacaggtctcagaa 30852
                                                                                  ********************             ***************************

chimp.LOC112206744-LOC107973052-GGT2.revcompl                                     ggcccctgactgtgactctgaccgcaaccctctggcacccacaaccttccgtggctcccc 43775
chimp54546-135457.revcompl.Pan.troglodytes.clone.rp43-41g5.GenBank:AC099533.36    GGCCCCTGACTGTGGCTCTGACCACAACCCTCTGGCACCCACAACCTTCCGTGGCTCCCC 45818
human.FAM30B-LOC105372935-GGT2.NCBI.                                              ggcccttgactgtgactctgaccgcaaccctctggcacccacaaccttccgtggctcccc 42773
human.FAM230E-LOC105377182-GGT3P.NCBI.                                            ggcccttgactgtgactctgaccgcaaccctctggcacccacaaccttccgtggctcccc 41432
human.FAM230J-LOC105372942-GGTLC5P.NCBI.rev.compl                                 ggcccttgactgtgactctgaccgcaaccctctggcacccacaaccttccgtggctcccc 42151
human.FAM230A.ncRNA.18487127.18500594-GGTLC3.18516335.18518165.rev.compl.REF      ggcccttgactgtgactctgaccgcaaccctctggcacccacaaccttccgtggctcccc 30912
                                                                                  ***** ******** ******** ************************************

chimp.LOC112206744-LOC107973052-GGT2.revcompl                                     aggacccaagggcaggcccaggaccttgcacaaccagtctgactccctgtctctgtcgcg 43835
chimp54546-135457.revcompl.Pan.troglodytes.clone.rp43-41g5.GenBank:AC099533.36    AGGACCCAAGGGCAGGCCCAGGATCTTGCATGACCAGTCTGACTCCCTGTCTCTGTCGCG 45878
human.FAM30B-LOC105372935-GGT2.NCBI.                                              aggacccaagggcaggcccaagaccttgcatgaccagtctgactccctgtctctgtcgcg 42833
human.FAM230E-LOC105377182-GGT3P.NCBI.                                            aggacccaagggcaggcccaggaccttgcatgaccagtctgactccctgtctctgtcgcg 41492
human.FAM230J-LOC105372942-GGTLC5P.NCBI.rev.compl                                 aggacccaagggcaggcccaggaccttgcatgaccagtctgactccctgtctctgtcgcg 42211
human.FAM230A.ncRNA.18487127.18500594-GGTLC3.18516335.18518165.rev.compl.REF      aggacccaagggcaggcccaagaccttgcatgaccagtctgactccctgtctctgtcgcg 30972
                                                                                  ******************** ** ******  ****************************

chimp.LOC112206744-LOC107973052-GGT2.revcompl                                     ttccagcaactctgaatgtctgtctgcgtggtcctcagcctccagacccttgccacattc 43895
chimp54546-135457.revcompl.Pan.troglodytes.clone.rp43-41g5.GenBank:AC099533.36    TTCCAGCAACTCTGAATGTCTGTCTGCCTGGTCCTCAGCCTCCAGACCCTTGCTGCATTC 45938
human.FAM30B-LOC105372935-GGT2.NCBI.                                              ttccagcaactctgaatgtctgtctgcctggtcctcagcctccagacccttgccgcattc 42893
human.FAM230E-LOC105377182-GGT3P.NCBI.                                            ttccagcaactctgaatgtctgtctgcctggtcctcagcctccagacccttgccacattc 41552
human.FAM230J-LOC105372942-GGTLC5P.NCBI.rev.compl                                 ttccagcaactctgaatgtctgtctgcctggtcctcagcctccagacccttgccgcattc 42271
human.FAM230A.ncRNA.18487127.18500594-GGTLC3.18516335.18518165.rev.compl.REF      ttccagcaactctgaatgtctgtctgcctggtcctcagcctccagacccttgccgcattc 31032
                                                                                  *************************** *************************  *****

chimp.LOC112206744-LOC107973052-GGT2.revcompl                                     aatcactcattctttcatgcaaaaaatatttctagaatttgcactgcatgcctggcactg 43955
chimp54546-135457.revcompl.Pan.troglodytes.clone.rp43-41g5.GenBank:AC099533.36    AATCACTCATTCCTTCATGCAAAAAATATTTCTAGAGTTTGCACTGCATGCCTGGAACTG 45998
human.FAM30B-LOC105372935-GGT2.NCBI.                                              aatcactcattccttcatgcaaaaaatatttctagaatttgcactgcatgcctggcactg 42953
human.FAM230E-LOC105377182-GGT3P.NCBI.                                            aatcactcattctttcatgcaaaaaatatttctagaatttgcactgcatgcctggcactg 41612
human.FAM230J-LOC105372942-GGTLC5P.NCBI.rev.compl                                 aatcactcattctttcatgcaaaaaatatttctagaatttgcactgcatgcctggcactg 42331
human.FAM230A.ncRNA.18487127.18500594-GGTLC3.18516335.18518165.rev.compl.REF      aatcact----------------------------------------------------- 31039
                                                                                  *******

chimp.LOC112206744-LOC107973052-GGT2.revcompl                                     gggaatcaacagggaacagacatttaggtcctgccctcatgccaagaaaaacaaacacac 44015
chimp54546-135457.revcompl.Pan.troglodytes.clone.rp43-41g5.GenBank:AC099533.36    GGGAATCAACAGGGAACAGACACTTAGGTCCTGCCCTCACGCCAAGAAAAACAAACACAC 46058
human.FAM30B-LOC105372935-GGT2.NCBI.                                              gggaatcaacagggaacagacacttaagtcctgccctcatgccaagaaaaacaaacacac 43013
human.FAM230E-LOC105377182-GGT3P.NCBI.                                            gggaatcaacagggaacagacacttaagtcctgccctcatgccaagaaaaacaaacacac 41672
human.FAM230J-LOC105372942-GGTLC5P.NCBI.rev.compl                                 gggaatcaacagggaacagacacttaagtcctgccctcatgccaagaaaaacaaacacac 42391
human.FAM230A.ncRNA.18487127.18500594-GGTLC3.18516335.18518165.rev.compl.REF      ------------------------------------------------------------ 31039

chimp.LOC112206744-LOC107973052-GGT2.revcompl                                     acagggaaagtgctgaaaccacaagccaggtaaggggaatcaagaggcatgaggtatggg 44075
chimp54546-135457.revcompl.Pan.troglodytes.clone.rp43-41g5.GenBank:AC099533.36    ACAGGGAAAGTGCTGAAACCACAAGCCAGGTAAGGGGAATCAAGAGGCATGAGGTATGGG 46118
human.FAM30B-LOC105372935-GGT2.NCBI.                                              ccagggaaagtgctgaaaccacaggccaggtaaggggaatcaagaggcatgaggtatggg 43073
human.FAM230E-LOC105377182-GGT3P.NCBI.                                            acagggaaagtgctgaaaccacaggccaggtaaggggaatcaagaggcatgaggtatggg 41732
human.FAM230J-LOC105372942-GGTLC5P.NCBI.rev.compl                                 acagggaaagtgctgaaaccacaggccaggtaaggggaatcaagaggcatgaggtatggg 42451
human.FAM230A.ncRNA.18487127.18500594-GGTLC3.18516335.18518165.rev.compl.REF      ------------------------------------------------------------ 31039

chimp.LOC112206744-LOC107973052-GGT2.revcompl                                     cagagtggtcagggagggcttctcagaggaggcagcgtgtgaaaagagcctggaatgtgg 44135
chimp54546-135457.revcompl.Pan.troglodytes.clone.rp43-41g5.GenBank:AC099533.36    CAGAGTGGTCAGGGAGGGCTTCTCAGAGGAGGCAACATGTGAAAAGAGCCTGGAATGTGG 46178
human.FAM30B-LOC105372935-GGT2.NCBI.                                              cagagtggtcagggagggcttctcagaggaggcaacgtgtgaaaagagcctggaatgtgg 43133
```

```
human.FAM230E-LOC105377182-GGT3P.NCBI.                                            cagagtggtcagggagggcttctcagaggaggcaacgtgtgaaaagagcctggaaagtgg 41792
human.FAM230J-LOC105372942-GGTLC5P.NCBI.rev.compl                                 cagagtggtcagggagggcttctcagaggaggcaacgtgtgaaaagagcctggaatgttg 42511
human.FAM230A.ncRNA.18487127.18500594-GGTLC3.18516335.18518165.rev.compl.REF      ------------------------------------------------------------ 31039

chimp.LOC112206744-LOC107973052-GGT2.revcompl                                     cctaaatggtcagtgcaaaggccctgaggcaggtggtataggctggtgagcgataggcag 44195
chimp54546-135457.revcompl.Pan.troglodytes.clone.rp43-41g5.GenBank:AC099533.36    CCTAAATGGTCAGTGCAAAGGCCCTGAGGCAGGTGGTATAGGCTGGTGAGCGATAGGCAG 46238
human.FAM30B-LOC105372935-GGT2.NCBI.                                              cctaaatggtcagtgcaaaggccctgaggcaggtggtataggctggtgagcgataggcag 43193
human.FAM230E-LOC105377182-GGT3P.NCBI.                                            cctaaatggtcagtgcaaaggccctgaggcaggtggcataggctggtgagcgataggcag 41852
human.FAM230J-LOC105372942-GGTLC5P.NCBI.rev.compl                                 cctaaatggtcagtgcaaaggccctgaggcaggtggtataggctggtgagcgataggcag 42571
human.FAM230A.ncRNA.18487127.18500594-GGTLC3.18516335.18518165.rev.compl.REF      ------------------------------------------------------------ 31039

chimp.LOC112206744-LOC107973052-GGT2.revcompl                                     agagtgaatggagtaggatggggagaagaggatgaagatgcaggctggggcccatcccac 44255
chimp54546-135457.revcompl.Pan.troglodytes.clone.rp43-41g5.GenBank:AC099533.36    AGAGTGAATGGAGTGGGGTGGGGAGAAGAGGATGAAGATGCAGGCTGGGGCCCATCCCAC 46298
human.FAM30B-LOC105372935-GGT2.NCBI.                                              agagtgaatggagtagggtggggagaagaggatgaagatgcaggctggggcccatcccac 43253
human.FAM230E-LOC105377182-GGT3P.NCBI.                                            agagtgaatggagtagggtggggagaagaggatgaagatgcaggctggggcccatcccac 41912
human.FAM230J-LOC105372942-GGTLC5P.NCBI.rev.compl                                 agagtgaatggagtggggtggggagaagaggatgaagatgcaggctggggcccatcccac 42631
human.FAM230A.ncRNA.18487127.18500594-GGTLC3.18516335.18518165.rev.compl.REF      ------------------------------------------------------------ 31039

chimp.LOC112206744-LOC107973052-GGT2.revcompl                                     aggacctcctaggtcccataacaactggcttttgctctgtgccatgcaggcttagggcag 44315
chimp54546-135457.revcompl.Pan.troglodytes.clone.rp43-41g5.GenBank:AC099533.36    AGGACCTCCTAGGTCCCATAACAACTGGCTTTTGCTCTGTGCCATGCAGGCTTAGGGCAG 46358
human.FAM30B-LOC105372935-GGT2.NCBI.                                              aggacctcctaggtcccataacaactggctttttctctgtgccatgcaggcttagggcag 43313
human.FAM230E-LOC105377182-GGT3P.NCBI.                                            aggacctcctaggtcccataacaactggcttttgctctgtgccatgcaggcttagggcag 41972
human.FAM230J-LOC105372942-GGTLC5P.NCBI.rev.compl                                 aggacctcctaggtcccataacaactggcttttgctctgtgccatgcaggcttagggcag 42691
human.FAM230A.ncRNA.18487127.18500594-GGTLC3.18516335.18518165.rev.compl.REF      ------------------------------------------------------------ 31039

chimp.LOC112206744-LOC107973052-GGT2.revcompl                                     aggaatgaggagggtggggagtgttttcacagggtccctctggcagctatgacggggata 44375
chimp54546-135457.revcompl.Pan.troglodytes.clone.rp43-41g5.GenBank:AC099533.36    AGGAATGAGCAGGCTGGGGAGTGTTTTCACAGGGTCCCTCTGGCAGCTATGACGGGGATA 46418
human.FAM30B-LOC105372935-GGT2.NCBI.                                              aggaatgaggaggctggggagtgttttcacagggtccctctggcagctatgacggggata 43373
human.FAM230E-LOC105377182-GGT3P.NCBI.                                            aggaatgaggaggctggggagtgttttcacagggtccctctggcagctatgacggggata 42032
human.FAM230J-LOC105372942-GGTLC5P.NCBI.rev.compl                                 aagaatgaggaggctggggagtgttttcacagggtccctctggcagctatgacggggata 42751
human.FAM230A.ncRNA.18487127.18500594-GGTLC3.18516335.18518165.rev.compl.REF      ------------------------------------------------------------ 31039

chimp.LOC112206744-LOC107973052-GGT2.revcompl                                     aggataaagcccaaaggggaggctgtgggtatcaaccaggcaagagatgatggcctgggt 44435
chimp54546-135457.revcompl.Pan.troglodytes.clone.rp43-41g5.GenBank:AC099533.36    AGGATAAAGCCAAAAGGGGAGGCTGTGGGTATCAACCAGGCAAGAGATGATGGCCTGGGT 46478
human.FAM30B-LOC105372935-GGT2.NCBI.                                              aggataaagcccaaaggggaggctgtgggtatcaaccaggcaagagatgatggcctgggt 43433
human.FAM230E-LOC105377182-GGT3P.NCBI.                                            aggataaagcccaaaggggaggctgtgggtatcaaccaggcaagagatgatggcctgggt 42092
human.FAM230J-LOC105372942-GGTLC5P.NCBI.rev.compl                                 aggataaagcccaaaggggaggctgtgggtatcaaccaggcaagagatgatggcctgggt 42811
human.FAM230A.ncRNA.18487127.18500594-GGTLC3.18516335.18518165.rev.compl.REF      ------------------------------------------------------------ 31039

chimp.LOC112206744-LOC107973052-GGT2.revcompl                                     gggagaaagagaagaatcaaggatggtgccgactagcgaggtaaaccctgcagaaggggc 44495
chimp54546-135457.revcompl.Pan.troglodytes.clone.rp43-41g5.GenBank:AC099533.36    GGGAGAAAGAGAAGAATCAAGGATGGTGCCGACTAGCGAGGTAAACCCTGCAGAAGGGGC 46538
human.FAM30B-LOC105372935-GGT2.NCBI.                                              gggagaaagagaagaatcaaggatggtgccgactagcgagg-----ccggcagaaggggc 43488
human.FAM230E-LOC105377182-GGT3P.NCBI.                                            gggagaaagagaagaatcaagcatggtgccgactagcgagg-----ccggcagaaggggc 42147
human.FAM230J-LOC105372942-GGTLC5P.NCBI.rev.compl                                 gggagaaagagaagaatcaagcatggtgccgactagcgagg-----ccggcagaaggggc 42866
human.FAM230A.ncRNA.18487127.18500594-GGTLC3.18516335.18518165.rev.compl.REF      ------------------------------------------------------------ 31039

chimp.LOC112206744-LOC107973052-GGT2.revcompl                                     aggtttggggatggtcaggagcttgattttggatacttcatcagacctgaagagcatggg 44555
chimp54546-135457.revcompl.Pan.troglodytes.clone.rp43-41g5.GenBank:AC099533.36    AGGTTTGGGGATGGTCAGGAGCTTGATTTTGGATACTTCATCAGACCCGAAGAGCATGGG 46598
human.FAM30B-LOC105372935-GGT2.NCBI.                                              cggtttggggatggtcaggagcttgattttggatacttcatcagacccaaagagcatggg 43548
human.FAM230E-LOC105377182-GGT3P.NCBI.                                            cggtttggggatggtcaggagcttgattttggatacttcatcagacccaaagagcatggg 42207
human.FAM230J-LOC105372942-GGTLC5P.NCBI.rev.compl                                 cggtttggggatggtcaggagcttgattttggatacttcatcagacccaaagagcatggg 42926
human.FAM230A.ncRNA.18487127.18500594-GGTLC3.18516335.18518165.rev.compl.REF      ------------------------------------------------------------ 31039

chimp.LOC112206744-LOC107973052-GGT2.revcompl                                     tgcacgtataaaaaaaataaataaataagcatgggttcacgggcaagggcgggctgagag 44615
chimp54546-135457.revcompl.Pan.troglodytes.clone.rp43-41g5.GenBank:AC099533.36    TGCAAGTATAAAAAAAATAAATAAATAAGCATGGGTTCACGGGCAAGGGCGGGCTGAGAG 46658
human.FAM30B-LOC105372935-GGT2.NCBI.                                              tgcacgtataaaaaaaataaataaataagcatgggttcacgggcaagggcgggctgagag 43608
human.FAM230E-LOC105377182-GGT3P.NCBI.                                            tgcacgtataaaaaaaataaataaataagcatgggttcacgggcaagggcgggctgagag 42267
human.FAM230J-LOC105372942-GGTLC5P.NCBI.rev.compl                                 tgcacgtataaaaaaaataaataaataagcatgggttcacgggcaagggcgggctgagag 42986
human.FAM230A.ncRNA.18487127.18500594-GGTLC3.18516335.18518165.rev.compl.REF      ------------------------------------------------------------ 31039

chimp.LOC112206744-LOC107973052-GGT2.revcompl                                     atgaacatggaggtattgacattgagtggctgctggatgccatgagcctggccaaggtcc 44675
chimp54546-135457.revcompl.Pan.troglodytes.clone.rp43-41g5.GenBank:AC099533.36    ATGAACACGGAGGTATTGACATTGAGTGGCTGCTGGATGCCATGAGCCTGGCCAAGGTCC 46718
human.FAM30B-LOC105372935-GGT2.NCBI.                                              atgaacatggaggtattgacattgagtggctgctggatgccatgagcctggccaaggtcc 43668
human.FAM230E-LOC105377182-GGT3P.NCBI.                                            atgaacatggaggtattgacattgagtggctgctggatgccatgagcctggccaaggtcc 42327
human.FAM230J-LOC105372942-GGTLC5P.NCBI.rev.compl                                 atgaacatggaggtattgacattgagtggctgctggatgccatgagcctggccaaggtcc 43046
human.FAM230A.ncRNA.18487127.18500594-GGTLC3.18516335.18518165.rev.compl.REF      ------------------------------------------------------------ 31039

chimp.LOC112206744-LOC107973052-GGT2.revcompl                                     ccaaggcagtggcgaggaggagatgaggaggtcaaggaggagac--agaggatggacccg 44733
chimp54546-135457.revcompl.Pan.troglodytes.clone.rp43-41g5.GenBank:AC099533.36    CCAAGGCAGTGGCGAGGAGGAGATGAGGAGGTCAAGGAGGAGAC--AGAGGATGGACCCG 46776
human.FAM30B-LOC105372935-GGT2.NCBI.                                              ccaaggcagtggcgaggaggagatgaggaggtcaaagaggagacagagaggatggacccg 43728
human.FAM230E-LOC105377182-GGT3P.NCBI.                                            ccaaggcagtggcgaggaggagatgaggaggtcaaagaggagacagagaggatggacccg 42387
human.FAM230J-LOC105372942-GGTLC5P.NCBI.rev.compl                                 ccaaggcagtggcgaggaggagatgaggaggtcaaagaggagacagagaggatggacccg 43106
human.FAM230A.ncRNA.18487127.18500594-GGTLC3.18516335.18518165.rev.compl.REF      ------------------------------------------------------------ 31039

chimp.LOC112206744-LOC107973052-GGT2.revcompl                                     aaggccaaagaaaatgcctcaagagagtttcaacaccgggcgcggtggctcacgcctgta 44793
chimp54546-135457.revcompl.Pan.troglodytes.clone.rp43-41g5.GenBank:AC099533.36    AAGGCCGAAGAAAATGCCTCAAGAGAGTTTCAACACCGGGCGCGGTGGCTCACGCCTGTA 46836
human.FAM30B-LOC105372935-GGT2.NCBI.                                              aaggccgaagaaaatgcctcaagagagtttcaacaccgggcgcggtggctcacgcctgta 43788
human.FAM230E-LOC105377182-GGT3P.NCBI.                                            aaggccgaagaaaatgcctcaagagagtttcaacaccgggcgcggtggctcacgcctgta 42447
human.FAM230J-LOC105372942-GGTLC5P.NCBI.rev.compl                                 aaggccgaagaaaatgcctcaagagagtttcaacaccgggcgcggtggctcacgcttgta 43166
human.FAM230A.ncRNA.18487127.18500594-GGTLC3.18516335.18518165.rev.compl.REF      ------------------------------------------------------------ 31039

chimp.LOC112206744-LOC107973052-GGT2.revcompl                                     atcccagcactttgggaggccgaggcctgtaatcccagcactttgggagggcggatcacg 44853
chimp54546-135457.revcompl.Pan.troglodytes.clone.rp43-41g5.GenBank:AC099533.36    ATCCCAGCACTTTGGGAGGCCGAGGCCTGTAATCCCAGCACTTTGGGAGGG-GGATCACG 46895
human.FAM30B-LOC105372935-GGT2.NCBI.                                              atcccagcactttgggaggccgaggcctgtaatcccagcactttgggagggcggatcatg 43848
human.FAM230E-LOC105377182-GGT3P.NCBI.                                            atcccagcactttgggaggccgaggcctgtaatcccagcactttgggagggcggatcatg 42507
human.FAM230J-LOC105372942-GGTLC5P.NCBI.rev.compl                                 atcccagcactttgggaggccgaggcctgtaatcccagcactttgggagggcggatcatg 43226
human.FAM230A.ncRNA.18487127.18500594-GGTLC3.18516335.18518165.rev.compl.REF      ------------------------------------------------------------ 31039

chimp.LOC112206744-LOC107973052-GGT2.revcompl                                     acgtcaggagatcaagaccatcctggctaacacagtgaaaccccgtctctactaaaaata 44913
chimp54546-135457.revcompl.Pan.troglodytes.clone.rp43-41g5.GenBank:AC099533.36    ACGTCAGGAGATGGAGATCATCCTGGCTAACAGGGTGAAACTCCGTCTCTACTAAAAATA 46955
human.FAM30B-LOC105372935-GGT2.NCBI.                                              acgtcaggagatcgagaccatcctggctaacacagtgaaaccccgtctctactaaaaata 43908
human.FAM230E-LOC105377182-GGT3P.NCBI.                                            acgtcaggagatcgagaccatcctggctaacacagtgaaaccccgtctctactaaaaata 42567
human.FAM230J-LOC105372942-GGTLC5P.NCBI.rev.compl                                 acgtcaggagatcgagaccatcctggctaacacagtgaaaccccgtctctactaaaaata 43286
human.FAM230A.ncRNA.18487127.18500594-GGTLC3.18516335.18518165.rev.compl.REF      ------------------------------------------------------------ 31039

chimp.LOC112206744-LOC107973052-GGT2.revcompl                                     caaaaaattagctgggcgcggtggtgggcacctgtagtcccagctacttgggggggctga 44973
chimp54546-135457.revcompl.Pan.troglodytes.clone.rp43-41g5.GenBank:AC099533.36    CAAAAAATTAGCCGGGTGTGGTGGCGGGCGCCTGTAGTCCCAACTACTC-CAGAGGCTGA 47014
human.FAM30B-LOC105372935-GGT2.NCBI.                                              caaaaaattagctgggcgcggtggtgggcacctgtagtcccagctactt-ggggggctga 43967
human.FAM230E-LOC105377182-GGT3P.NCBI.                                            caaaaaattagctgggcgcggtggtgggcacctgtagtcccagctactt-ggggggctga 42626
human.FAM230J-LOC105372942-GGTLC5P.NCBI.rev.compl                                 caaaaaattagctgggcgcggtggtgggcacctgtagtcccagctactt-ggggggctga 43345
human.FAM230A.ncRNA.18487127.18500594-GGTLC3.18516335.18518165.rev.compl.REF      ------------------------------------------------------------ 31039

chimp.LOC112206744-LOC107973052-GGT2.revcompl                                     ggcaggagaatggcatgaacctgggaggcggagcttgcagtgagccgagatcccgccact 45033
chimp54546-135457.revcompl.Pan.troglodytes.clone.rp43-41g5.GenBank:AC099533.36    GGCAGGAGAATGGCGTGAACCCGGTAGGCGGAGCTTGCGGTGAGCCGAGATCAGGCCACT 47074
human.FAM30B-LOC105372935-GGT2.NCBI.                                              ggcaggagaatggcgtgaacctgggaggcggagcttgcagtgagcggagatcctgccact 44027
human.FAM230E-LOC105377182-GGT3P.NCBI.                                            ggcaggagaatggcgtgaacctgggaggcagagcttgcagtgagccgagatcccgccact 42686
human.FAM230J-LOC105372942-GGTLC5P.NCBI.rev.compl                                 ggcaggagaatggcgtgaacctgggaggcggagcttgcagtgagccgagatcccgccact 43405
human.FAM230A.ncRNA.18487127.18500594-GGTLC3.18516335.18518165.rev.compl.REF      ------------------------------------------------------------ 31039

chimp.LOC112206744-LOC107973052-GGT2.revcompl                                     gcactccagcctgggggacagcctgggggacagagcgagactccgtctca---------- 45083
chimp54546-135457.revcompl.Pan.troglodytes.clone.rp43-41g5.GenBank:AC099533.36    GGAATC------------CAGCCTGGGCGATAGAGGGAGACTCCATCTCA---------A 47113
human.FAM30B-LOC105372935-GGT2.NCBI.                                              gcactccagcctgggggacagcctgggggacagagcgagactccgtctca---------- 44077
human.FAM230E-LOC105377182-GGT3P.NCBI.                                            gcactccagcctgggggacagcctgggggacagagcgagactccgtctca---------- 42736
human.FAM230J-LOC105372942-GGTLC5P.NCBI.rev.compl                                 gcactccagcctgggggacagcctgggggacagagcgagactccgtctcaaaaataaaat 43465
human.FAM230A.ncRNA.18487127.18500594-GGTLC3.18516335.18518165.rev.compl.REF      ------------------------------------------------------------ 31039

chimp.LOC112206744-LOC107973052-GGT2.revcompl                                     aaaataaaataaaataaaataaaataaagtttcagcaacaccccacggatt--agttgac 45141
chimp54546-135457.revcompl.Pan.troglodytes.clone.rp43-41g5.GenBank:AC099533.36    AAAAAAAAAAAAAAAAAAAAAGAGTCTGGCTCTGTTGCCCAGGCTGGAGTGCAGTGGCG 47173
human.FAM30B-LOC105372935-GGT2.NCBI.                                              aaaataaaataaaataaaataaaataaagtttcagcaacaccccacggatt--agttgac 44135
human.FAM230E-LOC105377182-GGT3P.NCBI.                                            -----aaaataaaataaaataaaataaagtttcagcaacaccccacggatt--agctgac 42789
human.FAM230J-LOC105372942-GGTLC5P.NCBI.rev.compl                                 aaaataaaataaaataaaataaaataaagtttcagcaacaccccacggatt--agctgac 43523
human.FAM230A.ncRNA.18487127.18500594-GGTLC3.18516335.18518165.rev.compl.REF      ------------------------------------------------------------ 31039

chimp.LOC112206744-LOC107973052-GGT2.revcompl                                     caatcccagggaaaggtgttctgtctgaatctgccctcaaggaaacagaaaggcaaatcc 45201
chimp54546-135457.revcompl.Pan.troglodytes.clone.rp43-41g5.GenBank:AC099533.36    CGATCTCGGGGCATCACAATCCCTTCCCCGCCCGCCCCCGGGTTCAAGTGATTCTCCTGT 47233
human.FAM30B-LOC105372935-GGT2.NCBI.                                              caatcccagggaaaggtgttctgtctgaatctgccctcaaggaaacagaaaggcaaatcc 44195
human.FAM230E-LOC105377182-GGT3P.NCBI.                                            caatcccagggaaaggtgttctgtctgaatctgccctcaaggaaacagaaaggcaaatcc 42849
```

```
human.FAM230J-LOC105372942-GGTLC5P.NCBI.rev.compl                     caatcccagggaaaggtgttctgtctgaatctgccctcaaggaaacagaaaggcaaatcc 43583
human.FAM230A.ncRNA.18487127.18500594-GGTLC3.18516335.18518165.rev.compl.REF   ------------------------------------------------------------ 31039

chimp.LOC112206744-LOC107973052-GGT2.revcompl                         acgatgtgggacattttccaagactactgccctgggctttaaaaatcaacaaaacagg-- 45259
chimp54546-135457.revcompl.Pan.troglodytes.clone.rp43-41g5.GenBank:AC099533.36   CTCAGCCGCCGGAGTAGCTGGGACTACAGGCGCGTGCCACCATGTCTGACTAAATTTGTA 47293
human.FAM30B-LOC105372935-GGT2.NCBI.                                  acgatgtgggacattttccaagactactgccctgggctttaaaaatcaacaaaacagg-- 44253
human.FAM230E-LOC105377182-GGT3P.NCBI.                                acgatgtgggacattttccaagactactgccctgggctttaaaaatcaacaaaacagg-- 42907
human.FAM230J-LOC105372942-GGTLC5P.NCBI.rev.compl                     acgatgtgggacattttccaagactactgccctgggctttaaaaatcaacaaaacagg-- 43641
human.FAM230A.ncRNA.18487127.18500594-GGTLC3.18516335.18518165.rev.compl.REF   ------------------------------------------------------------ 31039

chimp.LOC112206744-LOC107973052-GGT2.revcompl                         ------ccaggcacggtggctcatgcctgtaatcccagcactttgggaggccgaggcagg 45313
chimp54546-135457.revcompl.Pan.troglodytes.clone.rp43-41g5.GenBank:AC099533.36   TTTTTACTAGAGACGGGGTTTCACTATGTT-----------------------GGCCAGG 47330
human.FAM30B-LOC105372935-GGT2.NCBI.                                  ------ccaggcacggtggttcatgcctgtaatcccagcactttgggaggccgaggcagg 44307
human.FAM230E-LOC105377182-GGT3P.NCBI.                                ------ccaggcacggtggctcatgcctgtaatcccagcactttgggaggccgaggcagg 42961
human.FAM230J-LOC105372942-GGTLC5P.NCBI.rev.compl                     ------ccaggcacggtggctcatgcctgtaatcccagcactttgggaggccgaggcagg 43695
human.FAM230A.ncRNA.18487127.18500594-GGTLC3.18516335.18518165.rev.compl.REF   ------------------------------------------------------------ 31039

chimp.LOC112206744-LOC107973052-GGT2.revcompl                         cggatcacgaggtcaggagatcgcgatcacggtgaaaccccgtctctactaaaaatacaa 45373
chimp54546-135457.revcompl.Pan.troglodytes.clone.rp43-41g5.GenBank:AC099533.36   CTGGTCTCCAACTCCTGATCTCGTGATCCGTCTGCCCCGACCTCCCAAAGTGCTAGGATT 47390
human.FAM30B-LOC105372935-GGT2.NCBI.                                  cggatcacgaggtcaggagatcgcgatcacggtgaaaccccgtctctactaaaaatacaa 44367
human.FAM230E-LOC105377182-GGT3P.NCBI.                                cggatcacgaggtcaggagatcgcgatcacggtgaaaccccgtctctactaaaaatacaa 43021
human.FAM230J-LOC105372942-GGTLC5P.NCBI.rev.compl                     cggatcacgaggtcaggagatcgcgatcacggtgaaaccccgtctctactaaaaatacaa 43755
human.FAM230A.ncRNA.18487127.18500594-GGTLC3.18516335.18518165.rev.compl.REF   ------------------------------------------------------------ 31039

chimp.LOC112206744-LOC107973052-GGT2.revcompl                         aaaattagctgggcgcagtggcgggtgcctgtagtcccagcta--------ctcgggagg 45425
chimp54546-135457.revcompl.Pan.troglodytes.clone.rp43-41g5.GenBank:AC099533.36   ATAGGCATAAGCCGCCACGCCCGGCCTCTTTTTTTTCTTTTTCTTTTTTTTATCTGGAGA 47450
human.FAM30B-LOC105372935-GGT2.NCBI.                                  aaaattagctgggcgcagtgtcgggcgcctgtagtcccagcta--------ctcgggagg 44419
human.FAM230E-LOC105377182-GGT3P.NCBI.                                aaaattagctgggcgcagtgtcgggcgcctgtagtcccagcta--------ctcgggagg 43073
human.FAM230J-LOC105372942-GGTLC5P.NCBI.rev.compl                     aaaattagctgggcgcagtgtcgggcgcctgtagtcccagcta--------ctcgggagg 43807
human.FAM230A.ncRNA.18487127.18500594-GGTLC3.18516335.18518165.rev.compl.REF   ------------------------------------------------------------ 31039

chimp.LOC112206744-LOC107973052-GGT2.revcompl                         ctgaggcaggagaatggcgtgaacccaggaggcggagcttgcagtgagccgagatagcac 45485
chimp54546-135457.revcompl.Pan.troglodytes.clone.rp43-41g5.GenBank:AC099533.36   CTGAGTTTTGCACTTGTTGCCCAGGCTGGAG--------TGCAATGGTGCGATCTCAGCT 47502
human.FAM30B-LOC105372935-GGT2.NCBI.                                  ctgaggcaggagaatggcgtgaacccaggaggcggagcttgcagtgagccgagatagcgc 44479
human.FAM230E-LOC105377182-GGT3P.NCBI.                                ctgaggcaggagaatggcgtgaacccaggaggcagagcttgcagtgagccgagatagcac 43133
human.FAM230J-LOC105372942-GGTLC5P.NCBI.rev.compl                     ctgaggcaggagaatggcgtgaacccaggaggcggagcttgcagtgagccgagatagcgc 43867
human.FAM230A.ncRNA.18487127.18500594-GGTLC3.18516335.18518165.rev.compl.REF   ------------------------------------------------------------ 31039

chimp.LOC112206744-LOC107973052-GGT2.revcompl                         cactgcactc-cagcctgggcgacagagcgacaccctgtctcacaaaaaagccacacctg 45544
chimp54546-135457.revcompl.Pan.troglodytes.clone.rp43-41g5.GenBank:AC099533.36   CACTGCAATCTCCACCTCAGCAGGAGAGCAGGAATCTTCAGTGATCCACCGGCAGATCTG 47562
human.FAM30B-LOC105372935-GGT2.NCBI.                                  cactgcactc-cagcctgggcaacagagcgagactccgtctcaaagaaaaaacacacctg 44538
human.FAM230E-LOC105377182-GGT3P.NCBI.                                cactgcactc-cagcctgggcaacagagcgagactccgtctcaaagaaaaaacacacctg 43192
human.FAM230J-LOC105372942-GGTLC5P.NCBI.rev.compl                     cactgcactc-cagcctgggcaacagagcgagactccgtctcaaagaaaaaacacacctg 43926
human.FAM230A.ncRNA.18487127.18500594-GGTLC3.18516335.18518165.rev.compl.REF   ------------------------------------------------------------ 31039

chimp.LOC112206744-LOC107973052-GGT2.revcompl                         taatcccagcgctttgggaggctgaggtgggaagatagtttgatcccaggagtttgagac 45604
chimp54546-135457.revcompl.Pan.troglodytes.clone.rp43-41g5.GenBank:AC099533.36   -----CCGCCATTGTGGGCACCTGTTCCTCCCGCGACCTTTGTGCCCGCCTCTCTCCTTC 47617
human.FAM30B-LOC105372935-GGT2.NCBI.                                  taatcccagcgctttgggaggctgaggtgggaagacagtttgatcccaggagtttgagac 44598
human.FAM230E-LOC105377182-GGT3P.NCBI.                                taatcccagcgctttgggaggctgaggtgggaagacagtttgatcccaggagtttgagac 43252
human.FAM230J-LOC105372942-GGTLC5P.NCBI.rev.compl                     taatcccagcgctttgggaggctgaggtgggaagacagtttgatcccaggagtttgagac 43986
human.FAM230A.ncRNA.18487127.18500594-GGTLC3.18516335.18518165.rev.compl.REF   ------------------------------------------------------------ 31039

chimp.LOC112206744-LOC107973052-GGT2.revcompl                         cagtctgggcaagaccctgtctctaaaaaaatatataaaaattatccaggtttggtggca 45664
chimp54546-135457.revcompl.Pan.troglodytes.clone.rp43-41g5.GenBank:AC099533.36   CAGT----------ACCTATTGCATGACCCC-CCACGTCCGCCTCCCGCCATTGCCAGCA 47666
human.FAM30B-LOC105372935-GGT2.NCBI.                                  cagtctgggcaagaccctgtctctaaaaaaa-atacaaaaattatccaggtttggtggca 44657
human.FAM230E-LOC105377182-GGT3P.NCBI.                                cagtctgggcaagaccctgtctctaaaaaaa-atacaaaaattatccaggtttggtggca 43311
human.FAM230J-LOC105372942-GGTLC5P.NCBI.rev.compl                     cagtctgggcaagaccctgtctctaaaaaaa-atacaaaaattatccaggtttggtggca 44045
human.FAM230A.ncRNA.18487127.18500594-GGTLC3.18516335.18518165.rev.compl.REF   ------------------------------------------------------------ 31039

chimp.LOC112206744-LOC107973052-GGT2.revcompl                         cgtgcctctggt------------------------------------------------ 45676
chimp54546-135457.revcompl.Pan.troglodytes.clone.rp43-41g5.GenBank:AC099533.36   AGTGCCTCGCGCGGGTACCTGGCTGCGCTTATTAATCCGTTAAGCTCGCTCTGTCACGGG 47726
human.FAM30B-LOC105372935-GGT2.NCBI.                                  cgtgcctctggt------------------------------------------------ 44669
human.FAM230E-LOC105377182-GGT3P.NCBI.                                cctgcctctggt------------------------------------------------ 43323
human.FAM230J-LOC105372942-GGTLC5P.NCBI.rev.compl                     cgtgcctctggt------------------------------------------------ 44057
human.FAM230A.ncRNA.18487127.18500594-GGTLC3.18516335.18518165.rev.compl.REF   ------------------------------------------------------------ 31039

chimp.LOC112206744-LOC107973052-GGT2.revcompl                         ------------------------------------------------------------ 45676
chimp54546-135457.revcompl.Pan.troglodytes.clone.rp43-41g5.GenBank:AC099533.36   CGCCATGATGTGCTCACGCGCCCGCTCCCTCAGGTTTAAAAGGCGCGTTGCCCGGCAACA 47786
human.FAM30B-LOC105372935-GGT2.NCBI.                                  ------------------------------------------------------------ 44669
human.FAM230E-LOC105377182-GGT3P.NCBI.                                ------------------------------------------------------------ 43323
human.FAM230J-LOC105372942-GGTLC5P.NCBI.rev.compl                     ------------------------------------------------------------ 44057
human.FAM230A.ncRNA.18487127.18500594-GGTLC3.18516335.18518165.rev.compl.REF   ------------------------------------------------------------ 31039

chimp.LOC112206744-LOC107973052-GGT2.revcompl                         ------------------cccagctgctcagggggctgaagtgggaggattgcttgagcc 45718
chimp54546-135457.revcompl.Pan.troglodytes.clone.rp43-41g5.GenBank:AC099533.36   GAAGAAACTGCTGGCTTAGCCTTTGGCCGAGTTGGCGGCTGGACGAGGACGCTCAGAGCC 47846
human.FAM30B-LOC105372935-GGT2.NCBI.                                  ------------------cccagctgctcagggggctgaagtgggaggattgcttgagcc 44711
human.FAM230E-LOC105377182-GGT3P.NCBI.                                ------------------cccagctgctcagggggctgaagtgggaggattgcttgagcc 43365
human.FAM230J-LOC105372942-GGTLC5P.NCBI.rev.compl                     ------------------cccagctgctcagggggctgaagtgggaggattgcttgagcc 44099
human.FAM230A.ncRNA.18487127.18500594-GGTLC3.18516335.18518165.rev.compl.REF   ------------------------------------------------------------ 31039

chimp.LOC112206744-LOC107973052-GGT2.revcompl                         ctggaggttgaggctgcagtgagccaagatcacaccactgcactccaacctggatgacag 45778
chimp54546-135457.revcompl.Pan.troglodytes.clone.rp43-41g5.GenBank:AC099533.36   C-AGCTCTCGAGAGTTCAAGCAACCGACGGTTCCCCACTGCTCCCAGGAGCGGTTACCTG 47905
human.FAM30B-LOC105372935-GGT2.NCBI.                                  ctggaggttgaggctgcagtgagccaagatcacaccactgcactccaacctggatgacag 44771
human.FAM230E-LOC105377182-GGT3P.NCBI.                                ctggaggttgaggctgcagtgagccaagatcacaccactgcactccaacctggatgacag 43425
human.FAM230J-LOC105372942-GGTLC5P.NCBI.rev.compl                     ctggaggttgaggctgcagtgagccaagatcacaccactgcactccaacctggatgacag 44159
human.FAM230A.ncRNA.18487127.18500594-GGTLC3.18516335.18518165.rev.compl.REF   ------------------------------------------------------------ 31039

chimp.LOC112206744-LOC107973052-GGT2.revcompl                         ag-actctgtgtccaaacaaaaacacaacaaaaacaaaacagccaagagagcctactgta 45837
chimp54546-135457.revcompl.Pan.troglodytes.clone.rp43-41g5.GenBank:AC099533.36   GGCACTCTGTGCCC-------CTCCTTCCTGTTCGGGCCCAGGCCGAGGACCTGCCAGTA 47958
human.FAM30B-LOC105372935-GGT2.NCBI.                                  ag-actctgtgtacaaacaaaaacacaacaaaaacaaaacagccaagggagcctactgta 44830
human.FAM230E-LOC105377182-GGT3P.NCBI.                                ag-actctgtgtacaaacaaaaacacaacaaaaacaaaacagccaagggagcctactgta 43484
human.FAM230J-LOC105372942-GGTLC5P.NCBI.rev.compl                     ag-actctgtgtacaaacaaaaacacaacaaaaacaaaacagccaagggagcctactgta 44218
human.FAM230A.ncRNA.18487127.18500594-GGTLC3.18516335.18518165.rev.compl.REF   ------------------------------------------------------------ 31039

chimp.LOC112206744-LOC107973052-GGT2.revcompl                         ggtaaagagaagggacaacagcaggttgtgtcaccccgacatcctgctgtgctatgttca 45897
chimp54546-135457.revcompl.Pan.troglodytes.clone.rp43-41g5.GenBank:AC099533.36   GGGCTCAGTTGCCT---GGAGCCCGTTCAGCCCATCCCCCAGTTCACTTTGC-------- 48007
human.FAM30B-LOC105372935-GGT2.NCBI.                                  ggtaaagagaaggg---acagcaggttgtgtcaccccgacatcctgctgtgctatgttca 44887
human.FAM230E-LOC105377182-GGT3P.NCBI.                                ggtaaagagaaggg---acagcaggttgtgtcaccccgacatcctgctgtgctatgttca 43541
human.FAM230J-LOC105372942-GGTLC5P.NCBI.rev.compl                     ggtaaagagaaggg---acagcaggttgtgtcaccccgacatcctgctgtgctatgttca 44275
human.FAM230A.ncRNA.18487127.18500594-GGTLC3.18516335.18518165.rev.compl.REF   ------------------------------------------------------------ 31039

chimp.LOC112206744-LOC107973052-GGT2.revcompl                         agtctcacttttgagacacaccctgaggtgcgacatcagtaacctactgtggaatccctc 45957
chimp54546-135457.revcompl.Pan.troglodytes.clone.rp43-41g5.GenBank:AC099533.36   ------------------------------------------------------------ 48007
human.FAM30B-LOC105372935-GGT2.NCBI.                                  agtctcacttttgagacacaccctgaggtgcgacatcagtaacctactgtggaatccctc 44947
human.FAM230E-LOC105377182-GGT3P.NCBI.                                agtctcacttttgagacacaccctgaggtgcgacatcagtaacctactgtggaatccctc 43601
human.FAM230J-LOC105372942-GGTLC5P.NCBI.rev.compl                     agtctcacttttgagacacaccctgaggtgcgacatcagtaacctactgtggaatccctc 44335
human.FAM230A.ncRNA.18487127.18500594-GGTLC3.18516335.18518165.rev.compl.REF   ------------------------------------------------------------ 31039

chimp.LOC112206744-LOC107973052-GGT2.revcompl                         agaaaaacacgaatcccaatagacgtgggtggagacggagagagttaggaaatccggcag 46017
chimp54546-135457.revcompl.Pan.troglodytes.clone.rp43-41g5.GenBank:AC099533.36   ------------------------------------------------------------ 48007
human.FAM30B-LOC105372935-GGT2.NCBI.                                  agaaaaacacgaatcccaatagatgtgggtggagacggagagagttaggaaatccggcag 45007
human.FAM230E-LOC105377182-GGT3P.NCBI.                                agaaaaacacgaatcccaatagatgtgggtggagacggagagagttaggaaatccggcag 43661
human.FAM230J-LOC105372942-GGTLC5P.NCBI.rev.compl                     agaaaaacacgaatcccaatagatgtgggtggagacggagagagttaggaaatccggcag 44395
human.FAM230A.ncRNA.18487127.18500594-GGTLC3.18516335.18518165.rev.compl.REF   ------------------------------------------------------------ 31039

chimp.LOC112206744-LOC107973052-GGT2.revcompl                         aaacgtccacactgcagactccaggaaaagggaacattgatgcttgggcagttttggggt 46077
chimp54546-135457.revcompl.Pan.troglodytes.clone.rp43-41g5.GenBank:AC099533.36   ------------------------------------------------------------ 48007
human.FAM30B-LOC105372935-GGT2.NCBI.                                  aaatgtccacactgcagaatccaggaaaagggaacattgatgcttgggcagttttgggtt 45067
human.FAM230E-LOC105377182-GGT3P.NCBI.                                aaatgtccacactgcagaatccaggaaaagggaacattgatgcttgggcagttttgggtt 43721
human.FAM230J-LOC105372942-GGTLC5P.NCBI.rev.compl                     aaatgtccacactgcagaatccaggaaaagggaacattgatgcttgggcagttttgggtt 44455
human.FAM230A.ncRNA.18487127.18500594-GGTLC3.18516335.18518165.rev.compl.REF   ------------------------------------------------------------ 31039

chimp.LOC112206744-LOC107973052-GGT2.revcompl                         atttttacatttttgtaggtgtgaaaatttgcaaaatgaaaactcgag--gagtgtggtg 46135
chimp54546-135457.revcompl.Pan.troglodytes.clone.rp43-41g5.GenBank:AC099533.36   -------------------------------------------------------TTGTG 48012
human.FAM30B-LOC105372935-GGT2.NCBI.                                  ttttttacatttttgtaggtgcgaaaatttgcaaaatgaaaactcgaggagagtgtggtg 45127
human.FAM230E-LOC105377182-GGT3P.NCBI.                                ttttttacatttttgtaggtgcgaaaatttgcaaaatgaaaactcgaggagagtgtggtg 43781
human.FAM230J-LOC105372942-GGTLC5P.NCBI.rev.compl                     ttttttacatttttgtaggtgcgaaaatttgcaaaatgaaaactcgaggagagtgtggtg 44515
```

```
human.FAM230A.ncRNA.18487127.18500594-GGTLC3.18516335.18518165.rev.compl.REF        ------------------------------------------------------------ 31039

chimp.LOC112206744-LOC107973052-GGT2.revcompl                                        agctgtgtgagatgctgctgagtggggcctgatggggaaactgaggctggacatggcgat 46195
chimp54546-135457.revcompl.Pan.troglodytes.clone.rp43-41g5.GenBank:AC099533.36       GGATCTCCCCGTTGCTCCTGCCCGTGGACTGAGTGGCAGGCCATCCTACAAGCACCCGGA 48072
human.FAM30B-LOC105372935-GGT2.NCBI.                                                 agctgtgtgagatgctgctgagtggggcctgatggggaaactgaggctggacatggcgat 45187
human.FAM230E-LOC105377182-GGT3P.NCBI.                                               agctgtgtgagatgctgctgagtggggcctgatggggaaactgaggctggacatggcgat 43841
human.FAM230J-LOC105372942-GGTLC5P.NCBI.rev.compl                                    agctgtgtgagatgctgctgagtggggcctgatggggaaactgaggctggacatggcgat 44575
human.FAM230A.ncRNA.18487127.18500594-GGTLC3.18516335.18518165.rev.compl.REF        ------------------------------------------------------------ 31039

chimp.LOC112206744-LOC107973052-GGT2.revcompl                                        ctggtggcatggggatagagcaggggaggggataccctgaagggagagaggacatagccc 46255
chimp54546-135457.revcompl.Pan.troglodytes.clone.rp43-41g5.GenBank:AC099533.36       CACTTGACATCAGTGGTGTCAAGACAACGCTA---------------------------- 48104
human.FAM30B-LOC105372935-GGT2.NCBI.                                                 ctggtggcatggggatagagcaggggaggggataccctgaagggagagaggacatagccc 45247
human.FAM230E-LOC105377182-GGT3P.NCBI.                                               ctggtggcatggggatagagcaggggaggggataccctgaagggagagaggacatagccc 43901
human.FAM230J-LOC105372942-GGTLC5P.NCBI.rev.compl                                    ctggtggcatggggatagagcaggggaggggataccctgaagggagagaggacatagccc 44635
human.FAM230A.ncRNA.18487127.18500594-GGTLC3.18516335.18518165.rev.compl.REF        ------------------------------------------------------------ 31039

chimp.LOC112206744-LOC107973052-GGT2.revcompl                                        agccatgtttgctctaagaggagcaaaggacagaaggaggcagcagatagaaatttctag 46315
chimp54546-135457.revcompl.Pan.troglodytes.clone.rp43-41g5.GenBank:AC099533.36       --------------------------------------------AGAAGGTTTTCCGTGA 48120
human.FAM30B-LOC105372935-GGT2.NCBI.                                                 agccatgtttgctctaagaggagcaaaggacagaaggaggcagcagatagaagtttctag 45307
human.FAM230E-LOC105377182-GGT3P.NCBI.                                               agccatgtttgctctaagaggagcaaaggacagaaggaggcagcagatagaagtttctag 43961
human.FAM230J-LOC105372942-GGTLC5P.NCBI.rev.compl                                    agccatgtttgctctaagaggagcaaaggacagaaggaggcagcagatagaagtctctag 44695
human.FAM230A.ncRNA.18487127.18500594-GGTLC3.18516335.18518165.rev.compl.REF        ------------------------------------------------------------ 31039

chimp.LOC112206744-LOC107973052-GGT2.revcompl                                        agcaacaacagctgcctttcttttgggaataatccgtgataaagaagtaaatcgtcagag 46375
chimp54546-135457.revcompl.Pan.troglodytes.clone.rp43-41g5.GenBank:AC099533.36       TCCTGCAAGCCCTGCGTTCCTTCCTGGGATCCTGCCTT---------------------- 48158
human.FAM30B-LOC105372935-GGT2.NCBI.                                                 agcaacaacagctgcctttctttcgggaataatccgtgataaagaaataaatcatcagag 45367
human.FAM230E-LOC105377182-GGT3P.NCBI.                                               agcaacaacagctgcctttttttttgggaataatccgtgataaagaaataaatcatcagag 44021
human.FAM230J-LOC105372942-GGTLC5P.NCBI.rev.compl                                    agcaataacagctgcctttttttttgggaataatccgtgataaagaaataaatcatcagag 44755
human.FAM230A.ncRNA.18487127.18500594-GGTLC3.18516335.18518165.rev.compl.REF        ------------------------------------------------------------ 31039

chimp.LOC112206744-LOC107973052-GGT2.revcompl                                        gcggaaaggagcattgtaggaacagcaccccgagccagcgagtagatgaaggagctggcc 46435
chimp54546-135457.revcompl.Pan.troglodytes.clone.rp43-41g5.GenBank:AC099533.36       -----CAATTTGATTGCACAGGTACCACAGCAAGCCAGTGCTGTGTGCTCCGAGTTCCAG 48213
human.FAM30B-LOC105372935-GGT2.NCBI.                                                 gcagacaggagcattgtaggaacagcaccccgagccagtgagtagatgaaggagctggcc 45427
human.FAM230E-LOC105377182-GGT3P.NCBI.                                               gcagacaggagcattgtaggaacagcaccccgagccagcgagtagatgaaggagctggcc 44081
human.FAM230J-LOC105372942-GGTLC5P.NCBI.rev.compl                                    gcagacaggagcattgtaggaacagcaccccgagccagcgagtagatgaaggacctggcc 44815
human.FAM230A.ncRNA.18487127.18500594-GGTLC3.18516335.18518165.rev.compl.REF        ------------------------------------------------------------ 31039

chimp.LOC112206744-LOC107973052-GGT2.revcompl                                        ttagccaaaagggagggcagagggacacgctgcagtggctctgtcccctcagagagacaa 46495
chimp54546-135457.revcompl.Pan.troglodytes.clone.rp43-41g5.GenBank:AC099533.36       GGCGTCCTCCAGCTCAGCCACTGCGCTGAGAACATGGACTCTCT---------------- 48257
human.FAM30B-LOC105372935-GGT2.NCBI.                                                 ttagccaaaaaggagggcagagggacacgctgcagtggctctgtcccctcagagagacaa 45487
human.FAM230E-LOC105377182-GGT3P.NCBI.                                               ttagccaaaaaggagggcagagggacacgctgcagtggctctgtcccctcagagagacaa 44141
human.FAM230J-LOC105372942-GGTLC5P.NCBI.rev.compl                                    ttagccaaaaaggagggcagagggacacgctgcagtggctctgtccccttagagagacaa 44875
human.FAM230A.ncRNA.18487127.18500594-GGTLC3.18516335.18518165.rev.compl.REF        ------------------------------------------------------------ 31039

chimp.LOC112206744-LOC107973052-GGT2.revcompl                                        gacaccaggtcactggctgcagcgggagtcagaggtgcagaatgctcacggggaaagaga 46555
chimp54546-135457.revcompl.Pan.troglodytes.clone.rp43-41g5.GenBank:AC099533.36       ---------------------GTGGGGCCCAGGAGCCGGGAGTCACCCCTTTGGGGTCCA 48296
human.FAM30B-LOC105372935-GGT2.NCBI.                                                 gacaccaggtcactggctgcagcgggagtcagaggtgcagaatgctcacagggaaagaga 45547
human.FAM230E-LOC105377182-GGT3P.NCBI.                                               gacaccaggtcactggctgcagcgggagtcagaggtgcagaatgctcacagggaaagaga 44201
human.FAM230J-LOC105372942-GGTLC5P.NCBI.rev.compl                                    gacaccaggtcactggctgcagcgggagtcagaggtgcagaatgctcacagggaaagaga 44935
human.FAM230A.ncRNA.18487127.18500594-GGTLC3.18516335.18518165.rev.compl.REF        ------------------------------------------------------------ 31039

chimp.LOC112206744-LOC107973052-GGT2.revcompl                                        agacaccacccgggcagctgacgcccctcctgggaggtcactggtcagtgt------ggg 46609
chimp54546-135457.revcompl.Pan.troglodytes.clone.rp43-41g5.GenBank:AC099533.36       CAACACCCGGCTGTCCCCAGACTTGTGTCCAGGGAAGATAGTGTTGAGGGCCCTCAAGGA 48356
human.FAM30B-LOC105372935-GGT2.NCBI.                                                 agacaccacccgggcagctgacgcccctcctgggaggtcactggtcagtgt------ggg 45601
human.FAM230E-LOC105377182-GGT3P.NCBI.                                               agacaccacccgggcagctgacgcccctcctgggaggtcactggtcagtgt------ggg 44255
human.FAM230J-LOC105372942-GGTLC5P.NCBI.rev.compl                                    agacaccacccgggcagctgacgcccctcctgggaggtcactggtcagtgt------ggg 44989
human.FAM230A.ncRNA.18487127.18500594-GGTLC3.18516335.18518165.rev.compl.REF        ------------------------------------------------------------ 31039

chimp.LOC112206744-LOC107973052-GGT2.revcompl                                        ggggtctgcagatccgcccaggaatgccaaggacccaagtaggtaaggagggatgtgagg 46669
chimp54546-135457.revcompl.Pan.troglodytes.clone.rp43-41g5.GenBank:AC099533.36       GAGCGGGGCAGGGATGCCTGAGCAGGACAAGGACCCTAGAGTCCAAGAGAATCCTGGTGA 48416
human.FAM30B-LOC105372935-GGT2.NCBI.                                                 ggggtctgcagatccgcccaggaatgccaaggacccaagtaggtaaggagggatgtgagg 45661
human.FAM230E-LOC105377182-GGT3P.NCBI.                                               ggggtctgcagatccgcccaggaatgccaaggacccaagtaggtaaggagggatgtgagg 44315
human.FAM230J-LOC105372942-GGTLC5P.NCBI.rev.compl                                    ggggtctgcagatccgcccaggaatgccaaggacccaagtaggtaaggagggatgtgagg 45049
human.FAM230A.ncRNA.18487127.18500594-GGTLC3.18516335.18518165.rev.compl.REF        ------------------------------------------------------------ 31039

chimp.LOC112206744-LOC107973052-GGT2.revcompl                                        atcctcagcaggaagggatatgacagggtcttatagggacccagcgtggagctggggcaa 46729
chimp54546-135457.revcompl.Pan.troglodytes.clone.rp43-41g5.GenBank:AC099533.36       TCAGAGAAGGGTCCCCGAGGTCACCGGGGATGCACGGTCTGCATTTCGGCCCCTGCGGGA 48476
human.FAM30B-LOC105372935-GGT2.NCBI.                                                 atcctcagtgggaagggatatgacagggtcttatagggacccagcatggagctggggcaa 45721
human.FAM230E-LOC105377182-GGT3P.NCBI.                                               atcctcagcgggaagggatatgacagggtcttatagggacccagcatggagctggggcaa 44375
human.FAM230J-LOC105372942-GGTLC5P.NCBI.rev.compl                                    atcctcagcgggaagggatatgacagggtcttatagggacccagcatggagctggggcaa 45109
human.FAM230A.ncRNA.18487127.18500594-GGTLC3.18516335.18518165.rev.compl.REF        ------------------------------------------------------------ 31039

chimp.LOC112206744-LOC107973052-GGT2.revcompl                                        ctactgagtgagtgggtcaggtggtgcatggctgagggtggcatctgggaagcgttagtt 46789
chimp54546-135457.revcompl.Pan.troglodytes.clone.rp43-41g5.GenBank:AC099533.36       C-AATGGAGGCCTCTCTCCCTTTGTGCCCAGGCCCGGGCCTCTGCAGACAGACCTCCATG 48535
human.FAM30B-LOC105372935-GGT2.NCBI.                                                 ctactgggtgagtgggtcaggtggtgcagggctgagggtggcatctgggaagcattagtt 45781
human.FAM230E-LOC105377182-GGT3P.NCBI.                                               ctactgggtgagtgggtcaggtggtgcagggctgagggtggcatctgggaagcgttagtt 44435
human.FAM230J-LOC105372942-GGTLC5P.NCBI.rev.compl                                    ctactgggtgagtgggtcaggtggtgcagggctgagggtggcatctgggaagcgttagtt 45169
human.FAM230A.ncRNA.18487127.18500594-GGTLC3.18516335.18518165.rev.compl.REF        ------------------------------------------------------------ 31039

chimp.LOC112206744-LOC107973052-GGT2.revcompl                                        tggagtgacagggagtggg----tggccaaggtctctgttcacctggccctctccctttc 46845
chimp54546-135457.revcompl.Pan.troglodytes.clone.rp43-41g5.GenBank:AC099533.36       CCCAGAGGTCAGAAATCAGATATAAGCAGACATCCCAGACCTCCTGGACGAGCTCCT--- 48592
human.FAM30B-LOC105372935-GGT2.NCBI.                                                 tggagtgacagggagtggg----tggccgaggtctctgttcacctggccctctccctttc 45837
human.FAM230E-LOC105377182-GGT3P.NCBI.                                               tggagtgacagggagtggg----tggccgaggtctctgttcacctggccctctccctttc 44491
human.FAM230J-LOC105372942-GGTLC5P.NCBI.rev.compl                                    tggagtgacagggagtggg----tggccgaggtctctgttcacctggccctctccctttc 45225
human.FAM230A.ncRNA.18487127.18500594-GGTLC3.18516335.18518165.rev.compl.REF        ------------------------------------------------------------ 31039

chimp.LOC112206744-LOC107973052-GGT2.revcompl                                        acccatccatctcaccatgcccgagggctgcaatgcccccatgcattcctcatctcagca 46905
chimp54546-135457.revcompl.Pan.troglodytes.clone.rp43-41g5.GenBank:AC099533.36       ---------------------------------------GCACCAACCGAAATGCCATCT 48613
human.FAM30B-LOC105372935-GGT2.NCBI.                                                 acccatccatctcaccatgcccgagggctgcaatgcccccatgcattcctcatctcagca 45897
human.FAM230E-LOC105377182-GGT3P.NCBI.                                               acccatccatctcaccatgcccgagggctgcaatgcccccatgcattcctcatctcagca 44551
human.FAM230J-LOC105372942-GGTLC5P.NCBI.rev.compl                                    acccatccatctcaccatgcccgagggctgcaatgcccccatgcattcctcatctcagca 45285
human.FAM230A.ncRNA.18487127.18500594-GGTLC3.18516335.18518165.rev.compl.REF        ------------------------------------------------------------ 31039

chimp.LOC112206744-LOC107973052-GGT2.revcompl                                        ctgagcactcgagaaccacagcatgtgcaaaggacctgaggtaggagggtggccactaat 46965
chimp54546-135457.revcompl.Pan.troglodytes.clone.rp43-41g5.GenBank:AC099533.36       CCAGCTCCTACAGCTCCACGGGAGGCTTGCCGGGGCTAAAG-CGGAGGAGGGGGCCAGCC 48672
human.FAM30B-LOC105372935-GGT2.NCBI.                                                 ctgagcactcaagaaccacagcatgtgcaaaggacctgaggtaggagggtggccactaat 45957
human.FAM230E-LOC105377182-GGT3P.NCBI.                                               ctgagcactcaagaaccacagcatgtgcaaaggacctgaggtaggagggtggccactaat 44611
human.FAM230J-LOC105372942-GGTLC5P.NCBI.rev.compl                                    ctgagcactcaagaaccacagcatgtgcaaaggacctgaggtaggagggtggccactaat 45345
human.FAM230A.ncRNA.18487127.18500594-GGTLC3.18516335.18518165.rev.compl.REF        ------------------------------------------------------------ 31039

chimp.LOC112206744-LOC107973052-GGT2.revcompl                                        tccccacactgtcagttctgtgcggcagaagccaagactgggggtcacccacgagtccat 47025
chimp54546-135457.revcompl.Pan.troglodytes.clone.rp43-41g5.GenBank:AC099533.36       TCATCCCACTGCCAGCTGACCCTCAGTTCCTCAAAGACAGTGAGTGA------------- 48719
human.FAM30B-LOC105372935-GGT2.NCBI.                                                 tccccgcactgtcagttctgtggggcagaagccaagactgggggtcacccacgagtccat 46017
human.FAM230E-LOC105377182-GGT3P.NCBI.                                               tccccacactgtcagttctgtggggcagaagccaagactgggggtcacccacaagtccat 44671
human.FAM230J-LOC105372942-GGTLC5P.NCBI.rev.compl                                    tccccacactgtcagttctgtggggcagaagccaagactgggggtcacccacgagtccat 45405
human.FAM230A.ncRNA.18487127.18500594-GGTLC3.18516335.18518165.rev.compl.REF        ------------------------------------------------------------ 31039

chimp.LOC112206744-LOC107973052-GGT2.revcompl                                        ccgagcacacagtaggcccgcaatcaaagtctgtggcaggaatggattcataaagcatat 47085
chimp54546-135457.revcompl.Pan.troglodytes.clone.rp43-41g5.GenBank:AC099533.36       ------------------------------------------------------------ 48719
human.FAM30B-LOC105372935-GGT2.NCBI.                                                 ccgagcacacagtaggcccgcaatcgaattctgtggcaggaatggattcataaagcatat 46077
human.FAM230E-LOC105377182-GGT3P.NCBI.                                               ccgagcacacagtaggcccgcaatcaaattctgtggcaggaatggattcataaagcatat 44731
human.FAM230J-LOC105372942-GGTLC5P.NCBI.rev.compl                                    ccgagcacacagtaggcccgcaatcaaattctgtggcaggaatggattcataaagcatat 45465
human.FAM230A.ncRNA.18487127.18500594-GGTLC3.18516335.18518165.rev.compl.REF        ------------------------------------------------------------ 31039

chimp.LOC112206744-LOC107973052-GGT2.revcompl                                        gtgaggctgtagccaccctggggagcccacctgaggcctctacagcaggc-cccacaccc 47144
chimp54546-135457.revcompl.Pan.troglodytes.clone.rp43-41g5.GenBank:AC099533.36       -------------------GGACAGGCCTCAGGCTGTCTCTTCGGGTCACACCCAGTGTG 48760
human.FAM30B-LOC105372935-GGT2.NCBI.                                                 gtgaggctgtagccaccctggggagcccacctgatgcctctacggcaggc-cccacaccc 46136
human.FAM230E-LOC105377182-GGT3P.NCBI.                                               gtgaggctgtagccaccctggggagcccacctgatgcctctacggcaggc-cccacaccc 44790
human.FAM230J-LOC105372942-GGTLC5P.NCBI.rev.compl                                    gtgaggctgtagccaccctggggagcccacctgatgcctctacggcaggc-cccacaccc 45524
human.FAM230A.ncRNA.18487127.18500594-GGTLC3.18516335.18518165.rev.compl.REF        ------------------------------------------------------------ 31039

chimp.LOC112206744-LOC107973052-GGT2.revcompl                                        acagtgggccagcccctgccccttaccccagtcacatccacaaacttggggtcccca-ag 47203
chimp54546-135457.revcompl.Pan.troglodytes.clone.rp43-41g5.GenBank:AC099533.36       AAAAGGCAGCAGATATAGCACCAGGGCAGACACTCGCCCTCAGGAATGACTCCTCCACAT 48820
human.FAM30B-LOC105372935-GGT2.NCBI.                                                 acagtgagccagcccctgccccttacctcagtcacatccacaaacttggggtcccca-ag 46195
human.FAM230E-LOC105377182-GGT3P.NCBI.                                               acagtgggccagcccctgccccttacctcagtcacatccacaaacttggggtcccca-ag 44849
human.FAM230J-LOC105372942-GGTLC5P.NCBI.rev.compl                                    acagtgagccagcccctgccccttacctcagtcacatccacaaacttggggtcccca-ag 45583
human.FAM230A.ncRNA.18487127.18500594-GGTLC3.18516335.18518165.rev.compl.REF        ------------------------------------------------------------ 31039
```

```
chimp.LOC112206744-LOC107973052-GGT2.revcompl                                         cagggtcctcttggcgtaggcaaaccggaaaggcctctacgatgtggtggtacgtcaggc 47263
chimp54546-135457.revcompl.Pan.troglodytes.clone.rp43-41g5.GenBank:AC099533.36        CCGAGGCCTCTAGGCCCAGTACACGCAAGTT-TCCCCTGCTGCCACGCAGGCGAGGGGAG 48879
human.FAM30B-LOC105372935-GGT2.NCBI.                                                  cagggtcctcttggcgtaggcaaaccggaaa-gcctctacgatgcggtggtacgtcaggc 46254
human.FAM230E-LOC105377182-GGT3P.NCBI.                                                cagggtcctcttggcgtaggcaaaccggaaa-gcctctacgatgcggtggtacgtcaggc 44908
human.FAM230J-LOC105372942-GGTLC5P.NCBI.rev.compl                                     cagggtcctcttggcgtaggcaaaccggaaa-gcctctacgatgcggtggtacgtcaggc 45642
human.FAM230A.ncRNA.18487127.18500594-GGTLC3.18516335.18518165.rev.compl.REF          ------------------------------------------------------------ 31039

chimp.LOC112206744-LOC107973052-GGT2.revcompl                                         ccttctgcttggggtctccacgctctcctgggagaagttgtaccctggttgatcagaac 47323
chimp54546-135457.revcompl.Pan.troglodytes.clone.rp43-41g5.GenBank:AC099533.36        CCTTTGATGCTGCCACCTCCCGTAGAGCTGGGGTACCGGGTCACTGTTGAAGACCTGGAC 48939
human.FAM30B-LOC105372935-GGT2.NCBI.                                                  ccttctgctcggggtctccacgctctcccgggagaagttgtaccctggttgatcagagc 46314
human.FAM230E-LOC105377182-GGT3P.NCBI.                                                ccttctgctcggggtctccacgctctcccgggagaagttgtaccctggttgatcagagc 44968
human.FAM230J-LOC105372942-GGTLC5P.NCBI.rev.compl                                     ccttctgctcggggtctccacgctctcccgggagaagttgtaccctggttgatcagagc 45702
human.FAM230A.ncRNA.18487127.18500594-GGTLC3.18516335.18518165.rev.compl.REF          ------------------------------------------------------------ 31039

chimp.LOC112206744-LOC107973052-GGT2.revcompl                                         caggtgcatgttgctgagccccagaggctctgaggggctcagagggtt-aacacctgcct 47382
chimp54546-135457.revcompl.Pan.troglodytes.clone.rp43-41g5.GenBank:AC099533.36        CGGGAGAAGGAGGCGGCCTTCCAGCGCATCAACAGTGCACTGCAGGTTGAGGACAAGGCC 48999
human.FAM30B-LOC105372935-GGT2.NCBI.                                                  caggtgcatgttgctgagccccagaggctctgaggggctcagagggtt-aacacctgcct 46373
human.FAM230E-LOC105377182-GGT3P.NCBI.                                                caggtgcatgttgctgagccccagaggctctgaggggctcagagggtt-aacacctgcct 45027
human.FAM230J-LOC105372942-GGTLC5P.NCBI.rev.compl                                     caggtgcatgttgctgagccccagaggctctgaggggctcagagggtt-aacacctgcct 45761
human.FAM230A.ncRNA.18487127.18500594-GGTLC3.18516335.18518165.rev.compl.REF          ------------------------------------------------------------ 31039

chimp.LOC112206744-LOC107973052-GGT2.revcompl                                         gagccactttgcccacctcaaggagcgtttaataaccaatagcagcagctgcttcagaag 47442
chimp54546-135457.revcompl.Pan.troglodytes.clone.rp43-41g5.GenBank:AC099533.36        ATCTCGGACTGCAGACCCTCACGGCC--TTCCCACACTTTGTCCTCACTTGCAACAGGGG 49057
human.FAM30B-LOC105372935-GGT2.NCBI.                                                  gagccactttgcccacctcaaggagcgtttaataaccaatagcagcagctgcttcagaag 46433
human.FAM230E-LOC105377182-GGT3P.NCBI.                                                gagccactttgcccacctcaaggagcgtttaataatcaatagcagcagctgcttcagaag 45087
human.FAM230J-LOC105372942-GGTLC5P.NCBI.rev.compl                                     gagccactttgcccacctcaaggagcgtttaataaccaatagcagcagctgcttcagaag 45821
human.FAM230A.ncRNA.18487127.18500594-GGTLC3.18516335.18518165.rev.compl.REF          ------------------------------------------------------------ 31039

chimp.LOC112206744-LOC107973052-GGT2.revcompl                                         gctgtggtgagagtgaagtaaggtgagggctcccga-gcctgggcctcacgtcatgcatc 47501
chimp54546-135457.revcompl.Pan.troglodytes.clone.rp43-41g5.GenBank:AC099533.36        CTTCTGGTCTGCCTGCCGTTTCTAAAGCACCCAGTATGGATGCACAGCAGGAGACACACA 49117
human.FAM30B-LOC105372935-GGT2.NCBI.                                                  gctgtggtgagagtgaagtaaggtgagggctcccga-gcctggacctcacgtcacgcatc 46492
human.FAM230E-LOC105377182-GGT3P.NCBI.                                                gctgtggtgagagtgaagtaaggtgagggctcccga-gcctggacctcacgtcacgcatc 45146
human.FAM230J-LOC105372942-GGTLC5P.NCBI.rev.compl                                     gctgtggtgagagtgaagtaaggtgagggctcctga-gcctggacctcacgtcacgcatc 45880
human.FAM230A.ncRNA.18487127.18500594-GGTLC3.18516335.18518165.rev.compl.REF          ------------------------------------------------------------ 31039

chimp.LOC112206744-LOC107973052-GGT2.revcompl                                         agctctggccattcagtgaccaagtggcagggtcacccactgg-accagggtaccctctg 47560
chimp54546-135457.revcompl.Pan.troglodytes.clone.rp43-41g5.GenBank:AC099533.36        AGTCCCAAGACTGCGTGGGCCTAGTGGCCCCCTTAGCATCTGCTGCAGAGGTCCCCTCTA 49177
human.FAM30B-LOC105372935-GGT2.NCBI.                                                  agctctggccattcaatgaccaagtggcagggtcacccactgg-accggggtaccctctg 46551
human.FAM230E-LOC105377182-GGT3P.NCBI.                                                agctctggccattcaatgaccaagtggcagggtcacccactgg-actggggtaccctctg 45205
human.FAM230J-LOC105372942-GGTLC5P.NCBI.rev.compl                                     agctctggccattcaatgaccaagtggcagggtcacccactgg-accggggtaccctctg 45939
human.FAM230A.ncRNA.18487127.18500594-GGTLC3.18516335.18518165.rev.compl.REF          ------------------------------------------------------------ 31039

chimp.LOC112206744-LOC107973052-GGT2.revcompl                                         gaccagccctg----------------------cagccctgagctcctgcacctctctcc 47598
chimp54546-135457.revcompl.Pan.troglodytes.clone.rp43-41g5.GenBank:AC099533.36        CAGCTCCCATGTCTGGGAAGAAGCACAGACCACCAGGCCCCCTGTTCTCCTCCTCAGGTC 49237
human.FAM30B-LOC105372935-GGT2.NCBI.                                                  gaccggccctg----------------------cagccctgagctcctgcacctctctcc 46589
human.FAM230E-LOC105377182-GGT3P.NCBI.                                                gaccggccctg----------------------cagccctgagctcctgcacctctctcc 45243
human.FAM230J-LOC105372942-GGTLC5P.NCBI.rev.compl                                     gaccggccctg----------------------cagccctgagctcctgcacctctctcc 45977
human.FAM230A.ncRNA.18487127.18500594-GGTLC3.18516335.18518165.rev.compl.REF          ------------------------------------------------------------ 31039

chimp.LOC112206744-LOC107973052-GGT2.revcompl                                         ctcctgatgactcctgttcctcctccaacccttgagcattgcccgcttcagccctgtcgc 47658
chimp54546-135457.revcompl.Pan.troglodytes.clone.rp43-41g5.GenBank:AC099533.36        CCCTTCCTGCCACCTCTTCCCATTCCCAGGACTCAGCCCAGGTCACCTC--GCTGATTCC 49295
human.FAM30B-LOC105372935-GGT2.NCBI.                                                  ctcctgatgactcctgttcctcctccaacccttgagcattgcccgcttcagccctgttgc 46649
human.FAM230E-LOC105377182-GGT3P.NCBI.                                                ctcctgatgactcctgttcctcctccaacccttgagcattgcccgcttcagccctgttgc 45303
human.FAM230J-LOC105372942-GGTLC5P.NCBI.rev.compl                                     ctcctgatgactcctgttcctcctccaacccttgagcattgcccgcttcagccctgttgc 46037
human.FAM230A.ncRNA.18487127.18500594-GGTLC3.18516335.18518165.rev.compl.REF          ------------------------------------------------------------ 31039

chimp.LOC112206744-LOC107973052-GGT2.revcompl                                         tctgcactgcctccttcaggacatgg---------------------------------- 47684
chimp54546-135457.revcompl.Pan.troglodytes.clone.rp43-41g5.GenBank:AC099533.36        TGCCCCCTTCCCAGCTGCAAGCATGGATGCGGGCATGAGAAGAACAAGGCCTGGCACTTC 49355
human.FAM30B-LOC105372935-GGT2.NCBI.                                                  tctgcactgcctccttcaggacatgg---------------------------------- 46675
human.FAM230E-LOC105377182-GGT3P.NCBI.                                                tctgcactgcctccttcaggacatgg---------------------------------- 45329
human.FAM230J-LOC105372942-GGTLC5P.NCBI.rev.compl                                     tctgcactgcctccttcaggacatgg---------------------------------- 46063
human.FAM230A.ncRNA.18487127.18500594-GGTLC3.18516335.18518165.rev.compl.REF          ------------------------------------------------------------ 31039

chimp.LOC112206744-LOC107973052-GGT2.revcompl                                         ------------------------------------------------------------ 47684
chimp54546-135457.revcompl.Pan.troglodytes.clone.rp43-41g5.GenBank:AC099533.36        TGCTCCTGCAGCTGCCGCAGCAGCCCCTCCCCCCTCCACATTGAACCCCACGTTGGGGTC 49415
human.FAM30B-LOC105372935-GGT2.NCBI.                                                  ------------------------------------------------------------ 46675
human.FAM230E-LOC105377182-GGT3P.NCBI.                                                ------------------------------------------------------------ 45329
human.FAM230J-LOC105372942-GGTLC5P.NCBI.rev.compl                                     ------------------------------------------------------------ 46063
human.FAM230A.ncRNA.18487127.18500594-GGTLC3.18516335.18518165.rev.compl.REF          ------------------------------------------------------------ 31039

chimp.LOC112206744-LOC107973052-GGT2.revcompl                                         ------------------------------------------------------------ 47684
chimp54546-135457.revcompl.Pan.troglodytes.clone.rp43-41g5.GenBank:AC099533.36        ACTACTGGAGTGGATGGAGGCCCTTCACATTTCTGGGCCTCAGCCACAGCTGCAGCAGGT 49475
human.FAM30B-LOC105372935-GGT2.NCBI.                                                  ------------------------------------------------------------ 46675
human.FAM230E-LOC105377182-GGT3P.NCBI.                                                ------------------------------------------------------------ 45329
human.FAM230J-LOC105372942-GGTLC5P.NCBI.rev.compl                                     ------------------------------------------------------------ 46063
human.FAM230A.ncRNA.18487127.18500594-GGTLC3.18516335.18518165.rev.compl.REF          ------------------------------------------------------------ 31039

chimp.LOC112206744-LOC107973052-GGT2.revcompl                                         -----------tgagctgtgatgcagggacacgcctcgggagctcagtgatggaaagacg 47733
chimp54546-135457.revcompl.Pan.troglodytes.clone.rp43-41g5.GenBank:AC099533.36        GCCCAGAGGTCAGAACCAGAGATCCCAGACCTCCTGGACCAGCTCGTGCCCCAAACGAAA 49535
human.FAM30B-LOC105372935-GGT2.NCBI.                                                  -----------tgagctgtgacgcagggacacacctcaggagctcagtgatggaaagacg 46724
human.FAM230E-LOC105377182-GGT3P.NCBI.                                                -----------tgagctgtgacgcagggacacacctcaggagctcagtgatggaaagacg 45378
human.FAM230J-LOC105372942-GGTLC5P.NCBI.rev.compl                                     -----------tgagctgtgacgcagggacacacctcaggagctcagtgatggaaagacg 46112
human.FAM230A.ncRNA.18487127.18500594-GGTLC3.18516335.18518165.rev.compl.REF          ------------------------------------------------------------ 31039

chimp.LOC112206744-LOC107973052-GGT2.revcompl                                         tggcatgggggcgagcagagatgcagcaggggtggggcgtggggagagagaggcagtgt 47793
chimp54546-135457.revcompl.Pan.troglodytes.clone.rp43-41g5.GenBank:AC099533.36        TGCCATCTCGAGCTCCTGCAGCTCTACGGGAGGCCTCCCGGGACGAAAGCGGAAGAGGGG 49595
human.FAM30B-LOC105372935-GGT2.NCBI.                                                  tggcatgggggcgagcagagatgcaggaggggtggggcgtggggagagagaggcagtgt 46784
human.FAM230E-LOC105377182-GGT3P.NCBI.                                                tggcatgggggcgagcagagatgcaggaggggtggggcgtggggagagagaggcagtgt 45438
human.FAM230J-LOC105372942-GGTLC5P.NCBI.rev.compl                                     tggcatgggggcgagcagagatgcaggaggggtggggcgtggggagagagaggcagtgt 46172
human.FAM230A.ncRNA.18487127.18500594-GGTLC3.18516335.18518165.rev.compl.REF          ------------------------------------------------------------ 31039

chimp.LOC112206744-LOC107973052-GGT2.revcompl                                         catgggtcctaccacacggctgtggtgcgaccacccacctttgaggatgttgaggatgag 47853
chimp54546-135457.revcompl.Pan.troglodytes.clone.rp43-41g5.GenBank:AC099533.36        CCAGCCTCATCCCACTGCCA---GCTGACCCTCAGTTCCTCAAACACAGTGAGTGAGGAC 49652
human.FAM30B-LOC105372935-GGT2.NCBI.                                                  catgggtcctaccacacggctgtggtgcgaccactcacctttgaggatgttgaggatgag 46844
human.FAM230E-LOC105377182-GGT3P.NCBI.                                                catgggtcctaccacacggctgtggtgcgaccactcacctttgaggatgttgaggatgag 45498
human.FAM230J-LOC105372942-GGTLC5P.NCBI.rev.compl                                     catgggtcctaccacacggctgtggtgcgaccactcacctttgaggatgttgaggatgag 46232
human.FAM230A.ncRNA.18487127.18500594-GGTLC3.18516335.18518165.rev.compl.REF          ------------------------------------------------------------ 31039

chimp.LOC112206744-LOC107973052-GGT2.revcompl                                         ggccagcacgggcccgctgagcggcgcact-------------gggcatgtacagcaccg 47900
chimp54546-135457.revcompl.Pan.troglodytes.clone.rp43-41g5.GenBank:AC099533.36        GGACCTCAGGCTGTCTCTTCGGGTCACACCCAGTGTGAAAAGACGGCAGATACAGCACCA 49712
human.FAM30B-LOC105372935-GGT2.NCBI.                                                  ggccagcacgggcccgctgagccgcgcact-------------gggcatgtacagcaccg 46891
human.FAM230E-LOC105377182-GGT3P.NCBI.                                                ggccagcacgggcccgctgagccgcgcact-------------gggcatgtacagcaccg 45545
human.FAM230J-LOC105372942-GGTLC5P.NCBI.rev.compl                                     ggccagcacgggcccgctgagccgcgcact-------------gggcatgtacagcaccg 46279
human.FAM230A.ncRNA.18487127.18500594-GGTLC3.18516335.18518165.rev.compl.REF          ------------------------------------------------------------ 31039

chimp.LOC112206744-LOC107973052-GGT2.revcompl                                         cgtc---------tcccaggctgatgttcagcgggtgctcgatcagctcagcacggtagt 47951
chimp54546-135457.revcompl.Pan.troglodytes.clone.rp43-41g5.GenBank:AC099533.36        GGGCAGACACTCGCCCCCAGGGGTGGCTCCCCCAGATCCAGGCCTCTAGGCCCTGTAGA 49772
human.FAM30B-LOC105372935-GGT2.NCBI.                                                  cgtc---------tcccaggctgatgttcagcgggtgctcgatcagctcagcacggtagt 46942
human.FAM230E-LOC105377182-GGT3P.NCBI.                                                cgtc---------tcccaggctgatgttcagcgggtgctcgatcagctcagcacagtagt 45596
human.FAM230J-LOC105372942-GGTLC5P.NCBI.rev.compl                                     cgtc---------tcccaggctgatgttcagcgggtgctcgatcagctcagcacggtagt 46330
human.FAM230A.ncRNA.18487127.18500594-GGTLC3.18516335.18518165.rev.compl.REF          ------------------------------------------------------------ 31039

chimp.LOC112206744-LOC107973052-GGT2.revcompl                                         tgttcaggtcctcagctgtcacaatgccccctgcaatgggacagcagctcgaatgggcgc 48011
chimp54546-135457.revcompl.Pan.troglodytes.clone.rp43-41g5.GenBank:AC099533.36        TGCAAGTTTCCCCTGCTGCCACG------------------------------------- 49795
human.FAM30B-LOC105372935-GGT2.NCBI.                                                  tgttcaggtcctcagctgtcacaatgccccctgcaatgggacagcagctcgaatgggtgc 47002
human.FAM230E-LOC105377182-GGT3P.NCBI.                                                tgttcaggtcctcagctgtcacaatgccccctgcaatgggacagcagctcgaatgggcgc 45656
human.FAM230J-LOC105372942-GGTLC5P.NCBI.rev.compl                                     tgttcaggtcctcagctgtcacaatgccccctgc-------------------------- 46364
human.FAM230A.ncRNA.18487127.18500594-GGTLC3.18516335.18518165.rev.compl.REF          ------------------------------------------------------------ 31039

chimp.LOC112206744-LOC107973052-GGT2.revcompl                                         tgggatggggctgcaccactgcgtggaggatggagctgcaccagtggtgttgggggcagg 48071
chimp54546-135457.revcompl.Pan.troglodytes.clone.rp43-41g5.GenBank:AC099533.36        ------CAGGCGAGGGGAGCCTTTGATGCTGCCACCTCCCTTAGAGCTGGGGTACCGGGT 49849
human.FAM30B-LOC105372935-GGT2.NCBI.                                                  tgggatggggctgcaccactgcgtggaggatggaactgcaccagtggtgttgggggcagg 47062
human.FAM230E-LOC105377182-GGT3P.NCBI.                                                tgggatggggctgcaccactgcgtggaggatggagctgcaccagtggtgttgggggcagg 45716
human.FAM230J-LOC105372942-GGTLC5P.NCBI.rev.compl                                     ------------------------------------------------------------ 46364
human.FAM230A.ncRNA.18487127.18500594-GGTLC3.18516335.18518165.rev.compl.REF          ------------------------------------------------------------ 31039
```

```
chimp.LOC112206744-LOC107973052-GGT2.revcompl                              catggctgcaccatggtggtggggaaaagcctgtacctaccagggaggacagagtgcact 48131
chimp54546-135457.revcompl.Pan.troglodytes.clone.rp43-41g5.GenBank:AC099533.36    CACTGCTGAAGACCTGGACCCGGAGAAGGAGGCGGCATTCCAGCGCATCAACAGTGCACT 49909
human.FAM30B-LOC105372935-GGT2.NCBI.                                       catggctgcaccatggtggtggggaaaagcctgtacctaccagggaggacagagtgcact 47122
human.FAM230E-LOC105377182-GGT3P.NCBI.                                     catggctgcaccatggtggtggggaaaagcctgtacctaccagggaggacagagtgcact 45776
human.FAM230J-LOC105372942-GGTLC5P.NCBI.rev.compl                          ------------------------------------------------------------ 46364
human.FAM230A.ncRNA.18487127.18500594-GGTLC3.18516335.18518165.rev.compl.REF    ------------------------------------------------------------ 31039

chimp.LOC112206744-LOC107973052-GGT2.revcompl                              actggaggggtgggactgtgccctgggaggggccacaggcaacctcacctc-cttggga 48190
chimp54546-135457.revcompl.Pan.troglodytes.clone.rp43-41g5.GenBank:AC099533.36    GCAGGTTGAGGACAAGGCCATCTCGGACTGCAGACCCTCACGGCCTTCCCACACTTTGTC 49969
human.FAM30B-LOC105372935-GGT2.NCBI.                                       actggaggggtgggactgtgccctgggaggggccacaggcaacctcacctc-cttggga 47181
human.FAM230E-LOC105377182-GGT3P.NCBI.                                     actggaggggtgggactgtgccctgggaggggccacaggcaacctcacctc-cttggga 45835
human.FAM230J-LOC105372942-GGTLC5P.NCBI.rev.compl                          ------------------------------------------------------------ 46364
human.FAM230A.ncRNA.18487127.18500594-GGTLC3.18516335.18518165.rev.compl.REF    ------------------------------------------------------------ 31039

chimp.LOC112206744-LOC107973052-GGT2.revcompl                              acctcaccagctccggcactcctgtctccctgacactgctcaccaccc------------ 48238
chimp54546-135457.revcompl.Pan.troglodytes.clone.rp43-41g5.GenBank:AC099533.36    CTCACTTGCAACAGGGGCTTCTGGTCTGCCTGCCGTTTCTAAAGCACCCAGTATGGATGC 50029
human.FAM30B-LOC105372935-GGT2.NCBI.                                       acctcaccagctccggcactcctgtctccctgacactgctcaccaccc------------ 47229
human.FAM230E-LOC105377182-GGT3P.NCBI.                                     acctcaccagctccggcactcctgtctccctgacactgctcaccaccc------------ 45883
human.FAM230J-LOC105372942-GGTLC5P.NCBI.rev.compl                          ------------------------------------------------------------ 46364
human.FAM230A.ncRNA.18487127.18500594-GGTLC3.18516335.18518165.rev.compl.REF    ------------------------------------------------------------ 31039

chimp.LOC112206744-LOC107973052-GGT2.revcompl                              ------------------------------------------------------------ 48238
chimp54546-135457.revcompl.Pan.troglodytes.clone.rp43-41g5.GenBank:AC099533.36    ACAGCAGGAGACACACAAGTCCCAAGACTGCCTGGGCCTAGTGGCCCCCCTGCATCTGCT 50089
human.FAM30B-LOC105372935-GGT2.NCBI.                                       ------------------------------------------------------------ 47229
human.FAM230E-LOC105377182-GGT3P.NCBI.                                     ------------------------------------------------------------ 45883
human.FAM230J-LOC105372942-GGTLC5P.NCBI.rev.compl                          ------------------------------------------------------------ 46364
human.FAM230A.ncRNA.18487127.18500594-GGTLC3.18516335.18518165.rev.compl.REF    ------------------------------------------------------------ 31039

chimp.LOC112206744-LOC107973052-GGT2.revcompl                              ------------------------------------------------------------ 48238
chimp54546-135457.revcompl.Pan.troglodytes.clone.rp43-41g5.GenBank:AC099533.36    GCACAGGCCTGTAGTCCCAGCTACTTGGGAGGCTGAGGCAGGAGAACGGCATAAACCCGG 50149
human.FAM30B-LOC105372935-GGT2.NCBI.                                       ------------------------------------------------------------ 47229
human.FAM230E-LOC105377182-GGT3P.NCBI.                                     ------------------------------------------------------------ 45883
human.FAM230J-LOC105372942-GGTLC5P.NCBI.rev.compl                          ------------------------------------------------------------ 46364
human.FAM230A.ncRNA.18487127.18500594-GGTLC3.18516335.18518165.rev.compl.REF    ------------------------------------------------------------ 31039

chimp.LOC112206744-LOC107973052-GGT2.revcompl                              ---------------gacagctgggctggggccacctgccctctgcctgcttggcttact 48283
chimp54546-135457.revcompl.Pan.troglodytes.clone.rp43-41g5.GenBank:AC099533.36    GAGGCAGAGCTTGCAGTGAGCTGAGATCGCGCCACTGCACTCCAGCCTGGGTGACAGAGC 50209
human.FAM30B-LOC105372935-GGT2.NCBI.                                       ---------------gacagctgggctggggccacctgccctctgcctgcttggcttact 47274
human.FAM230E-LOC105377182-GGT3P.NCBI.                                     ---------------gacagctgggctggggccacctgccctctgcctgcttggcttact 45928
human.FAM230J-LOC105372942-GGTLC5P.NCBI.rev.compl                          ------------------------------------------------------------ 46364
human.FAM230A.ncRNA.18487127.18500594-GGTLC3.18516335.18518165.rev.compl.REF    ------------------------------------------------------------ 31039

chimp.LOC112206744-LOC107973052-GGT2.revcompl                              ggcttcctgtctg----------------------------------------------- 48296
chimp54546-135457.revcompl.Pan.troglodytes.clone.rp43-41g5.GenBank:AC099533.36    GAGACTCCGTCTCAAAAAAAAAGAAAAAGAAAAAAAGTTCTTGTGACATTTGTGTATGAA 50269
human.FAM30B-LOC105372935-GGT2.NCBI.                                       ggcttcctgtctg----------------------------------------------- 47287
human.FAM230E-LOC105377182-GGT3P.NCBI.                                     ggcttcctgtctg----------------------------------------------- 45941
human.FAM230J-LOC105372942-GGTLC5P.NCBI.rev.compl                          ------------------------------------------------------------ 46364
human.FAM230A.ncRNA.18487127.18500594-GGTLC3.18516335.18518165.rev.compl.REF    ------------------------------------------------------------ 31039

chimp.LOC112206744-LOC107973052-GGT2.revcompl                              -------------------------ccttctctcatctgtggccggagagtg-------- 48323
chimp54546-135457.revcompl.Pan.troglodytes.clone.rp43-41g5.GenBank:AC099533.36    ATCAGCCTTCACTACATGGATAGGACCAGCACGCTTCTGCGGCACGACTCTGCCATCTTA 50329
human.FAM30B-LOC105372935-GGT2.NCBI.                                       -------------------------ccttctctcatctgtggccagagagtg-------- 47314
human.FAM230E-LOC105377182-GGT3P.NCBI.                                     -------------------------ccttctctcatctgtggccagagagtg-------- 45968
human.FAM230J-LOC105372942-GGTLC5P.NCBI.rev.compl                          ------------------------------------------------------------ 46364
human.FAM230A.ncRNA.18487127.18500594-GGTLC3.18516335.18518165.rev.compl.REF    ------------------------------------------------------------ 31039

chimp.LOC112206744-LOC107973052-GGT2.revcompl                              -----------tttcctttttctttttttttttttagagatacgatcttgctctgttacc 48372
chimp54546-135457.revcompl.Pan.troglodytes.clone.rp43-41g5.GenBank:AC099533.36    CTACATTTTTTTATTTGTATTTTTTTTATTCCTTTTGAGACAGTCTCACTCTGTCACC 50389
human.FAM30B-LOC105372935-GGT2.NCBI.                                       -----------ttttcttttttcttttttttttttagagatacgatcttgctctgttacc 47363
human.FAM230E-LOC105377182-GGT3P.NCBI.                                     -----------ttttcttttttc-tttttttttttagagatacgatcttgctctgttacc 46016
human.FAM230J-LOC105372942-GGTLC5P.NCBI.rev.compl                          ------------------------------------------------------------ 46364
human.FAM230A.ncRNA.18487127.18500594-GGTLC3.18516335.18518165.rev.compl.REF    ------------------------------------------------------------ 31039

chimp.LOC112206744-LOC107973052-GGT2.revcompl                              caggctggagtgcag--------tggcttaatcacagccttgaactcccgggctcaagtg 48424
chimp54546-135457.revcompl.Pan.troglodytes.clone.rp43-41g5.GenBank:AC099533.36    CAGGCTGAAGTGCAGCCGAGATCTCGGCTCACTGCAACCTCCACCTCTTGGGTTCAAGCA 50449
human.FAM30B-LOC105372935-GGT2.NCBI.                                       caggctggagtgcag--------tggcttaatcacagccttgaacccctgggctcaagtg 47415
human.FAM230E-LOC105377182-GGT3P.NCBI.                                     caggctggagtgcag--------tggcttaatcacagccttgaacccctgggctcaagtg 46068
human.FAM230J-LOC105372942-GGTLC5P.NCBI.rev.compl                          ------------------------------------------------------------ 46364
human.FAM230A.ncRNA.18487127.18500594-GGTLC3.18516335.18518165.rev.compl.REF    ------------------------------------------------------------ 31039

chimp.LOC112206744-LOC107973052-GGT2.revcompl                              atcctccag-cagaccctccccagtagctgagactaaaggcacaact----acacccagc 48479
chimp54546-135457.revcompl.Pan.troglodytes.clone.rp43-41g5.GenBank:AC099533.36    ATTCTCCTGTCTCAGCCTCCCAAGTAGCTGGGACTACAGGCACACATCAAAACGCCTGGC 50509
human.FAM30B-LOC105372935-GGT2.NCBI.                                       atcctccag-cagaccctccccagtagctgagactaaaggcacaact----acacccagc 47470
human.FAM230E-LOC105377182-GGT3P.NCBI.                                     atcctccag-cagaccctccccagtagctgagactaaaggcacaact----acacccagc 46123
human.FAM230J-LOC105372942-GGTLC5P.NCBI.rev.compl                          ------------------------------------------------------------ 46364
human.FAM230A.ncRNA.18487127.18500594-GGTLC3.18516335.18518165.rev.compl.REF    ------------------------------------------------------------ 31039

chimp.LOC112206744-LOC107973052-GGT2.revcompl                              aaattttaattttttt-------gttgtgttttgctatatttct---------------- 48516
chimp54546-135457.revcompl.Pan.troglodytes.clone.rp43-41g5.GenBank:AC099533.36    TAATTTTTGTATTTTTAGTAGAGATGGAGTTTTGCCATATTGGTCAGGCTGGTCTCGAAC 50569
human.FAM30B-LOC105372935-GGT2.NCBI.                                       aaattttaattttttt-------gttgtgttttgctatatttct---------------- 47507
human.FAM230E-LOC105377182-GGT3P.NCBI.                                     aaattttaattttttt-------gttgtgttttgctatatttct---------------- 46160
human.FAM230J-LOC105372942-GGTLC5P.NCBI.rev.compl                          ------------------------------------------------------------ 46364
human.FAM230A.ncRNA.18487127.18500594-GGTLC3.18516335.18518165.rev.compl.REF    ------------------------------------------------------------ 31039

chimp.LOC112206744-LOC107973052-GGT2.revcompl                              ------------------------------------------------------------ 48516
chimp54546-135457.revcompl.Pan.troglodytes.clone.rp43-41g5.GenBank:AC099533.36    TCCTGACCTCAGGTGATCGACCTGTCTTAGCCTCCCGAACTGCTAGGATTACAGGTGTAC 50629
human.FAM30B-LOC105372935-GGT2.NCBI.                                       ------------------------------------------------------------ 47507
human.FAM230E-LOC105377182-GGT3P.NCBI.                                     ------------------------------------------------------------ 46160
human.FAM230J-LOC105372942-GGTLC5P.NCBI.rev.compl                          ------------------------------------------------------------ 46364
human.FAM230A.ncRNA.18487127.18500594-GGTLC3.18516335.18518165.rev.compl.REF    ------------------------------------------------------------ 31039

chimp.LOC112206744-LOC107973052-GGT2.revcompl                              --------------------------ttctttctttctttcttttttttttgtgagacgg 48550
chimp54546-135457.revcompl.Pan.troglodytes.clone.rp43-41g5.GenBank:AC099533.36    GTTTATTTATTTGAGATGGAATCTTGCTCTGTATTT-ATTAATTTTTTATCTGAGATGG 50688
human.FAM30B-LOC105372935-GGT2.NCBI.                                       --------------------------ttctttcttt-ttttttttttttttgtgagacgg 47540
human.FAM230E-LOC105377182-GGT3P.NCBI.                                     --------------------------ttctttc-tt-ttttttttttttttgtgagacgg 46192
human.FAM230J-LOC105372942-GGTLC5P.NCBI.rev.compl                          ------------------------------------------------------------ 46364
human.FAM230A.ncRNA.18487127.18500594-GGTLC3.18516335.18518165.rev.compl.REF    ------------------------------------------------------------ 31039

chimp.LOC112206744-LOC107973052-GGT2.revcompl                              agtctcactctatcgcccaggctggagtgcagtggcgcgatatcggctcactgcaagctc 48610
chimp54546-135457.revcompl.Pan.troglodytes.clone.rp43-41g5.GenBank:AC099533.36    AGTCTTTCTCCATCGCCCAGGCTAGAGTGCAGTGGTGCAATCTCGGCTCCCTGCAACCTC 50748
human.FAM30B-LOC105372935-GGT2.NCBI.                                       agtctcgctctatcgcccaggctggagtgcagtggcgcgatatcggctcactgcaagctc 47600
human.FAM230E-LOC105377182-GGT3P.NCBI.                                     agtctcgctctatcgcccaggctggagtgcagtggcgcgatatcggctcactgcaagctc 46252
human.FAM230J-LOC105372942-GGTLC5P.NCBI.rev.compl                          ------------------------------------------------------------ 46364
human.FAM230A.ncRNA.18487127.18500594-GGTLC3.18516335.18518165.rev.compl.REF    ------------------------------------------------------------ 31039

chimp.LOC112206744-LOC107973052-GGT2.revcompl                              cgcctcccaggttcacgccattctcctgcctcagcctcccaagtagctgggactgcaggc 48670
chimp54546-135457.revcompl.Pan.troglodytes.clone.rp43-41g5.GenBank:AC099533.36    TGACTTCCAGTTTCAAGTGATTCTCCTGCCTCAGTGTCCCAAGTAGCTGGGATTACAGGT 50808
human.FAM30B-LOC105372935-GGT2.NCBI.                                       cgcctcccaggttcacgccattctcctgcctcagcctcccaagtagctgggactgcaggc 47660
human.FAM230E-LOC105377182-GGT3P.NCBI.                                     cgcctcccaggttcacgccattctcctgcctcagcctcccaagtagctgggactgcaggc 46312
human.FAM230J-LOC105372942-GGTLC5P.NCBI.rev.compl                          ------------------------------------------------------------ 46364
human.FAM230A.ncRNA.18487127.18500594-GGTLC3.18516335.18518165.rev.compl.REF    ------------------------------------------------------------ 31039

chimp.LOC112206744-LOC107973052-GGT2.revcompl                              acccgccaccacacctggctaatttttttaaatttttagtagagacag-gtttcactgtg 48729
chimp54546-135457.revcompl.Pan.troglodytes.clone.rp43-41g5.GenBank:AC099533.36    GCCTGCCACCACAGCTGGCTAATTTT--TGTATTTTTAGTAGAGACAGTGTTTCACCATC 50866
human.FAM30B-LOC105372935-GGT2.NCBI.                                       acccgccaccacatgtggctaatttttctgtatttttagtagagacag-gtttcactgtg 47719
human.FAM230E-LOC105377182-GGT3P.NCBI.                                     acccgccaccacatgtggctaatttttctgtatttttagtagagacag-gtttcactgtg 46371
human.FAM230J-LOC105372942-GGTLC5P.NCBI.rev.compl                          ------------------------------------------------------------ 46364
human.FAM230A.ncRNA.18487127.18500594-GGTLC3.18516335.18518165.rev.compl.REF    ------------------------------------------------------------ 31039

chimp.LOC112206744-LOC107973052-GGT2.revcompl                              ttagccaggatggtctcgatctcctgaccctgtgatccacacgcctcggcctctcaaagt 48789
chimp54546-135457.revcompl.Pan.troglodytes.clone.rp43-41g5.GenBank:AC099533.36    TTGGCCAGGCTGGTCTCGGGCTCCTGACCTCATGAACCACCTGCCTCAGCCTCCCAAAGT 50926
human.FAM30B-LOC105372935-GGT2.NCBI.                                       ttagccaggatggtctcgatctcctgaccccgtgatccacacgcctcggcctctcaaagt 47779
human.FAM230E-LOC105377182-GGT3P.NCBI.                                     ttagccaggatggtctcgatctcctgaccccgtgatccacacgcctcggcctctcaaagt 46431
human.FAM230J-LOC105372942-GGTLC5P.NCBI.rev.compl                          ------------------------------------------------------------ 46364
human.FAM230A.ncRNA.18487127.18500594-GGTLC3.18516335.18518165.rev.compl.REF    ------------------------------------------------------------ 31039
```

```
chimp.LOC112206744-LOC107973052-GGT2.revcompl                                    gctgggattacaggcgtgaggcaccatgcccggccgggttttgctatatttcttttcact 48849
chimp54546-135457.revcompl.Pan.troglodytes.clone.rp43-41g5.GenBank:AC099533.36   GTTGGGATTACAGGCCTAAGGCACCATGCTCGGCCATATTTATTTAT------------- 50973
human.FAM30B-LOC105372935-GGT2.NCBI.                                             gctgagattacaggcgtgaggcaccatgcccggccgggttttgctatatttcttttcact 47839
human.FAM230E-LOC105377182-GGT3P.NCBI.                                           gctgagattacaggcgtgaggcaccatgcccggccgggttttgctatatttcttttcact 46491
human.FAM230J-LOC105372942-GGTLC5P.NCBI.rev.compl                                ------------------------------------------------------------ 46364
human.FAM230A.ncRNA.18487127.18500594-GGTLC3.18516335.18518165.rev.compl.REF     ------------------------------------------------------------ 31039

chimp.LOC112206744-LOC107973052-GGT2.revcompl                                    atgctttgaatttttttgttttcttgttccccacccccacccccactatatttatgtagat 48909
chimp54546-135457.revcompl.Pan.troglodytes.clone.rp43-41g5.GenBank:AC099533.36   ------------------------------------------------------------ 50973
human.FAM30B-LOC105372935-GGT2.NCBI.                                             atgctttgaatttttttgttttcttgttccccacccccacccccactatatttatgtagat 47899
human.FAM230E-LOC105377182-GGT3P.NCBI.                                           atgctttgaatttttttgttttcttgttccccacccccacccccactatatttatgtagat 46551
human.FAM230J-LOC105372942-GGTLC5P.NCBI.rev.compl                                ------------------------------------------------------------ 46364
human.FAM230A.ncRNA.18487127.18500594-GGTLC3.18516335.18518165.rev.compl.REF     ------------------------------------------------------------ 31039

chimp.LOC112206744-LOC107973052-GGT2.revcompl                                    tctcaattttttttttgtagactcactatgttgcccaggcttgtcttggacctcctgacc 48969
chimp54546-135457.revcompl.Pan.troglodytes.clone.rp43-41g5.GenBank:AC099533.36   ------------------------------------------------------------ 50973
human.FAM30B-LOC105372935-GGT2.NCBI.                                             tctcaatattttttttgtagactcactatgttgcccaggcttgtcttggacccccctgacc 47959
human.FAM230E-LOC105377182-GGT3P.NCBI.                                           tctcaatattttttttgtagactcactatgttgcccaggcttgtcttggacctcctgacc 46611
human.FAM230J-LOC105372942-GGTLC5P.NCBI.rev.compl                                ------------------------------------------------------------ 46364
human.FAM230A.ncRNA.18487127.18500594-GGTLC3.18516335.18518165.rev.compl.REF     ------------------------------------------------------------ 31039

chimp.LOC112206744-LOC107973052-GGT2.revcompl                                    tcaacttctacctcagcctcccaaagtgttgggattacaggcatgagacaccaggctttg 49029
chimp54546-135457.revcompl.Pan.troglodytes.clone.rp43-41g5.GenBank:AC099533.36   ------------------------------------------------------------ 50973
human.FAM30B-LOC105372935-GGT2.NCBI.                                             tcaacttctacctcagcctcccaaagtgttgggattacaggcatgagacaccaagcttgg 48019
human.FAM230E-LOC105377182-GGT3P.NCBI.                                           tcaacttctacctcaggctcccaaagtgttgggattacaggcatgagacaccaagcttgg 46671
human.FAM230J-LOC105372942-GGTLC5P.NCBI.rev.compl                                ------------------------------------------------------------ 46364
human.FAM230A.ncRNA.18487127.18500594-GGTLC3.18516335.18518165.rev.compl.REF     ------------------------------------------------------------ 31039

chimp.LOC112206744-LOC107973052-GGT2.revcompl                                    cctcagacggcctttctttcttttttttttttttgagatggagtctcactctg------- 49082
chimp54546-135457.revcompl.Pan.troglodytes.clone.rp43-41g5.GenBank:AC099533.36   -----------------------TTAATTATTTAGAGACAAAGTCTTGCTCTGTCACCCA 51010
human.FAM30B-LOC105372935-GGT2.NCBI.                                             cctcagagggcctttctctc--ttttttttttttgagatggagtctcactctg------- 48070
human.FAM230E-LOC105377182-GGT3P.NCBI.                                           cctcagagggcctttctctc-tttttttttttttgagatggagtctcactctg------- 46723
human.FAM230J-LOC105372942-GGTLC5P.NCBI.rev.compl                                ------------------------------------------------------------ 46364
human.FAM230A.ncRNA.18487127.18500594-GGTLC3.18516335.18518165.rev.compl.REF     ------------------------------------------------------------ 31039

chimp.LOC112206744-LOC107973052-GGT2.revcompl                                    ---tggagtgcagtggtgcaatctcggctcactgcaacctcagcccccaggttctagcg 49139
chimp54546-135457.revcompl.Pan.troglodytes.clone.rp43-41g5.GenBank:AC099533.36   GGCTGGAGTGCAGTGGCGCCATCTCAGCTCACTGCAGCCTCCGCCTCCAAGGTTTAAGCG 51070
human.FAM30B-LOC105372935-GGT2.NCBI.                                             ---tggagtgcagtggtgcaatctcggctcactgcaacctcagccccacaggttctagcg 48127
human.FAM230E-LOC105377182-GGT3P.NCBI.                                           ---tggagtgcagtggtgcaatctcggctcactgcaacctcagccccacaggttctagcg 46780
human.FAM230J-LOC105372942-GGTLC5P.NCBI.rev.compl                                ------------------------------------------------------------ 46364
human.FAM230A.ncRNA.18487127.18500594-GGTLC3.18516335.18518165.rev.compl.REF     ------------------------------------------------------------ 31039

chimp.LOC112206744-LOC107973052-GGT2.revcompl                                    attctcctgccacagcctcccacgtagctgggattacaggcacaacccaccatgcctgac 49199
chimp54546-135457.revcompl.Pan.troglodytes.clone.rp43-41g5.GenBank:AC099533.36   ATTCTCATGCCTCAGCCTCCTGACTAACTGGGACTACAGGTACTCACCACCATGCAGGGA 51130
human.FAM30B-LOC105372935-GGT2.NCBI.                                             attctcctgccacagcctcccaagtagctgggattacaggcacaacccaccatgcctgac 48187
human.FAM230E-LOC105377182-GGT3P.NCBI.                                           attctcctgccacagcctcccaagtagctgggattacaggcacaacccaccatgcctgac 46840
human.FAM230J-LOC105372942-GGTLC5P.NCBI.rev.compl                                ------------------------------------------------------------ 46364
human.FAM230A.ncRNA.18487127.18500594-GGTLC3.18516335.18518165.rev.compl.REF     ------------------------------------------------------------ 31039

chimp.LOC112206744-LOC107973052-GGT2.revcompl                                    ta--atttgcatttttagtagagacagggtttcaccatgttggccaggctggtcttgaa 49257
chimp54546-135457.revcompl.Pan.troglodytes.clone.rp43-41g5.GenBank:AC099533.36   TATTTTTTTCTATTTTTTTTACAGACACAGTTTCACCATGTTGGCCAGGCTGGTCTCGAA 51190
human.FAM30B-LOC105372935-GGT2.NCBI.                                             ta--atttgcatttttagtagagacagggtttcaccatgttggccaggctggtcttgaa 48245
human.FAM230E-LOC105377182-GGT3P.NCBI.                                           ta--atttgcatttttagtagagacagggtttcaccatgttggccaggctggtcttgaa 46898
human.FAM230J-LOC105372942-GGTLC5P.NCBI.rev.compl                                ------------------------------------------------------------ 46364
human.FAM230A.ncRNA.18487127.18500594-GGTLC3.18516335.18518165.rev.compl.REF     ------------------------------------------------------------ 31039

chimp.LOC112206744-LOC107973052-GGT2.revcompl                                    ctcctgagctcaggtgttccacccgcctcggcctcccaaagtgctgggattacagacgtg 49317
chimp54546-135457.revcompl.Pan.troglodytes.clone.rp43-41g5.GenBank:AC099533.36   CTCCTGACCTTAGGTGATCTGACAGCCTCGTCCCCTCAAAGTACTGGGATTACAGGCATG 51250
human.FAM30B-LOC105372935-GGT2.NCBI.                                             ctcctgacctcaggtgatccacccgcctcggcctcccaaagtgctgggattacagacgtg 48305
human.FAM230E-LOC105377182-GGT3P.NCBI.                                           ctcctgacctcaggtgatccacccgcctcggcctcccaaagtgctgggattacagacgtg 46958
human.FAM230J-LOC105372942-GGTLC5P.NCBI.rev.compl                                ------------------------------------------------------------ 46364
human.FAM230A.ncRNA.18487127.18500594-GGTLC3.18516335.18518165.rev.compl.REF     ------------------------------------------------------------ 31039

chimp.LOC112206744-LOC107973052-GGT2.revcompl                                    agccactgcgcctggctcagagggcctttt--ctaactggagaattcctgccggtgtccc 49375
chimp54546-135457.revcompl.Pan.troglodytes.clone.rp43-41g5.GenBank:AC099533.36   AGCCACTGTGCCCGGCCTCTCACTACATTTAAGTGACGCCATGGCTCATGCCTGTAATCC 51310
human.FAM30B-LOC105372935-GGT2.NCBI.                                             agccaccgcacctggctcagagggcctttt--ctaactggagaattcctgccggtgtccc 48363
human.FAM230E-LOC105377182-GGT3P.NCBI.                                           agccaccgcacctggctcagagggcctttt--ctaactggagaattcctgccggtgtccc 47016
human.FAM230J-LOC105372942-GGTLC5P.NCBI.rev.compl                                ------------------------------------------------------------ 46364
human.FAM230A.ncRNA.18487127.18500594-GGTLC3.18516335.18518165.rev.compl.REF     ------------------------------------------------------------ 31039

chimp.LOC112206744-LOC107973052-GGT2.revcompl                                    tgctgct----------------------------------------------------- 49382
chimp54546-135457.revcompl.Pan.troglodytes.clone.rp43-41g5.GenBank:AC099533.36   TAGCACTTTGGGAGGCCAAGGCAGGTGGATCACTTCAGGTCAGAAGTTCGACACGAGCCT 51370
human.FAM30B-LOC105372935-GGT2.NCBI.                                             tgctgct----------------------------------------------------- 48370
human.FAM230E-LOC105377182-GGT3P.NCBI.                                           tgctgct----------------------------------------------------- 47023
human.FAM230J-LOC105372942-GGTLC5P.NCBI.rev.compl                                ------------------------------------------------------------ 46364
human.FAM230A.ncRNA.18487127.18500594-GGTLC3.18516335.18518165.rev.compl.REF     ------------------------------------------------------------ 31039

chimp.LOC112206744-LOC107973052-GGT2.revcompl                                    ------------------------------------------------------------ 49382
chimp54546-135457.revcompl.Pan.troglodytes.clone.rp43-41g5.GenBank:AC099533.36   GGCCAACATGGGGAAACCCCGTCTCTAGTAAAAATGCACAAATTAGTCAGGCGTGGTGGT 51430
human.FAM30B-LOC105372935-GGT2.NCBI.                                             ------------------------------------------------------------ 48370
human.FAM230E-LOC105377182-GGT3P.NCBI.                                           ------------------------------------------------------------ 47023
human.FAM230J-LOC105372942-GGTLC5P.NCBI.rev.compl                                ------------------------------------------------------------ 46364
human.FAM230A.ncRNA.18487127.18500594-GGTLC3.18516335.18518165.rev.compl.REF     ------------------------------------------------------------ 31039

chimp.LOC112206744-LOC107973052-GGT2.revcompl                                    ------------------------------------------------------------ 49382
chimp54546-135457.revcompl.Pan.troglodytes.clone.rp43-41g5.GenBank:AC099533.36   ACAAGCCTGTAGGCCCAGCTACTTGGAAGGCTGAGGCAGGAGAATCACTTTAACCGGGAG 51490
human.FAM30B-LOC105372935-GGT2.NCBI.                                             ------------------------------------------------------------ 48370
human.FAM230E-LOC105377182-GGT3P.NCBI.                                           ------------------------------------------------------------ 47023
human.FAM230J-LOC105372942-GGTLC5P.NCBI.rev.compl                                ------------------------------------------------------------ 46364
human.FAM230A.ncRNA.18487127.18500594-GGTLC3.18516335.18518165.rev.compl.REF     ------------------------------------------------------------ 31039

chimp.LOC112206744-LOC107973052-GGT2.revcompl                                    ------------------------------------------------------------ 49382
chimp54546-135457.revcompl.Pan.troglodytes.clone.rp43-41g5.GenBank:AC099533.36   GCAGAGGTTGCAGTGAGCCAATATCATGCCACTGCACTCCAGCTTGGGTGACAGAGTGAG 51550
human.FAM30B-LOC105372935-GGT2.NCBI.                                             ------------------------------------------------------------ 48370
human.FAM230E-LOC105377182-GGT3P.NCBI.                                           ------------------------------------------------------------ 47023
human.FAM230J-LOC105372942-GGTLC5P.NCBI.rev.compl                                ------------------------------------------------------------ 46364
human.FAM230A.ncRNA.18487127.18500594-GGTLC3.18516335.18518165.rev.compl.REF     ------------------------------------------------------------ 31039

chimp.LOC112206744-LOC107973052-GGT2.revcompl                                    ------------------------------------------------------------ 49382
chimp54546-135457.revcompl.Pan.troglodytes.clone.rp43-41g5.GenBank:AC099533.36   ACACCGTCTCAAAAAAAAAAAAAAAAAAGGAAAGAAAAACATATGATGCCGGGGCATCTC 51610
human.FAM30B-LOC105372935-GGT2.NCBI.                                             ------------------------------------------------------------ 48370
human.FAM230E-LOC105377182-GGT3P.NCBI.                                           ------------------------------------------------------------ 47023
human.FAM230J-LOC105372942-GGTLC5P.NCBI.rev.compl                                ------------------------------------------------------------ 46364
human.FAM230A.ncRNA.18487127.18500594-GGTLC3.18516335.18518165.rev.compl.REF     ------------------------------------------------------------ 31039

chimp.LOC112206744-LOC107973052-GGT2.revcompl                                    ---------------------------------------------------------tgg 49385
chimp54546-135457.revcompl.Pan.troglodytes.clone.rp43-41g5.GenBank:AC099533.36   GGCCTCAATACCTGCGTGAGCACAGTGATGTCCAGGCCAGGGCTGCTGGTCGAGGTCTGG 51670
human.FAM30B-LOC105372935-GGT2.NCBI.                                             ---------------------------------------------------------tgg 48373
human.FAM230E-LOC105377182-GGT3P.NCBI.                                           ---------------------------------------------------------tgg 47026
human.FAM230J-LOC105372942-GGTLC5P.NCBI.rev.compl                                ------------------------------------------------------------ 46364
human.FAM230A.ncRNA.18487127.18500594-GGTLC3.18516335.18518165.rev.compl.REF     ------------------------------------------------------------ 31039

chimp.LOC112206744-LOC107973052-GGT2.revcompl                                    cctcttctcctcatgaggaatggatagagggagggagggaggct---------------- 49429
chimp54546-135457.revcompl.Pan.troglodytes.clone.rp43-41g5.GenBank:AC099533.36   CCCCATCTCTTCCAGCAGAAAGGGAGTAAGCTTGCAGGGAGGCTGGGGGACAAGATCCCA 51730
human.FAM30B-LOC105372935-GGT2.NCBI.                                             cctcttctcctcacgatgaatggatagagggagggagggaggct---------------- 48417
human.FAM230E-LOC105377182-GGT3P.NCBI.                                           cctcttctcctcacgatgaatggatagagggagggagggaggct---------------- 47070
human.FAM230J-LOC105372942-GGTLC5P.NCBI.rev.compl                                ------------------------------------------------------------ 46364
human.FAM230A.ncRNA.18487127.18500594-GGTLC3.18516335.18518165.rev.compl.REF     ------------------------------------------------------------ 31039

chimp.LOC112206744-LOC107973052-GGT2.revcompl                                    ------------------------------------------------------------ 49429
chimp54546-135457.revcompl.Pan.troglodytes.clone.rp43-41g5.GenBank:AC099533.36   GGATCTCAGCCTCTGCTCATGGATCAGCTCTGAGACCCCGAGTGAGCTGGGGGTGCTCTG 51790
human.FAM30B-LOC105372935-GGT2.NCBI.                                             ------------------------------------------------------------ 48417
human.FAM230E-LOC105377182-GGT3P.NCBI.                                           ------------------------------------------------------------ 47070
human.FAM230J-LOC105372942-GGTLC5P.NCBI.rev.compl                                ------------------------------------------------------------ 46364
human.FAM230A.ncRNA.18487127.18500594-GGTLC3.18516335.18518165.rev.compl.REF     ------------------------------------------------------------ 31039

chimp.LOC112206744-LOC107973052-GGT2.revcompl                                    ------------------------------------------------------------ 49429
```

```
chimp54546-135457.revcompl.Pan.troglodytes.clone.rp43-41g5.GenBank:AC099533.36      TGCGCATTGGTTTCCCCAGCTGTCAAGTAAAGGGATTGGATGAGGAAGTCTTGTCAAGGT 51850
human.FAM30B-LOC105372935-GGT2.NCBI.                                                 ------------------------------------------------------------ 48417
human.FAM230E-LOC105377182-GGT3P.NCBI.                                               ------------------------------------------------------------ 47070
human.FAM230J-LOC105372942-GGTLC5P.NCBI.rev.compl                                    ------------------------------------------------------------ 46364
human.FAM230A.ncRNA.18487127.18500594-GGTLC3.18516335.18518165.rev.compl.REF         ------------------------------------------------------------ 31039

chimp.LOC112206744-LOC107973052-GGT2.revcompl                                        ------------------------------------------------------------ 49429
chimp54546-135457.revcompl.Pan.troglodytes.clone.rp43-41g5.GenBank:AC099533.36      GGAATGATCTCAGATTTGGGGCAGCAGTGAATGATCCCGCTCCCTGGGCCATGCCAGTGG 51910
human.FAM30B-LOC105372935-GGT2.NCBI.                                                 ------------------------------------------------------------ 48417
human.FAM230E-LOC105377182-GGT3P.NCBI.                                               ------------------------------------------------------------ 47070
human.FAM230J-LOC105372942-GGTLC5P.NCBI.rev.compl                                    ------------------------------------------------------------ 46364
human.FAM230A.ncRNA.18487127.18500594-GGTLC3.18516335.18518165.rev.compl.REF         ------------------------------------------------------------ 31039

chimp.LOC112206744-LOC107973052-GGT2.revcompl                                        ------------------------------------------------------------ 49429
chimp54546-135457.revcompl.Pan.troglodytes.clone.rp43-41g5.GenBank:AC099533.36      CCCGGCCTCGGCTCAACACAGCCCCAACACTCTGGAATGGGGATGAGGGGGCAGTCAGCT 51970
human.FAM30B-LOC105372935-GGT2.NCBI.                                                 ------------------------------------------------------------ 48417
human.FAM230E-LOC105377182-GGT3P.NCBI.                                               ------------------------------------------------------------ 47070
human.FAM230J-LOC105372942-GGTLC5P.NCBI.rev.compl                                    ------------------------------------------------------------ 46364
human.FAM230A.ncRNA.18487127.18500594-GGTLC3.18516335.18518165.rev.compl.REF         ------------------------------------------------------------ 31039

chimp.LOC112206744-LOC107973052-GGT2.revcompl                                        ------------------------------------------------------------ 49429
chimp54546-135457.revcompl.Pan.troglodytes.clone.rp43-41g5.GenBank:AC099533.36      CTTGCTCCTAGTAAGAGAGATGCAATAGGGCTCTGTGGCTGAGCTGGGTGCCTTGCCTCA 52030
human.FAM30B-LOC105372935-GGT2.NCBI.                                                 ------------------------------------------------------------ 48417
human.FAM230E-LOC105377182-GGT3P.NCBI.                                               ------------------------------------------------------------ 47070
human.FAM230J-LOC105372942-GGTLC5P.NCBI.rev.compl                                    ------------------------------------------------------------ 46364
human.FAM230A.ncRNA.18487127.18500594-GGTLC3.18516335.18518165.rev.compl.REF         ------------------------------------------------------------ 31039

chimp.LOC112206744-LOC107973052-GGT2.revcompl                                        ------------------------------------------------------------ 49429
chimp54546-135457.revcompl.Pan.troglodytes.clone.rp43-41g5.GenBank:AC099533.36      CACCTGTAATCCCAACCTTTGAGAGGCCGAGGCGGGAGGATTGCTTGAGGCCGGGAATTT 52090
human.FAM30B-LOC105372935-GGT2.NCBI.                                                 ------------------------------------------------------------ 48417
human.FAM230E-LOC105377182-GGT3P.NCBI.                                               ------------------------------------------------------------ 47070
human.FAM230J-LOC105372942-GGTLC5P.NCBI.rev.compl                                    ------------------------------------------------------------ 46364
human.FAM230A.ncRNA.18487127.18500594-GGTLC3.18516335.18518165.rev.compl.REF         ------------------------------------------------------------ 31039

chimp.LOC112206744-LOC107973052-GGT2.revcompl                                        ------------------------------------------------------------ 49429
chimp54546-135457.revcompl.Pan.troglodytes.clone.rp43-41g5.GenBank:AC099533.36      TGAGAATAGCCTGGACAACATAGCCAGACCCCATGTCTGCAAAATAATAAGAAAACACAC 52150
human.FAM30B-LOC105372935-GGT2.NCBI.                                                 ------------------------------------------------------------ 48417
human.FAM230E-LOC105377182-GGT3P.NCBI.                                               ------------------------------------------------------------ 47070
human.FAM230J-LOC105372942-GGTLC5P.NCBI.rev.compl                                    ------------------------------------------------------------ 46364
human.FAM230A.ncRNA.18487127.18500594-GGTLC3.18516335.18518165.rev.compl.REF         ------------------------------------------------------------ 31039

chimp.LOC112206744-LOC107973052-GGT2.revcompl                                        ------------------------------------------------------------ 49429
chimp54546-135457.revcompl.Pan.troglodytes.clone.rp43-41g5.GenBank:AC099533.36      AGCTATAGTCCAAGCTACTTGGCAGGCTGAGGCAGGAAGGTCCCTTGAGTCCGGGAATTG 52210
human.FAM30B-LOC105372935-GGT2.NCBI.                                                 ------------------------------------------------------------ 48417
human.FAM230E-LOC105377182-GGT3P.NCBI.                                               ------------------------------------------------------------ 47070
human.FAM230J-LOC105372942-GGTLC5P.NCBI.rev.compl                                    ------------------------------------------------------------ 46364
human.FAM230A.ncRNA.18487127.18500594-GGTLC3.18516335.18518165.rev.compl.REF         ------------------------------------------------------------ 31039

chimp.LOC112206744-LOC107973052-GGT2.revcompl                                        ------------------------------------------------------------ 49429
chimp54546-135457.revcompl.Pan.troglodytes.clone.rp43-41g5.GenBank:AC099533.36      GAGGCTGCATTGAGCTATAATCACACCACTGCACTCCAGCTTGGGTGACAAAGTGAGACC 52270
human.FAM30B-LOC105372935-GGT2.NCBI.                                                 ------------------------------------------------------------ 48417
human.FAM230E-LOC105377182-GGT3P.NCBI.                                               ------------------------------------------------------------ 47070
human.FAM230J-LOC105372942-GGTLC5P.NCBI.rev.compl                                    ------------------------------------------------------------ 46364
human.FAM230A.ncRNA.18487127.18500594-GGTLC3.18516335.18518165.rev.compl.REF         ------------------------------------------------------------ 31039

chimp.LOC112206744-LOC107973052-GGT2.revcompl                                        ------------------------------------------------------------ 49429
chimp54546-135457.revcompl.Pan.troglodytes.clone.rp43-41g5.GenBank:AC099533.36      CTGTCTCTAAAAGAAAAAAAAATTGGCCTGTGAGCATGGGTTTGATTTTCAAACAGGACC 52330
human.FAM30B-LOC105372935-GGT2.NCBI.                                                 ------------------------------------------------------------ 48417
human.FAM230E-LOC105377182-GGT3P.NCBI.                                               ------------------------------------------------------------ 47070
human.FAM230J-LOC105372942-GGTLC5P.NCBI.rev.compl                                    ------------------------------------------------------------ 46364
human.FAM230A.ncRNA.18487127.18500594-GGTLC3.18516335.18518165.rev.compl.REF         ------------------------------------------------------------ 31039

chimp.LOC112206744-LOC107973052-GGT2.revcompl                                        ------------------------------------------------------------ 49429
chimp54546-135457.revcompl.Pan.troglodytes.clone.rp43-41g5.GenBank:AC099533.36      CGGAGGGTAGGGACAGACAGTGCTGTCACCCTTAGGTGCTGAACACTCAGAAACGGGCCA 52390
human.FAM30B-LOC105372935-GGT2.NCBI.                                                 ------------------------------------------------------------ 48417
human.FAM230E-LOC105377182-GGT3P.NCBI.                                               ------------------------------------------------------------ 47070
human.FAM230J-LOC105372942-GGTLC5P.NCBI.rev.compl                                    ------------------------------------------------------------ 46364
human.FAM230A.ncRNA.18487127.18500594-GGTLC3.18516335.18518165.rev.compl.REF         ------------------------------------------------------------ 31039

chimp.LOC112206744-LOC107973052-GGT2.revcompl                                        ---------cttaattctcctggagtcagctccagacagacagggtattggcatgccaat 49480
chimp54546-135457.revcompl.Pan.troglodytes.clone.rp43-41g5.GenBank:AC099533.36      GCGGCAGCCCTTTCCTCACCTGCAGACACCAG----ATTGGGCAGAACAGCACGTGGCAC 52446
human.FAM30B-LOC105372935-GGT2.NCBI.                                                 ---------cttaattctcctggagtcagctc----cagacagggtattggcatgccaat 48464
human.FAM230E-LOC105377182-GGT3P.NCBI.                                               ---------cttaattctcctggagtcagctc----cagacagggtattggcatgccaat 47117
human.FAM230J-LOC105372942-GGTLC5P.NCBI.rev.compl                                    ------------------------------------------------------------ 46364
human.FAM230A.ncRNA.18487127.18500594-GGTLC3.18516335.18518165.rev.compl.REF         ------------------------------------------------------------ 31039

chimp.LOC112206744-LOC107973052-GGT2.revcompl                                        ttccagcctcagtggtaaaggtcg-acacgctaatcaccctcctccatgaagcagtgaca 49539
chimp54546-135457.revcompl.Pan.troglodytes.clone.rp43-41g5.GenBank:AC099533.36      TTGCAGCTCTTGCAGTGAGGACAGAACCCAGTGTCAACCCTTCTGCCTGTGGGAGGGGCT 52506
human.FAM30B-LOC105372935-GGT2.NCBI.                                                 ttccagcctcagtggtaaaggtcg-acacgctaatcaccctcctccatgaaacagtgaca 48523
human.FAM230E-LOC105377182-GGT3P.NCBI.                                               ttccagcctcagtggtaaaggtcg-acacgctaatcaccctcctccatgaaacagtgaca 47176
human.FAM230J-LOC105372942-GGTLC5P.NCBI.rev.compl                                    ------------------------------------------------------------ 46364
human.FAM230A.ncRNA.18487127.18500594-GGTLC3.18516335.18518165.rev.compl.REF         ------------------------------------------------------------ 31039

chimp.LOC112206744-LOC107973052-GGT2.revcompl                                        aaaattacctgaagaaagccacagccaagctccaggcccctgccccacaaatccccttcc 49599
chimp54546-135457.revcompl.Pan.troglodytes.clone.rp43-41g5.GenBank:AC099533.36      GCTGAGGCCTGCAGAGAG-----GCCAGGGTGGAGGCTCATCCCCTTGTCCAGCCCTTGG 52561
human.FAM30B-LOC105372935-GGT2.NCBI.                                                 aaaattacctgaagaaagccacagccaagctccaggcccctgccccacaaatccccttcc 48583
human.FAM230E-LOC105377182-GGT3P.NCBI.                                               aaaattacctgaagaaagccacagccaagctccaggcccctgccccacaaatccccttcc 47236
human.FAM230J-LOC105372942-GGTLC5P.NCBI.rev.compl                                    ------------------------------------------------------------ 46364
human.FAM230A.ncRNA.18487127.18500594-GGTLC3.18516335.18518165.rev.compl.REF         ------------------------------------------------------------ 31039

chimp.LOC112206744-LOC107973052-GGT2.revcompl                                        ccatgcctctctcaaggcgaccctcatcccttgtaaccctcttggtgaatcaaagcgccc 49659
chimp54546-135457.revcompl.Pan.troglodytes.clone.rp43-41g5.GenBank:AC099533.36      CGTGGTCTCCACCAGGTCCCCAGCCCACCAGTGTAGGGC---------------GCCCCT 52606
human.FAM30B-LOC105372935-GGT2.NCBI.                                                 ccatgcctctctcaaggcgaccctcatcccttgtaaccctcttggtgaatcaaagcgccc 48643
human.FAM230E-LOC105377182-GGT3P.NCBI.                                               ccatgcctctctcaaggcgaccctcatcccttgtaaccctcttggtgaatcaaagcgccc 47296
human.FAM230J-LOC105372942-GGTLC5P.NCBI.rev.compl                                    ------------------------------------------------------------ 46364
human.FAM230A.ncRNA.18487127.18500594-GGTLC3.18516335.18518165.rev.compl.REF         ------------------------------------------------------------ 31039

chimp.LOC112206744-LOC107973052-GGT2.revcompl                                        tctactgggccttagcccagctgttcctct-gctaggaatcccttcctctctctgcctaa 49718
chimp54546-135457.revcompl.Pan.troglodytes.clone.rp43-41g5.GenBank:AC099533.36      GAACCTGCTGCTGCCACAGGCCCTGTCTCTACCCAGGACGTCCCCCCACCCTCGCAGTGT 52666
human.FAM30B-LOC105372935-GGT2.NCBI.                                                 tctactgggccttagcccagctgttcctct-gctaggaatcccttcctctctctgcctaa 48702
human.FAM230E-LOC105377182-GGT3P.NCBI.                                               tctactgggccttagcccagctgttcctct-gctaggaatcccttcctctctctgcctaa 47355
human.FAM230J-LOC105372942-GGTLC5P.NCBI.rev.compl                                    ------------------------------------------------------------ 46364
human.FAM230A.ncRNA.18487127.18500594-GGTLC3.18516335.18518165.rev.compl.REF         ------------------------------------------------------------ 31039

chimp.LOC112206744-LOC107973052-GGT2.revcompl                                        cgaagttatctgcagcccagccgccacctcctccaagaagtcctcct------------- 49765
chimp54546-135457.revcompl.Pan.troglodytes.clone.rp43-41g5.GenBank:AC099533.36      CAGGGAAATGATCATGGTGGCGGTGACACTCCGCAGGCAGGGCTGCTGAGAGAAGCTGAG 52726
human.FAM30B-LOC105372935-GGT2.NCBI.                                                 cgaagttatctgcagcccagccgccacctcctccaagaagtcctcct------------- 48749
human.FAM230E-LOC105377182-GGT3P.NCBI.                                               cgaagttatctgcagcccagccgccacctcctccaagaagtcctcct------------- 47402
human.FAM230J-LOC105372942-GGTLC5P.NCBI.rev.compl                                    ------------------------------------------------------------ 46364
human.FAM230A.ncRNA.18487127.18500594-GGTLC3.18516335.18518165.rev.compl.REF         ------------------------------------------------------------ 31039

chimp.LOC112206744-LOC107973052-GGT2.revcompl                                        ------------------------------------------------------------ 49765
chimp54546-135457.revcompl.Pan.troglodytes.clone.rp43-41g5.GenBank:AC099533.36      AAGGGTCACACTGCAGGCAGGGGCCCGTGTGACAAGCCCCTCTCACCCCGAGAGAGCTGA 52786
human.FAM30B-LOC105372935-GGT2.NCBI.                                                 ------------------------------------------------------------ 48749
human.FAM230E-LOC105377182-GGT3P.NCBI.                                               ------------------------------------------------------------ 47402
human.FAM230J-LOC105372942-GGTLC5P.NCBI.rev.compl                                    ------------------------------------------------------------ 46364
human.FAM230A.ncRNA.18487127.18500594-GGTLC3.18516335.18518165.rev.compl.REF         ------------------------------------------------------------ 31039

chimp.LOC112206744-LOC107973052-GGT2.revcompl                                        ------------------------------------------------------------ 49765
chimp54546-135457.revcompl.Pan.troglodytes.clone.rp43-41g5.GenBank:AC099533.36      CCAGGCAGCTCACGAGCAGAGCCACATCCCGGGAGTCCGAGAAAGGTCCTGGCTGGGCTC 52846
human.FAM30B-LOC105372935-GGT2.NCBI.                                                 ------------------------------------------------------------ 48749
human.FAM230E-LOC105377182-GGT3P.NCBI.                                               ------------------------------------------------------------ 47402
human.FAM230J-LOC105372942-GGTLC5P.NCBI.rev.compl                                    ------------------------------------------------------------ 46364
human.FAM230A.ncRNA.18487127.18500594-GGTLC3.18516335.18518165.rev.compl.REF         ------------------------------------------------------------ 31039

chimp.LOC112206744-LOC107973052-GGT2.revcompl                                        -----------------------------------------------------ggatctt 49772
chimp54546-135457.revcompl.Pan.troglodytes.clone.rp43-41g5.GenBank:AC099533.36      AGCCACCTCATTGGCCACGGGCAGCCTTTGTCGTGTGAGCCTTGCTCTCCTGGGGAGGCT 52906
```

```
human.FAM30B-LOC105372935-GGT2.NCBI.                                       -----------------------------------------------------ggatctt 48756
human.FAM230E-LOC105377182-GGT3P.NCBI.                                     -----------------------------------------------------ggatctt 47409
human.FAM230J-LOC105372942-GGTLC5P.NCBI.rev.compl                          ------------------------------------------------------------ 46364
human.FAM230A.ncRNA.18487127.18500594-GGTLC3.18516335.18518165.rev.compl.REF ------------------------------------------------------------ 31039

chimp.LOC112206744-LOC107973052-GGT2.revcompl                              caggctgtattctagtgcttccctagccctggctcttgcctacacctgcattta------ 49826
chimp54546-135457.revcompl.Pan.troglodytes.clone.rp43-41g5.GenBank:AC099533.36 CAGGCTGATGGCTGATGTGGGCATTGCCGAGGGTAACCTGTGGCCCAGTGTATATGGCCG 52966
human.FAM30B-LOC105372935-GGT2.NCBI.                                       caggctgtattctagtgcttccctagccctggctcttgcctacacctgcattta------ 48810
human.FAM230E-LOC105377182-GGT3P.NCBI.                                     caggctgtattctagtgcttccctagccctggctcttgcctacacctgcattta------ 47463
human.FAM230J-LOC105372942-GGTLC5P.NCBI.rev.compl                          ------------------------------------------------------------ 46364
human.FAM230A.ncRNA.18487127.18500594-GGTLC3.18516335.18518165.rev.compl.REF ------------------------------------------------------------ 31039

chimp.LOC112206744-LOC107973052-GGT2.revcompl                              --------------------------------------------------ccccaacagg 49836
chimp54546-135457.revcompl.Pan.troglodytes.clone.rp43-41g5.GenBank:AC099533.36 GCTCTCCTCAAGCTGCATTCATTCAAGTAGGACCCAGGGTGCATGCCCATCTCCAGCCCA 53026
human.FAM30B-LOC105372935-GGT2.NCBI.                                       --------------------------------------------------ccccaacagg 48820
human.FAM230E-LOC105377182-GGT3P.NCBI.                                     --------------------------------------------------ccccaacagg 47473
human.FAM230J-LOC105372942-GGTLC5P.NCBI.rev.compl                          ------------------------------------------------------------ 46364
human.FAM230A.ncRNA.18487127.18500594-GGTLC3.18516335.18518165.rev.compl.REF ------------------------------------------------------------ 31039

chimp.LOC112206744-LOC107973052-GGT2.revcompl                              gacttgctctcctggactgtgtggcctctcttggttttgatataagcaggagctgtgga- 49895
chimp54546-135457.revcompl.Pan.troglodytes.clone.rp43-41g5.GenBank:AC099533.36 GGGCAGCTCCCCTGTAAGCTGGGTGAGCTACTGAAGCCAAGGTGGGAGGCAGCTGACAAC 53086
human.FAM30B-LOC105372935-GGT2.NCBI.                                       gacttgctctcctggactgtgtggcctctcttggttttgatataagcaggagctgtgga- 48879
human.FAM230E-LOC105377182-GGT3P.NCBI.                                     gacttgctctcctggactgtgtggcctctcttggttttgatataagcaggagctgtgga- 47532
human.FAM230J-LOC105372942-GGTLC5P.NCBI.rev.compl                          ------------------------------------------------------------ 46364
human.FAM230A.ncRNA.18487127.18500594-GGTLC3.18516335.18518165.rev.compl.REF ------------------------------------------------------------ 31039

chimp.LOC112206744-LOC107973052-GGT2.revcompl                              --cccacatggcc--------agtcactgaccctcctccaccaggaacttcctgcaggct 49945
chimp54546-135457.revcompl.Pan.troglodytes.clone.rp43-41g5.GenBank:AC099533.36 ACCACCCATGGCCCATGCGGAGGTGGCAGAAAGGCTGGACTCAGCAGGAACACCAAATCC 53146
human.FAM30B-LOC105372935-GGT2.NCBI.                                       --cccacatggcc--------agtcactgaccctcctccaccaggaacttcctgcaggct 48929
human.FAM230E-LOC105377182-GGT3P.NCBI.                                     --cccacatggcc--------agtcactgaccctcctccaccaggaacttcctgcaggct 47582
human.FAM230J-LOC105372942-GGTLC5P.NCBI.rev.compl                          ------------------------------------------------------------ 46364
human.FAM230A.ncRNA.18487127.18500594-GGTLC3.18516335.18518165.rev.compl.REF ------------------------------------------------------------ 31039

chimp.LOC112206744-LOC107973052-GGT2.revcompl                              caggcaagacaggaggaccccatggctctgggctacagctcagggtttccactgcagagt 50005
chimp54546-135457.revcompl.Pan.troglodytes.clone.rp43-41g5.GenBank:AC099533.36 CGGACCAGGCAGAAACCACCCAAGACTGAGGGGCTCGTGCCAGAGCGGTGGC--CACAGG 53204
human.FAM30B-LOC105372935-GGT2.NCBI.                                       caggcaagacaggaggaccccatggctctgggctacagctcagggtttccactgcagagt 48989
human.FAM230E-LOC105377182-GGT3P.NCBI.                                     caggcaagacaggaggaccccatggctctgggctacagctcagggtttccactgcagagt 47642
human.FAM230J-LOC105372942-GGTLC5P.NCBI.rev.compl                          ------------------------------------------------------------ 46364
human.FAM230A.ncRNA.18487127.18500594-GGTLC3.18516335.18518165.rev.compl.REF ------------------------------------------------------------ 31039

chimp.LOC112206744-LOC107973052-GGT2.revcompl                              tcctcacccaggtccttgaggttacccactcaccagccgcctggatgtccttcacaatct 50065
chimp54546-135457.revcompl.Pan.troglodytes.clone.rp43-41g5.GenBank:AC099533.36 TAAGAACCCGGGCCCAGGTTGT-------------GTGTGGCGGGAATCCTCCATGTCCC 53251
human.FAM30B-LOC105372935-GGT2.NCBI.                                       tcctcacccaggtccttgaggttacccactcaccagccgcctggatgtccttcacaatct 49049
human.FAM230E-LOC105377182-GGT3P.NCBI.                                     tcctcacccaggtccttgaggttacccactcaccagccgcctggatgtccttcacaatct 47702
human.FAM230J-LOC105372942-GGTLC5P.NCBI.rev.compl                          ------------------------------------------------------------ 46364
human.FAM230A.ncRNA.18487127.18500594-GGTLC3.18516335.18518165.rev.compl.REF ------------------------------------------------------------ 31039

chimp.LOC112206744-LOC107973052-GGT2.revcompl                              gggccatgaggctgccgttgtagaaggcctgggcgccctcgatggccagcatctcgtagg 50125
chimp54546-135457.revcompl.Pan.troglodytes.clone.rp43-41g5.GenBank:AC099533.36 AGGGCTTAGCGTAGCAAAGGAAGACCAGCCGGGTCACCCTGGTGGCCATC----TGTCCC 53307
human.FAM30B-LOC105372935-GGT2.NCBI.                                       gggccatgaggctgccgttgtagaaggcctgggcaccctcgatggccagcatctcatagg 49109
human.FAM230E-LOC105377182-GGT3P.NCBI.                                     gggccatgaggctgccgttgtagaaggcctgggcaccctcgatggccagcatctcgtagg 47762
human.FAM230J-LOC105372942-GGTLC5P.NCBI.rev.compl                          ------------------------------------------------------------ 46364
human.FAM230A.ncRNA.18487127.18500594-GGTLC3.18516335.18518165.rev.compl.REF ------------------------------------------------------------ 31039

chimp.LOC112206744-LOC107973052-GGT2.revcompl                              tgtcagccagccgcggcagggtcagtctctccccctcccgaagcacctttctatcccggc 50185
chimp54546-135457.revcompl.Pan.troglodytes.clone.rp43-41g5.GenBank:AC099533.36 TGTCCCACCTGCAGAGTCAGAACAGCCTCTCCCCAGTGGGGATCATCTCTCTCTGCCAAA 53367
human.FAM30B-LOC105372935-GGT2.NCBI.                                       tgtcagccagccgcggcagggtcagtctctccccctcccgaagcacctttctatcccggc 49169
human.FAM230E-LOC105377182-GGT3P.NCBI.                                     tgtcagccagccgcggcagggtcagtctctccccctcccgaagcacctttctatcccggc 47822
human.FAM230J-LOC105372942-GGTLC5P.NCBI.rev.compl                          ------------------------------------------------------------ 46364
human.FAM230A.ncRNA.18487127.18500594-GGTLC3.18516335.18518165.rev.compl.REF ------------------------------------------------------------ 31039

chimp.LOC112206744-LOC107973052-GGT2.revcompl                              agaacacctcactggggcagaggggg---------------------------------- 50211
chimp54546-135457.revcompl.Pan.troglodytes.clone.rp43-41g5.GenBank:AC099533.36 GCAATAGCGGTCCCTGCCCCAACCAGACTACCCCACTCAGTGGAGTTACGGATGCTGCTC 53427
human.FAM30B-LOC105372935-GGT2.NCBI.                                       agaacacctcaccggggcagaggggg---------------------------------- 49195
human.FAM230E-LOC105377182-GGT3P.NCBI.                                     agaacacctcactggggcagaggggg---------------------------------- 47848
human.FAM230J-LOC105372942-GGTLC5P.NCBI.rev.compl                          ------------------------------------------------------------ 46364
human.FAM230A.ncRNA.18487127.18500594-GGTLC3.18516335.18518165.rev.compl.REF ------------------------------------------------------------ 31039

chimp.LOC112206744-LOC107973052-GGT2.revcompl                              -------ctcatgtgaggcagcaggtggggtggactcagctagaccacccccacacccg 50264
chimp54546-135457.revcompl.Pan.troglodytes.clone.rp43-41g5.GenBank:AC099533.36 CAGCATCCTAACACTGCCCAGCTGGTGCCTGCCTGTGCTCACCCACACCCCCCAGGCCAG 53487
human.FAM30B-LOC105372935-GGT2.NCBI.                                       -------ctcatgtgaggcagcaggtggggtggactcagctagaccacccccacacctg 49248
human.FAM230E-LOC105377182-GGT3P.NCBI.                                     -------ctcatgtgaggcagcaggtggggtggactcagctagaccacccccacacctg 47901
human.FAM230J-LOC105372942-GGTLC5P.NCBI.rev.compl                          ------------------------------------------------------------ 46364
human.FAM230A.ncRNA.18487127.18500594-GGTLC3.18516335.18518165.rev.compl.REF ------------------------------------------------------------ 31039

chimp.LOC112206744-LOC107973052-GGT2.revcompl                              tccatacaggagaccagcacaaagcaggggcagcgcctgtcacaggtgggtggccctgtc 50324
chimp54546-135457.revcompl.Pan.troglodytes.clone.rp43-41g5.GenBank:AC099533.36 CCTTCCCTGCAGCCTGGGCTTGGCCACCTTGGCCTGATTGAGCACTGAGGACTCCTGGGC 53547
human.FAM30B-LOC105372935-GGT2.NCBI.                                       cccacacaggagaccagcacaaagcaggggcagcacctgtcacaggtgggtggccctgtc 49308
human.FAM230E-LOC105377182-GGT3P.NCBI.                                     cccacacaggagaacggagcagaacaggggcagcgcctgtcacaggtgggtggccctgtc 47961
human.FAM230J-LOC105372942-GGTLC5P.NCBI.rev.compl                          ------------------------------------------------------------ 46364
human.FAM230A.ncRNA.18487127.18500594-GGTLC3.18516335.18518165.rev.compl.REF ------------------------------------------------------------ 31039

chimp.LOC112206744-LOC107973052-GGT2.revcompl                              actcagcgctcatcctcctagtgtcccttccgggaacctcctagtgtc------------ 50372
chimp54546-135457.revcompl.Pan.troglodytes.clone.rp43-41g5.GenBank:AC099533.36 ACCCAGCCCCATCACTGCACCTGCTGCTTCCAGCCCCACCCCACCGGCTCTGGGGTTCTT 53607
human.FAM30B-LOC105372935-GGT2.NCBI.                                       actcagcgctcatcctcctagtgtcccttccgggagcctcctagtgtc------------ 49356
human.FAM230E-LOC105377182-GGT3P.NCBI.                                     actcagcgctcgtcctcctagtgtcccttccgggagcctcctagtgtc------------ 48009
human.FAM230J-LOC105372942-GGTLC5P.NCBI.rev.compl                          ------------------------------------------------------------ 46364
human.FAM230A.ncRNA.18487127.18500594-GGTLC3.18516335.18518165.rev.compl.REF ------------------------------------------------------------ 31039

chimp.LOC112206744-LOC107973052-GGT2.revcompl                              ------------------------------------------------------------ 50372
chimp54546-135457.revcompl.Pan.troglodytes.clone.rp43-41g5.GenBank:AC099533.36 CCCAGTGGCGCTGATCATGAAATCAACATGCACACAAGTCGTCTCAGGAAACTTTTTAAT 53667
human.FAM30B-LOC105372935-GGT2.NCBI.                                       ------------------------------------------------------------ 49356
human.FAM230E-LOC105377182-GGT3P.NCBI.                                     ------------------------------------------------------------ 48009
human.FAM230J-LOC105372942-GGTLC5P.NCBI.rev.compl                          ------------------------------------------------------------ 46364
human.FAM230A.ncRNA.18487127.18500594-GGTLC3.18516335.18518165.rev.compl.REF ------------------------------------------------------------ 31039

chimp.LOC112206744-LOC107973052-GGT2.revcompl                              ------------------------------------------------------------ 50372
chimp54546-135457.revcompl.Pan.troglodytes.clone.rp43-41g5.GenBank:AC099533.36 GAAAGTGTCGGCCATGGTGGTGTGTAGGTGGCTGAGCTCAGATTGCAGCTGCTAAGACAC 53727
human.FAM30B-LOC105372935-GGT2.NCBI.                                       ------------------------------------------------------------ 49356
human.FAM230E-LOC105377182-GGT3P.NCBI.                                     ------------------------------------------------------------ 48009
human.FAM230J-LOC105372942-GGTLC5P.NCBI.rev.compl                          ------------------------------------------------------------ 46364
human.FAM230A.ncRNA.18487127.18500594-GGTLC3.18516335.18518165.rev.compl.REF ------------------------------------------------------------ 31039

chimp.LOC112206744-LOC107973052-GGT2.revcompl                              ----------------------------ccttgccactcaggacacatggccagccacag 50404
chimp54546-135457.revcompl.Pan.troglodytes.clone.rp43-41g5.GenBank:AC099533.36 CAGCCACTTACCAAGAGAAAGCCAGGCTGCTTCAAACTCAGGGCCCGAGGCAAAAAAGCA 53787
human.FAM30B-LOC105372935-GGT2.NCBI.                                       ----------------------------ccttgccactcaggacacgtggccagccacag 49388
human.FAM230E-LOC105377182-GGT3P.NCBI.                                     ----------------------------ccttgccactcaggacacgtggccagccacag 48041
human.FAM230J-LOC105372942-GGTLC5P.NCBI.rev.compl                          ------------------------------------------------------------ 46364
human.FAM230A.ncRNA.18487127.18500594-GGTLC3.18516335.18518165.rev.compl.REF ------------------------------------------------------------ 31039

chimp.LOC112206744-LOC107973052-GGT2.revcompl                              tggccactgggaccccacgctcagaatgtgtccccacacgtggtgggaagggtctgtatc 50464
chimp54546-135457.revcompl.Pan.troglodytes.clone.rp43-41g5.GenBank:AC099533.36 TCACTTCTGGTCGGGGAGTCTGGAAGCCATGCCTTGTGGGAGGTCACACTGGCATCTAGG 53847
human.FAM30B-LOC105372935-GGT2.NCBI.                                       tggccactgggaccccatgctcagaatgtgtccccacacgtggtgggaagggtctgtatc 49448
human.FAM230E-LOC105377182-GGT3P.NCBI.                                     tggccactgggaccccacgctcagaatgtgtccccacacgtggtgggaagggtctgtatc 48101
human.FAM230J-LOC105372942-GGTLC5P.NCBI.rev.compl                          ------------------------------------------------------------ 46364
human.FAM230A.ncRNA.18487127.18500594-GGTLC3.18516335.18518165.rev.compl.REF ------------------------------------------------------------ 31039

chimp.LOC112206744-LOC107973052-GGT2.revcompl                              tcctcatcccattatcagcacagg------------gtcctgaaggcagagggccactcc 50512
chimp54546-135457.revcompl.Pan.troglodytes.clone.rp43-41g5.GenBank:AC099533.36 CCTTCGCCTGCATTGCAGAAGGAGAGCCGGGTCCCCCTCCTGGAGAACGCTGCGTTCCCC 53907
human.FAM30B-LOC105372935-GGT2.NCBI.                                       tcctcatcccattatcagcacagg------------gtcctgaaggcagagggccgctcc 49496
human.FAM230E-LOC105377182-GGT3P.NCBI.                                     tcctcatcccattatcagcacagg------------gtcctgaaggcagagggccgctcc 48149
human.FAM230J-LOC105372942-GGTLC5P.NCBI.rev.compl                          ------------------------------------------------------------ 46364
human.FAM230A.ncRNA.18487127.18500594-GGTLC3.18516335.18518165.rev.compl.REF ------------------------------------------------------------ 31039

chimp.LOC112206744-LOC107973052-GGT2.revcompl                              actgctgctacggcctgcaaggtccttgggctgtgcctgccctgcctgtgtcagggg--g 50570
chimp54546-135457.revcompl.Pan.troglodytes.clone.rp43-41g5.GenBank:AC099533.36 AGCCCCACACCGGCTTTGCCACCACACAGGCTTTTGAGGCAGGAGGTGGGTAAGACGTAG 53967
human.FAM30B-LOC105372935-GGT2.NCBI.                                       actgctgctacggcctgcaaggtccttgggctgtgcctgccctgcctgtgtcagggg--g 49554
```

```
human.FAM230E-LOC105377182-GGT3P.NCBI.                                           actgctgctacggcctgcaaggtccttgggctgtgcctgccctgcctgtgtcagggg--g 48207
human.FAM230J-LOC105372942-GGTLC5P.NCBI.rev.compl                                ------------------------------------------------------------ 46364
human.FAM230A.ncRNA.18487127.18500594-GGTLC3.18516335.18518165.rev.compl.REF     ------------------------------------------------------------ 31039

chimp.LOC112206744-LOC107973052-GGT2.revcompl                                    ccgcacccacagacatgccacaagacaggc------------------------------ 50600
chimp54546-135457.revcompl.Pan.troglodytes.clone.rp43-41g5.GenBank:AC099533.36   CTGTAGACCCAAAGCAACCACCAGCCCTGGGACCCTGCGGGAGAGGAGCACTTTTAGAAA 54027
human.FAM30B-LOC105372935-GGT2.NCBI.                                             ccgcacccacagacataccacaagacaggc------------------------------ 49584
human.FAM230E-LOC105377182-GGT3P.NCBI.                                           ccgcacccacagacataccacaagacaggc------------------------------ 48237
human.FAM230J-LOC105372942-GGTLC5P.NCBI.rev.compl                                ------------------------------------------------------------ 46364
human.FAM230A.ncRNA.18487127.18500594-GGTLC3.18516335.18518165.rev.compl.REF     ------------------------------------------------------------ 31039

chimp.LOC112206744-LOC107973052-GGT2.revcompl                                    ------------------------------------------------------------ 50600
chimp54546-135457.revcompl.Pan.troglodytes.clone.rp43-41g5.GenBank:AC099533.36   ATGGAAAAGTATGGTCATCCCATCATTAGACAGCACACATCCTACATAAATAAAAAGTCG 54087
human.FAM30B-LOC105372935-GGT2.NCBI.                                             ------------------------------------------------------------ 49584
human.FAM230E-LOC105377182-GGT3P.NCBI.                                           ------------------------------------------------------------ 48237
human.FAM230J-LOC105372942-GGTLC5P.NCBI.rev.compl                                ------------------------------------------------------------ 46364
human.FAM230A.ncRNA.18487127.18500594-GGTLC3.18516335.18518165.rev.compl.REF     ------------------------------------------------------------ 31039

chimp.LOC112206744-LOC107973052-GGT2.revcompl                                    ------------------------------------------------------------ 50600
chimp54546-135457.revcompl.Pan.troglodytes.clone.rp43-41g5.GenBank:AC099533.36   TATGGGGAAGGAGGTTGGGGAGGGAATAAAAAATTGGCACAGACATTGATAGACTGGTTT 54147
human.FAM30B-LOC105372935-GGT2.NCBI.                                             ------------------------------------------------------------ 49584
human.FAM230E-LOC105377182-GGT3P.NCBI.                                           ------------------------------------------------------------ 48237
human.FAM230J-LOC105372942-GGTLC5P.NCBI.rev.compl                                ------------------------------------------------------------ 46364
human.FAM230A.ncRNA.18487127.18500594-GGTLC3.18516335.18518165.rev.compl.REF     ------------------------------------------------------------ 31039

chimp.LOC112206744-LOC107973052-GGT2.revcompl                                    ------------------------------------------------------------ 50600
chimp54546-135457.revcompl.Pan.troglodytes.clone.rp43-41g5.GenBank:AC099533.36   CCAGTTTCAAGGTAACAGATGCACATCATGAGACCAGAGGAGGCAGAGACAAGGCTGGAT 54207
human.FAM30B-LOC105372935-GGT2.NCBI.                                             ------------------------------------------------------------ 49584
human.FAM230E-LOC105377182-GGT3P.NCBI.                                           ------------------------------------------------------------ 48237
human.FAM230J-LOC105372942-GGTLC5P.NCBI.rev.compl                                ------------------------------------------------------------ 46364
human.FAM230A.ncRNA.18487127.18500594-GGTLC3.18516335.18518165.rev.compl.REF     ------------------------------------------------------------ 31039

chimp.LOC112206744-LOC107973052-GGT2.revcompl                                    --tgctgctcgatgacggtccgcttgttttccagggctgccgccaagcccttgcccacag 50658
chimp54546-135457.revcompl.Pan.troglodytes.clone.rp43-41g5.GenBank:AC099533.36   TTGGCTTTTCTAAGCAACATGTGTTCCTGTGCAGGGCTGAATGGTCGCTGAGACAGAGAT 54267
human.FAM30B-LOC105372935-GGT2.NCBI.                                             --tgctgctcgatgacagtccgcttgttttccaggactgccgccaagcccttgcccacag 49642
human.FAM230E-LOC105377182-GGT3P.NCBI.                                           --tgctgctcgatgacggtccgcttgttttccaggactgccgccaagcccttgcccacgg 48295
human.FAM230J-LOC105372942-GGTLC5P.NCBI.rev.compl                                ------------------------------------------------------------ 46364
human.FAM230A.ncRNA.18487127.18500594-GGTLC3.18516335.18518165.rev.compl.REF     ------------------------------------------------------------ 31039

chimp.LOC112206744-LOC107973052-GGT2.revcompl                                    ggaagccctggcgggccagctg-------------------------------------- 50680
chimp54546-135457.revcompl.Pan.troglodytes.clone.rp43-41g5.GenBank:AC099533.36   GGAAGCCAGGACAAGGGAGCCCACCGGGCCCAGATAGGTACAGAGAGCAGAGGCTCCTGT 54327
human.FAM30B-LOC105372935-GGT2.NCBI.                                             ggaagccctggcgggccagctg-------------------------------------- 49664
human.FAM230E-LOC105377182-GGT3P.NCBI.                                           ggaagccctggcgggccagctg-------------------------------------- 48317
human.FAM230J-LOC105372942-GGTLC5P.NCBI.rev.compl                                ------------------------------------------------------------ 46364
human.FAM230A.ncRNA.18487127.18500594-GGTLC3.18516335.18518165.rev.compl.REF     ------------------------------------------------------------ 31039

chimp.LOC112206744-LOC107973052-GGT2.revcompl                                    ------------------------------------------------------------ 50680
chimp54546-135457.revcompl.Pan.troglodytes.clone.rp43-41g5.GenBank:AC099533.36   TCTGTCCTCGCCACCCACGAGGGTGACACTGCTTGTAAATGGTGGCTGTACTCTCCCAGC 54387
human.FAM30B-LOC105372935-GGT2.NCBI.                                             ------------------------------------------------------------ 49664
human.FAM230E-LOC105377182-GGT3P.NCBI.                                           ------------------------------------------------------------ 48317
human.FAM230J-LOC105372942-GGTLC5P.NCBI.rev.compl                                ------------------------------------------------------------ 46364
human.FAM230A.ncRNA.18487127.18500594-GGTLC3.18516335.18518165.rev.compl.REF     ------------------------------------------------------------ 31039

chimp.LOC112206744-LOC107973052-GGT2.revcompl                                    ------------------------------------------------------------ 50680
chimp54546-135457.revcompl.Pan.troglodytes.clone.rp43-41g5.GenBank:AC099533.36   AAGAAAAAAGCACAACTAAATCCACACTGCACACAGACGCAGACAGAAAGCCTTCAAGTG 54447
human.FAM30B-LOC105372935-GGT2.NCBI.                                             ------------------------------------------------------------ 49664
human.FAM230E-LOC105377182-GGT3P.NCBI.                                           ------------------------------------------------------------ 48317
human.FAM230J-LOC105372942-GGTLC5P.NCBI.rev.compl                                ------------------------------------------------------------ 46364
human.FAM230A.ncRNA.18487127.18500594-GGTLC3.18516335.18518165.rev.compl.REF     ------------------------------------------------------------ 31039

chimp.LOC112206744-LOC107973052-GGT2.revcompl                                    ------------------------------------------gatgctgggctggaagag 50698
chimp54546-135457.revcompl.Pan.troglodytes.clone.rp43-41g5.GenBank:AC099533.36   GCTCTGTTTTCTGCTCCCTGCCTTGCCAGGTCCACAAGCAGAGAGGAGTGTCAGGCGCAT 54507
human.FAM30B-LOC105372935-GGT2.NCBI.                                             ------------------------------------------gatgctgggctggaagag 49682
human.FAM230E-LOC105377182-GGT3P.NCBI.                                           ------------------------------------------gatgctgggctggaagag 48335
human.FAM230J-LOC105372942-GGTLC5P.NCBI.rev.compl                                ------------------------------------------------------------ 46364
human.FAM230A.ncRNA.18487127.18500594-GGTLC3.18516335.18518165.rev.compl.REF     ------------------------------------------------------------ 31039

chimp.LOC112206744-LOC107973052-GGT2.revcompl                                    gcgagcccagggcagctgcccatgccactgatgtgccagctcatagcctcggatctcccc 50758
chimp54546-135457.revcompl.Pan.troglodytes.clone.rp43-41g5.GenBank:AC099533.36   GGCCCCACTGTCAGGCTCCCCAGTGAGCTGCGGGTTCAGCAGGAGCTGCCCACTGACACA 54567
human.FAM30B-LOC105372935-GGT2.NCBI.                                             gcgagcccagggcagccgcccatgccgctggtgtgccagctcatagcctcggatctcccc 49742
human.FAM230E-LOC105377182-GGT3P.NCBI.                                           gcgagcccagggcagccgcccatgccgctggtgtgccagctcatagcctcggatctcccc 48395
human.FAM230J-LOC105372942-GGTLC5P.NCBI.rev.compl                                ------------------------------------------------------------ 46364
human.FAM230A.ncRNA.18487127.18500594-GGTLC3.18516335.18518165.rev.compl.REF     ------------------------------------------------------------ 31039

chimp.LOC112206744-LOC107973052-GGT2.revcompl                                    aggcaccgccaccgacagccctcctggggagagagagccacagttagtgaccctgagtgg 50818
chimp54546-135457.revcompl.Pan.troglodytes.clone.rp43-41g5.GenBank:AC099533.36   CAGGGGACACCCACTCCTGCCACCTTGGGAGCGGTTGCCAGACAGAGCCGCACTGGGTGC 54627
human.FAM30B-LOC105372935-GGT2.NCBI.                                             aggcaccgccaccgacagccctcctggggagagagagccacagttagtgaccctgagtgg 49802
human.FAM230E-LOC105377182-GGT3P.NCBI.                                           aggcaccgccaccgacagccctcctggggagagagagccacagttagtgaccctgagtgg 48455
human.FAM230J-LOC105372942-GGTLC5P.NCBI.rev.compl                                ------------------------------------------------------------ 46364
human.FAM230A.ncRNA.18487127.18500594-GGTLC3.18516335.18518165.rev.compl.REF     ------------------------------------------------------------ 31039

chimp.LOC112206744-LOC107973052-GGT2.revcompl                                    gggacatccggatctctcgcaggcagcatcccaggcacagtccctgactcgctttacaga 50878
chimp54546-135457.revcompl.Pan.troglodytes.clone.rp43-41g5.GenBank:AC099533.36   TG---------------------------------------------------------- 54629
human.FAM30B-LOC105372935-GGT2.NCBI.                                             gggacatcgggatctctagcaggcagcatcccaggcacagtccctgactcatttacaga 49862
human.FAM230E-LOC105377182-GGT3P.NCBI.                                           gggacatcgggatctctcgcaggcagcatcccaggcacagtccctgactcgctttacaga 48515
human.FAM230J-LOC105372942-GGTLC5P.NCBI.rev.compl                                ------------------------------------------------------------ 46364
human.FAM230A.ncRNA.18487127.18500594-GGTLC3.18516335.18518165.rev.compl.REF     ------------------------------------------------------------ 31039

chimp.LOC112206744-LOC107973052-GGT2.revcompl                                    tggggcaatgaggcttaggaggaaagatttttttttctttttttgagttggggtcttgcca 50938
chimp54546-135457.revcompl.Pan.troglodytes.clone.rp43-41g5.GenBank:AC099533.36   ---------GTGTCATCCAGGGACCCCACACACTTCCTTAAATGTGATCCTGCTTCCCTC 54680
human.FAM30B-LOC105372935-GGT2.NCBI.                                             tggggcaatgaggcttaggaggaaagatttttttttctttttttgagttggggtcttgcca 49922
human.FAM230E-LOC105377182-GGT3P.NCBI.                                           tggggcaatgaggcttaggaggaaagatttttttttctttttttgagttggggtcttgcca 48575
human.FAM230J-LOC105372942-GGTLC5P.NCBI.rev.compl                                ------------------------------------------------------------ 46364
human.FAM230A.ncRNA.18487127.18500594-GGTLC3.18516335.18518165.rev.compl.REF     ------------------------------------------------------------ 31039

chimp.LOC112206744-LOC107973052-GGT2.revcompl                                    tcttgcccagcctgatctcgaactcctggattcaagcaatcctcccacctcagcctcctg 50998
chimp54546-135457.revcompl.Pan.troglodytes.clone.rp43-41g5.GenBank:AC099533.36   TGGGCAGCTGCATCCTCTCCTCCTGCAGGACTGTCTGGAAACTTGGCTCTCAGTTTGCT- 54739
human.FAM30B-LOC105372935-GGT2.NCBI.                                             tcttgcccagcctgatctcgaactcctggattcaagcaatcctcccacctcagcctcctg 49982
human.FAM230E-LOC105377182-GGT3P.NCBI.                                           tcttgcccagcccaatctcgaactcctggattcaagcaatcctcccacctcagcctcctg 48635
human.FAM230J-LOC105372942-GGTLC5P.NCBI.rev.compl                                ------------------------------------------------------------ 46364
human.FAM230A.ncRNA.18487127.18500594-GGTLC3.18516335.18518165.rev.compl.REF     ------------------------------------------------------------ 31039

chimp.LOC112206744-LOC107973052-GGT2.revcompl                                    agtagctgagattacaggcgtgaacaccacac---ccagcagaaggggatttttaatttt 51055
chimp54546-135457.revcompl.Pan.troglodytes.clone.rp43-41g5.GenBank:AC099533.36   -CTCCCTTCTCTCCTCTGCCTGCCCAAGCCCCTCTTTCTAAAAAAGTGATGCCATGTTCA 54798
human.FAM30B-LOC105372935-GGT2.NCBI.                                             agtcgctgagattacaggcgtgaacagcacac---ctagcagaaggggatttttaatttt 50039
human.FAM230E-LOC105377182-GGT3P.NCBI.                                           agtcgctgagattacaggtgtgaacagcacac---ctagcagaaggggatttttaatttt 48692
human.FAM230J-LOC105372942-GGTLC5P.NCBI.rev.compl                                ------------------------------------------------------------ 46364
human.FAM230A.ncRNA.18487127.18500594-GGTLC3.18516335.18518165.rev.compl.REF     ------------------------------------------------------------ 31039

chimp.LOC112206744-LOC107973052-GGT2.revcompl                                    ttaatttatttttaattttaatttttttttttttaggaggggatgtttaattttttttt- 51114
chimp54546-135457.revcompl.Pan.troglodytes.clone.rp43-41g5.GenBank:AC099533.36   TGGGGTTATTTCTTGAAAATAC--------TTGGCGGCCTCCATGCTTCTGTTTTCTTTG 54850
human.FAM30B-LOC105372935-GGT2.NCBI.                                             taaatttatttttaattttaatttttttttttttaggaggggatgtttaatttttttttt 50099
human.FAM230E-LOC105377182-GGT3P.NCBI.                                           taaatttatttttaattttaattgtttttttttaggaggggatgtttaatttttttttt 48752
human.FAM230J-LOC105372942-GGTLC5P.NCBI.rev.compl                                ------------------------------------------------------------ 46364
human.FAM230A.ncRNA.18487127.18500594-GGTLC3.18516335.18518165.rev.compl.REF     ------------------------------------------------------------ 31039

chimp.LOC112206744-LOC107973052-GGT2.revcompl                                    -aggaggggctcagca-gtaggagtgtacatggacaagggatgtctgaggagggcacagg 51172
chimp54546-135457.revcompl.Pan.troglodytes.clone.rp43-41g5.GenBank:AC099533.36   AGCCAGGTGGTCAGGAGGGCTTACAAAGAATGCCTGGGCTCCCCCGCAGGTGTCAGCAGA 54910
human.FAM30B-LOC105372935-GGT2.NCBI.                                             taggaggggctcagcaggtaggagtgtacatggaccagggatgtctgaggagggcacagc 50159
human.FAM230E-LOC105377182-GGT3P.NCBI.                                           taggaggggctcagcaggtaggagtgtacatggaccagggatgtctgaggagggcacagc 48812
human.FAM230J-LOC105372942-GGTLC5P.NCBI.rev.compl                                ------------------------------------------------------------ 46364
human.FAM230A.ncRNA.18487127.18500594-GGTLC3.18516335.18518165.rev.compl.REF     ------------------------------------------------------------ 31039

chimp.LOC112206744-LOC107973052-GGT2.revcompl                                    aggggaagcagtagcatgcggctgggttttgctgtcccaggatgaggtgtctgtctgtgc 51232
chimp54546-135457.revcompl.Pan.troglodytes.clone.rp43-41g5.GenBank:AC099533.36   TGGGGTAGC----GAATGGTCCTGTGCCTCCACCTGCTCCGGGAGGAGTCT-------- 54958
human.FAM30B-LOC105372935-GGT2.NCBI.                                             aggggaagcagtagcatgcggctgggttttgctgtcccaggatgaggtgtctgtctgtac 50219
human.FAM230E-LOC105377182-GGT3P.NCBI.                                           aggggaagcagtagcatgcggctgggttttgctgtcccaggatgaggtgtctgtctgtgc 48872
```

```
human.FAM230J-LOC105372942-GGTLC5P.NCBI.rev.compl                                 ------------------------------------------------------------ 46364
human.FAM230A.ncRNA.18487127.18500594-GGTLC3.18516335.18518165.rev.compl.REF       ------------------------------------------------------------ 31039

chimp.LOC112206744-LOC107973052-GGT2.revcompl                                     aggtgcctgcatgtctaaaatcctgtgccaggccagacccccctcccatctcgctgaccac 51292
chimp54546-135457.revcompl.Pan.troglodytes.clone.rp43-41g5.GenBank:AC099533.36   ---------CCTGTCTCTAGGCCTGGCCCCTTCCTAACCCTCCACGTATCCTGTTCTCCA 55009
human.FAM30B-LOC105372935-GGT2.NCBI.                                              aggtgcctgcatgtctaaaatcctgtgccaggccagacccccctcccatctcgctgaccac 50279
human.FAM230E-LOC105377182-GGT3P.NCBI.                                            aggtgcctgcatgtctaaaatcctgtgccaggccagacccccctcccatctcgctgaccac 48932
human.FAM230J-LOC105372942-GGTLC5P.NCBI.rev.compl                                 ------------------------------------------------------------ 46364
human.FAM230A.ncRNA.18487127.18500594-GGTLC3.18516335.18518165.rev.compl.REF       ------------------------------------------------------------ 31039

chimp.LOC112206744-LOC107973052-GGT2.revcompl                                     aaggccttatcctgtaagactcatgggctccaccagaatgtgccaaaacaagagcagatc 51352
chimp54546-135457.revcompl.Pan.troglodytes.clone.rp43-41g5.GenBank:AC099533.36   GAGACTTCAGAATCCACTCCTGAGAACAGCGGAGCCAGGCGCTTAGAGGAAGACCAAATG 55069
human.FAM30B-LOC105372935-GGT2.NCBI.                                              aaggccttatcctgtaagactcatgggctccaccagaatgtgccaaaacaagagcagatc 50339
human.FAM230E-LOC105377182-GGT3P.NCBI.                                            aaggccttatcctgtaagactcatgggctccaccagaatgtgccaaaacaagagcagatc 48992
human.FAM230J-LOC105372942-GGTLC5P.NCBI.rev.compl                                 ------------------------------------------------------------ 46364
human.FAM230A.ncRNA.18487127.18500594-GGTLC3.18516335.18518165.rev.compl.REF       ------------------------------------------------------------ 31039

chimp.LOC112206744-LOC107973052-GGT2.revcompl                                     ccaccctgacccaggtcaagcacaggccaccttcaagacac------------------- 51393
chimp54546-135457.revcompl.Pan.troglodytes.clone.rp43-41g5.GenBank:AC099533.36   CTGCCAGGACACGGATTGTCCAGGGATTACATTCCAGCATCTTATTAGGTATCTGGATCT 55129
human.FAM30B-LOC105372935-GGT2.NCBI.                                              ccaccctgacccaggtcaagcacaggccaccttcaagacac------------------- 50380
human.FAM230E-LOC105377182-GGT3P.NCBI.                                            ccaccttgaccgaggtcaagcacaggccaccttcaagacac------------------- 49033
human.FAM230J-LOC105372942-GGTLC5P.NCBI.rev.compl                                 ------------------------------------------------------------ 46364
human.FAM230A.ncRNA.18487127.18500594-GGTLC3.18516335.18518165.rev.compl.REF       ------------------------------------------------------------ 31039

chimp.LOC112206744-LOC107973052-GGT2.revcompl                                     ------------------------------------------------------------ 51393
chimp54546-135457.revcompl.Pan.troglodytes.clone.rp43-41g5.GenBank:AC099533.36   GTTGGGGAAAAAATTAGAAACTACGTATAAAACTTAAAAATATTCAAGCATCAAAAGGTT 55189
human.FAM30B-LOC105372935-GGT2.NCBI.                                              ------------------------------------------------------------ 50380
human.FAM230E-LOC105377182-GGT3P.NCBI.                                            ------------------------------------------------------------ 49033
human.FAM230J-LOC105372942-GGTLC5P.NCBI.rev.compl                                 ------------------------------------------------------------ 46364
human.FAM230A.ncRNA.18487127.18500594-GGTLC3.18516335.18518165.rev.compl.REF       ------------------------------------------------------------ 31039

chimp.LOC112206744-LOC107973052-GGT2.revcompl                                     ------------------------------------------------------------ 51393
chimp54546-135457.revcompl.Pan.troglodytes.clone.rp43-41g5.GenBank:AC099533.36   ATTTAGGATGAAAGTTTTAAAACAAGTCATCAGCAAGCTGCTGCCACCAAGTGGAGACTT 55249
human.FAM30B-LOC105372935-GGT2.NCBI.                                              ------------------------------------------------------------ 50380
human.FAM230E-LOC105377182-GGT3P.NCBI.                                            ------------------------------------------------------------ 49033
human.FAM230J-LOC105372942-GGTLC5P.NCBI.rev.compl                                 ------------------------------------------------------------ 46364
human.FAM230A.ncRNA.18487127.18500594-GGTLC3.18516335.18518165.rev.compl.REF       ------------------------------------------------------------ 31039

chimp.LOC112206744-LOC107973052-GGT2.revcompl                                     ------------------------------------------------------------ 51393
chimp54546-135457.revcompl.Pan.troglodytes.clone.rp43-41g5.GenBank:AC099533.36   ATACAAAAGTTGAGCGAGTCCACTGAGCTGAGAGGACAGAAATGAAGTCACCTGTGCTGG 55309
human.FAM30B-LOC105372935-GGT2.NCBI.                                              ------------------------------------------------------------ 50380
human.FAM230E-LOC105377182-GGT3P.NCBI.                                            ------------------------------------------------------------ 49033
human.FAM230J-LOC105372942-GGTLC5P.NCBI.rev.compl                                 ------------------------------------------------------------ 46364
human.FAM230A.ncRNA.18487127.18500594-GGTLC3.18516335.18518165.rev.compl.REF       ------------------------------------------------------------ 31039

chimp.LOC112206744-LOC107973052-GGT2.revcompl                                     ------------------------------------------------------------ 51393
chimp54546-135457.revcompl.Pan.troglodytes.clone.rp43-41g5.GenBank:AC099533.36   GGCAGGGGCAGGGACACTGGGAGCAGGGAGTGTGTGGGCAGAGAAGCCAGAGAAGTCCAG 55369
human.FAM30B-LOC105372935-GGT2.NCBI.                                              ------------------------------------------------------------ 50380
human.FAM230E-LOC105377182-GGT3P.NCBI.                                            ------------------------------------------------------------ 49033
human.FAM230J-LOC105372942-GGTLC5P.NCBI.rev.compl                                 ------------------------------------------------------------ 46364
human.FAM230A.ncRNA.18487127.18500594-GGTLC3.18516335.18518165.rev.compl.REF       ------------------------------------------------------------ 31039

chimp.LOC112206744-LOC107973052-GGT2.revcompl                                     ------------------------------------------------------------ 51393
chimp54546-135457.revcompl.Pan.troglodytes.clone.rp43-41g5.GenBank:AC099533.36   GCCTGTGGAAGCCAAACAGGAGAGCGTGGGCCGGAAGGGCGGTCAGAATCGGGGGATGAG 55429
human.FAM30B-LOC105372935-GGT2.NCBI.                                              ------------------------------------------------------------ 50380
human.FAM230E-LOC105377182-GGT3P.NCBI.                                            ------------------------------------------------------------ 49033
human.FAM230J-LOC105372942-GGTLC5P.NCBI.rev.compl                                 ------------------------------------------------------------ 46364
human.FAM230A.ncRNA.18487127.18500594-GGTLC3.18516335.18518165.rev.compl.REF       ------------------------------------------------------------ 31039

chimp.LOC112206744-LOC107973052-GGT2.revcompl                                     ------------------------------------------------------------ 51393
chimp54546-135457.revcompl.Pan.troglodytes.clone.rp43-41g5.GenBank:AC099533.36   GTCGCTCTCCCTGGAGAATGAACCCTAAAGTGCGTAGCCTGGGATTCCCTCCCTGGGGGT 55489
human.FAM30B-LOC105372935-GGT2.NCBI.                                              ------------------------------------------------------------ 50380
human.FAM230E-LOC105377182-GGT3P.NCBI.                                            ------------------------------------------------------------ 49033
human.FAM230J-LOC105372942-GGTLC5P.NCBI.rev.compl                                 ------------------------------------------------------------ 46364
human.FAM230A.ncRNA.18487127.18500594-GGTLC3.18516335.18518165.rev.compl.REF       ------------------------------------------------------------ 31039

chimp.LOC112206744-LOC107973052-GGT2.revcompl                                     ------------------------------------------------------------ 51393
chimp54546-135457.revcompl.Pan.troglodytes.clone.rp43-41g5.GenBank:AC099533.36   CCTGTCCCCCGACATTTCACGGGCCTTCTGAGCTGCTTTCCAAGGAGGACTAACACAGCA 55549
human.FAM30B-LOC105372935-GGT2.NCBI.                                              ------------------------------------------------------------ 50380
human.FAM230E-LOC105377182-GGT3P.NCBI.                                            ------------------------------------------------------------ 49033
human.FAM230J-LOC105372942-GGTLC5P.NCBI.rev.compl                                 ------------------------------------------------------------ 46364
human.FAM230A.ncRNA.18487127.18500594-GGTLC3.18516335.18518165.rev.compl.REF       ------------------------------------------------------------ 31039

chimp.LOC112206744-LOC107973052-GGT2.revcompl                                     ------------------------------------------------------------ 51393
chimp54546-135457.revcompl.Pan.troglodytes.clone.rp43-41g5.GenBank:AC099533.36   ACAAAACATCCATTTCTGCACAAAAATCCTTCTGGAAAGAAAAAGAAGCAAGCCAAGAAT 55609
human.FAM30B-LOC105372935-GGT2.NCBI.                                              ------------------------------------------------------------ 50380
human.FAM230E-LOC105377182-GGT3P.NCBI.                                            ------------------------------------------------------------ 49033
human.FAM230J-LOC105372942-GGTLC5P.NCBI.rev.compl                                 ------------------------------------------------------------ 46364
human.FAM230A.ncRNA.18487127.18500594-GGTLC3.18516335.18518165.rev.compl.REF       ------------------------------------------------------------ 31039

chimp.LOC112206744-LOC107973052-GGT2.revcompl                                     ------------------------------------------------------------ 51393
chimp54546-135457.revcompl.Pan.troglodytes.clone.rp43-41g5.GenBank:AC099533.36   GGAGTGAAAACGCTACCCAGTGCTGACTAAGCCTCTCAAACCCTGTTCTAAGTGGAGTGT 55669
human.FAM30B-LOC105372935-GGT2.NCBI.                                              ------------------------------------------------------------ 50380
human.FAM230E-LOC105377182-GGT3P.NCBI.                                            ------------------------------------------------------------ 49033
human.FAM230J-LOC105372942-GGTLC5P.NCBI.rev.compl                                 ------------------------------------------------------------ 46364
human.FAM230A.ncRNA.18487127.18500594-GGTLC3.18516335.18518165.rev.compl.REF       ------------------------------------------------------------ 31039

chimp.LOC112206744-LOC107973052-GGT2.revcompl                                     ------------------------------------------------------------ 51393
chimp54546-135457.revcompl.Pan.troglodytes.clone.rp43-41g5.GenBank:AC099533.36   GGTTTCTAAGTCAGGGAAATGGAAGAGGCCCCACCCACACAGGGACAGGGCCACGGCCCC 55729
human.FAM30B-LOC105372935-GGT2.NCBI.                                              ------------------------------------------------------------ 50380
human.FAM230E-LOC105377182-GGT3P.NCBI.                                            ------------------------------------------------------------ 49033
human.FAM230J-LOC105372942-GGTLC5P.NCBI.rev.compl                                 ------------------------------------------------------------ 46364
human.FAM230A.ncRNA.18487127.18500594-GGTLC3.18516335.18518165.rev.compl.REF       ------------------------------------------------------------ 31039

chimp.LOC112206744-LOC107973052-GGT2.revcompl                                     ------------------------agccagccccaagaaagggctcccttcctcttttct 51429
chimp54546-135457.revcompl.Pan.troglodytes.clone.rp43-41g5.GenBank:AC099533.36   CACAGGATGAAGCAGCAGCGTTTATTCAAGATACAAGTGAGGGAATCCGGTCACGTTCCC 55789
human.FAM30B-LOC105372935-GGT2.NCBI.                                              ------------------------agccagccccaagaaagggctcccttcctcttttct 50416
human.FAM230E-LOC105377182-GGT3P.NCBI.                                            ------------------------agccagccccaagaaagggctccctgcctcttttct 49069
human.FAM230J-LOC105372942-GGTLC5P.NCBI.rev.compl                                 ------------------------------------------------------------ 46364
human.FAM230A.ncRNA.18487127.18500594-GGTLC3.18516335.18518165.rev.compl.REF       ------------------------------------------------------------ 31039

chimp.LOC112206744-LOC107973052-GGT2.revcompl                                     actgccccagagaggcaagactgagccttaacctccg----------------------- 51466
chimp54546-135457.revcompl.Pan.troglodytes.clone.rp43-41g5.GenBank:AC099533.36   TTCTCCCCAGAGAGGGCACTTCTTGACAAGTGATTCAGTAGAAATCTTTTAGACTCTATA 55849
human.FAM30B-LOC105372935-GGT2.NCBI.                                              actgccccagagaggcaagactgagccttaacctcca----------------------- 50453
human.FAM230E-LOC105377182-GGT3P.NCBI.                                            actgccccagagaggcaagactgagccttaatctccg----------------------- 49106
human.FAM230J-LOC105372942-GGTLC5P.NCBI.rev.compl                                 ------------------------------------------------------------ 46364
human.FAM230A.ncRNA.18487127.18500594-GGTLC3.18516335.18518165.rev.compl.REF       ------------------------------------------------------------ 31039

chimp.LOC112206744-LOC107973052-GGT2.revcompl                                     ------------------------tcctgtcccctctcccagcctcagtttctccatcca 51502
chimp54546-135457.revcompl.Pan.troglodytes.clone.rp43-41g5.GenBank:AC099533.36   AGTTAAGTTCATAAAAACCACTGCTTTCACCCTGTCTCCCAGGGCCAGGCCTGGACTCCA 55909
human.FAM30B-LOC105372935-GGT2.NCBI.                                              ------------------------tcctgtcccctctcccagcctcagtttctccaacca 50489
human.FAM230E-LOC105377182-GGT3P.NCBI.                                            ------------------------tcctgtcccctctcccagcctcagtttctgcatcca 49142
human.FAM230J-LOC105372942-GGTLC5P.NCBI.rev.compl                                 ------------------------------------------------------------ 46364
human.FAM230A.ncRNA.18487127.18500594-GGTLC3.18516335.18518165.rev.compl.REF       ------------------------------------------------------------ 31039

chimp.LOC112206744-LOC107973052-GGT2.revcompl                                     actataagggtttttgtttg----tctgttttgaga-cagggtctcattctgttgtccca 51557
chimp54546-135457.revcompl.Pan.troglodytes.clone.rp43-41g5.GenBank:AC099533.36   AGATGAACTGGCTTGGGGTGCCCTCGGGTGGCCACATAAAAAACTCACAGTCTGAGGCCA 55969
human.FAM30B-LOC105372935-GGT2.NCBI.                                              actataagggtttttgtttgtttgtctgttttgaga-cagggtctcactctgttgtccca 50548
human.FAM230E-LOC105377182-GGT3P.NCBI.                                            actataagggtttttgtttgtttgtctgttttgagaccagggtctcattctgttgtccca 49202
human.FAM230J-LOC105372942-GGTLC5P.NCBI.rev.compl                                 ------------------------------------------------------------ 46364
human.FAM230A.ncRNA.18487127.18500594-GGTLC3.18516335.18518165.rev.compl.REF       ------------------------------------------------------------ 31039

chimp.LOC112206744-LOC107973052-GGT2.revcompl                                     gctggaa----------------------------------------------------- 51564
chimp54546-135457.revcompl.Pan.troglodytes.clone.rp43-41g5.GenBank:AC099533.36   GCCTGGGGCTTTCAGACCTGGGCGGGATTTGCCCAGGCCACCTGTCCTTCTGCTTTGGGC 56029
human.FAM30B-LOC105372935-GGT2.NCBI.                                              gctggag----------------------------------------------------- 50555
human.FAM230E-LOC105377182-GGT3P.NCBI.                                            gctggag----------------------------------------------------- 49209
human.FAM230J-LOC105372942-GGTLC5P.NCBI.rev.compl                                 ------------------------------------------------------------ 46364
```

```
human.FAM230A.ncRNA.18487127.18500594-GGTLC3.18516335.18518165.rev.compl.REF        ------------------------------------------------------------ 31039

chimp.LOC112206744-LOC107973052-GGT2.revcompl                                       ------------------------------------------------------------ 51564
chimp54546-135457.revcompl.Pan.troglodytes.clone.rp43-41g5.GenBank:AC099533.36      CGCTGTCTCTTGGCAGATGGCCTGACACCTGGGGGTGGCCCAAGGATGCCTCAGAAAATC 56089
human.FAM30B-LOC105372935-GGT2.NCBI.                                                ------------------------------------------------------------ 50555
human.FAM230E-LOC105377182-GGT3P.NCBI.                                              ------------------------------------------------------------ 49209
human.FAM230J-LOC105372942-GGTLC5P.NCBI.rev.compl                                   ------------------------------------------------------------ 46364
human.FAM230A.ncRNA.18487127.18500594-GGTLC3.18516335.18518165.rev.compl.REF        ------------------------------------------------------------ 31039

chimp.LOC112206744-LOC107973052-GGT2.revcompl                                       ------------------------------------------------------------ 51564
chimp54546-135457.revcompl.Pan.troglodytes.clone.rp43-41g5.GenBank:AC099533.36      TTGATTCCCACTCTACAGATGGCCTGATTAGCCAGAGGTTTCCAGGCCGTCTGTCCGCCT 56149
human.FAM30B-LOC105372935-GGT2.NCBI.                                                ------------------------------------------------------------ 50555
human.FAM230E-LOC105377182-GGT3P.NCBI.                                              ------------------------------------------------------------ 49209
human.FAM230J-LOC105372942-GGTLC5P.NCBI.rev.compl                                   ------------------------------------------------------------ 46364
human.FAM230A.ncRNA.18487127.18500594-GGTLC3.18516335.18518165.rev.compl.REF        ------------------------------------------------------------ 31039

chimp.LOC112206744-LOC107973052-GGT2.revcompl                                       --------tgcagtggtgcaatcatggctcactgcagccttggcttcccaggctcaagcg 51616
chimp54546-135457.revcompl.Pan.troglodytes.clone.rp43-41g5.GenBank:AC099533.36      CCAGGAGATGGACTGGGACCTTTAGACATCGGTGGAGAACAGGATGCTCTGTCCCTTGCT 56209
human.FAM30B-LOC105372935-GGT2.NCBI.                                                --------tgcagtggtgcaatcatggctcactgcagccttggcttcccaggctcaagcg 50607
human.FAM230E-LOC105377182-GGT3P.NCBI.                                              --------tgcagtggtgcaatcatggctcactgcagccttggcttcccaggctcaagcg 49261
human.FAM230J-LOC105372942-GGTLC5P.NCBI.rev.compl                                   ------------------------------------------------------------ 46364
human.FAM230A.ncRNA.18487127.18500594-GGTLC3.18516335.18518165.rev.compl.REF        ------------------------------------------------------------ 31039

chimp.LOC112206744-LOC107973052-GGT2.revcompl                                       atcctcccacctcagcctctgaagtacctaagactacagacacaccccac---------- 51666
chimp54546-135457.revcompl.Pan.troglodytes.clone.rp43-41g5.GenBank:AC099533.36      GTCCAGGGCAGGGATGGCCTCCAGCCGCAAGAAGTACAGCAGCACCTCGACCTGCCCTCG 56269
human.FAM30B-LOC105372935-GGT2.NCBI.                                                atcctcccacttcagcctctgaagtacctaagactacagacataccccac---------- 50657
human.FAM230E-LOC105377182-GGT3P.NCBI.                                              atcctcccacctcagcctctgaagtacctaagactacagacataccccac---------- 49311
human.FAM230J-LOC105372942-GGTLC5P.NCBI.rev.compl                                   ------------------------------------------------------------ 46364
human.FAM230A.ncRNA.18487127.18500594-GGTLC3.18516335.18518165.rev.compl.REF        ------------------------------------------------------------ 31039

chimp.LOC112206744-LOC107973052-GGT2.revcompl                                       ------------------------------------------------------------ 51666
chimp54546-135457.revcompl.Pan.troglodytes.clone.rp43-41g5.GenBank:AC099533.36      CAGAGTGGGGAAGAGGAGAGTGGCTCGGAGGGGGGCTCACAGCTGCTGGTGGGGAGGTCT 56329
human.FAM30B-LOC105372935-GGT2.NCBI.                                                ------------------------------------------------------------ 50657
human.FAM230E-LOC105377182-GGT3P.NCBI.                                              ------------------------------------------------------------ 49311
human.FAM230J-LOC105372942-GGTLC5P.NCBI.rev.compl                                   ------------------------------------------------------------ 46364
human.FAM230A.ncRNA.18487127.18500594-GGTLC3.18516335.18518165.rev.compl.REF        ------------------------------------------------------------ 31039

chimp.LOC112206744-LOC107973052-GGT2.revcompl                                       ------------------------------------------------------------ 51666
chimp54546-135457.revcompl.Pan.troglodytes.clone.rp43-41g5.GenBank:AC099533.36      TTGGGGCCCAAGCCCCCAAGTCCACCTCAGGTGCTAGAAACCCCTGCTGGTGTCATCAAC 56389
human.FAM30B-LOC105372935-GGT2.NCBI.                                                ------------------------------------------------------------ 50657
human.FAM230E-LOC105377182-GGT3P.NCBI.                                              ------------------------------------------------------------ 49311
human.FAM230J-LOC105372942-GGTLC5P.NCBI.rev.compl                                   ------------------------------------------------------------ 46364
human.FAM230A.ncRNA.18487127.18500594-GGTLC3.18516335.18518165.rev.compl.REF        ------------------------------------------------------------ 31039

chimp.LOC112206744-LOC107973052-GGT2.revcompl                                       ------------------------------------------------tacacgtggctt 51678
chimp54546-135457.revcompl.Pan.troglodytes.clone.rp43-41g5.GenBank:AC099533.36      CCCTTACAGTGAGACGGGGGTGGGGTGGGGTCCTGACAAGGCATGACTTGTTGGGTGGGG 56449
human.FAM30B-LOC105372935-GGT2.NCBI.                                                ------------------------------------------------tgcacgtggctt 50669
human.FAM230E-LOC105377182-GGT3P.NCBI.                                              ------------------------------------------------tgcacgtggc-t 49322
human.FAM230J-LOC105372942-GGTLC5P.NCBI.rev.compl                                   ------------------------------------------------------------ 46364
human.FAM230A.ncRNA.18487127.18500594-GGTLC3.18516335.18518165.rev.compl.REF        ------------------------------------------------------------ 31039

chimp.LOC112206744-LOC107973052-GGT2.revcompl                                       tttttttttttttttt--tgagatggagtttcactcttgttgcccaggctggagtgcaat 51736
chimp54546-135457.revcompl.Pan.troglodytes.clone.rp43-41g5.GenBank:AC099533.36      GGTGGTTATTTATTTTAGAGATGCACAGGGCCTTGCTCTGTCCCCAGGCTGGAGTACAGT 56509
human.FAM30B-LOC105372935-GGT2.NCBI.                                                tttttttttttttttttttgagatggagtttcactcttgttgcccaggctggagtgcaat 50729
human.FAM230E-LOC105377182-GGT3P.NCBI.                                              ttttttttttttttttctgagatggagtttcactcttgttgcccaggctggagtgcaat 49382
human.FAM230J-LOC105372942-GGTLC5P.NCBI.rev.compl                                   ------------------------------------------------------------ 46364
human.FAM230A.ncRNA.18487127.18500594-GGTLC3.18516335.18518165.rev.compl.REF        ------------------------------------------------------------ 31039

chimp.LOC112206744-LOC107973052-GGT2.revcompl                                       ggcacaatcttggctcactccaacctccacctcccgagttcaagcaattctcctgcctca 51796
chimp54546-135457.revcompl.Pan.troglodytes.clone.rp43-41g5.GenBank:AC099533.36      GGCTCCATCATGGATCACTGCAGCCTCTAACTCCTGGGCTCAAGCAATCCTCCTGTGTCA 56569
human.FAM30B-LOC105372935-GGT2.NCBI.                                                ggcacaatcttggctcactccaacctccacctcccgagttcaagtaattctcctgcctca 50789
human.FAM230E-LOC105377182-GGT3P.NCBI.                                              ggcacaatcttggctcactccaacctccacctcccgagttcaagtaattctcctgcctca 49442
human.FAM230J-LOC105372942-GGTLC5P.NCBI.rev.compl                                   ------------------------------------------------------------ 46364
human.FAM230A.ncRNA.18487127.18500594-GGTLC3.18516335.18518165.rev.compl.REF        ------------------------------------------------------------ 31039

chimp.LOC112206744-LOC107973052-GGT2.revcompl                                       gcctcccaagtagctgggattacaggcatgcaccaccacgccttgctaatt----tttgt 51852
chimp54546-135457.revcompl.Pan.troglodytes.clone.rp43-41g5.GenBank:AC099533.36      GCCTCCCAGATACCTAGGATTACAGATGTGTGCCACAATGCCTGCCTAATTTTTCTTTGT 56629
human.FAM30B-LOC105372935-GGT2.NCBI.                                                gcctcccaagtacctgggattacaggcatgtaccaccacgccttgctaatt----tttgt 50845
human.FAM230E-LOC105377182-GGT3P.NCBI.                                              gcctcccaagtagctgggattacaggcatgtaccaccacgccttgctaatt----tttgt 49498
human.FAM230J-LOC105372942-GGTLC5P.NCBI.rev.compl                                   ------------------------------------------------------------ 46364
human.FAM230A.ncRNA.18487127.18500594-GGTLC3.18516335.18518165.rev.compl.REF        ------------------------------------------------------------ 31039

chimp.LOC112206744-LOC107973052-GGT2.revcompl                                       atttttagtggagacggggtttcaccaggttggtcaggctggtcttgaactcctaacatc 51912
chimp54546-135457.revcompl.Pan.troglodytes.clone.rp43-41g5.GenBank:AC099533.36      ATTTTTTCTGGAGATGGGGTTTG-CTGCATTGCCCAGACTGATCTCAAACACTTGGGTTC 56688
human.FAM30B-LOC105372935-GGT2.NCBI.                                                atttttagtagagacggggtttcaccaggttggtcaggctggtcttgaactcctaacatc 50905
human.FAM230E-LOC105377182-GGT3P.NCBI.                                              atttttagtagagacggggtttcaccaggttggtcaggctggtcttgaactcctaacatc 49558
human.FAM230J-LOC105372942-GGTLC5P.NCBI.rev.compl                                   ------------------------------------------------------------ 46364
human.FAM230A.ncRNA.18487127.18500594-GGTLC3.18516335.18518165.rev.compl.REF        ------------------------------------------------------------ 31039

chimp.LOC112206744-LOC107973052-GGT2.revcompl                                       aggtgatccatcctcctcagcctctcagagttctgggattacaggcgtgagccaccactc 51972
chimp54546-135457.revcompl.Pan.troglodytes.clone.rp43-41g5.GenBank:AC099533.36      GGTTGTCCTG----CCTCGGCCTCCCAAAGTGCTGGGATTACAGGCATGAGCCACCACAC 56744
human.FAM30B-LOC105372935-GGT2.NCBI.                                                aggtgatccatcctcctcagcttctcagagttctgggattacaggcgtgagccaccactc 50965
human.FAM230E-LOC105377182-GGT3P.NCBI.                                              aggtgatccatcctcctcagcctctcagagttctgggattacaggcgtgagccaccactc 49618
human.FAM230J-LOC105372942-GGTLC5P.NCBI.rev.compl                                   ------------------------------------------------------------ 46364
human.FAM230A.ncRNA.18487127.18500594-GGTLC3.18516335.18518165.rev.compl.REF        ------------------------------------------------------------ 31039

chimp.LOC112206744-LOC107973052-GGT2.revcompl                                       ccagcctaatttttttatttta-------------------------------------- 51994
chimp54546-135457.revcompl.Pan.troglodytes.clone.rp43-41g5.GenBank:AC099533.36      CCGAACACTTGGGTTGGTTTTAAGCCCCCAGCAAGGTGCACCAGCAGGACCAGGAGGTGG 56804
human.FAM30B-LOC105372935-GGT2.NCBI.                                                ccagcctaatttttttatttta-------------------------------------- 50987
human.FAM230E-LOC105377182-GGT3P.NCBI.                                              ccagcctaatttttttatttta-------------------------------------- 49640
human.FAM230J-LOC105372942-GGTLC5P.NCBI.rev.compl                                   ------------------------------------------------------------ 46364
human.FAM230A.ncRNA.18487127.18500594-GGTLC3.18516335.18518165.rev.compl.REF        ------------------------------------------------------------ 31039

chimp.LOC112206744-LOC107973052-GGT2.revcompl                                       ------------------------------------------------------------ 51994
chimp54546-135457.revcompl.Pan.troglodytes.clone.rp43-41g5.GenBank:AC099533.36      CCTGGGCACCCCCTATCACTCCCATCCATGCAAACCTAGGCAAGTCCCTGTCTCTGAATC 56864
human.FAM30B-LOC105372935-GGT2.NCBI.                                                ------------------------------------------------------------ 50987
human.FAM230E-LOC105377182-GGT3P.NCBI.                                              ------------------------------------------------------------ 49640
human.FAM230J-LOC105372942-GGTLC5P.NCBI.rev.compl                                   ------------------------------------------------------------ 46364
human.FAM230A.ncRNA.18487127.18500594-GGTLC3.18516335.18518165.rev.compl.REF        ------------------------------------------------------------ 31039

chimp.LOC112206744-LOC107973052-GGT2.revcompl                                       ------------------------------------------------------------ 51994
chimp54546-135457.revcompl.Pan.troglodytes.clone.rp43-41g5.GenBank:AC099533.36      TCAGCCACCACCACATACAATGCAAGTCGGAAGATGGGCAGGACTGGGGGTGGGGCAGGC 56924
human.FAM30B-LOC105372935-GGT2.NCBI.                                                ------------------------------------------------------------ 50987
human.FAM230E-LOC105377182-GGT3P.NCBI.                                              ------------------------------------------------------------ 49640
human.FAM230J-LOC105372942-GGTLC5P.NCBI.rev.compl                                   ------------------------------------------------------------ 46364
human.FAM230A.ncRNA.18487127.18500594-GGTLC3.18516335.18518165.rev.compl.REF        ------------------------------------------------------------ 31039

chimp.LOC112206744-LOC107973052-GGT2.revcompl                                       ------------------------------------------------------------ 51994
chimp54546-135457.revcompl.Pan.troglodytes.clone.rp43-41g5.GenBank:AC099533.36      AGAGGCCACCTCTGTCAGGCTGGGGTTGCATGGGCTGGAGACTGTCTTCCCATACCTGGG 56984
human.FAM30B-LOC105372935-GGT2.NCBI.                                                ------------------------------------------------------------ 50987
human.FAM230E-LOC105377182-GGT3P.NCBI.                                              ------------------------------------------------------------ 49640
human.FAM230J-LOC105372942-GGTLC5P.NCBI.rev.compl                                   ------------------------------------------------------------ 46364
human.FAM230A.ncRNA.18487127.18500594-GGTLC3.18516335.18518165.rev.compl.REF        ------------------------------------------------------------ 31039

chimp.LOC112206744-LOC107973052-GGT2.revcompl                                       -----------------------------------------ttttttgtagagacagggg 52013
chimp54546-135457.revcompl.Pan.troglodytes.clone.rp43-41g5.GenBank:AC099533.36      ACATGACCTCCAAGGACCTGCTGTCAGTCATGGTGATGGGCTGGTTGGGGTTGGCAGGGA 57044
human.FAM30B-LOC105372935-GGT2.NCBI.                                                -----------------------------------------ttttttgtagagacagggg 51006
human.FAM230E-LOC105377182-GGT3P.NCBI.                                              -----------------------------------------ttttttgtagagacagggg 49659
human.FAM230J-LOC105372942-GGTLC5P.NCBI.rev.compl                                   ------------------------------------------------------------ 46364
human.FAM230A.ncRNA.18487127.18500594-GGTLC3.18516335.18518165.rev.compl.REF        ------------------------------------------------------------ 31039

chimp.LOC112206744-LOC107973052-GGT2.revcompl                                       tcttgctacgttgccaaga----------------------------------------- 52032
chimp54546-135457.revcompl.Pan.troglodytes.clone.rp43-41g5.GenBank:AC099533.36      GCTTGCTCTCCTTCTCGGAGGGCCAGAGCAGCGTGGGGCCAAAGACAGTGGCGAGGTTGT 57104
human.FAM30B-LOC105372935-GGT2.NCBI.                                                tcttgctacattgccaaga----------------------------------------- 51025
human.FAM230E-LOC105377182-GGT3P.NCBI.                                              tcttgctacgttgccaaga----------------------------------------- 49678
human.FAM230J-LOC105372942-GGTLC5P.NCBI.rev.compl                                   ------------------------------------------------------------ 46364
human.FAM230A.ncRNA.18487127.18500594-GGTLC3.18516335.18518165.rev.compl.REF        ------------------------------------------------------------ 31039
```

```
chimp.LOC112206744-LOC107973052-GGT2.revcompl                                      ------------------------------------------------------------ 52032
chimp54546-135457.revcompl.Pan.troglodytes.clone.rp43-41g5.GenBank:AC099533.36     GCAGGGACATCTTATTGACTGCCTCCTTCTCTGCCACCCTGTAGAGGACCGAAGCAGAGG 57164
human.FAM30B-LOC105372935-GGT2.NCBI.                                               ------------------------------------------------------------ 51025
human.FAM230E-LOC105377182-GGT3P.NCBI.                                             ------------------------------------------------------------ 49678
human.FAM230J-LOC105372942-GGTLC5P.NCBI.rev.compl                                  ------------------------------------------------------------ 46364
human.FAM230A.ncRNA.18487127.18500594-GGTLC3.18516335.18518165.rev.compl.REF       ------------------------------------------------------------ 31039

chimp.LOC112206744-LOC107973052-GGT2.revcompl                                      ------------------------------------------------------------ 52032
chimp54546-135457.revcompl.Pan.troglodytes.clone.rp43-41g5.GenBank:AC099533.36     GTGCTGTTTCAATGCCACCACCAGGAGAGAGGCAGAGGGGCTGTGCCGTGCTAGAGTCCT 57224
human.FAM30B-LOC105372935-GGT2.NCBI.                                               ------------------------------------------------------------ 51025
human.FAM230E-LOC105377182-GGT3P.NCBI.                                             ------------------------------------------------------------ 49678
human.FAM230J-LOC105372942-GGTLC5P.NCBI.rev.compl                                  ------------------------------------------------------------ 46364
human.FAM230A.ncRNA.18487127.18500594-GGTLC3.18516335.18518165.rev.compl.REF       ------------------------------------------------------------ 31039

chimp.LOC112206744-LOC107973052-GGT2.revcompl                                      ------------------------------------------------------------ 52032
chimp54546-135457.revcompl.Pan.troglodytes.clone.rp43-41g5.GenBank:AC099533.36     CAGGGAGGGAGTGACCTCGACCCTGGCTGTGCTGCAAGCTGACTCCAGCCTTGGTACTTC 57284
human.FAM30B-LOC105372935-GGT2.NCBI.                                               ------------------------------------------------------------ 51025
human.FAM230E-LOC105377182-GGT3P.NCBI.                                             ------------------------------------------------------------ 49678
human.FAM230J-LOC105372942-GGTLC5P.NCBI.rev.compl                                  ------------------------------------------------------------ 46364
human.FAM230A.ncRNA.18487127.18500594-GGTLC3.18516335.18518165.rev.compl.REF       ------------------------------------------------------------ 31039

chimp.LOC112206744-LOC107973052-GGT2.revcompl                                      ------------------------------------------------------------ 52032
chimp54546-135457.revcompl.Pan.troglodytes.clone.rp43-41g5.GenBank:AC099533.36     TGGGTCTCAGTGGCCCGGGACAAGGGGCCAGCTCTGGGCTGATGGGGAGGTCTTCATGAT 57344
human.FAM30B-LOC105372935-GGT2.NCBI.                                               ------------------------------------------------------------ 51025
human.FAM230E-LOC105377182-GGT3P.NCBI.                                             ------------------------------------------------------------ 49678
human.FAM230J-LOC105372942-GGTLC5P.NCBI.rev.compl                                  ------------------------------------------------------------ 46364
human.FAM230A.ncRNA.18487127.18500594-GGTLC3.18516335.18518165.rev.compl.REF       ------------------------------------------------------------ 31039

chimp.LOC112206744-LOC107973052-GGT2.revcompl                                      ------------------------------------------------------------ 52032
chimp54546-135457.revcompl.Pan.troglodytes.clone.rp43-41g5.GenBank:AC099533.36     GTGCTTGGGAGGGAAGAGGCGGGTCCAAGTGCACTGCTGGCCACGGCCAAAGCTCTGAGC 57404
human.FAM30B-LOC105372935-GGT2.NCBI.                                               ------------------------------------------------------------ 51025
human.FAM230E-LOC105377182-GGT3P.NCBI.                                             ------------------------------------------------------------ 49678
human.FAM230J-LOC105372942-GGTLC5P.NCBI.rev.compl                                  ------------------------------------------------------------ 46364
human.FAM230A.ncRNA.18487127.18500594-GGTLC3.18516335.18518165.rev.compl.REF       ------------------------------------------------------------ 31039

chimp.LOC112206744-LOC107973052-GGT2.revcompl                                      ------------------------------------------------------------ 52032
chimp54546-135457.revcompl.Pan.troglodytes.clone.rp43-41g5.GenBank:AC099533.36     TCTTTGTTAAGGCCACAGTGCAGAGGGAGGAGGGTGGCAAAGAGGAGAGGCAGTGGTGGG 57464
human.FAM30B-LOC105372935-GGT2.NCBI.                                               ------------------------------------------------------------ 51025
human.FAM230E-LOC105377182-GGT3P.NCBI.                                             ------------------------------------------------------------ 49678
human.FAM230J-LOC105372942-GGTLC5P.NCBI.rev.compl                                  ------------------------------------------------------------ 46364
human.FAM230A.ncRNA.18487127.18500594-GGTLC3.18516335.18518165.rev.compl.REF       ------------------------------------------------------------ 31039

chimp.LOC112206744-LOC107973052-GGT2.revcompl                                      ------------------------------------------------------------ 52032
chimp54546-135457.revcompl.Pan.troglodytes.clone.rp43-41g5.GenBank:AC099533.36     GGTGGCAGTGGTGCTAGTCCTTAGAAGCAGTGAGTTACTGCAGACAGGGGTCGGGGGAAA 57524
human.FAM30B-LOC105372935-GGT2.NCBI.                                               ------------------------------------------------------------ 51025
human.FAM230E-LOC105377182-GGT3P.NCBI.                                             ------------------------------------------------------------ 49678
human.FAM230J-LOC105372942-GGTLC5P.NCBI.rev.compl                                  ------------------------------------------------------------ 46364
human.FAM230A.ncRNA.18487127.18500594-GGTLC3.18516335.18518165.rev.compl.REF       ------------------------------------------------------------ 31039

chimp.LOC112206744-LOC107973052-GGT2.revcompl                                      ------------------------------------------------------------ 52032
chimp54546-135457.revcompl.Pan.troglodytes.clone.rp43-41g5.GenBank:AC099533.36     AGGTCCTTGGTGCTGGGGGTCTGGTGGGAGCAAAGCGGCACCCCAACGGCCTGGAGACCT 57584
human.FAM30B-LOC105372935-GGT2.NCBI.                                               ------------------------------------------------------------ 51025
human.FAM230E-LOC105377182-GGT3P.NCBI.                                             ------------------------------------------------------------ 49678
human.FAM230J-LOC105372942-GGTLC5P.NCBI.rev.compl                                  ------------------------------------------------------------ 46364
human.FAM230A.ncRNA.18487127.18500594-GGTLC3.18516335.18518165.rev.compl.REF       ------------------------------------------------------------ 31039

chimp.LOC112206744-LOC107973052-GGT2.revcompl                                      ------------------------------------------------------------ 52032
chimp54546-135457.revcompl.Pan.troglodytes.clone.rp43-41g5.GenBank:AC099533.36     GGAGTCCTGGGCAGCCACAAGAGAGCTGGGCTACCTTTCAAGGTGGTCTAGAAGGAAAAG 57644
human.FAM30B-LOC105372935-GGT2.NCBI.                                               ------------------------------------------------------------ 51025
human.FAM230E-LOC105377182-GGT3P.NCBI.                                             ------------------------------------------------------------ 49678
human.FAM230J-LOC105372942-GGTLC5P.NCBI.rev.compl                                  ------------------------------------------------------------ 46364
human.FAM230A.ncRNA.18487127.18500594-GGTLC3.18516335.18518165.rev.compl.REF       ------------------------------------------------------------ 31039

chimp.LOC112206744-LOC107973052-GGT2.revcompl                                      ------------------------------------------------------------ 52032
chimp54546-135457.revcompl.Pan.troglodytes.clone.rp43-41g5.GenBank:AC099533.36     GAAGGTGAGCAGGTTGGCCTCCGGCAGGGACAACAACAGGTTGAACATGCAGCTCTTCTT 57704
human.FAM30B-LOC105372935-GGT2.NCBI.                                               ------------------------------------------------------------ 51025
human.FAM230E-LOC105377182-GGT3P.NCBI.                                             ------------------------------------------------------------ 49678
human.FAM230J-LOC105372942-GGTLC5P.NCBI.rev.compl                                  ------------------------------------------------------------ 46364
human.FAM230A.ncRNA.18487127.18500594-GGTLC3.18516335.18518165.rev.compl.REF       ------------------------------------------------------------ 31039

chimp.LOC112206744-LOC107973052-GGT2.revcompl                                      ------------------------------------------------------------ 52032
chimp54546-135457.revcompl.Pan.troglodytes.clone.rp43-41g5.GenBank:AC099533.36     TGCAACTGGGTCTGAAAGAGCTGCAGGAGGCAGTGGGTTACTCCTCTGGGTTACGACAAG 57764
human.FAM30B-LOC105372935-GGT2.NCBI.                                               ------------------------------------------------------------ 51025
human.FAM230E-LOC105377182-GGT3P.NCBI.                                             ------------------------------------------------------------ 49678
human.FAM230J-LOC105372942-GGTLC5P.NCBI.rev.compl                                  ------------------------------------------------------------ 46364
human.FAM230A.ncRNA.18487127.18500594-GGTLC3.18516335.18518165.rev.compl.REF       ------------------------------------------------------------ 31039

chimp.LOC112206744-LOC107973052-GGT2.revcompl                                      ------------------------------------------------------------ 52032
chimp54546-135457.revcompl.Pan.troglodytes.clone.rp43-41g5.GenBank:AC099533.36     CCAGAGACCTCTCCCGAGGTGGTCACATGGAGCACCTAGGACACGAGTCCTTGCGCAGTT 57824
human.FAM30B-LOC105372935-GGT2.NCBI.                                               ------------------------------------------------------------ 51025
human.FAM230E-LOC105377182-GGT3P.NCBI.                                             ------------------------------------------------------------ 49678
human.FAM230J-LOC105372942-GGTLC5P.NCBI.rev.compl                                  ------------------------------------------------------------ 46364
human.FAM230A.ncRNA.18487127.18500594-GGTLC3.18516335.18518165.rev.compl.REF       ------------------------------------------------------------ 31039

chimp.LOC112206744-LOC107973052-GGT2.revcompl                                      ------------------------------------------------------------ 52032
chimp54546-135457.revcompl.Pan.troglodytes.clone.rp43-41g5.GenBank:AC099533.36     TGGGCTTGTCATCATCGTCACACCCACAGCGCTGGCCACCAGTGAGGACCCTGTGAGGGG 57884
human.FAM30B-LOC105372935-GGT2.NCBI.                                               ------------------------------------------------------------ 51025
human.FAM230E-LOC105377182-GGT3P.NCBI.                                             ------------------------------------------------------------ 49678
human.FAM230J-LOC105372942-GGTLC5P.NCBI.rev.compl                                  ------------------------------------------------------------ 46364
human.FAM230A.ncRNA.18487127.18500594-GGTLC3.18516335.18518165.rev.compl.REF       ------------------------------------------------------------ 31039

chimp.LOC112206744-LOC107973052-GGT2.revcompl                                      ------------------------------------------------------------ 52032
chimp54546-135457.revcompl.Pan.troglodytes.clone.rp43-41g5.GenBank:AC099533.36     CACCTGTGTGGGGTGTGAACCACCTGAATGCCTTTTCTCTACCTCACAGGGGTCAGCAGC 57944
human.FAM30B-LOC105372935-GGT2.NCBI.                                               ------------------------------------------------------------ 51025
human.FAM230E-LOC105377182-GGT3P.NCBI.                                             ------------------------------------------------------------ 49678
human.FAM230J-LOC105372942-GGTLC5P.NCBI.rev.compl                                  ------------------------------------------------------------ 46364
human.FAM230A.ncRNA.18487127.18500594-GGTLC3.18516335.18518165.rev.compl.REF       ------------------------------------------------------------ 31039

chimp.LOC112206744-LOC107973052-GGT2.revcompl                                      ----------------------------------------ctggtctcaaactctggcct 52052
chimp54546-135457.revcompl.Pan.troglodytes.clone.rp43-41g5.GenBank:AC099533.36     ACCCGGTGAACAGCAGCAGGAGGAGCCGCTAGAGCAGCTGCTGATGGGCAGAGCTGCCCT 58004
human.FAM30B-LOC105372935-GGT2.NCBI.                                               ----------------------------------------ctggtctcaaactctggcct 51045
human.FAM230E-LOC105377182-GGT3P.NCBI.                                             ----------------------------------------ctggtctcaaactctggcct 49698
human.FAM230J-LOC105372942-GGTLC5P.NCBI.rev.compl                                  ------------------------------------------------------------ 46364
human.FAM230A.ncRNA.18487127.18500594-GGTLC3.18516335.18518165.rev.compl.REF       ------------------------------------------------------------ 31039

chimp.LOC112206744-LOC107973052-GGT2.revcompl                                      caagcaa-tcctcccacctcagcctcccaacatgctgggattacaggtgcactcagccta 52111
chimp54546-135457.revcompl.Pan.troglodytes.clone.rp43-41g5.GenBank:AC099533.36     TGGACAACTCCTGCCACCACCCCCTCCCCAGGGAGCCCAAGGCAGGGGAGGCTCAGCATG 58064
human.FAM30B-LOC105372935-GGT2.NCBI.                                               caagcaa-tcctcccacctcagcctcccaacatgctgggattacaggtgcactcagccta 51104
human.FAM230E-LOC105377182-GGT3P.NCBI.                                             caagcaa-tcctcccacctcagcctcccaacatgctgggattacaggtgcactcagccta 49757
human.FAM230J-LOC105372942-GGTLC5P.NCBI.rev.compl                                  ------------------------------------------------------------ 46364
human.FAM230A.ncRNA.18487127.18500594-GGTLC3.18516335.18518165.rev.compl.REF       ------------------------------------------------------------ 31039

chimp.LOC112206744-LOC107973052-GGT2.revcompl                                      taagg------------------------------------------------------- 52116
chimp54546-135457.revcompl.Pan.troglodytes.clone.rp43-41g5.GenBank:AC099533.36     GAATGAAACAGGGGAGTGAGGGACACAAGGAGGTGGGAAGTGGGAGGGTCCCAGCCCCAC 58124
human.FAM30B-LOC105372935-GGT2.NCBI.                                               taagg------------------------------------------------------- 51109
human.FAM230E-LOC105377182-GGT3P.NCBI.                                             taagg------------------------------------------------------- 49762
human.FAM230J-LOC105372942-GGTLC5P.NCBI.rev.compl                                  ------------------------------------------------------------ 46364
human.FAM230A.ncRNA.18487127.18500594-GGTLC3.18516335.18518165.rev.compl.REF       ------------------------------------------------------------ 31039

chimp.LOC112206744-LOC107973052-GGT2.revcompl                                      ------------------------------------------------------------ 52116
chimp54546-135457.revcompl.Pan.troglodytes.clone.rp43-41g5.GenBank:AC099533.36     CAAGTACGCAGAGACCCCCTCGTCGTCCTGGACACCACAGGGGCACCTGCAGGCTGGGAG 58184
human.FAM30B-LOC105372935-GGT2.NCBI.                                               ------------------------------------------------------------ 51109
human.FAM230E-LOC105377182-GGT3P.NCBI.                                             ------------------------------------------------------------ 49762
human.FAM230J-LOC105372942-GGTLC5P.NCBI.rev.compl                                  ------------------------------------------------------------ 46364
human.FAM230A.ncRNA.18487127.18500594-GGTLC3.18516335.18518165.rev.compl.REF       ------------------------------------------------------------ 31039
```

```
chimp.LOC112206744-LOC107973052-GGT2.revcompl                                      ------------------------------------------------------------ 52116
chimp54546-135457.revcompl.Pan.troglodytes.clone.rp43-41g5.GenBank:AC099533.36     ACCAGGTCCTCTGTGCATGGGCCCGGGAGGCAGACCTGCCCTAAGGGTGATGCAGAGGCC 58244
human.FAM30B-LOC105372935-GGT2.NCBI.                                               ------------------------------------------------------------ 51109
human.FAM230E-LOC105377182-GGT3P.NCBI.                                             ------------------------------------------------------------ 49762
human.FAM230J-LOC105372942-GGTLC5P.NCBI.rev.compl                                  ------------------------------------------------------------ 46364
human.FAM230A.ncRNA.18487127.18500594-GGTLC3.18516335.18518165.rev.compl.REF       ------------------------------------------------------------ 31039

chimp.LOC112206744-LOC107973052-GGT2.revcompl                                      ------------------------------------------------------------ 52116
chimp54546-135457.revcompl.Pan.troglodytes.clone.rp43-41g5.GenBank:AC099533.36     ACAGGTGCTGCATGCATCAGCACCCACTCTAGACATCAGCCTCCAGGTTGACTAAGGGTC 58304
human.FAM30B-LOC105372935-GGT2.NCBI.                                               ------------------------------------------------------------ 51109
human.FAM230E-LOC105377182-GGT3P.NCBI.                                             ------------------------------------------------------------ 49762
human.FAM230J-LOC105372942-GGTLC5P.NCBI.rev.compl                                  ------------------------------------------------------------ 46364
human.FAM230A.ncRNA.18487127.18500594-GGTLC3.18516335.18518165.rev.compl.REF       ------------------------------------------------------------ 31039

chimp.LOC112206744-LOC107973052-GGT2.revcompl                                      ------------------------------------------------------------ 52116
chimp54546-135457.revcompl.Pan.troglodytes.clone.rp43-41g5.GenBank:AC099533.36     AGGTCATGTTTGAAACCATGCTTGGCTGGACCAGGACCCATGGCAAGAGCACCTGGGCAC 58364
human.FAM30B-LOC105372935-GGT2.NCBI.                                               ------------------------------------------------------------ 51109
human.FAM230E-LOC105377182-GGT3P.NCBI.                                             ------------------------------------------------------------ 49762
human.FAM230J-LOC105372942-GGTLC5P.NCBI.rev.compl                                  ------------------------------------------------------------ 46364
human.FAM230A.ncRNA.18487127.18500594-GGTLC3.18516335.18518165.rev.compl.REF       ------------------------------------------------------------ 31039

chimp.LOC112206744-LOC107973052-GGT2.revcompl                                      ------------------------------------------------------------ 52116
chimp54546-135457.revcompl.Pan.troglodytes.clone.rp43-41g5.GenBank:AC099533.36     CAGTGTTTAGCCCTGGTCTGCAGGAAGGAGGACAGCAGACTTTAGGACCCCACAGCACGG 58424
human.FAM30B-LOC105372935-GGT2.NCBI.                                               ------------------------------------------------------------ 51109
human.FAM230E-LOC105377182-GGT3P.NCBI.                                             ------------------------------------------------------------ 49762
human.FAM230J-LOC105372942-GGTLC5P.NCBI.rev.compl                                  ------------------------------------------------------------ 46364
human.FAM230A.ncRNA.18487127.18500594-GGTLC3.18516335.18518165.rev.compl.REF       ------------------------------------------------------------ 31039

chimp.LOC112206744-LOC107973052-GGT2.revcompl                                      ------------------------------------------------------------ 52116
chimp54546-135457.revcompl.Pan.troglodytes.clone.rp43-41g5.GenBank:AC099533.36     CAGTGCTGACCATTTCACCCACTTGGCCTCCTTGAGGAATAGGGATGGGGAGCCCTCTGG 58484
human.FAM30B-LOC105372935-GGT2.NCBI.                                               ------------------------------------------------------------ 51109
human.FAM230E-LOC105377182-GGT3P.NCBI.                                             ------------------------------------------------------------ 49762
human.FAM230J-LOC105372942-GGTLC5P.NCBI.rev.compl                                  ------------------------------------------------------------ 46364
human.FAM230A.ncRNA.18487127.18500594-GGTLC3.18516335.18518165.rev.compl.REF       ------------------------------------------------------------ 31039

chimp.LOC112206744-LOC107973052-GGT2.revcompl                                      --------------------------------------------------ggttttgcct 52126
chimp54546-135457.revcompl.Pan.troglodytes.clone.rp43-41g5.GenBank:AC099533.36     GGATGGGCAAGGCCTTCCAGGACAGGCTCGGTTCTGGTCCCCTGCTTTTTGAGGTTGGGT 58544
human.FAM30B-LOC105372935-GGT2.NCBI.                                               --------------------------------------------------ggttttgcct 51119
human.FAM230E-LOC105377182-GGT3P.NCBI.                                             --------------------------------------------------ggttttgcct 49772
human.FAM230J-LOC105372942-GGTLC5P.NCBI.rev.compl                                  ------------------------------------------------------------ 46364
human.FAM230A.ncRNA.18487127.18500594-GGTLC3.18516335.18518165.rev.compl.REF       ------------------------------------------------------------ 31039

chimp.LOC112206744-LOC107973052-GGT2.revcompl                                      tccagttctgacttttgaggaggtcattggaaacagacccctgggcctgctt-------- 52178
chimp54546-135457.revcompl.Pan.troglodytes.clone.rp43-41g5.GenBank:AC099533.36     TAAAATGCCGACCATGGCAGAGGGGTTACAGCTCAGGTTCCCACACCTCGCTTTTCACAG 58604
human.FAM30B-LOC105372935-GGT2.NCBI.                                               tccagttctgacttttgaggaggtcattggaaacagacccctgggcctgctt-------- 51171
human.FAM230E-LOC105377182-GGT3P.NCBI.                                             tccagttctgacttttgaggaggtcattggaaacagacccctgggcctgctt-------- 49824
human.FAM230J-LOC105372942-GGTLC5P.NCBI.rev.compl                                  ------------------------------------------------------------ 46364
human.FAM230A.ncRNA.18487127.18500594-GGTLC3.18516335.18518165.rev.compl.REF       ------------------------------------------------------------ 31039

chimp.LOC112206744-LOC107973052-GGT2.revcompl                                      tctccctgagcctcactgcccatatggacactacagacactgaccctttgcccagaaagg 52238
chimp54546-135457.revcompl.Pan.troglodytes.clone.rp43-41g5.GenBank:AC099533.36     CCTCCGAGGGCAGCAGTGCACATGGAGGAGACGTCTCCCACGAGGCCAAGGCCTCCAGTG 58664
human.FAM30B-LOC105372935-GGT2.NCBI.                                               cccccctgagccccactgcccatatggacactacagacactgaccctttgcccagaaagg 51231
human.FAM230E-LOC105377182-GGT3P.NCBI.                                             cccccc-gagccccactgcccatatggacactacagacactgaccctttgcccagaaagg 49883
human.FAM230J-LOC105372942-GGTLC5P.NCBI.rev.compl                                  ------------------------------------------------------------ 46364
human.FAM230A.ncRNA.18487127.18500594-GGTLC3.18516335.18518165.rev.compl.REF       ------------------------------------------------------------ 31039

chimp.LOC112206744-LOC107973052-GGT2.revcompl                                      tacaactatggcctctgcc----------------------------------------- 52257
chimp54546-135457.revcompl.Pan.troglodytes.clone.rp43-41g5.GenBank:AC099533.36     CTCACCGATGCCCTCTGCGAAGTTGGGGTAGAACTCGTCAGTGAAGAGGGGCTCGGGCAG 58724
human.FAM30B-LOC105372935-GGT2.NCBI.                                               tacaactatggcctctgcc----------------------------------------- 51250
human.FAM230E-LOC105377182-GGT3P.NCBI.                                             tacaactatggcctctgcc----------------------------------------- 49902
human.FAM230J-LOC105372942-GGTLC5P.NCBI.rev.compl                                  ------------------------------------------------------------ 46364
human.FAM230A.ncRNA.18487127.18500594-GGTLC3.18516335.18518165.rev.compl.REF       ------------------------------------------------------------ 31039

chimp.LOC112206744-LOC107973052-GGT2.revcompl                                      ------------------------------------------------------------ 52257
chimp54546-135457.revcompl.Pan.troglodytes.clone.rp43-41g5.GenBank:AC099533.36     CTCACGGAAGTACAGCTTCAGCGTGCCTGCGATGGCGTTCACGTCCATCTCACTCATCAT 58784
human.FAM30B-LOC105372935-GGT2.NCBI.                                               ------------------------------------------------------------ 51250
human.FAM230E-LOC105377182-GGT3P.NCBI.                                             ------------------------------------------------------------ 49902
human.FAM230J-LOC105372942-GGTLC5P.NCBI.rev.compl                                  ------------------------------------------------------------ 46364
human.FAM230A.ncRNA.18487127.18500594-GGTLC3.18516335.18518165.rev.compl.REF       ------------------------------------------------------------ 31039

chimp.LOC112206744-LOC107973052-GGT2.revcompl                                      ------------------------------------------------------------ 52257
chimp54546-135457.revcompl.Pan.troglodytes.clone.rp43-41g5.GenBank:AC099533.36     CACCGACATGTCCTTGTTATCTGGAAAGAGCATGGAAATGCAGCGGCCTCCTTGAAGATC 58844
human.FAM30B-LOC105372935-GGT2.NCBI.                                               ------------------------------------------------------------ 51250
human.FAM230E-LOC105377182-GGT3P.NCBI.                                             ------------------------------------------------------------ 49902
human.FAM230J-LOC105372942-GGTLC5P.NCBI.rev.compl                                  ------------------------------------------------------------ 46364
human.FAM230A.ncRNA.18487127.18500594-GGTLC3.18516335.18518165.rev.compl.REF       ------------------------------------------------------------ 31039

chimp.LOC112206744-LOC107973052-GGT2.revcompl                                      ------------------------------------------------------------ 52257
chimp54546-135457.revcompl.Pan.troglodytes.clone.rp43-41g5.GenBank:AC099533.36     CTGAGTGAGTCACCCGCCACCCCTGCCTTGGCTAAAGCACCGTCCCTGCCATGCTGACCA 58904
human.FAM30B-LOC105372935-GGT2.NCBI.                                               ------------------------------------------------------------ 51250
human.FAM230E-LOC105377182-GGT3P.NCBI.                                             ------------------------------------------------------------ 49902
human.FAM230J-LOC105372942-GGTLC5P.NCBI.rev.compl                                  ------------------------------------------------------------ 46364
human.FAM230A.ncRNA.18487127.18500594-GGTLC3.18516335.18518165.rev.compl.REF       ------------------------------------------------------------ 31039

chimp.LOC112206744-LOC107973052-GGT2.revcompl                                      ------------------------------------------------------------ 52257
chimp54546-135457.revcompl.Pan.troglodytes.clone.rp43-41g5.GenBank:AC099533.36     CTGTATGGGTCCCTCCTGGGCTTTGAGCAGCTCATCTGACTCCTCCCAAGAGCTGTGCAT 58964
human.FAM30B-LOC105372935-GGT2.NCBI.                                               ------------------------------------------------------------ 51250
human.FAM230E-LOC105377182-GGT3P.NCBI.                                             ------------------------------------------------------------ 49902
human.FAM230J-LOC105372942-GGTLC5P.NCBI.rev.compl                                  ------------------------------------------------------------ 46364
human.FAM230A.ncRNA.18487127.18500594-GGTLC3.18516335.18518165.rev.compl.REF       ------------------------------------------------------------ 31039

chimp.LOC112206744-LOC107973052-GGT2.revcompl                                      ------------------------------------------------------------ 52257
chimp54546-135457.revcompl.Pan.troglodytes.clone.rp43-41g5.GenBank:AC099533.36     GGTTCTGTGTCTGCAGAGTTGATAGGGGTGCGTGGGCATTCCCATTCCTCTCCCCTGCTT 59024
human.FAM30B-LOC105372935-GGT2.NCBI.                                               ------------------------------------------------------------ 51250
human.FAM230E-LOC105377182-GGT3P.NCBI.                                             ------------------------------------------------------------ 49902
human.FAM230J-LOC105372942-GGTLC5P.NCBI.rev.compl                                  ------------------------------------------------------------ 46364
human.FAM230A.ncRNA.18487127.18500594-GGTLC3.18516335.18518165.rev.compl.REF       ------------------------------------------------------------ 31039

chimp.LOC112206744-LOC107973052-GGT2.revcompl                                      ------------------------------------------------------------ 52257
chimp54546-135457.revcompl.Pan.troglodytes.clone.rp43-41g5.GenBank:AC099533.36     GGCCTGATGTGATGGCCAGGAGGAGGCCAGCATGGCAGGACACAGCACCTGCGTGGGGAT 59084
human.FAM30B-LOC105372935-GGT2.NCBI.                                               ------------------------------------------------------------ 51250
human.FAM230E-LOC105377182-GGT3P.NCBI.                                             ------------------------------------------------------------ 49902
human.FAM230J-LOC105372942-GGTLC5P.NCBI.rev.compl                                  ------------------------------------------------------------ 46364
human.FAM230A.ncRNA.18487127.18500594-GGTLC3.18516335.18518165.rev.compl.REF       ------------------------------------------------------------ 31039

chimp.LOC112206744-LOC107973052-GGT2.revcompl                                      ------------------------------------------------------------ 52257
chimp54546-135457.revcompl.Pan.troglodytes.clone.rp43-41g5.GenBank:AC099533.36     TGGGTGGCTCTGCCCTGTACATAGCAACCACCCCTGCACCAGTGTCGTCTGATAGCAGGA 59144
human.FAM30B-LOC105372935-GGT2.NCBI.                                               ------------------------------------------------------------ 51250
human.FAM230E-LOC105377182-GGT3P.NCBI.                                             ------------------------------------------------------------ 49902
human.FAM230J-LOC105372942-GGTLC5P.NCBI.rev.compl                                  ------------------------------------------------------------ 46364
human.FAM230A.ncRNA.18487127.18500594-GGTLC3.18516335.18518165.rev.compl.REF       ------------------------------------------------------------ 31039

chimp.LOC112206744-LOC107973052-GGT2.revcompl                                      ------------------------------------------------------------ 52257
chimp54546-135457.revcompl.Pan.troglodytes.clone.rp43-41g5.GenBank:AC099533.36     AGGCCGTGGGAGAATCTGATTGGTTTCAGTGTTTGAACTGGTGTCTTCATTTGGACCCAA 59204
human.FAM30B-LOC105372935-GGT2.NCBI.                                               ------------------------------------------------------------ 51250
human.FAM230E-LOC105377182-GGT3P.NCBI.                                             ------------------------------------------------------------ 49902
human.FAM230J-LOC105372942-GGTLC5P.NCBI.rev.compl                                  ------------------------------------------------------------ 46364
human.FAM230A.ncRNA.18487127.18500594-GGTLC3.18516335.18518165.rev.compl.REF       ------------------------------------------------------------ 31039

chimp.LOC112206744-LOC107973052-GGT2.revcompl                                      ------------------------------------------------------------ 52257
chimp54546-135457.revcompl.Pan.troglodytes.clone.rp43-41g5.GenBank:AC099533.36     TTGGCCACTGGTGCTTACATCCTCACCACAGGCCAGGTTCATTCTGGGCCCCCAGAGGGA 59264
human.FAM30B-LOC105372935-GGT2.NCBI.                                               ------------------------------------------------------------ 51250
human.FAM230E-LOC105377182-GGT3P.NCBI.                                             ------------------------------------------------------------ 49902
human.FAM230J-LOC105372942-GGTLC5P.NCBI.rev.compl                                  ------------------------------------------------------------ 46364
human.FAM230A.ncRNA.18487127.18500594-GGTLC3.18516335.18518165.rev.compl.REF       ------------------------------------------------------------ 31039
```

```
chimp.LOC112206744-LOC107973052-GGT2.revcompl                                 ----------cccagggactctcctgctcttgcgagagatgatggggccatttgccttgg 52307
chimp54546-135457.revcompl.Pan.troglodytes.clone.rp43-41g5.GenBank:AC099533.36 GCTGAAACTACCACAGGGCCCTCCCAGGATGCTGGGCATTCTAGGGGTCCTGGTCAGGG 59324
human.FAM30B-LOC105372935-GGT2.NCBI.                                          ----------cccagggactctcctgctcttgcgagagatgatggggccatttgccttgg 51300
human.FAM230E-LOC105377182-GGT3P.NCBI.                                        ----------cccagggactctcctgctcttgcgagagatgatggggccatttgccttgg 49952
human.FAM230J-LOC105372942-GGTLC5P.NCBI.rev.compl                             ------------------------------------------------------------ 46364
human.FAM230A.ncRNA.18487127.18500594-GGTLC3.18516335.18518165.rev.compl.REF  ------------------------------------------------------------ 31039

chimp.LOC112206744-LOC107973052-GGT2.revcompl                                 cttggcggctgtggctctagaactgcctctcccaccctgaagcctggcac---------- 52357
chimp54546-135457.revcompl.Pan.troglodytes.clone.rp43-41g5.GenBank:AC099533.36 TGGGTGGTGTGTGCTGCAAAGAAGGGTCTGCAGGCACAAAATCCTGTTGCTTTGAAGATG 59384
human.FAM30B-LOC105372935-GGT2.NCBI.                                          cttggcggctgtggctctagaactgcctctcccaccctgaagcctggcac---------- 51350
human.FAM230E-LOC105377182-GGT3P.NCBI.                                        cttggcggctgtggctctagaactgcctctcccaccctgaagcctggcac---------- 50002
human.FAM230J-LOC105372942-GGTLC5P.NCBI.rev.compl                             ------------------------------------------------------------ 46364
human.FAM230A.ncRNA.18487127.18500594-GGTLC3.18516335.18518165.rev.compl.REF  ------------------------------------------------------------ 31039

chimp.LOC112206744-LOC107973052-GGT2.revcompl                                 ------------------------------------------------------------ 52357
chimp54546-135457.revcompl.Pan.troglodytes.clone.rp43-41g5.GenBank:AC099533.36 CTGGGAAGGACCCTCTGGGGTCTCAGTGCCCTCCCCTGGCATTTGAGGCAGGTCTGGGTC 59444
human.FAM30B-LOC105372935-GGT2.NCBI.                                          ------------------------------------------------------------ 51350
human.FAM230E-LOC105377182-GGT3P.NCBI.                                        ------------------------------------------------------------ 50002
human.FAM230J-LOC105372942-GGTLC5P.NCBI.rev.compl                             ------------------------------------------------------------ 46364
human.FAM230A.ncRNA.18487127.18500594-GGTLC3.18516335.18518165.rev.compl.REF  ------------------------------------------------------------ 31039

chimp.LOC112206744-LOC107973052-GGT2.revcompl                                 -----aagtttccaagagctggtggtttcaattcctagaagctgcacgtacatcccggaa 52412
chimp54546-135457.revcompl.Pan.troglodytes.clone.rp43-41g5.GenBank:AC099533.36 CTTCAAAGCCTGTGAGGGTTGGTGAGATGGAGGCGGAGATGCTGCAG----CCCCGGCCT 59500
human.FAM30B-LOC105372935-GGT2.NCBI.                                          -----aagtttccaagagctggtggtttcaattcctagaagctgcacatatatcccggaa 51405
human.FAM230E-LOC105377182-GGT3P.NCBI.                                        -----aagtttccaagagctggtggtttcaattcctagaagctgcacatatatcccggaa 50057
human.FAM230J-LOC105372942-GGTLC5P.NCBI.rev.compl                             ------------------------------------------------------------ 46364
human.FAM230A.ncRNA.18487127.18500594-GGTLC3.18516335.18518165.rev.compl.REF  ------------------------------------------------------------ 31039

chimp.LOC112206744-LOC107973052-GGT2.revcompl                                 ggtctgacacccagcacatgattccttccaccttgtagttagacag-------------- 52458
chimp54546-135457.revcompl.Pan.troglodytes.clone.rp43-41g5.GenBank:AC099533.36 GCGCTGAATTTCATCAGTGCCCTCTGCCCACCACATCCTCATACAGGGCAGTGGACAGAC 59560
human.FAM30B-LOC105372935-GGT2.NCBI.                                          ggtctgacacccagcatatgattccttccaccttgtagttagacag-------------- 51451
human.FAM230E-LOC105377182-GGT3P.NCBI.                                        ggtctgacacccagcatatgattccttccaccttgtagttagacag-------------- 50103
human.FAM230J-LOC105372942-GGTLC5P.NCBI.rev.compl                             ------------------------------------------------------------ 46364
human.FAM230A.ncRNA.18487127.18500594-GGTLC3.18516335.18518165.rev.compl.REF  ------------------------------------------------------------ 31039

chimp.LOC112206744-LOC107973052-GGT2.revcompl                                 ------------------------------------------------------------ 52458
chimp54546-135457.revcompl.Pan.troglodytes.clone.rp43-41g5.GenBank:AC099533.36 CGCACTGAGTCCTGGGCTTCCACCTCCTGTCCACCCCCAAGGCAGGAAGGTCAAGTCCCC 59620
human.FAM30B-LOC105372935-GGT2.NCBI.                                          ------------------------------------------------------------ 51451
human.FAM230E-LOC105377182-GGT3P.NCBI.                                        ------------------------------------------------------------ 50103
human.FAM230J-LOC105372942-GGTLC5P.NCBI.rev.compl                             ------------------------------------------------------------ 46364
human.FAM230A.ncRNA.18487127.18500594-GGTLC3.18516335.18518165.rev.compl.REF  ------------------------------------------------------------ 31039

chimp.LOC112206744-LOC107973052-GGT2.revcompl                                 ------------------------------------------------------------ 52458
chimp54546-135457.revcompl.Pan.troglodytes.clone.rp43-41g5.GenBank:AC099533.36 ACAGAAGCCCCTGGTCCACTCCACCAAGTGGCATGAGTGGGTACGATGGTGTAAAAACTG 59680
human.FAM30B-LOC105372935-GGT2.NCBI.                                          ------------------------------------------------------------ 51451
human.FAM230E-LOC105377182-GGT3P.NCBI.                                        ------------------------------------------------------------ 50103
human.FAM230J-LOC105372942-GGTLC5P.NCBI.rev.compl                             ------------------------------------------------------------ 46364
human.FAM230A.ncRNA.18487127.18500594-GGTLC3.18516335.18518165.rev.compl.REF  ------------------------------------------------------------ 31039

chimp.LOC112206744-LOC107973052-GGT2.revcompl                                 ------------------------------------------------------------ 52458
chimp54546-135457.revcompl.Pan.troglodytes.clone.rp43-41g5.GenBank:AC099533.36 GCTTCTATAGAAGCTGTTTGTACAACTCTTGTTTTCTCTTTTTTAAAAATAATAAAACAG 59740
human.FAM30B-LOC105372935-GGT2.NCBI.                                          ------------------------------------------------------------ 51451
human.FAM230E-LOC105377182-GGT3P.NCBI.                                        ------------------------------------------------------------ 50103
human.FAM230J-LOC105372942-GGTLC5P.NCBI.rev.compl                             ------------------------------------------------------------ 46364
human.FAM230A.ncRNA.18487127.18500594-GGTLC3.18516335.18518165.rev.compl.REF  ------------------------------------------------------------ 31039

chimp.LOC112206744-LOC107973052-GGT2.revcompl                                 ------------------------------------------------------------ 52458
chimp54546-135457.revcompl.Pan.troglodytes.clone.rp43-41g5.GenBank:AC099533.36 TAAATGAAGAAAAGACACAGAGAAGGATGTGACATGCCTGGGCCATGGAGCACTCTGAGA 59800
human.FAM30B-LOC105372935-GGT2.NCBI.                                          ------------------------------------------------------------ 51451
human.FAM230E-LOC105377182-GGT3P.NCBI.                                        ------------------------------------------------------------ 50103
human.FAM230J-LOC105372942-GGTLC5P.NCBI.rev.compl                             ------------------------------------------------------------ 46364
human.FAM230A.ncRNA.18487127.18500594-GGTLC3.18516335.18518165.rev.compl.REF  ------------------------------------------------------------ 31039

chimp.LOC112206744-LOC107973052-GGT2.revcompl                                 ------------------------------------------------------------ 52458
chimp54546-135457.revcompl.Pan.troglodytes.clone.rp43-41g5.GenBank:AC099533.36 TCTCATCGCGGACACCACTGCCCACACCTCCATCCCGTCCTGCGCAGGCCGACACTCACT 59860
human.FAM30B-LOC105372935-GGT2.NCBI.                                          ------------------------------------------------------------ 51451
human.FAM230E-LOC105377182-GGT3P.NCBI.                                        ------------------------------------------------------------ 50103
human.FAM230J-LOC105372942-GGTLC5P.NCBI.rev.compl                             ------------------------------------------------------------ 46364
human.FAM230A.ncRNA.18487127.18500594-GGTLC3.18516335.18518165.rev.compl.REF  ------------------------------------------------------------ 31039

chimp.LOC112206744-LOC107973052-GGT2.revcompl                                 ------------------------------------------------------------ 52458
chimp54546-135457.revcompl.Pan.troglodytes.clone.rp43-41g5.GenBank:AC099533.36 GACGTGGAAGCCTGCCTTCAGTGCCTGGATGTCTGCAGCCACACCGGACACGCGGTAGAT 59920
human.FAM30B-LOC105372935-GGT2.NCBI.                                          ------------------------------------------------------------ 51451
human.FAM230E-LOC105377182-GGT3P.NCBI.                                        ------------------------------------------------------------ 50103
human.FAM230J-LOC105372942-GGTLC5P.NCBI.rev.compl                             ------------------------------------------------------------ 46364
human.FAM230A.ncRNA.18487127.18500594-GGTLC3.18516335.18518165.rev.compl.REF  ------------------------------------------------------------ 31039

chimp.LOC112206744-LOC107973052-GGT2.revcompl                                 ------------------------------------------------------------ 52458
chimp54546-135457.revcompl.Pan.troglodytes.clone.rp43-41g5.GenBank:AC099533.36 GCCCACCTCCTCCATGCCTCGGTGCTCGATCTCCTCCACGCACTGGCGCACGATGTAGGG 59980
human.FAM30B-LOC105372935-GGT2.NCBI.                                          ------------------------------------------------------------ 51451
human.FAM230E-LOC105377182-GGT3P.NCBI.                                        ------------------------------------------------------------ 50103
human.FAM230J-LOC105372942-GGTLC5P.NCBI.rev.compl                             ------------------------------------------------------------ 46364
human.FAM230A.ncRNA.18487127.18500594-GGTLC3.18516335.18518165.rev.compl.REF  ------------------------------------------------------------ 31039

chimp.LOC112206744-LOC107973052-GGT2.revcompl                                 ------------------------------------------------------------ 52458
chimp54546-135457.revcompl.Pan.troglodytes.clone.rp43-41g5.GenBank:AC099533.36 CACCTTGGACCTCTCTCTCCTGCGGGAGGAGGGAATGCTCTCAGTGTCCTAAGAGCCCTG 60040
human.FAM30B-LOC105372935-GGT2.NCBI.                                          ------------------------------------------------------------ 51451
human.FAM230E-LOC105377182-GGT3P.NCBI.                                        ------------------------------------------------------------ 50103
human.FAM230J-LOC105372942-GGTLC5P.NCBI.rev.compl                             ------------------------------------------------------------ 46364
human.FAM230A.ncRNA.18487127.18500594-GGTLC3.18516335.18518165.rev.compl.REF  ------------------------------------------------------------ 31039

chimp.LOC112206744-LOC107973052-GGT2.revcompl                                 ------------------------------------------------------------ 52458
chimp54546-135457.revcompl.Pan.troglodytes.clone.rp43-41g5.GenBank:AC099533.36 CTTGGGCCATAACACAGGAGACCTGCTCCCTATCTGCACACCCGGAGGTGGGGTGAGGAC 60100
human.FAM30B-LOC105372935-GGT2.NCBI.                                          ------------------------------------------------------------ 51451
human.FAM230E-LOC105377182-GGT3P.NCBI.                                        ------------------------------------------------------------ 50103
human.FAM230J-LOC105372942-GGTLC5P.NCBI.rev.compl                             ------------------------------------------------------------ 46364
human.FAM230A.ncRNA.18487127.18500594-GGTLC3.18516335.18518165.rev.compl.REF  ------------------------------------------------------------ 31039

chimp.LOC112206744-LOC107973052-GGT2.revcompl                                 ------------------------------------------------------------ 52458
chimp54546-135457.revcompl.Pan.troglodytes.clone.rp43-41g5.GenBank:AC099533.36 GGTGACGAAGGTACCCAGGTCTGGGGCTGCACACAGAGCCTGCTGCATGCCTGTCCTCCC 60160
human.FAM30B-LOC105372935-GGT2.NCBI.                                          ------------------------------------------------------------ 51451
human.FAM230E-LOC105377182-GGT3P.NCBI.                                        ------------------------------------------------------------ 50103
human.FAM230J-LOC105372942-GGTLC5P.NCBI.rev.compl                             ------------------------------------------------------------ 46364
human.FAM230A.ncRNA.18487127.18500594-GGTLC3.18516335.18518165.rev.compl.REF  ------------------------------------------------------------ 31039

chimp.LOC112206744-LOC107973052-GGT2.revcompl                                 ------------------------------------------------------------ 52458
chimp54546-135457.revcompl.Pan.troglodytes.clone.rp43-41g5.GenBank:AC099533.36 TCTGCAAGCTCTGTCCTCATTGCATGTACTTTCTCAGGAACCTTTCAAGCGGCCAGAACC 60220
human.FAM30B-LOC105372935-GGT2.NCBI.                                          ------------------------------------------------------------ 51451
human.FAM230E-LOC105377182-GGT3P.NCBI.                                        ------------------------------------------------------------ 50103
human.FAM230J-LOC105372942-GGTLC5P.NCBI.rev.compl                             ------------------------------------------------------------ 46364
human.FAM230A.ncRNA.18487127.18500594-GGTLC3.18516335.18518165.rev.compl.REF  ------------------------------------------------------------ 31039

chimp.LOC112206744-LOC107973052-GGT2.revcompl                                 ------------------------------------------------------------ 52458
chimp54546-135457.revcompl.Pan.troglodytes.clone.rp43-41g5.GenBank:AC099533.36 CCTGCGAATCACACATGATCTTTGCGGGAAGGTCAGGAGGCCTTTCTAAGTCAAGTCAGC 60280
human.FAM30B-LOC105372935-GGT2.NCBI.                                          ------------------------------------------------------------ 51451
human.FAM230E-LOC105377182-GGT3P.NCBI.                                        ------------------------------------------------------------ 50103
human.FAM230J-LOC105372942-GGTLC5P.NCBI.rev.compl                             ------------------------------------------------------------ 46364
human.FAM230A.ncRNA.18487127.18500594-GGTLC3.18516335.18518165.rev.compl.REF  ------------------------------------------------------------ 31039

chimp.LOC112206744-LOC107973052-GGT2.revcompl                                 ------------------------------------------------------------ 52458
chimp54546-135457.revcompl.Pan.troglodytes.clone.rp43-41g5.GenBank:AC099533.36 ACAGGAAGGGCATCTCACAGATTCCAGGCCTGGGGTTAGCAGCCTGTGCCCCCGGCTGGG 60340
human.FAM30B-LOC105372935-GGT2.NCBI.                                          ------------------------------------------------------------ 51451
human.FAM230E-LOC105377182-GGT3P.NCBI.                                        ------------------------------------------------------------ 50103
human.FAM230J-LOC105372942-GGTLC5P.NCBI.rev.compl                             ------------------------------------------------------------ 46364
human.FAM230A.ncRNA.18487127.18500594-GGTLC3.18516335.18518165.rev.compl.REF  ------------------------------------------------------------ 31039

chimp.LOC112206744-LOC107973052-GGT2.revcompl                                 ------------------------------------------------------------ 52458
```

```
chimp54546-135457.revcompl.Pan.troglodytes.clone.rp43-41g5.GenBank:AC099533.36      AGATCAGACCCAGTGTTGGTCCTGCCACCCACGTGCTGTGTGAGAGGAGAATCCCTGACC 60400
human.FAM30B-LOC105372935-GGT2.NCBI.                                                 ------------------------------------------------------------ 51451
human.FAM230E-LOC105377182-GGT3P.NCBI.                                               ------------------------------------------------------------ 50103
human.FAM230J-LOC105372942-GGTLC5P.NCBI.rev.compl                                    ------------------------------------------------------------ 46364
human.FAM230A.ncRNA.18487127.18500594-GGTLC3.18516335.18518165.rev.compl.REF         ------------------------------------------------------------ 31039

chimp.LOC112206744-LOC107973052-GGT2.revcompl                                        ------------------------------------------------------------ 52458
chimp54546-135457.revcompl.Pan.troglodytes.clone.rp43-41g5.GenBank:AC099533.36      CCTGCGCTGGGCCTTAACACACATCCGACGAATGAATGAAGGGTTGCCTCAGCACCGGTG 60460
human.FAM30B-LOC105372935-GGT2.NCBI.                                                 ------------------------------------------------------------ 51451
human.FAM230E-LOC105377182-GGT3P.NCBI.                                               ------------------------------------------------------------ 50103
human.FAM230J-LOC105372942-GGTLC5P.NCBI.rev.compl                                    ------------------------------------------------------------ 46364
human.FAM230A.ncRNA.18487127.18500594-GGTLC3.18516335.18518165.rev.compl.REF         ------------------------------------------------------------ 31039

chimp.LOC112206744-LOC107973052-GGT2.revcompl                                        ----aagttcttttttgttttgttttgtt-ttctttgtttgtttttgagatggagtcttg 52513
chimp54546-135457.revcompl.Pan.troglodytes.clone.rp43-41g5.GenBank:AC099533.36      CTCCAAGTCCTGCGATGCTAAGTGCTTTTCTCCTCTG--AGTCTTAGCAATGGAAAATTC 60518
human.FAM30B-LOC105372935-GGT2.NCBI.                                                 ----aagttcttttttgttttgttttgtttttctttgtttgtttttgagatggagtcttg 51507
human.FAM230E-LOC105377182-GGT3P.NCBI.                                               ----aagttcttttttgttttgttttttttttctttgtttgtttttgagatggagtcttg 50159
human.FAM230J-LOC105372942-GGTLC5P.NCBI.rev.compl                                    ------------------------------------------------------------ 46364
human.FAM230A.ncRNA.18487127.18500594-GGTLC3.18516335.18518165.rev.compl.REF         ------------------------------------------------------------ 31039

chimp.LOC112206744-LOC107973052-GGT2.revcompl                                        ctctgtctcc--------------------------------------cagactgcagtg 52535
chimp54546-135457.revcompl.Pan.troglodytes.clone.rp43-41g5.GenBank:AC099533.36      CAATACCTCCACACAGGACACTAGAGTAAGAATCCTTCACAGTTAGAACGCAGTGCTGTG 60578
human.FAM30B-LOC105372935-GGT2.NCBI.                                                 ctctgtctcc--------------------------------------cagactgcagtg 51529
human.FAM230E-LOC105377182-GGT3P.NCBI.                                               ctctgtctcc--------------------------------------cagactgcagtg 50181
human.FAM230J-LOC105372942-GGTLC5P.NCBI.rev.compl                                    ------------------------------------------------------------ 46364
human.FAM230A.ncRNA.18487127.18500594-GGTLC3.18516335.18518165.rev.compl.REF         ------------------------------------------------------------ 31039

chimp.LOC112206744-LOC107973052-GGT2.revcompl                                        cagtggcatgatc-tcagctcactataacctccgcctcccaggttcaagcgattctcctg 52594
chimp54546-135457.revcompl.Pan.troglodytes.clone.rp43-41g5.GenBank:AC099533.36      CGGAGGCCTTAACTTGAGTTCTGTTTTGCACCTGGATTTACCAGCACATCAAAGCTGCTT 60638
human.FAM30B-LOC105372935-GGT2.NCBI.                                                 cagtggcatgatc-tcagctcactataacctccgcctcccaggttcaagcgattctcctg 51588
human.FAM230E-LOC105377182-GGT3P.NCBI.                                               cagtggcatgatc-tcagctcactataacctccgcctcccaggttcaagcgattctcctg 50240
human.FAM230J-LOC105372942-GGTLC5P.NCBI.rev.compl                                    ------------------------------------------------------------ 46364
human.FAM230A.ncRNA.18487127.18500594-GGTLC3.18516335.18518165.rev.compl.REF         ------------------------------------------------------------ 31039

chimp.LOC112206744-LOC107973052-GGT2.revcompl                                        cctcagcctcccgagtagctgggatta--------------------------------- 52621
chimp54546-135457.revcompl.Pan.troglodytes.clone.rp43-41g5.GenBank:AC099533.36      CGCAAGCCCCCTCATCAGCAGGGCTTATCTGGGGGAGCTGCTGATGGAGTCCTCGCTGCT 60698
human.FAM30B-LOC105372935-GGT2.NCBI.                                                 cctcagcctcccgagtagctgggatta--------------------------------- 51615
human.FAM230E-LOC105377182-GGT3P.NCBI.                                               cctcagcctcccgagtagctgggatta--------------------------------- 50267
human.FAM230J-LOC105372942-GGTLC5P.NCBI.rev.compl                                    ------------------------------------------------------------ 46364
human.FAM230A.ncRNA.18487127.18500594-GGTLC3.18516335.18518165.rev.compl.REF         ------------------------------------------------------------ 31039

chimp.LOC112206744-LOC107973052-GGT2.revcompl                                        ------------------------------------------------------------ 52621
chimp54546-135457.revcompl.Pan.troglodytes.clone.rp43-41g5.GenBank:AC099533.36      CATGCCCACAGCCCTCCCAGAGTGCTATGCGAGTGGCTGCCATGCAGTTGGGGGTGGGGT 60758
human.FAM30B-LOC105372935-GGT2.NCBI.                                                 ------------------------------------------------------------ 51615
human.FAM230E-LOC105377182-GGT3P.NCBI.                                               ------------------------------------------------------------ 50267
human.FAM230J-LOC105372942-GGTLC5P.NCBI.rev.compl                                    ------------------------------------------------------------ 46364
human.FAM230A.ncRNA.18487127.18500594-GGTLC3.18516335.18518165.rev.compl.REF         ------------------------------------------------------------ 31039

chimp.LOC112206744-LOC107973052-GGT2.revcompl                                        ------------------------------------------caggcacaggccagcacg 52639
chimp54546-135457.revcompl.Pan.troglodytes.clone.rp43-41g5.GenBank:AC099533.36      GTGGTGTTTAGACACAGATGGGAGTCCAGGGTATGACTGATGGAGGCCCCGGCCCACGTG 60818
human.FAM30B-LOC105372935-GGT2.NCBI.                                                 ------------------------------------------caggcacaggccagcacg 51633
human.FAM230E-LOC105377182-GGT3P.NCBI.                                               ------------------------------------------caggcacaggccagcacg 50285
human.FAM230J-LOC105372942-GGTLC5P.NCBI.rev.compl                                    ------------------------------------------------------------ 46364
human.FAM230A.ncRNA.18487127.18500594-GGTLC3.18516335.18518165.rev.compl.REF         ------------------------------------------------------------ 31039

chimp.LOC112206744-LOC107973052-GGT2.revcompl                                        ccaggctaa--------------------------------------------------- 52648
chimp54546-135457.revcompl.Pan.troglodytes.clone.rp43-41g5.GenBank:AC099533.36      ACCAGCAAGGTCAGAGGCCCAGCCAGATTCCATCCTAGGGAAGCAAATGAATTCTCAGAG 60878
human.FAM30B-LOC105372935-GGT2.NCBI.                                                 ccaggctaa--------------------------------------------------- 51642
human.FAM230E-LOC105377182-GGT3P.NCBI.                                               ccaggctaa--------------------------------------------------- 50294
human.FAM230J-LOC105372942-GGTLC5P.NCBI.rev.compl                                    ------------------------------------------------------------ 46364
human.FAM230A.ncRNA.18487127.18500594-GGTLC3.18516335.18518165.rev.compl.REF         ------------------------------------------------------------ 31039

chimp.LOC112206744-LOC107973052-GGT2.revcompl                                        ------------------------------------------------------------ 52648
chimp54546-135457.revcompl.Pan.troglodytes.clone.rp43-41g5.GenBank:AC099533.36      GAAGTGGTCTGTGTCTGCATGAACTGCTCTCAAACCAACAAACAGGCTTCTCTTGGCAAC 60938
human.FAM30B-LOC105372935-GGT2.NCBI.                                                 ------------------------------------------------------------ 51642
human.FAM230E-LOC105377182-GGT3P.NCBI.                                               ------------------------------------------------------------ 50294
human.FAM230J-LOC105372942-GGTLC5P.NCBI.rev.compl                                    ------------------------------------------------------------ 46364
human.FAM230A.ncRNA.18487127.18500594-GGTLC3.18516335.18518165.rev.compl.REF         ------------------------------------------------------------ 31039

chimp.LOC112206744-LOC107973052-GGT2.revcompl                                        ------------------------------------------------------------ 52648
chimp54546-135457.revcompl.Pan.troglodytes.clone.rp43-41g5.GenBank:AC099533.36      TGACTCATGACAAAGGGGCAGGGGTTACAAACATTCAACAGGAATCTGGGACCGTGACTA 60998
human.FAM30B-LOC105372935-GGT2.NCBI.                                                 ------------------------------------------------------------ 51642
human.FAM230E-LOC105377182-GGT3P.NCBI.                                               ------------------------------------------------------------ 50294
human.FAM230J-LOC105372942-GGTLC5P.NCBI.rev.compl                                    ------------------------------------------------------------ 46364
human.FAM230A.ncRNA.18487127.18500594-GGTLC3.18516335.18518165.rev.compl.REF         ------------------------------------------------------------ 31039

chimp.LOC112206744-LOC107973052-GGT2.revcompl                                        ------------------------------------------------------------ 52648
chimp54546-135457.revcompl.Pan.troglodytes.clone.rp43-41g5.GenBank:AC099533.36      ATTCCAAGCAAGCGCAGAAAACCCTGCACTTGGGCAGTTCCTGGAGCCAGCCTACTGCTT 61058
human.FAM30B-LOC105372935-GGT2.NCBI.                                                 ------------------------------------------------------------ 51642
human.FAM230E-LOC105377182-GGT3P.NCBI.                                               ------------------------------------------------------------ 50294
human.FAM230J-LOC105372942-GGTLC5P.NCBI.rev.compl                                    ------------------------------------------------------------ 46364
human.FAM230A.ncRNA.18487127.18500594-GGTLC3.18516335.18518165.rev.compl.REF         ------------------------------------------------------------ 31039

chimp.LOC112206744-LOC107973052-GGT2.revcompl                                        ------------------------------------------------------------ 52648
chimp54546-135457.revcompl.Pan.troglodytes.clone.rp43-41g5.GenBank:AC099533.36      CCAAGGAAGGTGGCAAGGACATTCCTGAGCTGGTTTGCAGGAAACACATGAGGTAGCTGA 61118
human.FAM30B-LOC105372935-GGT2.NCBI.                                                 ------------------------------------------------------------ 51642
human.FAM230E-LOC105377182-GGT3P.NCBI.                                               ------------------------------------------------------------ 50294
human.FAM230J-LOC105372942-GGTLC5P.NCBI.rev.compl                                    ------------------------------------------------------------ 46364
human.FAM230A.ncRNA.18487127.18500594-GGTLC3.18516335.18518165.rev.compl.REF         ------------------------------------------------------------ 31039

chimp.LOC112206744-LOC107973052-GGT2.revcompl                                        ------------------------------------------------------------ 52648
chimp54546-135457.revcompl.Pan.troglodytes.clone.rp43-41g5.GenBank:AC099533.36      ACACGCCGTGTTCCCTGGGGCCTGACCTCACCGTGCAGTGGCCACACACAGAGAGGAGAG 61178
human.FAM30B-LOC105372935-GGT2.NCBI.                                                 ------------------------------------------------------------ 51642
human.FAM230E-LOC105377182-GGT3P.NCBI.                                               ------------------------------------------------------------ 50294
human.FAM230J-LOC105372942-GGTLC5P.NCBI.rev.compl                                    ------------------------------------------------------------ 46364
human.FAM230A.ncRNA.18487127.18500594-GGTLC3.18516335.18518165.rev.compl.REF         ------------------------------------------------------------ 31039

chimp.LOC112206744-LOC107973052-GGT2.revcompl                                        ------------------------------------------------------------ 52648
chimp54546-135457.revcompl.Pan.troglodytes.clone.rp43-41g5.GenBank:AC099533.36      ATCCACACATGCTCTTTTCTGCAGAAACAGCTGAAGGGACAGGCACCCTCAGTGCGTGAG 61238
human.FAM30B-LOC105372935-GGT2.NCBI.                                                 ------------------------------------------------------------ 51642
human.FAM230E-LOC105377182-GGT3P.NCBI.                                               ------------------------------------------------------------ 50294
human.FAM230J-LOC105372942-GGTLC5P.NCBI.rev.compl                                    ------------------------------------------------------------ 46364
human.FAM230A.ncRNA.18487127.18500594-GGTLC3.18516335.18518165.rev.compl.REF         ------------------------------------------------------------ 31039

chimp.LOC112206744-LOC107973052-GGT2.revcompl                                        ------------------------------------------------------------ 52648
chimp54546-135457.revcompl.Pan.troglodytes.clone.rp43-41g5.GenBank:AC099533.36      CCCAAGCCCCTCCCCACTCACTTGGTGACCACAGCAATCTTGACTCTGAAGACCCCTGTC 61298
human.FAM30B-LOC105372935-GGT2.NCBI.                                                 ------------------------------------------------------------ 51642
human.FAM230E-LOC105377182-GGT3P.NCBI.                                               ------------------------------------------------------------ 50294
human.FAM230J-LOC105372942-GGTLC5P.NCBI.rev.compl                                    ------------------------------------------------------------ 46364
human.FAM230A.ncRNA.18487127.18500594-GGTLC3.18516335.18518165.rev.compl.REF         ------------------------------------------------------------ 31039

chimp.LOC112206744-LOC107973052-GGT2.revcompl                                        ------------------------------------------------------------ 52648
chimp54546-135457.revcompl.Pan.troglodytes.clone.rp43-41g5.GenBank:AC099533.36      TGTTTCTGGGATGGCATCCTCTTCAAGCTGAACTCCCTGCTGTTGAACTTGACCGAGAGC 61358
human.FAM30B-LOC105372935-GGT2.NCBI.                                                 ------------------------------------------------------------ 51642
human.FAM230E-LOC105377182-GGT3P.NCBI.                                               ------------------------------------------------------------ 50294
human.FAM230J-LOC105372942-GGTLC5P.NCBI.rev.compl                                    ------------------------------------------------------------ 46364
human.FAM230A.ncRNA.18487127.18500594-GGTLC3.18516335.18518165.rev.compl.REF         ------------------------------------------------------------ 31039

chimp.LOC112206744-LOC107973052-GGT2.revcompl                                        ------------------------------------------------------------ 52648
chimp54546-135457.revcompl.Pan.troglodytes.clone.rp43-41g5.GenBank:AC099533.36      TTTACTTCAATCTACAGTGGGGAGAAGGGGCACAGTCCTCTAATCCTTCTGGTCCTCATA 61418
human.FAM30B-LOC105372935-GGT2.NCBI.                                                 ------------------------------------------------------------ 51642
human.FAM230E-LOC105377182-GGT3P.NCBI.                                               ------------------------------------------------------------ 50294
human.FAM230J-LOC105372942-GGTLC5P.NCBI.rev.compl                                    ------------------------------------------------------------ 46364
human.FAM230A.ncRNA.18487127.18500594-GGTLC3.18516335.18518165.rev.compl.REF         ------------------------------------------------------------ 31039

chimp.LOC112206744-LOC107973052-GGT2.revcompl                                        ------------------------------------------------------------ 52648
chimp54546-135457.revcompl.Pan.troglodytes.clone.rp43-41g5.GenBank:AC099533.36      GCCCAAGGTCCCAAGGACCCCTGAGAGTCTCATCAAATATTTCCACCCCCAAGGCATGCA 61478
```

```
human.FAM30B-LOC105372935-GGT2.NCBI.                                    ------------------------------------------------------------ 51642
human.FAM230E-LOC105377182-GGT3P.NCBI.                                  ------------------------------------------------------------ 50294
human.FAM230J-LOC105372942-GGTLC5P.NCBI.rev.compl                       ------------------------------------------------------------ 46364
human.FAM230A.ncRNA.18487127.18500594-GGTLC3.18516335.18518165.rev.compl.REF ------------------------------------------------------------ 31039

chimp.LOC112206744-LOC107973052-GGT2.revcompl                           ------------------------------------------------------------ 52648
chimp54546-135457.revcompl.Pan.troglodytes.clone.rp43-41g5.GenBank:AC099533.36 GAGGTCTCAGCATGAGGTGGAGCCGCCCCTCGGGGAAGCCCCTAGTCTACTTCAGCAAGA 61538
human.FAM30B-LOC105372935-GGT2.NCBI.                                    ------------------------------------------------------------ 51642
human.FAM230E-LOC105377182-GGT3P.NCBI.                                  ------------------------------------------------------------ 50294
human.FAM230J-LOC105372942-GGTLC5P.NCBI.rev.compl                       ------------------------------------------------------------ 46364
human.FAM230A.ncRNA.18487127.18500594-GGTLC3.18516335.18518165.rev.compl.REF ------------------------------------------------------------ 31039

chimp.LOC112206744-LOC107973052-GGT2.revcompl                           ------------------------------------------------------------ 52648
chimp54546-135457.revcompl.Pan.troglodytes.clone.rp43-41g5.GenBank:AC099533.36 GCCAGGATTGTTGAAGATGATGTAAACGACTGGTTTCTGCAGAACCATCTCCTAAAATCC 61598
human.FAM30B-LOC105372935-GGT2.NCBI.                                    ------------------------------------------------------------ 51642
human.FAM230E-LOC105377182-GGT3P.NCBI.                                  ------------------------------------------------------------ 50294
human.FAM230J-LOC105372942-GGTLC5P.NCBI.rev.compl                       ------------------------------------------------------------ 46364
human.FAM230A.ncRNA.18487127.18500594-GGTLC3.18516335.18518165.rev.compl.REF ------------------------------------------------------------ 31039

chimp.LOC112206744-LOC107973052-GGT2.revcompl                           -----------------tttttgtatttttagtacagatggtgttttgccatgttggcca 52691
chimp54546-135457.revcompl.Pan.troglodytes.clone.rp43-41g5.GenBank:AC099533.36 CCGCTGCTTCTCTTGCTTGCTTACTTTTTTTAAGATATGGATCTTGCAATGTTGCCCA 61658
human.FAM30B-LOC105372935-GGT2.NCBI.                                    -----------------tttttgtatttttagtacagatggggttttgccatgttggcca 51685
human.FAM230E-LOC105377182-GGT3P.NCBI.                                  -----------------tttttgtatttttagtacagatggggttttgccatgttggcca 50337
human.FAM230J-LOC105372942-GGTLC5P.NCBI.rev.compl                       ------------------------------------------------------------ 46364
human.FAM230A.ncRNA.18487127.18500594-GGTLC3.18516335.18518165.rev.compl.REF ------------------------------------------------------------ 31039

chimp.LOC112206744-LOC107973052-GGT2.revcompl                           ggctgttttcaaactcttgacctcaggtcatccacccacctcagccttccaaggtgctgg 52751
chimp54546-135457.revcompl.Pan.troglodytes.clone.rp43-41g5.GenBank:AC099533.36 GGCTGTAATCAAACTCTTGGGCTCCAGTGATTCTCCTGGCTCAGCCTCCAGAGTAGCTGA 61718
human.FAM30B-LOC105372935-GGT2.NCBI.                                    ggctgttttcaaactcctgacctcaggtcatccacccacctcagccttccaaggtgctgg 51745
human.FAM230E-LOC105377182-GGT3P.NCBI.                                  ggctgttttcagactcctgacctcaggtcatccacccacctcagccttccaaggtgctgg 50397
human.FAM230J-LOC105372942-GGTLC5P.NCBI.rev.compl                       ------------------------------------------------------------ 46364
human.FAM230A.ncRNA.18487127.18500594-GGTLC3.18516335.18518165.rev.compl.REF ------------------------------------------------------------ 31039

chimp.LOC112206744-LOC107973052-GGT2.revcompl                           gatgacaggcgtgagccaccgtgcccagccaagacaggagaagttctaatctttgatagc 52811
chimp54546-135457.revcompl.Pan.troglodytes.clone.rp43-41g5.GenBank:AC099533.36 GATTATAGGTGT-----ACTGTGACCAGCTTTCTCTTTCTTAGTAACAATTAAAAAT--- 61770
human.FAM30B-LOC105372935-GGT2.NCBI.                                    gatgacaggcgtgagccaccgtgcccagccaagacaggagaagttctaatctttgatagc 51805
human.FAM230E-LOC105377182-GGT3P.NCBI.                                  gatgacaggcgtgagccaccgtgcccagccaagacaggagaagttctaatctttgatagc 50457
human.FAM230J-LOC105372942-GGTLC5P.NCBI.rev.compl                       ------------------------------------------------------------ 46364
human.FAM230A.ncRNA.18487127.18500594-GGTLC3.18516335.18518165.rev.compl.REF ------------------------------------------------------------ 31039

chimp.LOC112206744-LOC107973052-GGT2.revcompl                           agaccagggtgacgatgcttagcaacagtattttgtatatttcaaagtaacgaagagagg 52871
chimp54546-135457.revcompl.Pan.troglodytes.clone.rp43-41g5.GenBank:AC099533.36 -------------------------------------------TAAGTAGGAAAGACAGA 61787
human.FAM30B-LOC105372935-GGT2.NCBI.                                    agaccagggtgacgatgcttagcaacagtattttgtatatttcaaagtaacgaagagagg 51865
human.FAM230E-LOC105377182-GGT3P.NCBI.                                  agaccagggtgacgatgcttagcaacagtattttgtatatttcaaagtaacgaagagagc 50517
human.FAM230J-LOC105372942-GGTLC5P.NCBI.rev.compl                       ------------------------------------------------------------ 46364
human.FAM230A.ncRNA.18487127.18500594-GGTLC3.18516335.18518165.rev.compl.REF ------------------------------------------------------------ 31039

chimp.LOC112206744-LOC107973052-GGT2.revcompl                           actatggtgctaacacccagaaatgaaaaatattcaaggt-------------------- 52911
chimp54546-135457.revcompl.Pan.troglodytes.clone.rp43-41g5.GenBank:AC099533.36 ATAATATAAGAAAAAGCCAAGAATCTCAAATATTAACATTTTACCATTTATACCCGGGCC 61847
human.FAM30B-LOC105372935-GGT2.NCBI.                                    actatggtgctaacacccggaaatgaaaaatattcaaggt-------------------- 51905
human.FAM230E-LOC105377182-GGT3P.NCBI.                                  actatggtgctaacacccagaaatgaaaaatattcaaggt-------------------- 50557
human.FAM230J-LOC105372942-GGTLC5P.NCBI.rev.compl                       ------------------------------------------------------------ 46364
human.FAM230A.ncRNA.18487127.18500594-GGTLC3.18516335.18518165.rev.compl.REF ------------------------------------------------------------ 31039

chimp.LOC112206744-LOC107973052-GGT2.revcompl                           ------------------------------------------------------------ 52911
chimp54546-135457.revcompl.Pan.troglodytes.clone.rp43-41g5.GenBank:AC099533.36 TTTTTGGGTGTGTTATTGCTTTTAAGGTCTAAACCTTTCTGGGGTACCTAAAGACAAATC 61907
human.FAM30B-LOC105372935-GGT2.NCBI.                                    ------------------------------------------------------------ 51905
human.FAM230E-LOC105377182-GGT3P.NCBI.                                  ------------------------------------------------------------ 50557
human.FAM230J-LOC105372942-GGTLC5P.NCBI.rev.compl                       ------------------------------------------------------------ 46364
human.FAM230A.ncRNA.18487127.18500594-GGTLC3.18516335.18518165.rev.compl.REF ------------------------------------------------------------ 31039

chimp.LOC112206744-LOC107973052-GGT2.revcompl                           --------------------------------------gacggagactccaaataccctg 52933
chimp54546-135457.revcompl.Pan.troglodytes.clone.rp43-41g5.GenBank:AC099533.36 AGGGGTCTCCATTTCTATGGGCAGCCCCCAGGTCCCTGAGGGGAGGCCCCACCTGTCCTG 61967
human.FAM30B-LOC105372935-GGT2.NCBI.                                    --------------------------------------gacggagactccaaataccctg 51927
human.FAM230E-LOC105377182-GGT3P.NCBI.                                  --------------------------------------gacggagactccaaataccctg 50579
human.FAM230J-LOC105372942-GGTLC5P.NCBI.rev.compl                       ------------------------------------------------------------ 46364
human.FAM230A.ncRNA.18487127.18500594-GGTLC3.18516335.18518165.rev.compl.REF ------------------------------------------------------------ 31039

chimp.LOC112206744-LOC107973052-GGT2.revcompl                           ccttgatcattatacactatatgcatggaacaagcactcacatgtacccataaatataga 52993
chimp54546-135457.revcompl.Pan.troglodytes.clone.rp43-41g5.GenBank:AC099533.36 GTGGTGCCATGAGACCTTTGTTTGAGGAGGGATGCTCAGGGCCTGGCTGGCAACTGCATT 62027
human.FAM30B-LOC105372935-GGT2.NCBI.                                    ccttgatcattatacactctatgcatggaacaagcactcacatgtacccataaatataga 51987
human.FAM230E-LOC105377182-GGT3P.NCBI.                                  ccttgatcattatacactctatgcatggaacaagcactcacatgtacccataaatataga 50639
human.FAM230J-LOC105372942-GGTLC5P.NCBI.rev.compl                       ------------------------------------------------------------ 46364
human.FAM230A.ncRNA.18487127.18500594-GGTLC3.18516335.18518165.rev.compl.REF ------------------------------------------------------------ 31039

chimp.LOC112206744-LOC107973052-GGT2.revcompl                           aaatatcatgtatcaatatcagaaaaaaatctcctcctgacctcagcccaatcaggctct 53053
chimp54546-135457.revcompl.Pan.troglodytes.clone.rp43-41g5.GenBank:AC099533.36 CCAGCCAATGAATGACAATTCCGGTGCATGGTGATTCTGCTCTTGGGGGCCCTCCCTTCT 62087
human.FAM30B-LOC105372935-GGT2.NCBI.                                    aaatatcatgtatcaatatcagaaaaaaatctcctcctgacctcagcccaatcaggctct 52047
human.FAM230E-LOC105377182-GGT3P.NCBI.                                  aaatatcatgtatcaatatcagaaaaaaatctcctcctgacctcagcccaatcaggctct 50699
human.FAM230J-LOC105372942-GGTLC5P.NCBI.rev.compl                       ------------------------------------------------------------ 46364
human.FAM230A.ncRNA.18487127.18500594-GGTLC3.18516335.18518165.rev.compl.REF ------------------------------------------------------------ 31039

chimp.LOC112206744-LOC107973052-GGT2.revcompl                           catgccaccacacttgccaagttctctg-------------------------------- 53081
chimp54546-135457.revcompl.Pan.troglodytes.clone.rp43-41g5.GenBank:AC099533.36 CCAACCAGGCTCCCTCTCAGATGCTGCCTCAGGCTCAGCCACCTCTGACTGATTTGAAAT 62147
human.FAM30B-LOC105372935-GGT2.NCBI.                                    catgccaccacacttgccaagttctctg-------------------------------- 52075
human.FAM230E-LOC105377182-GGT3P.NCBI.                                  catgccaccacacttgccaagttctctg-------------------------------- 50727
human.FAM230J-LOC105372942-GGTLC5P.NCBI.rev.compl                       ------------------------------------------------------------ 46364
human.FAM230A.ncRNA.18487127.18500594-GGTLC3.18516335.18518165.rev.compl.REF ------------------------------------------------------------ 31039

chimp.LOC112206744-LOC107973052-GGT2.revcompl                           ------------------------------------------------------------ 53081
chimp54546-135457.revcompl.Pan.troglodytes.clone.rp43-41g5.GenBank:AC099533.36 GAACAGAGGCCACCCTATCCTTCTGGGGGAAGTGGGCACCAGGAGAGTCCCACGGACACG 62207
human.FAM30B-LOC105372935-GGT2.NCBI.                                    ------------------------------------------------------------ 52075
human.FAM230E-LOC105377182-GGT3P.NCBI.                                  ------------------------------------------------------------ 50727
human.FAM230J-LOC105372942-GGTLC5P.NCBI.rev.compl                       ------------------------------------------------------------ 46364
human.FAM230A.ncRNA.18487127.18500594-GGTLC3.18516335.18518165.rev.compl.REF ------------------------------------------------------------ 31039

chimp.LOC112206744-LOC107973052-GGT2.revcompl                           ------------------------------------------------------------ 53081
chimp54546-135457.revcompl.Pan.troglodytes.clone.rp43-41g5.GenBank:AC099533.36 TACCCCATTCATGGCGATGACGGTGCGCTGCCAGTCTCTGTCCTGCAGGGCCTGCGGGTC 62267
human.FAM30B-LOC105372935-GGT2.NCBI.                                    ------------------------------------------------------------ 52075
human.FAM230E-LOC105377182-GGT3P.NCBI.                                  ------------------------------------------------------------ 50727
human.FAM230J-LOC105372942-GGTLC5P.NCBI.rev.compl                       ------------------------------------------------------------ 46364
human.FAM230A.ncRNA.18487127.18500594-GGTLC3.18516335.18518165.rev.compl.REF ------------------------------------------------------------ 31039

chimp.LOC112206744-LOC107973052-GGT2.revcompl                           ------------------------------------------------------------ 53081
chimp54546-135457.revcompl.Pan.troglodytes.clone.rp43-41g5.GenBank:AC099533.36 CAGCTGAAAGCAAGGAGTCATGATTGGCAGAGGCCCTGGGGTCTGTTCAGAGGCCCCACG 62327
human.FAM30B-LOC105372935-GGT2.NCBI.                                    ------------------------------------------------------------ 52075
human.FAM230E-LOC105377182-GGT3P.NCBI.                                  ------------------------------------------------------------ 50727
human.FAM230J-LOC105372942-GGTLC5P.NCBI.rev.compl                       ------------------------------------------------------------ 46364
human.FAM230A.ncRNA.18487127.18500594-GGTLC3.18516335.18518165.rev.compl.REF ------------------------------------------------------------ 31039

chimp.LOC112206744-LOC107973052-GGT2.revcompl                           -----------------------------------------------gtgacccccacac 53094
chimp54546-135457.revcompl.Pan.troglodytes.clone.rp43-41g5.GenBank:AC099533.36 GGGCAGCAGCCCACATCCGACACTTCCCCAACCCTTAGTCAGAGGCAGAGGCCCGACTGC 62387
human.FAM30B-LOC105372935-GGT2.NCBI.                                    -----------------------------------------------gtgacccccacac 52088
human.FAM230E-LOC105377182-GGT3P.NCBI.                                  -----------------------------------------------gtgacccccacac 50740
human.FAM230J-LOC105372942-GGTLC5P.NCBI.rev.compl                       ------------------------------------------------------------ 46364
human.FAM230A.ncRNA.18487127.18500594-GGTLC3.18516335.18518165.rev.compl.REF ------------------------------------------------------------ 31039

chimp.LOC112206744-LOC107973052-GGT2.revcompl                           tgccagacccagtgccccctctcagctttactgggctcatcactctccctgagagccgcc 53154
chimp54546-135457.revcompl.Pan.troglodytes.clone.rp43-41g5.GenBank:AC099533.36 GGCCAGCACCTCCATCCAGAGGCCGAGAGAGGTAGAGTAAGGAGGGGCCTGCCAGACACC 62447
human.FAM30B-LOC105372935-GGT2.NCBI.                                    tgccagacccagtgccccctctcagctttactgggctcatcactctccctgagagccgcc 52148
human.FAM230E-LOC105377182-GGT3P.NCBI.                                  tgccagacccagtgccccctctcagctttactgggctcatcactctccctgagagccgcc 50800
human.FAM230J-LOC105372942-GGTLC5P.NCBI.rev.compl                       ------------------------------------------------------------ 46364
human.FAM230A.ncRNA.18487127.18500594-GGTLC3.18516335.18518165.rev.compl.REF ------------------------------------------------------------ 31039

chimp.LOC112206744-LOC107973052-GGT2.revcompl                           cctgcatcccagcacctggctccacccgagtctcccccgcctgccatcctagctcctact 53214
chimp54546-135457.revcompl.Pan.troglodytes.clone.rp43-41g5.GenBank:AC099533.36 CCTGGAACCCAGGGCCCCACTGCTCCCAGAGCCCTGCTGCCTGGGTCC--------CCA 62499
human.FAM30B-LOC105372935-GGT2.NCBI.                                    cctgcatcccagcacctggctccacccgagtctcccccgcctgccatcctagctcctact 52208
```

```
human.FAM230E-LOC105377182-GGT3P.NCBI.                                  cctgcatcccagcacctggctccacccgagtctcccccgcctgccatcctagctcctact 50860
human.FAM230J-LOC105372942-GGTLC5P.NCBI.rev.compl                       ------------------------------------------------------------ 46364
human.FAM230A.ncRNA.18487127.18500594-GGTLC3.18516335.18518165.rev.compl.REF ------------------------------------------------------------ 31039

chimp.LOC112206744-LOC107973052-GGT2.revcompl                           ctcccctctgtctttgctctctctcctggtggtctgcttgacatctgagcttcagcctcc 53274
chimp54546-135457.revcompl.Pan.troglodytes.clone.rp43-41g5.GenBank:AC099533.36 CCCCCTCCTCCCTCTGCCCTCCCAGTTGGGAGCTGATGTGCAGGCTGGCAGGCTCTCTCC 62559
human.FAM30B-LOC105372935-GGT2.NCBI.                                    ctcccctctgtctttgctctctctcctggtggtctgcttgacatctgagcttcagcctcc 52268
human.FAM230E-LOC105377182-GGT3P.NCBI.                                  ctcccctctgtctttgctctctctcctggtggtctgcttgacatctgagcttcagcctcc 50920
human.FAM230J-LOC105372942-GGTLC5P.NCBI.rev.compl                       ------------------------------------------------------------ 46364
human.FAM230A.ncRNA.18487127.18500594-GGTLC3.18516335.18518165.rev.compl.REF ------------------------------------------------------------ 31039

chimp.LOC112206744-LOC107973052-GGT2.revcompl                           atttatgcactgacaactcccaaattgacctgctggcctggactgctcctctgatcacca 53334
chimp54546-135457.revcompl.Pan.troglodytes.clone.rp43-41g5.GenBank:AC099533.36 ATCCGGCTCCCATCACCACGGGTGCTATGTGGAGCTCAGTGAAAGGTGAGCTGTCCAGCA 62619
human.FAM30B-LOC105372935-GGT2.NCBI.                                    atttatgcactgacaactctcaaattgacctgctggcctggactgctcctccgatcacca 52328
human.FAM230E-LOC105377182-GGT3P.NCBI.                                  atttatgcactgacaactctcaaattgacctgctggcctggactgctcctccgatcacca 50980
human.FAM230J-LOC105372942-GGTLC5P.NCBI.rev.compl                       ------------------------------------------------------------ 46364
human.FAM230A.ncRNA.18487127.18500594-GGTLC3.18516335.18518165.rev.compl.REF ------------------------------------------------------------ 31039

chimp.LOC112206744-LOC107973052-GGT2.revcompl                           g----------------------------------------------------------- 53335
chimp54546-135457.revcompl.Pan.troglodytes.clone.rp43-41g5.GenBank:AC099533.36 GCAACTTCAGTAACAACTCCTGGGGCCTGGCATCCACAGCTGCTGTCTGGGGAACTGTG 62679
human.FAM30B-LOC105372935-GGT2.NCBI.                                    g----------------------------------------------------------- 52329
human.FAM230E-LOC105377182-GGT3P.NCBI.                                  g----------------------------------------------------------- 50981
human.FAM230J-LOC105372942-GGTLC5P.NCBI.rev.compl                       ------------------------------------------------------------ 46364
human.FAM230A.ncRNA.18487127.18500594-GGTLC3.18516335.18518165.rev.compl.REF ------------------------------------------------------------ 31039

chimp.LOC112206744-LOC107973052-GGT2.revcompl                           ------------------------------------------------------------ 53335
chimp54546-135457.revcompl.Pan.troglodytes.clone.rp43-41g5.GenBank:AC099533.36 GACACTGTTCTCCCCCTGCACCATGAACAGCTTGTGGCGGGGGGAGCCTGCAGGCCACAC 62739
human.FAM30B-LOC105372935-GGT2.NCBI.                                    ------------------------------------------------------------ 52329
human.FAM230E-LOC105377182-GGT3P.NCBI.                                  ------------------------------------------------------------ 50981
human.FAM230J-LOC105372942-GGTLC5P.NCBI.rev.compl                       ------------------------------------------------------------ 46364
human.FAM230A.ncRNA.18487127.18500594-GGTLC3.18516335.18518165.rev.compl.REF ------------------------------------------------------------ 31039

chimp.LOC112206744-LOC107973052-GGT2.revcompl                           ------------------------------------------------------------ 53335
chimp54546-135457.revcompl.Pan.troglodytes.clone.rp43-41g5.GenBank:AC099533.36 CAGTCCCATGGTTCAGAATCAGGCCCAGAAGGACCCAGAAACAAGCCCTAAATCAAGCAG 62799
human.FAM30B-LOC105372935-GGT2.NCBI.                                    ------------------------------------------------------------ 52329
human.FAM230E-LOC105377182-GGT3P.NCBI.                                  ------------------------------------------------------------ 50981
human.FAM230J-LOC105372942-GGTLC5P.NCBI.rev.compl                       ------------------------------------------------------------ 46364
human.FAM230A.ncRNA.18487127.18500594-GGTLC3.18516335.18518165.rev.compl.REF ------------------------------------------------------------ 31039

chimp.LOC112206744-LOC107973052-GGT2.revcompl                           ------------------------------------------------------------ 53335
chimp54546-135457.revcompl.Pan.troglodytes.clone.rp43-41g5.GenBank:AC099533.36 CTGATGAGGGGGCCGGGAGGAAGCCCCTCTGTGACCCCTCTAACTTCTACCCCCAGCCAG 62859
human.FAM30B-LOC105372935-GGT2.NCBI.                                    ------------------------------------------------------------ 52329
human.FAM230E-LOC105377182-GGT3P.NCBI.                                  ------------------------------------------------------------ 50981
human.FAM230J-LOC105372942-GGTLC5P.NCBI.rev.compl                       ------------------------------------------------------------ 46364
human.FAM230A.ncRNA.18487127.18500594-GGTLC3.18516335.18518165.rev.compl.REF ------------------------------------------------------------ 31039

chimp.LOC112206744-LOC107973052-GGT2.revcompl                           ------------------------------------------------------------ 53335
chimp54546-135457.revcompl.Pan.troglodytes.clone.rp43-41g5.GenBank:AC099533.36 CACTATCCAGGCAGCTAGCCTGCAGGAGCTTGTCCTTAATGTCACCCCAACACAGTCTGT 62919
human.FAM30B-LOC105372935-GGT2.NCBI.                                    ------------------------------------------------------------ 52329
human.FAM230E-LOC105377182-GGT3P.NCBI.                                  ------------------------------------------------------------ 50981
human.FAM230J-LOC105372942-GGTLC5P.NCBI.rev.compl                       ------------------------------------------------------------ 46364
human.FAM230A.ncRNA.18487127.18500594-GGTLC3.18516335.18518165.rev.compl.REF ------------------------------------------------------------ 31039

chimp.LOC112206744-LOC107973052-GGT2.revcompl                           ------------------------------------------------------------ 53335
chimp54546-135457.revcompl.Pan.troglodytes.clone.rp43-41g5.GenBank:AC099533.36 CCTGGTATGTAACACCCTGACTGTCATGACCAATCCACATGTCCCCCACCCAGGAACGCT 62979
human.FAM30B-LOC105372935-GGT2.NCBI.                                    ------------------------------------------------------------ 52329
human.FAM230E-LOC105377182-GGT3P.NCBI.                                  ------------------------------------------------------------ 50981
human.FAM230J-LOC105372942-GGTLC5P.NCBI.rev.compl                       ------------------------------------------------------------ 46364
human.FAM230A.ncRNA.18487127.18500594-GGTLC3.18516335.18518165.rev.compl.REF ------------------------------------------------------------ 31039

chimp.LOC112206744-LOC107973052-GGT2.revcompl                           ------------------------------------------------------------ 53335
chimp54546-135457.revcompl.Pan.troglodytes.clone.rp43-41g5.GenBank:AC099533.36 CTGTTTCCCACTTTATGGGGAGCACAAGCTCTATGTCAGCGGTTCCTGGGTATGCCAAGA 63039
human.FAM30B-LOC105372935-GGT2.NCBI.                                    ------------------------------------------------------------ 52329
human.FAM230E-LOC105377182-GGT3P.NCBI.                                  ------------------------------------------------------------ 50981
human.FAM230J-LOC105372942-GGTLC5P.NCBI.rev.compl                       ------------------------------------------------------------ 46364
human.FAM230A.ncRNA.18487127.18500594-GGTLC3.18516335.18518165.rev.compl.REF ------------------------------------------------------------ 31039

chimp.LOC112206744-LOC107973052-GGT2.revcompl                           --acctgagtatctacctgcctgcttgaagaaagcatctcaaacttcaacgtgcccaaaa 53393
chimp54546-135457.revcompl.Pan.troglodytes.clone.rp43-41g5.GenBank:AC099533.36 CCACTGGCTGGGCTGGGTGAATGAGCAGTATCATCTTGTTAAAAATACACCTTCCTGGGT 63099
human.FAM30B-LOC105372935-GGT2.NCBI.                                    --acctgaggatctacctgcctgctcgaagaaagcatctcaaacttcaacgtgcccaaaa 52387
human.FAM230E-LOC105377182-GGT3P.NCBI.                                  --acctgaggatctacctgcctgcttgaagaaagcatctcaaacttcaacgtgcccaaaa 51039
human.FAM230J-LOC105372942-GGTLC5P.NCBI.rev.compl                       ------------------------------------------------------------ 46364
human.FAM230A.ncRNA.18487127.18500594-GGTLC3.18516335.18518165.rev.compl.REF ------------------------------------------------------------ 31039

chimp.LOC112206744-LOC107973052-GGT2.revcompl                           ccgagctcctgagtgtctgctcaacctgcttcctgagaaccctgcctgtctccattaggg 53453
chimp54546-135457.revcompl.Pan.troglodytes.clone.rp43-41g5.GenBank:AC099533.36 TCCACCTGCAGAGAACTGGACTCATAATCTGAGTGGGACCTGGGATTTACACAGCTAGAA 63159
human.FAM30B-LOC105372935-GGT2.NCBI.                                    ccgagctcctgagtgtctgctcaacctgcttcctgagaaccctgcctgtctccattaggg 52447
human.FAM230E-LOC105377182-GGT3P.NCBI.                                  ccgagctcctgagtgtctgctcaacctgcttcctgagaaccctgcctgtctccattaggg 51099
human.FAM230J-LOC105372942-GGTLC5P.NCBI.rev.compl                       ------------------------------------------------------------ 46364
human.FAM230A.ncRNA.18487127.18500594-GGTLC3.18516335.18518165.rev.compl.REF ------------------------------------------------------------ 31039

chimp.LOC112206744-LOC107973052-GGT2.revcompl                           tcacccccatccttccaggtacagacaaaagatcaggggtccccggggactccctacacaa 53513
chimp54546-135457.revcompl.Pan.troglodytes.clone.rp43-41g5.GenBank:AC099533.36 ACCCCAGGTAGTTCTGGAGTGAAGCCAGGTCACAGAGCCACCGTCTGACTCCATAGAAGG 63219
human.FAM30B-LOC105372935-GGT2.NCBI.                                    tcaccccatccttccaggtacagacaaaagatcaggggtccccggggactccctacacaa 52507
human.FAM230E-LOC105377182-GGT3P.NCBI.                                  tcaccccatccttccaggtacagacaaaagatcaggggtccccggggactccctacacaa 51159
human.FAM230J-LOC105372942-GGTLC5P.NCBI.rev.compl                       ------------------------------------------------------------ 46364
human.FAM230A.ncRNA.18487127.18500594-GGTLC3.18516335.18518165.rev.compl.REF ------------------------------------------------------------ 31039

chimp.LOC112206744-LOC107973052-GGT2.revcompl                           gcgtcacacccaacccatcctcaaatccgcaggctccacttccaagtgtgtctgtccagc 53573
chimp54546-135457.revcompl.Pan.troglodytes.clone.rp43-41g5.GenBank:AC099533.36 CCACTGCATCTGGCCTTGTGTTTCCATCCCTTGATTCAGGTGATGGTTCACTTCAGCCAC 63279
human.FAM30B-LOC105372935-GGT2.NCBI.                                    gcgtcacacccaacccatcctcaaatccacaggctccacttccaagtgtgtctgtgcagc 52567
human.FAM230E-LOC105377182-GGT3P.NCBI.                                  gcgtcacacccaacccatcctcaaatccacaggctccacttccaagtgtgtctgtgcagc 51219
human.FAM230J-LOC105372942-GGTLC5P.NCBI.rev.compl                       ------------------------------------------------------------ 46364
human.FAM230A.ncRNA.18487127.18500594-GGTLC3.18516335.18518165.rev.compl.REF ------------------------------------------------------------ 31039

chimp.LOC112206744-LOC107973052-GGT2.revcompl                           gtcagccacttcccagcac-----------tctctccacgaattactgcagtgacctccg 53622
chimp54546-135457.revcompl.Pan.troglodytes.clone.rp43-41g5.GenBank:AC099533.36 AAGGCAAGCTGCCTAACAGCAGGGACCACACCTCATGATGTCTGCACACATTCACATGCG 63339
human.FAM30B-LOC105372935-GGT2.NCBI.                                    gtcagccacttcccagcac-----------cctctccacgaattactgcagtgacctccg 52616
human.FAM230E-LOC105377182-GGT3P.NCBI.                                  gtcagccacttcccagcac-----------cctctccacgaattactgcagtgacctccg 51268
human.FAM230J-LOC105372942-GGTLC5P.NCBI.rev.compl                       ------------------------------------------------------------ 46364
human.FAM230A.ncRNA.18487127.18500594-GGTLC3.18516335.18518165.rev.compl.REF ------------------------------------------------------------ 31039

chimp.LOC112206744-LOC107973052-GGT2.revcompl                           gacaggcccccacatgctccctgcccccttacacagcaacccaaggggtccatagaccaga 53682
chimp54546-135457.revcompl.Pan.troglodytes.clone.rp43-41g5.GenBank:AC099533.36 TGCACATATACATACACACACGACGGTGCTGAGCACAGCGCCAGGCCTAGAGAGGGGC 63399
human.FAM30B-LOC105372935-GGT2.NCBI.                                    gacaggtccccacatgctccctgcccccttacacagcaatcaaggggtccatagaccaga 52676
human.FAM230E-LOC105377182-GGT3P.NCBI.                                  gacaggtccccacatgctccctgcccccttacacagcaatccaaggggtccatagaccaga 51328
human.FAM230J-LOC105372942-GGTLC5P.NCBI.rev.compl                       ------------------------------------------------------------ 46364
human.FAM230A.ncRNA.18487127.18500594-GGTLC3.18516335.18518165.rev.compl.REF ------------------------------------------------------------ 31039

chimp.LOC112206744-LOC107973052-GGT2.revcompl                           t--------ccatcgccttccactcacatactccacgagccccccacttccctcagacagg 53734
chimp54546-135457.revcompl.Pan.troglodytes.clone.rp43-41g5.GenBank:AC099533.36 TGAGGACAGGCATCATTTCCACTGCAAGGGACCCTAGCGGCCACAGCCACCTGGGCCTTC 63459
human.FAM30B-LOC105372935-GGT2.NCBI.                                    t--------ccatccccttccagtcacacactccacgagccccccacttccctcagacagg 52728
human.FAM230E-LOC105377182-GGT3P.NCBI.                                  t--------ccatccccttccagtcacacactccacgagccccccacttccctcagacagg 51380
human.FAM230J-LOC105372942-GGTLC5P.NCBI.rev.compl                       ------------------------------------------------------------ 46364
human.FAM230A.ncRNA.18487127.18500594-GGTLC3.18516335.18518165.rev.compl.REF ------------------------------------------------------------ 31039

chimp.LOC112206744-LOC107973052-GGT2.revcompl                           aagcagaggcttcaccataacctaagagatcccacacaacct----gggccattcccctt 53790
chimp54546-135457.revcompl.Pan.troglodytes.clone.rp43-41g5.GenBank:AC099533.36 TGCAGGTGAAGTGGCCAGAAAATAAGAAGCTCTGGAGCAGGTCAAGGAGCCCAGCCAGCT 63519
human.FAM30B-LOC105372935-GGT2.NCBI.                                    aagcagaggcttcaccataacctaagagatcccgcacaacct----gggccattcccctt 52784
human.FAM230E-LOC105377182-GGT3P.NCBI.                                  aagcagaggcttcaccataacctaagagatcccgcacaacct----gggccattcccctt 51436
human.FAM230J-LOC105372942-GGTLC5P.NCBI.rev.compl                       ------------------------------------------------------------ 46364
human.FAM230A.ncRNA.18487127.18500594-GGTLC3.18516335.18518165.rev.compl.REF ------------------------------------------------------------ 31039

chimp.LOC112206744-LOC107973052-GGT2.revcompl                           gggtcacttgctgcagcctccccagctccccacagggctctgtccctgc----catcaca 53846
chimp54546-135457.revcompl.Pan.troglodytes.clone.rp43-41g5.GenBank:AC099533.36 GCCCCTACTTCACAGGTGTGTGCACCTGGGCCTCCCTCCCTGCACCTGCTCAGCATGAAG 63579
human.FAM30B-LOC105372935-GGT2.NCBI.                                    ggatcacttgctgcagcctccccagctccccacagggctctgtccctgc----catcaca 52840
human.FAM230E-LOC105377182-GGT3P.NCBI.                                  ggatcacttgctgcagcctccccagctccccacagggctctgtccctgc----catcaca 51492
```

```
human.FAM230J-LOC105372942-GGTLC5P.NCBI.rev.compl                          ------------------------------------------------------------ 46364
human.FAM230A.ncRNA.18487127.18500594-GGTLC3.18516335.18518165.rev.compl.REF ------------------------------------------------------------ 31039

chimp.LOC112206744-LOC107973052-GGT2.revcompl                              cctggatagcagaccaggagatagctcccctgaccccatctctgcctctgggtctt---- 53902
chimp54546-135457.revcompl.Pan.troglodytes.clone.rp43-41g5.GenBank:AC099533.36 GCTCAGCACAGGCACACGTGGGGGGTGACCCCTTCTCACCTCCCCTTCTAGGTCCTACTG 63639
human.FAM30B-LOC105372935-GGT2.NCBI.                                       cctggatagcagaccaggagataactcccctgaccccatctctgcctctgggtctt---- 52896
human.FAM230E-LOC105377182-GGT3P.NCBI.                                     cctggatagcagaccaggagataactcccctgaccccatctctgcctctgggtctt---- 51548
human.FAM230J-LOC105372942-GGTLC5P.NCBI.rev.compl                          ------------------------------------------------------------ 46364
human.FAM230A.ncRNA.18487127.18500594-GGTLC3.18516335.18518165.rev.compl.REF ------------------------------------------------------------ 31039

chimp.LOC112206744-LOC107973052-GGT2.revcompl                              ------------------------------------------------------------ 53902
chimp54546-135457.revcompl.Pan.troglodytes.clone.rp43-41g5.GenBank:AC099533.36 GTTAACATTTCCCAAGGCTCTGAACTGTGCAGCCAGAGTGGCACAGAGAAGCAAAGGGCG 63699
human.FAM30B-LOC105372935-GGT2.NCBI.                                       ------------------------------------------------------------ 52896
human.FAM230E-LOC105377182-GGT3P.NCBI.                                     ------------------------------------------------------------ 51548
human.FAM230J-LOC105372942-GGTLC5P.NCBI.rev.compl                          ------------------------------------------------------------ 46364
human.FAM230A.ncRNA.18487127.18500594-GGTLC3.18516335.18518165.rev.compl.REF ------------------------------------------------------------ 31039

chimp.LOC112206744-LOC107973052-GGT2.revcompl                              ------------------------------------------------------------ 53902
chimp54546-135457.revcompl.Pan.troglodytes.clone.rp43-41g5.GenBank:AC099533.36 GCGGGACGTCTCCCACCAGGGGCCATGTGGCCACAGGCTCTCATTCCGCACCTGGCCCTG 63759
human.FAM30B-LOC105372935-GGT2.NCBI.                                       ------------------------------------------------------------ 52896
human.FAM230E-LOC105377182-GGT3P.NCBI.                                     ------------------------------------------------------------ 51548
human.FAM230J-LOC105372942-GGTLC5P.NCBI.rev.compl                          ------------------------------------------------------------ 46364
human.FAM230A.ncRNA.18487127.18500594-GGTLC3.18516335.18518165.rev.compl.REF ------------------------------------------------------------ 31039

chimp.LOC112206744-LOC107973052-GGT2.revcompl                              ----tgctcagatgtccccttccctgactaggtcaccctccatagagtcccagattttga 53958
chimp54546-135457.revcompl.Pan.troglodytes.clone.rp43-41g5.GenBank:AC099533.36 AAGGTTGTCAGAGGCCCACACTCCAGGCCTCAGGGACCCCACCCTGAGAAGCAGCATCTGT 63819
human.FAM30B-LOC105372935-GGT2.NCBI.                                       ----tgctcagatgtccccttccctgactaggtcaccctccatagagtcccagattttga 52952
human.FAM230E-LOC105377182-GGT3P.NCBI.                                     ----tgctcagatgtccccttccctgactaggtcaccctccatagagtcccagattttga 51604
human.FAM230J-LOC105372942-GGTLC5P.NCBI.rev.compl                          ------------------------------------------------------------ 46364
human.FAM230A.ncRNA.18487127.18500594-GGTLC3.18516335.18518165.rev.compl.REF ------------------------------------------------------------ 31039

chimp.LOC112206744-LOC107973052-GGT2.revcompl                              ggccctccaggtctgtttttctacagcctgtaacacacccgcacctgcctagtttctttt 54018
chimp54546-135457.revcompl.Pan.troglodytes.clone.rp43-41g5.GenBank:AC099533.36 GAGCAGGGAGCGGTCAGTCAGAACCACACATAAAAAGGAAACTCTCAAGTGGTACTTTCC 63879
human.FAM30B-LOC105372935-GGT2.NCBI.                                       ggccctccaggtctgtttttctacagcccgtaacacacccgcacttgcctagtttctcct 53012
human.FAM230E-LOC105377182-GGT3P.NCBI.                                     ggccctccaggtctgtttttctacagcccgtaacacacccgcacttgcctagtttctcct 51664
human.FAM230J-LOC105372942-GGTLC5P.NCBI.rev.compl                          ------------------------------------------------------------ 46364
human.FAM230A.ncRNA.18487127.18500594-GGTLC3.18516335.18518165.rev.compl.REF ------------------------------------------------------------ 31039

chimp.LOC112206744-LOC107973052-GGT2.revcompl                              cccacctggagtgtcacagatttcatctgccgtctttgtttttcaccccagcttcaggaa 54078
chimp54546-135457.revcompl.Pan.troglodytes.clone.rp43-41g5.GenBank:AC099533.36 CCCGGAGTG--------------------------------------------------- 63888
human.FAM30B-LOC105372935-GGT2.NCBI.                                       cccacctggagtgtcacagatttcatctgccgtctttgtttttcaccccagcttcaggaa 53072
human.FAM230E-LOC105377182-GGT3P.NCBI.                                     cccacctggagtgtcacagatttcatctgccgtctttgtttttcaccccagcttcaggaa 51724
human.FAM230J-LOC105372942-GGTLC5P.NCBI.rev.compl                          ------------------------------------------------------------ 46364
human.FAM230A.ncRNA.18487127.18500594-GGTLC3.18516335.18518165.rev.compl.REF ------------------------------------------------------------ 31039

chimp.LOC112206744-LOC107973052-GGT2.revcompl                              caacagctgattctttaagacaatgctcaatacattctagttaaataaatgattctaagc 54138
chimp54546-135457.revcompl.Pan.troglodytes.clone.rp43-41g5.GenBank:AC099533.36 --------------------------------------AACCCTAAGCAGAGGCAGGAGA 63910
human.FAM30B-LOC105372935-GGT2.NCBI.                                       caacagctgattctttaagacaatgctcaatacattctagtcaaataaatggttttaagc 53132
human.FAM230E-LOC105377182-GGT3P.NCBI.                                     caacagctgattctttaagacaatgctcaatacattctagtcaaataaatggttttaagc 51784
human.FAM230J-LOC105372942-GGTLC5P.NCBI.rev.compl                          ------------------------------------------------------------ 46364
human.FAM230A.ncRNA.18487127.18500594-GGTLC3.18516335.18518165.rev.compl.REF ------------------------------------------------------------ 31039

chimp.LOC112206744-LOC107973052-GGT2.revcompl                              gtccacaaggtgccaagcctgtgattcccgcattgtcttaccctgagcaacttcacatct 54198
chimp54546-135457.revcompl.Pan.troglodytes.clone.rp43-41g5.GenBank:AC099533.36 GGACAACATGAGGGATGCGGACAGAGTGCGCTTTCTCTGGAAAGGAGGAGTTAATACCA 63970
human.FAM30B-LOC105372935-GGT2.NCBI.                                       gtccacaaggtgccaagcctatggttcccgcattctcttaccctcagcaacttcatgtct 53192
human.FAM230E-LOC105377182-GGT3P.NCBI.                                     gtccacaaggtgccaagcctatgattcccgcattctcttaccctcagcaacttcatgtct 51844
human.FAM230J-LOC105372942-GGTLC5P.NCBI.rev.compl                          ------------------------------------------------------------ 46364
human.FAM230A.ncRNA.18487127.18500594-GGTLC3.18516335.18518165.rev.compl.REF ------------------------------------------------------------ 31039

chimp.LOC112206744-LOC107973052-GGT2.revcompl                              acagatgctgagtttctcaatgagtattaaaaacaaatgaaagattggtcgggcacagtg 54258
chimp54546-135457.revcompl.Pan.troglodytes.clone.rp43-41g5.GenBank:AC099533.36 CTGCTGGGAGCTTTAAAAAATGGAAA---------------------------------- 63996
human.FAM30B-LOC105372935-GGT2.NCBI.                                       acagatgctgagtttctcaatgagtattaaaaacaaatgaaagattggtcgggcacagtg 53252
human.FAM230E-LOC105377182-GGT3P.NCBI.                                     acagatgctgagtttctcagtgagtattaaaaacaaatgaaagattggtcgggcacagtg 51904
human.FAM230J-LOC105372942-GGTLC5P.NCBI.rev.compl                          ------------------------------------------------------------ 46364
human.FAM230A.ncRNA.18487127.18500594-GGTLC3.18516335.18518165.rev.compl.REF ------------------------------------------------------------ 31039

chimp.LOC112206744-LOC107973052-GGT2.revcompl                              gctcacgcctgtaattccagcactttgggaggctgaggcaggggggatcacgaggtcagga 54318
chimp54546-135457.revcompl.Pan.troglodytes.clone.rp43-41g5.GenBank:AC099533.36 --CCCTGCCTGTGATATCAGCACTTTGAAAGGCTGAGGCAGGAGGATCACTTGAGCCCAT 64054
human.FAM30B-LOC105372935-GGT2.NCBI.                                       gctcacgcctgtaattccagcactttgggaggctgaggcaggggggatcacgaggtcagga 53312
human.FAM230E-LOC105377182-GGT3P.NCBI.                                     gctcacgcctgtaattccagcactttgggaggctgaggcaggggggatcacgaggtcagga 51964
human.FAM230J-LOC105372942-GGTLC5P.NCBI.rev.compl                          ------------------------------------------------------------ 46364
human.FAM230A.ncRNA.18487127.18500594-GGTLC3.18516335.18518165.rev.compl.REF ------------------------------------------------------------ 31039

chimp.LOC112206744-LOC107973052-GGT2.revcompl                              gatcgagggaccagcctgggcaacatagtgaaaacccgtctctactaaaaaatacaaaaa 54378
chimp54546-135457.revcompl.Pan.troglodytes.clone.rp43-41g5.GenBank:AC099533.36 GAGTTTAAGACCATCCTGGGAAATATAGTGAGACCTTGTCTGTATTAAAAAAAAAAAAAA 64114
human.FAM30B-LOC105372935-GGT2.NCBI.                                       gatcgagggaccagcctgggcaacatagtgaaaacccgtctctactaaaaaatacaaaaa 53372
human.FAM230E-LOC105377182-GGT3P.NCBI.                                     gatcgagggaccagcctgggcaacacagtgaaaacccgtctctactaaaaaatacaaaaa 52024
human.FAM230J-LOC105372942-GGTLC5P.NCBI.rev.compl                          ------------------------------------------------------------ 46364
human.FAM230A.ncRNA.18487127.18500594-GGTLC3.18516335.18518165.rev.compl.REF ------------------------------------------------------------ 31039

chimp.LOC112206744-LOC107973052-GGT2.revcompl                              att--------agccgggtgtggtggcaggcgcctgtagtcccagctactcaggaggctg 54430
chimp54546-135457.revcompl.Pan.troglodytes.clone.rp43-41g5.GenBank:AC099533.36 TTTAAAAAGGCAGCCAAGCATGGTGGCACATACCTGCAGTCCCAGCTGCTTGAGAGGCTG 64174
human.FAM30B-LOC105372935-GGT2.NCBI.                                       att--------agccgggcgcggtggcgggcgcctgtagtcccagctactcgggaggctg 53424
human.FAM230E-LOC105377182-GGT3P.NCBI.                                     att--------agccgggtgtggtggcgggcgcctgtagtcccagctactcgggaggctg 52076
human.FAM230J-LOC105372942-GGTLC5P.NCBI.rev.compl                          ------------------------------------------------------------ 46364
human.FAM230A.ncRNA.18487127.18500594-GGTLC3.18516335.18518165.rev.compl.REF ------------------------------------------------------------ 31039

chimp.LOC112206744-LOC107973052-GGT2.revcompl                              aggcaggagaatggcgtgaactcaggaggcagagcttgcagtgagccgagatcgcaccac 54490
chimp54546-135457.revcompl.Pan.troglodytes.clone.rp43-41g5.GenBank:AC099533.36 AGGTGGGAGCATTGCTCGAGCCTGGAAGGTGGAGGCTGCTGTGAGCCACGATCATGCCAT 64234
human.FAM30B-LOC105372935-GGT2.NCBI.                                       aggcaggagaatggcgcgaactcaggaggcagagcttgcagtgagccgagatcgcaccac 53484
human.FAM230E-LOC105377182-GGT3P.NCBI.                                     aggcaggagaatggcgtgaactcaggaggcggaacttgcagtgagctgagatcgcaccac 52136
human.FAM230J-LOC105372942-GGTLC5P.NCBI.rev.compl                          ------------------------------------------------------------ 46364
human.FAM230A.ncRNA.18487127.18500594-GGTLC3.18516335.18518165.rev.compl.REF ------------------------------------------------------------ 31039

chimp.LOC112206744-LOC107973052-GGT2.revcompl                              tgcactccaacctgggcgacagagtaagactccgtctcaaaaaaaaaaaaaagaaaaaaa 54550
chimp54546-135457.revcompl.Pan.troglodytes.clone.rp43-41g5.GenBank:AC099533.36 TGTACTCCAGCCTGCGCAACAGAGCGAGAGACCCTGGCTCTAACTAACTAACCAAGTAAC 64294
human.FAM30B-LOC105372935-GGT2.NCBI.                                       tgcactccaacctgggcgacagagtgagactctgtctcaaaaaaaaaaaaaagaaaaaaa 53544
human.FAM230E-LOC105377182-GGT3P.NCBI.                                     tgcactccaacctgggcgacagagtgagactccgtctcaaaaaaaaaaaaaagaaaaaaa 52196
human.FAM230J-LOC105372942-GGTLC5P.NCBI.rev.compl                          ------------------------------------------------------------ 46364
human.FAM230A.ncRNA.18487127.18500594-GGTLC3.18516335.18518165.rev.compl.REF ------------------------------------------------------------ 31039

chimp.LOC112206744-LOC107973052-GGT2.revcompl                              aatcaaagattgagtatgttgcagaagactccaaagggcaccacccaggacccccacctg 54610
chimp54546-135457.revcompl.Pan.troglodytes.clone.rp43-41g5.GenBank:AC099533.36 TGGAAACATAGAAACATGTT---------------------------------------- 64314
human.FAM30B-LOC105372935-GGT2.NCBI.                                       aatcaaagattgagtatgttgcagaagactccaaagggcaccacccaggacccccacctg 53604
human.FAM230E-LOC105377182-GGT3P.NCBI.                                     aatcaaagattgagtatgttgcagaagactccaaagggcaccacccaggacccccacctg 52256
human.FAM230J-LOC105372942-GGTLC5P.NCBI.rev.compl                          ------------------------------------------------------------ 46364
human.FAM230A.ncRNA.18487127.18500594-GGTLC3.18516335.18518165.rev.compl.REF ------------------------------------------------------------ 31039

chimp.LOC112206744-LOC107973052-GGT2.revcompl                              aagtctaagacctgctgtggtaagtgtgtcctgcccctccatcctccaac---ttttttt 54667
chimp54546-135457.revcompl.Pan.troglodytes.clone.rp43-41g5.GenBank:AC099533.36 ------------------------------------------------------------ 64314
human.FAM30B-LOC105372935-GGT2.NCBI.                                       aagtctaagacctgctatggtgagtgtgtcctgcccctccatcctccaactttttttttt 53664
human.FAM230E-LOC105377182-GGT3P.NCBI.                                     aagtctaagacctgctatggtgagtgtgtcctgcccctccatcctccaactttttttttt 52316
human.FAM230J-LOC105372942-GGTLC5P.NCBI.rev.compl                          ------------------------------------------------------------ 46364
human.FAM230A.ncRNA.18487127.18500594-GGTLC3.18516335.18518165.rev.compl.REF ------------------------------------------------------------ 31039

chimp.LOC112206744-LOC107973052-GGT2.revcompl                              tttttttttttttgagacggagcttcgttcttgttgcccaggctggagtgcagtggcatg 54727
chimp54546-135457.revcompl.Pan.troglodytes.clone.rp43-41g5.GenBank:AC099533.36 ------------------------------------------------------------ 64314
human.FAM30B-LOC105372935-GGT2.NCBI.                                       tttttttttttttgagacggagcttcgttcttgttgcccaggctggagtgcagtggcatg 53724
human.FAM230E-LOC105377182-GGT3P.NCBI.                                     tttttttttttttgagacggagcttcgttcttgttgcccaggctggagtgcagtggcatg 52376
human.FAM230J-LOC105372942-GGTLC5P.NCBI.rev.compl                          ------------------------------------------------------------ 46364
human.FAM230A.ncRNA.18487127.18500594-GGTLC3.18516335.18518165.rev.compl.REF ------------------------------------------------------------ 31039

chimp.LOC112206744-LOC107973052-GGT2.revcompl                              atctcggctcactgcaatctccacctcgtgggttcaagcgattctcctgcctcagcctcc 54787
chimp54546-135457.revcompl.Pan.troglodytes.clone.rp43-41g5.GenBank:AC099533.36 ------------------------------------------------------------ 64314
human.FAM30B-LOC105372935-GGT2.NCBI.                                       atctcggctcactgcaatctctacctcgtgggttcaagcgattctcctgccttagcctcc 53784
human.FAM230E-LOC105377182-GGT3P.NCBI.                                     atctcggctcacagcaatctctacctcgtgggttcaagcgattctcctgccttagcctcc 52436
human.FAM230J-LOC105372942-GGTLC5P.NCBI.rev.compl                          ------------------------------------------------------------ 46364
```

```
human.FAM230A.ncRNA.18487127.18500594-GGTLC3.18516335.18518165.rev.compl.REF        ------------------------------------------------------------ 31039

chimp.LOC112206744-LOC107973052-GGT2.revcompl                                       tgagtagctgggatattacaggcctgtgccaccacgcccaactaattatatacttttag 54847
chimp54546-135457.revcompl.Pan.troglodytes.clone.rp43-41g5.GenBank:AC099533.36     ---------------------------------AAGCAGGAGACATTCTTCCCTTTCCTT 64341
human.FAM30B-LOC105372935-GGT2.NCBI.                                                tgagtagctgggatattacaggcctgtgccaccacgcccgactaattattgtacttttag 53844
human.FAM230E-LOC105377182-GGT3P.NCBI.                                              tgagtagctgggatattacaggcctgtgccaccacgcccgactaattattgtacttttag 52496
human.FAM230J-LOC105372942-GGTLC5P.NCBI.rev.compl                                   ------------------------------------------------------------ 46364
human.FAM230A.ncRNA.18487127.18500594-GGTLC3.18516335.18518165.rev.compl.REF        ------------------------------------------------------------ 31039

chimp.LOC112206744-LOC107973052-GGT2.revcompl                                       tagagaaagggtttcactatgttggccaggctggtcttgaactc-ctgacctaggtgatc 54906
chimp54546-135457.revcompl.Pan.troglodytes.clone.rp43-41g5.GenBank:AC099533.36     CAAAGAACAGCCATCACTG----GGCCCCTCTCCCCTTGAAAACTCAGAACTGGCAGAAG 64397
human.FAM30B-LOC105372935-GGT2.NCBI.                                                tagagaaagggtttcactatgttggccaggctggtcttggactc-ctgacctaggtgatc 53903
human.FAM230E-LOC105377182-GGT3P.NCBI.                                              tagagaaagggtttcactatgttggccaggctggtcttggactc-ctgacctaggtgatc 52555
human.FAM230J-LOC105372942-GGTLC5P.NCBI.rev.compl                                   ------------------------------------------------------------ 46364
human.FAM230A.ncRNA.18487127.18500594-GGTLC3.18516335.18518165.rev.compl.REF        ------------------------------------------------------------ 31039

chimp.LOC112206744-LOC107973052-GGT2.revcompl                                       catctgcctcagcctcccaaagtgctgggattacaggcgtgagcctgtgaaaaaaaggcc 54966
chimp54546-135457.revcompl.Pan.troglodytes.clone.rp43-41g5.GenBank:AC099533.36     CAGCTAAC-AAGCAACCCAGAGTCCAGGTAAGAAAA------------------------ 64432
human.FAM30B-LOC105372935-GGT2.NCBI.                                                catctgcctcagcctcccaaagtgctgggattacaggcgtgagcctgtgaaaaaaaggcc 53963
human.FAM230E-LOC105377182-GGT3P.NCBI.                                              catctgcctcagcctcccaaagtgctgggattacaggcgtgagcctgtgaaaaaaaggcc 52615
human.FAM230J-LOC105372942-GGTLC5P.NCBI.rev.compl                                   ------------------------------------------------------------ 46364
human.FAM230A.ncRNA.18487127.18500594-GGTLC3.18516335.18518165.rev.compl.REF        ------------------------------------------------------------ 31039

chimp.LOC112206744-LOC107973052-GGT2.revcompl                                       cagccttttttttttttttttttttgacagcgtctcactttgttgcccaagctagagt 55026
chimp54546-135457.revcompl.Pan.troglodytes.clone.rp43-41g5.GenBank:AC099533.36     ------------------------------------------------------------ 64432
human.FAM30B-LOC105372935-GGT2.NCBI.                                                cagcc----ttttttttttttttttttgacagggtctcactttgttgcccaagctagagt 54019
human.FAM230E-LOC105377182-GGT3P.NCBI.                                              cagcc---tttttttttttttttttttgacagggtctcactttgttgcccaagctagagt 52672
human.FAM230J-LOC105372942-GGTLC5P.NCBI.rev.compl                                   ------------------------------------------------------------ 46364
human.FAM230A.ncRNA.18487127.18500594-GGTLC3.18516335.18518165.rev.compl.REF        ------------------------------------------------------------ 31039

chimp.LOC112206744-LOC107973052-GGT2.revcompl                                       gtagtggtataatcatggctgactgcagcctcaacctcctgggctcaagtgatcctccca 55086
chimp54546-135457.revcompl.Pan.troglodytes.clone.rp43-41g5.GenBank:AC099533.36     --------------------GACAGAAAGAGAAGCTGTCCAGACATCAGGCAGCTGCCCC 64472
human.FAM30B-LOC105372935-GGT2.NCBI.                                                gtagtggtataatcatggctgactgcagcctcaacctcctgggctcaagtgatcctccca 54079
human.FAM230E-LOC105377182-GGT3P.NCBI.                                              gtagtggtataatcatggctgactgcagcctcaacctcctgggctcaagtgatcctccca 52732
human.FAM230J-LOC105372942-GGTLC5P.NCBI.rev.compl                                   ------------------------------------------------------------ 46364
human.FAM230A.ncRNA.18487127.18500594-GGTLC3.18516335.18518165.rev.compl.REF        ------------------------------------------------------------ 31039

chimp.LOC112206744-LOC107973052-GGT2.revcompl                                       ccttagcctcccgagtagctgggaccataagcacacaccgccatacctagctaa-ttttt 55145
chimp54546-135457.revcompl.Pan.troglodytes.clone.rp43-41g5.GenBank:AC099533.36     TCTGAGCCCGCCTGGAACCATGGCCCACATGCCAGCCCATTAATAAAATAATAA------ 64526
human.FAM30B-LOC105372935-GGT2.NCBI.                                                ccttagcctcccgagtaactgggaccataagcacacaccgccatacct-gctaattttt 54138
human.FAM230E-LOC105377182-GGT3P.NCBI.                                              ccttagcctcccgagtaactgggaccataagcacacaccgccatacctagctaattttt 52792
human.FAM230J-LOC105372942-GGTLC5P.NCBI.rev.compl                                   ------------------------------------------------------------ 46364
human.FAM230A.ncRNA.18487127.18500594-GGTLC3.18516335.18518165.rev.compl.REF        ------------------------------------------------------------ 31039

chimp.LOC112206744-LOC107973052-GGT2.revcompl                                       tttccatttttgtagagatggagtcttgctatgttgtccaggcaggtctcctgggctca 55205
chimp54546-135457.revcompl.Pan.troglodytes.clone.rp43-41g5.GenBank:AC099533.36     ------------------------------------------------------------ 64526
human.FAM30B-LOC105372935-GGT2.NCBI.                                                tttccatttttgtagagatggagtcttgctatgttgtccaggcaggtctcctgggctca 54198
human.FAM230E-LOC105377182-GGT3P.NCBI.                                              tttccatttttgtagagatggagtcttgctatgttgtccaggcaggtctcctgggctca 52852
human.FAM230J-LOC105372942-GGTLC5P.NCBI.rev.compl                                   ------------------------------------------------------------ 46364
human.FAM230A.ncRNA.18487127.18500594-GGTLC3.18516335.18518165.rev.compl.REF        ------------------------------------------------------------ 31039

chimp.LOC112206744-LOC107973052-GGT2.revcompl                                       tgcgctcctcctgacttggcctcctaaagtgctaggattagaggcgtgagttgttgagac 55265
chimp54546-135457.revcompl.Pan.troglodytes.clone.rp43-41g5.GenBank:AC099533.36     ------------------------------------------------------------ 64526
human.FAM30B-LOC105372935-GGT2.NCBI.                                                tgcgctcctcctgacttggcctcctaaagtgctaggattagaggcgtgagttgttgagac 54258
human.FAM230E-LOC105377182-GGT3P.NCBI.                                              tgcgctcctcctgacttggcctcctaaagtgctaggattagaggtgtgagttgttgagac 52912
human.FAM230J-LOC105372942-GGTLC5P.NCBI.rev.compl                                   ------------------------------------------------------------ 46364
human.FAM230A.ncRNA.18487127.18500594-GGTLC3.18516335.18518165.rev.compl.REF        ------------------------------------------------------------ 31039

chimp.LOC112206744-LOC107973052-GGT2.revcompl                                       cctcccatcctccaacttttatctcacaatctattgtgcctcctttggggacagacagtg 55325
chimp54546-135457.revcompl.Pan.troglodytes.clone.rp43-41g5.GenBank:AC099533.36     ------------------------------------------------------------ 64526
human.FAM30B-LOC105372935-GGT2.NCBI.                                                cctcccatcctccaacttttatctcacaatctattgtgcctcctttggggacagacagtg 54318
human.FAM230E-LOC105377182-GGT3P.NCBI.                                              cctcccatcctccaacttttatctcacactctattgtgcctcctttggggacagacagtg 52972
human.FAM230J-LOC105372942-GGTLC5P.NCBI.rev.compl                                   ------------------------------------------------------------ 46364
human.FAM230A.ncRNA.18487127.18500594-GGTLC3.18516335.18518165.rev.compl.REF        ------------------------------------------------------------ 31039

chimp.LOC112206744-LOC107973052-GGT2.revcompl                                       gcttcctggatggacagtggcttcccttcaggtacctggggaatttgggggcctcctctc 55385
chimp54546-135457.revcompl.Pan.troglodytes.clone.rp43-41g5.GenBank:AC099533.36     ------------------------------------------------------------ 64526
human.FAM30B-LOC105372935-GGT2.NCBI.                                                gcttcctggatggacagtggcttcccctcaggtacctggggaatttgggggcctcctccc 54378
human.FAM230E-LOC105377182-GGT3P.NCBI.                                              gcttcctggatggacagtggcttcccctcaggtacctggggaatttgggggcctcctccc 53032
human.FAM230J-LOC105372942-GGTLC5P.NCBI.rev.compl                                   ------------------------------------------------------------ 46364
human.FAM230A.ncRNA.18487127.18500594-GGTLC3.18516335.18518165.rev.compl.REF        ------------------------------------------------------------ 31039

chimp.LOC112206744-LOC107973052-GGT2.revcompl                                       cacttaagaccagattagaaaagagggactccacctcacattctagagcaccatccccac 55445
chimp54546-135457.revcompl.Pan.troglodytes.clone.rp43-41g5.GenBank:AC099533.36     ------------------------------------------------------------ 64526
human.FAM30B-LOC105372935-GGT2.NCBI.                                                cacttaagaccagattagaaaagagagactccacctcacattctagagcgccatccccac 54438
human.FAM230E-LOC105377182-GGT3P.NCBI.                                              cacttaagaccagattagaaaagagagactccacctcacattctagagcgccatccccac 53092
human.FAM230J-LOC105372942-GGTLC5P.NCBI.rev.compl                                   ------------------------------------------------------------ 46364
human.FAM230A.ncRNA.18487127.18500594-GGTLC3.18516335.18518165.rev.compl.REF        ------------------------------------------------------------ 31039

chimp.LOC112206744-LOC107973052-GGT2.revcompl                                       aaatgaacaaatgagtgaatgggatgcctgttgaaaaagcaggatatagacagcctgggt 55505
chimp54546-135457.revcompl.Pan.troglodytes.clone.rp43-41g5.GenBank:AC099533.36     ------------------------------------------------------------ 64526
human.FAM30B-LOC105372935-GGT2.NCBI.                                                aaatgaacaaatgagtgaatgggatgcctgttgaaaaggcaggatatagacagcctgggt 54498
human.FAM230E-LOC105377182-GGT3P.NCBI.                                              aaatgaacaaatgagtgaatgggatgcctgttgaaaaggcaggatatagacagcctgggt 53152
human.FAM230J-LOC105372942-GGTLC5P.NCBI.rev.compl                                   ------------------------------------------------------------ 46364
human.FAM230A.ncRNA.18487127.18500594-GGTLC3.18516335.18518165.rev.compl.REF        ------------------------------------------------------------ 31039

chimp.LOC112206744-LOC107973052-GGT2.revcompl                                       tcaattttagctttaccacctcccagctgtgtgacctcaactgatttgcatgacctgtct 55565
chimp54546-135457.revcompl.Pan.troglodytes.clone.rp43-41g5.GenBank:AC099533.36     ------------------------------------------------------------ 64526
human.FAM30B-LOC105372935-GGT2.NCBI.                                                tcaattttagcttcaccacctcccagctgtgtgacctcagctgatttgcatgacctgtct 54558
human.FAM230E-LOC105377182-GGT3P.NCBI.                                              tcaattttagcttcaccacctcccagctgtgtgacctcagctgatttgcatgacctgtct 53212
human.FAM230J-LOC105372942-GGTLC5P.NCBI.rev.compl                                   ------------------------------------------------------------ 46364
human.FAM230A.ncRNA.18487127.18500594-GGTLC3.18516335.18518165.rev.compl.REF        ------------------------------------------------------------ 31039

chimp.LOC112206744-LOC107973052-GGT2.revcompl                                       gagcctcagcatccccaccctgtaaaatgggaatccacacagcatcccctagcccaaagg 55625
chimp54546-135457.revcompl.Pan.troglodytes.clone.rp43-41g5.GenBank:AC099533.36     ------------------------------------------------------------ 64526
human.FAM30B-LOC105372935-GGT2.NCBI.                                                gagcctcagcatccccaccctgtaaaatgggaatccacacagcatcccctagcccaaagg 54618
human.FAM230E-LOC105377182-GGT3P.NCBI.                                              gagcctcagcatccccaccctgtaaaatgggaatccacacagcatccccttgcccaaagg 53272
human.FAM230J-LOC105372942-GGTLC5P.NCBI.rev.compl                                   ------------------------------------------------------------ 46364
human.FAM230A.ncRNA.18487127.18500594-GGTLC3.18516335.18518165.rev.compl.REF        ------------------------------------------------------------ 31039

chimp.LOC112206744-LOC107973052-GGT2.revcompl                                       agcagggagggttgtgagaggctcgtgggtgaaaagcacagagcagagcgtgggccccag 55685
chimp54546-135457.revcompl.Pan.troglodytes.clone.rp43-41g5.GenBank:AC099533.36     ------------------------------------------------------------ 64526
human.FAM30B-LOC105372935-GGT2.NCBI.                                                agcagggagggttgtgagaggctcgtgggtgaaaagcacagagcagagcatgggccccag 54678
human.FAM230E-LOC105377182-GGT3P.NCBI.                                              agcagggagggttgtgagaggctcgtgggtgaaaagcacagagcagagcatgggccccag 53332
human.FAM230J-LOC105372942-GGTLC5P.NCBI.rev.compl                                   ------------------------------------------------------------ 46364
human.FAM230A.ncRNA.18487127.18500594-GGTLC3.18516335.18518165.rev.compl.REF        ------------------------------------------------------------ 31039

chimp.LOC112206744-LOC107973052-GGT2.revcompl                                       tgagccctggtccatgaggtctgctagcataataattattctttccatgtgctgcacaga 55745
chimp54546-135457.revcompl.Pan.troglodytes.clone.rp43-41g5.GenBank:AC099533.36     ------------------------------------------------------------ 64526
human.FAM30B-LOC105372935-GGT2.NCBI.                                                tgagccctgatccatgaggtctgctagcataataattattctttccatgtgctgcacaga 54738
human.FAM230E-LOC105377182-GGT3P.NCBI.                                              tgagccctgatccatgaggtctgctagcataataattattctttccatgtgctgcacaga 53392
human.FAM230J-LOC105372942-GGTLC5P.NCBI.rev.compl                                   ------------------------------------------------------------ 46364
human.FAM230A.ncRNA.18487127.18500594-GGTLC3.18516335.18518165.rev.compl.REF        ------------------------------------------------------------ 31039

chimp.LOC112206744-LOC107973052-GGT2.revcompl                                       gtggccccggaggccttagcagaaataacagaagctcccggccctttaccgtggtgatga 55805
chimp54546-135457.revcompl.Pan.troglodytes.clone.rp43-41g5.GenBank:AC099533.36     ------------------------------------------------------------ 64526
human.FAM30B-LOC105372935-GGT2.NCBI.                                                gtggccccggaggccttagcagaaataacagaagctcccggccctttaccgtggtgatga 54798
human.FAM230E-LOC105377182-GGT3P.NCBI.                                              gtggccccggaggccttagcagaaataacagaagctcccggccctttaccatggtgatga 53452
human.FAM230J-LOC105372942-GGTLC5P.NCBI.rev.compl                                   ------------------------------------------------------------ 46364
human.FAM230A.ncRNA.18487127.18500594-GGTLC3.18516335.18518165.rev.compl.REF        ------------------------------------------------------------ 31039

chimp.LOC112206744-LOC107973052-GGT2.revcompl                                       tggtcctgaccactcattgtgggagggtgctatggggcaaggaagggatgggggtgcta 55865
chimp54546-135457.revcompl.Pan.troglodytes.clone.rp43-41g5.GenBank:AC099533.36     ------------------------------------------------------------ 64526
human.FAM30B-LOC105372935-GGT2.NCBI.                                                tggtcctgaccactcattgtgggagggtgctatggggccaggaagggatgggggtgaca 54858
human.FAM230E-LOC105377182-GGT3P.NCBI.                                              tggtcctgaccactcactgtgggagggtgctatggggacaggaagggatgggggtgcta 53512
human.FAM230J-LOC105372942-GGTLC5P.NCBI.rev.compl                                   ------------------------------------------------------------ 46364
human.FAM230A.ncRNA.18487127.18500594-GGTLC3.18516335.18518165.rev.compl.REF        ------------------------------------------------------------ 31039
```

```
chimp.LOC112206744-LOC107973052-GGT2.revcompl                                       gaactgcccctgaaccctgacgggagcaggctcctgtgggcaaggccccttcccggtggc 55925
chimp54546-135457.revcompl.Pan.troglodytes.clone.rp43-41g5.GenBank:AC099533.36      ----------TGAAAACTACTCACATCTAACAAGAGATTAGACATCCAGATAGAGGAGGC 64576
human.FAM30B-LOC105372935-GGT2.NCBI.                                                gaactgcccctgaaccctgacgggagcaggcttctgtgggcaaggccccttcccggtggc 54918
human.FAM230E-LOC105377182-GGT3P.NCBI.                                              gaactgcccctgaaccctgacgggagcaggctcctgtgggcaaggccccttcccggtggc 53572
human.FAM230J-LOC105372942-GGTLC5P.NCBI.rev.compl                                   ------------------------------------------------------------ 46364
human.FAM230A.ncRNA.18487127.18500594-GGTLC3.18516335.18518165.rev.compl.REF        ------------------------------------------------------------ 31039

chimp.LOC112206744-LOC107973052-GGT2.revcompl                                       tcagccagctctgcac----ccatgccccaagtctgcagcatggcttacccttctgggac 55981
chimp54546-135457.revcompl.Pan.troglodytes.clone.rp43-41g5.GenBank:AC099533.36      TCAAATATCCCCAAACAATGCAATGCAAAGGGTCTTTGCCGTGGCAAACTATAATCAGAC 64636
human.FAM30B-LOC105372935-GGT2.NCBI.                                                tcagccagctctgcac----ccatgccccaagtctgcagcatggcttacccttctgggac 54974
human.FAM230E-LOC105377182-GGT3P.NCBI.                                              tcagccagctctgcac----ccacgccccaagtctgccgcatggcttacccttctgggac 53628
human.FAM230J-LOC105372942-GGTLC5P.NCBI.rev.compl                                   ------------------------------------------------------------ 46364
human.FAM230A.ncRNA.18487127.18500594-GGTLC3.18516335.18518165.rev.compl.REF        ------------------------------------------------------------ 31039

chimp.LOC112206744-LOC107973052-GGT2.revcompl                                       tgctccgagctgttg----aacatgctggcaaaggccagcctgggggccacctcgcgggc 56037
chimp54546-135457.revcompl.Pan.troglodytes.clone.rp43-41g5.GenBank:AC099533.36      TATTTAAGGTCAATGTGAAAATGCGAATTCTAAAACCACCAAGAGAAAAGAATCTAGTCA 64696
human.FAM30B-LOC105372935-GGT2.NCBI.                                                tgctccgagctgttg----aacatgctggcaaaggccagcctgggggccacctcgcgggc 55030
human.FAM230E-LOC105377182-GGT3P.NCBI.                                              tgctccgagctgttg----aacatgctggcaaaggccagcctgggggccacctcgcgggc 53684
human.FAM230J-LOC105372942-GGTLC5P.NCBI.rev.compl                                   ------------------------------------------------------------ 46364
human.FAM230A.ncRNA.18487127.18500594-GGTLC3.18516335.18518165.rev.compl.REF        ------------------------------------------------------------ 31039

chimp.LOC112206744-LOC107973052-GGT2.revcompl                                       gttgatgacctcagcttttcctagaaggagaagcaggtaggcaggcccacccacccaaac 56097
chimp54546-135457.revcompl.Pan.troglodytes.clone.rp43-41g5.GenBank:AC099533.36      CCTATAAAGCAAACCCCATCAGATG---AAGAGCAGATTTCACAGCCCAGCAAATAGAAA 64753
human.FAM30B-LOC105372935-GGT2.NCBI.                                                attgatgacctcagcttttcctaga---agaagcaggtaggcaggcccacccaccaaaac 55087
human.FAM230E-LOC105377182-GGT3P.NCBI.                                              attgatgacctcagcttttcctaga---agaagcaggtaggcaggcccacccaccaaaac 53741
human.FAM230J-LOC105372942-GGTLC5P.NCBI.rev.compl                                   ------------------------------------------------------------ 46364
human.FAM230A.ncRNA.18487127.18500594-GGTLC3.18516335.18518165.rev.compl.REF        ------------------------------------------------------------ 31039

chimp.LOC112206744-LOC107973052-GGT2.revcompl                                       ------cctttatgccacgtgagcctgggggccacccagctgtgcctcggcccaacccac 56151
chimp54546-135457.revcompl.Pan.troglodytes.clone.rp43-41g5.GenBank:AC099533.36      ACATGTGCTTAAAGGCCTAGGAGGCAGAGAGGTACAGGGCTGGAGACAGGCAAGGCCAGC 64813
human.FAM30B-LOC105372935-GGT2.NCBI.                                                ------cctttatgccacgtgagcctgggggccacccagctgtgcctcggcccaacccac 55141
human.FAM230E-LOC105377182-GGT3P.NCBI.                                              ------cctttatgccacgtgagcctgggggccacccagctgtgcctcggcccaacccac 53795
human.FAM230J-LOC105372942-GGTLC5P.NCBI.rev.compl                                   ------------------------------------------------------------ 46364
human.FAM230A.ncRNA.18487127.18500594-GGTLC3.18516335.18518165.rev.compl.REF        ------------------------------------------------------------ 31039

chimp.LOC112206744-LOC107973052-GGT2.revcompl                                       acccccctgcccctctccctctcctcttccgaggcactcatgagtggtgctgttgtagatg 56211
chimp54546-135457.revcompl.Pan.troglodytes.clone.rp43-41g5.GenBank:AC099533.36      ATTCCCCTCAAGGCAACGGCCAATCCTGGAAGAGGAGGCTGAGAGACAGAGCAGT----- 64868
human.FAM30B-LOC105372935-GGT2.NCBI.                                                acccccctgcccctctccctctcctcttctgaggcactcatgagtggtgctgttgtagatg 55201
human.FAM230E-LOC105377182-GGT3P.NCBI.                                              acccccctgcccctctccctctccgcttccgaggcactcatgagtggtgctgttgtagatg 53855
human.FAM230J-LOC105372942-GGTLC5P.NCBI.rev.compl                                   ------------------------------------------------------------ 46364
human.FAM230A.ncRNA.18487127.18500594-GGTLC3.18516335.18518165.rev.compl.REF        ------------------------------------------------------------ 31039

chimp.LOC112206744-LOC107973052-GGT2.revcompl                                       gtgaggaagaggccacccccgatgcccatgctgtgggcattcatgagccccacacacaac 56271
chimp54546-135457.revcompl.Pan.troglodytes.clone.rp43-41g5.GenBank:AC099533.36      ------------------------------------------------------------ 64868
human.FAM30B-LOC105372935-GGT2.NCBI.                                                gtgaggaagaggccaaccccgatgcccatgctgtgggcattcatgagccccacacacaac 55261
human.FAM230E-LOC105377182-GGT3P.NCBI.                                              gtgaggaagaggccaaccccgatgcccatgctgtgggcattcatgagccccacacacaac 53915
human.FAM230J-LOC105372942-GGTLC5P.NCBI.rev.compl                                   ------------------------------------------------------------ 46364
human.FAM230A.ncRNA.18487127.18500594-GGTLC3.18516335.18518165.rev.compl.REF        ------------------------------------------------------------ 31039

chimp.LOC112206744-LOC107973052-GGT2.revcompl                                       agggctgcaatggctgcatccaccgcagagccaccgtcccgcagtgcgtccctgccatgt 56331
chimp54546-135457.revcompl.Pan.troglodytes.clone.rp43-41g5.GenBank:AC099533.36      ------------------------------------------------------------ 64868
human.FAM30B-LOC105372935-GGT2.NCBI.                                                agggctgcaatggctgcatccactgcagagccaccgtcccgcagtgtgtccctgccatgt 55321
human.FAM230E-LOC105377182-GGT3P.NCBI.                                              agggctgcaatggctgcatccactgcagagccaccgtcccgcagtgtgtccctgccatgt 53975
human.FAM230J-LOC105372942-GGTLC5P.NCBI.rev.compl                                   ------------------------------------------------------------ 46364
human.FAM230A.ncRNA.18487127.18500594-GGTLC3.18516335.18518165.rev.compl.REF        ------------------------------------------------------------ 31039

chimp.LOC112206744-LOC107973052-GGT2.revcompl                                       ggcacataaaggcatgagaacctgcaggcttccaccctggccccgcatacacaccctgct 56391
chimp54546-135457.revcompl.Pan.troglodytes.clone.rp43-41g5.GenBank:AC099533.36      ------------------------------------------------------------ 64868
human.FAM30B-LOC105372935-GGT2.NCBI.                                                ggcacataaaggcatgagaacctgcaggcttccaccctggccccgcatacacaccctgct 55381
human.FAM230E-LOC105377182-GGT3P.NCBI.                                              ggcacataaaggcatgagaacctgcaggcttccaccctggccccgcatacacaccctgct 54035
human.FAM230J-LOC105372942-GGTLC5P.NCBI.rev.compl                                   ------------------------------------------------------------ 46364
human.FAM230A.ncRNA.18487127.18500594-GGTLC3.18516335.18518165.rev.compl.REF        ------------------------------------------------------------ 31039

chimp.LOC112206744-LOC107973052-GGT2.revcompl                                       gcccacctgcccaaaggaggatggaagagaagtccattcgagttttggggtttttgtttt 56451
chimp54546-135457.revcompl.Pan.troglodytes.clone.rp43-41g5.GenBank:AC099533.36      ------------------------------------------------------------ 64868
human.FAM30B-LOC105372935-GGT2.NCBI.                                                gcccacctgcccaaaggaggatggaagagaagtccattcgagttttggggtttttgtttt 55441
human.FAM230E-LOC105377182-GGT3P.NCBI.                                              gcccacctgcccaaaggaggatggaagagaagtccattcgagttttggggtttttgtttt 54095
human.FAM230J-LOC105372942-GGTLC5P.NCBI.rev.compl                                   ------------------------------------------------------------ 46364
human.FAM230A.ncRNA.18487127.18500594-GGTLC3.18516335.18518165.rev.compl.REF        ------------------------------------------------------------ 31039

chimp.LOC112206744-LOC107973052-GGT2.revcompl                                       tagtttctttctcttttttttgagatggagtcttgctctatt-gccaggctggagtgcag 56510
chimp54546-135457.revcompl.Pan.troglodytes.clone.rp43-41g5.GenBank:AC099533.36      --GATTCTTTTTTTTTTTTTTGAGACGGAGTCTCACTCTGTCACCCAGGCTGGAGTGCAG 64926
human.FAM30B-LOC105372935-GGT2.NCBI.                                                tagtttctttctcttttttttgagatggagtcttgctctatt-gccaggctggagtgcag 55500
human.FAM230E-LOC105377182-GGT3P.NCBI.                                              tagtttctttctcttttttttgagatggagtcttgctctatt-gccaggctggagtgcag 54154
human.FAM230J-LOC105372942-GGTLC5P.NCBI.rev.compl                                   ------------------------------------------------------------ 46364
human.FAM230A.ncRNA.18487127.18500594-GGTLC3.18516335.18518165.rev.compl.REF        ------------------------------------------------------------ 31039

chimp.LOC112206744-LOC107973052-GGT2.revcompl                                       tggcacgatctcagctcactgcaacctctgcctcccgggttcaagccattctcctgcctc 56570
chimp54546-135457.revcompl.Pan.troglodytes.clone.rp43-41g5.GenBank:AC099533.36      TGGCATGATCTCGGCTCACTGCAAGCTCCACCTCCTGGGTTCATGCCATTCTCCTGCCTC 64986
human.FAM30B-LOC105372935-GGT2.NCBI.                                                tggcacgatctcagctcactgcaacctctgcctcccgggttcaagccattctcctgcctc 55560
human.FAM230E-LOC105377182-GGT3P.NCBI.                                              tggcacgatctcagctcactgcaacctctgcctcccgggttcaagccattctcctgcctc 54214
human.FAM230J-LOC105372942-GGTLC5P.NCBI.rev.compl                                   ------------------------------------------------------------ 46364
human.FAM230A.ncRNA.18487127.18500594-GGTLC3.18516335.18518165.rev.compl.REF        ------------------------------------------------------------ 31039

chimp.LOC112206744-LOC107973052-GGT2.revcompl                                       agcctcctgagtagctgggactacaggcgcacaccaccacgcccggctaa-tttttgtat 56629
chimp54546-135457.revcompl.Pan.troglodytes.clone.rp43-41g5.GenBank:AC099533.36      AGCCTCCTGAGTAGCTGGGACTACAGGCGCCTGCCACCACGCCCAGCTAATTTTTTGTAT 65046
human.FAM30B-LOC105372935-GGT2.NCBI.                                                agcctcctgagtagctgggactacaggtgcatgccaccacgcccagctaa-tttttgtat 55619
human.FAM230E-LOC105377182-GGT3P.NCBI.                                              agcctcctgagtagctgggactacaggtgcatgccaccacgcccagctaa-tttttgtat 54273
human.FAM230J-LOC105372942-GGTLC5P.NCBI.rev.compl                                   ------------------------------------------------------------ 46364
human.FAM230A.ncRNA.18487127.18500594-GGTLC3.18516335.18518165.rev.compl.REF        ------------------------------------------------------------ 31039

chimp.LOC112206744-LOC107973052-GGT2.revcompl                                       ttttagtagagacggggtttcaccatgttggccaggatggtctctatctcttgacttcat 56689
chimp54546-135457.revcompl.Pan.troglodytes.clone.rp43-41g5.GenBank:AC099533.36      TTTTAGTAGAAATGGGGTTTCACCGTGTTAGCCAGGATGGTCTCGATCTCCTGATCTCAT 65106
human.FAM30B-LOC105372935-GGT2.NCBI.                                                ttttagtagagatggggtttcaccatgttggccaggatggtctctatctcttgacttcat 55679
human.FAM230E-LOC105377182-GGT3P.NCBI.                                              ttttagtagagacggggtttcaccatgttggccaggatggtctctatctcttgacttcat 54333
human.FAM230J-LOC105372942-GGTLC5P.NCBI.rev.compl                                   ------------------------------------------------------------ 46364
human.FAM230A.ncRNA.18487127.18500594-GGTLC3.18516335.18518165.rev.compl.REF        ------------------------------------------------------------ 31039

chimp.LOC112206744-LOC107973052-GGT2.revcompl                                       gatccacctgccttggcctc--------tttttttttttttttggagacaaagtctctct 56741
chimp54546-135457.revcompl.Pan.troglodytes.clone.rp43-41g5.GenBank:AC099533.36      GATCTGCCCGCCTTG--------------------------------------------- 65121
human.FAM30B-LOC105372935-GGT2.NCBI.                                                gatccgcctgccttggcctcttttttttttttttttttttttggagacaaagtctctct 55739
human.FAM230E-LOC105377182-GGT3P.NCBI.                                              gatccacctgccttggcctcttttttttttttttttttttttggagacaaagtctctct 54393
human.FAM230J-LOC105372942-GGTLC5P.NCBI.rev.compl                                   ------------------------------------------------------------ 46364
human.FAM230A.ncRNA.18487127.18500594-GGTLC3.18516335.18518165.rev.compl.REF        ------------------------------------------------------------ 31039

chimp.LOC112206744-LOC107973052-GGT2.revcompl                                       ttgttgctcaggctggagtgcagtggtgtgatctcggatcactgcagcctcagtctcctg 56801
chimp54546-135457.revcompl.Pan.troglodytes.clone.rp43-41g5.GenBank:AC099533.36      ------------------------------------------------------------ 65121
human.FAM30B-LOC105372935-GGT2.NCBI.                                                ttgttgctcaggctggagtgcagtggtgtgatcttggatcactgcagcctcagtctcctg 55799
human.FAM230E-LOC105377182-GGT3P.NCBI.                                              ttgttgctcaggctggagtgcagtggtgtgatcttggatcactgcagcctcagtctcctg 54453
human.FAM230J-LOC105372942-GGTLC5P.NCBI.rev.compl                                   ------------------------------------------------------------ 46364
human.FAM230A.ncRNA.18487127.18500594-GGTLC3.18516335.18518165.rev.compl.REF        ------------------------------------------------------------ 31039

chimp.LOC112206744-LOC107973052-GGT2.revcompl                                       ggctcaattgaccctcccatctcagcctcccaagtagctgggactatgggcacatgccac 56861
chimp54546-135457.revcompl.Pan.troglodytes.clone.rp43-41g5.GenBank:AC099533.36      ------------------------GCCTCCCAAAGTGCTGGGATTAGAGGCGTGAGCCAC 65157
human.FAM30B-LOC105372935-GGT2.NCBI.                                                ggctcaattgaccctcccatctcagcctcccaagtagctgggactatgggcacatgccac 55859
human.FAM230E-LOC105377182-GGT3P.NCBI.                                              ggctcaattgaccctcccatctcagcctcccaagtagctgggactatgggcacatgccac 54513
human.FAM230J-LOC105372942-GGTLC5P.NCBI.rev.compl                                   ------------------------------------------------------------ 46364
human.FAM230A.ncRNA.18487127.18500594-GGTLC3.18516335.18518165.rev.compl.REF        ------------------------------------------------------------ 31039

chimp.LOC112206744-LOC107973052-GGT2.revcompl                                       catgcccagccaatt---------------------ttgtttgtgtgttttgtagagatg 56900
chimp54546-135457.revcompl.Pan.troglodytes.clone.rp43-41g5.GenBank:AC099533.36      CAAGCCCAGCTGATTCTTTTTTTTAATTCTTCCTTCCTTTTTTTTTTTTTTTTTTTTTTT 65217
human.FAM30B-LOC105372935-GGT2.NCBI.                                                catgcccagccaatt-----------------ttgtttgtttgtgtattttgtagagatg 55902
human.FAM230E-LOC105377182-GGT3P.NCBI.                                              catgcccagccaatt-----------------ttgtttgtttgtgtattttgtagagatg 54556
human.FAM230J-LOC105372942-GGTLC5P.NCBI.rev.compl                                   ------------------------------------------------------------ 46364
human.FAM230A.ncRNA.18487127.18500594-GGTLC3.18516335.18518165.rev.compl.REF        ------------------------------------------------------------ 31039
```

```
chimp.LOC112206744-LOC107973052-GGT2.revcompl                              gggtttcatcatgttgcccaggctggtcaagaactcctgtgctcaagtgatccacccacc 56960
chimp54546-135457.revcompl.Pan.troglodytes.clone.rp43-41g5.GenBank:AC099533.36   GCTTTTTATTTTGCTGAAACAGGGTTGCCATTACTTTTAAAGGCAAAAACCGCAATTACT 65277
human.FAM30B-LOC105372935-GGT2.NCBI.                                       gggtttcatcatgttgcccaggctggtcaagaactcctgtgctcaagtaatccacccacc 55962
human.FAM230E-LOC105377182-GGT3P.NCBI.                                     gggtttcatcatgttgcccaggctggtcaagaactcctgtgctcaagtaatccacccacc 54616
human.FAM230J-LOC105372942-GGTLC5P.NCBI.rev.compl                          ------------------------------------------------------------ 46364
human.FAM230A.ncRNA.18487127.18500594-GGTLC3.18516335.18518165.rev.compl.REF   ------------------------------------------------------------ 31039

chimp.LOC112206744-LOC107973052-GGT2.revcompl                              ttggcctcccaaagtgctggtattacaggcatgagccactgtgctcagcctttgttttat 57020
chimp54546-135457.revcompl.Pan.troglodytes.clone.rp43-41g5.GenBank:AC099533.36   TTTG--TACCAACCTAATAATAT------------------------------------- 65298
human.FAM30B-LOC105372935-GGT2.NCBI.                                       ttggcctcccaaagtgctggtattacaggcatgagccactgtgcccagcctttgttttat 56022
human.FAM230E-LOC105377182-GGT3P.NCBI.                                     ttggcctcccaaagtgctggtattacaggcatgagccactgtgcccagcctttgttttat 54676
human.FAM230J-LOC105372942-GGTLC5P.NCBI.rev.compl                          ------------------------------------------------------------ 46364
human.FAM230A.ncRNA.18487127.18500594-GGTLC3.18516335.18518165.rev.compl.REF   ------------------------------------------------------------ 31039

chimp.LOC112206744-LOC107973052-GGT2.revcompl                              gagacagggtctcactttgtcacccaggatgaagtgcagtggcacagtcttggctcaatg 57080
chimp54546-135457.revcompl.Pan.troglodytes.clone.rp43-41g5.GenBank:AC099533.36   -------------------------------------------------ATTGCTCTGGC 65309
human.FAM30B-LOC105372935-GGT2.NCBI.                                       gagacagggtctcactttgtcacccaggatgaagtgcagtggcacagtcttggctcaatg 56082
human.FAM230E-LOC105377182-GGT3P.NCBI.                                     gagacagggtctcactttgtcacccaggatgaagtgcagtggcacagtcttggctcaatg 54736
human.FAM230J-LOC105372942-GGTLC5P.NCBI.rev.compl                          ------------------------------------------------------------ 46364
human.FAM230A.ncRNA.18487127.18500594-GGTLC3.18516335.18518165.rev.compl.REF   ------------------------------------------------------------ 31039

chimp.LOC112206744-LOC107973052-GGT2.revcompl                              cagctttgacctcctgggctcaagcaatcctcccacttcagtctcctgagtagctgggac 57140
chimp54546-135457.revcompl.Pan.troglodytes.clone.rp43-41g5.GenBank:AC099533.36   TGGTTTCAAATTCCTGGCCTGAAGCAATCCTCCTGCCTCAGTCTCCCGAGTAGCTGGGAC 65369
human.FAM30B-LOC105372935-GGT2.NCBI.                                       cagctttgacctcctgggctcaagcaatcctcccacttcagtctcctgagtagctgggac 56142
human.FAM230E-LOC105377182-GGT3P.NCBI.                                     cagctttgacctcctgggctcaagcaatcctcccacttcagtctcctgagtagctgggac 54796
human.FAM230J-LOC105372942-GGTLC5P.NCBI.rev.compl                          ------------------------------------------------------------ 46364
human.FAM230A.ncRNA.18487127.18500594-GGTLC3.18516335.18518165.rev.compl.REF   ------------------------------------------------------------ 31039

chimp.LOC112206744-LOC107973052-GGT2.revcompl                              tacaggtaagaaccaccacaccgggcaattttttgtcttttttgtagagatagggtcttt 57200
chimp54546-135457.revcompl.Pan.troglodytes.clone.rp43-41g5.GenBank:AC099533.36   TATAGGCGTGAGCCACCACACCTGGTCTAGAGCAGT------------------------ 65405
human.FAM30B-LOC105372935-GGT2.NCBI.                                       tacaggtaagaaccaccacaccgggcaattttttgtcttttttgtagagatagggtcttt 56202
human.FAM230E-LOC105377182-GGT3P.NCBI.                                     tacaggtaagaaccaccacaccgggcaattttttgtcttttttcgtagagatagggtcttt 54856
human.FAM230J-LOC105372942-GGTLC5P.NCBI.rev.compl                          ------------------------------------------------------------ 46364
human.FAM230A.ncRNA.18487127.18500594-GGTLC3.18516335.18518165.rev.compl.REF   ------------------------------------------------------------ 31039

chimp.LOC112206744-LOC107973052-GGT2.revcompl                              ctatgttgcccaggctggtctcaaactcatggtctaaagcaatcttatctcctcaacctc 57260
chimp54546-135457.revcompl.Pan.troglodytes.clone.rp43-41g5.GenBank:AC099533.36   -------------GCTGTTGACTTACTCATGTGGCTGAGGGAGGGGATGCCATAATGCCC 65452
human.FAM30B-LOC105372935-GGT2.NCBI.                                       ctatgttgcccaggctggtctcaaactcatggtctaaagcaatcctatcgcctcaacctc 56262
human.FAM230E-LOC105377182-GGT3P.NCBI.                                     ctatgttgcccaggctggtctcaaactcatggtctaaagcaatcctatcgcctcaacctc 54916
human.FAM230J-LOC105372942-GGTLC5P.NCBI.rev.compl                          ------------------------------------------------------------ 46364
human.FAM230A.ncRNA.18487127.18500594-GGTLC3.18516335.18518165.rev.compl.REF   ------------------------------------------------------------ 31039

chimp.LOC112206744-LOC107973052-GGT2.revcompl                              ccaaagtgctgggattacagtttcttcttttt----------------------------- 57291
chimp54546-135457.revcompl.Pan.troglodytes.clone.rp43-41g5.GenBank:AC099533.36   CACACG---TGGGGTGCAGGTTTCTTACTCTAGGAGTCACAGTCAACCAGAAATAAAGCA 65509
human.FAM30B-LOC105372935-GGT2.NCBI.                                       ccaaagtgctgggattacagtttcttcttttt----------------------------- 56293
human.FAM230E-LOC105377182-GGT3P.NCBI.                                     ccaaagtgctgggattacagtttcttcttttt----------------------------- 54947
human.FAM230J-LOC105372942-GGTLC5P.NCBI.rev.compl                          ------------------------------------------------------------ 46364
human.FAM230A.ncRNA.18487127.18500594-GGTLC3.18516335.18518165.rev.compl.REF   ------------------------------------------------------------ 31039

chimp.LOC112206744-LOC107973052-GGT2.revcompl                              ------------------------------------------------------------ 57291
chimp54546-135457.revcompl.Pan.troglodytes.clone.rp43-41g5.GenBank:AC099533.36   GCCCTCAGGAGCACTGGATCCAGCTTCAGATCATCTCAGTCACTCAAATGCATAAAGGAG 65569
human.FAM30B-LOC105372935-GGT2.NCBI.                                       ------------------------------------------------------------ 56293
human.FAM230E-LOC105377182-GGT3P.NCBI.                                     ------------------------------------------------------------ 54947
human.FAM230J-LOC105372942-GGTLC5P.NCBI.rev.compl                          ------------------------------------------------------------ 46364
human.FAM230A.ncRNA.18487127.18500594-GGTLC3.18516335.18518165.rev.compl.REF   ------------------------------------------------------------ 31039

chimp.LOC112206744-LOC107973052-GGT2.revcompl                              -----------------------------------------tcttttcttttttttttct 57310
chimp54546-135457.revcompl.Pan.troglodytes.clone.rp43-41g5.GenBank:AC099533.36   ATCTCAGAATTTCAGACCCCTAGGCCTCAAATTATTTTATTTTTATTTATTTATTTATTT 65629
human.FAM30B-LOC105372935-GGT2.NCBI.                                       -------------------------------tcttttcttttttctttttttttttttttct 56322
human.FAM230E-LOC105377182-GGT3P.NCBI.                                     -------------------------------tcttttcttttttctttttttttttttttct 54976
human.FAM230J-LOC105372942-GGTLC5P.NCBI.rev.compl                          ------------------------------------------------------------ 46364
human.FAM230A.ncRNA.18487127.18500594-GGTLC3.18516335.18518165.rev.compl.REF   ------------------------------------------------------------ 31039

chimp.LOC112206744-LOC107973052-GGT2.revcompl                              gagacagagtttcactcagttgcccaggctggagtgcagtggcatgatcttagctcactg 57370
chimp54546-135457.revcompl.Pan.troglodytes.clone.rp43-41g5.GenBank:AC099533.36   GAGACAGCATCTCACTCTGTTGCCCAGGCTGGAGTACAGTCTCAGGATCATGGCTCACTG 65689
human.FAM30B-LOC105372935-GGT2.NCBI.                                       gagacagagtttcactcagttgcccaggctggagtgcagtggcatgatcttagctcactg 56382
human.FAM230E-LOC105377182-GGT3P.NCBI.                                     gagacagagtttcactcagttgcccaggctggagtgcagtggcatgatcttagctcactg 55036
human.FAM230J-LOC105372942-GGTLC5P.NCBI.rev.compl                          ------------------------------------------------------------ 46364
human.FAM230A.ncRNA.18487127.18500594-GGTLC3.18516335.18518165.rev.compl.REF   ------------------------------------------------------------ 31039

chimp.LOC112206744-LOC107973052-GGT2.revcompl                              caacctctgcctcctgggttcaagcgattctcctgcctcagcctcctgagtagctgggat 57430
chimp54546-135457.revcompl.Pan.troglodytes.clone.rp43-41g5.GenBank:AC099533.36   CAGCCTCGATCACCCAGGCTCAAGTGAGCCTCCCACCTCAGCGTCCCAAACAGCTGGGAC 65749
human.FAM30B-LOC105372935-GGT2.NCBI.                                       caacctctgcctcctgggttcaagcgattctcctgcctcagcctcctgagtagctgggat 56442
human.FAM230E-LOC105377182-GGT3P.NCBI.                                     caacctctgcctcctgggttcaagcgattctcctgcctcagcctcctgagtagctgggat 55096
human.FAM230J-LOC105372942-GGTLC5P.NCBI.rev.compl                          ------------------------------------------------------------ 46364
human.FAM230A.ncRNA.18487127.18500594-GGTLC3.18516335.18518165.rev.compl.REF   ------------------------------------------------------------ 31039

chimp.LOC112206744-LOC107973052-GGT2.revcompl                              tacaggcgcacaccaccatg---cccggctaactttttatatttttagtagggacagag- 57486
chimp54546-135457.revcompl.Pan.troglodytes.clone.rp43-41g5.GenBank:AC099533.36   TACAGGCACACACCACCACCACACCCAGCTAATTTTT--TATTTTTTGTAGAGATGGGGT 65807
human.FAM30B-LOC105372935-GGT2.NCBI.                                       tacaggcgcacaccaccatg---cccggctaactttttatatttttagtagggacagag- 56498
human.FAM230E-LOC105377182-GGT3P.NCBI.                                     tacaggcgcacaccaccatg---cccggctaactttttatatttttagtagggacagag- 55152
human.FAM230J-LOC105372942-GGTLC5P.NCBI.rev.compl                          ------------------------------------------------------------ 46364
human.FAM230A.ncRNA.18487127.18500594-GGTLC3.18516335.18518165.rev.compl.REF   ------------------------------------------------------------ 31039

chimp.LOC112206744-LOC107973052-GGT2.revcompl                              tgcaccatgttggccaggctggtctcgaactcctgacctcgggtggtctgcccgcctcaa 57546
chimp54546-135457.revcompl.Pan.troglodytes.clone.rp43-41g5.GenBank:AC099533.36   TTCACCATGTTGCGCAGGCTGATCTTGAACTCCTAGGCTCAAGCAATCTGCCTGCCTCAG 65867
human.FAM30B-LOC105372935-GGT2.NCBI.                                       tgcaccatgttggccaggctgatctcgaactcctgacctc-agtggtctgcccgcctcaa 56557
human.FAM230E-LOC105377182-GGT3P.NCBI.                                     tgcaccatgttggccaggctggtctcgaactcctgacctc-agtggtctgcccgcctcaa 55211
human.FAM230J-LOC105372942-GGTLC5P.NCBI.rev.compl                          ------------------------------------------------------------ 46364
human.FAM230A.ncRNA.18487127.18500594-GGTLC3.18516335.18518165.rev.compl.REF   ------------------------------------------------------------ 31039

chimp.LOC112206744-LOC107973052-GGT2.revcompl                              ccccccaaagtgctgggattactggtgtggaccaccttgcccagacagtttcctctttat 57606
chimp54546-135457.revcompl.Pan.troglodytes.clone.rp43-41g5.GenBank:AC099533.36   CCTCCCAAAGTGATGGGATTACAAGCATGAGCCATCATGCTTGG---------------- 65911
human.FAM30B-LOC105372935-GGT2.NCBI.                                       ccccccaaagtgctgggattacaggtgtggaccaccttgcccagacagtttcctctttat 56617
human.FAM230E-LOC105377182-GGT3P.NCBI.                                     ccccccaaagtgctgggattacaggtgtggaccaccttgcccagacagtttcctctttat 55271
human.FAM230J-LOC105372942-GGTLC5P.NCBI.rev.compl                          ------------------------------------------------------------ 46364
human.FAM230A.ncRNA.18487127.18500594-GGTLC3.18516335.18518165.rev.compl.REF   ------------------------------------------------------------ 31039

chimp.LOC112206744-LOC107973052-GGT2.revcompl                              caagcaaacaaatgtacatgacttttataattgggacaaaaagggaaattgctatacttt 57666
chimp54546-135457.revcompl.Pan.troglodytes.clone.rp43-41g5.GenBank:AC099533.36   --------------------------------------------------GCTCGAAATT 65921
human.FAM30B-LOC105372935-GGT2.NCBI.                                       taagcaaacaaatgtacatgacttttataattgggacaaaaagggaaattgctatacttt 56677
human.FAM230E-LOC105377182-GGT3P.NCBI.                                     taagcaaacaaatgtacatgacttttataattgggacaaaaagggaaattgctatacttt 55331
human.FAM230J-LOC105372942-GGTLC5P.NCBI.rev.compl                          ------------------------------------------------------------ 46364
human.FAM230A.ncRNA.18487127.18500594-GGTLC3.18516335.18518165.rev.compl.REF   ------------------------------------------------------------ 31039

chimp.LOC112206744-LOC107973052-GGT2.revcompl                              attaataacattttttttttccttgctagagatggtggcttacacctgtaatctcagcac 57726
chimp54546-135457.revcompl.Pan.troglodytes.clone.rp43-41g5.GenBank:AC099533.36   ATTTTTAAAAATTATTTCT----GGCTGGGCGCAGTGGCTCACGCCTGTAATCCCAGCAC 65977
human.FAM30B-LOC105372935-GGT2.NCBI.                                       attaataacatttttttttccctgctagagatggtggcttacacctgtaatctcagcac 56737
human.FAM230E-LOC105377182-GGT3P.NCBI.                                     attaataacatttttttttccctgctagagatggtggcttacacctgtaatctcagcac 55391
human.FAM230J-LOC105372942-GGTLC5P.NCBI.rev.compl                          ------------------------------------------------------------ 46364
human.FAM230A.ncRNA.18487127.18500594-GGTLC3.18516335.18518165.rev.compl.REF   ------------------------------------------------------------ 31039

chimp.LOC112206744-LOC107973052-GGT2.revcompl                              tctgggaggccaaggtggaggatcacttgaggccaggagttcaagaccagcctgggcaat 57786
chimp54546-135457.revcompl.Pan.troglodytes.clone.rp43-41g5.GenBank:AC099533.36   TTTGGGAAGCTGAGGTGGA-CAGATCACGAGGTCAGGAGTCTGAGACCAGCCTGACCAAC 66036
human.FAM30B-LOC105372935-GGT2.NCBI.                                       tttgggaggccaaggtggaggatcacttgaggccaggagttcaagaccagcctgggcaat 56797
human.FAM230E-LOC105377182-GGT3P.NCBI.                                     tttgggaggccaaggtggaggatcacttgaggccaggagttcaagaccagcctgggcaat 55451
human.FAM230J-LOC105372942-GGTLC5P.NCBI.rev.compl                          ------------------------------------------------------------ 46364
human.FAM230A.ncRNA.18487127.18500594-GGTLC3.18516335.18518165.rev.compl.REF   ------------------------------------------------------------ 31039

chimp.LOC112206744-LOC107973052-GGT2.revcompl                              agagtgaga--ccatctctagaaaacagtcttttttaattagtcaggtgtgatgcacgcc 57844
chimp54546-135457.revcompl.Pan.troglodytes.clone.rp43-41g5.GenBank:AC099533.36   ATGGTGAAAACCCATCTCTACTAAAAATACAAAAATTAGCCAGGCGTGCTGGCACCTGCC 66096
human.FAM30B-LOC105372935-GGT2.NCBI.                                       agagtgaga--ccatctctacaaaacagtcttttttaattagtcaggtatgatgcacgcc 56855
human.FAM230E-LOC105377182-GGT3P.NCBI.                                     agagtgaga--ccatctctacaaaacagtcttttttaattagtcaggtatgatgcacgcc 55509
human.FAM230J-LOC105372942-GGTLC5P.NCBI.rev.compl                          ------------------------------------------------------------ 46364
human.FAM230A.ncRNA.18487127.18500594-GGTLC3.18516335.18518165.rev.compl.REF   ------------------------------------------------------------ 31039
```

```
chimp.LOC112206744-LOC107973052-GGT2.revcompl    tgtagtcctagctactcaggggggctgaggtgagaggatcgcttaagcccaagagttcaag 57904
chimp54546-135457.revcompl.Pan.troglodytes.clone.rp43-41g5.GenBank:AC099533.36    TGCAATCCCAGCTACTTGGGAGGCTGAGATGGGAGAAT---TGAACCCTGGGATGCAGAG 66153
human.FAM30B-LOC105372935-GGT2.NCBI.    tgtagtcctagctactcaggggggctgaggtgagaggatcgcttaagcccaagagttcaag 56915
human.FAM230E-LOC105377182-GGT3P.NCBI.    tgtagtcctagctactcaggggggctgaggtgagaggatcgcttaagcccaagagttcaag 55569
human.FAM230J-LOC105372942-GGTLC5P.NCBI.rev.compl    ------------------------------------------------------------ 46364
human.FAM230A.ncRNA.18487127.18500594-GGTLC3.18516335.18518165.rev.compl.REF    ------------------------------------------------------------ 31039

chimp.LOC112206744-LOC107973052-GGT2.revcompl    gctgcagtgagctatgatcatgccactgcactccagcctgggtgacagaacaagaccctg 57964
chimp54546-135457.revcompl.Pan.troglodytes.clone.rp43-41g5.GenBank:AC099533.36    GTTGCAGTGAGCCGAGATCACATCATTGCACTCCAGCTTGGGCAACAAAGCGAGACTCCA 66213
human.FAM30B-LOC105372935-GGT2.NCBI.    gctgcagtgagctatgatcatgccactgcactccagcctgggtgacagaacaagaccctg 56975
human.FAM230E-LOC105377182-GGT3P.NCBI.    gctgcagtgagctatgatcatgccactgcactccagcctgggtgacagaacaagaccctg 55629
human.FAM230J-LOC105372942-GGTLC5P.NCBI.rev.compl    ------------------------------------------------------------ 46364
human.FAM230A.ncRNA.18487127.18500594-GGTLC3.18516335.18518165.rev.compl.REF    ------------------------------------------------------------ 31039

chimp.LOC112206744-LOC107973052-GGT2.revcompl    tctcaaaatatgaaaaacataatattttttctgtttaagtctttaagagggaactgatct 58024
chimp54546-135457.revcompl.Pan.troglodytes.clone.rp43-41g5.GenBank:AC099533.36    TCTCAAAAAAAAAATTATTTGTAACAATTTTTATACACAACTTTCATTGCCAAATTAAAA 66273
human.FAM30B-LOC105372935-GGT2.NCBI.    tctcaaaatatgaaaaacataatattttttctgtttaagtctttaggagggaacttatct 57035
human.FAM230E-LOC105377182-GGT3P.NCBI.    tctcaaaatatgaaaaacataatattttttctgtttaagtctttaggagggaacttatct 55689
human.FAM230J-LOC105372942-GGTLC5P.NCBI.rev.compl    ------------------------------------------------------------ 46364
human.FAM230A.ncRNA.18487127.18500594-GGTLC3.18516335.18518165.rev.compl.REF    ------------------------------------------------------------ 31039

chimp.LOC112206744-LOC107973052-GGT2.revcompl    ttatatataacatgagataagagtctaaaatagaataaaacagtagaagccaggcaccat 58084
chimp54546-135457.revcompl.Pan.troglodytes.clone.rp43-41g5.GenBank:AC099533.36    ACAAGCAAACAAGGAAATAAGA--CAGCAGGAACAAAACTCAAGAGAAAGCACAC----- 66326
human.FAM30B-LOC105372935-GGT2.NCBI.    ttatatataacatgagataagagtctaaaatagaataacacagtagaagccaggcaccat 57095
human.FAM230E-LOC105377182-GGT3P.NCBI.    ttatatataacatgagataagagtctaaaatagaataacacagtagaagccaggcaccat 55749
human.FAM230J-LOC105372942-GGTLC5P.NCBI.rev.compl    ------------------------------------------------------------ 46364
human.FAM230A.ncRNA.18487127.18500594-GGTLC3.18516335.18518165.rev.compl.REF    ------------------------------------------------------------ 31039

chimp.LOC112206744-LOC107973052-GGT2.revcompl    ggctcacacctgtaatcccagcactttaggaagctgaggcgggaggatcacttaaaccca 58144
chimp54546-135457.revcompl.Pan.troglodytes.clone.rp43-41g5.GenBank:AC099533.36    ------------------------------------------------------------ 66326
human.FAM30B-LOC105372935-GGT2.NCBI.    ggctcacacctgtaatcccagcactttaggaagctgaggcgggaggatcacttaaaccca 57155
human.FAM230E-LOC105377182-GGT3P.NCBI.    ggctcacacctgtaatcccagcactttaggaagctgaggcgggaggatcacttaaaccca 55809
human.FAM230J-LOC105372942-GGTLC5P.NCBI.rev.compl    ------------------------------------------------------------ 46364
human.FAM230A.ncRNA.18487127.18500594-GGTLC3.18516335.18518165.rev.compl.REF    ------------------------------------------------------------ 31039

chimp.LOC112206744-LOC107973052-GGT2.revcompl    agagtttgaggctgcagtgggctatgatcgctcccctatactccagcctgggtgacagag 58204
chimp54546-135457.revcompl.Pan.troglodytes.clone.rp43-41g5.GenBank:AC099533.36    ------------------------------------------------------------ 66326
human.FAM30B-LOC105372935-GGT2.NCBI.    agagtttgaggctgcagtgggctatgatcgctcccctatactccagcctgggtgacagag 57215
human.FAM230E-LOC105377182-GGT3P.NCBI.    agagtttgaggctgcagtgggctatgatcgctcccctatactccagcctgggtgacagag 55869
human.FAM230J-LOC105372942-GGTLC5P.NCBI.rev.compl    ------------------------------------------------------------ 46364
human.FAM230A.ncRNA.18487127.18500594-GGTLC3.18516335.18518165.rev.compl.REF    ------------------------------------------------------------ 31039

chimp.LOC112206744-LOC107973052-GGT2.revcompl    taagactccaccttaaaaaaagaataagcccttcatgtccctgtttgggcaacagcgttc 58264
chimp54546-135457.revcompl.Pan.troglodytes.clone.rp43-41g5.GenBank:AC099533.36    ------------------------------------------------------------ 66326
human.FAM30B-LOC105372935-GGT2.NCBI.    taagactccaccttaaaaaaagaataagcccttcatgtccctgtttgggcaacagcgttc 57275
human.FAM230E-LOC105377182-GGT3P.NCBI.    taagactccaccttaaaaaaagaataagcccttcatgtccctgtttgggcaacagcgttc 55929
human.FAM230J-LOC105372942-GGTLC5P.NCBI.rev.compl    ------------------------------------------------------------ 46364
human.FAM230A.ncRNA.18487127.18500594-GGTLC3.18516335.18518165.rev.compl.REF    ------------------------------------------------------------ 31039

chimp.LOC112206744-LOC107973052-GGT2.revcompl    ctt-------ggggggatgggggagggatggcattgggagatatacataatgctaaatga 58317
chimp54546-135457.revcompl.Pan.troglodytes.clone.rp43-41g5.GenBank:AC099533.36    ------------------------------------------------------------ 66326
human.FAM30B-LOC105372935-GGT2.NCBI.    cttggggggaggggggaggggggagggatggcattgggagatatacctaatgctaaatga 57335
human.FAM230E-LOC105377182-GGT3P.NCBI.    cttggggggaggggggaggggggagggatggcattgggagatatacctaatgctaaatga 55989
human.FAM230J-LOC105372942-GGTLC5P.NCBI.rev.compl    ------------------------------------------------------------ 46364
human.FAM230A.ncRNA.18487127.18500594-GGTLC3.18516335.18518165.rev.compl.REF    ------------------------------------------------------------ 31039

chimp.LOC112206744-LOC107973052-GGT2.revcompl    tgagttggtgggtgcagcacaccaacatggcacatgtgtacatatgtaacaaacctgcac 58377
chimp54546-135457.revcompl.Pan.troglodytes.clone.rp43-41g5.GenBank:AC099533.36    ------------------------------------------------ACAACTTTGAAA 66338
human.FAM30B-LOC105372935-GGT2.NCBI.    cgagttggtgggtgcagcacaccaacatggcacatgtgtacatatgtaacaaacctgcac 57395
human.FAM230E-LOC105377182-GGT3P.NCBI.    cgagttggtgggtgcagcacaccaacatggcacatgtgtacatatgtaacaaacctgcac 56049
human.FAM230J-LOC105372942-GGTLC5P.NCBI.rev.compl    ------------------------------------------------------------ 46364
human.FAM230A.ncRNA.18487127.18500594-GGTLC3.18516335.18518165.rev.compl.REF    ------------------------------------------------------------ 31039

chimp.LOC112206744-LOC107973052-GGT2.revcompl    attgtgcacatgtaccctaaaacttaaagtataataataataaaataaaataaaaaaaag 58437
chimp54546-135457.revcompl.Pan.troglodytes.clone.rp43-41g5.GenBank:AC099533.36    AAGGCCTATAGGATCTCCAGAATATAAAAT---CATCAGACCATGTACTTTAAAATAAAT 66395
human.FAM30B-LOC105372935-GGT2.NCBI.    attgtgcacatgtaccctaaaacttaaagt---ataataataaaataaaataaaaaaaag 57452
human.FAM230E-LOC105377182-GGT3P.NCBI.    attgtgcacatgtaccctaaaacttaaagt---ataataataaaataaaataaaaaaaag 56106
human.FAM230J-LOC105372942-GGTLC5P.NCBI.rev.compl    ------------------------------------------------------------ 46364
human.FAM230A.ncRNA.18487127.18500594-GGTLC3.18516335.18518165.rev.compl.REF    ------------------------------------------------------------ 31039

chimp.LOC112206744-LOC107973052-GGT2.revcompl    aaagaaagaaattgtctcctaaccaaaaaaaaaaaaaaaaaaaagaaaagaaaag----- 58492
chimp54546-135457.revcompl.Pan.troglodytes.clone.rp43-41g5.GenBank:AC099533.36    TCTTTACATGTGTATCAAGGATATAAAGAAATAATTAAGAATTTCATCAGAAAACTGGCA 66455
human.FAM30B-LOC105372935-GGT2.NCBI.    aaagaaagaaattgtctcctaaccaaaaaaaaaaaaaaaaaaaagaaaagaaaag----- 57507
human.FAM230E-LOC105377182-GGT3P.NCBI.    aaagaaagaaattgtctcctaacc-------aaaaaaaaaaaaagaaaagaaaag----- 56154
human.FAM230J-LOC105372942-GGTLC5P.NCBI.rev.compl    ------------------------------------------------------------ 46364
human.FAM230A.ncRNA.18487127.18500594-GGTLC3.18516335.18518165.rev.compl.REF    ------------------------------------------------------------ 31039

chimp.LOC112206744-LOC107973052-GGT2.revcompl    -------------aataagccgggcgtggtggctcacacctgtaatcccaacacttcggg 58539
chimp54546-135457.revcompl.Pan.troglodytes.clone.rp43-41g5.GenBank:AC099533.36    GCTACAAAAAATGGACGCACTAGGTGTGGTGACTCACGCCTGTAATCCCAGCACTATGGG 66515
human.FAM30B-LOC105372935-GGT2.NCBI.    --------aaaagaataagccgggcgtggtggctcatacctgtaatcccaacactttggg 57559
human.FAM230E-LOC105377182-GGT3P.NCBI.    --------aaaagaataagccgggcgtggtggctcatacctgtaatcccaacactttggg 56206
human.FAM230J-LOC105372942-GGTLC5P.NCBI.rev.compl    ------------------------------------------------------------ 46364
human.FAM230A.ncRNA.18487127.18500594-GGTLC3.18516335.18518165.rev.compl.REF    ------------------------------------------------------------ 31039

chimp.LOC112206744-LOC107973052-GGT2.revcompl    aggcccaggtgggtggattacctgaggtcaggagttcgagaccagcctgaccaacatggt 58599
chimp54546-135457.revcompl.Pan.troglodytes.clone.rp43-41g5.GenBank:AC099533.36    AGGCTGAGGTGGGTGGATCACCTGAGGTCAGAAGTTCAAGACCAGCCTGACCAATATGGT 66575
human.FAM30B-LOC105372935-GGT2.NCBI.    aggcccaggtgggtggattacctgaggtcaggagttcgagaccagcctgaccaacatggt 57619
human.FAM230E-LOC105377182-GGT3P.NCBI.    aggcccaggtgggtggattacctgaggtcaggagttcgagaccagcctgaccaacatggt 56266
human.FAM230J-LOC105372942-GGTLC5P.NCBI.rev.compl    ------------------------------------------------------------ 46364
human.FAM230A.ncRNA.18487127.18500594-GGTLC3.18516335.18518165.rev.compl.REF    ------------------------------------------------------------ 31039

chimp.LOC112206744-LOC107973052-GGT2.revcompl    gaaaccccatctctactaaaaacaccataattaggccaggcgcagtggctcacacctgta 58659
chimp54546-135457.revcompl.Pan.troglodytes.clone.rp43-41g5.GenBank:AC099533.36    GAAACCCCA--------------------------------------------------- 66584
human.FAM30B-LOC105372935-GGT2.NCBI.    gaaaccccatctctactaaaaacaccataattaggccaggcgcagtggctcacacctgta 57679
human.FAM230E-LOC105377182-GGT3P.NCBI.    gaaaccccatctctactaaaaacaccataattaggccaggcgcagtggctcacacctgta 56326
human.FAM230J-LOC105372942-GGTLC5P.NCBI.rev.compl    ------------------------------------------------------------ 46364
human.FAM230A.ncRNA.18487127.18500594-GGTLC3.18516335.18518165.rev.compl.REF    ------------------------------------------------------------ 31039

chimp.LOC112206744-LOC107973052-GGT2.revcompl    atcccagcactttgagaggcagaggtgggaggatcacaaggtcaggagttcaagaccagc 58719
chimp54546-135457.revcompl.Pan.troglodytes.clone.rp43-41g5.GenBank:AC099533.36    ------------------------------------------------------------ 66584
human.FAM30B-LOC105372935-GGT2.NCBI.    atcccagcactttgagaggcagaggtgggaggatcacaaggtcaggagttcaagaccagc 57739
human.FAM230E-LOC105377182-GGT3P.NCBI.    atcccagcactttgagaggcagaggtgggaggatcacaaggtcaggagttcaagaccagc 56386
human.FAM230J-LOC105372942-GGTLC5P.NCBI.rev.compl    ------------------------------------------------------------ 46364
human.FAM230A.ncRNA.18487127.18500594-GGTLC3.18516335.18518165.rev.compl.REF    ------------------------------------------------------------ 31039

chimp.LOC112206744-LOC107973052-GGT2.revcompl    ctggccaacatagcaaaccctgtctctactaaaaatacaaaaattagctgggcatggtg 58779
chimp54546-135457.revcompl.Pan.troglodytes.clone.rp43-41g5.GenBank:AC099533.36    -----------------------TCTCTACTAAAATTACAAAAATTAGCCAGGCATGGTG 66621
human.FAM30B-LOC105372935-GGT2.NCBI.    ctggccaacatagcaaaccctgtctctactaaaaatacaaaaattagctgggcatggtg 57799
human.FAM230E-LOC105377182-GGT3P.NCBI.    ctggccaacatagcaaaccctgtctctactaaaaatacaaaaattagctgggcatggtg 56446
human.FAM230J-LOC105372942-GGTLC5P.NCBI.rev.compl    ------------------------------------------------------------ 46364
human.FAM230A.ncRNA.18487127.18500594-GGTLC3.18516335.18518165.rev.compl.REF    ------------------------------------------------------------ 31039

chimp.LOC112206744-LOC107973052-GGT2.revcompl    gcacatgcctgtagtctcagctactgtggaggctgaggcaggagaatcacttgaacctgg 58839
chimp54546-135457.revcompl.Pan.troglodytes.clone.rp43-41g5.GenBank:AC099533.36    GCATGCTCCTGTAGTCCCAGCTACTCGGGAGGCTGAGGCAGGAGAATCACTTGAA-CTGG 66680
human.FAM30B-LOC105372935-GGT2.NCBI.    gcacatgcctgtagtctcagctactgtggaggctgaggcaggagaatcacttgaacctgg 57859
human.FAM230E-LOC105377182-GGT3P.NCBI.    gcacatgcctgtagtctcagctactgtggaggctgaggcaggagaatcacttgaacctgg 56506
human.FAM230J-LOC105372942-GGTLC5P.NCBI.rev.compl    ------------------------------------------------------------ 46364
human.FAM230A.ncRNA.18487127.18500594-GGTLC3.18516335.18518165.rev.compl.REF    ------------------------------------------------------------ 31039

chimp.LOC112206744-LOC107973052-GGT2.revcompl    gaggtggaggctgcagtaagccaagattgcaccactgccctccagcctgggtaacagagc 58899
chimp54546-135457.revcompl.Pan.troglodytes.clone.rp43-41g5.GenBank:AC099533.36    GAAGGGGAGGTTGCAGTGAGCCGAGATCGCACCACTCTACTCCAGCCTGGGTGACAGAGC 66740
human.FAM30B-LOC105372935-GGT2.NCBI.    gaggtggaggctgcagtaagccaagattgcaccactgccctccagcctgggtaacagagc 57919
human.FAM230E-LOC105377182-GGT3P.NCBI.    gaggtggaggctgcagtaagccaagattgcaccactgccctccagcctgggtaacagagc 56566
human.FAM230J-LOC105372942-GGTLC5P.NCBI.rev.compl    ------------------------------------------------------------ 46364
human.FAM230A.ncRNA.18487127.18500594-GGTLC3.18516335.18518165.rev.compl.REF    ------------------------------------------------------------ 31039

chimp.LOC112206744-LOC107973052-GGT2.revcompl    gatactctgtctcagaaaacacacaca--------------cacacacacacacacatat 58945
```

```
chimp54546-135457.revcompl.Pan.troglodytes.clone.rp43-41g5.GenBank:AC099533.36      GAGACTCCGTCTCAAAAAAAAAAAAAAAAAAAAAAAAAATGGA----------------- 66783
human.FAM30B-LOC105372935-GGT2.NCBI.                                                 gatactctgtctcagaaaacacacacacacacacacacacacacacacacacacacatat 57979
human.FAM230E-LOC105377182-GGT3P.NCBI.                                               gatactctgtctcagaaa----------acacacacacacacacacacacacacacatat 56616
human.FAM230J-LOC105372942-GGTLC5P.NCBI.rev.compl                                    ------------------------------------------------------------ 46364
human.FAM230A.ncRNA.18487127.18500594-GGTLC3.18516335.18518165.rev.compl.REF         ------------------------------------------------------------ 31039

chimp.LOC112206744-LOC107973052-GGT2.revcompl                                        acacacacacaaatttgttgggcgtggtggcacactcctgtaatcccagctacttgggag 59005
chimp54546-135457.revcompl.Pan.troglodytes.clone.rp43-41g5.GenBank:AC099533.36      ------------------------------------------------------------ 66783
human.FAM30B-LOC105372935-GGT2.NCBI.                                                 acacacacacaaatttgttgggcgtggtggcgcactcctataatcccagctacttgggag 58039
human.FAM230E-LOC105377182-GGT3P.NCBI.                                               acacacacacaaatttgttgggcgtggtggcgcactcctataatcccagctacttgggag 56676
human.FAM230J-LOC105372942-GGTLC5P.NCBI.rev.compl                                    ------------------------------------------------------------ 46364
human.FAM230A.ncRNA.18487127.18500594-GGTLC3.18516335.18518165.rev.compl.REF         ------------------------------------------------------------ 31039

chimp.LOC112206744-LOC107973052-GGT2.revcompl                                        gcttaggcatgagaatcacttgaacttgggaggcggaggttgcagttaactgagattgca 59065
chimp54546-135457.revcompl.Pan.troglodytes.clone.rp43-41g5.GenBank:AC099533.36      ------------------------------------------------------------ 66783
human.FAM30B-LOC105372935-GGT2.NCBI.                                                 gcttaggcatgagaatcgcttgaacttgggaggcggaggttgcagttaactgagattgca 58099
human.FAM230E-LOC105377182-GGT3P.NCBI.                                               gcttaggcatgagaatcgctttaacttgggaggcggaggttgcagttaactgagattgca 56736
human.FAM230J-LOC105372942-GGTLC5P.NCBI.rev.compl                                    ------------------------------------------------------------ 46364
human.FAM230A.ncRNA.18487127.18500594-GGTLC3.18516335.18518165.rev.compl.REF         ------------------------------------------------------------ 31039

chimp.LOC112206744-LOC107973052-GGT2.revcompl                                        ccactgcactccagcctggcaacagagcaaaactccatctctaaatagat-tagatagat 59124
chimp54546-135457.revcompl.Pan.troglodytes.clone.rp43-41g5.GenBank:AC099533.36      ------------------------------------------------------------ 66783
human.FAM30B-LOC105372935-GGT2.NCBI.                                                 ccactgcactccagcctggcaacagagcaaaactccatctctaaatagatatagatagat 58159
human.FAM230E-LOC105377182-GGT3P.NCBI.                                               ccactgcactccagcctggcaacagagcaaaactccatctctaaatagatatagatagat 56796
human.FAM230J-LOC105372942-GGTLC5P.NCBI.rev.compl                                    ------------------------------------------------------------ 46364
human.FAM230A.ncRNA.18487127.18500594-GGTLC3.18516335.18518165.rev.compl.REF         ------------------------------------------------------------ 31039

chimp.LOC112206744-LOC107973052-GGT2.revcompl                                        agatagatagatagata-atagata-------------------------------gata 59152
chimp54546-135457.revcompl.Pan.troglodytes.clone.rp43-41g5.GenBank:AC099533.36      ------------------------------------------------------------ 66783
human.FAM30B-LOC105372935-GGT2.NCBI.                                                 agatagatagatagatagatagatagatagatagat--------------------gata 58199
human.FAM230E-LOC105377182-GGT3P.NCBI.                                               agatagatagatagatagatagatagatagatagataatagatagatagatagatagata 56856
human.FAM230J-LOC105372942-GGTLC5P.NCBI.rev.compl                                    ------------------------------------------------------------ 46364
human.FAM230A.ncRNA.18487127.18500594-GGTLC3.18516335.18518165.rev.compl.REF         ------------------------------------------------------------ 31039

chimp.LOC112206744-LOC107973052-GGT2.revcompl                                        gatagatagatagttagataggaagaaataagtaaaaataacaaccaaacaacaaaacag 59212
chimp54546-135457.revcompl.Pan.troglodytes.clone.rp43-41g5.GenBank:AC099533.36      ------------------------------------------------------------ 66783
human.FAM30B-LOC105372935-GGT2.NCBI.                                                 gatagatagatagatagataggaagaaataagtaaaaataacaaccaaacaacaaaacag 58259
human.FAM230E-LOC105377182-GGT3P.NCBI.                                               gatagatagatagatagataggaagaaataagtaaaaataacaaccaaacaacaaaacag 56916
human.FAM230J-LOC105372942-GGTLC5P.NCBI.rev.compl                                    ------------------------------------------------------------ 46364
human.FAM230A.ncRNA.18487127.18500594-GGTLC3.18516335.18518165.rev.compl.REF         ------------------------------------------------------------ 31039

chimp.LOC112206744-LOC107973052-GGT2.revcompl                                        tggagtttatccaaagagacagagactcttagaactgagaaaggggccctgtttggctc 59272
chimp54546-135457.revcompl.Pan.troglodytes.clone.rp43-41g5.GenBank:AC099533.36      ------------------------------------------------------------ 66783
human.FAM30B-LOC105372935-GGT2.NCBI.                                                 tggagtttatccaaagaaacagagactcttagaactgagaaaggggccctgtttggctc 58319
human.FAM230E-LOC105377182-GGT3P.NCBI.                                               tggagtttatccaaagaaacagagactcttagaactgagaaaggggccctgtttggctc 56976
human.FAM230J-LOC105372942-GGTLC5P.NCBI.rev.compl                                    ------------------------------------------------------------ 46364
human.FAM230A.ncRNA.18487127.18500594-GGTLC3.18516335.18518165.rev.compl.REF         ------------------------------------------------------------ 31039

chimp.LOC112206744-LOC107973052-GGT2.revcompl                                        taggagacccacaccctgctctcggagtcaccgtccccttcccaaaggctactgagagag 59332
chimp54546-135457.revcompl.Pan.troglodytes.clone.rp43-41g5.GenBank:AC099533.36      ------------------------------------------------------------ 66783
human.FAM30B-LOC105372935-GGT2.NCBI.                                                 ta-gagacccacaccctgctctcggagtcaccgtccccttcccaaaggctactgagagag 58378
human.FAM230E-LOC105377182-GGT3P.NCBI.                                               ta-gagacccacaccctgctctcggagtcaccgtccccttcccaaaggctactgagagag 57035
human.FAM230J-LOC105372942-GGTLC5P.NCBI.rev.compl                                    ------------------------------------------------------------ 46364
human.FAM230A.ncRNA.18487127.18500594-GGTLC3.18516335.18518165.rev.compl.REF         ------------------------------------------------------------ 31039

chimp.LOC112206744-LOC107973052-GGT2.revcompl                                        tccaagcaaagcttacatgtggggaaactgagtctcagaggggtgaaggtattgctcagg 59392
chimp54546-135457.revcompl.Pan.troglodytes.clone.rp43-41g5.GenBank:AC099533.36      ------------------------------------------------------------ 66783
human.FAM30B-LOC105372935-GGT2.NCBI.                                                 tccaagcgaagcttgcatgtggggaaactgagtctcagaggggtgaaggtattgctcagg 58438
human.FAM230E-LOC105377182-GGT3P.NCBI.                                               tccaagcgaagcttacatgtggggaaactgagtctcagaggggtgaaggtattgctcagg 57095
human.FAM230J-LOC105372942-GGTLC5P.NCBI.rev.compl                                    ------------------------------------------------------------ 46364
human.FAM230A.ncRNA.18487127.18500594-GGTLC3.18516335.18518165.rev.compl.REF         ------------------------------------------------------------ 31039

chimp.LOC112206744-LOC107973052-GGT2.revcompl                                        tccacttgcccagttttcagggcccatgtcccatgccctgccccgctcacctcccaatct 59452
chimp54546-135457.revcompl.Pan.troglodytes.clone.rp43-41g5.GenBank:AC099533.36      ------------------------------------------------------------ 66783
human.FAM30B-LOC105372935-GGT2.NCBI.                                                 tccacttgcccagttttcagggcccatgtcccatgccctgccccgctcacctcccaatct 58498
human.FAM230E-LOC105377182-GGT3P.NCBI.                                               tccacttgcccagttttcagggcccatgtcccatgccctgccccgctcacctcccaatct 57155
human.FAM230J-LOC105372942-GGTLC5P.NCBI.rev.compl                                    ------------------------------------------------------------ 46364
human.FAM230A.ncRNA.18487127.18500594-GGTLC3.18516335.18518165.rev.compl.REF         ------------------------------------------------------------ 31039

chimp.LOC112206744-LOC107973052-GGT2.revcompl                                        ccgagcactgcttggcatccgcgcccacggcagccctggtgtacacatggttgtcaggtt 59512
chimp54546-135457.revcompl.Pan.troglodytes.clone.rp43-41g5.GenBank:AC099533.36      ------------------------------------------------------------ 66783
human.FAM30B-LOC105372935-GGT2.NCBI.                                                 ccaagcactgcttggcatccgcggccatggcagccctggtgtacacatggttgtcaggtt 58558
human.FAM230E-LOC105377182-GGT3P.NCBI.                                               ccaagcactgcttggcatccgcggccacggcagccctggtgtacacatggttgtcaggtt 57215
human.FAM230J-LOC105372942-GGTLC5P.NCBI.rev.compl                                    ------------------------------------------------------------ 46364
human.FAM230A.ncRNA.18487127.18500594-GGTLC3.18516335.18518165.rev.compl.REF         ------------------------------------------------------------ 31039

chimp.LOC112206744-LOC107973052-GGT2.revcompl                                        ccttggaggctgagggcagccagagacagaggccgacaatgaccagcacgaggaccacgg 59572
chimp54546-135457.revcompl.Pan.troglodytes.clone.rp43-41g5.GenBank:AC099533.36      ------------------------------------------------------------ 66783
human.FAM30B-LOC105372935-GGT2.NCBI.                                                 ccttggaggccgagggcagccagagacagaggccgacaatgaccagcaccaggaccacgg 58618
human.FAM230E-LOC105377182-GGT3P.NCBI.                                               ccttggaggccgagggcagccagagacagaggccgacaatgaccagcaccaggaccacgg 57275
human.FAM230J-LOC105372942-GGTLC5P.NCBI.rev.compl                                    ------------------------------------------------------------ 46364
human.FAM230A.ncRNA.18487127.18500594-GGTLC3.18516335.18518165.rev.compl.REF         ------------------------------------------------------------ 31039

chimp.LOC112206744-LOC107973052-GGT2.revcompl                                        ccagcaggcccagcaccactaacttcttcttcatggctctgctgcacccacggggtaagg 59632
chimp54546-135457.revcompl.Pan.troglodytes.clone.rp43-41g5.GenBank:AC099533.36      ------------------------------------------------------------ 66783
human.FAM30B-LOC105372935-GGT2.NCBI.                                                 ccagcaggcccagcaccactaacttcttcttcatggctctgctgcacccacggggtaagg 58678
human.FAM230E-LOC105377182-GGT3P.NCBI.                                               ccagcaggcccagcaccactaacttcttcttcatg------------------------- 57310
human.FAM230J-LOC105372942-GGTLC5P.NCBI.rev.compl                                    ------------------------------------------------------------ 46364
human.FAM230A.ncRNA.18487127.18500594-GGTLC3.18516335.18518165.rev.compl.REF         ------------------------------------------------------------ 31039

chimp.LOC112206744-LOC107973052-GGT2.revcompl                                        agcagggtcaggcccagcctcagacattccctggcccctccccaacagggcacagtctaa 59692
chimp54546-135457.revcompl.Pan.troglodytes.clone.rp43-41g5.GenBank:AC099533.36      ------------------------------------------------------------ 66783
human.FAM30B-LOC105372935-GGT2.NCBI.                                                 agcagggtcaggcccagcctcagacatgccctggcccctccccaacagggcacagtctaa 58738
human.FAM230E-LOC105377182-GGT3P.NCBI.                                               ------------------------------------------------------------ 57310
human.FAM230J-LOC105372942-GGTLC5P.NCBI.rev.compl                                    ------------------------------------------------------------ 46364
human.FAM230A.ncRNA.18487127.18500594-GGTLC3.18516335.18518165.rev.compl.REF         ------------------------------------------------------------ 31039

chimp.LOC112206744-LOC107973052-GGT2.revcompl                                        agtcaggcctcagaaacacaaggcctgtgtctccttcccgcttcccagaatacgtgcagg 59752
chimp54546-135457.revcompl.Pan.troglodytes.clone.rp43-41g5.GenBank:AC099533.36      ------------------------------------------------------------ 66783
human.FAM30B-LOC105372935-GGT2.NCBI.                                                 agtcgggcctcagaaacacaaggccggagtctccttcccgcttcccagaatacgtgcagg 58798
human.FAM230E-LOC105377182-GGT3P.NCBI.                                               ------------------------------------------------------------ 57310
human.FAM230J-LOC105372942-GGTLC5P.NCBI.rev.compl                                    ------------------------------------------------------------ 46364
human.FAM230A.ncRNA.18487127.18500594-GGTLC3.18516335.18518165.rev.compl.REF         ------------------------------------------------------------ 31039

chimp.LOC112206744-LOC107973052-GGT2.revcompl                                        ctgtccggcccccagacctttgcgcaggccatgccctctgccagaagctctgggcctcat 59812
chimp54546-135457.revcompl.Pan.troglodytes.clone.rp43-41g5.GenBank:AC099533.36      ------------------------------------------------------------ 66783
human.FAM30B-LOC105372935-GGT2.NCBI.                                                 ctgtccggcccccagacctttgcgcaggccatgccctctgccagaagctctgggcctcat 58858
human.FAM230E-LOC105377182-GGT3P.NCBI.                                               ------------------------------------------------------------ 57310
human.FAM230J-LOC105372942-GGTLC5P.NCBI.rev.compl                                    ------------------------------------------------------------ 46364
human.FAM230A.ncRNA.18487127.18500594-GGTLC3.18516335.18518165.rev.compl.REF         ------------------------------------------------------------ 31039

chimp.LOC112206744-LOC107973052-GGT2.revcompl                                        ctctgccctcccaaatcctccctgcttatcttcagagcccatcctggtaagaaccccatc 59872
chimp54546-135457.revcompl.Pan.troglodytes.clone.rp43-41g5.GenBank:AC099533.36      ------------------------------------------------------------ 66783
human.FAM30B-LOC105372935-GGT2.NCBI.                                                 ctctgccctcccaaatcctccctgcttatcttcagagcccatcctggtaagaaccccatc 58918
human.FAM230E-LOC105377182-GGT3P.NCBI.                                               ------------------------------------------------------------ 57310
human.FAM230J-LOC105372942-GGTLC5P.NCBI.rev.compl                                    ------------------------------------------------------------ 46364
human.FAM230A.ncRNA.18487127.18500594-GGTLC3.18516335.18518165.rev.compl.REF         ------------------------------------------------------------ 31039

chimp.LOC112206744-LOC107973052-GGT2.revcompl                                        tccaccagcggcccttcctgggagcccccagatttccacacccctctttctgcagggcct 59932
chimp54546-135457.revcompl.Pan.troglodytes.clone.rp43-41g5.GenBank:AC099533.36      ------------------------------------------------------------ 66783
human.FAM30B-LOC105372935-GGT2.NCBI.                                                 tccagcagcggcccttcctgggagcccccagatttccacacccctctttctgcagggcct 58978
human.FAM230E-LOC105377182-GGT3P.NCBI.                                               ------------------------------------------------------------ 57310
human.FAM230J-LOC105372942-GGTLC5P.NCBI.rev.compl                                    ------------------------------------------------------------ 46364
human.FAM230A.ncRNA.18487127.18500594-GGTLC3.18516335.18518165.rev.compl.REF         ------------------------------------------------------------ 31039

chimp.LOC112206744-LOC107973052-GGT2.revcompl                                        ggcctacctcctcacagtggctgagcctccactgcttagggagaagctccagtagggatg 59992
chimp54546-135457.revcompl.Pan.troglodytes.clone.rp43-41g5.GenBank:AC099533.36      ------------------------------------------------------------ 66783
```

```
human.FAM30B-LOC105372935-GGT2.NCBI.                                                    ggcctacctcctcacagtggctgagcctccactgcttagggagaagctccagcagggatg 59038
human.FAM230E-LOC105377182-GGT3P.NCBI.                                                  ------------------------------------------------------------ 57310
human.FAM230J-LOC105372942-GGTLC5P.NCBI.rev.compl                                       ------------------------------------------------------------ 46364
human.FAM230A.ncRNA.18487127.18500594-GGTLC3.18516335.18518165.rev.compl.REF            ------------------------------------------------------------ 31039

chimp.LOC112206744-LOC107973052-GGT2.revcompl                                           ggcctggcctggtttctcctgtgtcccccacctcagcctagagcctggcactgtccagga 60052
chimp54546-135457.revcompl.Pan.troglodytes.clone.rp43-41g5.GenBank:AC099533.36          ------------------------------------------------------------ 66783
human.FAM30B-LOC105372935-GGT2.NCBI.                                                    ggcctggcctggtttctcctgcgtcccccaccccagcctagagcctggcactgtccagga 59098
human.FAM230E-LOC105377182-GGT3P.NCBI.                                                  ------------------------------------------------------------ 57310
human.FAM230J-LOC105372942-GGTLC5P.NCBI.rev.compl                                       ------------------------------------------------------------ 46364
human.FAM230A.ncRNA.18487127.18500594-GGTLC3.18516335.18518165.rev.compl.REF            ------------------------------------------------------------ 31039

chimp.LOC112206744-LOC107973052-GGT2.revcompl                                           gtcctctgaagaccctccaccccacctggagcatggggtttagcttccatagtgcccaca 60112
chimp54546-135457.revcompl.Pan.troglodytes.clone.rp43-41g5.GenBank:AC099533.36          ------------------------------------------------------------ 66783
human.FAM30B-LOC105372935-GGT2.NCBI.                                                    gtcctctgaagaccctccaccccacctggagcatggggtttagcttccatagtgcccaca 59158
human.FAM230E-LOC105377182-GGT3P.NCBI.                                                  ------------------------------------------------------------ 57310
human.FAM230J-LOC105372942-GGTLC5P.NCBI.rev.compl                                       ------------------------------------------------------------ 46364
human.FAM230A.ncRNA.18487127.18500594-GGTLC3.18516335.18518165.rev.compl.REF            ------------------------------------------------------------ 31039

chimp.LOC112206744-LOC107973052-GGT2.revcompl                                           atcagagcgccccacagattcactgccacggggccaggacttaccgtccagcagcagaca 60172
chimp54546-135457.revcompl.Pan.troglodytes.clone.rp43-41g5.GenBank:AC099533.36          ------------------------------------------------------------ 66783
human.FAM30B-LOC105372935-GGT2.NCBI.                                                    atcagagcgccccacagattcactgccacggggccaggacttaccgtccagcagcagacg 59218
human.FAM230E-LOC105377182-GGT3P.NCBI.                                                  ------------------------------------------------------------ 57310
human.FAM230J-LOC105372942-GGTLC5P.NCBI.rev.compl                                       ------------------------------------------------------------ 46364
human.FAM230A.ncRNA.18487127.18500594-GGTLC3.18516335.18518165.rev.compl.REF            ------------------------------------------------------------ 31039

chimp.LOC112206744-LOC107973052-GGT2.revcompl                                           ggggccccaagccttgcctggggtgttggccacgaaagacaggaggatttggtggaaaca 60232
chimp54546-135457.revcompl.Pan.troglodytes.clone.rp43-41g5.GenBank:AC099533.36          ------------------------------------------------------------ 66783
human.FAM30B-LOC105372935-GGT2.NCBI.                                                    ggggccccaagccttgcctggggtgttggccacgaaagacaggaggatttggtggaaaca 59278
human.FAM230E-LOC105377182-GGT3P.NCBI.                                                  ------------------------------------------------------------ 57310
human.FAM230J-LOC105372942-GGTLC5P.NCBI.rev.compl                                       ------------------------------------------------------------ 46364
human.FAM230A.ncRNA.18487127.18500594-GGTLC3.18516335.18518165.rev.compl.REF            ------------------------------------------------------------ 31039

chimp.LOC112206744-LOC107973052-GGT2.revcompl                                           gctgaggaaataaccggggtctccctcacactctgctgaagcctgtagccacagaatctt 60292
chimp54546-135457.revcompl.Pan.troglodytes.clone.rp43-41g5.GenBank:AC099533.36          ------------------------------------------------------------ 66783
human.FAM30B-LOC105372935-GGT2.NCBI.                                                    gctgaggaaataaccggggtctccctcacactctgctgaagcctgtagccacagaatctt 59338
human.FAM230E-LOC105377182-GGT3P.NCBI.                                                  ------------------------------------------------------------ 57310
human.FAM230J-LOC105372942-GGTLC5P.NCBI.rev.compl                                       ------------------------------------------------------------ 46364
human.FAM230A.ncRNA.18487127.18500594-GGTLC3.18516335.18518165.rev.compl.REF            ------------------------------------------------------------ 31039

chimp.LOC112206744-LOC107973052-GGT2.revcompl                                           cttcagagactccctgatcaggcagccttctcgttctcctgaaggtcaaaggaggttacc 60352
chimp54546-135457.revcompl.Pan.troglodytes.clone.rp43-41g5.GenBank:AC099533.36          ------------------------------------------------------------ 66783
human.FAM30B-LOC105372935-GGT2.NCBI.                                                    cttcagagactctctgatcaggcagccttctcgttctcctgaaggtcaagggaggttacc 59398
human.FAM230E-LOC105377182-GGT3P.NCBI.                                                  ------------------------------------------------------------ 57310
human.FAM230J-LOC105372942-GGTLC5P.NCBI.rev.compl                                       ------------------------------------------------------------ 46364
human.FAM230A.ncRNA.18487127.18500594-GGTLC3.18516335.18518165.rev.compl.REF            ------------------------------------------------------------ 31039

chimp.LOC112206744-LOC107973052-GGT2.revcompl                                           tgaagcacgcacagctcagacctttctggggtactccgtgttacctccctctgcctctag 60412
chimp54546-135457.revcompl.Pan.troglodytes.clone.rp43-41g5.GenBank:AC099533.36          ------------------------------------------------------------ 66783
human.FAM30B-LOC105372935-GGT2.NCBI.                                                    tgaagcacgcacagcccagacctttctggggggactccgtgttacctccctctgcctctag 59458
human.FAM230E-LOC105377182-GGT3P.NCBI.                                                  ------------------------------------------------------------ 57310
human.FAM230J-LOC105372942-GGTLC5P.NCBI.rev.compl                                       ------------------------------------------------------------ 46364
human.FAM230A.ncRNA.18487127.18500594-GGTLC3.18516335.18518165.rev.compl.REF            ------------------------------------------------------------ 31039

chimp.LOC112206744-LOC107973052-GGT2.revcompl                                           ctggtttctctgtctccagttgaactctggaggcaaagaggctgtcagtaacacatttgt 60472
chimp54546-135457.revcompl.Pan.troglodytes.clone.rp43-41g5.GenBank:AC099533.36          ------------------------------------------------------------ 66783
human.FAM30B-LOC105372935-GGT2.NCBI.                                                    ctggtttctctgtctccagttgaactctggaggcaaagaggctgtcagtaacacatttgt 59518
human.FAM230E-LOC105377182-GGT3P.NCBI.                                                  ------------------------------------------------------------ 57310
human.FAM230J-LOC105372942-GGTLC5P.NCBI.rev.compl                                       ------------------------------------------------------------ 46364
human.FAM230A.ncRNA.18487127.18500594-GGTLC3.18516335.18518165.rev.compl.REF            ------------------------------------------------------------ 31039

chimp.LOC112206744-LOC107973052-GGT2.revcompl                                           ttccatgaattctctcagcatgtctcccaggcacagctcaaagagggttttgcacggagc 60532
chimp54546-135457.revcompl.Pan.troglodytes.clone.rp43-41g5.GenBank:AC099533.36          ------------------------------------------------------------ 66783
human.FAM30B-LOC105372935-GGT2.NCBI.                                                    ttccatgaattctctcagcatgtctcccaggcac---------agggttttgcacggagc 59569
human.FAM230E-LOC105377182-GGT3P.NCBI.                                                  ------------------------------------------------------------ 57310
human.FAM230J-LOC105372942-GGTLC5P.NCBI.rev.compl                                       ------------------------------------------------------------ 46364
human.FAM230A.ncRNA.18487127.18500594-GGTLC3.18516335.18518165.rev.compl.REF            ------------------------------------------------------------ 31039

chimp.LOC112206744-LOC107973052-GGT2.revcompl                                           agggcaggtaggggacagggcattcctgcacaagcccaggatgtccatgcggtaagcatg 60592
chimp54546-135457.revcompl.Pan.troglodytes.clone.rp43-41g5.GenBank:AC099533.36          ------------------------------------------------------------ 66783
human.FAM30B-LOC105372935-GGT2.NCBI.                                                    agggcaggtaggggacagggcattcctgcacaagcccaggatgtgcatgcggtaagcatg 59629
human.FAM230E-LOC105377182-GGT3P.NCBI.                                                  ------------------------------------------------------------ 57310
human.FAM230J-LOC105372942-GGTLC5P.NCBI.rev.compl                                       ------------------------------------------------------------ 46364
human.FAM230A.ncRNA.18487127.18500594-GGTLC3.18516335.18518165.rev.compl.REF            ------------------------------------------------------------ 31039

chimp.LOC112206744-LOC107973052-GGT2.revcompl                                           gcaaagggggctcagggggcaccgccagcctgccctgctctgacgctggacttgccactc 60652
chimp54546-135457.revcompl.Pan.troglodytes.clone.rp43-41g5.GenBank:AC099533.36          ------------------------------------------------------------ 66783
human.FAM30B-LOC105372935-GGT2.NCBI.                                                    gcaaagggggctcaggaggcaccgccagcctgccctgctctgacgctggacttgccactc 59689
human.FAM230E-LOC105377182-GGT3P.NCBI.                                                  ------------------------------------------------------------ 57310
human.FAM230J-LOC105372942-GGTLC5P.NCBI.rev.compl                                       ------------------------------------------------------------ 46364
human.FAM230A.ncRNA.18487127.18500594-GGTLC3.18516335.18518165.rev.compl.REF            ------------------------------------------------------------ 31039

chimp.LOC112206744-LOC107973052-GGT2.revcompl                                           acctgctgtggggcctcaggcaaatcactgaactgtccagcctggatgacggcagcacct 60712
chimp54546-135457.revcompl.Pan.troglodytes.clone.rp43-41g5.GenBank:AC099533.36          ------------------------------------------------------------ 66783
human.FAM30B-LOC105372935-GGT2.NCBI.                                                    acctgctgtggggcctcaggcaaatcactgaactgtccagcctggatgacggcagcacct 59749
human.FAM230E-LOC105377182-GGT3P.NCBI.                                                  ------------------------------------------------------------ 57310
human.FAM230J-LOC105372942-GGTLC5P.NCBI.rev.compl                                       ------------------------------------------------------------ 46364
human.FAM230A.ncRNA.18487127.18500594-GGTLC3.18516335.18518165.rev.compl.REF            ------------------------------------------------------------ 31039

chimp.LOC112206744-LOC107973052-GGT2.revcompl                                           cacttgccttgctgctgggagtgttgtgaatagagtaggttagactgtgggcagggcttg 60772
chimp54546-135457.revcompl.Pan.troglodytes.clone.rp43-41g5.GenBank:AC099533.36          ------------------------------------------------------------ 66783
human.FAM30B-LOC105372935-GGT2.NCBI.                                                    cacttgccttgctgctgggagtgttgtgaatagagtaggttagactgtgggcagggcttg 59809
human.FAM230E-LOC105377182-GGT3P.NCBI.                                                  ------------------------------------------------------------ 57310
human.FAM230J-LOC105372942-GGTLC5P.NCBI.rev.compl                                       ------------------------------------------------------------ 46364
human.FAM230A.ncRNA.18487127.18500594-GGTLC3.18516335.18518165.rev.compl.REF            ------------------------------------------------------------ 31039

chimp.LOC112206744-LOC107973052-GGT2.revcompl                                           gtgaatggtagctgtgattatcatcatggctgcaccggggacacccccaggaggcctgag 60832
chimp54546-135457.revcompl.Pan.troglodytes.clone.rp43-41g5.GenBank:AC099533.36          ------------------------------------------------------------ 66783
human.FAM30B-LOC105372935-GGT2.NCBI.                                                    gtgaatggtagctgtgattatcatcatggctgcactggggacacccccaggaggcctgag 59869
human.FAM230E-LOC105377182-GGT3P.NCBI.                                                  ------------------------------------------------------------ 57310
human.FAM230J-LOC105372942-GGTLC5P.NCBI.rev.compl                                       ------------------------------------------------------------ 46364
human.FAM230A.ncRNA.18487127.18500594-GGTLC3.18516335.18518165.rev.compl.REF            ------------------------------------------------------------ 31039

chimp.LOC112206744-LOC107973052-GGT2.revcompl                                           tggcacaggtctcttgctcactgtatgtcccctgtggactcccttccaggctgtgcagtg 60892
chimp54546-135457.revcompl.Pan.troglodytes.clone.rp43-41g5.GenBank:AC099533.36          ------------------------------------------------------------ 66783
human.FAM30B-LOC105372935-GGT2.NCBI.                                                    tggcacaggtctcttgctcactgtatgtcccctgtggactcccttccaggctgtgcagtg 59929
human.FAM230E-LOC105377182-GGT3P.NCBI.                                                  ------------------------------------------------------------ 57310
human.FAM230J-LOC105372942-GGTLC5P.NCBI.rev.compl                                       ------------------------------------------------------------ 46364
human.FAM230A.ncRNA.18487127.18500594-GGTLC3.18516335.18518165.rev.compl.REF            ------------------------------------------------------------ 31039

chimp.LOC112206744-LOC107973052-GGT2.revcompl                                           agtggcagcagtgacccttgggaagcctcatggctatggcagcaggtgacaggtgtgaca 60952
chimp54546-135457.revcompl.Pan.troglodytes.clone.rp43-41g5.GenBank:AC099533.36          ------------------------------------------------------------ 66783
human.FAM30B-LOC105372935-GGT2.NCBI.                                                    agtggcagcagtgacccttgggaagtctcatggctacggcagcaggtgacaggtgtgaca 59989
human.FAM230E-LOC105377182-GGT3P.NCBI.                                                  ------------------------------------------------------------ 57310
human.FAM230J-LOC105372942-GGTLC5P.NCBI.rev.compl                                       ------------------------------------------------------------ 46364
human.FAM230A.ncRNA.18487127.18500594-GGTLC3.18516335.18518165.rev.compl.REF            ------------------------------------------------------------ 31039

chimp.LOC112206744-LOC107973052-GGT2.revcompl                                           acagggaagaaggatgtggtgacagaggttggggttcccctctcccacagtcagtttccc 61012
chimp54546-135457.revcompl.Pan.troglodytes.clone.rp43-41g5.GenBank:AC099533.36          ------------------------------------------------------------ 66783
human.FAM30B-LOC105372935-GGT2.NCBI.                                                    acagggaagagggatgtggtgacagaggttggggttcccctctcccacagtcagtttccc 60049
human.FAM230E-LOC105377182-GGT3P.NCBI.                                                  ------------------------------------------------------------ 57310
human.FAM230J-LOC105372942-GGTLC5P.NCBI.rev.compl                                       ------------------------------------------------------------ 46364
human.FAM230A.ncRNA.18487127.18500594-GGTLC3.18516335.18518165.rev.compl.REF            ------------------------------------------------------------ 31039

chimp.LOC112206744-LOC107973052-GGT2.revcompl                                           acaaagggcggtgtctgccagcaagcccctccaatgagccccaagctgggtttccctcca 61072
chimp54546-135457.revcompl.Pan.troglodytes.clone.rp43-41g5.GenBank:AC099533.36          ------------------------------------------------------------ 66783
human.FAM30B-LOC105372935-GGT2.NCBI.                                                    acaaagggcggtgtctgccagcaagcccctccaatgagccccaagcttggtttccctcca 60109
```

```
human.FAM230E-LOC105377182-GGT3P.NCBI.                                        ------------------------------------------------------------ 57310
human.FAM230J-LOC105372942-GGTLC5P.NCBI.rev.compl                             ------------------------------------------------------------ 46364
human.FAM230A.ncRNA.18487127.18500594-GGTLC3.18516335.18518165.rev.compl.REF  ------------------------------------------------------------ 31039

chimp.LOC112206744-LOC107973052-GGT2.revcompl                                 ctccaccctgtcccagtgcagagcgtctgacctcagaggcagacacactgtcccagaggt 61132
chimp54546-135457.revcompl.Pan.troglodytes.clone.rp43-41g5.GenBank:AC099533.36 ------------------------------------------------------------ 66783
human.FAM30B-LOC105372935-GGT2.NCBI.                                          ctccacgctgtcccagtgcagagcatctgacctcagaggcagacacactgtcccagaggt 60169
human.FAM230E-LOC105377182-GGT3P.NCBI.                                        ------------------------------------------------------------ 57310
human.FAM230J-LOC105372942-GGTLC5P.NCBI.rev.compl                             ------------------------------------------------------------ 46364
human.FAM230A.ncRNA.18487127.18500594-GGTLC3.18516335.18518165.rev.compl.REF  ------------------------------------------------------------ 31039

chimp.LOC112206744-LOC107973052-GGT2.revcompl                                 ggtctacgaatggagtccctgtgccctccccacacacagggaacatccaaatgccatcgt 61192
chimp54546-135457.revcompl.Pan.troglodytes.clone.rp43-41g5.GenBank:AC099533.36 ------------------------------------------------------------ 66783
human.FAM30B-LOC105372935-GGT2.NCBI.                                          ggtctacgaatggagtccccgtgccctccccacacacagggaacatccaaatgccatcgt 60229
human.FAM230E-LOC105377182-GGT3P.NCBI.                                        ------------------------------------------------------------ 57310
human.FAM230J-LOC105372942-GGTLC5P.NCBI.rev.compl                             ------------------------------------------------------------ 46364
human.FAM230A.ncRNA.18487127.18500594-GGTLC3.18516335.18518165.rev.compl.REF  ------------------------------------------------------------ 31039

chimp.LOC112206744-LOC107973052-GGT2.revcompl                                 ggaagggtgg-cacctccccaggcttggtgggcctggggccgatagtgtgatacatttga 61251
chimp54546-135457.revcompl.Pan.troglodytes.clone.rp43-41g5.GenBank:AC099533.36 ------------------------------------------------------------ 66783
human.FAM30B-LOC105372935-GGT2.NCBI.                                          ggaagggtggccacctccccaggcttggtgggcctggggccgatagtgtgatacatttga 60289
human.FAM230E-LOC105377182-GGT3P.NCBI.                                        ------------------------------------------------------------ 57310
human.FAM230J-LOC105372942-GGTLC5P.NCBI.rev.compl                             ------------------------------------------------------------ 46364
human.FAM230A.ncRNA.18487127.18500594-GGTLC3.18516335.18518165.rev.compl.REF  ------------------------------------------------------------ 31039

chimp.LOC112206744-LOC107973052-GGT2.revcompl                                 ccccctcccagccctggatgcagacaccaagagcagagagacctggcagtagtcatgcag 61311
chimp54546-135457.revcompl.Pan.troglodytes.clone.rp43-41g5.GenBank:AC099533.36 ------------------------------------------------------------ 66783
human.FAM30B-LOC105372935-GGT2.NCBI.                                          ccccctcccagccctggatgcagacaccaagagcagagagacctggcagtagccatgcag 60349
human.FAM230E-LOC105377182-GGT3P.NCBI.                                        ------------------------------------------------------------ 57310
human.FAM230J-LOC105372942-GGTLC5P.NCBI.rev.compl                             ------------------------------------------------------------ 46364
human.FAM230A.ncRNA.18487127.18500594-GGTLC3.18516335.18518165.rev.compl.REF  ------------------------------------------------------------ 31039

chimp.LOC112206744-LOC107973052-GGT2.revcompl                                 cagcgcaccaccccacatccccagcacaatccaaagcagcccgtcatccccaccgtgacc 61371
chimp54546-135457.revcompl.Pan.troglodytes.clone.rp43-41g5.GenBank:AC099533.36 ------------------------------------------------------------ 66783
human.FAM30B-LOC105372935-GGT2.NCBI.                                          cagcgcaccaccccacatccccagcacaatccaaagcagcccctcatccccaccgtgacc 60409
human.FAM230E-LOC105377182-GGT3P.NCBI.                                        ------------------------------------------------------------ 57310
human.FAM230J-LOC105372942-GGTLC5P.NCBI.rev.compl                             ------------------------------------------------------------ 46364
human.FAM230A.ncRNA.18487127.18500594-GGTLC3.18516335.18518165.rev.compl.REF  ------------------------------------------------------------ 31039

chimp.LOC112206744-LOC107973052-GGT2.revcompl                                 accacagcctgaatccaggtgccacctgtttctgacctgaactccctcacagcccctgcc 61431
chimp54546-135457.revcompl.Pan.troglodytes.clone.rp43-41g5.GenBank:AC099533.36 ------------------------------------------------------------ 66783
human.FAM30B-LOC105372935-GGT2.NCBI.                                          accacagcctgaatccaggcgccacctgtttctgacctgaactccctcacagcccctgcc 60469
human.FAM230E-LOC105377182-GGT3P.NCBI.                                        ------------------------------------------------------------ 57310
human.FAM230J-LOC105372942-GGTLC5P.NCBI.rev.compl                             ------------------------------------------------------------ 46364
human.FAM230A.ncRNA.18487127.18500594-GGTLC3.18516335.18518165.rev.compl.REF  ------------------------------------------------------------ 31039

chimp.LOC112206744-LOC107973052-GGT2.revcompl                                 tgcactccctccctccaacatcgcctgcccttcagtcttccagaaagcagctagagggct 61491
chimp54546-135457.revcompl.Pan.troglodytes.clone.rp43-41g5.GenBank:AC099533.36 ------------------------------------------------------------ 66783
human.FAM30B-LOC105372935-GGT2.NCBI.                                          tgcactccctccctccaacatcacctgcccttcagtcttccagaaagcagctagagggct 60529
human.FAM230E-LOC105377182-GGT3P.NCBI.                                        ------------------------------------------------------------ 57310
human.FAM230J-LOC105372942-GGTLC5P.NCBI.rev.compl                             ------------------------------------------------------------ 46364
human.FAM230A.ncRNA.18487127.18500594-GGTLC3.18516335.18518165.rev.compl.REF  ------------------------------------------------------------ 31039

chimp.LOC112206744-LOC107973052-GGT2.revcompl                                 ctgtctgtccaactgcagaacaggccctgcctcctccctgtcccgttggacagctcacac 61551
chimp54546-135457.revcompl.Pan.troglodytes.clone.rp43-41g5.GenBank:AC099533.36 ------------------------------------------------------------ 66783
human.FAM30B-LOC105372935-GGT2.NCBI.                                          ctgtctgtccaactgcagaacaggccctgcctcctccctgccctgttggacagctcacac 60589
human.FAM230E-LOC105377182-GGT3P.NCBI.                                        ------------------------------------------------------------ 57310
human.FAM230J-LOC105372942-GGTLC5P.NCBI.rev.compl                             ------------------------------------------------------------ 46364
human.FAM230A.ncRNA.18487127.18500594-GGTLC3.18516335.18518165.rev.compl.REF  ------------------------------------------------------------ 31039

chimp.LOC112206744-LOC107973052-GGT2.revcompl                                 ccttcaccaggcctgacagcactcttgccactccaacaccctgggtcccagccgggagtc 61611
chimp54546-135457.revcompl.Pan.troglodytes.clone.rp43-41g5.GenBank:AC099533.36 ------------------------------------------------------------ 66783
human.FAM30B-LOC105372935-GGT2.NCBI.                                          ccttcaccaggcctgacagcgctcttgccactccaacaccccgggtcccagccgggagtc 60649
human.FAM230E-LOC105377182-GGT3P.NCBI.                                        ------------------------------------------------------------ 57310
human.FAM230J-LOC105372942-GGTLC5P.NCBI.rev.compl                             ------------------------------------------------------------ 46364
human.FAM230A.ncRNA.18487127.18500594-GGTLC3.18516335.18518165.rev.compl.REF  ------------------------------------------------------------ 31039

chimp.LOC112206744-LOC107973052-GGT2.revcompl                                 caggtcagggttaagggttcctgatagagacaccgattcctggaggtccaaagagcctca 61671
chimp54546-135457.revcompl.Pan.troglodytes.clone.rp43-41g5.GenBank:AC099533.36 ------------------------------------------------------------ 66783
human.FAM30B-LOC105372935-GGT2.NCBI.                                          cgggtcagggttaagggttcctggtagagacaccgattcctggaggtccaaagagcctca 60709
human.FAM230E-LOC105377182-GGT3P.NCBI.                                        ------------------------------------------------------------ 57310
human.FAM230J-LOC105372942-GGTLC5P.NCBI.rev.compl                             ------------------------------------------------------------ 46364
human.FAM230A.ncRNA.18487127.18500594-GGTLC3.18516335.18518165.rev.compl.REF  ------------------------------------------------------------ 31039

chimp.LOC112206744-LOC107973052-GGT2.revcompl                                 ggagctgggccagcaatatgcagcatctattatggacacagaacattcccatcacatggc 61731
chimp54546-135457.revcompl.Pan.troglodytes.clone.rp43-41g5.GenBank:AC099533.36 ---------------------------------------------------------GGC 66786
human.FAM30B-LOC105372935-GGT2.NCBI.                                          ggagctgggccagcaatatgcagcatctattatggacacagaacattcccatcacatggc 60769
human.FAM230E-LOC105377182-GGT3P.NCBI.                                        ------------------------------------------------------------ 57310
human.FAM230J-LOC105372942-GGTLC5P.NCBI.rev.compl                             ------------------------------------------------------------ 46364
human.FAM230A.ncRNA.18487127.18500594-GGTLC3.18516335.18518165.rev.compl.REF  ------------------------------------------------------------ 31039

chimp.LOC112206744-LOC107973052-GGT2.revcompl                                 cgggtgcagtggctcacgcctataatcccag----------------------------- 61762
chimp54546-135457.revcompl.Pan.troglodytes.clone.rp43-41g5.GenBank:AC099533.36 TGGGAGTGGTGGCTTAGGCCTATAATCCCAGCACTTTGGGAGACAGAGGTAGGCCAATCA 66846
human.FAM30B-LOC105372935-GGT2.NCBI.                                          cgggtgcagtggctcacgcctttaatcccagcactttgggaggcagaggcaggtggatca 60829
human.FAM230E-LOC105377182-GGT3P.NCBI.                                        ------------------------------------------------------------ 57310
human.FAM230J-LOC105372942-GGTLC5P.NCBI.rev.compl                             ------------------------------------------------------------ 46364
human.FAM230A.ncRNA.18487127.18500594-GGTLC3.18516335.18518165.rev.compl.REF  ------------------------------------------------------------ 31039

chimp.LOC112206744-LOC107973052-GGT2.revcompl                                 ------------------------------------------------------------ 61762
chimp54546-135457.revcompl.Pan.troglodytes.clone.rp43-41g5.GenBank:AC099533.36 CATGAGGCCAGGAGTTCAAGACCAGCCTGGCCAACATGG-CAGAACCCCATCTCTACTAA 66905
human.FAM30B-LOC105372935-GGT2.NCBI.                                          cctgaggtgaagagttcgagaccagtctggccaacatggtgaaaccccatctccactaa 60889
human.FAM230E-LOC105377182-GGT3P.NCBI.                                        ------------------------------------------------------------ 57310
human.FAM230J-LOC105372942-GGTLC5P.NCBI.rev.compl                             ------------------------------------------------------------ 46364
human.FAM230A.ncRNA.18487127.18500594-GGTLC3.18516335.18518165.rev.compl.REF  ------------------------------------------------------------ 31039

chimp.LOC112206744-LOC107973052-GGT2.revcompl                                 ------------------------------------------------------------ 61762
chimp54546-135457.revcompl.Pan.troglodytes.clone.rp43-41g5.GenBank:AC099533.36 AAATAC-AAAAATGAGTTAGGCGTGGTG----------GTAATCCCAGTTACTCGAGAGG 66954
human.FAM30B-LOC105372935-GGT2.NCBI.                                          aaatacaaaaaattagccaggcatggtggcaggtgcctgtaatcccagctactc-aggag 60948
human.FAM230E-LOC105377182-GGT3P.NCBI.                                        ------------------------------------------------------------ 57310
human.FAM230J-LOC105372942-GGTLC5P.NCBI.rev.compl                             ------------------------------------------------------------ 46364
human.FAM230A.ncRNA.18487127.18500594-GGTLC3.18516335.18518165.rev.compl.REF  ------------------------------------------------------------ 31039

chimp.LOC112206744-LOC107973052-GGT2.revcompl                                 ------------------------------------------------------------ 61762
chimp54546-135457.revcompl.Pan.troglodytes.clone.rp43-41g5.GenBank:AC099533.36 CTGAGGCATGAGAATCGCTTGAACCTGGGAGGTGGAGGCTACAGTGAGTTGAGATTGCAC 67014
human.FAM30B-LOC105372935-GGT2.NCBI.                                          ctgaggcaggagaattgcttgaacccgggaggtggaggttgcagtgaggcaagattgcac 61008
human.FAM230E-LOC105377182-GGT3P.NCBI.                                        ------------------------------------------------------------ 57310
human.FAM230J-LOC105372942-GGTLC5P.NCBI.rev.compl                             ------------------------------------------------------------ 46364
human.FAM230A.ncRNA.18487127.18500594-GGTLC3.18516335.18518165.rev.compl.REF  ------------------------------------------------------------ 31039

chimp.LOC112206744-LOC107973052-GGT2.revcompl                                 ------------------------------------------------------------ 61762
chimp54546-135457.revcompl.Pan.troglodytes.clone.rp43-41g5.GenBank:AC099533.36 CACTGAACTCCAGCCTGGGC-------------------AACAGATAGAGACTCAAAAAA 67055
human.FAM30B-LOC105372935-GGT2.NCBI.                                          cactgcactccagcctgggccacaagagtgaaactccgtcacacacacacacacacacac 61068
human.FAM230E-LOC105377182-GGT3P.NCBI.                                        ------------------------------------------------------------ 57310
human.FAM230J-LOC105372942-GGTLC5P.NCBI.rev.compl                             ------------------------------------------------------------ 46364
human.FAM230A.ncRNA.18487127.18500594-GGTLC3.18516335.18518165.rev.compl.REF  ------------------------------------------------------------ 31039

chimp.LOC112206744-LOC107973052-GGT2.revcompl                                 ------------------------------------------------------------ 61762
chimp54546-135457.revcompl.Pan.troglodytes.clone.rp43-41g5.GenBank:AC099533.36 AAAAAGTCTCAAAAAAAAATGGAATTAGGAACTCAATTTGAACAGCACACTAGACACAAA 67115
human.FAM30B-LOC105372935-GGT2.NCBI.                                          acacaaaaaaaaaaaaaagaaaaaaaaggtctcctgctggaacacagactagttagaga 61128
human.FAM230E-LOC105377182-GGT3P.NCBI.                                        ------------------------------------------------------------ 57310
human.FAM230J-LOC105372942-GGTLC5P.NCBI.rev.compl                             ------------------------------------------------------------ 46364
human.FAM230A.ncRNA.18487127.18500594-GGTLC3.18516335.18518165.rev.compl.REF  ------------------------------------------------------------ 31039

chimp.LOC112206744-LOC107973052-GGT2.revcompl                                 ------------------------------------------------------------ 61762
chimp54546-135457.revcompl.Pan.troglodytes.clone.rp43-41g5.GenBank:AC099533.36 CATAAAAAAAAATAATGAACTGGTATTTATTTATTTA----------------------- 67152
human.FAM30B-LOC105372935-GGT2.NCBI.                                          aaggaaaaataaacaaatgatagtatgtgtattaataaaagaactagcaacacccactgc 61188
human.FAM230E-LOC105377182-GGT3P.NCBI.                                        ------------------------------------------------------------ 57310
```

```
human.FAM230J-LOC105372942-GGTLC5P.NCBI.rev.compl  ------------------------------------------------------------ 46364
human.FAM230A.ncRNA.18487127.18500594-GGTLC3.18516335.18518165.rev.compl.REF  ------------------------------------------------------------ 31039

chimp.LOC112206744-LOC107973052-GGT2.revcompl  ------------------------------------------------------------ 61762
chimp54546-135457.revcompl.Pan.troglodytes.clone.rp43-41g5.GenBank:AC099533.36  ------------------------------------------------------------ 67152
human.FAM30B-LOC105372935-GGT2.NCBI.  ttagttgtgataataaaaactggacattaatataggcagaaaaacagaacagccttgata 61248
human.FAM230E-LOC105377182-GGT3P.NCBI.  ------------------------------------------------------------ 57310
human.FAM230J-LOC105372942-GGTLC5P.NCBI.rev.compl  ------------------------------------------------------------ 46364
human.FAM230A.ncRNA.18487127.18500594-GGTLC3.18516335.18518165.rev.compl.REF  ------------------------------------------------------------ 31039

chimp.LOC112206744-LOC107973052-GGT2.revcompl  ------------------------------------------------------------ 61762
chimp54546-135457.revcompl.Pan.troglodytes.clone.rp43-41g5.GenBank:AC099533.36  ------------------------------------------------------------ 67152
human.FAM30B-LOC105372935-GGT2.NCBI.  tggttaaccctttgacactggcaaacactgtgaccagtgctgcccacactggaagctcct 61308
human.FAM230E-LOC105377182-GGT3P.NCBI.  ------------------------------------------------------------ 57310
human.FAM230J-LOC105372942-GGTLC5P.NCBI.rev.compl  ------------------------------------------------------------ 46364
human.FAM230A.ncRNA.18487127.18500594-GGTLC3.18516335.18518165.rev.compl.REF  ------------------------------------------------------------ 31039

chimp.LOC112206744-LOC107973052-GGT2.revcompl  ------------------------------------------------------------ 61762
chimp54546-135457.revcompl.Pan.troglodytes.clone.rp43-41g5.GenBank:AC099533.36  ------------------------------------------------------------ 67152
human.FAM30B-LOC105372935-GGT2.NCBI.  gcatccctcttcctcatgtttccttcagcattaaggagcaacaagggagacagccagttc 61368
human.FAM230E-LOC105377182-GGT3P.NCBI.  ------------------------------------------------------------ 57310
human.FAM230J-LOC105372942-GGTLC5P.NCBI.rev.compl  ------------------------------------------------------------ 46364
human.FAM230A.ncRNA.18487127.18500594-GGTLC3.18516335.18518165.rev.compl.REF  ------------------------------------------------------------ 31039

chimp.LOC112206744-LOC107973052-GGT2.revcompl  ------------------------------------------------------------ 61762
chimp54546-135457.revcompl.Pan.troglodytes.clone.rp43-41g5.GenBank:AC099533.36  ------------------------------------------------------------ 67152
human.FAM30B-LOC105372935-GGT2.NCBI.  atagttccttacgcatggagccaaaggaccttcaatgtacaaggtctgagcaaggacccg 61428
human.FAM230E-LOC105377182-GGT3P.NCBI.  ------------------------------------------------------------ 57310
human.FAM230J-LOC105372942-GGTLC5P.NCBI.rev.compl  ------------------------------------------------------------ 46364
human.FAM230A.ncRNA.18487127.18500594-GGTLC3.18516335.18518165.rev.compl.REF  ------------------------------------------------------------ 31039

chimp.LOC112206744-LOC107973052-GGT2.revcompl  ------------------------------------------------------------ 61762
chimp54546-135457.revcompl.Pan.troglodytes.clone.rp43-41g5.GenBank:AC099533.36  ------------------------------------------------------------ 67152
human.FAM30B-LOC105372935-GGT2.NCBI.  cagccacatgtgcttcctgcttcagcagtgccccgtgggtctcagagctacccaaaggcc 61488
human.FAM230E-LOC105377182-GGT3P.NCBI.  ------------------------------------------------------------ 57310
human.FAM230J-LOC105372942-GGTLC5P.NCBI.rev.compl  ------------------------------------------------------------ 46364
human.FAM230A.ncRNA.18487127.18500594-GGTLC3.18516335.18518165.rev.compl.REF  ------------------------------------------------------------ 31039

chimp.LOC112206744-LOC107973052-GGT2.revcompl  ------------------------------------------------------------ 61762
chimp54546-135457.revcompl.Pan.troglodytes.clone.rp43-41g5.GenBank:AC099533.36  ------------------------------------------------------------ 67152
human.FAM30B-LOC105372935-GGT2.NCBI.  tatccttccgagaggtttctcctctctcccaactgatgtcatacgtctcatcttccttgt 61548
human.FAM230E-LOC105377182-GGT3P.NCBI.  ------------------------------------------------------------ 57310
human.FAM230J-LOC105372942-GGTLC5P.NCBI.rev.compl  ------------------------------------------------------------ 46364
human.FAM230A.ncRNA.18487127.18500594-GGTLC3.18516335.18518165.rev.compl.REF  ------------------------------------------------------------ 31039

chimp.LOC112206744-LOC107973052-GGT2.revcompl  ------------------------------------------------------------ 61762
chimp54546-135457.revcompl.Pan.troglodytes.clone.rp43-41g5.GenBank:AC099533.36  ------------------------------------------------------------ 67152
human.FAM30B-LOC105372935-GGT2.NCBI.  cattcagatcatcaaccacaaacgtccttgccttatattttcatatcccttttcaccagt 61608
human.FAM230E-LOC105377182-GGT3P.NCBI.  ------------------------------------------------------------ 57310
human.FAM230J-LOC105372942-GGTLC5P.NCBI.rev.compl  ------------------------------------------------------------ 46364
human.FAM230A.ncRNA.18487127.18500594-GGTLC3.18516335.18518165.rev.compl.REF  ------------------------------------------------------------ 31039

chimp.LOC112206744-LOC107973052-GGT2.revcompl  ------------------------------------------------------------ 61762
chimp54546-135457.revcompl.Pan.troglodytes.clone.rp43-41g5.GenBank:AC099533.36  ------------------------------------------------------------ 67152
human.FAM30B-LOC105372935-GGT2.NCBI.  tacagggttagttaacttacgaaattcttaacatctgcattagatctttttaaagtttca 61668
human.FAM230E-LOC105377182-GGT3P.NCBI.  ------------------------------------------------------------ 57310
human.FAM230J-LOC105372942-GGTLC5P.NCBI.rev.compl  ------------------------------------------------------------ 46364
human.FAM230A.ncRNA.18487127.18500594-GGTLC3.18516335.18518165.rev.compl.REF  ------------------------------------------------------------ 31039

chimp.LOC112206744-LOC107973052-GGT2.revcompl  ------------------------------------------------------------ 61762
chimp54546-135457.revcompl.Pan.troglodytes.clone.rp43-41g5.GenBank:AC099533.36  ------------------------------------------------------------ 67152
human.FAM30B-LOC105372935-GGT2.NCBI.  ccctcaaccacctattatttagaagtgaacacagaaatttagtttccttgtgctatctgt 61728
human.FAM230E-LOC105377182-GGT3P.NCBI.  ------------------------------------------------------------ 57310
human.FAM230J-LOC105372942-GGTLC5P.NCBI.rev.compl  ------------------------------------------------------------ 46364
human.FAM230A.ncRNA.18487127.18500594-GGTLC3.18516335.18518165.rev.compl.REF  ------------------------------------------------------------ 31039

chimp.LOC112206744-LOC107973052-GGT2.revcompl  ------------------------------------------------------------ 61762
chimp54546-135457.revcompl.Pan.troglodytes.clone.rp43-41g5.GenBank:AC099533.36  ------------------------------------------------------------ 67152
human.FAM30B-LOC105372935-GGT2.NCBI.  tgacccctaaaatatgctgggagttttgggatttttttaaagtcaaatgcatggcataa 61788
human.FAM230E-LOC105377182-GGT3P.NCBI.  ------------------------------------------------------------ 57310
human.FAM230J-LOC105372942-GGTLC5P.NCBI.rev.compl  ------------------------------------------------------------ 46364
human.FAM230A.ncRNA.18487127.18500594-GGTLC3.18516335.18518165.rev.compl.REF  ------------------------------------------------------------ 31039

chimp.LOC112206744-LOC107973052-GGT2.revcompl  ------------------------------------------------------------ 61762
chimp54546-135457.revcompl.Pan.troglodytes.clone.rp43-41g5.GenBank:AC099533.36  ------------------------------------------------------------ 67152
human.FAM30B-LOC105372935-GGT2.NCBI.  agcaaaattacactactaaagaactgagtcaggccagacgctggcaacgtgaagacagct 61848
human.FAM230E-LOC105377182-GGT3P.NCBI.  ------------------------------------------------------------ 57310
human.FAM230J-LOC105372942-GGTLC5P.NCBI.rev.compl  ------------------------------------------------------------ 46364
human.FAM230A.ncRNA.18487127.18500594-GGTLC3.18516335.18518165.rev.compl.REF  ------------------------------------------------------------ 31039

chimp.LOC112206744-LOC107973052-GGT2.revcompl  ------------------------------------------------------------ 61762
chimp54546-135457.revcompl.Pan.troglodytes.clone.rp43-41g5.GenBank:AC099533.36  ------------------------------------------------------------ 67152
human.FAM30B-LOC105372935-GGT2.NCBI.  tctccttacccactaggttcatctttgcactgttaactcccaaagtattgcttctcggtg 61908
human.FAM230E-LOC105377182-GGT3P.NCBI.  ------------------------------------------------------------ 57310
human.FAM230J-LOC105372942-GGTLC5P.NCBI.rev.compl  ------------------------------------------------------------ 46364
human.FAM230A.ncRNA.18487127.18500594-GGTLC3.18516335.18518165.rev.compl.REF  ------------------------------------------------------------ 31039

chimp.LOC112206744-LOC107973052-GGT2.revcompl  ------------------------------------------------------------ 61762
chimp54546-135457.revcompl.Pan.troglodytes.clone.rp43-41g5.GenBank:AC099533.36  ------------------------------------------------------------ 67152
human.FAM30B-LOC105372935-GGT2.NCBI.  ctgggtgtgtggcattcatattccccggcatcccgggcctgaagatctgtgatgtgcaat 61968
human.FAM230E-LOC105377182-GGT3P.NCBI.  ------------------------------------------------------------ 57310
human.FAM230J-LOC105372942-GGTLC5P.NCBI.rev.compl  ------------------------------------------------------------ 46364
human.FAM230A.ncRNA.18487127.18500594-GGTLC3.18516335.18518165.rev.compl.REF  ------------------------------------------------------------ 31039

chimp.LOC112206744-LOC107973052-GGT2.revcompl  ------------------------------------------------------------ 61762
chimp54546-135457.revcompl.Pan.troglodytes.clone.rp43-41g5.GenBank:AC099533.36  ------------------------------------------------------------ 67152
human.FAM30B-LOC105372935-GGT2.NCBI.  agggttgggttcccctggactctttctatgaagatcttccctccgcggacgcgctgggtg 62028
human.FAM230E-LOC105377182-GGT3P.NCBI.  ------------------------------------------------------------ 57310
human.FAM230J-LOC105372942-GGTLC5P.NCBI.rev.compl  ------------------------------------------------------------ 46364
human.FAM230A.ncRNA.18487127.18500594-GGTLC3.18516335.18518165.rev.compl.REF  ------------------------------------------------------------ 31039

chimp.LOC112206744-LOC107973052-GGT2.revcompl  ------------------------------------------------------------ 61762
chimp54546-135457.revcompl.Pan.troglodytes.clone.rp43-41g5.GenBank:AC099533.36  ------------------------------------------------------------ 67152
human.FAM30B-LOC105372935-GGT2.NCBI.  tagatggcataggggaaggaagagtccatggtgctgacgatctgcacctctcgctctggc 62088
human.FAM230E-LOC105377182-GGT3P.NCBI.  ------------------------------------------------------------ 57310
human.FAM230J-LOC105372942-GGTLC5P.NCBI.rev.compl  ------------------------------------------------------------ 46364
human.FAM230A.ncRNA.18487127.18500594-GGTLC3.18516335.18518165.rev.compl.REF  ------------------------------------------------------------ 31039

chimp.LOC112206744-LOC107973052-GGT2.revcompl  ------------------------------------------------------------ 61762
chimp54546-135457.revcompl.Pan.troglodytes.clone.rp43-41g5.GenBank:AC099533.36  ------------------------------------------------------------ 67152
human.FAM30B-LOC105372935-GGT2.NCBI.  gacgaaggcaggtaaatggaccactggaaattctgttcagaaggtccctggtagccactc 62148
human.FAM230E-LOC105377182-GGT3P.NCBI.  ------------------------------------------------------------ 57310
human.FAM230J-LOC105372942-GGTLC5P.NCBI.rev.compl  ------------------------------------------------------------ 46364
human.FAM230A.ncRNA.18487127.18500594-GGTLC3.18516335.18518165.rev.compl.REF  ------------------------------------------------------------ 31039

chimp.LOC112206744-LOC107973052-GGT2.revcompl  ------------------------------------------------------------ 61762
chimp54546-135457.revcompl.Pan.troglodytes.clone.rp43-41g5.GenBank:AC099533.36  ------------------------------------------------------------ 67152
human.FAM30B-LOC105372935-GGT2.NCBI.  acattgcaccagatagtgatgtgggagccctccgtgcggtacaagggtccttcctgaacg 62208
human.FAM230E-LOC105377182-GGT3P.NCBI.  ------------------------------------------------------------ 57310
human.FAM230J-LOC105372942-GGTLC5P.NCBI.rev.compl  ------------------------------------------------------------ 46364
human.FAM230A.ncRNA.18487127.18500594-GGTLC3.18516335.18518165.rev.compl.REF  ------------------------------------------------------------ 31039

chimp.LOC112206744-LOC107973052-GGT2.revcompl  ------------------------------------------------------------ 61762
chimp54546-135457.revcompl.Pan.troglodytes.clone.rp43-41g5.GenBank:AC099533.36  ------------------------------------------------------------ 67152
human.FAM30B-LOC105372935-GGT2.NCBI.  gtgacctgccgctgtgctgacaccacacctacgagggagagaaacacacgcaacatgctc 62268
human.FAM230E-LOC105377182-GGT3P.NCBI.  ------------------------------------------------------------ 57310
human.FAM230J-LOC105372942-GGTLC5P.NCBI.rev.compl  ------------------------------------------------------------ 46364
```

```
human.FAM230A.ncRNA.18487127.18500594-GGTLC3.18516335.18518165.rev.compl.REF        ------------------------------------------------------------ 31039

chimp.LOC112206744-LOC107973052-GGT2.revcompl                                       ------------------------------------------------------------ 61762
chimp54546-135457.revcompl.Pan.troglodytes.clone.rp43-41g5.GenBank:AC099533.36      ------------------------------------------------------------ 67152
human.FAM30B-LOC105372935-GGT2.NCBI.                                                acttacttctcagaccaaaatgcaaagtaggcagcaatctccaaagggtttgttatttgt 62328
human.FAM230E-LOC105377182-GGT3P.NCBI.                                              ------------------------------------------------------------ 57310
human.FAM230J-LOC105372942-GGTLC5P.NCBI.rev.compl                                   ------------------------------------------------------------ 46364
human.FAM230A.ncRNA.18487127.18500594-GGTLC3.18516335.18518165.rev.compl.REF        ------------------------------------------------------------ 31039

chimp.LOC112206744-LOC107973052-GGT2.revcompl                                       ------------------------------------------------------------ 61762
chimp54546-135457.revcompl.Pan.troglodytes.clone.rp43-41g5.GenBank:AC099533.36      ------------------------------------------------------------ 67152
human.FAM30B-LOC105372935-GGT2.NCBI.                                                gtgacatgataataatataaacagcttactggccctttcaaaggcactctgtgatttata 62388
human.FAM230E-LOC105377182-GGT3P.NCBI.                                              ------------------------------------------------------------ 57310
human.FAM230J-LOC105372942-GGTLC5P.NCBI.rev.compl                                   ------------------------------------------------------------ 46364
human.FAM230A.ncRNA.18487127.18500594-GGTLC3.18516335.18518165.rev.compl.REF        ------------------------------------------------------------ 31039

chimp.LOC112206744-LOC107973052-GGT2.revcompl                                       ------------------------------------------------------------ 61762
chimp54546-135457.revcompl.Pan.troglodytes.clone.rp43-41g5.GenBank:AC099533.36      ------------------------------------------------------------ 67152
human.FAM30B-LOC105372935-GGT2.NCBI.                                                aatattaattaaaacactctgtggtaaagcactgtccctaatttgcatatatgggaattt 62448
human.FAM230E-LOC105377182-GGT3P.NCBI.                                              ------------------------------------------------------------ 57310
human.FAM230J-LOC105372942-GGTLC5P.NCBI.rev.compl                                   ------------------------------------------------------------ 46364
human.FAM230A.ncRNA.18487127.18500594-GGTLC3.18516335.18518165.rev.compl.REF        ------------------------------------------------------------ 31039

chimp.LOC112206744-LOC107973052-GGT2.revcompl                                       ------------------------------------------------------------ 61762
chimp54546-135457.revcompl.Pan.troglodytes.clone.rp43-41g5.GenBank:AC099533.36      ------------------------------------------------------------ 67152
human.FAM30B-LOC105372935-GGT2.NCBI.                                                gatcatgccaagatgtgcttttgttacttattcaacagctgacccttgggaatttcattt 62508
human.FAM230E-LOC105377182-GGT3P.NCBI.                                              ------------------------------------------------------------ 57310
human.FAM230J-LOC105372942-GGTLC5P.NCBI.rev.compl                                   ------------------------------------------------------------ 46364
human.FAM230A.ncRNA.18487127.18500594-GGTLC3.18516335.18518165.rev.compl.REF        ------------------------------------------------------------ 31039

chimp.LOC112206744-LOC107973052-GGT2.revcompl                                       ------------------------------------------------------------ 61762
chimp54546-135457.revcompl.Pan.troglodytes.clone.rp43-41g5.GenBank:AC099533.36      ------------------------------------------------------------ 67152
human.FAM30B-LOC105372935-GGT2.NCBI.                                                agcccctctgtttgtcatgtatacggcagttttatatctatttattaatgctgtgtataa 62568
human.FAM230E-LOC105377182-GGT3P.NCBI.                                              ------------------------------------------------------------ 57310
human.FAM230J-LOC105372942-GGTLC5P.NCBI.rev.compl                                   ------------------------------------------------------------ 46364
human.FAM230A.ncRNA.18487127.18500594-GGTLC3.18516335.18518165.rev.compl.REF        ------------------------------------------------------------ 31039

chimp.LOC112206744-LOC107973052-GGT2.revcompl                                       ------------------------------------------------------------ 61762
chimp54546-135457.revcompl.Pan.troglodytes.clone.rp43-41g5.GenBank:AC099533.36      ------------------------------------------------------------ 67152
human.FAM30B-LOC105372935-GGT2.NCBI.                                                agtttttaataaaataagcaggaaaagttgaatccttggcacaaattcagatgaaaacaa 62628
human.FAM230E-LOC105377182-GGT3P.NCBI.                                              ------------------------------------------------------------ 57310
human.FAM230J-LOC105372942-GGTLC5P.NCBI.rev.compl                                   ------------------------------------------------------------ 46364
human.FAM230A.ncRNA.18487127.18500594-GGTLC3.18516335.18518165.rev.compl.REF        ------------------------------------------------------------ 31039

chimp.LOC112206744-LOC107973052-GGT2.revcompl                                       ------------------------------------------------------------ 61762
chimp54546-135457.revcompl.Pan.troglodytes.clone.rp43-41g5.GenBank:AC099533.36      ------------------------------------------------------------ 67152
human.FAM30B-LOC105372935-GGT2.NCBI.                                                ataaaagcaaccatctaggaatctggctgaaattaaatgcttcttccttggtccgagtgc 62688
human.FAM230E-LOC105377182-GGT3P.NCBI.                                              ------------------------------------------------------------ 57310
human.FAM230J-LOC105372942-GGTLC5P.NCBI.rev.compl                                   ------------------------------------------------------------ 46364
human.FAM230A.ncRNA.18487127.18500594-GGTLC3.18516335.18518165.rev.compl.REF        ------------------------------------------------------------ 31039

chimp.LOC112206744-LOC107973052-GGT2.revcompl                                       ------------------------------------------------------------ 61762
chimp54546-135457.revcompl.Pan.troglodytes.clone.rp43-41g5.GenBank:AC099533.36      ------------------------------------------------------------ 67152
human.FAM30B-LOC105372935-GGT2.NCBI.                                                tggtgggagggagtcctgggagcacctttttactggtgctgcacccttcattttcccatg 62748
human.FAM230E-LOC105377182-GGT3P.NCBI.                                              ------------------------------------------------------------ 57310
human.FAM230J-LOC105372942-GGTLC5P.NCBI.rev.compl                                   ------------------------------------------------------------ 46364
human.FAM230A.ncRNA.18487127.18500594-GGTLC3.18516335.18518165.rev.compl.REF        ------------------------------------------------------------ 31039

chimp.LOC112206744-LOC107973052-GGT2.revcompl                                       ------------------------------------------------------------ 61762
chimp54546-135457.revcompl.Pan.troglodytes.clone.rp43-41g5.GenBank:AC099533.36      ------------------------------------------------------------ 67152
human.FAM30B-LOC105372935-GGT2.NCBI.                                                tagctgctccccaacaccaccccaaggagcattccttagtttctcttccctccataaatg 62808
human.FAM230E-LOC105377182-GGT3P.NCBI.                                              ------------------------------------------------------------ 57310
human.FAM230J-LOC105372942-GGTLC5P.NCBI.rev.compl                                   ------------------------------------------------------------ 46364
human.FAM230A.ncRNA.18487127.18500594-GGTLC3.18516335.18518165.rev.compl.REF        ------------------------------------------------------------ 31039

chimp.LOC112206744-LOC107973052-GGT2.revcompl                                       ------------------------------------------------------------ 61762
chimp54546-135457.revcompl.Pan.troglodytes.clone.rp43-41g5.GenBank:AC099533.36      ------------------------------------------------------------ 67152
human.FAM30B-LOC105372935-GGT2.NCBI.                                                gaggcagagctgagctcctgcaggctgctatcgttttaagctgagcacacccaaatacaa 62868
human.FAM230E-LOC105377182-GGT3P.NCBI.                                              ------------------------------------------------------------ 57310
human.FAM230J-LOC105372942-GGTLC5P.NCBI.rev.compl                                   ------------------------------------------------------------ 46364
human.FAM230A.ncRNA.18487127.18500594-GGTLC3.18516335.18518165.rev.compl.REF        ------------------------------------------------------------ 31039

chimp.LOC112206744-LOC107973052-GGT2.revcompl                                       ------------------------------------------------------------ 61762
chimp54546-135457.revcompl.Pan.troglodytes.clone.rp43-41g5.GenBank:AC099533.36      ------------------------------------------------------------ 67152
human.FAM30B-LOC105372935-GGT2.NCBI.                                                atgtgaagggctgaaacagatccctgatgcccagattcccacaaactgacaaagggggga 62928
human.FAM230E-LOC105377182-GGT3P.NCBI.                                              ------------------------------------------------------------ 57310
human.FAM230J-LOC105372942-GGTLC5P.NCBI.rev.compl                                   ------------------------------------------------------------ 46364
human.FAM230A.ncRNA.18487127.18500594-GGTLC3.18516335.18518165.rev.compl.REF        ------------------------------------------------------------ 31039

chimp.LOC112206744-LOC107973052-GGT2.revcompl                                       ------------------------------------------------------------ 61762
chimp54546-135457.revcompl.Pan.troglodytes.clone.rp43-41g5.GenBank:AC099533.36      ------------------------------------------------------------ 67152
human.FAM30B-LOC105372935-GGT2.NCBI.                                                actaacagtgcccagctaaacctgtgacaaggacagtcaggctacccttgagtttcccac 62988
human.FAM230E-LOC105377182-GGT3P.NCBI.                                              ------------------------------------------------------------ 57310
human.FAM230J-LOC105372942-GGTLC5P.NCBI.rev.compl                                   ------------------------------------------------------------ 46364
human.FAM230A.ncRNA.18487127.18500594-GGTLC3.18516335.18518165.rev.compl.REF        ------------------------------------------------------------ 31039

chimp.LOC112206744-LOC107973052-GGT2.revcompl                                       ------------------------------------------------------------ 61762
chimp54546-135457.revcompl.Pan.troglodytes.clone.rp43-41g5.GenBank:AC099533.36      ------------------------------------------------------------ 67152
human.FAM30B-LOC105372935-GGT2.NCBI.                                                tcaagactcccccgctgacctcctgccttctggctcagatactgttctccaaggtcactt 63048
human.FAM230E-LOC105377182-GGT3P.NCBI.                                              ------------------------------------------------------------ 57310
human.FAM230J-LOC105372942-GGTLC5P.NCBI.rev.compl                                   ------------------------------------------------------------ 46364
human.FAM230A.ncRNA.18487127.18500594-GGTLC3.18516335.18518165.rev.compl.REF        ------------------------------------------------------------ 31039

chimp.LOC112206744-LOC107973052-GGT2.revcompl                                       ------------------------------------------------------------ 61762
chimp54546-135457.revcompl.Pan.troglodytes.clone.rp43-41g5.GenBank:AC099533.36      ------------------------------------------------------------ 67152
human.FAM30B-LOC105372935-GGT2.NCBI.                                                ctgcctgccttctccatgttcttcttaaagtggcatggtgagtttaatttctcatttcgt 63108
human.FAM230E-LOC105377182-GGT3P.NCBI.                                              ------------------------------------------------------------ 57310
human.FAM230J-LOC105372942-GGTLC5P.NCBI.rev.compl                                   ------------------------------------------------------------ 46364
human.FAM230A.ncRNA.18487127.18500594-GGTLC3.18516335.18518165.rev.compl.REF        ------------------------------------------------------------ 31039

chimp.LOC112206744-LOC107973052-GGT2.revcompl                                       ------------------------------------------------------------ 61762
chimp54546-135457.revcompl.Pan.troglodytes.clone.rp43-41g5.GenBank:AC099533.36      ------------------------------------------------------------ 67152
human.FAM30B-LOC105372935-GGT2.NCBI.                                                gccaactcaacattgattgtgcctgagccactcagaaggtgggagggggcacacccctgc 63168
human.FAM230E-LOC105377182-GGT3P.NCBI.                                              ------------------------------------------------------------ 57310
human.FAM230J-LOC105372942-GGTLC5P.NCBI.rev.compl                                   ------------------------------------------------------------ 46364
human.FAM230A.ncRNA.18487127.18500594-GGTLC3.18516335.18518165.rev.compl.REF        ------------------------------------------------------------ 31039

chimp.LOC112206744-LOC107973052-GGT2.revcompl                                       ------------------------------------------------------------ 61762
chimp54546-135457.revcompl.Pan.troglodytes.clone.rp43-41g5.GenBank:AC099533.36      ------------------------------------------------------------ 67152
human.FAM30B-LOC105372935-GGT2.NCBI.                                                ctcttcatttcggcagtggtcacgtactgcaggttgtgaaaagcgcaggtgagttatagg 63228
human.FAM230E-LOC105377182-GGT3P.NCBI.                                              ------------------------------------------------------------ 57310
human.FAM230J-LOC105372942-GGTLC5P.NCBI.rev.compl                                   ------------------------------------------------------------ 46364
human.FAM230A.ncRNA.18487127.18500594-GGTLC3.18516335.18518165.rev.compl.REF        ------------------------------------------------------------ 31039

chimp.LOC112206744-LOC107973052-GGT2.revcompl                                       ------------------------------------------------------------ 61762
chimp54546-135457.revcompl.Pan.troglodytes.clone.rp43-41g5.GenBank:AC099533.36      ------------------------------------------------------------ 67152
human.FAM30B-LOC105372935-GGT2.NCBI.                                                cgcagctttgccaccaattagagctggttgtgtcatctcagacaacttgcttttcttctc 63288
human.FAM230E-LOC105377182-GGT3P.NCBI.                                              ------------------------------------------------------------ 57310
human.FAM230J-LOC105372942-GGTLC5P.NCBI.rev.compl                                   ------------------------------------------------------------ 46364
human.FAM230A.ncRNA.18487127.18500594-GGTLC3.18516335.18518165.rev.compl.REF        ------------------------------------------------------------ 31039

chimp.LOC112206744-LOC107973052-GGT2.revcompl                                       ------------------------------------------------------------ 61762
chimp54546-135457.revcompl.Pan.troglodytes.clone.rp43-41g5.GenBank:AC099533.36      ------------------------------------------------------------ 67152
human.FAM30B-LOC105372935-GGT2.NCBI.                                                cagaccttgatcttctcatctatcacatgatttctaagaacgctttagagagcactagca 63348
human.FAM230E-LOC105377182-GGT3P.NCBI.                                              ------------------------------------------------------------ 57310
human.FAM230J-LOC105372942-GGTLC5P.NCBI.rev.compl                                   ------------------------------------------------------------ 46364
human.FAM230A.ncRNA.18487127.18500594-GGTLC3.18516335.18518165.rev.compl.REF        ------------------------------------------------------------ 31039
```

```
chimp.LOC112206744-LOC107973052-GGT2.revcompl                                      ------------------------------------------------------------ 61762
chimp54546-135457.revcompl.Pan.troglodytes.clone.rp43-41g5.GenBank:AC099533.36     ------------------------------------------------------------ 67152
human.FAM30B-LOC105372935-GGT2.NCBI.                                               tgggttccgatgctggctttgtcacttattaactgtgatcttggataagctgcttaacct 63408
human.FAM230E-LOC105377182-GGT3P.NCBI.                                             ------------------------------------------------------------ 57310
human.FAM230J-LOC105372942-GGTLC5P.NCBI.rev.compl                                  ------------------------------------------------------------ 46364
human.FAM230A.ncRNA.18487127.18500594-GGTLC3.18516335.18518165.rev.compl.REF       ------------------------------------------------------------ 31039

chimp.LOC112206744-LOC107973052-GGT2.revcompl                                      ------------------------------------------------------------ 61762
chimp54546-135457.revcompl.Pan.troglodytes.clone.rp43-41g5.GenBank:AC099533.36     ------------------------------------------------------------ 67152
human.FAM30B-LOC105372935-GGT2.NCBI.                                               ctctaagcctcagttttctcatcggtaagatggggataataatgcctgtctcatagggtt 63468
human.FAM230E-LOC105377182-GGT3P.NCBI.                                             ------------------------------------------------------------ 57310
human.FAM230J-LOC105372942-GGTLC5P.NCBI.rev.compl                                  ------------------------------------------------------------ 46364
human.FAM230A.ncRNA.18487127.18500594-GGTLC3.18516335.18518165.rev.compl.REF       ------------------------------------------------------------ 31039

chimp.LOC112206744-LOC107973052-GGT2.revcompl                                      ------------------------------------------------------------ 61762
chimp54546-135457.revcompl.Pan.troglodytes.clone.rp43-41g5.GenBank:AC099533.36     ------------------------------------------------------------ 67152
human.FAM30B-LOC105372935-GGT2.NCBI.                                               attgagaagatttagtaacatctgtaaagaccctagcacagagcctagcccacaatcagt 63528
human.FAM230E-LOC105377182-GGT3P.NCBI.                                             ------------------------------------------------------------ 57310
human.FAM230J-LOC105372942-GGTLC5P.NCBI.rev.compl                                  ------------------------------------------------------------ 46364
human.FAM230A.ncRNA.18487127.18500594-GGTLC3.18516335.18518165.rev.compl.REF       ------------------------------------------------------------ 31039

chimp.LOC112206744-LOC107973052-GGT2.revcompl                                      ------------------------------------------------------------ 61762
chimp54546-135457.revcompl.Pan.troglodytes.clone.rp43-41g5.GenBank:AC099533.36     ------------------------------------------------------------ 67152
human.FAM30B-LOC105372935-GGT2.NCBI.                                               cctcaataaagagctgctggctaagacatatttctgctggaacattctgtggctctggat 63588
human.FAM230E-LOC105377182-GGT3P.NCBI.                                             ------------------------------------------------------------ 57310
human.FAM230J-LOC105372942-GGTLC5P.NCBI.rev.compl                                  ------------------------------------------------------------ 46364
human.FAM230A.ncRNA.18487127.18500594-GGTLC3.18516335.18518165.rev.compl.REF       ------------------------------------------------------------ 31039

chimp.LOC112206744-LOC107973052-GGT2.revcompl                                      ------------------------------------------------------------ 61762
chimp54546-135457.revcompl.Pan.troglodytes.clone.rp43-41g5.GenBank:AC099533.36     ------------------------------------------------------------ 67152
human.FAM30B-LOC105372935-GGT2.NCBI.                                               ccatagaggacaggatttggccaatgactgcttaaggacatttcaacggccccttgacaa 63648
human.FAM230E-LOC105377182-GGT3P.NCBI.                                             ------------------------------------------------------------ 57310
human.FAM230J-LOC105372942-GGTLC5P.NCBI.rev.compl                                  ------------------------------------------------------------ 46364
human.FAM230A.ncRNA.18487127.18500594-GGTLC3.18516335.18518165.rev.compl.REF       ------------------------------------------------------------ 31039

chimp.LOC112206744-LOC107973052-GGT2.revcompl                                      ------------------------------------------------------------ 61762
chimp54546-135457.revcompl.Pan.troglodytes.clone.rp43-41g5.GenBank:AC099533.36     ------------------------------------------------------------ 67152
human.FAM30B-LOC105372935-GGT2.NCBI.                                               tgacaccatcactgggctgcccacctcatcccacagctgcagcccattcttgtgatagct 63708
human.FAM230E-LOC105377182-GGT3P.NCBI.                                             ------------------------------------------------------------ 57310
human.FAM230J-LOC105372942-GGTLC5P.NCBI.rev.compl                                  ------------------------------------------------------------ 46364
human.FAM230A.ncRNA.18487127.18500594-GGTLC3.18516335.18518165.rev.compl.REF       ------------------------------------------------------------ 31039

chimp.LOC112206744-LOC107973052-GGT2.revcompl                                      ------------------------------------------------------------ 61762
chimp54546-135457.revcompl.Pan.troglodytes.clone.rp43-41g5.GenBank:AC099533.36     ------------------------------------------------------------ 67152
human.FAM30B-LOC105372935-GGT2.NCBI.                                               tccctttccccacaaatgggaatggcggctgccctggctatgcctatctgaagtggat 63768
human.FAM230E-LOC105377182-GGT3P.NCBI.                                             ------------------------------------------------------------ 57310
human.FAM230J-LOC105372942-GGTLC5P.NCBI.rev.compl                                  ------------------------------------------------------------ 46364
human.FAM230A.ncRNA.18487127.18500594-GGTLC3.18516335.18518165.rev.compl.REF       ------------------------------------------------------------ 31039

chimp.LOC112206744-LOC107973052-GGT2.revcompl                                      ------------------------------------------------------------ 61762
chimp54546-135457.revcompl.Pan.troglodytes.clone.rp43-41g5.GenBank:AC099533.36     ------------------------------------------------------------ 67152
human.FAM30B-LOC105372935-GGT2.NCBI.                                               ccagctggtgaacagccgggacagctcaaatactagtggagtgcagctctctgcccagcc 63828
human.FAM230E-LOC105377182-GGT3P.NCBI.                                             ------------------------------------------------------------ 57310
human.FAM230J-LOC105372942-GGTLC5P.NCBI.rev.compl                                  ------------------------------------------------------------ 46364
human.FAM230A.ncRNA.18487127.18500594-GGTLC3.18516335.18518165.rev.compl.REF       ------------------------------------------------------------ 31039

chimp.LOC112206744-LOC107973052-GGT2.revcompl                                      ------------------------------------------------------------ 61762
chimp54546-135457.revcompl.Pan.troglodytes.clone.rp43-41g5.GenBank:AC099533.36     ------------------------------------------------------------ 67152
human.FAM30B-LOC105372935-GGT2.NCBI.                                               cagcaggtgactgtgcctcagtgtaagctaatctaatctgcaagatgacagacctcaaaa 63888
human.FAM230E-LOC105377182-GGT3P.NCBI.                                             ------------------------------------------------------------ 57310
human.FAM230J-LOC105372942-GGTLC5P.NCBI.rev.compl                                  ------------------------------------------------------------ 46364
human.FAM230A.ncRNA.18487127.18500594-GGTLC3.18516335.18518165.rev.compl.REF       ------------------------------------------------------------ 31039

chimp.LOC112206744-LOC107973052-GGT2.revcompl                                      ------------------------------------------------------------ 61762
chimp54546-135457.revcompl.Pan.troglodytes.clone.rp43-41g5.GenBank:AC099533.36     ------------------------------------------------------------ 67152
human.FAM30B-LOC105372935-GGT2.NCBI.                                               cggcccatacaatgaaatgcccatggcttgcaaaactttaattcatgcagaacctttcct 63948
human.FAM230E-LOC105377182-GGT3P.NCBI.                                             ------------------------------------------------------------ 57310
human.FAM230J-LOC105372942-GGTLC5P.NCBI.rev.compl                                  ------------------------------------------------------------ 46364
human.FAM230A.ncRNA.18487127.18500594-GGTLC3.18516335.18518165.rev.compl.REF       ------------------------------------------------------------ 31039

chimp.LOC112206744-LOC107973052-GGT2.revcompl                                      ------------------------------------------------------------ 61762
chimp54546-135457.revcompl.Pan.troglodytes.clone.rp43-41g5.GenBank:AC099533.36     ------------------------------------------------------------ 67152
human.FAM30B-LOC105372935-GGT2.NCBI.                                               cacaaacaaaatcttacacagcaccccatcttggcttcagcagaaccaggaaaaataaaa 64008
human.FAM230E-LOC105377182-GGT3P.NCBI.                                             ------------------------------------------------------------ 57310
human.FAM230J-LOC105372942-GGTLC5P.NCBI.rev.compl                                  ------------------------------------------------------------ 46364
human.FAM230A.ncRNA.18487127.18500594-GGTLC3.18516335.18518165.rev.compl.REF       ------------------------------------------------------------ 31039

chimp.LOC112206744-LOC107973052-GGT2.revcompl                                      ------------------------------------------------------------ 61762
chimp54546-135457.revcompl.Pan.troglodytes.clone.rp43-41g5.GenBank:AC099533.36     ------------------------------------------------------------ 67152
human.FAM30B-LOC105372935-GGT2.NCBI.                                               aattggcaggggaaggaaataaacagaacaagtggtatctgaatttatttgatatattct 64068
human.FAM230E-LOC105377182-GGT3P.NCBI.                                             ------------------------------------------------------------ 57310
human.FAM230J-LOC105372942-GGTLC5P.NCBI.rev.compl                                  ------------------------------------------------------------ 46364
human.FAM230A.ncRNA.18487127.18500594-GGTLC3.18516335.18518165.rev.compl.REF       ------------------------------------------------------------ 31039

chimp.LOC112206744-LOC107973052-GGT2.revcompl                                      ------------------------------------------------------------ 61762
chimp54546-135457.revcompl.Pan.troglodytes.clone.rp43-41g5.GenBank:AC099533.36     ------------------------------------------------------------ 67152
human.FAM30B-LOC105372935-GGT2.NCBI.                                               ggttgatggcattaattatgacatttaaagtcaccatgaggacaactcattagtaatatc 64128
human.FAM230E-LOC105377182-GGT3P.NCBI.                                             ------------------------------------------------------------ 57310
human.FAM230J-LOC105372942-GGTLC5P.NCBI.rev.compl                                  ------------------------------------------------------------ 46364
human.FAM230A.ncRNA.18487127.18500594-GGTLC3.18516335.18518165.rev.compl.REF       ------------------------------------------------------------ 31039

chimp.LOC112206744-LOC107973052-GGT2.revcompl                                      ------------------------------------------------------------ 61762
chimp54546-135457.revcompl.Pan.troglodytes.clone.rp43-41g5.GenBank:AC099533.36     ------------------------------------------------------------ 67152
human.FAM30B-LOC105372935-GGT2.NCBI.                                               agtatgttaaaatatggtgcataacatttttaatatatgtacagtcatgcactgaataac 64188
human.FAM230E-LOC105377182-GGT3P.NCBI.                                             ------------------------------------------------------------ 57310
human.FAM230J-LOC105372942-GGTLC5P.NCBI.rev.compl                                  ------------------------------------------------------------ 46364
human.FAM230A.ncRNA.18487127.18500594-GGTLC3.18516335.18518165.rev.compl.REF       ------------------------------------------------------------ 31039

chimp.LOC112206744-LOC107973052-GGT2.revcompl                                      ------------------------------------------------------------ 61762
chimp54546-135457.revcompl.Pan.troglodytes.clone.rp43-41g5.GenBank:AC099533.36     ------------------------------------------------------------ 67152
human.FAM30B-LOC105372935-GGT2.NCBI.                                               atttcagtcaatgaggaaacacatgtgcaacagtgatcctgtaagattataatggagcat 64248
human.FAM230E-LOC105377182-GGT3P.NCBI.                                             ------------------------------------------------------------ 57310
human.FAM230J-LOC105372942-GGTLC5P.NCBI.rev.compl                                  ------------------------------------------------------------ 46364
human.FAM230A.ncRNA.18487127.18500594-GGTLC3.18516335.18518165.rev.compl.REF       ------------------------------------------------------------ 31039

chimp.LOC112206744-LOC107973052-GGT2.revcompl                                      ------------------------------------------------------------ 61762
chimp54546-135457.revcompl.Pan.troglodytes.clone.rp43-41g5.GenBank:AC099533.36     ------------------------------------------------------------ 67152
human.FAM30B-LOC105372935-GGT2.NCBI.                                               atatagagatctaatatatggcacttaatgttggcatggcagatcaagtaggggaaatga 64308
human.FAM230E-LOC105377182-GGT3P.NCBI.                                             ------------------------------------------------------------ 57310
human.FAM230J-LOC105372942-GGTLC5P.NCBI.rev.compl                                  ------------------------------------------------------------ 46364
human.FAM230A.ncRNA.18487127.18500594-GGTLC3.18516335.18518165.rev.compl.REF       ------------------------------------------------------------ 31039

chimp.LOC112206744-LOC107973052-GGT2.revcompl                                      ------------------------------------------------------------ 61762
chimp54546-135457.revcompl.Pan.troglodytes.clone.rp43-41g5.GenBank:AC099533.36     ------------------------------------------------------------ 67152
human.FAM30B-LOC105372935-GGT2.NCBI.                                               ctgatatttagtaacagtgctgggacatttgattttccataataaaatatataaatgaaa 64368
human.FAM230E-LOC105377182-GGT3P.NCBI.                                             ------------------------------------------------------------ 57310
human.FAM230J-LOC105372942-GGTLC5P.NCBI.rev.compl                                  ------------------------------------------------------------ 46364
human.FAM230A.ncRNA.18487127.18500594-GGTLC3.18516335.18518165.rev.compl.REF       ------------------------------------------------------------ 31039

chimp.LOC112206744-LOC107973052-GGT2.revcompl                                      ------------------------------------------------------------ 61762
chimp54546-135457.revcompl.Pan.troglodytes.clone.rp43-41g5.GenBank:AC099533.36     ------------------------------------------------------------ 67152
human.FAM30B-LOC105372935-GGT2.NCBI.                                               atatatatcccatctaggtttgttgaaatacaccctatgatgttcacacaagaatgaaat 64428
human.FAM230E-LOC105377182-GGT3P.NCBI.                                             ------------------------------------------------------------ 57310
human.FAM230J-LOC105372942-GGTLC5P.NCBI.rev.compl                                  ------------------------------------------------------------ 46364
human.FAM230A.ncRNA.18487127.18500594-GGTLC3.18516335.18518165.rev.compl.REF       ------------------------------------------------------------ 31039
```

```
chimp.LOC112206744-LOC107973052-GGT2.revcompl                                    ------------------------------------------------------------ 61762
chimp54546-135457.revcompl.Pan.troglodytes.clone.rp43-41g5.GenBank:AC099533.36   ------------------------------------------------------------ 67152
human.FAM30B-LOC105372935-GGT2.NCBI.                                             tgcctaatgatgcatttcttaaaacatgtccccatcattaagtgacccatgactgtatac 64488
human.FAM230E-LOC105377182-GGT3P.NCBI.                                           ------------------------------------------------------------ 57310
human.FAM230J-LOC105372942-GGTLC5P.NCBI.rev.compl                                ------------------------------------------------------------ 46364
human.FAM230A.ncRNA.18487127.18500594-GGTLC3.18516335.18518165.rev.compl.REF     ------------------------------------------------------------ 31039

chimp.LOC112206744-LOC107973052-GGT2.revcompl                                    ------------------------------------------------------------ 61762
chimp54546-135457.revcompl.Pan.troglodytes.clone.rp43-41g5.GenBank:AC099533.36   ------------------------------------------------------------ 67152
human.FAM30B-LOC105372935-GGT2.NCBI.                                             acacacacatatggtgacacttaaatcaaatgattgcagtattcctgaaacacagaacat 64548
human.FAM230E-LOC105377182-GGT3P.NCBI.                                           ------------------------------------------------------------ 57310
human.FAM230J-LOC105372942-GGTLC5P.NCBI.rev.compl                                ------------------------------------------------------------ 46364
human.FAM230A.ncRNA.18487127.18500594-GGTLC3.18516335.18518165.rev.compl.REF     ------------------------------------------------------------ 31039

chimp.LOC112206744-LOC107973052-GGT2.revcompl                                    ------------------------------------------------------------ 61762
chimp54546-135457.revcompl.Pan.troglodytes.clone.rp43-41g5.GenBank:AC099533.36   ------------------------------------------------------------ 67152
human.FAM30B-LOC105372935-GGT2.NCBI.                                             tttgcaaacaatttacctacattcataacattaggcatccttagaattgcagtgttctat 64608
human.FAM230E-LOC105377182-GGT3P.NCBI.                                           ------------------------------------------------------------ 57310
human.FAM230J-LOC105372942-GGTLC5P.NCBI.rev.compl                                ------------------------------------------------------------ 46364
human.FAM230A.ncRNA.18487127.18500594-GGTLC3.18516335.18518165.rev.compl.REF     ------------------------------------------------------------ 31039

chimp.LOC112206744-LOC107973052-GGT2.revcompl                                    ------------------------------------------------------------ 61762
chimp54546-135457.revcompl.Pan.troglodytes.clone.rp43-41g5.GenBank:AC099533.36   ------------------------------------------------------------ 67152
human.FAM30B-LOC105372935-GGT2.NCBI.                                             gacagagcaactacaaccaacccccatccatctcctgcaaagaaggctggaggcaggtca 64668
human.FAM230E-LOC105377182-GGT3P.NCBI.                                           ------------------------------------------------------------ 57310
human.FAM230J-LOC105372942-GGTLC5P.NCBI.rev.compl                                ------------------------------------------------------------ 46364
human.FAM230A.ncRNA.18487127.18500594-GGTLC3.18516335.18518165.rev.compl.REF     ------------------------------------------------------------ 31039

chimp.LOC112206744-LOC107973052-GGT2.revcompl                                    ------------------------------------------------------------ 61762
chimp54546-135457.revcompl.Pan.troglodytes.clone.rp43-41g5.GenBank:AC099533.36   ------------------------------------------------------------ 67152
human.FAM30B-LOC105372935-GGT2.NCBI.                                             gggtacaccagcatcttcaagaactgcttctcaatcctggacctacactggaatcacctg 64728
human.FAM230E-LOC105377182-GGT3P.NCBI.                                           ------------------------------------------------------------ 57310
human.FAM230J-LOC105372942-GGTLC5P.NCBI.rev.compl                                ------------------------------------------------------------ 46364
human.FAM230A.ncRNA.18487127.18500594-GGTLC3.18516335.18518165.rev.compl.REF     ------------------------------------------------------------ 31039

chimp.LOC112206744-LOC107973052-GGT2.revcompl                                    ------------------------------------------------------------ 61762
chimp54546-135457.revcompl.Pan.troglodytes.clone.rp43-41g5.GenBank:AC099533.36   ------------------------------------------------------------ 67152
human.FAM30B-LOC105372935-GGT2.NCBI.                                             gggagctttaaaaaaaataacagtgcctggaccccacctgacataactggtcttaggatt 64788
human.FAM230E-LOC105377182-GGT3P.NCBI.                                           ------------------------------------------------------------ 57310
human.FAM230J-LOC105372942-GGTLC5P.NCBI.rev.compl                                ------------------------------------------------------------ 46364
human.FAM230A.ncRNA.18487127.18500594-GGTLC3.18516335.18518165.rev.compl.REF     ------------------------------------------------------------ 31039

chimp.LOC112206744-LOC107973052-GGT2.revcompl                                    ------------------------------------------------------------ 61762
chimp54546-135457.revcompl.Pan.troglodytes.clone.rp43-41g5.GenBank:AC099533.36   ------------------------------------------------------------ 67152
human.FAM30B-LOC105372935-GGT2.NCBI.                                             aattagggtttttttgaagctccacagatgattcaaatcaggtgtagcaaagcactgtt 64848
human.FAM230E-LOC105377182-GGT3P.NCBI.                                           ------------------------------------------------------------ 57310
human.FAM230J-LOC105372942-GGTLC5P.NCBI.rev.compl                                ------------------------------------------------------------ 46364
human.FAM230A.ncRNA.18487127.18500594-GGTLC3.18516335.18518165.rev.compl.REF     ------------------------------------------------------------ 31039

chimp.LOC112206744-LOC107973052-GGT2.revcompl                                    ------------------------------------------------------------ 61762
chimp54546-135457.revcompl.Pan.troglodytes.clone.rp43-41g5.GenBank:AC099533.36   ------------------------------------------------------------ 67152
human.FAM30B-LOC105372935-GGT2.NCBI.                                             actttaaagtgtctctacctacatggtgccagccaagtgctcaaatgaaatatcttaagg 64908
human.FAM230E-LOC105377182-GGT3P.NCBI.                                           ------------------------------------------------------------ 57310
human.FAM230J-LOC105372942-GGTLC5P.NCBI.rev.compl                                ------------------------------------------------------------ 46364
human.FAM230A.ncRNA.18487127.18500594-GGTLC3.18516335.18518165.rev.compl.REF     ------------------------------------------------------------ 31039

chimp.LOC112206744-LOC107973052-GGT2.revcompl                                    ------------------------------------------------------------ 61762
chimp54546-135457.revcompl.Pan.troglodytes.clone.rp43-41g5.GenBank:AC099533.36   ------------------------------------------------------------ 67152
human.FAM30B-LOC105372935-GGT2.NCBI.                                             ctctcactggctttaagtttctctctttggtagagacccaatcactggttcaaggaagtt 64968
human.FAM230E-LOC105377182-GGT3P.NCBI.                                           ------------------------------------------------------------ 57310
human.FAM230J-LOC105372942-GGTLC5P.NCBI.rev.compl                                ------------------------------------------------------------ 46364
human.FAM230A.ncRNA.18487127.18500594-GGTLC3.18516335.18518165.rev.compl.REF     ------------------------------------------------------------ 31039

chimp.LOC112206744-LOC107973052-GGT2.revcompl                                    ------------------------------------------------------------ 61762
chimp54546-135457.revcompl.Pan.troglodytes.clone.rp43-41g5.GenBank:AC099533.36   ------------------------------------------------------------ 67152
human.FAM30B-LOC105372935-GGT2.NCBI.                                             tcattctcctccagtcttccccagtgcaaaagaaaacagctgagacccatcagatactgg 65028
human.FAM230E-LOC105377182-GGT3P.NCBI.                                           ------------------------------------------------------------ 57310
human.FAM230J-LOC105372942-GGTLC5P.NCBI.rev.compl                                ------------------------------------------------------------ 46364
human.FAM230A.ncRNA.18487127.18500594-GGTLC3.18516335.18518165.rev.compl.REF     ------------------------------------------------------------ 31039

chimp.LOC112206744-LOC107973052-GGT2.revcompl                                    ------------------------------------------------------------ 61762
chimp54546-135457.revcompl.Pan.troglodytes.clone.rp43-41g5.GenBank:AC099533.36   ------------------------------------------------------------ 67152
human.FAM30B-LOC105372935-GGT2.NCBI.                                             cttttgtgatgcaataatgagtttaaccagaatgcatccgatttacagaccttacaaagg 65088
human.FAM230E-LOC105377182-GGT3P.NCBI.                                           ------------------------------------------------------------ 57310
human.FAM230J-LOC105372942-GGTLC5P.NCBI.rev.compl                                ------------------------------------------------------------ 46364
human.FAM230A.ncRNA.18487127.18500594-GGTLC3.18516335.18518165.rev.compl.REF     ------------------------------------------------------------ 31039

chimp.LOC112206744-LOC107973052-GGT2.revcompl                                    ------------------------------------------------------------ 61762
chimp54546-135457.revcompl.Pan.troglodytes.clone.rp43-41g5.GenBank:AC099533.36   ------------------------------------------------------------ 67152
human.FAM30B-LOC105372935-GGT2.NCBI.                                             cactaggccagcggatcatgcacgtcctccccacccaaaggtaaacagtgttaagtggcc 65148
human.FAM230E-LOC105377182-GGT3P.NCBI.                                           ------------------------------------------------------------ 57310
human.FAM230J-LOC105372942-GGTLC5P.NCBI.rev.compl                                ------------------------------------------------------------ 46364
human.FAM230A.ncRNA.18487127.18500594-GGTLC3.18516335.18518165.rev.compl.REF     ------------------------------------------------------------ 31039

chimp.LOC112206744-LOC107973052-GGT2.revcompl                                    ------------------------------------------------------------ 61762
chimp54546-135457.revcompl.Pan.troglodytes.clone.rp43-41g5.GenBank:AC099533.36   ------------------------------------------------------------ 67152
human.FAM30B-LOC105372935-GGT2.NCBI.                                             tgaatgagccaggacagcagggtcaaatcaacttgtctaggctggaagcaggttataaac 65208
human.FAM230E-LOC105377182-GGT3P.NCBI.                                           ------------------------------------------------------------ 57310
human.FAM230J-LOC105372942-GGTLC5P.NCBI.rev.compl                                ------------------------------------------------------------ 46364
human.FAM230A.ncRNA.18487127.18500594-GGTLC3.18516335.18518165.rev.compl.REF     ------------------------------------------------------------ 31039

chimp.LOC112206744-LOC107973052-GGT2.revcompl                                    ------------------------------------------------------------ 61762
chimp54546-135457.revcompl.Pan.troglodytes.clone.rp43-41g5.GenBank:AC099533.36   ------------------------------------------------------------ 67152
human.FAM30B-LOC105372935-GGT2.NCBI.                                             aatccggacaaataaataactactggattgcactttaaccacacacattcagctgcacct 65268
human.FAM230E-LOC105377182-GGT3P.NCBI.                                           ------------------------------------------------------------ 57310
human.FAM230J-LOC105372942-GGTLC5P.NCBI.rev.compl                                ------------------------------------------------------------ 46364
human.FAM230A.ncRNA.18487127.18500594-GGTLC3.18516335.18518165.rev.compl.REF     ------------------------------------------------------------ 31039

chimp.LOC112206744-LOC107973052-GGT2.revcompl                                    ------------------------------------------------------------ 61762
chimp54546-135457.revcompl.Pan.troglodytes.clone.rp43-41g5.GenBank:AC099533.36   ------------------------------------------------------------ 67152
human.FAM30B-LOC105372935-GGT2.NCBI.                                             gtttattaaatacctccttaatttccctctgccccacaaagggcttctgaccccctgaaaa 65328
human.FAM230E-LOC105377182-GGT3P.NCBI.                                           ------------------------------------------------------------ 57310
human.FAM230J-LOC105372942-GGTLC5P.NCBI.rev.compl                                ------------------------------------------------------------ 46364
human.FAM230A.ncRNA.18487127.18500594-GGTLC3.18516335.18518165.rev.compl.REF     ------------------------------------------------------------ 31039

chimp.LOC112206744-LOC107973052-GGT2.revcompl                                    ------------------------------------------------------------ 61762
chimp54546-135457.revcompl.Pan.troglodytes.clone.rp43-41g5.GenBank:AC099533.36   ------------------------------------------------------------ 67152
human.FAM30B-LOC105372935-GGT2.NCBI.                                             taatgcttctcaacataaaaagaattgttttcttcttggcagtttaattccccttccaca 65388
human.FAM230E-LOC105377182-GGT3P.NCBI.                                           ------------------------------------------------------------ 57310
human.FAM230J-LOC105372942-GGTLC5P.NCBI.rev.compl                                ------------------------------------------------------------ 46364
human.FAM230A.ncRNA.18487127.18500594-GGTLC3.18516335.18518165.rev.compl.REF     ------------------------------------------------------------ 31039

chimp.LOC112206744-LOC107973052-GGT2.revcompl                                    ------------------------------------------------------------ 61762
chimp54546-135457.revcompl.Pan.troglodytes.clone.rp43-41g5.GenBank:AC099533.36   ------------------------------------------------------------ 67152
human.FAM30B-LOC105372935-GGT2.NCBI.                                             catcccaccccaccatttccctataatggtaataccccgctgtgcagacctctgctgcct 65448
human.FAM230E-LOC105377182-GGT3P.NCBI.                                           ------------------------------------------------------------ 57310
human.FAM230J-LOC105372942-GGTLC5P.NCBI.rev.compl                                ------------------------------------------------------------ 46364
human.FAM230A.ncRNA.18487127.18500594-GGTLC3.18516335.18518165.rev.compl.REF     ------------------------------------------------------------ 31039

chimp.LOC112206744-LOC107973052-GGT2.revcompl                                    ------------------------------------------------------------ 61762
chimp54546-135457.revcompl.Pan.troglodytes.clone.rp43-41g5.GenBank:AC099533.36   ------------------------------------------------------------ 67152
human.FAM30B-LOC105372935-GGT2.NCBI.                                             ccagagagacacaggcgcccaaccctgaccagggcatcctctgacccacagcccctctat 65508
human.FAM230E-LOC105377182-GGT3P.NCBI.                                           ------------------------------------------------------------ 57310
human.FAM230J-LOC105372942-GGTLC5P.NCBI.rev.compl                                ------------------------------------------------------------ 46364
human.FAM230A.ncRNA.18487127.18500594-GGTLC3.18516335.18518165.rev.compl.REF     ------------------------------------------------------------ 31039
```

```
chimp.LOC112206744-LOC107973052-GGT2.revcompl                                   ------------------------------------------------------------ 61762
chimp54546-135457.revcompl.Pan.troglodytes.clone.rp43-41g5.GenBank:AC099533.36  ------------------------------------------------------------ 67152
human.FAM30B-LOC105372935-GGT2.NCBI.                                            ctcccctctcagcctacaatagaaagttcctcccaggcaaggatcatttttttaaataa 65568
human.FAM230E-LOC105377182-GGT3P.NCBI.                                          ------------------------------------------------------------ 57310
human.FAM230J-LOC105372942-GGTLC5P.NCBI.rev.compl                               ------------------------------------------------------------ 46364
human.FAM230A.ncRNA.18487127.18500594-GGTLC3.18516335.18518165.rev.compl.REF    ------------------------------------------------------------ 31039

chimp.LOC112206744-LOC107973052-GGT2.revcompl                                   ------------------------------------------------------------ 61762
chimp54546-135457.revcompl.Pan.troglodytes.clone.rp43-41g5.GenBank:AC099533.36  ------------------------------------------------------------ 67152
human.FAM30B-LOC105372935-GGT2.NCBI.                                            ctttttaactttagacattctgtactctttggattactctgcaataagcaaatatgact 65628
human.FAM230E-LOC105377182-GGT3P.NCBI.                                          ------------------------------------------------------------ 57310
human.FAM230J-LOC105372942-GGTLC5P.NCBI.rev.compl                               ------------------------------------------------------------ 46364
human.FAM230A.ncRNA.18487127.18500594-GGTLC3.18516335.18518165.rev.compl.REF    ------------------------------------------------------------ 31039

chimp.LOC112206744-LOC107973052-GGT2.revcompl                                   ------------------------------------------------------------ 61762
chimp54546-135457.revcompl.Pan.troglodytes.clone.rp43-41g5.GenBank:AC099533.36  ------------------------------------------------------------ 67152
human.FAM30B-LOC105372935-GGT2.NCBI.                                            tctgtaacataaaaagaaagatcaaaatgttatatataactgcatgaaaagaactagaaa 65688
human.FAM230E-LOC105377182-GGT3P.NCBI.                                          ------------------------------------------------------------ 57310
human.FAM230J-LOC105372942-GGTLC5P.NCBI.rev.compl                               ------------------------------------------------------------ 46364
human.FAM230A.ncRNA.18487127.18500594-GGTLC3.18516335.18518165.rev.compl.REF    ------------------------------------------------------------ 31039

chimp.LOC112206744-LOC107973052-GGT2.revcompl                                   ------------------------------------------------------------ 61762
chimp54546-135457.revcompl.Pan.troglodytes.clone.rp43-41g5.GenBank:AC099533.36  ------------------------------------------------------------ 67152
human.FAM30B-LOC105372935-GGT2.NCBI.                                            gaaaatactaggtgacaggatggcaagtgattttatttttttcttaatattttactcct 65748
human.FAM230E-LOC105377182-GGT3P.NCBI.                                          ------------------------------------------------------------ 57310
human.FAM230J-LOC105372942-GGTLC5P.NCBI.rev.compl                               ------------------------------------------------------------ 46364
human.FAM230A.ncRNA.18487127.18500594-GGTLC3.18516335.18518165.rev.compl.REF    ------------------------------------------------------------ 31039

chimp.LOC112206744-LOC107973052-GGT2.revcompl                                   ------------------------------------------------------------ 61762
chimp54546-135457.revcompl.Pan.troglodytes.clone.rp43-41g5.GenBank:AC099533.36  ------------------------------------------------------------ 67152
human.FAM30B-LOC105372935-GGT2.NCBI.                                            tcccccagtcctctgcaatgagtatgtaccagttttataactagaaaaaaatttagcca 65808
human.FAM230E-LOC105377182-GGT3P.NCBI.                                          ------------------------------------------------------------ 57310
human.FAM230J-LOC105372942-GGTLC5P.NCBI.rev.compl                               ------------------------------------------------------------ 46364
human.FAM230A.ncRNA.18487127.18500594-GGTLC3.18516335.18518165.rev.compl.REF    ------------------------------------------------------------ 31039

chimp.LOC112206744-LOC107973052-GGT2.revcompl                                   ------------------------------------------------------------ 61762
chimp54546-135457.revcompl.Pan.troglodytes.clone.rp43-41g5.GenBank:AC099533.36  ------------------------------------------------------------ 67152
human.FAM30B-LOC105372935-GGT2.NCBI.                                            aaaagaaaaataatgcatgtttgctatataaaattcactactgtagtatatatatataca 65868
human.FAM230E-LOC105377182-GGT3P.NCBI.                                          ------------------------------------------------------------ 57310
human.FAM230J-LOC105372942-GGTLC5P.NCBI.rev.compl                               ------------------------------------------------------------ 46364
human.FAM230A.ncRNA.18487127.18500594-GGTLC3.18516335.18518165.rev.compl.REF    ------------------------------------------------------------ 31039

chimp.LOC112206744-LOC107973052-GGT2.revcompl                                   ------------------------------------------------------------ 61762
chimp54546-135457.revcompl.Pan.troglodytes.clone.rp43-41g5.GenBank:AC099533.36  ------------------------------------------------------------ 67152
human.FAM30B-LOC105372935-GGT2.NCBI.                                            tacatacatatttggtctttgtccccagttcctggcactgagctcctaaaccccttggaa 65928
human.FAM230E-LOC105377182-GGT3P.NCBI.                                          ------------------------------------------------------------ 57310
human.FAM230J-LOC105372942-GGTLC5P.NCBI.rev.compl                               ------------------------------------------------------------ 46364
human.FAM230A.ncRNA.18487127.18500594-GGTLC3.18516335.18518165.rev.compl.REF    ------------------------------------------------------------ 31039

chimp.LOC112206744-LOC107973052-GGT2.revcompl                                   ------------------------------------------------------------ 61762
chimp54546-135457.revcompl.Pan.troglodytes.clone.rp43-41g5.GenBank:AC099533.36  ------------------------------------------------------------ 67152
human.FAM30B-LOC105372935-GGT2.NCBI.                                            cttcctaagcaatgggagtaccttttgttatttataagaagccccttttggccatcccag 65988
human.FAM230E-LOC105377182-GGT3P.NCBI.                                          ------------------------------------------------------------ 57310
human.FAM230J-LOC105372942-GGTLC5P.NCBI.rev.compl                               ------------------------------------------------------------ 46364
human.FAM230A.ncRNA.18487127.18500594-GGTLC3.18516335.18518165.rev.compl.REF    ------------------------------------------------------------ 31039

chimp.LOC112206744-LOC107973052-GGT2.revcompl                                   ------------------------------------------------------------ 61762
chimp54546-135457.revcompl.Pan.troglodytes.clone.rp43-41g5.GenBank:AC099533.36  ------------------------------------------------------------ 67152
human.FAM30B-LOC105372935-GGT2.NCBI.                                            agtttatgctaacgaggtgactgaaggtgagcacggaggcagtttcaaggaaggagctgg 66048
human.FAM230E-LOC105377182-GGT3P.NCBI.                                          ------------------------------------------------------------ 57310
human.FAM230J-LOC105372942-GGTLC5P.NCBI.rev.compl                               ------------------------------------------------------------ 46364
human.FAM230A.ncRNA.18487127.18500594-GGTLC3.18516335.18518165.rev.compl.REF    ------------------------------------------------------------ 31039

chimp.LOC112206744-LOC107973052-GGT2.revcompl                                   ------------------------------------------------------------ 61762
chimp54546-135457.revcompl.Pan.troglodytes.clone.rp43-41g5.GenBank:AC099533.36  ------------------------------------------------------------ 67152
human.FAM30B-LOC105372935-GGT2.NCBI.                                            ccacgcttagaattgtggagttttcagccccgcccttagacctccaaggacaagaggggg 66108
human.FAM230E-LOC105377182-GGT3P.NCBI.                                          ------------------------------------------------------------ 57310
human.FAM230J-LOC105372942-GGTLC5P.NCBI.rev.compl                               ------------------------------------------------------------ 46364
human.FAM230A.ncRNA.18487127.18500594-GGTLC3.18516335.18518165.rev.compl.REF    ------------------------------------------------------------ 31039

chimp.LOC112206744-LOC107973052-GGT2.revcompl                                   ------------------------------------------------------------ 61762
chimp54546-135457.revcompl.Pan.troglodytes.clone.rp43-41g5.GenBank:AC099533.36  ------------------------------------------------------------ 67152
human.FAM30B-LOC105372935-GGT2.NCBI.                                            accgcagattgatccaatcagcattggtcagtgatttgatcagtcatacccacgtaatga 66168
human.FAM230E-LOC105377182-GGT3P.NCBI.                                          ------------------------------------------------------------ 57310
human.FAM230J-LOC105372942-GGTLC5P.NCBI.rev.compl                               ------------------------------------------------------------ 46364
human.FAM230A.ncRNA.18487127.18500594-GGTLC3.18516335.18518165.rev.compl.REF    ------------------------------------------------------------ 31039

chimp.LOC112206744-LOC107973052-GGT2.revcompl                                   ------------------------------------------------------------ 61762
chimp54546-135457.revcompl.Pan.troglodytes.clone.rp43-41g5.GenBank:AC099533.36  ------------------------------------------------------------ 67152
human.FAM30B-LOC105372935-GGT2.NCBI.                                            aatcccatatgaaaaccctaaataatggagttaggagagcttctgggtgctggaagcggc 66228
human.FAM230E-LOC105377182-GGT3P.NCBI.                                          ------------------------------------------------------------ 57310
human.FAM230J-LOC105372942-GGTLC5P.NCBI.rev.compl                               ------------------------------------------------------------ 46364
human.FAM230A.ncRNA.18487127.18500594-GGTLC3.18516335.18518165.rev.compl.REF    ------------------------------------------------------------ 31039

chimp.LOC112206744-LOC107973052-GGT2.revcompl                                   ------------------------------------------------------------ 61762
chimp54546-135457.revcompl.Pan.troglodytes.clone.rp43-41g5.GenBank:AC099533.36  ------------------------------------------------------------ 67152
human.FAM30B-LOC105372935-GGT2.NCBI.                                            gtgcaccaggagagggcatgggcagtgagcactactcccctctcagaccttgccctatgc 66288
human.FAM230E-LOC105377182-GGT3P.NCBI.                                          ------------------------------------------------------------ 57310
human.FAM230J-LOC105372942-GGTLC5P.NCBI.rev.compl                               ------------------------------------------------------------ 46364
human.FAM230A.ncRNA.18487127.18500594-GGTLC3.18516335.18518165.rev.compl.REF    ------------------------------------------------------------ 31039

chimp.LOC112206744-LOC107973052-GGT2.revcompl                                   ------------------------------------------------------------ 61762
chimp54546-135457.revcompl.Pan.troglodytes.clone.rp43-41g5.GenBank:AC099533.36  ------------------------------------------------------------ 67152
human.FAM30B-LOC105372935-GGT2.NCBI.                                            actaaataggactgacctatgtgaccaataggaaatgcacaaatggtggaatgtgacttc 66348
human.FAM230E-LOC105377182-GGT3P.NCBI.                                          ------------------------------------------------------------ 57310
human.FAM230J-LOC105372942-GGTLC5P.NCBI.rev.compl                               ------------------------------------------------------------ 46364
human.FAM230A.ncRNA.18487127.18500594-GGTLC3.18516335.18518165.rev.compl.REF    ------------------------------------------------------------ 31039

chimp.LOC112206744-LOC107973052-GGT2.revcompl                                   ------------------------------------------------------------ 61762
chimp54546-135457.revcompl.Pan.troglodytes.clone.rp43-41g5.GenBank:AC099533.36  ------------------------------------------------------------ 67152
human.FAM30B-LOC105372935-GGT2.NCBI.                                            cagggctaggtcataaaaagacaagcacgatgttatgaggacatcaaagcagccatatgg 66408
human.FAM230E-LOC105377182-GGT3P.NCBI.                                          ------------------------------------------------------------ 57310
human.FAM230J-LOC105372942-GGTLC5P.NCBI.rev.compl                               ------------------------------------------------------------ 46364
human.FAM230A.ncRNA.18487127.18500594-GGTLC3.18516335.18518165.rev.compl.REF    ------------------------------------------------------------ 31039

chimp.LOC112206744-LOC107973052-GGT2.revcompl                                   ------------------------------------------------------------ 61762
chimp54546-135457.revcompl.Pan.troglodytes.clone.rp43-41g5.GenBank:AC099533.36  ------------------------------------------------------------ 67152
human.FAM30B-LOC105372935-GGT2.NCBI.                                            ggaggaccccatgaggcctcctgccagcagctagcactaacttggctagcatgtgactgg 66468
human.FAM230E-LOC105377182-GGT3P.NCBI.                                          ------------------------------------------------------------ 57310
human.FAM230J-LOC105372942-GGTLC5P.NCBI.rev.compl                               ------------------------------------------------------------ 46364
human.FAM230A.ncRNA.18487127.18500594-GGTLC3.18516335.18518165.rev.compl.REF    ------------------------------------------------------------ 31039

chimp.LOC112206744-LOC107973052-GGT2.revcompl                                   ------------------------------------------------------------ 61762
chimp54546-135457.revcompl.Pan.troglodytes.clone.rp43-41g5.GenBank:AC099533.36  -----------------------------------------------------------T 67153
human.FAM30B-LOC105372935-GGT2.NCBI.                                            aagtagattctccagcctcagtcaactcagtcaagccttcagataatgtcaaccccaggc 66528
human.FAM230E-LOC105377182-GGT3P.NCBI.                                          ------------------------------------------------------------ 57310
human.FAM230J-LOC105372942-GGTLC5P.NCBI.rev.compl                               ------------------------------------------------------------ 46364
human.FAM230A.ncRNA.18487127.18500594-GGTLC3.18516335.18518165.rev.compl.REF    ------------------------------------------------------------ 31039

chimp.LOC112206744-LOC107973052-GGT2.revcompl                                   ------------------------------------------------------------ 61762
chimp54546-135457.revcompl.Pan.troglodytes.clone.rp43-41g5.GenBank:AC099533.36  TTATTTATTTATTTATTGAGACAGGGTCTCACTCTGTTGCCCAGGCTAGAGTGCAGTGGC 67213
human.FAM30B-LOC105372935-GGT2.NCBI.                                            atctttttgttgttgttgagacagggtctcactctgtcacccaggctggagtgcagtggt 66588
human.FAM230E-LOC105377182-GGT3P.NCBI.                                          ------------------------------------------------------------ 57310
human.FAM230J-LOC105372942-GGTLC5P.NCBI.rev.compl                               ------------------------------------------------------------ 46364
human.FAM230A.ncRNA.18487127.18500594-GGTLC3.18516335.18518165.rev.compl.REF    ------------------------------------------------------------ 31039

chimp.LOC112206744-LOC107973052-GGT2.revcompl                                   ------------------------------------------------------------ 61762
```

```
chimp54546-135457.revcompl.Pan.troglodytes.clone.rp43-41g5.GenBank:AC099533.36       ATGATCTCGGCTCACTACAACTTCCACCTCCCGGGTTCAAGCAATTGTCCTGCCTCAGCC 67273
human.FAM30B-LOC105372935-GGT2.NCBI.                                                  gcaatcatagctcactgcagccttgacctctgaggctcaagtgaccctcccacatcagcc 66648
human.FAM230E-LOC105377182-GGT3P.NCBI.                                                ------------------------------------------------------------ 57310
human.FAM230J-LOC105372942-GGTLC5P.NCBI.rev.compl                                     ------------------------------------------------------------ 46364
human.FAM230A.ncRNA.18487127.18500594-GGTLC3.18516335.18518165.rev.compl.REF          ------------------------------------------------------------ 31039

chimp.LOC112206744-LOC107973052-GGT2.revcompl                                         ------------------------------------------------------------ 61762
chimp54546-135457.revcompl.Pan.troglodytes.clone.rp43-41g5.GenBank:AC099533.36       TCCCAAGTAGCTGGGATTACAGGCACCCACCACCACGCCCGGCTAATTGGGAGATAGTTT 67333
human.FAM30B-LOC105372935-GGT2.NCBI.                                                  tcctgagtatctgggaccaccagcacacaccaccatgcctggctaactttttatttctt 66708
human.FAM230E-LOC105377182-GGT3P.NCBI.                                                ------------------------------------------------------------ 57310
human.FAM230J-LOC105372942-GGTLC5P.NCBI.rev.compl                                     ------------------------------------------------------------ 46364
human.FAM230A.ncRNA.18487127.18500594-GGTLC3.18516335.18518165.rev.compl.REF          ------------------------------------------------------------ 31039

chimp.LOC112206744-LOC107973052-GGT2.revcompl                                         ------------------------------------------------------------ 61762
chimp54546-135457.revcompl.Pan.troglodytes.clone.rp43-41g5.GenBank:AC099533.36       G----------------------------------------------------------- 67334
human.FAM30B-LOC105372935-GGT2.NCBI.                                                  atagagatgggatgatatggttggggtctgtgtacccacctaaatctcacgttcaattat 66768
human.FAM230E-LOC105377182-GGT3P.NCBI.                                                ------------------------------------------------------------ 57310
human.FAM230J-LOC105372942-GGTLC5P.NCBI.rev.compl                                     ------------------------------------------------------------ 46364
human.FAM230A.ncRNA.18487127.18500594-GGTLC3.18516335.18518165.rev.compl.REF          ------------------------------------------------------------ 31039

chimp.LOC112206744-LOC107973052-GGT2.revcompl                                         ------------------------------------------------------------ 61762
chimp54546-135457.revcompl.Pan.troglodytes.clone.rp43-41g5.GenBank:AC099533.36       ------------------------------------------------------------ 67334
human.FAM30B-LOC105372935-GGT2.NCBI.                                                  aatccccaatgttagaggtggggcctggtgggaggtgactggatcacaggaatggatctt 66828
human.FAM230E-LOC105377182-GGT3P.NCBI.                                                ------------------------------------------------------------ 57310
human.FAM230J-LOC105372942-GGTLC5P.NCBI.rev.compl                                     ------------------------------------------------------------ 46364
human.FAM230A.ncRNA.18487127.18500594-GGTLC3.18516335.18518165.rev.compl.REF          ------------------------------------------------------------ 31039

chimp.LOC112206744-LOC107973052-GGT2.revcompl                                         ------------------------------------------------------------ 61762
chimp54546-135457.revcompl.Pan.troglodytes.clone.rp43-41g5.GenBank:AC099533.36       ------------------------------------------------------------ 67334
human.FAM30B-LOC105372935-GGT2.NCBI.                                                  tcatgaatgatttagcatcaccctttggtgctgtgcttgttagagttctcccagtatct 66888
human.FAM230E-LOC105377182-GGT3P.NCBI.                                                ------------------------------------------------------------ 57310
human.FAM230J-LOC105372942-GGTLC5P.NCBI.rev.compl                                     ------------------------------------------------------------ 46364
human.FAM230A.ncRNA.18487127.18500594-GGTLC3.18516335.18518165.rev.compl.REF          ------------------------------------------------------------ 31039

chimp.LOC112206744-LOC107973052-GGT2.revcompl                                         ------------------------------------------------------------ 61762
chimp54546-135457.revcompl.Pan.troglodytes.clone.rp43-41g5.GenBank:AC099533.36       ------------------------------------------------------------ 67334
human.FAM30B-LOC105372935-GGT2.NCBI.                                                  aattgtttaaaagtgtgtggcacctccccctctctctcttgctcctgctttggccatgt 66948
human.FAM230E-LOC105377182-GGT3P.NCBI.                                                ------------------------------------------------------------ 57310
human.FAM230J-LOC105372942-GGTLC5P.NCBI.rev.compl                                     ------------------------------------------------------------ 46364
human.FAM230A.ncRNA.18487127.18500594-GGTLC3.18516335.18518165.rev.compl.REF          ------------------------------------------------------------ 31039

chimp.LOC112206744-LOC107973052-GGT2.revcompl                                         ------------------------------------------------------------ 61762
chimp54546-135457.revcompl.Pan.troglodytes.clone.rp43-41g5.GenBank:AC099533.36       ------------------------------------------------------------ 67334
human.FAM30B-LOC105372935-GGT2.NCBI.                                                  aaggcgtgcctgcttccccttcaccttccaccatgattgaaagtttcctgtgcttgcccc 67008
human.FAM230E-LOC105377182-GGT3P.NCBI.                                                ------------------------------------------------------------ 57310
human.FAM230J-LOC105372942-GGTLC5P.NCBI.rev.compl                                     ------------------------------------------------------------ 46364
human.FAM230A.ncRNA.18487127.18500594-GGTLC3.18516335.18518165.rev.compl.REF          ------------------------------------------------------------ 31039

chimp.LOC112206744-LOC107973052-GGT2.revcompl                                         ------------------------------------------------------------ 61762
chimp54546-135457.revcompl.Pan.troglodytes.clone.rp43-41g5.GenBank:AC099533.36       ------------------------------------------------------------ 67334
human.FAM30B-LOC105372935-GGT2.NCBI.                                                  agaagctgagcagatgccagcattatgctccctgaacagcctgtggaactgcgagacaat 67068
human.FAM230E-LOC105377182-GGT3P.NCBI.                                                ------------------------------------------------------------ 57310
human.FAM230J-LOC105372942-GGTLC5P.NCBI.rev.compl                                     ------------------------------------------------------------ 46364
human.FAM230A.ncRNA.18487127.18500594-GGTLC3.18516335.18518165.rev.compl.REF          ------------------------------------------------------------ 31039

chimp.LOC112206744-LOC107973052-GGT2.revcompl                                         ------------------------------------------------------------ 61762
chimp54546-135457.revcompl.Pan.troglodytes.clone.rp43-41g5.GenBank:AC099533.36       ------------------------------------------------------------ 67334
human.FAM30B-LOC105372935-GGT2.NCBI.                                                  taaacctcttttctttataaattctccagtctcatgtatttatagcaatgtgagaactga 67128
human.FAM230E-LOC105377182-GGT3P.NCBI.                                                ------------------------------------------------------------ 57310
human.FAM230J-LOC105372942-GGTLC5P.NCBI.rev.compl                                     ------------------------------------------------------------ 46364
human.FAM230A.ncRNA.18487127.18500594-GGTLC3.18516335.18518165.rev.compl.REF          ------------------------------------------------------------ 31039

chimp.LOC112206744-LOC107973052-GGT2.revcompl                                         ------------------------------------------------------------ 61762
chimp54546-135457.revcompl.Pan.troglodytes.clone.rp43-41g5.GenBank:AC099533.36       ------------------------------------------------------------ 67334
human.FAM30B-LOC105372935-GGT2.NCBI.                                                  ctaatacatagggtctcactgtgtttcccaggctggtctcaaactcctgggctcaagtaa 67188
human.FAM230E-LOC105377182-GGT3P.NCBI.                                                ------------------------------------------------------------ 57310
human.FAM230J-LOC105372942-GGTLC5P.NCBI.rev.compl                                     ------------------------------------------------------------ 46364
human.FAM230A.ncRNA.18487127.18500594-GGTLC3.18516335.18518165.rev.compl.REF          ------------------------------------------------------------ 31039

chimp.LOC112206744-LOC107973052-GGT2.revcompl                                         ------------------------------------------------------------ 61762
chimp54546-135457.revcompl.Pan.troglodytes.clone.rp43-41g5.GenBank:AC099533.36       ------------------------------------------------------------ 67334
human.FAM30B-LOC105372935-GGT2.NCBI.                                                  tcctcctggctttgcttcccagagtgtagagattacaggcatgagccactgaacctgccc 67248
human.FAM230E-LOC105377182-GGT3P.NCBI.                                                ------------------------------------------------------------ 57310
human.FAM230J-LOC105372942-GGTLC5P.NCBI.rev.compl                                     ------------------------------------------------------------ 46364
human.FAM230A.ncRNA.18487127.18500594-GGTLC3.18516335.18518165.rev.compl.REF          ------------------------------------------------------------ 31039

chimp.LOC112206744-LOC107973052-GGT2.revcompl                                         ------------------------------------------------------------ 61762
chimp54546-135457.revcompl.Pan.troglodytes.clone.rp43-41g5.GenBank:AC099533.36       ------------------------------------------------------------ 67334
human.FAM30B-LOC105372935-GGT2.NCBI.                                                  ctttcccccaacaacatcttgattgcaacctcctaagagactccgagtcagaagtatcc 67308
human.FAM230E-LOC105377182-GGT3P.NCBI.                                                ------------------------------------------------------------ 57310
human.FAM230J-LOC105372942-GGTLC5P.NCBI.rev.compl                                     ------------------------------------------------------------ 46364
human.FAM230A.ncRNA.18487127.18500594-GGTLC3.18516335.18518165.rev.compl.REF          ------------------------------------------------------------ 31039

chimp.LOC112206744-LOC107973052-GGT2.revcompl                                         ------------------------------------------------------------ 61762
chimp54546-135457.revcompl.Pan.troglodytes.clone.rp43-41g5.GenBank:AC099533.36       ------------------------------------------------------------ 67334
human.FAM30B-LOC105372935-GGT2.NCBI.                                                  agttaagatgttactgactttctgaactacagaaactgtgagataataaacatctattgt 67368
human.FAM230E-LOC105377182-GGT3P.NCBI.                                                ------------------------------------------------------------ 57310
human.FAM230J-LOC105372942-GGTLC5P.NCBI.rev.compl                                     ------------------------------------------------------------ 46364
human.FAM230A.ncRNA.18487127.18500594-GGTLC3.18516335.18518165.rev.compl.REF          ------------------------------------------------------------ 31039

chimp.LOC112206744-LOC107973052-GGT2.revcompl                                         ------------------------------------------------------------ 61762
chimp54546-135457.revcompl.Pan.troglodytes.clone.rp43-41g5.GenBank:AC099533.36       ------------------------------------------------------------ 67334
human.FAM30B-LOC105372935-GGT2.NCBI.                                                  tcctttaagccttaagtttggggcttttttttttacaccaatatataactaatgtagac 67428
human.FAM230E-LOC105377182-GGT3P.NCBI.                                                ------------------------------------------------------------ 57310
human.FAM230J-LOC105372942-GGTLC5P.NCBI.rev.compl                                     ------------------------------------------------------------ 46364
human.FAM230A.ncRNA.18487127.18500594-GGTLC3.18516335.18518165.rev.compl.REF          ------------------------------------------------------------ 31039

chimp.LOC112206744-LOC107973052-GGT2.revcompl                                         ------------------------------------------------------------ 61762
chimp54546-135457.revcompl.Pan.troglodytes.clone.rp43-41g5.GenBank:AC099533.36       ------------------------------------------------------------ 67334
human.FAM30B-LOC105372935-GGT2.NCBI.                                                  acctgcagtccctaagattgaatgcaaaattgtgttacatacacatgagtgtgttcttct 67488
human.FAM230E-LOC105377182-GGT3P.NCBI.                                                ------------------------------------------------------------ 57310
human.FAM230J-LOC105372942-GGTLC5P.NCBI.rev.compl                                     ------------------------------------------------------------ 46364
human.FAM230A.ncRNA.18487127.18500594-GGTLC3.18516335.18518165.rev.compl.REF          ------------------------------------------------------------ 31039

chimp.LOC112206744-LOC107973052-GGT2.revcompl                                         ------------------------------------------------------------ 61762
chimp54546-135457.revcompl.Pan.troglodytes.clone.rp43-41g5.GenBank:AC099533.36       ------------------------------------------------------------ 67334
human.FAM30B-LOC105372935-GGT2.NCBI.                                                  gggaaaagagtccataaacttcaaaagattctcaaaggaatccaagacccaaaaagatta 67548
human.FAM230E-LOC105377182-GGT3P.NCBI.                                                ------------------------------------------------------------ 57310
human.FAM230J-LOC105372942-GGTLC5P.NCBI.rev.compl                                     ------------------------------------------------------------ 46364
human.FAM230A.ncRNA.18487127.18500594-GGTLC3.18516335.18518165.rev.compl.REF          ------------------------------------------------------------ 31039

chimp.LOC112206744-LOC107973052-GGT2.revcompl                                         ------------------------------------------------------------ 61762
chimp54546-135457.revcompl.Pan.troglodytes.clone.rp43-41g5.GenBank:AC099533.36       ------------------------------------------------------------ 67334
human.FAM30B-LOC105372935-GGT2.NCBI.                                                  agaaacattgcatcagacaataaattgtcaatgagcaaggtcctgtccttttagattggg 67608
human.FAM230E-LOC105377182-GGT3P.NCBI.                                                ------------------------------------------------------------ 57310
human.FAM230J-LOC105372942-GGTLC5P.NCBI.rev.compl                                     ------------------------------------------------------------ 46364
human.FAM230A.ncRNA.18487127.18500594-GGTLC3.18516335.18518165.rev.compl.REF          ------------------------------------------------------------ 31039

chimp.LOC112206744-LOC107973052-GGT2.revcompl                                         ------------------------------------------------------------ 61762
chimp54546-135457.revcompl.Pan.troglodytes.clone.rp43-41g5.GenBank:AC099533.36       ------------------------------------------------------------ 67334
human.FAM30B-LOC105372935-GGT2.NCBI.                                                  ttttgaaatcacccatcccacaaatatatgataccacttaactagtttttcaagatattc 67668
human.FAM230E-LOC105377182-GGT3P.NCBI.                                                ------------------------------------------------------------ 57310
human.FAM230J-LOC105372942-GGTLC5P.NCBI.rev.compl                                     ------------------------------------------------------------ 46364
human.FAM230A.ncRNA.18487127.18500594-GGTLC3.18516335.18518165.rev.compl.REF          ------------------------------------------------------------ 31039

chimp.LOC112206744-LOC107973052-GGT2.revcompl                                         ------------------------------------------------------------ 61762
chimp54546-135457.revcompl.Pan.troglodytes.clone.rp43-41g5.GenBank:AC099533.36       ------------------------------------------------------------ 67334
```

```
human.FAM30B-LOC105372935-GGT2.NCBI.    cttgttcaaagatgctcccactttttctttgtgctcaagtcctagataaacttggctagg 67728
human.FAM230E-LOC105377182-GGT3P.NCBI.    ------------------------------------------------------------ 57310
human.FAM230J-LOC105372942-GGTLC5P.NCBI.rev.compl    ------------------------------------------------------------ 46364
human.FAM230A.ncRNA.18487127.18500594-GGTLC3.18516335.18518165.rev.compl.REF    ------------------------------------------------------------ 31039

chimp.LOC112206744-LOC107973052-GGT2.revcompl    ------------------------------------------------------------ 61762
chimp54546-135457.revcompl.Pan.troglodytes.clone.rp43-41g5.GenBank:AC099533.36    ------------------------------------------------------------ 67334
human.FAM30B-LOC105372935-GGT2.NCBI.    ggcaggaggtatgattcaggagttagtgagccgggagggctcactgtgcctgggaaagcc 67788
human.FAM230E-LOC105377182-GGT3P.NCBI.    ------------------------------------------------------------ 57310
human.FAM230J-LOC105372942-GGTLC5P.NCBI.rev.compl    ------------------------------------------------------------ 46364
human.FAM230A.ncRNA.18487127.18500594-GGTLC3.18516335.18518165.rev.compl.REF    ------------------------------------------------------------ 31039

chimp.LOC112206744-LOC107973052-GGT2.revcompl    ------------------------------------------------------------ 61762
chimp54546-135457.revcompl.Pan.troglodytes.clone.rp43-41g5.GenBank:AC099533.36    ------------------------------------------------------------ 67334
human.FAM30B-LOC105372935-GGT2.NCBI.    agcttctcatgaatttttggaatatgccttggctcagaaacctacctctctaggcacagg 67848
human.FAM230E-LOC105377182-GGT3P.NCBI.    ------------------------------------------------------------ 57310
human.FAM230J-LOC105372942-GGTLC5P.NCBI.rev.compl    ------------------------------------------------------------ 46364
human.FAM230A.ncRNA.18487127.18500594-GGTLC3.18516335.18518165.rev.compl.REF    ------------------------------------------------------------ 31039

chimp.LOC112206744-LOC107973052-GGT2.revcompl    ------------------------------------------------------------ 61762
chimp54546-135457.revcompl.Pan.troglodytes.clone.rp43-41g5.GenBank:AC099533.36    ------------------------------------------------------------ 67334
human.FAM30B-LOC105372935-GGT2.NCBI.    atccaacttatacgttcaagatgcgtgactgagcatgtgaaagtgtcagagggatcgacc 67908
human.FAM230E-LOC105377182-GGT3P.NCBI.    ------------------------------------------------------------ 57310
human.FAM230J-LOC105372942-GGTLC5P.NCBI.rev.compl    ------------------------------------------------------------ 46364
human.FAM230A.ncRNA.18487127.18500594-GGTLC3.18516335.18518165.rev.compl.REF    ------------------------------------------------------------ 31039

chimp.LOC112206744-LOC107973052-GGT2.revcompl    ------------------------------------------------------------ 61762
chimp54546-135457.revcompl.Pan.troglodytes.clone.rp43-41g5.GenBank:AC099533.36    ------------------------------------------------------------ 67334
human.FAM30B-LOC105372935-GGT2.NCBI.    tcaccattttacatctacttttccagttgtggcccctacaatcatcaaggctcattcggc 67968
human.FAM230E-LOC105377182-GGT3P.NCBI.    ------------------------------------------------------------ 57310
human.FAM230J-LOC105372942-GGTLC5P.NCBI.rev.compl    ------------------------------------------------------------ 46364
human.FAM230A.ncRNA.18487127.18500594-GGTLC3.18516335.18518165.rev.compl.REF    ------------------------------------------------------------ 31039

chimp.LOC112206744-LOC107973052-GGT2.revcompl    ------------------------------------------------------------ 61762
chimp54546-135457.revcompl.Pan.troglodytes.clone.rp43-41g5.GenBank:AC099533.36    ------------------------------------------------------------ 67334
human.FAM30B-LOC105372935-GGT2.NCBI.    accatgctcctccaaattctgcatgcgactcagccacactaatatctttcttcatctata 68028
human.FAM230E-LOC105377182-GGT3P.NCBI.    ------------------------------------------------------------ 57310
human.FAM230J-LOC105372942-GGTLC5P.NCBI.rev.compl    ------------------------------------------------------------ 46364
human.FAM230A.ncRNA.18487127.18500594-GGTLC3.18516335.18518165.rev.compl.REF    ------------------------------------------------------------ 31039

chimp.LOC112206744-LOC107973052-GGT2.revcompl    ------------------------------------------------------------ 61762
chimp54546-135457.revcompl.Pan.troglodytes.clone.rp43-41g5.GenBank:AC099533.36    ------------------------------------------------------------ 67334
human.FAM30B-LOC105372935-GGT2.NCBI.    agatgaaagtaatacaacgcaccatgtagcaaactcacaggagaccctagagatcaaata 68088
human.FAM230E-LOC105377182-GGT3P.NCBI.    ------------------------------------------------------------ 57310
human.FAM230J-LOC105372942-GGTLC5P.NCBI.rev.compl    ------------------------------------------------------------ 46364
human.FAM230A.ncRNA.18487127.18500594-GGTLC3.18516335.18518165.rev.compl.REF    ------------------------------------------------------------ 31039

chimp.LOC112206744-LOC107973052-GGT2.revcompl    ------------------------------------------------------------ 61762
chimp54546-135457.revcompl.Pan.troglodytes.clone.rp43-41g5.GenBank:AC099533.36    -----------------------------------------------ATACAAACTATAC 67347
human.FAM30B-LOC105372935-GGT2.NCBI.    acactgtgaaagccaccaaatgaatattttaaaatttatctcatttctgccaaggtgag 68148
human.FAM230E-LOC105377182-GGT3P.NCBI.    ------------------------------------------------------------ 57310
human.FAM230J-LOC105372942-GGTLC5P.NCBI.rev.compl    ------------------------------------------------------------ 46364
human.FAM230A.ncRNA.18487127.18500594-GGTLC3.18516335.18518165.rev.compl.REF    ------------------------------------------------------------ 31039

chimp.LOC112206744-LOC107973052-GGT2.revcompl    ------------------------------------------------------------ 61762
chimp54546-135457.revcompl.Pan.troglodytes.clone.rp43-41g5.GenBank:AC099533.36    AAATTGAAGTATAAAGAGAAAAATAGATAGAACATACAGGAAATTGATTAAGAG----AC 67403
human.FAM30B-LOC105372935-GGT2.NCBI.    atatcaatctaaaaacaggataaatctttgtaagtacaagaacattaaaaagaacattaa 68208
human.FAM230E-LOC105377182-GGT3P.NCBI.    ------------------------------------------------------------ 57310
human.FAM230J-LOC105372942-GGTLC5P.NCBI.rev.compl    ------------------------------------------------------------ 46364
human.FAM230A.ncRNA.18487127.18500594-GGTLC3.18516335.18518165.rev.compl.REF    ------------------------------------------------------------ 31039

chimp.LOC112206744-LOC107973052-GGT2.revcompl    ------------------------------------------------------------ 61762
chimp54546-135457.revcompl.Pan.troglodytes.clone.rp43-41g5.GenBank:AC099533.36    ATGGGGATTAACAAGGCATAAGTC------------------------------------ 67427
human.FAM30B-LOC105372935-GGT2.NCBI.    gtttgggttaaaaatgcaaaactgcatatgcaggaaaattacgttaaaaaaaaaaaaatc 68268
human.FAM230E-LOC105377182-GGT3P.NCBI.    ------------------------------------------------------------ 57310
human.FAM230J-LOC105372942-GGTLC5P.NCBI.rev.compl    ------------------------------------------------------------ 46364
human.FAM230A.ncRNA.18487127.18500594-GGTLC3.18516335.18518165.rev.compl.REF    ------------------------------------------------------------ 31039

chimp.LOC112206744-LOC107973052-GGT2.revcompl    ------------------------------------------------------------ 61762
chimp54546-135457.revcompl.Pan.troglodytes.clone.rp43-41g5.GenBank:AC099533.36    ------------------------------------------------------------ 67427
human.FAM30B-LOC105372935-GGT2.NCBI.    ctacacctacaaaaacaaccaaacaaaacattgatgatactgggctgattcactgaatgg 68328
human.FAM230E-LOC105377182-GGT3P.NCBI.    ------------------------------------------------------------ 57310
human.FAM230J-LOC105372942-GGTLC5P.NCBI.rev.compl    ------------------------------------------------------------ 46364
human.FAM230A.ncRNA.18487127.18500594-GGTLC3.18516335.18518165.rev.compl.REF    ------------------------------------------------------------ 31039

chimp.LOC112206744-LOC107973052-GGT2.revcompl    ------------------------------------------------------------ 61762
chimp54546-135457.revcompl.Pan.troglodytes.clone.rp43-41g5.GenBank:AC099533.36    ------------------------------------------------------------ 67427
human.FAM30B-LOC105372935-GGT2.NCBI.    tgagaatatgagtgattttctcctctcctatttttctcaataccttttctctattatctc 68388
human.FAM230E-LOC105377182-GGT3P.NCBI.    ------------------------------------------------------------ 57310
human.FAM230J-LOC105372942-GGTLC5P.NCBI.rev.compl    ------------------------------------------------------------ 46364
human.FAM230A.ncRNA.18487127.18500594-GGTLC3.18516335.18518165.rev.compl.REF    ------------------------------------------------------------ 31039

chimp.LOC112206744-LOC107973052-GGT2.revcompl    ------------------------------------------------------------ 61762
chimp54546-135457.revcompl.Pan.troglodytes.clone.rp43-41g5.GenBank:AC099533.36    ------------------------------------------------------------ 67427
human.FAM30B-LOC105372935-GGT2.NCBI.    ctttttaaataaacaagtatatattgttctaaaccgaagataaatgatcatgagcagca 68448
human.FAM230E-LOC105377182-GGT3P.NCBI.    ------------------------------------------------------------ 57310
human.FAM230J-LOC105372942-GGTLC5P.NCBI.rev.compl    ------------------------------------------------------------ 46364
human.FAM230A.ncRNA.18487127.18500594-GGTLC3.18516335.18518165.rev.compl.REF    ------------------------------------------------------------ 31039

chimp.LOC112206744-LOC107973052-GGT2.revcompl    ------------------------------------------------------------ 61762
chimp54546-135457.revcompl.Pan.troglodytes.clone.rp43-41g5.GenBank:AC099533.36    ------------------------------------------------------------ 67427
human.FAM30B-LOC105372935-GGT2.NCBI.    ctttcccaggcactgtcctagttgctctgcatacatcatcgcatttaatcctcacaacag 68508
human.FAM230E-LOC105377182-GGT3P.NCBI.    ------------------------------------------------------------ 57310
human.FAM230J-LOC105372942-GGTLC5P.NCBI.rev.compl    ------------------------------------------------------------ 46364
human.FAM230A.ncRNA.18487127.18500594-GGTLC3.18516335.18518165.rev.compl.REF    ------------------------------------------------------------ 31039

chimp.LOC112206744-LOC107973052-GGT2.revcompl    ------------------------------------------------------------ 61762
chimp54546-135457.revcompl.Pan.troglodytes.clone.rp43-41g5.GenBank:AC099533.36    ------------------------------------------------------------ 67427
human.FAM30B-LOC105372935-GGT2.NCBI.    tgaatggggtcgatttcattacagccacatttaccagacaaggaaatggttttcaaggtt 68568
human.FAM230E-LOC105377182-GGT3P.NCBI.    ------------------------------------------------------------ 57310
human.FAM230J-LOC105372942-GGTLC5P.NCBI.rev.compl    ------------------------------------------------------------ 46364
human.FAM230A.ncRNA.18487127.18500594-GGTLC3.18516335.18518165.rev.compl.REF    ------------------------------------------------------------ 31039

chimp.LOC112206744-LOC107973052-GGT2.revcompl    ------------------------------------------------------------ 61762
chimp54546-135457.revcompl.Pan.troglodytes.clone.rp43-41g5.GenBank:AC099533.36    ------------------------------------------------------------ 67427
human.FAM30B-LOC105372935-GGT2.NCBI.    aagtaaccttctcatgacacagaatgtgactaagagttttcttattccatggctcacact 68628
human.FAM230E-LOC105377182-GGT3P.NCBI.    ------------------------------------------------------------ 57310
human.FAM230J-LOC105372942-GGTLC5P.NCBI.rev.compl    ------------------------------------------------------------ 46364
human.FAM230A.ncRNA.18487127.18500594-GGTLC3.18516335.18518165.rev.compl.REF    ------------------------------------------------------------ 31039

chimp.LOC112206744-LOC107973052-GGT2.revcompl    ------------------------------------------------------------ 61762
chimp54546-135457.revcompl.Pan.troglodytes.clone.rp43-41g5.GenBank:AC099533.36    -----------------------------------------ATGTGAAATTGGAGTCTCA 67446
human.FAM30B-LOC105372935-GGT2.NCBI.    aaatgtgctctgtgaatatcagtgagaagtaataaagcactgactggattttcttggtca 68688
human.FAM230E-LOC105377182-GGT3P.NCBI.    ------------------------------------------------------------ 57310
human.FAM230J-LOC105372942-GGTLC5P.NCBI.rev.compl    ------------------------------------------------------------ 46364
human.FAM230A.ncRNA.18487127.18500594-GGTLC3.18516335.18518165.rev.compl.REF    ------------------------------------------------------------ 31039

chimp.LOC112206744-LOC107973052-GGT2.revcompl    ------------------------------------------------------------ 61762
chimp54546-135457.revcompl.Pan.troglodytes.clone.rp43-41g5.GenBank:AC099533.36    TAAGGAGAGTAGAGAGAGGATGGGGAAGACGTAATACTTGAAAAGATAAAGCTGACAGTT 67506
human.FAM30B-LOC105372935-GGT2.NCBI.    ccgggtcctaaaagaaatgttggggaat--ctatgcagtcccaagttaaaaatgggaact 68746
human.FAM230E-LOC105377182-GGT3P.NCBI.    ------------------------------------------------------------ 57310
human.FAM230J-LOC105372942-GGTLC5P.NCBI.rev.compl    ------------------------------------------------------------ 46364
human.FAM230A.ncRNA.18487127.18500594-GGTLC3.18516335.18518165.rev.compl.REF    ------------------------------------------------------------ 31039

chimp.LOC112206744-LOC107973052-GGT2.revcompl    ------------------------------------------------------------ 61762
chimp54546-135457.revcompl.Pan.troglodytes.clone.rp43-41g5.GenBank:AC099533.36    TTCTAAAATGGACAAAAAAGAATCAAGTCATGGAATGAAGAGCCACCTAAATGCATCAGT 67566
human.FAM30B-LOC105372935-GGT2.NCBI.    gaactgagctgccacagcatgctgaataggtggagtccag-gccaactaagtgcttcatg 68805
```

```
human.FAM230E-LOC105377182-GGT3P.NCBI.                                          ------------------------------------------------------------ 57310
human.FAM230J-LOC105372942-GGTLC5P.NCBI.rev.compl                               ------------------------------------------------------------ 46364
human.FAM230A.ncRNA.18487127.18500594-GGTLC3.18516335.18518165.rev.compl.REF    ------------------------------------------------------------ 31039

chimp.LOC112206744-LOC107973052-GGT2.revcompl                                   ------------------------------------------------------------ 61762
chimp54546-135457.revcompl.Pan.troglodytes.clone.rp43-41g5.GenBank:AC099533.36  G---TAAACTGCTCAAAACAAAGACAAAACTTTTCAAAGCAGCCAGAGGAAAAAAGATAC 67623
human.FAM30B-LOC105372935-GGT2.NCBI.                                            gaacttaaaggctccagagggaaagcaatcgggaaggcacttccacaggcacatcggtga 68865
human.FAM230E-LOC105377182-GGT3P.NCBI.                                          ------------------------------------------------------------ 57310
human.FAM230J-LOC105372942-GGTLC5P.NCBI.rev.compl                               ------------------------------------------------------------ 46364
human.FAM230A.ncRNA.18487127.18500594-GGTLC3.18516335.18518165.rev.compl.REF    ------------------------------------------------------------ 31039

chimp.LOC112206744-LOC107973052-GGT2.revcompl                                   ------------------------------------------------------------ 61762
chimp54546-135457.revcompl.Pan.troglodytes.clone.rp43-41g5.GenBank:AC099533.36  TGCCTTCAGGCCAGGGTT------------------------------------------ 67641
human.FAM30B-LOC105372935-GGT2.NCBI.                                            caacttctggggattcttacctgtcagccttcagccaaggaaggacaggaagtggcatgg 68925
human.FAM230E-LOC105377182-GGT3P.NCBI.                                          ------------------------------------------------------------ 57310
human.FAM230J-LOC105372942-GGTLC5P.NCBI.rev.compl                               ------------------------------------------------------------ 46364
human.FAM230A.ncRNA.18487127.18500594-GGTLC3.18516335.18518165.rev.compl.REF    ------------------------------------------------------------ 31039

chimp.LOC112206744-LOC107973052-GGT2.revcompl                                   ------------------------------------------------------------ 61762
chimp54546-135457.revcompl.Pan.troglodytes.clone.rp43-41g5.GenBank:AC099533.36  ------------------------------------------------------------ 67641
human.FAM30B-LOC105372935-GGT2.NCBI.                                            gctaaaagatcctttgtttgacctctttccagatcaggtgacaggaaaacacaacacttg 68985
human.FAM230E-LOC105377182-GGT3P.NCBI.                                          ------------------------------------------------------------ 57310
human.FAM230J-LOC105372942-GGTLC5P.NCBI.rev.compl                               ------------------------------------------------------------ 46364
human.FAM230A.ncRNA.18487127.18500594-GGTLC3.18516335.18518165.rev.compl.REF    ------------------------------------------------------------ 31039

chimp.LOC112206744-LOC107973052-GGT2.revcompl                                   ------------------------------------------------------------ 61762
chimp54546-135457.revcompl.Pan.troglodytes.clone.rp43-41g5.GenBank:AC099533.36  ------------------------------------------------------------ 67641
human.FAM30B-LOC105372935-GGT2.NCBI.                                            aacctttccaataaggacagccagcacttactgagccccagccaggtaactgacactgtt 69045
human.FAM230E-LOC105377182-GGT3P.NCBI.                                          ------------------------------------------------------------ 57310
human.FAM230J-LOC105372942-GGTLC5P.NCBI.rev.compl                               ------------------------------------------------------------ 46364
human.FAM230A.ncRNA.18487127.18500594-GGTLC3.18516335.18518165.rev.compl.REF    ------------------------------------------------------------ 31039

chimp.LOC112206744-LOC107973052-GGT2.revcompl                                   ------------------------------------------------------------ 61762
chimp54546-135457.revcompl.Pan.troglodytes.clone.rp43-41g5.GenBank:AC099533.36  ------------------------------------------------------------ 67641
human.FAM30B-LOC105372935-GGT2.NCBI.                                            ccaagtactctgcttgtataaccagctttaaccttgcaacagccctctgaggtaaatact 69105
human.FAM230E-LOC105377182-GGT3P.NCBI.                                          ------------------------------------------------------------ 57310
human.FAM230J-LOC105372942-GGTLC5P.NCBI.rev.compl                               ------------------------------------------------------------ 46364
human.FAM230A.ncRNA.18487127.18500594-GGTLC3.18516335.18518165.rev.compl.REF    ------------------------------------------------------------ 31039

chimp.LOC112206744-LOC107973052-GGT2.revcompl                                   ------------------------------------------------------------ 61762
chimp54546-135457.revcompl.Pan.troglodytes.clone.rp43-41g5.GenBank:AC099533.36  ------------------------------------------------------------ 67641
human.FAM30B-LOC105372935-GGT2.NCBI.                                            cctagtgacctcactttacggatgcagaacctgaggcaaggagttagggagtaaatgaag 69165
human.FAM230E-LOC105377182-GGT3P.NCBI.                                          ------------------------------------------------------------ 57310
human.FAM230J-LOC105372942-GGTLC5P.NCBI.rev.compl                               ------------------------------------------------------------ 46364
human.FAM230A.ncRNA.18487127.18500594-GGTLC3.18516335.18518165.rev.compl.REF    ------------------------------------------------------------ 31039

chimp.LOC112206744-LOC107973052-GGT2.revcompl                                   ------------------------------------------------------------ 61762
chimp54546-135457.revcompl.Pan.troglodytes.clone.rp43-41g5.GenBank:AC099533.36  ------------------------------------------------------------ 67641
human.FAM30B-LOC105372935-GGT2.NCBI.                                            ctggctaaaggcacttgcactggccaggcacagtggctcatgcctgtaatcccagcactt 69225
human.FAM230E-LOC105377182-GGT3P.NCBI.                                          ------------------------------------------------------------ 57310
human.FAM230J-LOC105372942-GGTLC5P.NCBI.rev.compl                               ------------------------------------------------------------ 46364
human.FAM230A.ncRNA.18487127.18500594-GGTLC3.18516335.18518165.rev.compl.REF    ------------------------------------------------------------ 31039

chimp.LOC112206744-LOC107973052-GGT2.revcompl                                   ------------------------------------------------------------ 61762
chimp54546-135457.revcompl.Pan.troglodytes.clone.rp43-41g5.GenBank:AC099533.36  ------------------------------------------------------------ 67641
human.FAM30B-LOC105372935-GGT2.NCBI.                                            ccagatggatcacttgaggttaggggttcgagaccagcctggccaacatggtgaatcccc 69285
human.FAM230E-LOC105377182-GGT3P.NCBI.                                          ------------------------------------------------------------ 57310
human.FAM230J-LOC105372942-GGTLC5P.NCBI.rev.compl                               ------------------------------------------------------------ 46364
human.FAM230A.ncRNA.18487127.18500594-GGTLC3.18516335.18518165.rev.compl.REF    ------------------------------------------------------------ 31039

chimp.LOC112206744-LOC107973052-GGT2.revcompl                                   ------------------------------------------------------------ 61762
chimp54546-135457.revcompl.Pan.troglodytes.clone.rp43-41g5.GenBank:AC099533.36  ------------------------------GGGGGCTGGCTCATGCCTGTAATCCCAGCA 67671
human.FAM30B-LOC105372935-GGT2.NCBI.                                            agctctactaaaaatacaaaaattagccgggtgtgatggcacaagcctgtaatcccagca 69345
human.FAM230E-LOC105377182-GGT3P.NCBI.                                          ------------------------------------------------------------ 57310
human.FAM230J-LOC105372942-GGTLC5P.NCBI.rev.compl                               ------------------------------------------------------------ 46364
human.FAM230A.ncRNA.18487127.18500594-GGTLC3.18516335.18518165.rev.compl.REF    ------------------------------------------------------------ 31039

chimp.LOC112206744-LOC107973052-GGT2.revcompl                                   ------------------------------------------------------------ 61762
chimp54546-135457.revcompl.Pan.troglodytes.clone.rp43-41g5.GenBank:AC099533.36  -TTTGGGAGGCTGAGGCGGGCGGATCACCTGAAATGAGGAGTTTGGGATCAGCCTGACCA 67730
human.FAM30B-LOC105372935-GGT2.NCBI.                                            ctttgggaggccaaggcagatgggtcacttgaggttaggggttcgagaccagcctggcca 69405
human.FAM230E-LOC105377182-GGT3P.NCBI.                                          ------------------------------------------------------------ 57310
human.FAM230J-LOC105372942-GGTLC5P.NCBI.rev.compl                               ------------------------------------------------------------ 46364
human.FAM230A.ncRNA.18487127.18500594-GGTLC3.18516335.18518165.rev.compl.REF    ------------------------------------------------------------ 31039

chimp.LOC112206744-LOC107973052-GGT2.revcompl                                   ------------------------------------------------------------ 61762
chimp54546-135457.revcompl.Pan.troglodytes.clone.rp43-41g5.GenBank:AC099533.36  ACATGGCAAAAGTTCATCTCTATTAAAAATACAACAATTAGCCAGGTGTGATGGTGGACA 67790
human.FAM30B-LOC105372935-GGT2.NCBI.                                            acatggtgaatccccagctctactaaaaatacaaaaattagccgggtgtgatggcacacg 69465
human.FAM230E-LOC105377182-GGT3P.NCBI.                                          ------------------------------------------------------------ 57310
human.FAM230J-LOC105372942-GGTLC5P.NCBI.rev.compl                               ------------------------------------------------------------ 46364
human.FAM230A.ncRNA.18487127.18500594-GGTLC3.18516335.18518165.rev.compl.REF    ------------------------------------------------------------ 31039

chimp.LOC112206744-LOC107973052-GGT2.revcompl                                   ------------------------------------------------------------ 61762
chimp54546-135457.revcompl.Pan.troglodytes.clone.rp43-41g5.GenBank:AC099533.36  CCTGTAATCCCAGCTACTTGGGAGGCTGAGGCAGAAGAATCACTTGAACCCGGGAGGCAG 67850
human.FAM30B-LOC105372935-GGT2.NCBI.                                            cctgtaatcccagctactcaggaggctgaggcaggagaattgcttgaacctgggagatgg 69525
human.FAM230E-LOC105377182-GGT3P.NCBI.                                          ------------------------------------------------------------ 57310
human.FAM230J-LOC105372942-GGTLC5P.NCBI.rev.compl                               ------------------------------------------------------------ 46364
human.FAM230A.ncRNA.18487127.18500594-GGTLC3.18516335.18518165.rev.compl.REF    ------------------------------------------------------------ 31039

chimp.LOC112206744-LOC107973052-GGT2.revcompl                                   ------------------------------------------------------------ 61762
chimp54546-135457.revcompl.Pan.troglodytes.clone.rp43-41g5.GenBank:AC099533.36  AGGTTGCAGTGAGCTGAGATTGCGCTACTGCACTCCAGCCTGGGCGACAGAGTAAGACTC 67910
human.FAM30B-LOC105372935-GGT2.NCBI.                                            aggttgcagcaagccgagatcgcaccactgcactccagcccactcggtcaatgtcactgc 69585
human.FAM230E-LOC105377182-GGT3P.NCBI.                                          ------------------------------------------------------------ 57310
human.FAM230J-LOC105372942-GGTLC5P.NCBI.rev.compl                               ------------------------------------------------------------ 46364
human.FAM230A.ncRNA.18487127.18500594-GGTLC3.18516335.18518165.rev.compl.REF    ------------------------------------------------------------ 31039

chimp.LOC112206744-LOC107973052-GGT2.revcompl                                   ------------------------------------------------------------ 61762
chimp54546-135457.revcompl.Pan.troglodytes.clone.rp43-41g5.GenBank:AC099533.36  CATC-----TCAGAAAACACAACAGCATAAAAAGCCATTGCCTTCAAATGAGCAAAATTA 67965
human.FAM30B-LOC105372935-GGT2.NCBI.                                            catcaagggacagtctaaccaaggtccagatggcacaagtggagccataaaaccaggaca 69645
human.FAM230E-LOC105377182-GGT3P.NCBI.                                          ------------------------------------------------------------ 57310
human.FAM230J-LOC105372942-GGTLC5P.NCBI.rev.compl                               ------------------------------------------------------------ 46364
human.FAM230A.ncRNA.18487127.18500594-GGTLC3.18516335.18518165.rev.compl.REF    ------------------------------------------------------------ 31039

chimp.LOC112206744-LOC107973052-GGT2.revcompl                                   ------------------------------------------------------------ 61762
chimp54546-135457.revcompl.Pan.troglodytes.clone.rp43-41g5.GenBank:AC099533.36  GATTGAGAATTGGCTTCCCAATAAAAGCAGGAGAAATCAGATACAAGGGAATCATGTTTT 68025
human.FAM30B-LOC105372935-GGT2.NCBI.                                            gcttgaggttcagcttcctaagccca-caggtgcaaggaccttatctggatccatacttt 69704
human.FAM230E-LOC105377182-GGT3P.NCBI.                                          ------------------------------------------------------------ 57310
human.FAM230J-LOC105372942-GGTLC5P.NCBI.rev.compl                               ------------------------------------------------------------ 46364
human.FAM230A.ncRNA.18487127.18500594-GGTLC3.18516335.18518165.rev.compl.REF    ------------------------------------------------------------ 31039

chimp.LOC112206744-LOC107973052-GGT2.revcompl                                   ------------------------------------------------------------ 61762
chimp54546-135457.revcompl.Pan.troglodytes.clone.rp43-41g5.GenBank:AC099533.36  CAAACTGAGGAAAGAAAATATCTGCCAACCTAGAATTCTATGCTCACTAACAATATCCTT 68085
human.FAM30B-LOC105372935-GGT2.NCBI.                                            ca---------------------------------------------------------t 69707
human.FAM230E-LOC105377182-GGT3P.NCBI.                                          ------------------------------------------------------------ 57310
human.FAM230J-LOC105372942-GGTLC5P.NCBI.rev.compl                               ------------------------------------------------------------ 46364
human.FAM230A.ncRNA.18487127.18500594-GGTLC3.18516335.18518165.rev.compl.REF    ------------------------------------------------------------ 31039

chimp.LOC112206744-LOC107973052-GGT2.revcompl                                   ------------------------------------------------------------ 61762
chimp54546-135457.revcompl.Pan.troglodytes.clone.rp43-41g5.GenBank:AC099533.36  CAAAAGTGAAAAT---GGCTGGGCATGGTGGCTCATGCCTGTAATCCCAGCACTTTGGAA 68142
human.FAM30B-LOC105372935-GGT2.NCBI.                                            caaaaatgtcccttgaggccaggcacagtggctcatgcctataatcccagcactttggga 69767
human.FAM230E-LOC105377182-GGT3P.NCBI.                                          ------------------------------------------------------------ 57310
human.FAM230J-LOC105372942-GGTLC5P.NCBI.rev.compl                               ------------------------------------------------------------ 46364
human.FAM230A.ncRNA.18487127.18500594-GGTLC3.18516335.18518165.rev.compl.REF    ------------------------------------------------------------ 31039

chimp.LOC112206744-LOC107973052-GGT2.revcompl                                   ------------------------------------------------------------ 61762
chimp54546-135457.revcompl.Pan.troglodytes.clone.rp43-41g5.GenBank:AC099533.36  GGCCAAGGCAGGTGGATCACCTGAGGTCAGAAGTTCGAGACCAGCCTGGCCAACATGGTG 68202
human.FAM30B-LOC105372935-GGT2.NCBI.                                            gggcgaggcaggtggatcacctgaggtcaggagttcgagaccagcctggccaatatggtg 69827
human.FAM230E-LOC105377182-GGT3P.NCBI.                                          ------------------------------------------------------------ 57310
```

```
human.FAM230J-LOC105372942-GGTLC5P.NCBI.rev.compl                            ------------------------------------------------------------ 46364
human.FAM230A.ncRNA.18487127.18500594-GGTLC3.18516335.18518165.rev.compl.REF ------------------------------------------------------------ 31039

chimp.LOC112206744-LOC107973052-GGT2.revcompl                                ------------------------------------------------------------ 61762
chimp54546-135457.revcompl.Pan.troglodytes.clone.rp43-41g5.GenBank:AC099533.36 AAACCCCATCTCTACCAAAAATACACAAA-TTAGCCAGATGTGGTGGTGGATGCCTGTAA 68261
human.FAM30B-LOC105372935-GGT2.NCBI.                                         aaaccctgtccctactgaaaatacaaaaatttagcccagcatggtgatgtgcgcctgtaa 69887
human.FAM230E-LOC105377182-GGT3P.NCBI.                                       ------------------------------------------------------------ 57310
human.FAM230J-LOC105372942-GGTLC5P.NCBI.rev.compl                            ------------------------------------------------------------ 46364
human.FAM230A.ncRNA.18487127.18500594-GGTLC3.18516335.18518165.rev.compl.REF ------------------------------------------------------------ 31039

chimp.LOC112206744-LOC107973052-GGT2.revcompl                                ------------------------------------------------------------ 61762
chimp54546-135457.revcompl.Pan.troglodytes.clone.rp43-41g5.GenBank:AC099533.36 TCCCAGCTACTCAGGAGGCTGAGGCACAAGAATTGCTTGAACCCGAGAGGTGGAGGTTGC 68321
human.FAM30B-LOC105372935-GGT2.NCBI.                                         tcccagctactccggaggctgaggcaggagaatcgcttgaacccagcaggtggaggttgc 69947
human.FAM230E-LOC105377182-GGT3P.NCBI.                                       ------------------------------------------------------------ 57310
human.FAM230J-LOC105372942-GGTLC5P.NCBI.rev.compl                            ------------------------------------------------------------ 46364
human.FAM230A.ncRNA.18487127.18500594-GGTLC3.18516335.18518165.rev.compl.REF ------------------------------------------------------------ 31039

chimp.LOC112206744-LOC107973052-GGT2.revcompl                                ------------------------------------------------------------ 61762
chimp54546-135457.revcompl.Pan.troglodytes.clone.rp43-41g5.GenBank:AC099533.36 AGTGAGCCAAGATCTTGCCACTGCACTCCAGCCTGGGTGACAGAGCAAAACTCCATCTCA 68381
human.FAM30B-LOC105372935-GGT2.NCBI.                                         agtgagccgagatcatgtcattgcactgtgggctgggcaacagagcaacactctatgtaa 70007
human.FAM230E-LOC105377182-GGT3P.NCBI.                                       ------------------------------------------------------------ 57310
human.FAM230J-LOC105372942-GGTLC5P.NCBI.rev.compl                            ------------------------------------------------------------ 46364
human.FAM230A.ncRNA.18487127.18500594-GGTLC3.18516335.18518165.rev.compl.REF ------------------------------------------------------------ 31039

chimp.LOC112206744-LOC107973052-GGT2.revcompl                                ------------------------------------------------------------ 61762
chimp54546-135457.revcompl.Pan.troglodytes.clone.rp43-41g5.GenBank:AC099533.36 AAAAAAAAAACAGG--CCAGTCACGGTGGCTCATGCCT---------------------- 68417
human.FAM30B-LOC105372935-GGT2.NCBI.                                         aaaaaaaaaaaaaggtcccttcaggattcctcgagcctgggaggccgaggctgccatgaa 70067
human.FAM230E-LOC105377182-GGT3P.NCBI.                                       ------------------------------------------------------------ 57310
human.FAM230J-LOC105372942-GGTLC5P.NCBI.rev.compl                            ------------------------------------------------------------ 46364
human.FAM230A.ncRNA.18487127.18500594-GGTLC3.18516335.18518165.rev.compl.REF ------------------------------------------------------------ 31039

chimp.LOC112206744-LOC107973052-GGT2.revcompl                                ------------------------------------------------------------ 61762
chimp54546-135457.revcompl.Pan.troglodytes.clone.rp43-41g5.GenBank:AC099533.36 ------------------------------------------------------------ 68417
human.FAM30B-LOC105372935-GGT2.NCBI.                                         caatgattgtgccactgcactccagcctgggtgacagagcgagatcttgtctcaaaaaaa 70127
human.FAM230E-LOC105377182-GGT3P.NCBI.                                       ------------------------------------------------------------ 57310
human.FAM230J-LOC105372942-GGTLC5P.NCBI.rev.compl                            ------------------------------------------------------------ 46364
human.FAM230A.ncRNA.18487127.18500594-GGTLC3.18516335.18518165.rev.compl.REF ------------------------------------------------------------ 31039

chimp.LOC112206744-LOC107973052-GGT2.revcompl                                ------------------------------------------------------------ 61762
chimp54546-135457.revcompl.Pan.troglodytes.clone.rp43-41g5.GenBank:AC099533.36 ------------------------------------------------------------ 68417
human.FAM30B-LOC105372935-GGT2.NCBI.                                         aaaaaaaaaaaaaaaatggctcttgaaacagccatcaactcccatgtgataaatgcactg 70187
human.FAM230E-LOC105377182-GGT3P.NCBI.                                       ------------------------------------------------------------ 57310
human.FAM230J-LOC105372942-GGTLC5P.NCBI.rev.compl                            ------------------------------------------------------------ 46364
human.FAM230A.ncRNA.18487127.18500594-GGTLC3.18516335.18518165.rev.compl.REF ------------------------------------------------------------ 31039

chimp.LOC112206744-LOC107973052-GGT2.revcompl                                ------------------------------------------------------------ 61762
chimp54546-135457.revcompl.Pan.troglodytes.clone.rp43-41g5.GenBank:AC099533.36 ------------------------------------------------------------ 68417
human.FAM30B-LOC105372935-GGT2.NCBI.                                         ccacaatccagcagcaaacctaaagttcggagtgaattcctaagtcttcgccttcactga 70247
human.FAM230E-LOC105377182-GGT3P.NCBI.                                       ------------------------------------------------------------ 57310
human.FAM230J-LOC105372942-GGTLC5P.NCBI.rev.compl                            ------------------------------------------------------------ 46364
human.FAM230A.ncRNA.18487127.18500594-GGTLC3.18516335.18518165.rev.compl.REF ------------------------------------------------------------ 31039

chimp.LOC112206744-LOC107973052-GGT2.revcompl                                ------------------------------------------------------------ 61762
chimp54546-135457.revcompl.Pan.troglodytes.clone.rp43-41g5.GenBank:AC099533.36 ------------------------------------------------------------ 68417
human.FAM30B-LOC105372935-GGT2.NCBI.                                         cttcccacagcatgggcagtagctgtgagaccttgggctggttacttaacctctctgcac 70307
human.FAM230E-LOC105377182-GGT3P.NCBI.                                       ------------------------------------------------------------ 57310
human.FAM230J-LOC105372942-GGTLC5P.NCBI.rev.compl                            ------------------------------------------------------------ 46364
human.FAM230A.ncRNA.18487127.18500594-GGTLC3.18516335.18518165.rev.compl.REF ------------------------------------------------------------ 31039

chimp.LOC112206744-LOC107973052-GGT2.revcompl                                ------------------------------------------------------------ 61762
chimp54546-135457.revcompl.Pan.troglodytes.clone.rp43-41g5.GenBank:AC099533.36 ------------------------------------------------------------ 68417
human.FAM30B-LOC105372935-GGT2.NCBI.                                         ctcagtttcctcatctgtaaagtggtataatcgtcacagtgttatggaaggatttaatga 70367
human.FAM230E-LOC105377182-GGT3P.NCBI.                                       ------------------------------------------------------------ 57310
human.FAM230J-LOC105372942-GGTLC5P.NCBI.rev.compl                            ------------------------------------------------------------ 46364
human.FAM230A.ncRNA.18487127.18500594-GGTLC3.18516335.18518165.rev.compl.REF ------------------------------------------------------------ 31039

chimp.LOC112206744-LOC107973052-GGT2.revcompl                                ------------------------------------------------------------ 61762
chimp54546-135457.revcompl.Pan.troglodytes.clone.rp43-41g5.GenBank:AC099533.36 ------------------------------------------------------------ 68417
human.FAM30B-LOC105372935-GGT2.NCBI.                                         gttgctacatttaaaacttaattcctggcacacaataagtgctcaataaatgtttagccg 70427
human.FAM230E-LOC105377182-GGT3P.NCBI.                                       ------------------------------------------------------------ 57310
human.FAM230J-LOC105372942-GGTLC5P.NCBI.rev.compl                            ------------------------------------------------------------ 46364
human.FAM230A.ncRNA.18487127.18500594-GGTLC3.18516335.18518165.rev.compl.REF ------------------------------------------------------------ 31039

chimp.LOC112206744-LOC107973052-GGT2.revcompl                                ------------------------------------------------------------ 61762
chimp54546-135457.revcompl.Pan.troglodytes.clone.rp43-41g5.GenBank:AC099533.36 ------------------------------------------------------------ 68417
human.FAM30B-LOC105372935-GGT2.NCBI.                                         ctagttgcagtagtagcaatagtagtgatattaactctagctgtggaactaatttctccc 70487
human.FAM230E-LOC105377182-GGT3P.NCBI.                                       ------------------------------------------------------------ 57310
human.FAM230J-LOC105372942-GGTLC5P.NCBI.rev.compl                            ------------------------------------------------------------ 46364
human.FAM230A.ncRNA.18487127.18500594-GGTLC3.18516335.18518165.rev.compl.REF ------------------------------------------------------------ 31039

chimp.LOC112206744-LOC107973052-GGT2.revcompl                                ------------------------------------------------------------ 61762
chimp54546-135457.revcompl.Pan.troglodytes.clone.rp43-41g5.GenBank:AC099533.36 ------------------------------------------------------------ 68417
human.FAM30B-LOC105372935-GGT2.NCBI.                                         acacaccgtggcaggttgtattctccagacacagcagtaacaatatctccttcccacacg 70547
human.FAM230E-LOC105377182-GGT3P.NCBI.                                       ------------------------------------------------------------ 57310
human.FAM230J-LOC105372942-GGTLC5P.NCBI.rev.compl                            ------------------------------------------------------------ 46364
human.FAM230A.ncRNA.18487127.18500594-GGTLC3.18516335.18518165.rev.compl.REF ------------------------------------------------------------ 31039

chimp.LOC112206744-LOC107973052-GGT2.revcompl                                ------------------------------------------------------------ 61762
chimp54546-135457.revcompl.Pan.troglodytes.clone.rp43-41g5.GenBank:AC099533.36 ------------------------------------------------------------ 68417
human.FAM30B-LOC105372935-GGT2.NCBI.                                         ctcttttgcaatgtgaccatgccacacactcccatcaagaggtggagtttagaataattc 70607
human.FAM230E-LOC105377182-GGT3P.NCBI.                                       ------------------------------------------------------------ 57310
human.FAM230J-LOC105372942-GGTLC5P.NCBI.rev.compl                            ------------------------------------------------------------ 46364
human.FAM230A.ncRNA.18487127.18500594-GGTLC3.18516335.18518165.rev.compl.REF ------------------------------------------------------------ 31039

chimp.LOC112206744-LOC107973052-GGT2.revcompl                                ------------------------------------------------------------ 61762
chimp54546-135457.revcompl.Pan.troglodytes.clone.rp43-41g5.GenBank:AC099533.36 ------------------------------------------------------------ 68417
human.FAM30B-LOC105372935-GGT2.NCBI.                                         cctcctctgtggatctgggcaggaagtcactgtgtgacttccagtcctaggtcatgacaa 70667
human.FAM230E-LOC105377182-GGT3P.NCBI.                                       ------------------------------------------------------------ 57310
human.FAM230J-LOC105372942-GGTLC5P.NCBI.rev.compl                            ------------------------------------------------------------ 46364
human.FAM230A.ncRNA.18487127.18500594-GGTLC3.18516335.18518165.rev.compl.REF ------------------------------------------------------------ 31039

chimp.LOC112206744-LOC107973052-GGT2.revcompl                                ------------------------------------------------------------ 61762
chimp54546-135457.revcompl.Pan.troglodytes.clone.rp43-41g5.GenBank:AC099533.36 ------------------------------------------------------------ 68417
human.FAM30B-LOC105372935-GGT2.NCBI.                                         gtgatgtagcttccatttcactcagggtgccctgaaccaccacataagaaagccaacggt 70727
human.FAM230E-LOC105377182-GGT3P.NCBI.                                       ------------------------------------------------------------ 57310
human.FAM230J-LOC105372942-GGTLC5P.NCBI.rev.compl                            ------------------------------------------------------------ 46364
human.FAM230A.ncRNA.18487127.18500594-GGTLC3.18516335.18518165.rev.compl.REF ------------------------------------------------------------ 31039

chimp.LOC112206744-LOC107973052-GGT2.revcompl                                ------------------------------------------------------------ 61762
chimp54546-135457.revcompl.Pan.troglodytes.clone.rp43-41g5.GenBank:AC099533.36 ------------------------------------------------------------ 68417
human.FAM30B-LOC105372935-GGT2.NCBI.                                         cctgaggccatgctgtgagggggcccaaggcacatggagaggcccagggcaggtgctcca 70787
human.FAM230E-LOC105377182-GGT3P.NCBI.                                       ------------------------------------------------------------ 57310
human.FAM230J-LOC105372942-GGTLC5P.NCBI.rev.compl                            ------------------------------------------------------------ 46364
human.FAM230A.ncRNA.18487127.18500594-GGTLC3.18516335.18518165.rev.compl.REF ------------------------------------------------------------ 31039

chimp.LOC112206744-LOC107973052-GGT2.revcompl                                ------------------------------------------------------------ 61762
chimp54546-135457.revcompl.Pan.troglodytes.clone.rp43-41g5.GenBank:AC099533.36 ------------------------------------------------------------ 68417
human.FAM30B-LOC105372935-GGT2.NCBI.                                         gtccagagctggctccagccaggtccccctgccaccatgtgagtgaatgagccaatcacc 70847
human.FAM230E-LOC105377182-GGT3P.NCBI.                                       ------------------------------------------------------------ 57310
human.FAM230J-LOC105372942-GGTLC5P.NCBI.rev.compl                            ------------------------------------------------------------ 46364
human.FAM230A.ncRNA.18487127.18500594-GGTLC3.18516335.18518165.rev.compl.REF ------------------------------------------------------------ 31039

chimp.LOC112206744-LOC107973052-GGT2.revcompl                                ------------------------------------------------------------ 61762
chimp54546-135457.revcompl.Pan.troglodytes.clone.rp43-41g5.GenBank:AC099533.36 ------------------------------------------------------------ 68417
human.FAM30B-LOC105372935-GGT2.NCBI.                                         acccagctccacctgccaccatgtgagtgaacaagcctcccctgtgagctgcccccagcc 70907
human.FAM230E-LOC105377182-GGT3P.NCBI.                                       ------------------------------------------------------------ 57310
human.FAM230J-LOC105372942-GGTLC5P.NCBI.rev.compl                            ------------------------------------------------------------ 46364
```

```
human.FAM230A.ncRNA.18487127.18500594-GGTLC3.18516335.18518165.rev.compl.REF        ------------------------------------------------------------ 31039


chimp.LOC112206744-LOC107973052-GGT2.revcompl                                       ------------------------------------------------------------ 61762
chimp54546-135457.revcompl.Pan.troglodytes.clone.rp43-41g5.GenBank:AC099533.36      ------------------------------------------------------------ 68417
human.FAM30B-LOC105372935-GGT2.NCBI.                                                ttggaacttccagccatgcctgccgtgcctgtaaattctcaacccccagattctgtgagc 70967
human.FAM230E-LOC105377182-GGT3P.NCBI.                                              ------------------------------------------------------------ 57310
human.FAM230J-LOC105372942-GGTLC5P.NCBI.rev.compl                                   ------------------------------------------------------------ 46364
human.FAM230A.ncRNA.18487127.18500594-GGTLC3.18516335.18518165.rev.compl.REF        ------------------------------------------------------------ 31039


chimp.LOC112206744-LOC107973052-GGT2.revcompl                                       ------------------------------------------------------------ 61762
chimp54546-135457.revcompl.Pan.troglodytes.clone.rp43-41g5.GenBank:AC099533.36      ------------------------------------------------------------ 68417
human.FAM30B-LOC105372935-GGT2.NCBI.                                                ataacaaaattgttgttttaagccactgaggttgggaaacatttgttacatagaaaggta 71027
human.FAM230E-LOC105377182-GGT3P.NCBI.                                              ------------------------------------------------------------ 57310
human.FAM230J-LOC105372942-GGTLC5P.NCBI.rev.compl                                   ------------------------------------------------------------ 46364
human.FAM230A.ncRNA.18487127.18500594-GGTLC3.18516335.18518165.rev.compl.REF        ------------------------------------------------------------ 31039


chimp.LOC112206744-LOC107973052-GGT2.revcompl                                       ------------------------------------------------------------ 61762
chimp54546-135457.revcompl.Pan.troglodytes.clone.rp43-41g5.GenBank:AC099533.36      ------------------------------------------------------------ 68417
human.FAM30B-LOC105372935-GGT2.NCBI.                                                acaagaacatatgctatggcttcaatcaacatttttttcagtacctattatgatggacac 71087
human.FAM230E-LOC105377182-GGT3P.NCBI.                                              ------------------------------------------------------------ 57310
human.FAM230J-LOC105372942-GGTLC5P.NCBI.rev.compl                                   ------------------------------------------------------------ 46364
human.FAM230A.ncRNA.18487127.18500594-GGTLC3.18516335.18518165.rev.compl.REF        ------------------------------------------------------------ 31039


chimp.LOC112206744-LOC107973052-GGT2.revcompl                                       ------------------------------------------------------------ 61762
chimp54546-135457.revcompl.Pan.troglodytes.clone.rp43-41g5.GenBank:AC099533.36      ------------------------------------------------------------ 68417
human.FAM30B-LOC105372935-GGT2.NCBI.                                                taggacagaaactgtgagggagagattaataagacatagttgacaagttgtcctaagctg 71147
human.FAM230E-LOC105377182-GGT3P.NCBI.                                              ------------------------------------------------------------ 57310
human.FAM230J-LOC105372942-GGTLC5P.NCBI.rev.compl                                   ------------------------------------------------------------ 46364
human.FAM230A.ncRNA.18487127.18500594-GGTLC3.18516335.18518165.rev.compl.REF        ------------------------------------------------------------ 31039


chimp.LOC112206744-LOC107973052-GGT2.revcompl                                       ------------------------------------------------------------ 61762
chimp54546-135457.revcompl.Pan.troglodytes.clone.rp43-41g5.GenBank:AC099533.36      ------------------------------------------------------------ 68417
human.FAM30B-LOC105372935-GGT2.NCBI.                                                acaatcctaaaataaatacacataactaatcattttaatggtatacgatagcagtgaggt 71207
human.FAM230E-LOC105377182-GGT3P.NCBI.                                              ------------------------------------------------------------ 57310
human.FAM230J-LOC105372942-GGTLC5P.NCBI.rev.compl                                   ------------------------------------------------------------ 46364
human.FAM230A.ncRNA.18487127.18500594-GGTLC3.18516335.18518165.rev.compl.REF        ------------------------------------------------------------ 31039


chimp.LOC112206744-LOC107973052-GGT2.revcompl                                       ------------------------------------------------------------ 61762
chimp54546-135457.revcompl.Pan.troglodytes.clone.rp43-41g5.GenBank:AC099533.36      ------------------------------------------------------------ 68417
human.FAM30B-LOC105372935-GGT2.NCBI.                                                gcaaagtgcaaggttaagccctttcagaaagtgaaagcattcaaaaagccttcaaaaaag 71267
human.FAM230E-LOC105377182-GGT3P.NCBI.                                              ------------------------------------------------------------ 57310
human.FAM230J-LOC105372942-GGTLC5P.NCBI.rev.compl                                   ------------------------------------------------------------ 46364
human.FAM230A.ncRNA.18487127.18500594-GGTLC3.18516335.18518165.rev.compl.REF        ------------------------------------------------------------ 31039


chimp.LOC112206744-LOC107973052-GGT2.revcompl                                       ------------------------------------------------------------ 61762
chimp54546-135457.revcompl.Pan.troglodytes.clone.rp43-41g5.GenBank:AC099533.36      ------------------------------------------------------------ 68417
human.FAM30B-LOC105372935-GGT2.NCBI.                                                tagtgtcatctgaattggatcccaaagcgcaagggaaatatataccacatgaaaacaccg 71327
human.FAM230E-LOC105377182-GGT3P.NCBI.                                              ------------------------------------------------------------ 57310
human.FAM230J-LOC105372942-GGTLC5P.NCBI.rev.compl                                   ------------------------------------------------------------ 46364
human.FAM230A.ncRNA.18487127.18500594-GGTLC3.18516335.18518165.rev.compl.REF        ------------------------------------------------------------ 31039


chimp.LOC112206744-LOC107973052-GGT2.revcompl                                       ------------------------------------------------------------ 61762
chimp54546-135457.revcompl.Pan.troglodytes.clone.rp43-41g5.GenBank:AC099533.36      ------------------------------------------------------------ 68417
human.FAM30B-LOC105372935-GGT2.NCBI.                                                cagtgcattccaggattagcaagaagtttgcggaggtcgttttgtcacatgactggaggg 71387
human.FAM230E-LOC105377182-GGT3P.NCBI.                                              ------------------------------------------------------------ 57310
human.FAM230J-LOC105372942-GGTLC5P.NCBI.rev.compl                                   ------------------------------------------------------------ 46364
human.FAM230A.ncRNA.18487127.18500594-GGTLC3.18516335.18518165.rev.compl.REF        ------------------------------------------------------------ 31039


chimp.LOC112206744-LOC107973052-GGT2.revcompl                                       ------------------------------------------------------------ 61762
chimp54546-135457.revcompl.Pan.troglodytes.clone.rp43-41g5.GenBank:AC099533.36      ------------------------------------------------------------ 68417
human.FAM30B-LOC105372935-GGT2.NCBI.                                                cacatcagtgccaggcggaaaagggcaccgaaaggcacattaataaagtcaaacacctcc 71447
human.FAM230E-LOC105377182-GGT3P.NCBI.                                              ------------------------------------------------------------ 57310
human.FAM230J-LOC105372942-GGTLC5P.NCBI.rev.compl                                   ------------------------------------------------------------ 46364
human.FAM230A.ncRNA.18487127.18500594-GGTLC3.18516335.18518165.rev.compl.REF        ------------------------------------------------------------ 31039


chimp.LOC112206744-LOC107973052-GGT2.revcompl                                       ------------------------------------------------------------ 61762
chimp54546-135457.revcompl.Pan.troglodytes.clone.rp43-41g5.GenBank:AC099533.36      ------------------------------------------------------------ 68417
human.FAM30B-LOC105372935-GGT2.NCBI.                                                tctgggacaggggaagttgagaaagagcctaacagaggtaaggaagaagacggatcagat 71507
human.FAM230E-LOC105377182-GGT3P.NCBI.                                              ------------------------------------------------------------ 57310
human.FAM230J-LOC105372942-GGTLC5P.NCBI.rev.compl                                   ------------------------------------------------------------ 46364
human.FAM230A.ncRNA.18487127.18500594-GGTLC3.18516335.18518165.rev.compl.REF        ------------------------------------------------------------ 31039


chimp.LOC112206744-LOC107973052-GGT2.revcompl                                       ------------------------------------------------------------ 61762
chimp54546-135457.revcompl.Pan.troglodytes.clone.rp43-41g5.GenBank:AC099533.36      ------------------------------------------------------------ 68417
human.FAM30B-LOC105372935-GGT2.NCBI.                                                ttacactttcgaattatcactctggtggcaggtgaaggacagagcagagggcaggactgg 71567
human.FAM230E-LOC105377182-GGT3P.NCBI.                                              ------------------------------------------------------------ 57310
human.FAM230J-LOC105372942-GGTLC5P.NCBI.rev.compl                                   ------------------------------------------------------------ 46364
human.FAM230A.ncRNA.18487127.18500594-GGTLC3.18516335.18518165.rev.compl.REF        ------------------------------------------------------------ 31039


chimp.LOC112206744-LOC107973052-GGT2.revcompl                                       ------------------------------------------------------------ 61762
chimp54546-135457.revcompl.Pan.troglodytes.clone.rp43-41g5.GenBank:AC099533.36      ------------------------------------------------------------ 68417
human.FAM30B-LOC105372935-GGT2.NCBI.                                                aagcagacagaccaaattcacagtcaaatgcgacaagatgagggcctggcctggttttaa 71627
human.FAM230E-LOC105377182-GGT3P.NCBI.                                              ------------------------------------------------------------ 57310
human.FAM230J-LOC105372942-GGTLC5P.NCBI.rev.compl                                   ------------------------------------------------------------ 46364
human.FAM230A.ncRNA.18487127.18500594-GGTLC3.18516335.18518165.rev.compl.REF        ------------------------------------------------------------ 31039


chimp.LOC112206744-LOC107973052-GGT2.revcompl                                       ------------------------------------------------------------ 61762
chimp54546-135457.revcompl.Pan.troglodytes.clone.rp43-41g5.GenBank:AC099533.36      ------------------------------------------------------------ 68417
human.FAM30B-LOC105372935-GGT2.NCBI.                                                atccaaggacaaaaatatcattcattcactctttcattctttctcttatctgaatcccca 71687
human.FAM230E-LOC105377182-GGT3P.NCBI.                                              ------------------------------------------------------------ 57310
human.FAM230J-LOC105372942-GGTLC5P.NCBI.rev.compl                                   ------------------------------------------------------------ 46364
human.FAM230A.ncRNA.18487127.18500594-GGTLC3.18516335.18518165.rev.compl.REF        ------------------------------------------------------------ 31039


chimp.LOC112206744-LOC107973052-GGT2.revcompl                                       ------------------------------------------------------------ 61762
chimp54546-135457.revcompl.Pan.troglodytes.clone.rp43-41g5.GenBank:AC099533.36      ------------------------------------------------------------ 68417
human.FAM30B-LOC105372935-GGT2.NCBI.                                                gtggtacctaaaatagagctggtctaaagatgatgattaatgaatgttaaattattgata 71747
human.FAM230E-LOC105377182-GGT3P.NCBI.                                              ------------------------------------------------------------ 57310
human.FAM230J-LOC105372942-GGTLC5P.NCBI.rev.compl                                   ------------------------------------------------------------ 46364
human.FAM230A.ncRNA.18487127.18500594-GGTLC3.18516335.18518165.rev.compl.REF        ------------------------------------------------------------ 31039


chimp.LOC112206744-LOC107973052-GGT2.revcompl                                       ------------------------------------------------------------ 61762
chimp54546-135457.revcompl.Pan.troglodytes.clone.rp43-41g5.GenBank:AC099533.36      ------------------------------------------------------------ 68417
human.FAM30B-LOC105372935-GGT2.NCBI.                                                ctacttctaagaagatgtaactgccagctacagaaccagccaaaaggactgcaaacccag 71807
human.FAM230E-LOC105377182-GGT3P.NCBI.                                              ------------------------------------------------------------ 57310
human.FAM230J-LOC105372942-GGTLC5P.NCBI.rev.compl                                   ------------------------------------------------------------ 46364
human.FAM230A.ncRNA.18487127.18500594-GGTLC3.18516335.18518165.rev.compl.REF        ------------------------------------------------------------ 31039


chimp.LOC112206744-LOC107973052-GGT2.revcompl                                       ------------------------------------------------------------ 61762
chimp54546-135457.revcompl.Pan.troglodytes.clone.rp43-41g5.GenBank:AC099533.36      ------------------------------------------------------------ 68417
human.FAM30B-LOC105372935-GGT2.NCBI.                                                taattcatttcatcccaacctgactgatctctctgatgtctgcactgctgtgtcactaga 71867
human.FAM230E-LOC105377182-GGT3P.NCBI.                                              ------------------------------------------------------------ 57310
human.FAM230J-LOC105372942-GGTLC5P.NCBI.rev.compl                                   ------------------------------------------------------------ 46364
human.FAM230A.ncRNA.18487127.18500594-GGTLC3.18516335.18518165.rev.compl.REF        ------------------------------------------------------------ 31039


chimp.LOC112206744-LOC107973052-GGT2.revcompl                                       ------------------------------------------------------------ 61762
chimp54546-135457.revcompl.Pan.troglodytes.clone.rp43-41g5.GenBank:AC099533.36      ------------------------------------------------------------ 68417
human.FAM30B-LOC105372935-GGT2.NCBI.                                                ctgcccgattgtcctaggttgagacatttacctactgagccccagggatgggtggctgct 71927
human.FAM230E-LOC105377182-GGT3P.NCBI.                                              ------------------------------------------------------------ 57310
human.FAM230J-LOC105372942-GGTLC5P.NCBI.rev.compl                                   ------------------------------------------------------------ 46364
human.FAM230A.ncRNA.18487127.18500594-GGTLC3.18516335.18518165.rev.compl.REF        ------------------------------------------------------------ 31039


chimp.LOC112206744-LOC107973052-GGT2.revcompl                                       ------------------------------------------------------------ 61762
chimp54546-135457.revcompl.Pan.troglodytes.clone.rp43-41g5.GenBank:AC099533.36      ------------------------------------------------------------ 68417
human.FAM30B-LOC105372935-GGT2.NCBI.                                                tctaagctgctgcctaattccctcattactaaatgttgacgctgcttccttagaggactg 71987
human.FAM230E-LOC105377182-GGT3P.NCBI.                                              ------------------------------------------------------------ 57310
human.FAM230J-LOC105372942-GGTLC5P.NCBI.rev.compl                                   ------------------------------------------------------------ 46364
human.FAM230A.ncRNA.18487127.18500594-GGTLC3.18516335.18518165.rev.compl.REF        ------------------------------------------------------------ 31039
```

```
chimp.LOC112206744-LOC107973052-GGT2.revcompl                                 ------------------------------------------------------------ 61762
chimp54546-135457.revcompl.Pan.troglodytes.clone.rp43-41g5.GenBank:AC099533.36 ------------------------------------------------------------ 68417
human.FAM30B-LOC105372935-GGT2.NCBI.                                          ctaaaccataactcaaaaatgttttaaacatgctactatttacttagattagaatattta 72047
human.FAM230E-LOC105377182-GGT3P.NCBI.                                        ------------------------------------------------------------ 57310
human.FAM230J-LOC105372942-GGTLC5P.NCBI.rev.compl                             ------------------------------------------------------------ 46364
human.FAM230A.ncRNA.18487127.18500594-GGTLC3.18516335.18518165.rev.compl.REF  ------------------------------------------------------------ 31039

chimp.LOC112206744-LOC107973052-GGT2.revcompl                                 ------------------------------------------------------------ 61762
chimp54546-135457.revcompl.Pan.troglodytes.clone.rp43-41g5.GenBank:AC099533.36 ------------------------------------------------------------ 68417
human.FAM30B-LOC105372935-GGT2.NCBI.                                          cttaaatgttttaaaaaatacagtattttaagtcttttatctcttctgactactaccact 72107
human.FAM230E-LOC105377182-GGT3P.NCBI.                                        ------------------------------------------------------------ 57310
human.FAM230J-LOC105372942-GGTLC5P.NCBI.rev.compl                             ------------------------------------------------------------ 46364
human.FAM230A.ncRNA.18487127.18500594-GGTLC3.18516335.18518165.rev.compl.REF  ------------------------------------------------------------ 31039

chimp.LOC112206744-LOC107973052-GGT2.revcompl                                 ------------------------------------------------------------ 61762
chimp54546-135457.revcompl.Pan.troglodytes.clone.rp43-41g5.GenBank:AC099533.36 ------------------------------------------------------------ 68417
human.FAM30B-LOC105372935-GGT2.NCBI.                                          aaccctgcttcccctcattttccagagattagcactactgtgatttaaggatagagctc 72167
human.FAM230E-LOC105377182-GGT3P.NCBI.                                        ------------------------------------------------------------ 57310
human.FAM230J-LOC105372942-GGTLC5P.NCBI.rev.compl                             ------------------------------------------------------------ 46364
human.FAM230A.ncRNA.18487127.18500594-GGTLC3.18516335.18518165.rev.compl.REF  ------------------------------------------------------------ 31039

chimp.LOC112206744-LOC107973052-GGT2.revcompl                                 ------------------------------------------------------------ 61762
chimp54546-135457.revcompl.Pan.troglodytes.clone.rp43-41g5.GenBank:AC099533.36 ------------------------------------------------------------ 68417
human.FAM30B-LOC105372935-GGT2.NCBI.                                          tccagacttaaaaaaaaaccaaaaaactgcttttatatccagatatatgtaaccactgaa 72227
human.FAM230E-LOC105377182-GGT3P.NCBI.                                        ------------------------------------------------------------ 57310
human.FAM230J-LOC105372942-GGTLC5P.NCBI.rev.compl                             ------------------------------------------------------------ 46364
human.FAM230A.ncRNA.18487127.18500594-GGTLC3.18516335.18518165.rev.compl.REF  ------------------------------------------------------------ 31039

chimp.LOC112206744-LOC107973052-GGT2.revcompl                                 ------------------------------------------------------------ 61762
chimp54546-135457.revcompl.Pan.troglodytes.clone.rp43-41g5.GenBank:AC099533.36 ------------------------------------------------------------ 68417
human.FAM30B-LOC105372935-GGT2.NCBI.                                          aaatatacattatttttcagcatttttacataaacattatcatccctttcattgagtta 72287
human.FAM230E-LOC105377182-GGT3P.NCBI.                                        ------------------------------------------------------------ 57310
human.FAM230J-LOC105372942-GGTLC5P.NCBI.rev.compl                             ------------------------------------------------------------ 46364
human.FAM230A.ncRNA.18487127.18500594-GGTLC3.18516335.18518165.rev.compl.REF  ------------------------------------------------------------ 31039

chimp.LOC112206744-LOC107973052-GGT2.revcompl                                 ------------------------------------------------------------ 61762
chimp54546-135457.revcompl.Pan.troglodytes.clone.rp43-41g5.GenBank:AC099533.36 ------------------------------------------------------------ 68417
human.FAM30B-LOC105372935-GGT2.NCBI.                                          tgttcctcagttttcctcctcaaagtttttgagatcaaatatagttcccatttttcttct 72347
human.FAM230E-LOC105377182-GGT3P.NCBI.                                        ------------------------------------------------------------ 57310
human.FAM230J-LOC105372942-GGTLC5P.NCBI.rev.compl                             ------------------------------------------------------------ 46364
human.FAM230A.ncRNA.18487127.18500594-GGTLC3.18516335.18518165.rev.compl.REF  ------------------------------------------------------------ 31039

chimp.LOC112206744-LOC107973052-GGT2.revcompl                                 ------------------------------------------------------------ 61762
chimp54546-135457.revcompl.Pan.troglodytes.clone.rp43-41g5.GenBank:AC099533.36 ------------------------------------------------------------ 68417
human.FAM30B-LOC105372935-GGT2.NCBI.                                          ttatatagctgcacagtattccatcataggattacaccacattttatttattccaatatt 72407
human.FAM230E-LOC105377182-GGT3P.NCBI.                                        ------------------------------------------------------------ 57310
human.FAM230J-LOC105372942-GGTLC5P.NCBI.rev.compl                             ------------------------------------------------------------ 46364
human.FAM230A.ncRNA.18487127.18500594-GGTLC3.18516335.18518165.rev.compl.REF  ------------------------------------------------------------ 31039

chimp.LOC112206744-LOC107973052-GGT2.revcompl                                 ------------------------------------------------------------ 61762
chimp54546-135457.revcompl.Pan.troglodytes.clone.rp43-41g5.GenBank:AC099533.36 ------------------------------------------------------------ 68417
human.FAM30B-LOC105372935-GGT2.NCBI.                                          aatgaatatttaggttatttatacttttttttctataaccaacaagcccatgccttctac 72467
human.FAM230E-LOC105377182-GGT3P.NCBI.                                        ------------------------------------------------------------ 57310
human.FAM230J-LOC105372942-GGTLC5P.NCBI.rev.compl                             ------------------------------------------------------------ 46364
human.FAM230A.ncRNA.18487127.18500594-GGTLC3.18516335.18518165.rev.compl.REF  ------------------------------------------------------------ 31039

chimp.LOC112206744-LOC107973052-GGT2.revcompl                                 ------------------------------------------------------------ 61762
chimp54546-135457.revcompl.Pan.troglodytes.clone.rp43-41g5.GenBank:AC099533.36 ------------------------------------------------------------ 68417
human.FAM30B-LOC105372935-GGT2.NCBI.                                          tgcttatcctttccccacctgaatttgtacaatccccagcttctcttatccatgaccact 72527
human.FAM230E-LOC105377182-GGT3P.NCBI.                                        ------------------------------------------------------------ 57310
human.FAM230J-LOC105372942-GGTLC5P.NCBI.rev.compl                             ------------------------------------------------------------ 46364
human.FAM230A.ncRNA.18487127.18500594-GGTLC3.18516335.18518165.rev.compl.REF  ------------------------------------------------------------ 31039

chimp.LOC112206744-LOC107973052-GGT2.revcompl                                 ------------------------------------------------------------ 61762
chimp54546-135457.revcompl.Pan.troglodytes.clone.rp43-41g5.GenBank:AC099533.36 ------------------------------------------------------------ 68417
human.FAM30B-LOC105372935-GGT2.NCBI.                                          gaaaccccagggaagaccttatcactttcaggatgtccatggctctttcaccctatgcca 72587
human.FAM230E-LOC105377182-GGT3P.NCBI.                                        ------------------------------------------------------------ 57310
human.FAM230J-LOC105372942-GGTLC5P.NCBI.rev.compl                             ------------------------------------------------------------ 46364
human.FAM230A.ncRNA.18487127.18500594-GGTLC3.18516335.18518165.rev.compl.REF  ------------------------------------------------------------ 31039

chimp.LOC112206744-LOC107973052-GGT2.revcompl                                 ------------------------------------------------------------ 61762
chimp54546-135457.revcompl.Pan.troglodytes.clone.rp43-41g5.GenBank:AC099533.36 ------------------------------------------------------------ 68417
human.FAM30B-LOC105372935-GGT2.NCBI.                                          tgcttggaaccaggaaagcccaaaagaccccacttcctgcccttctattctgctcctcct 72647
human.FAM230E-LOC105377182-GGT3P.NCBI.                                        ------------------------------------------------------------ 57310
human.FAM230J-LOC105372942-GGTLC5P.NCBI.rev.compl                             ------------------------------------------------------------ 46364
human.FAM230A.ncRNA.18487127.18500594-GGTLC3.18516335.18518165.rev.compl.REF  ------------------------------------------------------------ 31039

chimp.LOC112206744-LOC107973052-GGT2.revcompl                                 ------------------------------------------------------------ 61762
chimp54546-135457.revcompl.Pan.troglodytes.clone.rp43-41g5.GenBank:AC099533.36 ------------------------------------------------------------ 68417
human.FAM30B-LOC105372935-GGT2.NCBI.                                          ctcctatcgctcatttctttgcctctatcctccactcctcccagcaaaggtagggatagc 72707
human.FAM230E-LOC105377182-GGT3P.NCBI.                                        ------------------------------------------------------------ 57310
human.FAM230J-LOC105372942-GGTLC5P.NCBI.rev.compl                             ------------------------------------------------------------ 46364
human.FAM230A.ncRNA.18487127.18500594-GGTLC3.18516335.18518165.rev.compl.REF  ------------------------------------------------------------ 31039

chimp.LOC112206744-LOC107973052-GGT2.revcompl                                 ------------------------------------------------------------ 61762
chimp54546-135457.revcompl.Pan.troglodytes.clone.rp43-41g5.GenBank:AC099533.36 ------------------------------------------------------------ 68417
human.FAM30B-LOC105372935-GGT2.NCBI.                                          acctgagtcctgccaccctctcacctgcattccccctcactctcatgcgaaggacagcac 72767
human.FAM230E-LOC105377182-GGT3P.NCBI.                                        ------------------------------------------------------------ 57310
human.FAM230J-LOC105372942-GGTLC5P.NCBI.rev.compl                             ------------------------------------------------------------ 46364
human.FAM230A.ncRNA.18487127.18500594-GGTLC3.18516335.18518165.rev.compl.REF  ------------------------------------------------------------ 31039

chimp.LOC112206744-LOC107973052-GGT2.revcompl                                 ------------------------------------------------------------ 61762
chimp54546-135457.revcompl.Pan.troglodytes.clone.rp43-41g5.GenBank:AC099533.36 ------------------------------------------------------------ 68417
human.FAM30B-LOC105372935-GGT2.NCBI.                                          agaagcagcatcaaaaaagaaaggcagcgagggagccctgcagcctgccctaggccagcc 72827
human.FAM230E-LOC105377182-GGT3P.NCBI.                                        ------------------------------------------------------------ 57310
human.FAM230J-LOC105372942-GGTLC5P.NCBI.rev.compl                             ------------------------------------------------------------ 46364
human.FAM230A.ncRNA.18487127.18500594-GGTLC3.18516335.18518165.rev.compl.REF  ------------------------------------------------------------ 31039

chimp.LOC112206744-LOC107973052-GGT2.revcompl                                 ------------------------------------------------------------ 61762
chimp54546-135457.revcompl.Pan.troglodytes.clone.rp43-41g5.GenBank:AC099533.36 ------------------------------------------------------------ 68417
human.FAM30B-LOC105372935-GGT2.NCBI.                                          atgcccacagccacaggaatagacactgttttcagccccacctagattcaaggcagcatc 72887
human.FAM230E-LOC105377182-GGT3P.NCBI.                                        ------------------------------------------------------------ 57310
human.FAM230J-LOC105372942-GGTLC5P.NCBI.rev.compl                             ------------------------------------------------------------ 46364
human.FAM230A.ncRNA.18487127.18500594-GGTLC3.18516335.18518165.rev.compl.REF  ------------------------------------------------------------ 31039

chimp.LOC112206744-LOC107973052-GGT2.revcompl                                 ------------------------------------------------------------ 61762
chimp54546-135457.revcompl.Pan.troglodytes.clone.rp43-41g5.GenBank:AC099533.36 ------------------------------------------------------------ 68417
human.FAM30B-LOC105372935-GGT2.NCBI.                                          taatagaaatgcttataagtaaatataagtgccagtatttgccttctattttctagaggg 72947
human.FAM230E-LOC105377182-GGT3P.NCBI.                                        ------------------------------------------------------------ 57310
human.FAM230J-LOC105372942-GGTLC5P.NCBI.rev.compl                             ------------------------------------------------------------ 46364
human.FAM230A.ncRNA.18487127.18500594-GGTLC3.18516335.18518165.rev.compl.REF  ------------------------------------------------------------ 31039

chimp.LOC112206744-LOC107973052-GGT2.revcompl                                 ------------------------------------------------------------ 61762
chimp54546-135457.revcompl.Pan.troglodytes.clone.rp43-41g5.GenBank:AC099533.36 ------------------------------------------------------------ 68417
human.FAM30B-LOC105372935-GGT2.NCBI.                                          gaaaaaaattaaaaggcaagatcagattacttgcctggaaatgttctccctgtaattact 73007
human.FAM230E-LOC105377182-GGT3P.NCBI.                                        ------------------------------------------------------------ 57310
human.FAM230J-LOC105372942-GGTLC5P.NCBI.rev.compl                             ------------------------------------------------------------ 46364
human.FAM230A.ncRNA.18487127.18500594-GGTLC3.18516335.18518165.rev.compl.REF  ------------------------------------------------------------ 31039

chimp.LOC112206744-LOC107973052-GGT2.revcompl                                 ------------------------------------------------------------ 61762
chimp54546-135457.revcompl.Pan.troglodytes.clone.rp43-41g5.GenBank:AC099533.36 ------------------------------------------------------------ 68417
human.FAM30B-LOC105372935-GGT2.NCBI.                                          ctgaccgctctttctttccctccggttttggagataatggaaataggctgagtgttgggc 73067
human.FAM230E-LOC105377182-GGT3P.NCBI.                                        ------------------------------------------------------------ 57310
human.FAM230J-LOC105372942-GGTLC5P.NCBI.rev.compl                             ------------------------------------------------------------ 46364
human.FAM230A.ncRNA.18487127.18500594-GGTLC3.18516335.18518165.rev.compl.REF  ------------------------------------------------------------ 31039
```

```
chimp.LOC112206744-LOC107973052-GGT2.revcompl  ------------------------------------------------------------ 61762
chimp54546-135457.revcompl.Pan.troglodytes.clone.rp43-41g5.GenBank:AC099533.36  ------------------------------------------------------------ 68417
human.FAM30B-LOC105372935-GGT2.NCBI.  atttttttttttaaagagggcagcttcatggcaacactccctctgagaagtccagaggct 73127
human.FAM230E-LOC105377182-GGT3P.NCBI.  ------------------------------------------------------------ 57310
human.FAM230J-LOC105372942-GGTLC5P.NCBI.rev.compl  ------------------------------------------------------------ 46364
human.FAM230A.ncRNA.18487127.18500594-GGTLC3.18516335.18518165.rev.compl.REF  ------------------------------------------------------------ 31039

chimp.LOC112206744-LOC107973052-GGT2.revcompl  ------------------------------------------------------------ 61762
chimp54546-135457.revcompl.Pan.troglodytes.clone.rp43-41g5.GenBank:AC099533.36  ------------------------------------------------------------ 68417
human.FAM30B-LOC105372935-GGT2.NCBI.  cagtgaaccgcagcttagagtgagttccagtaaaaggaccatcacttgaaccagtaagtg 73187
human.FAM230E-LOC105377182-GGT3P.NCBI.  ------------------------------------------------------------ 57310
human.FAM230J-LOC105372942-GGTLC5P.NCBI.rev.compl  ------------------------------------------------------------ 46364
human.FAM230A.ncRNA.18487127.18500594-GGTLC3.18516335.18518165.rev.compl.REF  ------------------------------------------------------------ 31039

chimp.LOC112206744-LOC107973052-GGT2.revcompl  ------------------------------------------------------------ 61762
chimp54546-135457.revcompl.Pan.troglodytes.clone.rp43-41g5.GenBank:AC099533.36  ------------------------------------------------------------ 68417
human.FAM30B-LOC105372935-GGT2.NCBI.  cctcattccttctcacagcatacatttcacagcaattactgttgctattatttacgctt 73247
human.FAM230E-LOC105377182-GGT3P.NCBI.  ------------------------------------------------------------ 57310
human.FAM230J-LOC105372942-GGTLC5P.NCBI.rev.compl  ------------------------------------------------------------ 46364
human.FAM230A.ncRNA.18487127.18500594-GGTLC3.18516335.18518165.rev.compl.REF  ------------------------------------------------------------ 31039

chimp.LOC112206744-LOC107973052-GGT2.revcompl  ------------------------------------------------------------ 61762
chimp54546-135457.revcompl.Pan.troglodytes.clone.rp43-41g5.GenBank:AC099533.36  ------------------------------------------------------------ 68417
human.FAM30B-LOC105372935-GGT2.NCBI.  catttctggacctaggatacctcagtttctccagactctcagtctggtccccaggttttc 73307
human.FAM230E-LOC105377182-GGT3P.NCBI.  ------------------------------------------------------------ 57310
human.FAM230J-LOC105372942-GGTLC5P.NCBI.rev.compl  ------------------------------------------------------------ 46364
human.FAM230A.ncRNA.18487127.18500594-GGTLC3.18516335.18518165.rev.compl.REF  ------------------------------------------------------------ 31039

chimp.LOC112206744-LOC107973052-GGT2.revcompl  ------------------------------------------------------------ 61762
chimp54546-135457.revcompl.Pan.troglodytes.clone.rp43-41g5.GenBank:AC099533.36  ------------------------------------------------------------ 68417
human.FAM30B-LOC105372935-GGT2.NCBI.  agtctgggagaaaggtgtctgctggatttcaaacagtccctctactggccgccaccacca 73367
human.FAM230E-LOC105377182-GGT3P.NCBI.  ------------------------------------------------------------ 57310
human.FAM230J-LOC105372942-GGTLC5P.NCBI.rev.compl  ------------------------------------------------------------ 46364
human.FAM230A.ncRNA.18487127.18500594-GGTLC3.18516335.18518165.rev.compl.REF  ------------------------------------------------------------ 31039

chimp.LOC112206744-LOC107973052-GGT2.revcompl  ------------------------------------------------------------ 61762
chimp54546-135457.revcompl.Pan.troglodytes.clone.rp43-41g5.GenBank:AC099533.36  ------------------------------------------------------------ 68417
human.FAM30B-LOC105372935-GGT2.NCBI.  ccctctcctaaagaggactggaataactcttccacagctcccgccacactcagcactgcc 73427
human.FAM230E-LOC105377182-GGT3P.NCBI.  ------------------------------------------------------------ 57310
human.FAM230J-LOC105372942-GGTLC5P.NCBI.rev.compl  ------------------------------------------------------------ 46364
human.FAM230A.ncRNA.18487127.18500594-GGTLC3.18516335.18518165.rev.compl.REF  ------------------------------------------------------------ 31039

chimp.LOC112206744-LOC107973052-GGT2.revcompl  ------------------------------------------------------------ 61762
chimp54546-135457.revcompl.Pan.troglodytes.clone.rp43-41g5.GenBank:AC099533.36  ------------------------------------------------------------ 68417
human.FAM30B-LOC105372935-GGT2.NCBI.  tgcacctcctcctggagctggcagggccccagctgtgcacccctcaatgctctcacagct 73487
human.FAM230E-LOC105377182-GGT3P.NCBI.  ------------------------------------------------------------ 57310
human.FAM230J-LOC105372942-GGTLC5P.NCBI.rev.compl  ------------------------------------------------------------ 46364
human.FAM230A.ncRNA.18487127.18500594-GGTLC3.18516335.18518165.rev.compl.REF  ------------------------------------------------------------ 31039

chimp.LOC112206744-LOC107973052-GGT2.revcompl  ------------------------------------------------------------ 61762
chimp54546-135457.revcompl.Pan.troglodytes.clone.rp43-41g5.GenBank:AC099533.36  ------------------------------------------------------------ 68417
human.FAM30B-LOC105372935-GGT2.NCBI.  tcaggaagaggactgctaagcccagggccctgctgacaacatgccgggagcgcatctgca 73547
human.FAM230E-LOC105377182-GGT3P.NCBI.  ------------------------------------------------------------ 57310
human.FAM230J-LOC105372942-GGTLC5P.NCBI.rev.compl  ------------------------------------------------------------ 46364
human.FAM230A.ncRNA.18487127.18500594-GGTLC3.18516335.18518165.rev.compl.REF  ------------------------------------------------------------ 31039

chimp.LOC112206744-LOC107973052-GGT2.revcompl  ------------------------------------------------------------ 61762
chimp54546-135457.revcompl.Pan.troglodytes.clone.rp43-41g5.GenBank:AC099533.36  ------------------------------------------------------------ 68417
human.FAM30B-LOC105372935-GGT2.NCBI.  ggcagggtcagaatgtcagctaccaggcagcagagagatgcggggcgccaacagcaaaag 73607
human.FAM230E-LOC105377182-GGT3P.NCBI.  ------------------------------------------------------------ 57310
human.FAM230J-LOC105372942-GGTLC5P.NCBI.rev.compl  ------------------------------------------------------------ 46364
human.FAM230A.ncRNA.18487127.18500594-GGTLC3.18516335.18518165.rev.compl.REF  ------------------------------------------------------------ 31039

chimp.LOC112206744-LOC107973052-GGT2.revcompl  ------------------------------------------------------------ 61762
chimp54546-135457.revcompl.Pan.troglodytes.clone.rp43-41g5.GenBank:AC099533.36  ------------------------------------------------------------ 68417
human.FAM30B-LOC105372935-GGT2.NCBI.  cacttgggggcttttttcaagtggcatcagaactagacctggatacaaaggggtggccct 73667
human.FAM230E-LOC105377182-GGT3P.NCBI.  ------------------------------------------------------------ 57310
human.FAM230J-LOC105372942-GGTLC5P.NCBI.rev.compl  ------------------------------------------------------------ 46364
human.FAM230A.ncRNA.18487127.18500594-GGTLC3.18516335.18518165.rev.compl.REF  ------------------------------------------------------------ 31039

chimp.LOC112206744-LOC107973052-GGT2.revcompl  ------------------------------------------------------------ 61762
chimp54546-135457.revcompl.Pan.troglodytes.clone.rp43-41g5.GenBank:AC099533.36  ------------------------------------------------------------ 68417
human.FAM30B-LOC105372935-GGT2.NCBI.  ggcttcaggaagcccatgataagtcagtctgctgtgtccttcaacacagtgtccccacct 73727
human.FAM230E-LOC105377182-GGT3P.NCBI.  ------------------------------------------------------------ 57310
human.FAM230J-LOC105372942-GGTLC5P.NCBI.rev.compl  ------------------------------------------------------------ 46364
human.FAM230A.ncRNA.18487127.18500594-GGTLC3.18516335.18518165.rev.compl.REF  ------------------------------------------------------------ 31039

chimp.LOC112206744-LOC107973052-GGT2.revcompl  ------------------------------------------------------------ 61762
chimp54546-135457.revcompl.Pan.troglodytes.clone.rp43-41g5.GenBank:AC099533.36  ------------------------------------------------------------ 68417
human.FAM30B-LOC105372935-GGT2.NCBI.  tctgtctcccaacactaggccatggagacacaaaactagaaaatggggctgcctagctac 73787
human.FAM230E-LOC105377182-GGT3P.NCBI.  ------------------------------------------------------------ 57310
human.FAM230J-LOC105372942-GGTLC5P.NCBI.rev.compl  ------------------------------------------------------------ 46364
human.FAM230A.ncRNA.18487127.18500594-GGTLC3.18516335.18518165.rev.compl.REF  ------------------------------------------------------------ 31039

chimp.LOC112206744-LOC107973052-GGT2.revcompl  ------------------------------------------------------------ 61762
chimp54546-135457.revcompl.Pan.troglodytes.clone.rp43-41g5.GenBank:AC099533.36  ------------------------------------------------------------ 68417
human.FAM30B-LOC105372935-GGT2.NCBI.  gaatgcccaggctgtgaattggggtctccaaaggaacctcagtgccgtgtcctactcaga 73847
human.FAM230E-LOC105377182-GGT3P.NCBI.  ------------------------------------------------------------ 57310
human.FAM230J-LOC105372942-GGTLC5P.NCBI.rev.compl  ------------------------------------------------------------ 46364
human.FAM230A.ncRNA.18487127.18500594-GGTLC3.18516335.18518165.rev.compl.REF  ------------------------------------------------------------ 31039

chimp.LOC112206744-LOC107973052-GGT2.revcompl  ------------------------------------------------------------ 61762
chimp54546-135457.revcompl.Pan.troglodytes.clone.rp43-41g5.GenBank:AC099533.36  ------------------------------------------------------------ 68417
human.FAM30B-LOC105372935-GGT2.NCBI.  gcagctccttcagctcttcaccctggaaagcactccggtacaaaacaggaatattcctct 73907
human.FAM230E-LOC105377182-GGT3P.NCBI.  ------------------------------------------------------------ 57310
human.FAM230J-LOC105372942-GGTLC5P.NCBI.rev.compl  ------------------------------------------------------------ 46364
human.FAM230A.ncRNA.18487127.18500594-GGTLC3.18516335.18518165.rev.compl.REF  ------------------------------------------------------------ 31039

chimp.LOC112206744-LOC107973052-GGT2.revcompl  ------------------------------------------------------------ 61762
chimp54546-135457.revcompl.Pan.troglodytes.clone.rp43-41g5.GenBank:AC099533.36  ------------------------------------------------------------ 68417
human.FAM30B-LOC105372935-GGT2.NCBI.  tctcctgactctgcccaaggcactgctgtgaccctctcagcagtggaagcactagccaat 73967
human.FAM230E-LOC105377182-GGT3P.NCBI.  ------------------------------------------------------------ 57310
human.FAM230J-LOC105372942-GGTLC5P.NCBI.rev.compl  ------------------------------------------------------------ 46364
human.FAM230A.ncRNA.18487127.18500594-GGTLC3.18516335.18518165.rev.compl.REF  ------------------------------------------------------------ 31039

chimp.LOC112206744-LOC107973052-GGT2.revcompl  ------------------------------------------------------------ 61762
chimp54546-135457.revcompl.Pan.troglodytes.clone.rp43-41g5.GenBank:AC099533.36  ------------------------------------------------------------ 68417
human.FAM30B-LOC105372935-GGT2.NCBI.  gtgccatcctcctggacgatagctggataatcgcaataagcagagccccaacaactagct 74027
human.FAM230E-LOC105377182-GGT3P.NCBI.  ------------------------------------------------------------ 57310
human.FAM230J-LOC105372942-GGTLC5P.NCBI.rev.compl  ------------------------------------------------------------ 46364
human.FAM230A.ncRNA.18487127.18500594-GGTLC3.18516335.18518165.rev.compl.REF  ------------------------------------------------------------ 31039

chimp.LOC112206744-LOC107973052-GGT2.revcompl  ------------------------------------------------------------ 61762
chimp54546-135457.revcompl.Pan.troglodytes.clone.rp43-41g5.GenBank:AC099533.36  ------------------------------------------------------------ 68417
human.FAM30B-LOC105372935-GGT2.NCBI.  ctagtgtgtggggagaagtggggaggatggagaaaggatacaaaaagacactgtgggatt 74087
human.FAM230E-LOC105377182-GGT3P.NCBI.  ------------------------------------------------------------ 57310
human.FAM230J-LOC105372942-GGTLC5P.NCBI.rev.compl  ------------------------------------------------------------ 46364
human.FAM230A.ncRNA.18487127.18500594-GGTLC3.18516335.18518165.rev.compl.REF  ------------------------------------------------------------ 31039

chimp.LOC112206744-LOC107973052-GGT2.revcompl  ------------------------------------------------------------ 61762
chimp54546-135457.revcompl.Pan.troglodytes.clone.rp43-41g5.GenBank:AC099533.36  ------------------------------------------------------------ 68417
human.FAM30B-LOC105372935-GGT2.NCBI.  tcagggcaacccatgagtgaggatcacagcagagctatgccttgcaaggaccagaactgg 74147
human.FAM230E-LOC105377182-GGT3P.NCBI.  ------------------------------------------------------------ 57310
human.FAM230J-LOC105372942-GGTLC5P.NCBI.rev.compl  ------------------------------------------------------------ 46364
human.FAM230A.ncRNA.18487127.18500594-GGTLC3.18516335.18518165.rev.compl.REF  ------------------------------------------------------------ 31039
```

```
chimp.LOC112206744-LOC107973052-GGT2.revcompl                                     ------------------------------------------------------------ 61762
chimp54546-135457.revcompl.Pan.troglodytes.clone.rp43-41g5.GenBank:AC099533.36    ------------------------------------------------------------ 68417
human.FAM30B-LOC105372935-GGT2.NCBI.                                              aaaacctttgtgacttagaccgccctcctcaggacagctcccgccactcttcagggacaa 74207
human.FAM230E-LOC105377182-GGT3P.NCBI.                                            ------------------------------------------------------------ 57310
human.FAM230J-LOC105372942-GGTLC5P.NCBI.rev.compl                                 ------------------------------------------------------------ 46364
human.FAM230A.ncRNA.18487127.18500594-GGTLC3.18516335.18518165.rev.compl.REF      ------------------------------------------------------------ 31039

chimp.LOC112206744-LOC107973052-GGT2.revcompl                                     ------------------------------------------------------------ 61762
chimp54546-135457.revcompl.Pan.troglodytes.clone.rp43-41g5.GenBank:AC099533.36    ------------------------------------------------------------ 68417
human.FAM30B-LOC105372935-GGT2.NCBI.                                              agttcctttactcagccatagtttctgtgctccaaatactcagcttacacatgcctgttg 74267
human.FAM230E-LOC105377182-GGT3P.NCBI.                                            ------------------------------------------------------------ 57310
human.FAM230J-LOC105372942-GGTLC5P.NCBI.rev.compl                                 ------------------------------------------------------------ 46364
human.FAM230A.ncRNA.18487127.18500594-GGTLC3.18516335.18518165.rev.compl.REF      ------------------------------------------------------------ 31039

chimp.LOC112206744-LOC107973052-GGT2.revcompl                                     ------------------------------------------------------------ 61762
chimp54546-135457.revcompl.Pan.troglodytes.clone.rp43-41g5.GenBank:AC099533.36    ------------------------------------------------------------ 68417
human.FAM30B-LOC105372935-GGT2.NCBI.                                              ctgctgcactcaccactgtggttctcagtgcaaattccccagaaagcagtctgactggct 74327
human.FAM230E-LOC105377182-GGT3P.NCBI.                                            ------------------------------------------------------------ 57310
human.FAM230J-LOC105372942-GGTLC5P.NCBI.rev.compl                                 ------------------------------------------------------------ 46364
human.FAM230A.ncRNA.18487127.18500594-GGTLC3.18516335.18518165.rev.compl.REF      ------------------------------------------------------------ 31039

chimp.LOC112206744-LOC107973052-GGT2.revcompl                                     ------------------------------------------------------------ 61762
chimp54546-135457.revcompl.Pan.troglodytes.clone.rp43-41g5.GenBank:AC099533.36    ------------------------------------------------------------ 68417
human.FAM30B-LOC105372935-GGT2.NCBI.                                              gggagacctttttcttttaaccaggcccccaaagttgtagattttggggtcatttacaa 74387
human.FAM230E-LOC105377182-GGT3P.NCBI.                                            ------------------------------------------------------------ 57310
human.FAM230J-LOC105372942-GGTLC5P.NCBI.rev.compl                                 ------------------------------------------------------------ 46364
human.FAM230A.ncRNA.18487127.18500594-GGTLC3.18516335.18518165.rev.compl.REF      ------------------------------------------------------------ 31039

chimp.LOC112206744-LOC107973052-GGT2.revcompl                                     ------------------------------------------------------------ 61762
chimp54546-135457.revcompl.Pan.troglodytes.clone.rp43-41g5.GenBank:AC099533.36    ------------------------------------------------------------ 68417
human.FAM30B-LOC105372935-GGT2.NCBI.                                              tatgcatttccccaacacactcagatctaggtggacttgaccagatgggcatctgtattc 74447
human.FAM230E-LOC105377182-GGT3P.NCBI.                                            ------------------------------------------------------------ 57310
human.FAM230J-LOC105372942-GGTLC5P.NCBI.rev.compl                                 ------------------------------------------------------------ 46364
human.FAM230A.ncRNA.18487127.18500594-GGTLC3.18516335.18518165.rev.compl.REF      ------------------------------------------------------------ 31039

chimp.LOC112206744-LOC107973052-GGT2.revcompl                                     ------------------------------------------------------------ 61762
chimp54546-135457.revcompl.Pan.troglodytes.clone.rp43-41g5.GenBank:AC099533.36    ------------------------------------------------------------ 68417
human.FAM30B-LOC105372935-GGT2.NCBI.                                              cattacccagcatataaccactttgggcaacagggaatgtgaagggccccaaaccgttcc 74507
human.FAM230E-LOC105377182-GGT3P.NCBI.                                            ------------------------------------------------------------ 57310
human.FAM230J-LOC105372942-GGTLC5P.NCBI.rev.compl                                 ------------------------------------------------------------ 46364
human.FAM230A.ncRNA.18487127.18500594-GGTLC3.18516335.18518165.rev.compl.REF      ------------------------------------------------------------ 31039

chimp.LOC112206744-LOC107973052-GGT2.revcompl                                     ------------------------------------------------------------ 61762
chimp54546-135457.revcompl.Pan.troglodytes.clone.rp43-41g5.GenBank:AC099533.36    ------------------------------------------------------------ 68417
human.FAM30B-LOC105372935-GGT2.NCBI.                                              ccaccaagagtgtgggtggcaggcaccctgaaatatgtctagcacactcataaagcatga 74567
human.FAM230E-LOC105377182-GGT3P.NCBI.                                            ------------------------------------------------------------ 57310
human.FAM230J-LOC105372942-GGTLC5P.NCBI.rev.compl                                 ------------------------------------------------------------ 46364
human.FAM230A.ncRNA.18487127.18500594-GGTLC3.18516335.18518165.rev.compl.REF      ------------------------------------------------------------ 31039

chimp.LOC112206744-LOC107973052-GGT2.revcompl                                     ------------------------------------------------------------ 61762
chimp54546-135457.revcompl.Pan.troglodytes.clone.rp43-41g5.GenBank:AC099533.36    ------------------------------------------------------------ 68417
human.FAM30B-LOC105372935-GGT2.NCBI.                                              acatctacacacaactaaaaataaatagtgagagccctgacactgcttctgggggttttc 74627
human.FAM230E-LOC105377182-GGT3P.NCBI.                                            ------------------------------------------------------------ 57310
human.FAM230J-LOC105372942-GGTLC5P.NCBI.rev.compl                                 ------------------------------------------------------------ 46364
human.FAM230A.ncRNA.18487127.18500594-GGTLC3.18516335.18518165.rev.compl.REF      ------------------------------------------------------------ 31039

chimp.LOC112206744-LOC107973052-GGT2.revcompl                                     ------------------------------------------------------------ 61762
chimp54546-135457.revcompl.Pan.troglodytes.clone.rp43-41g5.GenBank:AC099533.36    ------------------------------------------------------------ 68417
human.FAM30B-LOC105372935-GGT2.NCBI.                                              ccaagcacctgttacgaatggtaacaatggatggccgctccttctccctcagctttgact 74687
human.FAM230E-LOC105377182-GGT3P.NCBI.                                            ------------------------------------------------------------ 57310
human.FAM230J-LOC105372942-GGTLC5P.NCBI.rev.compl                                 ------------------------------------------------------------ 46364
human.FAM230A.ncRNA.18487127.18500594-GGTLC3.18516335.18518165.rev.compl.REF      ------------------------------------------------------------ 31039

chimp.LOC112206744-LOC107973052-GGT2.revcompl                                     ------------------------------------------------------------ 61762
chimp54546-135457.revcompl.Pan.troglodytes.clone.rp43-41g5.GenBank:AC099533.36    ------------------------------------------------------------ 68417
human.FAM30B-LOC105372935-GGT2.NCBI.                                              ttcattgctggtggctcaggctcctgtgcttggctgtcatcggtgggtagccctggattg 74747
human.FAM230E-LOC105377182-GGT3P.NCBI.                                            ------------------------------------------------------------ 57310
human.FAM230J-LOC105372942-GGTLC5P.NCBI.rev.compl                                 ------------------------------------------------------------ 46364
human.FAM230A.ncRNA.18487127.18500594-GGTLC3.18516335.18518165.rev.compl.REF      ------------------------------------------------------------ 31039

chimp.LOC112206744-LOC107973052-GGT2.revcompl                                     ------------------------------------------------------------ 61762
chimp54546-135457.revcompl.Pan.troglodytes.clone.rp43-41g5.GenBank:AC099533.36    ------------------------------------------------------------ 68417
human.FAM30B-LOC105372935-GGT2.NCBI.                                              gacattttccctattcaattcaaggaccatgttcttgagcataactatgaccaggcagta 74807
human.FAM230E-LOC105377182-GGT3P.NCBI.                                            ------------------------------------------------------------ 57310
human.FAM230J-LOC105372942-GGTLC5P.NCBI.rev.compl                                 ------------------------------------------------------------ 46364
human.FAM230A.ncRNA.18487127.18500594-GGTLC3.18516335.18518165.rev.compl.REF      ------------------------------------------------------------ 31039

chimp.LOC112206744-LOC107973052-GGT2.revcompl                                     ------------------------------------------------------------ 61762
chimp54546-135457.revcompl.Pan.troglodytes.clone.rp43-41g5.GenBank:AC099533.36    ------------------------------------------------------------ 68417
human.FAM30B-LOC105372935-GGT2.NCBI.                                              ggcagtgggccctgctctttctttccagggaggcacagataagccagcatcacagatgca 74867
human.FAM230E-LOC105377182-GGT3P.NCBI.                                            ------------------------------------------------------------ 57310
human.FAM230J-LOC105372942-GGTLC5P.NCBI.rev.compl                                 ------------------------------------------------------------ 46364
human.FAM230A.ncRNA.18487127.18500594-GGTLC3.18516335.18518165.rev.compl.REF      ------------------------------------------------------------ 31039

chimp.LOC112206744-LOC107973052-GGT2.revcompl                                     ------------------------------------------------------------ 61762
chimp54546-135457.revcompl.Pan.troglodytes.clone.rp43-41g5.GenBank:AC099533.36    ------------------------------------------------------------ 68417
human.FAM30B-LOC105372935-GGT2.NCBI.                                              aaaaacagaaccgtggaggtagagactaattcttcagcagcccagagaagagaactggga 74927
human.FAM230E-LOC105377182-GGT3P.NCBI.                                            ------------------------------------------------------------ 57310
human.FAM230J-LOC105372942-GGTLC5P.NCBI.rev.compl                                 ------------------------------------------------------------ 46364
human.FAM230A.ncRNA.18487127.18500594-GGTLC3.18516335.18518165.rev.compl.REF      ------------------------------------------------------------ 31039

chimp.LOC112206744-LOC107973052-GGT2.revcompl                                     ------------------------------------------------------------ 61762
chimp54546-135457.revcompl.Pan.troglodytes.clone.rp43-41g5.GenBank:AC099533.36    ------------------------------------------------------------ 68417
human.FAM30B-LOC105372935-GGT2.NCBI.                                              aacagggccctcgcagatacacaagaccaagctggagagttcagggacttccaaaagaag 74987
human.FAM230E-LOC105377182-GGT3P.NCBI.                                            ------------------------------------------------------------ 57310
human.FAM230J-LOC105372942-GGTLC5P.NCBI.rev.compl                                 ------------------------------------------------------------ 46364
human.FAM230A.ncRNA.18487127.18500594-GGTLC3.18516335.18518165.rev.compl.REF      ------------------------------------------------------------ 31039

chimp.LOC112206744-LOC107973052-GGT2.revcompl                                     ------------------------------------------------------------ 61762
chimp54546-135457.revcompl.Pan.troglodytes.clone.rp43-41g5.GenBank:AC099533.36    ------------------------------------------------------------ 68417
human.FAM30B-LOC105372935-GGT2.NCBI.                                              gtacaagaatgagcactataccatagtggggtgggtcacaaaagcaggaaaaggtcccca 75047
human.FAM230E-LOC105377182-GGT3P.NCBI.                                            ------------------------------------------------------------ 57310
human.FAM230J-LOC105372942-GGTLC5P.NCBI.rev.compl                                 ------------------------------------------------------------ 46364
human.FAM230A.ncRNA.18487127.18500594-GGTLC3.18516335.18518165.rev.compl.REF      ------------------------------------------------------------ 31039

chimp.LOC112206744-LOC107973052-GGT2.revcompl                                     ------------------------------------------------------------ 61762
chimp54546-135457.revcompl.Pan.troglodytes.clone.rp43-41g5.GenBank:AC099533.36    ------------------------------------------------------------ 68417
human.FAM30B-LOC105372935-GGT2.NCBI.                                              aacaaagaatccatcagtcagttacagcactctccacatgtggtgaaacagtatgcaacc 75107
human.FAM230E-LOC105377182-GGT3P.NCBI.                                            ------------------------------------------------------------ 57310
human.FAM230J-LOC105372942-GGTLC5P.NCBI.rev.compl                                 ------------------------------------------------------------ 46364
human.FAM230A.ncRNA.18487127.18500594-GGTLC3.18516335.18518165.rev.compl.REF      ------------------------------------------------------------ 31039

chimp.LOC112206744-LOC107973052-GGT2.revcompl                                     ------------------------------------------------------------ 61762
chimp54546-135457.revcompl.Pan.troglodytes.clone.rp43-41g5.GenBank:AC099533.36    ------------------------------------------------------------ 68417
human.FAM30B-LOC105372935-GGT2.NCBI.                                              atttaaaaactgcagagaacaatatttatgaacacctaaaaatgctgaaagcacactact 75167
human.FAM230E-LOC105377182-GGT3P.NCBI.                                            ------------------------------------------------------------ 57310
human.FAM230J-LOC105372942-GGTLC5P.NCBI.rev.compl                                 ------------------------------------------------------------ 46364
human.FAM230A.ncRNA.18487127.18500594-GGTLC3.18516335.18518165.rev.compl.REF      ------------------------------------------------------------ 31039

chimp.LOC112206744-LOC107973052-GGT2.revcompl                                     ------------------------------------------------------------ 61762
chimp54546-135457.revcompl.Pan.troglodytes.clone.rp43-41g5.GenBank:AC099533.36    ------------------------------------------------------------ 68417
human.FAM30B-LOC105372935-GGT2.NCBI.                                              aagtaaaaggggagaaaggctataaaaggatttgaaagtattttcgtaaagataaacacc 75227
human.FAM230E-LOC105377182-GGT3P.NCBI.                                            ------------------------------------------------------------ 57310
human.FAM230J-LOC105372942-GGTLC5P.NCBI.rev.compl                                 ------------------------------------------------------------ 46364
human.FAM230A.ncRNA.18487127.18500594-GGTLC3.18516335.18518165.rev.compl.REF      ------------------------------------------------------------ 31039

chimp.LOC112206744-LOC107973052-GGT2.revcompl                                     ------------------------------------------------------------ 61762
```

```
chimp54546-135457.revcompl.Pan.troglodytes.clone.rp43-41g5.GenBank:AC099533.36      ------------------------------------------------------------ 68417
human.FAM30B-LOC105372935-GGT2.NCBI.                                                cacacacacccacacacctctatatatgtgtggaaaaaggactgggctagtatatttaaa 75287
human.FAM230E-LOC105377182-GGT3P.NCBI.                                              ------------------------------------------------------------ 57310
human.FAM230J-LOC105372942-GGTLC5P.NCBI.rev.compl                                   ------------------------------------------------------------ 46364
human.FAM230A.ncRNA.18487127.18500594-GGTLC3.18516335.18518165.rev.compl.REF        ------------------------------------------------------------ 31039

chimp.LOC112206744-LOC107973052-GGT2.revcompl                                       ------------------------------------------------------------ 61762
chimp54546-135457.revcompl.Pan.troglodytes.clone.rp43-41g5.GenBank:AC099533.36      ------------------------------------------------------------ 68417
human.FAM30B-LOC105372935-GGT2.NCBI.                                                aaacgctaaagggaggtcgggtaataagggattgtttttccttgcactttgtgttttct 75347
human.FAM230E-LOC105377182-GGT3P.NCBI.                                              ------------------------------------------------------------ 57310
human.FAM230J-LOC105372942-GGTLC5P.NCBI.rev.compl                                   ------------------------------------------------------------ 46364
human.FAM230A.ncRNA.18487127.18500594-GGTLC3.18516335.18518165.rev.compl.REF        ------------------------------------------------------------ 31039

chimp.LOC112206744-LOC107973052-GGT2.revcompl                                       ------------------------------------------------------------ 61762
chimp54546-135457.revcompl.Pan.troglodytes.clone.rp43-41g5.GenBank:AC099533.36      ------------------------------------------------------------ 68417
human.FAM30B-LOC105372935-GGT2.NCBI.                                                aagtattctatagtctgaatgaattagattcataattaggaccaacagtcataagcaatt 75407
human.FAM230E-LOC105377182-GGT3P.NCBI.                                              ------------------------------------------------------------ 57310
human.FAM230J-LOC105372942-GGTLC5P.NCBI.rev.compl                                   ------------------------------------------------------------ 46364
human.FAM230A.ncRNA.18487127.18500594-GGTLC3.18516335.18518165.rev.compl.REF        ------------------------------------------------------------ 31039

chimp.LOC112206744-LOC107973052-GGT2.revcompl                                       ------------------------------------------------------------ 61762
chimp54546-135457.revcompl.Pan.troglodytes.clone.rp43-41g5.GenBank:AC099533.36      ------------------------------------------------------------ 68417
human.FAM30B-LOC105372935-GGT2.NCBI.                                                tctcagaccacctgcctcataaacccaaaaaaagaagtcagagaggggtgaatattttg 75467
human.FAM230E-LOC105377182-GGT3P.NCBI.                                              ------------------------------------------------------------ 57310
human.FAM230J-LOC105372942-GGTLC5P.NCBI.rev.compl                                   ------------------------------------------------------------ 46364
human.FAM230A.ncRNA.18487127.18500594-GGTLC3.18516335.18518165.rev.compl.REF        ------------------------------------------------------------ 31039

chimp.LOC112206744-LOC107973052-GGT2.revcompl                                       ------------------------------------------------------------ 61762
chimp54546-135457.revcompl.Pan.troglodytes.clone.rp43-41g5.GenBank:AC099533.36      ------------------------------------------------------------ 68417
human.FAM30B-LOC105372935-GGT2.NCBI.                                                gtacagatgaattcaggttgtgtaagtggagaggagggaggtggccaaggagcctgaagt 75527
human.FAM230E-LOC105377182-GGT3P.NCBI.                                              ------------------------------------------------------------ 57310
human.FAM230J-LOC105372942-GGTLC5P.NCBI.rev.compl                                   ------------------------------------------------------------ 46364
human.FAM230A.ncRNA.18487127.18500594-GGTLC3.18516335.18518165.rev.compl.REF        ------------------------------------------------------------ 31039

chimp.LOC112206744-LOC107973052-GGT2.revcompl                                       ------------------------------------------------------------ 61762
chimp54546-135457.revcompl.Pan.troglodytes.clone.rp43-41g5.GenBank:AC099533.36      ------------------------------------------------------------ 68417
human.FAM30B-LOC105372935-GGT2.NCBI.                                                gggggaggggggctgatggtgaaatagggaatgggtggagagagaaaatgattgcaaaat 75587
human.FAM230E-LOC105377182-GGT3P.NCBI.                                              ------------------------------------------------------------ 57310
human.FAM230J-LOC105372942-GGTLC5P.NCBI.rev.compl                                   ------------------------------------------------------------ 46364
human.FAM230A.ncRNA.18487127.18500594-GGTLC3.18516335.18518165.rev.compl.REF        ------------------------------------------------------------ 31039

chimp.LOC112206744-LOC107973052-GGT2.revcompl                                       ------------------------------------------------------------ 61762
chimp54546-135457.revcompl.Pan.troglodytes.clone.rp43-41g5.GenBank:AC099533.36      ------------------------------------------------------------ 68417
human.FAM30B-LOC105372935-GGT2.NCBI.                                                tggctggagccagcctggtcaggtcaactttgcaggtgtgtgaagtgtcaaatggttaat 75647
human.FAM230E-LOC105377182-GGT3P.NCBI.                                              ------------------------------------------------------------ 57310
human.FAM230J-LOC105372942-GGTLC5P.NCBI.rev.compl                                   ------------------------------------------------------------ 46364
human.FAM230A.ncRNA.18487127.18500594-GGTLC3.18516335.18518165.rev.compl.REF        ------------------------------------------------------------ 31039

chimp.LOC112206744-LOC107973052-GGT2.revcompl                                       ------------------------------------------------------------ 61762
chimp54546-135457.revcompl.Pan.troglodytes.clone.rp43-41g5.GenBank:AC099533.36      ------------------------------------------------------------ 68417
human.FAM30B-LOC105372935-GGT2.NCBI.                                                cagttgccctctgtttgagctgcttaatgcacaagggcctggttgaacaggtttgacctc 75707
human.FAM230E-LOC105377182-GGT3P.NCBI.                                              ------------------------------------------------------------ 57310
human.FAM230J-LOC105372942-GGTLC5P.NCBI.rev.compl                                   ------------------------------------------------------------ 46364
human.FAM230A.ncRNA.18487127.18500594-GGTLC3.18516335.18518165.rev.compl.REF        ------------------------------------------------------------ 31039

chimp.LOC112206744-LOC107973052-GGT2.revcompl                                       ------------------------------------------------------------ 61762
chimp54546-135457.revcompl.Pan.troglodytes.clone.rp43-41g5.GenBank:AC099533.36      ------------------------------------------------------------ 68417
human.FAM30B-LOC105372935-GGT2.NCBI.                                                atttacttttcttaaagctgtttagccactttcctaaaaactcctattttctgagttat 75767
human.FAM230E-LOC105377182-GGT3P.NCBI.                                              ------------------------------------------------------------ 57310
human.FAM230J-LOC105372942-GGTLC5P.NCBI.rev.compl                                   ------------------------------------------------------------ 46364
human.FAM230A.ncRNA.18487127.18500594-GGTLC3.18516335.18518165.rev.compl.REF        ------------------------------------------------------------ 31039

chimp.LOC112206744-LOC107973052-GGT2.revcompl                                       ------------------------------------------------------------ 61762
chimp54546-135457.revcompl.Pan.troglodytes.clone.rp43-41g5.GenBank:AC099533.36      ------------------------------------------------------------ 68417
human.FAM30B-LOC105372935-GGT2.NCBI.                                                taaactttccattccatgttgatcccttccgagccctctaacgcatctcttctcttgctt 75827
human.FAM230E-LOC105377182-GGT3P.NCBI.                                              ------------------------------------------------------------ 57310
human.FAM230J-LOC105372942-GGTLC5P.NCBI.rev.compl                                   ------------------------------------------------------------ 46364
human.FAM230A.ncRNA.18487127.18500594-GGTLC3.18516335.18518165.rev.compl.REF        ------------------------------------------------------------ 31039

chimp.LOC112206744-LOC107973052-GGT2.revcompl                                       ------------------------------------------------------------ 61762
chimp54546-135457.revcompl.Pan.troglodytes.clone.rp43-41g5.GenBank:AC099533.36      ------------------------------------------------------------ 68417
human.FAM30B-LOC105372935-GGT2.NCBI.                                                cttccaactggatgccttcatgaggcattaacaggtactgtgagccctgatgaaaatcat 75887
human.FAM230E-LOC105377182-GGT3P.NCBI.                                              ------------------------------------------------------------ 57310
human.FAM230J-LOC105372942-GGTLC5P.NCBI.rev.compl                                   ------------------------------------------------------------ 46364
human.FAM230A.ncRNA.18487127.18500594-GGTLC3.18516335.18518165.rev.compl.REF        ------------------------------------------------------------ 31039

chimp.LOC112206744-LOC107973052-GGT2.revcompl                                       ------------------------------------------------------------ 61762
chimp54546-135457.revcompl.Pan.troglodytes.clone.rp43-41g5.GenBank:AC099533.36      ------------------------------------------------------------ 68417
human.FAM30B-LOC105372935-GGT2.NCBI.                                                cctttcagggtacaaggaaggcacctgaatgacagatggttggaggaatcttgagaaagc 75947
human.FAM230E-LOC105377182-GGT3P.NCBI.                                              ------------------------------------------------------------ 57310
human.FAM230J-LOC105372942-GGTLC5P.NCBI.rev.compl                                   ------------------------------------------------------------ 46364
human.FAM230A.ncRNA.18487127.18500594-GGTLC3.18516335.18518165.rev.compl.REF        ------------------------------------------------------------ 31039

chimp.LOC112206744-LOC107973052-GGT2.revcompl                                       ------------------------------------------------------------ 61762
chimp54546-135457.revcompl.Pan.troglodytes.clone.rp43-41g5.GenBank:AC099533.36      ------------------------------------------------------------ 68417
human.FAM30B-LOC105372935-GGT2.NCBI.                                                agtttaggcattaacatgcaggctgtcgctgacttacacagagccaaggacgtgacatac 76007
human.FAM230E-LOC105377182-GGT3P.NCBI.                                              ------------------------------------------------------------ 57310
human.FAM230J-LOC105372942-GGTLC5P.NCBI.rev.compl                                   ------------------------------------------------------------ 46364
human.FAM230A.ncRNA.18487127.18500594-GGTLC3.18516335.18518165.rev.compl.REF        ------------------------------------------------------------ 31039

chimp.LOC112206744-LOC107973052-GGT2.revcompl                                       ------------------------------------------------------------ 61762
chimp54546-135457.revcompl.Pan.troglodytes.clone.rp43-41g5.GenBank:AC099533.36      ------------------------------------------------------------ 68417
human.FAM30B-LOC105372935-GGT2.NCBI.                                                agcatggggccatttctaaatccccactgaggacaagggtcctggagggaatgggatttg 76067
human.FAM230E-LOC105377182-GGT3P.NCBI.                                              ------------------------------------------------------------ 57310
human.FAM230J-LOC105372942-GGTLC5P.NCBI.rev.compl                                   ------------------------------------------------------------ 46364
human.FAM230A.ncRNA.18487127.18500594-GGTLC3.18516335.18518165.rev.compl.REF        ------------------------------------------------------------ 31039

chimp.LOC112206744-LOC107973052-GGT2.revcompl                                       ------------------------------------------------------------ 61762
chimp54546-135457.revcompl.Pan.troglodytes.clone.rp43-41g5.GenBank:AC099533.36      ------------------------------------------------------------ 68417
human.FAM30B-LOC105372935-GGT2.NCBI.                                                gaacaagatttgggctttgaacattcgctacaggctctgacaaagcagtgatggttcaaa 76127
human.FAM230E-LOC105377182-GGT3P.NCBI.                                              ------------------------------------------------------------ 57310
human.FAM230J-LOC105372942-GGTLC5P.NCBI.rev.compl                                   ------------------------------------------------------------ 46364
human.FAM230A.ncRNA.18487127.18500594-GGTLC3.18516335.18518165.rev.compl.REF        ------------------------------------------------------------ 31039

chimp.LOC112206744-LOC107973052-GGT2.revcompl                                       ------------------------------------------------------------ 61762
chimp54546-135457.revcompl.Pan.troglodytes.clone.rp43-41g5.GenBank:AC099533.36      ------------------------------------------------------------ 68417
human.FAM30B-LOC105372935-GGT2.NCBI.                                                agcctctgccccctggtggcaagccctgcccagatgctgttaaggggctctccaggcacg 76187
human.FAM230E-LOC105377182-GGT3P.NCBI.                                              ------------------------------------------------------------ 57310
human.FAM230J-LOC105372942-GGTLC5P.NCBI.rev.compl                                   ------------------------------------------------------------ 46364
human.FAM230A.ncRNA.18487127.18500594-GGTLC3.18516335.18518165.rev.compl.REF        ------------------------------------------------------------ 31039

chimp.LOC112206744-LOC107973052-GGT2.revcompl                                       ------------------------------------------------------------ 61762
chimp54546-135457.revcompl.Pan.troglodytes.clone.rp43-41g5.GenBank:AC099533.36      ------------------------------------------------------------ 68417
human.FAM30B-LOC105372935-GGT2.NCBI.                                                ccctcctaattaggggcaaaatcagagaagcctctttggaagaagacaaaaggcacaatg 76247
human.FAM230E-LOC105377182-GGT3P.NCBI.                                              ------------------------------------------------------------ 57310
human.FAM230J-LOC105372942-GGTLC5P.NCBI.rev.compl                                   ------------------------------------------------------------ 46364
human.FAM230A.ncRNA.18487127.18500594-GGTLC3.18516335.18518165.rev.compl.REF        ------------------------------------------------------------ 31039

chimp.LOC112206744-LOC107973052-GGT2.revcompl                                       ------------------------------------------------------------ 61762
chimp54546-135457.revcompl.Pan.troglodytes.clone.rp43-41g5.GenBank:AC099533.36      ------------------------------------------------------------ 68417
human.FAM30B-LOC105372935-GGT2.NCBI.                                                agatgcggtctcctgaagtagaggctttctctctggttatcacctgaaacaggaaataag 76307
human.FAM230E-LOC105377182-GGT3P.NCBI.                                              ------------------------------------------------------------ 57310
human.FAM230J-LOC105372942-GGTLC5P.NCBI.rev.compl                                   ------------------------------------------------------------ 46364
human.FAM230A.ncRNA.18487127.18500594-GGTLC3.18516335.18518165.rev.compl.REF        ------------------------------------------------------------ 31039

chimp.LOC112206744-LOC107973052-GGT2.revcompl                                       ------------------------------------------------------------ 61762
chimp54546-135457.revcompl.Pan.troglodytes.clone.rp43-41g5.GenBank:AC099533.36      ------------------------------------------------------------ 68417
```

```
human.FAM30B-LOC105372935-GGT2.NCBI.                                      gactgagtacctgctcaattccaggcactagccaaatctgcagcaaactagcatttccaa 76367
human.FAM230E-LOC105377182-GGT3P.NCBI.                                    ------------------------------------------------------------ 57310
human.FAM230J-LOC105372942-GGTLC5P.NCBI.rev.compl                         ------------------------------------------------------------ 46364
human.FAM230A.ncRNA.18487127.18500594-GGTLC3.18516335.18518165.rev.compl.REF ------------------------------------------------------------ 31039

chimp.LOC112206744-LOC107973052-GGT2.revcompl                             ------------------------------------------------------------ 61762
chimp54546-135457.revcompl.Pan.troglodytes.clone.rp43-41g5.GenBank:AC099533.36 ------------------------------------------------------------ 68417
human.FAM30B-LOC105372935-GGT2.NCBI.                                      aggtgtcttcttactttaatctcacagccaccctgtgagggaagtgtacattccccattt 76427
human.FAM230E-LOC105377182-GGT3P.NCBI.                                    ------------------------------------------------------------ 57310
human.FAM230J-LOC105372942-GGTLC5P.NCBI.rev.compl                         ------------------------------------------------------------ 46364
human.FAM230A.ncRNA.18487127.18500594-GGTLC3.18516335.18518165.rev.compl.REF ------------------------------------------------------------ 31039

chimp.LOC112206744-LOC107973052-GGT2.revcompl                             ------------------------------------------------------------ 61762
chimp54546-135457.revcompl.Pan.troglodytes.clone.rp43-41g5.GenBank:AC099533.36 ------------------------------------------------------------ 68417
human.FAM30B-LOC105372935-GGT2.NCBI.                                      cgctgatgagatgagcaaactgaggctcggagaaattgtcagtgggatacagagttggca 76487
human.FAM230E-LOC105377182-GGT3P.NCBI.                                    ------------------------------------------------------------ 57310
human.FAM230J-LOC105372942-GGTLC5P.NCBI.rev.compl                         ------------------------------------------------------------ 46364
human.FAM230A.ncRNA.18487127.18500594-GGTLC3.18516335.18518165.rev.compl.REF ------------------------------------------------------------ 31039

chimp.LOC112206744-LOC107973052-GGT2.revcompl                             ------------------------------------------------------------ 61762
chimp54546-135457.revcompl.Pan.troglodytes.clone.rp43-41g5.GenBank:AC099533.36 ------------------------------------------------------------ 68417
human.FAM30B-LOC105372935-GGT2.NCBI.                                      tccaaacccacaatctctgactctaagcttctgtttcattcattccacaaatatatatgg 76547
human.FAM230E-LOC105377182-GGT3P.NCBI.                                    ------------------------------------------------------------ 57310
human.FAM230J-LOC105372942-GGTLC5P.NCBI.rev.compl                         ------------------------------------------------------------ 46364
human.FAM230A.ncRNA.18487127.18500594-GGTLC3.18516335.18518165.rev.compl.REF ------------------------------------------------------------ 31039

chimp.LOC112206744-LOC107973052-GGT2.revcompl                             ------------------------------------------------------------ 61762
chimp54546-135457.revcompl.Pan.troglodytes.clone.rp43-41g5.GenBank:AC099533.36 ------------------------------------------------------------ 68417
human.FAM30B-LOC105372935-GGT2.NCBI.                                      tgtctcctctttgctggacactgtactgggctgatgagctacaagacccaatctttacag 76607
human.FAM230E-LOC105377182-GGT3P.NCBI.                                    ------------------------------------------------------------ 57310
human.FAM230J-LOC105372942-GGTLC5P.NCBI.rev.compl                         ------------------------------------------------------------ 46364
human.FAM230A.ncRNA.18487127.18500594-GGTLC3.18516335.18518165.rev.compl.REF ------------------------------------------------------------ 31039

chimp.LOC112206744-LOC107973052-GGT2.revcompl                             ------------------------------------------------------------ 61762
chimp54546-135457.revcompl.Pan.troglodytes.clone.rp43-41g5.GenBank:AC099533.36 ------------------------------------------------------------ 68417
human.FAM30B-LOC105372935-GGT2.NCBI.                                      tctaggaggcagagatagataagtaaacagaggcggcaaagcgtggcactgtgaaagatg 76667
human.FAM230E-LOC105377182-GGT3P.NCBI.                                    ------------------------------------------------------------ 57310
human.FAM230J-LOC105372942-GGTLC5P.NCBI.rev.compl                         ------------------------------------------------------------ 46364
human.FAM230A.ncRNA.18487127.18500594-GGTLC3.18516335.18518165.rev.compl.REF ------------------------------------------------------------ 31039

chimp.LOC112206744-LOC107973052-GGT2.revcompl                             ------------------------------------------------------------ 61762
chimp54546-135457.revcompl.Pan.troglodytes.clone.rp43-41g5.GenBank:AC099533.36 ------------------------------------------------------------ 68417
human.FAM30B-LOC105372935-GGT2.NCBI.                                      cactgaggacaccgagctaggtgttccacaggcgtcaggaaaggcttcacacaagagagg 76727
human.FAM230E-LOC105377182-GGT3P.NCBI.                                    ------------------------------------------------------------ 57310
human.FAM230J-LOC105372942-GGTLC5P.NCBI.rev.compl                         ------------------------------------------------------------ 46364
human.FAM230A.ncRNA.18487127.18500594-GGTLC3.18516335.18518165.rev.compl.REF ------------------------------------------------------------ 31039

chimp.LOC112206744-LOC107973052-GGT2.revcompl                             ------------------------------------------------------------ 61762
chimp54546-135457.revcompl.Pan.troglodytes.clone.rp43-41g5.GenBank:AC099533.36 ------------------------------------------------------------ 68417
human.FAM30B-LOC105372935-GGT2.NCBI.                                      ttgcttgtgcccagttctaattaggatgagtaggtatttctgctagatcgtggtggggtg 76787
human.FAM230E-LOC105377182-GGT3P.NCBI.                                    ------------------------------------------------------------ 57310
human.FAM230J-LOC105372942-GGTLC5P.NCBI.rev.compl                         ------------------------------------------------------------ 46364
human.FAM230A.ncRNA.18487127.18500594-GGTLC3.18516335.18518165.rev.compl.REF ------------------------------------------------------------ 31039

chimp.LOC112206744-LOC107973052-GGT2.revcompl                             ------------------------------------------------------------ 61762
chimp54546-135457.revcompl.Pan.troglodytes.clone.rp43-41g5.GenBank:AC099533.36 ------------------------------------------------------------ 68417
human.FAM30B-LOC105372935-GGT2.NCBI.                                      gggaggacatcacataggacctcagcagagagcagctcgaatctatgcagagctgcacga 76847
human.FAM230E-LOC105377182-GGT3P.NCBI.                                    ------------------------------------------------------------ 57310
human.FAM230J-LOC105372942-GGTLC5P.NCBI.rev.compl                         ------------------------------------------------------------ 46364
human.FAM230A.ncRNA.18487127.18500594-GGTLC3.18516335.18518165.rev.compl.REF ------------------------------------------------------------ 31039

chimp.LOC112206744-LOC107973052-GGT2.revcompl                             ------------------------------------------------------------ 61762
chimp54546-135457.revcompl.Pan.troglodytes.clone.rp43-41g5.GenBank:AC099533.36 ------------------------------------------------------------ 68417
human.FAM30B-LOC105372935-GGT2.NCBI.                                      ggagccggaggtccagagtactgcagcagtttgacattgatgggtggtagggtccatgaa 76907
human.FAM230E-LOC105377182-GGT3P.NCBI.                                    ------------------------------------------------------------ 57310
human.FAM230J-LOC105372942-GGTLC5P.NCBI.rev.compl                         ------------------------------------------------------------ 46364
human.FAM230A.ncRNA.18487127.18500594-GGTLC3.18516335.18518165.rev.compl.REF ------------------------------------------------------------ 31039

chimp.LOC112206744-LOC107973052-GGT2.revcompl                             ------------------------------------------------------------ 61762
chimp54546-135457.revcompl.Pan.troglodytes.clone.rp43-41g5.GenBank:AC099533.36 ------------------------------------------------------------ 68417
human.FAM30B-LOC105372935-GGT2.NCBI.                                      agtagaggaagaccaagaggaagcagcagacaggggccaggccagggcagggatagggtg 76967
human.FAM230E-LOC105377182-GGT3P.NCBI.                                    ------------------------------------------------------------ 57310
human.FAM230J-LOC105372942-GGTLC5P.NCBI.rev.compl                         ------------------------------------------------------------ 46364
human.FAM230A.ncRNA.18487127.18500594-GGTLC3.18516335.18518165.rev.compl.REF ------------------------------------------------------------ 31039

chimp.LOC112206744-LOC107973052-GGT2.revcompl                             ------------------------------------------------------------ 61762
chimp54546-135457.revcompl.Pan.troglodytes.clone.rp43-41g5.GenBank:AC099533.36 ------------------------------------------------------------ 68417
human.FAM30B-LOC105372935-GGT2.NCBI.                                      ctctactaaagagcatgggctttgaactatgagcagtggagagccactgggggtgtaaat 77027
human.FAM230E-LOC105377182-GGT3P.NCBI.                                    ------------------------------------------------------------ 57310
human.FAM230J-LOC105372942-GGTLC5P.NCBI.rev.compl                         ------------------------------------------------------------ 46364
human.FAM230A.ncRNA.18487127.18500594-GGTLC3.18516335.18518165.rev.compl.REF ------------------------------------------------------------ 31039

chimp.LOC112206744-LOC107973052-GGT2.revcompl                             ------------------------------------------------------------ 61762
chimp54546-135457.revcompl.Pan.troglodytes.clone.rp43-41g5.GenBank:AC099533.36 ------------------------------------------------------------ 68417
human.FAM30B-LOC105372935-GGT2.NCBI.                                      aaggaaacagatattccaaatcctgccttgggaagattgttctggcaggagtgatgggaa 77087
human.FAM230E-LOC105377182-GGT3P.NCBI.                                    ------------------------------------------------------------ 57310
human.FAM230J-LOC105372942-GGTLC5P.NCBI.rev.compl                         ------------------------------------------------------------ 46364
human.FAM230A.ncRNA.18487127.18500594-GGTLC3.18516335.18518165.rev.compl.REF ------------------------------------------------------------ 31039

chimp.LOC112206744-LOC107973052-GGT2.revcompl                             ------------------------------------------------------------ 61762
chimp54546-135457.revcompl.Pan.troglodytes.clone.rp43-41g5.GenBank:AC099533.36 ------------------------------------------------------------ 68417
human.FAM30B-LOC105372935-GGT2.NCBI.                                      aggggccctggcagaagattccttcatgctcctgcggtgatccaggtgagagctgctgag 77147
human.FAM230E-LOC105377182-GGT3P.NCBI.                                    ------------------------------------------------------------ 57310
human.FAM230J-LOC105372942-GGTLC5P.NCBI.rev.compl                         ------------------------------------------------------------ 46364
human.FAM230A.ncRNA.18487127.18500594-GGTLC3.18516335.18518165.rev.compl.REF ------------------------------------------------------------ 31039

chimp.LOC112206744-LOC107973052-GGT2.revcompl                             ------------------------------------------------------------ 61762
chimp54546-135457.revcompl.Pan.troglodytes.clone.rp43-41g5.GenBank:AC099533.36 ------------------------------------------------------------ 68417
human.FAM30B-LOC105372935-GGT2.NCBI.                                      gaccccaaccaaatcagatcactggacctccactgggagtgaaagggggtggtctggcca 77207
human.FAM230E-LOC105377182-GGT3P.NCBI.                                    ------------------------------------------------------------ 57310
human.FAM230J-LOC105372942-GGTLC5P.NCBI.rev.compl                         ------------------------------------------------------------ 46364
human.FAM230A.ncRNA.18487127.18500594-GGTLC3.18516335.18518165.rev.compl.REF ------------------------------------------------------------ 31039

chimp.LOC112206744-LOC107973052-GGT2.revcompl                             ------------------------------------------------------------ 61762
chimp54546-135457.revcompl.Pan.troglodytes.clone.rp43-41g5.GenBank:AC099533.36 ------------------------------------------------------------ 68417
human.FAM30B-LOC105372935-GGT2.NCBI.                                      gagagatagtaaagagatgtagttaacaggcaagagaatgtggggatatccagggagctg 77267
human.FAM230E-LOC105377182-GGT3P.NCBI.                                    ------------------------------------------------------------ 57310
human.FAM230J-LOC105372942-GGTLC5P.NCBI.rev.compl                         ------------------------------------------------------------ 46364
human.FAM230A.ncRNA.18487127.18500594-GGTLC3.18516335.18518165.rev.compl.REF ------------------------------------------------------------ 31039

chimp.LOC112206744-LOC107973052-GGT2.revcompl                             ------------------------------------------------------------ 61762
chimp54546-135457.revcompl.Pan.troglodytes.clone.rp43-41g5.GenBank:AC099533.36 ------------------------------------------------------------ 68417
human.FAM30B-LOC105372935-GGT2.NCBI.                                      tccacaattcgcaggtcttaggcaactgcttgggagaagttgatgaggtcatcactggac 77327
human.FAM230E-LOC105377182-GGT3P.NCBI.                                    ------------------------------------------------------------ 57310
human.FAM230J-LOC105372942-GGTLC5P.NCBI.rev.compl                         ------------------------------------------------------------ 46364
human.FAM230A.ncRNA.18487127.18500594-GGTLC3.18516335.18518165.rev.compl.REF ------------------------------------------------------------ 31039

chimp.LOC112206744-LOC107973052-GGT2.revcompl                             ------------------------------------------------------------ 61762
chimp54546-135457.revcompl.Pan.troglodytes.clone.rp43-41g5.GenBank:AC099533.36 ------------------------------------------------------------ 68417
human.FAM30B-LOC105372935-GGT2.NCBI.                                      agagaatctattccctgggcccaaccaactctctagaccatcctgccatgtgcctctgag 77387
human.FAM230E-LOC105377182-GGT3P.NCBI.                                    ------------------------------------------------------------ 57310
human.FAM230J-LOC105372942-GGTLC5P.NCBI.rev.compl                         ------------------------------------------------------------ 46364
human.FAM230A.ncRNA.18487127.18500594-GGTLC3.18516335.18518165.rev.compl.REF ------------------------------------------------------------ 31039

chimp.LOC112206744-LOC107973052-GGT2.revcompl                             ------------------------------------------------------------ 61762
chimp54546-135457.revcompl.Pan.troglodytes.clone.rp43-41g5.GenBank:AC099533.36 ------------------------------------------------------------ 68417
human.FAM30B-LOC105372935-GGT2.NCBI.                                      aagtgagcttccccagccaagcccacaggaggacatatggagcaaaggtcaacagtcact 77447
```

```
human.FAM230E-LOC105377182-GGT3P.NCBI.                                          ------------------------------------------------------------ 57310
human.FAM230J-LOC105372942-GGTLC5P.NCBI.rev.compl                               ------------------------------------------------------------ 46364
human.FAM230A.ncRNA.18487127.18500594-GGTLC3.18516335.18518165.rev.compl.REF    ------------------------------------------------------------ 31039

chimp.LOC112206744-LOC107973052-GGT2.revcompl                                   ------------------------------------------------------------ 61762
chimp54546-135457.revcompl.Pan.troglodytes.clone.rp43-41g5.GenBank:AC099533.36  ------------------------------------------------------------ 68417
human.FAM30B-LOC105372935-GGT2.NCBI.                                            cagctgatggatggcaagcattcagatgggaggtgcctccacaggccatgataacccatg 77507
human.FAM230E-LOC105377182-GGT3P.NCBI.                                          ------------------------------------------------------------ 57310
human.FAM230J-LOC105372942-GGTLC5P.NCBI.rev.compl                               ------------------------------------------------------------ 46364
human.FAM230A.ncRNA.18487127.18500594-GGTLC3.18516335.18518165.rev.compl.REF    ------------------------------------------------------------ 31039

chimp.LOC112206744-LOC107973052-GGT2.revcompl                                   ------------------------------------------------------------ 61762
chimp54546-135457.revcompl.Pan.troglodytes.clone.rp43-41g5.GenBank:AC099533.36  ------------------------------------------------------------ 68417
human.FAM30B-LOC105372935-GGT2.NCBI.                                            gatccatttgagaaagatgccaaaaaacagttaattcaaggccactttaaaatacctggc 77567
human.FAM230E-LOC105377182-GGT3P.NCBI.                                          ------------------------------------------------------------ 57310
human.FAM230J-LOC105372942-GGTLC5P.NCBI.rev.compl                               ------------------------------------------------------------ 46364
human.FAM230A.ncRNA.18487127.18500594-GGTLC3.18516335.18518165.rev.compl.REF    ------------------------------------------------------------ 31039

chimp.LOC112206744-LOC107973052-GGT2.revcompl                                   ------------------------------------------------------------ 61762
chimp54546-135457.revcompl.Pan.troglodytes.clone.rp43-41g5.GenBank:AC099533.36  ------------------------------------------------------------ 68417
human.FAM30B-LOC105372935-GGT2.NCBI.                                            attcaatcctcctcaatgtctccatcaactgtagaccctataactaccaaatgagctcca 77627
human.FAM230E-LOC105377182-GGT3P.NCBI.                                          ------------------------------------------------------------ 57310
human.FAM230J-LOC105372942-GGTLC5P.NCBI.rev.compl                               ------------------------------------------------------------ 46364
human.FAM230A.ncRNA.18487127.18500594-GGTLC3.18516335.18518165.rev.compl.REF    ------------------------------------------------------------ 31039

chimp.LOC112206744-LOC107973052-GGT2.revcompl                                   ------------------------------------------------------------ 61762
chimp54546-135457.revcompl.Pan.troglodytes.clone.rp43-41g5.GenBank:AC099533.36  ------------------------------------------------------------ 68417
human.FAM30B-LOC105372935-GGT2.NCBI.                                            ctcgccacaaaggtcaagacacagaaacggtcagagagcagagcctattcctgaaaagca 77687
human.FAM230E-LOC105377182-GGT3P.NCBI.                                          ------------------------------------------------------------ 57310
human.FAM230J-LOC105372942-GGTLC5P.NCBI.rev.compl                               ------------------------------------------------------------ 46364
human.FAM230A.ncRNA.18487127.18500594-GGTLC3.18516335.18518165.rev.compl.REF    ------------------------------------------------------------ 31039

chimp.LOC112206744-LOC107973052-GGT2.revcompl                                   ------------------------------------------------------------ 61762
chimp54546-135457.revcompl.Pan.troglodytes.clone.rp43-41g5.GenBank:AC099533.36  ------------------------------------------------------------ 68417
human.FAM30B-LOC105372935-GGT2.NCBI.                                            aaagactccaaactgccacaactcaaccagtgaagggctggggaagactgacaggttcac 77747
human.FAM230E-LOC105377182-GGT3P.NCBI.                                          ------------------------------------------------------------ 57310
human.FAM230J-LOC105372942-GGTLC5P.NCBI.rev.compl                               ------------------------------------------------------------ 46364
human.FAM230A.ncRNA.18487127.18500594-GGTLC3.18516335.18518165.rev.compl.REF    ------------------------------------------------------------ 31039

chimp.LOC112206744-LOC107973052-GGT2.revcompl                                   ------------------------------------------------------------ 61762
chimp54546-135457.revcompl.Pan.troglodytes.clone.rp43-41g5.GenBank:AC099533.36  ------------------------------------------------------------ 68417
human.FAM30B-LOC105372935-GGT2.NCBI.                                            taagaaatgagaggaaggacaaacaaggtgttcaaattagaaaatgattccattaatgag 77807
human.FAM230E-LOC105377182-GGT3P.NCBI.                                          ------------------------------------------------------------ 57310
human.FAM230J-LOC105372942-GGTLC5P.NCBI.rev.compl                               ------------------------------------------------------------ 46364
human.FAM230A.ncRNA.18487127.18500594-GGTLC3.18516335.18518165.rev.compl.REF    ------------------------------------------------------------ 31039

chimp.LOC112206744-LOC107973052-GGT2.revcompl                                   ------------------------------------------------------------ 61762
chimp54546-135457.revcompl.Pan.troglodytes.clone.rp43-41g5.GenBank:AC099533.36  ------------------------------------------------------------ 68417
human.FAM30B-LOC105372935-GGT2.NCBI.                                            ggctctgatgatatagttgtggaggttatcagtttgccagagctgcagatttaaaatact 77867
human.FAM230E-LOC105377182-GGT3P.NCBI.                                          ------------------------------------------------------------ 57310
human.FAM230J-LOC105372942-GGTLC5P.NCBI.rev.compl                               ------------------------------------------------------------ 46364
human.FAM230A.ncRNA.18487127.18500594-GGTLC3.18516335.18518165.rev.compl.REF    ------------------------------------------------------------ 31039

chimp.LOC112206744-LOC107973052-GGT2.revcompl                                   ------------------------------------------------------------ 61762
chimp54546-135457.revcompl.Pan.troglodytes.clone.rp43-41g5.GenBank:AC099533.36  ------------------------------------------------------------ 68417
human.FAM30B-LOC105372935-GGT2.NCBI.                                            tacatggtcatcgaaaccaaaaaatgctgaccatcccccaaaacctcccaccccacctag 77927
human.FAM230E-LOC105377182-GGT3P.NCBI.                                          ------------------------------------------------------------ 57310
human.FAM230J-LOC105372942-GGTLC5P.NCBI.rev.compl                               ------------------------------------------------------------ 46364
human.FAM230A.ncRNA.18487127.18500594-GGTLC3.18516335.18518165.rev.compl.REF    ------------------------------------------------------------ 31039

chimp.LOC112206744-LOC107973052-GGT2.revcompl                                   ------------------------------------------------------------ 61762
chimp54546-135457.revcompl.Pan.troglodytes.clone.rp43-41g5.GenBank:AC099533.36  ------------------------------------------------------------ 68417
human.FAM30B-LOC105372935-GGT2.NCBI.                                            tgctgataacaggagttatcagttgccttttgtgtacagctagatgagatatggatctgt 77987
human.FAM230E-LOC105377182-GGT3P.NCBI.                                          ------------------------------------------------------------ 57310
human.FAM230J-LOC105372942-GGTLC5P.NCBI.rev.compl                               ------------------------------------------------------------ 46364
human.FAM230A.ncRNA.18487127.18500594-GGTLC3.18516335.18518165.rev.compl.REF    ------------------------------------------------------------ 31039

chimp.LOC112206744-LOC107973052-GGT2.revcompl                                   ------------------------------------------------------------ 61762
chimp54546-135457.revcompl.Pan.troglodytes.clone.rp43-41g5.GenBank:AC099533.36  ------------------------------------------------------------ 68417
human.FAM30B-LOC105372935-GGT2.NCBI.                                            gggggaagaaggggtgggcgtgctagcatctctcagctagactgtcatcggatggtgac 78047
human.FAM230E-LOC105377182-GGT3P.NCBI.                                          ------------------------------------------------------------ 57310
human.FAM230J-LOC105372942-GGTLC5P.NCBI.rev.compl                               ------------------------------------------------------------ 46364
human.FAM230A.ncRNA.18487127.18500594-GGTLC3.18516335.18518165.rev.compl.REF    ------------------------------------------------------------ 31039

chimp.LOC112206744-LOC107973052-GGT2.revcompl                                   ------------------------------------------------------------ 61762
chimp54546-135457.revcompl.Pan.troglodytes.clone.rp43-41g5.GenBank:AC099533.36  ------------------------------------------------------------ 68417
human.FAM30B-LOC105372935-GGT2.NCBI.                                            attccccaggactcaagcagcatctgagaagccactttcagtatgactatggaacagcta 78107
human.FAM230E-LOC105377182-GGT3P.NCBI.                                          ------------------------------------------------------------ 57310
human.FAM230J-LOC105372942-GGTLC5P.NCBI.rev.compl                               ------------------------------------------------------------ 46364
human.FAM230A.ncRNA.18487127.18500594-GGTLC3.18516335.18518165.rev.compl.REF    ------------------------------------------------------------ 31039

chimp.LOC112206744-LOC107973052-GGT2.revcompl                                   ------------------------------------------------------------ 61762
chimp54546-135457.revcompl.Pan.troglodytes.clone.rp43-41g5.GenBank:AC099533.36  ------------------------------------------------------------ 68417
human.FAM30B-LOC105372935-GGT2.NCBI.                                            accacacaacacagcatactgaatgaggagcctcgcaggctgagaagaaagcaaacccgg 78167
human.FAM230E-LOC105377182-GGT3P.NCBI.                                          ------------------------------------------------------------ 57310
human.FAM230J-LOC105372942-GGTLC5P.NCBI.rev.compl                               ------------------------------------------------------------ 46364
human.FAM230A.ncRNA.18487127.18500594-GGTLC3.18516335.18518165.rev.compl.REF    ------------------------------------------------------------ 31039

chimp.LOC112206744-LOC107973052-GGT2.revcompl                                   ------------------------------------------------------------ 61762
chimp54546-135457.revcompl.Pan.troglodytes.clone.rp43-41g5.GenBank:AC099533.36  ------------------------------------------------------------ 68417
human.FAM30B-LOC105372935-GGT2.NCBI.                                            aaggcaacaaatggatacgtgcatgctgaaggactcgtgtttaattcttcccggtaatcc 78227
human.FAM230E-LOC105377182-GGT3P.NCBI.                                          ------------------------------------------------------------ 57310
human.FAM230J-LOC105372942-GGTLC5P.NCBI.rev.compl                               ------------------------------------------------------------ 46364
human.FAM230A.ncRNA.18487127.18500594-GGTLC3.18516335.18518165.rev.compl.REF    ------------------------------------------------------------ 31039

chimp.LOC112206744-LOC107973052-GGT2.revcompl                                   ------------------------------------------------------------ 61762
chimp54546-135457.revcompl.Pan.troglodytes.clone.rp43-41g5.GenBank:AC099533.36  ------------------------------------------------------------ 68417
human.FAM30B-LOC105372935-GGT2.NCBI.                                            tttgtcaggcatatatttttgtctcctcaaacttaagaaagaaatccatgctctggaggg 78287
human.FAM230E-LOC105377182-GGT3P.NCBI.                                          ------------------------------------------------------------ 57310
human.FAM230J-LOC105372942-GGTLC5P.NCBI.rev.compl                               ------------------------------------------------------------ 46364
human.FAM230A.ncRNA.18487127.18500594-GGTLC3.18516335.18518165.rev.compl.REF    ------------------------------------------------------------ 31039

chimp.LOC112206744-LOC107973052-GGT2.revcompl                                   ------------------------------------------------------------ 61762
chimp54546-135457.revcompl.Pan.troglodytes.clone.rp43-41g5.GenBank:AC099533.36  ------------------------------------------------------------ 68417
human.FAM30B-LOC105372935-GGT2.NCBI.                                            gagaaagagaggctgtactcattaactcagagctgacctcgtctggcaggattaactcgc 78347
human.FAM230E-LOC105377182-GGT3P.NCBI.                                          ------------------------------------------------------------ 57310
human.FAM230J-LOC105372942-GGTLC5P.NCBI.rev.compl                               ------------------------------------------------------------ 46364
human.FAM230A.ncRNA.18487127.18500594-GGTLC3.18516335.18518165.rev.compl.REF    ------------------------------------------------------------ 31039

chimp.LOC112206744-LOC107973052-GGT2.revcompl                                   ------------------------------------------------------------ 61762
chimp54546-135457.revcompl.Pan.troglodytes.clone.rp43-41g5.GenBank:AC099533.36  ------------------------------------------------------------ 68417
human.FAM30B-LOC105372935-GGT2.NCBI.                                            catacctgtccagcccacagaacctggcctggagcatggagggagaaccacagccaactc 78407
human.FAM230E-LOC105377182-GGT3P.NCBI.                                          ------------------------------------------------------------ 57310
human.FAM230J-LOC105372942-GGTLC5P.NCBI.rev.compl                               ------------------------------------------------------------ 46364
human.FAM230A.ncRNA.18487127.18500594-GGTLC3.18516335.18518165.rev.compl.REF    ------------------------------------------------------------ 31039

chimp.LOC112206744-LOC107973052-GGT2.revcompl                                   ------------------------------------------------------------ 61762
chimp54546-135457.revcompl.Pan.troglodytes.clone.rp43-41g5.GenBank:AC099533.36  ------------------------------------------------------------ 68417
human.FAM30B-LOC105372935-GGT2.NCBI.                                            cagaaaaataaaaggctgcagcgagaaacaagtggggccacagagagaaaggggttttga 78467
human.FAM230E-LOC105377182-GGT3P.NCBI.                                          ------------------------------------------------------------ 57310
human.FAM230J-LOC105372942-GGTLC5P.NCBI.rev.compl                               ------------------------------------------------------------ 46364
human.FAM230A.ncRNA.18487127.18500594-GGTLC3.18516335.18518165.rev.compl.REF    ------------------------------------------------------------ 31039

chimp.LOC112206744-LOC107973052-GGT2.revcompl                                   ------------------------------------------------------------ 61762
chimp54546-135457.revcompl.Pan.troglodytes.clone.rp43-41g5.GenBank:AC099533.36  ------------------------------------------------------------ 68417
human.FAM30B-LOC105372935-GGT2.NCBI.                                            aaaagttctaaatccccatggctagggacagtgacatcagtgtgtttaccaagtatttga 78527
human.FAM230E-LOC105377182-GGT3P.NCBI.                                          ------------------------------------------------------------ 57310
```

```
human.FAM230J-LOC105372942-GGTLC5P.NCBI.rev.compl                             ------------------------------------------------------------ 46364
human.FAM230A.ncRNA.18487127.18500594-GGTLC3.18516335.18518165.rev.compl.REF  ------------------------------------------------------------ 31039

chimp.LOC112206744-LOC107973052-GGT2.revcompl                                 ------------------------------------------------------------ 61762
chimp54546-135457.revcompl.Pan.troglodytes.clone.rp43-41g5.GenBank:AC099533.36 ------------------------------------------------------------ 68417
human.FAM30B-LOC105372935-GGT2.NCBI.                                          tttacttgctcaggcaaaattccagaagagacactgggacctgtgtggagagtttcttca 78587
human.FAM230E-LOC105377182-GGT3P.NCBI.                                        ------------------------------------------------------------ 57310
human.FAM230J-LOC105372942-GGTLC5P.NCBI.rev.compl                             ------------------------------------------------------------ 46364
human.FAM230A.ncRNA.18487127.18500594-GGTLC3.18516335.18518165.rev.compl.REF  ------------------------------------------------------------ 31039

chimp.LOC112206744-LOC107973052-GGT2.revcompl                                 ------------------------------------------------------------ 61762
chimp54546-135457.revcompl.Pan.troglodytes.clone.rp43-41g5.GenBank:AC099533.36 ------------------------------------------------------------ 68417
human.FAM30B-LOC105372935-GGT2.NCBI.                                          ctgcagctatcctgccaagctcattgtaaagtgacgaattttgtggacacttacagagaa 78647
human.FAM230E-LOC105377182-GGT3P.NCBI.                                        ------------------------------------------------------------ 57310
human.FAM230J-LOC105372942-GGTLC5P.NCBI.rev.compl                             ------------------------------------------------------------ 46364
human.FAM230A.ncRNA.18487127.18500594-GGTLC3.18516335.18518165.rev.compl.REF  ------------------------------------------------------------ 31039

chimp.LOC112206744-LOC107973052-GGT2.revcompl                                 ------------------------------------------------------------ 61762
chimp54546-135457.revcompl.Pan.troglodytes.clone.rp43-41g5.GenBank:AC099533.36 ------------------------------------------------------------ 68417
human.FAM30B-LOC105372935-GGT2.NCBI.                                          aaaatagcttcaaatgctatagaccaatgctgcacgctgcagagacacagtctttgggtt 78707
human.FAM230E-LOC105377182-GGT3P.NCBI.                                        ------------------------------------------------------------ 57310
human.FAM230J-LOC105372942-GGTLC5P.NCBI.rev.compl                             ------------------------------------------------------------ 46364
human.FAM230A.ncRNA.18487127.18500594-GGTLC3.18516335.18518165.rev.compl.REF  ------------------------------------------------------------ 31039

chimp.LOC112206744-LOC107973052-GGT2.revcompl                                 ------------------------------------------------------------ 61762
chimp54546-135457.revcompl.Pan.troglodytes.clone.rp43-41g5.GenBank:AC099533.36 ------------------------------------------------------------ 68417
human.FAM30B-LOC105372935-GGT2.NCBI.                                          ctgtgagctgtatttgggcattacgcagagttcagctgccatctgtccctttgaaggcca 78767
human.FAM230E-LOC105377182-GGT3P.NCBI.                                        ------------------------------------------------------------ 57310
human.FAM230J-LOC105372942-GGTLC5P.NCBI.rev.compl                             ------------------------------------------------------------ 46364
human.FAM230A.ncRNA.18487127.18500594-GGTLC3.18516335.18518165.rev.compl.REF  ------------------------------------------------------------ 31039

chimp.LOC112206744-LOC107973052-GGT2.revcompl                                 ------------------------------------------------------------ 61762
chimp54546-135457.revcompl.Pan.troglodytes.clone.rp43-41g5.GenBank:AC099533.36 ------------------------------------------------------------ 68417
human.FAM30B-LOC105372935-GGT2.NCBI.                                          gctaagaagttggtcccagcctgggcatggtaccactgtcctacattccagccatttggg 78827
human.FAM230E-LOC105377182-GGT3P.NCBI.                                        ------------------------------------------------------------ 57310
human.FAM230J-LOC105372942-GGTLC5P.NCBI.rev.compl                             ------------------------------------------------------------ 46364
human.FAM230A.ncRNA.18487127.18500594-GGTLC3.18516335.18518165.rev.compl.REF  ------------------------------------------------------------ 31039

chimp.LOC112206744-LOC107973052-GGT2.revcompl                                 ------------------------------------------------------------ 61762
chimp54546-135457.revcompl.Pan.troglodytes.clone.rp43-41g5.GenBank:AC099533.36 ------------------------------------------------------------ 68417
human.FAM30B-LOC105372935-GGT2.NCBI.                                          atatcctgggttcaagcattccctcccattgctggatcaagcacttcccacctcctgctc 78887
human.FAM230E-LOC105377182-GGT3P.NCBI.                                        ------------------------------------------------------------ 57310
human.FAM230J-LOC105372942-GGTLC5P.NCBI.rev.compl                             ------------------------------------------------------------ 46364
human.FAM230A.ncRNA.18487127.18500594-GGTLC3.18516335.18518165.rev.compl.REF  ------------------------------------------------------------ 31039

chimp.LOC112206744-LOC107973052-GGT2.revcompl                                 ------------------------------------------------------------ 61762
chimp54546-135457.revcompl.Pan.troglodytes.clone.rp43-41g5.GenBank:AC099533.36 ------------------------------------------------------------ 68417
human.FAM30B-LOC105372935-GGT2.NCBI.                                          tgggaaatcagatctcagtaggtcatcaaccatcccagaaagtccagcagagttggtttt 78947
human.FAM230E-LOC105377182-GGT3P.NCBI.                                        ------------------------------------------------------------ 57310
human.FAM230J-LOC105372942-GGTLC5P.NCBI.rev.compl                             ------------------------------------------------------------ 46364
human.FAM230A.ncRNA.18487127.18500594-GGTLC3.18516335.18518165.rev.compl.REF  ------------------------------------------------------------ 31039

chimp.LOC112206744-LOC107973052-GGT2.revcompl                                 ------------------------------------------------------------ 61762
chimp54546-135457.revcompl.Pan.troglodytes.clone.rp43-41g5.GenBank:AC099533.36 ------------------------------------------------------------ 68417
human.FAM30B-LOC105372935-GGT2.NCBI.                                          ggtttggggtgtttttgagttgggtggaagaagagaccttacataatcttttaacttt 79007
human.FAM230E-LOC105377182-GGT3P.NCBI.                                        ------------------------------------------------------------ 57310
human.FAM230J-LOC105372942-GGTLC5P.NCBI.rev.compl                             ------------------------------------------------------------ 46364
human.FAM230A.ncRNA.18487127.18500594-GGTLC3.18516335.18518165.rev.compl.REF  ------------------------------------------------------------ 31039

chimp.LOC112206744-LOC107973052-GGT2.revcompl                                 ------------------------------------------------------------ 61762
chimp54546-135457.revcompl.Pan.troglodytes.clone.rp43-41g5.GenBank:AC099533.36 ------------------------------------------------------------ 68417
human.FAM30B-LOC105372935-GGT2.NCBI.                                          ttctctaaggctgaattccctataagcaaaattttaaattttttttaaagaagaaaaac 79067
human.FAM230E-LOC105377182-GGT3P.NCBI.                                        ------------------------------------------------------------ 57310
human.FAM230J-LOC105372942-GGTLC5P.NCBI.rev.compl                             ------------------------------------------------------------ 46364
human.FAM230A.ncRNA.18487127.18500594-GGTLC3.18516335.18518165.rev.compl.REF  ------------------------------------------------------------ 31039

chimp.LOC112206744-LOC107973052-GGT2.revcompl                                 ------------------------------------------------------------ 61762
chimp54546-135457.revcompl.Pan.troglodytes.clone.rp43-41g5.GenBank:AC099533.36 ------------------------------------------------------------ 68417
human.FAM30B-LOC105372935-GGT2.NCBI.                                          acccagacaatccttaaaatcccctttcagctccaagactgtgtaattctgtgataccag 79127
human.FAM230E-LOC105377182-GGT3P.NCBI.                                        ------------------------------------------------------------ 57310
human.FAM230J-LOC105372942-GGTLC5P.NCBI.rev.compl                             ------------------------------------------------------------ 46364
human.FAM230A.ncRNA.18487127.18500594-GGTLC3.18516335.18518165.rev.compl.REF  ------------------------------------------------------------ 31039

chimp.LOC112206744-LOC107973052-GGT2.revcompl                                 ------------------------------------------------------------ 61762
chimp54546-135457.revcompl.Pan.troglodytes.clone.rp43-41g5.GenBank:AC099533.36 ------------------------------------------------------------ 68417
human.FAM30B-LOC105372935-GGT2.NCBI.                                          tattgaaagcccctgcatttccctaactatatattcccctatgccagggaaaggtgcatg 79187
human.FAM230E-LOC105377182-GGT3P.NCBI.                                        ------------------------------------------------------------ 57310
human.FAM230J-LOC105372942-GGTLC5P.NCBI.rev.compl                             ------------------------------------------------------------ 46364
human.FAM230A.ncRNA.18487127.18500594-GGTLC3.18516335.18518165.rev.compl.REF  ------------------------------------------------------------ 31039

chimp.LOC112206744-LOC107973052-GGT2.revcompl                                 ------------------------------------------------------------ 61762
chimp54546-135457.revcompl.Pan.troglodytes.clone.rp43-41g5.GenBank:AC099533.36 ------------------------------------------------------------ 68417
human.FAM30B-LOC105372935-GGT2.NCBI.                                          taagtgagagaacaaaaaatttcacaatcgtggggacctcttctttcggctttggagccc 79247
human.FAM230E-LOC105377182-GGT3P.NCBI.                                        ------------------------------------------------------------ 57310
human.FAM230J-LOC105372942-GGTLC5P.NCBI.rev.compl                             ------------------------------------------------------------ 46364
human.FAM230A.ncRNA.18487127.18500594-GGTLC3.18516335.18518165.rev.compl.REF  ------------------------------------------------------------ 31039

chimp.LOC112206744-LOC107973052-GGT2.revcompl                                 ------------------------------------------------------------ 61762
chimp54546-135457.revcompl.Pan.troglodytes.clone.rp43-41g5.GenBank:AC099533.36 ------------------------------------------------------------ 68417
human.FAM30B-LOC105372935-GGT2.NCBI.                                          ccttccctctgtctctgtacggggcagctttttccttctgtcttctcccttccttcttgt 79307
human.FAM230E-LOC105377182-GGT3P.NCBI.                                        ------------------------------------------------------------ 57310
human.FAM230J-LOC105372942-GGTLC5P.NCBI.rev.compl                             ------------------------------------------------------------ 46364
human.FAM230A.ncRNA.18487127.18500594-GGTLC3.18516335.18518165.rev.compl.REF  ------------------------------------------------------------ 31039

chimp.LOC112206744-LOC107973052-GGT2.revcompl                                 ------------------------------------------------------------ 61762
chimp54546-135457.revcompl.Pan.troglodytes.clone.rp43-41g5.GenBank:AC099533.36 ------------------------------------------------------------ 68417
human.FAM30B-LOC105372935-GGT2.NCBI.                                          ctactaaactctctgcgccttagaaccaaaaataaaaattcaaaatcatgatcttagggg 79367
human.FAM230E-LOC105377182-GGT3P.NCBI.                                        ------------------------------------------------------------ 57310
human.FAM230J-LOC105372942-GGTLC5P.NCBI.rev.compl                             ------------------------------------------------------------ 46364
human.FAM230A.ncRNA.18487127.18500594-GGTLC3.18516335.18518165.rev.compl.REF  ------------------------------------------------------------ 31039

chimp.LOC112206744-LOC107973052-GGT2.revcompl                                 ------------------------------------------------------------ 61762
chimp54546-135457.revcompl.Pan.troglodytes.clone.rp43-41g5.GenBank:AC099533.36 ------------------------------------------------------------ 68417
human.FAM30B-LOC105372935-GGT2.NCBI.                                          tgaaagaacactgagaaaccatagaaagcaaaaacgaaaagccaacatagaaaaagtctg 79427
human.FAM230E-LOC105377182-GGT3P.NCBI.                                        ------------------------------------------------------------ 57310
human.FAM230J-LOC105372942-GGTLC5P.NCBI.rev.compl                             ------------------------------------------------------------ 46364
human.FAM230A.ncRNA.18487127.18500594-GGTLC3.18516335.18518165.rev.compl.REF  ------------------------------------------------------------ 31039

chimp.LOC112206744-LOC107973052-GGT2.revcompl                                 ------------------------------------------------------------ 61762
chimp54546-135457.revcompl.Pan.troglodytes.clone.rp43-41g5.GenBank:AC099533.36 ------------------------------------------------------------ 68417
human.FAM30B-LOC105372935-GGT2.NCBI.                                          acttgaattagaagcagcacttaaaattacaggttcagaaaaagcttgaacagaatccag 79487
human.FAM230E-LOC105377182-GGT3P.NCBI.                                        ------------------------------------------------------------ 57310
human.FAM230J-LOC105372942-GGTLC5P.NCBI.rev.compl                             ------------------------------------------------------------ 46364
human.FAM230A.ncRNA.18487127.18500594-GGTLC3.18516335.18518165.rev.compl.REF  ------------------------------------------------------------ 31039

chimp.LOC112206744-LOC107973052-GGT2.revcompl                                 ------------------------------------------------------------ 61762
chimp54546-135457.revcompl.Pan.troglodytes.clone.rp43-41g5.GenBank:AC099533.36 ------------------------------------------------------------ 68417
human.FAM30B-LOC105372935-GGT2.NCBI.                                          ggctgaggtcagccctgagtctgcaggctctgccctctcacaaaggtgagtagatgggtc 79547
human.FAM230E-LOC105377182-GGT3P.NCBI.                                        ------------------------------------------------------------ 57310
human.FAM230J-LOC105372942-GGTLC5P.NCBI.rev.compl                             ------------------------------------------------------------ 46364
human.FAM230A.ncRNA.18487127.18500594-GGTLC3.18516335.18518165.rev.compl.REF  ------------------------------------------------------------ 31039

chimp.LOC112206744-LOC107973052-GGT2.revcompl                                 ------------------------------------------------------------ 61762
chimp54546-135457.revcompl.Pan.troglodytes.clone.rp43-41g5.GenBank:AC099533.36 ------------------------------------------------------------ 68417
human.FAM30B-LOC105372935-GGT2.NCBI.                                          tggagagggctgttcatgctatggaagggcctctatgccatcgaggctaaaaccaaagca 79607
human.FAM230E-LOC105377182-GGT3P.NCBI.                                        ------------------------------------------------------------ 57310
human.FAM230J-LOC105372942-GGTLC5P.NCBI.rev.compl                             ------------------------------------------------------------ 46364
```

```
human.FAM230A.ncRNA.18487127.18500594-GGTLC3.18516335.18518165.rev.compl.REF        ------------------------------------------------------------ 31039

chimp.LOC112206744-LOC107973052-GGT2.revcompl                                       ------------------------------------------------------------ 61762
chimp54546-135457.revcompl.Pan.troglodytes.clone.rp43-41g5.GenBank:AC099533.36      ------------------------------------------------------------ 68417
human.FAM30B-LOC105372935-GGT2.NCBI.                                                ggagcaaatcagccctgctagtgtgtacagtcggcatcaaagaggcaacaacggtagaga 79667
human.FAM230E-LOC105377182-GGT3P.NCBI.                                              ------------------------------------------------------------ 57310
human.FAM230J-LOC105372942-GGTLC5P.NCBI.rev.compl                                   ------------------------------------------------------------ 46364
human.FAM230A.ncRNA.18487127.18500594-GGTLC3.18516335.18518165.rev.compl.REF        ------------------------------------------------------------ 31039

chimp.LOC112206744-LOC107973052-GGT2.revcompl                                       ------------------------------------------------------------ 61762
chimp54546-135457.revcompl.Pan.troglodytes.clone.rp43-41g5.GenBank:AC099533.36      ------------------------------------------------------------ 68417
human.FAM30B-LOC105372935-GGT2.NCBI.                                                atattttctgaaagacacacatgaaaccagtaacggcgatggtctctggaagaattaggt 79727
human.FAM230E-LOC105377182-GGT3P.NCBI.                                              ------------------------------------------------------------ 57310
human.FAM230J-LOC105372942-GGTLC5P.NCBI.rev.compl                                   ------------------------------------------------------------ 46364
human.FAM230A.ncRNA.18487127.18500594-GGTLC3.18516335.18518165.rev.compl.REF        ------------------------------------------------------------ 31039

chimp.LOC112206744-LOC107973052-GGT2.revcompl                                       ------------------------------------------------------------ 61762
chimp54546-135457.revcompl.Pan.troglodytes.clone.rp43-41g5.GenBank:AC099533.36      ------------------------------------------------------------ 68417
human.FAM30B-LOC105372935-GGT2.NCBI.                                                cagaagcagacttaatttgcactgtgtacattctttggtatcctttttaattgaacgcat 79787
human.FAM230E-LOC105377182-GGT3P.NCBI.                                              ------------------------------------------------------------ 57310
human.FAM230J-LOC105372942-GGTLC5P.NCBI.rev.compl                                   ------------------------------------------------------------ 46364
human.FAM230A.ncRNA.18487127.18500594-GGTLC3.18516335.18518165.rev.compl.REF        ------------------------------------------------------------ 31039

chimp.LOC112206744-LOC107973052-GGT2.revcompl                                       ------------------------------------------------------------ 61762
chimp54546-135457.revcompl.Pan.troglodytes.clone.rp43-41g5.GenBank:AC099533.36      ------------------------------------------------------------ 68417
human.FAM30B-LOC105372935-GGT2.NCBI.                                                gtgaatgcattacctattcaaaaataaatgtgtaaaaacacatgccacgacaaagcaaaa 79847
human.FAM230E-LOC105377182-GGT3P.NCBI.                                              ------------------------------------------------------------ 57310
human.FAM230J-LOC105372942-GGTLC5P.NCBI.rev.compl                                   ------------------------------------------------------------ 46364
human.FAM230A.ncRNA.18487127.18500594-GGTLC3.18516335.18518165.rev.compl.REF        ------------------------------------------------------------ 31039

chimp.LOC112206744-LOC107973052-GGT2.revcompl                                       ------------------------------------------------------------ 61762
chimp54546-135457.revcompl.Pan.troglodytes.clone.rp43-41g5.GenBank:AC099533.36      ------------------------------------------------------------ 68417
human.FAM30B-LOC105372935-GGT2.NCBI.                                                tgaggagacagtataaaaaagacaaggataattttaaatttctcctaataaaacgctgtt 79907
human.FAM230E-LOC105377182-GGT3P.NCBI.                                              ------------------------------------------------------------ 57310
human.FAM230J-LOC105372942-GGTLC5P.NCBI.rev.compl                                   ------------------------------------------------------------ 46364
human.FAM230A.ncRNA.18487127.18500594-GGTLC3.18516335.18518165.rev.compl.REF        ------------------------------------------------------------ 31039

chimp.LOC112206744-LOC107973052-GGT2.revcompl                                       ------------------------------------------------------------ 61762
chimp54546-135457.revcompl.Pan.troglodytes.clone.rp43-41g5.GenBank:AC099533.36      ------------------------------------------------------------ 68417
human.FAM30B-LOC105372935-GGT2.NCBI.                                                taaaaatctaagaggaaaaaagggtaggctccagctagtggagagaacaactgaaaaaca 79967
human.FAM230E-LOC105377182-GGT3P.NCBI.                                              ------------------------------------------------------------ 57310
human.FAM230J-LOC105372942-GGTLC5P.NCBI.rev.compl                                   ------------------------------------------------------------ 46364
human.FAM230A.ncRNA.18487127.18500594-GGTLC3.18516335.18518165.rev.compl.REF        ------------------------------------------------------------ 31039

chimp.LOC112206744-LOC107973052-GGT2.revcompl                                       ------------------------------------------------------------ 61762
chimp54546-135457.revcompl.Pan.troglodytes.clone.rp43-41g5.GenBank:AC099533.36      ------------------------------------------------------------ 68417
human.FAM30B-LOC105372935-GGT2.NCBI.                                                agagccacgtgttcccatggtcctggccaaccccacttctttcccatcctcactcgttca 80027
human.FAM230E-LOC105377182-GGT3P.NCBI.                                              ------------------------------------------------------------ 57310
human.FAM230J-LOC105372942-GGTLC5P.NCBI.rev.compl                                   ------------------------------------------------------------ 46364
human.FAM230A.ncRNA.18487127.18500594-GGTLC3.18516335.18518165.rev.compl.REF        ------------------------------------------------------------ 31039

chimp.LOC112206744-LOC107973052-GGT2.revcompl                                       ------------------------------------------------------------ 61762
chimp54546-135457.revcompl.Pan.troglodytes.clone.rp43-41g5.GenBank:AC099533.36      ------------------------------------------------------------ 68417
human.FAM30B-LOC105372935-GGT2.NCBI.                                                catctgccttggagcacccacagcaagcctggcttcatgcccagaacatcacctggcttt 80087
human.FAM230E-LOC105377182-GGT3P.NCBI.                                              ------------------------------------------------------------ 57310
human.FAM230J-LOC105372942-GGTLC5P.NCBI.rev.compl                                   ------------------------------------------------------------ 46364
human.FAM230A.ncRNA.18487127.18500594-GGTLC3.18516335.18518165.rev.compl.REF        ------------------------------------------------------------ 31039

chimp.LOC112206744-LOC107973052-GGT2.revcompl                                       ------------------------------------------------------------ 61762
chimp54546-135457.revcompl.Pan.troglodytes.clone.rp43-41g5.GenBank:AC099533.36      ------------------------------------------------------------ 68417
human.FAM30B-LOC105372935-GGT2.NCBI.                                                cctcctctctccctgcccacatctcctcagtcactcgctgcctcctctcttgcttctctc 80147
human.FAM230E-LOC105377182-GGT3P.NCBI.                                              ------------------------------------------------------------ 57310
human.FAM230J-LOC105372942-GGTLC5P.NCBI.rev.compl                                   ------------------------------------------------------------ 46364
human.FAM230A.ncRNA.18487127.18500594-GGTLC3.18516335.18518165.rev.compl.REF        ------------------------------------------------------------ 31039

chimp.LOC112206744-LOC107973052-GGT2.revcompl                                       ------------------------------------------------------------ 61762
chimp54546-135457.revcompl.Pan.troglodytes.clone.rp43-41g5.GenBank:AC099533.36      ------------------------------------------------------------ 68417
human.FAM30B-LOC105372935-GGT2.NCBI.                                                ccagttctcagtttgcccatcaaatctcaggtggtactctccattctctgctatgaggga 80207
human.FAM230E-LOC105377182-GGT3P.NCBI.                                              ------------------------------------------------------------ 57310
human.FAM230J-LOC105372942-GGTLC5P.NCBI.rev.compl                                   ------------------------------------------------------------ 46364
human.FAM230A.ncRNA.18487127.18500594-GGTLC3.18516335.18518165.rev.compl.REF        ------------------------------------------------------------ 31039

chimp.LOC112206744-LOC107973052-GGT2.revcompl                                       ------------------------------------------------------------ 61762
chimp54546-135457.revcompl.Pan.troglodytes.clone.rp43-41g5.GenBank:AC099533.36      ------------------------------------------------------------ 68417
human.FAM30B-LOC105372935-GGT2.NCBI.                                                gttcatttgttcatgcaatttccccaccggagacaaaatgtatttcccaaagttggcttg 80267
human.FAM230E-LOC105377182-GGT3P.NCBI.                                              ------------------------------------------------------------ 57310
human.FAM230J-LOC105372942-GGTLC5P.NCBI.rev.compl                                   ------------------------------------------------------------ 46364
human.FAM230A.ncRNA.18487127.18500594-GGTLC3.18516335.18518165.rev.compl.REF        ------------------------------------------------------------ 31039

chimp.LOC112206744-LOC107973052-GGT2.revcompl                                       ------------------------------------------------------------ 61762
chimp54546-135457.revcompl.Pan.troglodytes.clone.rp43-41g5.GenBank:AC099533.36      ------------------------------------------------------------ 68417
human.FAM30B-LOC105372935-GGT2.NCBI.                                                caatgacatctcccatgcctgctctgcttacccatgactttgacactccttccatcgaga 80327
human.FAM230E-LOC105377182-GGT3P.NCBI.                                              ------------------------------------------------------------ 57310
human.FAM230J-LOC105372942-GGTLC5P.NCBI.rev.compl                                   ------------------------------------------------------------ 46364
human.FAM230A.ncRNA.18487127.18500594-GGTLC3.18516335.18518165.rev.compl.REF        ------------------------------------------------------------ 31039

chimp.LOC112206744-LOC107973052-GGT2.revcompl                                       ------------------------------------------------------------ 61762
chimp54546-135457.revcompl.Pan.troglodytes.clone.rp43-41g5.GenBank:AC099533.36      ------------------------------------------------------------ 68417
human.FAM30B-LOC105372935-GGT2.NCBI.                                                gggcagaacctgtgtttcctctctctgaatctaggtgggctacagctctggtagaagtga 80387
human.FAM230E-LOC105377182-GGT3P.NCBI.                                              ------------------------------------------------------------ 57310
human.FAM230J-LOC105372942-GGTLC5P.NCBI.rev.compl                                   ------------------------------------------------------------ 46364
human.FAM230A.ncRNA.18487127.18500594-GGTLC3.18516335.18518165.rev.compl.REF        ------------------------------------------------------------ 31039

chimp.LOC112206744-LOC107973052-GGT2.revcompl                                       ------------------------------------------------------------ 61762
chimp54546-135457.revcompl.Pan.troglodytes.clone.rp43-41g5.GenBank:AC099533.36      ------------------------------------------------------------ 68417
human.FAM30B-LOC105372935-GGT2.NCBI.                                                cactctgtgaccttaaggctaggtcacaaaaagggaatgcagcttctgccagcaatcctg 80447
human.FAM230E-LOC105377182-GGT3P.NCBI.                                              ------------------------------------------------------------ 57310
human.FAM230J-LOC105372942-GGTLC5P.NCBI.rev.compl                                   ------------------------------------------------------------ 46364
human.FAM230A.ncRNA.18487127.18500594-GGTLC3.18516335.18518165.rev.compl.REF        ------------------------------------------------------------ 31039

chimp.LOC112206744-LOC107973052-GGT2.revcompl                                       ------------------------------------------------------------ 61762
chimp54546-135457.revcompl.Pan.troglodytes.clone.rp43-41g5.GenBank:AC099533.36      ------------------------------------------------------------ 68417
human.FAM30B-LOC105372935-GGT2.NCBI.                                                ggacacacacttggagcccgagctgctgtgtaagccatctgcctgctctgagcccaccac 80507
human.FAM230E-LOC105377182-GGT3P.NCBI.                                              ------------------------------------------------------------ 57310
human.FAM230J-LOC105372942-GGTLC5P.NCBI.rev.compl                                   ------------------------------------------------------------ 46364
human.FAM230A.ncRNA.18487127.18500594-GGTLC3.18516335.18518165.rev.compl.REF        ------------------------------------------------------------ 31039

chimp.LOC112206744-LOC107973052-GGT2.revcompl                                       ------------------------------------------------------------ 61762
chimp54546-135457.revcompl.Pan.troglodytes.clone.rp43-41g5.GenBank:AC099533.36      ------------------------------------------------------------ 68417
human.FAM30B-LOC105372935-GGT2.NCBI.                                                actataaggaagcccaagctagcccacaggagagacgaggtgaagagaatgagatgccag 80567
human.FAM230E-LOC105377182-GGT3P.NCBI.                                              ------------------------------------------------------------ 57310
human.FAM230J-LOC105372942-GGTLC5P.NCBI.rev.compl                                   ------------------------------------------------------------ 46364
human.FAM230A.ncRNA.18487127.18500594-GGTLC3.18516335.18518165.rev.compl.REF        ------------------------------------------------------------ 31039

chimp.LOC112206744-LOC107973052-GGT2.revcompl                                       ------------------------------------------------------------ 61762
chimp54546-135457.revcompl.Pan.troglodytes.clone.rp43-41g5.GenBank:AC099533.36      ------------------------------------------------------------ 68417
human.FAM30B-LOC105372935-GGT2.NCBI.                                                tctccactgctccagctctctagtgtcccagccccagccactatctgactgcaaccccat 80627
human.FAM230E-LOC105377182-GGT3P.NCBI.                                              ------------------------------------------------------------ 57310
human.FAM230J-LOC105372942-GGTLC5P.NCBI.rev.compl                                   ------------------------------------------------------------ 46364
human.FAM230A.ncRNA.18487127.18500594-GGTLC3.18516335.18518165.rev.compl.REF        ------------------------------------------------------------ 31039

chimp.LOC112206744-LOC107973052-GGT2.revcompl                                       ------------------------------------------------------------ 61762
chimp54546-135457.revcompl.Pan.troglodytes.clone.rp43-41g5.GenBank:AC099533.36      ------------------------------------------------------------ 68417
human.FAM30B-LOC105372935-GGT2.NCBI.                                                gaaagactgagccagaaccacgagcccagcccttcctgatgttgacatacagacttcatg 80687
human.FAM230E-LOC105377182-GGT3P.NCBI.                                              ------------------------------------------------------------ 57310
human.FAM230J-LOC105372942-GGTLC5P.NCBI.rev.compl                                   ------------------------------------------------------------ 46364
human.FAM230A.ncRNA.18487127.18500594-GGTLC3.18516335.18518165.rev.compl.REF        ------------------------------------------------------------ 31039
```

```
chimp.LOC112206744-LOC107973052-GGT2.revcompl                                      ------------------------------------------------------------ 61762
chimp54546-135457.revcompl.Pan.troglodytes.clone.rp43-41g5.GenBank:AC099533.36     ------------------------------------------------------------ 68417
human.FAM30B-LOC105372935-GGT2.NCBI.                                               aaagagaataaactaatttttgctttaagccactatgatttgggatgatttgttatgcag 80747
human.FAM230E-LOC105377182-GGT3P.NCBI.                                             ------------------------------------------------------------ 57310
human.FAM230J-LOC105372942-GGTLC5P.NCBI.rev.compl                                  ------------------------------------------------------------ 46364
human.FAM230A.ncRNA.18487127.18500594-GGTLC3.18516335.18518165.rev.compl.REF       ------------------------------------------------------------ 31039

chimp.LOC112206744-LOC107973052-GGT2.revcompl                                      ------------------------------------------------------------ 61762
chimp54546-135457.revcompl.Pan.troglodytes.clone.rp43-41g5.GenBank:AC099533.36     ------------------------------------------------------------ 68417
human.FAM30B-LOC105372935-GGT2.NCBI.                                               cactggataactgcaacacctgcttctaaacagtagaatcccaaacgtctaggccccatc 80807
human.FAM230E-LOC105377182-GGT3P.NCBI.                                             ------------------------------------------------------------ 57310
human.FAM230J-LOC105372942-GGTLC5P.NCBI.rev.compl                                  ------------------------------------------------------------ 46364
human.FAM230A.ncRNA.18487127.18500594-GGTLC3.18516335.18518165.rev.compl.REF       ------------------------------------------------------------ 31039

chimp.LOC112206744-LOC107973052-GGT2.revcompl                                      ------------------------------------------------------------ 61762
chimp54546-135457.revcompl.Pan.troglodytes.clone.rp43-41g5.GenBank:AC099533.36     ------------------------------------------------------------ 68417
human.FAM30B-LOC105372935-GGT2.NCBI.                                               tctaacttacttcatgttcctgttctcactgatttttgccactgagcttgaccaagccc 80867
human.FAM230E-LOC105377182-GGT3P.NCBI.                                             ------------------------------------------------------------ 57310
human.FAM230J-LOC105372942-GGTLC5P.NCBI.rev.compl                                  ------------------------------------------------------------ 46364
human.FAM230A.ncRNA.18487127.18500594-GGTLC3.18516335.18518165.rev.compl.REF       ------------------------------------------------------------ 31039

chimp.LOC112206744-LOC107973052-GGT2.revcompl                                      ------------------------------------------------------------ 61762
chimp54546-135457.revcompl.Pan.troglodytes.clone.rp43-41g5.GenBank:AC099533.36     ------------------------------------------------------------ 68417
human.FAM30B-LOC105372935-GGT2.NCBI.                                               ttgagtcaccaattccagatatttgtttaaacaaattcactgtgactgtctacactatcc 80927
human.FAM230E-LOC105377182-GGT3P.NCBI.                                             ------------------------------------------------------------ 57310
human.FAM230J-LOC105372942-GGTLC5P.NCBI.rev.compl                                  ------------------------------------------------------------ 46364
human.FAM230A.ncRNA.18487127.18500594-GGTLC3.18516335.18518165.rev.compl.REF       ------------------------------------------------------------ 31039

chimp.LOC112206744-LOC107973052-GGT2.revcompl                                      ------------------------------------------------------------ 61762
chimp54546-135457.revcompl.Pan.troglodytes.clone.rp43-41g5.GenBank:AC099533.36     ------------------------------------------------------------ 68417
human.FAM30B-LOC105372935-GGT2.NCBI.                                               attcctccaaaacaatccctgttatagttggccttccccagttctcttcctggcatagag 80987
human.FAM230E-LOC105377182-GGT3P.NCBI.                                             ------------------------------------------------------------ 57310
human.FAM230J-LOC105372942-GGTLC5P.NCBI.rev.compl                                  ------------------------------------------------------------ 46364
human.FAM230A.ncRNA.18487127.18500594-GGTLC3.18516335.18518165.rev.compl.REF       ------------------------------------------------------------ 31039

chimp.LOC112206744-LOC107973052-GGT2.revcompl                                      ------------------------------------------------------------ 61762
chimp54546-135457.revcompl.Pan.troglodytes.clone.rp43-41g5.GenBank:AC099533.36     ------------------------------------------------------------ 68417
human.FAM30B-LOC105372935-GGT2.NCBI.                                               tagtctgtgcccttctctcatttcttaatatgaccctcacccatggagtgttcctagatt 81047
human.FAM230E-LOC105377182-GGT3P.NCBI.                                             ------------------------------------------------------------ 57310
human.FAM230J-LOC105372942-GGTLC5P.NCBI.rev.compl                                  ------------------------------------------------------------ 46364
human.FAM230A.ncRNA.18487127.18500594-GGTLC3.18516335.18518165.rev.compl.REF       ------------------------------------------------------------ 31039

chimp.LOC112206744-LOC107973052-GGT2.revcompl                                      ------------------------------------------------------------ 61762
chimp54546-135457.revcompl.Pan.troglodytes.clone.rp43-41g5.GenBank:AC099533.36     ------------------------------------------------------------ 68417
human.FAM30B-LOC105372935-GGT2.NCBI.                                               ctgctcattcataatgcttcttcacccaccctaacctctccctcctgtagccacagatca 81107
human.FAM230E-LOC105377182-GGT3P.NCBI.                                             ------------------------------------------------------------ 57310
human.FAM230J-LOC105372942-GGTLC5P.NCBI.rev.compl                                  ------------------------------------------------------------ 46364
human.FAM230A.ncRNA.18487127.18500594-GGTLC3.18516335.18518165.rev.compl.REF       ------------------------------------------------------------ 31039

chimp.LOC112206744-LOC107973052-GGT2.revcompl                                      ------------------------------------------------------------ 61762
chimp54546-135457.revcompl.Pan.troglodytes.clone.rp43-41g5.GenBank:AC099533.36     ------------------------------------------------------------ 68417
human.FAM30B-LOC105372935-GGT2.NCBI.                                               tccacttttccaggcctctgcactggaggctgcatcagccacctcccagtacgtcatcct 81167
human.FAM230E-LOC105377182-GGT3P.NCBI.                                             ------------------------------------------------------------ 57310
human.FAM230J-LOC105372942-GGTLC5P.NCBI.rev.compl                                  ------------------------------------------------------------ 46364
human.FAM230A.ncRNA.18487127.18500594-GGTLC3.18516335.18518165.rev.compl.REF       ------------------------------------------------------------ 31039

chimp.LOC112206744-LOC107973052-GGT2.revcompl                                      ------------------------------------------------------------ 61762
chimp54546-135457.revcompl.Pan.troglodytes.clone.rp43-41g5.GenBank:AC099533.36     ------------------------------------------------------------ 68417
human.FAM30B-LOC105372935-GGT2.NCBI.                                               ttccaatgccctacaacacggggcagaaaggactctttttgctgcacatccaggctgtt 81227
human.FAM230E-LOC105377182-GGT3P.NCBI.                                             ------------------------------------------------------------ 57310
human.FAM230J-LOC105372942-GGTLC5P.NCBI.rev.compl                                  ------------------------------------------------------------ 46364
human.FAM230A.ncRNA.18487127.18500594-GGTLC3.18516335.18518165.rev.compl.REF       ------------------------------------------------------------ 31039

chimp.LOC112206744-LOC107973052-GGT2.revcompl                                      ------------------------------------------------------------ 61762
chimp54546-135457.revcompl.Pan.troglodytes.clone.rp43-41g5.GenBank:AC099533.36     ------------------------------------------------------------ 68417
human.FAM30B-LOC105372935-GGT2.NCBI.                                               taagaaaaggcttaaggcagcttaagcacaaagaatgtcatcttcaaacattgtaacggc 81287
human.FAM230E-LOC105377182-GGT3P.NCBI.                                             ------------------------------------------------------------ 57310
human.FAM230J-LOC105372942-GGTLC5P.NCBI.rev.compl                                  ------------------------------------------------------------ 46364
human.FAM230A.ncRNA.18487127.18500594-GGTLC3.18516335.18518165.rev.compl.REF       ------------------------------------------------------------ 31039

chimp.LOC112206744-LOC107973052-GGT2.revcompl                                      ------------------------------------------------------------ 61762
chimp54546-135457.revcompl.Pan.troglodytes.clone.rp43-41g5.GenBank:AC099533.36     ------------------------------------------------------------ 68417
human.FAM30B-LOC105372935-GGT2.NCBI.                                               atcttctttttggggcactgaagtattcccatgattccaagaatcaaatgaacagcatct 81347
human.FAM230E-LOC105377182-GGT3P.NCBI.                                             ------------------------------------------------------------ 57310
human.FAM230J-LOC105372942-GGTLC5P.NCBI.rev.compl                                  ------------------------------------------------------------ 46364
human.FAM230A.ncRNA.18487127.18500594-GGTLC3.18516335.18518165.rev.compl.REF       ------------------------------------------------------------ 31039

chimp.LOC112206744-LOC107973052-GGT2.revcompl                                      ------------------------------------------------------------ 61762
chimp54546-135457.revcompl.Pan.troglodytes.clone.rp43-41g5.GenBank:AC099533.36     ------------------------------------------------------------ 68417
human.FAM30B-LOC105372935-GGT2.NCBI.                                               ggttatggggtctctggtctctgccaaggcaaactcataaagcatttcagagcccttcat 81407
human.FAM230E-LOC105377182-GGT3P.NCBI.                                             ------------------------------------------------------------ 57310
human.FAM230J-LOC105372942-GGTLC5P.NCBI.rev.compl                                  ------------------------------------------------------------ 46364
human.FAM230A.ncRNA.18487127.18500594-GGTLC3.18516335.18518165.rev.compl.REF       ------------------------------------------------------------ 31039

chimp.LOC112206744-LOC107973052-GGT2.revcompl                                      ------------------------------------------------------------ 61762
chimp54546-135457.revcompl.Pan.troglodytes.clone.rp43-41g5.GenBank:AC099533.36     ------------------------------------------------------------ 68417
human.FAM30B-LOC105372935-GGT2.NCBI.                                               ctggaaaggcctttctccaccccacgcccagccaatgcccactcatactttaaggccta 81467
human.FAM230E-LOC105377182-GGT3P.NCBI.                                             ------------------------------------------------------------ 57310
human.FAM230J-LOC105372942-GGTLC5P.NCBI.rev.compl                                  ------------------------------------------------------------ 46364
human.FAM230A.ncRNA.18487127.18500594-GGTLC3.18516335.18518165.rev.compl.REF       ------------------------------------------------------------ 31039

chimp.LOC112206744-LOC107973052-GGT2.revcompl                                      ------------------------------------------------------------ 61762
chimp54546-135457.revcompl.Pan.troglodytes.clone.rp43-41g5.GenBank:AC099533.36     ------------------------------------------------------------ 68417
human.FAM30B-LOC105372935-GGT2.NCBI.                                               gctcatccgtcatctctgctagggaaccttcctcagtgccttatgcaggcaaccattcct 81527
human.FAM230E-LOC105377182-GGT3P.NCBI.                                             ------------------------------------------------------------ 57310
human.FAM230J-LOC105372942-GGTLC5P.NCBI.rev.compl                                  ------------------------------------------------------------ 46364
human.FAM230A.ncRNA.18487127.18500594-GGTLC3.18516335.18518165.rev.compl.REF       ------------------------------------------------------------ 31039

chimp.LOC112206744-LOC107973052-GGT2.revcompl                                      ------------------------------------------------------------ 61762
chimp54546-135457.revcompl.Pan.troglodytes.clone.rp43-41g5.GenBank:AC099533.36     ------------------------------------------------------------ 68417
human.FAM30B-LOC105372935-GGT2.NCBI.                                               gccttgtgttgtgttgtgttgtgttgtgccttaagtcaaatccttaactcatctttgtgt 81587
human.FAM230E-LOC105377182-GGT3P.NCBI.                                             ------------------------------------------------------------ 57310
human.FAM230J-LOC105372942-GGTLC5P.NCBI.rev.compl                                  ------------------------------------------------------------ 46364
human.FAM230A.ncRNA.18487127.18500594-GGTLC3.18516335.18518165.rev.compl.REF       ------------------------------------------------------------ 31039

chimp.LOC112206744-LOC107973052-GGT2.revcompl                                      ------------------------------------------------------------ 61762
chimp54546-135457.revcompl.Pan.troglodytes.clone.rp43-41g5.GenBank:AC099533.36     ------------------------------------------------------------ 68417
human.FAM30B-LOC105372935-GGT2.NCBI.                                               gtccccttgcatttcagacagtaggtgcccaataaatgtttgtgggatgaatcaactggt 81647
human.FAM230E-LOC105377182-GGT3P.NCBI.                                             ------------------------------------------------------------ 57310
human.FAM230J-LOC105372942-GGTLC5P.NCBI.rev.compl                                  ------------------------------------------------------------ 46364
human.FAM230A.ncRNA.18487127.18500594-GGTLC3.18516335.18518165.rev.compl.REF       ------------------------------------------------------------ 31039

chimp.LOC112206744-LOC107973052-GGT2.revcompl                                      ------------------------------------------------------------ 61762
chimp54546-135457.revcompl.Pan.troglodytes.clone.rp43-41g5.GenBank:AC099533.36     ------------------------------------------------------------ 68417
human.FAM30B-LOC105372935-GGT2.NCBI.                                               tattctcctggaacaccctttcccagtacacctccactcgccagaacaactgtcttcctc 81707
human.FAM230E-LOC105377182-GGT3P.NCBI.                                             ------------------------------------------------------------ 57310
human.FAM230J-LOC105372942-GGTLC5P.NCBI.rev.compl                                  ------------------------------------------------------------ 46364
human.FAM230A.ncRNA.18487127.18500594-GGTLC3.18516335.18518165.rev.compl.REF       ------------------------------------------------------------ 31039

chimp.LOC112206744-LOC107973052-GGT2.revcompl                                      ------------------------------------------------------------ 61762
chimp54546-135457.revcompl.Pan.troglodytes.clone.rp43-41g5.GenBank:AC099533.36     ------------------------------------------------------------ 68417
human.FAM30B-LOC105372935-GGT2.NCBI.                                               acaggtgccagccatcaccccgaccacaccctctcaaaaggcctttgccagtcatcaaac 81767
human.FAM230E-LOC105377182-GGT3P.NCBI.                                             ------------------------------------------------------------ 57310
human.FAM230J-LOC105372942-GGTLC5P.NCBI.rev.compl                                  ------------------------------------------------------------ 46364
human.FAM230A.ncRNA.18487127.18500594-GGTLC3.18516335.18518165.rev.compl.REF       ------------------------------------------------------------ 31039
```

```
chimp.LOC112206744-LOC107973052-GGT2.revcompl                                         ------------------------------------------------------------ 61762
chimp54546-135457.revcompl.Pan.troglodytes.clone.rp43-41g5.GenBank:AC099533.36        ------------------------------------------------------------ 68417
human.FAM30B-LOC105372935-GGT2.NCBI.                                                  aaagcctcttgtctcctgagagccgctccacaatcaccttcccaagaagaattccccata 81827
human.FAM230E-LOC105377182-GGT3P.NCBI.                                                ------------------------------------------------------------ 57310
human.FAM230J-LOC105372942-GGTLC5P.NCBI.rev.compl                                     ------------------------------------------------------------ 46364
human.FAM230A.ncRNA.18487127.18500594-GGTLC3.18516335.18518165.rev.compl.REF          ------------------------------------------------------------ 31039

chimp.LOC112206744-LOC107973052-GGT2.revcompl                                         ------------------------------------------------------------ 61762
chimp54546-135457.revcompl.Pan.troglodytes.clone.rp43-41g5.GenBank:AC099533.36        ------------------------------------------------------------ 68417
human.FAM30B-LOC105372935-GGT2.NCBI.                                                  ccgagtccaccattccttccctgatcaaagttgctttcacaggcatcttttcagctatcc 81887
human.FAM230E-LOC105377182-GGT3P.NCBI.                                                ------------------------------------------------------------ 57310
human.FAM230J-LOC105372942-GGTLC5P.NCBI.rev.compl                                     ------------------------------------------------------------ 46364
human.FAM230A.ncRNA.18487127.18500594-GGTLC3.18516335.18518165.rev.compl.REF          ------------------------------------------------------------ 31039

chimp.LOC112206744-LOC107973052-GGT2.revcompl                                         ------------------------------------------------------------ 61762
chimp54546-135457.revcompl.Pan.troglodytes.clone.rp43-41g5.GenBank:AC099533.36        ------------------------------------------------------------ 68417
human.FAM30B-LOC105372935-GGT2.NCBI.                                                  actcttcacggcttggttttcatgcactctgagttggccctatctcccctttttggcctc 81947
human.FAM230E-LOC105377182-GGT3P.NCBI.                                                ------------------------------------------------------------ 57310
human.FAM230J-LOC105372942-GGTLC5P.NCBI.rev.compl                                     ------------------------------------------------------------ 46364
human.FAM230A.ncRNA.18487127.18500594-GGTLC3.18516335.18518165.rev.compl.REF          ------------------------------------------------------------ 31039

chimp.LOC112206744-LOC107973052-GGT2.revcompl                                         ------------------------------------------------------------ 61762
chimp54546-135457.revcompl.Pan.troglodytes.clone.rp43-41g5.GenBank:AC099533.36        ------------------------------------------------------------ 68417
human.FAM30B-LOC105372935-GGT2.NCBI.                                                  aagcacgtcaagaggaaggctggtgacagcctttcccgagctcaggacagttttcatgca 82007
human.FAM230E-LOC105377182-GGT3P.NCBI.                                                ------------------------------------------------------------ 57310
human.FAM230J-LOC105372942-GGTLC5P.NCBI.rev.compl                                     ------------------------------------------------------------ 46364
human.FAM230A.ncRNA.18487127.18500594-GGTLC3.18516335.18518165.rev.compl.REF          ------------------------------------------------------------ 31039

chimp.LOC112206744-LOC107973052-GGT2.revcompl                                         ------------------------------------------------------------ 61762
chimp54546-135457.revcompl.Pan.troglodytes.clone.rp43-41g5.GenBank:AC099533.36        ------------------------------------------------------------ 68417
human.FAM30B-LOC105372935-GGT2.NCBI.                                                  aactggttcttccaggttcttatgcctggaagaactgctgctctccaatagcatgaagta 82067
human.FAM230E-LOC105377182-GGT3P.NCBI.                                                ------------------------------------------------------------ 57310
human.FAM230J-LOC105372942-GGTLC5P.NCBI.rev.compl                                     ------------------------------------------------------------ 46364
human.FAM230A.ncRNA.18487127.18500594-GGTLC3.18516335.18518165.rev.compl.REF          ------------------------------------------------------------ 31039

chimp.LOC112206744-LOC107973052-GGT2.revcompl                                         ------------------------------------------------------------ 61762
chimp54546-135457.revcompl.Pan.troglodytes.clone.rp43-41g5.GenBank:AC099533.36        ------------------------------------------------------------ 68417
human.FAM30B-LOC105372935-GGT2.NCBI.                                                  gacaatgatgttcttagcatccagtcttaaaactagaaaaagtattggcactgctggatc 82127
human.FAM230E-LOC105377182-GGT3P.NCBI.                                                ------------------------------------------------------------ 57310
human.FAM230J-LOC105372942-GGTLC5P.NCBI.rev.compl                                     ------------------------------------------------------------ 46364
human.FAM230A.ncRNA.18487127.18500594-GGTLC3.18516335.18518165.rev.compl.REF          ------------------------------------------------------------ 31039

chimp.LOC112206744-LOC107973052-GGT2.revcompl                                         ------------------------------------------------------------ 61762
chimp54546-135457.revcompl.Pan.troglodytes.clone.rp43-41g5.GenBank:AC099533.36        ------------------------------------------------------------ 68417
human.FAM30B-LOC105372935-GGT2.NCBI.                                                  cccagctcctagaatggtgtgtggcacctaaacagagctcaaaaaatattagttgaatgg 82187
human.FAM230E-LOC105377182-GGT3P.NCBI.                                                ------------------------------------------------------------ 57310
human.FAM230J-LOC105372942-GGTLC5P.NCBI.rev.compl                                     ------------------------------------------------------------ 46364
human.FAM230A.ncRNA.18487127.18500594-GGTLC3.18516335.18518165.rev.compl.REF          ------------------------------------------------------------ 31039

chimp.LOC112206744-LOC107973052-GGT2.revcompl                                         ------------------------------------------------------------ 61762
chimp54546-135457.revcompl.Pan.troglodytes.clone.rp43-41g5.GenBank:AC099533.36        ------------------------------------------------------------ 68417
human.FAM30B-LOC105372935-GGT2.NCBI.                                                  aggaaagaatgaacagtcatctggcccagacttggagaagctgtgatctgtctagccctt 82247
human.FAM230E-LOC105377182-GGT3P.NCBI.                                                ------------------------------------------------------------ 57310
human.FAM230J-LOC105372942-GGTLC5P.NCBI.rev.compl                                     ------------------------------------------------------------ 46364
human.FAM230A.ncRNA.18487127.18500594-GGTLC3.18516335.18518165.rev.compl.REF          ------------------------------------------------------------ 31039

chimp.LOC112206744-LOC107973052-GGT2.revcompl                                         ------------------------------------------------------------ 61762
chimp54546-135457.revcompl.Pan.troglodytes.clone.rp43-41g5.GenBank:AC099533.36        ------------------------------------------------------------ 68417
human.FAM30B-LOC105372935-GGT2.NCBI.                                                  ctacaccaggagcgaggcctgaaaagctgcatgactcacccaaggtcacaagcaagccag 82307
human.FAM230E-LOC105377182-GGT3P.NCBI.                                                ------------------------------------------------------------ 57310
human.FAM230J-LOC105372942-GGTLC5P.NCBI.rev.compl                                     ------------------------------------------------------------ 46364
human.FAM230A.ncRNA.18487127.18500594-GGTLC3.18516335.18518165.rev.compl.REF          ------------------------------------------------------------ 31039

chimp.LOC112206744-LOC107973052-GGT2.revcompl                                         ------------------------------------------------------------ 61762
chimp54546-135457.revcompl.Pan.troglodytes.clone.rp43-41g5.GenBank:AC099533.36        ------------------------------------------------------------ 68417
human.FAM30B-LOC105372935-GGT2.NCBI.                                                  tgaagaggggctggaaccctgtcttttgcctcaaagcacagggccctccttctgtcccac 82367
human.FAM230E-LOC105377182-GGT3P.NCBI.                                                ------------------------------------------------------------ 57310
human.FAM230J-LOC105372942-GGTLC5P.NCBI.rev.compl                                     ------------------------------------------------------------ 46364
human.FAM230A.ncRNA.18487127.18500594-GGTLC3.18516335.18518165.rev.compl.REF          ------------------------------------------------------------ 31039

chimp.LOC112206744-LOC107973052-GGT2.revcompl                                         ------------------------------------------------------------ 61762
chimp54546-135457.revcompl.Pan.troglodytes.clone.rp43-41g5.GenBank:AC099533.36        ------------------------------------------------------------ 68417
human.FAM30B-LOC105372935-GGT2.NCBI.                                                  ccagctttctcaccacccaccccaccccttccacactccaaagacagcatctccgtgaga 82427
human.FAM230E-LOC105377182-GGT3P.NCBI.                                                ------------------------------------------------------------ 57310
human.FAM230J-LOC105372942-GGTLC5P.NCBI.rev.compl                                     ------------------------------------------------------------ 46364
human.FAM230A.ncRNA.18487127.18500594-GGTLC3.18516335.18518165.rev.compl.REF          ------------------------------------------------------------ 31039

chimp.LOC112206744-LOC107973052-GGT2.revcompl                                         ------------------------------------------------------------ 61762
chimp54546-135457.revcompl.Pan.troglodytes.clone.rp43-41g5.GenBank:AC099533.36        ------------------------------------------------------------ 68417
human.FAM30B-LOC105372935-GGT2.NCBI.                                                  agtgttgggcagtgtccaggagtgtttttcttcctggctttaaaacaaattgtgacaaat 82487
human.FAM230E-LOC105377182-GGT3P.NCBI.                                                ------------------------------------------------------------ 57310
human.FAM230J-LOC105372942-GGTLC5P.NCBI.rev.compl                                     ------------------------------------------------------------ 46364
human.FAM230A.ncRNA.18487127.18500594-GGTLC3.18516335.18518165.rev.compl.REF          ------------------------------------------------------------ 31039

chimp.LOC112206744-LOC107973052-GGT2.revcompl                                         ------------------------------------------------------------ 61762
chimp54546-135457.revcompl.Pan.troglodytes.clone.rp43-41g5.GenBank:AC099533.36        ------------------------------------------------------------ 68417
human.FAM30B-LOC105372935-GGT2.NCBI.                                                  ggctccacagatctaccgggccatggagctgagcagatgggactgcaattcacgctatta 82547
human.FAM230E-LOC105377182-GGT3P.NCBI.                                                ------------------------------------------------------------ 57310
human.FAM230J-LOC105372942-GGTLC5P.NCBI.rev.compl                                     ------------------------------------------------------------ 46364
human.FAM230A.ncRNA.18487127.18500594-GGTLC3.18516335.18518165.rev.compl.REF          ------------------------------------------------------------ 31039

chimp.LOC112206744-LOC107973052-GGT2.revcompl                                         ------------------------------------------------------------ 61762
chimp54546-135457.revcompl.Pan.troglodytes.clone.rp43-41g5.GenBank:AC099533.36        ------------------------------------------------------------ 68417
human.FAM30B-LOC105372935-GGT2.NCBI.                                                  actcaaattctacttgactttctttcaaatcagggcattttatctagaaattataaaaaa 82607
human.FAM230E-LOC105377182-GGT3P.NCBI.                                                ------------------------------------------------------------ 57310
human.FAM230J-LOC105372942-GGTLC5P.NCBI.rev.compl                                     ------------------------------------------------------------ 46364
human.FAM230A.ncRNA.18487127.18500594-GGTLC3.18516335.18518165.rev.compl.REF          ------------------------------------------------------------ 31039

chimp.LOC112206744-LOC107973052-GGT2.revcompl                                         ------------------------------------------------------------ 61762
chimp54546-135457.revcompl.Pan.troglodytes.clone.rp43-41g5.GenBank:AC099533.36        ------------------------------------------------------------ 68417
human.FAM30B-LOC105372935-GGT2.NCBI.                                                  ttgttggggggagtgcctttgtaaggaagtttgccagcacattcatttacaaatcgacttg 82667
human.FAM230E-LOC105377182-GGT3P.NCBI.                                                ------------------------------------------------------------ 57310
human.FAM230J-LOC105372942-GGTLC5P.NCBI.rev.compl                                     ------------------------------------------------------------ 46364
human.FAM230A.ncRNA.18487127.18500594-GGTLC3.18516335.18518165.rev.compl.REF          ------------------------------------------------------------ 31039

chimp.LOC112206744-LOC107973052-GGT2.revcompl                                         ------------------------------------------------------------ 61762
chimp54546-135457.revcompl.Pan.troglodytes.clone.rp43-41g5.GenBank:AC099533.36        ------------------------------------------------------------ 68417
human.FAM30B-LOC105372935-GGT2.NCBI.                                                  ctcctggccaccctcactgatcacggacttttactcacgctcccttactctctcccatca 82727
human.FAM230E-LOC105377182-GGT3P.NCBI.                                                ------------------------------------------------------------ 57310
human.FAM230J-LOC105372942-GGTLC5P.NCBI.rev.compl                                     ------------------------------------------------------------ 46364
human.FAM230A.ncRNA.18487127.18500594-GGTLC3.18516335.18518165.rev.compl.REF          ------------------------------------------------------------ 31039

chimp.LOC112206744-LOC107973052-GGT2.revcompl                                         ------------------------------------------------------------ 61762
chimp54546-135457.revcompl.Pan.troglodytes.clone.rp43-41g5.GenBank:AC099533.36        ------------------------------------------------------------ 68417
human.FAM30B-LOC105372935-GGT2.NCBI.                                                  aagagtccagcgcatggagagagtgatggaatcttgaggcccaccggtctaaccacccac 82787
human.FAM230E-LOC105377182-GGT3P.NCBI.                                                ------------------------------------------------------------ 57310
human.FAM230J-LOC105372942-GGTLC5P.NCBI.rev.compl                                     ------------------------------------------------------------ 46364
human.FAM230A.ncRNA.18487127.18500594-GGTLC3.18516335.18518165.rev.compl.REF          ------------------------------------------------------------ 31039

chimp.LOC112206744-LOC107973052-GGT2.revcompl                                         ------------------------------------------------------------ 61762
chimp54546-135457.revcompl.Pan.troglodytes.clone.rp43-41g5.GenBank:AC099533.36        ------------------------------------------------------------ 68417
human.FAM30B-LOC105372935-GGT2.NCBI.                                                  ctaagcttgactcccctgtgcctgtggacatcctgcgtgcagatgcacattcccactaat 82847
human.FAM230E-LOC105377182-GGT3P.NCBI.                                                ------------------------------------------------------------ 57310
human.FAM230J-LOC105372942-GGTLC5P.NCBI.rev.compl                                     ------------------------------------------------------------ 46364
human.FAM230A.ncRNA.18487127.18500594-GGTLC3.18516335.18518165.rev.compl.REF          ------------------------------------------------------------ 31039
```

```
chimp.LOC112206744-LOC107973052-GGT2.revcompl                                  ------------------------------------------------------------ 61762
chimp54546-135457.revcompl.Pan.troglodytes.clone.rp43-41g5.GenBank:AC099533.36 ------------------------------------------------------------ 68417
human.FAM30B-LOC105372935-GGT2.NCBI.                                           ggcagccaggggaaagcactattcgaaagttcttctttactttgaactcaaattgggctc 82907
human.FAM230E-LOC105377182-GGT3P.NCBI.                                         ------------------------------------------------------------ 57310
human.FAM230J-LOC105372942-GGTLC5P.NCBI.rev.compl                              ------------------------------------------------------------ 46364
human.FAM230A.ncRNA.18487127.18500594-GGTLC3.18516335.18518165.rev.compl.REF   ------------------------------------------------------------ 31039

chimp.LOC112206744-LOC107973052-GGT2.revcompl                                  ------------------------------------------------------------ 61762
chimp54546-135457.revcompl.Pan.troglodytes.clone.rp43-41g5.GenBank:AC099533.36 ------------------------------------------------------------ 68417
human.FAM30B-LOC105372935-GGT2.NCBI.                                           aggggtcctggatacaggacaaccccaatctcttatttattcaatggacaacaggtgagc 82967
human.FAM230E-LOC105377182-GGT3P.NCBI.                                         ------------------------------------------------------------ 57310
human.FAM230J-LOC105372942-GGTLC5P.NCBI.rev.compl                              ------------------------------------------------------------ 46364
human.FAM230A.ncRNA.18487127.18500594-GGTLC3.18516335.18518165.rev.compl.REF   ------------------------------------------------------------ 31039

chimp.LOC112206744-LOC107973052-GGT2.revcompl                                  ------------------------------------------------------------ 61762
chimp54546-135457.revcompl.Pan.troglodytes.clone.rp43-41g5.GenBank:AC099533.36 ------------------------------------------------------------ 68417
human.FAM30B-LOC105372935-GGT2.NCBI.                                           acatactaggcctggacactgatgtaggtaggcactaaagcttcagagagatgcaaatga 83027
human.FAM230E-LOC105377182-GGT3P.NCBI.                                         ------------------------------------------------------------ 57310
human.FAM230J-LOC105372942-GGTLC5P.NCBI.rev.compl                              ------------------------------------------------------------ 46364
human.FAM230A.ncRNA.18487127.18500594-GGTLC3.18516335.18518165.rev.compl.REF   ------------------------------------------------------------ 31039

chimp.LOC112206744-LOC107973052-GGT2.revcompl                                  ------------------------------------------------------------ 61762
chimp54546-135457.revcompl.Pan.troglodytes.clone.rp43-41g5.GenBank:AC099533.36 ------------------------------------------------------------ 68417
human.FAM30B-LOC105372935-GGT2.NCBI.                                           acaatctttaccctgaaggagctcatgttctatgggttcatttcctttccctgtgtctta 83087
human.FAM230E-LOC105377182-GGT3P.NCBI.                                         ------------------------------------------------------------ 57310
human.FAM230J-LOC105372942-GGTLC5P.NCBI.rev.compl                              ------------------------------------------------------------ 46364
human.FAM230A.ncRNA.18487127.18500594-GGTLC3.18516335.18518165.rev.compl.REF   ------------------------------------------------------------ 31039

chimp.LOC112206744-LOC107973052-GGT2.revcompl                                  ------------------------------------------------------------ 61762
chimp54546-135457.revcompl.Pan.troglodytes.clone.rp43-41g5.GenBank:AC099533.36 ------------------------------------------------------------ 68417
human.FAM30B-LOC105372935-GGT2.NCBI.                                           caaacaggttccagtttgtttccgtccttcagagtgtactgcctagaactgcatgctata 83147
human.FAM230E-LOC105377182-GGT3P.NCBI.                                         ------------------------------------------------------------ 57310
human.FAM230J-LOC105372942-GGTLC5P.NCBI.rev.compl                              ------------------------------------------------------------ 46364
human.FAM230A.ncRNA.18487127.18500594-GGTLC3.18516335.18518165.rev.compl.REF   ------------------------------------------------------------ 31039

chimp.LOC112206744-LOC107973052-GGT2.revcompl                                  ------------------------------------------------------------ 61762
chimp54546-135457.revcompl.Pan.troglodytes.clone.rp43-41g5.GenBank:AC099533.36 ------------------------------------------------------------ 68417
human.FAM30B-LOC105372935-GGT2.NCBI.                                           ctccacaagtggcctgaccagcttgtggacaaggatgaagcaagcagaccaccttctttc 83207
human.FAM230E-LOC105377182-GGT3P.NCBI.                                         ------------------------------------------------------------ 57310
human.FAM230J-LOC105372942-GGTLC5P.NCBI.rev.compl                              ------------------------------------------------------------ 46364
human.FAM230A.ncRNA.18487127.18500594-GGTLC3.18516335.18518165.rev.compl.REF   ------------------------------------------------------------ 31039

chimp.LOC112206744-LOC107973052-GGT2.revcompl                                  ------------------------------------------------------------ 61762
chimp54546-135457.revcompl.Pan.troglodytes.clone.rp43-41g5.GenBank:AC099533.36 ------------------------------------------------------------ 68417
human.FAM30B-LOC105372935-GGT2.NCBI.                                           agtgcagacaagatggctgtttagaggccacatcacgtggcagacacctgccaactaaaa 83267
human.FAM230E-LOC105377182-GGT3P.NCBI.                                         ------------------------------------------------------------ 57310
human.FAM230J-LOC105372942-GGTLC5P.NCBI.rev.compl                              ------------------------------------------------------------ 46364
human.FAM230A.ncRNA.18487127.18500594-GGTLC3.18516335.18518165.rev.compl.REF   ------------------------------------------------------------ 31039

chimp.LOC112206744-LOC107973052-GGT2.revcompl                                  ------------------------------------------------------------ 61762
chimp54546-135457.revcompl.Pan.troglodytes.clone.rp43-41g5.GenBank:AC099533.36 ------------------------------------------------------------ 68417
human.FAM30B-LOC105372935-GGT2.NCBI.                                           ccctccatccttcccacagcctcccctcctacactttgcagctggattcaaaaccttgc 83327
human.FAM230E-LOC105377182-GGT3P.NCBI.                                         ------------------------------------------------------------ 57310
human.FAM230J-LOC105372942-GGTLC5P.NCBI.rev.compl                              ------------------------------------------------------------ 46364
human.FAM230A.ncRNA.18487127.18500594-GGTLC3.18516335.18518165.rev.compl.REF   ------------------------------------------------------------ 31039

chimp.LOC112206744-LOC107973052-GGT2.revcompl                                  ------------------------------------------------------------ 61762
chimp54546-135457.revcompl.Pan.troglodytes.clone.rp43-41g5.GenBank:AC099533.36 ------------------------------------------------------------ 68417
human.FAM30B-LOC105372935-GGT2.NCBI.                                           attcacccttgttccatcaacagggccaggccatcagtctatctcatccggactgttttt 83387
human.FAM230E-LOC105377182-GGT3P.NCBI.                                         ------------------------------------------------------------ 57310
human.FAM230J-LOC105372942-GGTLC5P.NCBI.rev.compl                              ------------------------------------------------------------ 46364
human.FAM230A.ncRNA.18487127.18500594-GGTLC3.18516335.18518165.rev.compl.REF   ------------------------------------------------------------ 31039

chimp.LOC112206744-LOC107973052-GGT2.revcompl                                  ------------------------------------------------------------ 61762
chimp54546-135457.revcompl.Pan.troglodytes.clone.rp43-41g5.GenBank:AC099533.36 ------------------------------------------------------------ 68417
human.FAM30B-LOC105372935-GGT2.NCBI.                                           tatcttacttgccaacctacacacttgctgttccttcttgccttaaagtcatacagatga 83447
human.FAM230E-LOC105377182-GGT3P.NCBI.                                         ------------------------------------------------------------ 57310
human.FAM230J-LOC105372942-GGTLC5P.NCBI.rev.compl                              ------------------------------------------------------------ 46364
human.FAM230A.ncRNA.18487127.18500594-GGTLC3.18516335.18518165.rev.compl.REF   ------------------------------------------------------------ 31039

chimp.LOC112206744-LOC107973052-GGT2.revcompl                                  ------------------------------------------------------------ 61762
chimp54546-135457.revcompl.Pan.troglodytes.clone.rp43-41g5.GenBank:AC099533.36 ------------------------------------------------------------ 68417
human.FAM30B-LOC105372935-GGT2.NCBI.                                           ctttaggtgattcagatgagtgtgacctctactttcaccaaacacaatagtcaattcaga 83507
human.FAM230E-LOC105377182-GGT3P.NCBI.                                         ------------------------------------------------------------ 57310
human.FAM230J-LOC105372942-GGTLC5P.NCBI.rev.compl                              ------------------------------------------------------------ 46364
human.FAM230A.ncRNA.18487127.18500594-GGTLC3.18516335.18518165.rev.compl.REF   ------------------------------------------------------------ 31039

chimp.LOC112206744-LOC107973052-GGT2.revcompl                                  ------------------------------------------------------------ 61762
chimp54546-135457.revcompl.Pan.troglodytes.clone.rp43-41g5.GenBank:AC099533.36 ------------------------------------------------------------ 68417
human.FAM30B-LOC105372935-GGT2.NCBI.                                           aaggtcaaggacagaaccctccagcatgctaacgcagccaaagaagtccgaaaacaatac 83567
human.FAM230E-LOC105377182-GGT3P.NCBI.                                         ------------------------------------------------------------ 57310
human.FAM230J-LOC105372942-GGTLC5P.NCBI.rev.compl                              ------------------------------------------------------------ 46364
human.FAM230A.ncRNA.18487127.18500594-GGTLC3.18516335.18518165.rev.compl.REF   ------------------------------------------------------------ 31039

chimp.LOC112206744-LOC107973052-GGT2.revcompl                                  ------------------------------------------------------------ 61762
chimp54546-135457.revcompl.Pan.troglodytes.clone.rp43-41g5.GenBank:AC099533.36 ------------------------------------------------------------ 68417
human.FAM30B-LOC105372935-GGT2.NCBI.                                           ttggccacactgttttcctgcttacggatgtggactgctcttgtcctgcctgtaaatata 83627
human.FAM230E-LOC105377182-GGT3P.NCBI.                                         ------------------------------------------------------------ 57310
human.FAM230J-LOC105372942-GGTLC5P.NCBI.rev.compl                              ------------------------------------------------------------ 46364
human.FAM230A.ncRNA.18487127.18500594-GGTLC3.18516335.18518165.rev.compl.REF   ------------------------------------------------------------ 31039

chimp.LOC112206744-LOC107973052-GGT2.revcompl                                  ------------------------------------------------------------ 61762
chimp54546-135457.revcompl.Pan.troglodytes.clone.rp43-41g5.GenBank:AC099533.36 ------------------------------------------------------------ 68417
human.FAM30B-LOC105372935-GGT2.NCBI.                                           tcggatcgctctccttcttgctggcatccacagtaactatactcaataaacacaccaact 83687
human.FAM230E-LOC105377182-GGT3P.NCBI.                                         ------------------------------------------------------------ 57310
human.FAM230J-LOC105372942-GGTLC5P.NCBI.rev.compl                              ------------------------------------------------------------ 46364
human.FAM230A.ncRNA.18487127.18500594-GGTLC3.18516335.18518165.rev.compl.REF   ------------------------------------------------------------ 31039

chimp.LOC112206744-LOC107973052-GGT2.revcompl                                  ------------------------------------------------------------ 61762
chimp54546-135457.revcompl.Pan.troglodytes.clone.rp43-41g5.GenBank:AC099533.36 ------------------------------------------------------------ 68417
human.FAM30B-LOC105372935-GGT2.NCBI.                                           ttgatgccaaatcctaggccatcccacacatcttaaaaataaaacaaaaatatgctattc 83747
human.FAM230E-LOC105377182-GGT3P.NCBI.                                         ------------------------------------------------------------ 57310
human.FAM230J-LOC105372942-GGTLC5P.NCBI.rev.compl                              ------------------------------------------------------------ 46364
human.FAM230A.ncRNA.18487127.18500594-GGTLC3.18516335.18518165.rev.compl.REF   ------------------------------------------------------------ 31039

chimp.LOC112206744-LOC107973052-GGT2.revcompl                                  ------------------------------------------------------------ 61762
chimp54546-135457.revcompl.Pan.troglodytes.clone.rp43-41g5.GenBank:AC099533.36 ------------------------------------------------------------ 68417
human.FAM30B-LOC105372935-GGT2.NCBI.                                           cagctaccctcactctttttgtggtttcctgcagtccaactagctacaaatctcaaagcc 83807
human.FAM230E-LOC105377182-GGT3P.NCBI.                                         ------------------------------------------------------------ 57310
human.FAM230J-LOC105372942-GGTLC5P.NCBI.rev.compl                              ------------------------------------------------------------ 46364
human.FAM230A.ncRNA.18487127.18500594-GGTLC3.18516335.18518165.rev.compl.REF   ------------------------------------------------------------ 31039

chimp.LOC112206744-LOC107973052-GGT2.revcompl                                  ------------------------------------------------------------ 61762
chimp54546-135457.revcompl.Pan.troglodytes.clone.rp43-41g5.GenBank:AC099533.36 ------------------------------------------------------------ 68417
human.FAM30B-LOC105372935-GGT2.NCBI.                                           aaagattgtagctccctctcttttgttctttcatcaagccagggagtcctgctcctctct 83867
human.FAM230E-LOC105377182-GGT3P.NCBI.                                         ------------------------------------------------------------ 57310
human.FAM230J-LOC105372942-GGTLC5P.NCBI.rev.compl                              ------------------------------------------------------------ 46364
human.FAM230A.ncRNA.18487127.18500594-GGTLC3.18516335.18518165.rev.compl.REF   ------------------------------------------------------------ 31039

chimp.LOC112206744-LOC107973052-GGT2.revcompl                                  ------------------------------------------------------------ 61762
chimp54546-135457.revcompl.Pan.troglodytes.clone.rp43-41g5.GenBank:AC099533.36 ------------------------------------------------------------ 68417
human.FAM30B-LOC105372935-GGT2.NCBI.                                           ctctagtctctcctgtgtcagtgccacttgtcaactgcctaatccagccacgctgctctt 83927
human.FAM230E-LOC105377182-GGT3P.NCBI.                                         ------------------------------------------------------------ 57310
human.FAM230J-LOC105372942-GGTLC5P.NCBI.rev.compl                              ------------------------------------------------------------ 46364
human.FAM230A.ncRNA.18487127.18500594-GGTLC3.18516335.18518165.rev.compl.REF   ------------------------------------------------------------ 31039

chimp.LOC112206744-LOC107973052-GGT2.revcompl                                  ------------------------------------------------------------ 61762
```

```
chimp54546-135457.revcompl.Pan.troglodytes.clone.rp43-41g5.GenBank:AC099533.36      ------------------------------------------------------------ 68417
human.FAM30B-LOC105372935-GGT2.NCBI.                                                caatattaaaggtcttcagcatattttaaagttagtctcctactgccactcttttcccta 83987
human.FAM230E-LOC105377182-GGT3P.NCBI.                                              ------------------------------------------------------------ 57310
human.FAM230J-LOC105372942-GGTLC5P.NCBI.rev.compl                                   ------------------------------------------------------------ 46364
human.FAM230A.ncRNA.18487127.18500594-GGTLC3.18516335.18518165.rev.compl.REF        ------------------------------------------------------------ 31039


chimp.LOC112206744-LOC107973052-GGT2.revcompl                                       ------------------------------------------------------------ 61762
chimp54546-135457.revcompl.Pan.troglodytes.clone.rp43-41g5.GenBank:AC099533.36      ------------------------------------------------------------ 68417
human.FAM30B-LOC105372935-GGT2.NCBI.                                                ctcttgccagtccactcagtaaactactgccatacaggtctcccacagcagtttctcacc 84047
human.FAM230E-LOC105377182-GGT3P.NCBI.                                              ------------------------------------------------------------ 57310
human.FAM230J-LOC105372942-GGTLC5P.NCBI.rev.compl                                   ------------------------------------------------------------ 46364
human.FAM230A.ncRNA.18487127.18500594-GGTLC3.18516335.18518165.rev.compl.REF        ------------------------------------------------------------ 31039


chimp.LOC112206744-LOC107973052-GGT2.revcompl                                       ------------------------------------------------------------ 61762
chimp54546-135457.revcompl.Pan.troglodytes.clone.rp43-41g5.GenBank:AC099533.36      ------------------------------------------------------------ 68417
human.FAM30B-LOC105372935-GGT2.NCBI.                                                agataaacaatggccggagattccactttgccaatcacatcaagtccaaactccgtgtct 84107
human.FAM230E-LOC105377182-GGT3P.NCBI.                                              ------------------------------------------------------------ 57310
human.FAM230J-LOC105372942-GGTLC5P.NCBI.rev.compl                                   ------------------------------------------------------------ 46364
human.FAM230A.ncRNA.18487127.18500594-GGTLC3.18516335.18518165.rev.compl.REF        ------------------------------------------------------------ 31039


chimp.LOC112206744-LOC107973052-GGT2.revcompl                                       ------------------------------------------------------------ 61762
chimp54546-135457.revcompl.Pan.troglodytes.clone.rp43-41g5.GenBank:AC099533.36      ------------------------------------------------------------ 68417
human.FAM30B-LOC105372935-GGT2.NCBI.                                                tggtatccaaggcttctaaagtctggccccactccacctactcactctgacacccaccac 84167
human.FAM230E-LOC105377182-GGT3P.NCBI.                                              ------------------------------------------------------------ 57310
human.FAM230J-LOC105372942-GGTLC5P.NCBI.rev.compl                                   ------------------------------------------------------------ 46364
human.FAM230A.ncRNA.18487127.18500594-GGTLC3.18516335.18518165.rev.compl.REF        ------------------------------------------------------------ 31039


chimp.LOC112206744-LOC107973052-GGT2.revcompl                                       ------------------------------------------------------------ 61762
chimp54546-135457.revcompl.Pan.troglodytes.clone.rp43-41g5.GenBank:AC099533.36      ------------------------------------------------------------ 68417
human.FAM30B-LOC105372935-GGT2.NCBI.                                                tccctccaaacatccccggtacgtcttactggattttgccatttcctccaagttgaccag 84227
human.FAM230E-LOC105377182-GGT3P.NCBI.                                              ------------------------------------------------------------ 57310
human.FAM230J-LOC105372942-GGTLC5P.NCBI.rev.compl                                   ------------------------------------------------------------ 46364
human.FAM230A.ncRNA.18487127.18500594-GGTLC3.18516335.18518165.rev.compl.REF        ------------------------------------------------------------ 31039


chimp.LOC112206744-LOC107973052-GGT2.revcompl                                       ------------------------------------------------------------ 61762
chimp54546-135457.revcompl.Pan.troglodytes.clone.rp43-41g5.GenBank:AC099533.36      ------------------------------------------------------------ 68417
human.FAM30B-LOC105372935-GGT2.NCBI.                                                gagtcattcacacctaaaggctatgcttacatcgttttccatctcttgtgttggtgagct 84287
human.FAM230E-LOC105377182-GGT3P.NCBI.                                              ------------------------------------------------------------ 57310
human.FAM230J-LOC105372942-GGTLC5P.NCBI.rev.compl                                   ------------------------------------------------------------ 46364
human.FAM230A.ncRNA.18487127.18500594-GGTLC3.18516335.18518165.rev.compl.REF        ------------------------------------------------------------ 31039


chimp.LOC112206744-LOC107973052-GGT2.revcompl                                       ------------------------------------------------------------ 61762
chimp54546-135457.revcompl.Pan.troglodytes.clone.rp43-41g5.GenBank:AC099533.36      ------------------------------------------------------------ 68417
human.FAM30B-LOC105372935-GGT2.NCBI.                                                aaaaacaagatggaaaagaaaaaagaagaggaagagagaaaaagagaaaaggaagaaatg 84347
human.FAM230E-LOC105377182-GGT3P.NCBI.                                              ------------------------------------------------------------ 57310
human.FAM230J-LOC105372942-GGTLC5P.NCBI.rev.compl                                   ------------------------------------------------------------ 46364
human.FAM230A.ncRNA.18487127.18500594-GGTLC3.18516335.18518165.rev.compl.REF        ------------------------------------------------------------ 31039


chimp.LOC112206744-LOC107973052-GGT2.revcompl                                       ------------------------------------------------------------ 61762
chimp54546-135457.revcompl.Pan.troglodytes.clone.rp43-41g5.GenBank:AC099533.36      ------------------------------------------------------------ 68417
human.FAM30B-LOC105372935-GGT2.NCBI.                                                gggaaaacagcaataggaccagctaactcctacagcagagtcttgccaggccctccccta 84407
human.FAM230E-LOC105377182-GGT3P.NCBI.                                              ------------------------------------------------------------ 57310
human.FAM230J-LOC105372942-GGTLC5P.NCBI.rev.compl                                   ------------------------------------------------------------ 46364
human.FAM230A.ncRNA.18487127.18500594-GGTLC3.18516335.18518165.rev.compl.REF        ------------------------------------------------------------ 31039


chimp.LOC112206744-LOC107973052-GGT2.revcompl                                       ------------------------------------------------------------ 61762
chimp54546-135457.revcompl.Pan.troglodytes.clone.rp43-41g5.GenBank:AC099533.36      ------------------------------------------------------------ 68417
human.FAM30B-LOC105372935-GGT2.NCBI.                                                attgctttatatgtgttacaatgtttaaccctcacaactacaacaggagttgggtaccat 84467
human.FAM230E-LOC105377182-GGT3P.NCBI.                                              ------------------------------------------------------------ 57310
human.FAM230J-LOC105372942-GGTLC5P.NCBI.rev.compl                                   ------------------------------------------------------------ 46364
human.FAM230A.ncRNA.18487127.18500594-GGTLC3.18516335.18518165.rev.compl.REF        ------------------------------------------------------------ 31039


chimp.LOC112206744-LOC107973052-GGT2.revcompl                                       ------------------------------------------------------------ 61762
chimp54546-135457.revcompl.Pan.troglodytes.clone.rp43-41g5.GenBank:AC099533.36      ------------------------------------------------------------ 68417
human.FAM30B-LOC105372935-GGT2.NCBI.                                                tccatccccacctgcagatgagtgtggcaccccaagattaagtatctgcccagggtcacg 84527
human.FAM230E-LOC105377182-GGT3P.NCBI.                                              ------------------------------------------------------------ 57310
human.FAM230J-LOC105372942-GGTLC5P.NCBI.rev.compl                                   ------------------------------------------------------------ 46364
human.FAM230A.ncRNA.18487127.18500594-GGTLC3.18516335.18518165.rev.compl.REF        ------------------------------------------------------------ 31039


chimp.LOC112206744-LOC107973052-GGT2.revcompl                                       ------------------------------------------------------------ 61762
chimp54546-135457.revcompl.Pan.troglodytes.clone.rp43-41g5.GenBank:AC099533.36      ------------------------------------------------------------ 68417
human.FAM30B-LOC105372935-GGT2.NCBI.                                                tgaaaagtctgcatgcatgctgaaccattccagctgcaccactgcccaggaaaaaagtca 84587
human.FAM230E-LOC105377182-GGT3P.NCBI.                                              ------------------------------------------------------------ 57310
human.FAM230J-LOC105372942-GGTLC5P.NCBI.rev.compl                                   ------------------------------------------------------------ 46364
human.FAM230A.ncRNA.18487127.18500594-GGTLC3.18516335.18518165.rev.compl.REF        ------------------------------------------------------------ 31039


chimp.LOC112206744-LOC107973052-GGT2.revcompl                                       ------------------------------------------------------------ 61762
chimp54546-135457.revcompl.Pan.troglodytes.clone.rp43-41g5.GenBank:AC099533.36      ------------------------------------------------------------ 68417
human.FAM30B-LOC105372935-GGT2.NCBI.                                                caggtaggaagacaagcatagaattaagaacaaagcaactaactgtaccaggtgctacag 84647
human.FAM230E-LOC105377182-GGT3P.NCBI.                                              ------------------------------------------------------------ 57310
human.FAM230J-LOC105372942-GGTLC5P.NCBI.rev.compl                                   ------------------------------------------------------------ 46364
human.FAM230A.ncRNA.18487127.18500594-GGTLC3.18516335.18518165.rev.compl.REF        ------------------------------------------------------------ 31039


chimp.LOC112206744-LOC107973052-GGT2.revcompl                                       ------------------------------------------------------------ 61762
chimp54546-135457.revcompl.Pan.troglodytes.clone.rp43-41g5.GenBank:AC099533.36      ------------------------------------------------------------ 68417
human.FAM30B-LOC105372935-GGT2.NCBI.                                                aagggtggtcagcctggaaaatcaggaggtgcattccttgggagattgccgcaatcagag 84707
human.FAM230E-LOC105377182-GGT3P.NCBI.                                              ------------------------------------------------------------ 57310
human.FAM230J-LOC105372942-GGTLC5P.NCBI.rev.compl                                   ------------------------------------------------------------ 46364
human.FAM230A.ncRNA.18487127.18500594-GGTLC3.18516335.18518165.rev.compl.REF        ------------------------------------------------------------ 31039


chimp.LOC112206744-LOC107973052-GGT2.revcompl                                       ------------------------------------------------------------ 61762
chimp54546-135457.revcompl.Pan.troglodytes.clone.rp43-41g5.GenBank:AC099533.36      ------------------------------------------------------------ 68417
human.FAM30B-LOC105372935-GGT2.NCBI.                                                tctgggagtatgcaggagactggcaggcagcccagggacaggtgaggcacaggcactcat 84767
human.FAM230E-LOC105377182-GGT3P.NCBI.                                              ------------------------------------------------------------ 57310
human.FAM230J-LOC105372942-GGTLC5P.NCBI.rev.compl                                   ------------------------------------------------------------ 46364
human.FAM230A.ncRNA.18487127.18500594-GGTLC3.18516335.18518165.rev.compl.REF        ------------------------------------------------------------ 31039


chimp.LOC112206744-LOC107973052-GGT2.revcompl                                       ------------------------------------------------------------ 61762
chimp54546-135457.revcompl.Pan.troglodytes.clone.rp43-41g5.GenBank:AC099533.36      ------------------------------------------------------------ 68417
human.FAM30B-LOC105372935-GGT2.NCBI.                                                tcccttccatgtgagaagtcagggcacgtgcaaagccaaggccagaggacaaaggtgagc 84827
human.FAM230E-LOC105377182-GGT3P.NCBI.                                              ------------------------------------------------------------ 57310
human.FAM230J-LOC105372942-GGTLC5P.NCBI.rev.compl                                   ------------------------------------------------------------ 46364
human.FAM230A.ncRNA.18487127.18500594-GGTLC3.18516335.18518165.rev.compl.REF        ------------------------------------------------------------ 31039


chimp.LOC112206744-LOC107973052-GGT2.revcompl                                       ------------------------------------------------------------ 61762
chimp54546-135457.revcompl.Pan.troglodytes.clone.rp43-41g5.GenBank:AC099533.36      ------------------------------------------------------------ 68417
human.FAM30B-LOC105372935-GGT2.NCBI.                                                tcgccaatagggccaatactgggtctggctggaaggggagggtggacagagatgggtact 84887
human.FAM230E-LOC105377182-GGT3P.NCBI.                                              ------------------------------------------------------------ 57310
human.FAM230J-LOC105372942-GGTLC5P.NCBI.rev.compl                                   ------------------------------------------------------------ 46364
human.FAM230A.ncRNA.18487127.18500594-GGTLC3.18516335.18518165.rev.compl.REF        ------------------------------------------------------------ 31039


chimp.LOC112206744-LOC107973052-GGT2.revcompl                                       ------------------------------------------------------------ 61762
chimp54546-135457.revcompl.Pan.troglodytes.clone.rp43-41g5.GenBank:AC099533.36      ------------------------------------------------------------ 68417
human.FAM30B-LOC105372935-GGT2.NCBI.                                                aagaagtgaagatggacaggcaggaccaggtcttaaagggtttgtgtcttgctaaagatg 84947
human.FAM230E-LOC105377182-GGT3P.NCBI.                                              ------------------------------------------------------------ 57310
human.FAM230J-LOC105372942-GGTLC5P.NCBI.rev.compl                                   ------------------------------------------------------------ 46364
human.FAM230A.ncRNA.18487127.18500594-GGTLC3.18516335.18518165.rev.compl.REF        ------------------------------------------------------------ 31039


chimp.LOC112206744-LOC107973052-GGT2.revcompl                                       ------------------------------------------------------------ 61762
chimp54546-135457.revcompl.Pan.troglodytes.clone.rp43-41g5.GenBank:AC099533.36      ------------------------------------------------------------ 68417
human.FAM30B-LOC105372935-GGT2.NCBI.                                                gacccttaattccgtgacacatctattcaaccactcctcattatctacctaaattctact 85007
human.FAM230E-LOC105377182-GGT3P.NCBI.                                              ------------------------------------------------------------ 57310
human.FAM230J-LOC105372942-GGTLC5P.NCBI.rev.compl                                   ------------------------------------------------------------ 46364
human.FAM230A.ncRNA.18487127.18500594-GGTLC3.18516335.18518165.rev.compl.REF        ------------------------------------------------------------ 31039


chimp.LOC112206744-LOC107973052-GGT2.revcompl                                       ------------------------------------------------------------ 61762
chimp54546-135457.revcompl.Pan.troglodytes.clone.rp43-41g5.GenBank:AC099533.36      ------------------------------------------------------------ 68417
```

```
human.FAM30B-LOC105372935-GGT2.NCBI.                                                gtgcctttgcaaagcccagctcaagtcctttttcctctcaggcctgtcctccctactcct 85067
human.FAM230E-LOC105377182-GGT3P.NCBI.                                              ------------------------------------------------------------ 57310
human.FAM230J-LOC105372942-GGTLC5P.NCBI.rev.compl                                   ------------------------------------------------------------ 46364
human.FAM230A.ncRNA.18487127.18500594-GGTLC3.18516335.18518165.rev.compl.REF        ------------------------------------------------------------ 31039


chimp.LOC112206744-LOC107973052-GGT2.revcompl                                       ------------------------------------------------------------ 61762
chimp54546-135457.revcompl.Pan.troglodytes.clone.rp43-41g5.GenBank:AC099533.36      ------------------------------------------------------------ 68417
human.FAM30B-LOC105372935-GGT2.NCBI.                                                gagcaaaccacgggtgtggagacaggtcccaaccacctgcagatgacagaaggaagctgc 85127
human.FAM230E-LOC105377182-GGT3P.NCBI.                                              ------------------------------------------------------------ 57310
human.FAM230J-LOC105372942-GGTLC5P.NCBI.rev.compl                                   ------------------------------------------------------------ 46364
human.FAM230A.ncRNA.18487127.18500594-GGTLC3.18516335.18518165.rev.compl.REF        ------------------------------------------------------------ 31039


chimp.LOC112206744-LOC107973052-GGT2.revcompl                                       ------------------------------------------------------------ 61762
chimp54546-135457.revcompl.Pan.troglodytes.clone.rp43-41g5.GenBank:AC099533.36      ------------------------------------------------------------ 68417
human.FAM30B-LOC105372935-GGT2.NCBI.                                                atctaggatcggtctacacagaggcaagaagggatcctcctctctgccttctcctttca 85187
human.FAM230E-LOC105377182-GGT3P.NCBI.                                              ------------------------------------------------------------ 57310
human.FAM230J-LOC105372942-GGTLC5P.NCBI.rev.compl                                   ------------------------------------------------------------ 46364
human.FAM230A.ncRNA.18487127.18500594-GGTLC3.18516335.18518165.rev.compl.REF        ------------------------------------------------------------ 31039


chimp.LOC112206744-LOC107973052-GGT2.revcompl                                       ------------------------------------------------------------ 61762
chimp54546-135457.revcompl.Pan.troglodytes.clone.rp43-41g5.GenBank:AC099533.36      ------------------------------------------------------------ 68417
human.FAM30B-LOC105372935-GGT2.NCBI.                                                tgctgttcttactttctctcaaaaagaaagaaaatacagtgtcattgcttcttccctaag 85247
human.FAM230E-LOC105377182-GGT3P.NCBI.                                              ------------------------------------------------------------ 57310
human.FAM230J-LOC105372942-GGTLC5P.NCBI.rev.compl                                   ------------------------------------------------------------ 46364
human.FAM230A.ncRNA.18487127.18500594-GGTLC3.18516335.18518165.rev.compl.REF        ------------------------------------------------------------ 31039


chimp.LOC112206744-LOC107973052-GGT2.revcompl                                       ------------------------------------------------------------ 61762
chimp54546-135457.revcompl.Pan.troglodytes.clone.rp43-41g5.GenBank:AC099533.36      ------------------------------------------------------------ 68417
human.FAM30B-LOC105372935-GGT2.NCBI.                                                cttgttaagtcaaagcaagtcaaagcatcagagcttcattacttggctttttctagctat 85307
human.FAM230E-LOC105377182-GGT3P.NCBI.                                              ------------------------------------------------------------ 57310
human.FAM230J-LOC105372942-GGTLC5P.NCBI.rev.compl                                   ------------------------------------------------------------ 46364
human.FAM230A.ncRNA.18487127.18500594-GGTLC3.18516335.18518165.rev.compl.REF        ------------------------------------------------------------ 31039


chimp.LOC112206744-LOC107973052-GGT2.revcompl                                       ------------------------------------------------------------ 61762
chimp54546-135457.revcompl.Pan.troglodytes.clone.rp43-41g5.GenBank:AC099533.36      ------------------------------------------------------------ 68417
human.FAM30B-LOC105372935-GGT2.NCBI.                                                aagcgtctggaaaatatcaggagtcactggcctctaggaatccattaagttaaaggagct 85367
human.FAM230E-LOC105377182-GGT3P.NCBI.                                              ------------------------------------------------------------ 57310
human.FAM230J-LOC105372942-GGTLC5P.NCBI.rev.compl                                   ------------------------------------------------------------ 46364
human.FAM230A.ncRNA.18487127.18500594-GGTLC3.18516335.18518165.rev.compl.REF        ------------------------------------------------------------ 31039


chimp.LOC112206744-LOC107973052-GGT2.revcompl                                       ------------------------------------------------------------ 61762
chimp54546-135457.revcompl.Pan.troglodytes.clone.rp43-41g5.GenBank:AC099533.36      -------------------------------ATAATCCCAGCACTTTGAGAGGCCGAGGT 68446
human.FAM30B-LOC105372935-GGT2.NCBI.                                                ccatgatctactatcataagatgacaaagaaaatatgcactttctttgggaggtcaagtt 85427
human.FAM230E-LOC105377182-GGT3P.NCBI.                                              ------------------------------------------------------------ 57310
human.FAM230J-LOC105372942-GGTLC5P.NCBI.rev.compl                                   ------------------------------------------------------------ 46364
human.FAM230A.ncRNA.18487127.18500594-GGTLC3.18516335.18518165.rev.compl.REF        ------------------------------------------------------------ 31039


chimp.LOC112206744-LOC107973052-GGT2.revcompl                                       ------------------------------------------------------------ 61762
chimp54546-135457.revcompl.Pan.troglodytes.clone.rp43-41g5.GenBank:AC099533.36      GGGCGGATTGCCTGAAGTCAGAAGTTCAAGACTAGTCTGACCATCATAGGGAAACCCAGT 68506
human.FAM30B-LOC105372935-GGT2.NCBI.                                                gggcgaatcacttgaggtcaggagttcaagaccaatctggccaacatggtgaaaccccgt 85487
human.FAM230E-LOC105377182-GGT3P.NCBI.                                              ------------------------------------------------------------ 57310
human.FAM230J-LOC105372942-GGTLC5P.NCBI.rev.compl                                   ------------------------------------------------------------ 46364
human.FAM230A.ncRNA.18487127.18500594-GGTLC3.18516335.18518165.rev.compl.REF        ------------------------------------------------------------ 31039


chimp.LOC112206744-LOC107973052-GGT2.revcompl                                       ------------------------------------------------------------ 61762
chimp54546-135457.revcompl.Pan.troglodytes.clone.rp43-41g5.GenBank:AC099533.36      CTCTACTACAAATACAAAAAAATTATCCAGGCATGGTGGCGTGCACCTGTAATCCCAGCT 68566
human.FAM30B-LOC105372935-GGT2.NCBI.                                                ctctactgaaa--acacaaaaattagctgggcttggtggcaggcacctgtaatcccagct 85545
human.FAM230E-LOC105377182-GGT3P.NCBI.                                              ------------------------------------------------------------ 57310
human.FAM230J-LOC105372942-GGTLC5P.NCBI.rev.compl                                   ------------------------------------------------------------ 46364
human.FAM230A.ncRNA.18487127.18500594-GGTLC3.18516335.18518165.rev.compl.REF        ------------------------------------------------------------ 31039


chimp.LOC112206744-LOC107973052-GGT2.revcompl                                       ------------------------------------------------------------ 61762
chimp54546-135457.revcompl.Pan.troglodytes.clone.rp43-41g5.GenBank:AC099533.36      CCTTGGAAGGCTGCGGCAGGGGAATTGCTTGA------ACCAGGGAGATGGAGGTTGCAG 68620
human.FAM30B-LOC105372935-GGT2.NCBI.                                                actcaggagactgaggcaggagaatcgattgaacccttacccaggagacagaggttgcaa 85605
human.FAM230E-LOC105377182-GGT3P.NCBI.                                              ------------------------------------------------------------ 57310
human.FAM230J-LOC105372942-GGTLC5P.NCBI.rev.compl                                   ------------------------------------------------------------ 46364
human.FAM230A.ncRNA.18487127.18500594-GGTLC3.18516335.18518165.rev.compl.REF        ------------------------------------------------------------ 31039


chimp.LOC112206744-LOC107973052-GGT2.revcompl                                       ------------------------------------------------------------ 61762
chimp54546-135457.revcompl.Pan.troglodytes.clone.rp43-41g5.GenBank:AC099533.36      TGAGCTGAGATCACGCCACTGCACTCCAGCCTGGGCAACAGAGTGATACTCCGTCTCAAA 68680
human.FAM30B-LOC105372935-GGT2.NCBI.                                                tgagccgagatggcaccacggcactccagcctgagcaacagcgtg--actccatctcaaa 85663
human.FAM230E-LOC105377182-GGT3P.NCBI.                                              ------------------------------------------------------------ 57310
human.FAM230J-LOC105372942-GGTLC5P.NCBI.rev.compl                                   ------------------------------------------------------------ 46364
human.FAM230A.ncRNA.18487127.18500594-GGTLC3.18516335.18518165.rev.compl.REF        ------------------------------------------------------------ 31039


chimp.LOC112206744-LOC107973052-GGT2.revcompl                                       ------------------------------------------------------------ 61762
chimp54546-135457.revcompl.Pan.troglodytes.clone.rp43-41g5.GenBank:AC099533.36      AACAAACAAAC------------------------------------------------- 68691
human.FAM30B-LOC105372935-GGT2.NCBI.                                                aaaaaaaaaaagaaagaaagaaaggaaggaaggaagaaagaagaaaatatgcattttctc 85723
human.FAM230E-LOC105377182-GGT3P.NCBI.                                              ------------------------------------------------------------ 57310
human.FAM230J-LOC105372942-GGTLC5P.NCBI.rev.compl                                   ------------------------------------------------------------ 46364
human.FAM230A.ncRNA.18487127.18500594-GGTLC3.18516335.18518165.rev.compl.REF        ------------------------------------------------------------ 31039


chimp.LOC112206744-LOC107973052-GGT2.revcompl                                       ------------------------------------------------------------ 61762
chimp54546-135457.revcompl.Pan.troglodytes.clone.rp43-41g5.GenBank:AC099533.36      ------------------------------------------------------------ 68691
human.FAM30B-LOC105372935-GGT2.NCBI.                                                aggattaaataacaaaaaatgccaagtttcttagctccagtcaagaaaatgacatcatgt 85783
human.FAM230E-LOC105377182-GGT3P.NCBI.                                              ------------------------------------------------------------ 57310
human.FAM230J-LOC105372942-GGTLC5P.NCBI.rev.compl                                   ------------------------------------------------------------ 46364
human.FAM230A.ncRNA.18487127.18500594-GGTLC3.18516335.18518165.rev.compl.REF        ------------------------------------------------------------ 31039


chimp.LOC112206744-LOC107973052-GGT2.revcompl                                       ------------------------------------------------------------ 61762
chimp54546-135457.revcompl.Pan.troglodytes.clone.rp43-41g5.GenBank:AC099533.36      ------------------------------------------------------------ 68691
human.FAM30B-LOC105372935-GGT2.NCBI.                                                gacaacagcagccattgcaagtttatcagaagatgaacaattacagatttttcatttctg 85843
human.FAM230E-LOC105377182-GGT3P.NCBI.                                              ------------------------------------------------------------ 57310
human.FAM230J-LOC105372942-GGTLC5P.NCBI.rev.compl                                   ------------------------------------------------------------ 46364
human.FAM230A.ncRNA.18487127.18500594-GGTLC3.18516335.18518165.rev.compl.REF        ------------------------------------------------------------ 31039


chimp.LOC112206744-LOC107973052-GGT2.revcompl                                       ------------------------------------------------------------ 61762
chimp54546-135457.revcompl.Pan.troglodytes.clone.rp43-41g5.GenBank:AC099533.36      ------------------------------------------------------------ 68691
human.FAM30B-LOC105372935-GGT2.NCBI.                                                aataaggaaccatttacacagacacatgtataacaaatgtctgtgcacatccaggacttg 85903
human.FAM230E-LOC105377182-GGT3P.NCBI.                                              ------------------------------------------------------------ 57310
human.FAM230J-LOC105372942-GGTLC5P.NCBI.rev.compl                                   ------------------------------------------------------------ 46364
human.FAM230A.ncRNA.18487127.18500594-GGTLC3.18516335.18518165.rev.compl.REF        ------------------------------------------------------------ 31039


chimp.LOC112206744-LOC107973052-GGT2.revcompl                                       ------------------------------------------------------------ 61762
chimp54546-135457.revcompl.Pan.troglodytes.clone.rp43-41g5.GenBank:AC099533.36      ------------------------------------------------------------ 68691
human.FAM30B-LOC105372935-GGT2.NCBI.                                                tcaaatcttaacattttgccacacctggtttagtttttttatttttcctttcaagaaaga 85963
human.FAM230E-LOC105377182-GGT3P.NCBI.                                              ------------------------------------------------------------ 57310
human.FAM230J-LOC105372942-GGTLC5P.NCBI.rev.compl                                   ------------------------------------------------------------ 46364
human.FAM230A.ncRNA.18487127.18500594-GGTLC3.18516335.18518165.rev.compl.REF        ------------------------------------------------------------ 31039


chimp.LOC112206744-LOC107973052-GGT2.revcompl                                       ------------------------------------------------------------ 61762
chimp54546-135457.revcompl.Pan.troglodytes.clone.rp43-41g5.GenBank:AC099533.36      ------------------------------------------------------------ 68691
human.FAM30B-LOC105372935-GGT2.NCBI.                                                aagcattacaggctgtgcttcaatccctatttcacccccttcaagccccactattccac 86023
human.FAM230E-LOC105377182-GGT3P.NCBI.                                              ------------------------------------------------------------ 57310
human.FAM230J-LOC105372942-GGTLC5P.NCBI.rev.compl                                   ------------------------------------------------------------ 46364
human.FAM230A.ncRNA.18487127.18500594-GGTLC3.18516335.18518165.rev.compl.REF        ------------------------------------------------------------ 31039


chimp.LOC112206744-LOC107973052-GGT2.revcompl                                       ------------------------------------------------------------ 61762
chimp54546-135457.revcompl.Pan.troglodytes.clone.rp43-41g5.GenBank:AC099533.36      ------------------------------------------------------------ 68691
human.FAM30B-LOC105372935-GGT2.NCBI.                                                ccatcccagaaatgacaacatcccgaatttggtatttaccactcttgcacatttattttt 86083
human.FAM230E-LOC105377182-GGT3P.NCBI.                                              ------------------------------------------------------------ 57310
human.FAM230J-LOC105372942-GGTLC5P.NCBI.rev.compl                                   ------------------------------------------------------------ 46364
human.FAM230A.ncRNA.18487127.18500594-GGTLC3.18516335.18518165.rev.compl.REF        ------------------------------------------------------------ 31039


chimp.LOC112206744-LOC107973052-GGT2.revcompl                                       ------------------------------------------------------------ 61762
chimp54546-135457.revcompl.Pan.troglodytes.clone.rp43-41g5.GenBank:AC099533.36      ------------------------------------------------------------ 68691
human.FAM30B-LOC105372935-GGT2.NCBI.                                                atgcttttactacctatgtatgtagccataaacaatatacagagtttttggatgctataa 86143
```

```
human.FAM230E-LOC105377182-GGT3P.NCBI.                                    ------------------------------------------------------------ 57310
human.FAM230J-LOC105372942-GGTLC5P.NCBI.rev.compl                         ------------------------------------------------------------ 46364
human.FAM230A.ncRNA.18487127.18500594-GGTLC3.18516335.18518165.rev.compl.REF ------------------------------------------------------------ 31039

chimp.LOC112206744-LOC107973052-GGT2.revcompl                             ------------------------------------------------------------ 61762
chimp54546-135457.revcompl.Pan.troglodytes.clone.rp43-41g5.GenBank:AC099533.36 ------------------------------------------------------------ 68691
human.FAM30B-LOC105372935-GGT2.NCBI.                                      aactgtttaaaatggcatcacactgtatgcacccttttgcatctgttttaattgacaaaa 86203
human.FAM230E-LOC105377182-GGT3P.NCBI.                                    ------------------------------------------------------------ 57310
human.FAM230J-LOC105372942-GGTLC5P.NCBI.rev.compl                         ------------------------------------------------------------ 46364
human.FAM230A.ncRNA.18487127.18500594-GGTLC3.18516335.18518165.rev.compl.REF ------------------------------------------------------------ 31039

chimp.LOC112206744-LOC107973052-GGT2.revcompl                             ------------------------------------------------------------ 61762
chimp54546-135457.revcompl.Pan.troglodytes.clone.rp43-41g5.GenBank:AC099533.36 ------------------------------------------------------------ 68691
human.FAM30B-LOC105372935-GGT2.NCBI.                                      cattttgagggtaatccttgttgacatatgtcactccagggcattcctttcaactgctt 86263
human.FAM230E-LOC105377182-GGT3P.NCBI.                                    ------------------------------------------------------------ 57310
human.FAM230J-LOC105372942-GGTLC5P.NCBI.rev.compl                         ------------------------------------------------------------ 46364
human.FAM230A.ncRNA.18487127.18500594-GGTLC3.18516335.18518165.rev.compl.REF ------------------------------------------------------------ 31039

chimp.LOC112206744-LOC107973052-GGT2.revcompl                             ------------------------------------------------------------ 61762
chimp54546-135457.revcompl.Pan.troglodytes.clone.rp43-41g5.GenBank:AC099533.36 ------------------------------------------------------------ 68691
human.FAM30B-LOC105372935-GGT2.NCBI.                                      tataccatggtccactgtctggataatcattatctatagcacaatgtattcattttccta 86323
human.FAM230E-LOC105377182-GGT3P.NCBI.                                    ------------------------------------------------------------ 57310
human.FAM230J-LOC105372942-GGTLC5P.NCBI.rev.compl                         ------------------------------------------------------------ 46364
human.FAM230A.ncRNA.18487127.18500594-GGTLC3.18516335.18518165.rev.compl.REF ------------------------------------------------------------ 31039

chimp.LOC112206744-LOC107973052-GGT2.revcompl                             ------------------------------------------------------------ 61762
chimp54546-135457.revcompl.Pan.troglodytes.clone.rp43-41g5.GenBank:AC099533.36 ------------------------------------------------------------ 68691
human.FAM30B-LOC105372935-GGT2.NCBI.                                      ctgataaattttaagtggtttccaacttttcattgcaataaatattcttgtaaatcttc 86383
human.FAM230E-LOC105377182-GGT3P.NCBI.                                    ------------------------------------------------------------ 57310
human.FAM230J-LOC105372942-GGTLC5P.NCBI.rev.compl                         ------------------------------------------------------------ 46364
human.FAM230A.ncRNA.18487127.18500594-GGTLC3.18516335.18518165.rev.compl.REF ------------------------------------------------------------ 31039

chimp.LOC112206744-LOC107973052-GGT2.revcompl                             ------------------------------------------------------------ 61762
chimp54546-135457.revcompl.Pan.troglodytes.clone.rp43-41g5.GenBank:AC099533.36 ------------------------------------------------------------ 68691
human.FAM30B-LOC105372935-GGT2.NCBI.                                      gtccaaggttatcttcaaccacactaggcattgcccaaattgctctgcaacatcagtgca 86443
human.FAM230E-LOC105377182-GGT3P.NCBI.                                    ------------------------------------------------------------ 57310
human.FAM230J-LOC105372942-GGTLC5P.NCBI.rev.compl                         ------------------------------------------------------------ 46364
human.FAM230A.ncRNA.18487127.18500594-GGTLC3.18516335.18518165.rev.compl.REF ------------------------------------------------------------ 31039

chimp.LOC112206744-LOC107973052-GGT2.revcompl                             ------------------------------------------------------------ 61762
chimp54546-135457.revcompl.Pan.troglodytes.clone.rp43-41g5.GenBank:AC099533.36 ------------------------------------------------------------ 68691
human.FAM30B-LOC105372935-GGT2.NCBI.                                      ccaatttacattgccaagatcagtgtttaagtgtttctattctttgatgtctttgtcaac 86503
human.FAM230E-LOC105377182-GGT3P.NCBI.                                    ------------------------------------------------------------ 57310
human.FAM230J-LOC105372942-GGTLC5P.NCBI.rev.compl                         ------------------------------------------------------------ 46364
human.FAM230A.ncRNA.18487127.18500594-GGTLC3.18516335.18518165.rev.compl.REF ------------------------------------------------------------ 31039

chimp.LOC112206744-LOC107973052-GGT2.revcompl                             ------------------------------------------------------------ 61762
chimp54546-135457.revcompl.Pan.troglodytes.clone.rp43-41g5.GenBank:AC099533.36 ------------------------------------------------------------ 68691
human.FAM30B-LOC105372935-GGT2.NCBI.                                      cttggtactatttgactactacacttttcaccaatcagatataaggatgaaatggtgcct 86563
human.FAM230E-LOC105377182-GGT3P.NCBI.                                    ------------------------------------------------------------ 57310
human.FAM230J-LOC105372942-GGTLC5P.NCBI.rev.compl                         ------------------------------------------------------------ 46364
human.FAM230A.ncRNA.18487127.18500594-GGTLC3.18516335.18518165.rev.compl.REF ------------------------------------------------------------ 31039

chimp.LOC112206744-LOC107973052-GGT2.revcompl                             ------------------------------------------------------------ 61762
chimp54546-135457.revcompl.Pan.troglodytes.clone.rp43-41g5.GenBank:AC099533.36 ------------------------------------------------------------ 68691
human.FAM30B-LOC105372935-GGT2.NCBI.                                      tcttggtattctgaaaattaattttaaatgggaaaataaattactaataagtgcagctta 86623
human.FAM230E-LOC105377182-GGT3P.NCBI.                                    ------------------------------------------------------------ 57310
human.FAM230J-LOC105372942-GGTLC5P.NCBI.rev.compl                         ------------------------------------------------------------ 46364
human.FAM230A.ncRNA.18487127.18500594-GGTLC3.18516335.18518165.rev.compl.REF ------------------------------------------------------------ 31039

chimp.LOC112206744-LOC107973052-GGT2.revcompl                             ------------------------------------------------------------ 61762
chimp54546-135457.revcompl.Pan.troglodytes.clone.rp43-41g5.GenBank:AC099533.36 ------------------------------------------------------------ 68691
human.FAM30B-LOC105372935-GGT2.NCBI.                                      atcaacaagagttttcattaacatgtgggaatacaccaggtgttccttggattttcttga 86683
human.FAM230E-LOC105377182-GGT3P.NCBI.                                    ------------------------------------------------------------ 57310
human.FAM230J-LOC105372942-GGTLC5P.NCBI.rev.compl                         ------------------------------------------------------------ 46364
human.FAM230A.ncRNA.18487127.18500594-GGTLC3.18516335.18518165.rev.compl.REF ------------------------------------------------------------ 31039

chimp.LOC112206744-LOC107973052-GGT2.revcompl                             ------------------------------------------------------------ 61762
chimp54546-135457.revcompl.Pan.troglodytes.clone.rp43-41g5.GenBank:AC099533.36 ------------------------------------------------------------ 68691
human.FAM30B-LOC105372935-GGT2.NCBI.                                      aaattctacagggagggtacgggagtgacggtacagataagattggacatgagctgataa 86743
human.FAM230E-LOC105377182-GGT3P.NCBI.                                    ------------------------------------------------------------ 57310
human.FAM230J-LOC105372942-GGTLC5P.NCBI.rev.compl                         ------------------------------------------------------------ 46364
human.FAM230A.ncRNA.18487127.18500594-GGTLC3.18516335.18518165.rev.compl.REF ------------------------------------------------------------ 31039

chimp.LOC112206744-LOC107973052-GGT2.revcompl                             ------------------------------------------------------------ 61762
chimp54546-135457.revcompl.Pan.troglodytes.clone.rp43-41g5.GenBank:AC099533.36 ------------------------------------------------------------ 68691
human.FAM30B-LOC105372935-GGT2.NCBI.                                      ttgttggaactaggtgatgagtgtatgagggttaattattctgctctctccactttggta 86803
human.FAM230E-LOC105377182-GGT3P.NCBI.                                    ------------------------------------------------------------ 57310
human.FAM230J-LOC105372942-GGTLC5P.NCBI.rev.compl                         ------------------------------------------------------------ 46364
human.FAM230A.ncRNA.18487127.18500594-GGTLC3.18516335.18518165.rev.compl.REF ------------------------------------------------------------ 31039

chimp.LOC112206744-LOC107973052-GGT2.revcompl                             ------------------------------------------------------------ 61762
chimp54546-135457.revcompl.Pan.troglodytes.clone.rp43-41g5.GenBank:AC099533.36 ------------------------------------------------------------ 68691
human.FAM30B-LOC105372935-GGT2.NCBI.                                      catatttgaaatttttcacaatagaaattaaaagaagaacggggcgtggtggctcacgcc 86863
human.FAM230E-LOC105377182-GGT3P.NCBI.                                    ------------------------------------------------------------ 57310
human.FAM230J-LOC105372942-GGTLC5P.NCBI.rev.compl                         ------------------------------------------------------------ 46364
human.FAM230A.ncRNA.18487127.18500594-GGTLC3.18516335.18518165.rev.compl.REF ------------------------------------------------------------ 31039

chimp.LOC112206744-LOC107973052-GGT2.revcompl                             ------------------------------------------------------------ 61762
chimp54546-135457.revcompl.Pan.troglodytes.clone.rp43-41g5.GenBank:AC099533.36 ------------------------------------------------------------ 68691
human.FAM30B-LOC105372935-GGT2.NCBI.                                      tgtaattccagcactttggaaggccatggcagaggatcacttgaggtcaggaattaaaga 86923
human.FAM230E-LOC105377182-GGT3P.NCBI.                                    ------------------------------------------------------------ 57310
human.FAM230J-LOC105372942-GGTLC5P.NCBI.rev.compl                         ------------------------------------------------------------ 46364
human.FAM230A.ncRNA.18487127.18500594-GGTLC3.18516335.18518165.rev.compl.REF ------------------------------------------------------------ 31039

chimp.LOC112206744-LOC107973052-GGT2.revcompl                             ------------------------------------------------------------ 61762
chimp54546-135457.revcompl.Pan.troglodytes.clone.rp43-41g5.GenBank:AC099533.36 -----------------------------------AAACAAACAAAAATTAGCCAGACGT 68716
human.FAM30B-LOC105372935-GGT2.NCBI.                                      ccagctggtcaacatggggaaaccccacctctactaaaaatacaaaaattagctgggtat 86983
human.FAM230E-LOC105377182-GGT3P.NCBI.                                    ------------------------------------------------------------ 57310
human.FAM230J-LOC105372942-GGTLC5P.NCBI.rev.compl                         ------------------------------------------------------------ 46364
human.FAM230A.ncRNA.18487127.18500594-GGTLC3.18516335.18518165.rev.compl.REF ------------------------------------------------------------ 31039

chimp.LOC112206744-LOC107973052-GGT2.revcompl                             ------------------------------------------------------------ 61762
chimp54546-135457.revcompl.Pan.troglodytes.clone.rp43-41g5.GenBank:AC099533.36 GATGGCTAACACCTGTAATCCCAGCTACTCAGGAGGCTGAGACGGGAGAATCACCTGAAC 68776
human.FAM30B-LOC105372935-GGT2.NCBI.                                      ggtgg------tgtgtaatcccagctattgaggaggctgaggcacgagaatcgcttggac 87037
human.FAM230E-LOC105377182-GGT3P.NCBI.                                    ------------------------------------------------------------ 57310
human.FAM230J-LOC105372942-GGTLC5P.NCBI.rev.compl                         ------------------------------------------------------------ 46364
human.FAM230A.ncRNA.18487127.18500594-GGTLC3.18516335.18518165.rev.compl.REF ------------------------------------------------------------ 31039

chimp.LOC112206744-LOC107973052-GGT2.revcompl                             ------------------------------------------------------------ 61762
chimp54546-135457.revcompl.Pan.troglodytes.clone.rp43-41g5.GenBank:AC099533.36 CCAGGAGGTAGAAGTTGCAGTGAGCTGAGATCCCACCACCGCACTCCAGCCTGGGCCACA 68836
human.FAM30B-LOC105372935-GGT2.NCBI.                                      ccaggaggcagaggttgcagtgagccaagatcgtgccactgcactctagcctgggcaaca 87097
human.FAM230E-LOC105377182-GGT3P.NCBI.                                    ------------------------------------------------------------ 57310
human.FAM230J-LOC105372942-GGTLC5P.NCBI.rev.compl                         ------------------------------------------------------------ 46364
human.FAM230A.ncRNA.18487127.18500594-GGTLC3.18516335.18518165.rev.compl.REF ------------------------------------------------------------ 31039

chimp.LOC112206744-LOC107973052-GGT2.revcompl                             ------------------------------------------------------------ 61762
chimp54546-135457.revcompl.Pan.troglodytes.clone.rp43-41g5.GenBank:AC099533.36 GAGCAAGACTCGGTCTCAAAAAACAAACAAACAAAAAAGTGAAAGTGGAATATATATA-- 68894
human.FAM30B-LOC105372935-GGT2.NCBI.                                      gggcaagcctctgtctcaaaaaaaaaaaaaaaaaaaaaaaaaagaagaagaagaaagaaa 87157
human.FAM230E-LOC105377182-GGT3P.NCBI.                                    ------------------------------------------------------------ 57310
human.FAM230J-LOC105372942-GGTLC5P.NCBI.rev.compl                         ------------------------------------------------------------ 46364
human.FAM230A.ncRNA.18487127.18500594-GGTLC3.18516335.18518165.rev.compl.REF ------------------------------------------------------------ 31039

chimp.LOC112206744-LOC107973052-GGT2.revcompl                             ------------------------------------------------------------ 61762
chimp54546-135457.revcompl.Pan.troglodytes.clone.rp43-41g5.GenBank:AC099533.36 ------------------------------------------------------------ 68894
human.FAM30B-LOC105372935-GGT2.NCBI.                                      acaaagaagaaagaaagaaagaaaaagaaagaaagaaagaaaattaaaagaaggaaatgc 87217
human.FAM230E-LOC105377182-GGT3P.NCBI.                                    ------------------------------------------------------------ 57310
```

```
human.FAM230J-LOC105372942-GGTLC5P.NCBI.rev.compl                                 ------------------------------------------------------------ 46364
human.FAM230A.ncRNA.18487127.18500594-GGTLC3.18516335.18518165.rev.compl.REF      ------------------------------------------------------------ 31039

chimp.LOC112206744-LOC107973052-GGT2.revcompl                                     ------------------------------------------------------------ 61762
chimp54546-135457.revcompl.Pan.troglodytes.clone.rp43-41g5.GenBank:AC099533.36    ------------------------------------------------------------ 68894
human.FAM30B-LOC105372935-GGT2.NCBI.                                              cataaagcatctggagcaaagaacatccaaagaagagcactgatcactgcagcagacagc 87277
human.FAM230E-LOC105377182-GGT3P.NCBI.                                            ------------------------------------------------------------ 57310
human.FAM230J-LOC105372942-GGTLC5P.NCBI.rev.compl                                 ------------------------------------------------------------ 46364
human.FAM230A.ncRNA.18487127.18500594-GGTLC3.18516335.18518165.rev.compl.REF      ------------------------------------------------------------ 31039

chimp.LOC112206744-LOC107973052-GGT2.revcompl                                     ------------------------------------------------------------ 61762
chimp54546-135457.revcompl.Pan.troglodytes.clone.rp43-41g5.GenBank:AC099533.36    ------------------------------------------------------------ 68894
human.FAM30B-LOC105372935-GGT2.NCBI.                                              cattcagaatgggcagagagcagagctggcaggaagagaggacaaaggaagaccaggaa 87337
human.FAM230E-LOC105377182-GGT3P.NCBI.                                            ------------------------------------------------------------ 57310
human.FAM230J-LOC105372942-GGTLC5P.NCBI.rev.compl                                 ------------------------------------------------------------ 46364
human.FAM230A.ncRNA.18487127.18500594-GGTLC3.18516335.18518165.rev.compl.REF      ------------------------------------------------------------ 31039

chimp.LOC112206744-LOC107973052-GGT2.revcompl                                     ------------------------------------------------------------ 61762
chimp54546-135457.revcompl.Pan.troglodytes.clone.rp43-41g5.GenBank:AC099533.36    ------------------------------------------------------------ 68894
human.FAM30B-LOC105372935-GGT2.NCBI.                                              gggaagccaggaaaagagattttgcattgtgttattatttcaatcagaagacttaggaac 87397
human.FAM230E-LOC105377182-GGT3P.NCBI.                                            ------------------------------------------------------------ 57310
human.FAM230J-LOC105372942-GGTLC5P.NCBI.rev.compl                                 ------------------------------------------------------------ 46364
human.FAM230A.ncRNA.18487127.18500594-GGTLC3.18516335.18518165.rev.compl.REF      ------------------------------------------------------------ 31039

chimp.LOC112206744-LOC107973052-GGT2.revcompl                                     ------------------------------------------------------------ 61762
chimp54546-135457.revcompl.Pan.troglodytes.clone.rp43-41g5.GenBank:AC099533.36    ------------------------------------------------------------ 68894
human.FAM30B-LOC105372935-GGT2.NCBI.                                              attatttaattatcaaatttgtgtgtctgatgatcctttacaacactgcagcatcaaata 87457
human.FAM230E-LOC105377182-GGT3P.NCBI.                                            ------------------------------------------------------------ 57310
human.FAM230J-LOC105372942-GGTLC5P.NCBI.rev.compl                                 ------------------------------------------------------------ 46364
human.FAM230A.ncRNA.18487127.18500594-GGTLC3.18516335.18518165.rev.compl.REF      ------------------------------------------------------------ 31039

chimp.LOC112206744-LOC107973052-GGT2.revcompl                                     ------------------------------------------------------------ 61762
chimp54546-135457.revcompl.Pan.troglodytes.clone.rp43-41g5.GenBank:AC099533.36    ------------------------------------------------------------ 68894
human.FAM30B-LOC105372935-GGT2.NCBI.                                              aagtgttatcagatttctttctttttttttttttttgagactgggtctgattctgtagccc 87517
human.FAM230E-LOC105377182-GGT3P.NCBI.                                            ------------------------------------------------------------ 57310
human.FAM230J-LOC105372942-GGTLC5P.NCBI.rev.compl                                 ------------------------------------------------------------ 46364
human.FAM230A.ncRNA.18487127.18500594-GGTLC3.18516335.18518165.rev.compl.REF      ------------------------------------------------------------ 31039

chimp.LOC112206744-LOC107973052-GGT2.revcompl                                     ------------------------------------------------------------ 61762
chimp54546-135457.revcompl.Pan.troglodytes.clone.rp43-41g5.GenBank:AC099533.36    -------------------------------------------------TGTGTGTGTGT 68905
human.FAM30B-LOC105372935-GGT2.NCBI.                                              aggctggagtgcaggggtataataatagctcactgcagcttcaaactcctgagtttgttt 87577
human.FAM230E-LOC105377182-GGT3P.NCBI.                                            ------------------------------------------------------------ 57310
human.FAM230J-LOC105372942-GGTLC5P.NCBI.rev.compl                                 ------------------------------------------------------------ 46364
human.FAM230A.ncRNA.18487127.18500594-GGTLC3.18516335.18518165.rev.compl.REF      ------------------------------------------------------------ 31039

chimp.LOC112206744-LOC107973052-GGT2.revcompl                                     ------------------------------------------------------------ 61762
chimp54546-135457.revcompl.Pan.troglodytes.clone.rp43-41g5.GenBank:AC099533.36    GTATATGTGTATATGTGTGTGTGTATATG------------------------------- 68934
human.FAM30B-LOC105372935-GGT2.NCBI.                                              gtttgtttgtctgtttgtttgtttgtttggagagacagcgtctcgaactcctgggctcca 87637
human.FAM230E-LOC105377182-GGT3P.NCBI.                                            ------------------------------------------------------------ 57310
human.FAM230J-LOC105372942-GGTLC5P.NCBI.rev.compl                                 ------------------------------------------------------------ 46364
human.FAM230A.ncRNA.18487127.18500594-GGTLC3.18516335.18518165.rev.compl.REF      ------------------------------------------------------------ 31039

chimp.LOC112206744-LOC107973052-GGT2.revcompl                                     ------------------------------------------------------------ 61762
chimp54546-135457.revcompl.Pan.troglodytes.clone.rp43-41g5.GenBank:AC099533.36    ------------------------------------------------------------ 68934
human.FAM30B-LOC105372935-GGT2.NCBI.                                              gtgatcctcccacctcagcctcccaaagtgctgggattacaagcatgagccactgtgccc 87697
human.FAM230E-LOC105377182-GGT3P.NCBI.                                            ------------------------------------------------------------ 57310
human.FAM230J-LOC105372942-GGTLC5P.NCBI.rev.compl                                 ------------------------------------------------------------ 46364
human.FAM230A.ncRNA.18487127.18500594-GGTLC3.18516335.18518165.rev.compl.REF      ------------------------------------------------------------ 31039

chimp.LOC112206744-LOC107973052-GGT2.revcompl                                     ------------------------------------------------------------ 61762
chimp54546-135457.revcompl.Pan.troglodytes.clone.rp43-41g5.GenBank:AC099533.36    ------------------------------------------------------------ 68934
human.FAM30B-LOC105372935-GGT2.NCBI.                                              cagcccaacatctgatttcaaaagtactattgatattataatcaccattaaaaatcactg 87757
human.FAM230E-LOC105377182-GGT3P.NCBI.                                            ------------------------------------------------------------ 57310
human.FAM230J-LOC105372942-GGTLC5P.NCBI.rev.compl                                 ------------------------------------------------------------ 46364
human.FAM230A.ncRNA.18487127.18500594-GGTLC3.18516335.18518165.rev.compl.REF      ------------------------------------------------------------ 31039

chimp.LOC112206744-LOC107973052-GGT2.revcompl                                     ------------------------------------------------------------ 61762
chimp54546-135457.revcompl.Pan.troglodytes.clone.rp43-41g5.GenBank:AC099533.36    ------------------------------------------------------------ 68934
human.FAM30B-LOC105372935-GGT2.NCBI.                                              aagtttttgtcacccaaagttttacgttgaaatatagaacatttcatttttttcccacac 87817
human.FAM230E-LOC105377182-GGT3P.NCBI.                                            ------------------------------------------------------------ 57310
human.FAM230J-LOC105372942-GGTLC5P.NCBI.rev.compl                                 ------------------------------------------------------------ 46364
human.FAM230A.ncRNA.18487127.18500594-GGTLC3.18516335.18518165.rev.compl.REF      ------------------------------------------------------------ 31039

chimp.LOC112206744-LOC107973052-GGT2.revcompl                                     ------------------------------------------------------------ 61762
chimp54546-135457.revcompl.Pan.troglodytes.clone.rp43-41g5.GenBank:AC099533.36    ------------------------------------------------------------ 68934
human.FAM30B-LOC105372935-GGT2.NCBI.                                              acctcacatctgcttttgatttttaacatgtgcatgaattacctcttattagaaaaaaaa 87877
human.FAM230E-LOC105377182-GGT3P.NCBI.                                            ------------------------------------------------------------ 57310
human.FAM230J-LOC105372942-GGTLC5P.NCBI.rev.compl                                 ------------------------------------------------------------ 46364
human.FAM230A.ncRNA.18487127.18500594-GGTLC3.18516335.18518165.rev.compl.REF      ------------------------------------------------------------ 31039

chimp.LOC112206744-LOC107973052-GGT2.revcompl                                     ------------------------------------------------------------ 61762
chimp54546-135457.revcompl.Pan.troglodytes.clone.rp43-41g5.GenBank:AC099533.36    ------------------------------------------------------------ 68934
human.FAM30B-LOC105372935-GGT2.NCBI.                                              acacaaaatttttttggtaaaagaattttccttaagtgggtattttttccctaagtctgaa 87937
human.FAM230E-LOC105377182-GGT3P.NCBI.                                            ------------------------------------------------------------ 57310
human.FAM230J-LOC105372942-GGTLC5P.NCBI.rev.compl                                 ------------------------------------------------------------ 46364
human.FAM230A.ncRNA.18487127.18500594-GGTLC3.18516335.18518165.rev.compl.REF      ------------------------------------------------------------ 31039

chimp.LOC112206744-LOC107973052-GGT2.revcompl                                     ------------------------------------------------------------ 61762
chimp54546-135457.revcompl.Pan.troglodytes.clone.rp43-41g5.GenBank:AC099533.36    ------------------------------------------------------------ 68934
human.FAM30B-LOC105372935-GGT2.NCBI.                                              atgacttagtaaggataagaggtcaaaccatatgctttgtagtgaaactgcaactatgat 87997
human.FAM230E-LOC105377182-GGT3P.NCBI.                                            ------------------------------------------------------------ 57310
human.FAM230J-LOC105372942-GGTLC5P.NCBI.rev.compl                                 ------------------------------------------------------------ 46364
human.FAM230A.ncRNA.18487127.18500594-GGTLC3.18516335.18518165.rev.compl.REF      ------------------------------------------------------------ 31039

chimp.LOC112206744-LOC107973052-GGT2.revcompl                                     ------------------------------------------------------------ 61762
chimp54546-135457.revcompl.Pan.troglodytes.clone.rp43-41g5.GenBank:AC099533.36    ------------------------------------------------------------ 68934
human.FAM30B-LOC105372935-GGT2.NCBI.                                              gaataacattagaattctggatgccttccagcagctggaacagggcagacctgaggcggg 88057
human.FAM230E-LOC105377182-GGT3P.NCBI.                                            ------------------------------------------------------------ 57310
human.FAM230J-LOC105372942-GGTLC5P.NCBI.rev.compl                                 ------------------------------------------------------------ 46364
human.FAM230A.ncRNA.18487127.18500594-GGTLC3.18516335.18518165.rev.compl.REF      ------------------------------------------------------------ 31039

chimp.LOC112206744-LOC107973052-GGT2.revcompl                                     ------------------------------------------------------------ 61762
chimp54546-135457.revcompl.Pan.troglodytes.clone.rp43-41g5.GenBank:AC099533.36    ------------------------------------------------------------ 68934
human.FAM30B-LOC105372935-GGT2.NCBI.                                              agtcctcatgcttaggggaaggggctcaggcactgggagtcggtcagggctcccaggatg 88117
human.FAM230E-LOC105377182-GGT3P.NCBI.                                            ------------------------------------------------------------ 57310
human.FAM230J-LOC105372942-GGTLC5P.NCBI.rev.compl                                 ------------------------------------------------------------ 46364
human.FAM230A.ncRNA.18487127.18500594-GGTLC3.18516335.18518165.rev.compl.REF      ------------------------------------------------------------ 31039

chimp.LOC112206744-LOC107973052-GGT2.revcompl                                     ------------------------------------------------------------ 61762
chimp54546-135457.revcompl.Pan.troglodytes.clone.rp43-41g5.GenBank:AC099533.36    ------------------------------------------------------------ 68934
human.FAM30B-LOC105372935-GGT2.NCBI.                                              taaaagatacacatgaaagaaacaaacagtgggactcgggccagtgcaaaatttggcctg 88177
human.FAM230E-LOC105377182-GGT3P.NCBI.                                            ------------------------------------------------------------ 57310
human.FAM230J-LOC105372942-GGTLC5P.NCBI.rev.compl                                 ------------------------------------------------------------ 46364
human.FAM230A.ncRNA.18487127.18500594-GGTLC3.18516335.18518165.rev.compl.REF      ------------------------------------------------------------ 31039

chimp.LOC112206744-LOC107973052-GGT2.revcompl                                     ------------------------------------------------------------ 61762
chimp54546-135457.revcompl.Pan.troglodytes.clone.rp43-41g5.GenBank:AC099533.36    ------------------------------------------------------------ 68934
human.FAM30B-LOC105372935-GGT2.NCBI.                                              gaactgagctaaaaactgtcatactggtgggaagaagcagattggttcaacagggtaacc 88237
human.FAM230E-LOC105377182-GGT3P.NCBI.                                            ------------------------------------------------------------ 57310
human.FAM230J-LOC105372942-GGTLC5P.NCBI.rev.compl                                 ------------------------------------------------------------ 46364
human.FAM230A.ncRNA.18487127.18500594-GGTLC3.18516335.18518165.rev.compl.REF      ------------------------------------------------------------ 31039

chimp.LOC112206744-LOC107973052-GGT2.revcompl                                     ------------------------------------------------------------ 61762
chimp54546-135457.revcompl.Pan.troglodytes.clone.rp43-41g5.GenBank:AC099533.36    ------------------------------------------------------------ 68934
human.FAM30B-LOC105372935-GGT2.NCBI.                                              aaatgtgaaaaagaaaagcctcacaagaattaaaattgatggttggaactgggcgctccg 88297
human.FAM230E-LOC105377182-GGT3P.NCBI.                                            ------------------------------------------------------------ 57310
human.FAM230J-LOC105372942-GGTLC5P.NCBI.rev.compl                                 ------------------------------------------------------------ 46364
```

```
human.FAM230A.ncRNA.18487127.18500594-GGTLC3.18516335.18518165.rev.compl.REF         ------------------------------------------------------------ 31039

chimp.LOC112206744-LOC107973052-GGT2.revcompl                                          ------------------------------------------------------------ 61762
chimp54546-135457.revcompl.Pan.troglodytes.clone.rp43-41g5.GenBank:AC099533.36         ------------------------------------------------------------ 68934
human.FAM30B-LOC105372935-GGT2.NCBI.                                                   aggctgtgggccggtgtcctgccccgaagggtttggctgccagcaggtgcagcgctgccc 88357
human.FAM230E-LOC105377182-GGT3P.NCBI.                                                 ------------------------------------------------------------ 57310
human.FAM230J-LOC105372942-GGTLC5P.NCBI.rev.compl                                      ------------------------------------------------------------ 46364
human.FAM230A.ncRNA.18487127.18500594-GGTLC3.18516335.18518165.rev.compl.REF         ------------------------------------------------------------ 31039

chimp.LOC112206744-LOC107973052-GGT2.revcompl                                          ------------------------------------------------------------ 61762
chimp54546-135457.revcompl.Pan.troglodytes.clone.rp43-41g5.GenBank:AC099533.36         ------------------------------------------------------------ 68934
human.FAM30B-LOC105372935-GGT2.NCBI.                                                   tgggataccagctcctgggatgcgctgtagtatcatcaatcaccggtgctgctcgctggg 88417
human.FAM230E-LOC105377182-GGT3P.NCBI.                                                 ------------------------------------------------------------ 57310
human.FAM230J-LOC105372942-GGTLC5P.NCBI.rev.compl                                      ------------------------------------------------------------ 46364
human.FAM230A.ncRNA.18487127.18500594-GGTLC3.18516335.18518165.rev.compl.REF         ------------------------------------------------------------ 31039

chimp.LOC112206744-LOC107973052-GGT2.revcompl                                          ------------------------------------------------------------ 61762
chimp54546-135457.revcompl.Pan.troglodytes.clone.rp43-41g5.GenBank:AC099533.36         ------------------------------------------------------------ 68934
human.FAM30B-LOC105372935-GGT2.NCBI.                                                   ccttggaagcacacagtagtagcagatgtcagcagggggccaccagggccacctgccagt 88477
human.FAM230E-LOC105377182-GGT3P.NCBI.                                                 ------------------------------------------------------------ 57310
human.FAM230J-LOC105372942-GGTLC5P.NCBI.rev.compl                                      ------------------------------------------------------------ 46364
human.FAM230A.ncRNA.18487127.18500594-GGTLC3.18516335.18518165.rev.compl.REF         ------------------------------------------------------------ 31039

chimp.LOC112206744-LOC107973052-GGT2.revcompl                                          ------------------------------------------------------------ 61762
chimp54546-135457.revcompl.Pan.troglodytes.clone.rp43-41g5.GenBank:AC099533.36         ------------------------------------------------------------ 68934
human.FAM30B-LOC105372935-GGT2.NCBI.                                                   gagcccactgcttagctgagtcatggtcaagtttctctgcagattgttgcctgaaaatat 88537
human.FAM230E-LOC105377182-GGT3P.NCBI.                                                 ------------------------------------------------------------ 57310
human.FAM230J-LOC105372942-GGTLC5P.NCBI.rev.compl                                      ------------------------------------------------------------ 46364
human.FAM230A.ncRNA.18487127.18500594-GGTLC3.18516335.18518165.rev.compl.REF         ------------------------------------------------------------ 31039

chimp.LOC112206744-LOC107973052-GGT2.revcompl                                          ------------------------------------------------------------ 61762
chimp54546-135457.revcompl.Pan.troglodytes.clone.rp43-41g5.GenBank:AC099533.36         ------------------------------------------------------------ 68934
human.FAM30B-LOC105372935-GGT2.NCBI.                                                   ttcaccacctggaacctccctgtcccacccatcttcacttgcaatactcactcccctcac 88597
human.FAM230E-LOC105377182-GGT3P.NCBI.                                                 ------------------------------------------------------------ 57310
human.FAM230J-LOC105372942-GGTLC5P.NCBI.rev.compl                                      ------------------------------------------------------------ 46364
human.FAM230A.ncRNA.18487127.18500594-GGTLC3.18516335.18518165.rev.compl.REF         ------------------------------------------------------------ 31039

chimp.LOC112206744-LOC107973052-GGT2.revcompl                                          ------------------------------------------------------------ 61762
chimp54546-135457.revcompl.Pan.troglodytes.clone.rp43-41g5.GenBank:AC099533.36         ------------------------------------------------------------ 68934
human.FAM30B-LOC105372935-GGT2.NCBI.                                                   ctctgcccctaaagcaattattcattaaccacagggatccaccatccaggaacaaaagag 88657
human.FAM230E-LOC105377182-GGT3P.NCBI.                                                 ------------------------------------------------------------ 57310
human.FAM230J-LOC105372942-GGTLC5P.NCBI.rev.compl                                      ------------------------------------------------------------ 46364
human.FAM230A.ncRNA.18487127.18500594-GGTLC3.18516335.18518165.rev.compl.REF         ------------------------------------------------------------ 31039

chimp.LOC112206744-LOC107973052-GGT2.revcompl                                          ------------------------------------------------------------ 61762
chimp54546-135457.revcompl.Pan.troglodytes.clone.rp43-41g5.GenBank:AC099533.36         ------------------------------------------------------------ 68934
human.FAM30B-LOC105372935-GGT2.NCBI.                                                   gcttgaaagctctatctgtgcctgcacttcttggtccaaaaggcgcaggggggatgggga 88717
human.FAM230E-LOC105377182-GGT3P.NCBI.                                                 ------------------------------------------------------------ 57310
human.FAM230J-LOC105372942-GGTLC5P.NCBI.rev.compl                                      ------------------------------------------------------------ 46364
human.FAM230A.ncRNA.18487127.18500594-GGTLC3.18516335.18518165.rev.compl.REF         ------------------------------------------------------------ 31039

chimp.LOC112206744-LOC107973052-GGT2.revcompl                                          ------------------------------------------------------------ 61762
chimp54546-135457.revcompl.Pan.troglodytes.clone.rp43-41g5.GenBank:AC099533.36         ------------------------------------------------------------ 68934
human.FAM30B-LOC105372935-GGT2.NCBI.                                                   gaggcaccggggtcatggtgcaggctcggctgttgagcctcattccccaaacacaatgtg 88777
human.FAM230E-LOC105377182-GGT3P.NCBI.                                                 ------------------------------------------------------------ 57310
human.FAM230J-LOC105372942-GGTLC5P.NCBI.rev.compl                                      ------------------------------------------------------------ 46364
human.FAM230A.ncRNA.18487127.18500594-GGTLC3.18516335.18518165.rev.compl.REF         ------------------------------------------------------------ 31039

chimp.LOC112206744-LOC107973052-GGT2.revcompl                                          ------------------------------------------------------------ 61762
chimp54546-135457.revcompl.Pan.troglodytes.clone.rp43-41g5.GenBank:AC099533.36         ------------------------------------------------------------ 68934
human.FAM30B-LOC105372935-GGT2.NCBI.                                                   acaggtatcaatagttctggagttgatcatttcacagaaaggaaactgctgtcaacctgc 88837
human.FAM230E-LOC105377182-GGT3P.NCBI.                                                 ------------------------------------------------------------ 57310
human.FAM230J-LOC105372942-GGTLC5P.NCBI.rev.compl                                      ------------------------------------------------------------ 46364
human.FAM230A.ncRNA.18487127.18500594-GGTLC3.18516335.18518165.rev.compl.REF         ------------------------------------------------------------ 31039

chimp.LOC112206744-LOC107973052-GGT2.revcompl                                          ------------------------------------------------------------ 61762
chimp54546-135457.revcompl.Pan.troglodytes.clone.rp43-41g5.GenBank:AC099533.36         ------------------------------------------------------------ 68934
human.FAM30B-LOC105372935-GGT2.NCBI.                                                   ccaaggccacacagctagtcctggataaattgacttaaagaacccaggtctctgattccc 88897
human.FAM230E-LOC105377182-GGT3P.NCBI.                                                 ------------------------------------------------------------ 57310
human.FAM230J-LOC105372942-GGTLC5P.NCBI.rev.compl                                      ------------------------------------------------------------ 46364
human.FAM230A.ncRNA.18487127.18500594-GGTLC3.18516335.18518165.rev.compl.REF         ------------------------------------------------------------ 31039

chimp.LOC112206744-LOC107973052-GGT2.revcompl                                          ------------------------------------------------------------ 61762
chimp54546-135457.revcompl.Pan.troglodytes.clone.rp43-41g5.GenBank:AC099533.36         ------------------------------------------------------------ 68934
human.FAM30B-LOC105372935-GGT2.NCBI.                                                   aaatcagtgctcttacccctcccacgccacctctctcttctaaagatccacccaatggaa 88957
human.FAM230E-LOC105377182-GGT3P.NCBI.                                                 ------------------------------------------------------------ 57310
human.FAM230J-LOC105372942-GGTLC5P.NCBI.rev.compl                                      ------------------------------------------------------------ 46364
human.FAM230A.ncRNA.18487127.18500594-GGTLC3.18516335.18518165.rev.compl.REF         ------------------------------------------------------------ 31039

chimp.LOC112206744-LOC107973052-GGT2.revcompl                                          ------------------------------------------------------------ 61762
chimp54546-135457.revcompl.Pan.troglodytes.clone.rp43-41g5.GenBank:AC099533.36         ------------------------------------------------------------ 68934
human.FAM30B-LOC105372935-GGT2.NCBI.                                                   ctgccccagggctggtatcccagctacttctccattcatgccgcaggtcaatgattcagg 89017
human.FAM230E-LOC105377182-GGT3P.NCBI.                                                 ------------------------------------------------------------ 57310
human.FAM230J-LOC105372942-GGTLC5P.NCBI.rev.compl                                      ------------------------------------------------------------ 46364
human.FAM230A.ncRNA.18487127.18500594-GGTLC3.18516335.18518165.rev.compl.REF         ------------------------------------------------------------ 31039

chimp.LOC112206744-LOC107973052-GGT2.revcompl                                          ------------------------------------------------------------ 61762
chimp54546-135457.revcompl.Pan.troglodytes.clone.rp43-41g5.GenBank:AC099533.36         ------------------------------------------------------------ 68934
human.FAM30B-LOC105372935-GGT2.NCBI.                                                   gcggacactcctcaagtgaccttcagaggccctgactcactcctccgacagtcctttgtg 89077
human.FAM230E-LOC105377182-GGT3P.NCBI.                                                 ------------------------------------------------------------ 57310
human.FAM230J-LOC105372942-GGTLC5P.NCBI.rev.compl                                      ------------------------------------------------------------ 46364
human.FAM230A.ncRNA.18487127.18500594-GGTLC3.18516335.18518165.rev.compl.REF         ------------------------------------------------------------ 31039

chimp.LOC112206744-LOC107973052-GGT2.revcompl                                          ------------------------------------------------------------ 61762
chimp54546-135457.revcompl.Pan.troglodytes.clone.rp43-41g5.GenBank:AC099533.36         ------------------------------------------------------------ 68934
human.FAM30B-LOC105372935-GGT2.NCBI.                                                   tctggactcccagccgctcctcagcagctctgtcctcttttgtacatttattactgtcag 89137
human.FAM230E-LOC105377182-GGT3P.NCBI.                                                 ------------------------------------------------------------ 57310
human.FAM230J-LOC105372942-GGTLC5P.NCBI.rev.compl                                      ------------------------------------------------------------ 46364
human.FAM230A.ncRNA.18487127.18500594-GGTLC3.18516335.18518165.rev.compl.REF         ------------------------------------------------------------ 31039

chimp.LOC112206744-LOC107973052-GGT2.revcompl                                          ------------------------------------------------------------ 61762
chimp54546-135457.revcompl.Pan.troglodytes.clone.rp43-41g5.GenBank:AC099533.36         ------------------------------------------------------------ 68934
human.FAM30B-LOC105372935-GGT2.NCBI.                                                   cctgaggctagcaggaggtctcaatgcctggtgatctgaggccagcctgtcccaggccct 89197
human.FAM230E-LOC105377182-GGT3P.NCBI.                                                 ------------------------------------------------------------ 57310
human.FAM230J-LOC105372942-GGTLC5P.NCBI.rev.compl                                      ------------------------------------------------------------ 46364
human.FAM230A.ncRNA.18487127.18500594-GGTLC3.18516335.18518165.rev.compl.REF         ------------------------------------------------------------ 31039

chimp.LOC112206744-LOC107973052-GGT2.revcompl                                          ------------------------------------------------------------ 61762
chimp54546-135457.revcompl.Pan.troglodytes.clone.rp43-41g5.GenBank:AC099533.36         ------------------------------------------------------------ 68934
human.FAM30B-LOC105372935-GGT2.NCBI.                                                   cccagtcccgtgaggtcctcctggggatgcaggcacagcgactgacaaggataaggagca 89257
human.FAM230E-LOC105377182-GGT3P.NCBI.                                                 ------------------------------------------------------------ 57310
human.FAM230J-LOC105372942-GGTLC5P.NCBI.rev.compl                                      ------------------------------------------------------------ 46364
human.FAM230A.ncRNA.18487127.18500594-GGTLC3.18516335.18518165.rev.compl.REF         ------------------------------------------------------------ 31039

chimp.LOC112206744-LOC107973052-GGT2.revcompl                                          ------------------------------------------------------------ 61762
chimp54546-135457.revcompl.Pan.troglodytes.clone.rp43-41g5.GenBank:AC099533.36         ------------------------------------------------------------ 68934
human.FAM30B-LOC105372935-GGT2.NCBI.                                                   gaagggcaggtggctttctgggtgtgagcttggccttctgccacacccatgttcagaaag 89317
human.FAM230E-LOC105377182-GGT3P.NCBI.                                                 ------------------------------------------------------------ 57310
human.FAM230J-LOC105372942-GGTLC5P.NCBI.rev.compl                                      ------------------------------------------------------------ 46364
human.FAM230A.ncRNA.18487127.18500594-GGTLC3.18516335.18518165.rev.compl.REF         ------------------------------------------------------------ 31039

chimp.LOC112206744-LOC107973052-GGT2.revcompl                                          ------------------------------------------------------------ 61762
chimp54546-135457.revcompl.Pan.troglodytes.clone.rp43-41g5.GenBank:AC099533.36         ------------------------------------------------------------ 68934
human.FAM30B-LOC105372935-GGT2.NCBI.                                                   gtgtctacgggcacatcttgtgttgtcacaggtgtagaaagggctttgacacgcggcaga 89377
human.FAM230E-LOC105377182-GGT3P.NCBI.                                                 ------------------------------------------------------------ 57310
human.FAM230J-LOC105372942-GGTLC5P.NCBI.rev.compl                                      ------------------------------------------------------------ 46364
human.FAM230A.ncRNA.18487127.18500594-GGTLC3.18516335.18518165.rev.compl.REF         ------------------------------------------------------------ 31039
```

```
chimp.LOC112206744-LOC107973052-GGT2.revcompl                                      ------------------------------------------------------------ 61762
chimp54546-135457.revcompl.Pan.troglodytes.clone.rp43-41g5.GenBank:AC099533.36     ------------------------------------------------------------ 68934
human.FAM30B-LOC105372935-GGT2.NCBI.                                               tcctccaatccctgtgaaagagggatcattctttctattatacagtgaggacgctgaagt 89437
human.FAM230E-LOC105377182-GGT3P.NCBI.                                             ------------------------------------------------------------ 57310
human.FAM230J-LOC105372942-GGTLC5P.NCBI.rev.compl                                  ------------------------------------------------------------ 46364
human.FAM230A.ncRNA.18487127.18500594-GGTLC3.18516335.18518165.rev.compl.REF       ------------------------------------------------------------ 31039

chimp.LOC112206744-LOC107973052-GGT2.revcompl                                      ------------------------------------------------------------ 61762
chimp54546-135457.revcompl.Pan.troglodytes.clone.rp43-41g5.GenBank:AC099533.36     ------------------------------------------------------------ 68934
human.FAM30B-LOC105372935-GGT2.NCBI.                                               ccagaaaggtcaggtgactgatgaaaggtcataaaattagcaggaatcagggaaagggca 89497
human.FAM230E-LOC105377182-GGT3P.NCBI.                                             ------------------------------------------------------------ 57310
human.FAM230J-LOC105372942-GGTLC5P.NCBI.rev.compl                                  ------------------------------------------------------------ 46364
human.FAM230A.ncRNA.18487127.18500594-GGTLC3.18516335.18518165.rev.compl.REF       ------------------------------------------------------------ 31039

chimp.LOC112206744-LOC107973052-GGT2.revcompl                                      ------------------------------------------------------------ 61762
chimp54546-135457.revcompl.Pan.troglodytes.clone.rp43-41g5.GenBank:AC099533.36     ------------------------------------------------------------ 68934
human.FAM30B-LOC105372935-GGT2.NCBI.                                               ggaaagcagagcttccagttgcaaggtcaggactcttcagcccagcaggcgagtttgcca 89557
human.FAM230E-LOC105377182-GGT3P.NCBI.                                             ------------------------------------------------------------ 57310
human.FAM230J-LOC105372942-GGTLC5P.NCBI.rev.compl                                  ------------------------------------------------------------ 46364
human.FAM230A.ncRNA.18487127.18500594-GGTLC3.18516335.18518165.rev.compl.REF       ------------------------------------------------------------ 31039

chimp.LOC112206744-LOC107973052-GGT2.revcompl                                      ------------------------------------------------------------ 61762
chimp54546-135457.revcompl.Pan.troglodytes.clone.rp43-41g5.GenBank:AC099533.36     ------------------------------------------------------------ 68934
human.FAM30B-LOC105372935-GGT2.NCBI.                                               gccagtctgtgccctggtctctgccctccccaaacacgtgcaccaatttggaggggctgc 89617
human.FAM230E-LOC105377182-GGT3P.NCBI.                                             ------------------------------------------------------------ 57310
human.FAM230J-LOC105372942-GGTLC5P.NCBI.rev.compl                                  ------------------------------------------------------------ 46364
human.FAM230A.ncRNA.18487127.18500594-GGTLC3.18516335.18518165.rev.compl.REF       ------------------------------------------------------------ 31039

chimp.LOC112206744-LOC107973052-GGT2.revcompl                                      ------------------------------------------------------------ 61762
chimp54546-135457.revcompl.Pan.troglodytes.clone.rp43-41g5.GenBank:AC099533.36     ------------------------------------------------------------ 68934
human.FAM30B-LOC105372935-GGT2.NCBI.                                               tcccagcacgcatgctgcaagataaacttcagtgctcagccctctcactgtgacaacaat 89677
human.FAM230E-LOC105377182-GGT3P.NCBI.                                             ------------------------------------------------------------ 57310
human.FAM230J-LOC105372942-GGTLC5P.NCBI.rev.compl                                  ------------------------------------------------------------ 46364
human.FAM230A.ncRNA.18487127.18500594-GGTLC3.18516335.18518165.rev.compl.REF       ------------------------------------------------------------ 31039

chimp.LOC112206744-LOC107973052-GGT2.revcompl                                      ------------------------------------------------------------ 61762
chimp54546-135457.revcompl.Pan.troglodytes.clone.rp43-41g5.GenBank:AC099533.36     ------------------------------------------------------------ 68934
human.FAM30B-LOC105372935-GGT2.NCBI.                                               cccttagaaaggggtccacgctgtccatctgggcccctacaccaaagaacacctaattat 89737
human.FAM230E-LOC105377182-GGT3P.NCBI.                                             ------------------------------------------------------------ 57310
human.FAM230J-LOC105372942-GGTLC5P.NCBI.rev.compl                                  ------------------------------------------------------------ 46364
human.FAM230A.ncRNA.18487127.18500594-GGTLC3.18516335.18518165.rev.compl.REF       ------------------------------------------------------------ 31039

chimp.LOC112206744-LOC107973052-GGT2.revcompl                                      ------------------------------------------------------------ 61762
chimp54546-135457.revcompl.Pan.troglodytes.clone.rp43-41g5.GenBank:AC099533.36     ------------------------------------------------------------ 68934
human.FAM30B-LOC105372935-GGT2.NCBI.                                               cagtggtcctcactcaggaaacatgcccacaaataaagacaaaagtaaacccaactacac 89797
human.FAM230E-LOC105377182-GGT3P.NCBI.                                             ------------------------------------------------------------ 57310
human.FAM230J-LOC105372942-GGTLC5P.NCBI.rev.compl                                  ------------------------------------------------------------ 46364
human.FAM230A.ncRNA.18487127.18500594-GGTLC3.18516335.18518165.rev.compl.REF       ------------------------------------------------------------ 31039

chimp.LOC112206744-LOC107973052-GGT2.revcompl                                      ------------------------------------------------------------ 61762
chimp54546-135457.revcompl.Pan.troglodytes.clone.rp43-41g5.GenBank:AC099533.36     ------------------------------------------------------------ 68934
human.FAM30B-LOC105372935-GGT2.NCBI.                                               attcctcaaatcctcctctccatttcacagaaacaccacaacagagcagaactgcaacc 89857
human.FAM230E-LOC105377182-GGT3P.NCBI.                                             ------------------------------------------------------------ 57310
human.FAM230J-LOC105372942-GGTLC5P.NCBI.rev.compl                                  ------------------------------------------------------------ 46364
human.FAM230A.ncRNA.18487127.18500594-GGTLC3.18516335.18518165.rev.compl.REF       ------------------------------------------------------------ 31039

chimp.LOC112206744-LOC107973052-GGT2.revcompl                                      ------------------------------------------------------------ 61762
chimp54546-135457.revcompl.Pan.troglodytes.clone.rp43-41g5.GenBank:AC099533.36     ------------------------------------------------------------ 68934
human.FAM30B-LOC105372935-GGT2.NCBI.                                               tgatagcatccctgtctggttttggctctttcaaaacatcagggacctcacaggaaagca 89917
human.FAM230E-LOC105377182-GGT3P.NCBI.                                             ------------------------------------------------------------ 57310
human.FAM230J-LOC105372942-GGTLC5P.NCBI.rev.compl                                  ------------------------------------------------------------ 46364
human.FAM230A.ncRNA.18487127.18500594-GGTLC3.18516335.18518165.rev.compl.REF       ------------------------------------------------------------ 31039

chimp.LOC112206744-LOC107973052-GGT2.revcompl                                      ------------------------------------------------------------ 61762
chimp54546-135457.revcompl.Pan.troglodytes.clone.rp43-41g5.GenBank:AC099533.36     ------------------------------------------------------------ 68934
human.FAM30B-LOC105372935-GGT2.NCBI.                                               atgcagaaaactgcagtcatttggccgcttgttaaggagagccctgagcagcaagctcca 89977
human.FAM230E-LOC105377182-GGT3P.NCBI.                                             ------------------------------------------------------------ 57310
human.FAM230J-LOC105372942-GGTLC5P.NCBI.rev.compl                                  ------------------------------------------------------------ 46364
human.FAM230A.ncRNA.18487127.18500594-GGTLC3.18516335.18518165.rev.compl.REF       ------------------------------------------------------------ 31039

chimp.LOC112206744-LOC107973052-GGT2.revcompl                                      ------------------------------------------------------------ 61762
chimp54546-135457.revcompl.Pan.troglodytes.clone.rp43-41g5.GenBank:AC099533.36     ------------------------------------------------------------ 68934
human.FAM30B-LOC105372935-GGT2.NCBI.                                               gaccggagatgcgcatataacagcagcagttccacacagccctgcacagcagcaattcag 90037
human.FAM230E-LOC105377182-GGT3P.NCBI.                                             ------------------------------------------------------------ 57310
human.FAM230J-LOC105372942-GGTLC5P.NCBI.rev.compl                                  ------------------------------------------------------------ 46364
human.FAM230A.ncRNA.18487127.18500594-GGTLC3.18516335.18518165.rev.compl.REF       ------------------------------------------------------------ 31039

chimp.LOC112206744-LOC107973052-GGT2.revcompl                                      ------------------------------------------------------------ 61762
chimp54546-135457.revcompl.Pan.troglodytes.clone.rp43-41g5.GenBank:AC099533.36     ------------------------------------------------------------ 68934
human.FAM30B-LOC105372935-GGT2.NCBI.                                               cctctggcagctgtgatgatgctgaggcagtgtccccactgccccagtggcttctgcatg 90097
human.FAM230E-LOC105377182-GGT3P.NCBI.                                             ------------------------------------------------------------ 57310
human.FAM230J-LOC105372942-GGTLC5P.NCBI.rev.compl                                  ------------------------------------------------------------ 46364
human.FAM230A.ncRNA.18487127.18500594-GGTLC3.18516335.18518165.rev.compl.REF       ------------------------------------------------------------ 31039

chimp.LOC112206744-LOC107973052-GGT2.revcompl                                      ------------------------------------------------------------ 61762
chimp54546-135457.revcompl.Pan.troglodytes.clone.rp43-41g5.GenBank:AC099533.36     ------------------------------------------------------------ 68934
human.FAM30B-LOC105372935-GGT2.NCBI.                                               caggttcatttgggacaggacacctctaactccatctctggccaggagccagcatagtag 90157
human.FAM230E-LOC105377182-GGT3P.NCBI.                                             ------------------------------------------------------------ 57310
human.FAM230J-LOC105372942-GGTLC5P.NCBI.rev.compl                                  ------------------------------------------------------------ 46364
human.FAM230A.ncRNA.18487127.18500594-GGTLC3.18516335.18518165.rev.compl.REF       ------------------------------------------------------------ 31039

chimp.LOC112206744-LOC107973052-GGT2.revcompl                                      ------------------------------------------------------------ 61762
chimp54546-135457.revcompl.Pan.troglodytes.clone.rp43-41g5.GenBank:AC099533.36     ------------------------------------------------------------ 68934
human.FAM30B-LOC105372935-GGT2.NCBI.                                               gtgccgtgaaagccaggaagtgaactgcacagaatggactgttccagtgtccatgggctg 90217
human.FAM230E-LOC105377182-GGT3P.NCBI.                                             ------------------------------------------------------------ 57310
human.FAM230J-LOC105372942-GGTLC5P.NCBI.rev.compl                                  ------------------------------------------------------------ 46364
human.FAM230A.ncRNA.18487127.18500594-GGTLC3.18516335.18518165.rev.compl.REF       ------------------------------------------------------------ 31039

chimp.LOC112206744-LOC107973052-GGT2.revcompl                                      ------------------------------------------------------------ 61762
chimp54546-135457.revcompl.Pan.troglodytes.clone.rp43-41g5.GenBank:AC099533.36     ------------------------------------------------------------ 68934
human.FAM30B-LOC105372935-GGT2.NCBI.                                               caagtgggtcccacgtcacccacaggtgaatcttaattatgaaccaaggtgacggcagaa 90277
human.FAM230E-LOC105377182-GGT3P.NCBI.                                             ------------------------------------------------------------ 57310
human.FAM230J-LOC105372942-GGTLC5P.NCBI.rev.compl                                  ------------------------------------------------------------ 46364
human.FAM230A.ncRNA.18487127.18500594-GGTLC3.18516335.18518165.rev.compl.REF       ------------------------------------------------------------ 31039

chimp.LOC112206744-LOC107973052-GGT2.revcompl                                      ------------------------------------------------------------ 61762
chimp54546-135457.revcompl.Pan.troglodytes.clone.rp43-41g5.GenBank:AC099533.36     ------------------------------------------------------------ 68934
human.FAM30B-LOC105372935-GGT2.NCBI.                                               aggatgggcagcaaaggagggtgtgtgtggtgatcagccatcagaggagacggccctgtg 90337
human.FAM230E-LOC105377182-GGT3P.NCBI.                                             ------------------------------------------------------------ 57310
human.FAM230J-LOC105372942-GGTLC5P.NCBI.rev.compl                                  ------------------------------------------------------------ 46364
human.FAM230A.ncRNA.18487127.18500594-GGTLC3.18516335.18518165.rev.compl.REF       ------------------------------------------------------------ 31039

chimp.LOC112206744-LOC107973052-GGT2.revcompl                                      ------------------------------------------------------------ 61762
chimp54546-135457.revcompl.Pan.troglodytes.clone.rp43-41g5.GenBank:AC099533.36     ------------------------------------------------------------ 68934
human.FAM30B-LOC105372935-GGT2.NCBI.                                               tgatgaaaggacctcctcaggaaacctccccaccagcaactggctccaaatggtcagact 90397
human.FAM230E-LOC105377182-GGT3P.NCBI.                                             ------------------------------------------------------------ 57310
human.FAM230J-LOC105372942-GGTLC5P.NCBI.rev.compl                                  ------------------------------------------------------------ 46364
human.FAM230A.ncRNA.18487127.18500594-GGTLC3.18516335.18518165.rev.compl.REF       ------------------------------------------------------------ 31039

chimp.LOC112206744-LOC107973052-GGT2.revcompl                                      ------------------------------------------------------------ 61762
chimp54546-135457.revcompl.Pan.troglodytes.clone.rp43-41g5.GenBank:AC099533.36     ------------------------------------------------------------ 68934
human.FAM30B-LOC105372935-GGT2.NCBI.                                               ttccaagaaattcctgtgaaaaggacaggctcagggcatgtgaatgtgatagtcagcaat 90457
human.FAM230E-LOC105377182-GGT3P.NCBI.                                             ------------------------------------------------------------ 57310
human.FAM230J-LOC105372942-GGTLC5P.NCBI.rev.compl                                  ------------------------------------------------------------ 46364
human.FAM230A.ncRNA.18487127.18500594-GGTLC3.18516335.18518165.rev.compl.REF       ------------------------------------------------------------ 31039
```

```
chimp.LOC112206744-LOC107973052-GGT2.revcompl                                ------------------------------------------------------------ 61762
chimp54546-135457.revcompl.Pan.troglodytes.clone.rp43-41g5.GenBank:AC099533.36  ------------------------------------------------------------ 68934
human.FAM30B-LOC105372935-GGT2.NCBI.                                         tcgcacctcagcagccagtgacactatggtatagggaaaagaacttagattaggcatcaa 90517
human.FAM230E-LOC105377182-GGT3P.NCBI.                                       ------------------------------------------------------------ 57310
human.FAM230J-LOC105372942-GGTLC5P.NCBI.rev.compl                            ------------------------------------------------------------ 46364
human.FAM230A.ncRNA.18487127.18500594-GGTLC3.18516335.18518165.rev.compl.REF ------------------------------------------------------------ 31039

chimp.LOC112206744-LOC107973052-GGT2.revcompl                                ------------------------------------------------------------ 61762
chimp54546-135457.revcompl.Pan.troglodytes.clone.rp43-41g5.GenBank:AC099533.36  ---------------------------------------------------------TGT 68937
human.FAM30B-LOC105372935-GGT2.NCBI.                                         aagtacccggtccccttcatacatggctggtggggaggtaggatggtgcagccactttgg 90577
human.FAM230E-LOC105377182-GGT3P.NCBI.                                       ------------------------------------------------------------ 57310
human.FAM230J-LOC105372942-GGTLC5P.NCBI.rev.compl                            ------------------------------------------------------------ 46364
human.FAM230A.ncRNA.18487127.18500594-GGTLC3.18516335.18518165.rev.compl.REF ------------------------------------------------------------ 31039

chimp.LOC112206744-LOC107973052-GGT2.revcompl                                ------------------------------------------------------------ 61762
chimp54546-135457.revcompl.Pan.troglodytes.clone.rp43-41g5.GenBank:AC099533.36  GTATATATATGGAATATATATGTATATTTGCCCATGACATAGCCCTCAGG-----AGAT 68992
human.FAM30B-LOC105372935-GGT2.NCBI.                                         aaaacagtctggtagtttctttcaaagttaccaaatgacccagcaatcccactcctaagt 90637
human.FAM230E-LOC105377182-GGT3P.NCBI.                                       ------------------------------------------------------------ 57310
human.FAM230J-LOC105372942-GGTLC5P.NCBI.rev.compl                            ------------------------------------------------------------ 46364
human.FAM230A.ncRNA.18487127.18500594-GGTLC3.18516335.18518165.rev.compl.REF ------------------------------------------------------------ 31039

chimp.LOC112206744-LOC107973052-GGT2.revcompl                                ------------------------------------------------------------ 61762
chimp54546-135457.revcompl.Pan.troglodytes.clone.rp43-41g5.GenBank:AC099533.36  CCTTGACCATGTGCTGAAGGATATTTTTTCCCAAAGATATTTTTATAAATAAAAATAAAG 69052
human.FAM30B-LOC105372935-GGT2.NCBI.                                         atttacccaagagaaataaaacctgtgtccacacaaagacttgtacacatcctaatagct 90697
human.FAM230E-LOC105377182-GGT3P.NCBI.                                       ------------------------------------------------------------ 57310
human.FAM230J-LOC105372942-GGTLC5P.NCBI.rev.compl                            ------------------------------------------------------------ 46364
human.FAM230A.ncRNA.18487127.18500594-GGTLC3.18516335.18518165.rev.compl.REF ------------------------------------------------------------ 31039

chimp.LOC112206744-LOC107973052-GGT2.revcompl                                ------------------------------------------------------------ 61762
chimp54546-135457.revcompl.Pan.troglodytes.clone.rp43-41g5.GenBank:AC099533.36  GGTAACAGAAGAGGTAGATGTGTGGGTAAATCTAAATGAATGTTATTGGTATAAAA---- 69108
human.FAM30B-LOC105372935-GGT2.NCBI.                                         gtattcatagtagccccaaactggaaacaacccaaatgcccatcagtttgtgaagagatg 90757
human.FAM230E-LOC105377182-GGT3P.NCBI.                                       ------------------------------------------------------------ 57310
human.FAM230J-LOC105372942-GGTLC5P.NCBI.rev.compl                            ------------------------------------------------------------ 46364
human.FAM230A.ncRNA.18487127.18500594-GGTLC3.18516335.18518165.rev.compl.REF ------------------------------------------------------------ 31039

chimp.LOC112206744-LOC107973052-GGT2.revcompl                                ------------------------------------------------------------ 61762
chimp54546-135457.revcompl.Pan.troglodytes.clone.rp43-41g5.GenBank:AC099533.36  ------------------------------------------------------------ 69108
human.FAM30B-LOC105372935-GGT2.NCBI.                                         gacaaaatgtgctatatccataaaacggaaccctactcagcaataaaaaggtgtgaacta 90817
human.FAM230E-LOC105377182-GGT3P.NCBI.                                       ------------------------------------------------------------ 57310
human.FAM230J-LOC105372942-GGTLC5P.NCBI.rev.compl                            ------------------------------------------------------------ 46364
human.FAM230A.ncRNA.18487127.18500594-GGTLC3.18516335.18518165.rev.compl.REF ------------------------------------------------------------ 31039

chimp.LOC112206744-LOC107973052-GGT2.revcompl                                ------------------------------------------------------------ 61762
chimp54546-135457.revcompl.Pan.troglodytes.clone.rp43-41g5.GenBank:AC099533.36  ------------------------------------------------------------ 69108
human.FAM30B-LOC105372935-GGT2.NCBI.                                         gatacacacaagaacatggagggatctcaaaagcactgtgctgagcgagagaagccagat 90877
human.FAM230E-LOC105377182-GGT3P.NCBI.                                       ------------------------------------------------------------ 57310
human.FAM230J-LOC105372942-GGTLC5P.NCBI.rev.compl                            ------------------------------------------------------------ 46364
human.FAM230A.ncRNA.18487127.18500594-GGTLC3.18516335.18518165.rev.compl.REF ------------------------------------------------------------ 31039

chimp.LOC112206744-LOC107973052-GGT2.revcompl                                ------------------------------------------------------------ 61762
chimp54546-135457.revcompl.Pan.troglodytes.clone.rp43-41g5.GenBank:AC099533.36  ------------------------------------------------------------ 69108
human.FAM30B-LOC105372935-GGT2.NCBI.                                         tcaaagggctacacactgtaggatgccatgtgcatcacattctgtacagtgcaaaactgt 90937
human.FAM230E-LOC105377182-GGT3P.NCBI.                                       ------------------------------------------------------------ 57310
human.FAM230J-LOC105372942-GGTLC5P.NCBI.rev.compl                            ------------------------------------------------------------ 46364
human.FAM230A.ncRNA.18487127.18500594-GGTLC3.18516335.18518165.rev.compl.REF ------------------------------------------------------------ 31039

chimp.LOC112206744-LOC107973052-GGT2.revcompl                                ------------------------------------------------------------ 61762
chimp54546-135457.revcompl.Pan.troglodytes.clone.rp43-41g5.GenBank:AC099533.36  ------------------------------------------------------------ 69108
human.FAM30B-LOC105372935-GGT2.NCBI.                                         agtggcaggattcaggtcccctgaggctggggtcctggaggcaggagaggggctggtagc 90997
human.FAM230E-LOC105377182-GGT3P.NCBI.                                       ------------------------------------------------------------ 57310
human.FAM230J-LOC105372942-GGTLC5P.NCBI.rev.compl                            ------------------------------------------------------------ 46364
human.FAM230A.ncRNA.18487127.18500594-GGTLC3.18516335.18518165.rev.compl.REF ------------------------------------------------------------ 31039

chimp.LOC112206744-LOC107973052-GGT2.revcompl                                ------------------------------------------------------------ 61762
chimp54546-135457.revcompl.Pan.troglodytes.clone.rp43-41g5.GenBank:AC099533.36  ------------------------------------------------------------ 69108
human.FAM30B-LOC105372935-GGT2.NCBI.                                         aaggaggcacaagggaacttttttggggacagaggagtggtccatatttgataggtggtgg 91057
human.FAM230E-LOC105377182-GGT3P.NCBI.                                       ------------------------------------------------------------ 57310
human.FAM230J-LOC105372942-GGTLC5P.NCBI.rev.compl                            ------------------------------------------------------------ 46364
human.FAM230A.ncRNA.18487127.18500594-GGTLC3.18516335.18518165.rev.compl.REF ------------------------------------------------------------ 31039

chimp.LOC112206744-LOC107973052-GGT2.revcompl                                ------------------------------------------------------------ 61762
chimp54546-135457.revcompl.Pan.troglodytes.clone.rp43-41g5.GenBank:AC099533.36  ------------------------------------------------------------ 69108
human.FAM30B-LOC105372935-GGT2.NCBI.                                         ttgtgtggctgctcatgtttgtcaacaaagaattacatattttaaactggtgagttccgt 91117
human.FAM230E-LOC105377182-GGT3P.NCBI.                                       ------------------------------------------------------------ 57310
human.FAM230J-LOC105372942-GGTLC5P.NCBI.rev.compl                            ------------------------------------------------------------ 46364
human.FAM230A.ncRNA.18487127.18500594-GGTLC3.18516335.18518165.rev.compl.REF ------------------------------------------------------------ 31039

chimp.LOC112206744-LOC107973052-GGT2.revcompl                                ------------------------------------------------------------ 61762
chimp54546-135457.revcompl.Pan.troglodytes.clone.rp43-41g5.GenBank:AC099533.36  ------------------------------------------------------------ 69108
human.FAM30B-LOC105372935-GGT2.NCBI.                                         tgtatgcaagcaaattataccacaatgaagcaaatctttaaaacactaggttccttttct 91177
human.FAM230E-LOC105377182-GGT3P.NCBI.                                       ------------------------------------------------------------ 57310
human.FAM230J-LOC105372942-GGTLC5P.NCBI.rev.compl                            ------------------------------------------------------------ 46364
human.FAM230A.ncRNA.18487127.18500594-GGTLC3.18516335.18518165.rev.compl.REF ------------------------------------------------------------ 31039

chimp.LOC112206744-LOC107973052-GGT2.revcompl                                ------------------------------------------------------------ 61762
chimp54546-135457.revcompl.Pan.troglodytes.clone.rp43-41g5.GenBank:AC099533.36  ------------------------------------------------------------ 69108
human.FAM30B-LOC105372935-GGT2.NCBI.                                         ttatctagagctcaaagtagtggcccatgctggctatgacttggagaaatcacacaagct 91237
human.FAM230E-LOC105377182-GGT3P.NCBI.                                       ------------------------------------------------------------ 57310
human.FAM230J-LOC105372942-GGTLC5P.NCBI.rev.compl                            ------------------------------------------------------------ 46364
human.FAM230A.ncRNA.18487127.18500594-GGTLC3.18516335.18518165.rev.compl.REF ------------------------------------------------------------ 31039

chimp.LOC112206744-LOC107973052-GGT2.revcompl                                ------------------------------------------------------------ 61762
chimp54546-135457.revcompl.Pan.troglodytes.clone.rp43-41g5.GenBank:AC099533.36  ------------------------------------------------------------ 69108
human.FAM30B-LOC105372935-GGT2.NCBI.                                         tactgggtctcaactgcttcatttgtacaggtggctgcataacactccccgggcaggggga 91297
human.FAM230E-LOC105377182-GGT3P.NCBI.                                       ------------------------------------------------------------ 57310
human.FAM230J-LOC105372942-GGTLC5P.NCBI.rev.compl                            ------------------------------------------------------------ 46364
human.FAM230A.ncRNA.18487127.18500594-GGTLC3.18516335.18518165.rev.compl.REF ------------------------------------------------------------ 31039

chimp.LOC112206744-LOC107973052-GGT2.revcompl                                ------------------------------------------------------------ 61762
chimp54546-135457.revcompl.Pan.troglodytes.clone.rp43-41g5.GenBank:AC099533.36  ------------------------------------------------------------ 69108
human.FAM30B-LOC105372935-GGT2.NCBI.                                         actgtagggatgaaatcagcaaatgtgagtgaacacacagaggaagcaacacataggcct 91357
human.FAM230E-LOC105377182-GGT3P.NCBI.                                       ------------------------------------------------------------ 57310
human.FAM230J-LOC105372942-GGTLC5P.NCBI.rev.compl                            ------------------------------------------------------------ 46364
human.FAM230A.ncRNA.18487127.18500594-GGTLC3.18516335.18518165.rev.compl.REF ------------------------------------------------------------ 31039

chimp.LOC112206744-LOC107973052-GGT2.revcompl                                ------------------------------------------------------------ 61762
chimp54546-135457.revcompl.Pan.troglodytes.clone.rp43-41g5.GenBank:AC099533.36  ------------------------------------------------------------ 69108
human.FAM30B-LOC105372935-GGT2.NCBI.                                         tccccaaatacccacttctgttttctttcctcagtgtgagtaagtggctctaatgatcca 91417
human.FAM230E-LOC105377182-GGT3P.NCBI.                                       ------------------------------------------------------------ 57310
human.FAM230J-LOC105372942-GGTLC5P.NCBI.rev.compl                            ------------------------------------------------------------ 46364
human.FAM230A.ncRNA.18487127.18500594-GGTLC3.18516335.18518165.rev.compl.REF ------------------------------------------------------------ 31039

chimp.LOC112206744-LOC107973052-GGT2.revcompl                                ------------------------------------------------------------ 61762
chimp54546-135457.revcompl.Pan.troglodytes.clone.rp43-41g5.GenBank:AC099533.36  ------------------------------------------------------------ 69108
human.FAM30B-LOC105372935-GGT2.NCBI.                                         caaaccttccaaccagccaagtcaaggaccagcaggaggactcaggaaagtactgaaatg 91477
human.FAM230E-LOC105377182-GGT3P.NCBI.                                       ------------------------------------------------------------ 57310
human.FAM230J-LOC105372942-GGTLC5P.NCBI.rev.compl                            ------------------------------------------------------------ 46364
human.FAM230A.ncRNA.18487127.18500594-GGTLC3.18516335.18518165.rev.compl.REF ------------------------------------------------------------ 31039

chimp.LOC112206744-LOC107973052-GGT2.revcompl                                ------------------------------------------------------------ 61762
chimp54546-135457.revcompl.Pan.troglodytes.clone.rp43-41g5.GenBank:AC099533.36  ------------------------------------------------------------ 69108
human.FAM30B-LOC105372935-GGT2.NCBI.                                         ctggggagaaccaggacagcggctgagcacagccagacagggagggcagcctggtggggc 91537
human.FAM230E-LOC105377182-GGT3P.NCBI.                                       ------------------------------------------------------------ 57310
human.FAM230J-LOC105372942-GGTLC5P.NCBI.rev.compl                            ------------------------------------------------------------ 46364
human.FAM230A.ncRNA.18487127.18500594-GGTLC3.18516335.18518165.rev.compl.REF ------------------------------------------------------------ 31039
```

```
chimp.LOC112206744-LOC107973052-GGT2.revcompl                                    ------------------------------------------------------------ 61762
chimp54546-135457.revcompl.Pan.troglodytes.clone.rp43-41g5.GenBank:AC099533.36   ------------------------------------------------------------ 69108
human.FAM30B-LOC105372935-GGT2.NCBI.                                             cctgggagccatggacgcgactgcgctgggagtggggatgggagatagcaggagttggtg 91597
human.FAM230E-LOC105377182-GGT3P.NCBI.                                           ------------------------------------------------------------ 57310
human.FAM230J-LOC105372942-GGTLC5P.NCBI.rev.compl                                ------------------------------------------------------------ 46364
human.FAM230A.ncRNA.18487127.18500594-GGTLC3.18516335.18518165.rev.compl.REF     ------------------------------------------------------------ 31039

chimp.LOC112206744-LOC107973052-GGT2.revcompl                                    ------------------------------------------------------------ 61762
chimp54546-135457.revcompl.Pan.troglodytes.clone.rp43-41g5.GenBank:AC099533.36   ------------------------------------------------------------ 69108
human.FAM30B-LOC105372935-GGT2.NCBI.                                             ggtggggcacaaagtgggagccagggccaaattgctgctgatttgaccattttgcaaaag 91657
human.FAM230E-LOC105377182-GGT3P.NCBI.                                           ------------------------------------------------------------ 57310
human.FAM230J-LOC105372942-GGTLC5P.NCBI.rev.compl                                ------------------------------------------------------------ 46364
human.FAM230A.ncRNA.18487127.18500594-GGTLC3.18516335.18518165.rev.compl.REF     ------------------------------------------------------------ 31039

chimp.LOC112206744-LOC107973052-GGT2.revcompl                                    ------------------------------------------------------------ 61762
chimp54546-135457.revcompl.Pan.troglodytes.clone.rp43-41g5.GenBank:AC099533.36   -------------------------------TTACAGTAGTATAGGAAATGATATCTTGT 69137
human.FAM30B-LOC105372935-GGT2.NCBI.                                             gccagcctggctgtggcctcaggcctctcatttcctctaccataatgaggggtcacggaa 91717
human.FAM230E-LOC105377182-GGT3P.NCBI.                                           ------------------------------------------------------------ 57310
human.FAM230J-LOC105372942-GGTLC5P.NCBI.rev.compl                                ------------------------------------------------------------ 46364
human.FAM230A.ncRNA.18487127.18500594-GGTLC3.18516335.18518165.rev.compl.REF     ------------------------------------------------------------ 31039

chimp.LOC112206744-LOC107973052-GGT2.revcompl                                    ------------------------------------------------------------ 61762
chimp54546-135457.revcompl.Pan.troglodytes.clone.rp43-41g5.GenBank:AC099533.36   GGGGTTTAAAATAAATGAAACATACTGAAATATGTATGGGTACAGTTATATATCTGGGAT 69197
human.FAM30B-LOC105372935-GGT2.NCBI.                                             gtgccttcccacaactgcagcatctccctccacatttgggctcaaatacagagtggggcc 91777
human.FAM230E-LOC105377182-GGT3P.NCBI.                                           ------------------------------------------------------------ 57310
human.FAM230J-LOC105372942-GGTLC5P.NCBI.rev.compl                                ------------------------------------------------------------ 46364
human.FAM230A.ncRNA.18487127.18500594-GGTLC3.18516335.18518165.rev.compl.REF     ------------------------------------------------------------ 31039

chimp.LOC112206744-LOC107973052-GGT2.revcompl                                    ------------------------------------------------------------ 61762
chimp54546-135457.revcompl.Pan.troglodytes.clone.rp43-41g5.GenBank:AC099533.36   TTGCACTGAAATAGTGTGGG---------------------------------------- 69217
human.FAM30B-LOC105372935-GGT2.NCBI.                                             ggagacagaaaaagtctccgtgacacactgtgatttgtttctgcactgtgtgaacccttg 91837
human.FAM230E-LOC105377182-GGT3P.NCBI.                                           ------------------------------------------------------------ 57310
human.FAM230J-LOC105372942-GGTLC5P.NCBI.rev.compl                                ------------------------------------------------------------ 46364
human.FAM230A.ncRNA.18487127.18500594-GGTLC3.18516335.18518165.rev.compl.REF     ------------------------------------------------------------ 31039

chimp.LOC112206744-LOC107973052-GGT2.revcompl                                    ------------------------------------------------------------ 61762
chimp54546-135457.revcompl.Pan.troglodytes.clone.rp43-41g5.GenBank:AC099533.36   ------------------------------------------------------------ 69217
human.FAM30B-LOC105372935-GGT2.NCBI.                                             gctctgtgacacagaacacaggtttcagcagttcctgtcacttcccacaggcgaaacaga 91897
human.FAM230E-LOC105377182-GGT3P.NCBI.                                           ------------------------------------------------------------ 57310
human.FAM230J-LOC105372942-GGTLC5P.NCBI.rev.compl                                ------------------------------------------------------------ 46364
human.FAM230A.ncRNA.18487127.18500594-GGTLC3.18516335.18518165.rev.compl.REF     ------------------------------------------------------------ 31039

chimp.LOC112206744-LOC107973052-GGT2.revcompl                                    ------------------------------------------------------------ 61762
chimp54546-135457.revcompl.Pan.troglodytes.clone.rp43-41g5.GenBank:AC099533.36   ------------------------------------------------------------ 69217
human.FAM30B-LOC105372935-GGT2.NCBI.                                             cagagcattcttttgtccggcttctggtgaaacgccattatcgttatttttaggaggaag 91957
human.FAM230E-LOC105377182-GGT3P.NCBI.                                           ------------------------------------------------------------ 57310
human.FAM230J-LOC105372942-GGTLC5P.NCBI.rev.compl                                ------------------------------------------------------------ 46364
human.FAM230A.ncRNA.18487127.18500594-GGTLC3.18516335.18518165.rev.compl.REF     ------------------------------------------------------------ 31039

chimp.LOC112206744-LOC107973052-GGT2.revcompl                                    ------------------------------------------------------------ 61762
chimp54546-135457.revcompl.Pan.troglodytes.clone.rp43-41g5.GenBank:AC099533.36   ----------GTAGAGGGAAGCAGGAAAGAGGATACATGAAATTAGCTTGGCCATAAGAT 69267
human.FAM30B-LOC105372935-GGT2.NCBI.                                             atggagaattggggtgggtccccacaaagaggatggattcagaagacttccaggaacc-- 92015
human.FAM230E-LOC105377182-GGT3P.NCBI.                                           ------------------------------------------------------------ 57310
human.FAM230J-LOC105372942-GGTLC5P.NCBI.rev.compl                                ------------------------------------------------------------ 46364
human.FAM230A.ncRNA.18487127.18500594-GGTLC3.18516335.18518165.rev.compl.REF     ------------------------------------------------------------ 31039

chimp.LOC112206744-LOC107973052-GGT2.revcompl                                    ------------------------------------------------------------ 61762
chimp54546-135457.revcompl.Pan.troglodytes.clone.rp43-41g5.GenBank:AC099533.36   TGTTGTTGAAATTGAATGGATACTCGGGGCTTCATTACACAATTCTCTTTACTCTTACAT 69327
human.FAM30B-LOC105372935-GGT2.NCBI.                                             ------ttaaacggcttggaaactaattgtgccagttctttttgatgtctaattttattc 92069
human.FAM230E-LOC105377182-GGT3P.NCBI.                                           ------------------------------------------------------------ 57310
human.FAM230J-LOC105372942-GGTLC5P.NCBI.rev.compl                                ------------------------------------------------------------ 46364
human.FAM230A.ncRNA.18487127.18500594-GGTLC3.18516335.18518165.rev.compl.REF     ------------------------------------------------------------ 31039

chimp.LOC112206744-LOC107973052-GGT2.revcompl                                    ------------------------------------------------------------ 61762
chimp54546-135457.revcompl.Pan.troglodytes.clone.rp43-41g5.GenBank:AC099533.36   AGCTCTACACTCTCAACATAAATAAGAATAAAAACACAAAAAACACACAGATACATCTAT 69387
human.FAM30B-LOC105372935-GGT2.NCBI.                                             taaaatttcctctcagcacaatatattaatactgatcagccaagagaaggttaggttttg 92129
human.FAM230E-LOC105377182-GGT3P.NCBI.                                           ------------------------------------------------------------ 57310
human.FAM230J-LOC105372942-GGTLC5P.NCBI.rev.compl                                ------------------------------------------------------------ 46364
human.FAM230A.ncRNA.18487127.18500594-GGTLC3.18516335.18518165.rev.compl.REF     ------------------------------------------------------------ 31039

chimp.LOC112206744-LOC107973052-GGT2.revcompl                                    ------------------------------------------------------------ 61762
chimp54546-135457.revcompl.Pan.troglodytes.clone.rp43-41g5.GenBank:AC099533.36   GCACACACATATATTTAAAATACACAAAAATATTAGCATATAAGTCACTGGGGGTAAATT 69447
human.FAM30B-LOC105372935-GGT2.NCBI.                                             gctcacaaata-gtcaaggagagaatctccttcattcattgagaaggcagaagatggcgt 92188
human.FAM230E-LOC105377182-GGT3P.NCBI.                                           ------------------------------------------------------------ 57310
human.FAM230J-LOC105372942-GGTLC5P.NCBI.rev.compl                                ------------------------------------------------------------ 46364
human.FAM230A.ncRNA.18487127.18500594-GGTLC3.18516335.18518165.rev.compl.REF     ------------------------------------------------------------ 31039

chimp.LOC112206744-LOC107973052-GGT2.revcompl                                    ------------------------------------------------------------ 61762
chimp54546-135457.revcompl.Pan.troglodytes.clone.rp43-41g5.GenBank:AC099533.36   TAGTTCCTGTTCCAAGGTTCTTGTACTGACTAGGAAGAGGATAGAAGTACT--------- 69498
human.FAM30B-LOC105372935-GGT2.NCBI.                                             taaagagtattcacaggtaactaagattgccagagggagaatcgagtgagttgtcagtgg 92248
human.FAM230E-LOC105377182-GGT3P.NCBI.                                           ------------------------------------------------------------ 57310
human.FAM230J-LOC105372942-GGTLC5P.NCBI.rev.compl                                ------------------------------------------------------------ 46364
human.FAM230A.ncRNA.18487127.18500594-GGTLC3.18516335.18518165.rev.compl.REF     ------------------------------------------------------------ 31039

chimp.LOC112206744-LOC107973052-GGT2.revcompl                                    ------------------------------------------------------------ 61762
chimp54546-135457.revcompl.Pan.troglodytes.clone.rp43-41g5.GenBank:AC099533.36   ------------------------------------------------------------ 69498
human.FAM30B-LOC105372935-GGT2.NCBI.                                             tgaaatacagaggaggtagagagagctacactgagacaacaaaggcagcttaagggaaca 92308
human.FAM230E-LOC105377182-GGT3P.NCBI.                                           ------------------------------------------------------------ 57310
human.FAM230J-LOC105372942-GGTLC5P.NCBI.rev.compl                                ------------------------------------------------------------ 46364
human.FAM230A.ncRNA.18487127.18500594-GGTLC3.18516335.18518165.rev.compl.REF     ------------------------------------------------------------ 31039

chimp.LOC112206744-LOC107973052-GGT2.revcompl                                    ------------------------------------------------------------ 61762
chimp54546-135457.revcompl.Pan.troglodytes.clone.rp43-41g5.GenBank:AC099533.36   ------------------------------------------------------------ 69498
human.FAM30B-LOC105372935-GGT2.NCBI.                                             aagccagctaagggaacaaagccggggtccctcaggtctgtggccaccggctccagccag 92368
human.FAM230E-LOC105377182-GGT3P.NCBI.                                           ------------------------------------------------------------ 57310
human.FAM230J-LOC105372942-GGTLC5P.NCBI.rev.compl                                ------------------------------------------------------------ 46364
human.FAM230A.ncRNA.18487127.18500594-GGTLC3.18516335.18518165.rev.compl.REF     ------------------------------------------------------------ 31039

chimp.LOC112206744-LOC107973052-GGT2.revcompl                                    ------------------------------------------------------------ 61762
chimp54546-135457.revcompl.Pan.troglodytes.clone.rp43-41g5.GenBank:AC099533.36   ------------------------------------------------------------ 69498
human.FAM30B-LOC105372935-GGT2.NCBI.                                             gacagcctggtagctgaagagcacatccttgcccgttacatgcctgtgtttcaagtgatg 92428
human.FAM230E-LOC105377182-GGT3P.NCBI.                                           ------------------------------------------------------------ 57310
human.FAM230J-LOC105372942-GGTLC5P.NCBI.rev.compl                                ------------------------------------------------------------ 46364
human.FAM230A.ncRNA.18487127.18500594-GGTLC3.18516335.18518165.rev.compl.REF     ------------------------------------------------------------ 31039

chimp.LOC112206744-LOC107973052-GGT2.revcompl                                    ------------------------------------------------------------ 61762
chimp54546-135457.revcompl.Pan.troglodytes.clone.rp43-41g5.GenBank:AC099533.36   ------------------------------------------------------------ 69498
human.FAM30B-LOC105372935-GGT2.NCBI.                                             ctaacttgctaatgcagatatctccccaaaccctgcccctggctcaccacccagtgagca 92488
human.FAM230E-LOC105377182-GGT3P.NCBI.                                           ------------------------------------------------------------ 57310
human.FAM230J-LOC105372942-GGTLC5P.NCBI.rev.compl                                ------------------------------------------------------------ 46364
human.FAM230A.ncRNA.18487127.18500594-GGTLC3.18516335.18518165.rev.compl.REF     ------------------------------------------------------------ 31039

chimp.LOC112206744-LOC107973052-GGT2.revcompl                                    ------------------------------------------------------------ 61762
chimp54546-135457.revcompl.Pan.troglodytes.clone.rp43-41g5.GenBank:AC099533.36   ------------------------------------------------------------ 69498
human.FAM30B-LOC105372935-GGT2.NCBI.                                             atcactcactttccccaggacaccttcagggaggtactctgccagcaccgcacacttaaa 92548
human.FAM230E-LOC105377182-GGT3P.NCBI.                                           ------------------------------------------------------------ 57310
human.FAM230J-LOC105372942-GGTLC5P.NCBI.rev.compl                                ------------------------------------------------------------ 46364
human.FAM230A.ncRNA.18487127.18500594-GGTLC3.18516335.18518165.rev.compl.REF     ------------------------------------------------------------ 31039

chimp.LOC112206744-LOC107973052-GGT2.revcompl                                    ------------------------------------------------------------ 61762
chimp54546-135457.revcompl.Pan.troglodytes.clone.rp43-41g5.GenBank:AC099533.36   ------------------------------------------------------------ 69498
human.FAM30B-LOC105372935-GGT2.NCBI.                                             cagctactgcaggttcttctctcagacgcagcacggttcctgagatgacgtcagcgctgc 92608
human.FAM230E-LOC105377182-GGT3P.NCBI.                                           ------------------------------------------------------------ 57310
human.FAM230J-LOC105372942-GGTLC5P.NCBI.rev.compl                                ------------------------------------------------------------ 46364
human.FAM230A.ncRNA.18487127.18500594-GGTLC3.18516335.18518165.rev.compl.REF     ------------------------------------------------------------ 31039

chimp.LOC112206744-LOC107973052-GGT2.revcompl                                    ------------------------------------------------------------ 61762
```

```
chimp54546-135457.revcompl.Pan.troglodytes.clone.rp43-41g5.GenBank:AC099533.36      ------------------------------------------------------------ 69498
human.FAM30B-LOC105372935-GGT2.NCBI.                                                 tcgctgaaagcaccaggcaaggaggcaaaacaaaggaaatcctaaagagcaggcgagaga 92668
human.FAM230E-LOC105377182-GGT3P.NCBI.                                               ------------------------------------------------------------ 57310
human.FAM230J-LOC105372942-GGTLC5P.NCBI.rev.compl                                    ------------------------------------------------------------ 46364
human.FAM230A.ncRNA.18487127.18500594-GGTLC3.18516335.18518165.rev.compl.REF         ------------------------------------------------------------ 31039

chimp.LOC112206744-LOC107973052-GGT2.revcompl                                        ------------------------------------------------------------ 61762
chimp54546-135457.revcompl.Pan.troglodytes.clone.rp43-41g5.GenBank:AC099533.36      ------------------------------------------------------------ 69498
human.FAM30B-LOC105372935-GGT2.NCBI.                                                 acagtggaactggcagagagaggcactggtgcggatggagagggcagcaataaaaactgg 92728
human.FAM230E-LOC105377182-GGT3P.NCBI.                                               ------------------------------------------------------------ 57310
human.FAM230J-LOC105372942-GGTLC5P.NCBI.rev.compl                                    ------------------------------------------------------------ 46364
human.FAM230A.ncRNA.18487127.18500594-GGTLC3.18516335.18518165.rev.compl.REF         ------------------------------------------------------------ 31039

chimp.LOC112206744-LOC107973052-GGT2.revcompl                                        ------------------------------------------------------------ 61762
chimp54546-135457.revcompl.Pan.troglodytes.clone.rp43-41g5.GenBank:AC099533.36      ------------------------------------------------------------ 69498
human.FAM30B-LOC105372935-GGT2.NCBI.                                                 gggacacttagaggcaaaggtatggcaggtgacaaaaaagatgagctgaagctcaggaga 92788
human.FAM230E-LOC105377182-GGT3P.NCBI.                                               ------------------------------------------------------------ 57310
human.FAM230J-LOC105372942-GGTLC5P.NCBI.rev.compl                                    ------------------------------------------------------------ 46364
human.FAM230A.ncRNA.18487127.18500594-GGTLC3.18516335.18518165.rev.compl.REF         ------------------------------------------------------------ 31039

chimp.LOC112206744-LOC107973052-GGT2.revcompl                                        ------------------------------------------------------------ 61762
chimp54546-135457.revcompl.Pan.troglodytes.clone.rp43-41g5.GenBank:AC099533.36      ------------------------------------------------------------ 69498
human.FAM30B-LOC105372935-GGT2.NCBI.                                                 gacccaggcagcgatgcccaaatgaccggctgaggaatcacctcactgctgctgatgccc 92848
human.FAM230E-LOC105377182-GGT3P.NCBI.                                               ------------------------------------------------------------ 57310
human.FAM230J-LOC105372942-GGTLC5P.NCBI.rev.compl                                    ------------------------------------------------------------ 46364
human.FAM230A.ncRNA.18487127.18500594-GGTLC3.18516335.18518165.rev.compl.REF         ------------------------------------------------------------ 31039

chimp.LOC112206744-LOC107973052-GGT2.revcompl                                        ------------------------------------------------------------ 61762
chimp54546-135457.revcompl.Pan.troglodytes.clone.rp43-41g5.GenBank:AC099533.36      ------------------------------------------------------------ 69498
human.FAM30B-LOC105372935-GGT2.NCBI.                                                 ttgacagcgatgtctgaggctggcgtgagtgctccatgctcagagatacctagaacccag 92908
human.FAM230E-LOC105377182-GGT3P.NCBI.                                               ------------------------------------------------------------ 57310
human.FAM230J-LOC105372942-GGTLC5P.NCBI.rev.compl                                    ------------------------------------------------------------ 46364
human.FAM230A.ncRNA.18487127.18500594-GGTLC3.18516335.18518165.rev.compl.REF         ------------------------------------------------------------ 31039

chimp.LOC112206744-LOC107973052-GGT2.revcompl                                        ------------------------------------------------------------ 61762
chimp54546-135457.revcompl.Pan.troglodytes.clone.rp43-41g5.GenBank:AC099533.36      ------------------------------------------------------------ 69498
human.FAM30B-LOC105372935-GGT2.NCBI.                                                 ctacaatggccttatcagactgaaacaagaacaccaagatattccaagggaggtgacaat 92968
human.FAM230E-LOC105377182-GGT3P.NCBI.                                               ------------------------------------------------------------ 57310
human.FAM230J-LOC105372942-GGTLC5P.NCBI.rev.compl                                    ------------------------------------------------------------ 46364
human.FAM230A.ncRNA.18487127.18500594-GGTLC3.18516335.18518165.rev.compl.REF         ------------------------------------------------------------ 31039

chimp.LOC112206744-LOC107973052-GGT2.revcompl                                        ------------------------------------------------------------ 61762
chimp54546-135457.revcompl.Pan.troglodytes.clone.rp43-41g5.GenBank:AC099533.36      ------------------------------------------------------------ 69498
human.FAM30B-LOC105372935-GGT2.NCBI.                                                 ggggtccaaaggttgtgcatttatggatcacaactatgtgctgagtgtggcccaaatccc 93028
human.FAM230E-LOC105377182-GGT3P.NCBI.                                               ------------------------------------------------------------ 57310
human.FAM230J-LOC105372942-GGTLC5P.NCBI.rev.compl                                    ------------------------------------------------------------ 46364
human.FAM230A.ncRNA.18487127.18500594-GGTLC3.18516335.18518165.rev.compl.REF         ------------------------------------------------------------ 31039

chimp.LOC112206744-LOC107973052-GGT2.revcompl                                        ------------------------------------------------------------ 61762
chimp54546-135457.revcompl.Pan.troglodytes.clone.rp43-41g5.GenBank:AC099533.36      ------------------------------------------------------------ 69498
human.FAM30B-LOC105372935-GGT2.NCBI.                                                 agctccatatggtcacaccatgtctacagccaatcagctgaccggggtggcaacccctgg 93088
human.FAM230E-LOC105377182-GGT3P.NCBI.                                               ------------------------------------------------------------ 57310
human.FAM230J-LOC105372942-GGTLC5P.NCBI.rev.compl                                    ------------------------------------------------------------ 46364
human.FAM230A.ncRNA.18487127.18500594-GGTLC3.18516335.18518165.rev.compl.REF         ------------------------------------------------------------ 31039

chimp.LOC112206744-LOC107973052-GGT2.revcompl                                        ------------------------------------------------------------ 61762
chimp54546-135457.revcompl.Pan.troglodytes.clone.rp43-41g5.GenBank:AC099533.36      ------------------------------------------------------------ 69498
human.FAM30B-LOC105372935-GGT2.NCBI.                                                 ctctgctcaacaataatgacaaccatcatttactgagcacttactatgtgccagaccgat 93148
human.FAM230E-LOC105377182-GGT3P.NCBI.                                               ------------------------------------------------------------ 57310
human.FAM230J-LOC105372942-GGTLC5P.NCBI.rev.compl                                    ------------------------------------------------------------ 46364
human.FAM230A.ncRNA.18487127.18500594-GGTLC3.18516335.18518165.rev.compl.REF         ------------------------------------------------------------ 31039

chimp.LOC112206744-LOC107973052-GGT2.revcompl                                        ------------------------------------------------------------ 61762
chimp54546-135457.revcompl.Pan.troglodytes.clone.rp43-41g5.GenBank:AC099533.36      ------------------------------------------------------------ 69498
human.FAM30B-LOC105372935-GGT2.NCBI.                                                 gctaagtgctttcctcagatgtttccatttaactcttagaatcaccctaatgagtatctc 93208
human.FAM230E-LOC105377182-GGT3P.NCBI.                                               ------------------------------------------------------------ 57310
human.FAM230J-LOC105372942-GGTLC5P.NCBI.rev.compl                                    ------------------------------------------------------------ 46364
human.FAM230A.ncRNA.18487127.18500594-GGTLC3.18516335.18518165.rev.compl.REF         ------------------------------------------------------------ 31039

chimp.LOC112206744-LOC107973052-GGT2.revcompl                                        ------------------------------------------------------------ 61762
chimp54546-135457.revcompl.Pan.troglodytes.clone.rp43-41g5.GenBank:AC099533.36      ------------------------------------------------------------ 69498
human.FAM30B-LOC105372935-GGT2.NCBI.                                                 ctattattttacagatgaggaaatgagagtttagaaaagtaacttgcaaaatcatactgc 93268
human.FAM230E-LOC105377182-GGT3P.NCBI.                                               ------------------------------------------------------------ 57310
human.FAM230J-LOC105372942-GGTLC5P.NCBI.rev.compl                                    ------------------------------------------------------------ 46364
human.FAM230A.ncRNA.18487127.18500594-GGTLC3.18516335.18518165.rev.compl.REF         ------------------------------------------------------------ 31039

chimp.LOC112206744-LOC107973052-GGT2.revcompl                                        ------------------------------------------------------------ 61762
chimp54546-135457.revcompl.Pan.troglodytes.clone.rp43-41g5.GenBank:AC099533.36      ------------------------------------------------------------ 69498
human.FAM30B-LOC105372935-GGT2.NCBI.                                                 tggctccataacaaatgttgattgaatgttgtctgggtggacagaggcaaggatgaccgg 93328
human.FAM230E-LOC105377182-GGT3P.NCBI.                                               ------------------------------------------------------------ 57310
human.FAM230J-LOC105372942-GGTLC5P.NCBI.rev.compl                                    ------------------------------------------------------------ 46364
human.FAM230A.ncRNA.18487127.18500594-GGTLC3.18516335.18518165.rev.compl.REF         ------------------------------------------------------------ 31039

chimp.LOC112206744-LOC107973052-GGT2.revcompl                                        ------------------------------------------------------------ 61762
chimp54546-135457.revcompl.Pan.troglodytes.clone.rp43-41g5.GenBank:AC099533.36      ------------------------------------------------------------ 69498
human.FAM30B-LOC105372935-GGT2.NCBI.                                                 aacgagacaaacaaacaaaaggctgactgacagaccattcgggtagcacttttcctggga 93388
human.FAM230E-LOC105377182-GGT3P.NCBI.                                               ------------------------------------------------------------ 57310
human.FAM230J-LOC105372942-GGTLC5P.NCBI.rev.compl                                    ------------------------------------------------------------ 46364
human.FAM230A.ncRNA.18487127.18500594-GGTLC3.18516335.18518165.rev.compl.REF         ------------------------------------------------------------ 31039

chimp.LOC112206744-LOC107973052-GGT2.revcompl                                        ------------------------------------------------------------ 61762
chimp54546-135457.revcompl.Pan.troglodytes.clone.rp43-41g5.GenBank:AC099533.36      --------------------------------------------------AACTCATAGG 69508
human.FAM30B-LOC105372935-GGT2.NCBI.                                                 ttaagcttagaccttgtaggcctatcagccagaagcagtgtgtaaaataagagtggctga 93448
human.FAM230E-LOC105377182-GGT3P.NCBI.                                               ------------------------------------------------------------ 57310
human.FAM230J-LOC105372942-GGTLC5P.NCBI.rev.compl                                    ------------------------------------------------------------ 46364
human.FAM230A.ncRNA.18487127.18500594-GGTLC3.18516335.18518165.rev.compl.REF         ------------------------------------------------------------ 31039

chimp.LOC112206744-LOC107973052-GGT2.revcompl                                        ------------------------------------------------------------ 61762
chimp54546-135457.revcompl.Pan.troglodytes.clone.rp43-41g5.GenBank:AC099533.36      CTGGGCGTGGTGGCTCACGCCTGTAATCCCAACACTTTAGGACGCCGAGGTAAGCAGATC 69568
human.FAM30B-LOC105372935-GGT2.NCBI.                                                 ctgggcacggtggctcaagcctgtaatccaagcactttgggaggccaaggcgggcggatc 93508
human.FAM230E-LOC105377182-GGT3P.NCBI.                                               ------------------------------------------------------------ 57310
human.FAM230J-LOC105372942-GGTLC5P.NCBI.rev.compl                                    ------------------------------------------------------------ 46364
human.FAM230A.ncRNA.18487127.18500594-GGTLC3.18516335.18518165.rev.compl.REF         ------------------------------------------------------------ 31039

chimp.LOC112206744-LOC107973052-GGT2.revcompl                                        ------------------------------------------------------------ 61762
chimp54546-135457.revcompl.Pan.troglodytes.clone.rp43-41g5.GenBank:AC099533.36      TCTTAAGGTCAGGAGTTCAAGACCAGCCTGGCCAACATGGTGAAACCCTGTCTCTACTAA 69628
human.FAM30B-LOC105372935-GGT2.NCBI.                                                 acctgaggtcaggagttcaagaccagcctggccaacatgatgaaaccccgtctctactac 93568
human.FAM230E-LOC105377182-GGT3P.NCBI.                                               ------------------------------------------------------------ 57310
human.FAM230J-LOC105372942-GGTLC5P.NCBI.rev.compl                                    ------------------------------------------------------------ 46364
human.FAM230A.ncRNA.18487127.18500594-GGTLC3.18516335.18518165.rev.compl.REF         ------------------------------------------------------------ 31039

chimp.LOC112206744-LOC107973052-GGT2.revcompl                                        ------------------------------------------------------------ 61762
chimp54546-135457.revcompl.Pan.troglodytes.clone.rp43-41g5.GenBank:AC099533.36      AAAAGAATACAAAAATTGGCCGGGCATAGTGGTGCACACCTGTGGTCCCAGCTACTCAGG 69688
human.FAM30B-LOC105372935-GGT2.NCBI.                                                 a----aatacaaaaattagccgggtgtgatggcacacgcctgtaatcctagttactcggg 93624
human.FAM230E-LOC105377182-GGT3P.NCBI.                                               ------------------------------------------------------------ 57310
human.FAM230J-LOC105372942-GGTLC5P.NCBI.rev.compl                                    ------------------------------------------------------------ 46364
human.FAM230A.ncRNA.18487127.18500594-GGTLC3.18516335.18518165.rev.compl.REF         ------------------------------------------------------------ 31039

chimp.LOC112206744-LOC107973052-GGT2.revcompl                                        ------------------------------------------------------------ 61762
chimp54546-135457.revcompl.Pan.troglodytes.clone.rp43-41g5.GenBank:AC099533.36      TGACTGAAGCAGGAGAATTGCTTGAACCCAAGAAGTGGAGGTTGCAGTGAACCAAGATTG 69748
human.FAM30B-LOC105372935-GGT2.NCBI.                                                 aggctgaggcacgagaatcgcttgaacctgggaggcagaggttgcgataagctgagattg 93684
human.FAM230E-LOC105377182-GGT3P.NCBI.                                               ------------------------------------------------------------ 57310
human.FAM230J-LOC105372942-GGTLC5P.NCBI.rev.compl                                    ------------------------------------------------------------ 46364
human.FAM230A.ncRNA.18487127.18500594-GGTLC3.18516335.18518165.rev.compl.REF         ------------------------------------------------------------ 31039

chimp.LOC112206744-LOC107973052-GGT2.revcompl                                        ------------------------------------------------------------ 61762
chimp54546-135457.revcompl.Pan.troglodytes.clone.rp43-41g5.GenBank:AC099533.36      CTCCACTGCACTCCAGCCTGGGCAGGAGAGGAAGACTCTCTCTGTCTCAACAACAACAAA 69808
```

```
human.FAM30B-LOC105372935-GGT2.NCBI.                                            ggccactgcactccagcctgggcgacagagcaagactccctctgaaaaaaaaaaaaaaaa 93744
human.FAM230E-LOC105377182-GGT3P.NCBI.                                          ------------------------------------------------------------ 57310
human.FAM230J-LOC105372942-GGTLC5P.NCBI.rev.compl                               ------------------------------------------------------------ 46364
human.FAM230A.ncRNA.18487127.18500594-GGTLC3.18516335.18518165.rev.compl.REF    ------------------------------------------------------------ 31039

chimp.LOC112206744-LOC107973052-GGT2.revcompl                                   ------------------------------------------------------------ 61762
chimp54546-135457.revcompl.Pan.troglodytes.clone.rp43-41g5.GenBank:AC099533.36  A----------------------------------------------------------- 69809
human.FAM30B-LOC105372935-GGT2.NCBI.                                            agagtggctacccaagaacgcaacagcagcagcaaacactgatctgatgacaccccagtt 93804
human.FAM230E-LOC105377182-GGT3P.NCBI.                                          ------------------------------------------------------------ 57310
human.FAM230J-LOC105372942-GGTLC5P.NCBI.rev.compl                               ------------------------------------------------------------ 46364
human.FAM230A.ncRNA.18487127.18500594-GGTLC3.18516335.18518165.rev.compl.REF    ------------------------------------------------------------ 31039

chimp.LOC112206744-LOC107973052-GGT2.revcompl                                   ------------------------------------------------------------ 61762
chimp54546-135457.revcompl.Pan.troglodytes.clone.rp43-41g5.GenBank:AC099533.36  ------------------------------------------------------------ 69809
human.FAM30B-LOC105372935-GGT2.NCBI.                                            ttgcagaagaggaaggaaaaataagaatgtcaactctccattcaacagagaaacagtgcc 93864
human.FAM230E-LOC105377182-GGT3P.NCBI.                                          ------------------------------------------------------------ 57310
human.FAM230J-LOC105372942-GGTLC5P.NCBI.rev.compl                               ------------------------------------------------------------ 46364
human.FAM230A.ncRNA.18487127.18500594-GGTLC3.18516335.18518165.rev.compl.REF    ------------------------------------------------------------ 31039

chimp.LOC112206744-LOC107973052-GGT2.revcompl                                   ------------------------------------------------------------ 61762
chimp54546-135457.revcompl.Pan.troglodytes.clone.rp43-41g5.GenBank:AC099533.36  ------------------------------------------------------------ 69809
human.FAM30B-LOC105372935-GGT2.NCBI.                                            accactgtctttaaattttcaaacttatcgagaaaagctccagagaggccagtgccagca 93924
human.FAM230E-LOC105377182-GGT3P.NCBI.                                          ------------------------------------------------------------ 57310
human.FAM230J-LOC105372942-GGTLC5P.NCBI.rev.compl                               ------------------------------------------------------------ 46364
human.FAM230A.ncRNA.18487127.18500594-GGTLC3.18516335.18518165.rev.compl.REF    ------------------------------------------------------------ 31039

chimp.LOC112206744-LOC107973052-GGT2.revcompl                                   ------------------------------------------------------------ 61762
chimp54546-135457.revcompl.Pan.troglodytes.clone.rp43-41g5.GenBank:AC099533.36  ------------------------------------------------------------ 69809
human.FAM30B-LOC105372935-GGT2.NCBI.                                            atgccggctgtctcccaagtctgaagggttttcctctgggtattaggagttgaggtgttt 93984
human.FAM230E-LOC105377182-GGT3P.NCBI.                                          ------------------------------------------------------------ 57310
human.FAM230J-LOC105372942-GGTLC5P.NCBI.rev.compl                               ------------------------------------------------------------ 46364
human.FAM230A.ncRNA.18487127.18500594-GGTLC3.18516335.18518165.rev.compl.REF    ------------------------------------------------------------ 31039

chimp.LOC112206744-LOC107973052-GGT2.revcompl                                   ------------------------------------------------------------ 61762
chimp54546-135457.revcompl.Pan.troglodytes.clone.rp43-41g5.GenBank:AC099533.36  ------------------------------------------------------------ 69809
human.FAM30B-LOC105372935-GGT2.NCBI.                                            ggttacagaaggaaacaggatgcacgggagatcacacacagagtaagaggtgcctctcaa 94044
human.FAM230E-LOC105377182-GGT3P.NCBI.                                          ------------------------------------------------------------ 57310
human.FAM230J-LOC105372942-GGTLC5P.NCBI.rev.compl                               ------------------------------------------------------------ 46364
human.FAM230A.ncRNA.18487127.18500594-GGTLC3.18516335.18518165.rev.compl.REF    ------------------------------------------------------------ 31039

chimp.LOC112206744-LOC107973052-GGT2.revcompl                                   ------------------------------------------------------------ 61762
chimp54546-135457.revcompl.Pan.troglodytes.clone.rp43-41g5.GenBank:AC099533.36  ------------------------------------------------------------ 69809
human.FAM30B-LOC105372935-GGT2.NCBI.                                            atgcctttcccacgtttctaactctgctttttttaaagcacgatcagacccgccatctgc 94104
human.FAM230E-LOC105377182-GGT3P.NCBI.                                          ------------------------------------------------------------ 57310
human.FAM230J-LOC105372942-GGTLC5P.NCBI.rev.compl                               ------------------------------------------------------------ 46364
human.FAM230A.ncRNA.18487127.18500594-GGTLC3.18516335.18518165.rev.compl.REF    ------------------------------------------------------------ 31039

chimp.LOC112206744-LOC107973052-GGT2.revcompl                                   ------------------------------------------------------------ 61762
chimp54546-135457.revcompl.Pan.troglodytes.clone.rp43-41g5.GenBank:AC099533.36  ------------------------------------------------------------ 69809
human.FAM30B-LOC105372935-GGT2.NCBI.                                            acttttacctgctgcctgacgaaaaccagatgccaaaggcctcccagcactcacggggcc 94164
human.FAM230E-LOC105377182-GGT3P.NCBI.                                          ------------------------------------------------------------ 57310
human.FAM230J-LOC105372942-GGTLC5P.NCBI.rev.compl                               ------------------------------------------------------------ 46364
human.FAM230A.ncRNA.18487127.18500594-GGTLC3.18516335.18518165.rev.compl.REF    ------------------------------------------------------------ 31039

chimp.LOC112206744-LOC107973052-GGT2.revcompl                                   ------------------------------------------------------------ 61762
chimp54546-135457.revcompl.Pan.troglodytes.clone.rp43-41g5.GenBank:AC099533.36  ------------------------------------------------------------ 69809
human.FAM30B-LOC105372935-GGT2.NCBI.                                            ctttacccttaggaaaacaccgtgattcagccaggcgagattgatgcaacattgtaacaa 94224
human.FAM230E-LOC105377182-GGT3P.NCBI.                                          ------------------------------------------------------------ 57310
human.FAM230J-LOC105372942-GGTLC5P.NCBI.rev.compl                               ------------------------------------------------------------ 46364
human.FAM230A.ncRNA.18487127.18500594-GGTLC3.18516335.18518165.rev.compl.REF    ------------------------------------------------------------ 31039

chimp.LOC112206744-LOC107973052-GGT2.revcompl                                   ------------------------------------------------------------ 61762
chimp54546-135457.revcompl.Pan.troglodytes.clone.rp43-41g5.GenBank:AC099533.36  ------------------------------------------------------------ 69809
human.FAM30B-LOC105372935-GGT2.NCBI.                                            agcgtatgcaaagtgcctagctccggactgcttgcagaaaaggcacattttgactcttct 94284
human.FAM230E-LOC105377182-GGT3P.NCBI.                                          ------------------------------------------------------------ 57310
human.FAM230J-LOC105372942-GGTLC5P.NCBI.rev.compl                               ------------------------------------------------------------ 46364
human.FAM230A.ncRNA.18487127.18500594-GGTLC3.18516335.18518165.rev.compl.REF    ------------------------------------------------------------ 31039

chimp.LOC112206744-LOC107973052-GGT2.revcompl                                   ------------------------------------------------------------ 61762
chimp54546-135457.revcompl.Pan.troglodytes.clone.rp43-41g5.GenBank:AC099533.36  ------------------------------------------------------------ 69809
human.FAM30B-LOC105372935-GGT2.NCBI.                                            gagagtgggaaagatttgaaagatgggatttgcaccttattcttattcaagtctggagat 94344
human.FAM230E-LOC105377182-GGT3P.NCBI.                                          ------------------------------------------------------------ 57310
human.FAM230J-LOC105372942-GGTLC5P.NCBI.rev.compl                               ------------------------------------------------------------ 46364
human.FAM230A.ncRNA.18487127.18500594-GGTLC3.18516335.18518165.rev.compl.REF    ------------------------------------------------------------ 31039

chimp.LOC112206744-LOC107973052-GGT2.revcompl                                   ------------------------------------------------------------ 61762
chimp54546-135457.revcompl.Pan.troglodytes.clone.rp43-41g5.GenBank:AC099533.36  ------------------------------------------------------------ 69809
human.FAM30B-LOC105372935-GGT2.NCBI.                                            ctggaacttcttcctgaagctccatgtttagccagataattgagacgctccacctggtca 94404
human.FAM230E-LOC105377182-GGT3P.NCBI.                                          ------------------------------------------------------------ 57310
human.FAM230J-LOC105372942-GGTLC5P.NCBI.rev.compl                               ------------------------------------------------------------ 46364
human.FAM230A.ncRNA.18487127.18500594-GGTLC3.18516335.18518165.rev.compl.REF    ------------------------------------------------------------ 31039

chimp.LOC112206744-LOC107973052-GGT2.revcompl                                   ------------------------------------------------------------ 61762
chimp54546-135457.revcompl.Pan.troglodytes.clone.rp43-41g5.GenBank:AC099533.36  ------------------------------------------------------------ 69809
human.FAM30B-LOC105372935-GGT2.NCBI.                                            ccactggacaccagccacccaaacattctcctctgcaaggcacttggcattctggtgcct 94464
human.FAM230E-LOC105377182-GGT3P.NCBI.                                          ------------------------------------------------------------ 57310
human.FAM230J-LOC105372942-GGTLC5P.NCBI.rev.compl                               ------------------------------------------------------------ 46364
human.FAM230A.ncRNA.18487127.18500594-GGTLC3.18516335.18518165.rev.compl.REF    ------------------------------------------------------------ 31039

chimp.LOC112206744-LOC107973052-GGT2.revcompl                                   ------------------------------------------------------------ 61762
chimp54546-135457.revcompl.Pan.troglodytes.clone.rp43-41g5.GenBank:AC099533.36  ------------------------------------------------------------ 69809
human.FAM30B-LOC105372935-GGT2.NCBI.                                            gtggagagagcagccttctcactagcacagctcatgaacaaggatccccttccctggaca 94524
human.FAM230E-LOC105377182-GGT3P.NCBI.                                          ------------------------------------------------------------ 57310
human.FAM230J-LOC105372942-GGTLC5P.NCBI.rev.compl                               ------------------------------------------------------------ 46364
human.FAM230A.ncRNA.18487127.18500594-GGTLC3.18516335.18518165.rev.compl.REF    ------------------------------------------------------------ 31039

chimp.LOC112206744-LOC107973052-GGT2.revcompl                                   ------------------------------------------------------------ 61762
chimp54546-135457.revcompl.Pan.troglodytes.clone.rp43-41g5.GenBank:AC099533.36  ------------------------------------------------------------ 69809
human.FAM30B-LOC105372935-GGT2.NCBI.                                            gccaagatgtaaacctgcccccagaaacactgagagtcctagaggtagaatgagtcggcc 94584
human.FAM230E-LOC105377182-GGT3P.NCBI.                                          ------------------------------------------------------------ 57310
human.FAM230J-LOC105372942-GGTLC5P.NCBI.rev.compl                               ------------------------------------------------------------ 46364
human.FAM230A.ncRNA.18487127.18500594-GGTLC3.18516335.18518165.rev.compl.REF    ------------------------------------------------------------ 31039

chimp.LOC112206744-LOC107973052-GGT2.revcompl                                   ------------------------------------------------------------ 61762
chimp54546-135457.revcompl.Pan.troglodytes.clone.rp43-41g5.GenBank:AC099533.36  ------------------------------------------------------------ 69809
human.FAM30B-LOC105372935-GGT2.NCBI.                                            agcaagctccagcccattgaatgcagacccctgagcaaatcaagtcaggagaaggtggac 94644
human.FAM230E-LOC105377182-GGT3P.NCBI.                                          ------------------------------------------------------------ 57310
human.FAM230J-LOC105372942-GGTLC5P.NCBI.rev.compl                               ------------------------------------------------------------ 46364
human.FAM230A.ncRNA.18487127.18500594-GGTLC3.18516335.18518165.rev.compl.REF    ------------------------------------------------------------ 31039

chimp.LOC112206744-LOC107973052-GGT2.revcompl                                   ------------------------------------------------------------ 61762
chimp54546-135457.revcompl.Pan.troglodytes.clone.rp43-41g5.GenBank:AC099533.36  ------------------------------------------------------------ 69809
human.FAM30B-LOC105372935-GGT2.NCBI.                                            aactgtaccctgaagaccacataccacccagacccaacacaaatagctgtcagcactgcc 94704
human.FAM230E-LOC105377182-GGT3P.NCBI.                                          ------------------------------------------------------------ 57310
human.FAM230J-LOC105372942-GGTLC5P.NCBI.rev.compl                               ------------------------------------------------------------ 46364
human.FAM230A.ncRNA.18487127.18500594-GGTLC3.18516335.18518165.rev.compl.REF    ------------------------------------------------------------ 31039

chimp.LOC112206744-LOC107973052-GGT2.revcompl                                   ------------------------------------------------------------ 61762
chimp54546-135457.revcompl.Pan.troglodytes.clone.rp43-41g5.GenBank:AC099533.36  ------------------------------------------------------------ 69809
human.FAM30B-LOC105372935-GGT2.NCBI.                                            acccattacagcttcagatctcgcttctaccagcatttcccagctgagcaacaaccgtca 94764
human.FAM230E-LOC105377182-GGT3P.NCBI.                                          ------------------------------------------------------------ 57310
human.FAM230J-LOC105372942-GGTLC5P.NCBI.rev.compl                               ------------------------------------------------------------ 46364
human.FAM230A.ncRNA.18487127.18500594-GGTLC3.18516335.18518165.rev.compl.REF    ------------------------------------------------------------ 31039

chimp.LOC112206744-LOC107973052-GGT2.revcompl                                   ------------------------------------------------------------ 61762
chimp54546-135457.revcompl.Pan.troglodytes.clone.rp43-41g5.GenBank:AC099533.36  ------------------------------------------------------------ 69809
human.FAM30B-LOC105372935-GGT2.NCBI.                                            aagtctggtcctggaaggacagccctccccgtcttggcagccacactttctggccctagc 94824
```

```
human.FAM230E-LOC105377182-GGT3P.NCBI.                                             ------------------------------------------------------------ 57310
human.FAM230J-LOC105372942-GGTLC5P.NCBI.rev.compl                                  ------------------------------------------------------------ 46364
human.FAM230A.ncRNA.18487127.18500594-GGTLC3.18516335.18518165.rev.compl.REF       ------------------------------------------------------------ 31039

chimp.LOC112206744-LOC107973052-GGT2.revcompl                                      ------------------------------------------------------------ 61762
chimp54546-135457.revcompl.Pan.troglodytes.clone.rp43-41g5.GenBank:AC099533.36     ------------------------------------------------------------ 69809
human.FAM30B-LOC105372935-GGT2.NCBI.                                               ccaaggctgctgtggcatgggaggaatcaatggaggaatacatggctcttccctggttcc 94884
human.FAM230E-LOC105377182-GGT3P.NCBI.                                             ------------------------------------------------------------ 57310
human.FAM230J-LOC105372942-GGTLC5P.NCBI.rev.compl                                  ------------------------------------------------------------ 46364
human.FAM230A.ncRNA.18487127.18500594-GGTLC3.18516335.18518165.rev.compl.REF       ------------------------------------------------------------ 31039

chimp.LOC112206744-LOC107973052-GGT2.revcompl                                      ------------------------------------------------------------ 61762
chimp54546-135457.revcompl.Pan.troglodytes.clone.rp43-41g5.GenBank:AC099533.36     ------------------------------------------------------------ 69809
human.FAM30B-LOC105372935-GGT2.NCBI.                                               ttcctctcccacttcattgactaacactagtcacatggccccaactgaactgtgagggag 94944
human.FAM230E-LOC105377182-GGT3P.NCBI.                                             ------------------------------------------------------------ 57310
human.FAM230J-LOC105372942-GGTLC5P.NCBI.rev.compl                                  ------------------------------------------------------------ 46364
human.FAM230A.ncRNA.18487127.18500594-GGTLC3.18516335.18518165.rev.compl.REF       ------------------------------------------------------------ 31039

chimp.LOC112206744-LOC107973052-GGT2.revcompl                                      ------------------------------------------------------------ 61762
chimp54546-135457.revcompl.Pan.troglodytes.clone.rp43-41g5.GenBank:AC099533.36     ------------------------------------------------------------ 69809
human.FAM30B-LOC105372935-GGT2.NCBI.                                               gctggaaaatctggagtatgtgcatgcctagtgagttgcctctaccacatcctcttaacc 95004
human.FAM230E-LOC105377182-GGT3P.NCBI.                                             ------------------------------------------------------------ 57310
human.FAM230J-LOC105372942-GGTLC5P.NCBI.rev.compl                                  ------------------------------------------------------------ 46364
human.FAM230A.ncRNA.18487127.18500594-GGTLC3.18516335.18518165.rev.compl.REF       ------------------------------------------------------------ 31039

chimp.LOC112206744-LOC107973052-GGT2.revcompl                                      ------------------------------------------------------------ 61762
chimp54546-135457.revcompl.Pan.troglodytes.clone.rp43-41g5.GenBank:AC099533.36     ------------------------------------------------------------ 69809
human.FAM30B-LOC105372935-GGT2.NCBI.                                               cttggcctctcattcccaaagtctagccctattttcaaatgaattcaacgcctcattact 95064
human.FAM230E-LOC105377182-GGT3P.NCBI.                                             ------------------------------------------------------------ 57310
human.FAM230J-LOC105372942-GGTLC5P.NCBI.rev.compl                                  ------------------------------------------------------------ 46364
human.FAM230A.ncRNA.18487127.18500594-GGTLC3.18516335.18518165.rev.compl.REF       ------------------------------------------------------------ 31039

chimp.LOC112206744-LOC107973052-GGT2.revcompl                                      ------------------------------------------------------------ 61762
chimp54546-135457.revcompl.Pan.troglodytes.clone.rp43-41g5.GenBank:AC099533.36     ------------------------------------------------------------ 69809
human.FAM30B-LOC105372935-GGT2.NCBI.                                               tagatctttcctcattacctcaaagccatagtggcaaaaagcaaacagcatcttcctaaa 95124
human.FAM230E-LOC105377182-GGT3P.NCBI.                                             ------------------------------------------------------------ 57310
human.FAM230J-LOC105372942-GGTLC5P.NCBI.rev.compl                                  ------------------------------------------------------------ 46364
human.FAM230A.ncRNA.18487127.18500594-GGTLC3.18516335.18518165.rev.compl.REF       ------------------------------------------------------------ 31039

chimp.LOC112206744-LOC107973052-GGT2.revcompl                                      ------------------------------------------------------------ 61762
chimp54546-135457.revcompl.Pan.troglodytes.clone.rp43-41g5.GenBank:AC099533.36     ------------------------------------------------------------ 69809
human.FAM30B-LOC105372935-GGT2.NCBI.                                               atcttcctcttccatcttcctctcctagatatccatcttacccttctttgccagttaatt 95184
human.FAM230E-LOC105377182-GGT3P.NCBI.                                             ------------------------------------------------------------ 57310
human.FAM230J-LOC105372942-GGTLC5P.NCBI.rev.compl                                  ------------------------------------------------------------ 46364
human.FAM230A.ncRNA.18487127.18500594-GGTLC3.18516335.18518165.rev.compl.REF       ------------------------------------------------------------ 31039

chimp.LOC112206744-LOC107973052-GGT2.revcompl                                      ------------------------------------------------------------ 61762
chimp54546-135457.revcompl.Pan.troglodytes.clone.rp43-41g5.GenBank:AC099533.36     ------------------------------------------------------------ 69809
human.FAM30B-LOC105372935-GGT2.NCBI.                                               tattacaatcttcctttgacagcttcactccctccccgcactgctatcaactcaaatgtg 95244
human.FAM230E-LOC105377182-GGT3P.NCBI.                                             ------------------------------------------------------------ 57310
human.FAM230J-LOC105372942-GGTLC5P.NCBI.rev.compl                                  ------------------------------------------------------------ 46364
human.FAM230A.ncRNA.18487127.18500594-GGTLC3.18516335.18518165.rev.compl.REF       ------------------------------------------------------------ 31039

chimp.LOC112206744-LOC107973052-GGT2.revcompl                                      ------------------------------------------------------------ 61762
chimp54546-135457.revcompl.Pan.troglodytes.clone.rp43-41g5.GenBank:AC099533.36     ------------------------------------------------------------ 69809
human.FAM30B-LOC105372935-GGT2.NCBI.                                               agttatttcagtagcttgagcttgtcactggcctttactttttccccttccaaagaac 95304
human.FAM230E-LOC105377182-GGT3P.NCBI.                                             ------------------------------------------------------------ 57310
human.FAM230J-LOC105372942-GGTLC5P.NCBI.rev.compl                                  ------------------------------------------------------------ 46364
human.FAM230A.ncRNA.18487127.18500594-GGTLC3.18516335.18518165.rev.compl.REF       ------------------------------------------------------------ 31039

chimp.LOC112206744-LOC107973052-GGT2.revcompl                                      ------------------------------------------------------------ 61762
chimp54546-135457.revcompl.Pan.troglodytes.clone.rp43-41g5.GenBank:AC099533.36     ------------------------------------------------------------ 69809
human.FAM30B-LOC105372935-GGT2.NCBI.                                               ctggtacccagctacttaatccttatccttcttcccaaggcccactttcattcctttctc 95364
human.FAM230E-LOC105377182-GGT3P.NCBI.                                             ------------------------------------------------------------ 57310
human.FAM230J-LOC105372942-GGTLC5P.NCBI.rev.compl                                  ------------------------------------------------------------ 46364
human.FAM230A.ncRNA.18487127.18500594-GGTLC3.18516335.18518165.rev.compl.REF       ------------------------------------------------------------ 31039

chimp.LOC112206744-LOC107973052-GGT2.revcompl                                      ------------------------------------------------------------ 61762
chimp54546-135457.revcompl.Pan.troglodytes.clone.rp43-41g5.GenBank:AC099533.36     ------------------------------------------------------------ 69809
human.FAM30B-LOC105372935-GGT2.NCBI.                                               ccccattcatgagcctgcaatacttttcccgctgcataagtcacgtgtcaccatataaac 95424
human.FAM230E-LOC105377182-GGT3P.NCBI.                                             ------------------------------------------------------------ 57310
human.FAM230J-LOC105372942-GGTLC5P.NCBI.rev.compl                                  ------------------------------------------------------------ 46364
human.FAM230A.ncRNA.18487127.18500594-GGTLC3.18516335.18518165.rev.compl.REF       ------------------------------------------------------------ 31039

chimp.LOC112206744-LOC107973052-GGT2.revcompl                                      ------------------------------------------------------------ 61762
chimp54546-135457.revcompl.Pan.troglodytes.clone.rp43-41g5.GenBank:AC099533.36     ------------------------------------------------------------ 69809
human.FAM30B-LOC105372935-GGT2.NCBI.                                               tcacaatttgaggtgctctagaatctgtccacggaattagtcaggacataatctccctgt 95484
human.FAM230E-LOC105377182-GGT3P.NCBI.                                             ------------------------------------------------------------ 57310
human.FAM230J-LOC105372942-GGTLC5P.NCBI.rev.compl                                  ------------------------------------------------------------ 46364
human.FAM230A.ncRNA.18487127.18500594-GGTLC3.18516335.18518165.rev.compl.REF       ------------------------------------------------------------ 31039

chimp.LOC112206744-LOC107973052-GGT2.revcompl                                      ------------------------------------------------------------ 61762
chimp54546-135457.revcompl.Pan.troglodytes.clone.rp43-41g5.GenBank:AC099533.36     ------------------------------------------------------------ 69809
human.FAM30B-LOC105372935-GGT2.NCBI.                                               ttttcggcagtgccacatccatacctgtcttcacaggtttgctcatgccaccattccttt 95544
human.FAM230E-LOC105377182-GGT3P.NCBI.                                             ------------------------------------------------------------ 57310
human.FAM230J-LOC105372942-GGTLC5P.NCBI.rev.compl                                  ------------------------------------------------------------ 46364
human.FAM230A.ncRNA.18487127.18500594-GGTLC3.18516335.18518165.rev.compl.REF       ------------------------------------------------------------ 31039

chimp.LOC112206744-LOC107973052-GGT2.revcompl                                      ------------------------------------------------------------ 61762
chimp54546-135457.revcompl.Pan.troglodytes.clone.rp43-41g5.GenBank:AC099533.36     ------------------------------------------------------------ 69809
human.FAM30B-LOC105372935-GGT2.NCBI.                                               ccccagaacgcccttctgctcctcccacacaaatcctaatcctccctcaggacccattta 95604
human.FAM230E-LOC105377182-GGT3P.NCBI.                                             ------------------------------------------------------------ 57310
human.FAM230J-LOC105372942-GGTLC5P.NCBI.rev.compl                                  ------------------------------------------------------------ 46364
human.FAM230A.ncRNA.18487127.18500594-GGTLC3.18516335.18518165.rev.compl.REF       ------------------------------------------------------------ 31039

chimp.LOC112206744-LOC107973052-GGT2.revcompl                                      ------------------------------------------------------------ 61762
chimp54546-135457.revcompl.Pan.troglodytes.clone.rp43-41g5.GenBank:AC099533.36     ------------------------------------------------------------ 69809
human.FAM30B-LOC105372935-GGT2.NCBI.                                               aaatacaatctcctcactcaactctttcctaccaacttgctttcccctctcctttctgcc 95664
human.FAM230E-LOC105377182-GGT3P.NCBI.                                             ------------------------------------------------------------ 57310
human.FAM230J-LOC105372942-GGTLC5P.NCBI.rev.compl                                  ------------------------------------------------------------ 46364
human.FAM230A.ncRNA.18487127.18500594-GGTLC3.18516335.18518165.rev.compl.REF       ------------------------------------------------------------ 31039

chimp.LOC112206744-LOC107973052-GGT2.revcompl                                      ------------------------------------------------------------ 61762
chimp54546-135457.revcompl.Pan.troglodytes.clone.rp43-41g5.GenBank:AC099533.36     ------------------------------------------------------------ 69809
human.FAM30B-LOC105372935-GGT2.NCBI.                                               ttaccacaaagctaagctctgtctctgaataatgtgctttttgatgataatctgaatggt 95724
human.FAM230E-LOC105377182-GGT3P.NCBI.                                             ------------------------------------------------------------ 57310
human.FAM230J-LOC105372942-GGTLC5P.NCBI.rev.compl                                  ------------------------------------------------------------ 46364
human.FAM230A.ncRNA.18487127.18500594-GGTLC3.18516335.18518165.rev.compl.REF       ------------------------------------------------------------ 31039

chimp.LOC112206744-LOC107973052-GGT2.revcompl                                      ------------------------------------------------------------ 61762
chimp54546-135457.revcompl.Pan.troglodytes.clone.rp43-41g5.GenBank:AC099533.36     ---------------AGTACTAGCTCATGTTAGACTTTGATAAGTGAAGGATGCA----- 69849
human.FAM30B-LOC105372935-GGT2.NCBI.                                               acaagaagaaatcctatttcatgctcctttgagggagtgtctaatataggttgccttcga 95784
human.FAM230E-LOC105377182-GGT3P.NCBI.                                             ------------------------------------------------------------ 57310
human.FAM230J-LOC105372942-GGTLC5P.NCBI.rev.compl                                  ------------------------------------------------------------ 46364
human.FAM230A.ncRNA.18487127.18500594-GGTLC3.18516335.18518165.rev.compl.REF       ------------------------------------------------------------ 31039

chimp.LOC112206744-LOC107973052-GGT2.revcompl                                      ------------------------------------------------------------ 61762
chimp54546-135457.revcompl.Pan.troglodytes.clone.rp43-41g5.GenBank:AC099533.36     ------------------------------------------------------------ 69849
human.FAM30B-LOC105372935-GGT2.NCBI.                                               ctataagtagtagaaaattcccactcaactggcttcaacagtgaggacaaaatgtatgga 95844
human.FAM230E-LOC105377182-GGT3P.NCBI.                                             ------------------------------------------------------------ 57310
human.FAM230J-LOC105372942-GGTLC5P.NCBI.rev.compl                                  ------------------------------------------------------------ 46364
human.FAM230A.ncRNA.18487127.18500594-GGTLC3.18516335.18518165.rev.compl.REF       ------------------------------------------------------------ 31039

chimp.LOC112206744-LOC107973052-GGT2.revcompl                                      ------------------------------------------------------------ 61762
chimp54546-135457.revcompl.Pan.troglodytes.clone.rp43-41g5.GenBank:AC099533.36     ------------------------------------------------------------ 69849
human.FAM30B-LOC105372935-GGT2.NCBI.                                               gattggacgcttcagtcttcaggggcttggccctcagggtttttcccatctcttcccgtc 95904
human.FAM230E-LOC105377182-GGT3P.NCBI.                                             ------------------------------------------------------------ 57310
```

```
human.FAM230J-LOC105372942-GGTLC5P.NCBI.rev.compl                                  ------------------------------------------------------------ 46364
human.FAM230A.ncRNA.18487127.18500594-GGTLC3.18516335.18518165.rev.compl.REF       ------------------------------------------------------------ 31039

chimp.LOC112206744-LOC107973052-GGT2.revcompl                                      ------------------------------------------------------------ 61762
chimp54546-135457.revcompl.Pan.troglodytes.clone.rp43-41g5.GenBank:AC099533.36     ------------------------------------------------------------ 69849
human.FAM30B-LOC105372935-GGT2.NCBI.                                               tgctggcctcaacaaacctgctttcctcattgcaacaagatggctgcttcacgtccgaga 95964
human.FAM230E-LOC105377182-GGT3P.NCBI.                                             ------------------------------------------------------------ 57310
human.FAM230J-LOC105372942-GGTLC5P.NCBI.rev.compl                                  ------------------------------------------------------------ 46364
human.FAM230A.ncRNA.18487127.18500594-GGTLC3.18516335.18518165.rev.compl.REF       ------------------------------------------------------------ 31039

chimp.LOC112206744-LOC107973052-GGT2.revcompl                                      ------------------------------------------------------------ 61762
chimp54546-135457.revcompl.Pan.troglodytes.clone.rp43-41g5.GenBank:AC099533.36     ------------------------------------------------------------ 69849
human.FAM30B-LOC105372935-GGT2.NCBI.                                               atctagtgaaagaagctgtcccttctgcatttccttttacaaggaaaaacatctttccca 96024
human.FAM230E-LOC105377182-GGT3P.NCBI.                                             ------------------------------------------------------------ 57310
human.FAM230J-LOC105372942-GGTLC5P.NCBI.rev.compl                                  ------------------------------------------------------------ 46364
human.FAM230A.ncRNA.18487127.18500594-GGTLC3.18516335.18518165.rev.compl.REF       ------------------------------------------------------------ 31039

chimp.LOC112206744-LOC107973052-GGT2.revcompl                                      ------------------------------------------------------------ 61762
chimp54546-135457.revcompl.Pan.troglodytes.clone.rp43-41g5.GenBank:AC099533.36     ------------------------------------------------------------ 69849
human.FAM30B-LOC105372935-GGT2.NCBI.                                               gcatctcccagcagacttctcatacctcagtggccaaaaccatgtctaaactacgggcct 96084
human.FAM230E-LOC105377182-GGT3P.NCBI.                                             ------------------------------------------------------------ 57310
human.FAM230J-LOC105372942-GGTLC5P.NCBI.rev.compl                                  ------------------------------------------------------------ 46364
human.FAM230A.ncRNA.18487127.18500594-GGTLC3.18516335.18518165.rev.compl.REF       ------------------------------------------------------------ 31039

chimp.LOC112206744-LOC107973052-GGT2.revcompl                                      ------------------------------------------------------------ 61762
chimp54546-135457.revcompl.Pan.troglodytes.clone.rp43-41g5.GenBank:AC099533.36     ------------------------------------------------------------ 69849
human.FAM30B-LOC105372935-GGT2.NCBI.                                               ccaggatgggctcagctcaatcagcacataccctagagctggaggcagggtcacctttcc 96144
human.FAM230E-LOC105377182-GGT3P.NCBI.                                             ------------------------------------------------------------ 57310
human.FAM230J-LOC105372942-GGTLC5P.NCBI.rev.compl                                  ------------------------------------------------------------ 46364
human.FAM230A.ncRNA.18487127.18500594-GGTLC3.18516335.18518165.rev.compl.REF       ------------------------------------------------------------ 31039

chimp.LOC112206744-LOC107973052-GGT2.revcompl                                      ------------------------------------------------------------ 61762
chimp54546-135457.revcompl.Pan.troglodytes.clone.rp43-41g5.GenBank:AC099533.36     ------------------------------------------------------------ 69849
human.FAM30B-LOC105372935-GGT2.NCBI.                                               ttgagtggcgtgtgggagaggtgtggcatccaacccaactgggtcctgccagcaggaaga 96204
human.FAM230E-LOC105377182-GGT3P.NCBI.                                             ------------------------------------------------------------ 57310
human.FAM230J-LOC105372942-GGTLC5P.NCBI.rev.compl                                  ------------------------------------------------------------ 46364
human.FAM230A.ncRNA.18487127.18500594-GGTLC3.18516335.18518165.rev.compl.REF       ------------------------------------------------------------ 31039

chimp.LOC112206744-LOC107973052-GGT2.revcompl                                      ------------------------------------------------------------ 61762
chimp54546-135457.revcompl.Pan.troglodytes.clone.rp43-41g5.GenBank:AC099533.36     ------------------------------------------------------------ 69849
human.FAM30B-LOC105372935-GGT2.NCBI.                                               gaaaagaagagccactgtaggtaaccaatcacgtctaccacaggcagggaccaataccac 96264
human.FAM230E-LOC105377182-GGT3P.NCBI.                                             ------------------------------------------------------------ 57310
human.FAM230J-LOC105372942-GGTLC5P.NCBI.rev.compl                                  ------------------------------------------------------------ 46364
human.FAM230A.ncRNA.18487127.18500594-GGTLC3.18516335.18518165.rev.compl.REF       ------------------------------------------------------------ 31039

chimp.LOC112206744-LOC107973052-GGT2.revcompl                                      ------------------------------------------------------------ 61762
chimp54546-135457.revcompl.Pan.troglodytes.clone.rp43-41g5.GenBank:AC099533.36     ------------------------------------------------------------ 69849
human.FAM30B-LOC105372935-GGT2.NCBI.                                               atatctgtgtttcccctcacttgccacccccatcctgtgctggcagatgaggatgctcaa 96324
human.FAM230E-LOC105377182-GGT3P.NCBI.                                             ------------------------------------------------------------ 57310
human.FAM230J-LOC105372942-GGTLC5P.NCBI.rev.compl                                  ------------------------------------------------------------ 46364
human.FAM230A.ncRNA.18487127.18500594-GGTLC3.18516335.18518165.rev.compl.REF       ------------------------------------------------------------ 31039

chimp.LOC112206744-LOC107973052-GGT2.revcompl                                      ------------------------------------------------------------ 61762
chimp54546-135457.revcompl.Pan.troglodytes.clone.rp43-41g5.GenBank:AC099533.36     ------------------------------------------------------------ 69849
human.FAM30B-LOC105372935-GGT2.NCBI.                                               caaatagctgtggaactacaaattatcctgtgggtgtgagaaaagagtgtgcattataat 96384
human.FAM230E-LOC105377182-GGT3P.NCBI.                                             ------------------------------------------------------------ 57310
human.FAM230J-LOC105372942-GGTLC5P.NCBI.rev.compl                                  ------------------------------------------------------------ 46364
human.FAM230A.ncRNA.18487127.18500594-GGTLC3.18516335.18518165.rev.compl.REF       ------------------------------------------------------------ 31039

chimp.LOC112206744-LOC107973052-GGT2.revcompl                                      ------------------------------------------------------------ 61762
chimp54546-135457.revcompl.Pan.troglodytes.clone.rp43-41g5.GenBank:AC099533.36     ------------------------------------------------------------ 69849
human.FAM30B-LOC105372935-GGT2.NCBI.                                               agggtggtacaattggacatgtgtgcacggaaaatggaaaaacagccatgaaatccaaac 96444
human.FAM230E-LOC105377182-GGT3P.NCBI.                                             ------------------------------------------------------------ 57310
human.FAM230J-LOC105372942-GGTLC5P.NCBI.rev.compl                                  ------------------------------------------------------------ 46364
human.FAM230A.ncRNA.18487127.18500594-GGTLC3.18516335.18518165.rev.compl.REF       ------------------------------------------------------------ 31039

chimp.LOC112206744-LOC107973052-GGT2.revcompl                                      ------------------------------------------------------------ 61762
chimp54546-135457.revcompl.Pan.troglodytes.clone.rp43-41g5.GenBank:AC099533.36     ------------------------------------------------------------ 69849
human.FAM30B-LOC105372935-GGT2.NCBI.                                               agatgcaaaaacgttcttaacacagaaggtttgggaagctctgatctaagccctagcctg 96504
human.FAM230E-LOC105377182-GGT3P.NCBI.                                             ------------------------------------------------------------ 57310
human.FAM230J-LOC105372942-GGTLC5P.NCBI.rev.compl                                  ------------------------------------------------------------ 46364
human.FAM230A.ncRNA.18487127.18500594-GGTLC3.18516335.18518165.rev.compl.REF       ------------------------------------------------------------ 31039

chimp.LOC112206744-LOC107973052-GGT2.revcompl                                      ------------------------------------------------------------ 61762
chimp54546-135457.revcompl.Pan.troglodytes.clone.rp43-41g5.GenBank:AC099533.36     ------------------------------------------------------------ 69849
human.FAM30B-LOC105372935-GGT2.NCBI.                                               ggccctactcccagttccacagatagggagcaggccccaggccagagtgctgggcttctc 96564
human.FAM230E-LOC105377182-GGT3P.NCBI.                                             ------------------------------------------------------------ 57310
human.FAM230J-LOC105372942-GGTLC5P.NCBI.rev.compl                                  ------------------------------------------------------------ 46364
human.FAM230A.ncRNA.18487127.18500594-GGTLC3.18516335.18518165.rev.compl.REF       ------------------------------------------------------------ 31039

chimp.LOC112206744-LOC107973052-GGT2.revcompl                                      ------------------------------------------------------------ 61762
chimp54546-135457.revcompl.Pan.troglodytes.clone.rp43-41g5.GenBank:AC099533.36     ------------------------------------------------------------ 69849
human.FAM30B-LOC105372935-GGT2.NCBI.                                               ccagcaggatccagctggggctttctcctcagttcaacccttgcctctgaggaccagtgg 96624
human.FAM230E-LOC105377182-GGT3P.NCBI.                                             ------------------------------------------------------------ 57310
human.FAM230J-LOC105372942-GGTLC5P.NCBI.rev.compl                                  ------------------------------------------------------------ 46364
human.FAM230A.ncRNA.18487127.18500594-GGTLC3.18516335.18518165.rev.compl.REF       ------------------------------------------------------------ 31039

chimp.LOC112206744-LOC107973052-GGT2.revcompl                                      ------------------------------------------------------------ 61762
chimp54546-135457.revcompl.Pan.troglodytes.clone.rp43-41g5.GenBank:AC099533.36     ------------------------------------------------------------ 69849
human.FAM30B-LOC105372935-GGT2.NCBI.                                               tctacagtccagcccaccacccttcagacagaagttccaggcacaaaaaggaagctcaca 96684
human.FAM230E-LOC105377182-GGT3P.NCBI.                                             ------------------------------------------------------------ 57310
human.FAM230J-LOC105372942-GGTLC5P.NCBI.rev.compl                                  ------------------------------------------------------------ 46364
human.FAM230A.ncRNA.18487127.18500594-GGTLC3.18516335.18518165.rev.compl.REF       ------------------------------------------------------------ 31039

chimp.LOC112206744-LOC107973052-GGT2.revcompl                                      ------------------------------------------------------------ 61762
chimp54546-135457.revcompl.Pan.troglodytes.clone.rp43-41g5.GenBank:AC099533.36     ------------------------------------------------------------ 69849
human.FAM30B-LOC105372935-GGT2.NCBI.                                               ataaggaacgcccatgcctagccataactgagcgcatcacagactcacacaaatctgtgg 96744
human.FAM230E-LOC105377182-GGT3P.NCBI.                                             ------------------------------------------------------------ 57310
human.FAM230J-LOC105372942-GGTLC5P.NCBI.rev.compl                                  ------------------------------------------------------------ 46364
human.FAM230A.ncRNA.18487127.18500594-GGTLC3.18516335.18518165.rev.compl.REF       ------------------------------------------------------------ 31039

chimp.LOC112206744-LOC107973052-GGT2.revcompl                                      ------------------------------------------------------------ 61762
chimp54546-135457.revcompl.Pan.troglodytes.clone.rp43-41g5.GenBank:AC099533.36     ------------------------------------------------------------ 69849
human.FAM30B-LOC105372935-GGT2.NCBI.                                               aatgagtgcatgatctcatgagctctggtctggaaaagtaagtctccctacactaggact 96804
human.FAM230E-LOC105377182-GGT3P.NCBI.                                             ------------------------------------------------------------ 57310
human.FAM230J-LOC105372942-GGTLC5P.NCBI.rev.compl                                  ------------------------------------------------------------ 46364
human.FAM230A.ncRNA.18487127.18500594-GGTLC3.18516335.18518165.rev.compl.REF       ------------------------------------------------------------ 31039

chimp.LOC112206744-LOC107973052-GGT2.revcompl                                      ------------------------------------------------------------ 61762
chimp54546-135457.revcompl.Pan.troglodytes.clone.rp43-41g5.GenBank:AC099533.36     ------------------------------------------------------------ 69849
human.FAM30B-LOC105372935-GGT2.NCBI.                                               taaagggaaggcttctttggctatgccaatatctttcctgccactttcttcttcaggcca 96864
human.FAM230E-LOC105377182-GGT3P.NCBI.                                             ------------------------------------------------------------ 57310
human.FAM230J-LOC105372942-GGTLC5P.NCBI.rev.compl                                  ------------------------------------------------------------ 46364
human.FAM230A.ncRNA.18487127.18500594-GGTLC3.18516335.18518165.rev.compl.REF       ------------------------------------------------------------ 31039

chimp.LOC112206744-LOC107973052-GGT2.revcompl                                      ------------------------------------------------------------ 61762
chimp54546-135457.revcompl.Pan.troglodytes.clone.rp43-41g5.GenBank:AC099533.36     ------------------------------------------------------------ 69849
human.FAM30B-LOC105372935-GGT2.NCBI.                                               aacagctcctatccctgccttccatcttagctttccagcatactcttcccaggttctttg 96924
human.FAM230E-LOC105377182-GGT3P.NCBI.                                             ------------------------------------------------------------ 57310
human.FAM230J-LOC105372942-GGTLC5P.NCBI.rev.compl                                  ------------------------------------------------------------ 46364
human.FAM230A.ncRNA.18487127.18500594-GGTLC3.18516335.18518165.rev.compl.REF       ------------------------------------------------------------ 31039

chimp.LOC112206744-LOC107973052-GGT2.revcompl                                      ------------------------------------------------------------ 61762
chimp54546-135457.revcompl.Pan.troglodytes.clone.rp43-41g5.GenBank:AC099533.36     ------------------------------------------------------------ 69849
human.FAM30B-LOC105372935-GGT2.NCBI.                                               ggctactgacaagcactccagacccaaatacacatgctcccacctgaggctgggaaatgt 96984
human.FAM230E-LOC105377182-GGT3P.NCBI.                                             ------------------------------------------------------------ 57310
human.FAM230J-LOC105372942-GGTLC5P.NCBI.rev.compl                                  ------------------------------------------------------------ 46364
```

```
human.FAM230A.ncRNA.18487127.18500594-GGTLC3.18516335.18518165.rev.compl.REF        ------------------------------------------------------------ 31039

chimp.LOC112206744-LOC107973052-GGT2.revcompl                                        ------------------------------------------------------------ 61762
chimp54546-135457.revcompl.Pan.troglodytes.clone.rp43-41g5.GenBank:AC099533.36      ------------------------------------------------------------ 69849
human.FAM30B-LOC105372935-GGT2.NCBI.                                                 accaagaactgcatagaaatatactggacaaaaggcactccagtctcacagatctccag 97044
human.FAM230E-LOC105377182-GGT3P.NCBI.                                               ------------------------------------------------------------ 57310
human.FAM230J-LOC105372942-GGTLC5P.NCBI.rev.compl                                    ------------------------------------------------------------ 46364
human.FAM230A.ncRNA.18487127.18500594-GGTLC3.18516335.18518165.rev.compl.REF        ------------------------------------------------------------ 31039

chimp.LOC112206744-LOC107973052-GGT2.revcompl                                        ------------------------------------------------------------ 61762
chimp54546-135457.revcompl.Pan.troglodytes.clone.rp43-41g5.GenBank:AC099533.36      ------------------------------------------------------------ 69849
human.FAM30B-LOC105372935-GGT2.NCBI.                                                 tcccacagtctatatcaccagttttagctgtttgaagttttagaattccttttcaaaacc 97104
human.FAM230E-LOC105377182-GGT3P.NCBI.                                               ------------------------------------------------------------ 57310
human.FAM230J-LOC105372942-GGTLC5P.NCBI.rev.compl                                    ------------------------------------------------------------ 46364
human.FAM230A.ncRNA.18487127.18500594-GGTLC3.18516335.18518165.rev.compl.REF        ------------------------------------------------------------ 31039

chimp.LOC112206744-LOC107973052-GGT2.revcompl                                        ------------------------------------------------------------ 61762
chimp54546-135457.revcompl.Pan.troglodytes.clone.rp43-41g5.GenBank:AC099533.36      ------------------------------------------------------------ 69849
human.FAM30B-LOC105372935-GGT2.NCBI.                                                 atagctagcatttctttggctagagaaaactgtgtgaaatgcagcctaccataggatcac 97164
human.FAM230E-LOC105377182-GGT3P.NCBI.                                               ------------------------------------------------------------ 57310
human.FAM230J-LOC105372942-GGTLC5P.NCBI.rev.compl                                    ------------------------------------------------------------ 46364
human.FAM230A.ncRNA.18487127.18500594-GGTLC3.18516335.18518165.rev.compl.REF        ------------------------------------------------------------ 31039

chimp.LOC112206744-LOC107973052-GGT2.revcompl                                        ------------------------------------------------------------ 61762
chimp54546-135457.revcompl.Pan.troglodytes.clone.rp43-41g5.GenBank:AC099533.36      ------------------------------------------------------------ 69849
human.FAM30B-LOC105372935-GGT2.NCBI.                                                 tgattacaacatgcattcaacattcaatacatattgactgagcaactattaggttgaact 97224
human.FAM230E-LOC105377182-GGT3P.NCBI.                                               ------------------------------------------------------------ 57310
human.FAM230J-LOC105372942-GGTLC5P.NCBI.rev.compl                                    ------------------------------------------------------------ 46364
human.FAM230A.ncRNA.18487127.18500594-GGTLC3.18516335.18518165.rev.compl.REF        ------------------------------------------------------------ 31039

chimp.LOC112206744-LOC107973052-GGT2.revcompl                                        ------------------------------------------------------------ 61762
chimp54546-135457.revcompl.Pan.troglodytes.clone.rp43-41g5.GenBank:AC099533.36      ------------------------------------------------------------ 69849
human.FAM30B-LOC105372935-GGT2.NCBI.                                                 atgtgacattattgacattcgactgcttctgacctatcaaaatggcaataaagctcaact 97284
human.FAM230E-LOC105377182-GGT3P.NCBI.                                               ------------------------------------------------------------ 57310
human.FAM230J-LOC105372942-GGTLC5P.NCBI.rev.compl                                    ------------------------------------------------------------ 46364
human.FAM230A.ncRNA.18487127.18500594-GGTLC3.18516335.18518165.rev.compl.REF        ------------------------------------------------------------ 31039

chimp.LOC112206744-LOC107973052-GGT2.revcompl                                        ------------------------------------------------------------ 61762
chimp54546-135457.revcompl.Pan.troglodytes.clone.rp43-41g5.GenBank:AC099533.36      ------------------------------------------------------------ 69849
human.FAM30B-LOC105372935-GGT2.NCBI.                                                 taatatttactctatgccaggcagtaggctaaggaaggatttgggacaccatgacaaaca 97344
human.FAM230E-LOC105377182-GGT3P.NCBI.                                               ------------------------------------------------------------ 57310
human.FAM230J-LOC105372942-GGTLC5P.NCBI.rev.compl                                    ------------------------------------------------------------ 46364
human.FAM230A.ncRNA.18487127.18500594-GGTLC3.18516335.18518165.rev.compl.REF        ------------------------------------------------------------ 31039

chimp.LOC112206744-LOC107973052-GGT2.revcompl                                        ------------------------------------------------------------ 61762
chimp54546-135457.revcompl.Pan.troglodytes.clone.rp43-41g5.GenBank:AC099533.36      ------------------------------------------------------------ 69849
human.FAM30B-LOC105372935-GGT2.NCBI.                                                 acacagaagtaacccctatcttaatggaacttaaaatctagttgattaaaccattactta 97404
human.FAM230E-LOC105377182-GGT3P.NCBI.                                               ------------------------------------------------------------ 57310
human.FAM230J-LOC105372942-GGTLC5P.NCBI.rev.compl                                    ------------------------------------------------------------ 46364
human.FAM230A.ncRNA.18487127.18500594-GGTLC3.18516335.18518165.rev.compl.REF        ------------------------------------------------------------ 31039

chimp.LOC112206744-LOC107973052-GGT2.revcompl                                        ------------------------------------------------------------ 61762
chimp54546-135457.revcompl.Pan.troglodytes.clone.rp43-41g5.GenBank:AC099533.36      ------------------------------------------------------------ 69849
human.FAM30B-LOC105372935-GGT2.NCBI.                                                 caaaagtacaatctaacttcaggattgtaattaattataaatgcatttatttgtttactg 97464
human.FAM230E-LOC105377182-GGT3P.NCBI.                                               ------------------------------------------------------------ 57310
human.FAM230J-LOC105372942-GGTLC5P.NCBI.rev.compl                                    ------------------------------------------------------------ 46364
human.FAM230A.ncRNA.18487127.18500594-GGTLC3.18516335.18518165.rev.compl.REF        ------------------------------------------------------------ 31039

chimp.LOC112206744-LOC107973052-GGT2.revcompl                                        ------------------------------------------------------------ 61762
chimp54546-135457.revcompl.Pan.troglodytes.clone.rp43-41g5.GenBank:AC099533.36      ------------------------------------------------------------ 69849
human.FAM30B-LOC105372935-GGT2.NCBI.                                                 cctgacttctttgctagcctataaactacacaaagatgagggttatgtctcctccatttc 97524
human.FAM230E-LOC105377182-GGT3P.NCBI.                                               ------------------------------------------------------------ 57310
human.FAM230J-LOC105372942-GGTLC5P.NCBI.rev.compl                                    ------------------------------------------------------------ 46364
human.FAM230A.ncRNA.18487127.18500594-GGTLC3.18516335.18518165.rev.compl.REF        ------------------------------------------------------------ 31039

chimp.LOC112206744-LOC107973052-GGT2.revcompl                                        ------------------------------------------------------------ 61762
chimp54546-135457.revcompl.Pan.troglodytes.clone.rp43-41g5.GenBank:AC099533.36      ------------------------------------------------------------ 69849
human.FAM30B-LOC105372935-GGT2.NCBI.                                                 acacactggcaaagcatatagcagaatacatcatatatttgttaaatgtattgattagtg 97584
human.FAM230E-LOC105377182-GGT3P.NCBI.                                               ------------------------------------------------------------ 57310
human.FAM230J-LOC105372942-GGTLC5P.NCBI.rev.compl                                    ------------------------------------------------------------ 46364
human.FAM230A.ncRNA.18487127.18500594-GGTLC3.18516335.18518165.rev.compl.REF        ------------------------------------------------------------ 31039

chimp.LOC112206744-LOC107973052-GGT2.revcompl                                        ------------------------------------------------------------ 61762
chimp54546-135457.revcompl.Pan.troglodytes.clone.rp43-41g5.GenBank:AC099533.36      ------------------------------------------------------------ 69849
human.FAM30B-LOC105372935-GGT2.NCBI.                                                 ttacaacagacaattgcatgtgctatggaagcgtttggcagagagatctaatctaatctt 97644
human.FAM230E-LOC105377182-GGT3P.NCBI.                                               ------------------------------------------------------------ 57310
human.FAM230J-LOC105372942-GGTLC5P.NCBI.rev.compl                                    ------------------------------------------------------------ 46364
human.FAM230A.ncRNA.18487127.18500594-GGTLC3.18516335.18518165.rev.compl.REF        ------------------------------------------------------------ 31039

chimp.LOC112206744-LOC107973052-GGT2.revcompl                                        ------------------------------------------------------------ 61762
chimp54546-135457.revcompl.Pan.troglodytes.clone.rp43-41g5.GenBank:AC099533.36      ------------------------------------------------------------ 69849
human.FAM30B-LOC105372935-GGT2.NCBI.                                                 ggaaggtgaggatcctggccatcactaacttggaatttctgatactgatgatataaggtt 97704
human.FAM230E-LOC105377182-GGT3P.NCBI.                                               ------------------------------------------------------------ 57310
human.FAM230J-LOC105372942-GGTLC5P.NCBI.rev.compl                                    ------------------------------------------------------------ 46364
human.FAM230A.ncRNA.18487127.18500594-GGTLC3.18516335.18518165.rev.compl.REF        ------------------------------------------------------------ 31039

chimp.LOC112206744-LOC107973052-GGT2.revcompl                                        ------------------------------------------------------------ 61762
chimp54546-135457.revcompl.Pan.troglodytes.clone.rp43-41g5.GenBank:AC099533.36      ------------------------------------------------------------ 69849
human.FAM30B-LOC105372935-GGT2.NCBI.                                                 gagttatgtcttcgtgaaacaggtctctatacagcatataaacaactaccagcatgcaca 97764
human.FAM230E-LOC105377182-GGT3P.NCBI.                                               ------------------------------------------------------------ 57310
human.FAM230J-LOC105372942-GGTLC5P.NCBI.rev.compl                                    ------------------------------------------------------------ 46364
human.FAM230A.ncRNA.18487127.18500594-GGTLC3.18516335.18518165.rev.compl.REF        ------------------------------------------------------------ 31039

chimp.LOC112206744-LOC107973052-GGT2.revcompl                                        ------------------------------------------------------------ 61762
chimp54546-135457.revcompl.Pan.troglodytes.clone.rp43-41g5.GenBank:AC099533.36      ------------------------------------------------------------ 69849
human.FAM30B-LOC105372935-GGT2.NCBI.                                                 ggggttccgtagggactatttaaaatacttaagggaacaaacggttttcaagctcagtct 97824
human.FAM230E-LOC105377182-GGT3P.NCBI.                                               ------------------------------------------------------------ 57310
human.FAM230J-LOC105372942-GGTLC5P.NCBI.rev.compl                                    ------------------------------------------------------------ 46364
human.FAM230A.ncRNA.18487127.18500594-GGTLC3.18516335.18518165.rev.compl.REF        ------------------------------------------------------------ 31039

chimp.LOC112206744-LOC107973052-GGT2.revcompl                                        ------------------------------------------------------------ 61762
chimp54546-135457.revcompl.Pan.troglodytes.clone.rp43-41g5.GenBank:AC099533.36      ------------------------------------------------------------ 69849
human.FAM30B-LOC105372935-GGT2.NCBI.                                                 agtatttcagaaattccatttacataaacaattggtgcactaattgcaaatcattcttat 97884
human.FAM230E-LOC105377182-GGT3P.NCBI.                                               ------------------------------------------------------------ 57310
human.FAM230J-LOC105372942-GGTLC5P.NCBI.rev.compl                                    ------------------------------------------------------------ 46364
human.FAM230A.ncRNA.18487127.18500594-GGTLC3.18516335.18518165.rev.compl.REF        ------------------------------------------------------------ 31039

chimp.LOC112206744-LOC107973052-GGT2.revcompl                                        ------------------------------------------------------------ 61762
chimp54546-135457.revcompl.Pan.troglodytes.clone.rp43-41g5.GenBank:AC099533.36      ------------------------------------------------------------ 69849
human.FAM30B-LOC105372935-GGT2.NCBI.                                                 ttctctcctaaagcaggatcccctaaggatctttattaggcattctattagcctagtttg 97944
human.FAM230E-LOC105377182-GGT3P.NCBI.                                               ------------------------------------------------------------ 57310
human.FAM230J-LOC105372942-GGTLC5P.NCBI.rev.compl                                    ------------------------------------------------------------ 46364
human.FAM230A.ncRNA.18487127.18500594-GGTLC3.18516335.18518165.rev.compl.REF        ------------------------------------------------------------ 31039

chimp.LOC112206744-LOC107973052-GGT2.revcompl                                        ------------------------------------------------------------ 61762
chimp54546-135457.revcompl.Pan.troglodytes.clone.rp43-41g5.GenBank:AC099533.36      -------------------------------------TGTTGCAAGCTCTAAAATAATCC 69872
human.FAM30B-LOC105372935-GGT2.NCBI.                                                 agttacagtatttacttaaaagtaataacagtgaagttcttgaaaaatatttggtaattc 98004
human.FAM230E-LOC105377182-GGT3P.NCBI.                                               ------------------------------------------------------------ 57310
human.FAM230J-LOC105372942-GGTLC5P.NCBI.rev.compl                                    ------------------------------------------------------------ 46364
human.FAM230A.ncRNA.18487127.18500594-GGTLC3.18516335.18518165.rev.compl.REF        ------------------------------------------------------------ 31039

chimp.LOC112206744-LOC107973052-GGT2.revcompl                                        ------------------------------------------------------------ 61762
chimp54546-135457.revcompl.Pan.troglodytes.clone.rp43-41g5.GenBank:AC099533.36      AGTCATCTTTTAAAATAACTCTAAGACTGCACAGTTATGAAACTAATAGAGAAGGAGGAA 69932
human.FAM30B-LOC105372935-GGT2.NCBI.                                                 agacattttactaattcaaactaatcttcctcattagtctacgttgt--------tagag 98056
human.FAM230E-LOC105377182-GGT3P.NCBI.                                               ------------------------------------------------------------ 57310
human.FAM230J-LOC105372942-GGTLC5P.NCBI.rev.compl                                    ------------------------------------------------------------ 46364
human.FAM230A.ncRNA.18487127.18500594-GGTLC3.18516335.18518165.rev.compl.REF        ------------------------------------------------------------ 31039
```

```
chimp.LOC112206744-LOC107973052-GGT2.revcompl                                     ------------------------------------------------------------ 61762
chimp54546-135457.revcompl.Pan.troglodytes.clone.rp43-41g5.GenBank:AC099533.36    ATTAAATCATAAAAATAATAAATCCAAAACAAGATGTGAGAGGAGATAAGAAGAAATAGA 69992
human.FAM30B-LOC105372935-GGT2.NCBI.                                              gtttaaccattgtattaagagaaagtatcaaaggcttcac-ttagagtaatagtcaccaa 98115
human.FAM230E-LOC105377182-GGT3P.NCBI.                                            ------------------------------------------------------------ 57310
human.FAM230J-LOC105372942-GGTLC5P.NCBI.rev.compl                                 ------------------------------------------------------------ 46364
human.FAM230A.ncRNA.18487127.18500594-GGTLC3.18516335.18518165.rev.compl.REF      ------------------------------------------------------------ 31039

chimp.LOC112206744-LOC107973052-GGT2.revcompl                                     ------------------------------------------------------------ 61762
chimp54546-135457.revcompl.Pan.troglodytes.clone.rp43-41g5.GenBank:AC099533.36    ATAGGCATGGAAAACAAATTGGTGGTGGGTTTCAACCCAAATAAATCATTAGTTACATTT 70052
human.FAM30B-LOC105372935-GGT2.NCBI.                                              atttttttgcaagacaccttggtagcaggaacctgagcta--acatcaggaggtaggtag 98173
human.FAM230E-LOC105377182-GGT3P.NCBI.                                            ------------------------------------------------------------ 57310
human.FAM230J-LOC105372942-GGTLC5P.NCBI.rev.compl                                 ------------------------------------------------------------ 46364
human.FAM230A.ncRNA.18487127.18500594-GGTLC3.18516335.18518165.rev.compl.REF      ------------------------------------------------------------ 31039

chimp.LOC112206744-LOC107973052-GGT2.revcompl                                     ------------------------------------------------------------ 61762
chimp54546-135457.revcompl.Pan.troglodytes.clone.rp43-41g5.GenBank:AC099533.36    AAAAGGACAATAAAAGTTAAAATAATTGAAAATAAAGTAAAACCCAACTAATGCC----- 70107
human.FAM30B-LOC105372935-GGT2.NCBI.                                              aaatagaaaataaaggaatgacaatttgaagagcaactgcaatccaatttagtccaacag 98233
human.FAM230E-LOC105377182-GGT3P.NCBI.                                            ------------------------------------------------------------ 57310
human.FAM230J-LOC105372942-GGTLC5P.NCBI.rev.compl                                 ------------------------------------------------------------ 46364
human.FAM230A.ncRNA.18487127.18500594-GGTLC3.18516335.18518165.rev.compl.REF      ------------------------------------------------------------ 31039

chimp.LOC112206744-LOC107973052-GGT2.revcompl                                     ------------------------------------------------------------ 61762
chimp54546-135457.revcompl.Pan.troglodytes.clone.rp43-41g5.GenBank:AC099533.36    ------------------------------------------------------------ 70107
human.FAM30B-LOC105372935-GGT2.NCBI.                                              atgacacacatatgaagtcacattcaaattataagcagagcacaaaaatattttagatcc 98293
human.FAM230E-LOC105377182-GGT3P.NCBI.                                            ------------------------------------------------------------ 57310
human.FAM230J-LOC105372942-GGTLC5P.NCBI.rev.compl                                 ------------------------------------------------------------ 46364
human.FAM230A.ncRNA.18487127.18500594-GGTLC3.18516335.18518165.rev.compl.REF      ------------------------------------------------------------ 31039

chimp.LOC112206744-LOC107973052-GGT2.revcompl                                     ------------------------------------------------------------ 61762
chimp54546-135457.revcompl.Pan.troglodytes.clone.rp43-41g5.GenBank:AC099533.36    ------------------------------------------------------------ 70107
human.FAM30B-LOC105372935-GGT2.NCBI.                                              ttatcttttgattaacctggtatcattcagggaaaaatgtcagggaacaccactgaccag 98353
human.FAM230E-LOC105377182-GGT3P.NCBI.                                            ------------------------------------------------------------ 57310
human.FAM230J-LOC105372942-GGTLC5P.NCBI.rev.compl                                 ------------------------------------------------------------ 46364
human.FAM230A.ncRNA.18487127.18500594-GGTLC3.18516335.18518165.rev.compl.REF      ------------------------------------------------------------ 31039

chimp.LOC112206744-LOC107973052-GGT2.revcompl                                     ------------------------------------------------------------ 61762
chimp54546-135457.revcompl.Pan.troglodytes.clone.rp43-41g5.GenBank:AC099533.36    ------------------------------------------------------------ 70107
human.FAM30B-LOC105372935-GGT2.NCBI.                                              cagtcctctcttatctattttgcagatcttatctatttttctccctgagaaccagccctg 98413
human.FAM230E-LOC105377182-GGT3P.NCBI.                                            ------------------------------------------------------------ 57310
human.FAM230J-LOC105372942-GGTLC5P.NCBI.rev.compl                                 ------------------------------------------------------------ 46364
human.FAM230A.ncRNA.18487127.18500594-GGTLC3.18516335.18518165.rev.compl.REF      ------------------------------------------------------------ 31039

chimp.LOC112206744-LOC107973052-GGT2.revcompl                                     ------------------------------------------------------------ 61762
chimp54546-135457.revcompl.Pan.troglodytes.clone.rp43-41g5.GenBank:AC099533.36    ------------------------------------------------------------ 70107
human.FAM30B-LOC105372935-GGT2.NCBI.                                              gacacaaactgcataaacatcaatgagcctagacagggtcttcatgtgtgtttggcctgc 98473
human.FAM230E-LOC105377182-GGT3P.NCBI.                                            ------------------------------------------------------------ 57310
human.FAM230J-LOC105372942-GGTLC5P.NCBI.rev.compl                                 ------------------------------------------------------------ 46364
human.FAM230A.ncRNA.18487127.18500594-GGTLC3.18516335.18518165.rev.compl.REF      ------------------------------------------------------------ 31039

chimp.LOC112206744-LOC107973052-GGT2.revcompl                                     ------------------------------------------------------------ 61762
chimp54546-135457.revcompl.Pan.troglodytes.clone.rp43-41g5.GenBank:AC099533.36    ------------------------------------------------------------ 70107
human.FAM30B-LOC105372935-GGT2.NCBI.                                              cattgttctctgtaccagtacggagatttagtaacacactggtcacattcatggggcatc 98533
human.FAM230E-LOC105377182-GGT3P.NCBI.                                            ------------------------------------------------------------ 57310
human.FAM230J-LOC105372942-GGTLC5P.NCBI.rev.compl                                 ------------------------------------------------------------ 46364
human.FAM230A.ncRNA.18487127.18500594-GGTLC3.18516335.18518165.rev.compl.REF      ------------------------------------------------------------ 31039

chimp.LOC112206744-LOC107973052-GGT2.revcompl                                     ------------------------------------------------------------ 61762
chimp54546-135457.revcompl.Pan.troglodytes.clone.rp43-41g5.GenBank:AC099533.36    ------------------------------------------------------------ 70107
human.FAM30B-LOC105372935-GGT2.NCBI.                                              ttgaactctcttgacatggtaaatgaacagcaggtcctgttcccatgccaaaagcaaaca 98593
human.FAM230E-LOC105377182-GGT3P.NCBI.                                            ------------------------------------------------------------ 57310
human.FAM230J-LOC105372942-GGTLC5P.NCBI.rev.compl                                 ------------------------------------------------------------ 46364
human.FAM230A.ncRNA.18487127.18500594-GGTLC3.18516335.18518165.rev.compl.REF      ------------------------------------------------------------ 31039

chimp.LOC112206744-LOC107973052-GGT2.revcompl                                     ------------------------------------------------------------ 61762
chimp54546-135457.revcompl.Pan.troglodytes.clone.rp43-41g5.GenBank:AC099533.36    ------------------------------------------------------------ 70107
human.FAM30B-LOC105372935-GGT2.NCBI.                                              aaaacctgtcaccagcaaggaagacacccagtctctccaaaggatgccaaagtccacatc 98653
human.FAM230E-LOC105377182-GGT3P.NCBI.                                            ------------------------------------------------------------ 57310
human.FAM230J-LOC105372942-GGTLC5P.NCBI.rev.compl                                 ------------------------------------------------------------ 46364
human.FAM230A.ncRNA.18487127.18500594-GGTLC3.18516335.18518165.rev.compl.REF      ------------------------------------------------------------ 31039

chimp.LOC112206744-LOC107973052-GGT2.revcompl                                     ------------------------------------------------------------ 61762
chimp54546-135457.revcompl.Pan.troglodytes.clone.rp43-41g5.GenBank:AC099533.36    ------------------------------------------------------------ 70107
human.FAM30B-LOC105372935-GGT2.NCBI.                                              cagcctcctttgcagagcctcacaaatctctcggatgcaggcaggatgatcacgggctag 98713
human.FAM230E-LOC105377182-GGT3P.NCBI.                                            ------------------------------------------------------------ 57310
human.FAM230J-LOC105372942-GGTLC5P.NCBI.rev.compl                                 ------------------------------------------------------------ 46364
human.FAM230A.ncRNA.18487127.18500594-GGTLC3.18516335.18518165.rev.compl.REF      ------------------------------------------------------------ 31039

chimp.LOC112206744-LOC107973052-GGT2.revcompl                                     ------------------------------------------------------------ 61762
chimp54546-135457.revcompl.Pan.troglodytes.clone.rp43-41g5.GenBank:AC099533.36    ------------------------------------------------------------ 70107
human.FAM30B-LOC105372935-GGT2.NCBI.                                              ggaagattatgctctgaatgtcttaagagattagttacttaaaatcctccacatgacttc 98773
human.FAM230E-LOC105377182-GGT3P.NCBI.                                            ------------------------------------------------------------ 57310
human.FAM230J-LOC105372942-GGTLC5P.NCBI.rev.compl                                 ------------------------------------------------------------ 46364
human.FAM230A.ncRNA.18487127.18500594-GGTLC3.18516335.18518165.rev.compl.REF      ------------------------------------------------------------ 31039

chimp.LOC112206744-LOC107973052-GGT2.revcompl                                     ------------------------------------------------------------ 61762
chimp54546-135457.revcompl.Pan.troglodytes.clone.rp43-41g5.GenBank:AC099533.36    ------------------------------------------------------------ 70107
human.FAM30B-LOC105372935-GGT2.NCBI.                                              agtcatccccgttgcagatcacagagtaaaaagcctacctgattaaagctgaatgcctat 98833
human.FAM230E-LOC105377182-GGT3P.NCBI.                                            ------------------------------------------------------------ 57310
human.FAM230J-LOC105372942-GGTLC5P.NCBI.rev.compl                                 ------------------------------------------------------------ 46364
human.FAM230A.ncRNA.18487127.18500594-GGTLC3.18516335.18518165.rev.compl.REF      ------------------------------------------------------------ 31039

chimp.LOC112206744-LOC107973052-GGT2.revcompl                                     ------------------------------------------------------------ 61762
chimp54546-135457.revcompl.Pan.troglodytes.clone.rp43-41g5.GenBank:AC099533.36    ------------------------------------------------------------ 70107
human.FAM30B-LOC105372935-GGT2.NCBI.                                              acatttgcaaaagggaaaatatgcagaagaaaacgagagtcagatactgatgggcatagc 98893
human.FAM230E-LOC105377182-GGT3P.NCBI.                                            ------------------------------------------------------------ 57310
human.FAM230J-LOC105372942-GGTLC5P.NCBI.rev.compl                                 ------------------------------------------------------------ 46364
human.FAM230A.ncRNA.18487127.18500594-GGTLC3.18516335.18518165.rev.compl.REF      ------------------------------------------------------------ 31039

chimp.LOC112206744-LOC107973052-GGT2.revcompl                                     ------------------------------------------------------------ 61762
chimp54546-135457.revcompl.Pan.troglodytes.clone.rp43-41g5.GenBank:AC099533.36    ------------------------------------------------------------ 70107
human.FAM30B-LOC105372935-GGT2.NCBI.                                              accagctctaggacatcagaggtgttgaataaatgatgacaaggaggcacacatagttct 98953
human.FAM230E-LOC105377182-GGT3P.NCBI.                                            ------------------------------------------------------------ 57310
human.FAM230J-LOC105372942-GGTLC5P.NCBI.rev.compl                                 ------------------------------------------------------------ 46364
human.FAM230A.ncRNA.18487127.18500594-GGTLC3.18516335.18518165.rev.compl.REF      ------------------------------------------------------------ 31039

chimp.LOC112206744-LOC107973052-GGT2.revcompl                                     ------------------------------------------------------------ 61762
chimp54546-135457.revcompl.Pan.troglodytes.clone.rp43-41g5.GenBank:AC099533.36    ------------------------------------------------------------ 70107
human.FAM30B-LOC105372935-GGT2.NCBI.                                              gatatattcaccataaataccacagtcctcaaggctctctcaattacttacttcccccca 99013
human.FAM230E-LOC105377182-GGT3P.NCBI.                                            ------------------------------------------------------------ 57310
human.FAM230J-LOC105372942-GGTLC5P.NCBI.rev.compl                                 ------------------------------------------------------------ 46364
human.FAM230A.ncRNA.18487127.18500594-GGTLC3.18516335.18518165.rev.compl.REF      ------------------------------------------------------------ 31039

chimp.LOC112206744-LOC107973052-GGT2.revcompl                                     ------------------------------------------------------------ 61762
chimp54546-135457.revcompl.Pan.troglodytes.clone.rp43-41g5.GenBank:AC099533.36    ------------------------------------------------------------ 70107
human.FAM30B-LOC105372935-GGT2.NCBI.                                              aatcagagactgaatctggccttatcttcaaatgaggcccagctctccccaaattccaaa 99073
human.FAM230E-LOC105377182-GGT3P.NCBI.                                            ------------------------------------------------------------ 57310
human.FAM230J-LOC105372942-GGTLC5P.NCBI.rev.compl                                 ------------------------------------------------------------ 46364
human.FAM230A.ncRNA.18487127.18500594-GGTLC3.18516335.18518165.rev.compl.REF      ------------------------------------------------------------ 31039

chimp.LOC112206744-LOC107973052-GGT2.revcompl                                     ------------------------------------------------------------ 61762
chimp54546-135457.revcompl.Pan.troglodytes.clone.rp43-41g5.GenBank:AC099533.36    ------------------------------------------------------------ 70107
human.FAM30B-LOC105372935-GGT2.NCBI.                                              tatgggaaattacagctcctctgtggtaggggaagagtgggaaggcgaatgtttgtatt 99133
human.FAM230E-LOC105377182-GGT3P.NCBI.                                            ------------------------------------------------------------ 57310
human.FAM230J-LOC105372942-GGTLC5P.NCBI.rev.compl                                 ------------------------------------------------------------ 46364
human.FAM230A.ncRNA.18487127.18500594-GGTLC3.18516335.18518165.rev.compl.REF      ------------------------------------------------------------ 31039
```

```
chimp.LOC112206744-LOC107973052-GGT2.revcompl                                  ------------------------------------------------------------ 61762
chimp54546-135457.revcompl.Pan.troglodytes.clone.rp43-41g5.GenBank:AC099533.36 ------------------------------------------------------------ 70107
human.FAM30B-LOC105372935-GGT2.NCBI.                                           gcataacgttttaattaaaacaaagatgacgtgcaccagagtctctttctcatctcctgt 99193
human.FAM230E-LOC105377182-GGT3P.NCBI.                                         ------------------------------------------------------------ 57310
human.FAM230J-LOC105372942-GGTLC5P.NCBI.rev.compl                              ------------------------------------------------------------ 46364
human.FAM230A.ncRNA.18487127.18500594-GGTLC3.18516335.18518165.rev.compl.REF   ------------------------------------------------------------ 31039

chimp.LOC112206744-LOC107973052-GGT2.revcompl                                  ------------------------------------------------------------ 61762
chimp54546-135457.revcompl.Pan.troglodytes.clone.rp43-41g5.GenBank:AC099533.36 ------------------------------------------------------------ 70107
human.FAM30B-LOC105372935-GGT2.NCBI.                                           taggggcgttttccagtactacttccaccctttacccacccaggtcaccatggtcttttt 99253
human.FAM230E-LOC105377182-GGT3P.NCBI.                                         ------------------------------------------------------------ 57310
human.FAM230J-LOC105372942-GGTLC5P.NCBI.rev.compl                              ------------------------------------------------------------ 46364
human.FAM230A.ncRNA.18487127.18500594-GGTLC3.18516335.18518165.rev.compl.REF   ------------------------------------------------------------ 31039

chimp.LOC112206744-LOC107973052-GGT2.revcompl                                  ------------------------------------------------------------ 61762
chimp54546-135457.revcompl.Pan.troglodytes.clone.rp43-41g5.GenBank:AC099533.36 ------------------------------------------------------------ 70107
human.FAM30B-LOC105372935-GGT2.NCBI.                                           cttccctctaggacaattcacctttaaaataagaggattgggcaagaaatctacaaagat 99313
human.FAM230E-LOC105377182-GGT3P.NCBI.                                         ------------------------------------------------------------ 57310
human.FAM230J-LOC105372942-GGTLC5P.NCBI.rev.compl                              ------------------------------------------------------------ 46364
human.FAM230A.ncRNA.18487127.18500594-GGTLC3.18516335.18518165.rev.compl.REF   ------------------------------------------------------------ 31039

chimp.LOC112206744-LOC107973052-GGT2.revcompl                                  ------------------------------------------------------------ 61762
chimp54546-135457.revcompl.Pan.troglodytes.clone.rp43-41g5.GenBank:AC099533.36 ------------------------------------------------------------ 70107
human.FAM30B-LOC105372935-GGT2.NCBI.                                           ttttccaaaatgaactcaggcctagaggcatattagttctctatcttgttgctcccttct 99373
human.FAM230E-LOC105377182-GGT3P.NCBI.                                         ------------------------------------------------------------ 57310
human.FAM230J-LOC105372942-GGTLC5P.NCBI.rev.compl                              ------------------------------------------------------------ 46364
human.FAM230A.ncRNA.18487127.18500594-GGTLC3.18516335.18518165.rev.compl.REF   ------------------------------------------------------------ 31039

chimp.LOC112206744-LOC107973052-GGT2.revcompl                                  ------------------------------------------------------------ 61762
chimp54546-135457.revcompl.Pan.troglodytes.clone.rp43-41g5.GenBank:AC099533.36 ------------------------------------------------------------ 70107
human.FAM30B-LOC105372935-GGT2.NCBI.                                           agaagtgcacagttcagagggatgcccagccacagaggtgccaaaacgcagagctggaac 99433
human.FAM230E-LOC105377182-GGT3P.NCBI.                                         ------------------------------------------------------------ 57310
human.FAM230J-LOC105372942-GGTLC5P.NCBI.rev.compl                              ------------------------------------------------------------ 46364
human.FAM230A.ncRNA.18487127.18500594-GGTLC3.18516335.18518165.rev.compl.REF   ------------------------------------------------------------ 31039

chimp.LOC112206744-LOC107973052-GGT2.revcompl                                  ------------------------------------------------------------ 61762
chimp54546-135457.revcompl.Pan.troglodytes.clone.rp43-41g5.GenBank:AC099533.36 ------------------------------------------------------------ 70107
human.FAM30B-LOC105372935-GGT2.NCBI.                                           caatcacctgctccaacagagacaaggtgaaagcactgggcagggtgctcagaatgtgct 99493
human.FAM230E-LOC105377182-GGT3P.NCBI.                                         ------------------------------------------------------------ 57310
human.FAM230J-LOC105372942-GGTLC5P.NCBI.rev.compl                              ------------------------------------------------------------ 46364
human.FAM230A.ncRNA.18487127.18500594-GGTLC3.18516335.18518165.rev.compl.REF   ------------------------------------------------------------ 31039

chimp.LOC112206744-LOC107973052-GGT2.revcompl                                  ------------------------------------------------------------ 61762
chimp54546-135457.revcompl.Pan.troglodytes.clone.rp43-41g5.GenBank:AC099533.36 ------------------------------------------------------------ 70107
human.FAM30B-LOC105372935-GGT2.NCBI.                                           cttgcttctttgtaatgaaatatcagaagacagattgtaacattgttcagcaagctattt 99553
human.FAM230E-LOC105377182-GGT3P.NCBI.                                         ------------------------------------------------------------ 57310
human.FAM230J-LOC105372942-GGTLC5P.NCBI.rev.compl                              ------------------------------------------------------------ 46364
human.FAM230A.ncRNA.18487127.18500594-GGTLC3.18516335.18518165.rev.compl.REF   ------------------------------------------------------------ 31039

chimp.LOC112206744-LOC107973052-GGT2.revcompl                                  ------------------------------------------------------------ 61762
chimp54546-135457.revcompl.Pan.troglodytes.clone.rp43-41g5.GenBank:AC099533.36 ------------------------------------------------------------ 70107
human.FAM30B-LOC105372935-GGT2.NCBI.                                           tttgtactgaaacatcagaaggcattatgaaactgttcagcaaatcatctctaaacaatg 99613
human.FAM230E-LOC105377182-GGT3P.NCBI.                                         ------------------------------------------------------------ 57310
human.FAM230J-LOC105372942-GGTLC5P.NCBI.rev.compl                              ------------------------------------------------------------ 46364
human.FAM230A.ncRNA.18487127.18500594-GGTLC3.18516335.18518165.rev.compl.REF   ------------------------------------------------------------ 31039

chimp.LOC112206744-LOC107973052-GGT2.revcompl                                  ------------------------------------------------------------ 61762
chimp54546-135457.revcompl.Pan.troglodytes.clone.rp43-41g5.GenBank:AC099533.36 ------------------------------------------------------------ 70107
human.FAM30B-LOC105372935-GGT2.NCBI.                                           tcataaagggttacctagtttggggtaagcccggaatgaatatctgcaccatcattcctc 99673
human.FAM230E-LOC105377182-GGT3P.NCBI.                                         ------------------------------------------------------------ 57310
human.FAM230J-LOC105372942-GGTLC5P.NCBI.rev.compl                              ------------------------------------------------------------ 46364
human.FAM230A.ncRNA.18487127.18500594-GGTLC3.18516335.18518165.rev.compl.REF   ------------------------------------------------------------ 31039

chimp.LOC112206744-LOC107973052-GGT2.revcompl                                  ------------------------------------------------------------ 61762
chimp54546-135457.revcompl.Pan.troglodytes.clone.rp43-41g5.GenBank:AC099533.36 ------------------------------------------------------------ 70107
human.FAM30B-LOC105372935-GGT2.NCBI.                                           ccgctaacaaacatgtccatctatgtgaagcataattcccaacacaccaccatctccaaa 99733
human.FAM230E-LOC105377182-GGT3P.NCBI.                                         ------------------------------------------------------------ 57310
human.FAM230J-LOC105372942-GGTLC5P.NCBI.rev.compl                              ------------------------------------------------------------ 46364
human.FAM230A.ncRNA.18487127.18500594-GGTLC3.18516335.18518165.rev.compl.REF   ------------------------------------------------------------ 31039

chimp.LOC112206744-LOC107973052-GGT2.revcompl                                  ------------------------------------------------------------ 61762
chimp54546-135457.revcompl.Pan.troglodytes.clone.rp43-41g5.GenBank:AC099533.36 ------------------------------------------------------------ 70107
human.FAM30B-LOC105372935-GGT2.NCBI.                                           ccaacaccaaaccactgccaaacaaagctgaagaggaaaaccagaaaggtctccgagaag 99793
human.FAM230E-LOC105377182-GGT3P.NCBI.                                         ------------------------------------------------------------ 57310
human.FAM230J-LOC105372942-GGTLC5P.NCBI.rev.compl                              ------------------------------------------------------------ 46364
human.FAM230A.ncRNA.18487127.18500594-GGTLC3.18516335.18518165.rev.compl.REF   ------------------------------------------------------------ 31039

chimp.LOC112206744-LOC107973052-GGT2.revcompl                                  ------------------------------------------------------------ 61762
chimp54546-135457.revcompl.Pan.troglodytes.clone.rp43-41g5.GenBank:AC099533.36 ------------------------------------------------------------ 70107
human.FAM30B-LOC105372935-GGT2.NCBI.                                           tgctgcttgcagggagaagaatggccctgcacagaggctcctgagggcagcaggtgcctg 99853
human.FAM230E-LOC105377182-GGT3P.NCBI.                                         ------------------------------------------------------------ 57310
human.FAM230J-LOC105372942-GGTLC5P.NCBI.rev.compl                              ------------------------------------------------------------ 46364
human.FAM230A.ncRNA.18487127.18500594-GGTLC3.18516335.18518165.rev.compl.REF   ------------------------------------------------------------ 31039

chimp.LOC112206744-LOC107973052-GGT2.revcompl                                  ------------------------------------------------------------ 61762
chimp54546-135457.revcompl.Pan.troglodytes.clone.rp43-41g5.GenBank:AC099533.36 ------------------------------------------------------------ 70107
human.FAM30B-LOC105372935-GGT2.NCBI.                                           ctttggagccttggcaatttccaccaggcagaatttacaaccgtctgggacctgcagtct 99913
human.FAM230E-LOC105377182-GGT3P.NCBI.                                         ------------------------------------------------------------ 57310
human.FAM230J-LOC105372942-GGTLC5P.NCBI.rev.compl                              ------------------------------------------------------------ 46364
human.FAM230A.ncRNA.18487127.18500594-GGTLC3.18516335.18518165.rev.compl.REF   ------------------------------------------------------------ 31039

chimp.LOC112206744-LOC107973052-GGT2.revcompl                                  ------------------------------------------------------------ 61762
chimp54546-135457.revcompl.Pan.troglodytes.clone.rp43-41g5.GenBank:AC099533.36 ------------------------------------------------------------ 70107
human.FAM30B-LOC105372935-GGT2.NCBI.                                           cagagggctcacggagaagggggaagctcttggctacccgggaggtcaccagtgactctg 99973
human.FAM230E-LOC105377182-GGT3P.NCBI.                                         ------------------------------------------------------------ 57310
human.FAM230J-LOC105372942-GGTLC5P.NCBI.rev.compl                              ------------------------------------------------------------ 46364
human.FAM230A.ncRNA.18487127.18500594-GGTLC3.18516335.18518165.rev.compl.REF   ------------------------------------------------------------ 31039

chimp.LOC112206744-LOC107973052-GGT2.revcompl                                  ------------------------------------------------------------ 61762
chimp54546-135457.revcompl.Pan.troglodytes.clone.rp43-41g5.GenBank:AC099533.36 ------------------------------------------------------------ 70107
human.FAM30B-LOC105372935-GGT2.NCBI.                                           agcaaagtgcagatcaatgcctgcttagggttacccccaggctgggagagggcatctggc 100033
human.FAM230E-LOC105377182-GGT3P.NCBI.                                         ------------------------------------------------------------ 57310
human.FAM230J-LOC105372942-GGTLC5P.NCBI.rev.compl                              ------------------------------------------------------------ 46364
human.FAM230A.ncRNA.18487127.18500594-GGTLC3.18516335.18518165.rev.compl.REF   ------------------------------------------------------------ 31039

chimp.LOC112206744-LOC107973052-GGT2.revcompl                                  ------------------------------------------------------------ 61762
chimp54546-135457.revcompl.Pan.troglodytes.clone.rp43-41g5.GenBank:AC099533.36 ------------------------------------------------------------ 70107
human.FAM30B-LOC105372935-GGT2.NCBI.                                           aaggaacagacctggaagagcctgcccctctttgagctgctacagcagcttctgggaatt 100093
human.FAM230E-LOC105377182-GGT3P.NCBI.                                         ------------------------------------------------------------ 57310
human.FAM230J-LOC105372942-GGTLC5P.NCBI.rev.compl                              ------------------------------------------------------------ 46364
human.FAM230A.ncRNA.18487127.18500594-GGTLC3.18516335.18518165.rev.compl.REF   ------------------------------------------------------------ 31039

chimp.LOC112206744-LOC107973052-GGT2.revcompl                                  ------------------------------------------------------------ 61762
chimp54546-135457.revcompl.Pan.troglodytes.clone.rp43-41g5.GenBank:AC099533.36 ------------------------------------------------------------ 70107
human.FAM30B-LOC105372935-GGT2.NCBI.                                           tgtaagagccatactgatgagggtcccacaccctcccttgccagggcgccctcagggagg 100153
human.FAM230E-LOC105377182-GGT3P.NCBI.                                         ------------------------------------------------------------ 57310
human.FAM230J-LOC105372942-GGTLC5P.NCBI.rev.compl                              ------------------------------------------------------------ 46364
human.FAM230A.ncRNA.18487127.18500594-GGTLC3.18516335.18518165.rev.compl.REF   ------------------------------------------------------------ 31039

chimp.LOC112206744-LOC107973052-GGT2.revcompl                                  ------------------------------------------------------------ 61762
chimp54546-135457.revcompl.Pan.troglodytes.clone.rp43-41g5.GenBank:AC099533.36 ------------------------------------------------------------ 70107
human.FAM30B-LOC105372935-GGT2.NCBI.                                           ccacactgcaggaggcagagtatgaggtggggaagatggatcggccttatcctcaccacc 100213
human.FAM230E-LOC105377182-GGT3P.NCBI.                                         ------------------------------------------------------------ 57310
human.FAM230J-LOC105372942-GGTLC5P.NCBI.rev.compl                              ------------------------------------------------------------ 46364
human.FAM230A.ncRNA.18487127.18500594-GGTLC3.18516335.18518165.rev.compl.REF   ------------------------------------------------------------ 31039
```

```
chimp.LOC112206744-LOC107973052-GGT2.revcompl                                       ------------------------------------------------------------ 61762
chimp54546-135457.revcompl.Pan.troglodytes.clone.rp43-41g5.GenBank:AC099533.36      ------------------------------------------------------------ 70107
human.FAM30B-LOC105372935-GGT2.NCBI.                                                ccccatacgagtctcctctggtatggatggtgtctcctctctggtgggaagtatgtgatt 100273
human.FAM230E-LOC105377182-GGT3P.NCBI.                                              ------------------------------------------------------------ 57310
human.FAM230J-LOC105372942-GGTLC5P.NCBI.rev.compl                                   ------------------------------------------------------------ 46364
human.FAM230A.ncRNA.18487127.18500594-GGTLC3.18516335.18518165.rev.compl.REF        ------------------------------------------------------------ 31039

chimp.LOC112206744-LOC107973052-GGT2.revcompl                                       ------------------------------------------------------------ 61762
chimp54546-135457.revcompl.Pan.troglodytes.clone.rp43-41g5.GenBank:AC099533.36      ------------------------------------------------------------ 70107
human.FAM30B-LOC105372935-GGT2.NCBI.                                                tctctatttctatttctcttttgttcgggcttcctagaaattcagagctaggtgtgaaat 100333
human.FAM230E-LOC105377182-GGT3P.NCBI.                                              ------------------------------------------------------------ 57310
human.FAM230J-LOC105372942-GGTLC5P.NCBI.rev.compl                                   ------------------------------------------------------------ 46364
human.FAM230A.ncRNA.18487127.18500594-GGTLC3.18516335.18518165.rev.compl.REF        ------------------------------------------------------------ 31039

chimp.LOC112206744-LOC107973052-GGT2.revcompl                                       ------------------------------------------------------------ 61762
chimp54546-135457.revcompl.Pan.troglodytes.clone.rp43-41g5.GenBank:AC099533.36      ------------------------------------------------------------ 70107
human.FAM30B-LOC105372935-GGT2.NCBI.                                                tcccaattcccacagacaagcaacatccccagcagtaataaggacgttccacacacccag 100393
human.FAM230E-LOC105377182-GGT3P.NCBI.                                              ------------------------------------------------------------ 57310
human.FAM230J-LOC105372942-GGTLC5P.NCBI.rev.compl                                   ------------------------------------------------------------ 46364
human.FAM230A.ncRNA.18487127.18500594-GGTLC3.18516335.18518165.rev.compl.REF        ------------------------------------------------------------ 31039

chimp.LOC112206744-LOC107973052-GGT2.revcompl                                       ------------------------------------------------------------ 61762
chimp54546-135457.revcompl.Pan.troglodytes.clone.rp43-41g5.GenBank:AC099533.36      ------------------------------------------------------------ 70107
human.FAM30B-LOC105372935-GGT2.NCBI.                                                gacttgggatatttgtgtattttcaagaataaaagtttgccctttagcctcctcagagcc 100453
human.FAM230E-LOC105377182-GGT3P.NCBI.                                              ------------------------------------------------------------ 57310
human.FAM230J-LOC105372942-GGTLC5P.NCBI.rev.compl                                   ------------------------------------------------------------ 46364
human.FAM230A.ncRNA.18487127.18500594-GGTLC3.18516335.18518165.rev.compl.REF        ------------------------------------------------------------ 31039

chimp.LOC112206744-LOC107973052-GGT2.revcompl                                       ------------------------------------------------------------ 61762
chimp54546-135457.revcompl.Pan.troglodytes.clone.rp43-41g5.GenBank:AC099533.36      ------------------------------------------------------------ 70107
human.FAM30B-LOC105372935-GGT2.NCBI.                                                ccctcccttctcctcgctctccccttaccactgctattcatcatcttactcctacagata 100513
human.FAM230E-LOC105377182-GGT3P.NCBI.                                              ------------------------------------------------------------ 57310
human.FAM230J-LOC105372942-GGTLC5P.NCBI.rev.compl                                   ------------------------------------------------------------ 46364
human.FAM230A.ncRNA.18487127.18500594-GGTLC3.18516335.18518165.rev.compl.REF        ------------------------------------------------------------ 31039

chimp.LOC112206744-LOC107973052-GGT2.revcompl                                       ------------------------------------------------------------ 61762
chimp54546-135457.revcompl.Pan.troglodytes.clone.rp43-41g5.GenBank:AC099533.36      ------------------------------------------------------------ 70107
human.FAM30B-LOC105372935-GGT2.NCBI.                                                ccccctcttcgtggtggcaccgggcactagtaactcatggctgtaaagaagagctgagaa 100573
human.FAM230E-LOC105377182-GGT3P.NCBI.                                              ------------------------------------------------------------ 57310
human.FAM230J-LOC105372942-GGTLC5P.NCBI.rev.compl                                   ------------------------------------------------------------ 46364
human.FAM230A.ncRNA.18487127.18500594-GGTLC3.18516335.18518165.rev.compl.REF        ------------------------------------------------------------ 31039

chimp.LOC112206744-LOC107973052-GGT2.revcompl                                       ------------------------------------------------------------ 61762
chimp54546-135457.revcompl.Pan.troglodytes.clone.rp43-41g5.GenBank:AC099533.36      ------------------------------------------------------------ 70107
human.FAM30B-LOC105372935-GGT2.NCBI.                                                caaaacaggaagaaaagaggttaagtaatggcaagaaaaaagatgtaccttaggaataaa 100633
human.FAM230E-LOC105377182-GGT3P.NCBI.                                              ------------------------------------------------------------ 57310
human.FAM230J-LOC105372942-GGTLC5P.NCBI.rev.compl                                   ------------------------------------------------------------ 46364
human.FAM230A.ncRNA.18487127.18500594-GGTLC3.18516335.18518165.rev.compl.REF        ------------------------------------------------------------ 31039

chimp.LOC112206744-LOC107973052-GGT2.revcompl                                       ------------------------------------------------------------ 61762
chimp54546-135457.revcompl.Pan.troglodytes.clone.rp43-41g5.GenBank:AC099533.36      ------------------------------------------------------------ 70107
human.FAM30B-LOC105372935-GGT2.NCBI.                                                aaagagttcccgaatgtcacagtgtatttaattaatgcttctatttattaaagattaaaa 100693
human.FAM230E-LOC105377182-GGT3P.NCBI.                                              ------------------------------------------------------------ 57310
human.FAM230J-LOC105372942-GGTLC5P.NCBI.rev.compl                                   ------------------------------------------------------------ 46364
human.FAM230A.ncRNA.18487127.18500594-GGTLC3.18516335.18518165.rev.compl.REF        ------------------------------------------------------------ 31039

chimp.LOC112206744-LOC107973052-GGT2.revcompl                                       ------------------------------------------------------------ 61762
chimp54546-135457.revcompl.Pan.troglodytes.clone.rp43-41g5.GenBank:AC099533.36      ------------------------------------------------------------ 70107
human.FAM30B-LOC105372935-GGT2.NCBI.                                                atacatatcaggtgacgacttaaaatgttagtcatgattttaactgtgaattctgtggca 100753
human.FAM230E-LOC105377182-GGT3P.NCBI.                                              ------------------------------------------------------------ 57310
human.FAM230J-LOC105372942-GGTLC5P.NCBI.rev.compl                                   ------------------------------------------------------------ 46364
human.FAM230A.ncRNA.18487127.18500594-GGTLC3.18516335.18518165.rev.compl.REF        ------------------------------------------------------------ 31039

chimp.LOC112206744-LOC107973052-GGT2.revcompl                                       ------------------------------------------------------------ 61762
chimp54546-135457.revcompl.Pan.troglodytes.clone.rp43-41g5.GenBank:AC099533.36      ------------------------------------------------------------ 70107
human.FAM30B-LOC105372935-GGT2.NCBI.                                                ccggagataacagaacaatgagtgcttgtagaaggcactcggttatttgttgaatgaact 100813
human.FAM230E-LOC105377182-GGT3P.NCBI.                                              ------------------------------------------------------------ 57310
human.FAM230J-LOC105372942-GGTLC5P.NCBI.rev.compl                                   ------------------------------------------------------------ 46364
human.FAM230A.ncRNA.18487127.18500594-GGTLC3.18516335.18518165.rev.compl.REF        ------------------------------------------------------------ 31039

chimp.LOC112206744-LOC107973052-GGT2.revcompl                                       ------------------------------------------------------------ 61762
chimp54546-135457.revcompl.Pan.troglodytes.clone.rp43-41g5.GenBank:AC099533.36      ------------------------------------------------------------ 70107
human.FAM30B-LOC105372935-GGT2.NCBI.                                                tcttgggaaagccttcttaggaaagtattagaagtagattctgctaaaacaggggctacc 100873
human.FAM230E-LOC105377182-GGT3P.NCBI.                                              ------------------------------------------------------------ 57310
human.FAM230J-LOC105372942-GGTLC5P.NCBI.rev.compl                                   ------------------------------------------------------------ 46364
human.FAM230A.ncRNA.18487127.18500594-GGTLC3.18516335.18518165.rev.compl.REF        ------------------------------------------------------------ 31039

chimp.LOC112206744-LOC107973052-GGT2.revcompl                                       ------------------------------------------------------------ 61762
chimp54546-135457.revcompl.Pan.troglodytes.clone.rp43-41g5.GenBank:AC099533.36      ------------------------------------------------------------ 70107
human.FAM30B-LOC105372935-GGT2.NCBI.                                                ctcagaggtctcaggacaagggatagaaagtcaaacactgatgtttaagaagggatgcct 100933
human.FAM230E-LOC105377182-GGT3P.NCBI.                                              ------------------------------------------------------------ 57310
human.FAM230J-LOC105372942-GGTLC5P.NCBI.rev.compl                                   ------------------------------------------------------------ 46364
human.FAM230A.ncRNA.18487127.18500594-GGTLC3.18516335.18518165.rev.compl.REF        ------------------------------------------------------------ 31039

chimp.LOC112206744-LOC107973052-GGT2.revcompl                                       ------------------------------------------------------------ 61762
chimp54546-135457.revcompl.Pan.troglodytes.clone.rp43-41g5.GenBank:AC099533.36      ------------------------------------------------------------ 70107
human.FAM30B-LOC105372935-GGT2.NCBI.                                                atgacacatgaatcccacagccatctctccagaagagcaactatttgactgcgcatgcg 100993
human.FAM230E-LOC105377182-GGT3P.NCBI.                                              ------------------------------------------------------------ 57310
human.FAM230J-LOC105372942-GGTLC5P.NCBI.rev.compl                                   ------------------------------------------------------------ 46364
human.FAM230A.ncRNA.18487127.18500594-GGTLC3.18516335.18518165.rev.compl.REF        ------------------------------------------------------------ 31039

chimp.LOC112206744-LOC107973052-GGT2.revcompl                                       ------------------------------------------------------------ 61762
chimp54546-135457.revcompl.Pan.troglodytes.clone.rp43-41g5.GenBank:AC099533.36      ------------------------------------------------------------ 70107
human.FAM30B-LOC105372935-GGT2.NCBI.                                                ggaagatggcgggccgggcgacttgagatccgcgggtctccctgctccttttccgtctgc 101053
human.FAM230E-LOC105377182-GGT3P.NCBI.                                              ------------------------------------------------------------ 57310
human.FAM230J-LOC105372942-GGTLC5P.NCBI.rev.compl                                   ------------------------------------------------------------ 46364
human.FAM230A.ncRNA.18487127.18500594-GGTLC3.18516335.18518165.rev.compl.REF        ------------------------------------------------------------ 31039

chimp.LOC112206744-LOC107973052-GGT2.revcompl                                       ------------------------------------------------------------ 61762
chimp54546-135457.revcompl.Pan.troglodytes.clone.rp43-41g5.GenBank:AC099533.36      ------------------------------------------------------------ 70107
human.FAM30B-LOC105372935-GGT2.NCBI.                                                gtcgggagctcccgggcacgtgaggccgtgccgcgtttactggcgggagggacggcctag 101113
human.FAM230E-LOC105377182-GGT3P.NCBI.                                              ------------------------------------------------------------ 57310
human.FAM230J-LOC105372942-GGTLC5P.NCBI.rev.compl                                   ------------------------------------------------------------ 46364
human.FAM230A.ncRNA.18487127.18500594-GGTLC3.18516335.18518165.rev.compl.REF        ------------------------------------------------------------ 31039

chimp.LOC112206744-LOC107973052-GGT2.revcompl                                       ------------------------------------------------------------ 61762
chimp54546-135457.revcompl.Pan.troglodytes.clone.rp43-41g5.GenBank:AC099533.36      ------------------------------------------------------------ 70107
human.FAM30B-LOC105372935-GGT2.NCBI.                                                ccgggcgacgcctcggaggaggccgcggaccccttaggtgctgggcccttggaaatcggc 101173
human.FAM230E-LOC105377182-GGT3P.NCBI.                                              ------------------------------------------------------------ 57310
human.FAM230J-LOC105372942-GGTLC5P.NCBI.rev.compl                                   ------------------------------------------------------------ 46364
human.FAM230A.ncRNA.18487127.18500594-GGTLC3.18516335.18518165.rev.compl.REF        ------------------------------------------------------------ 31039

chimp.LOC112206744-LOC107973052-GGT2.revcompl                                       ------------------------------------------------------------ 61762
chimp54546-135457.revcompl.Pan.troglodytes.clone.rp43-41g5.GenBank:AC099533.36      ------------------------------------------------------------ 70107
human.FAM30B-LOC105372935-GGT2.NCBI.                                                gcgtgggggcggtgctcgagctgagcgcgagagggcgggagagctcgtggggtgcgagg 101233
human.FAM230E-LOC105377182-GGT3P.NCBI.                                              ------------------------------------------------------------ 57310
human.FAM230J-LOC105372942-GGTLC5P.NCBI.rev.compl                                   ------------------------------------------------------------ 46364
human.FAM230A.ncRNA.18487127.18500594-GGTLC3.18516335.18518165.rev.compl.REF        ------------------------------------------------------------ 31039

chimp.LOC112206744-LOC107973052-GGT2.revcompl                                       ------------------------------------------------------------ 61762
chimp54546-135457.revcompl.Pan.troglodytes.clone.rp43-41g5.GenBank:AC099533.36      ------------------------------------------------------------ 70107
human.FAM30B-LOC105372935-GGT2.NCBI.                                                ggagcaggacgcccggccgggcagcatgagtcagcagcggccggcgaggagattacccag 101293
human.FAM230E-LOC105377182-GGT3P.NCBI.                                              ------------------------------------------------------------ 57310
human.FAM230J-LOC105372942-GGTLC5P.NCBI.rev.compl                                   ------------------------------------------------------------ 46364
human.FAM230A.ncRNA.18487127.18500594-GGTLC3.18516335.18518165.rev.compl.REF        ------------------------------------------------------------ 31039

chimp.LOC112206744-LOC107973052-GGT2.revcompl                                       ------------------------------------------------------------ 61762
```

```
chimp54546-135457.revcompl.Pan.troglodytes.clone.rp43-41g5.GenBank:AC099533.36     ------------------------------------------------------------ 70107
human.FAM30B-LOC105372935-GGT2.NCBI.                                                tctcctcctgcacccgacggaggagaccccatcaacgtggagggcctgctgccatcaaaa 101353
human.FAM230E-LOC105377182-GGT3P.NCBI.                                              ------------------------------------------------------------ 57310
human.FAM230J-LOC105372942-GGTLC5P.NCBI.rev.compl                                   ------------------------------------------------------------ 46364
human.FAM230A.ncRNA.18487127.18500594-GGTLC3.18516335.18518165.rev.compl.REF        ------------------------------------------------------------ 31039

chimp.LOC112206744-LOC107973052-GGT2.revcompl                                       ------------------------------------------------------------ 61762
chimp54546-135457.revcompl.Pan.troglodytes.clone.rp43-41g5.GenBank:AC099533.36     ------------------------------------------------------------ 70107
human.FAM30B-LOC105372935-GGT2.NCBI.                                                ataaggattaatttagaagataatgtacaatatgtgtccatgagaaatctgctcccgggg 101413
human.FAM230E-LOC105377182-GGT3P.NCBI.                                              ------------------------------------------------------------ 57310
human.FAM230J-LOC105372942-GGTLC5P.NCBI.rev.compl                                   ------------------------------------------------------------ 46364
human.FAM230A.ncRNA.18487127.18500594-GGTLC3.18516335.18518165.rev.compl.REF        ------------------------------------------------------------ 31039

chimp.LOC112206744-LOC107973052-GGT2.revcompl                                       ------------------------------------------------------------ 61762
chimp54546-135457.revcompl.Pan.troglodytes.clone.rp43-41g5.GenBank:AC099533.36     ------------------------------------------------------------ 70107
human.FAM30B-LOC105372935-GGT2.NCBI.                                                gtattcttgacttaaacaaggttgcaacgaaactgggagtccgaaagcggagagtgtatg 101473
human.FAM230E-LOC105377182-GGT3P.NCBI.                                              ------------------------------------------------------------ 57310
human.FAM230J-LOC105372942-GGTLC5P.NCBI.rev.compl                                   ------------------------------------------------------------ 46364
human.FAM230A.ncRNA.18487127.18500594-GGTLC3.18516335.18518165.rev.compl.REF        ------------------------------------------------------------ 31039

chimp.LOC112206744-LOC107973052-GGT2.revcompl                                       ------------------------------------------------------------ 61762
chimp54546-135457.revcompl.Pan.troglodytes.clone.rp43-41g5.GenBank:AC099533.36     ------------------------------------------------------------ 70107
human.FAM30B-LOC105372935-GGT2.NCBI.                                                acatcaccgatgtcttagatggaatcgacctcgttgaaaagaaatccaagaaccatatta 101533
human.FAM230E-LOC105377182-GGT3P.NCBI.                                              ------------------------------------------------------------ 57310
human.FAM230J-LOC105372942-GGTLC5P.NCBI.rev.compl                                   ------------------------------------------------------------ 46364
human.FAM230A.ncRNA.18487127.18500594-GGTLC3.18516335.18518165.rev.compl.REF        ------------------------------------------------------------ 31039

chimp.LOC112206744-LOC107973052-GGT2.revcompl                                       ------------------------------------------------------------ 61762
chimp54546-135457.revcompl.Pan.troglodytes.clone.rp43-41g5.GenBank:AC099533.36     ------------------------------------------------------------ 70107
human.FAM30B-LOC105372935-GGT2.NCBI.                                                gatggataggatctgatcttagcaattttggagcagttccccaacaaaagaagctacagg 101593
human.FAM230E-LOC105377182-GGT3P.NCBI.                                              ------------------------------------------------------------ 57310
human.FAM230J-LOC105372942-GGTLC5P.NCBI.rev.compl                                   ------------------------------------------------------------ 46364
human.FAM230A.ncRNA.18487127.18500594-GGTLC3.18516335.18518165.rev.compl.REF        ------------------------------------------------------------ 31039

chimp.LOC112206744-LOC107973052-GGT2.revcompl                                       ------------------------------------------------------------ 61762
chimp54546-135457.revcompl.Pan.troglodytes.clone.rp43-41g5.GenBank:AC099533.36     ------------------------------------------------------------ 70107
human.FAM30B-LOC105372935-GGT2.NCBI.                                                aggaactttctgacttaccagcaatggaagatgctttggatgagttaattaaggattgtg 101653
human.FAM230E-LOC105377182-GGT3P.NCBI.                                              ------------------------------------------------------------ 57310
human.FAM230J-LOC105372942-GGTLC5P.NCBI.rev.compl                                   ------------------------------------------------------------ 46364
human.FAM230A.ncRNA.18487127.18500594-GGTLC3.18516335.18518165.rev.compl.REF        ------------------------------------------------------------ 31039

chimp.LOC112206744-LOC107973052-GGT2.revcompl                                       ------------------------------------------------------------ 61762
chimp54546-135457.revcompl.Pan.troglodytes.clone.rp43-41g5.GenBank:AC099533.36     ------------------------------------------------------------ 70107
human.FAM30B-LOC105372935-GGT2.NCBI.                                                ctcagcagctgtttgagttaacagatgacaaagaaaatgaaagactagcatatgtgacct 101713
human.FAM230E-LOC105377182-GGT3P.NCBI.                                              ------------------------------------------------------------ 57310
human.FAM230J-LOC105372942-GGTLC5P.NCBI.rev.compl                                   ------------------------------------------------------------ 46364
human.FAM230A.ncRNA.18487127.18500594-GGTLC3.18516335.18518165.rev.compl.REF        ------------------------------------------------------------ 31039

chimp.LOC112206744-LOC107973052-GGT2.revcompl                                       ------------------------------------------------------------ 61762
chimp54546-135457.revcompl.Pan.troglodytes.clone.rp43-41g5.GenBank:AC099533.36     ------------------------------------------------------------ 70107
human.FAM30B-LOC105372935-GGT2.NCBI.                                                atcaagacattcatagcattcaggccttccatgaacagatcgtcattgcagttaaagctc 101773
human.FAM230E-LOC105377182-GGT3P.NCBI.                                              ------------------------------------------------------------ 57310
human.FAM230J-LOC105372942-GGTLC5P.NCBI.rev.compl                                   ------------------------------------------------------------ 46364
human.FAM230A.ncRNA.18487127.18500594-GGTLC3.18516335.18518165.rev.compl.REF        ------------------------------------------------------------ 31039

chimp.LOC112206744-LOC107973052-GGT2.revcompl                                       ------------------------------------------------------------ 61762
chimp54546-135457.revcompl.Pan.troglodytes.clone.rp43-41g5.GenBank:AC099533.36     ------------------------------------------------------------ 70107
human.FAM30B-LOC105372935-GGT2.NCBI.                                                cagcagaaaccagattggatgttccagctcccagagaagactctatcacagtgcacataa 101833
human.FAM230E-LOC105377182-GGT3P.NCBI.                                              ------------------------------------------------------------ 57310
human.FAM230J-LOC105372942-GGTLC5P.NCBI.rev.compl                                   ------------------------------------------------------------ 46364
human.FAM230A.ncRNA.18487127.18500594-GGTLC3.18516335.18518165.rev.compl.REF        ------------------------------------------------------------ 31039

chimp.LOC112206744-LOC107973052-GGT2.revcompl                                       ------------------------------------------------------------ 61762
chimp54546-135457.revcompl.Pan.troglodytes.clone.rp43-41g5.GenBank:AC099533.36     ------------------------------------------------------------ 70107
human.FAM30B-LOC105372935-GGT2.NCBI.                                                ggagcaccaacggacctatcgatgtctatttgtgagaagtggagcagggtcagaccagta 101893
human.FAM230E-LOC105377182-GGT3P.NCBI.                                              ------------------------------------------------------------ 57310
human.FAM230J-LOC105372942-GGTLC5P.NCBI.rev.compl                                   ------------------------------------------------------------ 46364
human.FAM230A.ncRNA.18487127.18500594-GGTLC3.18516335.18518165.rev.compl.REF        ------------------------------------------------------------ 31039

chimp.LOC112206744-LOC107973052-GGT2.revcompl                                       ------------------------------------------------------------ 61762
chimp54546-135457.revcompl.Pan.troglodytes.clone.rp43-41g5.GenBank:AC099533.36     ------------------------------------------------------------ 70107
human.FAM30B-LOC105372935-GGT2.NCBI.                                                acaaaaggtctgaaggtgtcaggacctcttcatctgagagcactcatccagaaggccctg 101953
human.FAM230E-LOC105377182-GGT3P.NCBI.                                              ------------------------------------------------------------ 57310
human.FAM230J-LOC105372942-GGTLC5P.NCBI.rev.compl                                   ------------------------------------------------------------ 46364
human.FAM230A.ncRNA.18487127.18500594-GGTLC3.18516335.18518165.rev.compl.REF        ------------------------------------------------------------ 31039

chimp.LOC112206744-LOC107973052-GGT2.revcompl                                       ------------------------------------------------------------ 61762
chimp54546-135457.revcompl.Pan.troglodytes.clone.rp43-41g5.GenBank:AC099533.36     ------------------------------------------------------------ 70107
human.FAM30B-LOC105372935-GGT2.NCBI.                                                aggaagaagaaaatcctcagcaaagtgaagaattgcttgaagtaagcaactgatggcatt 102013
human.FAM230E-LOC105377182-GGT3P.NCBI.                                              ------------------------------------------------------------ 57310
human.FAM230J-LOC105372942-GGTLC5P.NCBI.rev.compl                                   ------------------------------------------------------------ 46364
human.FAM230A.ncRNA.18487127.18500594-GGTLC3.18516335.18518165.rev.compl.REF        ------------------------------------------------------------ 31039

chimp.LOC112206744-LOC107973052-GGT2.revcompl                                       ------------------------------------------------------------ 61762
chimp54546-135457.revcompl.Pan.troglodytes.clone.rp43-41g5.GenBank:AC099533.36     ------------------------------------------------------------ 70107
human.FAM30B-LOC105372935-GGT2.NCBI.                                                tgagaatttatgtatcactgagttttttgggaatatcttcctggagaattacgcatcaaa 102073
human.FAM230E-LOC105377182-GGT3P.NCBI.                                              ------------------------------------------------------------ 57310
human.FAM230J-LOC105372942-GGTLC5P.NCBI.rev.compl                                   ------------------------------------------------------------ 46364
human.FAM230A.ncRNA.18487127.18500594-GGTLC3.18516335.18518165.rev.compl.REF        ------------------------------------------------------------ 31039

chimp.LOC112206744-LOC107973052-GGT2.revcompl                                       ------------------------------------------------------------ 61762
chimp54546-135457.revcompl.Pan.troglodytes.clone.rp43-41g5.GenBank:AC099533.36     ------------------------------------------------------------ 70107
human.FAM30B-LOC105372935-GGT2.NCBI.                                                tttgattctcagagcaataaattatccatgaagtgctctcgttctcagtagcggcatcat 102133
human.FAM230E-LOC105377182-GGT3P.NCBI.                                              ------------------------------------------------------------ 57310
human.FAM230J-LOC105372942-GGTLC5P.NCBI.rev.compl                                   ------------------------------------------------------------ 46364
human.FAM230A.ncRNA.18487127.18500594-GGTLC3.18516335.18518165.rev.compl.REF        ------------------------------------------------------------ 31039

chimp.LOC112206744-LOC107973052-GGT2.revcompl                                       ------------------------------------------------------------ 61762
chimp54546-135457.revcompl.Pan.troglodytes.clone.rp43-41g5.GenBank:AC099533.36     ------------------------------------------------------------ 70107
human.FAM30B-LOC105372935-GGT2.NCBI.                                                ggccagtagtgtctttgaggagttcaccacttagattactgagtaattgtggtttccaca 102193
human.FAM230E-LOC105377182-GGT3P.NCBI.                                              ------------------------------------------------------------ 57310
human.FAM230J-LOC105372942-GGTLC5P.NCBI.rev.compl                                   ------------------------------------------------------------ 46364
human.FAM230A.ncRNA.18487127.18500594-GGTLC3.18516335.18518165.rev.compl.REF        ------------------------------------------------------------ 31039

chimp.LOC112206744-LOC107973052-GGT2.revcompl                                       ------------------------------------------------------------ 61762
chimp54546-135457.revcompl.Pan.troglodytes.clone.rp43-41g5.GenBank:AC099533.36     ------------------------------------------------------------ 70107
human.FAM30B-LOC105372935-GGT2.NCBI.                                                tttgaaaacaactccttttataattattcactgctttttgtcagtgaaatagacatcttg 102253
human.FAM230E-LOC105377182-GGT3P.NCBI.                                              ------------------------------------------------------------ 57310
human.FAM230J-LOC105372942-GGTLC5P.NCBI.rev.compl                                   ------------------------------------------------------------ 46364
human.FAM230A.ncRNA.18487127.18500594-GGTLC3.18516335.18518165.rev.compl.REF        ------------------------------------------------------------ 31039

chimp.LOC112206744-LOC107973052-GGT2.revcompl                                       ------------------------------------------------------------ 61762
chimp54546-135457.revcompl.Pan.troglodytes.clone.rp43-41g5.GenBank:AC099533.36     ------------------------------------------------------------ 70107
human.FAM30B-LOC105372935-GGT2.NCBI.                                                cctcctgaagtagcttcatcacagagtgtcatgaagacagacagtcaggctgaaaaggac 102313
human.FAM230E-LOC105377182-GGT3P.NCBI.                                              ------------------------------------------------------------ 57310
human.FAM230J-LOC105372942-GGTLC5P.NCBI.rev.compl                                   ------------------------------------------------------------ 46364
human.FAM230A.ncRNA.18487127.18500594-GGTLC3.18516335.18518165.rev.compl.REF        ------------------------------------------------------------ 31039

chimp.LOC112206744-LOC107973052-GGT2.revcompl                                       ------------------------------------------------------------ 61762
chimp54546-135457.revcompl.Pan.troglodytes.clone.rp43-41g5.GenBank:AC099533.36     ------------------------------------------------------------ 70107
human.FAM30B-LOC105372935-GGT2.NCBI.                                                agttctttgtggactctacccttcccttcaaggagtatgtcatatgtcacaaaagaaatt 102373
human.FAM230E-LOC105377182-GGT3P.NCBI.                                              ------------------------------------------------------------ 57310
human.FAM230J-LOC105372942-GGTLC5P.NCBI.rev.compl                                   ------------------------------------------------------------ 46364
human.FAM230A.ncRNA.18487127.18500594-GGTLC3.18516335.18518165.rev.compl.REF        ------------------------------------------------------------ 31039

chimp.LOC112206744-LOC107973052-GGT2.revcompl                                       ------------------------------------------------------------ 61762
chimp54546-135457.revcompl.Pan.troglodytes.clone.rp43-41g5.GenBank:AC099533.36     ------------------------------------------------------------ 70107
```

```
human.FAM30B-LOC105372935-GGT2.NCBI.                                               gccttacactggttcatgtttgcagttactgttgtacattgcatagatgtacacacgaat 102433
human.FAM230E-LOC105377182-GGT3P.NCBI.                                             ------------------------------------------------------------ 57310
human.FAM230J-LOC105372942-GGTLC5P.NCBI.rev.compl                                  ------------------------------------------------------------ 46364
human.FAM230A.ncRNA.18487127.18500594-GGTLC3.18516335.18518165.rev.compl.REF       ------------------------------------------------------------ 31039

chimp.LOC112206744-LOC107973052-GGT2.revcompl                                      ------------------------------------------------------------ 61762
chimp54546-135457.revcompl.Pan.troglodytes.clone.rp43-41g5.GenBank:AC099533.36     ---------------TTTTATATAAGAATACAGAGAGGTGGAAAATAATGAAAAATATGT 70152
human.FAM30B-LOC105372935-GGT2.NCBI.                                               ttaaatgtgatgtctttgtatatatctgtataatgttgagattacttacgaaatatgtct 102493
human.FAM230E-LOC105377182-GGT3P.NCBI.                                             ------------------------------------------------------------ 57310
human.FAM230J-LOC105372942-GGTLC5P.NCBI.rev.compl                                  ------------------------------------------------------------ 46364
human.FAM230A.ncRNA.18487127.18500594-GGTLC3.18516335.18518165.rev.compl.REF       ------------------------------------------------------------ 31039

chimp.LOC112206744-LOC107973052-GGT2.revcompl                                      ------------------------------------------------------------ 61762
chimp54546-135457.revcompl.Pan.troglodytes.clone.rp43-41g5.GenBank:AC099533.36     CA---------------------------------------------------------- 70154
human.FAM30B-LOC105372935-GGT2.NCBI.                                               gagtgacacttttcactctcgtacagccaaaataatgtatatatggaaagtgacagacaa 102553
human.FAM230E-LOC105377182-GGT3P.NCBI.                                             ------------------------------------------------------------ 57310
human.FAM230J-LOC105372942-GGTLC5P.NCBI.rev.compl                                  ------------------------------------------------------------ 46364
human.FAM230A.ncRNA.18487127.18500594-GGTLC3.18516335.18518165.rev.compl.REF       ------------------------------------------------------------ 31039

chimp.LOC112206744-LOC107973052-GGT2.revcompl                                      ------------------------------------------------------------ 61762
chimp54546-135457.revcompl.Pan.troglodytes.clone.rp43-41g5.GenBank:AC099533.36     ------------------------------------------------------------ 70154
human.FAM30B-LOC105372935-GGT2.NCBI.                                               attctctaatctctttggtatctataacttattagaatcctctggatgagggttagaaga 102613
human.FAM230E-LOC105377182-GGT3P.NCBI.                                             ------------------------------------------------------------ 57310
human.FAM230J-LOC105372942-GGTLC5P.NCBI.rev.compl                                  ------------------------------------------------------------ 46364
human.FAM230A.ncRNA.18487127.18500594-GGTLC3.18516335.18518165.rev.compl.REF       ------------------------------------------------------------ 31039

chimp.LOC112206744-LOC107973052-GGT2.revcompl                                      ------------------------------------------------------------ 61762
chimp54546-135457.revcompl.Pan.troglodytes.clone.rp43-41g5.GenBank:AC099533.36     ------------------------------------------------------------ 70154
human.FAM30B-LOC105372935-GGT2.NCBI.                                               gactttttccaaacttctacatgtagaagtatcataaatgtgctacacatttatgtttgt 102673
human.FAM230E-LOC105377182-GGT3P.NCBI.                                             ------------------------------------------------------------ 57310
human.FAM230J-LOC105372942-GGTLC5P.NCBI.rev.compl                                  ------------------------------------------------------------ 46364
human.FAM230A.ncRNA.18487127.18500594-GGTLC3.18516335.18518165.rev.compl.REF       ------------------------------------------------------------ 31039

chimp.LOC112206744-LOC107973052-GGT2.revcompl                                      ------------------------------------------------------------ 61762
chimp54546-135457.revcompl.Pan.troglodytes.clone.rp43-41g5.GenBank:AC099533.36     ------------------------------------------------------------ 70154
human.FAM30B-LOC105372935-GGT2.NCBI.                                               ggatttaattaaagtattttaatatggttttcagtgctaaaattggagtcagatacttct 102733
human.FAM230E-LOC105377182-GGT3P.NCBI.                                             ------------------------------------------------------------ 57310
human.FAM230J-LOC105372942-GGTLC5P.NCBI.rev.compl                                  ------------------------------------------------------------ 46364
human.FAM230A.ncRNA.18487127.18500594-GGTLC3.18516335.18518165.rev.compl.REF       ------------------------------------------------------------ 31039

chimp.LOC112206744-LOC107973052-GGT2.revcompl                                      ------------------------------------------------------------ 61762
chimp54546-135457.revcompl.Pan.troglodytes.clone.rp43-41g5.GenBank:AC099533.36     ------------------------------------------------------------ 70154
human.FAM30B-LOC105372935-GGT2.NCBI.                                               tggttttaagctgtctacctaattgctgtctcccagcagatcggtggcatgcccagtggc 102793
human.FAM230E-LOC105377182-GGT3P.NCBI.                                             ------------------------------------------------------------ 57310
human.FAM230J-LOC105372942-GGTLC5P.NCBI.rev.compl                                  ------------------------------------------------------------ 46364
human.FAM230A.ncRNA.18487127.18500594-GGTLC3.18516335.18518165.rev.compl.REF       ------------------------------------------------------------ 31039

chimp.LOC112206744-LOC107973052-GGT2.revcompl                                      ------------------------------------------------------------ 61762
chimp54546-135457.revcompl.Pan.troglodytes.clone.rp43-41g5.GenBank:AC099533.36     ------------------------------------------------------------ 70154
human.FAM30B-LOC105372935-GGT2.NCBI.                                               tttgggggcaaggatagaaatgtcatcaggaaatagctgaattcattgtgaaacatgaat 102853
human.FAM230E-LOC105377182-GGT3P.NCBI.                                             ------------------------------------------------------------ 57310
human.FAM230J-LOC105372942-GGTLC5P.NCBI.rev.compl                                  ------------------------------------------------------------ 46364
human.FAM230A.ncRNA.18487127.18500594-GGTLC3.18516335.18518165.rev.compl.REF       ------------------------------------------------------------ 31039

chimp.LOC112206744-LOC107973052-GGT2.revcompl                                      ------------------------------------------------------------ 61762
chimp54546-135457.revcompl.Pan.troglodytes.clone.rp43-41g5.GenBank:AC099533.36     ------------------------------------------------------------ 70154
human.FAM30B-LOC105372935-GGT2.NCBI.                                               tcagtcatggtgataattggaaactcctttcaggtttttgcaagtagattttgtaatgtt 102913
human.FAM230E-LOC105377182-GGT3P.NCBI.                                             ------------------------------------------------------------ 57310
human.FAM230J-LOC105372942-GGTLC5P.NCBI.rev.compl                                  ------------------------------------------------------------ 46364
human.FAM230A.ncRNA.18487127.18500594-GGTLC3.18516335.18518165.rev.compl.REF       ------------------------------------------------------------ 31039

chimp.LOC112206744-LOC107973052-GGT2.revcompl                                      ------------------------------------------------------------ 61762
chimp54546-135457.revcompl.Pan.troglodytes.clone.rp43-41g5.GenBank:AC099533.36     ------------------------------------------------------------ 70154
human.FAM30B-LOC105372935-GGT2.NCBI.                                               tgtgtatgcagccttgctgttgagtcagtccaaggggttttacttaggacaagttgtacc 102973
human.FAM230E-LOC105377182-GGT3P.NCBI.                                             ------------------------------------------------------------ 57310
human.FAM230J-LOC105372942-GGTLC5P.NCBI.rev.compl                                  ------------------------------------------------------------ 46364
human.FAM230A.ncRNA.18487127.18500594-GGTLC3.18516335.18518165.rev.compl.REF       ------------------------------------------------------------ 31039

chimp.LOC112206744-LOC107973052-GGT2.revcompl                                      ------------------------------------------------------------ 61762
chimp54546-135457.revcompl.Pan.troglodytes.clone.rp43-41g5.GenBank:AC099533.36     ------------------------------------------------------------ 70154
human.FAM30B-LOC105372935-GGT2.NCBI.                                               ttgccctctctccagctctgctcccacattttcacatacctagctatttctacctcattg 103033
human.FAM230E-LOC105377182-GGT3P.NCBI.                                             ------------------------------------------------------------ 57310
human.FAM230J-LOC105372942-GGTLC5P.NCBI.rev.compl                                  ------------------------------------------------------------ 46364
human.FAM230A.ncRNA.18487127.18500594-GGTLC3.18516335.18518165.rev.compl.REF       ------------------------------------------------------------ 31039

chimp.LOC112206744-LOC107973052-GGT2.revcompl                                      ------------------------------------------------------------ 61762
chimp54546-135457.revcompl.Pan.troglodytes.clone.rp43-41g5.GenBank:AC099533.36     ------------------------------------------------------------ 70154
human.FAM30B-LOC105372935-GGT2.NCBI.                                               ggtaagtcatttaccactctgtgcctcagtttactctgtagtttaccattagactgtgag 103093
human.FAM230E-LOC105377182-GGT3P.NCBI.                                             ------------------------------------------------------------ 57310
human.FAM230J-LOC105372942-GGTLC5P.NCBI.rev.compl                                  ------------------------------------------------------------ 46364
human.FAM230A.ncRNA.18487127.18500594-GGTLC3.18516335.18518165.rev.compl.REF       ------------------------------------------------------------ 31039

chimp.LOC112206744-LOC107973052-GGT2.revcompl                                      ------------------------------------------------------------ 61762
chimp54546-135457.revcompl.Pan.troglodytes.clone.rp43-41g5.GenBank:AC099533.36     ------------------------------------------------------------ 70154
human.FAM30B-LOC105372935-GGT2.NCBI.                                               ctccctgagggactttgtcataatcactgttacatcccagtgcctcacaccatgcctggc 103153
human.FAM230E-LOC105377182-GGT3P.NCBI.                                             ------------------------------------------------------------ 57310
human.FAM230J-LOC105372942-GGTLC5P.NCBI.rev.compl                                  ------------------------------------------------------------ 46364
human.FAM230A.ncRNA.18487127.18500594-GGTLC3.18516335.18518165.rev.compl.REF       ------------------------------------------------------------ 31039

chimp.LOC112206744-LOC107973052-GGT2.revcompl                                      ------------------------------------------------------------ 61762
chimp54546-135457.revcompl.Pan.troglodytes.clone.rp43-41g5.GenBank:AC099533.36     ------------------------------------------------------------ 70154
human.FAM30B-LOC105372935-GGT2.NCBI.                                               ccttaagaagtgctcaataaatgtctgaacaaaaaaaaaaaaaaaaaaaaaaaaaatcag 103213
human.FAM230E-LOC105377182-GGT3P.NCBI.                                             ------------------------------------------------------------ 57310
human.FAM230J-LOC105372942-GGTLC5P.NCBI.rev.compl                                  ------------------------------------------------------------ 46364
human.FAM230A.ncRNA.18487127.18500594-GGTLC3.18516335.18518165.rev.compl.REF       ------------------------------------------------------------ 31039

chimp.LOC112206744-LOC107973052-GGT2.revcompl                                      ------------------------------------------------------------ 61762
chimp54546-135457.revcompl.Pan.troglodytes.clone.rp43-41g5.GenBank:AC099533.36     ------------------------------------------------------------ 70154
human.FAM30B-LOC105372935-GGT2.NCBI.                                               gtctaaagatgagccatccaagaaactgctcttgctcccagaggttttaaagaaagtcat 103273
human.FAM230E-LOC105377182-GGT3P.NCBI.                                             ------------------------------------------------------------ 57310
human.FAM230J-LOC105372942-GGTLC5P.NCBI.rev.compl                                  ------------------------------------------------------------ 46364
human.FAM230A.ncRNA.18487127.18500594-GGTLC3.18516335.18518165.rev.compl.REF       ------------------------------------------------------------ 31039

chimp.LOC112206744-LOC107973052-GGT2.revcompl                                      ------------------------------------------------------------ 61762
chimp54546-135457.revcompl.Pan.troglodytes.clone.rp43-41g5.GenBank:AC099533.36     ------------------------------------------------------------ 70154
human.FAM30B-LOC105372935-GGT2.NCBI.                                               tcccatgaacgtgagaagtaacccaacactttgaaatgataatctttttacagcaataac 103333
human.FAM230E-LOC105377182-GGT3P.NCBI.                                             ------------------------------------------------------------ 57310
human.FAM230J-LOC105372942-GGTLC5P.NCBI.rev.compl                                  ------------------------------------------------------------ 46364
human.FAM230A.ncRNA.18487127.18500594-GGTLC3.18516335.18518165.rev.compl.REF       ------------------------------------------------------------ 31039

chimp.LOC112206744-LOC107973052-GGT2.revcompl                                      ------------------------------------------------------------ 61762
chimp54546-135457.revcompl.Pan.troglodytes.clone.rp43-41g5.GenBank:AC099533.36     ------TGCATGCACTAACCAAGAAAGCTGTATCGCTTT--------------------- 70187
human.FAM30B-LOC105372935-GGT2.NCBI.                                               tgttgctgcagggcttgagaaaaaaagttctcttcctttgggttaaagtcttaaatgctg 103393
human.FAM230E-LOC105377182-GGT3P.NCBI.                                             ------------------------------------------------------------ 57310
human.FAM230J-LOC105372942-GGTLC5P.NCBI.rev.compl                                  ------------------------------------------------------------ 46364
human.FAM230A.ncRNA.18487127.18500594-GGTLC3.18516335.18518165.rev.compl.REF       ------------------------------------------------------------ 31039

chimp.LOC112206744-LOC107973052-GGT2.revcompl                                      ------------------------------------------------------------ 61762
chimp54546-135457.revcompl.Pan.troglodytes.clone.rp43-41g5.GenBank:AC099533.36     ------------------------------------------------------------ 70187
human.FAM30B-LOC105372935-GGT2.NCBI.                                               agcccgtagccagcaaccttaaagactactcgggtccagttcattctatctggaggagtc 103453
human.FAM230E-LOC105377182-GGT3P.NCBI.                                             ------------------------------------------------------------ 57310
human.FAM230J-LOC105372942-GGTLC5P.NCBI.rev.compl                                  ------------------------------------------------------------ 46364
human.FAM230A.ncRNA.18487127.18500594-GGTLC3.18516335.18518165.rev.compl.REF       ------------------------------------------------------------ 31039

chimp.LOC112206744-LOC107973052-GGT2.revcompl                                      ------------------------------------------------------------ 61762
chimp54546-135457.revcompl.Pan.troglodytes.clone.rp43-41g5.GenBank:AC099533.36     ------------------------------------------------------------ 70187
human.FAM30B-LOC105372935-GGT2.NCBI.                                               acttgggaattgaaaacacaggaaaactttttttctggtccatgaacaaacaagaacgct 103513
```

```
human.FAM230E-LOC105377182-GGT3P.NCBI.                                      ------------------------------------------------------------ 57310
human.FAM230J-LOC105372942-GGTLC5P.NCBI.rev.compl                           ------------------------------------------------------------ 46364
human.FAM230A.ncRNA.18487127.18500594-GGTLC3.18516335.18518165.rev.compl.REF ------------------------------------------------------------ 31039

chimp.LOC112206744-LOC107973052-GGT2.revcompl                               ------------------------------------------------------------ 61762
chimp54546-135457.revcompl.Pan.troglodytes.clone.rp43-41g5.GenBank:AC099533.36 ------------------------------------------------------------ 70187
human.FAM30B-LOC105372935-GGT2.NCBI.                                        taactgaattagacacaaaactaagtatttatggttgtcatcttaacaggtttcaaacag 103573
human.FAM230E-LOC105377182-GGT3P.NCBI.                                      ------------------------------------------------------------ 57310
human.FAM230J-LOC105372942-GGTLC5P.NCBI.rev.compl                           ------------------------------------------------------------ 46364
human.FAM230A.ncRNA.18487127.18500594-GGTLC3.18516335.18518165.rev.compl.REF ------------------------------------------------------------ 31039

chimp.LOC112206744-LOC107973052-GGT2.revcompl                               ------------------------------------------------------------ 61762
chimp54546-135457.revcompl.Pan.troglodytes.clone.rp43-41g5.GenBank:AC099533.36 ------------------------------------------------------------ 70187
human.FAM30B-LOC105372935-GGT2.NCBI.                                        gcttcattttaatttaacataatttcaaaattatagcaaaatagtcaataaaatacgaac 103633
human.FAM230E-LOC105377182-GGT3P.NCBI.                                      ------------------------------------------------------------ 57310
human.FAM230J-LOC105372942-GGTLC5P.NCBI.rev.compl                           ------------------------------------------------------------ 46364
human.FAM230A.ncRNA.18487127.18500594-GGTLC3.18516335.18518165.rev.compl.REF ------------------------------------------------------------ 31039

chimp.LOC112206744-LOC107973052-GGT2.revcompl                               ------------------------------------------------------------ 61762
chimp54546-135457.revcompl.Pan.troglodytes.clone.rp43-41g5.GenBank:AC099533.36 ------------------------------------------------------------ 70187
human.FAM30B-LOC105372935-GGT2.NCBI.                                        gtgcttaatttgatgaatttatgtgtttttttgaagaaaatatgcagaataaacaagtcaa 103693
human.FAM230E-LOC105377182-GGT3P.NCBI.                                      ------------------------------------------------------------ 57310
human.FAM230J-LOC105372942-GGTLC5P.NCBI.rev.compl                           ------------------------------------------------------------ 46364
human.FAM230A.ncRNA.18487127.18500594-GGTLC3.18516335.18518165.rev.compl.REF ------------------------------------------------------------ 31039

chimp.LOC112206744-LOC107973052-GGT2.revcompl                               ------------------------------------------------------------ 61762
chimp54546-135457.revcompl.Pan.troglodytes.clone.rp43-41g5.GenBank:AC099533.36 ------------------------------------------------------------ 70187
human.FAM30B-LOC105372935-GGT2.NCBI.                                        ttattctcactaaatacattaaaatttaagggtaatatttttattatgacttgctctcct 103753
human.FAM230E-LOC105377182-GGT3P.NCBI.                                      ------------------------------------------------------------ 57310
human.FAM230J-LOC105372942-GGTLC5P.NCBI.rev.compl                           ------------------------------------------------------------ 46364
human.FAM230A.ncRNA.18487127.18500594-GGTLC3.18516335.18518165.rev.compl.REF ------------------------------------------------------------ 31039

chimp.LOC112206744-LOC107973052-GGT2.revcompl                               ------------------------------------------------------------ 61762
chimp54546-135457.revcompl.Pan.troglodytes.clone.rp43-41g5.GenBank:AC099533.36 ------------------------------------------------------------ 70187
human.FAM30B-LOC105372935-GGT2.NCBI.                                        gatatagcaaaatttattgttatatagttatggaaggaggctcactactccccaatgatt 103813
human.FAM230E-LOC105377182-GGT3P.NCBI.                                      ------------------------------------------------------------ 57310
human.FAM230J-LOC105372942-GGTLC5P.NCBI.rev.compl                           ------------------------------------------------------------ 46364
human.FAM230A.ncRNA.18487127.18500594-GGTLC3.18516335.18518165.rev.compl.REF ------------------------------------------------------------ 31039

chimp.LOC112206744-LOC107973052-GGT2.revcompl                               ------------------------------------------------------------ 61762
chimp54546-135457.revcompl.Pan.troglodytes.clone.rp43-41g5.GenBank:AC099533.36 ------------------------------------------------------------ 70187
human.FAM30B-LOC105372935-GGT2.NCBI.                                        tcacaaattacagcttcagccagctataaaataaccaaatcagttctgatatagccataa 103873
human.FAM230E-LOC105377182-GGT3P.NCBI.                                      ------------------------------------------------------------ 57310
human.FAM230J-LOC105372942-GGTLC5P.NCBI.rev.compl                           ------------------------------------------------------------ 46364
human.FAM230A.ncRNA.18487127.18500594-GGTLC3.18516335.18518165.rev.compl.REF ------------------------------------------------------------ 31039

chimp.LOC112206744-LOC107973052-GGT2.revcompl                               ------------------------------------------------------------ 61762
chimp54546-135457.revcompl.Pan.troglodytes.clone.rp43-41g5.GenBank:AC099533.36 ------------------------------------------------------------ 70187
human.FAM30B-LOC105372935-GGT2.NCBI.                                        aattaagcctgaaaaaatagtgaagtaaatcatccctttaaaatcaagttgttgaacaat 103933
human.FAM230E-LOC105377182-GGT3P.NCBI.                                      ------------------------------------------------------------ 57310
human.FAM230J-LOC105372942-GGTLC5P.NCBI.rev.compl                           ------------------------------------------------------------ 46364
human.FAM230A.ncRNA.18487127.18500594-GGTLC3.18516335.18518165.rev.compl.REF ------------------------------------------------------------ 31039

chimp.LOC112206744-LOC107973052-GGT2.revcompl                               ------------------------------------------------------------ 61762
chimp54546-135457.revcompl.Pan.troglodytes.clone.rp43-41g5.GenBank:AC099533.36 ------------------------------------------------------------ 70187
human.FAM30B-LOC105372935-GGT2.NCBI.                                        atgttctgcatgattccatttatgtcagaaaaaatatatttgtgtaagtatatacacaga 103993
human.FAM230E-LOC105377182-GGT3P.NCBI.                                      ------------------------------------------------------------ 57310
human.FAM230J-LOC105372942-GGTLC5P.NCBI.rev.compl                           ------------------------------------------------------------ 46364
human.FAM230A.ncRNA.18487127.18500594-GGTLC3.18516335.18518165.rev.compl.REF ------------------------------------------------------------ 31039

chimp.LOC112206744-LOC107973052-GGT2.revcompl                               ------------------------------------------------------------ 61762
chimp54546-135457.revcompl.Pan.troglodytes.clone.rp43-41g5.GenBank:AC099533.36 ------------------------------------------------------------ 70187
human.FAM30B-LOC105372935-GGT2.NCBI.                                        aaagggtctgggagaatatataccaaactataaagaatctatatcaaagtcaataagggg 104053
human.FAM230E-LOC105377182-GGT3P.NCBI.                                      ------------------------------------------------------------ 57310
human.FAM230J-LOC105372942-GGTLC5P.NCBI.rev.compl                           ------------------------------------------------------------ 46364
human.FAM230A.ncRNA.18487127.18500594-GGTLC3.18516335.18518165.rev.compl.REF ------------------------------------------------------------ 31039

chimp.LOC112206744-LOC107973052-GGT2.revcompl                               ------------------------------------------------------------ 61762
chimp54546-135457.revcompl.Pan.troglodytes.clone.rp43-41g5.GenBank:AC099533.36 ----------------------------------------TTTTTTTTTTTTTTTTGGAG 70207
human.FAM30B-LOC105372935-GGT2.NCBI.                                        catagacaaggaggggaactttcacattttactttccacattctttttttttttttttttg 104113
human.FAM230E-LOC105377182-GGT3P.NCBI.                                      ------------------------------------------------------------ 57310
human.FAM230J-LOC105372942-GGTLC5P.NCBI.rev.compl                           ------------------------------------------------------------ 46364
human.FAM230A.ncRNA.18487127.18500594-GGTLC3.18516335.18518165.rev.compl.REF ------------------------------------------------------------ 31039

chimp.LOC112206744-LOC107973052-GGT2.revcompl                               ------------------------------------------------------------ 61762
chimp54546-135457.revcompl.Pan.troglodytes.clone.rp43-41g5.GenBank:AC099533.36 ATAGAGCCTCACTCTGTCTCCCAGGCTGGAGTGCAATGATGTGATCTTGGGTTACAGCAA 70267
human.FAM30B-LOC105372935-GGT2.NCBI.                                        acggagtctcactctgttgcccaggctggagtgcagtggcgcaatcttggctcactgcaa 104173
human.FAM230E-LOC105377182-GGT3P.NCBI.                                      ------------------------------------------------------------ 57310
human.FAM230J-LOC105372942-GGTLC5P.NCBI.rev.compl                           ------------------------------------------------------------ 46364
human.FAM230A.ncRNA.18487127.18500594-GGTLC3.18516335.18518165.rev.compl.REF ------------------------------------------------------------ 31039

chimp.LOC112206744-LOC107973052-GGT2.revcompl                               ------------------------------------------------------------ 61762
chimp54546-135457.revcompl.Pan.troglodytes.clone.rp43-41g5.GenBank:AC099533.36 TCTCTCCCTTCTAGGCTCAAGCGATTCTCCCACCTCAGCATCCCAAGTAGCTGGGACTAC 70327
human.FAM30B-LOC105372935-GGT2.NCBI.                                        gctccgcctcccgggttcacaccattctcctgcctcagcctcccgagtagctgggactac 104233
human.FAM230E-LOC105377182-GGT3P.NCBI.                                      ------------------------------------------------------------ 57310
human.FAM230J-LOC105372942-GGTLC5P.NCBI.rev.compl                           ------------------------------------------------------------ 46364
human.FAM230A.ncRNA.18487127.18500594-GGTLC3.18516335.18518165.rev.compl.REF ------------------------------------------------------------ 31039

chimp.LOC112206744-LOC107973052-GGT2.revcompl                               ------------------------------------------------------------ 61762
chimp54546-135457.revcompl.Pan.troglodytes.clone.rp43-41g5.GenBank:AC099533.36 AAGTGTGCCAACTTAGAATTATATTAGCCACACCCAGCTAATTTTTGTATTTTTTGTAGA 70387
human.FAM30B-LOC105372935-GGT2.NCBI.                                        aggcacccgc---------------caccacgcccagctaatttttttatattt------ 104272
human.FAM230E-LOC105377182-GGT3P.NCBI.                                      ------------------------------------------------------------ 57310
human.FAM230J-LOC105372942-GGTLC5P.NCBI.rev.compl                           ------------------------------------------------------------ 46364
human.FAM230A.ncRNA.18487127.18500594-GGTLC3.18516335.18518165.rev.compl.REF ------------------------------------------------------------ 31039

chimp.LOC112206744-LOC107973052-GGT2.revcompl                               ------------------------------------------------------------ 61762
chimp54546-135457.revcompl.Pan.troglodytes.clone.rp43-41g5.GenBank:AC099533.36 GGCAGGGTCTCGCCATGTTGCCCAGGCTGGTCTTGAACTCCTGGGCTTCAGTGATCCACC 70447
human.FAM30B-LOC105372935-GGT2.NCBI.                                        ---ggggtttcactgcgttagccaggatggtctcaatctcctgacctt--gtgatctgcc 104327
human.FAM230E-LOC105377182-GGT3P.NCBI.                                      ------------------------------------------------------------ 57310
human.FAM230J-LOC105372942-GGTLC5P.NCBI.rev.compl                           ------------------------------------------------------------ 46364
human.FAM230A.ncRNA.18487127.18500594-GGTLC3.18516335.18518165.rev.compl.REF ------------------------------------------------------------ 31039

chimp.LOC112206744-LOC107973052-GGT2.revcompl                               ------------------------------------------------------------ 61762
chimp54546-135457.revcompl.Pan.troglodytes.clone.rp43-41g5.GenBank:AC099533.36 CACCTCAACCTCC-------AACAAAGTGCCAAGATTACAGCCATGAGCCACCATGCCCA 70500
human.FAM30B-LOC105372935-GGT2.NCBI.                                        cacctcacccacctcagcctcccaaagtgctgggattacaggcgtgactttctacattct 104387
human.FAM230E-LOC105377182-GGT3P.NCBI.                                      ------------------------------------------------------------ 57310
human.FAM230J-LOC105372942-GGTLC5P.NCBI.rev.compl                           ------------------------------------------------------------ 46364
human.FAM230A.ncRNA.18487127.18500594-GGTLC3.18516335.18518165.rev.compl.REF ------------------------------------------------------------ 31039

chimp.LOC112206744-LOC107973052-GGT2.revcompl                               ------------------------------------------------------------ 61762
chimp54546-135457.revcompl.Pan.troglodytes.clone.rp43-41g5.GenBank:AC099533.36 GCATAACTATTTTTAATGAAGTAGACTTTAAGAAGAAAAGTATTATTAGAGGTAAGAGAC 70560
human.FAM30B-LOC105372935-GGT2.NCBI.                                        tgtaacagaattatagtcaccgttacctccatcaaaaaattaaaattcaa---------- 104437
human.FAM230E-LOC105377182-GGT3P.NCBI.                                      ------------------------------------------------------------ 57310
human.FAM230J-LOC105372942-GGTLC5P.NCBI.rev.compl                           ------------------------------------------------------------ 46364
human.FAM230A.ncRNA.18487127.18500594-GGTLC3.18516335.18518165.rev.compl.REF ------------------------------------------------------------ 31039

chimp.LOC112206744-LOC107973052-GGT2.revcompl                               ------------------------------------------------------------ 61762
chimp54546-135457.revcompl.Pan.troglodytes.clone.rp43-41g5.GenBank:AC099533.36 ACATCATGGAAAAGAAGAATTTACTAGGAGCCAGGCGCGGTGGCTCATGCCTGTAATTCC 70620
human.FAM30B-LOC105372935-GGT2.NCBI.                                        ----------------------aaattaggccaggcacggtggctcacgcctgtaatccc 104475
human.FAM230E-LOC105377182-GGT3P.NCBI.                                      ------------------------------------------------------------ 57310
human.FAM230J-LOC105372942-GGTLC5P.NCBI.rev.compl                           ------------------------------------------------------------ 46364
human.FAM230A.ncRNA.18487127.18500594-GGTLC3.18516335.18518165.rev.compl.REF ------------------------------------------------------------ 31039

chimp.LOC112206744-LOC107973052-GGT2.revcompl                               ------------------------------------------------------------ 61762
chimp54546-135457.revcompl.Pan.troglodytes.clone.rp43-41g5.GenBank:AC099533.36 AGCACTTTGTGAGGCCAAGGCGGCGGATCACCTGAGGTTGGGAGTTCAACACCAGCCTGA 70680
human.FAM30B-LOC105372935-GGT2.NCBI.                                        agcactttgggagtctggagcagcagatcacctgaggtcaggagttggagagcagcctga 104535
human.FAM230E-LOC105377182-GGT3P.NCBI.                                      ------------------------------------------------------------ 57310
```

```
human.FAM230J-LOC105372942-GGTLC5P.NCBI.rev.compl                               ------------------------------------------------------------ 46364
human.FAM230A.ncRNA.18487127.18500594-GGTLC3.18516335.18518165.rev.compl.REF     ------------------------------------------------------------ 31039

chimp.LOC112206744-LOC107973052-GGT2.revcompl                                   ------------------------------------------------------------ 61762
chimp54546-135457.revcompl.Pan.troglodytes.clone.rp43-41g5.GenBank:AC099533.36  ----CCAACATGGAGAAGCCCTGTCTCTACTAAAAATACAAAAACTAGCCAAGCATGGTG 70736
human.FAM30B-LOC105372935-GGT2.NCBI.                                            ctggccaacatggtgaaaccccatctctactaaaaatacaaaaattagccgggcgtggtg 104595
human.FAM230E-LOC105377182-GGT3P.NCBI.                                          ------------------------------------------------------------ 57310
human.FAM230J-LOC105372942-GGTLC5P.NCBI.rev.compl                               ------------------------------------------------------------ 46364
human.FAM230A.ncRNA.18487127.18500594-GGTLC3.18516335.18518165.rev.compl.REF     ------------------------------------------------------------ 31039

chimp.LOC112206744-LOC107973052-GGT2.revcompl                                   ------------------------------------------------------------ 61762
chimp54546-135457.revcompl.Pan.troglodytes.clone.rp43-41g5.GenBank:AC099533.36  GCATATGCCTGTAATCCCAGCTACTCAGGAGGCTGAGGGAAGAGAATTGCTTGGACCCAG 70796
human.FAM30B-LOC105372935-GGT2.NCBI.                                            gagggcacctgtaatcccagctacttgggaggctaaggcaggagaatcacttaaacccgg 104655
human.FAM230E-LOC105377182-GGT3P.NCBI.                                          ------------------------------------------------------------ 57310
human.FAM230J-LOC105372942-GGTLC5P.NCBI.rev.compl                               ------------------------------------------------------------ 46364
human.FAM230A.ncRNA.18487127.18500594-GGTLC3.18516335.18518165.rev.compl.REF     ------------------------------------------------------------ 31039

chimp.LOC112206744-LOC107973052-GGT2.revcompl                                   ------------------------------------------------------------ 61762
chimp54546-135457.revcompl.Pan.troglodytes.clone.rp43-41g5.GenBank:AC099533.36  GAAGTGGAGATTGAGGTGAGCTGAGATTGTGCCATTGCATTCCAGCCTGGGCAACAAGAG 70856
human.FAM30B-LOC105372935-GGT2.NCBI.                                            gaggcggaggttgcagtgagccaagattgtgccactgcactccggcccgggtaac-agag 104714
human.FAM230E-LOC105377182-GGT3P.NCBI.                                          ------------------------------------------------------------ 57310
human.FAM230J-LOC105372942-GGTLC5P.NCBI.rev.compl                               ------------------------------------------------------------ 46364
human.FAM230A.ncRNA.18487127.18500594-GGTLC3.18516335.18518165.rev.compl.REF     ------------------------------------------------------------ 31039

chimp.LOC112206744-LOC107973052-GGT2.revcompl                                   ------------------------------------------------------------ 61762
chimp54546-135457.revcompl.Pan.troglodytes.clone.rp43-41g5.GenBank:AC099533.36  CAAAACTCTGTCTCAAAAAAAAAAAAAAAAAAAAAAAAAGGAA----------------- 70899
human.FAM30B-LOC105372935-GGT2.NCBI.                                            cgagactccatctcaaaaatgaatgaattaattaattaattaaagcaaacaaatataagc 104774
human.FAM230E-LOC105377182-GGT3P.NCBI.                                          ------------------------------------------------------------ 57310
human.FAM230J-LOC105372942-GGTLC5P.NCBI.rev.compl                               ------------------------------------------------------------ 46364
human.FAM230A.ncRNA.18487127.18500594-GGTLC3.18516335.18518165.rev.compl.REF     ------------------------------------------------------------ 31039

chimp.LOC112206744-LOC107973052-GGT2.revcompl                                   ------------------------------------------------------------ 61762
chimp54546-135457.revcompl.Pan.troglodytes.clone.rp43-41g5.GenBank:AC099533.36  ------------------------------------------------------------ 70899
human.FAM30B-LOC105372935-GGT2.NCBI.                                            tacctcctgaagtagaaaccagtatctcttcaccttacctttacataatactttacccta 104834
human.FAM230E-LOC105377182-GGT3P.NCBI.                                          ------------------------------------------------------------ 57310
human.FAM230J-LOC105372942-GGTLC5P.NCBI.rev.compl                               ------------------------------------------------------------ 46364
human.FAM230A.ncRNA.18487127.18500594-GGTLC3.18516335.18518165.rev.compl.REF     ------------------------------------------------------------ 31039

chimp.LOC112206744-LOC107973052-GGT2.revcompl                                   ------------------------------------------------------------ 61762
chimp54546-135457.revcompl.Pan.troglodytes.clone.rp43-41g5.GenBank:AC099533.36  ------------------------------------------------------------ 70899
human.FAM30B-LOC105372935-GGT2.NCBI.                                            atgcattttatctgctgttctaaatcatgacttacttcaaatcaaatcttaagtgaaaat 104894
human.FAM230E-LOC105377182-GGT3P.NCBI.                                          ------------------------------------------------------------ 57310
human.FAM230J-LOC105372942-GGTLC5P.NCBI.rev.compl                               ------------------------------------------------------------ 46364
human.FAM230A.ncRNA.18487127.18500594-GGTLC3.18516335.18518165.rev.compl.REF     ------------------------------------------------------------ 31039

chimp.LOC112206744-LOC107973052-GGT2.revcompl                                   ------------------------------------------------------------ 61762
chimp54546-135457.revcompl.Pan.troglodytes.clone.rp43-41g5.GenBank:AC099533.36  ------------------------------------------------------------ 70899
human.FAM30B-LOC105372935-GGT2.NCBI.                                            caagatttccttttggggggcattatacaaaaaagactgtaaaatttcattctctgaaagt 104954
human.FAM230E-LOC105377182-GGT3P.NCBI.                                          ------------------------------------------------------------ 57310
human.FAM230J-LOC105372942-GGTLC5P.NCBI.rev.compl                               ------------------------------------------------------------ 46364
human.FAM230A.ncRNA.18487127.18500594-GGTLC3.18516335.18518165.rev.compl.REF     ------------------------------------------------------------ 31039

chimp.LOC112206744-LOC107973052-GGT2.revcompl                                   ------------------------------------------------------------ 61762
chimp54546-135457.revcompl.Pan.troglodytes.clone.rp43-41g5.GenBank:AC099533.36  ------------------------------------------------------------ 70899
human.FAM30B-LOC105372935-GGT2.NCBI.                                            ctttttaaaaagggatagattgtatctggctgggatagggtaggtgtgatcggacccgga 105014
human.FAM230E-LOC105377182-GGT3P.NCBI.                                          ------------------------------------------------------------ 57310
human.FAM230J-LOC105372942-GGTLC5P.NCBI.rev.compl                               ------------------------------------------------------------ 46364
human.FAM230A.ncRNA.18487127.18500594-GGTLC3.18516335.18518165.rev.compl.REF     ------------------------------------------------------------ 31039

chimp.LOC112206744-LOC107973052-GGT2.revcompl                                   ------------------------------------------------------------ 61762
chimp54546-135457.revcompl.Pan.troglodytes.clone.rp43-41g5.GenBank:AC099533.36  ----------------------------------------GTTACTAGCTAGTTTCAGTA 70919
human.FAM30B-LOC105372935-GGT2.NCBI.                                            ggaaatgacggctgctcaactgtgcatctcctttcctcaccctgctgaaaagtgccacct 105074
human.FAM230E-LOC105377182-GGT3P.NCBI.                                          ------------------------------------------------------------ 57310
human.FAM230J-LOC105372942-GGTLC5P.NCBI.rev.compl                               ------------------------------------------------------------ 46364
human.FAM230A.ncRNA.18487127.18500594-GGTLC3.18516335.18518165.rev.compl.REF     ------------------------------------------------------------ 31039

chimp.LOC112206744-LOC107973052-GGT2.revcompl                                   ------------------------------------------------------------ 61762
chimp54546-135457.revcompl.Pan.troglodytes.clone.rp43-41g5.GenBank:AC099533.36  ATTCTTAACATCCAGGAAACTGGATGTGAAAGCTTTTCAGAGAAGCTAAACCAATAGGTT 70979
human.FAM30B-LOC105372935-GGT2.NCBI.                                            agcatggtcaatcaaggtcaccaatgcaaagcagtagtgggcaaacaccaccc--aggtt 105132
human.FAM230E-LOC105377182-GGT3P.NCBI.                                          ------------------------------------------------------------ 57310
human.FAM230J-LOC105372942-GGTLC5P.NCBI.rev.compl                               ------------------------------------------------------------ 46364
human.FAM230A.ncRNA.18487127.18500594-GGTLC3.18516335.18518165.rev.compl.REF     ------------------------------------------------------------ 31039

chimp.LOC112206744-LOC107973052-GGT2.revcompl                                   ------------------------------------------------------------ 61762
chimp54546-135457.revcompl.Pan.troglodytes.clone.rp43-41g5.GenBank:AC099533.36  ATACATAGAGAGAGATTTATTTAGGAATTGGCTTACATGATTGTGGGGACTAGCAAGTTT 71039
human.FAM30B-LOC105372935-GGT2.NCBI.                                            atccaaagcagcacatggggttgggaaactgatgaccttggagatttcaaaatcacctaa 105192
human.FAM230E-LOC105377182-GGT3P.NCBI.                                          ------------------------------------------------------------ 57310
human.FAM230J-LOC105372942-GGTLC5P.NCBI.rev.compl                               ------------------------------------------------------------ 46364
human.FAM230A.ncRNA.18487127.18500594-GGTLC3.18516335.18518165.rev.compl.REF     ------------------------------------------------------------ 31039

chimp.LOC112206744-LOC107973052-GGT2.revcompl                                   ------------------------------------------------------------ 61762
chimp54546-135457.revcompl.Pan.troglodytes.clone.rp43-41g5.GenBank:AC099533.36  AAAATCTGTAGGGCAAGCCAGCCGGCTATAAATTCAGGTAAGAGTTGATCTCGAAGTCTT 71099
human.FAM30B-LOC105372935-GGT2.NCBI.                                            caa----------taaccccaccttccctaaccacagggctaag--caacctaatggatg 105240
human.FAM230E-LOC105377182-GGT3P.NCBI.                                          ------------------------------------------------------------ 57310
human.FAM230J-LOC105372942-GGTLC5P.NCBI.rev.compl                               ------------------------------------------------------------ 46364
human.FAM230A.ncRNA.18487127.18500594-GGTLC3.18516335.18518165.rev.compl.REF     ------------------------------------------------------------ 31039

chimp.LOC112206744-LOC107973052-GGT2.revcompl                                   ------------------------------------------------------------ 61762
chimp54546-135457.revcompl.Pan.troglodytes.clone.rp43-41g5.GenBank:AC099533.36  GAACCTAAAATCTGTAGAGCAGTCAGCAGGCCAGAAACTCAGGCAGGGTTTG-------- 71151
human.FAM30B-LOC105372935-GGT2.NCBI.                                            gaacatatgtgtgctttggtcatccgcaacccagaccactagactaattttgctttagaa 105300
human.FAM230E-LOC105377182-GGT3P.NCBI.                                          ------------------------------------------------------------ 57310
human.FAM230J-LOC105372942-GGTLC5P.NCBI.rev.compl                               ------------------------------------------------------------ 46364
human.FAM230A.ncRNA.18487127.18500594-GGTLC3.18516335.18518165.rev.compl.REF     ------------------------------------------------------------ 31039

chimp.LOC112206744-LOC107973052-GGT2.revcompl                                   ------------------------------------------------------------ 61762
chimp54546-135457.revcompl.Pan.troglodytes.clone.rp43-41g5.GenBank:AC099533.36  ---------------------------------------TGTGTTACAGTCTTGAAGCAG 71172
human.FAM30B-LOC105372935-GGT2.NCBI.                                            ctggactatatcaagtgtgaacgatcatacttgttatcatgtgggaaaaaaggtttaatt 105360
human.FAM230E-LOC105377182-GGT3P.NCBI.                                          ------------------------------------------------------------ 57310
human.FAM230J-LOC105372942-GGTLC5P.NCBI.rev.compl                               ------------------------------------------------------------ 46364
human.FAM230A.ncRNA.18487127.18500594-GGTLC3.18516335.18518165.rev.compl.REF     ------------------------------------------------------------ 31039

chimp.LOC112206744-LOC107973052-GGT2.revcompl                                   ------------------------------------------------------------ 61762
chimp54546-135457.revcompl.Pan.troglodytes.clone.rp43-41g5.GenBank:AC099533.36  AATTCCTGCTTCTCTGGGAAACCTCAGTTTT-----------TGTTCTTAAGGCCTTCAA 71221
human.FAM30B-LOC105372935-GGT2.NCBI.                                            actccctaattgtctgggaaacaaaagatttcaaaatatggagctcctggggggaaggaga 105420
human.FAM230E-LOC105377182-GGT3P.NCBI.                                          ------------------------------------------------------------ 57310
human.FAM230J-LOC105372942-GGTLC5P.NCBI.rev.compl                               ------------------------------------------------------------ 46364
human.FAM230A.ncRNA.18487127.18500594-GGTLC3.18516335.18518165.rev.compl.REF     ------------------------------------------------------------ 31039

chimp.LOC112206744-LOC107973052-GGT2.revcompl                                   ------------------------------------------------------------ 61762
chimp54546-135457.revcompl.Pan.troglodytes.clone.rp43-41g5.GenBank:AC099533.36  CTGATTGGAGGTGGCCCACCCATATTATGGTGGGTAATCTGTTTTACTTAAAGTCAACTG 71281
human.FAM30B-LOC105372935-GGT2.NCBI.                                            gttaataaaatgggtccttgagaagtgaagtgggaagcctgcctcatgccatggcaccag 105480
human.FAM230E-LOC105377182-GGT3P.NCBI.                                          ------------------------------------------------------------ 57310
human.FAM230J-LOC105372942-GGTLC5P.NCBI.rev.compl                               ------------------------------------------------------------ 46364
human.FAM230A.ncRNA.18487127.18500594-GGTLC3.18516335.18518165.rev.compl.REF     ------------------------------------------------------------ 31039

chimp.LOC112206744-LOC107973052-GGT2.revcompl                                   ------------------------------------------------------------ 61762
chimp54546-135457.revcompl.Pan.troglodytes.clone.rp43-41g5.GenBank:AC099533.36  AC---------------------------------------------TGTCAGTGTTAAT 71296
human.FAM30B-LOC105372935-GGT2.NCBI.                                            cgatgccacaggtgtacccagtgccctgccgagagcagagacggtgttctcagagttgct 105540
human.FAM230E-LOC105377182-GGT3P.NCBI.                                          ------------------------------------------------------------ 57310
human.FAM230J-LOC105372942-GGTLC5P.NCBI.rev.compl                               ------------------------------------------------------------ 46364
human.FAM230A.ncRNA.18487127.18500594-GGTLC3.18516335.18518165.rev.compl.REF     ------------------------------------------------------------ 31039

chimp.LOC112206744-LOC107973052-GGT2.revcompl                                   ------------------------------------------------------------ 61762
chimp54546-135457.revcompl.Pan.troglodytes.clone.rp43-41g5.GenBank:AC099533.36  CACATCTATGAAATAACCTCC------CAGCAAGATATTGACAAGTATTTGACCAAACAA 71350
human.FAM30B-LOC105372935-GGT2.NCBI.                                            atcaaaaaggaggcagcagactgcattctgaagcatcagacaatgggtcacatgcggaaa 105600
human.FAM230E-LOC105377182-GGT3P.NCBI.                                          ------------------------------------------------------------ 57310
human.FAM230J-LOC105372942-GGTLC5P.NCBI.rev.compl                               ------------------------------------------------------------ 46364
```

```
human.FAM230A.ncRNA.18487127.18500594-GGTLC3.18516335.18518165.rev.compl.REF      ------------------------------------------------------------ 31039

chimp.LOC112206744-LOC107973052-GGT2.revcompl                                     ------------------------------------------------------------ 61762
chimp54546-135457.revcompl.Pan.troglodytes.clone.rp43-41g5.GenBank:AC099533.36    CGGGACACCATAGCTTAGCCAAGTTGACACATAAATTAACCATCAGGAGCAAGTAGAATA 71410
human.FAM30B-LOC105372935-GGT2.NCBI.                                              cagaattaaatacacaaaaatagcaccaacttaaggcaagcctgggcaaactgcactaca 105660
human.FAM230E-LOC105377182-GGT3P.NCBI.                                            ------------------------------------------------------------ 57310
human.FAM230J-LOC105372942-GGTLC5P.NCBI.rev.compl                                 ------------------------------------------------------------ 46364
human.FAM230A.ncRNA.18487127.18500594-GGTLC3.18516335.18518165.rev.compl.REF      ------------------------------------------------------------ 31039

chimp.LOC112206744-LOC107973052-GGT2.revcompl                                     ------------------------------------------------------------ 61762
chimp54546-135457.revcompl.Pan.troglodytes.clone.rp43-41g5.GenBank:AC099533.36    TCCAAAAAACAACATACTAGGGGTATTATGTCTTATATAGCTATTATAATTATATAAAAC 71470
human.FAM30B-LOC105372935-GGT2.NCBI.                                              tcaaagaaagaacacacaactggtatcagatttttttttctgttttaataaagagaaggc 105720
human.FAM230E-LOC105377182-GGT3P.NCBI.                                            ------------------------------------------------------------ 57310
human.FAM230J-LOC105372942-GGTLC5P.NCBI.rev.compl                                 ------------------------------------------------------------ 46364
human.FAM230A.ncRNA.18487127.18500594-GGTLC3.18516335.18518165.rev.compl.REF      ------------------------------------------------------------ 31039

chimp.LOC112206744-LOC107973052-GGT2.revcompl                                     ------------------------------------------------------------ 61762
chimp54546-135457.revcompl.Pan.troglodytes.clone.rp43-41g5.GenBank:AC099533.36    AT--------------------------------------------------ATAATTAT 71480
human.FAM30B-LOC105372935-GGT2.NCBI.                                              ctggacacctgaaagttgacgggaagcaggcatcactaatttctccgagttcacagtttt 105780
human.FAM230E-LOC105377182-GGT3P.NCBI.                                            ------------------------------------------------------------ 57310
human.FAM230J-LOC105372942-GGTLC5P.NCBI.rev.compl                                 ------------------------------------------------------------ 46364
human.FAM230A.ncRNA.18487127.18500594-GGTLC3.18516335.18518165.rev.compl.REF      ------------------------------------------------------------ 31039

chimp.LOC112206744-LOC107973052-GGT2.revcompl                                     ------------------------------------------------------------ 61762
chimp54546-135457.revcompl.Pan.troglodytes.clone.rp43-41g5.GenBank:AC099533.36    AGAATGAAGATATTAAGATAACCATTAGAACAAAAATATAAACTTTTCTTTCTTTCTTTT 71540
human.FAM30B-LOC105372935-GGT2.NCBI.                                              gttagaatgtctttgctctgattatggttttatttatttatttttttgagatggagtttt 105840
human.FAM230E-LOC105377182-GGT3P.NCBI.                                            ------------------------------------------------------------ 57310
human.FAM230J-LOC105372942-GGTLC5P.NCBI.rev.compl                                 ------------------------------------------------------------ 46364
human.FAM230A.ncRNA.18487127.18500594-GGTLC3.18516335.18518165.rev.compl.REF      ------------------------------------------------------------ 31039

chimp.LOC112206744-LOC107973052-GGT2.revcompl                                     ------------------------------------------------------------ 61762
chimp54546-135457.revcompl.Pan.troglodytes.clone.rp43-41g5.GenBank:AC099533.36    TTTTTTTTTGAGACCAAGTCTTGCTCTGTCACCCAGGCTGGAGTGCAGTGGTGCAATCTT 71600
human.FAM30B-LOC105372935-GGT2.NCBI.                                              ttttgttttgtttt---atttgcttttgttgccctggctggagtgcagtggtgcaatctc 105897
human.FAM230E-LOC105377182-GGT3P.NCBI.                                            ------------------------------------------------------------ 57310
human.FAM230J-LOC105372942-GGTLC5P.NCBI.rev.compl                                 ------------------------------------------------------------ 46364
human.FAM230A.ncRNA.18487127.18500594-GGTLC3.18516335.18518165.rev.compl.REF      ------------------------------------------------------------ 31039

chimp.LOC112206744-LOC107973052-GGT2.revcompl                                     ------------------------------------------------------------ 61762
chimp54546-135457.revcompl.Pan.troglodytes.clone.rp43-41g5.GenBank:AC099533.36    GGCTTACTGCAACCTTTGCCTCCTGGGTTCAAGTGATTCTCCTGTCTCAGCCTCCCAAGT 71660
human.FAM30B-LOC105372935-GGT2.NCBI.                                              ggctcaccgcaaactccgcctcccaggttcaagagattctcctgcctcagcctcccaagt 105957
human.FAM230E-LOC105377182-GGT3P.NCBI.                                            ------------------------------------------------------------ 57310
human.FAM230J-LOC105372942-GGTLC5P.NCBI.rev.compl                                 ------------------------------------------------------------ 46364
human.FAM230A.ncRNA.18487127.18500594-GGTLC3.18516335.18518165.rev.compl.REF      ------------------------------------------------------------ 31039

chimp.LOC112206744-LOC107973052-GGT2.revcompl                                     ------------------------------------------------------------ 61762
chimp54546-135457.revcompl.Pan.troglodytes.clone.rp43-41g5.GenBank:AC099533.36    AGCTGGGATTACAGGCACCCGCTACCATGCCCAAGCATCCACCATGTGGTGGTATTTGGA 71720
human.FAM30B-LOC105372935-GGT2.NCBI.                                              agctaggattacaggcacccactacca--------cgtctggcttctttgtgtttttagt 106009
human.FAM230E-LOC105377182-GGT3P.NCBI.                                            ------------------------------------------------------------ 57310
human.FAM230J-LOC105372942-GGTLC5P.NCBI.rev.compl                                 ------------------------------------------------------------ 46364
human.FAM230A.ncRNA.18487127.18500594-GGTLC3.18516335.18518165.rev.compl.REF      ------------------------------------------------------------ 31039

chimp.LOC112206744-LOC107973052-GGT2.revcompl                                     ------------------------------------------------------------ 61762
chimp54546-135457.revcompl.Pan.troglodytes.clone.rp43-41g5.GenBank:AC099533.36    TAAGATGTGCAAGCA--------------------------------------------- 71735
human.FAM30B-LOC105372935-GGT2.NCBI.                                              agagatggggtttcaccatgttggtcaggctggtcttgaactcttgacctcaagcgatcc 106069
human.FAM230E-LOC105377182-GGT3P.NCBI.                                            ------------------------------------------------------------ 57310
human.FAM230J-LOC105372942-GGTLC5P.NCBI.rev.compl                                 ------------------------------------------------------------ 46364
human.FAM230A.ncRNA.18487127.18500594-GGTLC3.18516335.18518165.rev.compl.REF      ------------------------------------------------------------ 31039

chimp.LOC112206744-LOC107973052-GGT2.revcompl                                     ------------------------------------------------------------ 61762
chimp54546-135457.revcompl.Pan.troglodytes.clone.rp43-41g5.GenBank:AC099533.36    ------------------------------------------------------------ 71735
human.FAM30B-LOC105372935-GGT2.NCBI.                                              acccgcctcagcccccaaagtgctggaattacaggcatgagccaccacgcctggcctga 106129
human.FAM230E-LOC105377182-GGT3P.NCBI.                                            ------------------------------------------------------------ 57310
human.FAM230J-LOC105372942-GGTLC5P.NCBI.rev.compl                                 ------------------------------------------------------------ 46364
human.FAM230A.ncRNA.18487127.18500594-GGTLC3.18516335.18518165.rev.compl.REF      ------------------------------------------------------------ 31039

chimp.LOC112206744-LOC107973052-GGT2.revcompl                                     ------------------------------------------------------------ 61762
chimp54546-135457.revcompl.Pan.troglodytes.clone.rp43-41g5.GenBank:AC099533.36    ------------------------------------------------------------ 71735
human.FAM30B-LOC105372935-GGT2.NCBI.                                              ttatgttttgatggatatacagaagctcttgctttggtacagaaacaggccccgctttgc 106189
human.FAM230E-LOC105377182-GGT3P.NCBI.                                            ------------------------------------------------------------ 57310
human.FAM230J-LOC105372942-GGTLC5P.NCBI.rev.compl                                 ------------------------------------------------------------ 46364
human.FAM230A.ncRNA.18487127.18500594-GGTLC3.18516335.18518165.rev.compl.REF      ------------------------------------------------------------ 31039

chimp.LOC112206744-LOC107973052-GGT2.revcompl                                     ------------------------------------------------------------ 61762
chimp54546-135457.revcompl.Pan.troglodytes.clone.rp43-41g5.GenBank:AC099533.36    ------------------------------------------------------------ 71735
human.FAM30B-LOC105372935-GGT2.NCBI.                                              ttcgacctctgggcccccaccttcagaaatgtctggtgccattcctggctcctcctctc 106249
human.FAM230E-LOC105377182-GGT3P.NCBI.                                            ------------------------------------------------------------ 57310
human.FAM230J-LOC105372942-GGTLC5P.NCBI.rev.compl                                 ------------------------------------------------------------ 46364
human.FAM230A.ncRNA.18487127.18500594-GGTLC3.18516335.18518165.rev.compl.REF      ------------------------------------------------------------ 31039

chimp.LOC112206744-LOC107973052-GGT2.revcompl                                     ------------------------------------------------------------ 61762
chimp54546-135457.revcompl.Pan.troglodytes.clone.rp43-41g5.GenBank:AC099533.36    ------------------------------------------------------------ 71735
human.FAM30B-LOC105372935-GGT2.NCBI.                                              agtggtcttccaggatatccactcacagctttctgatgtcacccacctgtctctagccca 106309
human.FAM230E-LOC105377182-GGT3P.NCBI.                                            ------------------------------------------------------------ 57310
human.FAM230J-LOC105372942-GGTLC5P.NCBI.rev.compl                                 ------------------------------------------------------------ 46364
human.FAM230A.ncRNA.18487127.18500594-GGTLC3.18516335.18518165.rev.compl.REF      ------------------------------------------------------------ 31039

chimp.LOC112206744-LOC107973052-GGT2.revcompl                                     ------------------------------------------------------------ 61762
chimp54546-135457.revcompl.Pan.troglodytes.clone.rp43-41g5.GenBank:AC099533.36    ------------------------------------------------------------ 71735
human.FAM30B-LOC105372935-GGT2.NCBI.                                              gatacccagtcctcatggcaagtgactgcagcaatttttacaaatctacgcccttagggt 106369
human.FAM230E-LOC105377182-GGT3P.NCBI.                                            ------------------------------------------------------------ 57310
human.FAM230J-LOC105372942-GGTLC5P.NCBI.rev.compl                                 ------------------------------------------------------------ 46364
human.FAM230A.ncRNA.18487127.18500594-GGTLC3.18516335.18518165.rev.compl.REF      ------------------------------------------------------------ 31039

chimp.LOC112206744-LOC107973052-GGT2.revcompl                                     ------------------------------------------------------------ 61762
chimp54546-135457.revcompl.Pan.troglodytes.clone.rp43-41g5.GenBank:AC099533.36    ------------------------------------------------------------ 71735
human.FAM30B-LOC105372935-GGT2.NCBI.                                              tcccatcccaggacgttcaggtttctccacccctttagaagtgcccagaatatcccctac 106429
human.FAM230E-LOC105377182-GGT3P.NCBI.                                            ------------------------------------------------------------ 57310
human.FAM230J-LOC105372942-GGTLC5P.NCBI.rev.compl                                 ------------------------------------------------------------ 46364
human.FAM230A.ncRNA.18487127.18500594-GGTLC3.18516335.18518165.rev.compl.REF      ------------------------------------------------------------ 31039

chimp.LOC112206744-LOC107973052-GGT2.revcompl                                     ------------------------------------------------------------ 61762
chimp54546-135457.revcompl.Pan.troglodytes.clone.rp43-41g5.GenBank:AC099533.36    ------------------------------------------------------------ 71735
human.FAM30B-LOC105372935-GGT2.NCBI.                                              cccacgattccaacaggacgtcactctcctcttacttaagaaaaatgcctaaaattccag 106489
human.FAM230E-LOC105377182-GGT3P.NCBI.                                            ------------------------------------------------------------ 57310
human.FAM230J-LOC105372942-GGTLC5P.NCBI.rev.compl                                 ------------------------------------------------------------ 46364
human.FAM230A.ncRNA.18487127.18500594-GGTLC3.18516335.18518165.rev.compl.REF      ------------------------------------------------------------ 31039

chimp.LOC112206744-LOC107973052-GGT2.revcompl                                     ------------------------------------------------------------ 61762
chimp54546-135457.revcompl.Pan.troglodytes.clone.rp43-41g5.GenBank:AC099533.36    ------------------------------------------------------------ 71735
human.FAM30B-LOC105372935-GGT2.NCBI.                                              cccttccaaaaggcctttctagattgaccactcaggagttactctggcctccccttcact 106549
human.FAM230E-LOC105377182-GGT3P.NCBI.                                            ------------------------------------------------------------ 57310
human.FAM230J-LOC105372942-GGTLC5P.NCBI.rev.compl                                 ------------------------------------------------------------ 46364
human.FAM230A.ncRNA.18487127.18500594-GGTLC3.18516335.18518165.rev.compl.REF      ------------------------------------------------------------ 31039

chimp.LOC112206744-LOC107973052-GGT2.revcompl                                     ------------------------------------------------------------ 61762
chimp54546-135457.revcompl.Pan.troglodytes.clone.rp43-41g5.GenBank:AC099533.36    ------------------------------------------------------------ 71735
human.FAM30B-LOC105372935-GGT2.NCBI.                                              cctgctagactgaaaacaaggacagacttgcctggcctctctcccccagtagacacctac 106609
human.FAM230E-LOC105377182-GGT3P.NCBI.                                            ------------------------------------------------------------ 57310
human.FAM230J-LOC105372942-GGTLC5P.NCBI.rev.compl                                 ------------------------------------------------------------ 46364
human.FAM230A.ncRNA.18487127.18500594-GGTLC3.18516335.18518165.rev.compl.REF      ------------------------------------------------------------ 31039

chimp.LOC112206744-LOC107973052-GGT2.revcompl                                     ------------------------------------------------------------ 61762
chimp54546-135457.revcompl.Pan.troglodytes.clone.rp43-41g5.GenBank:AC099533.36    ------------------------------------------------------------ 71735
human.FAM30B-LOC105372935-GGT2.NCBI.                                              ccccagtgaacacatcctgtcagtgtcctcccgtccatttgcaaattgattgactgtttc 106669
human.FAM230E-LOC105377182-GGT3P.NCBI.                                            ------------------------------------------------------------ 57310
human.FAM230J-LOC105372942-GGTLC5P.NCBI.rev.compl                                 ------------------------------------------------------------ 46364
human.FAM230A.ncRNA.18487127.18500594-GGTLC3.18516335.18518165.rev.compl.REF      ------------------------------------------------------------ 31039
```

```
chimp.LOC112206744-LOC107973052-GGT2.revcompl                                ------------------------------------------------------------ 61762
chimp54546-135457.revcompl.Pan.troglodytes.clone.rp43-41g5.GenBank:AC099533.36 ------------------------------------------------------------ 71735
human.FAM30B-LOC105372935-GGT2.NCBI.                                         attgctggccttggatgacctctcctagagggcagggacggtggtgttcggcaaagcctt 106729
human.FAM230E-LOC105377182-GGT3P.NCBI.                                       ------------------------------------------------------------ 57310
human.FAM230J-LOC105372942-GGTLC5P.NCBI.rev.compl                            ------------------------------------------------------------ 46364
human.FAM230A.ncRNA.18487127.18500594-GGTLC3.18516335.18518165.rev.compl.REF ------------------------------------------------------------ 31039

chimp.LOC112206744-LOC107973052-GGT2.revcompl                                ------------------------------------------------------------ 61762
chimp54546-135457.revcompl.Pan.troglodytes.clone.rp43-41g5.GenBank:AC099533.36 ---------------------------------GTGAAAGGTCATTGGAAGATTTGGGAG 71763
human.FAM30B-LOC105372935-GGT2.NCBI.                                         ggtcgtctggataatgggaaatggggtcttctggtgaatttaattacagggttaggtgga 106789
human.FAM230E-LOC105377182-GGT3P.NCBI.                                       ------------------------------------------------------------ 57310
human.FAM230J-LOC105372942-GGTLC5P.NCBI.rev.compl                            ------------------------------------------------------------ 46364
human.FAM230A.ncRNA.18487127.18500594-GGTLC3.18516335.18518165.rev.compl.REF ------------------------------------------------------------ 31039

chimp.LOC112206744-LOC107973052-GGT2.revcompl                                ------------------------------------------------------------ 61762
chimp54546-135457.revcompl.Pan.troglodytes.clone.rp43-41g5.GenBank:AC099533.36 ACCCATGCATTCA----------------------------------------------- 71776
human.FAM30B-LOC105372935-GGT2.NCBI.                                         aactttgcttcaactctcattattagaaagtctattgtaaagctttttaaaatacaactg 106849
human.FAM230E-LOC105377182-GGT3P.NCBI.                                       ------------------------------------------------------------ 57310
human.FAM230J-LOC105372942-GGTLC5P.NCBI.rev.compl                            ------------------------------------------------------------ 46364
human.FAM230A.ncRNA.18487127.18500594-GGTLC3.18516335.18518165.rev.compl.REF ------------------------------------------------------------ 31039

chimp.LOC112206744-LOC107973052-GGT2.revcompl                                ------------------------------------------------------------ 61762
chimp54546-135457.revcompl.Pan.troglodytes.clone.rp43-41g5.GenBank:AC099533.36 ------------------------------------------------------------ 71776
human.FAM30B-LOC105372935-GGT2.NCBI.                                         accactttccagcgagggtagagttaatatgccttaatctttgatttaggatcttgcagt 106909
human.FAM230E-LOC105377182-GGT3P.NCBI.                                       ------------------------------------------------------------ 57310
human.FAM230J-LOC105372942-GGTLC5P.NCBI.rev.compl                            ------------------------------------------------------------ 46364
human.FAM230A.ncRNA.18487127.18500594-GGTLC3.18516335.18518165.rev.compl.REF ------------------------------------------------------------ 31039

chimp.LOC112206744-LOC107973052-GGT2.revcompl                                ------------------------------------------------------------ 61762
chimp54546-135457.revcompl.Pan.troglodytes.clone.rp43-41g5.GenBank:AC099533.36 ------------------------------------------------------------ 71776
human.FAM30B-LOC105372935-GGT2.NCBI.                                         gccccaggctcaccattctcttctgctactataattatagatgcggaagacatacagact 106969
human.FAM230E-LOC105377182-GGT3P.NCBI.                                       ------------------------------------------------------------ 57310
human.FAM230J-LOC105372942-GGTLC5P.NCBI.rev.compl                            ------------------------------------------------------------ 46364
human.FAM230A.ncRNA.18487127.18500594-GGTLC3.18516335.18518165.rev.compl.REF ------------------------------------------------------------ 31039

chimp.LOC112206744-LOC107973052-GGT2.revcompl                                ------------------------------------------------------------ 61762
chimp54546-135457.revcompl.Pan.troglodytes.clone.rp43-41g5.GenBank:AC099533.36 ------------------------------------------------------------ 71776
human.FAM30B-LOC105372935-GGT2.NCBI.                                         gggctgacaactgcttggaagctaatttctagaataaacccttgctaccacatcttgtga 107029
human.FAM230E-LOC105377182-GGT3P.NCBI.                                       ------------------------------------------------------------ 57310
human.FAM230J-LOC105372942-GGTLC5P.NCBI.rev.compl                            ------------------------------------------------------------ 46364
human.FAM230A.ncRNA.18487127.18500594-GGTLC3.18516335.18518165.rev.compl.REF ------------------------------------------------------------ 31039

chimp.LOC112206744-LOC107973052-GGT2.revcompl                                ------------------------------------------------------------ 61762
chimp54546-135457.revcompl.Pan.troglodytes.clone.rp43-41g5.GenBank:AC099533.36 ------------------------------------------------------------ 71776
human.FAM30B-LOC105372935-GGT2.NCBI.                                         tttttgcccttctgaaggtggagttccaaaataaaaacaaaatcccgcttaactgagggt 107089
human.FAM230E-LOC105377182-GGT3P.NCBI.                                       ------------------------------------------------------------ 57310
human.FAM230J-LOC105372942-GGTLC5P.NCBI.rev.compl                            ------------------------------------------------------------ 46364
human.FAM230A.ncRNA.18487127.18500594-GGTLC3.18516335.18518165.rev.compl.REF ------------------------------------------------------------ 31039

chimp.LOC112206744-LOC107973052-GGT2.revcompl                                ------------------------------------------------------------ 61762
chimp54546-135457.revcompl.Pan.troglodytes.clone.rp43-41g5.GenBank:AC099533.36 ------------------------------------------------------------ 71776
human.FAM30B-LOC105372935-GGT2.NCBI.                                         tccagttgagggaggtctgctgccatggtctttgttaagtgcctgccaccatggcccaca 107149
human.FAM230E-LOC105377182-GGT3P.NCBI.                                       ------------------------------------------------------------ 57310
human.FAM230J-LOC105372942-GGTLC5P.NCBI.rev.compl                            ------------------------------------------------------------ 46364
human.FAM230A.ncRNA.18487127.18500594-GGTLC3.18516335.18518165.rev.compl.REF ------------------------------------------------------------ 31039

chimp.LOC112206744-LOC107973052-GGT2.revcompl                                ------------------------------------------------------------ 61762
chimp54546-135457.revcompl.Pan.troglodytes.clone.rp43-41g5.GenBank:AC099533.36 -------------------------GAGAATAGGAAAGCAAACCTAGGGAGTCATCCTTC 71811
human.FAM30B-LOC105372935-GGT2.NCBI.                                         attagtcagttaataaagggttcttgatgagagaatttggcaacatgtgattttccacaa 107209
human.FAM230E-LOC105377182-GGT3P.NCBI.                                       ------------------------------------------------------------ 57310
human.FAM230J-LOC105372942-GGTLC5P.NCBI.rev.compl                            ------------------------------------------------------------ 46364
human.FAM230A.ncRNA.18487127.18500594-GGTLC3.18516335.18518165.rev.compl.REF ------------------------------------------------------------ 31039

chimp.LOC112206744-LOC107973052-GGT2.revcompl                                ------------------------------------------------------------ 61762
chimp54546-135457.revcompl.Pan.troglodytes.clone.rp43-41g5.GenBank:AC099533.36 ATATGAGCAGAAATGTTTAGGGCCTTC----TTGTTTGGAACAGCTGAGCACCTCCCATA 71867
human.FAM30B-LOC105372935-GGT2.NCBI.                                         atataagcttgattttgatgcaaccactcatttctttgaaaatcaggggcattcacaca 107269
human.FAM230E-LOC105377182-GGT3P.NCBI.                                       ------------------------------------------------------------ 57310
human.FAM230J-LOC105372942-GGTLC5P.NCBI.rev.compl                            ------------------------------------------------------------ 46364
human.FAM230A.ncRNA.18487127.18500594-GGTLC3.18516335.18518165.rev.compl.REF ------------------------------------------------------------ 31039

chimp.LOC112206744-LOC107973052-GGT2.revcompl                                ------------------------------------------------------------ 61762
chimp54546-135457.revcompl.Pan.troglodytes.clone.rp43-41g5.GenBank:AC099533.36 GAAAAAAAGGACACATTATTACATCCTAACTTCGCATCAAGAAGAAGTATATATGGTCAT 71927
human.FAM30B-LOC105372935-GGT2.NCBI.                                         caggtgttctagatttccttacagtaaaaccaaccctttacaaggaatagctgtgttcac 107329
human.FAM230E-LOC105377182-GGT3P.NCBI.                                       ------------------------------------------------------------ 57310
human.FAM230J-LOC105372942-GGTLC5P.NCBI.rev.compl                            ------------------------------------------------------------ 46364
human.FAM230A.ncRNA.18487127.18500594-GGTLC3.18516335.18518165.rev.compl.REF ------------------------------------------------------------ 31039

chimp.LOC112206744-LOC107973052-GGT2.revcompl                                ------------------------------------------------------------ 61762
chimp54546-135457.revcompl.Pan.troglodytes.clone.rp43-41g5.GenBank:AC099533.36 GATGGATTCTGGAAGAAGCA---------------------------------------- 71947
human.FAM30B-LOC105372935-GGT2.NCBI.                                         tttgaagcagtaaagaagcaggtgtccactcatggcaaatgaggaactccctttttttt 107389
human.FAM230E-LOC105377182-GGT3P.NCBI.                                       ------------------------------------------------------------ 57310
human.FAM230J-LOC105372942-GGTLC5P.NCBI.rev.compl                            ------------------------------------------------------------ 46364
human.FAM230A.ncRNA.18487127.18500594-GGTLC3.18516335.18518165.rev.compl.REF ------------------------------------------------------------ 31039

chimp.LOC112206744-LOC107973052-GGT2.revcompl                                ------------------------------------------------------------ 61762
chimp54546-135457.revcompl.Pan.troglodytes.clone.rp43-41g5.GenBank:AC099533.36 -----------------------------------------TTATTCACTCTTGGGATGT 71966
human.FAM30B-LOC105372935-GGT2.NCBI.                                         ttttttttttttttttttttttgagacggggtctcgctctttttttgtgtgtgtgagat 107449
human.FAM230E-LOC105377182-GGT3P.NCBI.                                       ------------------------------------------------------------ 57310
human.FAM230J-LOC105372942-GGTLC5P.NCBI.rev.compl                            ------------------------------------------------------------ 46364
human.FAM230A.ncRNA.18487127.18500594-GGTLC3.18516335.18518165.rev.compl.REF ------------------------------------------------------------ 31039

chimp.LOC112206744-LOC107973052-GGT2.revcompl                                ------------------------------------------------------------ 61762
chimp54546-135457.revcompl.Pan.troglodytes.clone.rp43-41g5.GenBank:AC099533.36 TGCTCCAACCCAAATAGCAAAAACAAAGAAAAGTTGCATGTCTGTGGGTCCTAGAGCAGG 72026
human.FAM30B-LOC105372935-GGT2.NCBI.                                         gggggccacccaggctagagtgcagtagtgcaattacagctcactgtagcctcaaccttc 107509
human.FAM230E-LOC105377182-GGT3P.NCBI.                                       ------------------------------------------------------------ 57310
human.FAM230J-LOC105372942-GGTLC5P.NCBI.rev.compl                            ------------------------------------------------------------ 46364
human.FAM230A.ncRNA.18487127.18500594-GGTLC3.18516335.18518165.rev.compl.REF ------------------------------------------------------------ 31039

chimp.LOC112206744-LOC107973052-GGT2.revcompl                                ------------------------------------------------------------ 61762
chimp54546-135457.revcompl.Pan.troglodytes.clone.rp43-41g5.GenBank:AC099533.36 AAGGCT------------------------------------------------------ 72032
human.FAM30B-LOC105372935-GGT2.NCBI.                                         tgggctcaaccagttcttcccacctcaacttctgaagcagctgagactacaggtgtgtac 107569
human.FAM230E-LOC105377182-GGT3P.NCBI.                                       ------------------------------------------------------------ 57310
human.FAM230J-LOC105372942-GGTLC5P.NCBI.rev.compl                            ------------------------------------------------------------ 46364
human.FAM230A.ncRNA.18487127.18500594-GGTLC3.18516335.18518165.rev.compl.REF ------------------------------------------------------------ 31039

chimp.LOC112206744-LOC107973052-GGT2.revcompl                                ------------------------------------------------------------ 61762
chimp54546-135457.revcompl.Pan.troglodytes.clone.rp43-41g5.GenBank:AC099533.36 ------------------------------------------------------------ 72032
human.FAM30B-LOC105372935-GGT2.NCBI.                                         caccatgcccaggtaatgcttttgatttttagtaagatgaggtctcgttatattgcccag 107629
human.FAM230E-LOC105377182-GGT3P.NCBI.                                       ------------------------------------------------------------ 57310
human.FAM230J-LOC105372942-GGTLC5P.NCBI.rev.compl                            ------------------------------------------------------------ 46364
human.FAM230A.ncRNA.18487127.18500594-GGTLC3.18516335.18518165.rev.compl.REF ------------------------------------------------------------ 31039

chimp.LOC112206744-LOC107973052-GGT2.revcompl                                ------------------------------------------------------------ 61762
chimp54546-135457.revcompl.Pan.troglodytes.clone.rp43-41g5.GenBank:AC099533.36 ----------------------------------CTGCAGCAGGTCCACACAGTGGTGGG 72058
human.FAM30B-LOC105372935-GGT2.NCBI.                                         gctggtctcaaactcctgagctcaagtgatcctcctgcctcagcctcccaaagtgctagg 107689
human.FAM230E-LOC105377182-GGT3P.NCBI.                                       ------------------------------------------------------------ 57310
human.FAM230J-LOC105372942-GGTLC5P.NCBI.rev.compl                            ------------------------------------------------------------ 46364
human.FAM230A.ncRNA.18487127.18500594-GGTLC3.18516335.18518165.rev.compl.REF ------------------------------------------------------------ 31039

chimp.LOC112206744-LOC107973052-GGT2.revcompl                                ------------------------------------------------------------ 61762
chimp54546-135457.revcompl.Pan.troglodytes.clone.rp43-41g5.GenBank:AC099533.36 AGTGCCCTGCTGGTTGAGCCCTAGAACCCAGCTGGGTCTACTGCAATGCAGGAATTTATG 72118
human.FAM30B-LOC105372935-GGT2.NCBI.                                         attatcggcatgagccactatgcccagccagaagctaaaaatattaatcaaggagtcaca 107749
human.FAM230E-LOC105377182-GGT3P.NCBI.                                       ------------------------------------------------------------ 57310
human.FAM230J-LOC105372942-GGTLC5P.NCBI.rev.compl                            ------------------------------------------------------------ 46364
human.FAM230A.ncRNA.18487127.18500594-GGTLC3.18516335.18518165.rev.compl.REF ------------------------------------------------------------ 31039
```

```
chimp.LOC112206744-LOC107973052-GGT2.revcompl                                                 ------------------------------------------------------------ 61762
chimp54546-135457.revcompl.Pan.troglodytes.clone.rp43-41g5.GenBank:AC099533.36                ATGGGAGAAGAAACCTTGCATTTATGGCAAGACCCAATAGGAG----------------- 72161
human.FAM30B-LOC105372935-GGT2.NCBI.                                                          actaaggatgcactgcacacccgaagccagggaccacaagaggtcagcaactgggactgc 107809
human.FAM230E-LOC105377182-GGT3P.NCBI.                                                        ------------------------------------------------------------ 57310
human.FAM230J-LOC105372942-GGTLC5P.NCBI.rev.compl                                             ------------------------------------------------------------ 46364
human.FAM230A.ncRNA.18487127.18500594-GGTLC3.18516335.18518165.rev.compl.REF                  ------------------------------------------------------------ 31039

chimp.LOC112206744-LOC107973052-GGT2.revcompl                                                 ------------------------------------------------------------ 61762
chimp54546-135457.revcompl.Pan.troglodytes.clone.rp43-41g5.GenBank:AC099533.36                ------------------------------------------------------------ 72161
human.FAM30B-LOC105372935-GGT2.NCBI.                                                          ttcctccagttcccaggctggggctcagcagcagtggcctgggctctggggacttacttt 107869
human.FAM230E-LOC105377182-GGT3P.NCBI.                                                        ------------------------------------------------------------ 57310
human.FAM230J-LOC105372942-GGTLC5P.NCBI.rev.compl                                             ------------------------------------------------------------ 46364
human.FAM230A.ncRNA.18487127.18500594-GGTLC3.18516335.18518165.rev.compl.REF                  ------------------------------------------------------------ 31039

chimp.LOC112206744-LOC107973052-GGT2.revcompl                                                 ------------------------------------------------------------ 61762
chimp54546-135457.revcompl.Pan.troglodytes.clone.rp43-41g5.GenBank:AC099533.36                ------------------------------------------------------------ 72161
human.FAM30B-LOC105372935-GGT2.NCBI.                                                          aagccgaaatcccaaactcattatcattcaaaccaattttctgtgtccagcatcatccca 107929
human.FAM230E-LOC105377182-GGT3P.NCBI.                                                        ------------------------------------------------------------ 57310
human.FAM230J-LOC105372942-GGTLC5P.NCBI.rev.compl                                             ------------------------------------------------------------ 46364
human.FAM230A.ncRNA.18487127.18500594-GGTLC3.18516335.18518165.rev.compl.REF                  ------------------------------------------------------------ 31039

chimp.LOC112206744-LOC107973052-GGT2.revcompl                                                 ------------------------------------------------------------ 61762
chimp54546-135457.revcompl.Pan.troglodytes.clone.rp43-41g5.GenBank:AC099533.36                ------------------------------------------------------------ 72161
human.FAM30B-LOC105372935-GGT2.NCBI.                                                          cctaaactggtccttaaaaccactgacagtcatttctccaatctgtaaaaggcctgctga 107989
human.FAM230E-LOC105377182-GGT3P.NCBI.                                                        ------------------------------------------------------------ 57310
human.FAM230J-LOC105372942-GGTLC5P.NCBI.rev.compl                                             ------------------------------------------------------------ 46364
human.FAM230A.ncRNA.18487127.18500594-GGTLC3.18516335.18518165.rev.compl.REF                  ------------------------------------------------------------ 31039

chimp.LOC112206744-LOC107973052-GGT2.revcompl                                                 ------------------------------------------------------------ 61762
chimp54546-135457.revcompl.Pan.troglodytes.clone.rp43-41g5.GenBank:AC099533.36                ----------------------------------------------ATCCCAGTGCAGAC 72175
human.FAM30B-LOC105372935-GGT2.NCBI.                                                          ggcctttgcattatcaggacttaactgggagtagaatgccctccccatccaaatcctgac 108049
human.FAM230E-LOC105377182-GGT3P.NCBI.                                                        ------------------------------------------------------------ 57310
human.FAM230J-LOC105372942-GGTLC5P.NCBI.rev.compl                                             ------------------------------------------------------------ 46364
human.FAM230A.ncRNA.18487127.18500594-GGTLC3.18516335.18518165.rev.compl.REF                  ------------------------------------------------------------ 31039

chimp.LOC112206744-LOC107973052-GGT2.revcompl                                                 ------------------------------------------------------------ 61762
chimp54546-135457.revcompl.Pan.troglodytes.clone.rp43-41g5.GenBank:AC099533.36                ATGTGGGGTTGTGTAACAAGGCCCTGTCATCTTAATGAAATTTGCCCCT----------- 72224
human.FAM30B-LOC105372935-GGT2.NCBI.                                                          catttc---tgaacactcagcccatgtcctcttcctgagtgtcttccccaacaacaacca 108106
human.FAM230E-LOC105377182-GGT3P.NCBI.                                                        ------------------------------------------------------------ 57310
human.FAM230J-LOC105372942-GGTLC5P.NCBI.rev.compl                                             ------------------------------------------------------------ 46364
human.FAM230A.ncRNA.18487127.18500594-GGTLC3.18516335.18518165.rev.compl.REF                  ------------------------------------------------------------ 31039

chimp.LOC112206744-LOC107973052-GGT2.revcompl                                                 ------------------------------------------------------------ 61762
chimp54546-135457.revcompl.Pan.troglodytes.clone.rp43-41g5.GenBank:AC099533.36                TCTGGAAAGGTTTGTTCTGTCTTTGCTGCTCACACCTGAGAGTCGTGTGTGCACTTCTTT 72284
human.FAM30B-LOC105372935-GGT2.NCBI.                                                          cctacactgaactttactttgatctctgcaggaaactgtagatagagatggggtctcact 108166
human.FAM230E-LOC105377182-GGT3P.NCBI.                                                        ------------------------------------------------------------ 57310
human.FAM230J-LOC105372942-GGTLC5P.NCBI.rev.compl                                             ------------------------------------------------------------ 46364
human.FAM230A.ncRNA.18487127.18500594-GGTLC3.18516335.18518165.rev.compl.REF                  ------------------------------------------------------------ 31039

chimp.LOC112206744-LOC107973052-GGT2.revcompl                                                 ------------------------------------------------------------ 61762
chimp54546-135457.revcompl.Pan.troglodytes.clone.rp43-41g5.GenBank:AC099533.36                CTCATGTGCACGTGCAAGACAACTCCTGTTCTCAGACACCCAGAGAAGAAAAGGCTTGAA 72344
human.FAM30B-LOC105372935-GGT2.NCBI.                                                          atgttgcctagctggtcttgaactgctgagctcaa------------------------- 108201
human.FAM230E-LOC105377182-GGT3P.NCBI.                                                        ------------------------------------------------------------ 57310
human.FAM230J-LOC105372942-GGTLC5P.NCBI.rev.compl                                             ------------------------------------------------------------ 46364
human.FAM230A.ncRNA.18487127.18500594-GGTLC3.18516335.18518165.rev.compl.REF                  ------------------------------------------------------------ 31039

chimp.LOC112206744-LOC107973052-GGT2.revcompl                                                 ------------------------------------------------------------ 61762
chimp54546-135457.revcompl.Pan.troglodytes.clone.rp43-41g5.GenBank:AC099533.36                ACTGAGGAGCTGATCCCTAGGATTCCATTGCACCTGCCTGGCCCAGACAGAGCTCCTGAA 72404
human.FAM30B-LOC105372935-GGT2.NCBI.                                                          ------------------gtgatctccctgcttctgcctcccaaagtgctaggattacag 108243
human.FAM230E-LOC105377182-GGT3P.NCBI.                                                        ------------------------------------------------------------ 57310
human.FAM230J-LOC105372942-GGTLC5P.NCBI.rev.compl                                             ------------------------------------------------------------ 46364
human.FAM230A.ncRNA.18487127.18500594-GGTLC3.18516335.18518165.rev.compl.REF                  ------------------------------------------------------------ 31039

chimp.LOC112206744-LOC107973052-GGT2.revcompl                                                 ------------------------------------------------------------ 61762
chimp54546-135457.revcompl.Pan.troglodytes.clone.rp43-41g5.GenBank:AC099533.36                ATGAAAGGAGCTGGGCTGGGCCAGAGTAGCACATCTCATTCTGCGGGCACAATCCTACTG 72464
human.FAM30B-LOC105372935-GGT2.NCBI.                                                          cagtgagccaccacacctggcctatgtatg-cttttttttttctttcttttttttttttt 108302
human.FAM230E-LOC105377182-GGT3P.NCBI.                                                        ------------------------------------------------------------ 57310
human.FAM230J-LOC105372942-GGTLC5P.NCBI.rev.compl                                             ------------------------------------------------------------ 46364
human.FAM230A.ncRNA.18487127.18500594-GGTLC3.18516335.18518165.rev.compl.REF                  ------------------------------------------------------------ 31039

chimp.LOC112206744-LOC107973052-GGT2.revcompl                                                 ------------------------------------------------------------ 61762
chimp54546-135457.revcompl.Pan.troglodytes.clone.rp43-41g5.GenBank:AC099533.36                GTTCAAAATGCTTTCTTTTTCTCAAGCCCAA-----------------------GTTCTC 72501
human.FAM30B-LOC105372935-GGT2.NCBI.                                                          ttttgagacagagtcttgctctgtggcccagtccgcagtgcagtggtgcaatctcggctc 108362
human.FAM230E-LOC105377182-GGT3P.NCBI.                                                        ------------------------------------------------------------ 57310
human.FAM230J-LOC105372942-GGTLC5P.NCBI.rev.compl                                             ------------------------------------------------------------ 46364
human.FAM230A.ncRNA.18487127.18500594-GGTLC3.18516335.18518165.rev.compl.REF                  ------------------------------------------------------------ 31039

chimp.LOC112206744-LOC107973052-GGT2.revcompl                                                 ------------------------------------------------------------ 61762
chimp54546-135457.revcompl.Pan.troglodytes.clone.rp43-41g5.GenBank:AC099533.36                AGGGTTAGCTCAGAGACCTGGTTCCTTTCCTGCCACACTCTGTGTTGAAAGGGAGGGAAA 72561
human.FAM30B-LOC105372935-GGT2.NCBI.                                                          atggcaacctccaggttcaaggcattttcctgcctcagccttccaaggagctgggattac 108422
human.FAM230E-LOC105377182-GGT3P.NCBI.                                                        ------------------------------------------------------------ 57310
human.FAM230J-LOC105372942-GGTLC5P.NCBI.rev.compl                                             ------------------------------------------------------------ 46364
human.FAM230A.ncRNA.18487127.18500594-GGTLC3.18516335.18518165.rev.compl.REF                  ------------------------------------------------------------ 31039

chimp.LOC112206744-LOC107973052-GGT2.revcompl                                                 ------------------------------------------------------------ 61762
chimp54546-135457.revcompl.Pan.troglodytes.clone.rp43-41g5.GenBank:AC099533.36                CGACTGTCTCTAGTGCAGAAA---------CCAAGCTCTCTGGAGACATTTCTTATCTCA 72612
human.FAM30B-LOC105372935-GGT2.NCBI.                                                          aggcacgcaccatcacgcccagctaattggccaggctggtctcaaactcctgacctcagg 108482
human.FAM230E-LOC105377182-GGT3P.NCBI.                                                        ------------------------------------------------------------ 57310
human.FAM230J-LOC105372942-GGTLC5P.NCBI.rev.compl                                             ------------------------------------------------------------ 46364
human.FAM230A.ncRNA.18487127.18500594-GGTLC3.18516335.18518165.rev.compl.REF                  ------------------------------------------------------------ 31039

chimp.LOC112206744-LOC107973052-GGT2.revcompl                                                 ------------------------------------------------------------ 61762
chimp54546-135457.revcompl.Pan.troglodytes.clone.rp43-41g5.GenBank:AC099533.36                TGAACCTGTGCACTGGTCTCTGTAGGGCCTAGGTAATAAAGGCACAAGATGTTTGGATTT 72672
human.FAM30B-LOC105372935-GGT2.NCBI.                                                          tgatccacccaccttggcctcccaaagtgctgggattacaggtgtaagccaccacaccgg 108542
human.FAM230E-LOC105377182-GGT3P.NCBI.                                                        ------------------------------------------------------------ 57310
human.FAM230J-LOC105372942-GGTLC5P.NCBI.rev.compl                                             ------------------------------------------------------------ 46364
human.FAM230A.ncRNA.18487127.18500594-GGTLC3.18516335.18518165.rev.compl.REF                  ------------------------------------------------------------ 31039

chimp.LOC112206744-LOC107973052-GGT2.revcompl                                                 ------------------------------------------------------------ 61762
chimp54546-135457.revcompl.Pan.troglodytes.clone.rp43-41g5.GenBank:AC099533.36                ACAATGAATGTAATTGTAAAATTCATGGTTAAATAGAA---------------------- 72710
human.FAM30B-LOC105372935-GGT2.NCBI.                                                          gccacgtatgcattttaaaacgttatgctcagacggggtctataggtttcaccagactgc 108602
human.FAM230E-LOC105377182-GGT3P.NCBI.                                                        ------------------------------------------------------------ 57310
human.FAM230J-LOC105372942-GGTLC5P.NCBI.rev.compl                                             ------------------------------------------------------------ 46364
human.FAM230A.ncRNA.18487127.18500594-GGTLC3.18516335.18518165.rev.compl.REF                  ------------------------------------------------------------ 31039

chimp.LOC112206744-LOC107973052-GGT2.revcompl                                                 ------------------------------------------------------------ 61762
chimp54546-135457.revcompl.Pan.troglodytes.clone.rp43-41g5.GenBank:AC099533.36                ------------------------------------------------------------ 72710
human.FAM30B-LOC105372935-GGT2.NCBI.                                                          caaaaggatccatgacacaaaaaaagtactctcagttgcatccttcggaacctttcaaca 108662
human.FAM230E-LOC105377182-GGT3P.NCBI.                                                        ------------------------------------------------------------ 57310
human.FAM230J-LOC105372942-GGTLC5P.NCBI.rev.compl                                             ------------------------------------------------------------ 46364
human.FAM230A.ncRNA.18487127.18500594-GGTLC3.18516335.18518165.rev.compl.REF                  ------------------------------------------------------------ 31039

chimp.LOC112206744-LOC107973052-GGT2.revcompl                                                 ------------------------------------------------------------ 61762
chimp54546-135457.revcompl.Pan.troglodytes.clone.rp43-41g5.GenBank:AC099533.36                ------------------------------------------------------------ 72710
human.FAM30B-LOC105372935-GGT2.NCBI.                                                          catacacatatacctcactcatccttttgtgaagagtgtctattaatcactagattaaga 108722
human.FAM230E-LOC105377182-GGT3P.NCBI.                                                        ------------------------------------------------------------ 57310
human.FAM230J-LOC105372942-GGTLC5P.NCBI.rev.compl                                             ------------------------------------------------------------ 46364
human.FAM230A.ncRNA.18487127.18500594-GGTLC3.18516335.18518165.rev.compl.REF                  ------------------------------------------------------------ 31039

chimp.LOC112206744-LOC107973052-GGT2.revcompl                                                 ------------------------------------------------------------ 61762
chimp54546-135457.revcompl.Pan.troglodytes.clone.rp43-41g5.GenBank:AC099533.36                ------------------------------------------------------------ 72710
human.FAM30B-LOC105372935-GGT2.NCBI.                                                          ggatttggataaaatttataaatgtacaaactttgctaccaaggactgacacatttactt 108782
human.FAM230E-LOC105377182-GGT3P.NCBI.                                                        ------------------------------------------------------------ 57310
human.FAM230J-LOC105372942-GGTLC5P.NCBI.rev.compl                                             ------------------------------------------------------------ 46364
human.FAM230A.ncRNA.18487127.18500594-GGTLC3.18516335.18518165.rev.compl.REF                  ------------------------------------------------------------ 31039
```

```
chimp.LOC112206744-LOC107973052-GGT2.revcompl                                   ------------------------------------------------------------ 61762
chimp54546-135457.revcompl.Pan.troglodytes.clone.rp43-41g5.GenBank:AC099533.36  ------------------------------------------------------------ 72710
human.FAM30B-LOC105372935-GGT2.NCBI.                                            ttctagctcatcagtttatgtcctagaaaaattccagagccacattaaccagctgtttcc 108842
human.FAM230E-LOC105377182-GGT3P.NCBI.                                          ------------------------------------------------------------ 57310
human.FAM230J-LOC105372942-GGTLC5P.NCBI.rev.compl                               ------------------------------------------------------------ 46364
human.FAM230A.ncRNA.18487127.18500594-GGTLC3.18516335.18518165.rev.compl.REF    ------------------------------------------------------------ 31039

chimp.LOC112206744-LOC107973052-GGT2.revcompl                                   ------------------------------------------------------------ 61762
chimp54546-135457.revcompl.Pan.troglodytes.clone.rp43-41g5.GenBank:AC099533.36  --------ATCCGACAATTTCTTCTGCTGCTTTGGCTACCATGAGAAACCACCTGAGTGA 72762
human.FAM30B-LOC105372935-GGT2.NCBI.                                            tatttttcattttacgatggtgtttccactttcacctgcaaagtgactaatagtaatacc 108902
human.FAM230E-LOC105377182-GGT3P.NCBI.                                          ------------------------------------------------------------ 57310
human.FAM230J-LOC105372942-GGTLC5P.NCBI.rev.compl                               ------------------------------------------------------------ 46364
human.FAM230A.ncRNA.18487127.18500594-GGTLC3.18516335.18518165.rev.compl.REF    ------------------------------------------------------------ 31039

chimp.LOC112206744-LOC107973052-GGT2.revcompl                                   ------------------------------------------------------------ 61762
chimp54546-135457.revcompl.Pan.troglodytes.clone.rp43-41g5.GenBank:AC099533.36  ACTGAGACCAAAGTCATGCTCACTGCTTAGAGCAAGCTTGTCCAGCCTGCAGCCTGCAGG 72822
human.FAM30B-LOC105372935-GGT2.NCBI.                                            tattacagtaatttaagtaacaccatttagagtaatttcatctttagtactgtatttata 108962
human.FAM230E-LOC105377182-GGT3P.NCBI.                                          ------------------------------------------------------------ 57310
human.FAM230J-LOC105372942-GGTLC5P.NCBI.rev.compl                               ------------------------------------------------------------ 46364
human.FAM230A.ncRNA.18487127.18500594-GGTLC3.18516335.18518165.rev.compl.REF    ------------------------------------------------------------ 31039

chimp.LOC112206744-LOC107973052-GGT2.revcompl                                   ------------------------------------------------------------ 61762
chimp54546-135457.revcompl.Pan.troglodytes.clone.rp43-41g5.GenBank:AC099533.36  CCAC-------------------------------------------------------- 72826
human.FAM30B-LOC105372935-GGT2.NCBI.                                            ctactaagaaagccgccaacgattcaagctgcactcactggtgaaaccctatgctatata 109022
human.FAM230E-LOC105377182-GGT3P.NCBI.                                          ------------------------------------------------------------ 57310
human.FAM230J-LOC105372942-GGTLC5P.NCBI.rev.compl                               ------------------------------------------------------------ 46364
human.FAM230A.ncRNA.18487127.18500594-GGTLC3.18516335.18518165.rev.compl.REF    ------------------------------------------------------------ 31039

chimp.LOC112206744-LOC107973052-GGT2.revcompl                                   ------------------------------------------------------------ 61762
chimp54546-135457.revcompl.Pan.troglodytes.clone.rp43-41g5.GenBank:AC099533.36  ------------------------------------------------------------ 72826
human.FAM30B-LOC105372935-GGT2.NCBI.                                            tgtaattaatttatttcctaaattcttacctccctccacccccctccattggaaaataaac 109082
human.FAM230E-LOC105377182-GGT3P.NCBI.                                          ------------------------------------------------------------ 57310
human.FAM230J-LOC105372942-GGTLC5P.NCBI.rev.compl                               ------------------------------------------------------------ 46364
human.FAM230A.ncRNA.18487127.18500594-GGTLC3.18516335.18518165.rev.compl.REF    ------------------------------------------------------------ 31039

chimp.LOC112206744-LOC107973052-GGT2.revcompl                                   ------------------------------------------------------------ 61762
chimp54546-135457.revcompl.Pan.troglodytes.clone.rp43-41g5.GenBank:AC099533.36  ------------------------------------------------------------ 72826
human.FAM30B-LOC105372935-GGT2.NCBI.                                            tctaagaatgcaaaagtttttgtgtgtttttcttcaacgcttagagcagtgcctggcacc 109142
human.FAM230E-LOC105377182-GGT3P.NCBI.                                          ------------------------------------------------------------ 57310
human.FAM230J-LOC105372942-GGTLC5P.NCBI.rev.compl                               ------------------------------------------------------------ 46364
human.FAM230A.ncRNA.18487127.18500594-GGTLC3.18516335.18518165.rev.compl.REF    ------------------------------------------------------------ 31039

chimp.LOC112206744-LOC107973052-GGT2.revcompl                                   ------------------------------------------------------------ 61762
chimp54546-135457.revcompl.Pan.troglodytes.clone.rp43-41g5.GenBank:AC099533.36  ------------------------------------------------------------ 72826
human.FAM30B-LOC105372935-GGT2.NCBI.                                            taacagtcaatatctgctaactttgatgaatgaatggtatatttatattatgcaatatta 109202
human.FAM230E-LOC105377182-GGT3P.NCBI.                                          ------------------------------------------------------------ 57310
human.FAM230J-LOC105372942-GGTLC5P.NCBI.rev.compl                               ------------------------------------------------------------ 46364
human.FAM230A.ncRNA.18487127.18500594-GGTLC3.18516335.18518165.rev.compl.REF    ------------------------------------------------------------ 31039

chimp.LOC112206744-LOC107973052-GGT2.revcompl                                   ------------------------------------------------------------ 61762
chimp54546-135457.revcompl.Pan.troglodytes.clone.rp43-41g5.GenBank:AC099533.36  --ATGCAGCCTAGGATGGCTTTGAAT----GTGGCCCCACATAAATTTGTAAACTTTCTT 72880
human.FAM30B-LOC105372935-GGT2.NCBI.                                            ctatgcagccatgaaacacaatgagttagagtggtaccagctaatttaatggaatttcca 109262
human.FAM230E-LOC105377182-GGT3P.NCBI.                                          ------------------------------------------------------------ 57310
human.FAM230J-LOC105372942-GGTLC5P.NCBI.rev.compl                               ------------------------------------------------------------ 46364
human.FAM230A.ncRNA.18487127.18500594-GGTLC3.18516335.18518165.rev.compl.REF    ------------------------------------------------------------ 31039

chimp.LOC112206744-LOC107973052-GGT2.revcompl                                   ------------------------------------------------------------ 61762
chimp54546-135457.revcompl.Pan.troglodytes.clone.rp43-41g5.GenBank:AC099533.36  AAAACGTTATGAGAA--------------------------------------------- 72895
human.FAM30B-LOC105372935-GGT2.NCBI.                                            gaagtatgaaattaagaaaacaagaagaaaagtgtcaaaaatatcatcccactttttttt 109322
human.FAM230E-LOC105377182-GGT3P.NCBI.                                          ------------------------------------------------------------ 57310
human.FAM230J-LOC105372942-GGTLC5P.NCBI.rev.compl                               ------------------------------------------------------------ 46364
human.FAM230A.ncRNA.18487127.18500594-GGTLC3.18516335.18518165.rev.compl.REF    ------------------------------------------------------------ 31039

chimp.LOC112206744-LOC107973052-GGT2.revcompl                                   ------------------------------------------------------------ 61762
chimp54546-135457.revcompl.Pan.troglodytes.clone.rp43-41g5.GenBank:AC099533.36  ------------------------------------------------------------ 72895
human.FAM30B-LOC105372935-GGT2.NCBI.                                            tttcttccctgaaacggagtcgtgctctgacacccaggctacagtatagtgctgccatca 109382
human.FAM230E-LOC105377182-GGT3P.NCBI.                                          ------------------------------------------------------------ 57310
human.FAM230J-LOC105372942-GGTLC5P.NCBI.rev.compl                               ------------------------------------------------------------ 46364
human.FAM230A.ncRNA.18487127.18500594-GGTLC3.18516335.18518165.rev.compl.REF    ------------------------------------------------------------ 31039

chimp.LOC112206744-LOC107973052-GGT2.revcompl                                   ------------------------------------------------------------ 61762
chimp54546-135457.revcompl.Pan.troglodytes.clone.rp43-41g5.GenBank:AC099533.36  ---------------------------ATTTTTGCAATTTTTTAAGCTCATCAGCTATTG 72928
human.FAM30B-LOC105372935-GGT2.NCBI.                                            tggctcactgcaacctcaaccttctgggctcagggaatccttccacctcagcctcccatg 109442
human.FAM230E-LOC105377182-GGT3P.NCBI.                                          ------------------------------------------------------------ 57310
human.FAM230J-LOC105372942-GGTLC5P.NCBI.rev.compl                               ------------------------------------------------------------ 46364
human.FAM230A.ncRNA.18487127.18500594-GGTLC3.18516335.18518165.rev.compl.REF    ------------------------------------------------------------ 31039

chimp.LOC112206744-LOC107973052-GGT2.revcompl                                   ------------------------------------------------------------ 61762
chimp54546-135457.revcompl.Pan.troglodytes.clone.rp43-41g5.GenBank:AC099533.36  TTAGTGTTAATATATTTATGTGTGGCCCAAGACAATTTTTCTTCTTCAAA--------TG 72980
human.FAM30B-LOC105372935-GGT2.NCBI.                                            tagctgggactacaggtgtgtgccacaacgcccagctaatttattttaaattatttttg 109502
human.FAM230E-LOC105377182-GGT3P.NCBI.                                          ------------------------------------------------------------ 57310
human.FAM230J-LOC105372942-GGTLC5P.NCBI.rev.compl                               ------------------------------------------------------------ 46364
human.FAM230A.ncRNA.18487127.18500594-GGTLC3.18516335.18518165.rev.compl.REF    ------------------------------------------------------------ 31039

chimp.LOC112206744-LOC107973052-GGT2.revcompl                                   ------------------------------------------------------------ 61762
chimp54546-135457.revcompl.Pan.troglodytes.clone.rp43-41g5.GenBank:AC099533.36  TCGCCCAGGGAAGCCAAAATGTTGGACACCCTGGCTTAGAGGGGATGTGAGATATTGAGC 73040
human.FAM30B-LOC105372935-GGT2.NCBI.                                            tagagacagggtctccctatgttgcccaggctggtctcaaactactaggctcaagtgatc 109562
human.FAM230E-LOC105377182-GGT3P.NCBI.                                          ------------------------------------------------------------ 57310
human.FAM230J-LOC105372942-GGTLC5P.NCBI.rev.compl                               ------------------------------------------------------------ 46364
human.FAM230A.ncRNA.18487127.18500594-GGTLC3.18516335.18518165.rev.compl.REF    ------------------------------------------------------------ 31039

chimp.LOC112206744-LOC107973052-GGT2.revcompl                                   ------------------------------------------------------------ 61762
chimp54546-135457.revcompl.Pan.troglodytes.clone.rp43-41g5.GenBank:AC099533.36  CTAGTTATATTAGAGGTTCCTGTCCCACATCCCCATCTGTCTGTTTCAC---TGTGAGCC 73097
human.FAM30B-LOC105372935-GGT2.NCBI.                                            ctc---------------ccacctcagcttcccaaagtgttgggattacaggtgtgagcc 109607
human.FAM230E-LOC105377182-GGT3P.NCBI.                                          ------------------------------------------------------------ 57310
human.FAM230J-LOC105372942-GGTLC5P.NCBI.rev.compl                               ------------------------------------------------------------ 46364
human.FAM230A.ncRNA.18487127.18500594-GGTLC3.18516335.18518165.rev.compl.REF    ------------------------------------------------------------ 31039

chimp.LOC112206744-LOC107973052-GGT2.revcompl                                   ------------------------------------------------------------ 61762
chimp54546-135457.revcompl.Pan.troglodytes.clone.rp43-41g5.GenBank:AC099533.36  ACTGCCTGCTGCTCCCTCTGCCATCATAGGTGCTACAG--------TAATAGCCTTGTCC 73149
human.FAM30B-LOC105372935-GGT2.NCBI.                                            actgtgcccagcccaatcccacttttaaaaagtatacccccaaatccatacactggtga 109667
human.FAM230E-LOC105377182-GGT3P.NCBI.                                          ------------------------------------------------------------ 57310
human.FAM230J-LOC105372942-GGTLC5P.NCBI.rev.compl                               ------------------------------------------------------------ 46364
human.FAM230A.ncRNA.18487127.18500594-GGTLC3.18516335.18518165.rev.compl.REF    ------------------------------------------------------------ 31039

chimp.LOC112206744-LOC107973052-GGT2.revcompl                                   ------------------------------------------------------------ 61762
chimp54546-135457.revcompl.Pan.troglodytes.clone.rp43-41g5.GenBank:AC099533.36  CTGGCCTGGGCCCCACTGATGGTCAGAGTGGCCGTGTTCCACAAGTTGGAGCCAGAGAAT 73209
human.FAM30B-LOC105372935-GGT2.NCBI.                                            ggaagtggagaacgtggaatcctcacacacagaatgtaaaatggtacagctctggagaac 109727
human.FAM230E-LOC105377182-GGT3P.NCBI.                                          ------------------------------------------------------------ 57310
human.FAM230J-LOC105372942-GGTLC5P.NCBI.rev.compl                               ------------------------------------------------------------ 46364
human.FAM230A.ncRNA.18487127.18500594-GGTLC3.18516335.18518165.rev.compl.REF    ------------------------------------------------------------ 31039

chimp.LOC112206744-LOC107973052-GGT2.revcompl                                   ------------------------------------------------------------ 61762
chimp54546-135457.revcompl.Pan.troglodytes.clone.rp43-41g5.GenBank:AC099533.36  CACTCAGGGGTCCTTGAGGGCCACTTGCTGTCTTTATAGATGACCGGCACAGGGGTCTGG 73269
human.FAM30B-LOC105372935-GGT2.NCBI.                                            attttagcagtttctcaaaatgtttaacgtagctactacacgacccagcaattcaactcc 109787
human.FAM230E-LOC105377182-GGT3P.NCBI.                                          ------------------------------------------------------------ 57310
human.FAM230J-LOC105372942-GGTLC5P.NCBI.rev.compl                               ------------------------------------------------------------ 46364
human.FAM230A.ncRNA.18487127.18500594-GGTLC3.18516335.18518165.rev.compl.REF    ------------------------------------------------------------ 31039

chimp.LOC112206744-LOC107973052-GGT2.revcompl                                   ------------------------------------------------------------ 61762
chimp54546-135457.revcompl.Pan.troglodytes.clone.rp43-41g5.GenBank:AC099533.36  CCTGACTTCCGCTGGTACCAATAAACATATCCACCTCCAACAGTATTTCCACAGCAGGTG 73329
human.FAM30B-LOC105372935-GGT2.NCBI.                                            cagatatacacctgagagaactgaaaggatatgtccacacaaaaatttgcagaggaactt 109847
human.FAM230E-LOC105377182-GGT3P.NCBI.                                          ------------------------------------------------------------ 57310
human.FAM230J-LOC105372942-GGTLC5P.NCBI.rev.compl                               ------------------------------------------------------------ 46364
human.FAM230A.ncRNA.18487127.18500594-GGTLC3.18516335.18518165.rev.compl.REF    ------------------------------------------------------------ 31039

chimp.LOC112206744-LOC107973052-GGT2.revcompl                                   ------------------------------------------------------------ 61762
```

```
chimp54546-135457.revcompl.Pan.troglodytes.clone.rp43-41g5.GenBank:AC099533.36      ATCCTGGC---------------------------------------------------- 73337
human.FAM30B-LOC105372935-GGT2.NCBI.                                                  gtacacacttgttcacagtagcattattcataatagcccaaagtggcaaaaacgcaaatg 109907
human.FAM230E-LOC105377182-GGT3P.NCBI.                                                ------------------------------------------------------------ 57310
human.FAM230J-LOC105372942-GGTLC5P.NCBI.rev.compl                                     ------------------------------------------------------------ 46364
human.FAM230A.ncRNA.18487127.18500594-GGTLC3.18516335.18518165.rev.compl.REF          ------------------------------------------------------------ 31039

chimp.LOC112206744-LOC107973052-GGT2.revcompl                                         ------------------------------------------------------------ 61762
chimp54546-135457.revcompl.Pan.troglodytes.clone.rp43-41g5.GenBank:AC099533.36      ------------------------------------------------------------ 73337
human.FAM30B-LOC105372935-GGT2.NCBI.                                                  ttcatcaactgatggacaaataaagcatgaacaactgatggacaaataaagtatatcatt 109967
human.FAM230E-LOC105377182-GGT3P.NCBI.                                                ------------------------------------------------------------ 57310
human.FAM230J-LOC105372942-GGTLC5P.NCBI.rev.compl                                     ------------------------------------------------------------ 46364
human.FAM230A.ncRNA.18487127.18500594-GGTLC3.18516335.18518165.rev.compl.REF          ------------------------------------------------------------ 31039

chimp.LOC112206744-LOC107973052-GGT2.revcompl                                         ------------------------------------------------------------ 61762
chimp54546-135457.revcompl.Pan.troglodytes.clone.rp43-41g5.GenBank:AC099533.36      ------------------------------------------------------------ 73337
human.FAM30B-LOC105372935-GGT2.NCBI.                                                  cagcatataagaaatgaagtgctaatacatgctacaacaggaatagaccttaataacatt 110027
human.FAM230E-LOC105377182-GGT3P.NCBI.                                                ------------------------------------------------------------ 57310
human.FAM230J-LOC105372942-GGTLC5P.NCBI.rev.compl                                     ------------------------------------------------------------ 46364
human.FAM230A.ncRNA.18487127.18500594-GGTLC3.18516335.18518165.rev.compl.REF          ------------------------------------------------------------ 31039

chimp.LOC112206744-LOC107973052-GGT2.revcompl                                         ------------------------------------------------------------ 61762
chimp54546-135457.revcompl.Pan.troglodytes.clone.rp43-41g5.GenBank:AC099533.36      ------------------------------------------------------------ 73337
human.FAM30B-LOC105372935-GGT2.NCBI.                                                  ttgctaaatgaaaaaagccaaacaaaaggtcacatgttgtatgattctatttacataaaa 110087
human.FAM230E-LOC105377182-GGT3P.NCBI.                                                ------------------------------------------------------------ 57310
human.FAM230J-LOC105372942-GGTLC5P.NCBI.rev.compl                                     ------------------------------------------------------------ 46364
human.FAM230A.ncRNA.18487127.18500594-GGTLC3.18516335.18518165.rev.compl.REF          ------------------------------------------------------------ 31039

chimp.LOC112206744-LOC107973052-GGT2.revcompl                                         ------------------------------------------------------------ 61762
chimp54546-135457.revcompl.Pan.troglodytes.clone.rp43-41g5.GenBank:AC099533.36      ------------------------------------------------------------ 73337
human.FAM30B-LOC105372935-GGT2.NCBI.                                                  atacctagactaggcaaatccatagagacacaaagatcagtgactgcccagagccagagg 110147
human.FAM230E-LOC105377182-GGT3P.NCBI.                                                ------------------------------------------------------------ 57310
human.FAM230J-LOC105372942-GGTLC5P.NCBI.rev.compl                                     ------------------------------------------------------------ 46364
human.FAM230A.ncRNA.18487127.18500594-GGTLC3.18516335.18518165.rev.compl.REF          ------------------------------------------------------------ 31039

chimp.LOC112206744-LOC107973052-GGT2.revcompl                                         ------------------------------------------------------------ 61762
chimp54546-135457.revcompl.Pan.troglodytes.clone.rp43-41g5.GenBank:AC099533.36      ------------------------------------------------------------ 73337
human.FAM30B-LOC105372935-GGT2.NCBI.                                                  gagtggtgaatgtgaaatggctgctaatgagtacagacttctttttgaggtattgaacat 110207
human.FAM230E-LOC105377182-GGT3P.NCBI.                                                ------------------------------------------------------------ 57310
human.FAM230J-LOC105372942-GGTLC5P.NCBI.rev.compl                                     ------------------------------------------------------------ 46364
human.FAM230A.ncRNA.18487127.18500594-GGTLC3.18516335.18518165.rev.compl.REF          ------------------------------------------------------------ 31039

chimp.LOC112206744-LOC107973052-GGT2.revcompl                                         ------------------------------------------------------------ 61762
chimp54546-135457.revcompl.Pan.troglodytes.clone.rp43-41g5.GenBank:AC099533.36      ------------------------------------------------------------ 73337
human.FAM30B-LOC105372935-GGT2.NCBI.                                                  gttatggaatcagatggtggtgatcaatgcataactttgtgaatataccaaaaaccacat 110267
human.FAM230E-LOC105377182-GGT3P.NCBI.                                                ------------------------------------------------------------ 57310
human.FAM230J-LOC105372942-GGTLC5P.NCBI.rev.compl                                     ------------------------------------------------------------ 46364
human.FAM230A.ncRNA.18487127.18500594-GGTLC3.18516335.18518165.rev.compl.REF          ------------------------------------------------------------ 31039

chimp.LOC112206744-LOC107973052-GGT2.revcompl                                         ------------------------------------------------------------ 61762
chimp54546-135457.revcompl.Pan.troglodytes.clone.rp43-41g5.GenBank:AC099533.36      ------------------------------------------------------------ 73337
human.FAM30B-LOC105372935-GGT2.NCBI.                                                  gaaaggtacactttaaaagcctgaattttaaggtatgtgaattgtatctccataaacctg 110327
human.FAM230E-LOC105377182-GGT3P.NCBI.                                                ------------------------------------------------------------ 57310
human.FAM230J-LOC105372942-GGTLC5P.NCBI.rev.compl                                     ------------------------------------------------------------ 46364
human.FAM230A.ncRNA.18487127.18500594-GGTLC3.18516335.18518165.rev.compl.REF          ------------------------------------------------------------ 31039

chimp.LOC112206744-LOC107973052-GGT2.revcompl                                         ------------------------------------------------------------ 61762
chimp54546-135457.revcompl.Pan.troglodytes.clone.rp43-41g5.GenBank:AC099533.36      ------------------------------------------------------------ 73337
human.FAM30B-LOC105372935-GGT2.NCBI.                                                  tgattttgttaatatacgcttatttgcaaatgaatacatggataatgaggaataatatga 110387
human.FAM230E-LOC105377182-GGT3P.NCBI.                                                ------------------------------------------------------------ 57310
human.FAM230J-LOC105372942-GGTLC5P.NCBI.rev.compl                                     ------------------------------------------------------------ 46364
human.FAM230A.ncRNA.18487127.18500594-GGTLC3.18516335.18518165.rev.compl.REF          ------------------------------------------------------------ 31039

chimp.LOC112206744-LOC107973052-GGT2.revcompl                                         ------------------------------------------------------------ 61762
chimp54546-135457.revcompl.Pan.troglodytes.clone.rp43-41g5.GenBank:AC099533.36      ------------------------------------------------------------ 73337
human.FAM30B-LOC105372935-GGT2.NCBI.                                                  aaggttaatgacaaccttggtgatagggtacaggagactgggagaggaagataagtgttg 110447
human.FAM230E-LOC105377182-GGT3P.NCBI.                                                ------------------------------------------------------------ 57310
human.FAM230J-LOC105372942-GGTLC5P.NCBI.rev.compl                                     ------------------------------------------------------------ 46364
human.FAM230A.ncRNA.18487127.18500594-GGTLC3.18516335.18518165.rev.compl.REF          ------------------------------------------------------------ 31039

chimp.LOC112206744-LOC107973052-GGT2.revcompl                                         ------------------------------------------------------------ 61762
chimp54546-135457.revcompl.Pan.troglodytes.clone.rp43-41g5.GenBank:AC099533.36      ------------------------------------------------------------ 73337
human.FAM30B-LOC105372935-GGT2.NCBI.                                                  aggtagaagaagacaaaaaataaaaggctaaaacaataatccagggccagaggccatgag 110507
human.FAM230E-LOC105377182-GGT3P.NCBI.                                                ------------------------------------------------------------ 57310
human.FAM230J-LOC105372942-GGTLC5P.NCBI.rev.compl                                     ------------------------------------------------------------ 46364
human.FAM230A.ncRNA.18487127.18500594-GGTLC3.18516335.18518165.rev.compl.REF          ------------------------------------------------------------ 31039

chimp.LOC112206744-LOC107973052-GGT2.revcompl                                         ------------------------------------------------------------ 61762
chimp54546-135457.revcompl.Pan.troglodytes.clone.rp43-41g5.GenBank:AC099533.36      ------------------------------------------------------------ 73337
human.FAM30B-LOC105372935-GGT2.NCBI.                                                  ttgggagatggagcaagacgaagtgtacaaagaaatgagcataaatcataaaaatgaggg 110567
human.FAM230E-LOC105377182-GGT3P.NCBI.                                                ------------------------------------------------------------ 57310
human.FAM230J-LOC105372942-GGTLC5P.NCBI.rev.compl                                     ------------------------------------------------------------ 46364
human.FAM230A.ncRNA.18487127.18500594-GGTLC3.18516335.18518165.rev.compl.REF          ------------------------------------------------------------ 31039

chimp.LOC112206744-LOC107973052-GGT2.revcompl                                         ------------------------------------------------------------ 61762
chimp54546-135457.revcompl.Pan.troglodytes.clone.rp43-41g5.GenBank:AC099533.36      --------------------------CATCTGTAATGGGGCCACTGACACTGAGGGTGGC 73371
human.FAM30B-LOC105372935-GGT2.NCBI.                                                  aactagggttgaaaactccagcaagtagtttgtaatggggagatgaagcaaacctaagt 110627
human.FAM230E-LOC105377182-GGT3P.NCBI.                                                ------------------------------------------------------------ 57310
human.FAM230J-LOC105372942-GGTLC5P.NCBI.rev.compl                                     ------------------------------------------------------------ 46364
human.FAM230A.ncRNA.18487127.18500594-GGTLC3.18516335.18518165.rev.compl.REF          ------------------------------------------------------------ 31039

chimp.LOC112206744-LOC107973052-GGT2.revcompl                                         ------------------------------------------------------------ 61762
chimp54546-135457.revcompl.Pan.troglodytes.clone.rp43-41g5.GenBank:AC099533.36      TGAGACAGCTCATATGTGGCCATGAAACCTGCAGGAGATGAGGGAGAAAAGGCTTGAAGC 73431
human.FAM30B-LOC105372935-GGT2.NCBI.                                                  aactgcaatgattcaacggaatttgcagaaaatgcagaagggcgagaaaagcaatgaaaa 110687
human.FAM230E-LOC105377182-GGT3P.NCBI.                                                ------------------------------------------------------------ 57310
human.FAM230J-LOC105372942-GGTLC5P.NCBI.rev.compl                                     ------------------------------------------------------------ 46364
human.FAM230A.ncRNA.18487127.18500594-GGTLC3.18516335.18518165.rev.compl.REF          ------------------------------------------------------------ 31039

chimp.LOC112206744-LOC107973052-GGT2.revcompl                                         ------------------------------------------------------------ 61762
chimp54546-135457.revcompl.Pan.troglodytes.clone.rp43-41g5.GenBank:AC099533.36      TGAGGAGCTGA------------------------------------------------- 73442
human.FAM30B-LOC105372935-GGT2.NCBI.                                                  aatggatcagaagatttaaaattggctctaccccaccaaagttttaaaaagcaagcaaga 110747
human.FAM230E-LOC105377182-GGT3P.NCBI.                                                ------------------------------------------------------------ 57310
human.FAM230J-LOC105372942-GGTLC5P.NCBI.rev.compl                                     ------------------------------------------------------------ 46364
human.FAM230A.ncRNA.18487127.18500594-GGTLC3.18516335.18518165.rev.compl.REF          ------------------------------------------------------------ 31039

chimp.LOC112206744-LOC107973052-GGT2.revcompl                                         ------------------------------------------------------------ 61762
chimp54546-135457.revcompl.Pan.troglodytes.clone.rp43-41g5.GenBank:AC099533.36      ------------------------------------------------------------ 73442
human.FAM30B-LOC105372935-GGT2.NCBI.                                                  agaaaattcagttgctcctgggcaagaattcttgggcccagtggggtagaaccaaaccat 110807
human.FAM230E-LOC105377182-GGT3P.NCBI.                                                ------------------------------------------------------------ 57310
human.FAM230J-LOC105372942-GGTLC5P.NCBI.rev.compl                                     ------------------------------------------------------------ 46364
human.FAM230A.ncRNA.18487127.18500594-GGTLC3.18516335.18518165.rev.compl.REF          ------------------------------------------------------------ 31039

chimp.LOC112206744-LOC107973052-GGT2.revcompl                                         ------------------------------------------------------------ 61762
chimp54546-135457.revcompl.Pan.troglodytes.clone.rp43-41g5.GenBank:AC099533.36      ------------------------------------------------------------ 73442
human.FAM30B-LOC105372935-GGT2.NCBI.                                                  gataaggagacaccagtgttttgtaagcaagagaatatattaaggagagaaaaggggcat 110867
human.FAM230E-LOC105377182-GGT3P.NCBI.                                                ------------------------------------------------------------ 57310
human.FAM230J-LOC105372942-GGTLC5P.NCBI.rev.compl                                     ------------------------------------------------------------ 46364
human.FAM230A.ncRNA.18487127.18500594-GGTLC3.18516335.18518165.rev.compl.REF          ------------------------------------------------------------ 31039

chimp.LOC112206744-LOC107973052-GGT2.revcompl                                         ------------------------------------------------------------ 61762
chimp54546-135457.revcompl.Pan.troglodytes.clone.rp43-41g5.GenBank:AC099533.36      ----------------------------TCCTCAGGATTCCTTTGCACCTGCCTGGCTGG 73474
human.FAM30B-LOC105372935-GGT2.NCBI.                                                  ttattgggagaaaaggtctcactgtgattgttctggaataaagtagccccatcggactgc 110927
human.FAM230E-LOC105377182-GGT3P.NCBI.                                                ------------------------------------------------------------ 57310
human.FAM230J-LOC105372942-GGTLC5P.NCBI.rev.compl                                     ------------------------------------------------------------ 46364
human.FAM230A.ncRNA.18487127.18500594-GGTLC3.18516335.18518165.rev.compl.REF          ------------------------------------------------------------ 31039

chimp.LOC112206744-LOC107973052-GGT2.revcompl                                         ------------------------------------------------------------ 61762
chimp54546-135457.revcompl.Pan.troglodytes.clone.rp43-41g5.GenBank:AC099533.36      GACAGAGCTTCTGGATGAAAGGAGC----------------------------------- 73501
```

```
human.FAM30B-LOC105372935-GGT2.NCBI.                                          ctcaggaactcatggtggaagaggtgccactggaaaatggctaagtcgggatcttcccca 110987
human.FAM230E-LOC105377182-GGT3P.NCBI.                                        ------------------------------------------------------------ 57310
human.FAM230J-LOC105372942-GGTLC5P.NCBI.rev.compl                             ------------------------------------------------------------ 46364
human.FAM230A.ncRNA.18487127.18500594-GGTLC3.18516335.18518165.rev.compl.REF  ------------------------------------------------------------ 31039

chimp.LOC112206744-LOC107973052-GGT2.revcompl                                 ------------------------------------------------------------ 61762
chimp54546-135457.revcompl.Pan.troglodytes.clone.rp43-41g5.GenBank:AC099533.36 ------------------------------------------------------------ 73501
human.FAM30B-LOC105372935-GGT2.NCBI.                                          agaacactagaagagacagaggaagaatagaagagctttaaattttagcagatatttctt 111047
human.FAM230E-LOC105377182-GGT3P.NCBI.                                        ------------------------------------------------------------ 57310
human.FAM230J-LOC105372942-GGTLC5P.NCBI.rev.compl                             ------------------------------------------------------------ 46364
human.FAM230A.ncRNA.18487127.18500594-GGTLC3.18516335.18518165.rev.compl.REF  ------------------------------------------------------------ 31039

chimp.LOC112206744-LOC107973052-GGT2.revcompl                                 ------------------------------------------------------------ 61762
chimp54546-135457.revcompl.Pan.troglodytes.clone.rp43-41g5.GenBank:AC099533.36 ------------------------------------------------------CTGACT 73507
human.FAM30B-LOC105372935-GGT2.NCBI.                                          tttcttgtttaattaaaataaatcttattgacctaatattttgagactgggctcttgca 111107
human.FAM230E-LOC105377182-GGT3P.NCBI.                                        ------------------------------------------------------------ 57310
human.FAM230J-LOC105372942-GGTLC5P.NCBI.rev.compl                             ------------------------------------------------------------ 46364
human.FAM230A.ncRNA.18487127.18500594-GGTLC3.18516335.18518165.rev.compl.REF  ------------------------------------------------------------ 31039

chimp.LOC112206744-LOC107973052-GGT2.revcompl                                 ------------------------------------------------------------ 61762
chimp54546-135457.revcompl.Pan.troglodytes.clone.rp43-41g5.GenBank:AC099533.36 GTGCCCCTCAGAGTGCCCATGAGCACAGGAGTACAGGAAGATTTAACCTTACCAGCCCCT 73567
human.FAM30B-LOC105372935-GGT2.NCBI.                                          atgctccccagactgaactcaaactcctaggctcaagagattcttccacttcagcctcct 111167
human.FAM230E-LOC105377182-GGT3P.NCBI.                                        ------------------------------------------------------------ 57310
human.FAM230J-LOC105372942-GGTLC5P.NCBI.rev.compl                             ------------------------------------------------------------ 46364
human.FAM230A.ncRNA.18487127.18500594-GGTLC3.18516335.18518165.rev.compl.REF  ------------------------------------------------------------ 31039

chimp.LOC112206744-LOC107973052-GGT2.revcompl                                 ------------------------------------------------------------ 61762
chimp54546-135457.revcompl.Pan.troglodytes.clone.rp43-41g5.GenBank:AC099533.36 CACCCTGCAG-------------------------------------------------- 73577
human.FAM30B-LOC105372935-GGT2.NCBI.                                          gaaaagctggcactacaggtgcacaccactgggcccagcataatggctatttcttgacaa 111227
human.FAM230E-LOC105377182-GGT3P.NCBI.                                        ------------------------------------------------------------ 57310
human.FAM230J-LOC105372942-GGTLC5P.NCBI.rev.compl                             ------------------------------------------------------------ 46364
human.FAM230A.ncRNA.18487127.18500594-GGTLC3.18516335.18518165.rev.compl.REF  ------------------------------------------------------------ 31039

chimp.LOC112206744-LOC107973052-GGT2.revcompl                                 ------------------------------------------------------------ 61762
chimp54546-135457.revcompl.Pan.troglodytes.clone.rp43-41g5.GenBank:AC099533.36 -------CAAGGTCATGAGCATCCTTTCCACCACAGGCAGAGCCCTGCT---GAGCTCAC 73627
human.FAM30B-LOC105372935-GGT2.NCBI.                                          atcttgacaatgccaccagtatgtttttctttagagacagagtctcgctttgtagcccag 111287
human.FAM230E-LOC105377182-GGT3P.NCBI.                                        ------------------------------------------------------------ 57310
human.FAM230J-LOC105372942-GGTLC5P.NCBI.rev.compl                             ------------------------------------------------------------ 46364
human.FAM230A.ncRNA.18487127.18500594-GGTLC3.18516335.18518165.rev.compl.REF  ------------------------------------------------------------ 31039

chimp.LOC112206744-LOC107973052-GGT2.revcompl                                 ------------------------------------------------------------ 61762
chimp54546-135457.revcompl.Pan.troglodytes.clone.rp43-41g5.GenBank:AC099533.36 AATCAAGGCCAG-GCTGGTCCCACAGCTCGGGGCTGGGCGGGTGGCTGGGACTCTGGAGG 73686
human.FAM30B-LOC105372935-GGT2.NCBI.                                          gctggagaacagtggtgtgatcctggctcactgcagcctcaaacacctggactcaggaga 111347
human.FAM230E-LOC105377182-GGT3P.NCBI.                                        ------------------------------------------------------------ 57310
human.FAM230J-LOC105372942-GGTLC5P.NCBI.rev.compl                             ------------------------------------------------------------ 46364
human.FAM230A.ncRNA.18487127.18500594-GGTLC3.18516335.18518165.rev.compl.REF  ------------------------------------------------------------ 31039

chimp.LOC112206744-LOC107973052-GGT2.revcompl                                 ------------------------------------------------------------ 61762
chimp54546-135457.revcompl.Pan.troglodytes.clone.rp43-41g5.GenBank:AC099533.36 CAGCACCTGTGCAGTGAGCGAGGAGGCTGAGAAAGGTCCAGTCCATGACTGAGACATCCC 73746
human.FAM30B-LOC105372935-GGT2.NCBI.                                          tccccctgcctcagcctctcaggtagctgaga---------------------------- 111379
human.FAM230E-LOC105377182-GGT3P.NCBI.                                        ------------------------------------------------------------ 57310
human.FAM230J-LOC105372942-GGTLC5P.NCBI.rev.compl                             ------------------------------------------------------------ 46364
human.FAM230A.ncRNA.18487127.18500594-GGTLC3.18516335.18518165.rev.compl.REF  ------------------------------------------------------------ 31039

chimp.LOC112206744-LOC107973052-GGT2.revcompl                                 ------------------------------------------------------------ 61762
chimp54546-135457.revcompl.Pan.troglodytes.clone.rp43-41g5.GenBank:AC099533.36 TGATGCTGCTTCCTGTGGCCCAGGGCGGGTAGGTTTCTCCCAGGCCTCTCTTATCCCCTT 73806
human.FAM30B-LOC105372935-GGT2.NCBI.                                          ------------------------------------------------------------ 111379
human.FAM230E-LOC105377182-GGT3P.NCBI.                                        ------------------------------------------------------------ 57310
human.FAM230J-LOC105372942-GGTLC5P.NCBI.rev.compl                             ------------------------------------------------------------ 46364
human.FAM230A.ncRNA.18487127.18500594-GGTLC3.18516335.18518165.rev.compl.REF  ------------------------------------------------------------ 31039

chimp.LOC112206744-LOC107973052-GGT2.revcompl                                 ------------------------------------------------------------ 61762
chimp54546-135457.revcompl.Pan.troglodytes.clone.rp43-41g5.GenBank:AC099533.36 GATGGAGAAAGCGGCCCGTGATGCAAGTCAGCAGAATCAGGGGCTGCTACTGGAGTAGCT 73866
human.FAM30B-LOC105372935-GGT2.NCBI.                                          --------------ctacaggtgcgcaccaccacacttggcgaattttattttatttttt 111425
human.FAM230E-LOC105377182-GGT3P.NCBI.                                        ------------------------------------------------------------ 57310
human.FAM230J-LOC105372942-GGTLC5P.NCBI.rev.compl                             ------------------------------------------------------------ 46364
human.FAM230A.ncRNA.18487127.18500594-GGTLC3.18516335.18518165.rev.compl.REF  ------------------------------------------------------------ 31039

chimp.LOC112206744-LOC107973052-GGT2.revcompl                                 ------------------------------------------------------------ 61762
chimp54546-135457.revcompl.Pan.troglodytes.clone.rp43-41g5.GenBank:AC099533.36 ----TTGTAGTTTCCAGAGAAGGGCTCAGCCCCAAGGGACACAGAGAGGGGAGGGGAGGA 73922
human.FAM30B-LOC105372935-GGT2.NCBI.                                          tttattatactttaagttctagggtacatacacctggctaattataaaatgtttatctgt 111485
human.FAM230E-LOC105377182-GGT3P.NCBI.                                        ------------------------------------------------------------ 57310
human.FAM230J-LOC105372942-GGTLC5P.NCBI.rev.compl                             ------------------------------------------------------------ 46364
human.FAM230A.ncRNA.18487127.18500594-GGTLC3.18516335.18518165.rev.compl.REF  ------------------------------------------------------------ 31039

chimp.LOC112206744-LOC107973052-GGT2.revcompl                                 ------------------------------------------------------------ 61762
chimp54546-135457.revcompl.Pan.troglodytes.clone.rp43-41g5.GenBank:AC099533.36 CCCTACAGATCCACCT-------------------------------------------- 73938
human.FAM30B-LOC105372935-GGT2.NCBI.                                          acagacaggttctcgttatgttgcccaggctggtctcgaactctcggcctcaagcaatcc 111545
human.FAM230E-LOC105377182-GGT3P.NCBI.                                        ------------------------------------------------------------ 57310
human.FAM230J-LOC105372942-GGTLC5P.NCBI.rev.compl                             ------------------------------------------------------------ 46364
human.FAM230A.ncRNA.18487127.18500594-GGTLC3.18516335.18518165.rev.compl.REF  ------------------------------------------------------------ 31039

chimp.LOC112206744-LOC107973052-GGT2.revcompl                                 ------------------------------------------------------------ 61762
chimp54546-135457.revcompl.Pan.troglodytes.clone.rp43-41g5.GenBank:AC099533.36 -----------------------------------------CAGCCCACAGAATTCTCCT 73957
human.FAM30B-LOC105372935-GGT2.NCBI.                                          tctcctcctcaccttccaaagccccgaaatatgcccacatccagcctgccaccagtttct 111605
human.FAM230E-LOC105377182-GGT3P.NCBI.                                        ------------------------------------------------------------ 57310
human.FAM230J-LOC105372942-GGTLC5P.NCBI.rev.compl                             ------------------------------------------------------------ 46364
human.FAM230A.ncRNA.18487127.18500594-GGTLC3.18516335.18518165.rev.compl.REF  ------------------------------------------------------------ 31039

chimp.LOC112206744-LOC107973052-GGT2.revcompl                                 ------------------------------------------------------------ 61762
chimp54546-135457.revcompl.Pan.troglodytes.clone.rp43-41g5.GenBank:AC099533.36 TCACTTGCTGGTCCCAAAGTCTGGGCCAACATCTTTACCTTAAACTATGTCTTGGCCTCG 74017
human.FAM30B-LOC105372935-GGT2.NCBI.                                          ttgcatacttctttcctaggcttgaccaagacctttactttcttctgtgtacagctcagc 111665
human.FAM230E-LOC105377182-GGT3P.NCBI.                                        ------------------------------------------------------------ 57310
human.FAM230J-LOC105372942-GGTLC5P.NCBI.rev.compl                             ------------------------------------------------------------ 46364
human.FAM230A.ncRNA.18487127.18500594-GGTLC3.18516335.18518165.rev.compl.REF  ------------------------------------------------------------ 31039

chimp.LOC112206744-LOC107973052-GGT2.revcompl                                 ------------------------------------------------------------ 61762
chimp54546-135457.revcompl.Pan.troglodytes.clone.rp43-41g5.GenBank:AC099533.36 GTT--------------------------------------------------------- 74020
human.FAM30B-LOC105372935-GGT2.NCBI.                                          attgaaaaaaaacaaaaaagtttaggctgggcacaatgggtcacgcctgtaatcccaaca 111725
human.FAM230E-LOC105377182-GGT3P.NCBI.                                        ------------------------------------------------------------ 57310
human.FAM230J-LOC105372942-GGTLC5P.NCBI.rev.compl                             ------------------------------------------------------------ 46364
human.FAM230A.ncRNA.18487127.18500594-GGTLC3.18516335.18518165.rev.compl.REF  ------------------------------------------------------------ 31039

chimp.LOC112206744-LOC107973052-GGT2.revcompl                                 ------------------------------------------------------------ 61762
chimp54546-135457.revcompl.Pan.troglodytes.clone.rp43-41g5.GenBank:AC099533.36 --------------------------------------------CTCAGCTAGCCAGGCC 74036
human.FAM30B-LOC105372935-GGT2.NCBI.                                          cttagggtggccaaggtgggaggtttgcttgaacccaggagttccacaccagcctaggca 111785
human.FAM230E-LOC105377182-GGT3P.NCBI.                                        ------------------------------------------------------------ 57310
human.FAM230J-LOC105372942-GGTLC5P.NCBI.rev.compl                             ------------------------------------------------------------ 46364
human.FAM230A.ncRNA.18487127.18500594-GGTLC3.18516335.18518165.rev.compl.REF  ------------------------------------------------------------ 31039

chimp.LOC112206744-LOC107973052-GGT2.revcompl                                 ------------------------------------------------------------ 61762
chimp54546-135457.revcompl.Pan.troglodytes.clone.rp43-41g5.GenBank:AC099533.36 AGGTGGAGTTCACAGATCATTCTGTCTGCATTTT-------------------------- 74070
human.FAM30B-LOC105372935-GGT2.NCBI.                                          agatggcaagaccacatctctccaaaacaatttttgtaaaattagccaggtgtgctggtg 111845
human.FAM230E-LOC105377182-GGT3P.NCBI.                                        ------------------------------------------------------------ 57310
human.FAM230J-LOC105372942-GGTLC5P.NCBI.rev.compl                             ------------------------------------------------------------ 46364
human.FAM230A.ncRNA.18487127.18500594-GGTLC3.18516335.18518165.rev.compl.REF  ------------------------------------------------------------ 31039

chimp.LOC112206744-LOC107973052-GGT2.revcompl                                 ------------------------------------------------------------ 61762
chimp54546-135457.revcompl.Pan.troglodytes.clone.rp43-41g5.GenBank:AC099533.36 ------------------------------------GTGAGCCCATTCTATAGTTAACAG 74094
human.FAM30B-LOC105372935-GGT2.NCBI.                                          cgggcctgcagtctcagctagtgagaagactaaggcaagaggaacccttgaacccaggag 111905
human.FAM230E-LOC105377182-GGT3P.NCBI.                                        ------------------------------------------------------------ 57310
human.FAM230J-LOC105372942-GGTLC5P.NCBI.rev.compl                             ------------------------------------------------------------ 46364
human.FAM230A.ncRNA.18487127.18500594-GGTLC3.18516335.18518165.rev.compl.REF  ------------------------------------------------------------ 31039

chimp.LOC112206744-LOC107973052-GGT2.revcompl                                 ------------------------------------------------------------ 61762
chimp54546-135457.revcompl.Pan.troglodytes.clone.rp43-41g5.GenBank:AC099533.36 CAGAAGACTTTGATAGGTTCTGGGTCCTGCCCCAGAGCAGGTG-TGGATTCAGAACCTGC 74153
human.FAM30B-LOC105372935-GGT2.NCBI.                                          ttcaaggctacggtgagctgtgatcacctctccagcctgggtgatagggtgagacccagt 111965
```

```
human.FAM230E-LOC105377182-GGT3P.NCBI.                                            ------------------------------------------------------------ 57310
human.FAM230J-LOC105372942-GGTLC5P.NCBI.rev.compl                                 ------------------------------------------------------------ 46364
human.FAM230A.ncRNA.18487127.18500594-GGTLC3.18516335.18518165.rev.compl.REF      ------------------------------------------------------------ 31039

chimp.LOC112206744-LOC107973052-GGT2.revcompl                                     ------------------------------------------------------------ 61762
chimp54546-135457.revcompl.Pan.troglodytes.clone.rp43-41g5.GenBank:AC099533.36    TGCTCAGGTCCCT----------GATTCCCTTGTCCATGGACTGCTGGTTCCAAGAGAGG 74203
human.FAM30B-LOC105372935-GGT2.NCBI.                                              ctctgaaaacaattaaaaataaaaaaagttgaataatgagtcttctttttagaggcaatg 112025
human.FAM230E-LOC105377182-GGT3P.NCBI.                                            ------------------------------------------------------------ 57310
human.FAM230J-LOC105372942-GGTLC5P.NCBI.rev.compl                                 ------------------------------------------------------------ 46364
human.FAM230A.ncRNA.18487127.18500594-GGTLC3.18516335.18518165.rev.compl.REF      ------------------------------------------------------------ 31039

chimp.LOC112206744-LOC107973052-GGT2.revcompl                                     ------------------------------------------------------------ 61762
chimp54546-135457.revcompl.Pan.troglodytes.clone.rp43-41g5.GenBank:AC099533.36    AATTCCTCTTCTTTATTTGGCCTTATATGCCTCAATACCAGGCATAAGGGGCTCGGTTAC 74263
human.FAM30B-LOC105372935-GGT2.NCBI.                                              tatttttcctctttttgttttccttaggctatatgcacaatagacaaagttgaaatttaa 112085
human.FAM230E-LOC105377182-GGT3P.NCBI.                                            ------------------------------------------------------------ 57310
human.FAM230J-LOC105372942-GGTLC5P.NCBI.rev.compl                                 ------------------------------------------------------------ 46364
human.FAM230A.ncRNA.18487127.18500594-GGTLC3.18516335.18518165.rev.compl.REF      ------------------------------------------------------------ 31039

chimp.LOC112206744-LOC107973052-GGT2.revcompl                                     ------------------------------------------------------------ 61762
chimp54546-135457.revcompl.Pan.troglodytes.clone.rp43-41g5.GenBank:AC099533.36    CCAT-------------------------------------------------------- 74267
human.FAM30B-LOC105372935-GGT2.NCBI.                                              tcttaatctcgtaaacccttactgtccaaacacatcaaactcaaaaaaaaaaaaagacaa 112145
human.FAM230E-LOC105377182-GGT3P.NCBI.                                            ------------------------------------------------------------ 57310
human.FAM230J-LOC105372942-GGTLC5P.NCBI.rev.compl                                 ------------------------------------------------------------ 46364
human.FAM230A.ncRNA.18487127.18500594-GGTLC3.18516335.18518165.rev.compl.REF      ------------------------------------------------------------ 31039

chimp.LOC112206744-LOC107973052-GGT2.revcompl                                     ------------------------------------------------------------ 61762
chimp54546-135457.revcompl.Pan.troglodytes.clone.rp43-41g5.GenBank:AC099533.36    ---GAAATTATGAGCACGTCCTATTGGAGGAAAATGGGTAGGGCCC-------------- 74310
human.FAM30B-LOC105372935-GGT2.NCBI.                                              aaacaaaaaaggtgaaaggcgtacaagagaaaaaagggaagtaatctgtgccaaatgtac 112205
human.FAM230E-LOC105377182-GGT3P.NCBI.                                            ------------------------------------------------------------ 57310
human.FAM230J-LOC105372942-GGTLC5P.NCBI.rev.compl                                 ------------------------------------------------------------ 46364
human.FAM230A.ncRNA.18487127.18500594-GGTLC3.18516335.18518165.rev.compl.REF      ------------------------------------------------------------ 31039

chimp.LOC112206744-LOC107973052-GGT2.revcompl                                     ------------------------------------------------------------ 61762
chimp54546-135457.revcompl.Pan.troglodytes.clone.rp43-41g5.GenBank:AC099533.36    -------------------ACCTAAGACATCCCTCTACTGTCACTGGGTTGGTAAGTGGT 74351
human.FAM30B-LOC105372935-GGT2.NCBI.                                              attccttccacaatttgggacaaaaatgatgcctacctggtctctgggtgccaaaataag 112265
human.FAM230E-LOC105377182-GGT3P.NCBI.                                            ------------------------------------------------------------ 57310
human.FAM230J-LOC105372942-GGTLC5P.NCBI.rev.compl                                 ------------------------------------------------------------ 46364
human.FAM230A.ncRNA.18487127.18500594-GGTLC3.18516335.18518165.rev.compl.REF      ------------------------------------------------------------ 31039

chimp.LOC112206744-LOC107973052-GGT2.revcompl                                     ------------------------------------------------------------ 61762
chimp54546-135457.revcompl.Pan.troglodytes.clone.rp43-41g5.GenBank:AC099533.36    CCAATGCATCTTACCAAGGCTCTGCATATAGTGCCGCAGACCTCAGACACCAAGTTTCAT 74411
human.FAM30B-LOC105372935-GGT2.NCBI.                                              acattagagacactcccctc-cttcaagaaataaaattggaatcagacactaagatacaa 112324
human.FAM230E-LOC105377182-GGT3P.NCBI.                                            ------------------------------------------------------------ 57310
human.FAM230J-LOC105372942-GGTLC5P.NCBI.rev.compl                                 ------------------------------------------------------------ 46364
human.FAM230A.ncRNA.18487127.18500594-GGTLC3.18516335.18518165.rev.compl.REF      ------------------------------------------------------------ 31039

chimp.LOC112206744-LOC107973052-GGT2.revcompl                                     ------------------------------------------------------------ 61762
chimp54546-135457.revcompl.Pan.troglodytes.clone.rp43-41g5.GenBank:AC099533.36    GGCTCCAGGTGTAGAAAAACATGAGGCAGTGGTGGCTCCTGCCCCAGAAACAAAGGCAGG 74471
human.FAM30B-LOC105372935-GGT2.NCBI.                                              ttaaacagactttgaaagttggtatgcactactgggaggtgtctcacagct---ggcagg 112381
human.FAM230E-LOC105377182-GGT3P.NCBI.                                            ------------------------------------------------------------ 57310
human.FAM230J-LOC105372942-GGTLC5P.NCBI.rev.compl                                 ------------------------------------------------------------ 46364
human.FAM230A.ncRNA.18487127.18500594-GGTLC3.18516335.18518165.rev.compl.REF      ------------------------------------------------------------ 31039

chimp.LOC112206744-LOC107973052-GGT2.revcompl                                     ------------------------------------------------------------ 61762
chimp54546-135457.revcompl.Pan.troglodytes.clone.rp43-41g5.GenBank:AC099533.36    AGCATATCCACCCAGCCAATA--------------------------------------- 74492
human.FAM30B-LOC105372935-GGT2.NCBI.                                              gtcaactgcaactagaagaaaagctcctaagggcagaaactaagtggctcacttcctctg 112441
human.FAM230E-LOC105377182-GGT3P.NCBI.                                            ------------------------------------------------------------ 57310
human.FAM230J-LOC105372942-GGTLC5P.NCBI.rev.compl                                 ------------------------------------------------------------ 46364
human.FAM230A.ncRNA.18487127.18500594-GGTLC3.18516335.18518165.rev.compl.REF      ------------------------------------------------------------ 31039

chimp.LOC112206744-LOC107973052-GGT2.revcompl                                     ------------------------------------------------------------ 61762
chimp54546-135457.revcompl.Pan.troglodytes.clone.rp43-41g5.GenBank:AC099533.36    --------------------------------------------------TTTATTGAGA 74502
human.FAM30B-LOC105372935-GGT2.NCBI.                                              taacactgccgtgcccagcactgtaccagcaccccagcagagtcaacaactttccgtcaa 112501
human.FAM230E-LOC105377182-GGT3P.NCBI.                                            ------------------------------------------------------------ 57310
human.FAM230J-LOC105372942-GGTLC5P.NCBI.rev.compl                                 ------------------------------------------------------------ 46364
human.FAM230A.ncRNA.18487127.18500594-GGTLC3.18516335.18518165.rev.compl.REF      ------------------------------------------------------------ 31039

chimp.LOC112206744-LOC107973052-GGT2.revcompl                                     ------------------------------------------------------------ 61762
chimp54546-135457.revcompl.Pan.troglodytes.clone.rp43-41g5.GenBank:AC099533.36    CCCACTCTGATCCGGCAATATTCCATCCTGGGAATTCAGCATGAGAAAGACTGTCTCTGA 74562
human.FAM30B-LOC105372935-GGT2.NCBI.                                              gccacacacttacggtctcatcgtccagtggaggccgcagatgagacatagtgtgccaaa 112561
human.FAM230E-LOC105377182-GGT3P.NCBI.                                            ------------------------------------------------------------ 57310
human.FAM230J-LOC105372942-GGTLC5P.NCBI.rev.compl                                 ------------------------------------------------------------ 46364
human.FAM230A.ncRNA.18487127.18500594-GGTLC3.18516335.18518165.rev.compl.REF      ------------------------------------------------------------ 31039

chimp.LOC112206744-LOC107973052-GGT2.revcompl                                     ------------------------------------------------------------ 61762
chimp54546-135457.revcompl.Pan.troglodytes.clone.rp43-41g5.GenBank:AC099533.36    CCCCAGGAGTTTCCTACATGGGGGTTGGGGCTGTGTCATAGAAACAGTTTGTAGTAGAAG 74622
human.FAM30B-LOC105372935-GGT2.NCBI.                                              tcccaaatg--------------------------------------------------- 112570
human.FAM230E-LOC105377182-GGT3P.NCBI.                                            ------------------------------------------------------------ 57310
human.FAM230J-LOC105372942-GGTLC5P.NCBI.rev.compl                                 ------------------------------------------------------------ 46364
human.FAM230A.ncRNA.18487127.18500594-GGTLC3.18516335.18518165.rev.compl.REF      ------------------------------------------------------------ 31039

chimp.LOC112206744-LOC107973052-GGT2.revcompl                                     ------------------------------------------------------------ 61762
chimp54546-135457.revcompl.Pan.troglodytes.clone.rp43-41g5.GenBank:AC099533.36    CACCATCTTCTACCTGTGTATTTTCTCCTCTATTTATTTACAAAAAGATCCTTGTTTTCC 74682
human.FAM30B-LOC105372935-GGT2.NCBI.                                              ------------------------------------------------------------ 112570
human.FAM230E-LOC105377182-GGT3P.NCBI.                                            ------------------------------------------------------------ 57310
human.FAM230J-LOC105372942-GGTLC5P.NCBI.rev.compl                                 ------------------------------------------------------------ 46364
human.FAM230A.ncRNA.18487127.18500594-GGTLC3.18516335.18518165.rev.compl.REF      ------------------------------------------------------------ 31039

chimp.LOC112206744-LOC107973052-GGT2.revcompl                                     ------------------------------------------------------------ 61762
chimp54546-135457.revcompl.Pan.troglodytes.clone.rp43-41g5.GenBank:AC099533.36    CTGGTCACTCTTGGCTTGGACTGACCACATGGTCCCATGCTGGCCCCAGGGCTGTAAGGG 74742
human.FAM30B-LOC105372935-GGT2.NCBI.                                              -------------------------cacagggtgccagacaagtctgcgggaggcaggag 112605
human.FAM230E-LOC105377182-GGT3P.NCBI.                                            ------------------------------------------------------------ 57310
human.FAM230J-LOC105372942-GGTLC5P.NCBI.rev.compl                                 ------------------------------------------------------------ 46364
human.FAM230A.ncRNA.18487127.18500594-GGTLC3.18516335.18518165.rev.compl.REF      ------------------------------------------------------------ 31039

chimp.LOC112206744-LOC107973052-GGT2.revcompl                                     ------------------------------------------------------------ 61762
chimp54546-135457.revcompl.Pan.troglodytes.clone.rp43-41g5.GenBank:AC099533.36    TCAGGTGGTTG------------------------------------------------- 74753
human.FAM30B-LOC105372935-GGT2.NCBI.                                              gactagggaagccttccaggcaaggtgatgcctgagctgagttgtttttcttttgagttt 112665
human.FAM230E-LOC105377182-GGT3P.NCBI.                                            ------------------------------------------------------------ 57310
human.FAM230J-LOC105372942-GGTLC5P.NCBI.rev.compl                                 ------------------------------------------------------------ 46364
human.FAM230A.ncRNA.18487127.18500594-GGTLC3.18516335.18518165.rev.compl.REF      ------------------------------------------------------------ 31039

chimp.LOC112206744-LOC107973052-GGT2.revcompl                                     ------------------------------------------------------------ 61762
chimp54546-135457.revcompl.Pan.troglodytes.clone.rp43-41g5.GenBank:AC099533.36    ------------------------------------------------------------ 74753
human.FAM30B-LOC105372935-GGT2.NCBI.                                              ggaacattaatttaatcggaggaaaacaagggacatgaaaagggcaaggaaataagaaaa 112725
human.FAM230E-LOC105377182-GGT3P.NCBI.                                            ------------------------------------------------------------ 57310
human.FAM230J-LOC105372942-GGTLC5P.NCBI.rev.compl                                 ------------------------------------------------------------ 46364
human.FAM230A.ncRNA.18487127.18500594-GGTLC3.18516335.18518165.rev.compl.REF      ------------------------------------------------------------ 31039

chimp.LOC112206744-LOC107973052-GGT2.revcompl                                     ------------------------------------------------------------ 61762
chimp54546-135457.revcompl.Pan.troglodytes.clone.rp43-41g5.GenBank:AC099533.36    -------------------------GAGATCTTCTCAGGGAACATTTTCTCTTGTGATGG 74788
human.FAM30B-LOC105372935-GGT2.NCBI.                                              tgagctgagttttgaaggcaatgatcagatgaagtgagagaagaatattccagcagaggg 112785
human.FAM230E-LOC105377182-GGT3P.NCBI.                                            ------------------------------------------------------------ 57310
human.FAM230J-LOC105372942-GGTLC5P.NCBI.rev.compl                                 ------------------------------------------------------------ 46364
human.FAM230A.ncRNA.18487127.18500594-GGTLC3.18516335.18518165.rev.compl.REF      ------------------------------------------------------------ 31039

chimp.LOC112206744-LOC107973052-GGT2.revcompl                                     ------------------------------------------------------------ 61762
chimp54546-135457.revcompl.Pan.troglodytes.clone.rp43-41g5.GenBank:AC099533.36    AAATAAACAGACTG-------------------------TTTTCGTGCAGCACCTTTGCT 74823
human.FAM30B-LOC105372935-GGT2.NCBI.                                              agacaaggagactgcagagatttcaaggggccaagagctgttcggtgcagctcagtgaag 112845
human.FAM230E-LOC105377182-GGT3P.NCBI.                                            ------------------------------------------------------------ 57310
human.FAM230J-LOC105372942-GGTLC5P.NCBI.rev.compl                                 ------------------------------------------------------------ 46364
human.FAM230A.ncRNA.18487127.18500594-GGTLC3.18516335.18518165.rev.compl.REF      ------------------------------------------------------------ 31039

chimp.LOC112206744-LOC107973052-GGT2.revcompl                                     ------------------------------------------------------------ 61762
chimp54546-135457.revcompl.Pan.troglodytes.clone.rp43-41g5.GenBank:AC099533.36    CTTGAATATGGGTATAATAGCTGGAGCTGCAGCAGCCATCCTGAGGTCTTAAGTCCACAC 74883
human.FAM30B-LOC105372935-GGT2.NCBI.                                              cacctaaacagaggtgggggaaggagaacaggggtcaaagcccagtattgaagtgttctg 112905
human.FAM230E-LOC105377182-GGT3P.NCBI.                                            ------------------------------------------------------------ 57310
```

```
human.FAM230J-LOC105372942-GGTLC5P.NCBI.rev.compl                                  ------------------------------------------------------------ 46364
human.FAM230A.ncRNA.18487127.18500594-GGTLC3.18516335.18518165.rev.compl.REF       ------------------------------------------------------------ 31039

chimp.LOC112206744-LOC107973052-GGT2.revcompl                                      ------------------------------------------------------------ 61762
chimp54546-135457.revcompl.Pan.troglodytes.clone.rp43-41g5.GenBank:AC099533.36     AAATGAAAGAAAAGCCAACTGTGTCATGA------------TGGTGGAGCAGAAAGTTGG 74931
human.FAM30B-LOC105372935-GGT2.NCBI.                                               gataccaagctaaggagaccttattacgaaggtactggggctgctgaagggtttaaatgg 112965
human.FAM230E-LOC105377182-GGT3P.NCBI.                                             ------------------------------------------------------------ 57310
human.FAM230J-LOC105372942-GGTLC5P.NCBI.rev.compl                                  ------------------------------------------------------------ 46364
human.FAM230A.ncRNA.18487127.18500594-GGTLC3.18516335.18518165.rev.compl.REF       ------------------------------------------------------------ 31039

chimp.LOC112206744-LOC107973052-GGT2.revcompl                                      ------------------------------------------------------------ 61762
chimp54546-135457.revcompl.Pan.troglodytes.clone.rp43-41g5.GenBank:AC099533.36     GTAATTGCTGGGTTTCTCATTATTTGGTGAGTATGTGGGGAACAGCAGTGAA-CTCCACA 74990
human.FAM30B-LOC105372935-GGT2.NCBI.                                               gagagtgattgtgaacagattaatagtttggaaagatacaggcagaagtaaacctctgga 113025
human.FAM230E-LOC105377182-GGT3P.NCBI.                                             ------------------------------------------------------------ 57310
human.FAM230J-LOC105372942-GGTLC5P.NCBI.rev.compl                                  ------------------------------------------------------------ 46364
human.FAM230A.ncRNA.18487127.18500594-GGTLC3.18516335.18518165.rev.compl.REF       ------------------------------------------------------------ 31039

chimp.LOC112206744-LOC107973052-GGT2.revcompl                                      ------------------------------------------------------------ 61762
chimp54546-135457.revcompl.Pan.troglodytes.clone.rp43-41g5.GenBank:AC099533.36     CTTACTTTTTGAGGATAAAATAGCAAATGGTTCATGACTATTTCTCATGAGTGTCCAAGC 75050
human.FAM30B-LOC105372935-GGT2.NCBI.                                               tttaaccaaaaaataaaggacagaaccacaatgaagatgttggaaaagatcaggaaaaac 113085
human.FAM230E-LOC105377182-GGT3P.NCBI.                                             ------------------------------------------------------------ 57310
human.FAM230J-LOC105372942-GGTLC5P.NCBI.rev.compl                                  ------------------------------------------------------------ 46364
human.FAM230A.ncRNA.18487127.18500594-GGTLC3.18516335.18518165.rev.compl.REF       ------------------------------------------------------------ 31039

chimp.LOC112206744-LOC107973052-GGT2.revcompl                                      ------------------------------------------------------------ 61762
chimp54546-135457.revcompl.Pan.troglodytes.clone.rp43-41g5.GenBank:AC099533.36     AGAGGAGAAGCATATTTGATTGATTGTAGGAGTATTTAGCACTAGAAAATCAATTAATAA 75110
human.FAM30B-LOC105372935-GGT2.NCBI.                                               actgttaaaagatattcactctcggggccgggcgctgtggctcacgccttaaatcccggc 113145
human.FAM230E-LOC105377182-GGT3P.NCBI.                                             ------------------------------------------------------------ 57310
human.FAM230J-LOC105372942-GGTLC5P.NCBI.rev.compl                                  ------------------------------------------------------------ 46364
human.FAM230A.ncRNA.18487127.18500594-GGTLC3.18516335.18518165.rev.compl.REF       ------------------------------------------------------------ 31039

chimp.LOC112206744-LOC107973052-GGT2.revcompl                                      ------------------------------------------------------------ 61762
chimp54546-135457.revcompl.Pan.troglodytes.clone.rp43-41g5.GenBank:AC099533.36     ACAGTTAGGAAAAGAGGCATGAAGTTTAGGTGCTCTGCAGATCATTAATAGAATGCATAC 75170
human.FAM30B-LOC105372935-GGT2.NCBI.                                               actttggtaggccgaggcgggctgatcacgaggtcaggagatcgagaccatcctggctaa 113205
human.FAM230E-LOC105377182-GGT3P.NCBI.                                             ------------------------------------------------------------ 57310
human.FAM230J-LOC105372942-GGTLC5P.NCBI.rev.compl                                  ------------------------------------------------------------ 46364
human.FAM230A.ncRNA.18487127.18500594-GGTLC3.18516335.18518165.rev.compl.REF       ------------------------------------------------------------ 31039

chimp.LOC112206744-LOC107973052-GGT2.revcompl                                      ------------------------------------------------------------ 61762
chimp54546-135457.revcompl.Pan.troglodytes.clone.rp43-41g5.GenBank:AC099533.36     ACA--------------------------------------------------------- 75173
human.FAM30B-LOC105372935-GGT2.NCBI.                                               cacggtcaaaccccgtctctactaaaaatacaaaaaattagccgggtgtggtggcgggcg 113265
human.FAM230E-LOC105377182-GGT3P.NCBI.                                             ------------------------------------------------------------ 57310
human.FAM230J-LOC105372942-GGTLC5P.NCBI.rev.compl                                  ------------------------------------------------------------ 46364
human.FAM230A.ncRNA.18487127.18500594-GGTLC3.18516335.18518165.rev.compl.REF       ------------------------------------------------------------ 31039

chimp.LOC112206744-LOC107973052-GGT2.revcompl                                      ------------------------------------------------------------ 61762
chimp54546-135457.revcompl.Pan.troglodytes.clone.rp43-41g5.GenBank:AC099533.36     ---------------------------------------------GAAAATGACAGGCAG 75188
human.FAM30B-LOC105372935-GGT2.NCBI.                                               cctgtagtcccaactactccagaggctgaggcaggagaatggcgtgaacccggtaggcgg 113325
human.FAM230E-LOC105377182-GGT3P.NCBI.                                             ------------------------------------------------------------ 57310
human.FAM230J-LOC105372942-GGTLC5P.NCBI.rev.compl                                  ------------------------------------------------------------ 46364
human.FAM230A.ncRNA.18487127.18500594-GGTLC3.18516335.18518165.rev.compl.REF       ------------------------------------------------------------ 31039

chimp.LOC112206744-LOC107973052-GGT2.revcompl                                      ------------------------------------------------------------ 61762
chimp54546-135457.revcompl.Pan.troglodytes.clone.rp43-41g5.GenBank:AC099533.36     ACTGCTGTGTAATCGGAGGTTGGGGCGCCTCCCTGAAGAGGAGTCTCTGGAGCTGCTCTC 75248
human.FAM30B-LOC105372935-GGT2.NCBI.                                               agcttgcggtgagccgagatcaggccactggaatccagcctgggcgacagagggagactc 113385
human.FAM230E-LOC105377182-GGT3P.NCBI.                                             ------------------------------------------------------------ 57310
human.FAM230J-LOC105372942-GGTLC5P.NCBI.rev.compl                                  ------------------------------------------------------------ 46364
human.FAM230A.ncRNA.18487127.18500594-GGTLC3.18516335.18518165.rev.compl.REF       ------------------------------------------------------------ 31039

chimp.LOC112206744-LOC107973052-GGT2.revcompl                                      ------------------------------------------------------------ 61762
chimp54546-135457.revcompl.Pan.troglodytes.clone.rp43-41g5.GenBank:AC099533.36     AGGAAGGAAGAAACAGTGAAAGGAAGTGTGAGGAAGAAGAGTCTGTCACTGACCTGCAGG 75308
human.FAM30B-LOC105372935-GGT2.NCBI.                                               catctcaaaaaaaaaaaaaaaaaaaaagagag---agagagtctgactctgttgccaaga 113442
human.FAM230E-LOC105377182-GGT3P.NCBI.                                             ------------------------------------------------------------ 57310
human.FAM230J-LOC105372942-GGTLC5P.NCBI.rev.compl                                  ------------------------------------------------------------ 46364
human.FAM230A.ncRNA.18487127.18500594-GGTLC3.18516335.18518165.rev.compl.REF       ------------------------------------------------------------ 31039

chimp.LOC112206744-LOC107973052-GGT2.revcompl                                      ------------------------------------------------------------ 61762
chimp54546-135457.revcompl.Pan.troglodytes.clone.rp43-41g5.GenBank:AC099533.36     GAGGGGT---------------------------------------------------AG 75317
human.FAM30B-LOC105372935-GGT2.NCBI.                                               ctggagtgcagtggcgcgatctaggcacatcacaatcccttccccgcccccgggttcaag 113502
human.FAM230E-LOC105377182-GGT3P.NCBI.                                             ------------------------------------------------------------ 57310
human.FAM230J-LOC105372942-GGTLC5P.NCBI.rev.compl                                  ------------------------------------------------------------ 46364
human.FAM230A.ncRNA.18487127.18500594-GGTLC3.18516335.18518165.rev.compl.REF       ------------------------------------------------------------ 31039

chimp.LOC112206744-LOC107973052-GGT2.revcompl                                      ------------------------------------------------------------ 61762
chimp54546-135457.revcompl.Pan.troglodytes.clone.rp43-41g5.GenBank:AC099533.36     GTGTTTTCGAGGATCTGAAAGAGCAGTGACAAACATCAAGGGGGTCTGGGATCCAAACTC 75377
human.FAM30B-LOC105372935-GGT2.NCBI.                                               tgattctcctgtctcagccgccggagtagctgggactacaggcgcgtgccaccatgtctg 113562
human.FAM230E-LOC105377182-GGT3P.NCBI.                                             ------------------------------------------------------------ 57310
human.FAM230J-LOC105372942-GGTLC5P.NCBI.rev.compl                                  ------------------------------------------------------------ 46364
human.FAM230A.ncRNA.18487127.18500594-GGTLC3.18516335.18518165.rev.compl.REF       ------------------------------------------------------------ 31039

chimp.LOC112206744-LOC107973052-GGT2.revcompl                                      ------------------------------------------------------------ 61762
chimp54546-135457.revcompl.Pan.troglodytes.clone.rp43-41g5.GenBank:AC099533.36     ATTGCAATGCAGTGTATGATA--------------------------------------- 75398
human.FAM30B-LOC105372935-GGT2.NCBI.                                               attaaatctgtattttactagagacggggtttcactctgttggccaggctggtctccaa 113622
human.FAM230E-LOC105377182-GGT3P.NCBI.                                             ------------------------------------------------------------ 57310
human.FAM230J-LOC105372942-GGTLC5P.NCBI.rev.compl                                  ------------------------------------------------------------ 46364
human.FAM230A.ncRNA.18487127.18500594-GGTLC3.18516335.18518165.rev.compl.REF       ------------------------------------------------------------ 31039

chimp.LOC112206744-LOC107973052-GGT2.revcompl                                      ------------------------------------------------------------ 61762
chimp54546-135457.revcompl.Pan.troglodytes.clone.rp43-41g5.GenBank:AC099533.36     ------------------------------------------------------------ 75398
human.FAM30B-LOC105372935-GGT2.NCBI.                                               ctcctgatctcgtgatccgcccgccccgacctcccaaagtgctgggatgcgtgagcctcc 113682
human.FAM230E-LOC105377182-GGT3P.NCBI.                                             ------------------------------------------------------------ 57310
human.FAM230J-LOC105372942-GGTLC5P.NCBI.rev.compl                                  ------------------------------------------------------------ 46364
human.FAM230A.ncRNA.18487127.18500594-GGTLC3.18516335.18518165.rev.compl.REF       ------------------------------------------------------------ 31039

chimp.LOC112206744-LOC107973052-GGT2.revcompl                                      ------------------------------------------------------------ 61762
chimp54546-135457.revcompl.Pan.troglodytes.clone.rp43-41g5.GenBank:AC099533.36     ------------------------------------------------------------ 75398
human.FAM30B-LOC105372935-GGT2.NCBI.                                               acgcctggccactaattttctttctttctttcgtgtgtgtgtgtgtgtgtgtgtgtgtg 113742
human.FAM230E-LOC105377182-GGT3P.NCBI.                                             ------------------------------------------------------------ 57310
human.FAM230J-LOC105372942-GGTLC5P.NCBI.rev.compl                                  ------------------------------------------------------------ 46364
human.FAM230A.ncRNA.18487127.18500594-GGTLC3.18516335.18518165.rev.compl.REF       ------------------------------------------------------------ 31039

chimp.LOC112206744-LOC107973052-GGT2.revcompl                                      ------------------------------------------------------------ 61762
chimp54546-135457.revcompl.Pan.troglodytes.clone.rp43-41g5.GenBank:AC099533.36     -----------------------------------CTTTTAGCTGAGGCAGGAGTGCAGC 75423
human.FAM30B-LOC105372935-GGT2.NCBI.                                               tgtgtgtgtgtgtgtgtgtgtgacgaagtttcgctcttgttgcccaggttggagtgcaat 113802
human.FAM230E-LOC105377182-GGT3P.NCBI.                                             ------------------------------------------------------------ 57310
human.FAM230J-LOC105372942-GGTLC5P.NCBI.rev.compl                                  ------------------------------------------------------------ 46364
human.FAM230A.ncRNA.18487127.18500594-GGTLC3.18516335.18518165.rev.compl.REF       ------------------------------------------------------------ 31039

chimp.LOC112206744-LOC107973052-GGT2.revcompl                                      ------------------------------------------------------------ 61762
chimp54546-135457.revcompl.Pan.troglodytes.clone.rp43-41g5.GenBank:AC099533.36     GTATTAAGGTGTGGTTAGAGGAATTTCCAGGTGATGATGAATGATGTGACCTTGGCATCT 75483
human.FAM30B-LOC105372935-GGT2.NCBI.                                               ggtgtgatctcagctcactgcaatccccgcctcaacaggagagcaggaatctt------- 113855
human.FAM230E-LOC105377182-GGT3P.NCBI.                                             ------------------------------------------------------------ 57310
human.FAM230J-LOC105372942-GGTLC5P.NCBI.rev.compl                                  ------------------------------------------------------------ 46364
human.FAM230A.ncRNA.18487127.18500594-GGTLC3.18516335.18518165.rev.compl.REF       ------------------------------------------------------------ 31039

chimp.LOC112206744-LOC107973052-GGT2.revcompl                                      ------------------------------------------------------------ 61762
chimp54546-135457.revcompl.Pan.troglodytes.clone.rp43-41g5.GenBank:AC099533.36     GCTTAGCACATGGTATGGCCTCCAGGACGCTGCTGCTGGGGCGGGGGCAGGAGAGACACA 75543
human.FAM30B-LOC105372935-GGT2.NCBI.                                               ------------------------------------------------------------ 113855
human.FAM230E-LOC105377182-GGT3P.NCBI.                                             ------------------------------------------------------------ 57310
human.FAM230J-LOC105372942-GGTLC5P.NCBI.rev.compl                                  ------------------------------------------------------------ 46364
human.FAM230A.ncRNA.18487127.18500594-GGTLC3.18516335.18518165.rev.compl.REF       ------------------------------------------------------------ 31039

chimp.LOC112206744-LOC107973052-GGT2.revcompl                                      ------------------------------------------------------------ 61762
chimp54546-135457.revcompl.Pan.troglodytes.clone.rp43-41g5.GenBank:AC099533.36     AGTGGGGTCAGACCTGCCCCAGGCTGTTTCCTATTTTAGCAGGGTCTGAAGCTGCCCAGG 75603
human.FAM30B-LOC105372935-GGT2.NCBI.                                               -----------------------cagtgatccactggcggatctgcagccattgtgcgcg 113892
human.FAM230E-LOC105377182-GGT3P.NCBI.                                             ------------------------------------------------------------ 57310
human.FAM230J-LOC105372942-GGTLC5P.NCBI.rev.compl                                  ------------------------------------------------------------ 46364
```

```
human.FAM230A.ncRNA.18487127.18500594-GGTLC3.18516335.18518165.rev.compl.REF          ------------------------------------------------------------ 31039

chimp.LOC112206744-LOC107973052-GGT2.revcompl                                         ------------------------------------------------------------ 61762
chimp54546-135457.revcompl.Pan.troglodytes.clone.rp43-41g5.GenBank:AC099533.36        AGGGGGCTTACCCACCCCTTTGTTCCTCCATTACTTGTGCAACAAGCAGCCTTGTGGGGA 75663
human.FAM30B-LOC105372935-GGT2.NCBI.                                                  ccaggtcttcccaagtcttttgtgcgcgcctctccttccagtacc--------------- 113939
human.FAM230E-LOC105377182-GGT3P.NCBI.                                                ------------------------------------------------------------ 57310
human.FAM230J-LOC105372942-GGTLC5P.NCBI.rev.compl                                     ------------------------------------------------------------ 46364
human.FAM230A.ncRNA.18487127.18500594-GGTLC3.18516335.18518165.rev.compl.REF          ------------------------------------------------------------ 31039

chimp.LOC112206744-LOC107973052-GGT2.revcompl                                         ------------------------------------------------------------ 61762
chimp54546-135457.revcompl.Pan.troglodytes.clone.rp43-41g5.GenBank:AC099533.36        CATACAGTGCTCCATCCTGGGACCCCGTCGAGGGCCTCACATGGGTGACTGATCCCTAAT 75723
human.FAM30B-LOC105372935-GGT2.NCBI.                                                  tatgccgccagcgtcccctagcctcccgccattgccagcaggtgctgagatcgcgccatt 113999
human.FAM230E-LOC105377182-GGT3P.NCBI.                                                ------------------------------------------------------------ 57310
human.FAM230J-LOC105372942-GGTLC5P.NCBI.rev.compl                                     ------------------------------------------------------------ 46364
human.FAM230A.ncRNA.18487127.18500594-GGTLC3.18516335.18518165.rev.compl.REF          ------------------------------------------------------------ 31039

chimp.LOC112206744-LOC107973052-GGT2.revcompl                                         ------------------------------------------------------------ 61762
chimp54546-135457.revcompl.Pan.troglodytes.clone.rp43-41g5.GenBank:AC099533.36        GTCCCACAGACTGGCTGATCTGAGCTGATCACAGTGGGTGGGGAAAACACTGGACAGAGG 75783
human.FAM30B-LOC105372935-GGT2.NCBI.                                                  gcactccagcctggggacaagagcgaaactccatgtcaaaaaaaaaaaa----------- 114048
human.FAM230E-LOC105377182-GGT3P.NCBI.                                                ------------------------------------------------------------ 57310
human.FAM230J-LOC105372942-GGTLC5P.NCBI.rev.compl                                     ------------------------------------------------------------ 46364
human.FAM230A.ncRNA.18487127.18500594-GGTLC3.18516335.18518165.rev.compl.REF          ------------------------------------------------------------ 31039

chimp.LOC112206744-LOC107973052-GGT2.revcompl                                         ------------------------------------------------------------ 61762
chimp54546-135457.revcompl.Pan.troglodytes.clone.rp43-41g5.GenBank:AC099533.36        AGGGAAGGGCCAGAGGGTGAGTAAAGTTCCCAAGGTCGCCTGGGGAGGAAGAGGGGAGAG 75843
human.FAM30B-LOC105372935-GGT2.NCBI.                                                  -------------------------------aaggatgaaaattttgggaaaatatggaa 114077
human.FAM230E-LOC105377182-GGT3P.NCBI.                                                ------------------------------------------------------------ 57310
human.FAM230J-LOC105372942-GGTLC5P.NCBI.rev.compl                                     ------------------------------------------------------------ 46364
human.FAM230A.ncRNA.18487127.18500594-GGTLC3.18516335.18518165.rev.compl.REF          ------------------------------------------------------------ 31039

chimp.LOC112206744-LOC107973052-GGT2.revcompl                                         ------------------------------------------------------------ 61762
chimp54546-135457.revcompl.Pan.troglodytes.clone.rp43-41g5.GenBank:AC099533.36        CAAGACCAGGGAGGTCCTGGGCAGAAATAATGATCATTTCTAAGTGTGCAG---GAGCAA 75900
human.FAM30B-LOC105372935-GGT2.NCBI.                                                  gaaaccaaatggatttctagctcaactaaatcgtaattattcagtgtctagttttggcaa 114137
human.FAM230E-LOC105377182-GGT3P.NCBI.                                                ------------------------------------------------------------ 57310
human.FAM230J-LOC105372942-GGTLC5P.NCBI.rev.compl                                     ------------------------------------------------------------ 46364
human.FAM230A.ncRNA.18487127.18500594-GGTLC3.18516335.18518165.rev.compl.REF          ------------------------------------------------------------ 31039

chimp.LOC112206744-LOC107973052-GGT2.revcompl                                         ------------------------------------------------------------ 61762
chimp54546-135457.revcompl.Pan.troglodytes.clone.rp43-41g5.GenBank:AC099533.36        GCAAGCTGGATGGGAGTGTC---------GGGAACAATCCTGCACCACAATTGTGGGGC 75951
human.FAM30B-LOC105372935-GGT2.NCBI.                                                  gaaaccacatatttcatgtccacaatcagggacacagtcccagctctcaagtgtgggtct 114197
human.FAM230E-LOC105377182-GGT3P.NCBI.                                                ------------------------------------------------------------ 57310
human.FAM230J-LOC105372942-GGTLC5P.NCBI.rev.compl                                     ------------------------------------------------------------ 46364
human.FAM230A.ncRNA.18487127.18500594-GGTLC3.18516335.18518165.rev.compl.REF          ------------------------------------------------------------ 31039

chimp.LOC112206744-LOC107973052-GGT2.revcompl                                         ------------------------------------------------------------ 61762
chimp54546-135457.revcompl.Pan.troglodytes.clone.rp43-41g5.GenBank:AC099533.36        TCAGAAGG---------------------------------------------------- 75959
human.FAM30B-LOC105372935-GGT2.NCBI.                                                  ttcctaagcaaattgaagaacacaggcataaaagtacattaaatcaataaacctttttct 114257
human.FAM230E-LOC105377182-GGT3P.NCBI.                                                ------------------------------------------------------------ 57310
human.FAM230J-LOC105372942-GGTLC5P.NCBI.rev.compl                                     ------------------------------------------------------------ 46364
human.FAM230A.ncRNA.18487127.18500594-GGTLC3.18516335.18518165.rev.compl.REF          ------------------------------------------------------------ 31039

chimp.LOC112206744-LOC107973052-GGT2.revcompl                                         ------------------------------------------------------------ 61762
chimp54546-135457.revcompl.Pan.troglodytes.clone.rp43-41g5.GenBank:AC099533.36        --------ATATAGCTCTTCCCTTTCAGTTCTGCCAGCTGCCTGTTGGGGGAGCTGCTGC 76011
human.FAM30B-LOC105372935-GGT2.NCBI.                                                  ctctgtctctctctctctttttttttttttttttgagacggaggttggcactg-----tc 114312
human.FAM230E-LOC105377182-GGT3P.NCBI.                                                ------------------------------------------------------------ 57310
human.FAM230J-LOC105372942-GGTLC5P.NCBI.rev.compl                                     ------------------------------------------------------------ 46364
human.FAM230A.ncRNA.18487127.18500594-GGTLC3.18516335.18518165.rev.compl.REF          ------------------------------------------------------------ 31039

chimp.LOC112206744-LOC107973052-GGT2.revcompl                                         ------------------------------------------------------------ 61762
chimp54546-135457.revcompl.Pan.troglodytes.clone.rp43-41g5.GenBank:AC099533.36        ACCCAGACATGAGTGAGCCAGGCACCTGGGGGGTAGCTTCCAGCTC-------------- 76057
human.FAM30B-LOC105372935-GGT2.NCBI.                                                  acccaggctggagtgctgtggtgagatctcgggtcactgcaagctccgcctcccgagttc 114372
human.FAM230E-LOC105377182-GGT3P.NCBI.                                                ------------------------------------------------------------ 57310
human.FAM230J-LOC105372942-GGTLC5P.NCBI.rev.compl                                     ------------------------------------------------------------ 46364
human.FAM230A.ncRNA.18487127.18500594-GGTLC3.18516335.18518165.rev.compl.REF          ------------------------------------------------------------ 31039

chimp.LOC112206744-LOC107973052-GGT2.revcompl                                         ------------------------------------------------------------ 61762
chimp54546-135457.revcompl.Pan.troglodytes.clone.rp43-41g5.GenBank:AC099533.36        ------------------------------------------------------------ 76057
human.FAM30B-LOC105372935-GGT2.NCBI.                                                  acgccatcctcctgcctcagcctccagagtagctgggactacaagcgcccgccacgacgc 114432
human.FAM230E-LOC105377182-GGT3P.NCBI.                                                ------------------------------------------------------------ 57310
human.FAM230J-LOC105372942-GGTLC5P.NCBI.rev.compl                                     ------------------------------------------------------------ 46364
human.FAM230A.ncRNA.18487127.18500594-GGTLC3.18516335.18518165.rev.compl.REF          ------------------------------------------------------------ 31039

chimp.LOC112206744-LOC107973052-GGT2.revcompl                                         ------------------------------------------------------------ 61762
chimp54546-135457.revcompl.Pan.troglodytes.clone.rp43-41g5.GenBank:AC099533.36        ------------------------------------------------------------ 76057
human.FAM30B-LOC105372935-GGT2.NCBI.                                                  ccggctaatttttttgtagttttagtagagacggggtttcgctatgttggccaggctggtc 114492
human.FAM230E-LOC105377182-GGT3P.NCBI.                                                ------------------------------------------------------------ 57310
human.FAM230J-LOC105372942-GGTLC5P.NCBI.rev.compl                                     ------------------------------------------------------------ 46364
human.FAM230A.ncRNA.18487127.18500594-GGTLC3.18516335.18518165.rev.compl.REF          ------------------------------------------------------------ 31039

chimp.LOC112206744-LOC107973052-GGT2.revcompl                                         ------------------------------------------------------------ 61762
chimp54546-135457.revcompl.Pan.troglodytes.clone.rp43-41g5.GenBank:AC099533.36        ----------------------------------------------CCTGCACCCTTTGG 76071
human.FAM30B-LOC105372935-GGT2.NCBI.                                                  tccaactcgtgacatcgtgatccgcccgcctccgactcccaaagtgccacagcccaggcc 114552
human.FAM230E-LOC105377182-GGT3P.NCBI.                                                ------------------------------------------------------------ 57310
human.FAM230J-LOC105372942-GGTLC5P.NCBI.rev.compl                                     ------------------------------------------------------------ 46364
human.FAM230A.ncRNA.18487127.18500594-GGTLC3.18516335.18518165.rev.compl.REF          ------------------------------------------------------------ 31039

chimp.LOC112206744-LOC107973052-GGT2.revcompl                                         ------------------------------------------------------------ 61762
chimp54546-135457.revcompl.Pan.troglodytes.clone.rp43-41g5.GenBank:AC099533.36        TGCTCATAAAAACTCCGAGGGTTGGCTGGGCACGGTGGCTCAAGCCTGTAATCCCAGCAC 76131
human.FAM30B-LOC105372935-GGT2.NCBI.                                                  tttttttttaagacagagtctcggcccggcgcggtggctcacgcctttaatcccagcac 114612
human.FAM230E-LOC105377182-GGT3P.NCBI.                                                ------------------------------------------------------------ 57310
human.FAM230J-LOC105372942-GGTLC5P.NCBI.rev.compl                                     ------------------------------------------------------------ 46364
human.FAM230A.ncRNA.18487127.18500594-GGTLC3.18516335.18518165.rev.compl.REF          ------------------------------------------------------------ 31039

chimp.LOC112206744-LOC107973052-GGT2.revcompl                                         ------------------------------------------------------------ 61762
chimp54546-135457.revcompl.Pan.troglodytes.clone.rp43-41g5.GenBank:AC099533.36        TTTGGGAGGCCGAGGCGGGCGGATCACGAGGTCAGGAGATCGAGACCATCCTGGCTAACA 76191
human.FAM30B-LOC105372935-GGT2.NCBI.                                                  tttgggaggccgaggcgggctgatcacgaggtcaggaggtcgagaccatcctggctaaca 114672
human.FAM230E-LOC105377182-GGT3P.NCBI.                                                ------------------------------------------------------------ 57310
human.FAM230J-LOC105372942-GGTLC5P.NCBI.rev.compl                                     ------------------------------------------------------------ 46364
human.FAM230A.ncRNA.18487127.18500594-GGTLC3.18516335.18518165.rev.compl.REF          ------------------------------------------------------------ 31039

chimp.LOC112206744-LOC107973052-GGT2.revcompl                                         ------------------------------------------------------------ 61762
chimp54546-135457.revcompl.Pan.troglodytes.clone.rp43-41g5.GenBank:AC099533.36        CGGTGAAACCCCGTCTCTACTAAAAATACAAAAAATTAGCCAGGCGCGGTGGCGGGTGCC 76251
human.FAM30B-LOC105372935-GGT2.NCBI.                                                  cggtcaaaccccgtctctactaaaaatacaaaaaattagccgggtgtggtggcgggcgcc 114732
human.FAM230E-LOC105377182-GGT3P.NCBI.                                                ------------------------------------------------------------ 57310
human.FAM230J-LOC105372942-GGTLC5P.NCBI.rev.compl                                     ------------------------------------------------------------ 46364
human.FAM230A.ncRNA.18487127.18500594-GGTLC3.18516335.18518165.rev.compl.REF          ------------------------------------------------------------ 31039

chimp.LOC112206744-LOC107973052-GGT2.revcompl                                         ------------------------------------------------------------ 61762
chimp54546-135457.revcompl.Pan.troglodytes.clone.rp43-41g5.GenBank:AC099533.36        TGTAATCCCAGCTACTCAGGAGGCTGAGGCAGGAGAATGGCGTGAACCCGGGAGGCAGAG 76311
human.FAM30B-LOC105372935-GGT2.NCBI.                                                  tgtagtccgagctactcccgaggctgaggcaggagaatggcgtgaacccggtaggcggag 114792
human.FAM230E-LOC105377182-GGT3P.NCBI.                                                ------------------------------------------------------------ 57310
human.FAM230J-LOC105372942-GGTLC5P.NCBI.rev.compl                                     ------------------------------------------------------------ 46364
human.FAM230A.ncRNA.18487127.18500594-GGTLC3.18516335.18518165.rev.compl.REF          ------------------------------------------------------------ 31039

chimp.LOC112206744-LOC107973052-GGT2.revcompl                                         ------------------------------------------------------------ 61762
chimp54546-135457.revcompl.Pan.troglodytes.clone.rp43-41g5.GenBank:AC099533.36        CTTGCAGTGAGCCGAGATCACGCCACTGCACTCCAACCTGGGCGACAGAGCGAGACTCTG 76371
human.FAM30B-LOC105372935-GGT2.NCBI.                                                  cttgcggtgagccgagatcaggccactggaatccagcctgggcgacagagggagactccg 114852
human.FAM230E-LOC105377182-GGT3P.NCBI.                                                ------------------------------------------------------------ 57310
human.FAM230J-LOC105372942-GGTLC5P.NCBI.rev.compl                                     ------------------------------------------------------------ 46364
human.FAM230A.ncRNA.18487127.18500594-GGTLC3.18516335.18518165.rev.compl.REF          ------------------------------------------------------------ 31039

chimp.LOC112206744-LOC107973052-GGT2.revcompl                                         ------------------------------------------------------------ 61762
chimp54546-135457.revcompl.Pan.troglodytes.clone.rp43-41g5.GenBank:AC099533.36        ------------------------------------------------------------ 76371
human.FAM30B-LOC105372935-GGT2.NCBI.                                                  tctcaaaaaaaaaaaaaaagagtctggctctgttgcccaggctggagtgcagtggcgcga 114912
human.FAM230E-LOC105377182-GGT3P.NCBI.                                                ------------------------------------------------------------ 57310
human.FAM230J-LOC105372942-GGTLC5P.NCBI.rev.compl                                     ------------------------------------------------------------ 46364
human.FAM230A.ncRNA.18487127.18500594-GGTLC3.18516335.18518165.rev.compl.REF          ------------------------------------------------------------ 31039
```

```
chimp.LOC112206744-LOC107973052-GGT2.revcompl                                 ------------------------------------------------------------ 61762
chimp54546-135457.revcompl.Pan.troglodytes.clone.rp43-41g5.GenBank:AC099533.36 ------------------------------------------------------------ 76371
human.FAM30B-LOC105372935-GGT2.NCBI.                                          tctcggcgcatcgcaatcccttccccgcccccgggttcaagtgattctccagtctcagcc 114972
human.FAM230E-LOC105377182-GGT3P.NCBI.                                        ------------------------------------------------------------ 57310
human.FAM230J-LOC105372942-GGTLC5P.NCBI.rev.compl                             ------------------------------------------------------------ 46364
human.FAM230A.ncRNA.18487127.18500594-GGTLC3.18516335.18518165.rev.compl.REF  ------------------------------------------------------------ 31039

chimp.LOC112206744-LOC107973052-GGT2.revcompl                                 ------------------------------------------------------------ 61762
chimp54546-135457.revcompl.Pan.troglodytes.clone.rp43-41g5.GenBank:AC099533.36 ------------------------------------------------------------ 76371
human.FAM30B-LOC105372935-GGT2.NCBI.                                          gccggagtagctgggactacaggcgcgtgccaccatgtctgactaaatttgtatttttac 115032
human.FAM230E-LOC105377182-GGT3P.NCBI.                                        ------------------------------------------------------------ 57310
human.FAM230J-LOC105372942-GGTLC5P.NCBI.rev.compl                             ------------------------------------------------------------ 46364
human.FAM230A.ncRNA.18487127.18500594-GGTLC3.18516335.18518165.rev.compl.REF  ------------------------------------------------------------ 31039

chimp.LOC112206744-LOC107973052-GGT2.revcompl                                 ------------------------------------------------------------ 61762
chimp54546-135457.revcompl.Pan.troglodytes.clone.rp43-41g5.GenBank:AC099533.36 ------------------------------------------------------------ 76371
human.FAM30B-LOC105372935-GGT2.NCBI.                                          tagagacggggtttcactatgttggccaggctggtctccaactcctgatctcgtgatccg 115092
human.FAM230E-LOC105377182-GGT3P.NCBI.                                        ------------------------------------------------------------ 57310
human.FAM230J-LOC105372942-GGTLC5P.NCBI.rev.compl                             ------------------------------------------------------------ 46364
human.FAM230A.ncRNA.18487127.18500594-GGTLC3.18516335.18518165.rev.compl.REF  ------------------------------------------------------------ 31039

chimp.LOC112206744-LOC107973052-GGT2.revcompl                                 ------------------------------------------------------------ 61762
chimp54546-135457.revcompl.Pan.troglodytes.clone.rp43-41g5.GenBank:AC099533.36 ------------------------------------------------------------ 76371
human.FAM30B-LOC105372935-GGT2.NCBI.                                          tctgccccgacctcccaaagtgctaggattataggcataagccaccacgcccggcctctt 115152
human.FAM230E-LOC105377182-GGT3P.NCBI.                                        ------------------------------------------------------------ 57310
human.FAM230J-LOC105372942-GGTLC5P.NCBI.rev.compl                             ------------------------------------------------------------ 46364
human.FAM230A.ncRNA.18487127.18500594-GGTLC3.18516335.18518165.rev.compl.REF  ------------------------------------------------------------ 31039

chimp.LOC112206744-LOC107973052-GGT2.revcompl                                 ------------------------------------------------------------ 61762
chimp54546-135457.revcompl.Pan.troglodytes.clone.rp43-41g5.GenBank:AC099533.36 ------------------------------------------------------------ 76371
human.FAM30B-LOC105372935-GGT2.NCBI.                                          tttttcttttctttctttatctggagactgagttttgctcttgttgcccaggctgga 115212
human.FAM230E-LOC105377182-GGT3P.NCBI.                                        ------------------------------------------------------------ 57310
human.FAM230J-LOC105372942-GGTLC5P.NCBI.rev.compl                             ------------------------------------------------------------ 46364
human.FAM230A.ncRNA.18487127.18500594-GGTLC3.18516335.18518165.rev.compl.REF  ------------------------------------------------------------ 31039

chimp.LOC112206744-LOC107973052-GGT2.revcompl                                 ------------------------------------------------------------ 61762
chimp54546-135457.revcompl.Pan.troglodytes.clone.rp43-41g5.GenBank:AC099533.36 ------------------------------------------------------------ 76371
human.FAM30B-LOC105372935-GGT2.NCBI.                                          gtgcaatggtgcgatctcagctcactgcaatctccacctcagcaggagagcaggaatctt 115272
human.FAM230E-LOC105377182-GGT3P.NCBI.                                        ------------------------------------------------------------ 57310
human.FAM230J-LOC105372942-GGTLC5P.NCBI.rev.compl                             ------------------------------------------------------------ 46364
human.FAM230A.ncRNA.18487127.18500594-GGTLC3.18516335.18518165.rev.compl.REF  ------------------------------------------------------------ 31039

chimp.LOC112206744-LOC107973052-GGT2.revcompl                                 ------------------------------------------------------------ 61762
chimp54546-135457.revcompl.Pan.troglodytes.clone.rp43-41g5.GenBank:AC099533.36 ------------------------------------------------------------ 76371
human.FAM30B-LOC105372935-GGT2.NCBI.                                          cagtgatccacgggcagatctgccgccattgtgggcacctgtttctcccgcaacctttgt 115332
human.FAM230E-LOC105377182-GGT3P.NCBI.                                        ------------------------------------------------------------ 57310
human.FAM230J-LOC105372942-GGTLC5P.NCBI.rev.compl                             ------------------------------------------------------------ 46364
human.FAM230A.ncRNA.18487127.18500594-GGTLC3.18516335.18518165.rev.compl.REF  ------------------------------------------------------------ 31039

chimp.LOC112206744-LOC107973052-GGT2.revcompl                                 ------------------------------------------------------------ 61762
chimp54546-135457.revcompl.Pan.troglodytes.clone.rp43-41g5.GenBank:AC099533.36 ------------------------------------------------------------ 76371
human.FAM30B-LOC105372935-GGT2.NCBI.                                          gcccgcctctctccttcccgtacctattgcatgaccccccacgtccgcctcccgccattg 115392
human.FAM230E-LOC105377182-GGT3P.NCBI.                                        ------------------------------------------------------------ 57310
human.FAM230J-LOC105372942-GGTLC5P.NCBI.rev.compl                             ------------------------------------------------------------ 46364
human.FAM230A.ncRNA.18487127.18500594-GGTLC3.18516335.18518165.rev.compl.REF  ------------------------------------------------------------ 31039

chimp.LOC112206744-LOC107973052-GGT2.revcompl                                 ------------------------------------------------------------ 61762
chimp54546-135457.revcompl.Pan.troglodytes.clone.rp43-41g5.GenBank:AC099533.36 ------------------------------------------------------------ 76371
human.FAM30B-LOC105372935-GGT2.NCBI.                                          ccagcaagtgcctcgcgcgggtacctggctgcgcttattaatccgttaagctcgctctgt 115452
human.FAM230E-LOC105377182-GGT3P.NCBI.                                        ------------------------------------------------------------ 57310
human.FAM230J-LOC105372942-GGTLC5P.NCBI.rev.compl                             ------------------------------------------------------------ 46364
human.FAM230A.ncRNA.18487127.18500594-GGTLC3.18516335.18518165.rev.compl.REF  ------------------------------------------------------------ 31039

chimp.LOC112206744-LOC107973052-GGT2.revcompl                                 ------------------------------------------------------------ 61762
chimp54546-135457.revcompl.Pan.troglodytes.clone.rp43-41g5.GenBank:AC099533.36 ------------------------------------------------------------ 76371
human.FAM30B-LOC105372935-GGT2.NCBI.                                          cacgggcgccgtgatgtgctcacgcgcccgctccctcaggtttaaaaggcgcgttgcccg 115512
human.FAM230E-LOC105377182-GGT3P.NCBI.                                        ------------------------------------------------------------ 57310
human.FAM230J-LOC105372942-GGTLC5P.NCBI.rev.compl                             ------------------------------------------------------------ 46364
human.FAM230A.ncRNA.18487127.18500594-GGTLC3.18516335.18518165.rev.compl.REF  ------------------------------------------------------------ 31039

chimp.LOC112206744-LOC107973052-GGT2.revcompl                                 ------------------------------------------------------------ 61762
chimp54546-135457.revcompl.Pan.troglodytes.clone.rp43-41g5.GenBank:AC099533.36 ------------------------------------------------------------ 76371
human.FAM30B-LOC105372935-GGT2.NCBI.                                          gcaacagaagaaactgctggcttagccgttggccgagttggcggctggacgaggacgctc 115572
human.FAM230E-LOC105377182-GGT3P.NCBI.                                        ------------------------------------------------------------ 57310
human.FAM230J-LOC105372942-GGTLC5P.NCBI.rev.compl                             ------------------------------------------------------------ 46364
human.FAM230A.ncRNA.18487127.18500594-GGTLC3.18516335.18518165.rev.compl.REF  ------------------------------------------------------------ 31039

chimp.LOC112206744-LOC107973052-GGT2.revcompl                                 ------------------------------------------------------------ 61762
chimp54546-135457.revcompl.Pan.troglodytes.clone.rp43-41g5.GenBank:AC099533.36 ------------------------------------------------------------ 76371
human.FAM30B-LOC105372935-GGT2.NCBI.                                          agagcccagctctcgagagttcaagcaaccgacggttccccactgctcccaggagcggtt 115632
human.FAM230E-LOC105377182-GGT3P.NCBI.                                        ------------------------------------------------------------ 57310
human.FAM230J-LOC105372942-GGTLC5P.NCBI.rev.compl                             ------------------------------------------------------------ 46364
human.FAM230A.ncRNA.18487127.18500594-GGTLC3.18516335.18518165.rev.compl.REF  ------------------------------------------------------------ 31039

chimp.LOC112206744-LOC107973052-GGT2.revcompl                                 ------------------------------------------------------------ 61762
chimp54546-135457.revcompl.Pan.troglodytes.clone.rp43-41g5.GenBank:AC099533.36 ------------------------------------------------------------ 76371
human.FAM30B-LOC105372935-GGT2.NCBI.                                          acctgggcactctgtgcccctccttcctgttcgggcccaggccgaggacctgccagtagg 115692
human.FAM230E-LOC105377182-GGT3P.NCBI.                                        ------------------------------------------------------------ 57310
human.FAM230J-LOC105372942-GGTLC5P.NCBI.rev.compl                             ------------------------------------------------------------ 46364
human.FAM230A.ncRNA.18487127.18500594-GGTLC3.18516335.18518165.rev.compl.REF  ------------------------------------------------------------ 31039

chimp.LOC112206744-LOC107973052-GGT2.revcompl                                 ------------------------------------------------------------ 61762
chimp54546-135457.revcompl.Pan.troglodytes.clone.rp43-41g5.GenBank:AC099533.36 ------------------------------------------------------------ 76371
human.FAM30B-LOC105372935-GGT2.NCBI.                                          gctcagttgcctggagcccgttcagcccatcccccagttcactttgcttgtgggatctcc 115752
human.FAM230E-LOC105377182-GGT3P.NCBI.                                        ------------------------------------------------------------ 57310
human.FAM230J-LOC105372942-GGTLC5P.NCBI.rev.compl                             ------------------------------------------------------------ 46364
human.FAM230A.ncRNA.18487127.18500594-GGTLC3.18516335.18518165.rev.compl.REF  ------------------------------------------------------------ 31039

chimp.LOC112206744-LOC107973052-GGT2.revcompl                                 ------------------------------------------------------------ 61762
chimp54546-135457.revcompl.Pan.troglodytes.clone.rp43-41g5.GenBank:AC099533.36 ------------------------------------------------------------ 76371
human.FAM30B-LOC105372935-GGT2.NCBI.                                          ccgttgctcctgcccgtggactgagtggcaggccatcctacaagcacccggacacttgac 115812
human.FAM230E-LOC105377182-GGT3P.NCBI.                                        ------------------------------------------------------------ 57310
human.FAM230J-LOC105372942-GGTLC5P.NCBI.rev.compl                             ------------------------------------------------------------ 46364
human.FAM230A.ncRNA.18487127.18500594-GGTLC3.18516335.18518165.rev.compl.REF  ------------------------------------------------------------ 31039

chimp.LOC112206744-LOC107973052-GGT2.revcompl                                 ------------------------------------------------------------ 61762
chimp54546-135457.revcompl.Pan.troglodytes.clone.rp43-41g5.GenBank:AC099533.36 ------------------------------------------------------------ 76371
human.FAM30B-LOC105372935-GGT2.NCBI.                                          atcagtggtgtcaagacaactctaagaaggttttccgtgatcctgcaagccctgccttcc 115872
human.FAM230E-LOC105377182-GGT3P.NCBI.                                        ------------------------------------------------------------ 57310
human.FAM230J-LOC105372942-GGTLC5P.NCBI.rev.compl                             ------------------------------------------------------------ 46364
human.FAM230A.ncRNA.18487127.18500594-GGTLC3.18516335.18518165.rev.compl.REF  ------------------------------------------------------------ 31039

chimp.LOC112206744-LOC107973052-GGT2.revcompl                                 ------------------------------------------------------------ 61762
chimp54546-135457.revcompl.Pan.troglodytes.clone.rp43-41g5.GenBank:AC099533.36 ------------------------------------------------------------ 76371
human.FAM30B-LOC105372935-GGT2.NCBI.                                          ttcctgggatcctgccttcaatttgattgcacaggtaccacagcaagccagtgctgcgtg 115932
human.FAM230E-LOC105377182-GGT3P.NCBI.                                        ------------------------------------------------------------ 57310
human.FAM230J-LOC105372942-GGTLC5P.NCBI.rev.compl                             ------------------------------------------------------------ 46364
human.FAM230A.ncRNA.18487127.18500594-GGTLC3.18516335.18518165.rev.compl.REF  ------------------------------------------------------------ 31039

chimp.LOC112206744-LOC107973052-GGT2.revcompl                                 ------------------------------------------------------------ 61762
chimp54546-135457.revcompl.Pan.troglodytes.clone.rp43-41g5.GenBank:AC099533.36 ------------------------------------------------------------ 76371
human.FAM30B-LOC105372935-GGT2.NCBI.                                          ctccgagttccagggcgtcctccagctcagccactgcactgagaacatggactctctgtg 115992
human.FAM230E-LOC105377182-GGT3P.NCBI.                                        ------------------------------------------------------------ 57310
human.FAM230J-LOC105372942-GGTLC5P.NCBI.rev.compl                             ------------------------------------------------------------ 46364
human.FAM230A.ncRNA.18487127.18500594-GGTLC3.18516335.18518165.rev.compl.REF  ------------------------------------------------------------ 31039
```

```
chimp.LOC112206744-LOC107973052-GGT2.revcompl                                 ------------------------------------------------------------ 61762
chimp54546-135457.revcompl.Pan.troglodytes.clone.rp43-41g5.GenBank:AC099533.36 ------------------------------------------------------------ 76371
human.FAM30B-LOC105372935-GGT2.NCBI.                                          gggcccaggagccgggagtcacccctttggggtccacaacagccggctgtccccagactt 116052
human.FAM230E-LOC105377182-GGT3P.NCBI.                                        ------------------------------------------------------------ 57310
human.FAM230J-LOC105372942-GGTLC5P.NCBI.rev.compl                             ------------------------------------------------------------ 46364
human.FAM230A.ncRNA.18487127.18500594-GGTLC3.18516335.18518165.rev.compl.REF  ------------------------------------------------------------ 31039

chimp.LOC112206744-LOC107973052-GGT2.revcompl                                 ------------------------------------------------------------ 61762
chimp54546-135457.revcompl.Pan.troglodytes.clone.rp43-41g5.GenBank:AC099533.36 ------------------------------------------------------------ 76371
human.FAM30B-LOC105372935-GGT2.NCBI.                                          gtgtccagggaagatagtgttgagggccctcaaggagagcggggcagggatgcctgagca 116112
human.FAM230E-LOC105377182-GGT3P.NCBI.                                        ------------------------------------------------------------ 57310
human.FAM230J-LOC105372942-GGTLC5P.NCBI.rev.compl                             ------------------------------------------------------------ 46364
human.FAM230A.ncRNA.18487127.18500594-GGTLC3.18516335.18518165.rev.compl.REF  ------------------------------------------------------------ 31039

chimp.LOC112206744-LOC107973052-GGT2.revcompl                                 ---------  61762
chimp54546-135457.revcompl.Pan.troglodytes.clone.rp43-41g5.GenBank:AC099533.36 ---------  76371
human.FAM30B-LOC105372935-GGT2.NCBI.                                          ggacaagga  116121
human.FAM230E-LOC105377182-GGT3P.NCBI.                                        ---------  57310
human.FAM230J-LOC105372942-GGTLC5P.NCBI.rev.compl                             ---------  46364
human.FAM230A.ncRNA.18487127.18500594-GGTLC3.18516335.18518165.rev.compl.REF  ---------  31039
```
