## Supplementary figures and images for "Formation of human long intergenic non-coding RNA genes and pseudogenes: ancestral sequences are key players"

### Supplementary Figure S2

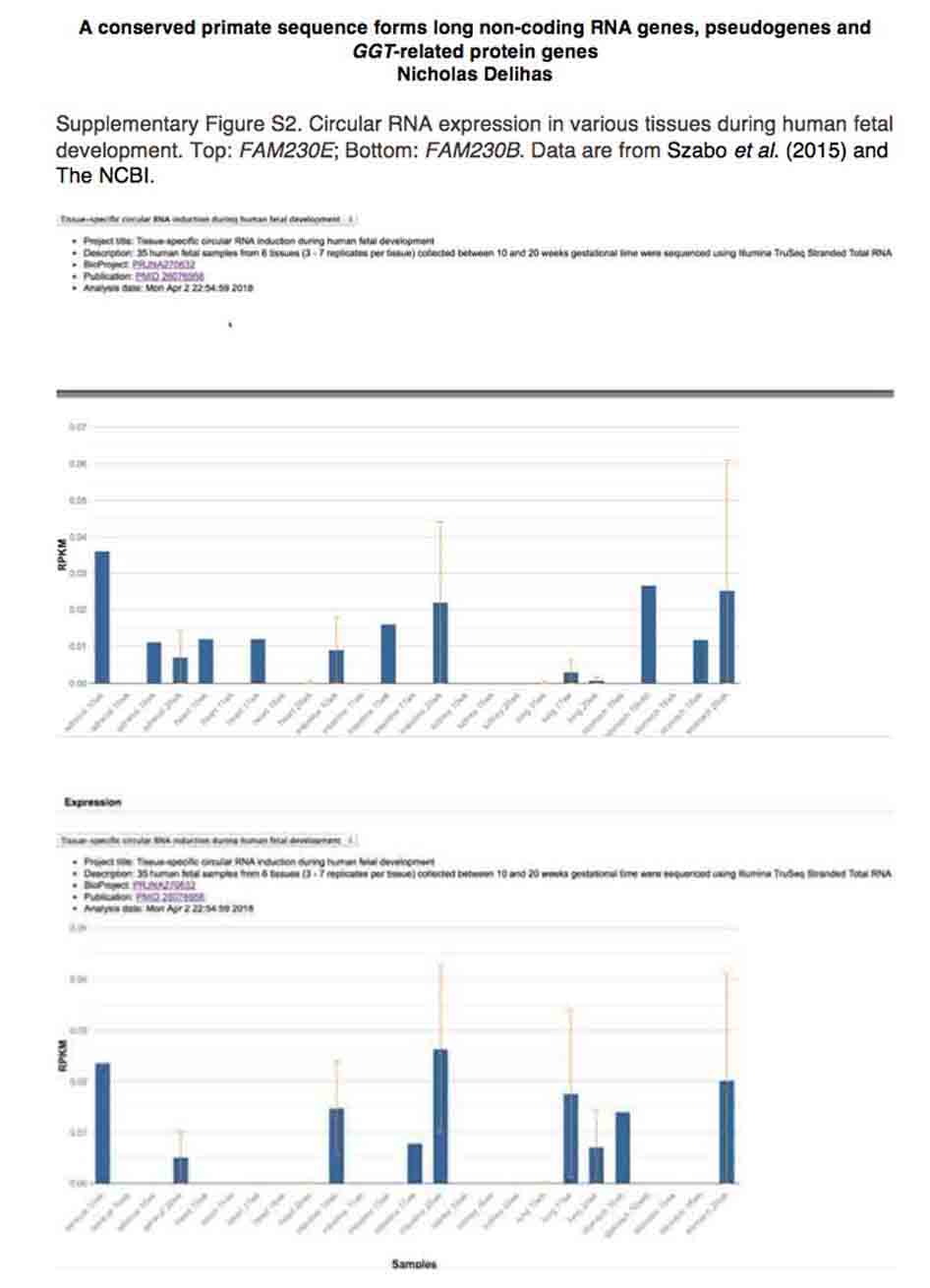

### Supplementary Figure S6

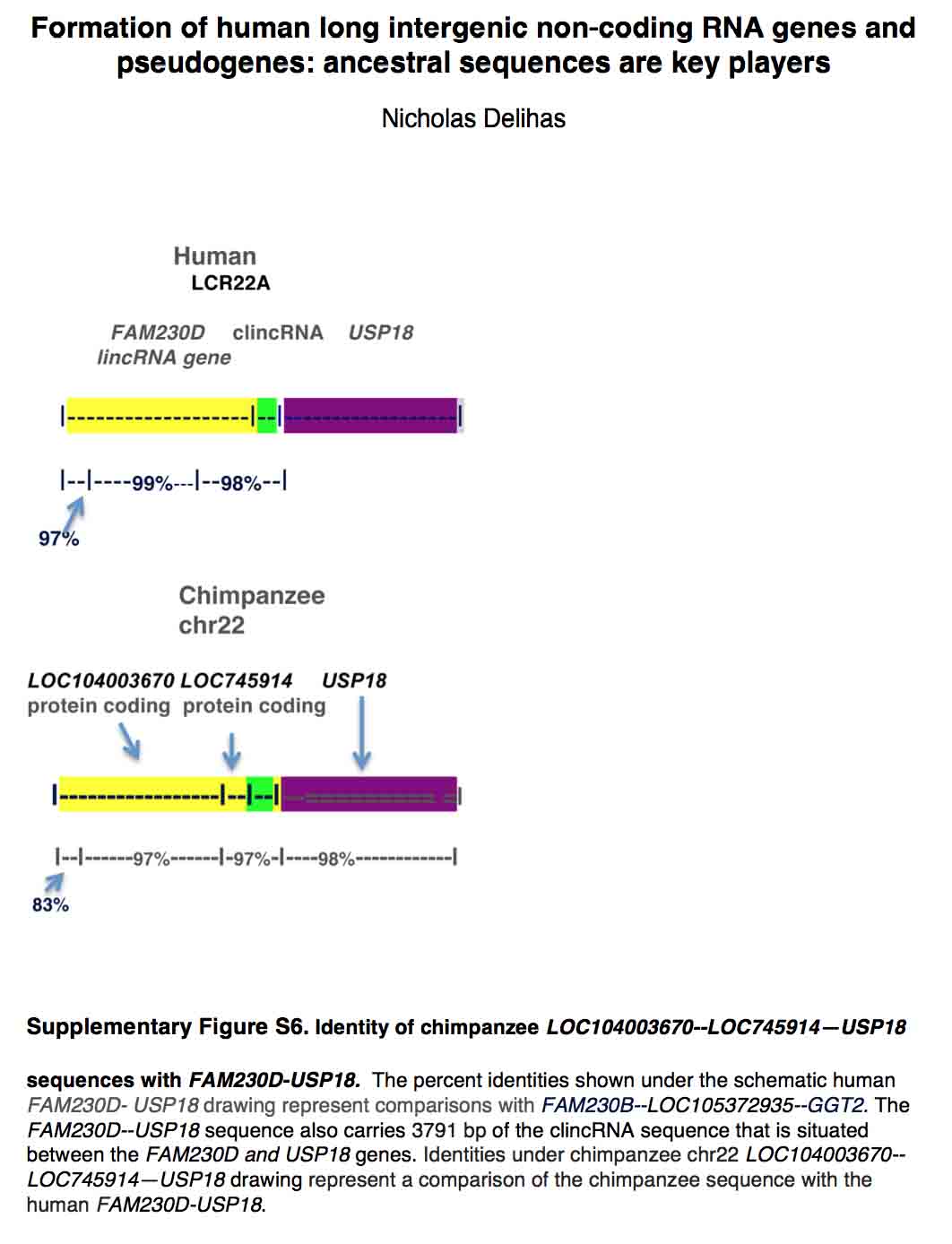

### Supplementary Figure S7

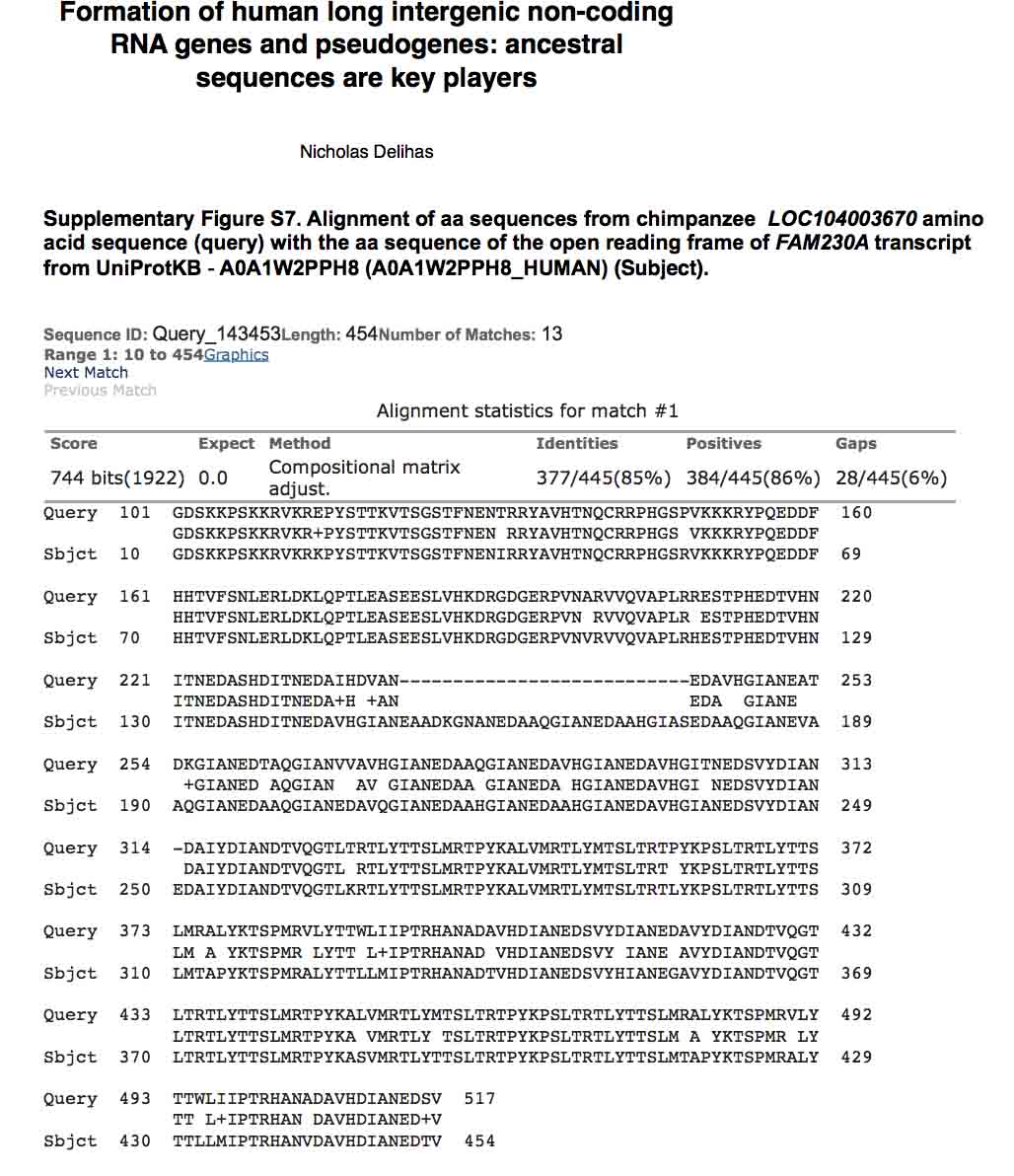

### Supplementary Figure S8

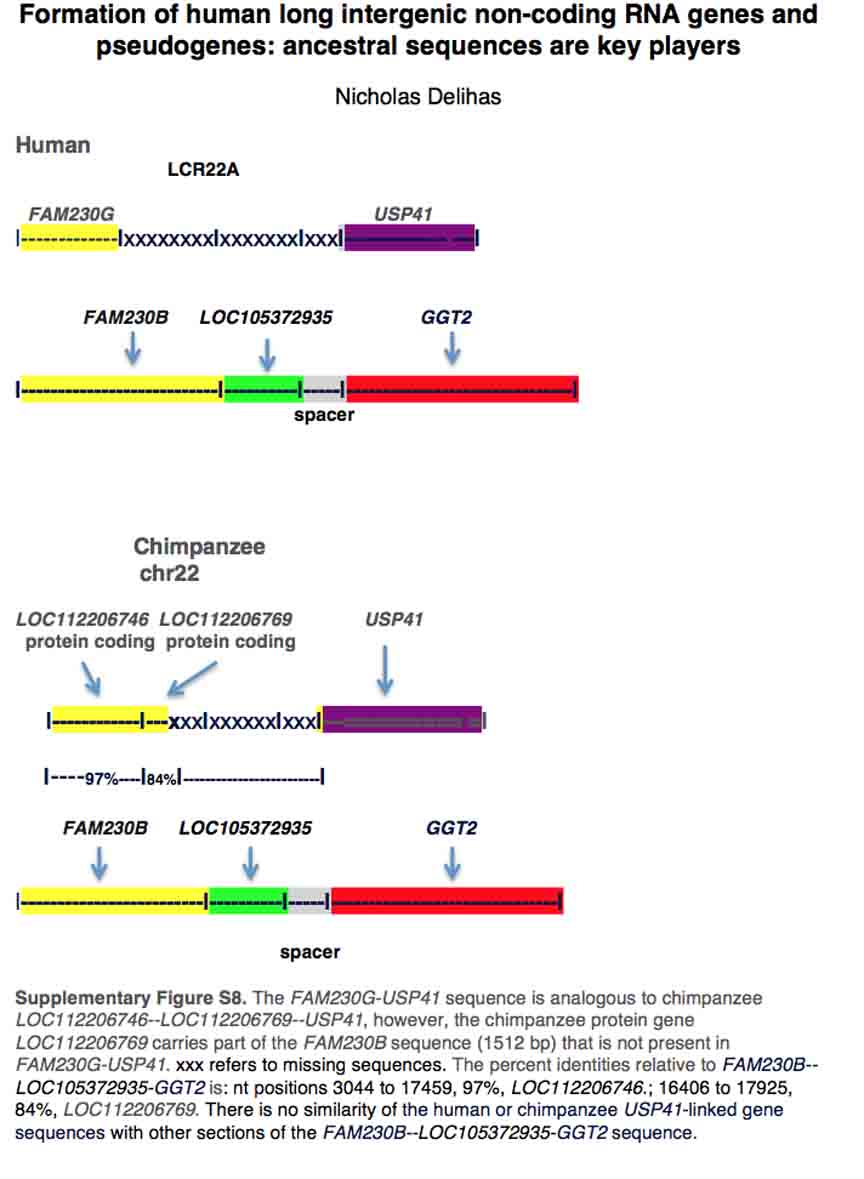
