## Supplementary Figure S3 for "Formation of human long intergenic non-coding RNA genes and pseudogenes: ancestral sequences are key players"

Nicholas Delihas
Department of Molecular Genetics and Microbiology
Renaissance School of Medicine
Stony Brook University
Stony Brook, N.Y. USA 11794

**Supplementary Figure S3. Alignment of human *GGT5* with *LOC105372935* clincRNA gene sequence.**

```
CLUSTAL O(1.2.4) multiple sequence alignment


GGT5.NCBI.REF                       tgcctggcctcctgccagcaacaagcaggagctgaaaaccagaagttgaggcgtgagttt    60
LOC105372935.clincRNA.NCBI.ref      --------------------------------tgaaaactagaagttgaggcatgagttt    28
                                                                    ******* ************ *******

GGT5.NCBI.REF                       ggccactccgtagtgtgcacttggtgagggcagcagctcgccacagctgccagccatctg    120
LOC105372935.clincRNA.NCBI.ref      ggccactccgtagtgtgcacttggtgagggcagcagctcgccacagctgccagccatctg    88
                                    ************************************************************

GGT5.NCBI.REF                       tccattcacccatctgtccatctggcagcccgctgttcagacctgtctgtctgtccgccc    180
LOC105372935.clincRNA.NCBI.ref      tccattcacccatctgtccatctggcagcccgctgttcagacctgtctgtctgtccgccc    148
                                    ************************************************************

GGT5.NCBI.REF                       atctgtaagcccatctctgtcccattgtctatctgaccatctttctcttactgtcctctt    240
LOC105372935.clincRNA.NCBI.ref      atctgtaagcccatctctgtcccattgtctatctgaccatctttctcttactgtcctctt    208
                                    ************************************************************

GGT5.NCBI.REF                       tgtctagctatctggcctatctgtcgatccatcttcgtgtctgtcttcagcccccacctg    300
LOC105372935.clincRNA.NCBI.ref      tgtctagctatctggcctgtctgtcgatccatcttcgtgtctgtcttcagcccccacctg    268
                                    ****************** *****************************************

GGT5.NCBI.REF                       tttgtccatctgtccaattacctgtga----ctctgtgcatcttcttgtccattcatctg    356
LOC105372935.clincRNA.NCBI.ref      tttgtccatctgtccaattacctgtgagtctatctatgcatcttcttgtccattcatctg    328
                                    ***************************     *** ************************

GGT5.NCBI.REF                       cccacccatccgtccctccgtctgcccaccagccgcccctctcctcctgggctgcagagc    416
LOC105372935.clincRNA.NCBI.ref      cccacccatctgtccctccatctgcccaccggcctcccctctccttctgggccgcagagc    388
                                    ********** ******** ********** *** ********** ****** *******

GGT5.NCBI.REF                       catggcccggggctacggggccacggtcagcctagtcct------gctgggtctggggct    470
LOC105372935.clincRNA.NCBI.ref      catggcccaggactacggagccatgggtgacctggtcctgctggggctggggctggggct    448
                                    ******** ** ****** **** **    *** *****      ****** ********

GGT5.NCBI.REF                       ggcgctggctgtcattgtgctggctgtggtcctctctcgacaccaggccccatgtggccc    530
LOC105372935.clincRNA.NCBI.ref      ggcgctggctgtcattgtgctggctgtggtcctctctcgacaccaggccccatttgaccc    508
                                    ***************************************************** ** ***

GGT5.NCBI.REF                       ccaggcctttgcccacgctgctgttgccgccgactccaaggtctgctcggatattggacg    590
LOC105372935.clincRNA.NCBI.ref      cc-ggcctttgcccacgccgctgttgctgctgactccaaggtcttctcaaatattgtacg    567
                                    ** *************** ******** ** ************* ***  ****** ***

GGT5.NCBI.REF                       gtgagtgagacgtgggaggaagctgggtggcccttggcagccagcccctcctggagaagg    650
LOC105372935.clincRNA.NCBI.ref      gtgagtgagacgtgggaggaagctgggtggcctttggcagccagcccctcctggagaagg    627
                                    ******************************** ***************************

GGT5.NCBI.REF                       cgtgtgtgtgt-------------------------------------------------    661
LOC105372935.clincRNA.NCBI.ref      cgtgtgtgtgtgagcatgtgtgtgtgtgagagattatgtgtgagtgtgtgtgggtatatg    687
                                    ***********

GGT5.NCBI.REF                       ----------------------------------------------gtgtgtgtgtgagt    675
LOC105372935.clincRNA.NCBI.ref      tgtgagtgtgtttgtggggtgtgggtgtgtgtgaatgtgtgtgatcgtgtttgggtgtgt    747
                                                                                  **** ** *** **

GGT5.NCBI.REF                       gaatgtatgtgtgggtgtgtgtgtgagtgtctgggtatgtgtgattgcatgcatgtgtgg    735
LOC105372935.clincRNA.NCBI.ref      gtatgtgtgagtgtgggtgtgtgtgaatgtgtgtgattgtgtttgtgtatgtgtgtgtgg    807
                                    * **** ** *** * ********** *** ** *  *****   ** ***  *******

GGT5.NCBI.REF                       gtgtgtgtgagtgtatgtgagtgtgagtgtg-----------------------------    766
LOC105372935.clincRNA.NCBI.ref      gtgtgtgtgagtatatgtgagtgtgagtgtgtggggtgtgggtgggtgtgaatgtgtgt    867
                                    ************ ******************
```

```
GGT5.NCBI.REF                       ------------------------------------------------------------    766
LOC105372935.clincRNA.NCBI.ref      gattgtgtttcgctgtgtgagggtgtgtgtgactgtgagtgtgtgagtgtgggtgtgtgg    927


GGT5.NCBI.REF                       ----------------------------gggagtgtgggtgtgtgtgaatgtgtgtgatt    798
LOC105372935.clincRNA.NCBI.ref      gtgtgtgtaaatgtgtgagtgtgagtatgggggggtgggtatgtgtgaatgtgtgtgatt    987
                                                                *** * ****** *******************

GGT5.NCBI.REF                       gtgtgtgggtatgtgtatgtgtgggtgggtgtgtgt----gtgtgtgtgtgtgcacgtgc    854
LOC105372935.clincRNA.NCBI.ref      gtgtgtgggtatatttgtgggggtgtgtgtgtgtgtgcacgtgtgtgtgtgtgcacgtgc    1047
                                    ************ * * ** * * *** ********    ********************

GGT5.NCBI.REF                       actggcccaggaagcaggagccgtgtgtgtgtgggcttcagcacctgcaggtcttgggcg    914
LOC105372935.clincRNA.NCBI.ref      actggcccaggaagcaggagcc--gtgtgtgtgggcttcagcacctgcagggcttgagcg    1105
                                    **********************  *************************** **** ***

GGT5.NCBI.REF                       caaggaggcagcctcagggcccttgcacagaacaggtggcagggtgtgctcgtggggcag    974
LOC105372935.clincRNA.NCBI.ref      caaggagacagcctcagggcccttgcacagaacaggcggcagggtgtgcccgtggggcag    1165
                                    ******* **************************** ************ **********

GGT5.NCBI.REF                       atggggacttggggacaatggtggtgtgtgagtccacgcctggctccaggattcaggagg    1034
LOC105372935.clincRNA.NCBI.ref      atggggacttggggacaatggtggtgtgtgagtccatacctggctccaggattcaggagg    1225
                                    ************************************  **********************

GGT5.NCBI.REF                       cccatttgcatatcccaggtgggaacctgtctggccccgcctgaccctgctggccggtgc    1094
LOC105372935.clincRNA.NCBI.ref      cccatttgcatatcccaggtgggaacctgtctggccccgcctgaccctgctggccggcgc    1285
                                    ********************************************************* **

GGT5.NCBI.REF                       aggccccttcagtgaggccaattctccaaggctgcggtcttctcccagggtcatgggtga    1154
LOC105372935.clincRNA.NCBI.ref      aggccccttcagtgaggccaattctccaaggctgcggtcttctcccagggtcatgggtga    1345
                                    ************************************************************

GGT5.NCBI.REF                       aggggtttggaggctccctgcgtgggtactggcctgctggggtacacacaatgctgccat    1214
LOC105372935.clincRNA.NCBI.ref      aggggtttggaggctccctgcgtgggtactggcctgctggggtacacacaatgctgccat    1405
                                    ************************************************************

GGT5.NCBI.REF                       agccagtctgccccaacacccagcctggggccacatctcgggtctctcagtcctgaggag    1274
LOC105372935.clincRNA.NCBI.ref      agccagtctgcccctacacccagcccggggccacatctcaggtctctcagtcctgaggag    1465
                                    ************** ********** ************* ********************

GGT5.NCBI.REF                       cccggtgccccacccctcacatcctctctccctgagtcagggcctgggtctcgtgagctg    1334
LOC105372935.clincRNA.NCBI.ref      cccggtgccccacccctcacatcctctctccctgagtcagggcctgggtctcgtgagctg    1525
                                    ************************************************************

GGT5.NCBI.REF                       agtgactgatacttggtgtcctggatgagggcgtggtggagaggggccacagcgggtgtt    1394
LOC105372935.clincRNA.NCBI.ref      agtgactgatacttggtgtcctggatgagggcgtggtggagaggggccacagcgggtgtt    1585
                                    ************************************************************

GGT5.NCBI.REF                       tcctgaccctcttccaggaagcccagcccaagggaggcctccgctgctgctgctgca--g    1452
LOC105372935.clincRNA.NCBI.ref      tcctgaccctcttccaggaag-------------------gtgctgctgccgctgcaggg    1626
                                    *********************                     ******** ******  *

GGT5.NCBI.REF                       aggacacatacaggatgccccttcctgctccctgcctcccactgggcccacaaaagccag    1512
LOC105372935.clincRNA.NCBI.ref      aggacacacacaggatgccccttcttgcccccctgcctcccattgggcccacaaaagccag    1686
                                    ******** *************** *** ************ ******************

GGT5.NCBI.REF                       ggcaagcctcccctccctgccagccacctggtctgcttcccagaagttctgtcttgcagg    1572
LOC105372935.clincRNA.NCBI.ref      ggcaagcctcccctccctgccagccacctggtctgcttcccagaaattctgtcttgcagg    1746
                                    ********************************************* **************

GGT5.NCBI.REF                       ctgttgggaggatcccagtgctttgtaaattaaagcaagggaggagtggctgctctctct    1632
LOC105372935.clincRNA.NCBI.ref      ctgttgggaggatcccagtactttgtaaactaaagcaagggaggagtggccgttctctct    1806
                                    ******************* ********* ******************** * *******

GGT5.NCBI.REF                       ctctgttcattcattcaccttttcattcattccttcttccctccattcccccatctgtcc    1692
LOC105372935.clincRNA.NCBI.ref      ctttgttcattcattcaccttttcattcattccttcttccctccattcccccatctgtgc    1866
                                    ** ******************************************************* *

GGT5.NCBI.REF                       atccttccctgccctgatttctcatgccacccccc----gcccctcctgacctggtcctt    1748
LOC105372935.clincRNA.NCBI.ref      atccttccctgccctgattgctcatgccacccccccccagcccctcctgacctggtcctt    1926
                                    ******************* ***************    *********************

GGT5.NCBI.REF                       tggtttctcttcaggtctttctgtctcctcccacagggctgagaatggcagctcagggac    1808
LOC105372935.clincRNA.NCBI.ref      tggtttctcttcagggctttctgtctcctcccacagggctgagaatggcagctcagggac    1986
                                    *************** ********************************************

GGT5.NCBI.REF                       aagtcggggctggggactgcttagtctccccagtggctctcaggggatttgagggattga    1868
LOC105372935.clincRNA.NCBI.ref      aagtaggggctggggactgcttagtctccccagtggctctcaggggatttgagggtttga    2046
                                    **** ************************************************** ****

GGT5.NCBI.REF                       cgccagccgccaccccaggctgtgcccctcctctgctcaggaggacattcag-gatgcga    1927
LOC105372935.clincRNA.NCBI.ref      cgccagctgccaccccaggctgtgcccctcctctgctcgggaggacatacagagatgcga    2106
                                    ******* ****************************** ********* *** *******
```

```
GGT5.NCBI.REF                    cacccacttaaactcgaagttgcaaagatgcaaatgagactggagtctcaggcaccagag    1987
LOC105372935.clincRNA.NCBI.ref   cacccacttaaactcgaagttgcaaagatgcaaatgagactggggtctcaggcaccagag    2166
                                 ******************************************* ****************

GGT5.NCBI.REF                    accacccgtgggcatgtggcctttgggagtggggacctgctgccacaaatttctaggtgg    2047
LOC105372935.clincRNA.NCBI.ref   accacccgtgggcacgtggcttttgggagtggggacctgctgccacagatctct--gaag    2224
                                 ************** ***** ************************** ** ***  *  *

GGT5.NCBI.REF                    agtctggacctgctgggtctccctgagtgactgtctgggggtctccatagcgtgccctgc    2107
LOC105372935.clincRNA.NCBI.ref   agtctggacctgctgggtctccccgagtgactgtctgggggtctccatagcatgccctgc    2284
                                 *********************** *************************** ********

GGT5.NCBI.REF                    tgtgtgcgtgacggtcactggttgggtaggggtctctactctaaagctccctctgctggc    2167
LOC105372935.clincRNA.NCBI.ref   tgtgtgcgtgacggtcactggttgggtaggggtctctactctaaagctccctctgccggc    2344
                                 ******************************************************** ***

GGT5.NCBI.REF                    atcccctcaaactgtcccttgaag-agagaggatgtggtttgccccagtgttttgtcaaa    2226
LOC105372935.clincRNA.NCBI.ref   atcccctccaactctcccttggtgaagagaggatgtggtttgccccagtgttttatcaaa    2404
                                 ******** **** *******  * ***************************** *****

GGT5.NCBI.REF                    caactctctccacttcctgtcttaagaagctgggagtggaagagagcctggggctggccc    2286
LOC105372935.clincRNA.NCBI.ref   caactctctccacttccagttttaagaagctgggagtggaagagagcctggggctggccc    2464
                                 ***************** ** ***************************************

GGT5.NCBI.REF                    cagctgctgctgcggaacaggggtcattggacactgggaccctggccggactggctgggg    2346
LOC105372935.clincRNA.NCBI.ref   cagctgctgctgtgaaacaggggtcactggacgctgggaccctggccgggctggct-gga    2523
                                 ************ * *********** ***** **************** ****** **

GGT5.NCBI.REF                    ggcctcaggaagaggcctgctgcagcgtcatcctggccaagatccctccttgcagaggcc    2406
LOC105372935.clincRNA.NCBI.ref   ggcctcaggaagaggcctgctacagcgtcatcctggccaagattcctccctgcagaggac    2583
                                 ********************* ********************* ***** ******** *

GGT5.NCBI.REF                    cctggccacactgccacagggtctgctggggccaccagaagcccatgctcctgactccat    2466
LOC105372935.clincRNA.NCBI.ref   cctggccacgctgccacagggtctgctggggccaccagaagcccatgctcctgcctc---    2640
                                 ********* ******************************************* ***

GGT5.NCBI.REF                    catctctcccctctgtgctcacctctcaccaggaggccctcccagagtccagtctcctgc    2526
LOC105372935.clincRNA.NCBI.ref   catctctcccctctgtgctcacctctcaccaggaggccctcccagagttcagtgtcctgc    2700
                                 ************************************************ **** ******

GGT5.NCBI.REF                    tcttttttctgttttgtttttgagatgctgtttcgctctgtcaccaggctggagtgcagt    2586
LOC105372935.clincRNA.NCBI.ref   ttttttt----ttttttttttgtgacggtgtctcactctgtcaccaggctggagtgcagt    2756
                                 * *****    **** ****** ** * *** ** *************************

GGT5.NCBI.REF                    ggcatgatctcggctcgctgcaacctctgcctccttggttcaaatgattctcctgcctca    2646
LOC105372935.clincRNA.NCBI.ref   ggcgcgatctcagcttactgcaacctctgcttcctcggttcaaatgattctcctgcctca    2816
                                 ***  ****** ***  ************* **** ************************

GGT5.NCBI.REF                    gcctcctgagtagctgggactacaggtgctagccaccacgcccagctaattttgtattt    2706
LOC105372935.clincRNA.NCBI.ref   gcctcctgagtagctgggactacaggtgccagccaccacgccccgctaatttctgtattt    2876
                                 ***************************** ************* ******** *******

GGT5.NCBI.REF                    ttagtagagatggggtttcaccatgttagccaggatggtctccaactctagacctcgtgc    2766
LOC105372935.clincRNA.NCBI.ref   ttagtagagacggggtttcaccgtgttggccaggatggtctctatctcttga--------    2928
                                 ********** *********** **** ************** * **** **

GGT5.NCBI.REF                    tttgcccacctccgcctcccaaagtgctgggattacaggagtgagtcacggcacccagcc    2826
LOC105372935.clincRNA.NCBI.ref   ttcacccgccttggccacccaaagtgctggcattacaggagtgagtcatggcacctggcc    2988
                                 **  *** ***  *** ************* ***************** ******  ***

GGT5.NCBI.REF                    ccatctcctactctttcagcactaggttttattcttgggattctgctacagccggagccc    2886
LOC105372935.clincRNA.NCBI.ref   tcatctcctactctttcagcaccaggttttattcttgggattctgctacagccggagccc    3048
                                  ********************* *************************************

GGT5.NCBI.REF                    ctgggtgcaagctcctaagctttctgtgagtgtggacccagcaccgtgcctagtagacat    2946
LOC105372935.clincRNA.NCBI.ref   ctgggtgcgagttcctaaggtttctgtgagtgtggacccagcaccgtgcctagtagacat    3108
                                 ******** ** ******* ****************************************

GGT5.NCBI.REF                    acaaaaggagcatggtgacagtgaggtctgttatctccagcataatgactgttttgatcc    3006
LOC105372935.clincRNA.NCBI.ref   acaaaaggagcatggtgacagtgaggtctgtcatctccagcataatgactgttttgatcc    3168
                                 ******************************* ****************************

GGT5.NCBI.REF                    ttgtaaaaaaggtgatttttggctgggtgtggtggctcacacctgtgatcccagcacttt    3066
LOC105372935.clincRNA.NCBI.ref   ttgtaaaaaaggtgatttttggctgggtgtggtggctcacacctgtaatcccagcacttt    3228
                                 ********************************************** *************

GGT5.NCBI.REF                    gggaggctgaggcgggtggatcatttaaggtcaggagttggagaccagcctgggcaacat    3126
LOC105372935.clincRNA.NCBI.ref   gggaggccgaggggggtggctcacttgaggtcaggagttggagaccagcctgggcaacat    3288
                                 ******* **** ****** *** ** *********************************

GGT5.NCBI.REF                    ggtgaaaccacgtctctactaaaaatacaaaaattagctgggcatggtagcgggtgcctg    3186
LOC105372935.clincRNA.NCBI.ref   ggtgaaaccatgtctctactaaaaatacaaaaattagctgggcatggtagcaggtgcctg    3348
```

```
                                   ********** ***************************************** ********

GGT5.NCBI.REF                      taatcccagctacttgggaggctgagacaggagaatcacttgaacccaggaggcaaaggt    3246
LOC105372935.clincRNA.NCBI.ref     taatcccagatacttgggaggctgagacaggagaatcacttgaacccaggaggcaaaggt    3408
                                   ********* **************************************************

GGT5.NCBI.REF                      tgcagtaagccaagattgcaccactgcactccagcctgggtgacagagcaagacttggtc    3306
LOC105372935.clincRNA.NCBI.ref     ttcagtaagccaagattgcaccactgcactccagcctgggtgacagagcaagacttggtc    3468
                                   * **********************************************************

GGT5.NCBI.REF                      tcaggaaaaaaaaaaaaagaaagaaagaaaagtttatatttttgttctaatggttatctt    3366
LOC105372935.clincRNA.NCBI.ref     tca--aaaaaaaaaaaaagaaagaaagaaaagtttatatttttgttctaatggttatctt    3526
                                   ***  *******************************************************

GGT5.NCBI.REF                      aatatcttcattctataattatatgttttatataattataatagctatataagatataat    3426
LOC105372935.clincRNA.NCBI.ref     aatatcgtcattctataattatatgttttatataattataatagctatataagatataat    3586
                                   ****** *****************************************************

GGT5.NCBI.REF                      acccctagtatgttgtttttggatattctactcgttcctgatggttaatttatgtgtca    3486
LOC105372935.clincRNA.NCBI.ref     acccctagtatgttgtttttggatattctacttgctcctgatggttaatttatgtgtca    3646
                                   ******************************** * ************************

GGT5.NCBI.REF                      acttggctaagctctggtgccccgttgtttggtcaaatacttgtcaatatcttgctggga    3546
LOC105372935.clincRNA.NCBI.ref     acttggctaagctatggtgtcctgttgtttggtcaaatacttgtcaatatcttgctggga    3706
                                   ************* ***** ** *************************************

GGT5.NCBI.REF                      ggttatttcatagatgtgattaacactgacagtcagttgactttaggtaaaacagattac    3606
LOC105372935.clincRNA.NCBI.ref     ggttatttcatagatgtgattaacactgacagtcaattgactttaagtaaaacagattac    3766
                                   *********************************** ********* **************

GGT5.NCBI.REF                      ccaccataatatgggtgggccacctccaatcagttgaaggccgtaagaacaaaaactgag    3666
LOC105372935.clincRNA.NCBI.ref     ccaccataatatgggtgggccacctccaatcagttgaaggccttaagaacaaaaactgag    3826
                                   ****************************************** *****************

GGT5.NCBI.REF                      gtttcccagagaagcaggaattctgcttcaacactataacacacaaaccctgcctgagtt    3726
LOC105372935.clincRNA.NCBI.ref     gtttcccagagaagcaggaattctgcttcaagactgtaacacacaaaccctgcctgagtt    3886
                                   ******************************* *** ************************

GGT5.NCBI.REF                      tctggcctgctgactgctctacagatgttaggttccagacttcgagatcaactcttacct    3786
LOC105372935.clincRNA.NCBI.ref     tctggcctgctgactgctctacagatttttaggttccagacttcgagatcaactcttacct    3946
                                   ************************** *********************************

GGT5.NCBI.REF                      gaatttatagcctgctggcttgccctacagattttt-aaacttgccagtccccaaaatcat    3845
LOC105372935.clincRNA.NCBI.ref     gaatttatagcctgctggcttgccctacagattttaaaacttgctagtccccacaatcat    4006
                                   *********************************** ******** ******** ******

GGT5.NCBI.REF                      gtgagccaattcctaaataaa----tctctatgtataacctattggtttagtttctctaa    3901
LOC105372935.clincRNA.NCBI.ref     gtgagccaattcctaaataaatctctctctatgtataatctattggtttagtttctctga    4066
                                   *********************    ************* ******************* *

GGT5.NCBI.REF                      aaaaacttctatatccagtttcctggatgttaagtaatactgaaactagctagtaacctc    3961
LOC105372935.clincRNA.NCBI.ref     aaaactttc-acatccagtttcctggttgttaagaattaccgaaactagctagtaacttc    4125
                                   ****  *** * ************** ******* * *** **************** **

GGT5.NCBI.REF                      gtgttttttttttttttttttttttgagatggagttttgttcttgttgcacaggctggagt    4021
LOC105372935.clincRNA.NCBI.ref     -----ttttttttttttttttttttgagacagagttttgctcttgttgcccaggctggaat    4180
                                        ***********************  ******** ********* ********* *

GGT5.NCBI.REF                      acaatagcacgatcttggctcaccgcaacctccacctcctgggttcaagcgattctcctg    4081
LOC105372935.clincRNA.NCBI.ref     gcaatggcacaatctcagctcaccgcaacctccacttcctgggtccaagcaattctcctc    4240
                                    **** **** ****  ****************** ******** ***** ********

GGT5.NCBI.REF                      cctcagcctcctgagtagctgggattacaggcatgcgccaccacacctggctaa-ttttg    4140
LOC105372935.clincRNA.NCBI.ref     cctcagcctcctgagtagctgggattacaggcatgtgccaccatgcttggctaatttttg    4300
                                   *********************************** *******  * ******* *****

GGT5.NCBI.REF                      tatttttggtagagacagggtttctccatgtgggtcaggctggtctcaaactcccgacct    4200
LOC105372935.clincRNA.NCBI.ref     tatttttagtagagacagggcttctccatgttggtcaggctggtcttgaactcccaacct    4360
                                   ******* ************ ********** **************  ******* ****

GGT5.NCBI.REF                      caggtgatctgcccgctttggcctcccaaagtgctgggattacaggcatgaactactgca    4260
LOC105372935.clincRNA.NCBI.ref     caggtgatccg-ccgccttggcctcacaaagtgctggaattacaggcacgagccattgcg    4419
                                   ********* * **** ******** *********** ********** ** * * ***

GGT5.NCBI.REF                      cccgtctcctagtaatttcttcttttccatgatgtgtctcttatctctaataatactttt    4320
LOC105372935.clincRNA.NCBI.ref     cctggctcctagtaaattcttcttttctgtgatgtgtctcttacctctaataatactttt    4479
                                   ** * ********** ***********  ************** ****************

GGT5.NCBI.REF                      cttcttaaagtctacttcattaaaaatagttatgctgggcatggtggctcatgcctgtaa    4380
LOC105372935.clincRNA.NCBI.ref     cttcttaaagtctacttcattaaaaatagttatgctgggcatggtggctcatggctgtaa    4539
                                   ***************************************************** ******

GGT5.NCBI.REF                      --ttggcactttg--ggaggtcaaggtgagtgggtcgctgaagcccaggagttcaagacc    4436
```

```
LOC105372935.clincRNA.NCBI.ref      tcttggcactttgctggaggtcgaggtgggtggatcactgaagcccaggagttcaagacc    4599
                                      ***********  ******* ***** **** ** ***********************

GGT5.NCBI.REF                       agcctgggcaacatggtgag-----------acaaaaagtacaaaaattagctgggtgtg    4485
LOC105372935.clincRNA.NCBI.ref      aacctgggcaacatggcgagaccctgcctctacaaaaaatacaaaaattagctgggtgtg    4659
                                    * ************** ***           ******* *********************

GGT5.NCBI.REF                       gctaatataattctaagttggcacacttgtagtcccagctacttgggaggctgagcgggg    4545
LOC105372935.clincRNA.NCBI.ref      gctaatataattctaagttggcacacttgtagtcccagctacttgggatgctgaggtggg    4719
                                    ************************************************ ******  ***

GGT5.NCBI.REF                       agaatcgcatgatgctagaagggagagattgctgtgagccaagatcacgtcgctgcactc    4605
LOC105372935.clincRNA.NCBI.ref      agaatcgcttgagcctagaagggagagattgctgtaagccaagatcacatcactgcactc    4779
                                    ******** ***  ******************** ************ ** ********

GGT5.NCBI.REF                       cggcctgggagacagagtgaggctctatctc-------------aaaaaaaaaaaaaaag    4652
LOC105372935.clincRNA.NCBI.ref      cagcctgggagacagagtgaggctctatctccaaaaaaaaaaaaaaaaaaaaaaaaaaag    4839
                                    * *****************************             ****************

GGT5.NCBI.REF                       ttatacagctttcttggttagtgcatgcatgatatatttttcattattttccacctttct    4712
LOC105372935.clincRNA.NCBI.ref      ttatacagctttcttggttagtacatgcatgacatatttttcatgatcttccacctctct    4899
                                    ********************** ********* *********** ** ******** ***

GGT5.NCBI.REF                       gtatccttacataaaaggca----ttgggttttactttatttccaattactttaattttt    4768
LOC105372935.clincRNA.NCBI.ref      gtacccttatataaaaggcattagttgggttttactttatttcaattatttaattttt    4959
                                    *** ***** **********    ****************** ****** **********

GGT5.NCBI.REF                       attgattgtccttttaaaagtaggtgatgatttatttgggttgaaacccaccaccaactt    4828
LOC105372935.clincRNA.NCBI.ref      ----attgtccttttaaatgtaactaatgatttatttgggttgaaacccaccaccaattt    5015
                                        ************** ***  * ******************************* **

GGT5.NCBI.REF                       gttttccatgcctattctgtttcttcttatctcctttcacatcttgttttgcatttatta    4888
LOC105372935.clincRNA.NCBI.ref      gttttccatgcctattctatttcttcttatctcctctcacatcttgttttggatttatta    5075
                                    ****************** **************** *************** ********

GGT5.NCBI.REF                       tttttattattcaatttcctttttctctattagtttcctaactgtgcagtcttggggtta    4948
LOC105372935.clincRNA.NCBI.ref      tttttattatttaatttcctccttctctattagtttcataactgtgcagtcttagagtta    5135
                                    *********** ********  *************** *************** * ****

GGT5.NCBI.REF                       ttttaaaagatgacagtagattattttagagcttacaacatgcatccttcacttatcaaa    5008
LOC105372935.clincRNA.NCBI.ref      ttttaaaagatgac--tggattattttagagcttacaacatgcatccttcccttatcaaa    5193
                                    **************  * ********************************* *********

GGT5.NCBI.REF                       gtctaacatgagctagtactttttttgtttatttgtttttcttgagatagagggagtc    5068
LOC105372935.clincRNA.NCBI.ref      gtctaacatgagctagtactttttgttg------------tggttgagatagagagagtc    5241
                                    ************************ ***            *  *********** *****

GGT5.NCBI.REF                       ttgctctgctgcccaggctggagtgcagtggagcaatcttggttcactgcaacctccacc    5128
LOC105372935.clincRNA.NCBI.ref      ttcctctgctgcccaggctggagtgcagtggagcaatcttggttcactgcaacctccact    5301
                                    ** ********************************************************

GGT5.NCBI.REF                       tcttgggttcaagcaattctcctgtctcagtctcctgagtagctgggaccactggtgtgc    5188
LOC105372935.clincRNA.NCBI.ref      tcttgggttcaagcaattctcctgcctcagtcacctgagtagctgggaccacaggtgtgc    5361
                                    ************************ ******* ******************* *******

GGT5.NCBI.REF                       accactatgcccggctaatttttgtattctttttagtagagacagggtttcaccatgtt    5248
LOC105372935.clincRNA.NCBI.ref      accactatgcccggccaatttttgtattctttttcagtagagacagggtttcaccatgtt    5421
                                    *************** ****************** ************************

GGT5.NCBI.REF                       ggccaggctggtcttgaactcctcaccttaagagatctgcttacctcggcgtcctaaagt    5308
LOC105372935.clincRNA.NCBI.ref      ggccaggctggtcttgaactccggaccttaagagatctgcctacctcggcatcctaaagt    5481
                                    **********************  **************** ********* *********

GGT5.NCBI.REF                       gttgggattacaggcgtgagccaccacgcccagcctatgagttagtacttctatcctctt    5368
LOC105372935.clincRNA.NCBI.ref      gttgggattacaggcgtgagccaccgcgcccagcctatgagttagtacttctatcctctt    5541
                                    ************************* **********************************

GGT5.NCBI.REF                       cctagtcagtagaagaaccttggaacaggaactaaatttacccccagtgacttatatgct    5428
LOC105372935.clincRNA.NCBI.ref      cctagtcagtacaagaaccttggaacaggaactaaatttacccccagtgacttatatgct    5601
                                    *********** ************************************************

GGT5.NCBI.REF                       aatatttttgtgtattttaaatatatgtgtgtgcatagatgtatctgtgtgttttttgtg    5488
LOC105372935.clincRNA.NCBI.ref      aatatttttgtgtattttaaatatgtgtgtgtgcatagatgtatctgtgtgttttttgtg    5661
                                    ************************ ***********************************

GGT5.NCBI.REF                       tttttattcttatttatgttgagagtgtagagctatgtaagagtaaagagaattgtgtaa    5548
LOC105372935.clincRNA.NCBI.ref      tttttattcttatttatgttgagagtgtagagctatgtaagagtaaagagaattgtgtaa    5721
                                    ************************************************************

GGT5.NCBI.REF                       tgaagccccgagtatccattcaatttcaacaacaatcttatggccaagctcatttcatgt    5608
LOC105372935.clincRNA.NCBI.ref      tgaagccccgagtatccattcaatttcaacaacaatcttatggccaagctcatttcatgt    5781
                                    ************************************************************
```

```
GGT5.NCBI.REF                     atactctttcctgcttccctctaccctacattatttcagtgcaaatcccagatatataac    5668
LOC105372935.clincRNA.NCBI.ref    atactctttcctgcttccctctaccccacattatttcagtgcaaatcccagatatataac    5841
                                  ************************** *********************************

GGT5.NCBI.REF                     tttacccatacatatttcagtatgt----tttattttaagccccacaagatatcattttc    5724
LOC105372935.clincRNA.NCBI.ref    tgtaccaatacatatttcagtatgttttatttattttaaaccccacaagatatcattttc    5901
                                  * **** ******************    ********** ********************

GGT5.NCBI.REF                     tatactactataattttataccaataacgttcatttagatttacccacacgtttacctct    5784
LOC105372935.clincRNA.NCBI.ref    tatactactgtaattttataccaataacattcatttagatttacccacacgtttacc---    5958
                                  ********* ****************** ****************************

GGT5.NCBI.REF                     tctgttaccctttttttttttgagacagagtctcgctctgtcgcccaggctggagtgcagt    5844
LOC105372935.clincRNA.NCBI.ref    ----ctaccctccgggtcctgtttgaaaatcaagcccatgctcacaggcca---------    6005
                                       ******     *  **    * * **  ** *   * * *****

GGT5.NCBI.REF                     ggcacaatctcggctcactgcaagctctgcctcctgggttcacgccattctcctgcttca    5904
LOC105372935.clincRNA.NCBI.ref    ------------------------------------------------------------    6005

GGT5.NCBI.REF                     gcctcccgagtagctgggactacagtcacccgccaccacgcctggctaatttttttgtatt    5964
LOC105372935.clincRNA.NCBI.ref    ------------------------------------------------------------    6005

GGT5.NCBI.REF                     ttttagtagagacgggtttcactgtgttagccaggatggtctcgatctcctaacctcatg    6024
LOC105372935.clincRNA.NCBI.ref    ------------------------------------------------------------    6005

GGT5.NCBI.REF                     atacacccgcctcggcctcccaaagtgctgggattacaggcgtgagccaccacgcctggc    6084
LOC105372935.clincRNA.NCBI.ref    ------------------------------------------------------------    6005

GGT5.NCBI.REF                     caccctttatttttatttataaaaatatctttgggaaaaaatatctttgggcacatggtc    6144
LOC105372935.clincRNA.NCBI.ref    ------------------------------------------------------------    6005

GGT5.NCBI.REF                     aaggatctcctgaggactgtgtcatgggctatatatatattatatatatatatatatatt    6204
LOC105372935.clincRNA.NCBI.ref    ------------------------------------------------------------    6005

GGT5.NCBI.REF                     atataatatatattatatatataatatatataatatatatatatataaaatatataatat    6264
LOC105372935.clincRNA.NCBI.ref    ------------------------------------------------------------    6005

GGT5.NCBI.REF                     atattatatatattatatatataatatatataatatatatatataaatatatatatatat    6324
LOC105372935.clincRNA.NCBI.ref    ------------------------------------------------------------    6005

GGT5.NCBI.REF                     atataatatattatatatatataaatatataatatatattatatatatatatatatattc    6384
LOC105372935.clincRNA.NCBI.ref    ------------------------------------------------------------    6005

GGT5.NCBI.REF                     catatatatacacacacatatatatatattccactttcactttttttttttttttgagac    6444
LOC105372935.clincRNA.NCBI.ref    ------------------------------------------------------------    6005

GGT5.NCBI.REF                     ggtgtcttgctctgtcgcccaggctggagtatagtggcgtgatctcagctcactgcaact    6504
LOC105372935.clincRNA.NCBI.ref    ------------------------------------------------------------    6005

GGT5.NCBI.REF                     tctgcctcccgggttcaagtgattctcctgtctcagccacctgagtagctgggattccag    6564
LOC105372935.clincRNA.NCBI.ref    ------------------------------------------------------------    6005

GGT5.NCBI.REF                     gcatgagccaccacgcctggctaatttttgtatttttttttttttgagatggggtctcact    6624
LOC105372935.clincRNA.NCBI.ref    -----------------------------attttttttcttttagagacagggtctcact    6036
                                                                * ****** **** ****  **********

GGT5.NCBI.REF                     gtgtcgcccaggctgaagtgcaatggcacgatatctgctcactgcaacctccacctccct    6684
LOC105372935.clincRNA.NCBI.ref    ttgtcacccaagctggagtgcagtggtgcgattatagctcaatgcagcctccaattcctg    6096
                                   **** **** **** ****** ***  ****    ***** **** ******  ***

GGT5.NCBI.REF                     ggttcaagcagttcccctgcctcagtctcccaagtagctgggattacaacaggcgcatga    6744
LOC105372935.clincRNA.NCBI.ref    gactcaagggaccctcctgcctcagcctgccaagtagcttggactatagctg--------    6148
                                  *  *****     * ********** ** ********** *** ** * * *

GGT5.NCBI.REF                     caccatgccctgctaatttttttgtatttgtagtagagacagagtttcaccatattcgcc    6804
LOC105372935.clincRNA.NCBI.ref    -------------tgtgttttcttattattttgtagacatggggtctggctatgttgtcc    6195
                                               *   **** *   * * * ***** *  * ** *  * ** **  **

GGT5.NCBI.REF                     agactggtctcaaac-tcctgacctcaagcaatttgcctgccttggcctcccaaagtgct    6863
LOC105372935.clincRNA.NCBI.ref    aggctattctcaaaattcccggcctcgagcaatcctcctgcctcggcctctcaaag-gtt    6254
                                  ** **  *******  *** * **** ******   ******* ****** ***** * *
```

```
GGT5.NCBI.REF                       gagattacaggcgtgagtcaccgcgcctggc-----------------------------   6894
LOC105372935.clincRNA.NCBI.ref      gggattacaggtgtgaggcaaggcacccagctcagccacagagccctgttgcatctctct   6314
                                    * ********* ***** **  ** **  **

GGT5.NCBI.REF                       ------------------------------------------------------------   6894
LOC105372935.clincRNA.NCBI.ref      tactaggagcaagagctgactgccccctcatccccattccagagtgttggggctgtgttc   6374

GGT5.NCBI.REF                       ------------------------------------------------------------   6894
LOC105372935.clincRNA.NCBI.ref      agccgaggccgggccactggcatggcccagggagcgggatcattcactgctgccccaaat   6434

GGT5.NCBI.REF                       ------------------------------------------------------------   6894
LOC105372935.clincRNA.NCBI.ref      ctgagatcattccaccttgacaagacttcctcatccaatccctttacttgacagctgggg   6494

GGT5.NCBI.REF                       ------------------------------------------------------------   6894
LOC105372935.clincRNA.NCBI.ref      aaaccaatgcgcacagagcacccccagctcactcggggtctcagagctgatccatgagca   6554

GGT5.NCBI.REF                       ------------------------------------------------------------   6894
LOC105372935.clincRNA.NCBI.ref      gaggctgagatcctgggatcttgtcccccagccgccctgcaagcttactccctttctgct   6614

GGT5.NCBI.REF                       ------------------------------------------------------------   6894
LOC105372935.clincRNA.NCBI.ref      ggaagagatggggccggacctcgaccagcagccctggcctggacatgactgtgctcaccc   6674

GGT5.NCBI.REF                       -----------------------------------------cttttttttttttttttt   6913
LOC105372935.clincRNA.NCBI.ref      aggtattgaggccgagatgccccggcatcatatgtttttctctctttttttctttttttt   6734
                                                                             ** ******* ********

GGT5.NCBI.REF                       tgagacggagcctcactctgtcccccaggctggggtgtggtgggaagatctcggcttcct   6973
LOC105372935.clincRNA.NCBI.ref      tgagacagtatctcactctgtcacccaagctggagtgcagtggcatgatattggctcact   6794
                                    ****** *   *********** **** ***** ***  **** * *** * ****  **

GGT5.NCBI.REF                       gcaacctccacctgctgggttcaaatgtttcttgtgcctcagcctcccaagtagatggga   7033
LOC105372935.clincRNA.NCBI.ref      gcaacctctgcctcc-cgcttaaagtgattctcctgcctcagtcttccaagtagctgggc   6853
                                    ********  *** *  * ** ** ** ****  ******** ** ******** ****

GGT5.NCBI.REF                       ttacaggtgcccacaaccacacctggctaatttttgtatttttggtagagatggggtttt   7093
LOC105372935.clincRNA.NCBI.ref      ctacaggcttgtaccaccacacctgactaatttttgtatttttactagagacgggg-ttt   6912
                                     ******     ** ********** *****************  ****** **** ***

GGT5.NCBI.REF                       aaccacgttggccaggctggtctcgaactcctgacctcaggtgacctacccaccttggcc   7153
LOC105372935.clincRNA.NCBI.ref      ccccatgttggccaggctcgtgtcgaactcctgacctcaggtgatccacctgccttggcc   6972
                                      *** ************ ** ********************** * ***  ********

GGT5.NCBI.REF                       ttccaaagtgatgggattacaggcgtgagccaccactcccagcctccatttttaatagat   7213
LOC105372935.clincRNA.NCBI.ref      tcccaaagtgctaggattacaggcatgagccatggcgtcac-------------------   7013
                                    * ******** * *********** *******   *  *

GGT5.NCBI.REF                       gacagttctttgataaaattcttatttctttatatccttgaatatagatacaaagtactt   7273
LOC105372935.clincRNA.NCBI.ref      ------------------------------------------------------------   7013

GGT5.NCBI.REF                       attttaaagtacatggtctgccaatttcataatctggagatcctatggacctttttaaaa   7333
LOC105372935.clincRNA.NCBI.ref      ------------------------------------------------------------   7013

GGT5.NCBI.REF                       gttgtctgtgctttctcatgagctttgttcctgctgtcttatttccttgtttgcttggtt   7393
LOC105372935.clincRNA.NCBI.ref      ------------------------------------------------------------   7013

GGT5.NCBI.REF                       gtttttaatttggcactggagagtatgtataaaaattgttagaggccgggtgcggtggct   7453
LOC105372935.clincRNA.NCBI.ref      ----------------------------ttaaatgtagtgagaggccgggcaagg-ggct   7044
                                                                 ****  * ** **********   ** ****

GGT5.NCBI.REF                       cacgcttgtaatcccagcactttgggaggccaaggcaggcagatcac--gaggtcaggag   7511
LOC105372935.clincRNA.NCBI.ref      catgcctgtaatcccagtgctttgagaggacgaggctgtcagatcacctaaggtcaggag   7104
                                    ** ** ***********  ***** **** * **** * ********   **********

GGT5.NCBI.REF                       tacaagaccagcctggccaacacagtgaaaccctgtctctactgaaaa--tacaaaaatt   7569
LOC105372935.clincRNA.NCBI.ref      ttcgagaccagcctggccaacatggtgaaactgtgtctctacaaaaaaatagaaaaaaaa   7164
                                    * * ******************  *******  *********  ****     *****

GGT5.NCBI.REF                       agctctgcgtgatggctggtggctgtaatcccagctacttgggaggctgagacaggagaa   7629
LOC105372935.clincRNA.NCBI.ref      atccctgcgtggtggcaagtatctgtagtcccagttactcaggaggctgaggcatgagaa   7224
                                    * * ******* ****  **  ***** ****** ****  ********** ** *****

GGT5.NCBI.REF                       ttgcttgaacccaggaggtgaaggttgcagtgagccgagatcgaggcactgcactccagc   7689
LOC105372935.clincRNA.NCBI.ref      ttgcttaaacctcggaggcggaggctgcagtgagctgagatggcgccactgcactccagc   7284
```

```
                                    ****** ****  ***** * *** ********** ***** * * **************

GGT5.NCBI.REF                       ctggacaacagagtgagatgctgtctc-aaataaataaataaataaatataaaataattt    7748
LOC105372935.clincRNA.NCBI.ref      ctgggtgacagagcaagactttgtctctaaataattaaataaataaatatggccgagcat    7344
                                    ****   ******  ***   ****** ****** ***************     *   *

GGT5.NCBI.REF                       gaggcctagggatctgaaattctgggatctcctttatgcatttgagtgactgagatgatt    7808
LOC105372935.clincRNA.NCBI.ref      ggtgccttaggcctgtaat-----------------------------------------    7363
                                    *  ****  **     **

GGT5.NCBI.REF                       tgaagctggatccagtgctcctgagggctgctctatttctagttgactcctagagtaaga    7868
LOC105372935.clincRNA.NCBI.ref      ------------------------------------------------------------    7363


GGT5.NCBI.REF                       aacctgcaccccacgtgtggggcattatggcatcccctccctcagcacatgagtaggtca    7928
LOC105372935.clincRNA.NCBI.ref      -------cccaacactttgggaggctgaggcaggtggttcatgaggtcaggagcccgaga    7416
                                            **     * ****    *  ****     * * * **  ** ***   *  *

GGT5.NCBI.REF                       acagc-------------------------------------actgctctagaccaggtg    7951
LOC105372935.clincRNA.NCBI.ref      ccagcctggccaagatggtgaaacactgtctctactaaaaatacaaaaattagccagctg    7476
                                     ****                                     **     *   **** **

GGT5.NCBI.REF                       tggtggctcacgcctatagtcccagctactcaggagactgaggcaggaggattgcttcag    8011
LOC105372935.clincRNA.NCBI.ref      tggtggcaggcacctgtaatcccagctacttgggacactgaggcaggagaatcgcttga-    7535
                                    *******   * *** ** ***********  *** ************* ** **** *

GGT5.NCBI.REF                       gccaagaatttgagaccagccagagcaatatattaggttggtacaaaagtaattgcggtt    8071
LOC105372935.clincRNA.NCBI.ref      ------acctggaaggcagaggttgcaatgagccgagattgcaccgctgcac-------t    7582
                                          *  * **   ***     *****       * * * **    * *        *

GGT5.NCBI.REF                       tttgccattaaaagtaatggtgaccctgtctcagaaaaataaa--aagcaaaaaaaaaaa    8129
LOC105372935.clincRNA.NCBI.ref      ctagcctgggcgatggagcaagactccatctcaaataaataaattaataaatacagagca    7642
                                     * ***      *   *    *** *  ***** * *******  **  ** * * *  *

GGT5.NCBI.REF                       aaaaaagagagaaaagaaaaaaagaatccctgatctatttctcagcctcctcttcaggat    8189
LOC105372935.clincRNA.NCBI.ref      agattccatctcaaataaataaataaatg-------------------------------    7671
                                    * *    *    *** *** *** **

GGT5.NCBI.REF                       tggccgtcaccttgaggggaatgctggccttgcctgtctccagccctgtacttctctgcc    8249
LOC105372935.clincRNA.NCBI.ref      ------------------------------------------------------------    7671


GGT5.NCBI.REF                       tcctatgcctttaagcacatgttttctatttgctgggcttttctagttctctgagggaag    8309
LOC105372935.clincRNA.NCBI.ref      ------------------------------------------------------------    7671


GGT5.NCBI.REF                       tgtggtctgaattacagtctcccattaccaggaagtctgatttacgtttaacagattttt    8369
LOC105372935.clincRNA.NCBI.ref      ------------------------------------------------------------    7671


GGT5.NCBI.REF                       ttttttttttgagatggagtctctctccgttgcccaggctggagtgcagtggtacaacc    8429
LOC105372935.clincRNA.NCBI.ref      ------------------------------------------------------------    7671


GGT5.NCBI.REF                       ttggctcactgcaagctccgacttccaggttcacaccattctcctgcctcagcctcctga    8489
LOC105372935.clincRNA.NCBI.ref      ------------------------------------------------------------    7671


GGT5.NCBI.REF                       gtagctgggactacaggcacccgccaccatgcccagctaatttttttgtatttttaataga    8549
LOC105372935.clincRNA.NCBI.ref      ------------------------------------------------------------    7671


GGT5.NCBI.REF                       gatggggtttcaccgtgttagccaggatggtctcaatctcctgacctcatgatctgcctg    8609
LOC105372935.clincRNA.NCBI.ref      ------------------------------------------------------------    7671


GGT5.NCBI.REF                       cctgggcctcccaaagtgctgggattacaggcatgagccaccgcgcccggccttaacaga    8669
LOC105372935.clincRNA.NCBI.ref      ------------------------------------------------------------    7671


GGT5.NCBI.REF                       ttatttttttaaaatgtaagggagcaaggccaacgctggggcctgggtaggaggccattg    8729
LOC105372935.clincRNA.NCBI.ref      ------------------------------------------------------------    7671


GGT5.NCBI.REF                       caataacccagctgagagaggttggtggcttggaccagtgtggtggagatggaggtagaa    8789
LOC105372935.clincRNA.NCBI.ref      ------------------------------------------------------------    7671


GGT5.NCBI.REF                       cctgcatccagagtcactggtgggttggatgtggggaaggaagaaagtgaccaatccagg    8849
LOC105372935.clincRNA.NCBI.ref      ------------------------------------------------------------    7671


GGT5.NCBI.REF                       atggctccagggtgtaggcgctgagcaaaaggggagtggttgtgtcaatttctaagaca    8909
```

```
LOC105372935.clincRNA.NCBI.ref      ------------------------------------------------------------    7671

GGT5.NCBI.REF                       acgaggactaggagaggagccatttcgcagggacggttggaaggaaaatcaaaagtacag    8969
LOC105372935.clincRNA.NCBI.ref      ------------------------------------------------------------    7671

GGT5.NCBI.REF                       tttaggtcatggtatgtttgaaatgtttcttagacacccagagtgggtaccttgtgcata    9029
LOC105372935.clincRNA.NCBI.ref      ------------------------------------------------------------    7671

GGT5.NCBI.REF                       gctgggtataagtgtctcgagtgtagggagaggactggcggagttaaagtcggggagtta    9089
LOC105372935.clincRNA.NCBI.ref      ------------------------------------------------------------    7671

GGT5.NCBI.REF                       ccatcacacaggttgcatctcatgccatgagcctggatggactcacccagggagaagata    9149
LOC105372935.clincRNA.NCBI.ref      ------------------------------------------------------------    7671

GGT5.NCBI.REF                       tagatgcacaaggaccatggaccaagcctggggccccaacatcaggagaccagggaggt    9209
LOC105372935.clincRNA.NCBI.ref      ------------------------------------------------------------    7671

GGT5.NCBI.REF                       gggacgaaaaccaaagcctggcagggtgggcgaggtgggtgcagggacctggcccattcc    9269
LOC105372935.clincRNA.NCBI.ref      ------------------------------------------------------------    7671

GGT5.NCBI.REF                       ctcttggggctttggtttccccatctgtaaccatttaccccattttgccctggaagccaa    9329
LOC105372935.clincRNA.NCBI.ref      ------------------------------------------------------------    7671

GGT5.NCBI.REF                       gacaactgactaggctggaagcagacaggttggggaagcaagggtgcactccatttagtt    9389
LOC105372935.clincRNA.NCBI.ref      ------------------------------------------------------------    7671

GGT5.NCBI.REF                       aggaagtgaggcctttgtctgggatgggagtttaaagtgttttttcccccagtttgtcat    9449
LOC105372935.clincRNA.NCBI.ref      ------------------------------------------------------------    7671

GGT5.NCBI.REF                       ttggctttgtttatggtgttttattcttataaaaattagtgtttatgcctgacttgttta    9509
LOC105372935.clincRNA.NCBI.ref      ------------------------------------------------------------    7671

GGT5.NCBI.REF                       tttcattttctggctttgtgctctctttcctgttccaagattacatgtgtccctgtggtg    9569
LOC105372935.clincRNA.NCBI.ref      ------------------------------------------------------------    7671

GGT5.NCBI.REF                       cccatgtgtgccacgtgtgggcccagagagggacatgtggctcaggagctggggtgggtt    9629
LOC105372935.clincRNA.NCBI.ref      ------------------------------------------------------------    7671

GGT5.NCBI.REF                       tgtgtccagatggacatgctaggtcaggacagacacctgagccaaaggagggtgctaggg    9689
LOC105372935.clincRNA.NCBI.ref      ------------------------------------------------------------    7671

GGT5.NCBI.REF                       ccagaggtcccacggggtgggtgcagggacccaggtccattccctctttgggccttggtt    9749
LOC105372935.clincRNA.NCBI.ref      ------------------------------------------------------------    7671

GGT5.NCBI.REF                       tccccatctgcaactggggcttaaatgctggcttctggccgggcacagtggctcacgcct    9809
LOC105372935.clincRNA.NCBI.ref      -----------------------------------------------------tacacct    7678
                                                                                          ** ***

GGT5.NCBI.REF                       gtaatcccagcactttgggaggccgaggtgagcagatcatctgatgtcgggagtttgaga    9869
LOC105372935.clincRNA.NCBI.ref      gtaatcctagcactttgggaggctaagacaggtcgatcacctgaggtcaggagttcgaga    7738
                                    ******* ***************  **    *  ***** **** *** ****** ****

GGT5.NCBI.REF                       ccagcctgaccaacatggagaaacctcgtctctactaaaaatac-aaaattagtcaggcg    9928
LOC105372935.clincRNA.NCBI.ref      ccagcctgaccaatatggcaaaactccatctctactaaaaatacaaaaattagccgggcg    7798
                                    ************* ****  ****  * **************** ******** * ****

GGT5.NCBI.REF                       tggtggtgcatgcctgtaatcccagctactcggcaggctgaagcaggagaatcgcttgaa    9988
LOC105372935.clincRNA.NCBI.ref      ttttgacgtgtgcctgtagtcccagctacttgggaggctgagacaagagaattgcttgaa    7858
                                    *  **  *  ******** *********** ** *******  ** ****** *******

GGT5.NCBI.REF                       tccgggagg-cgaggttgcaatgagccaagatagtgtcattgcactccagcctgggcaag    10047
LOC105372935.clincRNA.NCBI.ref      cccaagaggtggaggttgcagtgagccgagatctcggc--tgcacttcagcctgggtga-    7915
                                     **  ****  ********* ****** ****   * *  ****** *********  * 

GGT5.NCBI.REF                       aagattgaaactctgtctcaaaaaaaaaaaaaaaaaaaaaaaaaagctggcttcttgagg    10107
LOC105372935.clincRNA.NCBI.ref      cagagtgagactctgtctcaaaaggaataaataaaatacaaagtaa--------------    7961
                                     *** *** **************  ** *** **** * ***  *
```

```
GGT5.NCBI.REF                       tgctggggaggaaggattcctgggtgttagggaagaggttgacctcagtgagctgtgcag    10167
LOC105372935.clincRNA.NCBI.ref      ------------------------------------------------------------    7961

GGT5.NCBI.REF                       gcctgtgccccagggacagggatcatcataatgtaggccaggctgaccagccctcagtgc    10227
LOC105372935.clincRNA.NCBI.ref      -----------------------------aaaaaaatgtagtaagattgcagagtcgtgc    7992
                                                                 **   *    **   **          ****

GGT5.NCBI.REF                       taaagggcccacctcttgctgcccatggacagatggggaggggtaggccagagcattt--    10285
LOC105372935.clincRNA.NCBI.ref      cgcagaagcgtgctggtcctatccatgtagtgaaggctgatttcatacacaaatgtcaga    8052
                                       **   *   **  * **  ***** *  ** **        *  *   *   *

GGT5.NCBI.REF                       --attttctttctttcttttcttttcttttttttttcttttttgaggcagtcttgctttgt    10343
LOC105372935.clincRNA.NCBI.ref      agaacttttcttttctttttcttttcttttttttttttttgagacggagtctcgctctgt    8112
                                      *  ** * * **  ******************* ****  ** * ***** *** ***

GGT5.NCBI.REF                       tgcccaggctggagtgcagtggcatgatttcagctcactgcaacctctgcctgctgggtt    10403
LOC105372935.clincRNA.NCBI.ref      cacccaggctggagtgcagtggtgtgatctaggctcactgcaagctctgcctccctggtt    8172
                                      ********************  **** *  *********** ******** *  ****

GGT5.NCBI.REF                       caagcaattctcctgcctcagcctcctgcgtagctgggattacaggtgcccgccatcaca    10463
LOC105372935.clincRNA.NCBI.ref      tatgccattctcctgcctcagcctcctcagtagctgggactacaggcgcctgccacctag    8232
                                     * ** *********************  ********** ****** *** **** *

GGT5.NCBI.REF                       ttcagctaa--tttttgtatttttagtagagacggggtttcaccatgttggccaagctgg    10521
LOC105372935.clincRNA.NCBI.ref      cccagctaatttttttgtacttttagaggagatggggtttcaccgcgttagccaggatag    8292
                                      *******  ******** ******  **** ***********  *** **** * * *

GGT5.NCBI.REF                       tcttgaactcctgacctcaagtgatctatccgcctcggcctcccaaagtgcagggattat    10581
LOC105372935.clincRNA.NCBI.ref      tctcaatctcctgacctc--gtgatccgcccgtctcggcctcccaaagtgctgggattac    8350
                                    ***  * ***********  ******   *** ****************** *******

GGT5.NCBI.REF                       aggcatgagccaccacacccagcctaggccagagaatttccataggctcagagtcccaag    10641
LOC105372935.clincRNA.NCBI.ref      aggcatgagccaccacacccggcctt----------------------------------    8376
                                    ******************** ****

GGT5.NCBI.REF                       agaagttttggggtccatcagctcaagccctatgccctgctggcagggaaggatcctggg    10701
LOC105372935.clincRNA.NCBI.ref      ------------------------------------------------------------    8376

GGT5.NCBI.REF                       atcacctgccaaaggccagccttgaagtagttgtggccagagtgtccccagtgagatggg    10761
LOC105372935.clincRNA.NCBI.ref      ------------------------------------------------------------    8376

GGT5.NCBI.REF                       gagtggagtcccagggtgggctcctgaggccactgtgagggactgagaggtggccgaggc    10821
LOC105372935.clincRNA.NCBI.ref      ------------------------------------------------------------    8376

GGT5.NCBI.REF                       ctcgttctgcctgtagcatatgagctggtgctgctgggaacaatggcaacagaagtactt    10881
LOC105372935.clincRNA.NCBI.ref      ------------------------------------------------------------    8376

GGT5.NCBI.REF                       tgcaaactgagaagcaccttgtatatggcaaaggtcagtgtaaatggcatcacactttgc    10941
LOC105372935.clincRNA.NCBI.ref      ------------------------------------------------------------    8376

GGT5.NCBI.REF                       aaccgtagtgtgttgtggacctgtgagcagctggatttgatggggaagcctggttgtgtt    11001
LOC105372935.clincRNA.NCBI.ref      ------------------------------------------------------------    8376

GGT5.NCBI.REF                       ctgcaacctaatctgcggtttggtggcagccgacgtgccgcctccgtcaccattctccaa    11061
LOC105372935.clincRNA.NCBI.ref      ------------------------------------------------------------    8376

GGT5.NCBI.REF                       tgagtgccagctgagcaagcctggtggggaggcaggggggaggggagccccacagcgactc    11121
LOC105372935.clincRNA.NCBI.ref      ------------------------------------------------------------    8376

GGT5.NCBI.REF                       tgtgccatgtcccctagagccatcctccagcagcagggctcacccgtggatgccaccatc    11181
LOC105372935.clincRNA.NCBI.ref      ------------------------------------------------------------    8376

GGT5.NCBI.REF                       gcggctctggtctgcaccagcgtcgtcaaccctcagagcatgggcctgggcggaggggtc    11241
LOC105372935.clincRNA.NCBI.ref      ------------------------------------------------------------    8376

GGT5.NCBI.REF                       atcttcaccatctacaatgtgacaacaggtgggcccacatgtgaactgccatggggaaga    11301
LOC105372935.clincRNA.NCBI.ref      ------------------------------------------------------------    8376

GGT5.NCBI.REF                       tgggcttgtccacaccactgcgtaacccccttctccagccagtcccatcaactcaagcccg    11361
LOC105372935.clincRNA.NCBI.ref      ------------------------------------------------------------    8376
```

```
GGT5.NCBI.REF                    tttctgggtgttcttgggcccctgccctgccccatttgcctcagctgtgcaaagccctt    11421
LOC105372935.clincRNA.NCBI.ref   ------------------------------------------------------------    8376

GGT5.NCBI.REF                    gtgtcaactcctgtccctgtgttagcccagagtccatctcataatgcagagatggaaact    11481
LOC105372935.clincRNA.NCBI.ref   ------------------------------------------------------------    8376

GGT5.NCBI.REF                    gaggcctagagcaggccaggggctgtctgcagggtccatgaccactcacctgccctgccc    11541
LOC105372935.clincRNA.NCBI.ref   ------------------------------------------------------------    8376

GGT5.NCBI.REF                    tactccagggaaggtggaggtcatcaatgcccgggagacggtgccggccagccacgcccc    11601
LOC105372935.clincRNA.NCBI.ref   ------------------------------------------------------------    8376

GGT5.NCBI.REF                    gagcctgctggaccagtgtgcacaggctctgccactgggcacaggtgacgccccatggag    11661
LOC105372935.clincRNA.NCBI.ref   ------------------------------------------------------------    8376

GGT5.NCBI.REF                    gtcccccactcccacccccccaggccactaggactaatcagaacacccccgcaaaggagca    11721
LOC105372935.clincRNA.NCBI.ref   ------------------------------------------------------------    8376

GGT5.NCBI.REF                    gcgcctcacggacaggctgtgactccctggcacgggggacacggagcctgtggggaacc    11781
LOC105372935.clincRNA.NCBI.ref   ------------------------------------------------------------    8376

GGT5.NCBI.REF                    cccagccccaacctgcagggattgagaaggtccccttgggaacgggctctctgaggtga    11841
LOC105372935.clincRNA.NCBI.ref   ------------------------------------------------------------    8376

GGT5.NCBI.REF                    tccagggagagggaccaggctggacaaaggcctggagggttcagtggagatgggttggcg    11901
LOC105372935.clincRNA.NCBI.ref   ------------------------------------------------------------    8376

GGT5.NCBI.REF                    gggcagggtggggctcctggggccagtttgtgtagggccccttgggcttggccatggact    11961
LOC105372935.clincRNA.NCBI.ref   ------------------------------------------------------------    8376

GGT5.NCBI.REF                    ttgtggggagccacaggtgatttgagcagggagggacgtggtcgggtgtgggctccagaa    12021
LOC105372935.clincRNA.NCBI.ref   ------------------------------------------------------------    8376

GGT5.NCBI.REF                    agctcactctggctgctgtaactgggggccacatgtaggggtgatggcaaacctgggaa    12081
LOC105372935.clincRNA.NCBI.ref   ------------------------------------------------------------    8376

GGT5.NCBI.REF                    ggcctggaagccacagggttgagagctgggatgtgccatgcaggggcccagtggatcggg    12141
LOC105372935.clincRNA.NCBI.ref   ------------------------------------------------------------    8376

GGT5.NCBI.REF                    gtgcccggggagctccgtggctatgccgaggcccaccgccgccatggccgcctgccctgg    12201
LOC105372935.clincRNA.NCBI.ref   ------------------------------------------------------------    8376

GGT5.NCBI.REF                    gcgcagctgttccagcccaccatcgcgctgctccgagggggcatgtggtggcccctgtc    12261
LOC105372935.clincRNA.NCBI.ref   ------------------------------------------------------------    8376

GGT5.NCBI.REF                    ctcagccgtttcctgcacaacagcatcctgcggccttccttgcaggcgtcaaccctgcgg    12321
LOC105372935.clincRNA.NCBI.ref   ------------------------------------------------------------    8376

GGT5.NCBI.REF                    tgagccccacatggtggccctgggttcctgggttcaaggccatatcctgtctgggcccag    12381
LOC105372935.clincRNA.NCBI.ref   ------------------------------------------------------------    8376

GGT5.NCBI.REF                    tccacaccccagctctgcctcagtccctcttacactggaggattgaccctcggggtgctg    12441
LOC105372935.clincRNA.NCBI.ref   ------------------------------------------------------------    8376

GGT5.NCBI.REF                    gccaggacaaagagtacttttcatggcccttccagctccccgtggcagagcctaacttgg    12501
LOC105372935.clincRNA.NCBI.ref   ------------------------------------------------------------    8376

GGT5.NCBI.REF                    ggttaggctgcggggagtttattgcccaagtgataggactccaaggtgtggatttgggct    12561
LOC105372935.clincRNA.NCBI.ref   ------------------------------------------------------------    8376

GGT5.NCBI.REF                    ctgccacccgccctggggccccggcctccttcttcctctctctggcatggcaggagggct    12621
LOC105372935.clincRNA.NCBI.ref   ------------------------------------------------------------    8376
```

```
GGT5.NCBI.REF                    tcaactgccactctcacctgccccacccagcccactgacaggccctcccaggagtgagat    12681
LOC105372935.clincRNA.NCBI.ref   ------------------------------------------------------------    8376

GGT5.NCBI.REF                    gacaggccctaggtggacccaggccccactgggaaatttgccttccctggaccagggcca    12741
LOC105372935.clincRNA.NCBI.ref   ------------------------------------------------------------    8376

GGT5.NCBI.REF                    tgagctgggtgagtggtgtcagtcccaccacctcctagagaatcagttccccggtgtcca    12801
LOC105372935.clincRNA.NCBI.ref   ------------------------------------------------------------    8376

GGT5.NCBI.REF                    ccccagtgcctcgggaccctgggccagggcgccccccactgtagggtcctgaggactcca    12861
LOC105372935.clincRNA.NCBI.ref   ------------------------------------------------------------    8376

GGT5.NCBI.REF                    cccatgagagccccggaagagtgtggctgcctctgggacctggccccaccctgaggctcc    12921
LOC105372935.clincRNA.NCBI.ref   ------------------------------------------------------------    8376

GGT5.NCBI.REF                    ccccacccccccagccagctcttcttcaacgggacagaacccctgaggcctcaggacccac    12981
LOC105372935.clincRNA.NCBI.ref   ------------------------------------------------------------    8376

GGT5.NCBI.REF                    tcccatggcctgcactggccaccaccctggagaccgtggccacagagggcgtggaggtct    13041
LOC105372935.clincRNA.NCBI.ref   ------------------------------------------------------------    8376

GGT5.NCBI.REF                    tctacacggggaggctgggccagatgctggtggaggacattgccaaggaaggtcagcctc    13101
LOC105372935.clincRNA.NCBI.ref   ------------------------------------------------------------    8376

GGT5.NCBI.REF                    cctgaggttccataccccatccctgctggaggaagagccaagttctgggcactgggggtt    13161
LOC105372935.clincRNA.NCBI.ref   ------------------------------------------------------------    8376

GGT5.NCBI.REF                    gagggcaggctccttcccaggccctgagggagtgcaggcagtgcccgggaccccgagagg    13221
LOC105372935.clincRNA.NCBI.ref   ------------------------------------------------------------    8376

GGT5.NCBI.REF                    ccatgcacctagctctgccggcccaaggacccctgtggtccaggacctcagagagcagag    13281
LOC105372935.clincRNA.NCBI.ref   ------------------------------------------------------------    8376

GGT5.NCBI.REF                    cctgcgggttagggccagcagaggggcgttaccgatggtcacggcaggtctgggtagccg    13341
LOC105372935.clincRNA.NCBI.ref   ------------------------------------------------------------    8376

GGT5.NCBI.REF                    agctgaccattctcacaggctctcataagccagctccctctcccatcttccctttctttt    13401
LOC105372935.clincRNA.NCBI.ref   ------------------------------------------------------------    8376

GGT5.NCBI.REF                    ccccagcctccctgcctctggtctctgtccacccctccacgcctggccccaaggggccag    13461
LOC105372935.clincRNA.NCBI.ref   ------------------------------------------------------------    8376

GGT5.NCBI.REF                    tcctgaggccactgcctacagacctaccatcatcctgtgtccacaaggcaaaagcatgag    13521
LOC105372935.clincRNA.NCBI.ref   ------------------------------------------------------------    8376

GGT5.NCBI.REF                    tggaatcccattgtgtgacctgacagcagtggctattcctccccgcctcagtttcccatc    13581
LOC105372935.clincRNA.NCBI.ref   ------------------------------------------------------------    8376

GGT5.NCBI.REF                    tgttcatggagccctctccttctcatcccagggagccagctgacgctgcaggacctggcc    13641
LOC105372935.clincRNA.NCBI.ref   ------------------------------------------------------------    8376

GGT5.NCBI.REF                    aagttccagcccgaggtggtggatgccctggaggtgcccctggggactataccctgtac    13701
LOC105372935.clincRNA.NCBI.ref   ------------------------------------------------------------    8376

GGT5.NCBI.REF                    tcaccaccgccgcctgcaggggggtgccattctcagctttatcctcaacgtgctaagaggt    13761
LOC105372935.clincRNA.NCBI.ref   ------------------------------------------------------------    8376

GGT5.NCBI.REF                    aaagcccctgcccagagccctggcgcccacccccctcccccgcacccccggcccccgggaa    13821
LOC105372935.clincRNA.NCBI.ref   ------------------------------------------------------------    8376

GGT5.NCBI.REF                    ctcagactccaagcccaagcccagccaaggcccagctcagcctcctcctccataaaccgg    13881
```

```
LOC105372935.clincRNA.NCBI.ref      ------------------------------------------------------------    8376

GGT5.NCBI.REF                       gttcacagatgcacaatatgaagacagagctgtggaaaccaatcaggatggtgggtttta    13941
LOC105372935.clincRNA.NCBI.ref      ------------------------------------------------------------    8376

GGT5.NCBI.REF                       gagctggcctccagctcgacccatacgtgaggctttgctgaagggcgtgatatcattcgt    14001
LOC105372935.clincRNA.NCBI.ref      ------------------------------------------------------------    8376

GGT5.NCBI.REF                       ccagagtcctcctcccactgggtgaccagtcccagacacagaagcaggtgggctgtgtgg    14061
LOC105372935.clincRNA.NCBI.ref      ------------------------------------------------------------    8376

GGT5.NCBI.REF                       gtggtgagctgcctgtcaggggcgtgcacctgggggagcgaagccagcctagcccaaggc    14121
LOC105372935.clincRNA.NCBI.ref      ------------------------------------------------------------    8376

GGT5.NCBI.REF                       agcctgcaggggggttctggcagtgctgaggcctggggctgcctgcgctggcttaatatt    14181
LOC105372935.clincRNA.NCBI.ref      ------------------------------------------------------------    8376

GGT5.NCBI.REF                       agtgtcctgtgtctgccattacaaatcaccacaagcttggtggcttaaaataacagaaat    14241
LOC105372935.clincRNA.NCBI.ref      ------------------------------------------------------------    8376

GGT5.NCBI.REF                       gtgtcctctcacagttctggaagccagaagcccgatatctaggtgtccacagggctgcat    14301
LOC105372935.clincRNA.NCBI.ref      ------------------------------------------------------------    8376

GGT5.NCBI.REF                       tccattgggggctccaggggagaatctgttctttgccttgttgacctaaataaaaaacag    14361
LOC105372935.clincRNA.NCBI.ref      ------------------------------------------------------------    8376

GGT5.NCBI.REF                       aacaaggctctcaaagaaaatgatgtttgttccagagtaggcatcgcggtaggaataccc    14421
LOC105372935.clincRNA.NCBI.ref      ------------------------------------------------------------    8376

GGT5.NCBI.REF                       atgctagggtaaactatgtgtgtattcagggaggtgaaggaagacaaaagttttcaaaga    14481
LOC105372935.clincRNA.NCBI.ref      ------------------------------------------------------------    8376

GGT5.NCBI.REF                       gaaaatgaggaagattatataacttttttgtttacaatcaatgataatttattgtacatt    14541
LOC105372935.clincRNA.NCBI.ref      -------------cttatatgcttttattgcatttgagcgtacctcttttacagcgaaga    8423
                                                  ******   *** ***  *   * *     *  ****  *   *

GGT5.NCBI.REF                       ttatttttttcattgcattattattattattattattaattattgagacggtgtcttgct    14601
LOC105372935.clincRNA.NCBI.ref      tcttttttttaa------------------------------------------------    8435
                                    *  ******* *

GGT5.NCBI.REF                       ctgtcgcctagggtagagcacagtggcacaatctcggttcattgcaaactctgcttcctg    14661
LOC105372935.clincRNA.NCBI.ref      ------------------------------------------------------------    8435

GGT5.NCBI.REF                       ggttcaagtgattctcctgactcagcctcccaagtagctgggattacaggcatgggccac    14721
LOC105372935.clincRNA.NCBI.ref      ------------------------------------------------------------    8435

GGT5.NCBI.REF                       cacacgccgctgattttatcttttctttttttcctttttttttttttttctttttagaca    14781
LOC105372935.clincRNA.NCBI.ref      -------------------------------------attttattttatttttagagatg    8458
                                                                          **** **** * ***  ***

GGT5.NCBI.REF                       gagtctcaatctgtcacccaggctagagtgcaatggcacgatctcagctcactgcaacct    14841
LOC105372935.clincRNA.NCBI.ref      gagtcttgctttgttgcccaggctggagcactgtggtgtgatcatagctcactgcagcct    8518
                                    ******   * ***  ******** ***  *  ***   ****  *********** ***

GGT5.NCBI.REF                       ctgcctcccaggtacaagcaattctcctgcctcagtctcccaagtagctgggattacagg    14901
LOC105372935.clincRNA.NCBI.ref      tgaactcccgggcacaggtgatcctcccacctcagcctcctgaatagctgggactacagg    8578
                                        ***** ** *** *  ** ****  ****** ****  * ********* ******

GGT5.NCBI.REF                       cgcttgccacgacaccgagttaatttttgtatttttggtagagatggggttttgccacgt    14961
LOC105372935.clincRNA.NCBI.ref      catgcaccaccatgcctggcatattttaaagatgtttgtagagatgaggtctcgctatgt    8638
                                    *     **** *  **  *   *****     * ** ********* *** * ** * **

GGT5.NCBI.REF                       tggtcaggctggtcttgaacttctgatgtcaggtgatccgcccacctcggcatcccaaag    15021
LOC105372935.clincRNA.NCBI.ref      tgc-caggcttgtctcaaactactgggctcaagccatccatccatttcagcctcccaaag    8697
                                    **  ****** ****  **** ***   *** *  ****  ***  ** ** ********

GGT5.NCBI.REF                       tgctgggattacaggcatgagccaccgcgcccagatgattttatctttttagtagagac    15081
LOC105372935.clincRNA.NCBI.ref      tgcttggattataggcatgagc--------------------------------------    8719
                                    **** ****** **********
```

```
GGT5.NCBI.REF                    ggggttttaccatgttggccgggctggtcttgaactcttgaccttaactgacccacttgc    15141
LOC105372935.clincRNA.NCBI.ref   ------------------------------------------------------------    8719

GGT5.NCBI.REF                    ctcagcctcccaaagtgctgggattccaggcgtgagccacctctgcctggctattttatt    15201
LOC105372935.clincRNA.NCBI.ref   ---------------------------------------actgcgcctggccatcacact    8740
                                          **  ******* **   * *

GGT5.NCBI.REF                    tttatttcattagtttttggggaacaagtggtgttcggttgcatggaaaagttatttagt    15261
LOC105372935.clincRNA.NCBI.ref   gtttttttgtttgtttgt------------------------------------------    8758
                                  ** ***  ** **** *

GGT5.NCBI.REF                    ggtgatttctttttttttttcttttttttttgagatggaatctcactctgtggcccaggc    15321
LOC105372935.clincRNA.NCBI.ref   --------------------ttgtttgttttgagatggagtcttgctctgtcgcccaggc    8798
                                                     ** *** ************ ***  ****** ********

GGT5.NCBI.REF                    tggagagcagtggcatgatcttagctcactgcaacctctgcctctgaggttcaagcaatt    15381
LOC105372935.clincRNA.NCBI.ref   tggagtgcagtagtgggatctcacctcattgcaagctccgccttgtgggttcacgccatt    8858
                                 ***** ***** *   ***** * **** ***** *** ****    ****** ** ***

GGT5.NCBI.REF                    ctccagtctcagcctcctgagtagctgggattacaggtgcacgctaccaagcctggttaa    15441
LOC105372935.clincRNA.NCBI.ref   ctcctgcctcagcctcctgagtagctgggattacaggcgcccgccaccacgcctggctaa    8918
                                 **** * ****************************** ** *** **** ****** ***

GGT5.NCBI.REF                    ttaattaatttatttatttattttattttttttagtagagatggggtttcaccatgttggc    15501
LOC105372935.clincRNA.NCBI.ref   tt------------------ttttgtatttttagtagagacgggatttcaccgtgttagc    8960
                                 **                  **** * ************* *** ******* **** **

GGT5.NCBI.REF                    caggctggtcttaaactcctgacttcaagtgatctgcctgcctcagcctcccaaagtgct    15561
LOC105372935.clincRNA.NCBI.ref   caggatggtctcgatctcctgac----cgtgatctgcccgcctcggcctctcaaagtgct    9016
                                 **** ******  * ********     ********** ***** ***** *********

GGT5.NCBI.REF                    ggaattacaggcatgagccaccgcatctggtctttagtggtgatttctgagatcttgatg    15621
LOC105372935.clincRNA.NCBI.ref   gggattacaggcatgagccaataaatatttttataatcattaactaccgag---------    9067
                                 ** *****************    ** *  *  * *    * * * * ***

GGT5.NCBI.REF                    cacccatcacacaagcagtgtacactgttcctagtgtgtagtcttttatccctcattccc    15681
LOC105372935.clincRNA.NCBI.ref   ------------------------------------------------------------    9067

GGT5.NCBI.REF                    ctcccacccttccccttgagttcccagagtccattagagcatttttcttttcctctgaga    15741
LOC105372935.clincRNA.NCBI.ref   ------------------------------------------------------------    9067

GGT5.NCBI.REF                    cagagtctcgctctgtcgccaaggctggagtgcaatggtgcgtctcggttcactacaaac    15801
LOC105372935.clincRNA.NCBI.ref   ------------------------------------------------------------    9067

GGT5.NCBI.REF                    tctgcctcctgggttcaagcgattttcctgcctcagcctcccacgtagctgggattacag    15861
LOC105372935.clincRNA.NCBI.ref   ------------------------------------------------------------    9067

GGT5.NCBI.REF                    gcgcctgccaccacaactaatttttttttttttgtattattagtaaagatggggtttcacc    15921
LOC105372935.clincRNA.NCBI.ref   ------------------------------------------------------------    9067

GGT5.NCBI.REF                    gtgttgaccaggctgctcttgaactcctgacctcaggtgatctgcctgccttggcctccc    15981
LOC105372935.clincRNA.NCBI.ref   ------------------------------------------------------------    9067

GGT5.NCBI.REF                    aaattgctgggattacaggtgtgagccaccatgctgggccccattatatcattccttctt    16041
LOC105372935.clincRNA.NCBI.ref   ------------------------------------------------------------    9067

GGT5.NCBI.REF                    ttttctggagatggagttttgctcttgtcacccaggctggagtggagtgcaattgcgcga    16101
LOC105372935.clincRNA.NCBI.ref   ------------------------------------------------------------    9067

GGT5.NCBI.REF                    tctcagctcactgcaacctctgcctcccggttcaagcgattctcctgcctcagcctcccg    16161
LOC105372935.clincRNA.NCBI.ref   ------------------------------------------------------------    9067

GGT5.NCBI.REF                    agtagctaggactacccatgtgtgctaccatgcccggctaatttttgtattattagagac    16221
LOC105372935.clincRNA.NCBI.ref   ------------------------------------------------------------    9067

GGT5.NCBI.REF                    agggtttcatcatgttggccagactgctcttgaactcctgacctcaggtgatccacctgc    16281
LOC105372935.clincRNA.NCBI.ref   ------------------------------------------------------------    9067

GGT5.NCBI.REF                    ctcggcctcccaaagtgttgggattataggcataagccaccatgcctggcccattatatc    16341
LOC105372935.clincRNA.NCBI.ref   ------------------------------------------------------------    9067
```

```
GGT5.NCBI.REF                       atttttattcctttgcgtcctcatagcttagctttggtatataactgtttcgggcctggc    16401
LOC105372935.clincRNA.NCBI.ref      ------------------------------------------------------------    9067


GGT5.NCBI.REF                       atggtggctcacgcctgtaatcccagcactttgggaggccaaggcaggcggatcatgagt    16461
LOC105372935.clincRNA.NCBI.ref      ------------------------------------------------------------    9067


GGT5.NCBI.REF                       tcaggagatcgagaccatcctgcctaacacagtgaaaccctgtctctactaaaaatacaa    16521
LOC105372935.clincRNA.NCBI.ref      ------------------------------------------------------------    9067


GGT5.NCBI.REF                       aaaattagccaggcgtggtggcgggcacctgtagtcccagttgcttgggaggctgaggca    16581
LOC105372935.clincRNA.NCBI.ref      ----------------------gagaagctggagagaaaaaaggtggatgaagttaaggc    9105
                                                          * * * *** **    *   * * *      * * *

GGT5.NCBI.REF                       ggagaatggtgtgaacccaggagacagagcttgcagtgagctgagatcaca----ccact    16637
LOC105372935.clincRNA.NCBI.ref      agagagactttgtaagtttcttagctgagcttttggagactgtatatcagagatcctatt    9165
                                     ****    *   **         * ******   * **    * **** *    * * *

GGT5.NCBI.REF                       gcactccagcctgggcaacagagtgagactctgtctcaaaaaaaaaaaaaaactgttttg    16697
LOC105372935.clincRNA.NCBI.ref      tgaggtgattttcggctacagagataggcctggtcatttgaaaaataacgga--------    9217
                                      *    *   * *** ******  ** *   ***     ***** **   *

GGT5.NCBI.REF                       aaatgatcatccttggctagtaggatgcatggaaaagtgataagggtggcactggacagg    16757
LOC105372935.clincRNA.NCBI.ref      ------------------------------------------------------------    9217


GGT5.NCBI.REF                       ccactcatgggcagatgtctttgcagaagtattttttgtgtaagattgtgatggccttct    16817
LOC105372935.clincRNA.NCBI.ref      ------------------------------------------------------------    9217


GGT5.NCBI.REF                       tgtaaggttgtgagctgttgtgtttttgaggcagggtcttgctctgtcacctgggctgga    16877
LOC105372935.clincRNA.NCBI.ref      ------------------------------------------------------------    9217


GGT5.NCBI.REF                       gtgcagtggcactgtcaaggctcactgcagcctcaacctcctggactcaagcaatcctct    16937
LOC105372935.clincRNA.NCBI.ref      ------------------------------------------------------------    9217


GGT5.NCBI.REF                       cacctcagcctcccaaagttttgggattacaggcatgagccactgcacccagccgagaac    16997
LOC105372935.clincRNA.NCBI.ref      ------------------------------------------------------------    9217


GGT5.NCBI.REF                       ggggcacttgtgcacgagagtcctgtcttcatggcctttccctggctctatttttcagag    17057
LOC105372935.clincRNA.NCBI.ref      ------------------------------------------------------------    9217


GGT5.NCBI.REF                       tttttttttttttgttttgttttttttttttttttttgagacagagtcttgctctgttta    17117
LOC105372935.clincRNA.NCBI.ref      ------------------------------------------------------------    9217


GGT5.NCBI.REF                       ctaggctggagtgcagtggtgcaatcctgcctcactgcaacctcctcctcccgggttcaa    17177
LOC105372935.clincRNA.NCBI.ref      -ttaggtgaagcttgcgtttgaaatcctccttctacttctcatctttctctcttgtttat    9276
                                     *  * ** **        ** ****** * **       * ** * *** *  *** *

GGT5.NCBI.REF                       gcaattctccctcctcagcctccctagtagctggggttacaggcacccaccatcatgcct    17237
LOC105372935.clincRNA.NCBI.ref      tctgaacatccatcttagacaccc------------------------------------    9300
                                     *    *  **  ** ** * ***

GGT5.NCBI.REF                       ggataatttttgtatttttgtagagatggggtttcaccatgttagtcaggctggtc----    17293
LOC105372935.clincRNA.NCBI.ref      ----aatctttgtcactttgtggttgtcattgatttccctgttattgagattagaggttg    9356
                                        *** *****   ***** *   *      *  ** ***** * **  * *

GGT5.NCBI.REF                       ------------------------------------------------------------    17293
LOC105372935.clincRNA.NCBI.ref      gcagactttctctgtaaagggctggagagaaagtacttcagactttgtggtctgtgcagt    9416


GGT5.NCBI.REF                       ------ttgaactcctgacctcagcctcggcctcccaaagtgctgggactacaggtgtga    17347
LOC105372935.clincRNA.NCBI.ref      gtctgttgcaaccacttaactctgccctgtagagcaaaagcagccgtagacagtacatgg    9476
                                          *  ***  ** * *** ***  *     * ****     * *         **

GGT5.NCBI.REF                       gccactgtggctggctgccgctctattttgaaaattttcacagcctcttcccacttccgg    17407
LOC105372935.clincRNA.NCBI.ref      gcagatgagcatggctggggtccagttacatttacttgcaa-------------------    9517
                                    **   ** *  ******  *  *  **      * ** **

GGT5.NCBI.REF                       tggctgccagcattccttggcttgtggctgcctcactccagtctctgtctccatgatcat    17467
LOC105372935.clincRNA.NCBI.ref      ------------------------------------------------------------    9517


GGT5.NCBI.REF                       actggtttctcctctgctgtgtgtcctctcctctatgtgtctgtcttacaaggacactgt    17527
LOC105372935.clincRNA.NCBI.ref      ------------------------------------------------------------    9517
```

```
GGT5.NCBI.REF                       ggtcacattcagggcacatctagataatccaggatcatctcctcctctcaaaatctttaa    17587
LOC105372935.clincRNA.NCBI.ref      ----------------------------------------------------------aa    9519
                                                                                              **

GGT5.NCBI.REF                       cgtactttaggccgggtgtggtggctcatgcctgtaatcccagcactttgggaggctaag    17647
LOC105372935.clincRNA.NCBI.ref      agagggttggagactgggtgcgatctcccacctttaatcccggcactttgggaggccgag    9579
                                     *    ** *     * ***    ***   *** ******* **************  **

GGT5.NCBI.REF                       gagggtggatcacttgaggtcaggagttggagaccagcctggccaacacagtaaaacacc    17707
LOC105372935.clincRNA.NCBI.ref      gcaggaggatcacttcaagccaggagttcaagaccagcctgggcaacaaagcaagactcc    9639
                                    *  ** ********* * * ********  ************ ***** ** ** ** **

GGT5.NCBI.REF                       atctctactaaaaatacaaaaat----tagccgggcatgatggcctgtgcctgaagtcct    17763
LOC105372935.clincRNA.NCBI.ref      atctctacaaaaaaataaaaattattatagctaggcatggtggtacacacccgtagtcct    9699
                                    ******** *****   **** *    ****  ****** ***      ** * ******

GGT5.NCBI.REF                       agctacttgggagg------ctgaggcgggagaatcacttcaacccaggaggtggagatt    17817
LOC105372935.clincRNA.NCBI.ref      agctactcaggaagctaaaactgaggcaggagggtcagttgagcccaggagcacgaggct    9759
                                    *******  *** *      ******* ****  *** ** * ********   ***  *

GGT5.NCBI.REF                       gcagtaagctgagatcgtgccacacactccagcctgggtgacagaacaag-acttcctct    17876
LOC105372935.clincRNA.NCBI.ref      gtggtgagctattattgtgccagtgcactccagcctggtgacagagcaagaaccgtctca    9819
                                    *  ** ****   ** ******       *   *  ********* **** **   ***

GGT5.NCBI.REF                       ccaaaaaacacaaaaaaggcggggcatgatggcttacgcctgtaatcctagcactttggg    17936
LOC105372935.clincRNA.NCBI.ref      tcaaaaaataaacaaaaggctgggcacggtggctcacgcctgtaatccctgcactttggg    9879
                                     ******* * * ******* ***** * ***** *************  **********

GGT5.NCBI.REF                       aggccaaggcaggcagatcacttgaggccaggagttcaataccagcctgaacaacatggc    17996
LOC105372935.clincRNA.NCBI.ref      agactaaggtgggcagatca--tgaggtcaggagattgagaccatcctggctaacacggt    9937
                                    ** * ****  *********  ***** ****** *  * **** ****   **** **

GGT5.NCBI.REF                       gaaatcctgtctactaaaaatacaaaaataattagctgtacgtggtggcgaacacatgta    18056
LOC105372935.clincRNA.NCBI.ref      gaaaccctgtctctactaaaaatacaaaaagttagccgggcgtggtggtgggcgcctgta    9997
                                    **** *******     *** * * *** * ***** *  ******** *  * * ****

GGT5.NCBI.REF                       gtctcagctactcaggagactaaggaccaagaatcacttgaacccaggaggcagagattg    18116
LOC105372935.clincRNA.NCBI.ref      gtcccagccactcgggaggctgaggagggagaatcgtttgaacctgggaggcggaggttg    10057
                                    *** **** **** **** ** ****   ******  *******  ****** *** ***

GGT5.NCBI.REF                       cagtgagctgagactgcgccattgcactccagcctgggcaatagagtgagactctgtcta    18176
LOC105372935.clincRNA.NCBI.ref      cagtgagccaaggttgtgccactgcactctagcctgggctacagggcaagactccattaa    10117
                                    ********  **  ** **** ******* ********* * ** *  ******  *  *

GGT5.NCBI.REF                       aaaaaaaaaaaaaaaaaaaaaaaaatccttaacttacttttccacttaaggtattagtcac    18236
LOC105372935.clincRNA.NCBI.ref      aaaaaaaaaaaaaaaaccagcaaaaaccaaacaaaacat----------------------    10155
                                    ***************  *  **** **  *    ** *

GGT5.NCBI.REF                       tcttgtgctgtataaggtaatatccacaggttttgggaattaggatgtgggtggatcttt    18296
LOC105372935.clincRNA.NCBI.ref      ------------------------------------------------------------    10155

GGT5.NCBI.REF                       ctgtggtggggggtgggggcaacattcaacccattacatagggtgaccccaaccaacctg    18356
LOC105372935.clincRNA.NCBI.ref      ------------------------------------------------------------    10155

GGT5.NCBI.REF                       tgccccaacctctctccagggttcaacttctcaacagagtctatggccaggcctgaaggg    18416
LOC105372935.clincRNA.NCBI.ref      ------------------------------------------------------------    10155

GGT5.NCBI.REF                       agggtgaacgtgtaccaccaccttgtagagacgctcaagtttgccaaggggcagaggtgg    18476
LOC105372935.clincRNA.NCBI.ref      ------------------------------------------------------------    10155

GGT5.NCBI.REF                       aggctgggggaccctcgaagccacccgaagctccaggtgaggttgctgaggttgctgggc    18536
LOC105372935.clincRNA.NCBI.ref      ------------------------------------------------------------    10155

GGT5.NCBI.REF                       tggtgggccgtcctcctccctggctcaggacttggcatgaaatgagggtcaggcctggta    18596
LOC105372935.clincRNA.NCBI.ref      ------------------------------------------------------------    10155

GGT5.NCBI.REF                       gggggaagttggagggatatgtatgtggttctaggccagggcaggactgaaagggatccc    18656
LOC105372935.clincRNA.NCBI.ref      ------------------------------------------------------------    10155

GGT5.NCBI.REF                       ggggtggcaggtacaggggtcaggtgcaggagtggcaccatatctcaaaggacctggagg    18716
LOC105372935.clincRNA.NCBI.ref      ------------------------------------------------------------    10155

GGT5.NCBI.REF                       gtgagcagagtctagacctagctgggcttgagggagacctggccacaaggtagaggacag    18776
```

```
LOC105372935.clincRNA.NCBI.ref      ------------------------------------------------------------    10155

GGT5.NCBI.REF                       actggaggtggcccccatgggggctgatctcatcctgcccttggttctgcggattctgcc    18836
LOC105372935.clincRNA.NCBI.ref      ------------------------------------------------------------    10155

GGT5.NCBI.REF                       tggcccctcactgaccctgccacctgcccacccaccccagaatgcctcccgggacctgct    18896
LOC105372935.clincRNA.NCBI.ref      ------------------------------------------------------------    10155

GGT5.NCBI.REF                       gggggagaccctggcccagctcatccgccaacagatcgatggccgggggaccaccagct    18956
LOC105372935.clincRNA.NCBI.ref      ------------------------------------------------------------    10155

GGT5.NCBI.REF                       cagccactacagcttggccgaggcctggggccacgggacaggcacgtcccatgtgtctgt    19016
LOC105372935.clincRNA.NCBI.ref      ------------------------------------------------------------    10155

GGT5.NCBI.REF                       gctgggggaggatggcagcgccgtggctgccaccagcaccatcaacacaccgtgcgtagg    19076
LOC105372935.clincRNA.NCBI.ref      ------------------------------------------------------------    10155

GGT5.NCBI.REF                       gcctgggggaaggcggatggcttcactcctcctctcctagacctgcacaccccccagcccc    19136
LOC105372935.clincRNA.NCBI.ref      ------------------------------------------------------------    10155

GGT5.NCBI.REF                       atgtcccctcacttgtccccacggggcagcaccttgcttttgccctttttctcctcctct    19196
LOC105372935.clincRNA.NCBI.ref      ------------------------------------------------------------    10155

GGT5.NCBI.REF                       atttcaaaagaggcccccacccctgacatctctggctggaaaggctgctgctggggtggc    19256
LOC105372935.clincRNA.NCBI.ref      ------------------------------------------------------------    10155

GGT5.NCBI.REF                       cccgacccaagatttacctgggaatgggtagcctcactcagaagggtgccctgatgtggg    19316
LOC105372935.clincRNA.NCBI.ref      ----------------------aatgcatgttctctcttataaatgggagctaaacatgg    10193
                                                          ****  *   *** ** * **  * *  ** *    **

GGT5.NCBI.REF                       ggcacaggtgggtctttggggacccctcctgggtggtgccagggagagaatagcggcttc    19376
LOC105372935.clincRNA.NCBI.ref      ggactcattgacttaaagatggcaacaactgggaactgctggatggggag----------    10243
                                    **      **  *    *  * *  *  *****   ***  *   * **

GGT5.NCBI.REF                       agcatgcttcggggcagctgtaaaacgaggggggtcctgcaaagcgtgcagggtaagtggt    19436
LOC105372935.clincRNA.NCBI.ref      ------------------------------------------------------------    10243

GGT5.NCBI.REF                       gtctggtgggagccccgggtcctagcccaggctcttctgcctccacggctgcagctttgg    19496
LOC105372935.clincRNA.NCBI.ref      ------------------------------------------------------------    10243

GGT5.NCBI.REF                       agcgatggtgtattcaccacggacaggcatcatcctcaacaacgagctcctggacttatg    19556
LOC105372935.clincRNA.NCBI.ref      ------------------------------------------------------------    10243

GGT5.NCBI.REF                       cgagcgatgcccccggggttccggcaccacccccctcacctggtgagaacaaagcttccca    19616
LOC105372935.clincRNA.NCBI.ref      ------------------------------------------------------------    10243

GGT5.NCBI.REF                       cccggggtccacaagggccccccaccaggggagaggagggagggggctgggctggggttg    19676
LOC105372935.clincRNA.NCBI.ref      --------------------------ggaggggaggggtgaaaggccaactgttggggag    10277
                                                               * ** **** * **  ** *     * ***  *

GGT5.NCBI.REF                       catgctaacccctggatgggtcactgcacttgccaagacgctgtttgctcagcagtgagt    19736
LOC105372935.clincRNA.NCBI.ref      tatgctca----tatccatgtgacaaacctgcacatgtgcccgctgaatctaaaataaaa    10333
                                     ***** *    *      ** **    **   ** *   * * *   **   * * *

GGT5.NCBI.REF                       ggagacagggtgggtggagctcccggaaggtgctggcccccagttccaggcgagcgttcc    19796
LOC105372935.clincRNA.NCBI.ref      gttgaaagtagatttaaaaaaccccaagagggctgg------------------------    10369
                                    *  ** **      *  *   ***  *  * *****

GGT5.NCBI.REF                       ccatcctccatggtgccctccatcttgatcaacaaagcccaggggtcgaagctagtgatt    19856
LOC105372935.clincRNA.NCBI.ref      --------------------------------------------------------gttt    10373
                                                                                            * **

GGT5.NCBI.REF                       ggcggggctggcggggagctcatcatctctgctgtggcccaggtgagtctggggctcctg    19916
LOC105372935.clincRNA.NCBI.ref      ggcttgtgtgtccatagcttgttaacctccgct---------------------------    10406
                                    ***  *  ** *       *  * * *** ***

GGT5.NCBI.REF                       gctcgagtgtctcctctctgggcagcatactgtctgactgtctctggagtggggatgtga    19976
LOC105372935.clincRNA.NCBI.ref      ------------------------------------------------------------    10406
```

```
GGT5.NCBI.REF                       gggctgatgtagggtagcagggtgccccctttctccctgaaaccctcatctctcccccag    20036
LOC105372935.clincRNA.NCBI.ref      ------------------------------------------------------------    10406


GGT5.NCBI.REF                       gccatcatgagcaagctgtggcttggctttgacctgagagcggccattgcagcccccatc    20096
LOC105372935.clincRNA.NCBI.ref      ------------------------------------------------------------    10406


GGT5.NCBI.REF                       ctgcatgtcaacagcaagggctgtgtggagtacgagcccaacttcagccaggtgaggctg    20156
LOC105372935.clincRNA.NCBI.ref      ------------------------------------------------------------    10406


GGT5.NCBI.REF                       aggtccgagctggatgcctagggcagagcccactccccaaatccgtgctgctcaaagcca    20216
LOC105372935.clincRNA.NCBI.ref      ------------------------------------------------------------    10406


GGT5.NCBI.REF                       cctgggaggaactcagtcactgagattcttaggccaggtacacttcaactttgggggcca    20276
LOC105372935.clincRNA.NCBI.ref      -------------------------------------ttagatattaactaatagaaaca    10429
                                                                          ** *  * ****    *   **

GGT5.NCBI.REF                       taggagttggggaccttgatgggtgaggctgtcagtggcctccaggccagttctgtggcc    20336
LOC105372935.clincRNA.NCBI.ref      tagtgcttatcttcccaggccacctattttgttcctctccaaggtgatggatagatgaag    10489
                                    ***   **     **  *       *   ***   *  **     *   * *   **

GGT5.NCBI.REF                       tccaagacagagagcagggtattgtctatgctgctcccaggctgaggatctcagca-cct    20395
LOC105372935.clincRNA.NCBI.ref      gcctaatccagccgcctggaagtttgctgacgcttgtcctgtcacggattaatgaagcat    10549
                                     ** *  *     **  ** * * *     *   *  *  *    ****    * * * *

GGT5.NCBI.REF                       tggtctctggtctgtggttgatgccattttcagaagtgagttttcctggctggggcctc    20455
LOC105372935.clincRNA.NCBI.ref      tgttttctgatgaaggtttcatgccgctgtgctgatgtgtcttctcttctct--------    10601
                                    ** * **** *    * ** *****  * * * ** ***  ** ** *  **

GGT5.NCBI.REF                       tcagactctccctcatggtgactttttccttgtgtgatttgtaactaatatttgcaactt    20515
LOC105372935.clincRNA.NCBI.ref      ------------------------------------------------------------    10601


GGT5.NCBI.REF                       tatttgtgaaagaggttgtttttcctgtgacagtctgtaaagtgattttcttctggggca    20575
LOC105372935.clincRNA.NCBI.ref      -----------------------------ctaggcaggaaactgcatatcttctggttta    10632
                                                                   ** * * *** **  * ********   *

GGT5.NCBI.REF                       ctagggcattggtggtaatttttactatatatataatttttttttttttgagacagcat    20635
LOC105372935.clincRNA.NCBI.ref      catgaagatggagtgctaatggaaatgcccaaaacctt----------------------    10670
                                    *  *   ** *   *  * *   * *    * *   **

GGT5.NCBI.REF                       ctcactctgttgcccaggctggagtgcagtgttgcgatcacagctcactacagccttgac    20695
LOC105372935.clincRNA.NCBI.ref      ------------------------------------------------------------    10670


GGT5.NCBI.REF                       atcctgggctcaagtgatccttcctcctcaggctcccaaggaactaggactacaggtgtg    20755
LOC105372935.clincRNA.NCBI.ref      ------------------------------------------------------------    10670


GGT5.NCBI.REF                       tgccaccacacctggatagtttttatttttttgtttttcgtagaaacagggtctctcacc    20815
LOC105372935.clincRNA.NCBI.ref      ------------------------------------------------------------    10670


GGT5.NCBI.REF                       atgttgcctaggctggtcttgaactcctgagctcaagcaatcttcccgccttggcctccc    20875
LOC105372935.clincRNA.NCBI.ref      ------------------------------------------------------------    10670


GGT5.NCBI.REF                       aaaatgctgggattataggcgtgagccaccatgcccagcctacaaacatttttttgatt    20935
LOC105372935.clincRNA.NCBI.ref      ------------------------------------------------------------    10670


GGT5.NCBI.REF                       ccccacagaagcccatggtgcagctccaggggctgaattcttctggctgactccccctct    20995
LOC105372935.clincRNA.NCBI.ref      --------------------------------cagagattgacacgctgtcattttccat    10698
                                                                    * **  *   *  **** *     *  *

GGT5.NCBI.REF                       atccttagacagagactcagcttccttggagcatctttgggttgatggatgccctttgtt    21055
LOC105372935.clincRNA.NCBI.ref      ttccgttcctggatctacggagtcttctaagagattttgcaatgaggagaagcactgttt    10758
                                     *** *     **    * *  ** *   **    ****   *** *     *  *  **

GGT5.NCBI.REF                       aggacctatttaaaagattgatagtggccgggcatggtggctcatgcctgtaatcccagc    21115
LOC105372935.clincRNA.NCBI.ref      tcaaactatataactga-------------------------------------------    10775
                                       * **** ***  **

GGT5.NCBI.REF                       actttgggaggccgaggcgggtggatcacctgaggtcaggacttccaggccagcctggcc    21175
LOC105372935.clincRNA.NCBI.ref      ------------------------------------------------------------    10775


GGT5.NCBI.REF                       aacatggtaaaaccctgcctcaactaaaaatacaaaaattagccaggcgtggtggcacac    21235
LOC105372935.clincRNA.NCBI.ref      ----------------gccttatttataattagggatattatcaaatatg----------    10810
                                                    **** *  ** ** **   * **** * *    **
```

```
GGT5.NCBI.REF                       acctgtaatcccagctactcgggaggctgaagcaggagaaacacttgaacccaggaggta    21295
LOC105372935.clincRNA.NCBI.ref      ------------------------------------------------------------    10810

GGT5.NCBI.REF                       gaggttgcagtgagccaagatcgtgacattgcactccagcctgggtgacaagaacaaaac    21355
LOC105372935.clincRNA.NCBI.ref      ------------------------------------------------------------    10810

GGT5.NCBI.REF                       tccatctcaaaaaaaaaaaaaaaaaaaaaaaaaagattgatagcagctgaggttccagatc    21415
LOC105372935.clincRNA.NCBI.ref      ------------------------------------------------------------    10810

GGT5.NCBI.REF                       gtcaaagctgcctctcctgttatgagacttcttcggtctcaagtccaccttccaaagccc    21475
LOC105372935.clincRNA.NCBI.ref      ------------------------------------------------------------    10810

GGT5.NCBI.REF                       ttagaacctgagaaccatggccttagtccaggaacagagtgtagtcccaagagccttagt    21535
LOC105372935.clincRNA.NCBI.ref      ------------------------------------------------------------    10810

GGT5.NCBI.REF                       tttcagaccttgagactaaggacctcagcctgttaggctcaaactcttggagtttgcagg    21595
LOC105372935.clincRNA.NCBI.ref      ------------------------------------------------------------    10810

GGT5.NCBI.REF                       cctccctccgcatgtagggtccttcgcatgacaggaagaccctctcagggctctcgggtt    21655
LOC105372935.clincRNA.NCBI.ref      ------------------------------------------------------------    10810

GGT5.NCBI.REF                       ccagggatcttagatatggaggatttcaagatccagggcacatcctttggagccctcctc    21715
LOC105372935.clincRNA.NCBI.ref      ------------------------------------------------------------    10810

GGT5.NCBI.REF                       ccttgactcttttttttatttttatttttttagacagtctcactctgtcactcaggctgga    21775
LOC105372935.clincRNA.NCBI.ref      ------------------------------------------------------------    10810

GGT5.NCBI.REF                       gtgcagtggcatgatcttgcctcaccacaacctccgcttcccaggttcaagcaattctcc    21835
LOC105372935.clincRNA.NCBI.ref      ------------------------------------------------------------    10810

GGT5.NCBI.REF                       tgcctcagcctctgagtagctgggactacacatgtgcgccaccacacctggctaattttt    21895
LOC105372935.clincRNA.NCBI.ref      ------------------------------------------------------------    10810

GGT5.NCBI.REF                       gtatttttagtaaagatggggtttccccatgttggccaggctggtcttgagctcccaatc    21955
LOC105372935.clincRNA.NCBI.ref      ------------------------------------------------------------    10810

GGT5.NCBI.REF                       tcaagtgatctgtccaccttggcctcccaaagtgctgggcttataggcatgagccactgt    22015
LOC105372935.clincRNA.NCBI.ref      ------------------------------------------------------------    10810

GGT5.NCBI.REF                       gcctggccctcccttgactcttttttttttcttttttttgagacggagtctcgctctgttg    22075
LOC105372935.clincRNA.NCBI.ref      ------------------------------------------------------------    10810

GGT5.NCBI.REF                       cccaggctggcgtgcagtggcacgatcttggctcactgcaagctctgcctcccgagttca    22135
LOC105372935.clincRNA.NCBI.ref      ------------------------------------------------------------    10810

GGT5.NCBI.REF                       cgccattcccctgcctcatcctcccgcctcatcctcccaagtagctgggactacaggcac    22195
LOC105372935.clincRNA.NCBI.ref      ---------------taaccatgaggcccctcaggtcctgatcagtcagaatggatgctt    10855
                                                   * * * *   *** *      **   *   *  ** *  * **

GGT5.NCBI.REF                       ccaccaccacgctcggctaatttttgtatttttagtagagacagggtttcaccgtgtta    22255
LOC105372935.clincRNA.NCBI.ref      tcaccagcagacccggccatgt--------------------------------------    10877
                                     ***** **  * **** *  *

GGT5.NCBI.REF                       gccaggatggtctcgatctcctgaccttgtgatccgcccgccttggcctcccaaagtgct    22315
LOC105372935.clincRNA.NCBI.ref      ------------------------------------------------------------    10877

GGT5.NCBI.REF                       gggattataggcgtgagccaccgcgcccggcccctcccttgactcttgactgaaggacct    22375
LOC105372935.clincRNA.NCBI.ref      ------------------------------------------------------------    10877

GGT5.NCBI.REF                       ttgtctttgtgaacatcaattctcaggacctttcaccctggggacgtgaaatgctgagaa    22435
LOC105372935.clincRNA.NCBI.ref      ------------------------------------------------------------    10877

GGT5.NCBI.REF                       tttgggagatgacagtctggggactgggattaatggaatccagtgacccacaaacctaag    22495
LOC105372935.clincRNA.NCBI.ref      ------------------------------------------------------------    10877
```

```
GGT5.NCBI.REF                       gttctcagctcccttggggagttggaatgtcagctattcaggtctagggctttccatgga    22555
LOC105372935.clincRNA.NCBI.ref      ------------------------------------------------------------    10877


GGT5.NCBI.REF                       gtaaatcctaaactctgggttggagactttaagcctccaaggaccttcacagctaaggcc    22615
LOC105372935.clincRNA.NCBI.ref      ------------------------------------------------------------    10877


GGT5.NCBI.REF                       cagggactagggcgaggagagtctttgatcctcagagtcttggagtttggccagtggact    22675
LOC105372935.clincRNA.NCBI.ref      ------------------------------------------------------------    10877


GGT5.NCBI.REF                       ctgaggaatggagtctctgagcactgaaggtccaactttggcttcagcagtaaaggatct    22735
LOC105372935.clincRNA.NCBI.ref      ---------------------ggctgctcggtcctgggtgctcgctgctgtgcaagacat    10916
                                                           ***   *  *     **    * ** **  * **  *

GGT5.NCBI.REF                       tggccttcaagtctaaggacagtgggcaattagtaggtcaggcatggggaactcatagcc    22795
LOC105372935.clincRNA.NCBI.ref      tagccctttagttatgagcctgtgggaacttcaggggttcccagtggggagagcagtggc    10976
                                    * *** *  ***     * * ***** * **    ***      ******   **  * *

GGT5.NCBI.REF                       aaacgtgcagggctccaaagacctcatttgccctgtcagcagctcaggccatgtggcatc    22855
LOC105372935.clincRNA.NCBI.ref      ------------------------------------------------------------    10976


GGT5.NCBI.REF                       acccgatgcatctagatgtctccggaatctcaggccctgcagggtgaggttctcgggcca    22915
LOC105372935.clincRNA.NCBI.ref      ------------------------------------------------------------    10976


GGT5.NCBI.REF                       ttagtttttttttgttttgttttgttttgtttttttggcttgttgttgttgagacaaagtt    22975
LOC105372935.clincRNA.NCBI.ref      ------------------------------------------------------------    10976


GGT5.NCBI.REF                       tcactctgtcacccaggctggagtgcagtggcgcgatctcagcttattgcaacctccacc    23035
LOC105372935.clincRNA.NCBI.ref      ------------------------------------------------------------    10976


GGT5.NCBI.REF                       tcctgggttcaagcaattctcatgtctcagcctcccaagtagctgggattacaagtgtgt    23095
LOC105372935.clincRNA.NCBI.ref      ------------------------------------------------------------    10976


GGT5.NCBI.REF                       gccaccaagcctggctaatttttgtatttttagcagaaacagcgtttctccatgttggcc    23155
LOC105372935.clincRNA.NCBI.ref      ------------------------------------------------------------    10976


GGT5.NCBI.REF                       aggctggtctcaaactcctgacctcaggtgatctgcccaccttggcctcccaaagtgctg    23215
LOC105372935.clincRNA.NCBI.ref      ------------------------------------------------------------    10976


GGT5.NCBI.REF                       ggattacaggcatgaccaccgcgcctggctagggagcagtgtttttaaggacaacttggc    23275
LOC105372935.clincRNA.NCBI.ref      ------------------------------------------------------------    10976


GGT5.NCBI.REF                       gggttgggggaagccaatgagccaggagtgctgataggtcagggatgaaatcatagggag    23335
LOC105372935.clincRNA.NCBI.ref      ------------------------------------------------------------    10976


GGT5.NCBI.REF                       ttggctgcgcgcggtggctcccgcctacaatcccagcactttgggaggctgaggtgggtg    23395
LOC105372935.clincRNA.NCBI.ref      ------------------------------------------------------------    10976


GGT5.NCBI.REF                       gttcacctgaggtcaggagaccagcctggccaacatggcgaaaacctgtctctactaaaa    23455
LOC105372935.clincRNA.NCBI.ref      ------------------------------------------------------------    10976


GGT5.NCBI.REF                       ttacaaaaattagctgggcaaagtggcaggcatctgtaatcccagctactggggaggctg    23515
LOC105372935.clincRNA.NCBI.ref      ---------------------agtgggaggcatctgggggccaaaggtcag---------    11006
                                                         ***** *********    ** *    * *

GGT5.NCBI.REF                       aggcaagagaatcacttgaacctgggaggggaagttgcagtgagccaaggtcgtgccatt    23575
LOC105372935.clincRNA.NCBI.ref      ------------------------------------------------------------    11006


GGT5.NCBI.REF                       gcacgccagcatgggtgacagagcgggactccatctcagaaacagacacacacaaaaaac    23635
LOC105372935.clincRNA.NCBI.ref      ------------------------------------------------------------    11006


GGT5.NCBI.REF                       ccctggaaatcatagaatcatagggagttgaagctttcttcttgagctgagtccgttcct    23695
LOC105372935.clincRNA.NCBI.ref      ------------------------------------------------------------    11006


GGT5.NCBI.REF                       gggtgggggccacaaggtcagatgagccagttaatcgatctggatggtgccagctgatcc    23755
```

```
LOC105372935.clincRNA.NCBI.ref      ------------------------------------------------------------    11006

GGT5.NCBI.REF                       atcaagtgcagggtctgcaaaatttctcaagcactgatcttaagagcagtttagggaggg    23815
LOC105372935.clincRNA.NCBI.ref      ------------------------------------------------------------    11006

GGT5.NCBI.REF                       tcagaatctttagccttcagctgcgtgactcctaaaccataatttctaatcttgtggcta    23875
LOC105372935.clincRNA.NCBI.ref      ------------------------------------------------------------    11006

GGT5.NCBI.REF                       atgtgagtcctacgaaggcagtctggtgcccaggcaagaaggaagtctgctttgggaaag    23935
LOC105372935.clincRNA.NCBI.ref      ------------------------------------------------------------    11006

GGT5.NCBI.REF                       ggctgttaccgtctttgttttaaactataaactaagtttctctcaaagttagttcagcct    23995
LOC105372935.clincRNA.NCBI.ref      ------------------------------------------------------------    11006

GGT5.NCBI.REF                       acacccaggaatgagcaaggacaactgggaggttagaagcaagatggagtcgattaagtt    24055
LOC105372935.clincRNA.NCBI.ref      ------------------------------------------------------------    11006

GGT5.NCBI.REF                       agatctctttccctgtctcagtcataattttgcaaaggcggtttcagtctt-----gctg    24110
LOC105372935.clincRNA.NCBI.ref      -----------------------------tggcagggggtatttcagtattatacaactg    11037
                                                                 * ***  **   ******* **      ***

GGT5.NCBI.REF                       ttttgcccaggctggagtgcagtgtcatgatcaaagctcactgcagcctcaaactcctag    24170
LOC105372935.clincRNA.NCBI.ref      ctgtgaccagacttgtatactggctgaatatcagtgctgtttgtaatttttcactttgag    11097
                                     * ** **** ** *  * * *  * *  ****  ***   ** *   *   ***   **

GGT5.NCBI.REF                       cctcaagggatcctcctgcttcagcctcctgagtagctgggactataagcacagggccac    24230
LOC105372935.clincRNA.NCBI.ref      aacca---------------------------------------------acattaattc    11112
                                       **                                             ***      *

GGT5.NCBI.REF                       cacatccgactaattttttcttttttcttttcttttatttttttaataaagacgtgatctt    24290
LOC105372935.clincRNA.NCBI.ref      catatgaatcaagtgttttgtaactgctattcatttattcagcaaatatttattgatcat    11172
                                    ** **    * * * **** *   * ** *** ******    **  *     ***** *

GGT5.NCBI.REF                       g----ctacattgcccaggctggtcttgaactcctgagctcaaaagatcctcctgcgccg    24346
LOC105372935.clincRNA.NCBI.ref      ctcttctccataagatagtgtgataaacacagtcatgaataaagttattttccacaaaa-    11231
                                         ** ***     **  ** *    *    *     * **   **  ***

GGT5.NCBI.REF                       gcctcccaaagtgctgggattacaggtgtgagccaccacgcccagcccaattttttgattt    24406
LOC105372935.clincRNA.NCBI.ref      ------------------------------------------------------------    11231

GGT5.NCBI.REF                       tcttttagagacggtgtcttgctttgttgccctggctgttctcaaactcctggcctcaag    24466
LOC105372935.clincRNA.NCBI.ref      ------------------------------------------------------------    11231

GGT5.NCBI.REF                       tgatcctgccaccttggctcccaagtggctaagactgtaggtatgtgccaccacgcctgg    24526
LOC105372935.clincRNA.NCBI.ref      ------------------------------------------------------------    11231

GGT5.NCBI.REF                       ctttttttatttcttatttttttatttttttagaaacaggatcttgctgcattgcccaggc    24586
LOC105372935.clincRNA.NCBI.ref      ------------------------------------------------------------    11231

GGT5.NCBI.REF                       tggtctcaaactcctggcctcaagtgatcctccttcctcagcctcccaaattgctggcat    24646
LOC105372935.clincRNA.NCBI.ref      ------------------------------------------------------------    11231

GGT5.NCBI.REF                       tacaggtgtgaaccactgtgtctggcctcagtgtctaaatcttgagaagccacatcctcc    24706
LOC105372935.clincRNA.NCBI.ref      ------------------------------------------------------------    11231

GGT5.NCBI.REF                       atcgaaggccccctgaactcagatttgcacctcaaggccctggtgtccttggatccagaa    24766
LOC105372935.clincRNA.NCBI.ref      ------------------------------------------------------------    11231

GGT5.NCBI.REF                       cagacatgaacttgcatttggaggtcttcgggcctgggtggtgtccttcgtacatagtcc    24826
LOC105372935.clincRNA.NCBI.ref      ------------------------------------------------------------    11231

GGT5.NCBI.REF                       ctggaggctgagcttgtcctctatgggcctttggatctcttccatgctcccatctggtcc    24886
LOC105372935.clincRNA.NCBI.ref      ------------------------------------------------------------    11231

GGT5.NCBI.REF                       tcccctcttcctgtgtcttcaggccctcttcctctccgtcttgccttttgatcatttttc    24946
LOC105372935.clincRNA.NCBI.ref      ------------------------------------------------------------    11231
```

```
GGT5.NCBI.REF                    tcatttcttgcagaggcctctccctcctggagatgggctgagctgccctcccctgtgaac    25006
LOC105372935.clincRNA.NCBI.ref   ------------------------------------------------------------    11231

GGT5.NCBI.REF                    ctgctttctctcctctccaggaggtgcagaggggactccaagaccgtggccagaaccaga    25066
LOC105372935.clincRNA.NCBI.ref   ------------------------------------------------------------    11231

GGT5.NCBI.REF                    cccagaggcccttcttcctgaacgtggtccaggctgtgtcccaggagggggcctgtgtgt    25126
LOC105372935.clincRNA.NCBI.ref   ------------------------------------------------------------    11231

GGT5.NCBI.REF                    acgccgtctcggacctgaggaagagtggggaggccgcaggctactaagacactgctctgc    25186
LOC105372935.clincRNA.NCBI.ref   ------------------------------------------------------------    11231

GGT5.NCBI.REF                    ccagagctgaagtctggccccaccatgagtcctgtgtccaggccggacatggctgggga     25246
LOC105372935.clincRNA.NCBI.ref   ------------------------------------------------------------    11231

GGT5.NCBI.REF                    ccaactactctggcaggatctggacccctggcaggggagtccagctgagagtggaagagg    25306
LOC105372935.clincRNA.NCBI.ref   ------------------------------------------------------------    11231

GGT5.NCBI.REF                    tggcggggaccagctgggcagatgagaggctgagcctcatccctaacccccttcccaga    25366
LOC105372935.clincRNA.NCBI.ref   ------------------------------------------------------------    11231

GGT5.NCBI.REF                    gcccctggtggtcctgaaccggcccctctatccctccgcaggcctcttgcctggggccac    25426
LOC105372935.clincRNA.NCBI.ref   ------------------------------------------------------------    11231

GGT5.NCBI.REF                    tctcccaccctctcgatctgtatatcctccagtccaagattaaagaggcggactgtggcc    25486
LOC105372935.clincRNA.NCBI.ref   ------------------------------------------------------------    11231

GGT5.NCBI.REF                    tga 25489
LOC105372935.clincRNA.NCBI.ref   --- 11231
```
