## Supplementary Figure S4 for "Formation of human long intergenic non-coding RNA genes and pseudogenes: ancestral sequences are key players"

Nicholas Delihas

**Supplementary Figure S4.** **a.** Color highlighted sections represent the *FAM230B--LOC105372935* --*GGT2* sequences that are found in *FAM230C-LOC101060145-GGT4P* with the respective percent identities. xxx represents sequences from the clincRNA region (*LOC105372935*) of *FAM230B-LOC105372935* -*GGT2* that are missing in FAM230C---*LOC101060145--GGT4P.* The unhighlighted section, I-----I represents the 5' half sequence of *FAM230C* that does not form part of *FAM230B.* **b.** Schematic of *FAM230B--LOC105372935--spacer--GGT2* for comparisons. The % identities shown in Figure S3a are relative to the *FAM230B-LOC105372935* -*GGT2* sequence.

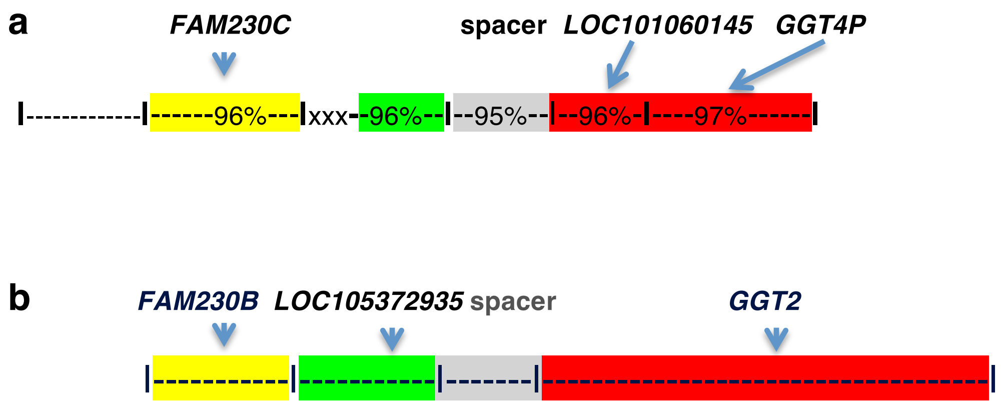
