## Supplementary Figure S5 for "Formation of human long intergenic non-coding RNA genes and pseudogenes: ancestral sequences are key players"

### A conserved primate sequence forms long non-coding RNA genes, pseudogenes and GGT-related protein genes

**Nicholas Delihas**

Supplementary Figure S5. Nt alignment of chimpanzee *LOC112206744-LOC107973052-GGT2* with human *FAM230B-LOC105372935GGT2.* The Clustal Omega, Multiple sequence alignment program was used for sequence alignment.

```
CLUSTAL O(1.2.4) multiple sequence alignment


chimp.LOC112206744.LOC107973052.GGT2-.646279-708040.rev.compl      ------------------------------------------------------------   0
FAM230B-LOC105372935-GGT2.NCBI.GGT2.rev.compl                      agcagttggacctcacagtgtggattgtgccttcaccctggaatgtttatgccctatcac   60
                                                                   

chimp.LOC112206744.LOC107973052.GGT2-.646279-708040.rev.compl      ------------------------------------------------------------   0
FAM230B-LOC105372935-GGT2.NCBI.GGT2.rev.compl                      catggtgatgggattagggatctcctgcccttggtcctaagtgccactatctgtgctgag   120
                                                                   

chimp.LOC112206744.LOC107973052.GGT2-.646279-708040.rev.compl      ------------------------------------------------------------   0
FAM230B-LOC105372935-GGT2.NCBI.GGT2.rev.compl                      tttttcaaaggtcagagcagattgaaccattgtggtttcattttccctgattttgatttt   180
                                                                   

chimp.LOC112206744.LOC107973052.GGT2-.646279-708040.rev.compl      ----atggggaacctgtgtggctgcattcaaggtatgttcatactggcctgtcaaatgcg   56
FAM230B-LOC105372935-GGT2.NCBI.GGT2.rev.compl                      tcttatggggaacctgtgtggctgcattcaaggtatgttcatactggcctgtcaaatgcg   240
                                                                       ********************************************************

chimp.LOC112206744.LOC107973052.GGT2-.646279-708040.rev.compl      atcttttcaaattactagttaatgctttcaaaatatgttatttaaaaaattatcctctgt   116
FAM230B-LOC105372935-GGT2.NCBI.GGT2.rev.compl                      atcttttcaaattactagttaatgctttcaaaatatgttatttaaaaaattagcctctgt   300
                                                                   **************************************************** *******

chimp.LOC112206744.LOC107973052.GGT2-.646279-708040.rev.compl      attttccatatggagttataaatatgtttcatggttatgttttattcctcaatttatata   176
FAM230B-LOC105372935-GGT2.NCBI.GGT2.rev.compl                      attttccatatgcagttataaatatgtttcatgattatgttttattcctcaatttatata   360
                                                                   ************ ******************** **************************

chimp.LOC112206744.LOC107973052.GGT2-.646279-708040.rev.compl      tttgattattgtaccaagcagagtacctttgaaatttttcttcatttaaaaaatatgtct   236
FAM230B-LOC105372935-GGT2.NCBI.GGT2.rev.compl                      tttgattattgtaccaagcagagtatctttgaaatttttcttcatttaaaaaatatgtat   420
                                                                   ************************* ******************************** *

chimp.LOC112206744.LOC107973052.GGT2-.646279-708040.rev.compl      cttggctcaggcctgtaatcccagcactttgggaggccaatgcaagaggatcacaaggtg   296
FAM230B-LOC105372935-GGT2.NCBI.GGT2.rev.compl                      cttgactcaggcctgtaatcccagcactttgggaggccaaggcaagaggatcacaaggtg   480
                                                                   **** *********************************** *******************

chimp.LOC112206744.LOC107973052.GGT2-.646279-708040.rev.compl      aggagatcaagaccatcctggtcaatacagtgaaaccctgtctgtacgaaaagtacaaaa   356
FAM230B-LOC105372935-GGT2.NCBI.GGT2.rev.compl                      aggagatcaagaccatcctggccaatacagtgaaaccctgtctctactacaaatacaaaa   540
                                                                   ********************* ********************* *** * ** *******

chimp.LOC112206744.LOC107973052.GGT2-.646279-708040.rev.compl      aattagccaggaatggtggcagctggtgtagtcccagtgtgaattgggattcagtttatt   416
FAM230B-LOC105372935-GGT2.NCBI.GGT2.rev.compl                      atttagccaggcatggtggcagctggtgtagtcccagtgtgaattgggattcagtttatt   600
                                                                   * ********* ************************************************

chimp.LOC112206744.LOC107973052.GGT2-.646279-708040.rev.compl      cccaaattcccaaattatatatat-tatatatataatatattatatatacagtatatata   475
FAM230B-LOC105372935-GGT2.NCBI.GGT2.rev.compl                      cccaaattcccaaattatatatatatatacatataaaata--------------------   640
                                                                   ************************ **** ****** ***

chimp.LOC112206744.LOC107973052.GGT2-.646279-708040.rev.compl      atatattatatatacagtatataatatactatgtatacagtatatattatatactgtatt   535
FAM230B-LOC105372935-GGT2.NCBI.GGT2.rev.compl                      ------------------------------------------------------------   640
                                                                   

chimp.LOC112206744.LOC107973052.GGT2-.646279-708040.rev.compl      atatacattatatactgtatacattatatactgtattatatattatatactgtatatata   595
FAM230B-LOC105372935-GGT2.NCBI.GGT2.rev.compl                      ------------------------------------------------------------   640
                                                                   

chimp.LOC112206744.LOC107973052.GGT2-.646279-708040.rev.compl      atatatattatatatacagtatatagtacatattatatatactatatgttatataaaata   655
FAM230B-LOC105372935-GGT2.NCBI.GGT2.rev.compl                      ------------------------------------------------------------   640
                                                                   

chimp.LOC112206744.LOC107973052.GGT2-.646279-708040.rev.compl      tataatataaaatatataatatataatatataatatataatatataatatgtataatata   715
FAM230B-LOC105372935-GGT2.NCBI.GGT2.rev.compl                      ------------------------------------------------------------   640
                                                                   

chimp.LOC112206744.LOC107973052.GGT2-.646279-708040.rev.compl      taatatataatatatagtatatatatcgtataatatatataatatatattatatataata   775
FAM230B-LOC105372935-GGT2.NCBI.GGT2.rev.compl                      ------------------------------------------------------------   640
                                                                   

chimp.LOC112206744.LOC107973052.GGT2-.646279-708040.rev.compl      tatattataaattataatttataatttatatatattatatatataataataaattataat   835
FAM230B-LOC105372935-GGT2.NCBI.GGT2.rev.compl                      ------------------------------------------------------------   640
                                                                   

chimp.LOC112206744.LOC107973052.GGT2-.646279-708040.rev.compl      tataaattataatttataatatatattatatataatatatattatatattataatttata   895
FAM230B-LOC105372935-GGT2.NCBI.GGT2.rev.compl                      ------------------------------------------------------------   640
                                                                   

chimp.LOC112206744.LOC107973052.GGT2-.646279-708040.rev.compl      atatacattatttatatataatatataatgtatattatatattatatatttaatatataa   955
FAM230B-LOC105372935-GGT2.NCBI.GGT2.rev.compl                      ------------------------------------------------------------   640
                                                                   

chimp.LOC112206744.LOC107973052.GGT2-.646279-708040.rev.compl      tatgttatatataatatataatatatattgtatataatatatattatattatatattata   1015
FAM230B-LOC105372935-GGT2.NCBI.GGT2.rev.compl                      ---catatataatatattaaatatattatatatataatatatattatattatatataatt   697
                                                                        ******  ****  *******  * ************************** **

chimp.LOC112206744.LOC107973052.GGT2-.646279-708040.rev.compl      atacataatataatatataatatattatatattatatatattatataatatattatatat   1075
FAM230B-LOC105372935-GGT2.NCBI.GGT2.rev.compl                      atatata-----------------------------------------------------   704
                                                                   *** ***

chimp.LOC112206744.LOC107973052.GGT2-.646279-708040.rev.compl      tatatatatatataatatataatatattatatatatgttatatattatatatatatgtta   1135
FAM230B-LOC105372935-GGT2.NCBI.GGT2.rev.compl                      ------------------------------------------------------------   704
                                                                   

chimp.LOC112206744.LOC107973052.GGT2-.646279-708040.rev.compl      tatattatagaatatgttatatattatatattatagaatatgttatatattatatattat   1195
FAM230B-LOC105372935-GGT2.NCBI.GGT2.rev.compl                      ------------------------------------------------------------   704
                                                                   

chimp.LOC112206744.LOC107973052.GGT2-.646279-708040.rev.compl      ataatatgttatatattatatattatagaatatgttatatattatataatatgttatata   1255
FAM230B-LOC105372935-GGT2.NCBI.GGT2.rev.compl                      ----------------tatatatttgtgggtgccctatttcccgtctcataacttatttt   748
                                                                                   ********   *  *    *** *    * * ***  **** *

chimp.LOC112206744.LOC107973052.GGT2-.646279-708040.rev.compl      ttatatattatataatatgttat-------------------------------------   1278
FAM230B-LOC105372935-GGT2.NCBI.GGT2.rev.compl                      aagaagccagcataataatgtgtgggcttgggattcagtttttgaaacaaaacactgagc   808
                                                                       *      ******   * *

chimp.LOC112206744.LOC107973052.GGT2-.646279-708040.rev.compl      ------------------------------------------------------------   1278
FAM230B-LOC105372935-GGT2.NCBI.GGT2.rev.compl                      ctttgatgaccttcctgtacttgtaaaagcccacctgtctgcatggcagcagttggacct   868
                                                                   

chimp.LOC112206744.LOC107973052.GGT2-.646279-708040.rev.compl      ---------------------------------------------------------ata   1281
FAM230B-LOC105372935-GGT2.NCBI.GGT2.rev.compl                      cacagtgtggattgtgccttcaccctggaatgtttatgccctatcgccatggtgatggga   928
                                                                                                                              *

chimp.LOC112206744.LOC107973052.GGT2-.646279-708040.rev.compl      ttatatatattatataatatgttatatattatataatatgttatatattatatatattat   1341
FAM230B-LOC105372935-GGT2.NCBI.GGT2.rev.compl                      ttagggatctcctgcccttggtcctaagtgccactatctgtgctgagtttttcaaaggtc   988
                                                                   ***   ** *  *    *  **  **  *   *  ** ***  *   ** *  * *

chimp.LOC112206744.LOC107973052.GGT2-.646279-708040.rev.compl      ataatatgttatatattatatatattatataatatgttatatattatatattatataata   1401
FAM230B-LOC105372935-GGT2.NCBI.GGT2.rev.compl                      agagcagattgaaccattgtggtttcattttccctgattttgatttt-----------tc   1037
                                                                   * *  *  **  *   *     * * ** *    ** * *  *** *           *

chimp.LOC112206744.LOC107973052.GGT2-.646279-708040.rev.compl      tgttatatattatatatattatataatatgttatatattatatattatataatatgttat   1461
FAM230B-LOC105372935-GGT2.NCBI.GGT2.rev.compl                      ttatggggaacctgtgtggctgcattcaaggtatgttcatactggcctgtcaaatgcgat   1097
                                                                   *  *    *   * * *          * * *** *      *    * * * ***  **

chimp.LOC112206744.LOC107973052.GGT2-.646279-708040.rev.compl      atattatatatattatataatatgtt--atatattatatattatataatatgttatatat   1519
FAM230B-LOC105372935-GGT2.NCBI.GGT2.rev.compl                      cttttcaaattactagttaatgctttcaaaatatgttatttaaaaaattagcctctgtat   1157
                                                                    * **  *  ** **  ****   **  * ****  *** * * * * **   * * ***

chimp.LOC112206744.LOC107973052.GGT2-.646279-708040.rev.compl      tatatatattatataatatatatt--------atatcttatataatataatttatatatt   1571
FAM230B-LOC105372935-GGT2.NCBI.GGT2.rev.compl                      tttccatatgcagttataaatatgtttcatgattatgttttattcctcaatttatatatt   1217
                                                                   * *  ****    * *** ****          *** ** ***     ************

chimp.LOC112206744.LOC107973052.GGT2-.646279-708040.rev.compl      ata-tattatacaatatatattatatattatattatatattatacaatatatat------   1624
FAM230B-LOC105372935-GGT2.NCBI.GGT2.rev.compl                      tgattattgtaccaagcagagtatctttgaaatttttcttcatttaaaaaatatgtatct   1277
                                                                     * **** *** *   * * *** * * * *** *   * **  ** * ****

chimp.LOC112206744.LOC107973052.GGT2-.646279-708040.rev.compl      ------------------------------------------------------------   1624
```

```
FAM230B-LOC105372935-GGT2.NCBI.GGT2.rev.compl                tgactcaggcctgtaatcccagcactttgggaggccaaggcaagaggatcacaaggtgag   1337

chimp.LOC112206744.LOC107973052.GGT2-.646279-708040.rev.compl   ------------------------------------------------------------   1624
FAM230B-LOC105372935-GGT2.NCBI.GGT2.rev.compl                gagatcaagaccatcctggccaatacagtgaaaccctgtctctactacaaatacaaaaat   1397

chimp.LOC112206744.LOC107973052.GGT2-.646279-708040.rev.compl   ------------------------tatatatatattatatattatacaatatatattata   1660
FAM230B-LOC105372935-GGT2.NCBI.GGT2.rev.compl                ttagccaggcatggtggcagctggtgtagtcccagtgtgaattgggattcagtttattca   1457
                                                                                     * **     * * *  ***       *  *  *  *

chimp.LOC112206744.LOC107973052.GGT2-.646279-708040.rev.compl   taatatataatatatattatattatataatatatatataatttccttttacatgctgcat   1720
FAM230B-LOC105372935-GGT2.NCBI.GGT2.rev.compl                gtttattcccaaattcccaaattatat-atatatatataatttccttttacatcctgcat   1516
                                                                   ***     *  *   * ******* ************************* ******

chimp.LOC112206744.LOC107973052.GGT2-.646279-708040.rev.compl   ccttcaacattccatcccccaccccacagattaagttattccccaggggagaatatggca   1780
FAM230B-LOC105372935-GGT2.NCBI.GGT2.rev.compl                ccttcaacgttacatcccccaccccacagattaagttattccccaggggagaatatggca   1576
                                                             ******** ** ************************************************

chimp.LOC112206744.LOC107973052.GGT2-.646279-708040.rev.compl   aactctattttaatgcagtttttagcctaattaagaa-ttatgaaatcattactttccaa   1839
FAM230B-LOC105372935-GGT2.NCBI.GGT2.rev.compl                aagtctattttaatgcagtttttaacccaattaagaacctatgaaatcattactttccaa   1636
                                                             ** ********************* ** *********  *********************

chimp.LOC112206744.LOC107973052.GGT2-.646279-708040.rev.compl   atatttggaacaaagccacagtagtatgcatgggttggaggcttttcacagaataaaatg   1899
FAM230B-LOC105372935-GGT2.NCBI.GGT2.rev.compl                aactttggaacaaagccacagtagtatggatccgttggaggcttttcacacaataaaatg   1696
                                                             *  ************************* **  ***************** *********

chimp.LOC112206744.LOC107973052.GGT2-.646279-708040.rev.compl   tacctatctttgtttttaacatgtttttcccttcctctcttctttctttgtgaaatgtgt   1959
FAM230B-LOC105372935-GGT2.NCBI.GGT2.rev.compl                tacctctctttgtttttaacatgtttttcccttcctctcttctttttttgtgaaatgtgt   1756
                                                             ***** *************************************** **************

chimp.LOC112206744.LOC107973052.GGT2-.646279-708040.rev.compl   atttactttaataaatttgtagtaagtcacttccattcacatattaattttttaaagtaa   2019
FAM230B-LOC105372935-GGT2.NCBI.GGT2.rev.compl                atttactttaatatatttgtagtaagtcacttccatgcacatattaattttttaaagtaa   1816
                                                             ************* ********************** ***********************

chimp.LOC112206744.LOC107973052.GGT2-.646279-708040.rev.compl   taagtatgtgtattgtctactagtgaaataaaacacacatttatttttattctttggaag   2079
FAM230B-LOC105372935-GGT2.NCBI.GGT2.rev.compl                taagtatgtgtattgtctacgtgtgaaagaaaacacacatttatttttatgctttggaag   1876
                                                             ********************  ****** ********************* *********

chimp.LOC112206744.LOC107973052.GGT2-.646279-708040.rev.compl   ttatccagaatcatggcattgtcaatcacagtcaatcacccaacctactcacctttccag   2139
FAM230B-LOC105372935-GGT2.NCBI.GGT2.rev.compl                ttatccagaatcatggaattgtcaatcacagtcaatcacccaacctactcacctttccag   1936
                                                             **************** *******************************************

chimp.LOC112206744.LOC107973052.GGT2-.646279-708040.rev.compl   tgtaatcttagtctaattttttttttgttatccaatgagatgcagtatttcaactcagag   2199
FAM230B-LOC105372935-GGT2.NCBI.GGT2.rev.compl                tgtaatcttagtcaaattttttttttgttatccaatgagatgcagtatttcaactcagaa   1996
                                                             ************* ********************************************* 

chimp.LOC112206744.LOC107973052.GGT2-.646279-708040.rev.compl   agataaatagaatgaattggtagagactattaactaagaatatacagttttatttatgct   2259
FAM230B-LOC105372935-GGT2.NCBI.GGT2.rev.compl                agataaatagagtgaatttatagagactattaactaagaacatacagttttatttatact   2056
                                                             *********** ******  ******************** **************** **

chimp.LOC112206744.LOC107973052.GGT2-.646279-708040.rev.compl   cagaagcaagtagattatgtatatatgtatgaagataaaaaattaaaggataattgtgta   2319
FAM230B-LOC105372935-GGT2.NCBI.GGT2.rev.compl                cagaagcaagtagattatgtacatatatatgaagataaaaaattaaaggataattgtgta   2116
                                                             ********************* **** *********************************

chimp.LOC112206744.LOC107973052.GGT2-.646279-708040.rev.compl   aatttgcatgtagagagctttgaaaacctgtttacttgttaatgctgttttgatgtattg   2379
FAM230B-LOC105372935-GGT2.NCBI.GGT2.rev.compl                aatttgcatgtagagggctttgaaaacctgtttacttgttaatgctgttttgatgtat--   2174
                                                             *************** ******************************************  

chimp.LOC112206744.LOC107973052.GGT2-.646279-708040.rev.compl   tgtgtctttgttctcccgacccagcatccagagctctctgctggagctaagtgctcatca   2439
FAM230B-LOC105372935-GGT2.NCBI.GGT2.rev.compl                tgtgtctttgttctcccgacccatcgtccagagctctctgcaggagctaagtgctcatca   2234
                                                             *********************** * *************** ******************

chimp.LOC112206744.LOC107973052.GGT2-.646279-708040.rev.compl   gttccatgacttggaaactgtctaagtttagaggcacttgtatttgttagtaaataagtc   2499
FAM230B-LOC105372935-GGT2.NCBI.GGT2.rev.compl                gttccatgacttggaaactgtctaagtttagaggcacttgtatttgttagtaaataaggc   2294
                                                             ********************************************************** *

chimp.LOC112206744.LOC107973052.GGT2-.646279-708040.rev.compl   aagatgatattgtttcacaggctttagtgcagaagactgaatagagaagctgctccaccc   2559
FAM230B-LOC105372935-GGT2.NCBI.GGT2.rev.compl                aagatgatattgtttcacaggttttagtgccgaagactgaatagataagctgctccaccc   2354
                                                             ********************* ******** ************** **************

chimp.LOC112206744.LOC107973052.GGT2-.646279-708040.rev.compl   agtacactggtgttcatttcatggtcatctcatctgttaaccatggatataaaacactta   2619
FAM230B-LOC105372935-GGT2.NCBI.GGT2.rev.compl                agtacactggtgttcatttcatggtcatctcatctgttaaccatggatataaaacattta   2414
                                                             ******************************************************** ***

chimp.LOC112206744.LOC107973052.GGT2-.646279-708040.rev.compl   tcttcaatgatcttcctgtacatgtaaaaacacacctgtctgcatggcagcagttggacc   2679
FAM230B-LOC105372935-GGT2.NCBI.GGT2.rev.compl                tcttcaat----------------------------------------------------   2422
                                                             ********

chimp.LOC112206744.LOC107973052.GGT2-.646279-708040.rev.compl   tcacagtgtggattgtgccttcaccctggaatgtctatgatgccctatcgaccatggtga   2739
FAM230B-LOC105372935-GGT2.NCBI.GGT2.rev.compl                ------------------------------------------------------------   2422

chimp.LOC112206744.LOC107973052.GGT2-.646279-708040.rev.compl   tgggattagggatctcttcccctttgctcggccactgtctgtgccagccaggccactggg   2799
FAM230B-LOC105372935-GGT2.NCBI.GGT2.rev.compl                ------------------------------------------------------------   2422

chimp.LOC112206744.LOC107973052.GGT2-.646279-708040.rev.compl   ccattgtggcggatggtgatgccctttgtgtggagcgcactgagtgtgcgctgaggcaga   2859
FAM230B-LOC105372935-GGT2.NCBI.GGT2.rev.compl                ------------------------------------------------------------   2422

chimp.LOC112206744.LOC107973052.GGT2-.646279-708040.rev.compl   cacagtacttttgacaactttatctgcttctttcaaaactgggtctgcattatgataaag   2919
FAM230B-LOC105372935-GGT2.NCBI.GGT2.rev.compl                ------------------------------------------------------------   2422

chimp.LOC112206744.LOC107973052.GGT2-.646279-708040.rev.compl   tggtcggttttaggccaagcaggtgtatgtcatagagcaatgacttgggagttcagaatc   2979
FAM230B-LOC105372935-GGT2.NCBI.GGT2.rev.compl                ------------------------------------------------------------   2422

chimp.LOC112206744.LOC107973052.GGT2-.646279-708040.rev.compl   ctgggcgtgaataagcctctgagccttattaagctgtgaaatctaaagcaagacatttca   3039
FAM230B-LOC105372935-GGT2.NCBI.GGT2.rev.compl                ------------------------------------------------------------   2422

chimp.LOC112206744.LOC107973052.GGT2-.646279-708040.rev.compl   ctgataattaagcctcatgagcacttatactgtacactttacctacatcaaatcattcgg   3099
FAM230B-LOC105372935-GGT2.NCBI.GGT2.rev.compl                ------------------------------------------------------------   2422

chimp.LOC112206744.LOC107973052.GGT2-.646279-708040.rev.compl   ccatcaccacaaacaaatggcatgattgatttactatcacctttcgctgggtagaaaaag   3159
FAM230B-LOC105372935-GGT2.NCBI.GGT2.rev.compl                ------------------------------------------------------------   2422

chimp.LOC112206744.LOC107973052.GGT2-.646279-708040.rev.compl   acagagatgaatgtgcccaagcttatgagcccctcagcaaaagagccaagatgggaaccc   3219
FAM230B-LOC105372935-GGT2.NCBI.GGT2.rev.compl                ------------------------------------------------------------   2422

chimp.LOC112206744.LOC107973052.GGT2-.646279-708040.rev.compl   aagcatacagccccaatgctgaggctctgaactactgacctgccctcagcactcagcctt   3279
FAM230B-LOC105372935-GGT2.NCBI.GGT2.rev.compl                ------------------------------------------------------------   2422

chimp.LOC112206744.LOC107973052.GGT2-.646279-708040.rev.compl   gggatcatgagtcactgtgcaagggagttccaacatctgcatgtatgtctggaatgatct   3339
FAM230B-LOC105372935-GGT2.NCBI.GGT2.rev.compl                ------------------------------------------------------------   2422

chimp.LOC112206744.LOC107973052.GGT2-.646279-708040.rev.compl   gagcctgtagagttcctacacactggccacattatagggtggtgtctgtggtcacacagc   3399
FAM230B-LOC105372935-GGT2.NCBI.GGT2.rev.compl                ------------------------------------------------------------   2422

chimp.LOC112206744.LOC107973052.GGT2-.646279-708040.rev.compl   tcagggcaggtattagtacatgaatagcttagatgtgtcatagtctttatgtgaaaggca   3459
FAM230B-LOC105372935-GGT2.NCBI.GGT2.rev.compl                ------------------------------------------------------------   2422

chimp.LOC112206744.LOC107973052.GGT2-.646279-708040.rev.compl   cataaaagcccaaatgcttttgaactggttgcggattttgagttggaggacaacacttgg   3519
FAM230B-LOC105372935-GGT2.NCBI.GGT2.rev.compl                ------------------------------------------------------------   2422

chimp.LOC112206744.LOC107973052.GGT2-.646279-708040.rev.compl   gacaaccggtcacattccttcagtgcgcggccactcctcagcacatatcccaaagccggt   3579
FAM230B-LOC105372935-GGT2.NCBI.GGT2.rev.compl                ------------------------------------------------------------   2422

chimp.LOC112206744.LOC107973052.GGT2-.646279-708040.rev.compl   aaataatgggaagaccctgtactgatgtttgatggtgttatgaacttcaaatgacccta   3639
FAM230B-LOC105372935-GGT2.NCBI.GGT2.rev.compl                ------------------------------------------------------------   2422

chimp.LOC112206744.LOC107973052.GGT2-.646279-708040.rev.compl   tagaaatgttaaaaaaaaaaaaaaaaaaaaagtctgtgagctacacaacctggggacaag   3699
FAM230B-LOC105372935-GGT2.NCBI.GGT2.rev.compl                ------------------------------------------------------------   2422
```

```
chimp.LOC112206744.LOC107973052.GGT2-.646279-708040.rev.compl      acaagtaagaacacaaaataccagtaaataatgagcctcatgtgtcaccctgcttgcaac   3759
FAM230B-LOC105372935-GGT2.NCBI.GGT2.rev.compl                      ------------------------------------------------------------   2422

chimp.LOC112206744.LOC107973052.GGT2-.646279-708040.rev.compl      tccctcggcagctggtttgaagagcctcagtgctcaccaggggccatctccccgctctgc   3819
FAM230B-LOC105372935-GGT2.NCBI.GGT2.rev.compl                      ------------------------------------------------------------   2422

chimp.LOC112206744.LOC107973052.GGT2-.646279-708040.rev.compl      cctctctcgctgtgggttgcctgggctgaacataggcagcccctcacatgatgactgcaa   3879
FAM230B-LOC105372935-GGT2.NCBI.GGT2.rev.compl                      ------------------------------------------------------------   2422

chimp.LOC112206744.LOC107973052.GGT2-.646279-708040.rev.compl      acacagggcaaaggaagactgaacggaaaatgtatggttatagtggttattttaaaatat   3939
FAM230B-LOC105372935-GGT2.NCBI.GGT2.rev.compl                      ------------------------------------------------------------   2422

chimp.LOC112206744.LOC107973052.GGT2-.646279-708040.rev.compl      aagtatttcagtattttctaatctttaataattgattattttaggggaggtttttttt   3999
FAM230B-LOC105372935-GGT2.NCBI.GGT2.rev.compl                      ------------------------------------------------------------   2422

chimp.LOC112206744.LOC107973052.GGT2-.646279-708040.rev.compl      tttttttttgagacagagtcttgcgctgttgcccaggctggagtgcagtggcatgatctc   4059
FAM230B-LOC105372935-GGT2.NCBI.GGT2.rev.compl                      ------------------------------------------------------------   2422

chimp.LOC112206744.LOC107973052.GGT2-.646279-708040.rev.compl      agctcactgcaacctctacctcctgtgttcaagcgattctcctgcctcagcctcccgagt   4119
FAM230B-LOC105372935-GGT2.NCBI.GGT2.rev.compl                      ------------------------------------------------------------   2422

chimp.LOC112206744.LOC107973052.GGT2-.646279-708040.rev.compl      agctgggactacaggcacgtgccaccacgcctggcaaagtttttgtgtttttagtagaga   4179
FAM230B-LOC105372935-GGT2.NCBI.GGT2.rev.compl                      ------------------------------------------------------------   2422

chimp.LOC112206744.LOC107973052.GGT2-.646279-708040.rev.compl      cggggtttcactctgttagccaggatggtctcaatcttctgacctcatgatccacccgcc   4239
FAM230B-LOC105372935-GGT2.NCBI.GGT2.rev.compl                      ------------------------------------------------------------   2422

chimp.LOC112206744.LOC107973052.GGT2-.646279-708040.rev.compl      tcggcctcccaaagtgctgggattacaggtgtgagccaccacacctggcctgatttttta   4299
FAM230B-LOC105372935-GGT2.NCBI.GGT2.rev.compl                      ------------------------------------------------------------   2422

chimp.LOC112206744.LOC107973052.GGT2-.646279-708040.rev.compl      ttttttatttttttgagatgaagtctctgtcacccaggctggagtgcagtggtgcaatct   4359
FAM230B-LOC105372935-GGT2.NCBI.GGT2.rev.compl                      ------------------------------------------------------------   2422

chimp.LOC112206744.LOC107973052.GGT2-.646279-708040.rev.compl      cagctcactgcaatctctgcctccaggttcaagggattgtcctgcctcagcctcctgagt   4419
FAM230B-LOC105372935-GGT2.NCBI.GGT2.rev.compl                      ------------------------------------------------------------   2422

chimp.LOC112206744.LOC107973052.GGT2-.646279-708040.rev.compl      agctgggttttttttttttttagttgagatggggttttaccatgttggtcaggctggtct   4479
FAM230B-LOC105372935-GGT2.NCBI.GGT2.rev.compl                      ---------------------------gatggggttttaccatgttggtcaggctggtct   2455
                                                                                              *********************************

chimp.LOC112206744.LOC107973052.GGT2-.646279-708040.rev.compl      cgaactcctgacctcaaatgatccgcccacctccaccttccaaactgctgggattacagg   4539
FAM230B-LOC105372935-GGT2.NCBI.GGT2.rev.compl                      cgaactcctgacctcaaatgatccacccacctccaccttccaaactgctgggattacagg   2515
                                                                   ************************ ***********************************

chimp.LOC112206744.LOC107973052.GGT2-.646279-708040.rev.compl      tgtgagccactatgcctggctgattattttcataaccaagaaaagaaataaatataatta   4599
FAM230B-LOC105372935-GGT2.NCBI.GGT2.rev.compl                      tgtgagccactatgcctgactgattattttcataaccaagaaaagaaataaatacaatta   2575
                                                                   ****************** *********************************** *****

chimp.LOC112206744.LOC107973052.GGT2-.646279-708040.rev.compl      atgctggtgcatggtattaaatctagtttttaaaaaattcacacataaaccaggcagaac   4659
FAM230B-LOC105372935-GGT2.NCBI.GGT2.rev.compl                      atgctggtgcatggtattaaatctagtttttaaaaaattcacacataaacaaggcagaac   2635
                                                                   ************************************************** *********

chimp.LOC112206744.LOC107973052.GGT2-.646279-708040.rev.compl      cctataccctccatgataaatgcagtagcagtgtatgtgggtctgtggaggttgaaaggg   4719
FAM230B-LOC105372935-GGT2.NCBI.GGT2.rev.compl                      cctataccctccatgataaatgcagtagcagtgtatgtgggtctgtggaggttgaaaggg   2695
                                                                   ************************************************************

chimp.LOC112206744.LOC107973052.GGT2-.646279-708040.rev.compl      acttggtagatgtcaagaaggtagtggcagtcctgctgggcttttaaagggtctgaagaa   4779
FAM230B-LOC105372935-GGT2.NCBI.GGT2.rev.compl                      acttggtagatgtcaagaagctagtggcagtcctgctgggcttttaaagggtctgaagaa   2755
                                                                   ******************** ***************************************

chimp.LOC112206744.LOC107973052.GGT2-.646279-708040.rev.compl      gtgacaggatgctgtggttgaatcctagcatgtattttagcatttgttcatttggagttt   4839
FAM230B-LOC105372935-GGT2.NCBI.GGT2.rev.compl                      gtgacaggatgctgtggttgaatcctagcatgtattttagcatttgttcatttggag-tt   2814
                                                                   ********************************************************* **

chimp.LOC112206744.LOC107973052.GGT2-.646279-708040.rev.compl      gattatttcacgttgctttcatttgccattacctggaaagccaagggctctactctcatt   4899
FAM230B-LOC105372935-GGT2.NCBI.GGT2.rev.compl                      gattatttcacgttgctttcatttgccattacctggaaagccaagggctctactctcatt   2874
                                                                   ************************************************************

chimp.LOC112206744.LOC107973052.GGT2-.646279-708040.rev.compl      tccttgctgctctttctttgccttccttggtccatgaagaagatggtccaggagaagctc   4959
FAM230B-LOC105372935-GGT2.NCBI.GGT2.rev.compl                      tccttgctgctctttctttgccttccttggtccgtgaagaagatggtccaggagaagctc   2934
                                                                   ********************************* **************************

chimp.LOC112206744.LOC107973052.GGT2-.646279-708040.rev.compl      attccatgcttgttaaccaggcacacccctaagttccagtccctgagtcattcatgagta   5019
FAM230B-LOC105372935-GGT2.NCBI.GGT2.rev.compl                      attccatgcttgttaaccaggcacgcccctaagttccagtccctgagtcattcatgagta   2994
                                                                   *********************** ************************************

chimp.LOC112206744.LOC107973052.GGT2-.646279-708040.rev.compl      gcactgccaatgaactgacagccatgctgtgtccctccacatcccctaggtgactcgaag   5079
FAM230B-LOC105372935-GGT2.NCBI.GGT2.rev.compl                      gcactgccaatgaactgacagccatgctgtgtccctccacatcccctaggtgactcgaag   3054
                                                                   ************************************************************

chimp.LOC112206744.LOC107973052.GGT2-.646279-708040.rev.compl      aagccttccaaaaagcatgtgaaaagggagccctcctctactaccaaggtaaagtagcct   5139
FAM230B-LOC105372935-GGT2.NCBI.GGT2.rev.compl                      aagccttccaaaaagcatgtgaaaaggaagccctactctactaccaaggtaaagtagcct   3114
                                                                   *************************** ****** *************************

chimp.LOC112206744.LOC107973052.GGT2-.646279-708040.rev.compl      gtctttgcctaagatgtaaatgttgttttcttggatcctttatttttcagttgatatctg   5199
FAM230B-LOC105372935-GGT2.NCBI.GGT2.rev.compl                      gtctttgcctaagatgtaaatgttgttttcttggatcctttatttttcagttgatatcag   3174
                                                                   ********************************************************** *

chimp.LOC112206744.LOC107973052.GGT2-.646279-708040.rev.compl      ctatgggaaaattatccactacattatagatgttagataatatttccttggggatggagg   5259
FAM230B-LOC105372935-GGT2.NCBI.GGT2.rev.compl                      ctatgggaaaattctccactacattataggtgttagataatatttccttggggatggagg   3234
                                                                   ************* *************** ******************************

chimp.LOC112206744.LOC107973052.GGT2-.646279-708040.rev.compl      aggtgtattttaccaactgacacctgattccagaggatgtgcaaaattggcagtgtcaga   5319
FAM230B-LOC105372935-GGT2.NCBI.GGT2.rev.compl                      aggtgtattttaccaactgacacctgattccagaggacgtgcaaaattggcagtgtcaga   3294
                                                                   ************************************ ***********************

chimp.LOC112206744.LOC107973052.GGT2-.646279-708040.rev.compl      tagtacactgcgtgttaagggatgttttcttcaggaacaagctttccactttagataaga   5379
FAM230B-LOC105372935-GGT2.NCBI.GGT2.rev.compl                      tagtacactgggtgttaagggatgttttcttcaggaacaagctttccactttagataaga   3354
                                                                   ********** *************************************************

chimp.LOC112206744.LOC107973052.GGT2-.646279-708040.rev.compl      attctgcaattgctactcaaaaattacctagacagaaacattcttcaagaaaagctcctg   5439
FAM230B-LOC105372935-GGT2.NCBI.GGT2.rev.compl                      attctgcaattgctactcaaaaattacctagacagaaacattcttcaagaaaagctcctg   3414
                                                                   ************************************************************

chimp.LOC112206744.LOC107973052.GGT2-.646279-708040.rev.compl      tgctttcctaagggaactctactctagagttggggcttttgacttgaaccttatttccaa   5499
FAM230B-LOC105372935-GGT2.NCBI.GGT2.rev.compl                      tgctttcctaagggaactctactctagagttggggcttttgacttgaaccttatttccag   3474
                                                                   ***********************************************************

chimp.LOC112206744.LOC107973052.GGT2-.646279-708040.rev.compl      tcttggttacccagagtttccaagtgaacaaaagacctgtgtgagccatccatagcatag   5559
FAM230B-LOC105372935-GGT2.NCBI.GGT2.rev.compl                      tcttggttacccagagtttccaagtgaacaaaagacctgtgtgagccatccatagcatag   3534
                                                                   ************************************************************

chimp.LOC112206744.LOC107973052.GGT2-.646279-708040.rev.compl      cctgattctcagagtgttttccttctctaattacaggtgacttcagggagcacattcaat   5619
FAM230B-LOC105372935-GGT2.NCBI.GGT2.rev.compl                      cctgattctcagagtgttttccttctctaattacaggtgacttcagggagcacattcaat   3594
                                                                   ************************************************************

chimp.LOC112206744.LOC107973052.GGT2-.646279-708040.rev.compl      ggtacgtattctggaatcactcactggttgttagaaaaggattctataggaaatctggag   5679
FAM230B-LOC105372935-GGT2.NCBI.GGT2.rev.compl                      ggtacgtattctggaatcactcactggttgttagaaaaggattctacaggaaatctggag   3654
                                                                   ********************************************** *************

chimp.LOC112206744.LOC107973052.GGT2-.646279-708040.rev.compl      cttaactgctggcttttgtctagagaggctccatgatccaagacatctggtgggaatgag   5739
FAM230B-LOC105372935-GGT2.NCBI.GGT2.rev.compl                      cttaactgctggcttttgtctggagagcctccatgatccaagacatctggtgggaatgag   3714
                                                                   ********************* ***** ********************************

chimp.LOC112206744.LOC107973052.GGT2-.646279-708040.rev.compl      gatgtgggatatagtaaaggaaactggttttccaggtgacatactctttttatctatgta   5799
FAM230B-LOC105372935-GGT2.NCBI.GGT2.rev.compl                      gatgtagggtatagtaaaagaaactggttttcctggtgacatactctttttatctatgta   3774
                                                                   ***** ** ********* ************** **************************

chimp.LOC112206744.LOC107973052.GGT2-.646279-708040.rev.compl      tagtttctgggaatgtgttcacattaggttgtgtgtgggtatgtgtgtattagggcgggg   5859
FAM230B-LOC105372935-GGT2.NCBI.GGT2.rev.compl                      tagtttctgggaacatgttcacattaggttgtgtgtgggtatgtgtgtattagggcgggg   3834
                                                                   *************  *********************************************
```

```
chimp.LOC112206744.LOC107973052.GGT2-.646279-708040.rev.compl       gtgggatgaggtggtctgtgtgcacatctgcatgatttgcttgtgaatgtgtgtctatgt   5919
FAM230B-LOC105372935-GGT2.NCBI.GGT2.rev.compl                       gtggggtgaggtggtctgtgtgcaagtctgcatgatttgcttgtgaatgtgtgtctatgt   3894
                                                                    ***** ******************  **********************************

chimp.LOC112206744.LOC107973052.GGT2-.646279-708040.rev.compl       gtgtttcccccaggaaaaaaatgttgtgtttacccagcacaactctcagtgccatttttc   5979
FAM230B-LOC105372935-GGT2.NCBI.GGT2.rev.compl                       gtgtttcccctaggaaaaaaatgttgtgtttacccagcacaactctcagtgccatttttc   3954
                                                                    ********** *************************************************

chimp.LOC112206744.LOC107973052.GGT2-.646279-708040.rev.compl       ttaatttaacaaatcagaccacatactttacttacattagttcacacctcatcatcatca   6039
FAM230B-LOC105372935-GGT2.NCBI.GGT2.rev.compl                       ttaatttaacaaatcagaccacatactttacttacattagttcacacctcatcatcatca   4014
                                                                    ************************************************************

chimp.LOC112206744.LOC107973052.GGT2-.646279-708040.rev.compl       tgcccatatgttgtgagcttgtttattgagcccacatgccagatggagaaactaagccac   6099
FAM230B-LOC105372935-GGT2.NCBI.GGT2.rev.compl                       tgcccatatgttgtgagcttgtttattgagcccacatgccagatggagaaactcagccac   4074
                                                                    ***************************************************** ******

chimp.LOC112206744.LOC107973052.GGT2-.646279-708040.rev.compl       ataaataaatgtgccctggttcacttgctgcatagtgaagagtcaaaatgtttactcata   6159
FAM230B-LOC105372935-GGT2.NCBI.GGT2.rev.compl                       ataaataaatgtgccctggttcacttgctgcatagtgaagagtcaaaatgtttactcata   4134
                                                                    ************************************************************

chimp.LOC112206744.LOC107973052.GGT2-.646279-708040.rev.compl       cggtgctaatgttgaaggcctgaactacaacctctatttatcagccagtgaagagatcac   6219
FAM230B-LOC105372935-GGT2.NCBI.GGT2.rev.compl                       cggtgctaatgttgaaggcctgaactacaacctctatttatcagccagtgaagagatcac   4194
                                                                    ************************************************************

chimp.LOC112206744.LOC107973052.GGT2-.646279-708040.rev.compl       tattcaccatgcaagggagttccagcaccctctatgcctggaattacccacgcctgcaga   6279
FAM230B-LOC105372935-GGT2.NCBI.GGT2.rev.compl                       tattccccatgcaagggagttccagcaccctctatgcctggaattacccacacctgcaga   4254
                                                                    ***** ********************************************* ********

chimp.LOC112206744.LOC107973052.GGT2-.646279-708040.rev.compl       gatcccaaacgccatccctcacataagagagcctcatgatctcataatccaggtagctat   6339
FAM230B-LOC105372935-GGT2.NCBI.GGT2.rev.compl                       gatcccaaacgccatccctcacataagacagcctcatgatctcataatccaggtagctat   4314
                                                                    **************************** *******************************

chimp.LOC112206744.LOC107973052.GGT2-.646279-708040.rev.compl       gtagacatcttcctgcaggtgtcacatagtccttagtgggaaaccaacatagaaagccca   6399
FAM230B-LOC105372935-GGT2.NCBI.GGT2.rev.compl                       gtagacatcttcctgcaggtgtcacatagtccttagtgtgaaaccaacatagaaagccca   4374
                                                                    ************************************** *********************

chimp.LOC112206744.LOC107973052.GGT2-.646279-708040.rev.compl       tgtttctgatcaaatcacaggttctgaaacactaagggaagcactaagtaggacaatgtg   6459
FAM230B-LOC105372935-GGT2.NCBI.GGT2.rev.compl                       tgtttctgatcaaatcacaggttctgaaacactaagggaggcactaagtaggacaacgtg   4434
                                                                    *************************************** **************** ***

chimp.LOC112206744.LOC107973052.GGT2-.646279-708040.rev.compl       gtgcctgtgtgtcatagctgggtctcctcaagacatggatcaagtccaataagaattggg   6519
FAM230B-LOC105372935-GGT2.NCBI.GGT2.rev.compl                       gtgcctgcgtgtcatagctgggtctcctcaagacatggatcaagtccagtaagaattggg   4494
                                                                    ******* **************************************** ***********

chimp.LOC112206744.LOC107973052.GGT2-.646279-708040.rev.compl       gagatgctttagagtcttgatggagttatcaccacaagccctctgagctacacactttag   6579
FAM230B-LOC105372935-GGT2.NCBI.GGT2.rev.compl                       gagatgctttagagtcttgatggagttatcaccacaagccctctgagctacacactttag   4554
                                                                    ************************************************************

chimp.LOC112206744.LOC107973052.GGT2-.646279-708040.rev.compl       ggatcatgaccattaagtactcaaattaccacttggttgttatccgggtatccgtcgtcc   6639
FAM230B-LOC105372935-GGT2.NCBI.GGT2.rev.compl                       ggatcatgaccattaagtactcaaattaccatttggttgctatccgggtatccgtcgtcc   4614
                                                                    ******************************* ******* ********************

chimp.LOC112206744.LOC107973052.GGT2-.646279-708040.rev.compl       ttgtggcaactctcttgtgaagctggtgtggacagcctcagtgctggagctgtgcctgcc   6699
FAM230B-LOC105372935-GGT2.NCBI.GGT2.rev.compl                       ttgtggcaaccctcttgtgaagctggtgtggacagcctcagtgctggagctgtgcctgcc   4674
                                                                    ********** *************************************************

chimp.LOC112206744.LOC107973052.GGT2-.646279-708040.rev.compl       ttctgaatggaccctttctgtgttagcaggtgggtacaagcatgggggtcagcacactca   6759
FAM230B-LOC105372935-GGT2.NCBI.GGT2.rev.compl                       ttctgagtggaccctttctgtgttagcaggtgggtacaagcgtgggggtcagcacactca   4734
                                                                    ****** ********************************** ******************

chimp.LOC112206744.LOC107973052.GGT2-.646279-708040.rev.compl       gtggatttacacacacagcattgaagagtaaggctgggcttcattatttatacattttca   6819
FAM230B-LOC105372935-GGT2.NCBI.GGT2.rev.compl                       gtggatttacacacacagcgttgaagagtaaggctgggcttcattatttatacattttca   4794
                                                                    ******************* ****************************************

chimp.LOC112206744.LOC107973052.GGT2-.646279-708040.rev.compl       ataaatgatgatcttcataacataaaatcaatgacgtagtacactagaatactgtcccta   6879
FAM230B-LOC105372935-GGT2.NCBI.GGT2.rev.compl                       ataaatgatgatcttcataacataaaatcaatgatgtagtacactagaatactgtcccta   4854
                                                                    ********************************** *************************

chimp.LOC112206744.LOC107973052.GGT2-.646279-708040.rev.compl       gtattgaatcttgtctctcaacaaagggttgcttaaagtcacatgacagattccattcaa   6939
FAM230B-LOC105372935-GGT2.NCBI.GGT2.rev.compl                       gtattgaatcttgtctctcagcaaagggttgcttaaagtcacgtgacagattccattcaa   4914
                                                                    ******************** ******************** ******************

chimp.LOC112206744.LOC107973052.GGT2-.646279-708040.rev.compl       ctgatgacacatgctgtagcagcagttaaagcagtcatttgaaaaggcttttactataaa   6999
FAM230B-LOC105372935-GGT2.NCBI.GGT2.rev.compl                       ctgatgacacatgctgtagcagcagttaaagcagtcatttgaaaaggcttttactataaa   4974
                                                                    ************************************************************

chimp.LOC112206744.LOC107973052.GGT2-.646279-708040.rev.compl       cttatgtgtgagcctgaagtggggataaaagaggtgattagctcccctgtgccatgttt   7059
FAM230B-LOC105372935-GGT2.NCBI.GGT2.rev.compl                       cttacgtgtgagcctgaagtggggataaaagaggcgattagctcccctgtgccatgttt   5034
                                                                    **** ****************************** ************************

chimp.LOC112206744.LOC107973052.GGT2-.646279-708040.rev.compl       ctattatgtgcatggtggaggaaaattacacaggaaggtgatggagagaacagagcaaag   7119
FAM230B-LOC105372935-GGT2.NCBI.GGT2.rev.compl                       ctattatgtgcgtggtggaggaaaattacacaggaaggtgatggagagaacagagcaaag   5094
                                                                    *********** ************************************************

chimp.LOC112206744.LOC107973052.GGT2-.646279-708040.rev.compl       gattggacaggtccattgaacccataagactatggtgaggttagtgaatgagattggtca   7179
FAM230B-LOC105372935-GGT2.NCBI.GGT2.rev.compl                       gattggacaggtccattgaacccataagactatggtgaggttagtgaatgagattggtca   5154
                                                                    ************************************************************

chimp.LOC112206744.LOC107973052.GGT2-.646279-708040.rev.compl       ttttaggtcaaattctacccagagctggtgcagccactgcccattcttagccagacctta   7239
FAM230B-LOC105372935-GGT2.NCBI.GGT2.rev.compl                       ttttaggtcaaattttacccagagctggtgcagccactgcccattcttagccagacctta   5214
                                                                    ************** *********************************************

chimp.LOC112206744.LOC107973052.GGT2-.646279-708040.rev.compl       ttgcaggcagctctgatcaatagtcaaggaggcagtggggcttgcagacttaattcatta   7299
FAM230B-LOC105372935-GGT2.NCBI.GGT2.rev.compl                       ttgcaggcagctctgatcaatagtcaaggaggcagtggggcttgcagacttaattcatta   5274
                                                                    ************************************************************

chimp.LOC112206744.LOC107973052.GGT2-.646279-708040.rev.compl       aatcaccaaagcaccagcccacacggccacttttccagttaattcacagtagcttgcata   7359
FAM230B-LOC105372935-GGT2.NCBI.GGT2.rev.compl                       aatcaccaaagcaccagcccaaacggccacttttccagttaattgacagtagcttgcata   5334
                                                                    ********************* ********************** ***************

chimp.LOC112206744.LOC107973052.GGT2-.646279-708040.rev.compl       ttcaggtttgatcagtggaagggaagttactctttgcagacccatcttttgacaatcatt   7419
FAM230B-LOC105372935-GGT2.NCBI.GGT2.rev.compl                       ttcaggtttgatcagtggaagggaagttactctttgcagacccatcttttgacaatcatt   5394
                                                                    ************************************************************

chimp.LOC112206744.LOC107973052.GGT2-.646279-708040.rev.compl       ttgcagtgtcagaaggtctgagcagcctcgggaggcaagcagtccctggtccctcagtgt   7479
FAM230B-LOC105372935-GGT2.NCBI.GGT2.rev.compl                       ttgtggtgtcggaaggtctgagcagcctcgggaggcaagcagtccctggtccctcagtgt   5454
                                                                    ***  ***** *************************************************

chimp.LOC112206744.LOC107973052.GGT2-.646279-708040.rev.compl       agtcactggaggagacagtcactgagaggcagctggcagggtgaagggaaaggggaggca   7539
FAM230B-LOC105372935-GGT2.NCBI.GGT2.rev.compl                       agtcactggaggagacagtcactgagaggcagctggcagggtgaagggaaaggggaggca   5514
                                                                    ************************************************************

chimp.LOC112206744.LOC107973052.GGT2-.646279-708040.rev.compl       ggccacagagatgacagcctttaagctgtcatactgggaggtcaaggatctgaaagagga   7599
FAM230B-LOC105372935-GGT2.NCBI.GGT2.rev.compl                       ggccacagagatgacagcctttaagctgtcatactgggaggtcaaggatctgaaagagga   5574
                                                                    ************************************************************

chimp.LOC112206744.LOC107973052.GGT2-.646279-708040.rev.compl       aggagaattctttatcattaaggacctgtccttatctcaggcatttcctccagagcatca   7659
FAM230B-LOC105372935-GGT2.NCBI.GGT2.rev.compl                       aggagaattctttatcattaaggacctgtccttatctcaggcatttcctccagagcatca   5634
                                                                    ************************************************************

chimp.LOC112206744.LOC107973052.GGT2-.646279-708040.rev.compl       cctttgtccacccacacaccttgggctaggaggactggagaaagacagtgaggggtctct   7719
FAM230B-LOC105372935-GGT2.NCBI.GGT2.rev.compl                       cctttgtccacccacacaccttgggctaggaggactggagaaagacagtgaggggtctct   5694
                                                                    ************************************************************

chimp.LOC112206744.LOC107973052.GGT2-.646279-708040.rev.compl       tgggtctctggcacagggcgtgatgaagaggtggcagtttttcaggaatctctctctcta   7779
FAM230B-LOC105372935-GGT2.NCBI.GGT2.rev.compl                       tgggtctctggcacagggcgtgatgaagaggtggcagtttttcaggaatctctctctcta   5754
                                                                    ************************************************************

chimp.LOC112206744.LOC107973052.GGT2-.646279-708040.rev.compl       gggaaccaaatacatttcccatctcagctccttcactcagcggggttgaggttctgctcg   7839
FAM230B-LOC105372935-GGT2.NCBI.GGT2.rev.compl                       gggaaccaaatacatttcccatctcaggtccttcactcagcggggttgaggttctgctcg   5814
                                                                    *************************** ********************************

chimp.LOC112206744.LOC107973052.GGT2-.646279-708040.rev.compl       tcacttatcatctctgaatgtcagcaccctcaagtgtaaaatctcagccacagcccttc   7899
FAM230B-LOC105372935-GGT2.NCBI.GGT2.rev.compl                       tcacttatcatctctgaatgtcagcaccctcaagtgtaaaatctcagccacagccctcc   5874
                                                                    ********************************************************** *

chimp.LOC112206744.LOC107973052.GGT2-.646279-708040.rev.compl       tctgcacccccctgcagggctgatgttctccataaaccataaggcatcatgcccacggaaa   7959
FAM230B-LOC105372935-GGT2.NCBI.GGT2.rev.compl                       tctgcacccccctgcagggctgatgttctccataaaccataaggcgtcatgcccacggaaa   5934
                                                                    ******************************************** ***************

chimp.LOC112206744.LOC107973052.GGT2-.646279-708040.rev.compl       agccgaacaggaaagcatgctccactgccccagagccatccaagttcccctccatattc   8019
FAM230B-LOC105372935-GGT2.NCBI.GGT2.rev.compl                       agccgaacaggaaagcatgctccactgccccggagtcatgcaagttcccctccatattc   5994
                                                                    ******************************* *** *** ********************
```

```
chimp.LOC112206744.LOC107973052.GGT2-.646279-708040.rev.compl      cgccactgctaagtgtccagcctattcctcctggcatgtagtaaacacttagagaacatt   8079
FAM230B-LOC105372935-GGT2.NCBI.GGT2.rev.compl                       cgccactgctaagtgtccagcttattcctcctggcatgtagtaaacacttagagaacatt   6054
                                                                    ********************* **************************************

chimp.LOC112206744.LOC107973052.GGT2-.646279-708040.rev.compl      actgaagtaccagtcctctccaaggttttcctgtatttagtgattttttagccctgtact   8139
FAM230B-LOC105372935-GGT2.NCBI.GGT2.rev.compl                       actgaagtaccagtcctctctaaggttttcctgtatttagtgattttttagccctgcact   6114
                                                                    ******************** *********************************** ***

chimp.LOC112206744.LOC107973052.GGT2-.646279-708040.rev.compl      gtgatactaagaagtagggcctaaatagggcctaaaaagtattgctaaaattacattatg   8199
FAM230B-LOC105372935-GGT2.NCBI.GGT2.rev.compl                       gtgatactaagaagtagggcttaaatagggcctaaaaagtattgctaaaattacattatg   6174
                                                                    ******************** ***************************************

chimp.LOC112206744.LOC107973052.GGT2-.646279-708040.rev.compl      acagtgcagagaactgagggcagagggaggacatgagcttgccaggtccacatggcttag   8259
FAM230B-LOC105372935-GGT2.NCBI.GGT2.rev.compl                       acagtgcagagaactgagggcagagggaggacatgagcttgccaggtccacatggcttag   6234
                                                                    ************************************************************

chimp.LOC112206744.LOC107973052.GGT2-.646279-708040.rev.compl      tggaatttgaatccgggcccccactctgcaccagccctgcactcacagtcatcctgctgt   8319
FAM230B-LOC105372935-GGT2.NCBI.GGT2.rev.compl                       tggaatttgaatccgggcccccactctgcaccagccctgcactcacagtcatccggctgt   6294
                                                                    ****************************************************** *****

chimp.LOC112206744.LOC107973052.GGT2-.646279-708040.rev.compl      gttctcctctccaggaaggcactgcccacgcagtctgtctgatagaggtgttgagtgctc   8379
FAM230B-LOC105372935-GGT2.NCBI.GGT2.rev.compl                       gttctcctctccaggaaggcactgcccacgcagtctgtctgatagaggtgttgagtgctc   6354
                                                                    ************************************************************

chimp.LOC112206744.LOC107973052.GGT2-.646279-708040.rev.compl      actgaactccgtgatcttcctgaaacccaactttgattcagtgggttctgcttggaagcc   8439
FAM230B-LOC105372935-GGT2.NCBI.GGT2.rev.compl                       actgaactccgtgatcttcctgaaacccaactttgattcagtgggctctgcttggaagcc   6414
                                                                    ********************************************* **************

chimp.LOC112206744.LOC107973052.GGT2-.646279-708040.rev.compl      tgtaaagaaaaggatcataagtttaaacttagaacagattatcactattttccctctggt   8499
FAM230B-LOC105372935-GGT2.NCBI.GGT2.rev.compl                       tgtaaagaaaaggatcataagtttaaacttagaacagattatcactattttccctctggt   6474
                                                                    ************************************************************

chimp.LOC112206744.LOC107973052.GGT2-.646279-708040.rev.compl      cttctgtcagcaagatgtcaacagccctatctattgtaaatgcattaaccagcatcttct   8559
FAM230B-LOC105372935-GGT2.NCBI.GGT2.rev.compl                       cttctgtcagcaagatgtcaacagccctatctattgtcaatgcattaaccagcatcttct   6534
                                                                    ************************************* **********************

chimp.LOC112206744.LOC107973052.GGT2-.646279-708040.rev.compl      ctgatagagaatataagaagatatgctgtgcacaccaaccagtgttggagacctcatggc   8619
FAM230B-LOC105372935-GGT2.NCBI.GGT2.rev.compl                       ctgatagagaatacaagaagatatgctgtgcacaccaaccagtgtaggagacctcatggc   6594
                                                                    ************* ******************************* **************

chimp.LOC112206744.LOC107973052.GGT2-.646279-708040.rev.compl      tcccgggtaaagaagaagaggtacccacaagaaggtactgtggaagttcattaattaagt   8679
FAM230B-LOC105372935-GGT2.NCBI.GGT2.rev.compl                       tcccgggtaaagaagaagaggtacccacaagaaggtactgtggaagttcattaattaagt   6654
                                                                    ************************************************************

chimp.LOC112206744.LOC107973052.GGT2-.646279-708040.rev.compl      tgattcaagaattgcaattgcggggagtattcagtgtcccatatgtaagaggaaactatg   8739
FAM230B-LOC105372935-GGT2.NCBI.GGT2.rev.compl                       tgattcaagaattgcagttgcggggagtattcagtgtcccatatgtaagaggaaactatg   6714
                                                                    **************** *******************************************

chimp.LOC112206744.LOC107973052.GGT2-.646279-708040.rev.compl      aagagactaagccatatttttaatgtgtcaggattctaatttgcctggtcagtaaatat   8799
FAM230B-LOC105372935-GGT2.NCBI.GGT2.rev.compl                       aagagactaagccatatttttaatgtgtcaggattctaatttgcctggtcagtaaatat   6774
                                                                    ************************************************************

chimp.LOC112206744.LOC107973052.GGT2-.646279-708040.rev.compl      tgctaccaccacaaaagtaaatatctacttaaaagtcaattttggttcatgttgaatgat   8859
FAM230B-LOC105372935-GGT2.NCBI.GGT2.rev.compl                       tgctaccaccacaaaagtaaatatctacttaaaagtcaattttggttcatgtttaatgat   6834
                                                                    ***************************************************** ******

chimp.LOC112206744.LOC107973052.GGT2-.646279-708040.rev.compl      agacaatgtttcaagctaatgtctagaacttacctggttgttaaacataagcatagatct   8919
FAM230B-LOC105372935-GGT2.NCBI.GGT2.rev.compl                       agacaatgtttcaagctaatgtctagaacttacctggttgttaaacataagcatagatct   6894
                                                                    ************************************************************

chimp.LOC112206744.LOC107973052.GGT2-.646279-708040.rev.compl      ccctgaaagagtggtgctatattattatttttcaattaatatatttctttagagagtttt   8979
FAM230B-LOC105372935-GGT2.NCBI.GGT2.rev.compl                       ccctgaaagagtggtgctatattattatttttcaattaatatatttctttagagagtttt   6954
                                                                    ************************************************************

chimp.LOC112206744.LOC107973052.GGT2-.646279-708040.rev.compl      aaattgacataaaaactgagcatatggccgggcgtggtggctcacacttataatcccagc   9039
FAM230B-LOC105372935-GGT2.NCBI.GGT2.rev.compl                       aaattgacataaaaactgagcatatggccgggcgtggtggctcacacttataatcccagc   7014
                                                                    ************************************************************

chimp.LOC112206744.LOC107973052.GGT2-.646279-708040.rev.compl      actttaggaggccaaggcaggcggatcatctgaggtcaagagttggagaccagcctggcc   9099
FAM230B-LOC105372935-GGT2.NCBI.GGT2.rev.compl                       actttaggaggccaaggcaggcggatcatctgaggtcaggagttggagaccagcctggcc   7074
                                                                    ************************************** *********************

chimp.LOC112206744.LOC107973052.GGT2-.646279-708040.rev.compl      aacatggtgaaaccccatctctactaaaatacaaaaaaattagccgggtgtggtggcag   9159
FAM230B-LOC105372935-GGT2.NCBI.GGT2.rev.compl                       aacatggtgaaaccccatctctactaaaatccaaaaaaattagccgggtgtggtggcag   7134
                                                                    ****************************** *****************************

chimp.LOC112206744.LOC107973052.GGT2-.646279-708040.rev.compl      gtgcctgtaatcccagctactcaggaggctgaggcaggagaatcgcttgaacccaggagg   9219
FAM230B-LOC105372935-GGT2.NCBI.GGT2.rev.compl                       gcgcctgtaatcccagctactcaggaggctgaggcaggagaatcgcttgaacccaggagg   7194
                                                                    * **********************************************************

chimp.LOC112206744.LOC107973052.GGT2-.646279-708040.rev.compl      cagaggttgcagtgagccaagatcatgccattgcactccagcctgggtgacaagagtgaa   9279
FAM230B-LOC105372935-GGT2.NCBI.GGT2.rev.compl                       cagaggttgcagtgagccaagatcatgccattgcactccagcctgggtgacaagagtgaa   7254
                                                                    ************************************************************

chimp.LOC112206744.LOC107973052.GGT2-.646279-708040.rev.compl      actccatctcaaaataaataaataaataaataaataaataaataaaaattgagtatataa   9339
FAM230B-LOC105372935-GGT2.NCBI.GGT2.rev.compl                       actccatctca--------aaataaataaataaataaataaataaaaattgagtatataa   7306
                                                                    ***********        *****************************************

chimp.LOC112206744.LOC107973052.GGT2-.646279-708040.rev.compl      tatacgaagttcccatattattctgtctcctcaccctcacttcctaatttcacctattag   9399
FAM230B-LOC105372935-GGT2.NCBI.GGT2.rev.compl                       tacacgaagttcccatattattctgtctcctcaccctcacttcctaatttcacctattag   7366
                                                                    ** *********************************************************

chimp.LOC112206744.LOC107973052.GGT2-.646279-708040.rev.compl      taacatcttacattactgtggtacatttgctagaataatgagaaaatattgacacattat   9459
FAM230B-LOC105372935-GGT2.NCBI.GGT2.rev.compl                       taacatcttacattactgtggtacatttgctagaataatgagaaaatattgacacattat   7426
                                                                    ************************************************************

chimp.LOC112206744.LOC107973052.GGT2-.646279-708040.rev.compl      tatctgaagtctacatttgcataatgttcattctttctgttatacatatatatgaatttt   9519
FAM230B-LOC105372935-GGT2.NCBI.GGT2.rev.compl                       tatctgaagtctacatttgcataatgttcattctttctgttatacatatatatgaatttt   7486
                                                                    ************************************************************

chimp.LOC112206744.LOC107973052.GGT2-.646279-708040.rev.compl      gaaatatttaaaacattatgttcacccttatggtctcataaagaaaatgttcacttccct   9579
FAM230B-LOC105372935-GGT2.NCBI.GGT2.rev.compl                       gaaatatttaaaacattatgttcacccttatggtctcataaagaaaatgttcacttccct   7546
                                                                    ************************************************************

chimp.LOC112206744.LOC107973052.GGT2-.646279-708040.rev.compl      aaaaatcctctcttctcattaatctctgtcctctttctccagaaaccttggcaactatta   9639
FAM230B-LOC105372935-GGT2.NCBI.GGT2.rev.compl                       aaaaatcctctcttctcattaatctctgtcctctttctccagaaaccttggcaactatta   7606
                                                                    ************************************************************

chimp.LOC112206744.LOC107973052.GGT2-.646279-708040.rev.compl      atatttttactatcacttcagctttgccttttccagaatgtcatatagttggaatcatat   9699
FAM230B-LOC105372935-GGT2.NCBI.GGT2.rev.compl                       atatttttactatcgcttcagctttgccttttccagaatgtcatatagttggaatcatat   7666
                                                                    ************** *********************************************

chimp.LOC112206744.LOC107973052.GGT2-.646279-708040.rev.compl      attatgtagtttttttcagatgaatttattgcactaaattgatgtacgctttagctgcttt   9759
FAM230B-LOC105372935-GGT2.NCBI.GGT2.rev.compl                       attatgtagtttttttcagatgaatttattgcactaaattgatgtacgctttagctgcttt   7726
                                                                    ************************************************************

chimp.LOC112206744.LOC107973052.GGT2-.646279-708040.rev.compl      catgtctttttttatgccttaatggcaaaaaatggcacattaaatcaccaaataatattgc   9819
FAM230B-LOC105372935-GGT2.NCBI.GGT2.rev.compl                       catgtctttttttatgccttaatggcaaaaaatggcacattaaatcaccaaataatattgc   7786
                                                                    ************************************************************

chimp.LOC112206744.LOC107973052.GGT2-.646279-708040.rev.compl      attaaatgaattttttgtctttttattcacctgttgaagaattcggtagatttcatgagag   9879
FAM230B-LOC105372935-GGT2.NCBI.GGT2.rev.compl                       attcaatgaattttttgtctttttattcacctgttgaagaattcggtagatttcatgagag   7846
                                                                    *** ********************************************************

chimp.LOC112206744.LOC107973052.GGT2-.646279-708040.rev.compl      aaaccatctgggcctggtgctttcttttcagaatgctcttaatgtgaattcaacttatt   9939
FAM230B-LOC105372935-GGT2.NCBI.GGT2.rev.compl                       aaaccatctgggcctggtgctttcttttcggaatgctcttaatgtgaattcaacttatt   7906
                                                                    ***************************** ******************************

chimp.LOC112206744.LOC107973052.GGT2-.646279-708040.rev.compl      taatagacataagtttattcaaattaggatcctagcatgaccttgggaagattgcctttc   9999
FAM230B-LOC105372935-GGT2.NCBI.GGT2.rev.compl                       taatagacataagtttattcaaattaggatcctagcatgaccttgggaagattgcctttc   7966
                                                                    ************************************************************

chimp.LOC112206744.LOC107973052.GGT2-.646279-708040.rev.compl      aaggaattgatacatttcactgaggttatcaaactgcggtcatagaactgttcatgatat   10059
FAM230B-LOC105372935-GGT2.NCBI.GGT2.rev.compl                       aaggaattgatacatttcactgaggttatcaaactgcggtcatagaactgttcatgatat   8026
                                                                    ************************************************************

chimp.LOC112206744.LOC107973052.GGT2-.646279-708040.rev.compl      tccttttaatgcctaacagttcagtagagatggctcctcttttatttctgaaattggtca   10119
FAM230B-LOC105372935-GGT2.NCBI.GGT2.rev.compl                       tccttttaatgcctaacagttcagtagagatggctcctcttttatttctgaaattggtca   8086
                                                                    ************************************************************

chimp.LOC112206744.LOC107973052.GGT2-.646279-708040.rev.compl      tttgtgttatcttctttttcttggttagcctgcatatcaattcattcattgtaatgagca   10179
FAM230B-LOC105372935-GGT2.NCBI.GGT2.rev.compl                       tttgtgttatcttctttttcttggttagcctgcatatcaattcattcattgtaatgagca   8146
                                                                    ************************************************************

chimp.LOC112206744.LOC107973052.GGT2-.646279-708040.rev.compl      tatcaaagaaccagcttttggttttattgattttctgatggtttcagtgttttaattta   10239
```

```
FAM230B-LOC105372935-GGT2.NCBI.GGT2.rev.compl                  tatcaaagaaccagcttttggttttattgattttctgatgatttcagtgttttaattta   8206
                                                               **************************************** *******************

chimp.LOC112206744.LOC107973052.GGT2-.646279-708040.rev.compl   ttgatttctgtgatgttgtttattacttttacttgctttccattgcattcctctattttc   10299
FAM230B-LOC105372935-GGT2.NCBI.GGT2.rev.compl                  ttgatttctgtgatgttgtttattacttttacttgctttccattgcattcctctattttc   8266
                                                               ************************************************************

chimp.LOC112206744.LOC107973052.GGT2-.646279-708040.rev.compl   tatagttccctaattgaaacatgatattactgattttaggtcttgtgattttttagtatat   10359
FAM230B-LOC105372935-GGT2.NCBI.GGT2.rev.compl                  tatagttccctaattgaaacatgatattactgattttaggtcttgtgattttttagtatat   8326
                                                               ************************************************************

chimp.LOC112206744.LOC107973052.GGT2-.646279-708040.rev.compl   tgcatccaatgctatagatttccctctaaggactgcttttgctacatccagaaatcttgc   10419
FAM230B-LOC105372935-GGT2.NCBI.GGT2.rev.compl                  tgcatccaatgctatagatttccctctaaggactgcttttgctacatccagaaatcttgc   8386
                                                               ************************************************************

chimp.LOC112206744.LOC107973052.GGT2-.646279-708040.rev.compl   caagtcacattttcttttaatgtagttaaaagtatttttaattttctattgagacttctt   10479
FAM230B-LOC105372935-GGT2.NCBI.GGT2.rev.compl                  caagtcacattttcttttaatgtagttaaaagtatttttaattttctattgagacttctt   8446
                                                               ************************************************************

chimp.LOC112206744.LOC107973052.GGT2-.646279-708040.rev.compl   ctttaacccatgagttatttaaaagtgcattgctaatttgcaaatatttggggattttgt   10539
FAM230B-LOC105372935-GGT2.NCBI.GGT2.rev.compl                  ctttaacccatgagttatttaaaagtgcattgctaatttgcaaatatttggggattttgt   8506
                                                               ************************************************************

chimp.LOC112206744.LOC107973052.GGT2-.646279-708040.rev.compl   ggctcttttacagttgttgatttttttgttgtcaggtatgtgttgcaaaagcagtcgtcta   10599
FAM230B-LOC105372935-GGT2.NCBI.GGT2.rev.compl                  ggctcttttacagttgttgaattttttgttgtcaggtatgtgttgcaaaagcagtcgtcta   8566
                                                               ******************** ***************************************

chimp.LOC112206744.LOC107973052.GGT2-.646279-708040.rev.compl   cctcatcttgccaccagccaagatggcccaggatgtgggctctccctgagtgaatctttg   10659
FAM230B-LOC105372935-GGT2.NCBI.GGT2.rev.compl                  cctcatcttgccaccacccaagatggcccaggatgtgggctctccctgagtgaatctttg   8626
                                                               **************** *******************************************

chimp.LOC112206744.LOC107973052.GGT2-.646279-708040.rev.compl   gcaatctgccaacctgatgtgttcggcctccttctttagtctgagtttcccttctgctta   10719
FAM230B-LOC105372935-GGT2.NCBI.GGT2.rev.compl                  gcaatctgccaacctgatgtgttcggcctccttctttagtctgagcttgccttctgctta   8686
                                                               ********************************************* ** ***********

chimp.LOC112206744.LOC107973052.GGT2-.646279-708040.rev.compl   gaaagggccattctcagttctggcagggagttttcccaacattgagaaggtggcattctt   10779
FAM230B-LOC105372935-GGT2.NCBI.GGT2.rev.compl                  gaaagggccattctcagttctggcagggagttttcccaacattgagaaggtggcattctt   8746
                                                               ************************************************************

chimp.LOC112206744.LOC107973052.GGT2-.646279-708040.rev.compl   actccccactgcagcctgcacctctgaccggtggtcagcagacaggacagaggtcctcag   10839
FAM230B-LOC105372935-GGT2.NCBI.GGT2.rev.compl                  actccccactgcagcctgcacctctgaccggtggtcagcagacaggacagaggtcctcat   8806
                                                               *********************************************************** 

chimp.LOC112206744.LOC107973052.GGT2-.646279-708040.rev.compl   tagacagagttcagcggggtctctgaccaaagtgcatcttcagagtctgcacctacccac   10899
FAM230B-LOC105372935-GGT2.NCBI.GGT2.rev.compl                  cagacagagttcagcagggtctctgaccaaagtgcatcttcagagtctgcacctacccac   8866
                                                                ************** ********************************************

chimp.LOC112206744.LOC107973052.GGT2-.646279-708040.rev.compl   tgtgaccacgggcaggctctgagtcctaaagcaggaggaactgtgcgaccatcctgattg   10959
FAM230B-LOC105372935-GGT2.NCBI.GGT2.rev.compl                  tgtgaccacgggcaggctctgagtcctaaagcaggaggaactgtgcgaccatcctgattg   8926
                                                               ************************************************************

chimp.LOC112206744.LOC107973052.GGT2-.646279-708040.rev.compl   gaaatttgtgaggatcaccgtgttactcaagtaaggtctttggaaagtgtcgtattacta   11019
FAM230B-LOC105372935-GGT2.NCBI.GGT2.rev.compl                  gaaatttgtgaggatcaccgtgttactcaagtaaggtctttggaaagtgtcgtattacta   8986
                                                               ************************************************************

chimp.LOC112206744.LOC107973052.GGT2-.646279-708040.rev.compl   ctgtttgtgaactgcttgttggtggcctggctgagccacacactttatgaaaaccaggac   11079
FAM230B-LOC105372935-GGT2.NCBI.GGT2.rev.compl                  ctgtttgtgaactgcttgttggtggcctggctgagccacacactttatgaaaaccaggac   9046
                                                               ************************************************************

chimp.LOC112206744.LOC107973052.GGT2-.646279-708040.rev.compl   ccctcagctggtgtgggtgtctatgcagactgagaccctcatgtgaacagccttgtggca   11139
FAM230B-LOC105372935-GGT2.NCBI.GGT2.rev.compl                  ccctcagctggtgtgggtgtctatgcagcctgagaccctcatgtgaacagcctcgtggca   9106
                                                               **************************** ************************ ******

chimp.LOC112206744.LOC107973052.GGT2-.646279-708040.rev.compl   gctgtctttgccccttgccaccatcagtgcctccttgttcctgggcactgctttctctga   11199
FAM230B-LOC105372935-GGT2.NCBI.GGT2.rev.compl                  gctgtctttgccccttgccaccatcagtgcctccttgttcctgggcactgctttctctga   9166
                                                               ************************************************************

chimp.LOC112206744.LOC107973052.GGT2-.646279-708040.rev.compl   tggtgctccattgttttcctgcacctcagtgtctacagctggatgtctcttctgcaatct   11259
FAM230B-LOC105372935-GGT2.NCBI.GGT2.rev.compl                  tggtgctccattgttttcctgcacctcagtgtctacagctggatgtctcttccgcaatct   9226
                                                               **************************************************** *******

chimp.LOC112206744.LOC107973052.GGT2-.646279-708040.rev.compl   aggcgagggggcatcaatggcagttctgctgtggcactgccctccttcttagcttgtctt   11319
FAM230B-LOC105372935-GGT2.NCBI.GGT2.rev.compl                  aggcgagggggcatcaacggcagttctgctgtggcactgccctccttcttagcttgtctt   9286
                                                               ***************** ******************************************

chimp.LOC112206744.LOC107973052.GGT2-.646279-708040.rev.compl   gctctgtcttaggctccctcaaagatcccacccttcaggttcttccacaagtttcttact   11379
FAM230B-LOC105372935-GGT2.NCBI.GGT2.rev.compl                  gctctgtcttaggctccctcaaagatcccacccttcaggttcttccacaagtttcttatt   9346
                                                               ********************************************************** *

chimp.LOC112206744.LOC107973052.GGT2-.646279-708040.rev.compl   gaaatccgagcagaaaactatgaccaatatgaccaatcctatacccactgaagacatgaa   11439
FAM230B-LOC105372935-GGT2.NCBI.GGT2.rev.compl                  gaaatccgagcagaaaactatgaccaatatgaccaatcctatacccactgaagacatgaa   9406
                                                               ************************************************************

chimp.LOC112206744.LOC107973052.GGT2-.646279-708040.rev.compl   tgaagaattaaaacaattctccatggactctaccatatagatccctagaagtaatttcta   11499
FAM230B-LOC105372935-GGT2.NCBI.GGT2.rev.compl                  tgaagaattaaaacaattctccatggactctaccatatagatccctagaagtaatttcta   9466
                                                               ************************************************************

chimp.LOC112206744.LOC107973052.GGT2-.646279-708040.rev.compl   aaaaaaaaaaaaaaa----caaggaagatgtaatagttttccataaattagaataccta   11555
FAM230B-LOC105372935-GGT2.NCBI.GGT2.rev.compl                  aaaaaaaaaaaaaaaaaatcaaggaagatgtaatagttttccataaattagaataccta   9526
                                                               ***************    *****************************************

chimp.LOC112206744.LOC107973052.GGT2-.646279-708040.rev.compl   catgtacaattaaatgaaatggctagtatagtcttgaaaccaaaaccagataaggtaaat   11615
FAM230B-LOC105372935-GGT2.NCBI.GGT2.rev.compl                  catgtacaattaaatgaaatggctagtatagtcttgaaaccaaaaccagataaggtaaat   9586
                                                               ************************************************************

chimp.LOC112206744.LOC107973052.GGT2-.646279-708040.rev.compl   taaattctgtgatattttaaaatactgtaaattctgactaatgtgagttaatctcaatat   11675
FAM230B-LOC105372935-GGT2.NCBI.GGT2.rev.compl                  taaattctgtgatattttaaaatactgtaaattctgactaatgtgagttaatctcaatat   9646
                                                               ************************************************************

chimp.LOC112206744.LOC107973052.GGT2-.646279-708040.rev.compl   atgaaatagtagattaacattgaaaatgcaataaataaaattagctacctcaagagttta   11735
FAM230B-LOC105372935-GGT2.NCBI.GGT2.rev.compl                  atgaaatagtagattaacattgaaaatgcaataaataaaattagctacctcaagagttta   9706
                                                               ************************************************************

chimp.LOC112206744.LOC107973052.GGT2-.646279-708040.rev.compl   atggaaaaaaatgtgattattgcaatagattcaggaaattcatgaataacattcaccctaa   11795
FAM230B-LOC105372935-GGT2.NCBI.GGT2.rev.compl                  atggaaaaaaatgtgattattgcaatagattcaggaaattcatgaataacattcaccctaa   9766
                                                               ************************************************************

chimp.LOC112206744.LOC107973052.GGT2-.646279-708040.rev.compl   tatttgtaggacaactatctgattaactttaagtgacctgtgacaaccatttgaattaat   11855
FAM230B-LOC105372935-GGT2.NCBI.GGT2.rev.compl                  tatttgtaggacagctatc----taactttaagtgacctgtgacaaccatttgaattaat   9822
                                                               ************* *****    *************************************

chimp.LOC112206744.LOC107973052.GGT2-.646279-708040.rev.compl   gctgctttcacagcatatctcttggcttgttaaaaacccgacaagaatttccgtaacatt   11915
FAM230B-LOC105372935-GGT2.NCBI.GGT2.rev.compl                  gctgctttcacagcatatctcttggcttgttaaaaacccgacaagaatttccgtaacatt   9882
                                                               ************************************************************

chimp.LOC112206744.LOC107973052.GGT2-.646279-708040.rev.compl   aatttattttaacacctatattgggtgtgaacccaccataaagtttgcccactgaaaag   11975
FAM230B-LOC105372935-GGT2.NCBI.GGT2.rev.compl                  aatttattttaacacctatattgggtgtgaacccaccataaagtttgcccactgaaaag   9942
                                                               ************************************************************

chimp.LOC112206744.LOC107973052.GGT2-.646279-708040.rev.compl   gtctacaatttgatgctttattaaattgatactgtgtgcacccatcaccaccatctaatt   12035
FAM230B-LOC105372935-GGT2.NCBI.GGT2.rev.compl                  gtctacaatttgatgctttattaaattgatactgtgtgcacccatcaccacgatctaatt   10002
                                                               *************************************************** ********

chimp.LOC112206744.LOC107973052.GGT2-.646279-708040.rev.compl   taaatatgtttccctcacccaaattctctcttgcgcactggcagttaatccccacccccca   12095
FAM230B-LOC105372935-GGT2.NCBI.GGT2.rev.compl                  taaatatgtttccctcacccaaattctctcttgcgcactggcagttaatccccactccca   10062
                                                               ******************************************************* ****

chimp.LOC112206744.LOC107973052.GGT2-.646279-708040.rev.compl   tctccagccctaagcaatactgctgtgacattccatctccataaatttcccacttgctta   12155
FAM230B-LOC105372935-GGT2.NCBI.GGT2.rev.compl                  tctccagccctaagcaatactgctgtgacattccatctccataaatttcccatttgctta   10122
                                                               **************************************************** *******

chimp.LOC112206744.LOC107973052.GGT2-.646279-708040.rev.compl   atagaaatggacatatatatatatttggaatctgacttccttcatttagcatactatgtt   12215
FAM230B-LOC105372935-GGT2.NCBI.GGT2.rev.compl                  atagaaatggacatatatatatatttggaatctgacttccttcatttagcatactatgtt   10182
                                                               ************************************************************

chimp.LOC112206744.LOC107973052.GGT2-.646279-708040.rev.compl   tgaagttaattgacgtgttagcacgtgctggtcatgtgttttccttcatagtctgctgtg   12275
FAM230B-LOC105372935-GGT2.NCBI.GGT2.rev.compl                  tgaagttaattgacgtgttagcacgtgctggtcatgtgttttccttcatagtctgctgtg   10242
                                                               ************************************************************

chimp.LOC112206744.LOC107973052.GGT2-.646279-708040.rev.compl   tttactcatacagatagtgtttattcgtttatcagttaatggacatttaattgttttgtt   12335
FAM230B-LOC105372935-GGT2.NCBI.GGT2.rev.compl                  tttactcatacagatagtgtttattcatttatcagttaatggacatttaattgttttgtt   10302
                                                               ************************** *********************************

chimp.LOC112206744.LOC107973052.GGT2-.646279-708040.rev.compl   attttctttgatgagtaacgtagcttcgagcattcatatacagtcaggtaatgcataatg   12395
FAM230B-LOC105372935-GGT2.NCBI.GGT2.rev.compl                  attttctttgatgagtaatgtagctttgagcattcatatacagtcatgtaatgcataatg   10362
```

```
                                                                 ****************** ******* ******************* *************

chimp.LOC112206744.LOC107973052.GGT2-.646279-708040.rev.compl    acattttggtcaaaaaaaatttttttttctgagacccaggctggagtgcagtggcacaa   12455
FAM230B-LOC105372935-GGT2.NCBI.GGT2.rev.compl                    acattttggtcaaaaaaattttttttttctgagacccaggctggagtgcagtggcacaa   10422
                                                                 ****************** *****************************************

chimp.LOC112206744.LOC107973052.GGT2-.646279-708040.rev.compl    tctcggctcactggaacctccacctcccaggtttaagcaattcttgtgcctcaacctccc   12515
FAM230B-LOC105372935-GGT2.NCBI.GGT2.rev.compl                    tctcggctcactggaacctccacctcccaggtttaagcaattctcatgcctcaacctccc   10482
                                                                 ********************************************  **************

chimp.LOC112206744.LOC107973052.GGT2-.646279-708040.rev.compl    gagtagctgggacaactggcacacgccaccatgcctggataattttgtattttcagtag   12575
FAM230B-LOC105372935-GGT2.NCBI.GGT2.rev.compl                    gagtagctgggacaactggcacacgccaccatgcctggataattttgtattttcagtag   10542
                                                                 ************************************************************

chimp.LOC112206744.LOC107973052.GGT2-.646279-708040.rev.compl    agacaggattttgctgtgttggtcaggctagtctcaaactcctagcctcaggtgatccac   12635
FAM230B-LOC105372935-GGT2.NCBI.GGT2.rev.compl                    agacaggattttgctgtgttggtcaggctagtctcaaactcccagcctcaggtgatccac   10602
                                                                 ****************************************** *****************

chimp.LOC112206744.LOC107973052.GGT2-.646279-708040.rev.compl    ccatctctgcctcccaaagtgctgggattataggcatgagccaccacacccagcctaatt   12695
FAM230B-LOC105372935-GGT2.NCBI.GGT2.rev.compl                    ctatctctgcctcccaaagtgctgggattataggcatgagccaccacacccagcctaatt   10662
                                                                 * **********************************************************

chimp.LOC112206744.LOC107973052.GGT2-.646279-708040.rev.compl    tttttaagaaacaagggaactattttctaaattacttttgccaatttatatttctaccat   12755
FAM230B-LOC105372935-GGT2.NCBI.GGT2.rev.compl                    tttttaagaaacaagggaactattttctaaattacttttgccaatttatatttctaccat   10722
                                                                 ************************************************************

chimp.LOC112206744.LOC107973052.GGT2-.646279-708040.rev.compl    gatgcatagcactaatttcaccgtacaatgtatggtaggccccaatatgtaagaaatgat   12815
FAM230B-LOC105372935-GGT2.NCBI.GGT2.rev.compl                    gatgcatagcactaatttcaccgtacaatgtatggtaggccccaatatgtaagaaatgat   10782
                                                                 ************************************************************

chimp.LOC112206744.LOC107973052.GGT2-.646279-708040.rev.compl    gaaagtaacacataaagattagtataaaacaaataaaattatcattgttgctatcatctt   12875
FAM230B-LOC105372935-GGT2.NCBI.GGT2.rev.compl                    gaaaataacacataaagattggtataaaacaaataagattatcattgttgctatcatctt   10842
                                                                 **** *************** *************** ***********************

chimp.LOC112206744.LOC107973052.GGT2-.646279-708040.rev.compl    tgtcaaattctgaaaacaatctgagtatattttatataaatatgcttggcaacatagct   12935
FAM230B-LOC105372935-GGT2.NCBI.GGT2.rev.compl                    tgtcaaattctgaaaacaatctgagtatattttatataaatatgcttggcaacatagct   10902
                                                                 ************************************************************

chimp.LOC112206744.LOC107973052.GGT2-.646279-708040.rev.compl    gaaaaacgcattatcagttacatttatcagtaacaaagacataaatttgaaggggaaaa   12995
FAM230B-LOC105372935-GGT2.NCBI.GGT2.rev.compl                    gaaaaacgcattatcagttacatttatcagtaacaaagacataaatttgaaggggaaca   10962
                                                                 ********************************************************** *

chimp.LOC112206744.LOC107973052.GGT2-.646279-708040.rev.compl    acacttgtactaacaacgcaatgtcagaattcacataaaaattctgctggtcactttgga   13055
FAM230B-LOC105372935-GGT2.NCBI.GGT2.rev.compl                    acacttgtactaacaacgcaatgtcagaattaacataaaaattctgctggtcactttgga   11022
                                                                 ******************************* ****************************

chimp.LOC112206744.LOC107973052.GGT2-.646279-708040.rev.compl    atatttaattgcctggggcagtgtttagtagacaaatgagcatctatggagcacccaaag   13115
FAM230B-LOC105372935-GGT2.NCBI.GGT2.rev.compl                    atatttaattgcctggggcagtgtttagtagacaaatgagcatctatggagcactcaaag   11082
                                                                 ****************************************************** *****

chimp.LOC112206744.LOC107973052.GGT2-.646279-708040.rev.compl    taggggaatcaacagaacttcggtttgaaaagttatctgggtttagagcgtgaaactttg   13175
FAM230B-LOC105372935-GGT2.NCBI.GGT2.rev.compl                    taggggaatcaacagaacttgggtttgaaaagttatctgggtttagagcgtgaaactttg   11142
                                                                 ******************** ***************************************

chimp.LOC112206744.LOC107973052.GGT2-.646279-708040.rev.compl    ttagaggacacacgccttgcatgagcgaggtgccttggtgtgtgtggacgtaccattatg   13235
FAM230B-LOC105372935-GGT2.NCBI.GGT2.rev.compl                    ttagaggacacgcaccttgcatgagcgaggtgccctggtgtgtgtggacgtaccattatg   11202
                                                                 *********** * ******************** *************************

chimp.LOC112206744.LOC107973052.GGT2-.646279-708040.rev.compl    cttggaggtacagcataatggtggcttcctccagaaagggacatttggggtggattcatt   13295
FAM230B-LOC105372935-GGT2.NCBI.GGT2.rev.compl                    cttggaggtacagcataatggtggcttcctccagaaagggacatttggggtggattcatt   11262
                                                                 ************************************************************

chimp.LOC112206744.LOC107973052.GGT2-.646279-708040.rev.compl    ccatctagacaacacagcctgatgtggcatggacatgaatggaggtgaaatggtcagtag   13355
FAM230B-LOC105372935-GGT2.NCBI.GGT2.rev.compl                    ccatctagacaacacagcctgatgtggcatggacatgaatggaggtgaaatggtcagtag   11322
                                                                 ************************************************************

chimp.LOC112206744.LOC107973052.GGT2-.646279-708040.rev.compl    ttgagaggatcagtcctgacaagggccaaggtgaaaaacctgggaaccccttcaggtaca   13415
FAM230B-LOC105372935-GGT2.NCBI.GGT2.rev.compl                    ttgagaggatcagtcctgacaagggccgaggtgaaaaacctgggaaccccttcaggtaca   11382
                                                                 *************************** ********************************

chimp.LOC112206744.LOC107973052.GGT2-.646279-708040.rev.compl    aagtcttcagttgaaaaaggaggtggtcacaggaaatactgagacaggacagcaatgcat   13475
FAM230B-LOC105372935-GGT2.NCBI.GGT2.rev.compl                    aagtcttcagttgaaaaaggaggtggtcacaggaaatactgagacaggacagcaaagcat   11442
                                                                 ******************************************************* ****

chimp.LOC112206744.LOC107973052.GGT2-.646279-708040.rev.compl    gggagacagagttcttggccctgcagggtgagtagtgtggattctcaagttttctcctct   13535
FAM230B-LOC105372935-GGT2.NCBI.GGT2.rev.compl                    gggagacagagttcttggccctgcagggtgagtagtgtggattctcaagttttctcctct   11502
                                                                 ************************************************************

chimp.LOC112206744.LOC107973052.GGT2-.646279-708040.rev.compl    ctccattaatttctttcccaatgcagatgacttccatcatacagtcttcagcaaccttga   13595
FAM230B-LOC105372935-GGT2.NCBI.GGT2.rev.compl                    ctccattaatttctttcccaatgcagatgacttccatcatacagtcttcagcaaccttga   11562
                                                                 ************************************************************

chimp.LOC112206744.LOC107973052.GGT2-.646279-708040.rev.compl    aagattggacaagcttcagcccactcttgaaggtaaaggaaggcagctaacaagactggc   13655
FAM230B-LOC105372935-GGT2.NCBI.GGT2.rev.compl                    aagattggacaagcttcagcccactcttgaaggtaaaggaaggcagctaacaagactggc   11622
                                                                 ************************************************************

chimp.LOC112206744.LOC107973052.GGT2-.646279-708040.rev.compl    atctgggcttggctgtgcgtgttttctattgtggggaaatatatataacacaatatttat   13715
FAM230B-LOC105372935-GGT2.NCBI.GGT2.rev.compl                    atctgggcttggctgtgcgtgttttctatcgtggggaaatatatataacacaatatttat   11682
                                                                 **************************** *******************************

chimp.LOC112206744.LOC107973052.GGT2-.646279-708040.rev.compl    catttgaaccttttaaccaaagtgtgcactccatggcattcaatatattcacagggttgc   13775
FAM230B-LOC105372935-GGT2.NCBI.GGT2.rev.compl                    catttgaaccttttaaccaaagtgtgcactccatggcattcaatatattcacagggttgc   11742
                                                                 ************************************************************

chimp.LOC112206744.LOC107973052.GGT2-.646279-708040.rev.compl    ataaccaacaccactatctacacccacaattttgatgatttcttacaaaaccttgtccac   13835
FAM230B-LOC105372935-GGT2.NCBI.GGT2.rev.compl                    ataaccaacaccactatctacacccacaattttgatgatttcttacgaaaccttgtccac   11802
                                                                 ********************************************** *************

chimp.LOC112206744.LOC107973052.GGT2-.646279-708040.rev.compl    tataagcaatatagcaccttcccccctatttccagctcatggtgattcctatcccactttc   13895
FAM230B-LOC105372935-GGT2.NCBI.GGT2.rev.compl                    aataagcaatatagcaccttcccccctatttccagcccatggtgattcctatcccactttc   11862
                                                                  ********************************* *************************

chimp.LOC112206744.LOC107973052.GGT2-.646279-708040.rev.compl    tcttgtatgaatttgactattctaggcacttcatgtaattacaattatacaatatattcc   13955
FAM230B-LOC105372935-GGT2.NCBI.GGT2.rev.compl                    tcttgtatgaatttgactattctaggcacttcatgtaattacaattatacaatatattcc   11922
                                                                 ************************************************************

chimp.LOC112206744.LOC107973052.GGT2-.646279-708040.rev.compl    ttttgtgtctggcttatttcactaagcataatgttctcaatgtccacccatgttgtatca   14015
FAM230B-LOC105372935-GGT2.NCBI.GGT2.rev.compl                    ttttgtgtctggcttatttcactaagcataatgttctcaatgtccacccatgttgtatca   11982
                                                                 ************************************************************

chimp.LOC112206744.LOC107973052.GGT2-.646279-708040.rev.compl    tctatcaaaattatgttcgtttttacagatggatgatgtagcattgcatgcagaccacc   14075
FAM230B-LOC105372935-GGT2.NCBI.GGT2.rev.compl                    tctttcaaaatgatgttcgtttttacagatggatgatgtagcattgcatgcagaccacc   12042
                                                                 *** ******* ************************************************

chimp.LOC112206744.LOC107973052.GGT2-.646279-708040.rev.compl    ttgcttttattacattcatttgttcactgatggttggattatttccaccttttggctcct   14135
FAM230B-LOC105372935-GGT2.NCBI.GGT2.rev.compl                    ttgcttttattacattcatttgttcactgatggttggattatttccaccttttggctcct   12102
                                                                 ************************************************************

chimp.LOC112206744.LOC107973052.GGT2-.646279-708040.rev.compl    gtgaaaagtgatgctacaaacattagtatacaaacatctgtttgatttctgttctctatt   14195
FAM230B-LOC105372935-GGT2.NCBI.GGT2.rev.compl                    gtgaaaagtgatgctacaaacattagtatacaaacatctgtttgatttccgttctctatt   12162
                                                                 ************************************************* **********

chimp.LOC112206744.LOC107973052.GGT2-.646279-708040.rev.compl    ctttggggtgcctaagagtagagttcctgggtccaacaggggttctatatttaaacttct   14255
FAM230B-LOC105372935-GGT2.NCBI.GGT2.rev.compl                    ctttggggtgcctaagagtagagttcctgggtccaacaggggttctatatttaaccttct   12222
                                                                 ****************************************************** *****

chimp.LOC112206744.LOC107973052.GGT2-.646279-708040.rev.compl    gagccactgcagactgtttttcagagtggctgcaactttatccatttctaccatcaatgt   14315
FAM230B-LOC105372935-GGT2.NCBI.GGT2.rev.compl                    gagccactgcagactgtttttcacagtggctgcaactttatccatttctaccatcaatgt   12282
                                                                 *********************** ************************************

chimp.LOC112206744.LOC107973052.GGT2-.646279-708040.rev.compl    atcagggttacaatttctttacgcccttcttcacacttattttcctttaaatcatcctag   14375
FAM230B-LOC105372935-GGT2.NCBI.GGT2.rev.compl                    atcagggttacaatttctttacgtccttcttcacacttattttcctttaaatcatcctag   12342
                                                                 *********************** ************************************

chimp.LOC112206744.LOC107973052.GGT2-.646279-708040.rev.compl    taggtgtatattggtggctgcttgtgtttctcatttgcatttccctaatgactaatgatc   14435
FAM230B-LOC105372935-GGT2.NCBI.GGT2.rev.compl                    taggtgtatattggtggctgcttgtgttttttcatttgcatttccctaatgactaatgatc   12402
                                                                 ***************************** ******************************

chimp.LOC112206744.LOC107973052.GGT2-.646279-708040.rev.compl    ctgagcagcttttcctgtgctactatctgtggctgtatcttctttagggaaatatatgtt   14495
FAM230B-LOC105372935-GGT2.NCBI.GGT2.rev.compl                    ctgagcagcttttcctgtgctactatctgtggctgtatcttctttagggaaatatatgtt   12462
                                                                 ************************************************************

chimp.LOC112206744.LOC107973052.GGT2-.646279-708040.rev.compl    gaagtcttttgcccattttaaagagttgtctgatttttatttagttagtttgttgttgt   14555
FAM230B-LOC105372935-GGT2.NCBI.GGT2.rev.compl                    gaagtcttttgcccattttaaagagttgtctgatttttatttagttagtttgttgttgt   12522
                                                                 ************************************************************
```

```
chimp.LOC112206744.LOC107973052.GGT2-.646279-708040.rev.compl      agattttgaatatatcttaaatatatttaaaatagattctaaattttagtctcttacaa   14615
FAM230B-LOC105372935-GGT2.NCBI.GGT2.rev.compl                      agattttgaatatatcttaaatatatttaaaatatattctaaattttagtctcttacaa   12582
                                                                   ********************************** *************************

chimp.LOC112206744.LOC107973052.GGT2-.646279-708040.rev.compl      gataaatgatttgcaaatatttccacctttgtgtagaactttagattcacaaacttcatt   14675
FAM230B-LOC105372935-GGT2.NCBI.GGT2.rev.compl                      gagaaatgatttgcaaatatttccacctttgtgtagaactttagattcacaaacttcgtt   12642
                                                                   ** ****************************************************** **

chimp.LOC112206744.LOC107973052.GGT2-.646279-708040.rev.compl      aatttgtgtgaaatcctcagcagttgaccccaaacagataagactgaagcagtattttag   14735
FAM230B-LOC105372935-GGT2.NCBI.GGT2.rev.compl                      aatttgtatgaaatcctcagcagttgaccccaaacagataagactgaagcagtattttag   12702
                                                                   ******* ****************************************************

chimp.LOC112206744.LOC107973052.GGT2-.646279-708040.rev.compl      gaatagttgaaagtatgatcaccacaaaacataagcgtaatcaaatcctgcaagctacat   14795
FAM230B-LOC105372935-GGT2.NCBI.GGT2.rev.compl                      gaatagttgaaagtatgatcaccacaaaacataagcgtaatcaaatcctgcaagctacat   12762
                                                                   ************************************************************

chimp.LOC112206744.LOC107973052.GGT2-.646279-708040.rev.compl      gtaaagcacaatgacaaataaggcagcaaagggccatctggtgattagttcaccacactt   14855
FAM230B-LOC105372935-GGT2.NCBI.GGT2.rev.compl                      gtaaggcacaatgacaaataaggcagcaaagggccatctggtgattagttcaccacactt   12822
                                                                   **** *******************************************************

chimp.LOC112206744.LOC107973052.GGT2-.646279-708040.rev.compl      gttgcaactgtttgcgctgcagagttaaaacacaccagcattcaacccatgtctcctctc   14915
FAM230B-LOC105372935-GGT2.NCBI.GGT2.rev.compl                      gttgcaactgtttgcgctgcagagttaaaacacaccagcattcaacccgtgtctcctctc   12882
                                                                   ************************************************ ***********

chimp.LOC112206744.LOC107973052.GGT2-.646279-708040.rev.compl      ttgaggtaaactgtcgtatgttggctggcctgaacaagcgtagatattctccatcctcaa   14975
FAM230B-LOC105372935-GGT2.NCBI.GGT2.rev.compl                      ttgaagtaaactgtcgtatgttggctggcctgaacaagcgtagatattctccatcctcaa   12942
                                                                   **** *******************************************************

chimp.LOC112206744.LOC107973052.GGT2-.646279-708040.rev.compl      ttaatatgcatgcatgacaaagaaaaggaggcctggatgaaaaaatattgtgtgattaat   15035
FAM230B-LOC105372935-GGT2.NCBI.GGT2.rev.compl                      ttaatatgcatgcatgacaaagaaaaggaggcctggatgaaaaaatattgtgtgattaat   13002
                                                                   ************************************************************

chimp.LOC112206744.LOC107973052.GGT2-.646279-708040.rev.compl      aattatgctttaattaattttaaaggatataatttcagtacttctaattctcccatcagc   15095
FAM230B-LOC105372935-GGT2.NCBI.GGT2.rev.compl                      aattatgctttaattaattttaaaggatataatttcagtacttctaattctcccatcagc   13062
                                                                   ************************************************************

chimp.LOC112206744.LOC107973052.GGT2-.646279-708040.rev.compl      agttataacaaaggattagtgaataaataccatagactgttttgcctggaattgaatcca   15155
FAM230B-LOC105372935-GGT2.NCBI.GGT2.rev.compl                      agttataacaaaggattagtgaataaataccatagactgttttgcctagaattgaatcca   13122
                                                                   *********************************************** ************

chimp.LOC112206744.LOC107973052.GGT2-.646279-708040.rev.compl      atctatctattaaactttgcttttattcaagtgcaaaatgctaaaacacataactgcagt   15215
FAM230B-LOC105372935-GGT2.NCBI.GGT2.rev.compl                      atctatctattaaactttgcttgtattcaagtgcaaaatgctaaaacacataactgcagt   13182
                                                                   ********************** *************************************

chimp.LOC112206744.LOC107973052.GGT2-.646279-708040.rev.compl      gacagccactgtggatcctcagaggtaaaagtagtcttgggacataaatcctgcaagcta   15275
FAM230B-LOC105372935-GGT2.NCBI.GGT2.rev.compl                      gacagccactgtggatcctcagaggtaaaagtagtcttgggacataaatcctgcaagcta   13242
                                                                   ************************************************************

chimp.LOC112206744.LOC107973052.GGT2-.646279-708040.rev.compl      atattgtttttacaggtttagaaaaccatttagctgggtttcaaacctcacagtgtgagc   15335
FAM230B-LOC105372935-GGT2.NCBI.GGT2.rev.compl                      atattgtttttacaggtttagaaaaccatttagctgggtttcaaacctcacagtgtgagc   13302
                                                                   ************************************************************

chimp.LOC112206744.LOC107973052.GGT2-.646279-708040.rev.compl      agtgggactctcatcaaactatagcatgtgcttcagtaccatttgtagactgattcattc   15395
FAM230B-LOC105372935-GGT2.NCBI.GGT2.rev.compl                      agtgggactctcatcaaactatagcatgtgcttcagtaccatttgtagactgactcattc   13362
                                                                   ***************************************************** ******

chimp.LOC112206744.LOC107973052.GGT2-.646279-708040.rev.compl      ccattttcttaagttgccatcagcaaaatgccagggactctatttcttgctccttagctc   15455
FAM230B-LOC105372935-GGT2.NCBI.GGT2.rev.compl                      ccattttcttcagttgccatcagcaaaatgccagggactctatttcttgctccttagctc   13422
                                                                   ********** *************************************************

chimp.LOC112206744.LOC107973052.GGT2-.646279-708040.rev.compl      ctcgttcttgcctgtctttccacgagggaagattttctagcaggagctcaagctgtgctt   15515
FAM230B-LOC105372935-GGT2.NCBI.GGT2.rev.compl                      ctcgttcttgcctgtctttccacgagggaagattttctagcaggagctcaagctgtgctt   13482
                                                                   ************************************************************

chimp.LOC112206744.LOC107973052.GGT2-.646279-708040.rev.compl      ttaatgaaacacatccacacacactgtcctgttgtccacattaagcagagctccctgaat   15575
FAM230B-LOC105372935-GGT2.NCBI.GGT2.rev.compl                      ttaatgaaacacatccacacacactgtcctgttgtccacattaagcagagctccctgaat   13542
                                                                   ************************************************************

chimp.LOC112206744.LOC107973052.GGT2-.646279-708040.rev.compl      aactcatgaacaaaagcatctatgactaactgttgctctgtgtcctcctagcctctgagg   15635
FAM230B-LOC105372935-GGT2.NCBI.GGT2.rev.compl                      aactcatgaacaaaagcatctatgactaactgttgctctgtgtcctcctagcctctgagg   13602
                                                                   ************************************************************

chimp.LOC112206744.LOC107973052.GGT2-.646279-708040.rev.compl      agtctctagttcacaaggacagaggagatggagagaggccagtcaacgcgagggtaaggt   15695
FAM230B-LOC105372935-GGT2.NCBI.GGT2.rev.compl                      agtctctagttcacaaggacagaggagatggagagaggccagtcaacgcgagggtaaggt   13662
                                                                   ************************************************************

chimp.LOC112206744.LOC107973052.GGT2-.646279-708040.rev.compl      tgccttgctttctctgaaatagaaatgttcctttcttggtgtctttctttttcaactgac   15755
FAM230B-LOC105372935-GGT2.NCBI.GGT2.rev.compl                      tgccttgctttctctgaaatagaaatgttcctttcttggtgtctttctttttcaactgac   13722
                                                                   ************************************************************

chimp.LOC112206744.LOC107973052.GGT2-.646279-708040.rev.compl      tttacatgtgaaaaaatgacaatgtccatgacaggtattaaatgcagttttctgaggggg   15815
FAM230B-LOC105372935-GGT2.NCBI.GGT2.rev.compl                      tttacatgtgaaaaaatgacaatgtccatgacaggtattaaatgcagttttctgaggggg   13782
                                                                   ************************************************************

chimp.LOC112206744.LOC107973052.GGT2-.646279-708040.rev.compl      aggaagaagtgactcttagcaactgatatgtaatccaaaatgacatttagctatgacggc   15875
FAM230B-LOC105372935-GGT2.NCBI.GGT2.rev.compl                      aggaagaagtgactcttagcaactgatatgtaatccaaaatggcatttagctatgacggc   13842
                                                                   ****************************************** *****************

chimp.LOC112206744.LOC107973052.GGT2-.646279-708040.rev.compl      ttcaggttgtagactgtatccttggggtccttgtccttggaagcaatgtcttctccttgg   15935
FAM230B-LOC105372935-GGT2.NCBI.GGT2.rev.compl                      ttcaggttgtggactgtatccttggggtccttgtccttggaagcaatgtcttctccttgg   13902
                                                                   ********** *************************************************

chimp.LOC112206744.LOC107973052.GGT2-.646279-708040.rev.compl      attcagtattttgcacttgccaacctacgtggacctgagagatccaccatccagaagctg   15995
FAM230B-LOC105372935-GGT2.NCBI.GGT2.rev.compl                      attcaggattttgcacttgccaacctacgtggacctgagagatccaccatccagaagctg   13962
                                                                   ****** *****************************************************

chimp.LOC112206744.LOC107973052.GGT2-.646279-708040.rev.compl      atgtcttttccagtgtgtagcctacccttgttttggaggccttgaagttgactacacttt   16055
FAM230B-LOC105372935-GGT2.NCBI.GGT2.rev.compl                      atgtcttttccagtgtgtatcctacccttgttttggaggccttgaagttgactacacttt   14022
                                                                   ******************* ****************************************

chimp.LOC112206744.LOC107973052.GGT2-.646279-708040.rev.compl      ctgatcaagttttcagtattcattgagagaaacagagccttgtgcaaacaatccacaaca   16115
FAM230B-LOC105372935-GGT2.NCBI.GGT2.rev.compl                      ctgatcaagttttcagtattcattgagagaaacagagccttgtgcaaacaatccacaaca   14082
                                                                   ************************************************************

chimp.LOC112206744.LOC107973052.GGT2-.646279-708040.rev.compl      tgacatacccctcaaaaagctttgtttctgtattgcaggtggtgcaggtggcccctctga   16175
FAM230B-LOC105372935-GGT2.NCBI.GGT2.rev.compl                      tgagatacccctgaaaaagctttgtttctgtattgcaggtggtgcaggtggcccctctga   14142
                                                                   *** ******** ***********************************************

chimp.LOC112206744.LOC107973052.GGT2-.646279-708040.rev.compl      ggtgtgaatctagtaagtattctggaatcacttgccaagaaaacaatctggatgccaaga   16235
FAM230B-LOC105372935-GGT2.NCBI.GGT2.rev.compl                      ggcttgaatctagtaagtattctggaatcacttgccaagaaaacaatctggatgccaaga   14202
                                                                   **  ********************************************************

chimp.LOC112206744.LOC107973052.GGT2-.646279-708040.rev.compl      aaggtgtggcatccttgcctggtttcaatgtgaagagcctccctgatcctgggattgtga   16295
FAM230B-LOC105372935-GGT2.NCBI.GGT2.rev.compl                      aaggtgtggcatccttgcctggtttcaatgtgaagagcctccctgatcctgggattgtga   14262
                                                                   ************************************************************

chimp.LOC112206744.LOC107973052.GGT2-.646279-708040.rev.compl      taggaataagtataggggaagtgtttttttaaaacctgaattccccagggaaaaattatg   16355
FAM230B-LOC105372935-GGT2.NCBI.GGT2.rev.compl                      taggaataagtataggggaagtgtttttttaaaacctgaattccccagggaaaaattatg   14322
                                                                   ************************************************************

chimp.LOC112206744.LOC107973052.GGT2-.646279-708040.rev.compl      gccaaattttgaggaagcagttgtgctcccttttgggtggtgccgagttgggtgcttgag   16415
FAM230B-LOC105372935-GGT2.NCBI.GGT2.rev.compl                      gccaaattttgaggaagcagctgtgctcccttttgggtggtgctgagttgggtgcttgag   14382
                                                                   ******************** ********************** ****************

chimp.LOC112206744.LOC107973052.GGT2-.646279-708040.rev.compl      gattggtggtgtcttgtgtgaggctgcatcgtgtggtgtgaatatgtgtgtttctgtaca   16475
FAM230B-LOC105372935-GGT2.NCBI.GGT2.rev.compl                      gattggtggtgtcttgtgtgaggctgcatcgtgtggtgtgaatgtgtgtgtttctgtaca   14442
                                                                   ******************************************* ****************

chimp.LOC112206744.LOC107973052.GGT2-.646279-708040.rev.compl      ggtgaggctgtgtgttttctcaggagagatttcccacttatacaacccaatcaccagtgt   16535
FAM230B-LOC105372935-GGT2.NCBI.GGT2.rev.compl                      ggtgaggctgtgtgttttctcaggagagatttcccacttatacaacccaatcaccagtgt   14502
                                                                   ************************************************************

chimp.LOC112206744.LOC107973052.GGT2-.646279-708040.rev.compl      ccacttctaacaataaaatccacccccgctctactctctctgtacagtgactcctccacc   16595
FAM230B-LOC105372935-GGT2.NCBI.GGT2.rev.compl                      ccacttctaacaataaaatccacccccgctctactctctctgtacagtgactcctccacc   14562
                                                                   ************************************************************

chimp.LOC112206744.LOC107973052.GGT2-.646279-708040.rev.compl      ctcactagagccatccccgggtctgccttattatccccactgctcaggtggagaacctga   16655
FAM230B-LOC105372935-GGT2.NCBI.GGT2.rev.compl                      ctcaccagagccatccccgggtctgccttattatccccactgctcaggtggagaacctga   14622
                                                                   ***** ******************************************************

chimp.LOC112206744.LOC107973052.GGT2-.646279-708040.rev.compl      agggccaagggagtggccccagctcccgagttcctgaatgaaaaagtgaaaacacgaacc   16715
FAM230B-LOC105372935-GGT2.NCBI.GGT2.rev.compl                      agggccaagggagtggccccagctcccgagttcctgaatgaaacagtgaaaacacgaacc   14682
                                                                   ******************************************* ****************
```

```
chimp.LOC112206744.LOC107973052.GGT2-.646279-708040.rev.compl     cagaagtgtgggccagtgctgatgctgacatgcacttagtcatggggtgttcaccaccac   16775
FAM230B-LOC105372935-GGT2.NCBI.GGT2.rev.compl                     caggagtgtggaccagtgctgacgctgacatgcacttagtcatggggtgttcaccaccac   14742
                                                                  *** ******* ********** *************************************

chimp.LOC112206744.LOC107973052.GGT2-.646279-708040.rev.compl     acagggagtccagcattcatgtataagccctaaagcaacgagcccaaaaggccccagaca   16835
FAM230B-LOC105372935-GGT2.NCBI.GGT2.rev.compl                     acacggagtccagcattcatgtataagccctaaagcaacgagcccaaaaggccccagaca   14802
                                                                  *** ********************************************************

chimp.LOC112206744.LOC107973052.GGT2-.646279-708040.rev.compl     ctgcccatcatcataaagtggcctccgtggtcacacaacccagggcagttataggctcat   16895
FAM230B-LOC105372935-GGT2.NCBI.GGT2.rev.compl                     ctgcccatcatcataaagtggcctccgtggtcacacaacccagggcagttatagcctcat   14862
                                                                  ****************************************************** *****

chimp.LOC112206744.LOC107973052.GGT2-.646279-708040.rev.compl     ctccccacggacaggcatagtcatcagtgtgtcaaaagcacaaagatccccaggtgtttg   16955
FAM230B-LOC105372935-GGT2.NCBI.GGT2.rev.compl                     ctccccacggaccgacatagtcatcagtgtgtcaaaagcacaaagatccccaggtgtttg   14922
                                                                  ************ * *********************************************

chimp.LOC112206744.LOC107973052.GGT2-.646279-708040.rev.compl     gctcagctcacagatccctttttttttttttaacttttaagttcaggggtacatgtgcagg   17015
FAM230B-LOC105372935-GGT2.NCBI.GGT2.rev.compl                     gctcagctcacagatccc-tttttttatttaacttttcagttcaggggtacatgtgcagg   14981
                                                                  ****************** ******* ********** **********************

chimp.LOC112206744.LOC107973052.GGT2-.646279-708040.rev.compl     atgtgcaggtttgctacatatataaatgtgtgtcatgagagtttgttgtacagattattg   17075
FAM230B-LOC105372935-GGT2.NCBI.GGT2.rev.compl                     atgtgcaggtttgctacatatataaatgtgtgttatgagagttttttgtacagattattg   15041
                                                                  ********************************* ********** ***************

chimp.LOC112206744.LOC107973052.GGT2-.646279-708040.rev.compl     catcacccatatattgagcctaatatcagttatttttcctgatcctcccctcctcccac   17135
FAM230B-LOC105372935-GGT2.NCBI.GGT2.rev.compl                     catcacccatatattgagcctaatatcagttatttttcctgatccacccctcctcccac   15101
                                                                  ********************************************* **************

chimp.LOC112206744.LOC107973052.GGT2-.646279-708040.rev.compl     ctcccaccctccagtaggccccacgctcacaaattctaagaggagtgggggaccacaaag   17195
FAM230B-LOC105372935-GGT2.NCBI.GGT2.rev.compl                     ctcccaccctccagtaggccccacgctcacaaattctaagaggagtgggggaccacaaag   15161
                                                                  ************************************************************

chimp.LOC112206744.LOC107973052.GGT2-.646279-708040.rev.compl     gccagtgtggcccacttcagttgtgaagttaatttgctcagcaactggccaaagtctata   17255
FAM230B-LOC105372935-GGT2.NCBI.GGT2.rev.compl                     gccagtgtggcccacttcagttgtgaagttaatttgctcagcaactggccaaagtctata   15221
                                                                  ************************************************************

chimp.LOC112206744.LOC107973052.GGT2-.646279-708040.rev.compl     aggatgggtgatgtattttagtagatttagtaatactatcttcccaagccctaaaatgct   17315
FAM230B-LOC105372935-GGT2.NCBI.GGT2.rev.compl                     aggatgggtgatgtattttagtagatttagtaatactatcttcccaagccctaaaatgct   15281
                                                                  ************************************************************

chimp.LOC112206744.LOC107973052.GGT2-.646279-708040.rev.compl     caaatcctgccagccaaaaatggtgaggagggacagataggaactctgtgtggcacttgg   17375
FAM230B-LOC105372935-GGT2.NCBI.GGT2.rev.compl                     caaatcctgccagccaaaaatggtgaggagggacagataggaactctgtgtggcacttgg   15341
                                                                  ************************************************************

chimp.LOC112206744.LOC107973052.GGT2-.646279-708040.rev.compl     ttattagcctggcttccatcccttagtggcaagtctcttgtatatgtgggttaaagaccc   17435
FAM230B-LOC105372935-GGT2.NCBI.GGT2.rev.compl                     ttattagcctggcttccatcccttagtggcaactctcttgtatatgtgggttaaagaccc   15401
                                                                  ******************************** ***************************

chimp.LOC112206744.LOC107973052.GGT2-.646279-708040.rev.compl     tcagcctcaagccaagcctcctccatgaggagccatctcactattgactggctagtgccg   17495
FAM230B-LOC105372935-GGT2.NCBI.GGT2.rev.compl                     tcagcctcaagccaagcctcctccatgaggagccatctcactattgaccggctagtgccg   15461
                                                                  ************************************************ ***********

chimp.LOC112206744.LOC107973052.GGT2-.646279-708040.rev.compl     ggtatggccaccagcccaactgaaacaaaatgttgctttaaaacaagtgtaaatctcata   17555
FAM230B-LOC105372935-GGT2.NCBI.GGT2.rev.compl                     ggtatggccaccagcccaactgaaacaaaatgttgctttaaaacaagtgtaaatctcata   15521
                                                                  ************************************************************

chimp.LOC112206744.LOC107973052.GGT2-.646279-708040.rev.compl     catacaacaggcaaatgcagaagcagtgtggtctcgcaagttgtaaagaggacagtcgca   17615
FAM230B-LOC105372935-GGT2.NCBI.GGT2.rev.compl                     catacaacaggcaaatgcagaagcagtgtggtctcgcaagttgtaaagaggacagtcgca   15581
                                                                  ************************************************************

chimp.LOC112206744.LOC107973052.GGT2-.646279-708040.rev.compl     attttgctggacttcaacctgggtagaagacatgagggaactctgtcactaaatcacggc   17675
FAM230B-LOC105372935-GGT2.NCBI.GGT2.rev.compl                     attttgctggacttcaacctgggtagaagacatgagggaac-ctgtcactaaatcacggc   15640
                                                                  **************************************** *******************

chimp.LOC112206744.LOC107973052.GGT2-.646279-708040.rev.compl     agagttcaaggccacttgcagactatttcatgttacagaaggtggcctttagctactaag   17735
FAM230B-LOC105372935-GGT2.NCBI.GGT2.rev.compl                     agagttcaaggccacttgtagactatttcatgttacagaaggtggcctttagctactaag   15700
                                                                  ****************** *****************************************

chimp.LOC112206744.LOC107973052.GGT2-.646279-708040.rev.compl     caaaggcctctgtttctcatttctttcctgttcatcttcttggtcatccttcttccgcaa   17795
FAM230B-LOC105372935-GGT2.NCBI.GGT2.rev.compl                     caaaggcctctgtttctcatttctttcctgttcatcttcttggtcatccttcttccgcaa   15760
                                                                  ************************************************************

chimp.LOC112206744.LOC107973052.GGT2-.646279-708040.rev.compl     gggaaacgagcccaagcaaaaggcagtttcaatattaatttgaccgaggttttgtgcagt   17855
FAM230B-LOC105372935-GGT2.NCBI.GGT2.rev.compl                     gggaaacgagcccaagcaaaaggcagtttcaatattaatttgaccgaggttttgtgcagt   15820
                                                                  ************************************************************

chimp.LOC112206744.LOC107973052.GGT2-.646279-708040.rev.compl     ttattatcatccaggtaatcaagtgcaacccagtctgcctagcagccccctatctctgc   17915
FAM230B-LOC105372935-GGT2.NCBI.GGT2.rev.compl                     ttattatcatccaggtaatcaggtgcaacccagtctgcctagccgcccccctatctctgc   15880
                                                                  ********************* ********************* ****************

chimp.LOC112206744.LOC107973052.GGT2-.646279-708040.rev.compl     tctgtgttttcatttaataaacattttggtctacttactatgtgctagattttctcgaga   17975
FAM230B-LOC105372935-GGT2.NCBI.GGT2.rev.compl                     tctgtgttttcatttaataaacattttggtctacttactatgtgctagattttctcgaga   15940
                                                                  ************************************************************

chimp.LOC112206744.LOC107973052.GGT2-.646279-708040.rev.compl     ccaagtaaatgagatagaatctttatgttggcagctaagttagatttaacataactgaca   18035
FAM230B-LOC105372935-GGT2.NCBI.GGT2.rev.compl                     ccaagtaaatgagatagaatctttatgctggcagctaagttagatttcacataactgaca   16000
                                                                  *************************** ******************* ************

chimp.LOC112206744.LOC107973052.GGT2-.646279-708040.rev.compl     aaaattaaaatttctgatttcttgtaaaaatattttgtatgtgtgaatgcatactgaatg   18095
FAM230B-LOC105372935-GGT2.NCBI.GGT2.rev.compl                     aaaattaaaatttctgatttcttgtaaaaatattttgtatgtgtgaatgcatactgaatg   16060
                                                                  ************************************************************

chimp.LOC112206744.LOC107973052.GGT2-.646279-708040.rev.compl     taaagtggataaaaaactcacacttgcactcatggaaggcttttcatgaatttgtcaatt   18155
FAM230B-LOC105372935-GGT2.NCBI.GGT2.rev.compl                     taaagtggataaaaaactcacacttgcactcatggaaggcttttcatgaatttgtcaatt   16120
                                                                  ************************************************************

chimp.LOC112206744.LOC107973052.GGT2-.646279-708040.rev.compl     tttatttttatatttccccacttcaccggataatgcatacctgaacctggaaattgatt   18215
FAM230B-LOC105372935-GGT2.NCBI.GGT2.rev.compl                     tttatttttatatttccccacttcaccagataatgcatacctgaacctggaaactgatt   16180
                                                                  **************************** ************************* *****

chimp.LOC112206744.LOC107973052.GGT2-.646279-708040.rev.compl     cccactgcagaaagtgttctgagccacatcccttagcttcactagtgcaggtccacctgg   18275
FAM230B-LOC105372935-GGT2.NCBI.GGT2.rev.compl                     cccactgcagaaagtgttctgagccacatcccttagcttcactagtgcaggtccacctgg   16240
                                                                  ************************************************************

chimp.LOC112206744.LOC107973052.GGT2-.646279-708040.rev.compl     gaggatgtcccagcatcagcttggcccatgctgtgatcagccacctccatgcaccacacc   18335
FAM230B-LOC105372935-GGT2.NCBI.GGT2.rev.compl                     gaggatgtcccagcatcagcttgacccatgctgtgatcagccacctccatgcatcacacc   16300
                                                                  ********************** ****************************** ******

chimp.LOC112206744.LOC107973052.GGT2-.646279-708040.rev.compl     aagcaagcccctgggtgattcacagtctccaccaccagggcactgaccttaactgtgtgt   18395
FAM230B-LOC105372935-GGT2.NCBI.GGT2.rev.compl                     aagcaagcccctgggtgattcacagtctccaccaccagggcactgaccttaactctgtgt   16360
                                                                  ****************************************************** *****

chimp.LOC112206744.LOC107973052.GGT2-.646279-708040.rev.compl     tcttctagctccccatgaagacaccgtacacgacatcactaacgaggacgccacacacga   18455
FAM230B-LOC105372935-GGT2.NCBI.GGT2.rev.compl                     ttttctagctccccatgaggacaccgtacacgacatcactaacgaggacgccacacacga   16420
                                                                  * **************** *****************************************

chimp.LOC112206744.LOC107973052.GGT2-.646279-708040.rev.compl     catcgccaacgaggacactgtacatgacaccgctaacgaggccgccgacaagggcatcgc   18515
FAM230B-LOC105372935-GGT2.NCBI.GGT2.rev.compl                     catcgctaacgaggatactgtacacgacatcgctaacgaggccgccgacaagggcatcgc   16480
                                                                  ****** ******** ******** **** ******************************

chimp.LOC112206744.LOC107973052.GGT2-.646279-708040.rev.compl     cagcgaggacgccgcccagggcatcgccaacgaggccgccgcccagggcatcgtcagcga   18575
FAM230B-LOC105372935-GGT2.NCBI.GGT2.rev.compl                     caacgaggatgccgcccacggcatcgccaacgagtacgccgcccacggcatcgccagcga   16540
                                                                  ** ****** ******** ***************  ********* ******* ******

chimp.LOC112206744.LOC107973052.GGT2-.646279-708040.rev.compl     ggacgccgcccacggaatcgccaacgaggacgccgtccagggcatcgccaacgtggtcgc   18635
FAM230B-LOC105372935-GGT2.NCBI.GGT2.rev.compl                     ggacgccgcccacggcatcgccagcgaggacgccgcccagggcatcgccagcgaggacgc   16600
                                                                  *************** ******* *********** ************** ** ** ***

chimp.LOC112206744.LOC107973052.GGT2-.646279-708040.rev.compl     cgcccacggaatcgccaacgaggacgccgcccagggcatcgc------------------   18677
FAM230B-LOC105372935-GGT2.NCBI.GGT2.rev.compl                     cgcccagggcatcgccaacgaggacaccatccagggcatcgccaaggagtacgccgtcca   16660
                                                                  ****** ** *************** **  ************

chimp.LOC112206744.LOC107973052.GGT2-.646279-708040.rev.compl     ------------------------------------------------------------   18677
FAM230B-LOC105372935-GGT2.NCBI.GGT2.rev.compl                     cggcatcgccaacgaggacgccgcccagggcatcgccaacgaggacgccgcccagggcat   16720

chimp.LOC112206744.LOC107973052.GGT2-.646279-708040.rev.compl     ------------------------------------------------------------   18677
FAM230B-LOC105372935-GGT2.NCBI.GGT2.rev.compl                     cgccaacgaggacgccgcccagggcatcgccaaggaggacgccgcccagggcatcgccaa   16780

chimp.LOC112206744.LOC107973052.GGT2-.646279-708040.rev.compl     ------------------------------------------------------------   18677
FAM230B-LOC105372935-GGT2.NCBI.GGT2.rev.compl                     cgagggcgccgcccagggcatcgccaaagaggacgccgcccagggcatcgctaacgaggc   16840

chimp.LOC112206744.LOC107973052.GGT2-.646279-708040.rev.compl     ------------------------------------------------------------   18677
```

```
FAM230B-LOC105372935-GGT2.NCBI.GGT2.rev.compl                      cgccgacaagggcatcgccaacgaggatgccgcccacggcatcgccaacgaggacgccgc   16900

chimp.LOC112206744.LOC107973052.GGT2-.646279-708040.rev.compl      ------------------------------------------------------------   18677
FAM230B-LOC105372935-GGT2.NCBI.GGT2.rev.compl                      ccacggcatcgccagcgaggacgccgcccacggcatcgccagcgaggacgccgcccacgg   16960

chimp.LOC112206744.LOC107973052.GGT2-.646279-708040.rev.compl      ---------------------------catcgctaatgaggatgccatatatgacatcgc   18710
FAM230B-LOC105372935-GGT2.NCBI.GGT2.rev.compl                      catcgccagcgaggacgccgcccagggcatcgccaacgaggacgccatccagggcatcgc   17020
                                                                                              ****** ** ***** *****  * * ******

chimp.LOC112206744.LOC107973052.GGT2-.646279-708040.rev.compl      taatg---acaccgtacaaggcat------------------------------------   18731
FAM230B-LOC105372935-GGT2.NCBI.GGT2.rev.compl                      caaggagtacgccgtccacggcatcgccaacgaggacgccgcccagggcatcgccaacga   17080
                                                                    ** *   ** **** ** *****

chimp.LOC112206744.LOC107973052.GGT2-.646279-708040.rev.compl      ------------------------------------------------------------   18731
FAM230B-LOC105372935-GGT2.NCBI.GGT2.rev.compl                      ggacgccgcccagggcatcgccaacgaggacgccgcccagggcatcgccaaggaggacgc   17140

chimp.LOC112206744.LOC107973052.GGT2-.646279-708040.rev.compl      ------------------------------------------------------------   18731
FAM230B-LOC105372935-GGT2.NCBI.GGT2.rev.compl                      cgcccagggcatcgccaaggaggacgccgcccagggcatcgccaacgagggcgccgccca   17200

chimp.LOC112206744.LOC107973052.GGT2-.646279-708040.rev.compl      ------------------------------------------------------------   18731
FAM230B-LOC105372935-GGT2.NCBI.GGT2.rev.compl                      gggcatcgccaaagaggacgccgcccacggcatcgccaacgaggacgccgcccagggcat   17260

chimp.LOC112206744.LOC107973052.GGT2-.646279-708040.rev.compl      ------------------------------------------------------------   18731
FAM230B-LOC105372935-GGT2.NCBI.GGT2.rev.compl                      cgccaacgaggacgccgcccacggaatcgccagtgaggacgccgcccacggcatcgccag   17320

chimp.LOC112206744.LOC107973052.GGT2-.646279-708040.rev.compl      ------------------------------------------------------------   18731
FAM230B-LOC105372935-GGT2.NCBI.GGT2.rev.compl                      cgaggacgccgtccagggcatcgacaaggaggatgccgcccagggcatcgccaacgagga   17380

chimp.LOC112206744.LOC107973052.GGT2-.646279-708040.rev.compl      ------------------------------------------------------------   18731
FAM230B-LOC105372935-GGT2.NCBI.GGT2.rev.compl                      cgccgcccagggcatcgccaacgaggacgccgcccagggcatcgccaaagaggacgccgc   17440

chimp.LOC112206744.LOC107973052.GGT2-.646279-708040.rev.compl      ---------------------------------gctaacgaagacatcgtacaagccgtc   18758
FAM230B-LOC105372935-GGT2.NCBI.GGT2.rev.compl                      ccacggcatcgccaacgagctgtatacgacatcgctaacgaggacaccctacaagccgtc   17500
                                                                                                    ******** **** * ***********

chimp.LOC112206744.LOC107973052.GGT2-.646279-708040.rev.compl      gctaacaagtacactgtacacaacatcgctaatgagggcactgtacaagacatcaccaat   18818
FAM230B-LOC105372935-GGT2.NCBI.GGT2.rev.compl                      gctaacaagtacactgtacacaacatcgctaatgagggcactgtacaagacatcaccaat   17560
                                                                   ************************************************************

chimp.LOC112206744.LOC107973052.GGT2-.646279-708040.rev.compl      gaggacgctttatacgacattgctaatggcaccgacaaggcacgctaacgtggatgctgt   18878
FAM230B-LOC105372935-GGT2.NCBI.GGT2.rev.compl                      gagggcgctttatacgacattgctaatggcaccgacaaggcacgctaacgtggacgctgt   17620
                                                                   **** ************************************************* *****

chimp.LOC112206744.LOC107973052.GGT2-.646279-708040.rev.compl      acacgacattgctaatgaggacaccgtataagacatcgctagtaactatcgcaagaacaa   18938
FAM230B-LOC105372935-GGT2.NCBI.GGT2.rev.compl                      acacgacattgctaatgaggacagcgtataagacatcgctagtaactatcgcaagaacaa   17680
                                                                   *********************** ************************************

chimp.LOC112206744.LOC107973052.GGT2-.646279-708040.rev.compl      aaaaccaaacaccgcatattctcactcataggtgggaattgaacaatgagatcacatgga   18998
FAM230B-LOC105372935-GGT2.NCBI.GGT2.rev.compl                      aaaaccaaacaccgcatattctcactcatagatgggaattgaacaatgagatcacatgga   17740
                                                                   ****************************** *****************************

chimp.LOC112206744.LOC107973052.GGT2-.646279-708040.rev.compl      cacaggaaggggaatatcacactctggggactgttgtggggtggggggaggggggaggga   19058
FAM230B-LOC105372935-GGT2.NCBI.GGT2.rev.compl                      cacaggaaggggaatatcacactctggggactgttgtggggttgg----ggggggaggga   17796
                                                                   ****************************************** **    ***********

chimp.LOC112206744.LOC107973052.GGT2-.646279-708040.rev.compl      tagcatcgggagatatacctaatgctagatgatgagttagtgggtgcagcgcaccagcat   19118
FAM230B-LOC105372935-GGT2.NCBI.GGT2.rev.compl                      tagcatcgggagatatacctaatgctagatgatgagttagtgggtgcagcgcaccagcac   17856
                                                                   ***********************************************************

chimp.LOC112206744.LOC107973052.GGT2-.646279-708040.rev.compl      ggcacatgtatacatatgtaactaacctgcacaatgtgcacatgtaccctaaaacttaaa   19178
FAM230B-LOC105372935-GGT2.NCBI.GGT2.rev.compl                      ggcacatgtatacatatgtaactaacctgcacaatgtgcacatgtaccctaaaacttaaa   17916
                                                                   ************************************************************

chimp.LOC112206744.LOC107973052.GGT2-.646279-708040.rev.compl      gtatatatataaaaaaaaagacatcgctagtgagcacgctgtatacgacatcactaatga   19238
FAM230B-LOC105372935-GGT2.NCBI.GGT2.rev.compl                      gtatatatataaaaaaaaagacatcgctagtgagcacgc---------------------   17955
                                                                   ***************************************

chimp.LOC112206744.LOC107973052.GGT2-.646279-708040.rev.compl      ggatgctatatatgacatcgctgatgaggacattgtatacgacatcactaatgaggacac   19298
FAM230B-LOC105372935-GGT2.NCBI.GGT2.rev.compl                      ---------------------------------tgtatacgacatcgctaatgaggacac   17982
                                                                                                    ************* *************

chimp.LOC112206744.LOC107973052.GGT2-.646279-708040.rev.compl      catacaaggcatcgctaacaatgatgctgtacacaacatcactaatgatgacaccgtata   19358
FAM230B-LOC105372935-GGT2.NCBI.GGT2.rev.compl                      catacaaggcatcgctaacgatgacgctgtacacaacatcactaatgatgacaccgtata   18042
                                                                   ****************** **** ************************************

chimp.LOC112206744.LOC107973052.GGT2-.646279-708040.rev.compl      acacatcgctaattatgacgctgtatacgacatcgctaatgacaccgtacaaggcacact   19418
FAM230B-LOC105372935-GGT2.NCBI.GGT2.rev.compl                      agacatcgctaattatgacgctgtatacgacatcgctaatgacaccgtacaaggcacgct   18102
                                                                   * ******************************************************* **

chimp.LOC112206744.LOC107973052.GGT2-.646279-708040.rev.compl      aatgaggatgctgtacacgacatcactaatgaggacagtgtacaagccatcgctaatgag   19478
FAM230B-LOC105372935-GGT2.NCBI.GGT2.rev.compl                      aacgaggatgctgtacacgacatcactaatgaggacagtgtacaagccatcactaatgag   18162
                                                                   ** ************************************************ ********

chimp.LOC112206744.LOC107973052.GGT2-.646279-708040.rev.compl      gacactgtatatgacatcgctaacgaggacactgtacaaggcattgctaacgaggacgct   19538
FAM230B-LOC105372935-GGT2.NCBI.GGT2.rev.compl                      gacactgtatatggcatcgctaacgaggacactgtacaagccattgctaacgaggacgct   18222
                                                                   ************* ************************** *******************

chimp.LOC112206744.LOC107973052.GGT2-.646279-708040.rev.compl      gtac---acatcgctaatgaggacaccatataagacatcaccaatgaggatgctgtatat   19595
FAM230B-LOC105372935-GGT2.NCBI.GGT2.rev.compl                      gtacacaacatcgctaatgaggacaccatataagacatcaccaatgaggatgctgtatat   18282
                                                                   ****   *****************************************************

chimp.LOC112206744.LOC107973052.GGT2-.646279-708040.rev.compl      gacatcactaatgacacccacaaggcatgctaacgaggacgctgtagacgacattgctta   19655
FAM230B-LOC105372935-GGT2.NCBI.GGT2.rev.compl                      gacatcgctaatggcacccacaaggcatgctaacgaggacgctgtagacgacattgctta   18342
                                                                   ****** ****** **********************************************

chimp.LOC112206744.LOC107973052.GGT2-.646279-708040.rev.compl      taaggacaccgtacaagacatcgctaacgaggacgctgtatacgacatcgctaatgagga   19715
FAM230B-LOC105372935-GGT2.NCBI.GGT2.rev.compl                      taaggacaccgtacaagacatcgctaacgaggacgctgtatacgacatcgctaatgagga   18402
                                                                   ************************************************************

chimp.LOC112206744.LOC107973052.GGT2-.646279-708040.rev.compl      cgttgtatatgacatcgctaatgaggatgctttacaagacatagctaatgaggttgctgt   19775
FAM230B-LOC105372935-GGT2.NCBI.GGT2.rev.compl                      cgttgtatatgacatcgctaatgaggatgctttacaagacatagctaatgaggttgctgt   18462
                                                                   ************************************************************

chimp.LOC112206744.LOC107973052.GGT2-.646279-708040.rev.compl      atatgatatcgctaatgaggacattgtatatgacatcgctaatgaggacgctctatacga   19835
FAM230B-LOC105372935-GGT2.NCBI.GGT2.rev.compl                      atatgacatcgctaatgaggacattgtatatgacatcgctaatgaggacgctctatacga   18522
                                                                   ****** *****************************************************

chimp.LOC112206744.LOC107973052.GGT2-.646279-708040.rev.compl      catcactaatgaggatgctgtatacaacatcgctaatgaggacgctgtatatggcatcgc   19895
FAM230B-LOC105372935-GGT2.NCBI.GGT2.rev.compl                      catcactaatgaggacgctgtatacaacatcgctaatgaggacgctgtatatggcatcgc   18582
                                                                   *************** ********************************************

chimp.LOC112206744.LOC107973052.GGT2-.646279-708040.rev.compl      taatgaggatgctgtatacgaattcgctaataaggacgctgtatatgacattgctaatga   19955
FAM230B-LOC105372935-GGT2.NCBI.GGT2.rev.compl                      taatgaggatgctgtatacgaattcgctaataaggacgctgtatatgacattgctaatga   18642
                                                                   ************************************************************

chimp.LOC112206744.LOC107973052.GGT2-.646279-708040.rev.compl      ggacactatacaagacatctgtaaaaaagaagatgctgccaatgtaagacacttttcttt   20015
FAM230B-LOC105372935-GGT2.NCBI.GGT2.rev.compl                      ggacactgtacaagacatctgtaaaaaagaagatgctgccaatgtaagacacttttcttt   18702
                                                                   ******* ****************************************************

chimp.LOC112206744.LOC107973052.GGT2-.646279-708040.rev.compl      gtcttgaacagaaatgttactttcctggcttctttccaatcagatgtagacatgaacatc   20075
FAM230B-LOC105372935-GGT2.NCBI.GGT2.rev.compl                      gtcttgaacagaaatgttactttcctggcttctttccaatcagatgtagacatgaacatc   18762
                                                                   ************************************************************

chimp.LOC112206744.LOC107973052.GGT2-.646279-708040.rev.compl      tgccagtgtgcattattgatgtcatctgcagtttaatcaaatgtagacatgaacatctgc   20135
FAM230B-LOC105372935-GGT2.NCBI.GGT2.rev.compl                      tgccagtgtgcattatcgatgtcatctgcagtttaatcaaatgtagacatgaacatctgc   18822
                                                                   **************** *******************************************

chimp.LOC112206744.LOC107973052.GGT2-.646279-708040.rev.compl      caatgtggactatttatgacatctgcaattcccttggtgtggtgctattgattggcagcc   20195
FAM230B-LOC105372935-GGT2.NCBI.GGT2.rev.compl                      caatgtggactatttatgacatctgcaattcccttggtgtggtgctattgattggcagcc   18882
                                                                   ************************************************************

chimp.LOC112206744.LOC107973052.GGT2-.646279-708040.rev.compl      tctcaccaacccatgccaggcacactggggtgtggtagatggcagcatccacgatccact   20255
FAM230B-LOC105372935-GGT2.NCBI.GGT2.rev.compl                      tctcaccaacccatgccaggcacactggggtgtggtagatggcagcatccacgatccact   18942
                                                                   ************************************************************

chimp.LOC112206744.LOC107973052.GGT2-.646279-708040.rev.compl      gcaatgcagaggtgtttccctccacagcagtttttccccatggattaagagttgtgaaac   20315
FAM230B-LOC105372935-GGT2.NCBI.GGT2.rev.compl                      gcaatgcagaggtgtttccctccacagcagtttttccccatggattaagagttgtgaaac   19002
```

```
                                                               ************************************************************

chimp.LOC112206744.LOC107973052.GGT2-.646279-708040.rev.compl      tgccaatctagatacactttaaagataaattctgtgggaaaaggtcttgtcttttccaca   20375
FAM230B-LOC105372935-GGT2.NCBI.GGT2.rev.compl                      tgccaatctagatacactttaaagataaattctgtgggaaaaggtcttgtcttttccaca   19062
                                                                   ************************************************************

chimp.LOC112206744.LOC107973052.GGT2-.646279-708040.rev.compl      ggtgtcttccgtgccagttttggggacttcgacctttgactcaatcactatacccttc   20435
FAM230B-LOC105372935-GGT2.NCBI.GGT2.rev.compl                      ggtgtcttccgtgccagttttggggacttcgacctttgactcaatcactatacccttc   19122
                                                                   ************************************************************

chimp.LOC112206744.LOC107973052.GGT2-.646279-708040.rev.compl      ttattttctctctcaagttgtcgagagactatcagatctgtgtgacgtgtatggcatcat   20495
FAM230B-LOC105372935-GGT2.NCBI.GGT2.rev.compl                      ctattttctctctcaagttgtcgagagactatcagatctgtgtgacgtgtatggcatcat   19182
                                                                    ***********************************************************

chimp.LOC112206744.LOC107973052.GGT2-.646279-708040.rev.compl      ttcaccctcctaatgttttcttttctataattgcaggagccattgacattggagaatgat   20555
FAM230B-LOC105372935-GGT2.NCBI.GGT2.rev.compl                      ttcaccctcctaatgttttcttttctataattgcaggagccattgacactggagaatgat   19242
                                                                   ************************************************ ***********

chimp.LOC112206744.LOC107973052.GGT2-.646279-708040.rev.compl      acgtaccctgaaataactcacttcctgaggaaaaagcgccatctctagggtacagaaacc   20615
FAM230B-LOC105372935-GGT2.NCBI.GGT2.rev.compl                      acgtaccctgaaataactcacttcctgaggaaaaagcgccatctctagggtacagaaacc   19302
                                                                   ************************************************************

chimp.LOC112206744.LOC107973052.GGT2-.646279-708040.rev.compl      tgattctgggctccttttgggaaggaggatttggggtctggtgagagcaaatgattttgc   20675
FAM230B-LOC105372935-GGT2.NCBI.GGT2.rev.compl                      tgattctgggctccttttgggaaggaggatttggggtctggtgagagcaaatgattttgc   19362
                                                                   ************************************************************

chimp.LOC112206744.LOC107973052.GGT2-.646279-708040.rev.compl      aagtataaaacaatgtccagagaggctgtagggatatctgtgagcccagaggaaacacca   20735
FAM230B-LOC105372935-GGT2.NCBI.GGT2.rev.compl                      aagtataaaaccatgtccagagaggctgtagggatatctgtgagcccagaggaaacacca   19422
                                                                   *********** ************************************************

chimp.LOC112206744.LOC107973052.GGT2-.646279-708040.rev.compl      ggggatcctgtgcgaagcaccatggcttcagctagggtgggaggagtgggtggcctctc   20795
FAM230B-LOC105372935-GGT2.NCBI.GGT2.rev.compl                      ggggatcctgtgcgaagcaccatggcttcagctagggtgggaggagtgggtggcctctc   19482
                                                                   ************************************************************

chimp.LOC112206744.LOC107973052.GGT2-.646279-708040.rev.compl      tctaatgacttatcctggtgtttgtgtttctaaagatttgattgtggagagcatatctga   20855
FAM230B-LOC105372935-GGT2.NCBI.GGT2.rev.compl                      tctaatgacttatcctggtgtttgtgtttctaaagatttgattgtggagagcatatctga   19542
                                                                   ************************************************************

chimp.LOC112206744.LOC107973052.GGT2-.646279-708040.rev.compl      tgatggggatttgtaggtaggtaactactttccacgtaagatccaattggagagagttcc   20915
FAM230B-LOC105372935-GGT2.NCBI.GGT2.rev.compl                      tgatggggatttgtaggtaggtaactgctttccacgtaagatccaattggagagagttcc   19602
                                                                   ************************** *********************************

chimp.LOC112206744.LOC107973052.GGT2-.646279-708040.rev.compl      caggggccttcggggtatccatgctgcttgggaggttaagggagggggcatgaaatcaaa   20975
FAM230B-LOC105372935-GGT2.NCBI.GGT2.rev.compl                      caggggccttcggggtatccatgctgcttgggaggttaagggagggggcatgaaatcaaa   19662
                                                                   ************************************************************

chimp.LOC112206744.LOC107973052.GGT2-.646279-708040.rev.compl      acgaaacaggaaatatgtgtcatattggatttggtcttttccgggtttattggcataata   21035
FAM230B-LOC105372935-GGT2.NCBI.GGT2.rev.compl                      acgaaacaggaaatatgtgtcatatcggatttggtcttttccgggtttattggcataata   19722
                                                                   ************************* **********************************

chimp.LOC112206744.LOC107973052.GGT2-.646279-708040.rev.compl      gtaagaactgtctctctgggctatgagggtgctgtgttatttaaaggtggtctttcccag   21095
FAM230B-LOC105372935-GGT2.NCBI.GGT2.rev.compl                      gttagaactgtctctctgggctatgagggtgctgtgttatttaaaggtggtctttcccag   19782
                                                                   ** *********************************************************

chimp.LOC112206744.LOC107973052.GGT2-.646279-708040.rev.compl      aacacctggccttttcttttccgcctctgccaaacatcacagactttgggttggattagt   21155
FAM230B-LOC105372935-GGT2.NCBI.GGT2.rev.compl                      aacacctggccttttcttttctgcctctgccaaacatcacagcctttgggttggattagt   19842
                                                                   ********************* ******************** *****************

chimp.LOC112206744.LOC107973052.GGT2-.646279-708040.rev.compl      cagcacccctttgggattgtgcagaagaggtttggggttgcatcgagtgtcacctgtggtg   21215
FAM230B-LOC105372935-GGT2.NCBI.GGT2.rev.compl                      cagcacccctttgggattgtgcagaagaggtttggggttgcattgagtgtcacctgtggtg   19902
                                                                   ****************************************** *****************

chimp.LOC112206744.LOC107973052.GGT2-.646279-708040.rev.compl      aacagaatctgagggacacaactctctcacaggcacttccttcaacctggagacagagtt   21275
FAM230B-LOC105372935-GGT2.NCBI.GGT2.rev.compl                      aacagaatctgagggacacaactctctcacaggcacttcctccaacctggagacagagtt   19962
                                                                   ***************************************** ******************

chimp.LOC112206744.LOC107973052.GGT2-.646279-708040.rev.compl      ctcctggtgtgtgcccaggggtgcaggagaaattgacagtctgcctctgaactttcagga   21335
FAM230B-LOC105372935-GGT2.NCBI.GGT2.rev.compl                      ctcctggtgtgtgcccaggggtggaggagaaattgacagtctgcctctgaactttcagga   20022
                                                                   *********************** ************************************

chimp.LOC112206744.LOC107973052.GGT2-.646279-708040.rev.compl      ctttaaaaagcactcatgtttccatcctcgctgttgactcctggcttaaagggatctcca   21395
FAM230B-LOC105372935-GGT2.NCBI.GGT2.rev.compl                      ctttaaaaagcactcatgtttccatcctcactgttgactcctggcttaaagggatctccc   20082
                                                                   ***************************** ******************************

chimp.LOC112206744.LOC107973052.GGT2-.646279-708040.rev.compl      ggggtgagtgaggaggagggatcggaccctggcagtctgacggcagcacctgtgttcctc   21455
FAM230B-LOC105372935-GGT2.NCBI.GGT2.rev.compl                      ggggtgagtgaggaggcgggatcggaccctggcagtctgacggcagcacctgtgttcctc   20142
                                                                   **************** *******************************************

chimp.LOC112206744.LOC107973052.GGT2-.646279-708040.rev.compl      tgcactgggccgtggatgacattacacaccttggtgagaatcaggaattgaggctaacca   21515
FAM230B-LOC105372935-GGT2.NCBI.GGT2.rev.compl                      tgcactgggccgtggatgacattacataccttggtgagaatcaggaattgaggctaacca   20202
                                                                   ************************** *********************************

chimp.LOC112206744.LOC107973052.GGT2-.646279-708040.rev.compl      catctgaaattgagatgggccttgagtcatataaatagtttggaaaagatgcattttact   21575
FAM230B-LOC105372935-GGT2.NCBI.GGT2.rev.compl                      catctgaaattgagatgggccttgagtcatataaatagtttggaaaagatgcattttact   20262
                                                                   ************************************************************

chimp.LOC112206744.LOC107973052.GGT2-.646279-708040.rev.compl      acgctattgaaagaaaccatttatttctcactccagcaggataaatggttttcagtatcc   21635
FAM230B-LOC105372935-GGT2.NCBI.GGT2.rev.compl                      acgctattgaaagaaaccatttatttctcactccagcaggataaatggttttcagtatcc   20322
                                                                   ************************************************************

chimp.LOC112206744.LOC107973052.GGT2-.646279-708040.rev.compl      atttaactgctcattgactcttactgtagatgaggaggtggccagcagcccctgccctcc   21695
FAM230B-LOC105372935-GGT2.NCBI.GGT2.rev.compl                      atttaactgctcattgactcttactgtagatgaggaggtggccagcagcccctgccctcc   20382
                                                                   ************************************************************

chimp.LOC112206744.LOC107973052.GGT2-.646279-708040.rev.compl      cccagtt-gtaggcccaaggtaaccagcaatggactggatataatggaagagtggtgcat   21754
FAM230B-LOC105372935-GGT2.NCBI.GGT2.rev.compl                      cccagttggtaggcccaaggtaaccagcaattgactggatataatggaagagtggtgcat   20442
                                                                   ******* *********************** ****************************

chimp.LOC112206744.LOC107973052.GGT2-.646279-708040.rev.compl      tcggaggtatctgtattaatgggacccacatgatatggatgagagctattagggtgagaa   21814
FAM230B-LOC105372935-GGT2.NCBI.GGT2.rev.compl                      tcgggggtatctgtattaatgggacccacatgatatggatgagagctattagggtgagaa   20502
                                                                   **** *******************************************************

chimp.LOC112206744.LOC107973052.GGT2-.646279-708040.rev.compl      aaagcctgggagcacaatgaaatatttaaatattaaacaaaacattgttgaaatctccat   21874
FAM230B-LOC105372935-GGT2.NCBI.GGT2.rev.compl                      aaagcctgggagcacaatgaaatatttaaatattaaacaaaacattgttgaaatctccat   20562
                                                                   ************************************************************

chimp.LOC112206744.LOC107973052.GGT2-.646279-708040.rev.compl      tgtactttagtagttgaagtcattcttgtggtcatcactgcctttcccaagcataacaag   21934
FAM230B-LOC105372935-GGT2.NCBI.GGT2.rev.compl                      tgtactttagtagttgaagtcattcttgtggtcatcactgcctttcccaagcataacaag   20622
                                                                   ************************************************************

chimp.LOC112206744.LOC107973052.GGT2-.646279-708040.rev.compl      ctacttaatatcacatagacccgtgccatgaggaatgatgatcagtttgtaaaatgccaa   21994
FAM230B-LOC105372935-GGT2.NCBI.GGT2.rev.compl                      ctacttaatatcacatggacccgtgccatgaggaatgatgatcagtttgtaaaatgccaa   20682
                                                                   **************** *******************************************

chimp.LOC112206744.LOC107973052.GGT2-.646279-708040.rev.compl      taaaacaattgcctatagaagccacaatgtttcatccatatatttcaatttccatgtgta   22054
FAM230B-LOC105372935-GGT2.NCBI.GGT2.rev.compl                      taaaacaattgcctatataagccacaatgtttcatccatatatttcaatttccatgtgta   20742
                                                                   ***************** ******************************************

chimp.LOC112206744.LOC107973052.GGT2-.646279-708040.rev.compl      agtatagttcaaatttcagaaatttattattatctaatagaatatgcatggtatatcaat   22114
FAM230B-LOC105372935-GGT2.NCBI.GGT2.rev.compl                      agtatagttcaaatttcagaaatttattattatctaatagaatatgcatggtatatcaat   20802
                                                                   ************************************************************

chimp.LOC112206744.LOC107973052.GGT2-.646279-708040.rev.compl      gagcaattatcatactgtttctattaacaattatttgtatgatgaaaaaagcagactccc   22174
FAM230B-LOC105372935-GGT2.NCBI.GGT2.rev.compl                      gagcaattatcatactgtttctattaacaattatttgtatgatgaaaaaagcagactccc   20862
                                                                   ************************************************************

chimp.LOC112206744.LOC107973052.GGT2-.646279-708040.rev.compl      attcttggatttttctcagtttgcacacattagcatgacagccccatttccacctgacat   22234
FAM230B-LOC105372935-GGT2.NCBI.GGT2.rev.compl                      attcttggatttttctcagtttgcacacattagcatgacagccccatttccacctgacat   20922
                                                                   ************************************************************

chimp.LOC112206744.LOC107973052.GGT2-.646279-708040.rev.compl      gtgccagcaagaggccaggaacagaggcttttcttat----taagatttctaaatgtatt   22290
FAM230B-LOC105372935-GGT2.NCBI.GGT2.rev.compl                      gtgccagcaagaggccaggaacagaggcttttcttattaactaagatttctaaatgtatt   20982
                                                                   *************************************    *******************

chimp.LOC112206744.LOC107973052.GGT2-.646279-708040.rev.compl      atgtattcacatttagaaactctaaatgtcataaaaggttagcaaggaagtttcccttcc   22350
FAM230B-LOC105372935-GGT2.NCBI.GGT2.rev.compl                      acgtattcacatttagaaactctaaatatcataaaaggttagcaaggaagtttcccttcc   21042
                                                                   * ************************* ********************************

chimp.LOC112206744.LOC107973052.GGT2-.646279-708040.rev.compl      actctgaacttccaaacaccaagtcaacatttttgtttgcatatcatccctgcaatctat   22410
FAM230B-LOC105372935-GGT2.NCBI.GGT2.rev.compl                      actctgaacttccaaacaccaagtcaacatttttgtttgcatatcatccctgcaatctat   21102
                                                                   ************************************************************

chimp.LOC112206744.LOC107973052.GGT2-.646279-708040.rev.compl      gtgcaaatagaagcatgcacctggaatgcaggctgatgtgtgatcgtgtttacacaaagt   22470
FAM230B-LOC105372935-GGT2.NCBI.GGT2.rev.compl                      gtgcaaatagaagcatgcacctggaatgcaggctgatgtgtgatcatgtttacacaaagt   21162
                                                                   ********************************************* **************
```

```
chimp.LOC112206744.LOC107973052.GGT2-.646279-708040.rev.compl      cctctgcacctctgcatatatcactgggcaatgcaccttagttatcattccacatttcaa   22530
FAM230B-LOC105372935-GGT2.NCBI.GGT2.rev.compl                      cctctgcacctctgcatatatcactgggcaatgcaccttagttatcattccacatttcaa   21222
                                                                   ************************************************************

chimp.LOC112206744.LOC107973052.GGT2-.646279-708040.rev.compl      atgtaaatccattgtattgtttcagagctatgaagtactgcaccccatgactattcccaa   22590
FAM230B-LOC105372935-GGT2.NCBI.GGT2.rev.compl                      atgtaaatccattgtattgtttcagagctataaagtactgcaccccatgactattcccaa   21282
                                                                   ******************************* ****************************

chimp.LOC112206744.LOC107973052.GGT2-.646279-708040.rev.compl      aattacttaagcaccccgctatgggtatccatttgttctgtttccagtcttgctcttata   22650
FAM230B-LOC105372935-GGT2.NCBI.GGT2.rev.compl                      aattacttaagcaccccgctatgggtatccatttgttctgtttccagtcttgctcttata   21342
                                                                   ************************************************************

chimp.LOC112206744.LOC107973052.GGT2-.646279-708040.rev.compl      accaatgctgtagtgaacagcactgtgttgagaagtggtgaacgtgggcatctttgtctt   22710
FAM230B-LOC105372935-GGT2.NCBI.GGT2.rev.compl                      accaatgctgtagtgaacagcactgtgttgagaagcggtgaacgtgggcatctttgtctt   21402
                                                                   *********************************** ************************

chimp.LOC112206744.LOC107973052.GGT2-.646279-708040.rev.compl      gttcccgtcctcagggggaatgctttcaactttcccccattcaggaaactgttggctgtg   22770
FAM230B-LOC105372935-GGT2.NCBI.GGT2.rev.compl                      gttcccttcctcaggcggaatgctttcaactctcccccattcaggaaaatgttggctgtg   21462
                                                                   ****** ******** *************** **************** ***********

chimp.LOC112206744.LOC107973052.GGT2-.646279-708040.rev.compl      ggtttgtcatagatagcttttattaccttaaggtatgtccattctatgctgattttgatg   22830
FAM230B-LOC105372935-GGT2.NCBI.GGT2.rev.compl                      ggtttgtcatagatagcttttattaccttaaggtatgtccgttctacgctgattttgatg   21522
                                                                   **************************************** ***** *************

chimp.LOC112206744.LOC107973052.GGT2-.646279-708040.rev.compl      aacggttttaatcataaagaaatgctggattttgtcaaaggctttttctgcatctattca   22890
FAM230B-LOC105372935-GGT2.NCBI.GGT2.rev.compl                      aacggttttaatcataaagaaatgctggattttgtcaaaggctttttctgcatctattca   21582
                                                                   ************************************************************

chimp.LOC112206744.LOC107973052.GGT2-.646279-708040.rev.compl      gattatcatgtgatttttgtttttagttttatttatgtgatgtatcacatttattgactt   22950
FAM230B-LOC105372935-GGT2.NCBI.GGT2.rev.compl                      gattatcatgtgatttttgtttttagttttattgatgtgatgtatcacatttattgactt   21642
                                                                   ********************************* **************************

chimp.LOC112206744.LOC107973052.GGT2-.646279-708040.rev.compl      gcgtatgttaaaccatccctgcatccctagtatgaaacccacttgaatcatggtggatta   23010
FAM230B-LOC105372935-GGT2.NCBI.GGT2.rev.compl                      gcgtatgttaaaccatccctgcatccctagtatgaaacccacttgaatcatggtggatta   21702
                                                                   ************************************************************

chimp.LOC112206744.LOC107973052.GGT2-.646279-708040.rev.compl      tctttttgatatgctgttggattcagttagcttggttgtagcatttctttttattccatc   23070
FAM230B-LOC105372935-GGT2.NCBI.GGT2.rev.compl                      tctttttgatatgctgttggattcagttagcttggttgtagcatttcttattattccatc   21762
                                                                   ************************************************* **********

chimp.LOC112206744.LOC107973052.GGT2-.646279-708040.rev.compl      tgtggaatgtattggtttaaataatgaaaacatgttctatcctcactgcttagcactttg   23130
FAM230B-LOC105372935-GGT2.NCBI.GGT2.rev.compl                      tgtggaatgtattggttgaaataatgaaaacatgttctatcctcactgcttagcactttg   21822
                                                                   ***************** ******************************************

chimp.LOC112206744.LOC107973052.GGT2-.646279-708040.rev.compl      tgtttctttaatagccttcccaacagggcaacataaaagcaggagccctgctagtcaccc   23190
FAM230B-LOC105372935-GGT2.NCBI.GGT2.rev.compl                      tgtttctttaatagccttcccaacagggcaacataaaagcaggagccctgctagtctccc   21882
                                                                   ******************************************************** ***

chimp.LOC112206744.LOC107973052.GGT2-.646279-708040.rev.compl      cttaacccggaatccccccttctccacagctcgcttattggacaggatagactgggagcc   23250
FAM230B-LOC105372935-GGT2.NCBI.GGT2.rev.compl                      cttaacccggaatccccccttctccacagctcgctcattggacaggatagactgggcgcc   21942
                                                                   *********************************** ******************** ***

chimp.LOC112206744.LOC107973052.GGT2-.646279-708040.rev.compl      caggcctcaaggtaaggacgtgctctgtcacctagaggtgcagtgcttgggaaggccaac   23310
FAM230B-LOC105372935-GGT2.NCBI.GGT2.rev.compl                      caggcttcaaggtaaggacgtgctctgtcacctagaggtgcagtgcttgggaaggccaac   22002
                                                                   ***** ******************************************************

chimp.LOC112206744.LOC107973052.GGT2-.646279-708040.rev.compl      cttggagggttgcctgccagctttacagtgacagaggtgttgggagggactgaccaccag   23370
FAM230B-LOC105372935-GGT2.NCBI.GGT2.rev.compl                      cttggagggttgcctgccagctttacagtgacagaggtgttgagagggactgaccaccag   22062
                                                                   ****************************************** *****************

chimp.LOC112206744.LOC107973052.GGT2-.646279-708040.rev.compl      tgcataaggctgtgctttgttggtgacataaaggattgtttcacaggttgttggggaggg   23430
FAM230B-LOC105372935-GGT2.NCBI.GGT2.rev.compl                      tgcataaggctgtgctttgttggtgacataaaggattgtttcacagattgttggggaggg   22122
                                                                   ********************************************** *************

chimp.LOC112206744.LOC107973052.GGT2-.646279-708040.rev.compl      acaatcccaaggcctcccctggccctggtgctggctctgcacaaaggcaataagagaggg   23490
FAM230B-LOC105372935-GGT2.NCBI.GGT2.rev.compl                      acaatcccaaggcctcccctggccctggtgctggctctgcacaaaggcaataagagaggg   22182
                                                                   ************************************************************

chimp.LOC112206744.LOC107973052.GGT2-.646279-708040.rev.compl      atgctggtaagggctgacctgttgctgtgctggggaggaaggtgctgggctgaaattcag   23550
FAM230B-LOC105372935-GGT2.NCBI.GGT2.rev.compl                      atgctggtaagggctgacctgttgctgtgctggggaggaaggtgctgggctgaaattcag   22242
                                                                   ************************************************************

chimp.LOC112206744.LOC107973052.GGT2-.646279-708040.rev.compl      gaggctgaggatgcagcagtcccataggaggtacgtgaccttcaggatacattttcttca   23610
FAM230B-LOC105372935-GGT2.NCBI.GGT2.rev.compl                      gaggctgaggatgcagcagtcccataggaggtacatgaccttcaggatacattttcttca   22302
                                                                   ********************************** *************************

chimp.LOC112206744.LOC107973052.GGT2-.646279-708040.rev.compl      ttgatgatcaatggaaatgagaaatcactgactatttttctatcattggaatctactct   23670
FAM230B-LOC105372935-GGT2.NCBI.GGT2.rev.compl                      ttgatgatcaatggaaatgagaaatcactgactatttttctatcattggaatctactct   22362
                                                                   ************************************************************

chimp.LOC112206744.LOC107973052.GGT2-.646279-708040.rev.compl      ctactgctcgtgctgttcctgtcttttggggaagatggaggatcaatcagtgtgcgctgc   23730
FAM230B-LOC105372935-GGT2.NCBI.GGT2.rev.compl                      ccactgctcatgctgttcctgtcttttggggaagatggaggatcaatcagtgtgcactgc   22422
                                                                   * ******* ********************************************* ****

chimp.LOC112206744.LOC107973052.GGT2-.646279-708040.rev.compl      actgagtggaaggaaggagaactgcgacaaaaattaaggaaggatgagagacgggagggc   23790
FAM230B-LOC105372935-GGT2.NCBI.GGT2.rev.compl                      actgagtggaaggaaggagaactgtgacaaaaattaaggaaggatgagagacgggagggc   22482
                                                                   ************************ ***********************************

chimp.LOC112206744.LOC107973052.GGT2-.646279-708040.rev.compl      ccttcatccagctgcttgcagagtcctcctgaggaggaaagccccgtggctccctggcga   23850
FAM230B-LOC105372935-GGT2.NCBI.GGT2.rev.compl                      ccttcatccagctgcttgcagagtcctcctgaggaggaaagccccgtggctccctggaga   22542
                                                                   ********************************************************* **

chimp.LOC112206744.LOC107973052.GGT2-.646279-708040.rev.compl      aggagcagtgagggctgcgtgactcccacagtgaagtgtgtggtatgtctgaggacaccc   23910
FAM230B-LOC105372935-GGT2.NCBI.GGT2.rev.compl                      aggagcattgagggctgcgtgactcccacagtgaagtgtgtggtatgtctgaggacaccc   22602
                                                                   ******* ****************************************************

chimp.LOC112206744.LOC107973052.GGT2-.646279-708040.rev.compl      aggctggtggtccatgaggagccagtggcagagtgagaagcagaaaggccaggagggtgg   23970
FAM230B-LOC105372935-GGT2.NCBI.GGT2.rev.compl                      aggctggtggtccatgaggagccagtggcagagtgagaagaagaaaggccaggagggtgg   22662
                                                                   **************************************** *******************

chimp.LOC112206744.LOC107973052.GGT2-.646279-708040.rev.compl      ctggaggccaggctctgagtcattctccatgtgatggaaacagccggagcccagtgggct   24030
FAM230B-LOC105372935-GGT2.NCBI.GGT2.rev.compl                      ctggaggccaggctctgagtcattctccatgtgatggaaacagccggagcccagtgggct   22722
                                                                   ************************************************************

chimp.LOC112206744.LOC107973052.GGT2-.646279-708040.rev.compl      tggaggtacaggatgcggtggctgatgacagaacaatgtggagagaggcgtcatttgtca   24090
FAM230B-LOC105372935-GGT2.NCBI.GGT2.rev.compl                      tggaggtacaggatgcggtggctgatgacagaacaatgtggagagaggcgtcatttgtca   22782
                                                                   ************************************************************

chimp.LOC112206744.LOC107973052.GGT2-.646279-708040.rev.compl      aatccttactttgttctgggcattgtgctaaaaattctgatggctcatcccatttagggg   24150
FAM230B-LOC105372935-GGT2.NCBI.GGT2.rev.compl                      aatccttactttgttctgggcattgtgctaaaaattctgatggctcatcccatttagggg   22842
                                                                   ************************************************************

chimp.LOC112206744.LOC107973052.GGT2-.646279-708040.rev.compl      ctgaaagttgcagaggtttaggaagctcacccatgatactggagcccccatctcctgccc   24210
FAM230B-LOC105372935-GGT2.NCBI.GGT2.rev.compl                      ctgaaagttgcagaggtttaggaagctcacccacgatactggagcccccatctcctgccc   22902
                                                                   ********************************* **************************

chimp.LOC112206744.LOC107973052.GGT2-.646279-708040.rev.compl      tagtgccgtccaccttctcacccagccaccacctgtttcaggggaacacacagaagtggt   24270
FAM230B-LOC105372935-GGT2.NCBI.GGT2.rev.compl                      tagtgctgtccaccttctcacccagccaccacctgtttcgggggaacacacagaagtggt   22962
                                                                   ****** ******************************** ********************

chimp.LOC112206744.LOC107973052.GGT2-.646279-708040.rev.compl      aacctcttatggagaggcaagtaaattctgctgtttttgttattcacagaaaaacactgg   24330
FAM230B-LOC105372935-GGT2.NCBI.GGT2.rev.compl                      aacctcttatggagaggcaagtaaattctgctgtttttgttattcacagaaaaacactgg   23022
                                                                   ************************************************************

chimp.LOC112206744.LOC107973052.GGT2-.646279-708040.rev.compl      ctcatgtgggttgggaaggtgaaataccagaagtatttcatctagttatttctacccatg   24390
FAM230B-LOC105372935-GGT2.NCBI.GGT2.rev.compl                      ctcgtgtgggttgggaaggtgaaataccagaagtatttcatctggttatttctacccatg   23082
                                                                   *** *************************************** ****************

chimp.LOC112206744.LOC107973052.GGT2-.646279-708040.rev.compl      caactcctatagtattgaaatgcataggttagcatttttggccaatttactcagcattct   24450
FAM230B-LOC105372935-GGT2.NCBI.GGT2.rev.compl                      cgactcctatagtattgaaatgcataggttagcatttttggccaatttactcagcattct   23142
                                                                   * **********************************************************

chimp.LOC112206744.LOC107973052.GGT2-.646279-708040.rev.compl      gggttaaaggct-------------tttatttatttatttatttattttcgagatggagt   24497
FAM230B-LOC105372935-GGT2.NCBI.GGT2.rev.compl                      gggttaaaggcttttatttattttatttatttatttatttatttttgagatggagt   23202
                                                                   ************             ************************ **********

chimp.LOC112206744.LOC107973052.GGT2-.646279-708040.rev.compl      ttcgctcttgttgcccaagctggagtgcaatggtgcgatcctggttcactgcaacctccg   24557
FAM230B-LOC105372935-GGT2.NCBI.GGT2.rev.compl                      ttcactcttgttgcccaagctggagtgcaatggtgcgatcctggttcattgcaacctccg   23262
                                                                   *** ******************************************** ***********

chimp.LOC112206744.LOC107973052.GGT2-.646279-708040.rev.compl      cctcccaggttcaaactattctcctgtctcagcctcccaagtagctgagattataggcac   24617
FAM230B-LOC105372935-GGT2.NCBI.GGT2.rev.compl                      cctcccaggttcaaactattctcctgtctcagcctcccaagtagctgagattataggcac   23322
                                                                   ************************************************************
```

```
chimp.LOC112206744.LOC107973052.GGT2-.646279-708040.rev.compl      gtgccaccacactgggctaatttttttgtatttttagtagaaatgagatttcaccatgtt   24677
FAM230B-LOC105372935-GGT2.NCBI.GGT2.rev.compl                      atgccaccacactgggctaatttttttgtatttttagtagaaatgagatttcaccatgtt   23382
                                                                    ***********************************************************

chimp.LOC112206744.LOC107973052.GGT2-.646279-708040.rev.compl      ggtcaggctggtttcaaaatcctgacctcaggtgatctgccctcctcggcctcttaaagt   24737
FAM230B-LOC105372935-GGT2.NCBI.GGT2.rev.compl                      ggtcaggctggtttcgaactcctgacctcaggtgatctgccccctcggcctcttaaagt   23442
                                                                   *************** ** *********************** *****************

chimp.LOC112206744.LOC107973052.GGT2-.646279-708040.rev.compl      gctgggattacaggcgtgagccaccatgcccaccctaaagtcttttaaaattcacttgta   24797
FAM230B-LOC105372935-GGT2.NCBI.GGT2.rev.compl                      gctgggattacaggcgtgagccaccacgcctgccctaaagtcttttaaaattcacttgta   23502
                                                                   ************************** ***  ****************************

chimp.LOC112206744.LOC107973052.GGT2-.646279-708040.rev.compl      taagttgacttagttttctttttttttttctttttctttttctttttttttttttttattgat   24857
FAM230B-LOC105372935-GGT2.NCBI.GGT2.rev.compl                      taagtt------------------------------------------------------   23508
                                                                   ******

chimp.LOC112206744.LOC107973052.GGT2-.646279-708040.rev.compl      cattcttggacttagttttctttaaccttgtagaaaaat-caaaaatggcaatctgtttt   24916
FAM230B-LOC105372935-GGT2.NCBI.GGT2.rev.compl                      --------gacttagttttctttaaccttgtagaaaaatacaaaaatggcaatctctttt   23560
                                                                           ******************************* *************** ****

chimp.LOC112206744.LOC107973052.GGT2-.646279-708040.rev.compl      atcacacaaataatgtctttttaatggagtgatttttttctaattgaggtattatgtact   24976
FAM230B-LOC105372935-GGT2.NCBI.GGT2.rev.compl                      atcacacaaataatgtctttttaatggagtgatttttttctaattgaggtattatgtact   23620
                                                                   ************************************************************

chimp.LOC112206744.LOC107973052.GGT2-.646279-708040.rev.compl      tttcatttactaattattgtttacatttgaagtgttttatgaattaatatttaattgcat   25036
FAM230B-LOC105372935-GGT2.NCBI.GGT2.rev.compl                      tttcatttactaattattgtttacatttgaagtgttttatgaattaatatttaattgcat   23680
                                                                   ************************************************************

chimp.LOC112206744.LOC107973052.GGT2-.646279-708040.rev.compl      agatgaagattactagttataggcatttagtaaccaatactcattaagcatagcgtgga   25096
FAM230B-LOC105372935-GGT2.NCBI.GGT2.rev.compl                      agatgaagattactagttataggcatttactaaccaatactcattaagcatagcgtgga   23740
                                                                   ***************************** ******************************

chimp.LOC112206744.LOC107973052.GGT2-.646279-708040.rev.compl      ttcatatgacatcaaggagctattttatttggtaaaacgaaaaagcacaagaatgaacga   25156
FAM230B-LOC105372935-GGT2.NCBI.GGT2.rev.compl                      ttcctatgacatcaaggagctattttatttggtaaaacgaaaaagcacaagaatgaacga   23800
                                                                   *** ********************************************************

chimp.LOC112206744.LOC107973052.GGT2-.646279-708040.rev.compl      acgcaagaactgaaacggtggagacacctagaatgacttgtctaagatctaaatcatttt   25216
FAM230B-LOC105372935-GGT2.NCBI.GGT2.rev.compl                      acgcaagaactgaaacagtggagacacctagaatgacttgtctaagatctaaatcatttt   23860
                                                                   **************** *******************************************

chimp.LOC112206744.LOC107973052.GGT2-.646279-708040.rev.compl      gttgtcttcccagcgtacttattatcctgatcattgtcatcagcattgtttgggtccttt   25276
FAM230B-LOC105372935-GGT2.NCBI.GGT2.rev.compl                      gttgtcttcccagcgtacttattatcctgatcattgtcatcagcattgtttgggtccttt   23920
                                                                   ************************************************************

chimp.LOC112206744.LOC107973052.GGT2-.646279-708040.rev.compl      tagcacagatttctcaaaatgggtaactccataacagttggaagcttacgaattcatata   25336
FAM230B-LOC105372935-GGT2.NCBI.GGT2.rev.compl                      tagcacagatttctcaaaatgggtaactccataacagttggaagcttacgaattcatata   23980
                                                                   ************************************************************

chimp.LOC112206744.LOC107973052.GGT2-.646279-708040.rev.compl      atttgtaagaggtcaatttggaagtacctatctattttaaaattccaataacctgggaat   25396
FAM230B-LOC105372935-GGT2.NCBI.GGT2.rev.compl                      atttgtaagaggtcaatttgggagtacctatctattttaaaattccaataacctgggaat   24040
                                                                   ********************* **************************************

chimp.LOC112206744.LOC107973052.GGT2-.646279-708040.rev.compl      ttcatcccatgtctagagtcttccatgtaaaatatttccacaattaggagaaatatgtgc   25456
FAM230B-LOC105372935-GGT2.NCBI.GGT2.rev.compl                      ttcatcccatgtctagagtcttttatgtaaaatatttccacaattaggagaaatatgtgc   24100
                                                                   **********************  ************************************

chimp.LOC112206744.LOC107973052.GGT2-.646279-708040.rev.compl      atggggattttctatgtagcggtgttttgatagaatagaaaaatgggataaaccaaattt   25516
FAM230B-LOC105372935-GGT2.NCBI.GGT2.rev.compl                      atggggattttctatgtagcggtgttttgatagaatagaaaattgggataaaccaaattt   24160
                                                                   ***************************************** ******************

chimp.LOC112206744.LOC107973052.GGT2-.646279-708040.rev.compl      ccatcacgaaggaaatagtaatatgctgaataataatacagcgaatattatgcaggcttt   25576
FAM230B-LOC105372935-GGT2.NCBI.GGT2.rev.compl                      ccatcacgaaggaaatagtaatatgctgaataataatacagcgaatattatgcaggcttt   24220
                                                                   ************************************************************

chimp.LOC112206744.LOC107973052.GGT2-.646279-708040.rev.compl      aaacatcaaaaaagagttcaacttctgacttccgatgatggtgttgaagcaggtcactgc   25636
FAM230B-LOC105372935-GGT2.NCBI.GGT2.rev.compl                      aaacatcaaaaaagagttcaacttctgacttccgatgatggtgttgaagcaggtcactgc   24280
                                                                   ************************************************************

chimp.LOC112206744.LOC107973052.GGT2-.646279-708040.rev.compl      tggtttacatttgattttcatgtgggaactctggaagtctgccttagtgattttacatgt   25696
FAM230B-LOC105372935-GGT2.NCBI.GGT2.rev.compl                      tggtttacatttgattttcatgtgggaactctggaagtccgccttagtgattttacatgt   24340
                                                                   *************************************** ********************

chimp.LOC112206744.LOC107973052.GGT2-.646279-708040.rev.compl      ggctaaattgagctaatgacaagctgttcgaagtatggcaaaatggaactttaaaacagt   25756
FAM230B-LOC105372935-GGT2.NCBI.GGT2.rev.compl                      ggctaaattgagctaatgacaagctgtttgcagtatggcaaaatggaactttaaaacagt   24400
                                                                   **************************** * *****************************

chimp.LOC112206744.LOC107973052.GGT2-.646279-708040.rev.compl      atcttgtcagcaaccaagtggacctgtttcacgtaaagcccatgcattcatctgcctgcc   25816
FAM230B-LOC105372935-GGT2.NCBI.GGT2.rev.compl                      atcttgtcaacaaccaagaggacctgtttcacataaagcccacgcattcatctgc-----   24455
                                                                   ********* ******** ************* ********* ************

chimp.LOC112206744.LOC107973052.GGT2-.646279-708040.rev.compl      catcattctgtctgtccatcattctgtctgtccacacgggcatcatttgttagtggaact   25876
FAM230B-LOC105372935-GGT2.NCBI.GGT2.rev.compl                      -----------ctgtccatcattctgtctgtccacacgggcatcattcgttagtggaact   24504
                                                                              ************************************ ************

chimp.LOC112206744.LOC107973052.GGT2-.646279-708040.rev.compl      gagtgcccactgtcgagctgacaagcccataacctccctgttcctagtcacacattaatt   25936
FAM230B-LOC105372935-GGT2.NCBI.GGT2.rev.compl                      gagtgcccgctgtcgagctgacaagcccataacctccctgttcctagtcacacattaatt   24564
                                                                   ******** ***************************************************

chimp.LOC112206744.LOC107973052.GGT2-.646279-708040.rev.compl      cttcaacaagttccttttgatagattgtgattaagcttagctgctatttccaattgcttc   25996
FAM230B-LOC105372935-GGT2.NCBI.GGT2.rev.compl                      cttcaacaagtcccttttgatagattgtgattaagcttagctactatttccaattgcttc   24624
                                                                   *********** ****************************** *****************

chimp.LOC112206744.LOC107973052.GGT2-.646279-708040.rev.compl      cccaaacgtacttctcactgttctcccatcacacccttcagcccatccatgcggggttcc   26056
FAM230B-LOC105372935-GGT2.NCBI.GGT2.rev.compl                      cccaaacgtacttctcactgttctcccatcacacccttcagcccatccatgcggggttcc   24684
                                                                   ************************************************************

chimp.LOC112206744.LOC107973052.GGT2-.646279-708040.rev.compl      tttgcttttcccaccttacaccaaactccctatttttactcccactttttacctcctctcc   26116
FAM230B-LOC105372935-GGT2.NCBI.GGT2.rev.compl                      tttgcttttcccaccttacaccaaactccctatttttactcccactttttacctcctctcc   24744
                                                                   ************************************************************

chimp.LOC112206744.LOC107973052.GGT2-.646279-708040.rev.compl      aagacaaaacaaacaaaactagcattttaaaacttagttgtaatctttcttccttcatga   26176
FAM230B-LOC105372935-GGT2.NCBI.GGT2.rev.compl                      aagacacaacaaacaaaactagcattttaaaacttagttgtaatctttcttccttcatga   24804
                                                                   ****** *****************************************************

chimp.LOC112206744.LOC107973052.GGT2-.646279-708040.rev.compl      aaatttctccaacagccactcccacggtcctgtgtgttccggatattttaaaataatggc   26236
FAM230B-LOC105372935-GGT2.NCBI.GGT2.rev.compl                      aaatttctccaacagccactcccacggtcctgtgtgttccggatattttaaaataatggc   24864
                                                                   ************************************************************

chimp.LOC112206744.LOC107973052.GGT2-.646279-708040.rev.compl      tataaggttgagcacttcaggatatgctgttttgctgtgtgcagatggaggcagtggctg   26296
FAM230B-LOC105372935-GGT2.NCBI.GGT2.rev.compl                      tataaggttgagcacttcaggatacgctgttttgctgtgtgcagatggaggcagtggctg   24924
                                                                   ************************ ***********************************

chimp.LOC112206744.LOC107973052.GGT2-.646279-708040.rev.compl      gagtgaatgaacggcaacacttgctggcaaccggcagaagctgagagacagggaacaggc   26356
FAM230B-LOC105372935-GGT2.NCBI.GGT2.rev.compl                      gagtgaatgaacggcaacacttgctggcaaccggcagaagctgagagacagggaacaggc   24984
                                                                   ************************************************************

chimp.LOC112206744.LOC107973052.GGT2-.646279-708040.rev.compl      tctcctccagagcctccaggagccaggcctttggacaccttgaatgtgggcttctgggag   26416
FAM230B-LOC105372935-GGT2.NCBI.GGT2.rev.compl                      tctcctccagagcctccaggagccaggcctttggacaacttgaatgtgggcttctgggag   25044
                                                                   ************************************* **********************

chimp.LOC112206744.LOC107973052.GGT2-.646279-708040.rev.compl      accatgcgtttctgttataagcagcccagtcgctggcagtttttacggctgcccccgaac   26476
FAM230B-LOC105372935-GGT2.NCBI.GGT2.rev.compl                      accatgcgtttctgttataagcagcccagtctctggcagtttttacggctgccccggaac   25104
                                                                   ******************************* *********************** ****

chimp.LOC112206744.LOC107973052.GGT2-.646279-708040.rev.compl      actcatctatacctgtctgacaaggtcaagctccaaggaagggattctctacatatctac   26536
FAM230B-LOC105372935-GGT2.NCBI.GGT2.rev.compl                      actcatctatacctgtctgacaaggtcaagctccaaggaagggactctctacatatctac   25164
                                                                   ******************************************** ***************

chimp.LOC112206744.LOC107973052.GGT2-.646279-708040.rev.compl      attgtttgcagattttacaataatcatttattcttgcatggctgatcattgttaaccaat   26596
FAM230B-LOC105372935-GGT2.NCBI.GGT2.rev.compl                      attgtttgcagattttacaataatcatttattcttgcatggctgatcattgttaaccaat   25224
                                                                   ************************************************************

chimp.LOC112206744.LOC107973052.GGT2-.646279-708040.rev.compl      acaaataaaataataaagaaatgacccacattttatgttgggagtttgatctgccattta   26656
FAM230B-LOC105372935-GGT2.NCBI.GGT2.rev.compl                      acaaataaaataataaagaaatgacccacattttatgttgggagtttgatctgccattta   25284
                                                                   ************************************************************

chimp.LOC112206744.LOC107973052.GGT2-.646279-708040.rev.compl      tcaagtatggaatcttgaacaaggggttaaacatctgaatgtctccatcacttcatctct   26716
FAM230B-LOC105372935-GGT2.NCBI.GGT2.rev.compl                      tcaagtatggaatcttgaacaaggggttaaacatctgaatgtctccatcacttcatctct   25344
                                                                   ************************************************************

chimp.LOC112206744.LOC107973052.GGT2-.646279-708040.rev.compl      aaagtgggggtgctcacacccactgggctccccct-ccaggttggtgccggactctccct   26775
FAM230B-LOC105372935-GGT2.NCBI.GGT2.rev.compl                      aaagtgggggtgctcacacccactgggctccccgcccaggttggtgccggactctccct   25404
                                                                   **********************************  ************************

chimp.LOC112206744.LOC107973052.GGT2-.646279-708040.rev.compl      gggcccccctgttctctcaccagccacatccattctccccccagaggcgctagtgactgt   26835
```

```
FAM230B-LOC105372935-GGT2.NCBI.GGT2.rev.compl                   gggccccctgttctctcaccagccacatccattctcccccagaggcgctagtgactgt   25464
                                                                ************************************************************

chimp.LOC112206744.LOC107973052.GGT2-.646279-708040.rev.compl   gcgtggctttccattcccaccacgtttgtctctaaccccagtggcagatcagtgtaagaa   26895
FAM230B-LOC105372935-GGT2.NCBI.GGT2.rev.compl                   gcgtggctttccattcccaccacgtttgtctctaaccccagtggcagatcagtgtaagaa   25524
                                                                ************************************************************

chimp.LOC112206744.LOC107973052.GGT2-.646279-708040.rev.compl   cacagctgagtgctcctcgcctccttgccccttcaagggctcctcaccacccaccagatc   26955
FAM230B-LOC105372935-GGT2.NCBI.GGT2.rev.compl                   cacagctgagtgctcctcgcctccttgccccttcaagggcccctcaccacccaccagatc   25584
                                                                **************************************** *******************

chimp.LOC112206744.LOC107973052.GGT2-.646279-708040.rev.compl   aggtgcaaacttccaagccttactggatccccttccacattctgagctccgcctgccttc   27015
FAM230B-LOC105372935-GGT2.NCBI.GGT2.rev.compl                   aggtgcaaacttccaagccttactggatccccttccacattctgagctccgcctgccttc   25644
                                                                ************************************************************

chimp.LOC112206744.LOC107973052.GGT2-.646279-708040.rev.compl   ccatcgctatccttccccacctgcctccctggtagagaaaagcagagtgtgtgatgctgt   27075
FAM230B-LOC105372935-GGT2.NCBI.GGT2.rev.compl                   ccatcgctatccttccccacctgcctccctggtagagaaaagcggagtgtgtgatgctgt   25704
                                                                ******************************************* ****************

chimp.LOC112206744.LOC107973052.GGT2-.646279-708040.rev.compl   ctgaatgctgagcacggccttttgcagccagtccactgtgtacgctgcccctatcggaga   27135
FAM230B-LOC105372935-GGT2.NCBI.GGT2.rev.compl                   ctgaatgctgagcacggccttttgcagccagtccactgtgtacgctgcccctatcggaga   25764
                                                                ************************************************************

chimp.LOC112206744.LOC107973052.GGT2-.646279-708040.rev.compl   cctccaccttaaccctttccagcctggaggctcctcccagggccccacaacagaagtgac   27195
FAM230B-LOC105372935-GGT2.NCBI.GGT2.rev.compl                   cctccaccttaaccctttccagcctggaggctcctcccagggccccacaacagaagtgac   25824
                                                                ************************************************************

chimp.LOC112206744.LOC107973052.GGT2-.646279-708040.rev.compl   ttcccttgcctttgaatttctatagcacaatccctactgcccccgttaaaactgcaaag   27255
FAM230B-LOC105372935-GGT2.NCBI.GGT2.rev.compl                   ttcccttgcctttgaatttctatagcacaatccctactgcccccgttaaaactgcaaag   25884
                                                                ************************************************************

chimp.LOC112206744.LOC107973052.GGT2-.646279-708040.rev.compl   tccttttgtggaaaataactttattcatgactgtgtttatcacactatcttatggagaag   27315
FAM230B-LOC105372935-GGT2.NCBI.GGT2.rev.compl                   tccttttgtggaaaataactttattcatgactgtgtttatcacactatcttatggagaag   25944
                                                                ************************************************************

chimp.LOC112206744.LOC107973052.GGT2-.646279-708040.rev.compl   agatgatcaataaatatttgcggaataaatgaatagcagttacaaaacacttgattcata   27375
FAM230B-LOC105372935-GGT2.NCBI.GGT2.rev.compl                   agatgatcaataaatatttgctgaataaatgaatagcagttacaaaacacttgattcata   26004
                                                                ********************* **************************************

chimp.LOC112206744.LOC107973052.GGT2-.646279-708040.rev.compl   tggaattaatgttggttctcaaagtgaaaaattacaaacagcactgatattcagccagta   27435
FAM230B-LOC105372935-GGT2.NCBI.GGT2.rev.compl                   tggaattaatgttggttctcaaagtgaaaaattacaaacagcactgatattcagccagta   26064
                                                                ************************************************************

chimp.LOC112206744.LOC107973052.GGT2-.646279-708040.rev.compl   tacaagtctggtcacagcagttgtataatactgaaataccccctgccactgacctttggc   27495
FAM230B-LOC105372935-GGT2.NCBI.GGT2.rev.compl                   tacaagtctggtcacagcagttgtataatactgaaataccccctgccactgacctttggc   26124
                                                                ************************************************************

chimp.LOC112206744.LOC107973052.GGT2-.646279-708040.rev.compl   ccccagatgcctcccactgccactgctctccccactgggaacccctgaagttcccacagg   27555
FAM230B-LOC105372935-GGT2.NCBI.GGT2.rev.compl                   ccccagatgcctcccactgccactgctctccccactgggaacccctgaagttcccacagg   26184
                                                                ************************************************************

chimp.LOC112206744.LOC107973052.GGT2-.646279-708040.rev.compl   ctcataactaaagggctaatgtcctgcacagcagcgagcacccaggaccgagcagccaca   27615
FAM230B-LOC105372935-GGT2.NCBI.GGT2.rev.compl                   ctcataactaaagggctaatgtcttgcacagcagcgagcacccaggaccgagcagccaca   26244
                                                                *********************** ************************************

chimp.LOC112206744.LOC107973052.GGT2-.646279-708040.rev.compl   tggccgggtctgctggtgaaagcatccattctgactgatcaggacctgaggggcctcatg   27675
FAM230B-LOC105372935-GGT2.NCBI.GGT2.rev.compl                   tggccgggtctgctggtgaaagcatccattctgactgatcaggacctgaggggcctcatg   26304
                                                                ************************************************************

chimp.LOC112206744.LOC107973052.GGT2-.646279-708040.rev.compl   gttacatattttgataatatccctaattataaataaggctcagttatatagtttgaaaac   27735
FAM230B-LOC105372935-GGT2.NCBI.GGT2.rev.compl                   gttacatattttgataatatccctaattataaataaggctcagttatatagtttgaaaac   26364
                                                                ************************************************************

chimp.LOC112206744.LOC107973052.GGT2-.646279-708040.rev.compl   aatgcttctcctcattgcaaaatctcttagaagactccgtagatccaggaacggaaatgg   27795
FAM230B-LOC105372935-GGT2.NCBI.GGT2.rev.compl                   agtgcttctcctcattgcaaaatctcttagaagactccgtagatccaggaacggaaatgg   26424
                                                                * **********************************************************

chimp.LOC112206744.LOC107973052.GGT2-.646279-708040.rev.compl   aaaatgacagcatgtcaatctctgaaggttttgggcatttccattagcactccatcttca   27855
FAM230B-LOC105372935-GGT2.NCBI.GGT2.rev.compl                   aaaatgacagcgtgtcaatctctgaaggttttgggcatttccattagcactccatcttca   26484
                                                                *********** ************************************************

chimp.LOC112206744.LOC107973052.GGT2-.646279-708040.rev.compl   tgtaaaccagaagatatgcagtttcctgcctagagagaagagaagacacatcagcacagc   27915
FAM230B-LOC105372935-GGT2.NCBI.GGT2.rev.compl                   tgtaaaccagaagatatgcagtttcctgcctagagagaagagaagacacatcagcacagc   26544
                                                                ************************************************************

chimp.LOC112206744.LOC107973052.GGT2-.646279-708040.rev.compl   ggcatgaaaccttcatcagaaaacaatgcttcattaatccttgacaggacaagcgtcagc   27975
FAM230B-LOC105372935-GGT2.NCBI.GGT2.rev.compl                   ggcatgaaaccttcatcagaaaacaatgcttcattaatccgtgacaggacaagcgtcagc   26604
                                                                **************************************** *******************

chimp.LOC112206744.LOC107973052.GGT2-.646279-708040.rev.compl   aaacttccaggccgctggattaagccttcatctatccatcaccttggagaggaacaaaat   28035
FAM230B-LOC105372935-GGT2.NCBI.GGT2.rev.compl                   aaacttccaggcggctggattaggccttcatctatccatcaccttggagaggaacaaaat   26664
                                                                *********** ********* **************************************

chimp.LOC112206744.LOC107973052.GGT2-.646279-708040.rev.compl   aggtggcctgggaagataagcactatgtttctattagttaatatctaaagcggaggttaa   28095
FAM230B-LOC105372935-GGT2.NCBI.GGT2.rev.compl                   aggtggcctgggaagataagcactatgtttctattagttaatatctaaagcggaggttaa   26724
                                                                ************************************************************

chimp.LOC112206744.LOC107973052.GGT2-.646279-708040.rev.compl   caagctatggacacacaagccaaacccagccctcttggggtttttaaatctactttcaa   28155
FAM230B-LOC105372935-GGT2.NCBI.GGT2.rev.compl                   caagctatggacacacaagccaaacccagccctcttggggtttttaaatctactttcaa   26784
                                                                ************************************************************

chimp.LOC112206744.LOC107973052.GGT2-.646279-708040.rev.compl   cttttattttagattcagcgggcacatgtgcaggtttgtcacgtggatatgagcatactc   28215
FAM230B-LOC105372935-GGT2.NCBI.GGT2.rev.compl                   cttttattttagattcagcgggcacatgtgcaggtttgtcacatggatatgagcatactc   26844
                                                                ****************************************** *****************

chimp.LOC112206744.LOC107973052.GGT2-.646279-708040.rev.compl   cccaacagttggcctttcacccctcccctccctccccatccagcagttcccagttgttgc   28275
FAM230B-LOC105372935-GGT2.NCBI.GGT2.rev.compl                   cccaacagttggcctttcacccctcccctccctccccatccagcagttcccagttgttgc   26904
                                                                ************************************************************

chimp.LOC112206744.LOC107973052.GGT2-.646279-708040.rev.compl   catctttaagtcaatgactgcccatgtttagctcccatttataagagagaatatgcatta   28335
FAM230B-LOC105372935-GGT2.NCBI.GGT2.rev.compl                   catctttaagtcaatgagtccccatgtttagctcccatttataagagagaacatgcatta   26964
                                                                ***************** * ******************************* ********

chimp.LOC112206744.LOC107973052.GGT2-.646279-708040.rev.compl   tgttttgtttggttttgctgg-----ttttttttttttaatggagtcttgccctgtagc   28390
FAM230B-LOC105372935-GGT2.NCBI.GGT2.rev.compl                   tgttttgtttggttttgctggttttttttttttttttttaatggagtcttgccctgtagc   27024
                                                                *********************     **********************************

chimp.LOC112206744.LOC107973052.GGT2-.646279-708040.rev.compl   ccaggctagagtgcagtggcacaatcttggctcactgcaacctccgcctcccaggttcaa   28450
FAM230B-LOC105372935-GGT2.NCBI.GGT2.rev.compl                   ccaggctagagtgcagtggcacaaccttggctcactgcaacctccgcctcccaggttcaa   27084
                                                                *********************** ************************************

chimp.LOC112206744.LOC107973052.GGT2-.646279-708040.rev.compl   acaattctccctcctcagcctcccgagtagctgggactacaggtgcctgccaccacgccc   28510
FAM230B-LOC105372935-GGT2.NCBI.GGT2.rev.compl                   acgattctccctcctcagcctcccgagtggctgggactacaggcgcccaccaccacgccc   27144
                                                                ** ************************* ************** ***  ***********

chimp.LOC112206744.LOC107973052.GGT2-.646279-708040.rev.compl   agctaacttttgtatttttagtagagacagggtttcactgtgttagccaggatggtctc   28570
FAM230B-LOC105372935-GGT2.NCBI.GGT2.rev.compl                   ggctaacttttgtatttttagtagagacagggtttcaccgtgttagccaggatggtctc   27204
                                                                 ************************************** ********************

chimp.LOC112206744.LOC107973052.GGT2-.646279-708040.rev.compl   aatctcctgacctcatgatctgcccacctcagtctcccaaagtgcaggattacaggcat   28630
FAM230B-LOC105372935-GGT2.NCBI.GGT2.rev.compl                   aatctcctgacctcatgatctgcccaccttagtctcccaaagtgcaggattacaggcgt   27264
                                                                ***************************** **************************** *

chimp.LOC112206744.LOC107973052.GGT2-.646279-708040.rev.compl   gagccaccgtgcccagtcttttgtttatttttgacgagaccgttcttgctctgtcacca   28690
FAM230B-LOC105372935-GGT2.NCBI.GGT2.rev.compl                   gagccaccgtgcccagccttttgtttatttttgatgagacggttcttgctctgtcacca   27324
                                                                **************** ****************** ***** ******************

chimp.LOC112206744.LOC107973052.GGT2-.646279-708040.rev.compl   ggctggagtgcactggcacagtaatagctcaccacagccttgtgctcctgggctcaactg   28750
FAM230B-LOC105372935-GGT2.NCBI.GGT2.rev.compl                   ggctggagtgcactggcacaataatagctcaccacagcctcgtgctcctgggctcaactg   27384
                                                                ******************** ******************* *******************

chimp.LOC112206744.LOC107973052.GGT2-.646279-708040.rev.compl   accctcctgcctcagttttagcttcctgagtagctaggactacgggtgtgtaccaccatg   28810
FAM230B-LOC105372935-GGT2.NCBI.GGT2.rev.compl                   accctcctgcctcagttttagcttcctgagtagctaggactacgggtgtgtaccaccatg   27444
                                                                ************************************************************

chimp.LOC112206744.LOC107973052.GGT2-.646279-708040.rev.compl   cctagctataataattttattttttgtagagatggagtcttgctttgttgcccaggct   28870
FAM230B-LOC105372935-GGT2.NCBI.GGT2.rev.compl                   cctagctataataattttattttttgtagagatggagtcttgctttgttgcccaggct   27504
                                                                ************************************************************

chimp.LOC112206744.LOC107973052.GGT2-.646279-708040.rev.compl   ggtcttgaactcctggcttaaagtgatcctcctgcctcggcctcccaaagtgctgggatt   28930
FAM230B-LOC105372935-GGT2.NCBI.GGT2.rev.compl                   ggtcttgaactcctggcttgaagtgatcctcctgcctcggcctcccaaagtgccgggatt   27564
                                                                ******************* ********************************* ******

chimp.LOC112206744.LOC107973052.GGT2-.646279-708040.rev.compl   aaaggtgtgagatcacacccagtctccaaccctctttttgcaagtaaatgtaactggacc   28990
FAM230B-LOC105372935-GGT2.NCBI.GGT2.rev.compl                   aaaggtgggagatcgcacccagtctccaaccctctttttgcaagtaaatgtaactggacc   27624
```

```
                                                                   ******* ****** *********************************************

chimp.LOC112206744.LOC107973052.GGT2-.646279-708040.rev.compl      ccagccatgctcatctgcccatgtactgtctacggctgcttttgctctatagggcagagt   29050
FAM230B-LOC105372935-GGT2.NCBI.GGT2.rev.compl                      ccagccatgctcatctgcccatgtactgtctacggctgcttttgctctacagggcagagt   27684
                                                                   ************************************************* **********

chimp.LOC112206744.LOC107973052.GGT2-.646279-708040.rev.compl      taagtggttgcaacagacactgcacagaccacaaagtctggaagtactttctctccagcc   29110
FAM230B-LOC105372935-GGT2.NCBI.GGT2.rev.compl                      taagtggttgcaacagacactgcacagaccacaaagtct-gaagtactttctctccagcc   27743
                                                                   *************************************** ********************

chimp.LOC112206744.LOC107973052.GGT2-.646279-708040.rev.compl      ctttacagagaaagtctgccaacctctaatctcaataacagggaaatcaatgacaaccac   29170
FAM230B-LOC105372935-GGT2.NCBI.GGT2.rev.compl                      ctttacagagaaagtctgccaacctctaatctcaataacagggaaatcaatgacaaccac   27803
                                                                   ************************************************************

chimp.LOC112206744.LOC107973052.GGT2-.646279-708040.rev.compl      aaagtgacaaagattgggtgtctaagatggatggtcagaataaacaagagagaaagatga   29230
FAM230B-LOC105372935-GGT2.NCBI.GGT2.rev.compl                      aaagtgacaaagattgggtgtctaagatggatgttcagaataaacaagagagaaagatga   27863
                                                                   ********************************* **************************

chimp.LOC112206744.LOC107973052.GGT2-.646279-708040.rev.compl      aaagtagaaggaggatttcaagcgcaagcttcacctaatccgttatttttcaaatgacca   29290
FAM230B-LOC105372935-GGT2.NCBI.GGT2.rev.compl                      gaagtagaaggaggatttcaaacgcaagcttcacctaatccgttatttttcaaatgacca   27923
                                                                    ******************** **************************************

chimp.LOC112206744.LOC107973052.GGT2-.646279-708040.rev.compl      ggcctatctctgtagctgaaatcacctcaaataggatctctgatatacagtctccaaaa   29350
FAM230B-LOC105372935-GGT2.NCBI.GGT2.rev.compl                      ggcctatctctgtagccgaaatcacctcaaataggatctctgatatacagtctccaaaa   27983
                                                                   **************** *******************************************

chimp.LOC112206744.LOC107973052.GGT2-.646279-708040.rev.compl      gctcagctaagaaacttac--agtctctctgccttaacttcatccacctttttctctcc   29408
FAM230B-LOC105372935-GGT2.NCBI.GGT2.rev.compl                      gctcagctaagaaacttacaaagtctctctgccttaacttcatccacctttttctctcc   28043
                                                                   *******************  ***************************************

chimp.LOC112206744.LOC107973052.GGT2-.646279-708040.rev.compl      agcttctcctcggtagttaatgattataaaaatatttattggctcatgcctgtaatccca   29468
FAM230B-LOC105372935-GGT2.NCBI.GGT2.rev.compl                      agcttctcctcggtagttaatgattataaaaatatttattggctcatgcctgtaatccca   28103
                                                                   ************************************************************

chimp.LOC112206744.LOC107973052.GGT2-.646279-708040.rev.compl      gcactttgagaggccaaggcgggcagatcacgaggtcaggagagcgagaccatcctggct   29528
FAM230B-LOC105372935-GGT2.NCBI.GGT2.rev.compl                      gcactttgagaggccgaggcgggcagatcac--ggtcaggagatcgagaccatcctggct   28161
                                                                   *************** ***************  ********** ****************

chimp.LOC112206744.LOC107973052.GGT2-.646279-708040.rev.compl      aacacggtgaaatcccgtctctactaaaaatacaaaaaattagccaggcgtggtggcggg   29588
FAM230B-LOC105372935-GGT2.NCBI.GGT2.rev.compl                      aacacggtgaaatcccgtctctactaaaaatacaaaaaattagccaggcgtggtggcggg   28221
                                                                   ************************************************************

chimp.LOC112206744.LOC107973052.GGT2-.646279-708040.rev.compl      cacctgtaatcccagctactcgggaggctgaggcaggagaatggcgtgaacccacaaggc   29648
FAM230B-LOC105372935-GGT2.NCBI.GGT2.rev.compl                      cgcctgtaatcccagctactcaggaggctgaggcaggagaatggcgtgaacccacaaggc   28281
                                                                   * ******************* **************************************

chimp.LOC112206744.LOC107973052.GGT2-.646279-708040.rev.compl      ggagcttgcaatgaggtgagatcccactactgcactccagcctgggcgacagagcaagac   29708
FAM230B-LOC105372935-GGT2.NCBI.GGT2.rev.compl                      ggagcttgcaatgaggtgagatcccactactgcactccagcctgggcgacagagcaagac   28341
                                                                   ************************************************************

chimp.LOC112206744.LOC107973052.GGT2-.646279-708040.rev.compl      tccatctca----aaacaaacaaacaaacaaaaaaacagtgtgatggccaggcgcggtgc   29764
FAM230B-LOC105372935-GGT2.NCBI.GGT2.rev.compl                      tccatctcaaaacaaacaaacaaacaaacaaaaaaacagtgtgatggccaggcgcagtgc   28401
                                                                   *********    ****************************************** ****

chimp.LOC112206744.LOC107973052.GGT2-.646279-708040.rev.compl      tcaagcctataatccaagcactttgggaggctgaaatggatcgatggcttgagcccagta   29824
FAM230B-LOC105372935-GGT2.NCBI.GGT2.rev.compl                      tcatgcctataatccaagcactttgggaggctgaaatggatggatggcttgagcccagta   28461
                                                                   *** ************************************* ******************

chimp.LOC112206744.LOC107973052.GGT2-.646279-708040.rev.compl      gtttgagacaagcctggcaacatagcgagacctcatctctacaaacatttttaaaatatg   29884
FAM230B-LOC105372935-GGT2.NCBI.GGT2.rev.compl                      gtttgagacaagcctggcaacatagcgagacctcatctctacaaacatctttaaaatatg   28521
                                                                   ************************************************ ***********

chimp.LOC112206744.LOC107973052.GGT2-.646279-708040.rev.compl      ccaggcatggtgtgcagactgcatgcctgtagtcccagctattcgggaggctgaggtggg   29944
FAM230B-LOC105372935-GGT2.NCBI.GGT2.rev.compl                      ccaggcatggtg------gtgcatgcctgtagtcccagctattcaggaggctgaggtggg   28575
                                                                   ************       ************************* ***************

chimp.LOC112206744.LOC107973052.GGT2-.646279-708040.rev.compl      aggatcacctgtgcccgggagttcaaggctgcagtgagctatgatcacaccacagtgctc   30004
FAM230B-LOC105372935-GGT2.NCBI.GGT2.rev.compl                      aggatcacctgtgcccgggagttcaaggctgcagtgagctatgatcacaccacagtgctc   28635
                                                                   ************************************************************

chimp.LOC112206744.LOC107973052.GGT2-.646279-708040.rev.compl      cagcctgggcaacaaagcaagactccatctctaaaaataaaataaaa-ttaaagaaaaga   30063
FAM230B-LOC105372935-GGT2.NCBI.GGT2.rev.compl                      cagcctgggcaacaaagcaagactccatctctaaaaataaaataaaatttaaaaaaaaga   28695
                                                                   *********************************************** ***** ******

chimp.LOC112206744.LOC107973052.GGT2-.646279-708040.rev.compl      tctttgctgtaaaagaggtacgctcaaatgcaataaaagcatataagaaggccgggtgtg   30123
FAM230B-LOC105372935-GGT2.NCBI.GGT2.rev.compl                      tcttcgctgtaaaagaggtacgctcaaatgcaataaaagcatataagaaggccgggtgtg   28755
                                                                   **** *******************************************************

chimp.LOC112206744.LOC107973052.GGT2-.646279-708040.rev.compl      gtggctcatgcctgtaatctcagcactttgggaggccaagacgggcggatcacgaggtca   30183
FAM230B-LOC105372935-GGT2.NCBI.GGT2.rev.compl                      gtggctcatgcctgtaatcccagcactttgggaggccgagacgggcggatcacgaggtca   28815
                                                                   ******************* ***************** **********************

chimp.LOC112206744.LOC107973052.GGT2-.646279-708040.rev.compl      ggagatcgagactatcctggctaacgcggagaaaccccatctcctctaaaagtacaaaaa   30243
FAM230B-LOC105372935-GGT2.NCBI.GGT2.rev.compl                      ggagattgagactatcctggctaacgcggtgaaaccccatctcctctaaaagtacaaaaa   28875
                                                                   ****** ********************** ******************************

chimp.LOC112206744.LOC107973052.GGT2-.646279-708040.rev.compl      aattagttgggctaggtggcaggcgcctgtagtcccagctactcaggaggctgaggcagg   30303
FAM230B-LOC105372935-GGT2.NCBI.GGT2.rev.compl                      aattagctgggctaggtggcaggcgcctgtagtcccagctactgaggaggctgaggcagg   28935
                                                                   ****** ************************************ ****************

chimp.LOC112206744.LOC107973052.GGT2-.646279-708040.rev.compl      agaatggcataaacccgggaggcagagcttgcagtgagccaagatcgcaccactgcactc   30363
FAM230B-LOC105372935-GGT2.NCBI.GGT2.rev.compl                      agaatggcataaaccagggaggcagagcttgcagtgagcctagatcacaccactgcactc   28995
                                                                   *************** ************************ ***** *************

chimp.LOC112206744.LOC107973052.GGT2-.646279-708040.rev.compl      cagcctgggtgacagagcgagactccgtctcaaaaaaaaaaaaa--aaaagaaaaagaaa   30421
FAM230B-LOC105372935-GGT2.NCBI.GGT2.rev.compl                      cagcctgggtgacagagcgagactccgtctcaaaaaaaaaaaaaagaaaagaaaaagaaa   29055
                                                                   ********************************************  **************

chimp.LOC112206744.LOC107973052.GGT2-.646279-708040.rev.compl      agaaaagttcttgtgacatttgtgtatgaaatcagccttcactacatggataggaccagc   30481
FAM230B-LOC105372935-GGT2.NCBI.GGT2.rev.compl                      agaaaagttcttctgacatttgtgtatgaaatcagccttcactacatggataggaccagc   29115
                                                                   ************ ***********************************************

chimp.LOC112206744.LOC107973052.GGT2-.646279-708040.rev.compl      acgcttccgcggcacgactctgcaatcttactacatttttttttactttgtattttattt   30541
FAM230B-LOC105372935-GGT2.NCBI.GGT2.rev.compl                      acgcttctgcggcacgactctgcaatcttactacatttttttttactttgtattttattt   29175
                                                                   ******* ****************************************************

chimp.LOC112206744.LOC107973052.GGT2-.646279-708040.rev.compl      attccttttgagacagagtctcactctgtcacccaggctgaagtgcagccgagatctcag   30601
FAM230B-LOC105372935-GGT2.NCBI.GGT2.rev.compl                      attccttttgagacagagtctcactctgtcacccaggctgaagtgcagccgagatctcgg   29235
                                                                   ********************************************************** *

chimp.LOC112206744.LOC107973052.GGT2-.646279-708040.rev.compl      ctcactgcaacctccacctcctgggttcaagcaattctcttgtctcagcctcccaagtag   30661
FAM230B-LOC105372935-GGT2.NCBI.GGT2.rev.compl                      ctcactgcaacctccacctcttgggttcaagcaattctcttgtctcagcctcccaagtag   29295
                                                                   ******************** ***************************************

chimp.LOC112206744.LOC107973052.GGT2-.646279-708040.rev.compl      ctgggactacaggcacacgtcaaaacgcccggctaatttttgtatttttagtagagatgg   30721
FAM230B-LOC105372935-GGT2.NCBI.GGT2.rev.compl                      ctgggactacaggcacacgtcaaaacgcccggctaatttttgtatttttagtagagatgg   29355
                                                                   ************************************************************

chimp.LOC112206744.LOC107973052.GGT2-.646279-708040.rev.compl      agtttagccatattggtcaggctggtctcgaactcctgacctcaggtgatcgacctgtct   30781
FAM230B-LOC105372935-GGT2.NCBI.GGT2.rev.compl                      agttttgccatattggtcaggctggtctcgaactcctgacctcaggtgatcgacctgtct   29415
                                                                   ***** ******************************************************

chimp.LOC112206744.LOC107973052.GGT2-.646279-708040.rev.compl      tagcctcccaaagtgctaggattacaggtgtacatttatttatttatttgagatggaatc   30841
FAM230B-LOC105372935-GGT2.NCBI.GGT2.rev.compl                      tagcctcccaaagtgctaggattacaggtgtacatttatttatttatttgagatggaatc   29475
                                                                   ************************************************************

chimp.LOC112206744.LOC107973052.GGT2-.646279-708040.rev.compl      ctgctctgtatttattaatttatttatttgagatggagtcttgctccattgcccaggcta   30901
FAM230B-LOC105372935-GGT2.NCBI.GGT2.rev.compl                      ttgctctgtatttattaatttatttatttgagatggagtcttgctccatcgcccaggcta   29535
                                                                    ************************************************ **********

chimp.LOC112206744.LOC107973052.GGT2-.646279-708040.rev.compl      gagtgcagtggtgcaatctcggctcattgcaacctctgacttccaggttcaagcgattct   30961
FAM230B-LOC105372935-GGT2.NCBI.GGT2.rev.compl                      gagtgcagcggtgcaatctcggctcattgcaacctctgccttccaggttcaagcgattct   29595
                                                                   ******** ***************************** *********************

chimp.LOC112206744.LOC107973052.GGT2-.646279-708040.rev.compl      cctgcctcagtgtcccaagtagctgggattacaggtgcctgccaccacagctggttaatt   31021
FAM230B-LOC105372935-GGT2.NCBI.GGT2.rev.compl                      cctgcctcagtgtcccaagtagctgggattacaggtgcctgccaccacagctggctaatt   29655
                                                                   ****************************************************** *****

chimp.LOC112206744.LOC107973052.GGT2-.646279-708040.rev.compl      tttgtatttttagtagagacagtgtttcaccatcttggccaggctggtctcgggctcctg   31081
FAM230B-LOC105372935-GGT2.NCBI.GGT2.rev.compl                      tttgtatttttagtagagacagtgtttcaccatcttggccaggctggtctcgggctcctg   29715
                                                                   ************************************************************

chimp.LOC112206744.LOC107973052.GGT2-.646279-708040.rev.compl      acctcatgaaccacctgcctcagcctcccaaagtgttgggattacaggcctaaggcacca   31141
FAM230B-LOC105372935-GGT2.NCBI.GGT2.rev.compl                      acctcatgaaccacctgcctcagcctcccaaagtgttgggattacaggcctaaggcacca   29775
                                                                   ************************************************************
```

```
chimp.LOC112206744.LOC107973052.GGT2-.646279-708040.rev.compl      tgttcggccatatttatttatttaattatttagagacaaagtcttgctctgtcacccagg   31201
FAM230B-LOC105372935-GGT2.NCBI.GGT2.rev.compl                      tgctcggccatatttatttatttaattatttagagacaaagtcttgctctgtcacccagg   29835
                                                                   ** *********************************************************

chimp.LOC112206744.LOC107973052.GGT2-.646279-708040.rev.compl      ctggagtgcagtggcgccatctcagctcactgcagcctccgcctctgaggtttaagcgat   31261
FAM230B-LOC105372935-GGT2.NCBI.GGT2.rev.compl                      ctggagtgcagtggcgccatctcagctcactgcagcctccgcctccgaggtttaagcaat   29895
                                                                   ********************************************* *********** **

chimp.LOC112206744.LOC107973052.GGT2-.646279-708040.rev.compl      tctcatgcctcagcctcctgaataactaggactacagatactcgccaccacgcagggatt   31321
FAM230B-LOC105372935-GGT2.NCBI.GGT2.rev.compl                      tctcatgcctcagcctcctgagtaactgggactacagatacttgccaccacgcaggga--   29953
                                                                   ********************* ***** ************** ***************

chimp.LOC112206744.LOC107973052.GGT2-.646279-708040.rev.compl      tttttttttctatttttttgtagagacacagtttcaccatgttggccaggctggtctcga   31381
FAM230B-LOC105372935-GGT2.NCBI.GGT2.rev.compl                      tttttttttctatttttttgtagagacacagtttcaccatgttggccaggctggtctcga   30013
                                                                   ************************************************************

chimp.LOC112206744.LOC107973052.GGT2-.646279-708040.rev.compl      actcctgaccttaggtgatctgacagcctcgtcctctctaagtactgggattacaggcat   31441
FAM230B-LOC105372935-GGT2.NCBI.GGT2.rev.compl                      actcctgaccttaggtgatctgacagcctcgtcctctcaaagcactgggattacaggcat   30073
                                                                   ************************************** *** *****************

chimp.LOC112206744.LOC107973052.GGT2-.646279-708040.rev.compl      gagccccttgcccggcctctcactacatttaagtgacgccatggctcatgcctgtaatcc   31501
FAM230B-LOC105372935-GGT2.NCBI.GGT2.rev.compl                      gagccccttgcccggcctctcactacatttaagtgacgccatggctcatgcctgtaatcc   30133
                                                                   ************************************************************

chimp.LOC112206744.LOC107973052.GGT2-.646279-708040.rev.compl      tagcactttgggaggccaaggcaggtggatcacctgaggtcaggagttcgacacgagcct   31561
FAM230B-LOC105372935-GGT2.NCBI.GGT2.rev.compl                      tagcactttgggaggccaaggcaggtggatcacctgaggtcaggagttcgacacgagcct   30193
                                                                   ************************************************************

chimp.LOC112206744.LOC107973052.GGT2-.646279-708040.rev.compl      ggccaacatggggaaaccccgtctctagtaaaaatacaaaaattagtcacgcgtggtggt   31621
FAM230B-LOC105372935-GGT2.NCBI.GGT2.rev.compl                      ggccaacatggggaaaccccgtctctagtaaaaatacaaaaattagtcaggtgtggtggt   30253
                                                                   ************************************************* * ********

chimp.LOC112206744.LOC107973052.GGT2-.646279-708040.rev.compl      acaagcctgtaggcccagctacttggaagactgaggcaggagaatcactttaagcgggag   31681
FAM230B-LOC105372935-GGT2.NCBI.GGT2.rev.compl                      acaagcctgtaggcccagctacttggaagactgaggcaggagaatcactttaagcgggag   30313
                                                                   ************************************************************

chimp.LOC112206744.LOC107973052.GGT2-.646279-708040.rev.compl      gcagaggttgcagtgagccaatatcatgccactgcactccagcttgggtgacggagtgag   31741
FAM230B-LOC105372935-GGT2.NCBI.GGT2.rev.compl                      gcagaggttgcagtgagccaatatcatgccactgcactccagcttgggtgacagagtgag   30373
                                                                   **************************************************** *******

chimp.LOC112206744.LOC107973052.GGT2-.646279-708040.rev.compl      acactgtctcaaa-------aaaaaaaaagagaaaaaaatatgatgccggggcatctcgg   31794
FAM230B-LOC105372935-GGT2.NCBI.GGT2.rev.compl                      atactgtctcaaaaaaaaagaaaaaaagagagaaaaacatatgatgccggggcatctcgg   30433
                                                                   * ***********       ******* ********* **********************

chimp.LOC112206744.LOC107973052.GGT2-.646279-708040.rev.compl      cctcaatacctgcgtgagcacagtcatgtccaggccagggctgctggtcgaggtccggtc   31854
FAM230B-LOC105372935-GGT2.NCBI.GGT2.rev.compl                      cctcaatacctgggtgagcacagtcatgtccaggccagggctgctggtcgaggtccggcc   30493
                                                                   ************ ********************************************* *

chimp.LOC112206744.LOC107973052.GGT2-.646279-708040.rev.compl      ccatctcttccagcagaaagggagtaagcttgcagggaggctgggggacaagatcccagg   31914
FAM230B-LOC105372935-GGT2.NCBI.GGT2.rev.compl                      ccatctcttccagcagaaagggagtaagcttgcagggcggctgggggacaagatcccagg   30553
                                                                   ************************************* **********************

chimp.LOC112206744.LOC107973052.GGT2-.646279-708040.rev.compl      atctcggcctccgctcatggatcagctctgataccccgagtgagctgggggtgctctgtg   31974
FAM230B-LOC105372935-GGT2.NCBI.GGT2.rev.compl                      atctcagcctctgctcatggatcagctctgagaccccgagtgagctgggggtgctctgtg   30613
                                                                   ***** ***** ******************* ****************************

chimp.LOC112206744.LOC107973052.GGT2-.646279-708040.rev.compl      cgcattggtttccccagctgtcaagtaaagggattgtatgaggaagtcttgtcaaggtgg   32034
FAM230B-LOC105372935-GGT2.NCBI.GGT2.rev.compl                      cgcattggtttccccagctgtcaagtaaagggattggatgaggaagtcttgtcaaggtgg   30673
                                                                   ************************************ ***********************

chimp.LOC112206744.LOC107973052.GGT2-.646279-708040.rev.compl      aatgatctcagatttggggcagcagtgaatgatcccgctccctgggccatgccagtggcc   32094
FAM230B-LOC105372935-GGT2.NCBI.GGT2.rev.compl                      aatgatctcagatttggggcagcagtgaatgatcccgctccctgggccatgccagtggcc   30733
                                                                   ************************************************************

chimp.LOC112206744.LOC107973052.GGT2-.646279-708040.rev.compl      cggcctcggctcaacacagccccaacactctggaatggggatgagggggcagtcagctct   32154
FAM230B-LOC105372935-GGT2.NCBI.GGT2.rev.compl                      cggcctcggctgaacacagccccaacactctggaatggggatgagggggcagtcagctct   30793
                                                                   *********** ************************************************

chimp.LOC112206744.LOC107973052.GGT2-.646279-708040.rev.compl      tgctcctagtaagagagatgcaacggggctctgtggctgagctgggtgccttgcctcaca   32214
FAM230B-LOC105372935-GGT2.NCBI.GGT2.rev.compl                      tgctcctagtaagagagatgcaacagggctctgtggctgagctgggtgccttgcctcaca   30853
                                                                   ************************ ***********************************

chimp.LOC112206744.LOC107973052.GGT2-.646279-708040.rev.compl      cctgtaatcccaacctttgagaggctgaggcaggaggattgctcgaggccgggaattttg   32274
FAM230B-LOC105372935-GGT2.NCBI.GGT2.rev.compl                      cctgtaatcccaacctttgagaggccgaggcaggaggattgctcgaggccgggaattttg   30913
                                                                   ************************* **********************************

chimp.LOC112206744.LOC107973052.GGT2-.646279-708040.rev.compl      agaatagcctggacaacatagccagaccccatgtctacaaaataataataaaacatacag   32334
FAM230B-LOC105372935-GGT2.NCBI.GGT2.rev.compl                      agaatagcctggacaacatagccagaccccatgtctacaaaataataagaaaacacacag   30973
                                                                   ************************************************ ****** ****

chimp.LOC112206744.LOC107973052.GGT2-.646279-708040.rev.compl      ctatagtccaggctacttggcaggctgaggcaggagggtcccttgagtccaggaattgga   32394
FAM230B-LOC105372935-GGT2.NCBI.GGT2.rev.compl                      ctatagtccaagctacttggcaggctgaggcaggagggtcccttgagtccaggaattgga   31033
                                                                   ********** *************************************************

chimp.LOC112206744.LOC107973052.GGT2-.646279-708040.rev.compl      ggctgcattgagctataatcgcaccactgcactccagcttgggtgacaaagtgagaccct   32454
FAM230B-LOC105372935-GGT2.NCBI.GGT2.rev.compl                      ggctgcattgagctataatcgcaccactgcactccagcttgggtgacaaagtgagaccct   31093
                                                                   ************************************************************

chimp.LOC112206744.LOC107973052.GGT2-.646279-708040.rev.compl      gtctctaaaagaaaaaacattggcctgtgagcatgggtttgagtttcaaacaggagccgg   32514
FAM230B-LOC105372935-GGT2.NCBI.GGT2.rev.compl                      gtctctaaaagaaaaaaaattggcctgtgagcatgggcttgattttcaaacaggacccgg   31153
                                                                   ***************** ******************* **** ************ ****

chimp.LOC112206744.LOC107973052.GGT2-.646279-708040.rev.compl      agggtagggtaaacttgtcggtaaatctaaatgaatgttattggtataaaattacaatag   32574
FAM230B-LOC105372935-GGT2.NCBI.GGT2.rev.compl                      agggtagggtaaacgtgtgggtaaatctaaatgaatgttattggtataaaattacagtag   31213
                                                                   ************** *** ************************************* ***

chimp.LOC112206744.LOC107973052.GGT2-.646279-708040.rev.compl      tatagaaaatgatatcttgtggggtttaaaataaataaaacattctgaaatatgtatggg   32634
FAM230B-LOC105372935-GGT2.NCBI.GGT2.rev.compl                      tatagaaaatgatatcttgtggggtttaaaataaataaaacatactgaaatatgtattgg   31273
                                                                   ******************************************* ************* **

chimp.LOC112206744.LOC107973052.GGT2-.646279-708040.rev.compl      tacagttatatatctgggatttgcactgaaataatgtggggtagagggaagcaggaaaga   32694
FAM230B-LOC105372935-GGT2.NCBI.GGT2.rev.compl                      tacagttatatatctgggatttgcactgaaataatgtggggtagagggaagcaggaaaga   31333
                                                                   ************************************************************

chimp.LOC112206744.LOC107973052.GGT2-.646279-708040.rev.compl      ggatacatgaaattagcttggccataagattgttgttgaaattgaatggatactcggggc   32754
FAM230B-LOC105372935-GGT2.NCBI.GGT2.rev.compl                      gtatacatgaaatgagcttggccataagattgttgttgaaattgaatggatactcggggc   31393
                                                                   * *********** **********************************************

chimp.LOC112206744.LOC107973052.GGT2-.646279-708040.rev.compl      ttcattacacaattcttttactcttacatagctctacactctcaacataaataagaata   32814
FAM230B-LOC105372935-GGT2.NCBI.GGT2.rev.compl                      ttcattacacaattctctttactcttacatagctctacactctcaacataaataagaata   31453
                                                                   *************** ********************************************

chimp.LOC112206744.LOC107973052.GGT2-.646279-708040.rev.compl      aaaacacaaaaaacacacagatacatctatgcacacacatgtattttaaatacacaaaaa   32874
FAM230B-LOC105372935-GGT2.NCBI.GGT2.rev.compl                      aaaacacaaaaaacacacagatacatctatgcacacacacatatttaaaatacacaaaaa   31513
                                                                   ***************************************  ***** *************

chimp.LOC112206744.LOC107973052.GGT2-.646279-708040.rev.compl      tattagcatataagtcactggggtaaatttagttcctgttccaaggttcttgtactgac   32934
FAM230B-LOC105372935-GGT2.NCBI.GGT2.rev.compl                      tattagcatataagtcactggggtaaatttagttcctgttccaaggttcttgtactgac   31573
                                                                   ************************************************************

chimp.LOC112206744.LOC107973052.GGT2-.646279-708040.rev.compl      taggaagaggatagaagtactaactcataggctgggcgtggtggctcacgcctgtaatcc   32994
FAM230B-LOC105372935-GGT2.NCBI.GGT2.rev.compl                      taggaagaggatagaagtactaactcataggctgggcgcggtggctcacgcctgtaatcc   31633
                                                                   ************************************** *********************

chimp.LOC112206744.LOC107973052.GGT2-.646279-708040.rev.compl      caacactttaggacgccgaggtaagcagatctcttaaggtcaggagttcaagaccagcct   33054
FAM230B-LOC105372935-GGT2.NCBI.GGT2.rev.compl                      caacactttaggatgccgaggtaggcagatctcttaaggtccggagttcaagaccagcct   31693
                                                                   ************* ********** **************** ******************

chimp.LOC112206744.LOC107973052.GGT2-.646279-708040.rev.compl      ggccaacatggtgaaaccctgtctctactaaaaaagaatacaaaaattggccgggcatag   33114
FAM230B-LOC105372935-GGT2.NCBI.GGT2.rev.compl                      ggccaacatggtgaaaccctgtctctactgaaaaagaatacaaaaattggccgggcatag   31753
                                                                   ***************************** ******************************

chimp.LOC112206744.LOC107973052.GGT2-.646279-708040.rev.compl      tggtgcacacctgtggtcccagctactcaggtgactgaggcaggagaattgcttgaaccc   33174
FAM230B-LOC105372935-GGT2.NCBI.GGT2.rev.compl                      tggtgcacacctgtggtcccagctactcaggtgactgaggcaggagaattgcttgaaccc   31813
                                                                   ************************************************************

chimp.LOC112206744.LOC107973052.GGT2-.646279-708040.rev.compl      aagaagtggaggttgcagtgaaccaagattgctccactgcactccagcctgggcagcaga   33234
FAM230B-LOC105372935-GGT2.NCBI.GGT2.rev.compl                      aagaagtggaggttgcagtgaaccaagattgctccactgcactccagcctgggcagcaga   31873
                                                                   ************************************************************

chimp.LOC112206744.LOC107973052.GGT2-.646279-708040.rev.compl      ggaagactctctctgtctcaacaacaacaaaaagtactagctcatgttagactttgataa   33294
FAM230B-LOC105372935-GGT2.NCBI.GGT2.rev.compl                      ggaagactctctctatctcaaccacaacaaaaagtactagctcatgttagactttgataa   31933
                                                                   ************** ******* *************************************
```

```
chimp.LOC112206744.LOC107973052.GGT2-.646279-708040.rev.compl      gtgaaggatgcatgttgtaagctctaaaataatccactgtcatcttttaaaataactcta   33354
FAM230B-LOC105372935-GGT2.NCBI.GGT2.rev.compl                     gggaaggatgcatgttgtaagctctaaaataatcca--gtcatcttttaaaataactcta   31991
                                                                  * **********************************  **********************

chimp.LOC112206744.LOC107973052.GGT2-.646279-708040.rev.compl      agactgcacagttatgaaactaatag---aggaggaaattaaataataaaaataataaat   33411
FAM230B-LOC105372935-GGT2.NCBI.GGT2.rev.compl                     agactgcacagttatgaaactaatagagaaggaggaaattaaataataaaaataataaat   32051
                                                                  **************************   *******************************

chimp.LOC112206744.LOC107973052.GGT2-.646279-708040.rev.compl      ccaaaacaagatgtgagaggagataagaagaaatagaataggcatggaaaacaaattggt   33471
FAM230B-LOC105372935-GGT2.NCBI.GGT2.rev.compl                     ccaaaacaagatgtgagaggagataagaagaaatagaataggcatggaaaacaaattggt   32111
                                                                  ************************************************************

chimp.LOC112206744.LOC107973052.GGT2-.646279-708040.rev.compl      ggtgggtttcaacccaaataaatcattagttacatttaaaaggacaataaaaattaaaat   33531
FAM230B-LOC105372935-GGT2.NCBI.GGT2.rev.compl                     ggtgggtttcaacccaaataaatcattagttacatttaaaaggacaataaaaattaaaat   32171
                                                                  ************************************************************

chimp.LOC112206744.LOC107973052.GGT2-.646279-708040.rev.compl      aatcgaaaataaagtaaaacccaactaatgccttttatataagaatacagagaggtggaa   33591
FAM230B-LOC105372935-GGT2.NCBI.GGT2.rev.compl                     aattgaaaataaagtaaaacccaactaatgccttttatataagggtacagagaggtggaa   32231
                                                                  *** ***************************************  ***************

chimp.LOC112206744.LOC107973052.GGT2-.646279-708040.rev.compl      gataatgaaaaatatgtcatgcatgcactaaccaagaaagctgtataac-----------   33640
FAM230B-LOC105372935-GGT2.NCBI.GGT2.rev.compl                     gatcatgaaaaatatgtcatgcatgtactaaccaagaaagctgtataacttttttttttt   32291
                                                                  *** ********************* ***********************

chimp.LOC112206744.LOC107973052.GGT2-.646279-708040.rev.compl      ttttttttttttttttggagatagagcctcactctgtctcccaggctggagtgcagtgat   33700
FAM230B-LOC105372935-GGT2.NCBI.GGT2.rev.compl                     ttttttttttttttttggagatagagcctcactctgtctcccaggctggagtgcagtgat   32351
                                                                  ************************************************************

chimp.LOC112206744.LOC107973052.GGT2-.646279-708040.rev.compl      gtgatcttggcttacagcaatctctcccttctaggctcaagcgattctcccacctcagca   33760
FAM230B-LOC105372935-GGT2.NCBI.GGT2.rev.compl                     gtgatcttggcttacagcaatctctcccttctaggctcaagcgattctcccacctcagca   32411
                                                                  ************************************************************

chimp.LOC112206744.LOC107973052.GGT2-.646279-708040.rev.compl      tcccaagtagctgggactacaagtgtgccaacttagaattatattagccacacccagcta   33820
FAM230B-LOC105372935-GGT2.NCBI.GGT2.rev.compl                     tcccaagtagctgggactacaagtgtgccaacttagaattatattagccacacccagcta   32471
                                                                  ************************************************************

chimp.LOC112206744.LOC107973052.GGT2-.646279-708040.rev.compl      atttttgtattttttgtagaggcagggtctcgccatgttgcccaggctggtcttgaactc   33880
FAM230B-LOC105372935-GGT2.NCBI.GGT2.rev.compl                     atttttgtattttttgtagaggcagggtctcgccatgttgcccaggttggtcttgaactc   32531
                                                                  ********************************************** *************

chimp.LOC112206744.LOC107973052.GGT2-.646279-708040.rev.compl      ctgggcttcagtgatccacccacctcgacatccagcaaagtgccaagattacagccatga   33940
FAM230B-LOC105372935-GGT2.NCBI.GGT2.rev.compl                     ctgggcttcagtgatccacccacctcgacctccagcaaagtgccaagattacagccatga   32591
                                                                  ***************************** ******************************

chimp.LOC112206744.LOC107973052.GGT2-.646279-708040.rev.compl      gccaccatgcccagcataaccatttttaatgaagtagactttaagaagaaaagtattatt   34000
FAM230B-LOC105372935-GGT2.NCBI.GGT2.rev.compl                     gccaccatgcccagcataactatttttaatgaagtagactttaagaagaaaagtattatt   32651
                                                                  ******************** ***************************************

chimp.LOC112206744.LOC107973052.GGT2-.646279-708040.rev.compl      agaggtaagagacacatcacggaaaagaagaatttactaggagccaggcgcagtggctca   34060
FAM230B-LOC105372935-GGT2.NCBI.GGT2.rev.compl                     agaggtaagagacacatcacagaaaagaagaatttactaggagccaggcgcaatggctcg   32711
                                                                  ******************** ******************************* ******

chimp.LOC112206744.LOC107973052.GGT2-.646279-708040.rev.compl      tgcctgtaattccagcactttgtgaggccaaggcggcggatcacctgaggttgggagttc   34120
FAM230B-LOC105372935-GGT2.NCBI.GGT2.rev.compl                     tgcctgtaattccagcactttgtgaggccaaggcggcggatcacctgaggttgggagttc   32771
                                                                  ************************************************************

chimp.LOC112206744.LOC107973052.GGT2-.646279-708040.rev.compl      aagaccagcctgaccaacatggagaagccctgtctctactaaaaatacaaaaattagcca   34180
FAM230B-LOC105372935-GGT2.NCBI.GGT2.rev.compl                     aagaccagcctgaccaacatggagaagccctgtctctactaaaaatacaaaaattagcca   32831
                                                                  ************************************************************

chimp.LOC112206744.LOC107973052.GGT2-.646279-708040.rev.compl      agcatggtggcacatgcctgtaatcccagctactcaggaggctgagggaggagaattgct   34240
FAM230B-LOC105372935-GGT2.NCBI.GGT2.rev.compl                     agcatggtggcacatgcctgtaatcccagctactcaggaggctgagggaggagaattgct   32891
                                                                  ************************************************************

chimp.LOC112206744.LOC107973052.GGT2-.646279-708040.rev.compl      tggacccaggaagtggaggttgcggtgagctgagattgtgccattgcattccagcctggg   34300
FAM230B-LOC105372935-GGT2.NCBI.GGT2.rev.compl                     tggacccaggaagtggaggttgcggtgagctgagattgtgccattgcattccagcctggg   32951
                                                                  ************************************************************

chimp.LOC112206744.LOC107973052.GGT2-.646279-708040.rev.compl      caacaagagcaaaactctgtctc----------aaaaaaaaagaagttactagctagtta   34350
FAM230B-LOC105372935-GGT2.NCBI.GGT2.rev.compl                     caacaagagcaaaactctgtctcaaaaaaaaaaaaaaaaaaagaagttactagctagttt   33011
                                                                  ***********************          **************************

chimp.LOC112206744.LOC107973052.GGT2-.646279-708040.rev.compl      cggtaattcttaacatccaggaaactggttgtgaaagtttttcagagaaactaaaccaat   34410
FAM230B-LOC105372935-GGT2.NCBI.GGT2.rev.compl                     cggtaattcttaacaaccaggaaactggatgtgaaagtttttcagagaaactaaaccaat   33071
                                                                  *************** ************ *******************************

chimp.LOC112206744.LOC107973052.GGT2-.646279-708040.rev.compl      agattatacatagagagagatttattgaggaattggctcacatgattgtggggactagca   34470
FAM230B-LOC105372935-GGT2.NCBI.GGT2.rev.compl                     agattatacatagagagagatttatttaggaattggctcacatgattgtggggactagca   33131
                                                                  ************************** *********************************

chimp.LOC112206744.LOC107973052.GGT2-.646279-708040.rev.compl      ag-tttaaaatctgtagggcaagccagcaggctataaattcaggtaagagttgatctcga   34529
FAM230B-LOC105372935-GGT2.NCBI.GGT2.rev.compl                     agttttaaaatctgtagggcaagccagcaggctataaattcaggtaagagttgatctcga   33191
                                                                  ** *********************************************************

chimp.LOC112206744.LOC107973052.GGT2-.646279-708040.rev.compl      agtctggaacctaaaatctgtagagcagtcagcaggccagaaactcaggcagggtttgtg   34589
FAM230B-LOC105372935-GGT2.NCBI.GGT2.rev.compl                     agtctggaacctaaaatctgtagagcagtcagcaggccagaaactcaggcagggtttgtg   33251
                                                                  ************************************************************

chimp.LOC112206744.LOC107973052.GGT2-.646279-708040.rev.compl      tgttacagtcttgaagcagaattcctgcttctttgggaaacct--gtttttgttcttaag   34647
FAM230B-LOC105372935-GGT2.NCBI.GGT2.rev.compl                     tgttacagtcttgaagcagaattcctgcttctctgggaaacctcagtttttgttcttaag   33311
                                                                  ******************************** **********  ***************

chimp.LOC112206744.LOC107973052.GGT2-.646279-708040.rev.compl      gccttcaagtgattggaggtggcccacccatattatggtgggtaatctgttttacttaaa   34707
FAM230B-LOC105372935-GGT2.NCBI.GGT2.rev.compl                     gccttcaactgattggaggtggcccacccatattatggtgggtaatctgttttacttaaa   33371
                                                                  ******** ***************************************************

chimp.LOC112206744.LOC107973052.GGT2-.646279-708040.rev.compl      gtcaattgactgtcagtgttaaccacatctatgaaataacctcccagcaagatattgaca   34767
FAM230B-LOC105372935-GGT2.NCBI.GGT2.rev.compl                     gtcaattgactgtcagtgttaatcacatctatgaaataacctcccagcaagatattgaca   33431
                                                                  ********************** *************************************

chimp.LOC112206744.LOC107973052.GGT2-.646279-708040.rev.compl      agtatttgaccaaacaacggggcaccatagcttagccaagttgacacataaattaaccat   34827
FAM230B-LOC105372935-GGT2.NCBI.GGT2.rev.compl                     agtatttgaccaaacaacaggacaccatagcttagccaagttgacacataaattaaccat   33491
                                                                  ****************** ** **************************************

chimp.LOC112206744.LOC107973052.GGT2-.646279-708040.rev.compl      caggagcgagtagaatatccaaaaaacaacatactaggggtattatatcttatatagc--   34885
FAM230B-LOC105372935-GGT2.NCBI.GGT2.rev.compl                     caggagcaagtagaatatccaaaaaacaacatactaggggtattatatcttatatagcta   33551
                                                                  ******* **************************************************

chimp.LOC112206744.LOC107973052.GGT2-.646279-708040.rev.compl      ---taattatataaaacatataattatagaatgaagatattaagataaccattagaacaa   34942
FAM230B-LOC105372935-GGT2.NCBI.GGT2.rev.compl                     ttataattatataaaacatataattatagaatgacgatattaagataaccattagaacaa   33611
                                                                     ******************************* *************************

chimp.LOC112206744.LOC107973052.GGT2-.646279-708040.rev.compl      aaatataaactttctt------ttctttttttttttgagaccaagtcttgctctgtcac   34996
FAM230B-LOC105372935-GGT2.NCBI.GGT2.rev.compl                     aaatataaactttctttctttctttttttttttttttgagaccaagtcttgctctgtcac   33671
                                                                  ****************      ** ***********************************

chimp.LOC112206744.LOC107973052.GGT2-.646279-708040.rev.compl      ccaggctggagtgcagtggtgcaatcttggcttactgcaacctttgcctcctgagttcaa   35056
FAM230B-LOC105372935-GGT2.NCBI.GGT2.rev.compl                     ccaggctggagtgcagtggtgcaatcttggcttactgaaacctttgcctcctgggttcaa   33731
                                                                  ************************************* *************** ******

chimp.LOC112206744.LOC107973052.GGT2-.646279-708040.rev.compl      gtgatcctcctgtctcagcctcccaagtagctgggattacaggcacctgctaccatgccc   35116
FAM230B-LOC105372935-GGT2.NCBI.GGT2.rev.compl                     gtgattctcctgtctcagcctcccaagtatctgggattacaggcacctgctaccatgccc   33791
                                                                  ***** *********************** ******************************

chimp.LOC112206744.LOC107973052.GGT2-.646279-708040.rev.compl      agctaatttttgtatttttagtagagacatggtttcaccatgttgcccaggctggtctcc   35176
FAM230B-LOC105372935-GGT2.NCBI.GGT2.rev.compl                     agctaatttttgtatttttagtagagacatggtttcaccatgttgcccaggctggtctcc   33851
                                                                  ************************************************************

chimp.LOC112206744.LOC107973052.GGT2-.646279-708040.rev.compl      aactcctgacctcaagtgagccagccccctcggcctcccaaagtgctgggattacaggtg   35236
FAM230B-LOC105372935-GGT2.NCBI.GGT2.rev.compl                     aactcctgacctcaagtgagccacccccctcggcctcccaaagtgctgggattacaggtg   33911
                                                                  *********************** ************************************

chimp.LOC112206744.LOC107973052.GGT2-.646279-708040.rev.compl      tgagccaccacacccagccaaaatcacctttttcacaaggatcaaaacagtcattacgc   35296
FAM230B-LOC105372935-GGT2.NCBI.GGT2.rev.compl                     tgagccaccacacccagccaaaatcaccttttttacaaggatcaaaacagtcattatgc   33971
                                                                  ********************************** ********************** **

chimp.LOC112206744.LOC107973052.GGT2-.646279-708040.rev.compl      tggagatgacagacctcactgtcaccatgctccttttgtatgtctactaggcatggtgct   35356
FAM230B-LOC105372935-GGT2.NCBI.GGT2.rev.compl                     tggagatgacagacctcactgtcaccatgctccttttgtatgtctactaggcacggtgct   34031
                                                                  ***************************************************** ******

chimp.LOC112206744.LOC107973052.GGT2-.646279-708040.rev.compl      gggtccacactcacagaaagcttaggagctcgcacccaggggctccggctgtagcagaat   35416
FAM230B-LOC105372935-GGT2.NCBI.GGT2.rev.compl                     gggtccacactcacagaaaccttaggaactcgcacccaggggctccggctgtagcagaat   34091
                                                                  ******************* ******* ********************************

chimp.LOC112206744.LOC107973052.GGT2-.646279-708040.rev.compl      cccaagaataaaacctggtgctgaaagagtaggagatgaggccggatgccatgactcact   35476
```

```
FAM230B-LOC105372935-GGT2.NCBI.GGT2.rev.compl                    cccaagaataaaacctggtgctgaaagagtaggagatgaggccaggtgccatgactcact   34151
                                                                 ******************************************* * **************

chimp.LOC112206744.LOC107973052.GGT2-.646279-708040.rev.compl    cctgtaatgccagcactttgggtggccaaggcgggtgaatcaagagatagagaccatcct   35536
FAM230B-LOC105372935-GGT2.NCBI.GGT2.rev.compl                    cctgtaatgccagcactttgggtggccaaggcgggtgaatcaagagatagagaccatcct   34211
                                                                 ************************************************************

chimp.LOC112206744.LOC107973052.GGT2-.646279-708040.rev.compl    ggccaacatggtgaaaccccgtctctactaaaaatacaaaaattagctgggcgtggtggc   35596
FAM230B-LOC105372935-GGT2.NCBI.GGT2.rev.compl                    ggccaacacggtgaaaccccgtctctactaaaaatacagaaattagcggggcgtggtggc   34271
                                                                 ******** ***************************** ******** ************

chimp.LOC112206744.LOC107973052.GGT2-.646279-708040.rev.compl    tggcacctgtagtcccagctactcaggaggctgaggcaggagaatcatttgaaccgagga   35656
FAM230B-LOC105372935-GGT2.NCBI.GGT2.rev.compl                    tggcacctgtagtcccagctactcaggaggctgaggcaggagaatcatttgaaccgagga   34331
                                                                 ************************************************************

chimp.LOC112206744.LOC107973052.GGT2-.646279-708040.rev.compl    agcagaggttgcagtaagctgagatcgtgccactgcactccagcctggtgacagagtgag   35716
FAM230B-LOC105372935-GGT2.NCBI.GGT2.rev.compl                    agcagaggttgcagtaagctgagatcgcgccactgcactccagcctggtgacagagtgag   34391
                                                                 *************************** ********************************

chimp.LOC112206744.LOC107973052.GGT2-.646279-708040.rev.compl    acaccgtctcaaaaaaaaaaaaaaaaaaaaaaaagcaggacactgaactctgggagggcc   35776
FAM230B-LOC105372935-GGT2.NCBI.GGT2.rev.compl                    acaccgtc-------acaaaaaaaaaaaaaaaaagcaggacactgaactctgggagggcc   34444
                                                                 ********       * *******************************************

chimp.LOC112206744.LOC107973052.GGT2-.646279-708040.rev.compl    tcctggtgagaggtgagcacagaggggagagatggaggcaggagcatgggcttctggtgg   35836
FAM230B-LOC105372935-GGT2.NCBI.GGT2.rev.compl                    tcctggtgagaggtgagcacagaggggagagatggaggcaggagcatgggcttctggtgg   34504
                                                                 ************************************************************

chimp.LOC112206744.LOC107973052.GGT2-.646279-708040.rev.compl    ccccagcagaccctgtggcagcgtggccagggtcctctgcagggaggaatcttggccagg   35896
FAM230B-LOC105372935-GGT2.NCBI.GGT2.rev.compl                    ccccagcagaccctgtggcagcgtggccagggtcctctgcagggaggaatcttggccagg   34564
                                                                 ************************************************************

chimp.LOC112206744.LOC107973052.GGT2-.646279-708040.rev.compl    atgacgctgtagcaggcctcttcctgaggcctccagccagcccggccagggacccagcgt   35956
FAM230B-LOC105372935-GGT2.NCBI.GGT2.rev.compl                    atgacgctgtagcaggcctcttcctgaggcctccagccagcccggccagggtcccagcgt   34624
                                                                 *************************************************** ********

chimp.LOC112206744.LOC107973052.GGT2-.646279-708040.rev.compl    ccagtgacccctgtttcacagcagcagctggggccagccccaggctctcttccactccca   36016
FAM230B-LOC105372935-GGT2.NCBI.GGT2.rev.compl                    ccagtgacccctgtttcacagcagcagctggggccagccccaggctctcttccactccca   34684
                                                                 ************************************************************

chimp.LOC112206744.LOC107973052.GGT2-.646279-708040.rev.compl    gcttcttaaaactggaagtggagagagttgtttgataaaacactggggcaaaccacatcc   36076
FAM230B-LOC105372935-GGT2.NCBI.GGT2.rev.compl                    gcttcttaaaactggaagtggagagagttgtttgataaaacactggggcaaaccacatcc   34744
                                                                 ************************************************************

chimp.LOC112206744.LOC107973052.GGT2-.646279-708040.rev.compl    tctcttcaccaagggagagttcgaggggatgccagcagagggcgctttagagtagagacc   36136
FAM230B-LOC105372935-GGT2.NCBI.GGT2.rev.compl                    tctcttcaccaagggagagttggaggggatgccggcagagggagctttagagtagagacc   34804
                                                                 ********************* *********** ******** *****************

chimp.LOC112206744.LOC107973052.GGT2-.646279-708040.rev.compl    cctacccaaccagtgaccgtcacgcacacagcagggcatgctatggagacccccagacag   36196
FAM230B-LOC105372935-GGT2.NCBI.GGT2.rev.compl                    cctacccaaccagtgaccgtcacgcacacagcagggcatgctatggagacccccagacag   34864
                                                                 ************************************************************

chimp.LOC112206744.LOC107973052.GGT2-.646279-708040.rev.compl    tcactcggggagacccagcaggtccagactcttcagagatctgtggcagcaggtccccac   36256
FAM230B-LOC105372935-GGT2.NCBI.GGT2.rev.compl                    tcactcggggagacccagcaggtccagactcttcagagatctgtggcagcaggtccccac   34924
                                                                 ************************************************************

chimp.LOC112206744.LOC107973052.GGT2-.646279-708040.rev.compl    tcccaaaagccacgtgcccacgggtggtctctggtgcctgagaccccagtctcatttgca   36316
FAM230B-LOC105372935-GGT2.NCBI.GGT2.rev.compl                    tcccaaaagccacgtgcccacgggtggtctctggtgcctgagaccccagtctcatttgca   34984
                                                                 ************************************************************

chimp.LOC112206744.LOC107973052.GGT2-.646279-708040.rev.compl    tctttgcaacttcgagtttaagtgggtgtcgcatctctgtatgtcctcctgagcagagga   36376
FAM230B-LOC105372935-GGT2.NCBI.GGT2.rev.compl                    tctttgcaacttcgagtttaagtgggtgtcgcatctctgtatgtcctcccgagcagagga   35044
                                                                 ************************************************* **********

chimp.LOC112206744.LOC107973052.GGT2-.646279-708040.rev.compl    ggggcacagcctggggtggcagctggcgtcaaaccctcaaatcccctgagagccactggg   36436
FAM230B-LOC105372935-GGT2.NCBI.GGT2.rev.compl                    ggggcacagcctggggtggcagctggcgtcaaaccctcaaatcccctgagagccactggg   35104
                                                                 ************************************************************

chimp.LOC112206744.LOC107973052.GGT2-.646279-708040.rev.compl    gagactaagcagtccccagcccccacttgtccctgagctgccattctcagccctgtggga   36496
FAM230B-LOC105372935-GGT2.NCBI.GGT2.rev.compl                    gagactaagcagtccccagcccctacttgtccctgagctgccattctcagccctgtggga   35164
                                                                 ********************** *************************************

chimp.LOC112206744.LOC107973052.GGT2-.646279-708040.rev.compl    ggagacagaaagccctgaagagaaaccaaaggaccaggtcaggaggggct--gggggggt   36554
FAM230B-LOC105372935-GGT2.NCBI.GGT2.rev.compl                    ggagacagaaagccctgaagagaaaccaaaggaccaggtcaggaggggctgggggggggt   35224
                                                                 **************************************************  ********

chimp.LOC112206744.LOC107973052.GGT2-.646279-708040.rev.compl    ggcatgagcaatcagggcagggaaggatggacagatggggaatggagggaagaaggaat   36614
FAM230B-LOC105372935-GGT2.NCBI.GGT2.rev.compl                    ggcatgagcaatcagggcagggaaggatgcacagatggggaatggagggaagaaggaat   35284
                                                                 **************************** *******************************

chimp.LOC112206744.LOC107973052.GGT2-.646279-708040.rev.compl    gaatgaaacggtgaatgaatgaacaaagagagagaacggccattcctcccttgctttagt   36674
FAM230B-LOC105372935-GGT2.NCBI.GGT2.rev.compl                    gaatgaaaaggtgaatgaatgaacaaagagagagaacggccactcctcccttgctttagt   35344
                                                                 ******** ********************************* *****************

chimp.LOC112206744.LOC107973052.GGT2-.646279-708040.rev.compl    ttacaaagtactgggatcctcccaacagcctgcaagacagaatttctgggaagcagacca   36734
FAM230B-LOC105372935-GGT2.NCBI.GGT2.rev.compl                    ttacaaagtactgggatcctcccaacagcctgcaagacagaatttctgggaagcagacca   35404
                                                                 ************************************************************

chimp.LOC112206744.LOC107973052.GGT2-.646279-708040.rev.compl    ggtggctggcagggaggggaggcttgccctggcttttgtgggcccaatgggaggcagggg   36794
FAM230B-LOC105372935-GGT2.NCBI.GGT2.rev.compl                    ggtggctggcagggaggggaggcttgccctggcttttgtgggcccaatgggaggcagggg   35464
                                                                 ************************************************************

chimp.LOC112206744.LOC107973052.GGT2-.646279-708040.rev.compl    gcaagaaggggcatcctgtgtgtgtcctccctgcagcggcagcagcaccttcctggaaga   36854
FAM230B-LOC105372935-GGT2.NCBI.GGT2.rev.compl                    gcaagaaggggcatcctgtgtgtgtcctccctgcagcggcagcagcaccttcctggaaga   35524
                                                                 ************************************************************

chimp.LOC112206744.LOC107973052.GGT2-.646279-708040.rev.compl    gggtcaggaaacactcactgtggcccctctccatcacgacctcatccaggacaccaaata   36914
FAM230B-LOC105372935-GGT2.NCBI.GGT2.rev.compl                    gggtcaggaaacacccgctgtggcccctctccaccacgccctcatccaggacaccaagta   35584
                                                                 ************** * **************** **** ****************** **

chimp.LOC112206744.LOC107973052.GGT2-.646279-708040.rev.compl    tcaagtcactcagctcacgagacccaggccctgactcagggagagaggatgtgaggggtg   36974
FAM230B-LOC105372935-GGT2.NCBI.GGT2.rev.compl                    tc-agtcactcagctcacgagacccaggccctgactcagggagagaggatgtgaggggtg   35643
                                                                 ** *********************************************************

chimp.LOC112206744.LOC107973052.GGT2-.646279-708040.rev.compl    gggcaccgggctcctcaggactgagagacctgagatgtggccccgggctgggtgttgggg   37034
FAM230B-LOC105372935-GGT2.NCBI.GGT2.rev.compl                    gggcaccgggctcctcaggactgagagacctgagatgtggccccgggctgggtgtagggg   35703
                                                                 ******************************************************* ****

chimp.LOC112206744.LOC107973052.GGT2-.646279-708040.rev.compl    cagactggctatggcagcattgtgtgtaccccagcaggccagtacccatgcagggagcct   37094
FAM230B-LOC105372935-GGT2.NCBI.GGT2.rev.compl                    cagactggctatggcagcattgtgtgtaccccagcaggccagtacccacgcagggagcct   35763
                                                                 ************************************************ ***********

chimp.LOC112206744.LOC107973052.GGT2-.646279-708040.rev.compl    ccaaaccccttcacccatgaccctgggagaagaccgcaggcttggagaattggcctcact   37154
FAM230B-LOC105372935-GGT2.NCBI.GGT2.rev.compl                    ccaaaccccttcacccatgaccctgggagaagaccgcagccttggagaattggcctcact   35823
                                                                 *************************************** ********************

chimp.LOC112206744.LOC107973052.GGT2-.646279-708040.rev.compl    gaagggccctgcaccggccagcagggtcaggcggggccagacaggttcccacctgggata   37214
FAM230B-LOC105372935-GGT2.NCBI.GGT2.rev.compl                    gaaggggcctgcgccggccagcagggtcaggcggggccagacaggttcccacctgggata   35883
                                                                 ****** ***** ***********************************************

chimp.LOC112206744.LOC107973052.GGT2-.646279-708040.rev.compl    tgcaaatgggcctcctgaatcctggagccaggtatggactcacacaccaccattgtcccc   37274
FAM230B-LOC105372935-GGT2.NCBI.GGT2.rev.compl                    tgcaaatgggcctcctgaatcctggagccaggtatggactcacacaccaccattgtcccc   35943
                                                                 ************************************************************

chimp.LOC112206744.LOC107973052.GGT2-.646279-708040.rev.compl    aagtccccatctgccccacgggcacaccctgccacctgttctgtgcaagagccctgaggc   37334
FAM230B-LOC105372935-GGT2.NCBI.GGT2.rev.compl                    aagtccccatctgccccacgggcacaccctgccgcctgttctgtgcaagggccctgaggc   36003
                                                                 ********************************* *************** **********

chimp.LOC112206744.LOC107973052.GGT2-.646279-708040.rev.compl    tgtctccttgcgctcaagccctgcaggtgctgaagcccacacacacagctcctgcttcct   37394
FAM230B-LOC105372935-GGT2.NCBI.GGT2.rev.compl                    tgtctccttgcgctcaagccctgcaggtgctgaagcccacacacacggctcctgcttcct   36063
                                                                 ********************************************** *************

chimp.LOC112206744.LOC107973052.GGT2-.646279-708040.rev.compl    gggccagtgcacgtgcacacacacacacgtgcacacacaca------cccacaaatatac   37448
FAM230B-LOC105372935-GGT2.NCBI.GGT2.rev.compl                    gggccagtgcacgtgcacacacacacacgtgcacacacacacacaccccacaaatatac   36123
                                                                 *****************************************      *************

chimp.LOC112206744.LOC107973052.GGT2-.646279-708040.rev.compl    ccacacacaatcacacacattcacacatacccacccccatacactcacacattcacaccc   37508
FAM230B-LOC105372935-GGT2.NCBI.GGT2.rev.compl                    ccacacacaatcacacacattcacacatacccacccccccatactcaca-----------   36172
                                                                 **************************************  * *******

chimp.LOC112206744.LOC107973052.GGT2-.646279-708040.rev.compl    acacccacacatacacacacacaatcacacacatttacacacacccacacacccacactc   37568
FAM230B-LOC105372935-GGT2.NCBI.GGT2.rev.compl                    -------------------------ctcacacatttacacacacccacacacccacactc   36207
                                                                                          * *********************************

chimp.LOC112206744.LOC107973052.GGT2-.646279-708040.rev.compl    acacact--cagtcacacacaccctcacacagcgaaacacaat--cacacattcacaccc   37624
FAM230B-LOC105372935-GGT2.NCBI.GGT2.rev.compl                    acacactcacagtcacacacaccctcacacagcgaaacacaatcacacacattcacaccc   36267
```

```
                                                                   *******  **********************************  ***************

chimp.LOC112206744.LOC107973052.GGT2-.646279-708040.rev.compl      acccacacccccacacactcacactcacatatactcacacacacccacacacacatacac   37684
FAM230B-LOC105372935-GGT2.NCBI.GGT2.rev.compl                      acccacacccccacacactcacactcacatatactcacacacacccacacacacatacac   36327
                                                                   ************************************************************

chimp.LOC112206744.LOC107973052.GGT2-.646279-708040.rev.compl      aaacacaatcacacacattcacacaca---------------------------------   37711
FAM230B-LOC105372935-GGT2.NCBI.GGT2.rev.compl                      aaacacaatcacacacattcacacacacccacactcacacatacacacacccaaacacga   36387
                                                                   ***************************

chimp.LOC112206744.LOC107973052.GGT2-.646279-708040.rev.compl      ---------------------caccccccacaaacacactcacacatatacccacacaca   37750
FAM230B-LOC105372935-GGT2.NCBI.GGT2.rev.compl                      tcacacacattcacacacacccacaccccacaaacacactcacacatatacccacacaca   36447
                                                                                        *** ***********************************

chimp.LOC112206744.LOC107973052.GGT2-.646279-708040.rev.compl      ctcacactcacacataatctcacacacacacacatgctcacacacacacgccttctcc   37810
FAM230B-LOC105372935-GGT2.NCBI.GGT2.rev.compl                      ctcacacata--------atctctcacacacacacatgctcacacacacgccttctcc   36499
                                                                   *******  *         ** * ************************************

chimp.LOC112206744.LOC107973052.GGT2-.646279-708040.rev.compl      aggaggggctggctgccaaaggccacccagcttcctcccacgtctcactcaccgtacaat   37870
FAM230B-LOC105372935-GGT2.NCBI.GGT2.rev.compl                      aggaggggctggctgccaaaggccacccagcttcctcccacgtctcactcaccgtacaat   36559
                                                                   ************************************************************

chimp.LOC112206744.LOC107973052.GGT2-.646279-708040.rev.compl      atttgagcagaccttggagtcagcagcaacagcagcgtgggcaaaggcctgggagccaca   37930
FAM230B-LOC105372935-GGT2.NCBI.GGT2.rev.compl                      atttgagaagaccttggagtcagcagcaacagcggcgtgggcaaaggcc-gggggtcaaa   36618
                                                                   ******* ************************* *************** *** * ** *

chimp.LOC112206744.LOC107973052.GGT2-.646279-708040.rev.compl      tggggcctggtgtcgagagaggaccacagccagcacaatgacagccagcgccagccccag   37990
FAM230B-LOC105372935-GGT2.NCBI.GGT2.rev.compl                      tggggcctggtgtcgagagaggaccacagccagcacaatgacagccagcgccagccccag   36678
                                                                   ************************************************************

chimp.LOC112206744.LOC107973052.GGT2-.646279-708040.rev.compl      ccccagccccagcaggaccaggtcacccatggctctgcagtcctgggccatggctctgcg   38050
FAM230B-LOC105372935-GGT2.NCBI.GGT2.rev.compl                      ccccagccccagcaggaccaggtcacccatggctccgtagtcctgggccatggctctgcg   36738
                                                                   *********************************** * **********************

chimp.LOC112206744.LOC107973052.GGT2-.646279-708040.rev.compl      gcccagaaggagaggggaggccggtgggcagacggagggacagatgggtgggcagatgaa   38110
FAM230B-LOC105372935-GGT2.NCBI.GGT2.rev.compl                      gcccagaaggagaggggaggccggtgggcagatggagggacagatgggtgggcagatgaa   36798
                                                                   ******************************** ***************************

chimp.LOC112206744.LOC107973052.GGT2-.646279-708040.rev.compl      tggacaagaagatgcatagatagactcacaggtaattggacagatggacaaacaggtggg   38170
FAM230B-LOC105372935-GGT2.NCBI.GGT2.rev.compl                      tggacaagaagatgcatagatagactcacaggtaattggacagatggacaaacaggtggg   36858
                                                                   ************************************************************

chimp.LOC112206744.LOC107973052.GGT2-.646279-708040.rev.compl      ggctgaagacagacacgaagatggatcgacagacaggccagatagctagacaaagaggac   38230
FAM230B-LOC105372935-GGT2.NCBI.GGT2.rev.compl                      ggctgaagacagacacgaagatggatcgacagacaggccagatagctagacaaagaggac   36918
                                                                   ************************************************************

chimp.LOC112206744.LOC107973052.GGT2-.646279-708040.rev.compl      agtaagagaaagatggtcagatagacaatggaacagagatgggcttacagatgggcggac   38290
FAM230B-LOC105372935-GGT2.NCBI.GGT2.rev.compl                      agtaagagaaagatggtcagatagacaatgggacagagatgggcttacagatgggcggac   36978
                                                                   ******************************* ****************************

chimp.LOC112206744.LOC107973052.GGT2-.646279-708040.rev.compl      agacagacaggtctgaacagcgggctgccagatggacagatgggtgaatggacagatggc   38350
FAM230B-LOC105372935-GGT2.NCBI.GGT2.rev.compl                      agacagacaggtctgaacagcgggctgccagatggacagatgggtgaatggacagatggc   37038
                                                                   ************************************************************

chimp.LOC112206744.LOC107973052.GGT2-.646279-708040.rev.compl      tggcagctgtggcgagctgctgccctcaccaagtgcacactacggagtggccaaactcat   38410
FAM230B-LOC105372935-GGT2.NCBI.GGT2.rev.compl                      tggcagctgtggcgagctgctgccctcaccaagtgcacactacggagtggccaaactcat   37098
                                                                   ************************************************************

chimp.LOC112206744.LOC107973052.GGT2-.646279-708040.rev.compl      gcctcaacttctagttttcagctcctgcttgttcctggcaggaggccaggcagcaaagcg   38470
FAM230B-LOC105372935-GGT2.NCBI.GGT2.rev.compl                      gcctcaacttctagttttcagctcctgcttgttcctggcaggaggccaggcagcaaagcg   37158
                                                                   ************************************************************

chimp.LOC112206744.LOC107973052.GGT2-.646279-708040.rev.compl      tctgaggggagttttctttgcctagagaagtcagctgctgtgttaactccctcactgctg   38530
FAM230B-LOC105372935-GGT2.NCBI.GGT2.rev.compl                      tctgaggggagttttctttgcctagagaagtcagctgctgtgttaactccctcactgctg   37218
                                                                   ************************************************************

chimp.LOC112206744.LOC107973052.GGT2-.646279-708040.rev.compl      gtaggtccaaaggccccacctaccgcccgccagagcccatggtcacactgtcgcaatgtg   38590
FAM230B-LOC105372935-GGT2.NCBI.GGT2.rev.compl                      gtaggtccaaaggccccacctactgcccgccagagcccatggtcacactgtcgcaatgtg   37278
                                                                   *********************** ************************************

chimp.LOC112206744.LOC107973052.GGT2-.646279-708040.rev.compl      caggagaacttggtgcctgctgcactgcgattgccaggtaggggcagggctccctggaac   38650
FAM230B-LOC105372935-GGT2.NCBI.GGT2.rev.compl                      caggagaacttggtgcgtgctgcactgcggttgccaggtaggggcagggctccctggaac   37338
                                                                   **************** ************ ******************************

chimp.LOC112206744.LOC107973052.GGT2-.646279-708040.rev.compl      ctccacaccattccccacgttctcagcagctctgggaaagcagagctggggccgcttaac   38710
FAM230B-LOC105372935-GGT2.NCBI.GGT2.rev.compl                      ctccacaccattccccaggttctcagcagctctgggaaagcagagctggggccgcttaac   37398
                                                                   ***************** ******************************************

chimp.LOC112206744.LOC107973052.GGT2-.646279-708040.rev.compl      tctgtcctggatccggcaaggctgccccactcccagagtggagccctgctccccagctcc   38770
FAM230B-LOC105372935-GGT2.NCBI.GGT2.rev.compl                      tctgccctggatccggcaaggctgcccacctcccagagtggagccctgctccccagctcc   37458
                                                                   **** **********************  *******************************

chimp.LOC112206744.LOC107973052.GGT2-.646279-708040.rev.compl      catctctatcccctaaccctctccgcatggcccagcctagtcagcatcaaggtggagctg   38830
FAM230B-LOC105372935-GGT2.NCBI.GGT2.rev.compl                      catctctatcccctaaccctctccgcatggcccagcctagtcagcatcaaggtggagctg   37518
                                                                   ************************************************************

chimp.LOC112206744.LOC107973052.GGT2-.646279-708040.rev.compl      aacagaggcagaaggaggaggacccaaggtggtgtcactcaggacccgggttcaagtcct   38890
FAM230B-LOC105372935-GGT2.NCBI.GGT2.rev.compl                      aacagaggcagaaggaggaggacccaaggtggtgtcactcaggacccgggttcaagtcct   37578
                                                                   ************************************************************

chimp.LOC112206744.LOC107973052.GGT2-.646279-708040.rev.compl      tatgcttctgcagcctggcctgggtcccccaatcccccccagggtgaccaagggcttccca   38950
FAM230B-LOC105372935-GGT2.NCBI.GGT2.rev.compl                      tatgtttctgcagcctggcctgggtcccccaaccccccccagggtgaccaagggcttccca   37638
                                                                   **** *************************** ***************************

chimp.LOC112206744.LOC107973052.GGT2-.646279-708040.rev.compl      gtctgcacagaggacagggggtcttgacagcatcaaatgctggtgactatgagacactta   39010
FAM230B-LOC105372935-GGT2.NCBI.GGT2.rev.compl                      gtctgcacagaggacagggggtcttgacagcatcaaatgctggtgactatgagacactta   37698
                                                                   ************************************************************

chimp.LOC112206744.LOC107973052.GGT2-.646279-708040.rev.compl      catgggaaatgcagacagaccatgcctctagcccttggtacccggcaccatccatccctg   39070
FAM230B-LOC105372935-GGT2.NCBI.GGT2.rev.compl                      catgggaaatgcagacagaccatgcctctagcccttggcaccaggcaccatccatccctg   37758
                                                                   ************************************** *** *****************

chimp.LOC112206744.LOC107973052.GGT2-.646279-708040.rev.compl      ggacttgctgtcctggaaatgcagcatggacctccagggagggggctgtgccatgtggg   39130
FAM230B-LOC105372935-GGT2.NCBI.GGT2.rev.compl                      ggacttgctgtcctggaaatgcagcatggacctccagggagggggctgtgccatgtggg   37818
                                                                   ************************************************************

chimp.LOC112206744.LOC107973052.GGT2-.646279-708040.rev.compl      ggcctcaccccacctgcagctctttcccaccctggctgcaggtctgcttccctgaatcca   39190
FAM230B-LOC105372935-GGT2.NCBI.GGT2.rev.compl                      ggccccaccccacctgcagctctttcccaccctggctgcaggtctgcttctctgaatcca   37878
                                                                   **** ********************************************* *********

chimp.LOC112206744.LOC107973052.GGT2-.646279-708040.rev.compl      aatccgctactactgtgctggcagcgcagcctctctggggacactggcctggctctgttc   39250
FAM230B-LOC105372935-GGT2.NCBI.GGT2.rev.compl                      aatccgctactactgtgctggcagcgcagcctctctggggacactggcctggctctgttc   37938
                                                                   ************************************************************

chimp.LOC112206744.LOC107973052.GGT2-.646279-708040.rev.compl      tccccaggcctcagggtgcctaaatgggaggcaaccaggggagtgaggacccactgaggg   39310
FAM230B-LOC105372935-GGT2.NCBI.GGT2.rev.compl                      tccccaggcctcagggtgcctaaatgggaggcaaccaggggagtgaggacccactgaggg   37998
                                                                   ************************************************************

chimp.LOC112206744.LOC107973052.GGT2-.646279-708040.rev.compl      gctccgttgaccaggctcagcaggggtgcaggtgatgtgggctggacttcttcccacgtg   39370
FAM230B-LOC105372935-GGT2.NCBI.GGT2.rev.compl                      gctccgttgaccaggctcagcaggggtgcaggtgatgtgggtggaatccttcccacatg   38058
                                                                   ***************************************** **** * ******** **

chimp.LOC112206744.LOC107973052.GGT2-.646279-708040.rev.compl      gcccccacagtcctccccgcttcctccccagctgaacactgcctgctccagatgtctaca   39430
FAM230B-LOC105372935-GGT2.NCBI.GGT2.rev.compl                      gcccccacagtcctccccgcttcctccccaactgaacactgcctgctccagatgtgtaca   38118
                                                                   ****************************** ************************ ****

chimp.LOC112206744.LOC107973052.GGT2-.646279-708040.rev.compl      cctggagtccgggcccctccatctgggcagcagagaaactgaggcacagagacagactgt   39490
FAM230B-LOC105372935-GGT2.NCBI.GGT2.rev.compl                      cctggagtctgggcccctccatctgggcagcagagaaactgaggcacagagacagactgt   38178
                                                                   ********* **************************************************

chimp.LOC112206744.LOC107973052.GGT2-.646279-708040.rev.compl      gtccttacaggccacacagcctgccaggcccctatgtccggccagagcccctggtcagcc   39550
FAM230B-LOC105372935-GGT2.NCBI.GGT2.rev.compl                      gtccttacagggcacacagcctgccaggcccctatgtccggccagagcccctggtcagcc   38238
                                                                   *********** ************************************************

chimp.LOC112206744.LOC107973052.GGT2-.646279-708040.rev.compl      tgggctgcagtgattgtttagaggtaggctgttcccacggctgcctctcacggtaggggg   39610
FAM230B-LOC105372935-GGT2.NCBI.GGT2.rev.compl                      tgggctgcagtgattgtttagaggtaggctgttcccacggctgcctctcacggtaggggg   38298
                                                                   ************************************************************

chimp.LOC112206744.LOC107973052.GGT2-.646279-708040.rev.compl      gcctgcggacgcctcctcccgtccccacccgactcccaagcctcagtgacattgctcaac   39670
FAM230B-LOC105372935-GGT2.NCBI.GGT2.rev.compl                      gcctgcggacgcctcctcccgcccccacccgactcccaagcctcagtgacattgctcaac   38358
                                                                   ********************* **************************************

chimp.LOC112206744.LOC107973052.GGT2-.646279-708040.rev.compl      caggagctgaagtgcattcctgggctcaggccagcccacccacccacccgctgcagtcct   39730
FAM230B-LOC105372935-GGT2.NCBI.GGT2.rev.compl                      caggaactgaagtgcattcctgggctcaggccagcccacccacccacccgctgcagtcct   38418
                                                                   ***** ******************************************************
```

```
chimp.LOC112206744.LOC107973052.GGT2-.646279-708040.rev.compl      ggaagcccagaggcctgggcagcaggaacagtggagacagcagtgtgggggacgtccccc   39790
FAM230B-LOC105372935-GGT2.NCBI.GGT2.rev.compl                      ggaagcccagaggcctgggcagcaggaacagtggagacagcagtgtgggggacgtccccc   38478
                                                                   ************************************************************

chimp.LOC112206744.LOC107973052.GGT2-.646279-708040.rev.compl      ctcctctccccaccatcctcgtcaggcagaggccagggtgcagggaccacccgagcaaag   39850
FAM230B-LOC105372935-GGT2.NCBI.GGT2.rev.compl                      ctcctctccccaccatcctcgtcaggcagaggccagggtgcagggaccaccggagcaaag   38538
                                                                   *************************************************** ********

chimp.LOC112206744.LOC107973052.GGT2-.646279-708040.rev.compl      gcccagggaaatgaatgggtgtcattttggtcctgacccgaggcacagccaggaaggtcc   39910
FAM230B-LOC105372935-GGT2.NCBI.GGT2.rev.compl                      gcccagggaaatgaatgggtgtcattctggtcctgacccgaggcacagccaggaaggtcc   38598
                                                                   ************************** *********************************

chimp.LOC112206744.LOC107973052.GGT2-.646279-708040.rev.compl      ctgtggggaaaagaaacagatatcagactgttactgtgtctatgtagaaagaagtagagg   39970
FAM230B-LOC105372935-GGT2.NCBI.GGT2.rev.compl                      ctgtggggaaaagaaagagatatcagactgttactgtgtctatgtagaaagaagtagacg   38658
                                                                   **************** ***************************************** *

chimp.LOC112206744.LOC107973052.GGT2-.646279-708040.rev.compl      taagaggctccatttgttgtgtagtaagaaaaattcttttgccttgagatactgttaat   40030
FAM230B-LOC105372935-GGT2.NCBI.GGT2.rev.compl                      taagaggctccatttgttgtgtagtaagaaaaattcttttgccttgagatgccgttaat   38718
                                                                   *************************************************** * ******

chimp.LOC112206744.LOC107973052.GGT2-.646279-708040.rev.compl      ctgtaaccctagccccaaccctgtgctcacagaaacatgtactgtgtcgactcaaggttt   40090
FAM230B-LOC105372935-GGT2.NCBI.GGT2.rev.compl                      ctgtaaccctagccccaaccctgtgctcacagaaacatgtactgtgtcgactcaaggttt   38778
                                                                   ************************************************************

chimp.LOC112206744.LOC107973052.GGT2-.646279-708040.rev.compl      aatggatttagggctgtgcaggatgtgctttgttaaacaaatgcttgaaggcagcatgct   40150
FAM230B-LOC105372935-GGT2.NCBI.GGT2.rev.compl                      aatggattcagggctgtgcaggatgtgctttgttaaacaaatgcttgaaggcagcatgct   38838
                                                                   ******** ***************************************************

chimp.LOC112206744.LOC107973052.GGT2-.646279-708040.rev.compl      tgttaaaagtcatcaccactccctaatctcaagtaagcagggacacaaaacactgcagaa   40210
FAM230B-LOC105372935-GGT2.NCBI.GGT2.rev.compl                      tgttaagagtcatcaccactccctaatctcaagtaagcagggacacaaaacactgcagaa   38898
                                                                   ****** *****************************************************

chimp.LOC112206744.LOC107973052.GGT2-.646279-708040.rev.compl      ggccgcagggacctctgcctaggaaagccaggtattgtccaaggtttctccccatgtgac   40270
FAM230B-LOC105372935-GGT2.NCBI.GGT2.rev.compl                      ggccgcagggacctctgcctaggaaagccaggtatcgtccaaggtttctccccaggtgac   38958
                                                                   *********************************** ****************** *****

chimp.LOC112206744.LOC107973052.GGT2-.646279-708040.rev.compl      agtctgaaatatggcctcgtgggaagggaaagacctgaccgtcccccagcccgacacctg   40330
FAM230B-LOC105372935-GGT2.NCBI.GGT2.rev.compl                      agtctgaaatatggcctcgtgggaagggaaagacctgaccgtcccccagcccgacacctg   39018
                                                                   ************************************************************

chimp.LOC112206744.LOC107973052.GGT2-.646279-708040.rev.compl      taaagggtctgtgctgaggaggattagtaaaggggaaggcctctttgcagttgagataa   40390
FAM230B-LOC105372935-GGT2.NCBI.GGT2.rev.compl                      taaagggtctgtgctgaggaggattagtaaaagaggaaggcctctttgcagttgagataa   39078
                                                                   ********************************* **************************

chimp.LOC112206744.LOC107973052.GGT2-.646279-708040.rev.compl      gaggaaggcatctctctcctgatcgtccctgggcaaaggaatgtctcagtgttgatggta   40450
FAM230B-LOC105372935-GGT2.NCBI.GGT2.rev.compl                      gaggaaggcatctctctcctgatcgtccctgggcaaaggaatgtctcagtgttgattgta   39138
                                                                   ******************************************************** ***

chimp.LOC112206744.LOC107973052.GGT2-.646279-708040.rev.compl      tattccatctactgagataggagaaaactgccttagggctggaggtgggacatgctggtg   40510
FAM230B-LOC105372935-GGT2.NCBI.GGT2.rev.compl                      tattccatctgctgagataggagaaaactgccttagggctggagatgggacatgctggtg   39198
                                                                   ********** ********************************** ***************

chimp.LOC112206744.LOC107973052.GGT2-.646279-708040.rev.compl      gcaatactgctatttaatgcattgagatgtttatgtatatgcacatcaaagcacagcacc   40570
FAM230B-LOC105372935-GGT2.NCBI.GGT2.rev.compl                      gcaatactgctctttaatgcattgagatgtttatgtatatgcacatcaaagcatagcacc   39258
                                                                   *********** ***************************************** ******

chimp.LOC112206744.LOC107973052.GGT2-.646279-708040.rev.compl      tttttcttaaccttgtttatgacacagagacatttgttcacgtgttttcctgctgaccct   40630
FAM230B-LOC105372935-GGT2.NCBI.GGT2.rev.compl                      tttttcttaaccttgtttatgacacagagacatttgttcacgtgttttcctgctgaccct   39318
                                                                   ************************************************************

chimp.LOC112206744.LOC107973052.GGT2-.646279-708040.rev.compl      ctccccactattaccctattgtcctgccacatcccccctctctgagatggtagagataatg   40690
FAM230B-LOC105372935-GGT2.NCBI.GGT2.rev.compl                      ctccccactattaccctattgtcctgccacatcccccctctctgagatggtagagataatg   39378
                                                                   ************************************************************

chimp.LOC112206744.LOC107973052.GGT2-.646279-708040.rev.compl      atcaataaatactagggaactcagagactggtgccagcgtgggtcctccatatgctgagt   40750
FAM230B-LOC105372935-GGT2.NCBI.GGT2.rev.compl                      atcaataaatactagggaactcagagactggtgccagcgtggggcctccgtatgctgagc   39438
                                                                   ******************************************* ***** ********* 

chimp.LOC112206744.LOC107973052.GGT2-.646279-708040.rev.compl      gcaggtcccctgggcccactttctttctctatactttgtctctgtgtctctttcttttc   40810
FAM230B-LOC105372935-GGT2.NCBI.GGT2.rev.compl                      gcaggtcccctgggcccactttctttctctatactttgtctctgtgtctctttcttttc   39498
                                                                   ************************************************************

chimp.LOC112206744.LOC107973052.GGT2-.646279-708040.rev.compl      tcagtctcttgtcccacctgatgagaaacacccacagatgtggaggggcaggccacccct   40870
FAM230B-LOC105372935-GGT2.NCBI.GGT2.rev.compl                      tcagtctcttgtcccagctgatgggaaacacccacaggtgtggaggggcaggccacccct   39558
                                                                   **************** ****** ************* **********************

chimp.LOC112206744.LOC107973052.GGT2-.646279-708040.rev.compl      tcaggtccctgaatgtccttcctcaggaaatgatgggggaaggggtgatgagaatgaagg   40930
FAM230B-LOC105372935-GGT2.NCBI.GGT2.rev.compl                      tcaggtccctgaatgtccttcctcaggaaatgatgggggaaggggtgatgagaatgaagg   39618
                                                                   ************************************************************

chimp.LOC112206744.LOC107973052.GGT2-.646279-708040.rev.compl      agaggatttaagtccctcacccccgaggtagtcctgggctgagccccatgggacctgga   40990
FAM230B-LOC105372935-GGT2.NCBI.GGT2.rev.compl                      agacgatttaagtccctcacccccgaggtagtcctgggctgagccccatgggacctaga   39678
                                                                   *** ***************************************************** **

chimp.LOC112206744.LOC107973052.GGT2-.646279-708040.rev.compl      gaaccagggtgtaccccaccagcgtgtcgggtccaggaagcctcgtggccagctcccact   41050
FAM230B-LOC105372935-GGT2.NCBI.GGT2.rev.compl                      gaaccagggtgtaccccaccagcgtgtcgggtccaggaagcctcgtggccagctcccact   39738
                                                                   ************************************************************

chimp.LOC112206744.LOC107973052.GGT2-.646279-708040.rev.compl      tctcttgctgctgtgcaacccagagcaaggcctgcccctccagcttcagtcttctcccct   41110
FAM230B-LOC105372935-GGT2.NCBI.GGT2.rev.compl                      tctcttcctgctgtgcaacccagagcaaggcctgcccctccagcttcagtcttctcccct   39798
                                                                   ****** *****************************************************

chimp.LOC112206744.LOC107973052.GGT2-.646279-708040.rev.compl      gcaaatggggccacggcctttcctctcaggccaaaataaggattgaggccgggtgcagtg   41170
FAM230B-LOC105372935-GGT2.NCBI.GGT2.rev.compl                      gcaaatggggccacggcctttcctctcaggccaatataaggattgaggccgggtgcagtg   39858
                                                                   ********************************** *************************

chimp.LOC112206744.LOC107973052.GGT2-.646279-708040.rev.compl      gctcacccctgtaatcctagcactttgggagactgagatgggggactgcttgaagtcag   41230
FAM230B-LOC105372935-GGT2.NCBI.GGT2.rev.compl                      gct-acccctgtaatcctagcactttgggagactgagatgggggactgcttgaagtcag   39917
                                                                   *** ********************************************************

chimp.LOC112206744.LOC107973052.GGT2-.646279-708040.rev.compl      gagttaagaccagcctggtcaacatagtgagaccccatctctattggtttaaattttgtt   41290
FAM230B-LOC105372935-GGT2.NCBI.GGT2.rev.compl                      gagttaagaccagcctggtcaacatagtgagaccccatctctattggtttaaattttttt   39977
                                                                   ********************************************************* **

chimp.LOC112206744.LOC107973052.GGT2-.646279-708040.rev.compl      t--tttaattaaataaataaaataaggattgaagagtgacttgtacaccagttgagccca   41348
FAM230B-LOC105372935-GGT2.NCBI.GGT2.rev.compl                      taaaaaaattaaataaataaaataaggattgaagagtgacttgtacaccagttgagccca   40037
                                                                   *     ******************************************************

chimp.LOC112206744.LOC107973052.GGT2-.646279-708040.rev.compl      cctccatctcacccttgcagagccccagagacacagccctccagggctcagacccagtgg   41408
FAM230B-LOC105372935-GGT2.NCBI.GGT2.rev.compl                      cctccatctcacccttgcagagccccagagacacagccctccagagctcagacccagtgg   40097
                                                                   ******************************************* ****************

chimp.LOC112206744.LOC107973052.GGT2-.646279-708040.rev.compl      gacttgactccacaggcataaaaccctgtttgtctatgggccctttggaatcaccaggtt   41468
FAM230B-LOC105372935-GGT2.NCBI.GGT2.rev.compl                      gacttgactccacaggcataaaaccctgtttgtctatgggccctttggaatcaccaggtt   40157
                                                                   ************************************************************

chimp.LOC112206744.LOC107973052.GGT2-.646279-708040.rev.compl      ttcggggatcctgaaggatagccccgacctggcctcacctggcccctggccccagtgccc   41528
FAM230B-LOC105372935-GGT2.NCBI.GGT2.rev.compl                      ttcggggctcctgaaggatagccccgacctggcctcacctggcccctggccccagtgccc   40217
                                                                   ******* ****************************************************

chimp.LOC112206744.LOC107973052.GGT2-.646279-708040.rev.compl      ctggtgatatccaggtgc------------------------------------------   41546
FAM230B-LOC105372935-GGT2.NCBI.GGT2.rev.compl                      ctggtgatatccaggtgctgggctgtgatcaccgcctcccaccagcccacctccaccagc   40277
                                                                   ******************

chimp.LOC112206744.LOC107973052.GGT2-.646279-708040.rev.compl      --------------------------tgggctgtgatctccgcctcccaccagcccacct   41580
FAM230B-LOC105372935-GGT2.NCBI.GGT2.rev.compl                      ccttcccagaaccctgctccaggtgttgggctgtgatcaccgcctcccaccagcccacct   40337
                                                                                             ************ *********************

chimp.LOC112206744.LOC107973052.GGT2-.646279-708040.rev.compl      ccaccagcccttcccagaaccctgccccaggtgttggaactgtgcacagaggagggagca   41640
FAM230B-LOC105372935-GGT2.NCBI.GGT2.rev.compl                      ccaccagcccttcccagaaccctgctccaggtgttggaactgtgcacagaggagggagca   40397
                                                                   ************************* **********************************

chimp.LOC112206744.LOC107973052.GGT2-.646279-708040.rev.compl      ggccctgagggaggcctggaggggctgcggatggtgaaggctgctgtgtctagctgtttc   41700
FAM230B-LOC105372935-GGT2.NCBI.GGT2.rev.compl                      ggccccgagggaggcctggaggggctgccaatggtgaaggctgctgtgtcgagctgtttc   40457
                                                                   ***** **********************  ******************** *********

chimp.LOC112206744.LOC107973052.GGT2-.646279-708040.rev.compl      cttctggacccactccctctgggctgcgtccccggctggtccaagccctgatccctggga   41760
FAM230B-LOC105372935-GGT2.NCBI.GGT2.rev.compl                      cttccggacccactccctctgggctgcgtccccggctggtccaagccctgatccctggga   40517
                                                                   **** *******************************************************

chimp.LOC112206744.LOC107973052.GGT2-.646279-708040.rev.compl      tctggggacatcttcccgtttgttgttccctgagaaccaggcctccctctggagaggatc   41820
FAM230B-LOC105372935-GGT2.NCBI.GGT2.rev.compl                      tctggggacatcttcccgtttgctgttccctgagaaccaggcctccctctggagaggatc   40577
                                                                   ********************** *************************************
```

```
chimp.LOC112206744.LOC107973052.GGT2-.646279-708040.rev.compl      acaagcttgggtttcactctgggcttgctcttgggaacccccaggggcgtggctctgac   41880
FAM230B-LOC105372935-GGT2.NCBI.GGT2.rev.compl                      acaagcttgggtttcactctgggcttgctcttgggaacccccaggggcatggctctgac   40637
                                                                   ************************************************ ***********

chimp.LOC112206744.LOC107973052.GGT2-.646279-708040.rev.compl      cgagatgttttcctccagcctgttgcctagtccccattcctcgga-cctcagcttcacct   41939
FAM230B-LOC105372935-GGT2.NCBI.GGT2.rev.compl                      cgagatgttttcctccagcctgttgcccagtccccattcctcggaccctcagcttcacct   40697
                                                                   ************************** ****************** **************

chimp.LOC112206744.LOC107973052.GGT2-.646279-708040.rev.compl      ccagtgtcatcggcagggtgagctggacgcctacgggtctgagaaggcacccgggttccc   41999
FAM230B-LOC105372935-GGT2.NCBI.GGT2.rev.compl                      ccagtgtcatcggcagggtgagctggacgcctacgggtctgagaaggcgcccgggttccc   40757
                                                                   ************************************************ ***********

chimp.LOC112206744.LOC107973052.GGT2-.646279-708040.rev.compl      agcatcagctggccacc-------------------------------------------   42016
FAM230B-LOC105372935-GGT2.NCBI.GGT2.rev.compl                      agcatcggctggccaccctctgcctaagaaagcgccagggtcgtgacacccctggtggc   40817
                                                                   ****** **********

chimp.LOC112206744.LOC107973052.GGT2-.646279-708040.rev.compl      ------------------------------------------------------------   42016
FAM230B-LOC105372935-GGT2.NCBI.GGT2.rev.compl                      tgatcctaggtagtgtcactgcccagccccagtaagggagggcctggccccaaagtctga   40877

chimp.LOC112206744.LOC107973052.GGT2-.646279-708040.rev.compl      ------------------------------------------------------------   42016
FAM230B-LOC105372935-GGT2.NCBI.GGT2.rev.compl                      gggatcagggtgggaaggggcagggtttggtgtgaaccttcccctggcccccagccatgt   40937

chimp.LOC112206744.LOC107973052.GGT2-.646279-708040.rev.compl      ------------------------------------------------------------   42016
FAM230B-LOC105372935-GGT2.NCBI.GGT2.rev.compl                      gccttgctctccccatgctgaagatgctgaggctagttccagtgtccgcattgtgaagat   40997

chimp.LOC112206744.LOC107973052.GGT2-.646279-708040.rev.compl      ---------------ctctgttcctccccagccagatggctccatttcacacacaataca   42061
FAM230B-LOC105372935-GGT2.NCBI.GGT2.rev.compl                      ctccgaatcccacctctctgttcctccccagccagatggctccatttcacacacaataca   41057
                                                                                  *********************************************

chimp.LOC112206744.LOC107973052.GGT2-.646279-708040.rev.compl      ctgaggcccagagagtggggagacaggccagggaggccacctggagcctggcacagtggc   42121
FAM230B-LOC105372935-GGT2.NCBI.GGT2.rev.compl                      ctgaggcccagagagtggggagacaggccagggaggccacctggagcctggcacagtggc   41117
                                                                   ************************************************************

chimp.LOC112206744.LOC107973052.GGT2-.646279-708040.rev.compl      ctcatttattatgctgctctgctgctcacaggggaagcccgtcccccaaagtcctcttcc   42181
FAM230B-LOC105372935-GGT2.NCBI.GGT2.rev.compl                      ctcatttattatgctgctctgctgctcacaggggaagcccgtcccccaaagtcctcttcc   41177
                                                                   ************************************************************

chimp.LOC112206744.LOC107973052.GGT2-.646279-708040.rev.compl      tcatcctggtgagtatcttgtccctggattgcttgtcagccttgtctgcctggagcaatc   42241
FAM230B-LOC105372935-GGT2.NCBI.GGT2.rev.compl                      tcatcctggtgagtatcttgtccctggattgcttgtcagccttgtctgcctggagcaatc   41237
                                                                   ************************************************************

chimp.LOC112206744.LOC107973052.GGT2-.646279-708040.rev.compl      agtagccagcaggttccccgcctttcctggagtccgaggcagctgcccagccaccagcca   42301
FAM230B-LOC105372935-GGT2.NCBI.GGT2.rev.compl                      agtagccagcaggttccccgcctttcctggagtccgaggcagctgcccagccaccagccg   41297
                                                                   ***********************************************************

chimp.LOC112206744.LOC107973052.GGT2-.646279-708040.rev.compl      tgcggacgatggcttgcaccacagcgatgaaggtggacgcgatctgggtgtgatggtgcc   42361
FAM230B-LOC105372935-GGT2.NCBI.GGT2.rev.compl                      tgcggacgatggcttgcaccacagcgatgaaggtggacgcgatctgggtgtgatggtgcc   41357
                                                                   ************************************************************

chimp.LOC112206744.LOC107973052.GGT2-.646279-708040.rev.compl      gggtctccagggctgcagtcactgcctgggggtgggaggagaggggaagcctgagcaggg   42421
FAM230B-LOC105372935-GGT2.NCBI.GGT2.rev.compl                      gggtctccagggctgcagtcactgcctgggggtgggaggagaggggaagcctgagcaggg   41417
                                                                   ************************************************************

chimp.LOC112206744.LOC107973052.GGT2-.646279-708040.rev.compl      ctccagatgccacctgaaccacgcctgtgtggtcacaggcctcagcccaggtggtgccat   42481
FAM230B-LOC105372935-GGT2.NCBI.GGT2.rev.compl                      ctccagatgccacctgaaccacacctgtgtggtcacaggcctcagcccaggtggtgccat   41477
                                                                   ********************** *************************************

chimp.LOC112206744.LOC107973052.GGT2-.646279-708040.rev.compl      ttcaggccaggtcatcaggaagagcaggttggggcctgctgggtctcactggagcagggg   42541
FAM230B-LOC105372935-GGT2.NCBI.GGT2.rev.compl                      ttcaggccaggtcatcaggaagagcaggttggggcctgctgggtctcactggagcagggg   41537
                                                                   ************************************************************

chimp.LOC112206744.LOC107973052.GGT2-.646279-708040.rev.compl      gcttggccctcatggcacaggggctccagatggcccaggcactagagagaggacaccaac   42601
FAM230B-LOC105372935-GGT2.NCBI.GGT2.rev.compl                      gcttggctctcatgtcacaggggctccagatggcccaggcactagagagaggacaccaac   41597
                                                                   ******* ****** *********************************************

chimp.LOC112206744.LOC107973052.GGT2-.646279-708040.rev.compl      cattgtccactctgtgatgatccaggcctccagcccaggatgccctggggccccacaccg   42661
FAM230B-LOC105372935-GGT2.NCBI.GGT2.rev.compl                      cattgtccactctgtgatgatccaggcctccagcccaggatgccctgggaccccacaccg   41657
                                                                   ************************************************* **********

chimp.LOC112206744.LOC107973052.GGT2-.646279-708040.rev.compl      tgactcagtttctccaacccccggcccacctggtcaatgtttctctccactgtcgtgacg   42721
FAM230B-LOC105372935-GGT2.NCBI.GGT2.rev.compl                      tgactcagtttctccaacccctggcccacctggtcaatgtttctctccactgtcgtgacg   41717
                                                                   ********************* **************************************

chimp.LOC112206744.LOC107973052.GGT2-.646279-708040.rev.compl      ttgggcagaagcttgttgtgcagccggggctcctccacggccttcttcacgtcatagccg   42781
FAM230B-LOC105372935-GGT2.NCBI.GGT2.rev.compl                      ttgggcagaagcttgttgtgcagccggggctcctccacggccctcttcacgtcatagccg   41777
                                                                   ****************************************** *****************

chimp.LOC112206744.LOC107973052.GGT2-.646279-708040.rev.compl      aaccagaggttgtagatgatggcctggggcatgggagtgtgatcagcgtggcttgggggc   42841
FAM230B-LOC105372935-GGT2.NCBI.GGT2.rev.compl                      aaccagaggttgtagatgatggcctggggcatgggagtgtgatcagggtggcttgggggc   41837
                                                                   ********************************************** *************

chimp.LOC112206744.LOC107973052.GGT2-.646279-708040.rev.compl      tgtgcagagtgggcagggccagggagaaaggggtgacacataccagtgcagtgtctgtg   42901
FAM230B-LOC105372935-GGT2.NCBI.GGT2.rev.compl                      tgtgcagagtgggcagggccagggagaaaggggtgacacataccagtgcagtgtctgtg   41897
                                                                   ************************************************************

chimp.LOC112206744.LOC107973052.GGT2-.646279-708040.rev.compl      gtgatctgcgtgcccccagcagctcccaccaccaaccggacctggccgtcctggcccacc   42961
FAM230B-LOC105372935-GGT2.NCBI.GGT2.rev.compl                      gtgatctgcgtgcccccagcagctcccaccaccatccggacctggccgtcctggcccacc   41957
                                                                   ********************************** *************************

chimp.LOC112206744.LOC107973052.GGT2-.646279-708040.rev.compl      atgatcgtcgggcacatggacgagagcggctgcttccctgcggctgatgggagaagacag   43021
FAM230B-LOC105372935-GGT2.NCBI.GGT2.rev.compl                      atgatcgtcaggcacatggacaagagcggctgcttccctgcggccgatgggagaagacag   42017
                                                                   ********* *********** ********************** ***************

chimp.LOC112206744.LOC107973052.GGT2-.646279-708040.rev.compl      ggatgcccgtcagctgcctgcccaggacacccgcccctctccaccccagtcccccca-ccc   43080
FAM230B-LOC105372935-GGT2.NCBI.GGT2.rev.compl                      ggatgcccgtcagctgcctgcccaggacacccgcccctctccaccccagtcccccaccccc   42077
                                                                   ******************************************************** ***

chimp.LOC112206744.LOC107973052.GGT2-.646279-708040.rev.compl      ccagacctccaccccatacctggctggatgaaattggcaggtgaggggggcaccccaaac   43140
FAM230B-LOC105372935-GGT2.NCBI.GGT2.rev.compl                      ccggacctccaccccatacctggctggatgaaattggcaggtgaggggggtgccccaaac   42137
                                                                   ** ***********************************************  ********

chimp.LOC112206744.LOC107973052.GGT2-.646279-708040.rev.compl      tcattggtg-atgctgggagagctgaagtcgtccattcattattgaacaggatcccactg   43199
FAM230B-LOC105372935-GGT2.NCBI.GGT2.rev.compl                      tcattggtgaatgctgggagagctgaagtcgtccattcattattgaacaggatcccactg   42197
                                                                   ********* **************************************************

chimp.LOC112206744.LOC107973052.GGT2-.646279-708040.rev.compl      accggggagcagaccttggagccaaagctaccgcccagccaggtcagacagcacctgacc   43259
FAM230B-LOC105372935-GGT2.NCBI.GGT2.rev.compl                      accggggagcagaccttggagccaaagctaccgcccagccaggtcagacagcacccgacc   42257
                                                                   ******************************************************* ****

chimp.LOC112206744.LOC107973052.GGT2-.646279-708040.rev.compl      ttgcctggcccagcctggtccctatccacccactgaggctcaaacatactcactgagagg   43319
FAM230B-LOC105372935-GGT2.NCBI.GGT2.rev.compl                      ttgcctggcccagcctggtccctatccacccactgaggctcaaacatactcactgagagg   42317
                                                                   ************************************************************

chimp.LOC112206744.LOC107973052.GGT2-.646279-708040.rev.compl      cccaggataagctaccaaggttgggcctcagtttcccaccaggaaaagaggtgatggagc   43379
FAM230B-LOC105372935-GGT2.NCBI.GGT2.rev.compl                      cccaggataagctaccaaggttgggcctcagtttcccaccaggaaaagaggtgatggagc   42377
                                                                   ************************************************************

chimp.LOC112206744.LOC107973052.GGT2-.646279-708040.rev.compl      caccttactggataagtgggcagtccctgggccacccgcccctggccctttcccacccag   43439
FAM230B-LOC105372935-GGT2.NCBI.GGT2.rev.compl                      caccttactggataagtgggcagtccctgggccacccgcccctggccctttcccacccag   42437
                                                                   ************************************************************

chimp.LOC112206744.LOC107973052.GGT2-.646279-708040.rev.compl      gcggcccagcagcccctactagaggttgatggtgctggtggccgacacagcactgccgtc   43499
FAM230B-LOC105372935-GGT2.NCBI.GGT2.rev.compl                      gcggcccagcagcccctactagaggttgatggtgctggtggcggacacagcactgccgtc   42497
                                                                   ****************************************** *****************

chimp.LOC112206744.LOC107973052.GGT2-.646279-708040.rev.compl      ctctgcgacgacagacaggtgagcagtgcccccgtcatccggcgtgtagaactcgggctt   43559
FAM230B-LOC105372935-GGT2.NCBI.GGT2.rev.compl                      ctctgcgacgacagacaggtgagcagtgcccccgtcatccggcgtgtagaactcgggctt   42557
                                                                   ************************************************************

chimp.LOC112206744.LOC107973052.GGT2-.646279-708040.rev.compl      gtagtaggagatcgggtgagtggtgtggtcagagatctgggcctggagctgggcagcgaa   43619
FAM230B-LOC105372935-GGT2.NCBI.GGT2.rev.compl                      gtagtaggagatcgggtgagtggtgtggtcagagatctgggaccggagctgggcagcgaa   42617
                                                                   ***************************************** * ****************

chimp.LOC112206744.LOC107973052.GGT2-.646279-708040.rev.compl      gaactcggaggtcatgttgcggaccacctgccgagaccccagagctggcctgaggaggtg   43679
FAM230B-LOC105372935-GGT2.NCBI.GGT2.rev.compl                      gaactcagaggtcatgttgcggaccacctgctgagaccccagagctggcctgaggaggtg   42677
                                                                   ****** ************************ ****************************

chimp.LOC112206744.LOC107973052.GGT2-.646279-708040.rev.compl      gggagggagggtggggagggggcacaggtctcagaaggcccctgactgtgactctgaccg   43739
FAM230B-LOC105372935-GGT2.NCBI.GGT2.rev.compl                      gggagggagggtggggagggggcacaggtctcagaaggcccttgactgtgactctgaccg   42737
                                                                   ***************************************** ******************

chimp.LOC112206744.LOC107973052.GGT2-.646279-708040.rev.compl      caaccctctggcacccacaaccttccgtggctccccaggacccaagggcaggcccaggac   43799
```

```
FAM230B-LOC105372935-GGT2.NCBI.GGT2.rev.compl                      caaccctctggcacccacaaccttccgtggctccccaggacccaagggcaggcccaagac   42797
                                                                   ******************************************************** ***

chimp.LOC112206744.LOC107973052.GGT2-.646279-708040.rev.compl      cttgcacaaccagtctgactccctgtctctgtcgcgttccagcaactctgaatgtctgtc   43859
FAM230B-LOC105372935-GGT2.NCBI.GGT2.rev.compl                      cttgcatgaccagtctgactccctgtctctgtcgcgttccagcaactctgaatgtctgtc   42857
                                                                   ******  ****************************************************

chimp.LOC112206744.LOC107973052.GGT2-.646279-708040.rev.compl      tgcgtggtcctcagcctccagacccttgccacattcaatcactcattctttcatgcaaaa   43919
FAM230B-LOC105372935-GGT2.NCBI.GGT2.rev.compl                      tgcctggtcctcagcctccagacccttgccgcattcaatcactcattccttcatgcaaaa   42917
                                                                   *** ************************** ***************** ***********

chimp.LOC112206744.LOC107973052.GGT2-.646279-708040.rev.compl      aatatttctagaatttgcactgcatgcctggcactggggaatcaacagggaacagacatt   43979
FAM230B-LOC105372935-GGT2.NCBI.GGT2.rev.compl                      aatatttctagaatttgcactgcatgcctggcactggggaatcaacagggaacagacact   42977
                                                                   ********************************************************** *

chimp.LOC112206744.LOC107973052.GGT2-.646279-708040.rev.compl      taggtcctgccctcatgccaagaaaaacaaacacacacagggaaagtgctgaaaccacaa   44039
FAM230B-LOC105372935-GGT2.NCBI.GGT2.rev.compl                      taagtcctgccctcatgccaagaaaaacaaacacacccagggaaagtgctgaaaccacag   43037
                                                                   ** ********************************* **********************

chimp.LOC112206744.LOC107973052.GGT2-.646279-708040.rev.compl      gccaggtaaggggaatcaagaggcatgaggtatgggcagagtggtcagggagggcttctc   44099
FAM230B-LOC105372935-GGT2.NCBI.GGT2.rev.compl                      gccaggtaaggggaatcaagaggcatgaggtatgggcagagtggtcagggagggcttctc   43097
                                                                   ************************************************************

chimp.LOC112206744.LOC107973052.GGT2-.646279-708040.rev.compl      agaggaggcagcgtgtgaaaagagcctggaatgtggcctaaatggtcagtgcaaaggccc   44159
FAM230B-LOC105372935-GGT2.NCBI.GGT2.rev.compl                      agaggaggcaacgtgtgaaaagagcctggaatgtggcctaaatggtcagtgcaaaggccc   43157
                                                                   ********** *************************************************

chimp.LOC112206744.LOC107973052.GGT2-.646279-708040.rev.compl      tgaggcaggtggtataggctggtgagcgataggcagagagtgaatggagtaggatgggga   44219
FAM230B-LOC105372935-GGT2.NCBI.GGT2.rev.compl                      tgaggcaggtggtataggctggtgagcgataggcagagagtgaatggagtagggtgggga   43217
                                                                   ***************************************************** ******

chimp.LOC112206744.LOC107973052.GGT2-.646279-708040.rev.compl      gaagaggatgaagatgcaggctggggcccatcccacaggacctcctaggtcccataacaa   44279
FAM230B-LOC105372935-GGT2.NCBI.GGT2.rev.compl                      gaagaggatgaagatgcaggctggggcccatcccacaggacctcctaggtcccataacaa   43277
                                                                   ************************************************************

chimp.LOC112206744.LOC107973052.GGT2-.646279-708040.rev.compl      ctggcttttgctctgtgccatgcaggcttagggcagaggaatgaggagggtggggagtgt   44339
FAM230B-LOC105372935-GGT2.NCBI.GGT2.rev.compl                      ctggcttttctctgtgccatgcaggcttagggcagaggaatgaggaggctggggagtgt   43337
                                                                   ********* *************************************** **********

chimp.LOC112206744.LOC107973052.GGT2-.646279-708040.rev.compl      tttcacagggtccctctggcagctatgacggggataaggataaagcccaaaggggaggct   44399
FAM230B-LOC105372935-GGT2.NCBI.GGT2.rev.compl                      tttcacagggtccctctggcagctatgacggggataaggataaagcccaaaggggaggct   43397
                                                                   ************************************************************

chimp.LOC112206744.LOC107973052.GGT2-.646279-708040.rev.compl      gtgggtatcaaccaggcaagagatgatggcctgggtgggagaaagagaagaatcaaggat   44459
FAM230B-LOC105372935-GGT2.NCBI.GGT2.rev.compl                      gtgggtatcaaccaggcaagagatgatggcctgggtgggagaaagagaagaatcaaggat   43457
                                                                   ************************************************************

chimp.LOC112206744.LOC107973052.GGT2-.646279-708040.rev.compl      ggtgccgactagcgaggtaaaccctgcagaaggggcaggtttggggatggtcaggagctt   44519
FAM230B-LOC105372935-GGT2.NCBI.GGT2.rev.compl                      ggtgccgactagcgagg-----ccggcagaaggggccggtttggggatggtcaggagctt   43512
                                                                   *****************     ** *********** ***********************

chimp.LOC112206744.LOC107973052.GGT2-.646279-708040.rev.compl      gattttggatacttcatcagacctgaagagcatgggtgcacgtataaaaaaaataaataa   44579
FAM230B-LOC105372935-GGT2.NCBI.GGT2.rev.compl                      gattttggatacttcatcagacccaaagagcatgggtgcacgtataaaaaaaataaataa   43572
                                                                   ***********************  ***********************************

chimp.LOC112206744.LOC107973052.GGT2-.646279-708040.rev.compl      ataagcatgggttcacgggcaagggcgggctgagagatgaacatggaggtattgacattg   44639
FAM230B-LOC105372935-GGT2.NCBI.GGT2.rev.compl                      ataagcatgggttcacgggcaagggcgggctgagagatgaacatggaggtattgacattg   43632
                                                                   ************************************************************

chimp.LOC112206744.LOC107973052.GGT2-.646279-708040.rev.compl      agtggctgctggatgccatgagcctggccaaggtccccaaggcagtggcgaggaggagat   44699
FAM230B-LOC105372935-GGT2.NCBI.GGT2.rev.compl                      agtggctgctggatgccatgagcctggccaaggtccccaaggcagtggcgaggaggagat   43692
                                                                   ************************************************************

chimp.LOC112206744.LOC107973052.GGT2-.646279-708040.rev.compl      gaggaggtcaaggaggagac--agaggatggacccgaaggccaaagaaaatgcctcaaga   44757
FAM230B-LOC105372935-GGT2.NCBI.GGT2.rev.compl                      gaggaggtcaaagaggagacagagaggatggacccgaaggccgaagaaaatgcctcaaga   43752
                                                                   *********** ********  ******************** *****************

chimp.LOC112206744.LOC107973052.GGT2-.646279-708040.rev.compl      gagtttcaacaccgggcgcggtggctcacgcctgtaatcccagcactttgggaggccgag   44817
FAM230B-LOC105372935-GGT2.NCBI.GGT2.rev.compl                      gagtttcaacaccgggcgcggtggctcacgcctgtaatcccagcactttgggaggccgag   43812
                                                                   ************************************************************

chimp.LOC112206744.LOC107973052.GGT2-.646279-708040.rev.compl      gcctgtaatcccagcactttgggaggcggatcacgacgtcaggagatcaagaccatcct   44877
FAM230B-LOC105372935-GGT2.NCBI.GGT2.rev.compl                      gcctgtaatcccagcactttgggaggcggatcatgacgtcaggagatcgagaccatcct   43872
                                                                   ********************************* ************** **********

chimp.LOC112206744.LOC107973052.GGT2-.646279-708040.rev.compl      ggctaacacagtgaaaccccgtctctactaaaaatacaaaaaattagctgggcgcggtgg   44937
FAM230B-LOC105372935-GGT2.NCBI.GGT2.rev.compl                      ggctaacacagtgaaaccccgtctctactaaaaatacaaaaaattagctgggcgcggtgg   43932
                                                                   ************************************************************

chimp.LOC112206744.LOC107973052.GGT2-.646279-708040.rev.compl      tgggcacctgtagtcccagctacttgggggggctgaggcaggagaatggcatgaacctgg   44997
FAM230B-LOC105372935-GGT2.NCBI.GGT2.rev.compl                      tgggcacctgtagtcccagctactt-gggggggctgaggcaggagaatggcgtgaacctgg   43991
                                                                   ************************* ************************ *********

chimp.LOC112206744.LOC107973052.GGT2-.646279-708040.rev.compl      gaggcggagcttgcagtgagccgagatcccgccactgcactccagcctgggggacagcct   45057
FAM230B-LOC105372935-GGT2.NCBI.GGT2.rev.compl                      gaggcggagcttgcagtgagcggagatcctgccactgcactccagcctgggggacagcct   44051
                                                                   ********************* ******* ******************************

chimp.LOC112206744.LOC107973052.GGT2-.646279-708040.rev.compl      gggggacagagcgagactccgtctcaaaaataaaataaaataaaataaaataaagtttca   45117
FAM230B-LOC105372935-GGT2.NCBI.GGT2.rev.compl                      gggggacagagcgagactccgtctcaaaaataaaataaaataaaataaaataaagtttca   44111
                                                                   ************************************************************

chimp.LOC112206744.LOC107973052.GGT2-.646279-708040.rev.compl      gcaacaccccacggattagttgaccaatcccagggaaaggtgttctgtctgaatctgccc   45177
FAM230B-LOC105372935-GGT2.NCBI.GGT2.rev.compl                      gcaacaccccacggattagttgaccaatcccagggaaaggtgttctgtctgaatctgccc   44171
                                                                   ************************************************************

chimp.LOC112206744.LOC107973052.GGT2-.646279-708040.rev.compl      tcaaggaaacagaaaggcaaatccacgatgtgggacattttccaagactactgccctggg   45237
FAM230B-LOC105372935-GGT2.NCBI.GGT2.rev.compl                      tcaaggaaacagaaaggcaaatccacgatgtgggacattttccaagactactgccctggg   44231
                                                                   ************************************************************

chimp.LOC112206744.LOC107973052.GGT2-.646279-708040.rev.compl      ctttaaaaatcaacaaaacaggccaggcacggtggctcatgcctgtaatcccagcacttt   45297
FAM230B-LOC105372935-GGT2.NCBI.GGT2.rev.compl                      ctttaaaaatcaacaaaacaggccaggcacggtggttcatgcctgtaatcccagcacttt   44291
                                                                   *********************************** ************************

chimp.LOC112206744.LOC107973052.GGT2-.646279-708040.rev.compl      gggaggccgaggcaggcggatcacgaggtcaggagatcgcgatcacggtgaaaccccgtc   45357
FAM230B-LOC105372935-GGT2.NCBI.GGT2.rev.compl                      gggaggccgaggcaggcggatcacgaggtcaggagatcgcgatcacggtgaaaccccgtc   44351
                                                                   ************************************************************

chimp.LOC112206744.LOC107973052.GGT2-.646279-708040.rev.compl      tctactaaaaatacaaaaaattagctgggcgcagtggcgggtgcctgtagtcccagctac   45417
FAM230B-LOC105372935-GGT2.NCBI.GGT2.rev.compl                      tctactaaaaatacaaaaaattagctgggcgcagtgtcgggcgcctgtagtcccagctac   44411
                                                                   ************************************ **** ******************

chimp.LOC112206744.LOC107973052.GGT2-.646279-708040.rev.compl      tcgggaggctgaggcaggagaatggcgtgaacccaggaggcggagcttgcagtgagccga   45477
FAM230B-LOC105372935-GGT2.NCBI.GGT2.rev.compl                      tcgggaggctgaggcaggagaatggcgtgaacccaggaggcggagcttgcagtgagccga   44471
                                                                   ************************************************************

chimp.LOC112206744.LOC107973052.GGT2-.646279-708040.rev.compl      gatagcaccactgcactccagcctgggcgacagagcgacaccctgtctcacaaaaaagcc   45537
FAM230B-LOC105372935-GGT2.NCBI.GGT2.rev.compl                      gatagcgccactgcactccagcctgggcaacagagcgagactccgtctcaaagaaaaaac   44531
                                                                   ****** ********************* ********* ** * ****** * ****  *

chimp.LOC112206744.LOC107973052.GGT2-.646279-708040.rev.compl      acacctgtaatcccagcgctttgggaggctgaggtgggaagatagtttgatcccaggagt   45597
FAM230B-LOC105372935-GGT2.NCBI.GGT2.rev.compl                      acacctgtaatcccagcgctttgggaggctgaggtgggaagacagtttgatcccaggagt   44591
                                                                   ****************************************** *****************

chimp.LOC112206744.LOC107973052.GGT2-.646279-708040.rev.compl      ttgagaccagtctgggcaagaccctgtctctaaaaaaatatataaaaattatccaggttt   45657
FAM230B-LOC105372935-GGT2.NCBI.GGT2.rev.compl                      ttgagaccagtctgggcaagaccctgtctctaaaaaaa-atacaaaaattatccaggttt   44650
                                                                   ************************************** *** *****************

chimp.LOC112206744.LOC107973052.GGT2-.646279-708040.rev.compl      ggtggcacgtgcctctggtcccagctgctcaggggggctgaagtgggaggattgcttgagc   45717
FAM230B-LOC105372935-GGT2.NCBI.GGT2.rev.compl                      ggtggcacgtgcctctggtcccagctgctcaggggggctgaagtgggaggattgcttgagc   44710
                                                                   ************************************************************

chimp.LOC112206744.LOC107973052.GGT2-.646279-708040.rev.compl      cctggaggttgaggctgcagtgagccaagatcacaccactgcactccaacctggatgaca   45777
FAM230B-LOC105372935-GGT2.NCBI.GGT2.rev.compl                      cctggaggttgaggctgcagtgagccaagatcacaccactgcactccaacctggatgaca   44770
                                                                   ************************************************************

chimp.LOC112206744.LOC107973052.GGT2-.646279-708040.rev.compl      gagactctgtgtccaaacaaaaacacaacaaaaacaaaacagccaagagagcctactgta   45837
FAM230B-LOC105372935-GGT2.NCBI.GGT2.rev.compl                      gagactctgtgtacaaacaaaaacacaacaaaaacaaaacagccaaggagcctactgta   44830
                                                                   ************ *********************************** ***********

chimp.LOC112206744.LOC107973052.GGT2-.646279-708040.rev.compl      ggtaaagagaagggacaacagcaggttgtgtcaccccgacatcctgctgtgctatgttca   45897
FAM230B-LOC105372935-GGT2.NCBI.GGT2.rev.compl                      ggtaaagagaaggg---acagcaggttgtgtcaccccgacatcctgctgtgctatgttca   44887
                                                                   **************   *******************************************

chimp.LOC112206744.LOC107973052.GGT2-.646279-708040.rev.compl      agtctcacttttgagacacaccctgaggtgcgacatcagtaacctactgtggaatccctc   45957
FAM230B-LOC105372935-GGT2.NCBI.GGT2.rev.compl                      agtctcacttttgagacacaccctgaggtgcgacatcagtaacctactgtggaatccctc   44947
```

```
                                                                   ************************************************************

chimp.LOC112206744.LOC107973052.GGT2-.646279-708040.rev.compl      agaaaaacacgaatcccaatagacgtgggtggagacggagagagttaggaaatccggcag   46017
FAM230B-LOC105372935-GGT2.NCBI.GGT2.rev.compl                      agaaaaacacgaatcccaatagatgtgggtggagacggagagagttaggaaatccggcag   45007
                                                                   *********************** ************************************

chimp.LOC112206744.LOC107973052.GGT2-.646279-708040.rev.compl      aaacgtccacactgcagactccaggaaaagggaacattgatgcttgggcagttttggggt   46077
FAM230B-LOC105372935-GGT2.NCBI.GGT2.rev.compl                      aaatgtccacactgcagaatccaggaaaagggaacattgatgcttgggcagttttgggtt   45067
                                                                   *** ************** *************************************** *

chimp.LOC112206744.LOC107973052.GGT2-.646279-708040.rev.compl      atttttacatttttgtaggtgtgaaaatttgcaaaatgaaaactcgag--gagtgtggtg   46135
FAM230B-LOC105372935-GGT2.NCBI.GGT2.rev.compl                      ttttttacatttttgtaggtgcgaaaatttgcaaaatgaaaactcgaggagagtgtggtg   45127
                                                                    ******************** **************************  **********

chimp.LOC112206744.LOC107973052.GGT2-.646279-708040.rev.compl      agctgtgtgagatgctgctgagtggggcctgatggggaaactgaggctggacatggcgat   46195
FAM230B-LOC105372935-GGT2.NCBI.GGT2.rev.compl                      agctgtgtgagatgctgctgagtggggcctgatggggaaactgaggctggacatggcgat   45187
                                                                   ************************************************************

chimp.LOC112206744.LOC107973052.GGT2-.646279-708040.rev.compl      ctggtggcatggggatagagcaggggaggggataccctgaagggagagaggacatagccc   46255
FAM230B-LOC105372935-GGT2.NCBI.GGT2.rev.compl                      ctggtggcatggggatagagcaggggaggggataccctgaagggagagaggacatagccc   45247
                                                                   ************************************************************

chimp.LOC112206744.LOC107973052.GGT2-.646279-708040.rev.compl      agccatgtttgctctaagaggagcaaaggacagaaggaggcagcagatagaaatttctag   46315
FAM230B-LOC105372935-GGT2.NCBI.GGT2.rev.compl                      agccatgtttgctctaagaggagcaaaggacagaaggaggcagcagatagaagtttctag   45307
                                                                   **************************************************** *******

chimp.LOC112206744.LOC107973052.GGT2-.646279-708040.rev.compl      agcaacaacagctgcctttcttttgggaataatccgtgataaagaagtaaatcgtcagag   46375
FAM230B-LOC105372935-GGT2.NCBI.GGT2.rev.compl                      agcaacaacagctgcctttctttcgggaataatccgtgataaagaaataaatcatcagag   45367
                                                                   *********************** ********************** ****** ******

chimp.LOC112206744.LOC107973052.GGT2-.646279-708040.rev.compl      gcggaaaggagcattgtaggaacagcaccccgagccagcgagtagatgaaggagctggcc   46435
FAM230B-LOC105372935-GGT2.NCBI.GGT2.rev.compl                      gcagacaggagcattgtaggaacagcaccccgagccagtgagtagatgaaggagctggcc   45427
                                                                   ** ** ******************************** *********************

chimp.LOC112206744.LOC107973052.GGT2-.646279-708040.rev.compl      ttagccaaaagggagggcagagggacacgctgcagtggctctgtcccctcagagagacaa   46495
FAM230B-LOC105372935-GGT2.NCBI.GGT2.rev.compl                      ttagccaaaaaggagggcagagggacacgctgcagtggctctgtcccctcagagagacaa   45487
                                                                   ********** *************************************************

chimp.LOC112206744.LOC107973052.GGT2-.646279-708040.rev.compl      gacaccaggtcactggctgcagcgggagtcagaggtgcagaatgctcacggggaaagaga   46555
FAM230B-LOC105372935-GGT2.NCBI.GGT2.rev.compl                      gacaccaggtcactggctgcagcgggagtcagaggtgcagaatgctcacagggaaagaga   45547
                                                                   ************************************************* **********

chimp.LOC112206744.LOC107973052.GGT2-.646279-708040.rev.compl      agacaccacccgggcagctgacgcccctcctgggaggtcactggtcagtgtggggggtc   46615
FAM230B-LOC105372935-GGT2.NCBI.GGT2.rev.compl                      agacaccacccgggcagctgacgcccctcctgggaggtcactggtcagtgtggggggtc   45607
                                                                   ************************************************************

chimp.LOC112206744.LOC107973052.GGT2-.646279-708040.rev.compl      tgcagatccgcccaggaatgccaaggacccaagtaggtaaggagggatgtgaggatcctc   46675
FAM230B-LOC105372935-GGT2.NCBI.GGT2.rev.compl                      tgcagatccgcccaggaatgccaaggacccaagtaggtaaggagggatgtgaggatcctc   45667
                                                                   ************************************************************

chimp.LOC112206744.LOC107973052.GGT2-.646279-708040.rev.compl      agcaggaagggatatgacagggtcttatagggacccagcgtggagctggggcaactactg   46735
FAM230B-LOC105372935-GGT2.NCBI.GGT2.rev.compl                      agtgggaagggatatgacagggtcttatagggacccagcatggagctggggcaactactg   45727
                                                                   **  ********************************* **********************

chimp.LOC112206744.LOC107973052.GGT2-.646279-708040.rev.compl      agtgagtgggtcaggtggtgcatggctgagggtggcatctgggaagcgttagtttggagt   46795
FAM230B-LOC105372935-GGT2.NCBI.GGT2.rev.compl                      ggtgagtgggtcaggtggtgcagggctgagggtggcatctgggaagcattagtttggagt   45787
                                                                    ********************* ************************ ************

chimp.LOC112206744.LOC107973052.GGT2-.646279-708040.rev.compl      gacagggagtgggtggccaaggtctctgttcacctggccctctccctttcacccatccat   46855
FAM230B-LOC105372935-GGT2.NCBI.GGT2.rev.compl                      gacagggagtgggtggccgaggtctctgttcacctggccctctccctttcacccatccat   45847
                                                                   ****************** *****************************************

chimp.LOC112206744.LOC107973052.GGT2-.646279-708040.rev.compl      ctcaccatgcccgagggctgcaatgcccccatgcattcctcatctcagcactgagcactc   46915
FAM230B-LOC105372935-GGT2.NCBI.GGT2.rev.compl                      ctcaccatgcccgagggctgcaatgcccccatgcattcctcatctcagcactgagcactc   45907
                                                                   ************************************************************

chimp.LOC112206744.LOC107973052.GGT2-.646279-708040.rev.compl      gagaaccacagcatgtgcaaaggacctgaggtaggagggtggccactaattccccacact   46975
FAM230B-LOC105372935-GGT2.NCBI.GGT2.rev.compl                      aagaaccacagcatgtgcaaaggacctgaggtaggagggtggccactaattccccgcact   45967
                                                                    ****************************************************** ****

chimp.LOC112206744.LOC107973052.GGT2-.646279-708040.rev.compl      gtcagttctgtgcggcagaagccaagactggggtcacccacgagtccatccgagcacac   47035
FAM230B-LOC105372935-GGT2.NCBI.GGT2.rev.compl                      gtcagttctgtggggcagaagccaagactggggtcacccacgagtccatccgagcacac   46027
                                                                   ************ ***********************************************

chimp.LOC112206744.LOC107973052.GGT2-.646279-708040.rev.compl      agtaggcccgcaatcaaagtctgtggcaggaatggattcataaagcatatgtgaggctgt   47095
FAM230B-LOC105372935-GGT2.NCBI.GGT2.rev.compl                      agtaggcccgcaatcgaattctgtggcaggaatggattcataaagcatatgtgaggctgt   46087
                                                                   *************** ** *****************************************

chimp.LOC112206744.LOC107973052.GGT2-.646279-708040.rev.compl      agccaccctggggagcccacctgaggcctctacagcaggccccacacccacagtgggcca   47155
FAM230B-LOC105372935-GGT2.NCBI.GGT2.rev.compl                      agccaccctggggagcccacctgatgcctctacggcaggccccacacccacagtgagcca   46147
                                                                   ************************ ******** ********************** ****

chimp.LOC112206744.LOC107973052.GGT2-.646279-708040.rev.compl      gcccctgccccttaccccagtcacatccacaaacttggggtccccaagcagggtcctctt   47215
FAM230B-LOC105372935-GGT2.NCBI.GGT2.rev.compl                      gcccctgccccttacctcagtcacatccacaaacttggggtccccaagcagggtcctctt   46207
                                                                   **************** *******************************************

chimp.LOC112206744.LOC107973052.GGT2-.646279-708040.rev.compl      ggcgtaggcaaaccggaaaggcctctacgatgtggtggtacgtcaggcccttctgcttgg   47275
FAM230B-LOC105372935-GGT2.NCBI.GGT2.rev.compl                      ggcgtaggcaaaccggaaa-gcctctacgatgcggtggtacgtcaggcccttctgctcgg   46266
                                                                   ******************* ************ ************************ **

chimp.LOC112206744.LOC107973052.GGT2-.646279-708040.rev.compl      gggtctccacgctctcctgggagaagttgtaccctggttgatcagaaccaggtgcatgtt   47335
FAM230B-LOC105372935-GGT2.NCBI.GGT2.rev.compl                      gggtctccacgctctcccgggagaagttgtaccctggttgatcagagccaggtgcatgtt   46326
                                                                   ***************** *************************** **************

chimp.LOC112206744.LOC107973052.GGT2-.646279-708040.rev.compl      gctgagccccagaggctctgaggggctcagagggttaacacctgcctgagccactttgcc   47395
FAM230B-LOC105372935-GGT2.NCBI.GGT2.rev.compl                      gctgagccccagaggctctgaggggctcagagggttaacacctgcctgagccactttgcc   46386
                                                                   ************************************************************

chimp.LOC112206744.LOC107973052.GGT2-.646279-708040.rev.compl      cacctcaaggagcgtttaataaccaatagcagcagctgcttcagaaggctgtggtgagag   47455
FAM230B-LOC105372935-GGT2.NCBI.GGT2.rev.compl                      cacctcaaggagcgtttaataaccaatagcagcagctgcttcagaaggctgtggtgagag   46446
                                                                   ************************************************************

chimp.LOC112206744.LOC107973052.GGT2-.646279-708040.rev.compl      tgaagtaaggtgagggctcccgagcctgggcctcacgtcatgcatcagctctggccattc   47515
FAM230B-LOC105372935-GGT2.NCBI.GGT2.rev.compl                      tgaagtaaggtgagggctcccgagcctggacctcacgtcacgcatcagctctggccattc   46506
                                                                   ***************************** ********** *******************

chimp.LOC112206744.LOC107973052.GGT2-.646279-708040.rev.compl      agtgaccaagtggcagggtcacccactggaccagggtaccctctggaccagccctgcagc   47575
FAM230B-LOC105372935-GGT2.NCBI.GGT2.rev.compl                      aatgaccaagtggcagggtcacccactggaccggggtaccctctggaccggccctgcagc   46566
                                                                   * ****************************** **************** **********

chimp.LOC112206744.LOC107973052.GGT2-.646279-708040.rev.compl      cctgagctcctgcacctctctccctcctgatgactcctgttcctcctccaacccttgagc   47635
FAM230B-LOC105372935-GGT2.NCBI.GGT2.rev.compl                      cctgagctcctgcacctctctccctcctgatgactcctgttcctcctccaacccttgagc   46626
                                                                   ************************************************************

chimp.LOC112206744.LOC107973052.GGT2-.646279-708040.rev.compl      attgcccgcttcagccctgtcgctctgcactgcctccttcaggacatggtgagctgtgat   47695
FAM230B-LOC105372935-GGT2.NCBI.GGT2.rev.compl                      attgcccgcttcagccctgttgctctgcactgcctccttcaggacatggtgagctgtgac   46686
                                                                   ******************** **************************************

chimp.LOC112206744.LOC107973052.GGT2-.646279-708040.rev.compl      gcagggacacgcctcgggagctcagtgatggaaagacgtggcatgggggcgagcagaga   47755
FAM230B-LOC105372935-GGT2.NCBI.GGT2.rev.compl                      gcagggacacacctcaggagctcagtgatggaaagacgtggcatgggggcgagcagaga   46746
                                                                   ********** **** ********************************************

chimp.LOC112206744.LOC107973052.GGT2-.646279-708040.rev.compl      tgcagcaggggtggggcgtggggagagagaggcagtgtcatgggtcctaccacacggctg   47815
FAM230B-LOC105372935-GGT2.NCBI.GGT2.rev.compl                      tgcaggaggggtggggcgtggggagagagaggcagtgtcatgggtcctaccacacggctg   46806
                                                                   ***** ******************************************************

chimp.LOC112206744.LOC107973052.GGT2-.646279-708040.rev.compl      tggtgcgaccacccacctttgaggatgttgaggatgagggccagcacgggcccgctgagc   47875
FAM230B-LOC105372935-GGT2.NCBI.GGT2.rev.compl                      tggtgcgaccactcacctttgaggatgttgaggatgagggccagcacgggcccgctgagc   46866
                                                                   ************ ***********************************************

chimp.LOC112206744.LOC107973052.GGT2-.646279-708040.rev.compl      ggcgcactgggcatgtacagcaccgcgtctcccaggctgatgttcagcgggtgctcgatc   47935
FAM230B-LOC105372935-GGT2.NCBI.GGT2.rev.compl                      cgcgcactgggcatgtacagcaccgcgtctcccaggctgatgttcagcgggtgctcgatc   46926
                                                                    ***********************************************************

chimp.LOC112206744.LOC107973052.GGT2-.646279-708040.rev.compl      agctcagcacggtagttgttcaggtcctcagctgtcacaatgcccccctgcaatgggacag   47995
FAM230B-LOC105372935-GGT2.NCBI.GGT2.rev.compl                      agctcagcacggtagttgttcaggtcctcagctgtcacaatgcccccctgcaatgggacag   46986
                                                                   ************************************************************

chimp.LOC112206744.LOC107973052.GGT2-.646279-708040.rev.compl      cagctcgaatgggcgctgggatggggctgcaccactgcgtggaggatggagctgcaccag   48055
FAM230B-LOC105372935-GGT2.NCBI.GGT2.rev.compl                      cagctcgaatgggtgctgggatggggctgcaccactgcgtggaggatggaactgcaccag   47046
                                                                   ************* ************************************ *********

chimp.LOC112206744.LOC107973052.GGT2-.646279-708040.rev.compl      tggtgttgggggcaggcatggctgcaccatggtggtggggaaaagcctgtacctaccagg   48115
FAM230B-LOC105372935-GGT2.NCBI.GGT2.rev.compl                      tggtgttgggggcaggcatggctgcaccatggtggtggggaaaagcctgtacctaccagg   47106
                                                                   ************************************************************
```

```
chimp.LOC112206744.LOC107973052.GGT2-.646279-708040.rev.compl      gaggacagagtgcactactggaggggtgggactgtgccctgggagggggccacaggcaac   48175
FAM230B-LOC105372935-GGT2.NCBI.GGT2.rev.compl                      gaggacagagtgcactactggaggggtgggactgtgccctgggagggggccacaggcaac   47166
                                                                   ************************************************************

chimp.LOC112206744.LOC107973052.GGT2-.646279-708040.rev.compl      ctcacctccttgggaacctcaccagctccggcactcctgtctccctgacactgctcacca   48235
FAM230B-LOC105372935-GGT2.NCBI.GGT2.rev.compl                      ctcacctccttgggaacctcaccagctccggcactcctgtctccctgacactgctcacca   47226
                                                                   ************************************************************

chimp.LOC112206744.LOC107973052.GGT2-.646279-708040.rev.compl      cccgacagctgggctggggccacctgccctctgcctgcttggcttactggcttcctgtct   48295
FAM230B-LOC105372935-GGT2.NCBI.GGT2.rev.compl                      cccgacagctgggctggggccacctgccctctgcctgcttggcttactggcttcctgtct   47286
                                                                   ************************************************************

chimp.LOC112206744.LOC107973052.GGT2-.646279-708040.rev.compl      gccttctctcatctgtggccggagagtgtttcctttttctttttttttttttagagatac   48355
FAM230B-LOC105372935-GGT2.NCBI.GGT2.rev.compl                      gccttctctcatctgtggccagagagtgttttctttttctttttttttttttagagatac   47346
                                                                   ******************** ********** ****************************

chimp.LOC112206744.LOC107973052.GGT2-.646279-708040.rev.compl      gatcttgctctgttacccaggctggagtgcagtggcttaatcacagccttgaactcccgg   48415
FAM230B-LOC105372935-GGT2.NCBI.GGT2.rev.compl                      gatcttgctctgttacccaggctggagtgcagtggcttaatcacagccttgaacccctgg   47406
                                                                   ****************************************************** ** **

chimp.LOC112206744.LOC107973052.GGT2-.646279-708040.rev.compl      gctcaagtgatcctccagcagaccctccccagtagctgagactaaaggcacaactacacc   48475
FAM230B-LOC105372935-GGT2.NCBI.GGT2.rev.compl                      gctcaagtgatcctccagcagaccctccccagtagctgagactaaaggcacaactacacc   47466
                                                                   ************************************************************

chimp.LOC112206744.LOC107973052.GGT2-.646279-708040.rev.compl      cagcaaatttaattttttttgttgtgttttgctatatttctttctttctttctttctttt   48535
FAM230B-LOC105372935-GGT2.NCBI.GGT2.rev.compl                      cagcaaatttaattttttttgttgtgttttgctatatttctttctttcttt-tttttttt   47525
                                                                   *************************************************** *** ****

chimp.LOC112206744.LOC107973052.GGT2-.646279-708040.rev.compl      ttttttgtgagacggagtctcactctatcgcccaggctggagtgcagtggcgcgatatcg   48595
FAM230B-LOC105372935-GGT2.NCBI.GGT2.rev.compl                      ttttttgtgagacggagtctcgctctatcgcccaggctggagtgcagtggcgcgatatcg   47585
                                                                   ********************* **************************************

chimp.LOC112206744.LOC107973052.GGT2-.646279-708040.rev.compl      gctcactgcaagctccgcctcccaggttcacgccattctcctgcctcagcctcccaagta   48655
FAM230B-LOC105372935-GGT2.NCBI.GGT2.rev.compl                      gctcactgcaagctccgcctcccaggttcacgccattctcctgcctcagcctcccaagta   47645
                                                                   ************************************************************

chimp.LOC112206744.LOC107973052.GGT2-.646279-708040.rev.compl      gctgggactgcaggcacccgccaccacacctggctaattttttttaaattttagtagaga   48715
FAM230B-LOC105372935-GGT2.NCBI.GGT2.rev.compl                      gctgggactgcaggcacccgccaccacatgtggctaattttttctgtattttagtagaga   47705
                                                                   ****************************  ************* *  *************

chimp.LOC112206744.LOC107973052.GGT2-.646279-708040.rev.compl      caggtttcactgtgttagccaggatggtctcgatctcctgaccctgtgatccacacgcct   48775
FAM230B-LOC105372935-GGT2.NCBI.GGT2.rev.compl                      caggtttcactgtgttagccaggatggtctcgatctcctgaccccgtgatccacacgcct   47765
                                                                   ******************************************** ***************

chimp.LOC112206744.LOC107973052.GGT2-.646279-708040.rev.compl      cggcctctcaaagtgctgggattacaggcgtgaggcaccatgcccggccgggttttgcta   48835
FAM230B-LOC105372935-GGT2.NCBI.GGT2.rev.compl                      cggcctctcaaagtgctgagattacaggcgtgaggcaccatgcccggccgggttttgcta   47825
                                                                   ***************** ******************************************

chimp.LOC112206744.LOC107973052.GGT2-.646279-708040.rev.compl      tatttcttttcactatgctttgaattttttgttttcttgttccccacccccacccccact   48895
FAM230B-LOC105372935-GGT2.NCBI.GGT2.rev.compl                      tatttcttttcactatgctttgaattttttgttttcttgttccccacccccacccccact   47885
                                                                   ************************************************************

chimp.LOC112206744.LOC107973052.GGT2-.646279-708040.rev.compl      atatttatgtagattctcaattttttttttgtagactcactatgttgcccaggcttgtct   48955
FAM230B-LOC105372935-GGT2.NCBI.GGT2.rev.compl                      atatttatgtagattctcaatattttttttgtagactcactatgttgcccaggcttgtct   47945
                                                                   ********************* **************************************

chimp.LOC112206744.LOC107973052.GGT2-.646279-708040.rev.compl      tggacctcctgacctcaacttctacctcagcctcccaaagtgttgggattacaggcatga   49015
FAM230B-LOC105372935-GGT2.NCBI.GGT2.rev.compl                      tggaccccctgacctcaacttctacctcagcctcccaaagtgttgggattacaggcatga   48005
                                                                   ****** *****************************************************

chimp.LOC112206744.LOC107973052.GGT2-.646279-708040.rev.compl      gacaccaggctttgcctcagacggccttttctttctttttttttttttgagatggagtct   49075
FAM230B-LOC105372935-GGT2.NCBI.GGT2.rev.compl                      gacaccaagcttggcctcagagggccttttc--tctcttttttttttttgagatggagtct   48063
                                                                   ******* **** ******** *********  *** ***********************

chimp.LOC112206744.LOC107973052.GGT2-.646279-708040.rev.compl      cactctgtggagtgcagtggtgcaatctcggctcactgcaacctcagccccccaggttct   49135
FAM230B-LOC105372935-GGT2.NCBI.GGT2.rev.compl                      cactctgtggagtgcagtggtgcaatctcggctcactgcaacctcagccccacaggttct   48123
                                                                   *************************************************** ********

chimp.LOC112206744.LOC107973052.GGT2-.646279-708040.rev.compl      agcgattctcctgccacagcctcccacgtagctgggattacaggcacaacccaccatgcc   49195
FAM230B-LOC105372935-GGT2.NCBI.GGT2.rev.compl                      agcgattctcctgccacagcctcccaagtagctgggattacaggcacaacccaccatgcc   48183
                                                                   ************************** *********************************

chimp.LOC112206744.LOC107973052.GGT2-.646279-708040.rev.compl      tgactaattttgcatttttagtagagacagggtttcaccatgttggccaggctggtcttg   49255
FAM230B-LOC105372935-GGT2.NCBI.GGT2.rev.compl                      tgactaattttgcatttttagtagagacagggtttcaccatgttggccaggctggtcttg   48243
                                                                   ************************************************************

chimp.LOC112206744.LOC107973052.GGT2-.646279-708040.rev.compl      aactcctgagctcaggtgttccacccgcctcggcctcccaaagtgctgggattacagacg   49315
FAM230B-LOC105372935-GGT2.NCBI.GGT2.rev.compl                      aactcctgacctcaggtgatccacccgcctcggcctcccaaagtgctgggattacagacg   48303
                                                                   ********* ******** *****************************************

chimp.LOC112206744.LOC107973052.GGT2-.646279-708040.rev.compl      tgagccactgcgcctggctcagagggccttttctaactggagaattcctgccggtgtccc   49375
FAM230B-LOC105372935-GGT2.NCBI.GGT2.rev.compl                      tgagccaccgcacctggctcagagggccttttctaactggagaattcctgccggtgtccc   48363
                                                                   ******** ** ************************************************

chimp.LOC112206744.LOC107973052.GGT2-.646279-708040.rev.compl      tgctgcttggcctcttctcctcatgaggaatggatagagggagggagggaggctcttaat   49435
FAM230B-LOC105372935-GGT2.NCBI.GGT2.rev.compl                      tgctgcttggcctcttctcctcacgatgaatggatagagggagggagggaggctcttaat   48423
                                                                   *********************** ** *********************************

chimp.LOC112206744.LOC107973052.GGT2-.646279-708040.rev.compl      tctcctggagtcagctccagacagacagggtattggcatgccaatttccagcctcagtgg   49495
FAM230B-LOC105372935-GGT2.NCBI.GGT2.rev.compl                      tctcctggagtcagctc----cagacagggtattggcatgccaatttccagcctcagtgg   48479
                                                                   *****************    ***************************************

chimp.LOC112206744.LOC107973052.GGT2-.646279-708040.rev.compl      taaaggtcgacacgctaatcaccctcctccatgaagcagtgacaaaaattacctgaagaa   49555
FAM230B-LOC105372935-GGT2.NCBI.GGT2.rev.compl                      taaaggtcgacacgctaatcaccctcctccatgaaacagtgacaaaaattacctgaagaa   48539
                                                                   *********************************** ************************

chimp.LOC112206744.LOC107973052.GGT2-.646279-708040.rev.compl      agccacagccaagctccaggcccctgccccacaaatccccttccccatgcctctctcaag   49615
FAM230B-LOC105372935-GGT2.NCBI.GGT2.rev.compl                      agccacagccaagctccaggcccctgccccacaaatccccttccccatgcctctctcaag   48599
                                                                   ************************************************************

chimp.LOC112206744.LOC107973052.GGT2-.646279-708040.rev.compl      gcgaccctcatcccttgtaaccctcttggtgaatcaaagcgccctctactgggccttagc   49675
FAM230B-LOC105372935-GGT2.NCBI.GGT2.rev.compl                      gcgaccctcatcccttgtaaccctcttggtgaatcaaagcgccctctactgggccttagc   48659
                                                                   ************************************************************

chimp.LOC112206744.LOC107973052.GGT2-.646279-708040.rev.compl      ccagctgttcctctgctaggaatcccttcctctctctgcctaacgaagttatctgcagcc   49735
FAM230B-LOC105372935-GGT2.NCBI.GGT2.rev.compl                      ccagctgttcctctgctaggaatcccttcctctctctgcctaacgaagttatctgcagcc   48719
                                                                   ************************************************************

chimp.LOC112206744.LOC107973052.GGT2-.646279-708040.rev.compl      cagccgccacctcctccaagaagtcctcctggatcttcaggctgtattctagtgcttccc   49795
FAM230B-LOC105372935-GGT2.NCBI.GGT2.rev.compl                      cagccgccacctcctccaagaagtcctcctggatcttcaggctgtattctagtgcttccc   48779
                                                                   ************************************************************

chimp.LOC112206744.LOC107973052.GGT2-.646279-708040.rev.compl      tagccctggctcttgcctacacctgcatttaccccaacagggacttgctctcctggactg   49855
FAM230B-LOC105372935-GGT2.NCBI.GGT2.rev.compl                      tagccctggctcttgcctacacctgcatttaccccaacagggacttgctctcctggactg   48839
                                                                   ************************************************************

chimp.LOC112206744.LOC107973052.GGT2-.646279-708040.rev.compl      tgtggcctctcttggttttgatataagcaggagctgtggacccacatggccagtcactga   49915
FAM230B-LOC105372935-GGT2.NCBI.GGT2.rev.compl                      tgtggcctctcttggttttgatataagcaggagctgtggacccacatggccagtcactga   48899
                                                                   ************************************************************

chimp.LOC112206744.LOC107973052.GGT2-.646279-708040.rev.compl      ccctcctccaccaggaacttcctgcaggctcaggcaagacaggaggaccccatggctctg   49975
FAM230B-LOC105372935-GGT2.NCBI.GGT2.rev.compl                      ccctcctccaccaggaacttcctgcaggctcaggcaagacaggaggaccccatggctctg   48959
                                                                   ************************************************************

chimp.LOC112206744.LOC107973052.GGT2-.646279-708040.rev.compl      ggctacagctcagggtttccactgcagagttcctcacccaggtccttgaggttacccact   50035
FAM230B-LOC105372935-GGT2.NCBI.GGT2.rev.compl                      ggctacagctcagggtttccactgcagagttcctcacccaggtccttgaggttacccact   49019
                                                                   ************************************************************

chimp.LOC112206744.LOC107973052.GGT2-.646279-708040.rev.compl      caccagccgcctggatgtccttcacaatctgggccatgaggctgccgttgtagaaggcct   50095
FAM230B-LOC105372935-GGT2.NCBI.GGT2.rev.compl                      caccagccgcctggatgtccttcacaatctgggccatgaggctgccgttgtagaaggcct   49079
                                                                   ************************************************************

chimp.LOC112206744.LOC107973052.GGT2-.646279-708040.rev.compl      gggcgccctcgatggccagcatctcgtaggtgtcagccagccgcggcagggtcagtctct   50155
FAM230B-LOC105372935-GGT2.NCBI.GGT2.rev.compl                      gggcaccctcgatggccagcatctcataggtgtcagccagccgcggcagggtcagtctct   49139
                                                                   **** ******************** **********************************

chimp.LOC112206744.LOC107973052.GGT2-.646279-708040.rev.compl      ccccctcccgaagcacctttctatcccggcagaacacctcactggggcagagggggctca   50215
FAM230B-LOC105372935-GGT2.NCBI.GGT2.rev.compl                      ccccctcccgaagcacctttctatcccggcagaacacctcaccggggcagagggggctca   49199
                                                                   ***************************************** ******************

chimp.LOC112206744.LOC107973052.GGT2-.646279-708040.rev.compl      tgtgaggcagcaggtggggtggactcagctagaccacccccca cacccgtccatacagga   50275
FAM230B-LOC105372935-GGT2.NCBI.GGT2.rev.compl                      tgtgaggcagcaggtggggtggactcagctagaccaccccccacacctgcccacacagga   49259
                                                                   *********************************************** * *** ******
```

```
chimp.LOC112206744.LOC107973052.GGT2-.646279-708040.rev.compl      gaccagcacaaagcaggggcagcgcctgtcacaggtgggtggccctgtcactcagcgctc   50335
FAM230B-LOC105372935-GGT2.NCBI.GGT2.rev.compl                      gaccagcacaaagcaggggcagcacctgtcacaggtgggtggccctgtcactcagcgctc   49319
                                                                   *********************** ************************************

chimp.LOC112206744.LOC107973052.GGT2-.646279-708040.rev.compl      atcctcctagtgtcccttccgggaacctcctagtgtcccttgccactcaggacacatggc   50395
FAM230B-LOC105372935-GGT2.NCBI.GGT2.rev.compl                      atcctcctagtgtcccttccgggagcctcctagtgtcccttgccactcaggacacgtggc   49379
                                                                   ************************ ****************************** ****

chimp.LOC112206744.LOC107973052.GGT2-.646279-708040.rev.compl      cagccacagtggccactgggaccccacgctcagaatgtgtccccacacgtggtgggaagg   50455
FAM230B-LOC105372935-GGT2.NCBI.GGT2.rev.compl                      cagccacagtggccactgggaccccatgctcagaatgtgtccccacacgtggtgggaagg   49439
                                                                   ************************** *********************************

chimp.LOC112206744.LOC107973052.GGT2-.646279-708040.rev.compl      gtctgtatctcctcatcccattatcagcacagggtcctgaaggcagagggccactccact   50515
FAM230B-LOC105372935-GGT2.NCBI.GGT2.rev.compl                      gtctgtatctcctcatcccattatcagcacagggtcctgaaggcagagggccgctccact   49499
                                                                   **************************************************** *******

chimp.LOC112206744.LOC107973052.GGT2-.646279-708040.rev.compl      gctgctacggcctgcaaggtccttgggctgtgcctgccctgcctgtgtcaggggggccgca   50575
FAM230B-LOC105372935-GGT2.NCBI.GGT2.rev.compl                      gctgctacggcctgcaaggtccttgggctgtgcctgccctgcctgtgtcaggggggccgca   49559
                                                                   ************************************************************

chimp.LOC112206744.LOC107973052.GGT2-.646279-708040.rev.compl      cccacagacatgccacaagacaggctgctgctcgatgacggtccgcttgttttccagggc   50635
FAM230B-LOC105372935-GGT2.NCBI.GGT2.rev.compl                      cccacagacataccacaagacaggctgctgctcgatgacagtccgcttgttttccaggac   49619
                                                                   *********** *************************** ****************** *

chimp.LOC112206744.LOC107973052.GGT2-.646279-708040.rev.compl      tgccgccaagcccttgcccacagggaagccctggcgggccagctggatgctgggctggaa   50695
FAM230B-LOC105372935-GGT2.NCBI.GGT2.rev.compl                      tgccgccaagcccttgcccacagggaagccctggcgggccagctggatgctgggctggaa   49679
                                                                   ************************************************************

chimp.LOC112206744.LOC107973052.GGT2-.646279-708040.rev.compl      gaggcgagcccagggcagctgcccatgccactgatgtgccagctcatagcctcggatctc   50755
FAM230B-LOC105372935-GGT2.NCBI.GGT2.rev.compl                      gaggcgagcccagggcagccgcccatgccgctggtgtgccagctcatagcctcggatctc   49739
                                                                   ******************* ********* *** **************************

chimp.LOC112206744.LOC107973052.GGT2-.646279-708040.rev.compl      cccaggcaccgccaccgacagccctcctggggagagagagccacagttagtgaccctgag   50815
FAM230B-LOC105372935-GGT2.NCBI.GGT2.rev.compl                      cccaggcaccgccaccgacagccctcctggggagagagagccacagttagtgaccctgag   49799
                                                                   ************************************************************

chimp.LOC112206744.LOC107973052.GGT2-.646279-708040.rev.compl      tgggggacatccggatctctcgcaggcagcatcccaggcacagtccctgactcgctttac   50875
FAM230B-LOC105372935-GGT2.NCBI.GGT2.rev.compl                      tgggggacatcgggatctctagcaggcagcatcccaggcacagtccctgactcattttac   49859
                                                                   *********** ******** ********************************  *****

chimp.LOC112206744.LOC107973052.GGT2-.646279-708040.rev.compl      agatggggcaatgaggcttaggaggaaagatttttttttctttttgagttggggtcttg   50935
FAM230B-LOC105372935-GGT2.NCBI.GGT2.rev.compl                      agatggggcaatgaggcttaggaggaaagatttttttttctttttgagttggggtcttg   49919
                                                                   ************************************************************

chimp.LOC112206744.LOC107973052.GGT2-.646279-708040.rev.compl      ccatcttgcccagcctgatctcgaactcctggattcaagcaatcctcccacctcagcctc   50995
FAM230B-LOC105372935-GGT2.NCBI.GGT2.rev.compl                      ccatcttgcccagcctgatctcgaactcctggattcaagcaatcctcccacctcagcctc   49979
                                                                   ************************************************************

chimp.LOC112206744.LOC107973052.GGT2-.646279-708040.rev.compl      ctgagtagctgagattacaggcgtgaacaccacacccagcagaaggggatttttaatttt   51055
FAM230B-LOC105372935-GGT2.NCBI.GGT2.rev.compl                      ctgagtcgctgagattacaggcgtgaacagcacacctagcagaaggggatttttaatttt   50039
                                                                   ****** ********************** ****** ***********************

chimp.LOC112206744.LOC107973052.GGT2-.646279-708040.rev.compl      ttaatttattttttaattttaattttttttttttaggaggggatgtttaa--tttttttt   51113
FAM230B-LOC105372935-GGT2.NCBI.GGT2.rev.compl                      taaatttatttttaattttaatttttttttttttaggaggggatgtttaatttttttttt   50099
                                                                   * *********** * *** * ****************************  ********

chimp.LOC112206744.LOC107973052.GGT2-.646279-708040.rev.compl      taggaggggctcagca-gtaggagtgtacatggacaagggatgtctgaggagggcacagg   51172
FAM230B-LOC105372935-GGT2.NCBI.GGT2.rev.compl                      taggaggggctcagcaggtaggagtgtacatggaccagggatgtctgaggagggcacagc   50159
                                                                   **************** ****************** ***********************

chimp.LOC112206744.LOC107973052.GGT2-.646279-708040.rev.compl      aggggaagcagtagcatgcggctgggttttgctgtcccaggatgaggtgtctgtctgtgc   51232
FAM230B-LOC105372935-GGT2.NCBI.GGT2.rev.compl                      aggggaagcagtagcatgcggctgggttttgctgtcccaggatgaggtgtctgtctgtac   50219
                                                                   ********************************************************** *

chimp.LOC112206744.LOC107973052.GGT2-.646279-708040.rev.compl      aggtgcctgcatgtctaaaatcctgtgccaggccagacccccctcccatctcgctgaccac   51292
FAM230B-LOC105372935-GGT2.NCBI.GGT2.rev.compl                      aggtgcctgcatgtctaaaatcctgtgccaggccagacccccctcccatctcgctgaccac   50279
                                                                   ************************************************************

chimp.LOC112206744.LOC107973052.GGT2-.646279-708040.rev.compl      aaggccttatcctgtaagactcatgggctccaccagaatgtgccaaaacaagagcagatc   51352
FAM230B-LOC105372935-GGT2.NCBI.GGT2.rev.compl                      aaggccttatcctgtaagactcatgggctccaccagaatgtgccaaaacaagagcagatc   50339
                                                                   ************************************************************

chimp.LOC112206744.LOC107973052.GGT2-.646279-708040.rev.compl      ccaccctgacccaggtcaagcacaggccaccttcaagacacagccagccccaagaaaggg   51412
FAM230B-LOC105372935-GGT2.NCBI.GGT2.rev.compl                      ccaccctgacccaggtcaagcacaggccaccttcaagacacagccagccccaagaaaggg   50399
                                                                   ************************************************************

chimp.LOC112206744.LOC107973052.GGT2-.646279-708040.rev.compl      ctcccttcctcttttctactgccccagagaggcaagactgagccttaacctccgtcctgt   51472
FAM230B-LOC105372935-GGT2.NCBI.GGT2.rev.compl                      ctcccttcctcttttctactgccccagagaggcaagactgagccttaacctccatcctgt   50459
                                                                   ***************************************************** ******

chimp.LOC112206744.LOC107973052.GGT2-.646279-708040.rev.compl      cccctctcccagcctcagtttctccatccaactataagggtt----tttgtttgtctgtt   51528
FAM230B-LOC105372935-GGT2.NCBI.GGT2.rev.compl                      cccctctcccagcctcagtttctccaaccaactataagggtttttgtttgtttgtctgtt   50519
                                                                   ************************** ***************    **************

chimp.LOC112206744.LOC107973052.GGT2-.646279-708040.rev.compl      ttgagacagggtctcattctgttgtcccagctggaatgcagtggtgcaatcatggctcac   51588
FAM230B-LOC105372935-GGT2.NCBI.GGT2.rev.compl                      ttgagacagggtctcactctgttgtcccagctggagtgcagtggtgcaatcatggctcac   50579
                                                                   **************** ****************** ************************

chimp.LOC112206744.LOC107973052.GGT2-.646279-708040.rev.compl      tgcagccttggcttcccaggctcaagcgatcctcccacctcagcctctgaagtacctaag   51648
FAM230B-LOC105372935-GGT2.NCBI.GGT2.rev.compl                      tgcagccttggcttcccaggctcaagcgatcctcccacttcagcctctgaagtacctaag   50639
                                                                   ************************************** *********************

chimp.LOC112206744.LOC107973052.GGT2-.646279-708040.rev.compl      actacagacacaccccactacacgtggc--ttttttttttttttttttgagatggagtt   51706
FAM230B-LOC105372935-GGT2.NCBI.GGT2.rev.compl                      actacagacataccccactgcacgtggcttttttttttttttttttttgagatggagtt   50699
                                                                   ********** ******** ********  ******************************

chimp.LOC112206744.LOC107973052.GGT2-.646279-708040.rev.compl      tcactcttgttgcccaggctggagtgcaatggcacaatcttggctcactccaacctccac   51766
FAM230B-LOC105372935-GGT2.NCBI.GGT2.rev.compl                      tcactcttgttgcccaggctggagtgcaatggcacaatcttggctcactccaacctccac   50759
                                                                   ************************************************************

chimp.LOC112206744.LOC107973052.GGT2-.646279-708040.rev.compl      ctcccgagttcaagcaattctcctgcctcagcctcccaagtagctgggattacaggcatg   51826
FAM230B-LOC105372935-GGT2.NCBI.GGT2.rev.compl                      ctcccgagttcaagtaattctcctgcctcagcctcccaagtacctgggattacaggcatg   50819
                                                                   ************** *************************** *****************

chimp.LOC112206744.LOC107973052.GGT2-.646279-708040.rev.compl      caccaccacgccttgctaattttgtatttttagtggagacggggtttcaccaggttggt   51886
FAM230B-LOC105372935-GGT2.NCBI.GGT2.rev.compl                      taccaccacgccttgctaattttgtatttttagtagagacggggtttcaccaggttggt   50879
                                                                    ********************************* ************************

chimp.LOC112206744.LOC107973052.GGT2-.646279-708040.rev.compl      caggctggtcttgaactcctaacatcaggtgatccatcctcctcagcctctcagagttct   51946
FAM230B-LOC105372935-GGT2.NCBI.GGT2.rev.compl                      caggctggtcttgaactcctaacatcaggtgatccatcctcctcagcttctcagagttct   50939
                                                                   *********************************************** ************

chimp.LOC112206744.LOC107973052.GGT2-.646279-708040.rev.compl      gggattacaggcgtgagccaccactcccagcctaatttttatttttatttttgtagag   52006
FAM230B-LOC105372935-GGT2.NCBI.GGT2.rev.compl                      gggattacaggcgtgagccaccactcccagcctaatttttatttttatttttgtagag   50999
                                                                   ************************************************************

chimp.LOC112206744.LOC107973052.GGT2-.646279-708040.rev.compl      acaggggtcttgctacgttgccaagactggtctcaaactctggcctcaagcaatcctccc   52066
FAM230B-LOC105372935-GGT2.NCBI.GGT2.rev.compl                      acaggggtcttgctacattgccaagactggtctcaaactctggcctcaagcaatcctccc   51059
                                                                   **************** *******************************************

chimp.LOC112206744.LOC107973052.GGT2-.646279-708040.rev.compl      acctcagcctcccaacatgctgggattacaggtgcactcagcctataaggggttttgcct   52126
FAM230B-LOC105372935-GGT2.NCBI.GGT2.rev.compl                      acctcagcctcccaacatgctgggattacaggtgcactcagcctataaggggttttgcct   51119
                                                                   ************************************************************

chimp.LOC112206744.LOC107973052.GGT2-.646279-708040.rev.compl      tccagttctgacttttgaggaggtcattggaaacagacccctgggcctgctttctccctg   52186
FAM230B-LOC105372935-GGT2.NCBI.GGT2.rev.compl                      tccagttctgacttttgaggaggtcattggaaacagacccctgggcctgcttcccccctg   51179
                                                                   **************************************************** * *****

chimp.LOC112206744.LOC107973052.GGT2-.646279-708040.rev.compl      agcctcactgcccatatggacactacagacactgaccctttgcccagaaaggtacaacta   52246
FAM230B-LOC105372935-GGT2.NCBI.GGT2.rev.compl                      agccccactgcccatatggacactacagacactgaccctttgcccagaaaggtacaacta   51239
                                                                   **** *******************************************************

chimp.LOC112206744.LOC107973052.GGT2-.646279-708040.rev.compl      tggcctctgcccccagggactctcctgctcttgcgagagatgatggggccatttgccttg   52306
FAM230B-LOC105372935-GGT2.NCBI.GGT2.rev.compl                      tggcctctgcccccagggactctcctgctcttgcgagagatgatggggccatttgccttg   51299
                                                                   ************************************************************

chimp.LOC112206744.LOC107973052.GGT2-.646279-708040.rev.compl      gcttggcggctgtggctctagaactgcctctcccaccctgaagcctggcacaagtttcca   52366
FAM230B-LOC105372935-GGT2.NCBI.GGT2.rev.compl                      gcttggcggctgtggctctagaactgcctctcccaccctgaagcctggcacaagtttcca   51359
                                                                   ************************************************************

chimp.LOC112206744.LOC107973052.GGT2-.646279-708040.rev.compl      agagctggtggtttcaattcctagaagctgcacgtacatcccggaaggtctgacacccag   52426
FAM230B-LOC105372935-GGT2.NCBI.GGT2.rev.compl                      agagctggtggtttcaattcctagaagctgcacatatatcccggaaggtctgacacccag   51419
                                                                   ********************************* ** ***********************

chimp.LOC112206744.LOC107973052.GGT2-.646279-708040.rev.compl      cacatgattccttccaccttgtagttagacagaagttctttttgttttgttttg-tttt   52485
```

```
FAM230B-LOC105372935-GGT2.NCBI.GGT2.rev.compl                  catatgattccttccaccttgtagttagacagaagttctttttgttttgttttgttttt   51479
                                                               ** **************************************************** ****

chimp.LOC112206744.LOC107973052.GGT2-.646279-708040.rev.compl  ctttgtttgtttttgagatggagtcttgctctgtctcccagactgcagtgcagtggcatg   52545
FAM230B-LOC105372935-GGT2.NCBI.GGT2.rev.compl                  ctttgtttgtttttgagatggagtcttgctctgtctcccagactgcagtgcagtggcatg   51539
                                                               ************************************************************

chimp.LOC112206744.LOC107973052.GGT2-.646279-708040.rev.compl  atctcagctcactataacctccgcctcccaggttcaagcgattctcctgcctcagcctcc   52605
FAM230B-LOC105372935-GGT2.NCBI.GGT2.rev.compl                  atctcagctcactataacctccgcctcccaggttcaagcgattctcctgcctcagcctcc   51599
                                                               ************************************************************

chimp.LOC112206744.LOC107973052.GGT2-.646279-708040.rev.compl  cgagtagctgggattacaggcacaggccagcacgccaggctaattttgtatttttagta   52665
FAM230B-LOC105372935-GGT2.NCBI.GGT2.rev.compl                  cgagtagctgggattacaggcacaggccagcacgccaggctaattttgtatttttagta   51659
                                                               ************************************************************

chimp.LOC112206744.LOC107973052.GGT2-.646279-708040.rev.compl  cagatggtgttttgccatgttggccaggctgttttcaaactcttgacctcaggtcatcca   52725
FAM230B-LOC105372935-GGT2.NCBI.GGT2.rev.compl                  cagatggggttttgccatgttggccaggctgttttcaaactcctgacctcaggtcatcca   51719
                                                               ******* ********************************** *****************

chimp.LOC112206744.LOC107973052.GGT2-.646279-708040.rev.compl  cccacctcagccttccaaggtgctgggatgacaggcgtgagccaccgtgcccagccaaga   52785
FAM230B-LOC105372935-GGT2.NCBI.GGT2.rev.compl                  cccacctcagccttccaaggtgctgggatgacaggcgtgagccaccgtgcccagccaaga   51779
                                                               ************************************************************

chimp.LOC112206744.LOC107973052.GGT2-.646279-708040.rev.compl  caggagaagttctaatctttgatagcagaccagggtgacgatgcttagcaacagtatttt   52845
FAM230B-LOC105372935-GGT2.NCBI.GGT2.rev.compl                  caggagaagttctaatctttgatagcagaccagggtgacgatgcttagcaacagtatttt   51839
                                                               ************************************************************

chimp.LOC112206744.LOC107973052.GGT2-.646279-708040.rev.compl  gtatatttcaaagtaacgaagagaggactatggtgctaacacccagaaatgaaaaatatt   52905
FAM230B-LOC105372935-GGT2.NCBI.GGT2.rev.compl                  gtatatttcaaagtaacgaagagaggactatggtgctaacacccggaaatgaaaaatatt   51899
                                                               ******************************************** ***************

chimp.LOC112206744.LOC107973052.GGT2-.646279-708040.rev.compl  caaggtgacggagactccaaataccctgccttgatcattatacactatatgcatggaaca   52965
FAM230B-LOC105372935-GGT2.NCBI.GGT2.rev.compl                  caaggtgacggagactccaaataccctgccttgatcattatacactctatgcatggaaca   51959
                                                               ********************************************** *************

chimp.LOC112206744.LOC107973052.GGT2-.646279-708040.rev.compl  agcactcacatgtacccataaatatagaaaatatcatgtatcaatatcagaaaaaaatct   53025
FAM230B-LOC105372935-GGT2.NCBI.GGT2.rev.compl                  agcactcacatgtacccataaatatagaaaatatcatgtatcaatatcagaaaaaaatct   52019
                                                               ************************************************************

chimp.LOC112206744.LOC107973052.GGT2-.646279-708040.rev.compl  cctcctgacctcagcccaatcaggctctcatgccaccacacttgccaagttctctggtga   53085
FAM230B-LOC105372935-GGT2.NCBI.GGT2.rev.compl                  cctcctgacctcagcccaatcaggctctcatgccaccacacttgccaagttctctggtga   52079
                                                               ************************************************************

chimp.LOC112206744.LOC107973052.GGT2-.646279-708040.rev.compl  cccccacactgccagacccagtgccccctctcagctttactgggctcatcactctccctg   53145
FAM230B-LOC105372935-GGT2.NCBI.GGT2.rev.compl                  cccccacactgccagacccagtgccccctctcagctttactgggctcatcactctccctg   52139
                                                               ************************************************************

chimp.LOC112206744.LOC107973052.GGT2-.646279-708040.rev.compl  agagccgcccctgcatcccagcacctggctccacccgagtctcccccgcctgccatccta   53205
FAM230B-LOC105372935-GGT2.NCBI.GGT2.rev.compl                  agagccgcccctgcatcccagcacctggctccacccgagtctcccccgcctgccatccta   52199
                                                               ************************************************************

chimp.LOC112206744.LOC107973052.GGT2-.646279-708040.rev.compl  gctcctactctcccctctgtctttgctctctctcctggtggtctgcttgacatctgagct   53265
FAM230B-LOC105372935-GGT2.NCBI.GGT2.rev.compl                  gctcctactctcccctctgtctttgctctctctcctggtggtctgcttgacatctgagct   52259
                                                               ************************************************************

chimp.LOC112206744.LOC107973052.GGT2-.646279-708040.rev.compl  tcagcctccatttatgcactgacaactcccaaattgacctgctggcctggactgctcctc   53325
FAM230B-LOC105372935-GGT2.NCBI.GGT2.rev.compl                  tcagcctccatttatgcactgacaactctcaaattgacctgctggcctggactgctcctc   52319
                                                               **************************** *******************************

chimp.LOC112206744.LOC107973052.GGT2-.646279-708040.rev.compl  tgatcaccagacctgagtatctacctgcctgcttgaagaaagcatctcaaacttcaacgt   53385
FAM230B-LOC105372935-GGT2.NCBI.GGT2.rev.compl                  cgatcaccagacctgaggatctacctgcctgctcgaagaaagcatctcaaacttcaacgt   52379
                                                                **************** *************** **************************

chimp.LOC112206744.LOC107973052.GGT2-.646279-708040.rev.compl  gcccaaaaccgagctcctgagtgtctgctcaacctgcttcctgagaaccctgcctgtctc   53445
FAM230B-LOC105372935-GGT2.NCBI.GGT2.rev.compl                  gcccaaaaccgagctcctgagtgtctgctcaacctgcttcctgagaaccctgcctgtctc   52439
                                                               ************************************************************

chimp.LOC112206744.LOC107973052.GGT2-.646279-708040.rev.compl  cattagggtcaccccatccttccaggtacagacaaaagatcaggggtccccggggactcc   53505
FAM230B-LOC105372935-GGT2.NCBI.GGT2.rev.compl                  cattagggtcaccccatccttccaggtacagacaaaagatcaggggtccccggggactcc   52499
                                                               ************************************************************

chimp.LOC112206744.LOC107973052.GGT2-.646279-708040.rev.compl  ctacacaagcgtcacacccaacccatcctcaaatccgcaggctccacttccaagtgtgtc   53565
FAM230B-LOC105372935-GGT2.NCBI.GGT2.rev.compl                  ctacacaagcgtcacacccaacccatcctcaaatccacaggctccacttccaagtgtgtc   52559
                                                               ************************************ ***********************

chimp.LOC112206744.LOC107973052.GGT2-.646279-708040.rev.compl  tgtccagcgtcagccacttcccagcactctctccacgaattactgcagtgacctccggac   53625
FAM230B-LOC105372935-GGT2.NCBI.GGT2.rev.compl                  tgtgcagcgtcagccacttcccagcaccctctccacgaattactgcagtgacctccggac   52619
                                                               *** *********************** ********************************

chimp.LOC112206744.LOC107973052.GGT2-.646279-708040.rev.compl  aggccccacatgctccctgccccttacacagcaacccaaggggtccatagaccagatcc   53685
FAM230B-LOC105372935-GGT2.NCBI.GGT2.rev.compl                  aggtccccacatgctccctgccccttacacagcaatccaaggggtccatagaccagatcc   52679
                                                               *** ****************************** *************************

chimp.LOC112206744.LOC107973052.GGT2-.646279-708040.rev.compl  atcgccttccactcacatactccacgagcccccacttccctcagacaggaagcagaggct   53745
FAM230B-LOC105372935-GGT2.NCBI.GGT2.rev.compl                  atccccttccagtcacacactccacgagcccccacttccctcagacaggaagcagaggct   52739
                                                               *** ******* ***** ******************************************

chimp.LOC112206744.LOC107973052.GGT2-.646279-708040.rev.compl  tcaccataacctaagagatcccacacaacctgggccattccccttgggtcacttgctgca   53805
FAM230B-LOC105372935-GGT2.NCBI.GGT2.rev.compl                  tcaccataacctaagagatcccgcacaacctgggccattccccttggatcacttgctgca   52799
                                                               ********************** ************************ ************

chimp.LOC112206744.LOC107973052.GGT2-.646279-708040.rev.compl  gcctccccagctccccacagggctctgtccctgccatcacacctggatagcagaccagga   53865
FAM230B-LOC105372935-GGT2.NCBI.GGT2.rev.compl                  gcctccccagctccccacagggctctgtccctgccatcacacctggatagcagaccagga   52859
                                                               ************************************************************

chimp.LOC112206744.LOC107973052.GGT2-.646279-708040.rev.compl  gatagctcccctgaccccatctctgcctctgggtctttgctcagatgtccccttccctga   53925
FAM230B-LOC105372935-GGT2.NCBI.GGT2.rev.compl                  gataactcccctgaccccatctctgcctctgggtctttgctcagatgtccccttccctga   52919
                                                               **** *******************************************************

chimp.LOC112206744.LOC107973052.GGT2-.646279-708040.rev.compl  ctaggtcaccctccatagagtcccagattttgaggccctccaggtctgtttttctacagc   53985
FAM230B-LOC105372935-GGT2.NCBI.GGT2.rev.compl                  ctaggtcaccctccatagagtcccagattttgaggccctccaggtctgtttttctacagc   52979
                                                               ************************************************************

chimp.LOC112206744.LOC107973052.GGT2-.646279-708040.rev.compl  ctgtaacacacccgcacctgcctagtttcttttcccacctggagtgtcacagatttcatc   54045
FAM230B-LOC105372935-GGT2.NCBI.GGT2.rev.compl                  ccgtaacacacccgcacttgcctagtttctcctcccacctggagtgtcacagatttcatc   53039
                                                               * *************** ************  ****************************

chimp.LOC112206744.LOC107973052.GGT2-.646279-708040.rev.compl  tgccgtctttgtttttcaccccagcttcaggaacaacagctgattctttaagacaatgct   54105
FAM230B-LOC105372935-GGT2.NCBI.GGT2.rev.compl                  tgccgtctttgtttttcaccccagcttcaggaacaacagctgattctttaagacaatgct   53099
                                                               ************************************************************

chimp.LOC112206744.LOC107973052.GGT2-.646279-708040.rev.compl  caatacattctagttaaataaatgattctaagcgtccacaaggtgccaagcctgtgattc   54165
FAM230B-LOC105372935-GGT2.NCBI.GGT2.rev.compl                  caatacattctagtcaaataaatggttttaagcgtccacaaggtgccaagcctatggttc   53159
                                                               ************** ********* ** ************************* ** ***

chimp.LOC112206744.LOC107973052.GGT2-.646279-708040.rev.compl  ccgcattgtcttaccctgagcaacttcacatctacagatgctgagtttctcaatgagtat   54225
FAM230B-LOC105372935-GGT2.NCBI.GGT2.rev.compl                  ccgcattctcttaccctcagcaacttcatgtctacagatgctgagtttctcaatgagtat   53219
                                                               ******* ********* **********  ******************************

chimp.LOC112206744.LOC107973052.GGT2-.646279-708040.rev.compl  taaaaacaaatgaaagattggtcgggcacagtggctcacgcctgtaattccagcactttg   54285
FAM230B-LOC105372935-GGT2.NCBI.GGT2.rev.compl                  taaaaacaaatgaaagattggtcgggcacagtggctcacgcctgtaattccagcactttg   53279
                                                               ************************************************************

chimp.LOC112206744.LOC107973052.GGT2-.646279-708040.rev.compl  ggaggctgaggcaggggatcacgaggtcaggagatcgagggaccagcctgggcaacata   54345
FAM230B-LOC105372935-GGT2.NCBI.GGT2.rev.compl                  ggaggctgaggcaggggatcacgaggtcaggagatcgagggaccagcctgggcaacata   53339
                                                               ************************************************************

chimp.LOC112206744.LOC107973052.GGT2-.646279-708040.rev.compl  gtgaaaacccgtctctactaaaaaatacaaaaaattagccgggtgtggtggcaggcgcct   54405
FAM230B-LOC105372935-GGT2.NCBI.GGT2.rev.compl                  gtgaaaacccgtctctactaaaaaatacaaaaaattagccgggcgcggtggcgggcgcct   53399
                                                               ******************************************* * ****** *******

chimp.LOC112206744.LOC107973052.GGT2-.646279-708040.rev.compl  gtagtcccagctactcaggaggctgaggcaggagaatggcgtgaactcaggaggcagagc   54465
FAM230B-LOC105372935-GGT2.NCBI.GGT2.rev.compl                  gtagtcccagctactcgggaggctgaggcaggagaatggcgcgaactcaggaggcagagc   53459
                                                               **************** *********************** ******************

chimp.LOC112206744.LOC107973052.GGT2-.646279-708040.rev.compl  ttgcagtgagccgagatcgcaccactgcactccaacctgggcgacagagtaagactccgt   54525
FAM230B-LOC105372935-GGT2.NCBI.GGT2.rev.compl                  ttgcagtgagccgagatcgcaccactgcactccaacctgggcgacagagtgagactctgt   53519
                                                               ************************************************** ****** **

chimp.LOC112206744.LOC107973052.GGT2-.646279-708040.rev.compl  ctcaaaaaaaaaaaaagaaaaaaaaaatcaaagattgagtatgttgcagaagactccaaa   54585
FAM230B-LOC105372935-GGT2.NCBI.GGT2.rev.compl                  ctcaaaaaaaaaaaagaaaaaaaaaaatcaaagattgagtatgttgcagaagactccaaa   53579
                                                               ***************  *******************************************

chimp.LOC112206744.LOC107973052.GGT2-.646279-708040.rev.compl  gggcaccacccaggacccccacctgaagtctaagacctgctgtggtaagtgtgtcctgcc   54645
FAM230B-LOC105372935-GGT2.NCBI.GGT2.rev.compl                  gggcaccacccaggacccccacctgaagtctaagacctgctatggtgagtgtgtcctgcc   53639
```

```
                                                                   ***************************************** **** *************

chimp.LOC112206744.LOC107973052.GGT2-.646279-708040.rev.compl      cctccatcctccaac---ttttttttttttttttttttgagacggagcttcgttcttgtt   54702
FAM230B-LOC105372935-GGT2.NCBI.GGT2.rev.compl                      cctccatcctccaacttttttttttttttttttttttgagacggagcttcgttcttgtt   53699
                                                                   ***************   ******************************************

chimp.LOC112206744.LOC107973052.GGT2-.646279-708040.rev.compl      gcccaggctggagtgcagtggcatgatctcggctcactgcaatctccacctcgtgggttc   54762
FAM230B-LOC105372935-GGT2.NCBI.GGT2.rev.compl                      gcccaggctggagtgcagtggcatgatctcggctcactgcaatctctacctcgtgggttc   53759
                                                                   ********************************************** *************

chimp.LOC112206744.LOC107973052.GGT2-.646279-708040.rev.compl      aagcgattctcctgcctcagcctcctgagtagctgggatattacaggcctgtgccaccac   54822
FAM230B-LOC105372935-GGT2.NCBI.GGT2.rev.compl                      aagcgattctcctgccttagcctcctgagtagctgggatattacaggcctgtgccaccac   53819
                                                                   ***************** ******************************************

chimp.LOC112206744.LOC107973052.GGT2-.646279-708040.rev.compl      gcccaactaattattatacttttagtagagaaagggtttcactatgttggccaggctggt   54882
FAM230B-LOC105372935-GGT2.NCBI.GGT2.rev.compl                      gcccgactaattattgtacttttagtagagaaagggtttcactatgttggccaggctggt   53879
                                                                   **** ********** ********************************************

chimp.LOC112206744.LOC107973052.GGT2-.646279-708040.rev.compl      cttgaactcctgacctaggtgatccatctgcctcagcctcccaaagtgctgggattacag   54942
FAM230B-LOC105372935-GGT2.NCBI.GGT2.rev.compl                      cttggactcctgacctaggtgatccatctgcctcagcctcccaaagtgctgggattacag   53939
                                                                   **** *******************************************************

chimp.LOC112206744.LOC107973052.GGT2-.646279-708040.rev.compl      gcgtgagcctgtgaaaaaaaggcccagcctttttttttttttttttttttttgacagcgtc   55002
FAM230B-LOC105372935-GGT2.NCBI.GGT2.rev.compl                      gcgtgagcctgtgaaaaaaaggcccagcc----ttttttttttttttttttgacagggtc   53995
                                                                   *****************************    *********************** ***

chimp.LOC112206744.LOC107973052.GGT2-.646279-708040.rev.compl      tcactttgttgcccaagctagagtgtagtggtataatcatggctgactgcagcctcaacc   55062
FAM230B-LOC105372935-GGT2.NCBI.GGT2.rev.compl                      tcactttgttgcccaagctagagtgtagtggtataatcatggctgactgcagcctcaacc   54055
                                                                   ************************************************************

chimp.LOC112206744.LOC107973052.GGT2-.646279-708040.rev.compl      tcctgggctcaagtgatcctcccaccttagcctcccgagtagctgggaccataagcacac   55122
FAM230B-LOC105372935-GGT2.NCBI.GGT2.rev.compl                      tcctgggctcaagtgatcctcccaccttagcctcccgagtaactgggaccataagcacac   54115
                                                                   ***************************************** ******************

chimp.LOC112206744.LOC107973052.GGT2-.646279-708040.rev.compl      accgccatacctagctaatttttttttccatttttgtagagatggagtcttgctatgttg   55182
FAM230B-LOC105372935-GGT2.NCBI.GGT2.rev.compl                      accgccatacctgctaattttttttttccatttttgtagagatggagtcttgctatgttg   54175
                                                                   ************    * ******************************************

chimp.LOC112206744.LOC107973052.GGT2-.646279-708040.rev.compl      tccaggcaggtctcctgggctcatgcgctcctcctgacttggcctcctaaagtgctagga   55242
FAM230B-LOC105372935-GGT2.NCBI.GGT2.rev.compl                      tccaggcaggtctcctgggctcatgcgctcctcctgacttggcctcctaaagtgctagga   54235
                                                                   ************************************************************

chimp.LOC112206744.LOC107973052.GGT2-.646279-708040.rev.compl      ttagaggcgtgagttgttgagaccctcccatcctccaacttttatctcacaatctattgt   55302
FAM230B-LOC105372935-GGT2.NCBI.GGT2.rev.compl                      ttagaggcgtgagttgttgagaccctcccatcctccaacttttatctcacaatctattgt   54295
                                                                   ************************************************************

chimp.LOC112206744.LOC107973052.GGT2-.646279-708040.rev.compl      gcctcctttggggacagacagtggcttcctggatggacagtggcttcccttcaggtacct   55362
FAM230B-LOC105372935-GGT2.NCBI.GGT2.rev.compl                      gcctcctttggggacagacagtggcttcctggatggacagtggcttcccctcaggtacct   54355
                                                                   ************************************************* **********

chimp.LOC112206744.LOC107973052.GGT2-.646279-708040.rev.compl      ggggaatttggggcctcctctccacttaagaccagattagaaaagagggactccacctc   55422
FAM230B-LOC105372935-GGT2.NCBI.GGT2.rev.compl                      ggggaatttggggcctcctccccacttaagaccagattagaaaagagagactccacctc   54415
                                                                   ******************** ************************** ************

chimp.LOC112206744.LOC107973052.GGT2-.646279-708040.rev.compl      acattctagagcaccatccccacaaatgaacaaatgagtgaatgggatgcctgttgaaaa   55482
FAM230B-LOC105372935-GGT2.NCBI.GGT2.rev.compl                      acattctagagcgccatccccacaaatgaacaaatgagtgaatgggatgcctgttgaaaa   54475
                                                                   ************ ***********************************************

chimp.LOC112206744.LOC107973052.GGT2-.646279-708040.rev.compl      agcaggatatagacagcctgggttcaattttagctttaccacctcccagctgtgtgacct   55542
FAM230B-LOC105372935-GGT2.NCBI.GGT2.rev.compl                      ggcaggatatagacagcctgggttcaattttagcttcaccacctcccagctgtgtgacct   54535
                                                                    *********************************** ***********************

chimp.LOC112206744.LOC107973052.GGT2-.646279-708040.rev.compl      caactgatttgcatgacctgtctgagcctcagcatccccaccctgtaaaatgggaatcca   55602
FAM230B-LOC105372935-GGT2.NCBI.GGT2.rev.compl                      cagctgatttgcatgacctgtctgagcctcagcatccccaccctgtaaaatgggaatcca   54595
                                                                   ** *********************************************************

chimp.LOC112206744.LOC107973052.GGT2-.646279-708040.rev.compl      cacagcatcccctagcccaaaggagcagggagggttgtgagaggctcgtgggtgaaaagc   55662
FAM230B-LOC105372935-GGT2.NCBI.GGT2.rev.compl                      cacagcatcccctagcccaaaggagcagggagggttgtgagaggctcgtgggtgaaaagc   54655
                                                                   ************************************************************

chimp.LOC112206744.LOC107973052.GGT2-.646279-708040.rev.compl      acagagcagagcgtgggccccagtgagccctggtccatgaggtctgctagcataataatt   55722
FAM230B-LOC105372935-GGT2.NCBI.GGT2.rev.compl                      acagagcagagcatgggccccagtgagccctgatccatgaggtctgctagcataataatt   54715
                                                                   ************ ******************* ***************************

chimp.LOC112206744.LOC107973052.GGT2-.646279-708040.rev.compl      attctttccatgtgctgcacagagtggccccggaggccttagcagaaataacagaagctc   55782
FAM230B-LOC105372935-GGT2.NCBI.GGT2.rev.compl                      attctttccatgtgctgcacagagtggccccggaggccttagcagaaataacagaagctc   54775
                                                                   ************************************************************

chimp.LOC112206744.LOC107973052.GGT2-.646279-708040.rev.compl      ccggccctttaccgtggtgatgatggtcctgaccactcattgtgggagggtgctatgggg   55842
FAM230B-LOC105372935-GGT2.NCBI.GGT2.rev.compl                      ccggccctttaccgtggtgatgatggtcctgaccactcattgtgggagggtgctatgggg   54835
                                                                   ************************************************************

chimp.LOC112206744.LOC107973052.GGT2-.646279-708040.rev.compl      caaggaagggatggggggtgctagaactgcccctgaaccctgacgggagcaggctcctgt   55902
FAM230B-LOC105372935-GGT2.NCBI.GGT2.rev.compl                      ccaggaagggatggggggtgacagaactgcccctgaaccctgacgggagcaggcttctgt   54895
                                                                   * ******************  ********************************* ****

chimp.LOC112206744.LOC107973052.GGT2-.646279-708040.rev.compl      gggcaaggccccttcccggtggctcagccagctctgcacccatgccccaagtctgcagca   55962
FAM230B-LOC105372935-GGT2.NCBI.GGT2.rev.compl                      gggcaaggccccttcccggtggctcagccagctctgcacccatgccccaagtctgcagca   54955
                                                                   ************************************************************

chimp.LOC112206744.LOC107973052.GGT2-.646279-708040.rev.compl      tggcttacccttctgggactgctccgagctgttgaacatgctggcaaaggccagcctggg   56022
FAM230B-LOC105372935-GGT2.NCBI.GGT2.rev.compl                      tggcttacccttctgggactgctccgagctgttgaacatgctggcaaaggccagcctggg   55015
                                                                   ************************************************************

chimp.LOC112206744.LOC107973052.GGT2-.646279-708040.rev.compl      ggccacctcgcgggcgttgatgacctcagcttttcctagaaggagaagcaggtaggcagg   56082
FAM230B-LOC105372935-GGT2.NCBI.GGT2.rev.compl                      ggccacctcgcgggcattgatgacctcagcttttcctaga---agaagcaggtaggcagg   55072
                                                                   *************** ************************   *****************

chimp.LOC112206744.LOC107973052.GGT2-.646279-708040.rev.compl      cccacccacccaaaccctttatgccacgtgagcctgggggccacccagctgtgcctcggc   56142
FAM230B-LOC105372935-GGT2.NCBI.GGT2.rev.compl                      cccacccaccaaaaccctttatgccacgtgagcctgggggccacccagctgtgcctcggc   55132
                                                                   ********** *************************************************

chimp.LOC112206744.LOC107973052.GGT2-.646279-708040.rev.compl      ccaacccacacccccctgcccctctccctctcctcttccgaggcactcatgagtggtgctg   56202
FAM230B-LOC105372935-GGT2.NCBI.GGT2.rev.compl                      ccaacccacacccccctgcccctctccctctcctcttctgaggcactcatgagtggtgctg   55192
                                                                   ************************************* **********************

chimp.LOC112206744.LOC107973052.GGT2-.646279-708040.rev.compl      ttgtagatggtgaggaagaggccacccccgatgcccatgctgtgggcattcatgagcccc   56262
FAM230B-LOC105372935-GGT2.NCBI.GGT2.rev.compl                      ttgtagatggtgaggaagaggccaaccccgatgcccatgctgtgggcattcatgagcccc   55252
                                                                   ************************ ***********************************

chimp.LOC112206744.LOC107973052.GGT2-.646279-708040.rev.compl      acacacaacagggctgcaatggctgcatccaccgcagagccaccgtcccgcagtgcgtcc   56322
FAM230B-LOC105372935-GGT2.NCBI.GGT2.rev.compl                      acacacaacagggctgcaatggctgcatccactgcagagccaccgtcccgcagtgtgtcc   55312
                                                                   ******************************** ********************** ****

chimp.LOC112206744.LOC107973052.GGT2-.646279-708040.rev.compl      ctgccatgtggcacataaaggcatgagaacctgcaggcttccaccctggccccgcataca   56382
FAM230B-LOC105372935-GGT2.NCBI.GGT2.rev.compl                      ctgccatgtggcacataaaggcatgagaacctgcaggcttccaccctggccccgcataca   55372
                                                                   ************************************************************

chimp.LOC112206744.LOC107973052.GGT2-.646279-708040.rev.compl      caccctgctgcccacctgcccaaaggaggatggaagagaagtccattcgagttttggggt   56442
FAM230B-LOC105372935-GGT2.NCBI.GGT2.rev.compl                      caccctgctgcccacctgcccaaaggaggatggaagagaagtccattcgagttttggggt   55432
                                                                   ************************************************************

chimp.LOC112206744.LOC107973052.GGT2-.646279-708040.rev.compl      ttttgtttttagtttctttctctttttttttgagatggagtcttgctctattgccaggctg   56502
FAM230B-LOC105372935-GGT2.NCBI.GGT2.rev.compl                      ttttgtttttagtttctttctctttttttttgagatggagtcttgctctattgccaggctg   55492
                                                                   ************************************************************

chimp.LOC112206744.LOC107973052.GGT2-.646279-708040.rev.compl      gagtgcagtggcacgatctcagctcactgcaacctctgcctcccgggttcaagccattct   56562
FAM230B-LOC105372935-GGT2.NCBI.GGT2.rev.compl                      gagtgcagtggcacgatctcagctcactgcaacctctgcctcccgggttcaagccattct   55552
                                                                   ************************************************************

chimp.LOC112206744.LOC107973052.GGT2-.646279-708040.rev.compl      cctgcctcagcctcctgagtagctgggactacaggcgcacaccaccacgcccggctaatt   56622
FAM230B-LOC105372935-GGT2.NCBI.GGT2.rev.compl                      cctgcctcagcctcctgagtagctgggactacaggtgcatgccaccacgcccagctaatt   55612
                                                                   *********************************** ***  *********** *******

chimp.LOC112206744.LOC107973052.GGT2-.646279-708040.rev.compl      tttgtatttttagtagagacggggtttcaccatgttggccaggatggtctctatctcttg   56682
FAM230B-LOC105372935-GGT2.NCBI.GGT2.rev.compl                      tttgtatttttagtagagatggggtttcaccatgttggccaggatggtctctatctcttg   55672
                                                                   ****************** *****************************************

chimp.LOC112206744.LOC107973052.GGT2-.646279-708040.rev.compl      acttcatgatccacctgccttggcctc--------tttttttttttttttggagacaaag   56734
FAM230B-LOC105372935-GGT2.NCBI.GGT2.rev.compl                      acttcatgatccgcctgccttggcctcttttttttttttttttttttttggagacaaag   55732
                                                                   ************ **************        *************************

chimp.LOC112206744.LOC107973052.GGT2-.646279-708040.rev.compl      tctctctttgttgctcaggctggagtgcagtggtgtgatctcggatcactgcagcctcag   56794
FAM230B-LOC105372935-GGT2.NCBI.GGT2.rev.compl                      tctctctttgttgctcaggctggagtgcagtggtgtgatcttggatcactgcagcctcag   55792
                                                                   ***************************************** ******************
```

```
chimp.LOC112206744.LOC107973052.GGT2-.646279-708040.rev.compl      tctcctgggctcaattgaccctcccatctcagcctcccaagtagctgggactatgggcac   56854
FAM230B-LOC105372935-GGT2.NCBI.GGT2.rev.compl                      tctcctgggctcaattgaccctcccatctcagcctcccaagtagctgggactatgggcac   55852
                                                                   ************************************************************

chimp.LOC112206744.LOC107973052.GGT2-.646279-708040.rev.compl      atgccaccatgcccagccaat----tttgtttgtgtgttttgtagagatggggtttcatc   56910
FAM230B-LOC105372935-GGT2.NCBI.GGT2.rev.compl                      atgccaccatgcccagccaattttgtttgtttgtgtattttgtagagatggggtttcatc   55912
                                                                   *********************    *********** ***********************

chimp.LOC112206744.LOC107973052.GGT2-.646279-708040.rev.compl      atgttgcccaggctggtcaagaactcctgtgctcaagtgatccacccaccttggcctccc   56970
FAM230B-LOC105372935-GGT2.NCBI.GGT2.rev.compl                      atgttgcccaggctggtcaagaactcctgtgctcaagtaatccacccaccttggcctccc   55972
                                                                   ************************************* **********************

chimp.LOC112206744.LOC107973052.GGT2-.646279-708040.rev.compl      aaagtgctggtattacaggcatgagccactgtgctcagcctttgttttatgagacagggt   57030
FAM230B-LOC105372935-GGT2.NCBI.GGT2.rev.compl                      aaagtgctggtattacaggcatgagccactgtgcccagcctttgttttatgagacagggt   56032
                                                                   ********************************** *************************

chimp.LOC112206744.LOC107973052.GGT2-.646279-708040.rev.compl      ctcactttgtcacccaggatgaagtgcagtggcacagtcttggctcaatgcagctttgac   57090
FAM230B-LOC105372935-GGT2.NCBI.GGT2.rev.compl                      ctcactttgtcacccaggatgaagtgcagtggcacagtcttggctcaatgcagctttgac   56092
                                                                   ************************************************************

chimp.LOC112206744.LOC107973052.GGT2-.646279-708040.rev.compl      ctcctgggctcaagcaatcctcccacttcagtctcctgagtagctgggactacaggtaag   57150
FAM230B-LOC105372935-GGT2.NCBI.GGT2.rev.compl                      ctcctgggctcaagcaatcctcccacttcagtctcctgagtagctgggactacaggtaag   56152
                                                                   ************************************************************

chimp.LOC112206744.LOC107973052.GGT2-.646279-708040.rev.compl      aaccaccacaccgggcaattttttgtcttttttgtagagatagggtctttctatgttgcc   57210
FAM230B-LOC105372935-GGT2.NCBI.GGT2.rev.compl                      aaccaccacaccgggcaattttttgtcttttttgtagagatagggtctttctatgttgcc   56212
                                                                   ************************************************************

chimp.LOC112206744.LOC107973052.GGT2-.646279-708040.rev.compl      caggctggtctcaaactcatggtctaaagcaatcttatctcctcaacctcccaaagtgct   57270
FAM230B-LOC105372935-GGT2.NCBI.GGT2.rev.compl                      caggctggtctcaaactcatggtctaaagcaatcctatcgcctcaacctcccaaagtgct   56272
                                                                   ********************************** **** ********************

chimp.LOC112206744.LOC107973052.GGT2-.646279-708040.rev.compl      gggattacagtttcttctttttcttttc----------ttttttttttctgagacagagt   57320
FAM230B-LOC105372935-GGT2.NCBI.GGT2.rev.compl                      gggattacagtttcttctttttcttttcttttttcttttttttttttctgagacagagt   56332
                                                                   ****************************          **********************

chimp.LOC112206744.LOC107973052.GGT2-.646279-708040.rev.compl      ttcactcagttgcccaggctggagtgcagtggcatgatcttagctcactgcaacctctgc   57380
FAM230B-LOC105372935-GGT2.NCBI.GGT2.rev.compl                      ttcactcagttgcccaggctggagtgcagtggcatgatcttagctcactgcaacctctgc   56392
                                                                   ************************************************************

chimp.LOC112206744.LOC107973052.GGT2-.646279-708040.rev.compl      ctcctgggttcaagcgattctcctgcctcagcctcctgagtagctgggattacaggcgca   57440
FAM230B-LOC105372935-GGT2.NCBI.GGT2.rev.compl                      ctcctgggttcaagcgattctcctgcctcagcctcctgagtagctgggattacaggcgca   56452
                                                                   ************************************************************

chimp.LOC112206744.LOC107973052.GGT2-.646279-708040.rev.compl      caccaccatgcccggctaactttttatatttttagtagggacagagtgcaccatgttggc   57500
FAM230B-LOC105372935-GGT2.NCBI.GGT2.rev.compl                      caccaccatgcccggctaactttttatatttttagtagggacagagtgcaccatgttggc   56512
                                                                   ************************************************************

chimp.LOC112206744.LOC107973052.GGT2-.646279-708040.rev.compl      caggctggtctcgaactcctgacctcgggtggtctgcccgcctcaaccccccaaagtgct   57560
FAM230B-LOC105372935-GGT2.NCBI.GGT2.rev.compl                      caggctgatctcgaactcctgacctc-agtggtctgcccgcctcaaccccccaaagtgct   56571
                                                                   ******* ******************  ********************************

chimp.LOC112206744.LOC107973052.GGT2-.646279-708040.rev.compl      gggattactggtgtggaccaccttgcccagacagtttcctctttatcaagcaaacaaatg   57620
FAM230B-LOC105372935-GGT2.NCBI.GGT2.rev.compl                      gggattacaggtgtggaccaccttgcccagacagtttcctctttattaagcaaacaaatg   56631
                                                                   ******** ************************************* *************

chimp.LOC112206744.LOC107973052.GGT2-.646279-708040.rev.compl      tacatgacttttataattgggacaaaaagggaaattgctatactttattaataacatttt   57680
FAM230B-LOC105372935-GGT2.NCBI.GGT2.rev.compl                      tacatgacttttataattgggacaaaaagggaaattgctatactttattaataacatttt   56691
                                                                   ************************************************************

chimp.LOC112206744.LOC107973052.GGT2-.646279-708040.rev.compl      ttttttccttgctagagatggtggcttacacctgtaatctcagcactctgggaggccaag   57740
FAM230B-LOC105372935-GGT2.NCBI.GGT2.rev.compl                      ttttttccctgctagagatggtggcttacacctgtaatctcagcactttgggaggccaag   56751
                                                                   ******** ************************************** ************

chimp.LOC112206744.LOC107973052.GGT2-.646279-708040.rev.compl      gtggaggatcacttgaggccaggagttcaagaccagcctgggcaatagagtgagaccatc   57800
FAM230B-LOC105372935-GGT2.NCBI.GGT2.rev.compl                      gtggaggatcacttgaggccaggagttcaagaccagcctgggcaatagagtgagaccatc   56811
                                                                   ************************************************************

chimp.LOC112206744.LOC107973052.GGT2-.646279-708040.rev.compl      tctagaaaacagtcttttttaattagtcaggtgtgatgcacgcctgtagtcctagctact   57860
FAM230B-LOC105372935-GGT2.NCBI.GGT2.rev.compl                      tctacaaaacagtcttttttaattagtcaggtatgatgcacgcctgtagtcctagctact   56871
                                                                   **** *************************** ***************************

chimp.LOC112206744.LOC107973052.GGT2-.646279-708040.rev.compl      cagggggctgaggtgagaggatcgcttaagcccaagagttcaaggctgcagtgagctatg   57920
FAM230B-LOC105372935-GGT2.NCBI.GGT2.rev.compl                      cagggggctgaggtgagaggatcgcttaagcccaagagttcaaggctgcagtgagctatg   56931
                                                                   ************************************************************

chimp.LOC112206744.LOC107973052.GGT2-.646279-708040.rev.compl      atcatgccactgcactccagcctgggtgacagaacaagaccctgtctcaaaatatgaaaa   57980
FAM230B-LOC105372935-GGT2.NCBI.GGT2.rev.compl                      atcatgccactgcactccagcctgggtgacagaacaagaccctgtctcaaaatatgaaaa   56991
                                                                   ************************************************************

chimp.LOC112206744.LOC107973052.GGT2-.646279-708040.rev.compl      acataatattttttctgtttaagtctttaagagggaactgatctttatatataacatgag   58040
FAM230B-LOC105372935-GGT2.NCBI.GGT2.rev.compl                      acataatattttttctgtttaagtctttaggagggaacttatctttatatataacatgag   57051
                                                                   ***************************** ********* ********************

chimp.LOC112206744.LOC107973052.GGT2-.646279-708040.rev.compl      ataagagtctaaaatagaataaaacagtagaagccaggcaccatggctcacacctgtaat   58100
FAM230B-LOC105372935-GGT2.NCBI.GGT2.rev.compl                      ataagagtctaaaatagaataacacagtagaagccaggcaccatggctcacacctgtaat   57111
                                                                   ********************** *************************************

chimp.LOC112206744.LOC107973052.GGT2-.646279-708040.rev.compl      cccagcactttaggaagctgaggcgggaggatcacttaaacccaagagtttgaggctgca   58160
FAM230B-LOC105372935-GGT2.NCBI.GGT2.rev.compl                      cccagcactttaggaagctgaggcgggaggatcacttaaacccaagagtttgaggctgca   57171
                                                                   ************************************************************

chimp.LOC112206744.LOC107973052.GGT2-.646279-708040.rev.compl      gtgggctatgatcgctcccctatactccagcctgggtgacagagtaagactccaccttaa   58220
FAM230B-LOC105372935-GGT2.NCBI.GGT2.rev.compl                      gtgggctatgatcgctcccctatactccagcctgggtgacagagtaagactccaccttaa   57231
                                                                   ************************************************************

chimp.LOC112206744.LOC107973052.GGT2-.646279-708040.rev.compl      aaaaagaataagcccttcatgtccctgtttgggcaacagcgttcctt-------gggggg   58273
FAM230B-LOC105372935-GGT2.NCBI.GGT2.rev.compl                      aaaaagaataagcccttcatgtccctgtttgggcaacagcgttccttggggggagggggg   57291
                                                                   ***********************************************       ******

chimp.LOC112206744.LOC107973052.GGT2-.646279-708040.rev.compl      atgggggagggatggcattgggagatatacataatgctaaatgatgagttggtgggtgca   58333
FAM230B-LOC105372935-GGT2.NCBI.GGT2.rev.compl                      aggggggagggatggcattgggagatatacctaatgctaaatgacgagttggtgggtgca   57351
                                                                   * **************************** ************* ***************

chimp.LOC112206744.LOC107973052.GGT2-.646279-708040.rev.compl      gcacaccaacatggcacatgtgtacatatgtaacaaacctgcacattgtgcacatgtacc   58393
FAM230B-LOC105372935-GGT2.NCBI.GGT2.rev.compl                      gcacaccaacatggcacatgtgtacatatgtaacaaacctgcacattgtgcacatgtacc   57411
                                                                   ************************************************************

chimp.LOC112206744.LOC107973052.GGT2-.646279-708040.rev.compl      ctaaacttaaagtataataataataaaataaaataaaaaaaaagaaagaaagaaattgtc   58453
FAM230B-LOC105372935-GGT2.NCBI.GGT2.rev.compl                      ctaaacttaaagt---ataataataaaataaaataaaaaaaaagaaagaaagaaattgtc   57468
                                                                   *************   ********************************************

chimp.LOC112206744.LOC107973052.GGT2-.646279-708040.rev.compl      tcctaaccaaaaaaaaaaaaaaaa-----aaaagaaaagaaaagaataagccgggcgtgg   58508
FAM230B-LOC105372935-GGT2.NCBI.GGT2.rev.compl                      tcctaaccaaaaaaaaaaaaaaaaaaaagaaaagaaaagaaaagaataagccgggcgtgg   57528
                                                                   ************************     *******************************

chimp.LOC112206744.LOC107973052.GGT2-.646279-708040.rev.compl      tggctcacacctgtaatcccaacacttcgggaggcccaggtgggtggattacctgaggtc   58568
FAM230B-LOC105372935-GGT2.NCBI.GGT2.rev.compl                      tggctcatacctgtaatcccaacactttgggaggcccaggtgggtggattacctgaggtc   57588
                                                                   ******* ******************* ********************************

chimp.LOC112206744.LOC107973052.GGT2-.646279-708040.rev.compl      aggagttcgagaccagcctgaccaacatggtgaaaccccatctctactaaaaacaccata   58628
FAM230B-LOC105372935-GGT2.NCBI.GGT2.rev.compl                      aggagttcgagaccagcctgaccaacatggtgaaaccccatctctactaaaaacaccata   57648
                                                                   ************************************************************

chimp.LOC112206744.LOC107973052.GGT2-.646279-708040.rev.compl      attaggccaggcgcagtggctcacacctgtaatcccagcactttgagaggcagaggtggg   58688
FAM230B-LOC105372935-GGT2.NCBI.GGT2.rev.compl                      attaggccaggcgcagtggctcacacctgtaatcccagcactttgagaggcagaggtggg   57708
                                                                   ************************************************************

chimp.LOC112206744.LOC107973052.GGT2-.646279-708040.rev.compl      aggatcacaaggtcaggagttcaagaccagcctggccaacatagcaaaaccctgtctcta   58748
FAM230B-LOC105372935-GGT2.NCBI.GGT2.rev.compl                      aggatcacaaggtcaggagttcaagaccagcctggccaacatagcaaaaccctgtctcta   57768
                                                                   ************************************************************

chimp.LOC112206744.LOC107973052.GGT2-.646279-708040.rev.compl      ctaaaaatacaaaaattagctgggcatggtggcacatgcctgtagtctcagctactgtgg   58808
FAM230B-LOC105372935-GGT2.NCBI.GGT2.rev.compl                      ctaaaaatacaaaaattagctgggcatggtggcacatgcctgtagtctcagctactgtgg   57828
                                                                   ************************************************************

chimp.LOC112206744.LOC107973052.GGT2-.646279-708040.rev.compl      aggctgaggcaggagaatcacttgaacctgggaggtggaggctgcagtaagccaagattg   58868
FAM230B-LOC105372935-GGT2.NCBI.GGT2.rev.compl                      aggctgaggcaggagaatcacttgaacctgggaggtggaggctgcagtaagccaagattg   57888
                                                                   ************************************************************

chimp.LOC112206744.LOC107973052.GGT2-.646279-708040.rev.compl      caccactgccctccagcctgggtaacagagcgatactctgtctcagaaa-----------   58917
FAM230B-LOC105372935-GGT2.NCBI.GGT2.rev.compl                      caccactgccctccagcctgggtaacagagcgatactctgtctcagaaaacacacacaca   57948
                                                                   *************************************************
```

```
chimp.LOC112206744.LOC107973052.GGT2-.646279-708040.rev.compl     ---acacacacacacacacacacacacatatacacacacacaaatttgttgggcgtggtg   58974
FAM230B-LOC105372935-GGT2.NCBI.GGT2.rev.compl                     cacacacacacacacacacacacacacatatacacacacacaaatttgttgggcgtggtg   58008
                                                                     *********************************************************

chimp.LOC112206744.LOC107973052.GGT2-.646279-708040.rev.compl     gcacactcctgtaatcccagctacttgggaggcttaggcatgagaatcacttgaacttgg   59034
FAM230B-LOC105372935-GGT2.NCBI.GGT2.rev.compl                     gcgcactcctataatcccagctacttgggaggcttaggcatgagaatcgcttgaacttgg   58068
                                                                  ** ******* ************************************ ************

chimp.LOC112206744.LOC107973052.GGT2-.646279-708040.rev.compl     gaggcggaggttgcagttaactgagattgcaccactgcactccagcctggcaacagagca   59094
FAM230B-LOC105372935-GGT2.NCBI.GGT2.rev.compl                     gaggcggaggttgcagttaactgagattgcaccactgcactccagcctggcaacagagca   58128
                                                                  ************************************************************

chimp.LOC112206744.LOC107973052.GGT2-.646279-708040.rev.compl     aaactccatctctaaatagat-------------tagatagatagatagatagatagata   59141
FAM230B-LOC105372935-GGT2.NCBI.GGT2.rev.compl                     aaactccatctctaaatagatatagatagatagatagatagatagatagatagatagata   58188
                                                                  *********************             **************************

chimp.LOC112206744.LOC107973052.GGT2-.646279-708040.rev.compl     atagatagatagatagatagatagttagataggaagaaataagtaaaaataacaaccaaa   59201
FAM230B-LOC105372935-GGT2.NCBI.GGT2.rev.compl                     gatagatgatagatagatagatagatagataggaagaaataagtaaaaataacaaccaaa   58248
                                                                         ***************** ***********************************

chimp.LOC112206744.LOC107973052.GGT2-.646279-708040.rev.compl     caacaaaacagtggagtttatccaaagagacagagactcttagaactgagaaaggggcc   59261
FAM230B-LOC105372935-GGT2.NCBI.GGT2.rev.compl                     caacaaaacagtggagtttatccaaagaaacagagactcttagaactgagaaaggggcc   58308
                                                                  **************************** *******************************

chimp.LOC112206744.LOC107973052.GGT2-.646279-708040.rev.compl     ctgtttggctctaggagacccacaccctgctctcggagtcaccgtccccttcccaaaggc   59321
FAM230B-LOC105372935-GGT2.NCBI.GGT2.rev.compl                     ctgtttggctcta-gagacccacaccctgctctcggagtcaccgtccccttcccaaaggc   58367
                                                                  ************* **********************************************

chimp.LOC112206744.LOC107973052.GGT2-.646279-708040.rev.compl     tactgagagagtccaagcaaagcttacatgtggggaaactgagtctcagaggggtgaagg   59381
FAM230B-LOC105372935-GGT2.NCBI.GGT2.rev.compl                     tactgagagagtccaagcgaagcttgcatgtggggaaactgagtctcagaggggtgaagg   58427
                                                                  ****************** ****** **********************************

chimp.LOC112206744.LOC107973052.GGT2-.646279-708040.rev.compl     tattgctcaggtccacttgcccagttttcagggcccatgtcccatgccctgccccgctca   59441
FAM230B-LOC105372935-GGT2.NCBI.GGT2.rev.compl                     tattgctcaggtccacttgcccagttttcagggcccatgtcccatgccctgccccgctca   58487
                                                                  ************************************************************

chimp.LOC112206744.LOC107973052.GGT2-.646279-708040.rev.compl     cctcccaatctccgagcactgcttggcatccgcgcccacggcagccctggtgtacacatg   59501
FAM230B-LOC105372935-GGT2.NCBI.GGT2.rev.compl                     cctcccaatctccaagcactgcttggcatccgcggccatggcagccctggtgtacacatg   58547
                                                                  ************* ******************** *** *********************

chimp.LOC112206744.LOC107973052.GGT2-.646279-708040.rev.compl     gttgtcaggttccttggaggctgagggcagccagagacagaggccgacaatgaccagcac   59561
FAM230B-LOC105372935-GGT2.NCBI.GGT2.rev.compl                     gttgtcaggttccttggaggccgagggcagccagagacagaggccgacaatgaccagcac   58607
                                                                  ********************* **************************************

chimp.LOC112206744.LOC107973052.GGT2-.646279-708040.rev.compl     gaggaccacggccagcaggcccagcaccactaacttcttcttcatggctctgctgcaccc   59621
FAM230B-LOC105372935-GGT2.NCBI.GGT2.rev.compl                     caggaccacggccagcaggcccagcaccactaacttcttcttcatggctctgctgcaccc   58667
                                                                   ***********************************************************

chimp.LOC112206744.LOC107973052.GGT2-.646279-708040.rev.compl     acggggtaaggagcagggtcaggcccagcctcagacattccctggcccctccccaacagg   59681
FAM230B-LOC105372935-GGT2.NCBI.GGT2.rev.compl                     acggggtaaggagcagggtcaggcccagcctcagacatgccctggcccctccccaacagg   58727
                                                                  ************************************** *********************

chimp.LOC112206744.LOC107973052.GGT2-.646279-708040.rev.compl     gcacagtctaaagtcaggcctcagaaacacaaggcctgtgtctccttcccgcttcccaga   59741
FAM230B-LOC105372935-GGT2.NCBI.GGT2.rev.compl                     gcacagtctaaagtcgggcctcagaaacacaaggccggagtctccttcccgcttcccaga   58787
                                                                  *************** ******************** * *********************

chimp.LOC112206744.LOC107973052.GGT2-.646279-708040.rev.compl     atacgtgcaggctgtccggcccccagacctttgcgcaggccatgccctctgccagaagct   59801
FAM230B-LOC105372935-GGT2.NCBI.GGT2.rev.compl                     atacgtgcaggctgtccggcccccagacctttgcgcaggccatgccctctgccagaagct   58847
                                                                  ************************************************************

chimp.LOC112206744.LOC107973052.GGT2-.646279-708040.rev.compl     ctgggcctcatctctgccctcccaaatcctccctgcttatcttcagagcccatcctggta   59861
FAM230B-LOC105372935-GGT2.NCBI.GGT2.rev.compl                     ctgggcctcatctctgccctcccaaatcctccctgcttatcttcagagcccatcctggta   58907
                                                                  ************************************************************

chimp.LOC112206744.LOC107973052.GGT2-.646279-708040.rev.compl     agaaccccatctccaccagcggcccttcctgggagcccccagatttccacacccctcttt   59921
FAM230B-LOC105372935-GGT2.NCBI.GGT2.rev.compl                     agaaccccatctccagcagcggcccttcctgggagcccccagatttccacacccctcttt   58967
                                                                  *************** ********************************************

chimp.LOC112206744.LOC107973052.GGT2-.646279-708040.rev.compl     ctgcagggcctggcctacctcctcacagtggctgagcctccactgcttagggagaagctc   59981
FAM230B-LOC105372935-GGT2.NCBI.GGT2.rev.compl                     ctgcagggcctggcctacctcctcacagtggctgagcctccactgcttagggagaagctc   59027
                                                                  ************************************************************

chimp.LOC112206744.LOC107973052.GGT2-.646279-708040.rev.compl     cagtagggatgggcctggcctggtttctcctgtgtcccccacctcagcctagagcctggc   60041
FAM230B-LOC105372935-GGT2.NCBI.GGT2.rev.compl                     cagcagggatgggcctggcctggtttctcctgcgtcccccaccccagcctagagcctggc   59087
                                                                  *** **************************** ********** ****************

chimp.LOC112206744.LOC107973052.GGT2-.646279-708040.rev.compl     actgtccaggagtcctctgaagaccctccaccccacctggagcatggggtttagcttcca   60101
FAM230B-LOC105372935-GGT2.NCBI.GGT2.rev.compl                     actgtccaggagtcctctgaagaccctccaccccacctggagcatggggtttagcttcca   59147
                                                                  ************************************************************

chimp.LOC112206744.LOC107973052.GGT2-.646279-708040.rev.compl     tagtgcccacaatcagagcgccccacagattcactgccacggggccaggacttaccgtcc   60161
FAM230B-LOC105372935-GGT2.NCBI.GGT2.rev.compl                     tagtgcccacaatcagagcgccccacagattcactgccacggggccaggacttaccgtcc   59207
                                                                  ************************************************************

chimp.LOC112206744.LOC107973052.GGT2-.646279-708040.rev.compl     agcagcagacaggggccccaagccttgcctggggtgttggccacgaaagacaggaggatt   60221
FAM230B-LOC105372935-GGT2.NCBI.GGT2.rev.compl                     agcagcagacgggggccccaagccttgcctggggtgttggccacgaaagacaggaggatt   59267
                                                                  ********** *************************************************

chimp.LOC112206744.LOC107973052.GGT2-.646279-708040.rev.compl     tggtggaaacagctgaggaaataaccggggtctccctcacactctgctgaagcctgtagc   60281
FAM230B-LOC105372935-GGT2.NCBI.GGT2.rev.compl                     tggtggaaacagctgaggaaataaccggggtctccctcacactctgctgaagcctgtagc   59327
                                                                  ************************************************************

chimp.LOC112206744.LOC107973052.GGT2-.646279-708040.rev.compl     cacagaatcttcttcagagactccctgatcaggcagccttctcgttctcctgaaggtcaa   60341
FAM230B-LOC105372935-GGT2.NCBI.GGT2.rev.compl                     cacagaatcttcttcagagactctctgatcaggcagccttctcgttctcctgaaggtcaa   59387
                                                                  *********************** ************************************

chimp.LOC112206744.LOC107973052.GGT2-.646279-708040.rev.compl     aggaggttacctgaagcacgcacagctcagacctttctggggtactccgtgttacctccc   60401
FAM230B-LOC105372935-GGT2.NCBI.GGT2.rev.compl                     gggaggttacctgaagcacgcacagcccagacctttctgggggactccgtgttacctccc   59447
                                                                   ************************* *************** *****************

chimp.LOC112206744.LOC107973052.GGT2-.646279-708040.rev.compl     tctgcctctagctggtttctctgtctccagttgaactctggaggcaaagaggctgtcagt   60461
FAM230B-LOC105372935-GGT2.NCBI.GGT2.rev.compl                     tctgcctctagctggtttctctgtctccagttgaactctggaggcaaagaggctgtcagt   59507
                                                                  ************************************************************

chimp.LOC112206744.LOC107973052.GGT2-.646279-708040.rev.compl     aacacatttgtttccatgaattctctcagcatgtctcccaggcacagctcaaagagggtt   60521
FAM230B-LOC105372935-GGT2.NCBI.GGT2.rev.compl                     aacacatttgtttccatgaattctctcagcatgtctcccaggcac---------agggtt   59558
                                                                  *********************************************         ******

chimp.LOC112206744.LOC107973052.GGT2-.646279-708040.rev.compl     ttgcacggagcagggcaggtaggggacagggcattcctgcacaagcccaggatgtccatg   60581
FAM230B-LOC105372935-GGT2.NCBI.GGT2.rev.compl                     ttgcacggagcagggcaggtaggggacagggcattcctgcacaagcccaggatgtgcatg   59618
                                                                  ******************************************************* ****

chimp.LOC112206744.LOC107973052.GGT2-.646279-708040.rev.compl     cggtaagcatggcaaagggggctcagggggcaccgccagcctgccctgctctgacgctgg   60641
FAM230B-LOC105372935-GGT2.NCBI.GGT2.rev.compl                     cggtaagcatggcaaagggggctcaggaggcaccgccagcctgccctgctctgacgctgg   59678
                                                                  *************************** ********************************

chimp.LOC112206744.LOC107973052.GGT2-.646279-708040.rev.compl     acttgccactcacctgctgtggggcctcaggcaaatcactgaactgtccagcctggatga   60701
FAM230B-LOC105372935-GGT2.NCBI.GGT2.rev.compl                     acttgccactcacctgctgtggggcctcaggcaaatcactgaactgtccagcctggatga   59738
                                                                  ************************************************************

chimp.LOC112206744.LOC107973052.GGT2-.646279-708040.rev.compl     cggcagcacctcacttgccttgctgctgggagtgttgtgaatagagtaggttagactgtg   60761
FAM230B-LOC105372935-GGT2.NCBI.GGT2.rev.compl                     cggcagcacctcacttgccttgctgctgggagtgttgtgaatagagtaggttagactgtg   59798
                                                                  ************************************************************

chimp.LOC112206744.LOC107973052.GGT2-.646279-708040.rev.compl     ggcagggcttggtgaatggtagctgtgattatcatcatggctgcaccggggacaccccca   60821
FAM230B-LOC105372935-GGT2.NCBI.GGT2.rev.compl                     ggcagggcttggtgaatggtagctgtgattatcatcatggctgcactggggacaccccca   59858
                                                                  ********************************************** *************

chimp.LOC112206744.LOC107973052.GGT2-.646279-708040.rev.compl     ggaggcctgagtggcacaggtctcttgctcactgtatgtcccctgtggactcccttccag   60881
FAM230B-LOC105372935-GGT2.NCBI.GGT2.rev.compl                     ggaggcctgagtggcacaggtctcttgctcactgtatgtcccctgtggactcccttccag   59918
                                                                  ************************************************************

chimp.LOC112206744.LOC107973052.GGT2-.646279-708040.rev.compl     gctgtgcagtgagtggcagcagtgacccttgggaagcctcatggctatggcagcaggtga   60941
FAM230B-LOC105372935-GGT2.NCBI.GGT2.rev.compl                     gctgtgcagtgagtggcagcagtgacccttgggaagtctcatggctacggcagcaggtga   59978
                                                                  ************************************ ********** ************

chimp.LOC112206744.LOC107973052.GGT2-.646279-708040.rev.compl     caggtgtgacaacagggaagaaggatgtggtgacagaggttggggttcccctctcccaca   61001
FAM230B-LOC105372935-GGT2.NCBI.GGT2.rev.compl                     caggtgtgacaacagggaagagggatgtggtgacagaggttggggttcccctctcccaca   60038
                                                                  ********************* **************************************

chimp.LOC112206744.LOC107973052.GGT2-.646279-708040.rev.compl     gtcagtttcccacaaagggcggtgtctgccagcaagcccctccaatgagccccaagctgg   61061
FAM230B-LOC105372935-GGT2.NCBI.GGT2.rev.compl                     gtcagtttcccacaaagggcggtgtctgccagcaagcccctccaatgagccccaagcttg   60098
                                                                  ********************************************************** *

chimp.LOC112206744.LOC107973052.GGT2-.646279-708040.rev.compl     gtttccctccactccaccctgtcccagtgcagagcgtctgacctcagaggcagacacact   61121
```

```
FAM230B-LOC105372935-GGT2.NCBI.GGT2.rev.compl                  gtttccctccactccacgctgtcccagtgcagagcatctgacctcagaggcagacacact   60158
                                                               ***************** ***************** ************************

chimp.LOC112206744.LOC107973052.GGT2-.646279-708040.rev.compl  gtcccagaggtggtctacgaatggagtccctgtgccctccccacacacagggaacatcca   61181
FAM230B-LOC105372935-GGT2.NCBI.GGT2.rev.compl                  gtcccagaggtggtctacgaatggagtccccgtgccctccccacacacagggaacatcca   60218
                                                               ****************************** *****************************

chimp.LOC112206744.LOC107973052.GGT2-.646279-708040.rev.compl  aatgccatcgtggaagggtgg-cacctccccaggcttggtgggcctggggccgatagtgt   61240
FAM230B-LOC105372935-GGT2.NCBI.GGT2.rev.compl                  aatgccatcgtggaagggtggccacctccccaggcttggtgggcctggggccgatagtgt   60278
                                                               ********************* **************************************

chimp.LOC112206744.LOC107973052.GGT2-.646279-708040.rev.compl  gatacatttgacccctcccagccctggatgcagacaccaagagcagagagacctggcag   61300
FAM230B-LOC105372935-GGT2.NCBI.GGT2.rev.compl                  gatacatttgacccctcccagccctggatgcagacaccaagagcagagagacctggcag   60338
                                                               ************************************************************

chimp.LOC112206744.LOC107973052.GGT2-.646279-708040.rev.compl  tagtcatgcagcagcgcaccaccccacatccccagcacaatccaaagcagcccgtcatcc   61360
FAM230B-LOC105372935-GGT2.NCBI.GGT2.rev.compl                  tagccatgcagcagcgcaccaccccacatccccagcacaatccaaagcagcccctcatcc   60398
                                                               *** ************************************************* ******

chimp.LOC112206744.LOC107973052.GGT2-.646279-708040.rev.compl  ccaccgtgaccaccacagcctgaatccaggtgccacctgtttctgacctgaactccctca   61420
FAM230B-LOC105372935-GGT2.NCBI.GGT2.rev.compl                  ccaccgtgaccaccacagcctgaatccaggcgccacctgtttctgacctgaactccctca   60458
                                                               ****************************** *****************************

chimp.LOC112206744.LOC107973052.GGT2-.646279-708040.rev.compl  cagcccctgcctgcactccctccctccaacatcgcctgcccttcagtcttccagaaagca   61480
FAM230B-LOC105372935-GGT2.NCBI.GGT2.rev.compl                  cagcccctgcctgcactccctccctccaacatcacctgcccttcagtcttccagaaagca   60518
                                                               ********************************* **************************

chimp.LOC112206744.LOC107973052.GGT2-.646279-708040.rev.compl  gctagagggctctgtctgtccaactgcagaacaggccctgcctcctccctgtcccgttgg   61540
FAM230B-LOC105372935-GGT2.NCBI.GGT2.rev.compl                  gctagagggctctgtctgtccaactgcagaacaggccctgcctcctccctgccctgttgg   60578
                                                               *************************************************** ** *****

chimp.LOC112206744.LOC107973052.GGT2-.646279-708040.rev.compl  acagctcacacccttcaccaggcctgacagcactcttgccactccaacaccctgggtccc   61600
FAM230B-LOC105372935-GGT2.NCBI.GGT2.rev.compl                  acagctcacacccttcaccaggcctgacagcgctcttgccactccaacaccccgggtccc   60638
                                                               ******************************* ******************** *******

chimp.LOC112206744.LOC107973052.GGT2-.646279-708040.rev.compl  agccgggagtccaggtcagggttaagggttcctgatagagacaccgattcctggaggtcc   61660
FAM230B-LOC105372935-GGT2.NCBI.GGT2.rev.compl                  agccgggagtccgggtcagggttaagggttcctggtagagacaccgattcctggaggtcc   60698
                                                               ************ ********************* *************************

chimp.LOC112206744.LOC107973052.GGT2-.646279-708040.rev.compl  aaagagcctcaggagctgggccagcaatatgcagcatctattatggacacagaacattcc   61720
FAM230B-LOC105372935-GGT2.NCBI.GGT2.rev.compl                  aaagagcctcaggagctgggccagcaatatgcagcatctattatggacacagaacattcc   60758
                                                               ************************************************************

chimp.LOC112206744.LOC107973052.GGT2-.646279-708040.rev.compl  catcacatggccgggtgcagtggctcacgcctataatcccag------------------   61762
FAM230B-LOC105372935-GGT2.NCBI.GGT2.rev.compl                  catcacatggccgggtgcagtggctcacgcctttaatcccagcactttgggaggcagagg   60818
                                                               ******************************** *********

chimp.LOC112206744.LOC107973052.GGT2-.646279-708040.rev.compl  ------------------------------------------------------------   61762
FAM230B-LOC105372935-GGT2.NCBI.GGT2.rev.compl                  caggtggatcacctgaggtgaagagttcgagaccagtctggccaacatggtgaaacccc   60878

chimp.LOC112206744.LOC107973052.GGT2-.646279-708040.rev.compl  ------------------------------------------------------------   61762
FAM230B-LOC105372935-GGT2.NCBI.GGT2.rev.compl                  atctccactaaaaatacaaaaaattagccaggcatggtggcaggtgcctgtaatcccagc   60938

chimp.LOC112206744.LOC107973052.GGT2-.646279-708040.rev.compl  ------------------------------------------------------------   61762
FAM230B-LOC105372935-GGT2.NCBI.GGT2.rev.compl                  tactcaggagctgaggcaggagaattgcttgaacccgggaggtggaggttgcagtgaggc   60998

chimp.LOC112206744.LOC107973052.GGT2-.646279-708040.rev.compl  ------------------------------------------------------------   61762
FAM230B-LOC105372935-GGT2.NCBI.GGT2.rev.compl                  aagattgcaccactgcactccagcctgggccacaagagtgaaactccgtcacacacacac   61058

chimp.LOC112206744.LOC107973052.GGT2-.646279-708040.rev.compl  ------------------------------------------------------------   61762
FAM230B-LOC105372935-GGT2.NCBI.GGT2.rev.compl                  acacacacacacacaaaaaaaaaaaaaaagaaaaaaaaggtctcctgctggaacacagac   61118

chimp.LOC112206744.LOC107973052.GGT2-.646279-708040.rev.compl  ------------------------------------------------------------   61762
FAM230B-LOC105372935-GGT2.NCBI.GGT2.rev.compl                  tagttagagaaaggaaaaataaacaaatgatagtatgtgtattaataaaagaactagcaa   61178

chimp.LOC112206744.LOC107973052.GGT2-.646279-708040.rev.compl  ------------------------------------------------------------   61762
FAM230B-LOC105372935-GGT2.NCBI.GGT2.rev.compl                  cacccactgcttagttgtgataataaaaactggacattaatataggcagaaaaacagaac   61238

chimp.LOC112206744.LOC107973052.GGT2-.646279-708040.rev.compl  ------------------------------------------------------------   61762
FAM230B-LOC105372935-GGT2.NCBI.GGT2.rev.compl                  agccttgatatggttaaccctttgacactggcaaacactgtgaccagtgctgcccacact   61298

chimp.LOC112206744.LOC107973052.GGT2-.646279-708040.rev.compl  ------------------------------------------------------------   61762
FAM230B-LOC105372935-GGT2.NCBI.GGT2.rev.compl                  ggaagctcctgcatccctcttcctcatgtttccttcagcattaaggagcaacaagggaga   61358

chimp.LOC112206744.LOC107973052.GGT2-.646279-708040.rev.compl  ------------------------------------------------------------   61762
FAM230B-LOC105372935-GGT2.NCBI.GGT2.rev.compl                  cagccagttcatagttccttacgcatggagccaaaggaccttcaatgtacaaggtctgag   61418

chimp.LOC112206744.LOC107973052.GGT2-.646279-708040.rev.compl  ------------------------------------------------------------   61762
FAM230B-LOC105372935-GGT2.NCBI.GGT2.rev.compl                  caaggacccgcagccacatgtgcttcctgcttcagcagtgccccgtgggtctcagagcta   61478

chimp.LOC112206744.LOC107973052.GGT2-.646279-708040.rev.compl  ------------------------------------------------------------   61762
FAM230B-LOC105372935-GGT2.NCBI.GGT2.rev.compl                  cccaaaggcctatccttccgagaggtttctcctctctcccaactgatgtcatacgtctca   61538

chimp.LOC112206744.LOC107973052.GGT2-.646279-708040.rev.compl  ------------------------------------------------------------   61762
FAM230B-LOC105372935-GGT2.NCBI.GGT2.rev.compl                  tcttccttgtcattcagatcatcaaccacaaacgtccttgccttatattttcatatccct   61598

chimp.LOC112206744.LOC107973052.GGT2-.646279-708040.rev.compl  ------------------------------------------------------------   61762
FAM230B-LOC105372935-GGT2.NCBI.GGT2.rev.compl                  tttcaccagttacagggttagttaacttacgaaattcttaacatctgcattagatctttt   61658

chimp.LOC112206744.LOC107973052.GGT2-.646279-708040.rev.compl  ------------------------------------------------------------   61762
FAM230B-LOC105372935-GGT2.NCBI.GGT2.rev.compl                  taaagtttcaccctcaaccacctattatttagaagtgaacacagaaatttagtttccttg   61718

chimp.LOC112206744.LOC107973052.GGT2-.646279-708040.rev.compl  ------------------------------------------------------------   61762
FAM230B-LOC105372935-GGT2.NCBI.GGT2.rev.compl                  tgctatctgttgacccctaaaatatgctgggagttttgggatttttttaaagtcaaatg   61778

chimp.LOC112206744.LOC107973052.GGT2-.646279-708040.rev.compl  ------------------------------------------------------------   61762
FAM230B-LOC105372935-GGT2.NCBI.GGT2.rev.compl                  catggcataaagcaaaattacactactaaagaactgagtcaggccagacgctggcaacgt   61838

chimp.LOC112206744.LOC107973052.GGT2-.646279-708040.rev.compl  ------------------------------------------------------------   61762
FAM230B-LOC105372935-GGT2.NCBI.GGT2.rev.compl                  gaagacagcttctccttacccactaggttcatctttgcactgttaactcccaaagtattg   61898

chimp.LOC112206744.LOC107973052.GGT2-.646279-708040.rev.compl  ------------------------------------------------------------   61762
FAM230B-LOC105372935-GGT2.NCBI.GGT2.rev.compl                  cttctcggtgctgggtgtgtggcattcatattccccggcatcccgggcctgaagatctgt   61958

chimp.LOC112206744.LOC107973052.GGT2-.646279-708040.rev.compl  ------------------------------------------------------------   61762
FAM230B-LOC105372935-GGT2.NCBI.GGT2.rev.compl                  gatgtgcaatagggttgggttcccctggactctttctatgaagatcttccctccgcggac   62018

chimp.LOC112206744.LOC107973052.GGT2-.646279-708040.rev.compl  ------------------------------------------------------------   61762
FAM230B-LOC105372935-GGT2.NCBI.GGT2.rev.compl                  gcgctgggtgtagatggcataggggaaggaagagtccatggtgctgacgatctgcacctc   62078

chimp.LOC112206744.LOC107973052.GGT2-.646279-708040.rev.compl  ------------------------------------------------------------   61762
FAM230B-LOC105372935-GGT2.NCBI.GGT2.rev.compl                  tcgctctggcgacgaaggcaggtaaatggaccactggaaattctgttcagaaggtccctg   62138

chimp.LOC112206744.LOC107973052.GGT2-.646279-708040.rev.compl  ------------------------------------------------------------   61762
FAM230B-LOC105372935-GGT2.NCBI.GGT2.rev.compl                  gtagccactcacattgcaccagatagtgatgtgggagccctccgtgcggtacaagggtcc   62198

chimp.LOC112206744.LOC107973052.GGT2-.646279-708040.rev.compl  ------------------------------------------------------------   61762
FAM230B-LOC105372935-GGT2.NCBI.GGT2.rev.compl                  ttcctgaacggtgacctgccgctgtgctgacaccacacctacgagggagagaaacacacg   62258

chimp.LOC112206744.LOC107973052.GGT2-.646279-708040.rev.compl  ------------------------------------------------------------   61762
FAM230B-LOC105372935-GGT2.NCBI.GGT2.rev.compl                  caacatgctcacttacttctcagaccaaaatgcaaagtaggcagcaatctccaaagggtt   62318
```

```
chimp.LOC112206744.LOC107973052.GGT2-.646279-708040.rev.compl     ------------------------------------------------------------   61762
FAM230B-LOC105372935-GGT2.NCBI.GGT2.rev.compl                     tgttatttgtgtgacatgataataatataaacagcttactggccctttcaaaggcactct   62378

chimp.LOC112206744.LOC107973052.GGT2-.646279-708040.rev.compl     ------------------------------------------------------------   61762
FAM230B-LOC105372935-GGT2.NCBI.GGT2.rev.compl                     gtgatttataaatattaattaaaacactctgtggtaaagcactgtccctaatttgcatat   62438

chimp.LOC112206744.LOC107973052.GGT2-.646279-708040.rev.compl     ------------------------------------------------------------   61762
FAM230B-LOC105372935-GGT2.NCBI.GGT2.rev.compl                     atgggaatttgatcatgccaagatgtgcttttgttacttattcaacagctgacccttggg   62498

chimp.LOC112206744.LOC107973052.GGT2-.646279-708040.rev.compl     ------------------------------------------------------------   61762
FAM230B-LOC105372935-GGT2.NCBI.GGT2.rev.compl                     aatttcatttagcccctctgtttgtcatgtatacggcagttttatatctatttattaatg   62558

chimp.LOC112206744.LOC107973052.GGT2-.646279-708040.rev.compl     ------------------------------------------------------------   61762
FAM230B-LOC105372935-GGT2.NCBI.GGT2.rev.compl                     ctgtgtataaagtttttaataaaataagcaggaaaagttgaatccttggcacaaattcag   62618

chimp.LOC112206744.LOC107973052.GGT2-.646279-708040.rev.compl     ------------------------------------------------------------   61762
FAM230B-LOC105372935-GGT2.NCBI.GGT2.rev.compl                     atgaaaacaaataaaagcaaccatctaggaatctggctgaaattaaatgcttcttccttg   62678

chimp.LOC112206744.LOC107973052.GGT2-.646279-708040.rev.compl     ------------------------------------------------------------   61762
FAM230B-LOC105372935-GGT2.NCBI.GGT2.rev.compl                     gtccgagtgctggtgggagggagtcctgggagcaccttttactggtgctgcacccttca   62738

chimp.LOC112206744.LOC107973052.GGT2-.646279-708040.rev.compl     ------------------------------------------------------------   61762
FAM230B-LOC105372935-GGT2.NCBI.GGT2.rev.compl                     ttttcccatgtagctgctccccaacaccaccccaaggagcattccttagtttctcttccc   62798

chimp.LOC112206744.LOC107973052.GGT2-.646279-708040.rev.compl     ------------------------------------------------------------   61762
FAM230B-LOC105372935-GGT2.NCBI.GGT2.rev.compl                     tccataaatggaggcagagctgagctcctgcaggctgctatcgttttaagctgagcacac   62858

chimp.LOC112206744.LOC107973052.GGT2-.646279-708040.rev.compl     ------------------------------------------------------------   61762
FAM230B-LOC105372935-GGT2.NCBI.GGT2.rev.compl                     ccaaatacaaatgtgaagggctgaaacagatccctgatgcccagattcccacaaactgac   62918

chimp.LOC112206744.LOC107973052.GGT2-.646279-708040.rev.compl     ------------------------------------------------------------   61762
FAM230B-LOC105372935-GGT2.NCBI.GGT2.rev.compl                     aaagggggaactaacagtgcccagctaaacctgtgacaaggacagtcaggctacccttg   62978

chimp.LOC112206744.LOC107973052.GGT2-.646279-708040.rev.compl     ------------------------------------------------------------   61762
FAM230B-LOC105372935-GGT2.NCBI.GGT2.rev.compl                     agtttcccactcaagactcccccgctgacctcctgccttctggctcagatactgttctcc   63038

chimp.LOC112206744.LOC107973052.GGT2-.646279-708040.rev.compl     ------------------------------------------------------------   61762
FAM230B-LOC105372935-GGT2.NCBI.GGT2.rev.compl                     aaggtcacttctgcctgccttctccatgttcttcttaaagtggcatggtgagtttaattt   63098

chimp.LOC112206744.LOC107973052.GGT2-.646279-708040.rev.compl     ------------------------------------------------------------   61762
FAM230B-LOC105372935-GGT2.NCBI.GGT2.rev.compl                     ctcatttcgtgccaactcaacattgattgtgcctgagccactcagaaggtgggagggggc   63158

chimp.LOC112206744.LOC107973052.GGT2-.646279-708040.rev.compl     ------------------------------------------------------------   61762
FAM230B-LOC105372935-GGT2.NCBI.GGT2.rev.compl                     acacccctgcctcttcatttcggcagtggtcacgtactgcaggttgtgaaaagcgcaggt   63218

chimp.LOC112206744.LOC107973052.GGT2-.646279-708040.rev.compl     ------------------------------------------------------------   61762
FAM230B-LOC105372935-GGT2.NCBI.GGT2.rev.compl                     gagttataggcgcagctttgccaccaattagagctggttgtgtcatctcagacaacttgc   63278

chimp.LOC112206744.LOC107973052.GGT2-.646279-708040.rev.compl     ------------------------------------------------------------   61762
FAM230B-LOC105372935-GGT2.NCBI.GGT2.rev.compl                     ttttcttctccagaccttgatcttctcatctatcacatgatttctaagaacgctttagag   63338

chimp.LOC112206744.LOC107973052.GGT2-.646279-708040.rev.compl     ------------------------------------------------------------   61762
FAM230B-LOC105372935-GGT2.NCBI.GGT2.rev.compl                     agcactagcatgggttccgatgctggctttgtcacttattaactgtgatcttggataagc   63398

chimp.LOC112206744.LOC107973052.GGT2-.646279-708040.rev.compl     ------------------------------------------------------------   61762
FAM230B-LOC105372935-GGT2.NCBI.GGT2.rev.compl                     tgcttaacctctctaagcctcagttttctcatcggtaagatggggataataatgcctgtc   63458

chimp.LOC112206744.LOC107973052.GGT2-.646279-708040.rev.compl     ------------------------------------------------------------   61762
FAM230B-LOC105372935-GGT2.NCBI.GGT2.rev.compl                     tcatagggttattgagaagatttagtaacatctgtaaagaccctagcacagagcctagcc   63518

chimp.LOC112206744.LOC107973052.GGT2-.646279-708040.rev.compl     ------------------------------------------------------------   61762
FAM230B-LOC105372935-GGT2.NCBI.GGT2.rev.compl                     cacaatcagtcctcaataaagagctgctggctaagacatatttctgctggaacattctgt   63578

chimp.LOC112206744.LOC107973052.GGT2-.646279-708040.rev.compl     ------------------------------------------------------------   61762
FAM230B-LOC105372935-GGT2.NCBI.GGT2.rev.compl                     ggctctggatccatagaggacaggatttggccaatgactgcttaaggacatttcaacggc   63638

chimp.LOC112206744.LOC107973052.GGT2-.646279-708040.rev.compl     ------------------------------------------------------------   61762
FAM230B-LOC105372935-GGT2.NCBI.GGT2.rev.compl                     cccttgacaatgacaccatcactgggctgcccacctcatcccacagctgcagcccattct   63698

chimp.LOC112206744.LOC107973052.GGT2-.646279-708040.rev.compl     ------------------------------------------------------------   61762
FAM230B-LOC105372935-GGT2.NCBI.GGT2.rev.compl                     tgtgatagcttcccttttcccccacaaatgggaatggcggctgccctggctatgcctatc   63758

chimp.LOC112206744.LOC107973052.GGT2-.646279-708040.rev.compl     ------------------------------------------------------------   61762
FAM230B-LOC105372935-GGT2.NCBI.GGT2.rev.compl                     tgaagtggatccagctggtgaacagccgggacagctcaaatactagtggagtgcagctct   63818

chimp.LOC112206744.LOC107973052.GGT2-.646279-708040.rev.compl     ------------------------------------------------------------   61762
FAM230B-LOC105372935-GGT2.NCBI.GGT2.rev.compl                     ctgcccagcccagcaggtgactgtgcctcagtgtaagctaatctaatctgcaagatgaca   63878

chimp.LOC112206744.LOC107973052.GGT2-.646279-708040.rev.compl     ------------------------------------------------------------   61762
FAM230B-LOC105372935-GGT2.NCBI.GGT2.rev.compl                     gacctcaaaacggcccatacaatgaaatgcccatggcttgcaaaactttaattcatgcag   63938

chimp.LOC112206744.LOC107973052.GGT2-.646279-708040.rev.compl     ------------------------------------------------------------   61762
FAM230B-LOC105372935-GGT2.NCBI.GGT2.rev.compl                     aacctttcctcacaaacaaaatcttacacagcaccccatcttggcttcagcagaaccagg   63998

chimp.LOC112206744.LOC107973052.GGT2-.646279-708040.rev.compl     ------------------------------------------------------------   61762
FAM230B-LOC105372935-GGT2.NCBI.GGT2.rev.compl                     aaaaataaaaaattggcaggggaaggaaataaacagaacaagtggtatctgaatttattt   64058

chimp.LOC112206744.LOC107973052.GGT2-.646279-708040.rev.compl     ------------------------------------------------------------   61762
FAM230B-LOC105372935-GGT2.NCBI.GGT2.rev.compl                     gatatattctggttgatggcattaattatgacatttaaagtcaccatgaggacaactcat   64118

chimp.LOC112206744.LOC107973052.GGT2-.646279-708040.rev.compl     ------------------------------------------------------------   61762
FAM230B-LOC105372935-GGT2.NCBI.GGT2.rev.compl                     tagtaatatcagtatgttaaaatatggtgcataacatttttaatatatgtacagtcatgc   64178

chimp.LOC112206744.LOC107973052.GGT2-.646279-708040.rev.compl     ------------------------------------------------------------   61762
FAM230B-LOC105372935-GGT2.NCBI.GGT2.rev.compl                     actgaataacatttcagtcaatgaggaaacacatgtgcaacagtgatcctgtaagattat   64238

chimp.LOC112206744.LOC107973052.GGT2-.646279-708040.rev.compl     ------------------------------------------------------------   61762
FAM230B-LOC105372935-GGT2.NCBI.GGT2.rev.compl                     aatggagcatatatagagatctaatatatggcacttaatgttggcatggcagatcaagta   64298

chimp.LOC112206744.LOC107973052.GGT2-.646279-708040.rev.compl     ------------------------------------------------------------   61762
FAM230B-LOC105372935-GGT2.NCBI.GGT2.rev.compl                     ggggaaatgactgatatttagtaacagtgctgggacatttgattttccataataaaatat   64358

chimp.LOC112206744.LOC107973052.GGT2-.646279-708040.rev.compl     ------------------------------------------------------------   61762
FAM230B-LOC105372935-GGT2.NCBI.GGT2.rev.compl                     ataaatgaaaatatatatcccatctaggtttgttgaaatacaccctatgatgttcacaca   64418

chimp.LOC112206744.LOC107973052.GGT2-.646279-708040.rev.compl     ------------------------------------------------------------   61762
FAM230B-LOC105372935-GGT2.NCBI.GGT2.rev.compl                     agaatgaaattgcctaatgatgcatttcttaaaacatgtccccatcattaagtgacccat   64478
```

```
chimp.LOC112206744.LOC107973052.GGT2-.646279-708040.rev.compl      ------------------------------------------------------------   61762
FAM230B-LOC105372935-GGT2.NCBI.GGT2.rev.compl                      gactgtatacacacacacatatggtgacacttaaatcaaatgattgcagtattcctgaaa   64538

chimp.LOC112206744.LOC107973052.GGT2-.646279-708040.rev.compl      ------------------------------------------------------------   61762
FAM230B-LOC105372935-GGT2.NCBI.GGT2.rev.compl                      cacagaacattttgcaaacaatttacctacattcataacattaggcatccttagaattgc   64598

chimp.LOC112206744.LOC107973052.GGT2-.646279-708040.rev.compl      ------------------------------------------------------------   61762
FAM230B-LOC105372935-GGT2.NCBI.GGT2.rev.compl                      agtgttctatgacagagcaactacaaccaacccccatccatctcctgcaaagaaggctgg   64658

chimp.LOC112206744.LOC107973052.GGT2-.646279-708040.rev.compl      ------------------------------------------------------------   61762
FAM230B-LOC105372935-GGT2.NCBI.GGT2.rev.compl                      aggcaggtcagggtacaccagcatcttcaagaactgcttctcaatcctggacctacactg   64718

chimp.LOC112206744.LOC107973052.GGT2-.646279-708040.rev.compl      ------------------------------------------------------------   61762
FAM230B-LOC105372935-GGT2.NCBI.GGT2.rev.compl                      gaatcacctggggagctttaaaaaaaataacagtgcctggaccccacctgacataactgg   64778

chimp.LOC112206744.LOC107973052.GGT2-.646279-708040.rev.compl      ------------------------------------------------------------   61762
FAM230B-LOC105372935-GGT2.NCBI.GGT2.rev.compl                      tcttaggattaattagggtttttttgaagctccacagatgattcaaatcaggtgtagca   64838

chimp.LOC112206744.LOC107973052.GGT2-.646279-708040.rev.compl      ------------------------------------------------------------   61762
FAM230B-LOC105372935-GGT2.NCBI.GGT2.rev.compl                      aagcactgttactttaaagtgtctctacctacatggtgccagccaagtgctcaaatgaaa   64898

chimp.LOC112206744.LOC107973052.GGT2-.646279-708040.rev.compl      ------------------------------------------------------------   61762
FAM230B-LOC105372935-GGT2.NCBI.GGT2.rev.compl                      tatcttaaggctctcactggctttaagtttctctctttggtagagacccaatcactggtt   64958

chimp.LOC112206744.LOC107973052.GGT2-.646279-708040.rev.compl      ------------------------------------------------------------   61762
FAM230B-LOC105372935-GGT2.NCBI.GGT2.rev.compl                      caaggaagtttcattctcctccagtcttccccagtgcaaaagaaaacagctgagacccat   65018

chimp.LOC112206744.LOC107973052.GGT2-.646279-708040.rev.compl      ------------------------------------------------------------   61762
FAM230B-LOC105372935-GGT2.NCBI.GGT2.rev.compl                      cagatactggcttttgtgatgcaataatgagtttaaccagaatgcatccgatttacagac   65078

chimp.LOC112206744.LOC107973052.GGT2-.646279-708040.rev.compl      ------------------------------------------------------------   61762
FAM230B-LOC105372935-GGT2.NCBI.GGT2.rev.compl                      cttacaaaggcactaggccagcggatcatgcacgtcctccccacccaaaggtaaacagtg   65138

chimp.LOC112206744.LOC107973052.GGT2-.646279-708040.rev.compl      ------------------------------------------------------------   61762
FAM230B-LOC105372935-GGT2.NCBI.GGT2.rev.compl                      ttaagtggcctgaatgagccaggacagcagggtcaaatcaacttgtctaggctggaagca   65198

chimp.LOC112206744.LOC107973052.GGT2-.646279-708040.rev.compl      ------------------------------------------------------------   61762
FAM230B-LOC105372935-GGT2.NCBI.GGT2.rev.compl                      ggttataaacaatccggacaaataaataactactggattgcactttaaccacacacattc   65258

chimp.LOC112206744.LOC107973052.GGT2-.646279-708040.rev.compl      ------------------------------------------------------------   61762
FAM230B-LOC105372935-GGT2.NCBI.GGT2.rev.compl                      agctgcacctgtttattaaatacctccttaatttccctctgccccacaaagggcttctga   65318

chimp.LOC112206744.LOC107973052.GGT2-.646279-708040.rev.compl      ------------------------------------------------------------   61762
FAM230B-LOC105372935-GGT2.NCBI.GGT2.rev.compl                      cccctgaaaataatgcttctcaacataaaaagaattgttttcttcttggcagtttaattc   65378

chimp.LOC112206744.LOC107973052.GGT2-.646279-708040.rev.compl      ------------------------------------------------------------   61762
FAM230B-LOC105372935-GGT2.NCBI.GGT2.rev.compl                      cccttccacacatcccaccccaccatttccctataatggtaataccccgctgtgcagacc   65438

chimp.LOC112206744.LOC107973052.GGT2-.646279-708040.rev.compl      ------------------------------------------------------------   61762
FAM230B-LOC105372935-GGT2.NCBI.GGT2.rev.compl                      tctgctgcctccagagagacacaggcgcccaaccctgaccagggcatcctctgacccaca   65498

chimp.LOC112206744.LOC107973052.GGT2-.646279-708040.rev.compl      ------------------------------------------------------------   61762
FAM230B-LOC105372935-GGT2.NCBI.GGT2.rev.compl                      gcccctctatctcccctctcagcctacaatagaaagttcctcccaggcaaggatcatttt   65558

chimp.LOC112206744.LOC107973052.GGT2-.646279-708040.rev.compl      ------------------------------------------------------------   61762
FAM230B-LOC105372935-GGT2.NCBI.GGT2.rev.compl                      ttttaaataactttttaactttagacattctgtactctttggattactctgcaataagc   65618

chimp.LOC112206744.LOC107973052.GGT2-.646279-708040.rev.compl      ------------------------------------------------------------   61762
FAM230B-LOC105372935-GGT2.NCBI.GGT2.rev.compl                      aaatatgacttctgtaacataaaaagaaagatcaaaatgttatatataactgcatgaaaa   65678

chimp.LOC112206744.LOC107973052.GGT2-.646279-708040.rev.compl      ------------------------------------------------------------   61762
FAM230B-LOC105372935-GGT2.NCBI.GGT2.rev.compl                      gaactagaaagaaaatactaggtgacaggatggcaagtgattttatttttttcttaata   65738

chimp.LOC112206744.LOC107973052.GGT2-.646279-708040.rev.compl      ------------------------------------------------------------   61762
FAM230B-LOC105372935-GGT2.NCBI.GGT2.rev.compl                      ttttactccttcccccagtcctctgcaatgagtatgtaccagttttataactagaaaaaa   65798

chimp.LOC112206744.LOC107973052.GGT2-.646279-708040.rev.compl      ------------------------------------------------------------   61762
FAM230B-LOC105372935-GGT2.NCBI.GGT2.rev.compl                      attttagccaaaaagaaaaataatgcatgtttgctatataaaattcactactgtagtata   65858

chimp.LOC112206744.LOC107973052.GGT2-.646279-708040.rev.compl      ------------------------------------------------------------   61762
FAM230B-LOC105372935-GGT2.NCBI.GGT2.rev.compl                      tatatatacatacatacatatttggtctttgtccccagttcctggcactgagctcctaaa   65918

chimp.LOC112206744.LOC107973052.GGT2-.646279-708040.rev.compl      ------------------------------------------------------------   61762
FAM230B-LOC105372935-GGT2.NCBI.GGT2.rev.compl                      ccccttggaacttcctaagcaatgggagtaccttttgttatttataagaagccccttttg   65978

chimp.LOC112206744.LOC107973052.GGT2-.646279-708040.rev.compl      ------------------------------------------------------------   61762
FAM230B-LOC105372935-GGT2.NCBI.GGT2.rev.compl                      gccatcccagagtttatgctaacgaggtgactgaaggtgagcacggaggcagtttcaagg   66038

chimp.LOC112206744.LOC107973052.GGT2-.646279-708040.rev.compl      ------------------------------------------------------------   61762
FAM230B-LOC105372935-GGT2.NCBI.GGT2.rev.compl                      aaggagctggccacgcttagaattgtggagttttcagccccgcccttagacctccaagga   66098

chimp.LOC112206744.LOC107973052.GGT2-.646279-708040.rev.compl      ------------------------------------------------------------   61762
FAM230B-LOC105372935-GGT2.NCBI.GGT2.rev.compl                      caagagggggaccgcagattgatccaatcagcattggtcagtgatttgatcagtcatacc   66158

chimp.LOC112206744.LOC107973052.GGT2-.646279-708040.rev.compl      ------------------------------------------------------------   61762
FAM230B-LOC105372935-GGT2.NCBI.GGT2.rev.compl                      cacgtaatgaaatcccatatgaaaaccctaaataatggagttaggagagcttctgggtgc   66218

chimp.LOC112206744.LOC107973052.GGT2-.646279-708040.rev.compl      ------------------------------------------------------------   61762
FAM230B-LOC105372935-GGT2.NCBI.GGT2.rev.compl                      tggaagcggcgtgcaccaggagagggcatgggcagtgagcactactcccctctcagacct   66278

chimp.LOC112206744.LOC107973052.GGT2-.646279-708040.rev.compl      ------------------------------------------------------------   61762
FAM230B-LOC105372935-GGT2.NCBI.GGT2.rev.compl                      tgccctatgcactaaataggactgacctatgtgaccaataggaaatgcacaaatggtgga   66338

chimp.LOC112206744.LOC107973052.GGT2-.646279-708040.rev.compl      ------------------------------------------------------------   61762
FAM230B-LOC105372935-GGT2.NCBI.GGT2.rev.compl                      atgtgacttccagggctaggtcataaaaagacaagcacgatgttatgaggacatcaaagc   66398

chimp.LOC112206744.LOC107973052.GGT2-.646279-708040.rev.compl      ------------------------------------------------------------   61762
FAM230B-LOC105372935-GGT2.NCBI.GGT2.rev.compl                      agccatatggggaggaccccatgaggcctcctgccagcagctagcactaacttggctagc   66458

chimp.LOC112206744.LOC107973052.GGT2-.646279-708040.rev.compl      ------------------------------------------------------------   61762
FAM230B-LOC105372935-GGT2.NCBI.GGT2.rev.compl                      atgtgactggaagtagattctccagcctcagtcaactcagtcaagccttcagataatgtc   66518

chimp.LOC112206744.LOC107973052.GGT2-.646279-708040.rev.compl      ------------------------------------------------------------   61762
FAM230B-LOC105372935-GGT2.NCBI.GGT2.rev.compl                      aaccccaggcatctttttgttgttgttgagacagggtctcactctgtcacccaggctgga   66578

chimp.LOC112206744.LOC107973052.GGT2-.646279-708040.rev.compl      ------------------------------------------------------------   61762
FAM230B-LOC105372935-GGT2.NCBI.GGT2.rev.compl                      gtgcagtggtgcaatcatagctcactgcagccttgacctctgaggctcaagtgaccctcc   66638
```

```
chimp.LOC112206744.LOC107973052.GGT2-.646279-708040.rev.compl      ------------------------------------------------------------   61762
FAM230B-LOC105372935-GGT2.NCBI.GGT2.rev.compl                      cacatcagcctcctgagtatctgggaccaccagcacacaccaccatgcctggctaacttt   66698

chimp.LOC112206744.LOC107973052.GGT2-.646279-708040.rev.compl      ------------------------------------------------------------   61762
FAM230B-LOC105372935-GGT2.NCBI.GGT2.rev.compl                      tttatttcttatagagatgggatgatatggttggggtctgtgtacccacctaaatctcac   66758

chimp.LOC112206744.LOC107973052.GGT2-.646279-708040.rev.compl      ------------------------------------------------------------   61762
FAM230B-LOC105372935-GGT2.NCBI.GGT2.rev.compl                      gttcaattataatccccaatgttagaggtggggcctggtgggaggtgactggatcacagg   66818

chimp.LOC112206744.LOC107973052.GGT2-.646279-708040.rev.compl      ------------------------------------------------------------   61762
FAM230B-LOC105372935-GGT2.NCBI.GGT2.rev.compl                      aatggatctttcatgaatgatttagcatcacccttttggtgctgtgcttgttagagttct   66878

chimp.LOC112206744.LOC107973052.GGT2-.646279-708040.rev.compl      ------------------------------------------------------------   61762
FAM230B-LOC105372935-GGT2.NCBI.GGT2.rev.compl                      cccagtatctaattgtttaaaagtgtgtggcacctccccctctctctcttgctcctgct   66938

chimp.LOC112206744.LOC107973052.GGT2-.646279-708040.rev.compl      ------------------------------------------------------------   61762
FAM230B-LOC105372935-GGT2.NCBI.GGT2.rev.compl                      ttggccatgtaaggcgtgcctgcttccccttcaccttccaccatgattgaaagtttcctg   66998

chimp.LOC112206744.LOC107973052.GGT2-.646279-708040.rev.compl      ------------------------------------------------------------   61762
FAM230B-LOC105372935-GGT2.NCBI.GGT2.rev.compl                      tgcttgccccagaagctgagcagatgccagcattatgctccctgaacagcctgtggaact   67058

chimp.LOC112206744.LOC107973052.GGT2-.646279-708040.rev.compl      ------------------------------------------------------------   61762
FAM230B-LOC105372935-GGT2.NCBI.GGT2.rev.compl                      gcgagacaattaaacctcttttctttataaattctccagtctcatgtatttatagcaatg   67118

chimp.LOC112206744.LOC107973052.GGT2-.646279-708040.rev.compl      ------------------------------------------------------------   61762
FAM230B-LOC105372935-GGT2.NCBI.GGT2.rev.compl                      tgagaactgactaatacatagggtctcactgtgtttcccaggctggtctcaaactcctgg   67178

chimp.LOC112206744.LOC107973052.GGT2-.646279-708040.rev.compl      ------------------------------------------------------------   61762
FAM230B-LOC105372935-GGT2.NCBI.GGT2.rev.compl                      gctcaagtaatcctcctggctttgcttcccagagtgtagagattacaggcatgagccact   67238

chimp.LOC112206744.LOC107973052.GGT2-.646279-708040.rev.compl      ------------------------------------------------------------   61762
FAM230B-LOC105372935-GGT2.NCBI.GGT2.rev.compl                      gaacctgcccctttcccccaacaacatcttgattgcaacctcctaagagactccgagtc   67298

chimp.LOC112206744.LOC107973052.GGT2-.646279-708040.rev.compl      ------------------------------------------------------------   61762
FAM230B-LOC105372935-GGT2.NCBI.GGT2.rev.compl                      agaagtatccagttaagatgttactgactttctgaactacagaaactgtgagataataaa   67358

chimp.LOC112206744.LOC107973052.GGT2-.646279-708040.rev.compl      ------------------------------------------------------------   61762
FAM230B-LOC105372935-GGT2.NCBI.GGT2.rev.compl                      catctattgttcctttaagccttaagtttggggctttttttttttacaccaatatataac   67418

chimp.LOC112206744.LOC107973052.GGT2-.646279-708040.rev.compl      ------------------------------------------------------------   61762
FAM230B-LOC105372935-GGT2.NCBI.GGT2.rev.compl                      taatgtagacacctgcagtccctaagattgaatgcaaaattgtgttacatacacatgagt   67478

chimp.LOC112206744.LOC107973052.GGT2-.646279-708040.rev.compl      ------------------------------------------------------------   61762
FAM230B-LOC105372935-GGT2.NCBI.GGT2.rev.compl                      gtgttcttctgggaaaagagtccataaacttcaaaagattctcaaaggaatccaagaccc   67538

chimp.LOC112206744.LOC107973052.GGT2-.646279-708040.rev.compl      ------------------------------------------------------------   61762
FAM230B-LOC105372935-GGT2.NCBI.GGT2.rev.compl                      aaaaagattaagaaacattgcatcagacaataaattgtcaatgagcaaggtcctgtcctt   67598

chimp.LOC112206744.LOC107973052.GGT2-.646279-708040.rev.compl      ------------------------------------------------------------   61762
FAM230B-LOC105372935-GGT2.NCBI.GGT2.rev.compl                      ttagattgggttttgaaatcacccatcccacaaatatatgataccacttaactagttttt   67658

chimp.LOC112206744.LOC107973052.GGT2-.646279-708040.rev.compl      ------------------------------------------------------------   61762
FAM230B-LOC105372935-GGT2.NCBI.GGT2.rev.compl                      caagatattccttgttcaaagatgctcccacttttttctttgtgctcaagtcctagataaa   67718

chimp.LOC112206744.LOC107973052.GGT2-.646279-708040.rev.compl      ------------------------------------------------------------   61762
FAM230B-LOC105372935-GGT2.NCBI.GGT2.rev.compl                      cttggctaggggcaggaggtatgattcaggagttagtgagccgggagggctcactgtgcc   67778

chimp.LOC112206744.LOC107973052.GGT2-.646279-708040.rev.compl      ------------------------------------------------------------   61762
FAM230B-LOC105372935-GGT2.NCBI.GGT2.rev.compl                      tgggaaagccagcttctcatgaatttttggaatatgccttggctcagaaacctacctctc   67838

chimp.LOC112206744.LOC107973052.GGT2-.646279-708040.rev.compl      ------------------------------------------------------------   61762
FAM230B-LOC105372935-GGT2.NCBI.GGT2.rev.compl                      taggcacaggatccaacttatacgttcaagatgcgtgactgagcatgtgaaagtgtcaga   67898

chimp.LOC112206744.LOC107973052.GGT2-.646279-708040.rev.compl      ------------------------------------------------------------   61762
FAM230B-LOC105372935-GGT2.NCBI.GGT2.rev.compl                      gggatcgacctcaccattttacatctacttttccagttgtggcccctacaatcatcaagg   67958

chimp.LOC112206744.LOC107973052.GGT2-.646279-708040.rev.compl      ------------------------------------------------------------   61762
FAM230B-LOC105372935-GGT2.NCBI.GGT2.rev.compl                      ctcattcggcaccatgctcctccaaattctgcatgcgactcagccacactaatatctttc   68018

chimp.LOC112206744.LOC107973052.GGT2-.646279-708040.rev.compl      ------------------------------------------------------------   61762
FAM230B-LOC105372935-GGT2.NCBI.GGT2.rev.compl                      ttcatctataagatgaaagtaatacaacgcaccatgtagcaaactcacaggagaccctag   68078

chimp.LOC112206744.LOC107973052.GGT2-.646279-708040.rev.compl      ------------------------------------------------------------   61762
FAM230B-LOC105372935-GGT2.NCBI.GGT2.rev.compl                      agatcaaataacactgtgaaagccaccaaatgaatattttaaaattttatctcatttctg   68138

chimp.LOC112206744.LOC107973052.GGT2-.646279-708040.rev.compl      ------------------------------------------------------------   61762
FAM230B-LOC105372935-GGT2.NCBI.GGT2.rev.compl                      ccaaggtgagatatcaatctaaaaacaggataaatctttgtaagtacaagaacattaaaa   68198

chimp.LOC112206744.LOC107973052.GGT2-.646279-708040.rev.compl      ------------------------------------------------------------   61762
FAM230B-LOC105372935-GGT2.NCBI.GGT2.rev.compl                      agaacattaagtttgggttaaaaatgcaaaactgcatatgcaggaaaattacgttaaaaa   68258

chimp.LOC112206744.LOC107973052.GGT2-.646279-708040.rev.compl      ------------------------------------------------------------   61762
FAM230B-LOC105372935-GGT2.NCBI.GGT2.rev.compl                      aaaaaaaatcctacacctacaaaaacaaccaaacaaaacattgatgatactgggctgatt   68318

chimp.LOC112206744.LOC107973052.GGT2-.646279-708040.rev.compl      ------------------------------------------------------------   61762
FAM230B-LOC105372935-GGT2.NCBI.GGT2.rev.compl                      cactgaatggtgagaatatgagtgattttctcctctcctatttttctcaatacctttttct   68378

chimp.LOC112206744.LOC107973052.GGT2-.646279-708040.rev.compl      ------------------------------------------------------------   61762
FAM230B-LOC105372935-GGT2.NCBI.GGT2.rev.compl                      ctattatctcctttttttaaataaacaagtatatattgttctaaaccgaagataaatgatc   68438

chimp.LOC112206744.LOC107973052.GGT2-.646279-708040.rev.compl      ------------------------------------------------------------   61762
FAM230B-LOC105372935-GGT2.NCBI.GGT2.rev.compl                      atgagcagcactttcccaggcactgtcctagttgctctgcatacatcatcgcatttaatc   68498

chimp.LOC112206744.LOC107973052.GGT2-.646279-708040.rev.compl      ------------------------------------------------------------   61762
FAM230B-LOC105372935-GGT2.NCBI.GGT2.rev.compl                      ctcacaacagtgaatggggtcgatttcattacagccacatttaccagacaaggaaatggt   68558

chimp.LOC112206744.LOC107973052.GGT2-.646279-708040.rev.compl      ------------------------------------------------------------   61762
FAM230B-LOC105372935-GGT2.NCBI.GGT2.rev.compl                      tttcaaggttaagtaaccttctcatgacacagaatgtgactaagagttttcttattccat   68618

chimp.LOC112206744.LOC107973052.GGT2-.646279-708040.rev.compl      ------------------------------------------------------------   61762
FAM230B-LOC105372935-GGT2.NCBI.GGT2.rev.compl                      ggctcacactaaatgtgctctgtgaatatcagtgagaagtaataaagcactgactggatt   68678

chimp.LOC112206744.LOC107973052.GGT2-.646279-708040.rev.compl      ------------------------------------------------------------   61762
FAM230B-LOC105372935-GGT2.NCBI.GGT2.rev.compl                      ttcttggtcaccgggtcctaaaagaaatgttggggaatctatgcagtcccaagttaaaaa   68738

chimp.LOC112206744.LOC107973052.GGT2-.646279-708040.rev.compl      ------------------------------------------------------------   61762
FAM230B-LOC105372935-GGT2.NCBI.GGT2.rev.compl                      tgggaactgaactgagctgccacagcatgctgaataggtggagtccaggccaactaagtg   68798

chimp.LOC112206744.LOC107973052.GGT2-.646279-708040.rev.compl      ------------------------------------------------------------   61762
```

```
FAM230B-LOC105372935-GGT2.NCBI.GGT2.rev.compl                    cttcatggaacttaaaggctccagagggaaagcaatcgggaaggcacttccacaggcaca   68858

chimp.LOC112206744.LOC107973052.GGT2-.646279-708040.rev.compl    ------------------------------------------------------------   61762
FAM230B-LOC105372935-GGT2.NCBI.GGT2.rev.compl                    tcggtgacaacttctggggattcttacctgtcagccttcagccaaggaaggacaggaagt   68918

chimp.LOC112206744.LOC107973052.GGT2-.646279-708040.rev.compl    ------------------------------------------------------------   61762
FAM230B-LOC105372935-GGT2.NCBI.GGT2.rev.compl                    ggcatgggctaaaagatcctttgtttgacctctttccagatcaggtgacaggaaaacaca   68978

chimp.LOC112206744.LOC107973052.GGT2-.646279-708040.rev.compl    ------------------------------------------------------------   61762
FAM230B-LOC105372935-GGT2.NCBI.GGT2.rev.compl                    acacttgaacctttccaataaggacagccagcacttactgagccccagccaggtaactga   69038

chimp.LOC112206744.LOC107973052.GGT2-.646279-708040.rev.compl    ------------------------------------------------------------   61762
FAM230B-LOC105372935-GGT2.NCBI.GGT2.rev.compl                    cactgttccaagtactctgcttgtataaccagctttaaccttgcaacagccctctgaggt   69098

chimp.LOC112206744.LOC107973052.GGT2-.646279-708040.rev.compl    ------------------------------------------------------------   61762
FAM230B-LOC105372935-GGT2.NCBI.GGT2.rev.compl                    aaatactcctagtgacctcactttacggatgcagaacctgaggcaaggagttagggagta   69158

chimp.LOC112206744.LOC107973052.GGT2-.646279-708040.rev.compl    ------------------------------------------------------------   61762
FAM230B-LOC105372935-GGT2.NCBI.GGT2.rev.compl                    aatgaagctggctaaaggcacttgcactggccaggcacagtggctcatgcctgtaatccc   69218

chimp.LOC112206744.LOC107973052.GGT2-.646279-708040.rev.compl    ------------------------------------------------------------   61762
FAM230B-LOC105372935-GGT2.NCBI.GGT2.rev.compl                    agcacttccagatggatcacttgaggttaggggttcgagaccagcctggccaacatggtg   69278

chimp.LOC112206744.LOC107973052.GGT2-.646279-708040.rev.compl    ------------------------------------------------------------   61762
FAM230B-LOC105372935-GGT2.NCBI.GGT2.rev.compl                    aatccccagctctactaaaaatacaaaaattagccgggtgtgatggcacaagcctgtaat   69338

chimp.LOC112206744.LOC107973052.GGT2-.646279-708040.rev.compl    ------------------------------------------------------------   61762
FAM230B-LOC105372935-GGT2.NCBI.GGT2.rev.compl                    cccagcactttgggaggccaaggcagatgggtcacttgaggttaggggttcgagaccagc   69398

chimp.LOC112206744.LOC107973052.GGT2-.646279-708040.rev.compl    ------------------------------------------------------------   61762
FAM230B-LOC105372935-GGT2.NCBI.GGT2.rev.compl                    ctggccaacatggtgaatccccagctctactaaaaatacaaaaattagccgggtgtgatg   69458

chimp.LOC112206744.LOC107973052.GGT2-.646279-708040.rev.compl    ------------------------------------------------------------   61762
FAM230B-LOC105372935-GGT2.NCBI.GGT2.rev.compl                    gcacacgcctgtaatcccagctactcaggaggctgaggcaggagaattgcttgaacctgg   69518

chimp.LOC112206744.LOC107973052.GGT2-.646279-708040.rev.compl    ------------------------------------------------------------   61762
FAM230B-LOC105372935-GGT2.NCBI.GGT2.rev.compl                    gagatggaggttgcagcaagccgagatcgcaccactgcactccagcccactcggtcaatg   69578

chimp.LOC112206744.LOC107973052.GGT2-.646279-708040.rev.compl    ------------------------------------------------------------   61762
FAM230B-LOC105372935-GGT2.NCBI.GGT2.rev.compl                    tcactgccatcaagggacagtctaaccaaggtccagatggcacaagtggagccataaaac   69638

chimp.LOC112206744.LOC107973052.GGT2-.646279-708040.rev.compl    ------------------------------------------------------------   61762
FAM230B-LOC105372935-GGT2.NCBI.GGT2.rev.compl                    caggacagcttgaggttcagcttcctaagcccacaggtgcaaggaccttatctggatcca   69698

chimp.LOC112206744.LOC107973052.GGT2-.646279-708040.rev.compl    ------------------------------------------------------------   61762
FAM230B-LOC105372935-GGT2.NCBI.GGT2.rev.compl                    tactttcatcaaaaatgtcccttgaggccaggcacagtggctcatgcctataatcccagc   69758

chimp.LOC112206744.LOC107973052.GGT2-.646279-708040.rev.compl    ------------------------------------------------------------   61762
FAM230B-LOC105372935-GGT2.NCBI.GGT2.rev.compl                    actttgggagggcgaggcaggtggatcacctgaggtcaggagttcgagaccagcctggcc   69818

chimp.LOC112206744.LOC107973052.GGT2-.646279-708040.rev.compl    ------------------------------------------------------------   61762
FAM230B-LOC105372935-GGT2.NCBI.GGT2.rev.compl                    aatatggtgaaaccctgtccctactgaaaatacaaaaatttagcccagcatggtgatgtg   69878

chimp.LOC112206744.LOC107973052.GGT2-.646279-708040.rev.compl    ------------------------------------------------------------   61762
FAM230B-LOC105372935-GGT2.NCBI.GGT2.rev.compl                    cgcctgtaatcccagctactccggaggctgaggcaggagaatcgcttgaacccagcaggt   69938

chimp.LOC112206744.LOC107973052.GGT2-.646279-708040.rev.compl    ------------------------------------------------------------   61762
FAM230B-LOC105372935-GGT2.NCBI.GGT2.rev.compl                    ggaggttgcagtgagccgagatcatgtcattgcactgtgggctgggcaacagagcaacac   69998

chimp.LOC112206744.LOC107973052.GGT2-.646279-708040.rev.compl    ------------------------------------------------------------   61762
FAM230B-LOC105372935-GGT2.NCBI.GGT2.rev.compl                    tctatgtaaaaaaaaaaaaaaaggtcccttcaggattcctcgagcctgggaggccgaggc   70058

chimp.LOC112206744.LOC107973052.GGT2-.646279-708040.rev.compl    ------------------------------------------------------------   61762
FAM230B-LOC105372935-GGT2.NCBI.GGT2.rev.compl                    tgccatgaacaatgattgtgccactgcactccagcctgggtgacagagcgagatcttgtc   70118

chimp.LOC112206744.LOC107973052.GGT2-.646279-708040.rev.compl    ------------------------------------------------------------   61762
FAM230B-LOC105372935-GGT2.NCBI.GGT2.rev.compl                    tcaaaaaaaaaaaaaaaaaaaaaaatggctcttgaaacagccatcaactcccatgtgata   70178

chimp.LOC112206744.LOC107973052.GGT2-.646279-708040.rev.compl    ------------------------------------------------------------   61762
FAM230B-LOC105372935-GGT2.NCBI.GGT2.rev.compl                    aatgcactgccacaatccagcagcaaacctaaagttcggagtgaattcctaagtcttcgc   70238

chimp.LOC112206744.LOC107973052.GGT2-.646279-708040.rev.compl    ------------------------------------------------------------   61762
FAM230B-LOC105372935-GGT2.NCBI.GGT2.rev.compl                    cttcactgacttcccacagcatgggcagtagctgtgagaccttgggctggttacttaacc   70298

chimp.LOC112206744.LOC107973052.GGT2-.646279-708040.rev.compl    ------------------------------------------------------------   61762
FAM230B-LOC105372935-GGT2.NCBI.GGT2.rev.compl                    tctctgcacctcagtttcctcatctgtaaagtggtataatcgtcacagtgttatggaagg   70358

chimp.LOC112206744.LOC107973052.GGT2-.646279-708040.rev.compl    ------------------------------------------------------------   61762
FAM230B-LOC105372935-GGT2.NCBI.GGT2.rev.compl                    atttaatgagttgctacatttaaaacttaattcctggcacacaataagtgctcaataaat   70418

chimp.LOC112206744.LOC107973052.GGT2-.646279-708040.rev.compl    ------------------------------------------------------------   61762
FAM230B-LOC105372935-GGT2.NCBI.GGT2.rev.compl                    gtttagccgctagttgcagtagtagcaatagtagtgatattaactctagctgtggaacta   70478

chimp.LOC112206744.LOC107973052.GGT2-.646279-708040.rev.compl    ------------------------------------------------------------   61762
FAM230B-LOC105372935-GGT2.NCBI.GGT2.rev.compl                    atttctcccacacaccgtggcaggttgtattctccagacacagcagtaacaatatctcct   70538

chimp.LOC112206744.LOC107973052.GGT2-.646279-708040.rev.compl    ------------------------------------------------------------   61762
FAM230B-LOC105372935-GGT2.NCBI.GGT2.rev.compl                    tcccacacgctcttttgcaatgtgaccatgccacacactcccatcaagaggtggagttta   70598

chimp.LOC112206744.LOC107973052.GGT2-.646279-708040.rev.compl    ------------------------------------------------------------   61762
FAM230B-LOC105372935-GGT2.NCBI.GGT2.rev.compl                    gaataattccctcctctgtggatctgggcaggaagtcactgtgtgacttccagtcctagg   70658

chimp.LOC112206744.LOC107973052.GGT2-.646279-708040.rev.compl    ------------------------------------------------------------   61762
FAM230B-LOC105372935-GGT2.NCBI.GGT2.rev.compl                    tcatgacaagtgatgtagcttccatttcactcagggtgccctgaaccaccacataagaaa   70718

chimp.LOC112206744.LOC107973052.GGT2-.646279-708040.rev.compl    ------------------------------------------------------------   61762
FAM230B-LOC105372935-GGT2.NCBI.GGT2.rev.compl                    gccaacggtcctgaggccatgctgtgagggggcccaaggcacatggagaggcccagggca   70778

chimp.LOC112206744.LOC107973052.GGT2-.646279-708040.rev.compl    ------------------------------------------------------------   61762
FAM230B-LOC105372935-GGT2.NCBI.GGT2.rev.compl                    ggtgctccagtccagagctggctccagccaggtcccccctgccaccatgtgagtgaatgag   70838

chimp.LOC112206744.LOC107973052.GGT2-.646279-708040.rev.compl    ------------------------------------------------------------   61762
FAM230B-LOC105372935-GGT2.NCBI.GGT2.rev.compl                    ccaatcaccacccagctccacctgccaccatgtgagtgaacaagcctcccctgtgagctg   70898

chimp.LOC112206744.LOC107973052.GGT2-.646279-708040.rev.compl    ------------------------------------------------------------   61762
FAM230B-LOC105372935-GGT2.NCBI.GGT2.rev.compl                    cccccagccttggaacttccagccatgcctgccgtgcctgtaaattctcaacccccagat   70958

chimp.LOC112206744.LOC107973052.GGT2-.646279-708040.rev.compl    ------------------------------------------------------------   61762
FAM230B-LOC105372935-GGT2.NCBI.GGT2.rev.compl                    tctgtgagcataacaaaattgttgttttaagccactgaggttgggaaacatttgttacat   71018
```

```
chimp.LOC112206744.LOC107973052.GGT2-.646279-708040.rev.compl      ------------------------------------------------------------   61762
FAM230B-LOC105372935-GGT2.NCBI.GGT2.rev.compl                      agaaaggtaacaagaacatatgctatggcttcaatcaacatttttttcagtacctattat   71078

chimp.LOC112206744.LOC107973052.GGT2-.646279-708040.rev.compl      ------------------------------------------------------------   61762
FAM230B-LOC105372935-GGT2.NCBI.GGT2.rev.compl                      gatggacactaggacagaaactgtgagggagagattaataagacatagttgacaagttgt   71138

chimp.LOC112206744.LOC107973052.GGT2-.646279-708040.rev.compl      ------------------------------------------------------------   61762
FAM230B-LOC105372935-GGT2.NCBI.GGT2.rev.compl                      cctaagctgacaatcctaaaataaatacacataactaatcattttaatggtatacgatag   71198

chimp.LOC112206744.LOC107973052.GGT2-.646279-708040.rev.compl      ------------------------------------------------------------   61762
FAM230B-LOC105372935-GGT2.NCBI.GGT2.rev.compl                      cagtgaggtgcaaagtgcaaggttaagccctttcagaaagtgaaagcattcaaaaagcct   71258

chimp.LOC112206744.LOC107973052.GGT2-.646279-708040.rev.compl      ------------------------------------------------------------   61762
FAM230B-LOC105372935-GGT2.NCBI.GGT2.rev.compl                      tcaaaaaagtagtgtcatctgaattggatcccaaagcgcaagggaaatatataccacatg   71318

chimp.LOC112206744.LOC107973052.GGT2-.646279-708040.rev.compl      ------------------------------------------------------------   61762
FAM230B-LOC105372935-GGT2.NCBI.GGT2.rev.compl                      aaaacaccgcagtgcattccaggattagcaagaagtttgcggaggtcgttttgtcacatg   71378

chimp.LOC112206744.LOC107973052.GGT2-.646279-708040.rev.compl      ------------------------------------------------------------   61762
FAM230B-LOC105372935-GGT2.NCBI.GGT2.rev.compl                      actggagggcacatcagtgccaggcggaaaagggcaccgaaaggcacattaataaagtca   71438

chimp.LOC112206744.LOC107973052.GGT2-.646279-708040.rev.compl      ------------------------------------------------------------   61762
FAM230B-LOC105372935-GGT2.NCBI.GGT2.rev.compl                      aacacctcctctgggacaggggaagttgagaaagagcctaacagaggtaaggaagaagac   71498

chimp.LOC112206744.LOC107973052.GGT2-.646279-708040.rev.compl      ------------------------------------------------------------   61762
FAM230B-LOC105372935-GGT2.NCBI.GGT2.rev.compl                      ggatcagatttacactttcgaattatcactctggtggcaggtgaaggacagagcagaggg   71558

chimp.LOC112206744.LOC107973052.GGT2-.646279-708040.rev.compl      ------------------------------------------------------------   61762
FAM230B-LOC105372935-GGT2.NCBI.GGT2.rev.compl                      caggactggaagcagacagaccaaattcacagtcaaatgcgacaagatgagggcctggcc   71618

chimp.LOC112206744.LOC107973052.GGT2-.646279-708040.rev.compl      ------------------------------------------------------------   61762
FAM230B-LOC105372935-GGT2.NCBI.GGT2.rev.compl                      tggttttaaatccaaggacaaaaatatcattcattcactctttcattctttctcttatct   71678

chimp.LOC112206744.LOC107973052.GGT2-.646279-708040.rev.compl      ------------------------------------------------------------   61762
FAM230B-LOC105372935-GGT2.NCBI.GGT2.rev.compl                      gaatccccagtggtacctaaaatagagctggtctaaagatgatgattaatgaatgttaaa   71738

chimp.LOC112206744.LOC107973052.GGT2-.646279-708040.rev.compl      ------------------------------------------------------------   61762
FAM230B-LOC105372935-GGT2.NCBI.GGT2.rev.compl                      ttattgatactacttctaagaagatgtaactgccagctacagaaccagccaaaaggactg   71798

chimp.LOC112206744.LOC107973052.GGT2-.646279-708040.rev.compl      ------------------------------------------------------------   61762
FAM230B-LOC105372935-GGT2.NCBI.GGT2.rev.compl                      caaacccagtaattcatttcatcccaacctgactgatctctctgatgtctgcactgctgt   71858

chimp.LOC112206744.LOC107973052.GGT2-.646279-708040.rev.compl      ------------------------------------------------------------   61762
FAM230B-LOC105372935-GGT2.NCBI.GGT2.rev.compl                      gtcactagactgcccgattgtcctaggttgagacatttacctactgagccccagggatgg   71918

chimp.LOC112206744.LOC107973052.GGT2-.646279-708040.rev.compl      ------------------------------------------------------------   61762
FAM230B-LOC105372935-GGT2.NCBI.GGT2.rev.compl                      gtggctgcttctaagctgctgcctaattccctcattactaaatgttgacgctgcttcctt   71978

chimp.LOC112206744.LOC107973052.GGT2-.646279-708040.rev.compl      ------------------------------------------------------------   61762
FAM230B-LOC105372935-GGT2.NCBI.GGT2.rev.compl                      agaggactgctaaaccataactcaaaaatgttttaaacatgctactatttacttagatta   72038

chimp.LOC112206744.LOC107973052.GGT2-.646279-708040.rev.compl      ------------------------------------------------------------   61762
FAM230B-LOC105372935-GGT2.NCBI.GGT2.rev.compl                      gaatatttacttaaatgttttaaaaaatacagtattttaagtcttttatctcttctgact   72098

chimp.LOC112206744.LOC107973052.GGT2-.646279-708040.rev.compl      ------------------------------------------------------------   61762
FAM230B-LOC105372935-GGT2.NCBI.GGT2.rev.compl                      actaccactaaccctgcttcccctcatttttccagagattagcactactgtgatttaagg   72158

chimp.LOC112206744.LOC107973052.GGT2-.646279-708040.rev.compl      ------------------------------------------------------------   61762
FAM230B-LOC105372935-GGT2.NCBI.GGT2.rev.compl                      atagagctctccagacttaaaaaaaaaccaaaaaactgcttttatatccagatatatgta   72218

chimp.LOC112206744.LOC107973052.GGT2-.646279-708040.rev.compl      ------------------------------------------------------------   61762
FAM230B-LOC105372935-GGT2.NCBI.GGT2.rev.compl                      accactgaaaaatatacattatttttcagcatttttacataaacattatcatccctttc   72278

chimp.LOC112206744.LOC107973052.GGT2-.646279-708040.rev.compl      ------------------------------------------------------------   61762
FAM230B-LOC105372935-GGT2.NCBI.GGT2.rev.compl                      attgagttatgttcctcagttttcctcctcaaagtttttgagatcaaatatagttcccat   72338

chimp.LOC112206744.LOC107973052.GGT2-.646279-708040.rev.compl      ------------------------------------------------------------   61762
FAM230B-LOC105372935-GGT2.NCBI.GGT2.rev.compl                      ttttcttctttatatagctgcacagtattccatcataggattacaccacattttatttat   72398

chimp.LOC112206744.LOC107973052.GGT2-.646279-708040.rev.compl      ------------------------------------------------------------   61762
FAM230B-LOC105372935-GGT2.NCBI.GGT2.rev.compl                      tccaatattaatgaatatttaggttatttatactttttttttctataaccaacaagcccat   72458

chimp.LOC112206744.LOC107973052.GGT2-.646279-708040.rev.compl      ------------------------------------------------------------   61762
FAM230B-LOC105372935-GGT2.NCBI.GGT2.rev.compl                      gccttctactgcttatcctttccccacctgaatttgtacaatccccagcttctcttatcc   72518

chimp.LOC112206744.LOC107973052.GGT2-.646279-708040.rev.compl      ------------------------------------------------------------   61762
FAM230B-LOC105372935-GGT2.NCBI.GGT2.rev.compl                      atgaccactgaaaccccagggaagaccttatcactttcaggatgtccatggctctttcac   72578

chimp.LOC112206744.LOC107973052.GGT2-.646279-708040.rev.compl      ------------------------------------------------------------   61762
FAM230B-LOC105372935-GGT2.NCBI.GGT2.rev.compl                      cctatgccatgcttggaaccaggaaagcccaaaagaccccacttcctgcccttctattct   72638

chimp.LOC112206744.LOC107973052.GGT2-.646279-708040.rev.compl      ------------------------------------------------------------   61762
FAM230B-LOC105372935-GGT2.NCBI.GGT2.rev.compl                      gctcctcctctcctatcgctcatttctttgcctctatcctccactcctcccagcaaaggt   72698

chimp.LOC112206744.LOC107973052.GGT2-.646279-708040.rev.compl      ------------------------------------------------------------   61762
FAM230B-LOC105372935-GGT2.NCBI.GGT2.rev.compl                      agggatagcacctgagtcctgccaccctctcacctgcattccccctcactctcatgcgaa   72758

chimp.LOC112206744.LOC107973052.GGT2-.646279-708040.rev.compl      ------------------------------------------------------------   61762
FAM230B-LOC105372935-GGT2.NCBI.GGT2.rev.compl                      ggacagcacagaagcagcatcaaaaaagaaaggcagcgagggagccctgcagcctgccct   72818

chimp.LOC112206744.LOC107973052.GGT2-.646279-708040.rev.compl      ------------------------------------------------------------   61762
FAM230B-LOC105372935-GGT2.NCBI.GGT2.rev.compl                      aggccagccatgcccacagccacaggaatagacactgttttcagccccacctagattcaa   72878

chimp.LOC112206744.LOC107973052.GGT2-.646279-708040.rev.compl      ------------------------------------------------------------   61762
FAM230B-LOC105372935-GGT2.NCBI.GGT2.rev.compl                      ggcagcatctaatagaaatgcttataagtaaatataagtgccagtatttgccttctattt   72938

chimp.LOC112206744.LOC107973052.GGT2-.646279-708040.rev.compl      ------------------------------------------------------------   61762
FAM230B-LOC105372935-GGT2.NCBI.GGT2.rev.compl                      tctagaggggaaaaaaattaaaaggcaagatcagattacttgcctggaaatgttctccct   72998

chimp.LOC112206744.LOC107973052.GGT2-.646279-708040.rev.compl      ------------------------------------------------------------   61762
FAM230B-LOC105372935-GGT2.NCBI.GGT2.rev.compl                      gtaattactctgaccgctctttctttccctccggttttggagataatggaaataggctga   73058

chimp.LOC112206744.LOC107973052.GGT2-.646279-708040.rev.compl      ------------------------------------------------------------   61762
FAM230B-LOC105372935-GGT2.NCBI.GGT2.rev.compl                      gtgttgggcatttttttttttaaagagggcagcttcatggcaacactccctctgagaagt   73118

chimp.LOC112206744.LOC107973052.GGT2-.646279-708040.rev.compl      ------------------------------------------------------------   61762
FAM230B-LOC105372935-GGT2.NCBI.GGT2.rev.compl                      ccagaggctcagtgaaccgcagcttagagtgagttccagtaaaaggaccatcacttgaac   73178
```

```
chimp.LOC112206744.LOC107973052.GGT2-.646279-708040.rev.compl      ------------------------------------------------------------   61762
FAM230B-LOC105372935-GGT2.NCBI.GGT2.rev.compl                      cagtaagtgcctcattccttctcacagcatacatttcacagcaattactgttgctattat   73238

chimp.LOC112206744.LOC107973052.GGT2-.646279-708040.rev.compl      ------------------------------------------------------------   61762
FAM230B-LOC105372935-GGT2.NCBI.GGT2.rev.compl                      tttacgcttcatttctggacctaggatacctcagtttctccagactctcagtctggtccc   73298

chimp.LOC112206744.LOC107973052.GGT2-.646279-708040.rev.compl      ------------------------------------------------------------   61762
FAM230B-LOC105372935-GGT2.NCBI.GGT2.rev.compl                      caggttttcagtctgggagaaaggtgtctgctggatttcaaacagtccctctactggccg   73358

chimp.LOC112206744.LOC107973052.GGT2-.646279-708040.rev.compl      ------------------------------------------------------------   61762
FAM230B-LOC105372935-GGT2.NCBI.GGT2.rev.compl                      ccaccaccaccctctcctaaagaggactggaataactcttccacagctcccgccacactc   73418

chimp.LOC112206744.LOC107973052.GGT2-.646279-708040.rev.compl      ------------------------------------------------------------   61762
FAM230B-LOC105372935-GGT2.NCBI.GGT2.rev.compl                      agcactgcctgcacctcctcctggagctggcagggccccagctgtgcacccctcaatgct   73478

chimp.LOC112206744.LOC107973052.GGT2-.646279-708040.rev.compl      ------------------------------------------------------------   61762
FAM230B-LOC105372935-GGT2.NCBI.GGT2.rev.compl                      ctcacagcttcaggaagaggactgctaagcccagggccctgctgacaacatgccgggagc   73538

chimp.LOC112206744.LOC107973052.GGT2-.646279-708040.rev.compl      ------------------------------------------------------------   61762
FAM230B-LOC105372935-GGT2.NCBI.GGT2.rev.compl                      gcatctgcaggcagggtcagaatgtcagctaccaggcagcagagagatgcggggcgccaa   73598

chimp.LOC112206744.LOC107973052.GGT2-.646279-708040.rev.compl      ------------------------------------------------------------   61762
FAM230B-LOC105372935-GGT2.NCBI.GGT2.rev.compl                      cagcaaaagcacttgggggcttttcaagtggcatcagaactagacctggatacaaagg   73658

chimp.LOC112206744.LOC107973052.GGT2-.646279-708040.rev.compl      ------------------------------------------------------------   61762
FAM230B-LOC105372935-GGT2.NCBI.GGT2.rev.compl                      ggtggccctggcttcaggaagcccatgataagtcagtctgctgtgtccttcaacacagtg   73718

chimp.LOC112206744.LOC107973052.GGT2-.646279-708040.rev.compl      ------------------------------------------------------------   61762
FAM230B-LOC105372935-GGT2.NCBI.GGT2.rev.compl                      tccccaccttctgtctcccaacactaggccatggagacacaaaactagaaaatggggctg   73778

chimp.LOC112206744.LOC107973052.GGT2-.646279-708040.rev.compl      ------------------------------------------------------------   61762
FAM230B-LOC105372935-GGT2.NCBI.GGT2.rev.compl                      cctagctacgaatgcccaggctgtgaattggggtctccaaaggaacctcagtgccgtgtc   73838

chimp.LOC112206744.LOC107973052.GGT2-.646279-708040.rev.compl      ------------------------------------------------------------   61762
FAM230B-LOC105372935-GGT2.NCBI.GGT2.rev.compl                      ctactcagagcagctccttcagctcttcaccctggaaagcactccggtacaaaacaggaa   73898

chimp.LOC112206744.LOC107973052.GGT2-.646279-708040.rev.compl      ------------------------------------------------------------   61762
FAM230B-LOC105372935-GGT2.NCBI.GGT2.rev.compl                      tattcctcttctcctgactctgcccaaggcactgctgtgaccctctcagcagtggaagca   73958

chimp.LOC112206744.LOC107973052.GGT2-.646279-708040.rev.compl      ------------------------------------------------------------   61762
FAM230B-LOC105372935-GGT2.NCBI.GGT2.rev.compl                      ctagccaatgtgccatcctcctggacgatagctggataatcgcaataagcagagccccaa   74018

chimp.LOC112206744.LOC107973052.GGT2-.646279-708040.rev.compl      ------------------------------------------------------------   61762
FAM230B-LOC105372935-GGT2.NCBI.GGT2.rev.compl                      caactagctctagtgtgtggggagaagtggggaggatggagaaaggatacaaaaagacac   74078

chimp.LOC112206744.LOC107973052.GGT2-.646279-708040.rev.compl      ------------------------------------------------------------   61762
FAM230B-LOC105372935-GGT2.NCBI.GGT2.rev.compl                      tgtgggatttcagggcaacccatgagtgaggatcacagcagagctatgccttgcaaggac   74138

chimp.LOC112206744.LOC107973052.GGT2-.646279-708040.rev.compl      ------------------------------------------------------------   61762
FAM230B-LOC105372935-GGT2.NCBI.GGT2.rev.compl                      cagaactggaaaacctttgtgacttagaccgccctcctcaggacagctcccgccactctt   74198

chimp.LOC112206744.LOC107973052.GGT2-.646279-708040.rev.compl      ------------------------------------------------------------   61762
FAM230B-LOC105372935-GGT2.NCBI.GGT2.rev.compl                      cagggacaaagttcctttactcagccatagtttctgtgctccaaatactcagcttacaca   74258

chimp.LOC112206744.LOC107973052.GGT2-.646279-708040.rev.compl      ------------------------------------------------------------   61762
FAM230B-LOC105372935-GGT2.NCBI.GGT2.rev.compl                      tgcctgttgctgctgcactcaccactgtggttctcagtgcaaattccccagaaagcagtc   74318

chimp.LOC112206744.LOC107973052.GGT2-.646279-708040.rev.compl      ------------------------------------------------------------   61762
FAM230B-LOC105372935-GGT2.NCBI.GGT2.rev.compl                      tgactggctgggagacctttttcttttaaccaggcccccaaagttgtagattttggggt   74378

chimp.LOC112206744.LOC107973052.GGT2-.646279-708040.rev.compl      ------------------------------------------------------------   61762
FAM230B-LOC105372935-GGT2.NCBI.GGT2.rev.compl                      catttacaatatgcatttccccaacacactcagatctaggtggacttgaccagatgggca   74438

chimp.LOC112206744.LOC107973052.GGT2-.646279-708040.rev.compl      ------------------------------------------------------------   61762
FAM230B-LOC105372935-GGT2.NCBI.GGT2.rev.compl                      tctgtattccattacccagcatataaccactttgggcaacagggaatgtgaagggcccca   74498

chimp.LOC112206744.LOC107973052.GGT2-.646279-708040.rev.compl      ------------------------------------------------------------   61762
FAM230B-LOC105372935-GGT2.NCBI.GGT2.rev.compl                      aaccgttccccaccaagagtgtgggtggcaggcaccctgaaatatgtctagcacactcat   74558

chimp.LOC112206744.LOC107973052.GGT2-.646279-708040.rev.compl      ------------------------------------------------------------   61762
FAM230B-LOC105372935-GGT2.NCBI.GGT2.rev.compl                      aaagcatgaacatctacacacaactaaaaataaatagtgagagccctgacactgcttctg   74618

chimp.LOC112206744.LOC107973052.GGT2-.646279-708040.rev.compl      ------------------------------------------------------------   61762
FAM230B-LOC105372935-GGT2.NCBI.GGT2.rev.compl                      ggggttttcccaagcacctgttacgaatggtaacaatggatggccgctccttctccctca   74678

chimp.LOC112206744.LOC107973052.GGT2-.646279-708040.rev.compl      ------------------------------------------------------------   61762
FAM230B-LOC105372935-GGT2.NCBI.GGT2.rev.compl                      gctttgactttcattgctggtggctcaggctcctgtgcttggctgtcatcggtgggtagc   74738

chimp.LOC112206744.LOC107973052.GGT2-.646279-708040.rev.compl      ------------------------------------------------------------   61762
FAM230B-LOC105372935-GGT2.NCBI.GGT2.rev.compl                      cctggattggacattttccctattcaattcaaggaccatgttcttgagcataactatgac   74798

chimp.LOC112206744.LOC107973052.GGT2-.646279-708040.rev.compl      ------------------------------------------------------------   61762
FAM230B-LOC105372935-GGT2.NCBI.GGT2.rev.compl                      caggcagtaggcagtgggccctgctctttctttccagggaggcacagataagccagcatc   74858

chimp.LOC112206744.LOC107973052.GGT2-.646279-708040.rev.compl      ------------------------------------------------------------   61762
FAM230B-LOC105372935-GGT2.NCBI.GGT2.rev.compl                      acagatgcaaaaaacagaaccgtggaggtagagactaattcttcagcagcccagagaaga   74918

chimp.LOC112206744.LOC107973052.GGT2-.646279-708040.rev.compl      ------------------------------------------------------------   61762
FAM230B-LOC105372935-GGT2.NCBI.GGT2.rev.compl                      gaactgggaaacagggccctcgcagatacacaagaccaagctggagagttcagggacttc   74978

chimp.LOC112206744.LOC107973052.GGT2-.646279-708040.rev.compl      ------------------------------------------------------------   61762
FAM230B-LOC105372935-GGT2.NCBI.GGT2.rev.compl                      caaaagaaggtacaagaatgagcactataccatagtggggtgggtcacaaaagcaggaaa   75038

chimp.LOC112206744.LOC107973052.GGT2-.646279-708040.rev.compl      ------------------------------------------------------------   61762
FAM230B-LOC105372935-GGT2.NCBI.GGT2.rev.compl                      aggtccccaaacaaagaatccatcagtcagttacagcactctccacatgtggtgaaacag   75098

chimp.LOC112206744.LOC107973052.GGT2-.646279-708040.rev.compl      ------------------------------------------------------------   61762
FAM230B-LOC105372935-GGT2.NCBI.GGT2.rev.compl                      tatgcaaccatttaaaaactgcagagaacaatatttatgaacacctaaaaatgctgaaag   75158

chimp.LOC112206744.LOC107973052.GGT2-.646279-708040.rev.compl      ------------------------------------------------------------   61762
FAM230B-LOC105372935-GGT2.NCBI.GGT2.rev.compl                      cacactactaagtaaaaggggagaaaggctataaaaggatttgaaagtattttcgtaaag   75218

chimp.LOC112206744.LOC107973052.GGT2-.646279-708040.rev.compl      ------------------------------------------------------------   61762
FAM230B-LOC105372935-GGT2.NCBI.GGT2.rev.compl                      ataaacacccacacacacccacacacctctatatatgtgtggaaaaaggactgggctagt   75278

chimp.LOC112206744.LOC107973052.GGT2-.646279-708040.rev.compl      ------------------------------------------------------------   61762
FAM230B-LOC105372935-GGT2.NCBI.GGT2.rev.compl                      atatttaaaaaacgctaaagggaggtcgggtaataagggattgtttttccttgcacttt   75338
```

```
chimp.LOC112206744.LOC107973052.GGT2-.646279-708040.rev.compl      ------------------------------------------------------------   61762
FAM230B-LOC105372935-GGT2.NCBI.GGT2.rev.compl                      gtgttttctaagtattctatagtctgaatgaattagattcataattaggaccaacagtca   75398

chimp.LOC112206744.LOC107973052.GGT2-.646279-708040.rev.compl      ------------------------------------------------------------   61762
FAM230B-LOC105372935-GGT2.NCBI.GGT2.rev.compl                      taagcaatttctcagaccacctgcctcataaaacccaaaaaaagaagtcagagaggggtg   75458

chimp.LOC112206744.LOC107973052.GGT2-.646279-708040.rev.compl      ------------------------------------------------------------   61762
FAM230B-LOC105372935-GGT2.NCBI.GGT2.rev.compl                      aatattttggtacagatgaattcaggttgtgtaagtggagaggagggaggtggccaagga   75518

chimp.LOC112206744.LOC107973052.GGT2-.646279-708040.rev.compl      ------------------------------------------------------------   61762
FAM230B-LOC105372935-GGT2.NCBI.GGT2.rev.compl                      gcctgaagtgggggagggggctgatggtgaaatagggaatgggtggagagagaaaatga   75578

chimp.LOC112206744.LOC107973052.GGT2-.646279-708040.rev.compl      ------------------------------------------------------------   61762
FAM230B-LOC105372935-GGT2.NCBI.GGT2.rev.compl                      ttgcaaaattggctggagccagcctggtcaggtcaactttgcaggtgtgtgaagtgtcaa   75638

chimp.LOC112206744.LOC107973052.GGT2-.646279-708040.rev.compl      ------------------------------------------------------------   61762
FAM230B-LOC105372935-GGT2.NCBI.GGT2.rev.compl                      atggttaatcagttgccctctgtttgagctgcttaatgcacaagggcctggttgaacagg   75698

chimp.LOC112206744.LOC107973052.GGT2-.646279-708040.rev.compl      ------------------------------------------------------------   61762
FAM230B-LOC105372935-GGT2.NCBI.GGT2.rev.compl                      tttgacctcatttactttttcttaaagctgtttagccactttcctaaaaactcctatttt   75758

chimp.LOC112206744.LOC107973052.GGT2-.646279-708040.rev.compl      ------------------------------------------------------------   61762
FAM230B-LOC105372935-GGT2.NCBI.GGT2.rev.compl                      ctgagttattaaactttccattccatgttgatcccttccgagccctctaacgcatctctt   75818

chimp.LOC112206744.LOC107973052.GGT2-.646279-708040.rev.compl      ------------------------------------------------------------   61762
FAM230B-LOC105372935-GGT2.NCBI.GGT2.rev.compl                      ctcttgcttcttccaactggatgccttcatgaggcattaacaggtactgtgagccctgat   75878

chimp.LOC112206744.LOC107973052.GGT2-.646279-708040.rev.compl      ------------------------------------------------------------   61762
FAM230B-LOC105372935-GGT2.NCBI.GGT2.rev.compl                      gaaaatcatcctttcagggtacaaggaaggcacctgaatgacagatggttggaggaatct   75938

chimp.LOC112206744.LOC107973052.GGT2-.646279-708040.rev.compl      ------------------------------------------------------------   61762
FAM230B-LOC105372935-GGT2.NCBI.GGT2.rev.compl                      tgagaaagcagtttaggcattaacatgcaggctgtcgctgacttacacagagccaaggac   75998

chimp.LOC112206744.LOC107973052.GGT2-.646279-708040.rev.compl      ------------------------------------------------------------   61762
FAM230B-LOC105372935-GGT2.NCBI.GGT2.rev.compl                      gtgacatacagcatggggccatttctaaatccccactgaggacaagggtcctggagggaa   76058

chimp.LOC112206744.LOC107973052.GGT2-.646279-708040.rev.compl      ------------------------------------------------------------   61762
FAM230B-LOC105372935-GGT2.NCBI.GGT2.rev.compl                      tgggatttggaacaagatttgggctttgaacattcgctacaggctctgacaaagcagtga   76118

chimp.LOC112206744.LOC107973052.GGT2-.646279-708040.rev.compl      ------------------------------------------------------------   61762
FAM230B-LOC105372935-GGT2.NCBI.GGT2.rev.compl                      tggttcaaaagcctctgcccctggtggcaagccctgcccagatgctgttaaggggctct   76178

chimp.LOC112206744.LOC107973052.GGT2-.646279-708040.rev.compl      ------------------------------------------------------------   61762
FAM230B-LOC105372935-GGT2.NCBI.GGT2.rev.compl                      ccaggcacgccctcctaattaggggcaaaatcagagaagcctctttggaagaagacaaaa   76238

chimp.LOC112206744.LOC107973052.GGT2-.646279-708040.rev.compl      ------------------------------------------------------------   61762
FAM230B-LOC105372935-GGT2.NCBI.GGT2.rev.compl                      ggcacaatgagatgcggtctcctgaagtagaggctttctctctggttatcacctgaaaca   76298

chimp.LOC112206744.LOC107973052.GGT2-.646279-708040.rev.compl      ------------------------------------------------------------   61762
FAM230B-LOC105372935-GGT2.NCBI.GGT2.rev.compl                      ggaaataaggactgagtacctgctcaattccaggcactagccaaatctgcagcaaactag   76358

chimp.LOC112206744.LOC107973052.GGT2-.646279-708040.rev.compl      ------------------------------------------------------------   61762
FAM230B-LOC105372935-GGT2.NCBI.GGT2.rev.compl                      catttccaaaggtgtcttcttactttaatctcacagccaccctgtgagggaagtgtacat   76418

chimp.LOC112206744.LOC107973052.GGT2-.646279-708040.rev.compl      ------------------------------------------------------------   61762
FAM230B-LOC105372935-GGT2.NCBI.GGT2.rev.compl                      tccccatttcgctgatgagatgagcaaactgaggctcggagaaattgtcagtgggataca   76478

chimp.LOC112206744.LOC107973052.GGT2-.646279-708040.rev.compl      ------------------------------------------------------------   61762
FAM230B-LOC105372935-GGT2.NCBI.GGT2.rev.compl                      gagttggcatccaaacccacaatctctgactctaagcttctgtttcattcattccacaaa   76538

chimp.LOC112206744.LOC107973052.GGT2-.646279-708040.rev.compl      ------------------------------------------------------------   61762
FAM230B-LOC105372935-GGT2.NCBI.GGT2.rev.compl                      tatatatggtgtctcctctttgctggacactgtactgggctgatgagctacaagacccaa   76598

chimp.LOC112206744.LOC107973052.GGT2-.646279-708040.rev.compl      ------------------------------------------------------------   61762
FAM230B-LOC105372935-GGT2.NCBI.GGT2.rev.compl                      tctttacagtctaggaggcagagatagataagtaaacagaggcggcaaagcgtggcactg   76658

chimp.LOC112206744.LOC107973052.GGT2-.646279-708040.rev.compl      ------------------------------------------------------------   61762
FAM230B-LOC105372935-GGT2.NCBI.GGT2.rev.compl                      tgaaagatgcactgaggacaccgagctaggtgttccacaggcgtcaggaaaggcttcaca   76718

chimp.LOC112206744.LOC107973052.GGT2-.646279-708040.rev.compl      ------------------------------------------------------------   61762
FAM230B-LOC105372935-GGT2.NCBI.GGT2.rev.compl                      caagagaggttgcttgtgcccagttctaattaggatgagtaggtatttctgctagatcgt   76778

chimp.LOC112206744.LOC107973052.GGT2-.646279-708040.rev.compl      ------------------------------------------------------------   61762
FAM230B-LOC105372935-GGT2.NCBI.GGT2.rev.compl                      ggtggggtggggaggacatcacataggacctcagcagagagcagctcgaatctatgcaga   76838

chimp.LOC112206744.LOC107973052.GGT2-.646279-708040.rev.compl      ------------------------------------------------------------   61762
FAM230B-LOC105372935-GGT2.NCBI.GGT2.rev.compl                      gctgcacgaggagccggaggtccagagtactgcagcagtttgacattgatgggtggtagg   76898

chimp.LOC112206744.LOC107973052.GGT2-.646279-708040.rev.compl      ------------------------------------------------------------   61762
FAM230B-LOC105372935-GGT2.NCBI.GGT2.rev.compl                      gtccatgaaagtagaggaagaccaagaggaagcagcagacaggggccaggccagggcagg   76958

chimp.LOC112206744.LOC107973052.GGT2-.646279-708040.rev.compl      ------------------------------------------------------------   61762
FAM230B-LOC105372935-GGT2.NCBI.GGT2.rev.compl                      gatagggtgctctactaaagagcatgggctttgaactatgagcagtggagagccactggg   77018

chimp.LOC112206744.LOC107973052.GGT2-.646279-708040.rev.compl      ------------------------------------------------------------   61762
FAM230B-LOC105372935-GGT2.NCBI.GGT2.rev.compl                      ggtgtaaataaggaaacagatattccaaatcctgccttgggaagattgttctggcaggag   77078

chimp.LOC112206744.LOC107973052.GGT2-.646279-708040.rev.compl      ------------------------------------------------------------   61762
FAM230B-LOC105372935-GGT2.NCBI.GGT2.rev.compl                      tgatgggaaaggggccctggcagaagattccttcatgctcctgcggtgatccaggtgaga   77138

chimp.LOC112206744.LOC107973052.GGT2-.646279-708040.rev.compl      ------------------------------------------------------------   61762
FAM230B-LOC105372935-GGT2.NCBI.GGT2.rev.compl                      gctgctgaggaccccaaccaaatcagatcactggacctccactgggagtgaaaggggtg   77198

chimp.LOC112206744.LOC107973052.GGT2-.646279-708040.rev.compl      ------------------------------------------------------------   61762
FAM230B-LOC105372935-GGT2.NCBI.GGT2.rev.compl                      gtctggccagagagatagtaaagagatgtagttaacaggcaagagaatgtggggatatcc   77258

chimp.LOC112206744.LOC107973052.GGT2-.646279-708040.rev.compl      ------------------------------------------------------------   61762
FAM230B-LOC105372935-GGT2.NCBI.GGT2.rev.compl                      agggagctgtccacaattcgcaggtcttaggcaactgcttgggagaagttgatgaggtca   77318

chimp.LOC112206744.LOC107973052.GGT2-.646279-708040.rev.compl      ------------------------------------------------------------   61762
FAM230B-LOC105372935-GGT2.NCBI.GGT2.rev.compl                      tcactggacagagaatctattccctgggcccaaccaactctctagaccatcctgccatgt   77378

chimp.LOC112206744.LOC107973052.GGT2-.646279-708040.rev.compl      ------------------------------------------------------------   61762
FAM230B-LOC105372935-GGT2.NCBI.GGT2.rev.compl                      gcctctgagaagtgagcttccccagccaagcccacaggaggacatatggagcaaaggtca   77438

chimp.LOC112206744.LOC107973052.GGT2-.646279-708040.rev.compl      ------------------------------------------------------------   61762
FAM230B-LOC105372935-GGT2.NCBI.GGT2.rev.compl                      acagtcactcagctgatggatggcaagcattcagatgggaggtgcctccacaggccatga   77498

chimp.LOC112206744.LOC107973052.GGT2-.646279-708040.rev.compl      ------------------------------------------------------------   61762
```

```
FAM230B-LOC105372935-GGT2.NCBI.GGT2.rev.compl                 taacccatggatccatttgagaaagatgccaaaaaacagttaattcaaggccactttaaa   77558

chimp.LOC112206744.LOC107973052.GGT2-.646279-708040.rev.compl   ------------------------------------------------------------   61762
FAM230B-LOC105372935-GGT2.NCBI.GGT2.rev.compl                 atacctggcattcaatcctcctcaatgtctccatcaactgtagaccctataactaccaaa   77618

chimp.LOC112206744.LOC107973052.GGT2-.646279-708040.rev.compl   ------------------------------------------------------------   61762
FAM230B-LOC105372935-GGT2.NCBI.GGT2.rev.compl                 tgagctccactcgccacaaaggtcaagacacagaaacggtcagagagcagagcctattcc   77678

chimp.LOC112206744.LOC107973052.GGT2-.646279-708040.rev.compl   ------------------------------------------------------------   61762
FAM230B-LOC105372935-GGT2.NCBI.GGT2.rev.compl                 tgaaaagcaaaagactccaaactgccacaactcaaccagtgaagggctggggaagactga   77738

chimp.LOC112206744.LOC107973052.GGT2-.646279-708040.rev.compl   ------------------------------------------------------------   61762
FAM230B-LOC105372935-GGT2.NCBI.GGT2.rev.compl                 caggttcactaagaaatgagaggaaggacaaacaaggtgttcaaattagaaaatgattcc   77798

chimp.LOC112206744.LOC107973052.GGT2-.646279-708040.rev.compl   ------------------------------------------------------------   61762
FAM230B-LOC105372935-GGT2.NCBI.GGT2.rev.compl                 attaatgagggctctgatgatatagttgtggaggttatcagtttgccagagctgcagatt   77858

chimp.LOC112206744.LOC107973052.GGT2-.646279-708040.rev.compl   ------------------------------------------------------------   61762
FAM230B-LOC105372935-GGT2.NCBI.GGT2.rev.compl                 taaaatacttacatggtcatcgaaaccaaaaaatgctgaccatcccccaaaacctcccac   77918

chimp.LOC112206744.LOC107973052.GGT2-.646279-708040.rev.compl   ------------------------------------------------------------   61762
FAM230B-LOC105372935-GGT2.NCBI.GGT2.rev.compl                 cccacctagtgctgataacaggagttatcagttgccttttgtgtacagctagatgagata   77978

chimp.LOC112206744.LOC107973052.GGT2-.646279-708040.rev.compl   ------------------------------------------------------------   61762
FAM230B-LOC105372935-GGT2.NCBI.GGT2.rev.compl                 tggatctgtgggggaagaaggggtgggcgtgctagcatctctcagctagactgtcatcg   78038

chimp.LOC112206744.LOC107973052.GGT2-.646279-708040.rev.compl   ------------------------------------------------------------   61762
FAM230B-LOC105372935-GGT2.NCBI.GGT2.rev.compl                 gatggtgacattccccaggactcaagcagcatctgagaagccactttcagtatgactatg   78098

chimp.LOC112206744.LOC107973052.GGT2-.646279-708040.rev.compl   ------------------------------------------------------------   61762
FAM230B-LOC105372935-GGT2.NCBI.GGT2.rev.compl                 gaacagctaaccacacaacacagcatactgaatgaggagcctcgcaggctgagaagaaag   78158

chimp.LOC112206744.LOC107973052.GGT2-.646279-708040.rev.compl   ------------------------------------------------------------   61762
FAM230B-LOC105372935-GGT2.NCBI.GGT2.rev.compl                 caaacccggaaggcaacaaatggatacgtgcatgctgaaggactcgtgtttaattcttcc   78218

chimp.LOC112206744.LOC107973052.GGT2-.646279-708040.rev.compl   ------------------------------------------------------------   61762
FAM230B-LOC105372935-GGT2.NCBI.GGT2.rev.compl                 cggtaatcctttgtcaggcatatatttttgtctcctcaaacttaagaaagaaatccatgc   78278

chimp.LOC112206744.LOC107973052.GGT2-.646279-708040.rev.compl   ------------------------------------------------------------   61762
FAM230B-LOC105372935-GGT2.NCBI.GGT2.rev.compl                 tctggaggggagaaagagaggctgtactcattaactcagagctgacctcgtctggcagga   78338

chimp.LOC112206744.LOC107973052.GGT2-.646279-708040.rev.compl   ------------------------------------------------------------   61762
FAM230B-LOC105372935-GGT2.NCBI.GGT2.rev.compl                 ttaactcgccatacctgtccagcccacagaacctggcctggagcatggagggagaaccac   78398

chimp.LOC112206744.LOC107973052.GGT2-.646279-708040.rev.compl   ------------------------------------------------------------   61762
FAM230B-LOC105372935-GGT2.NCBI.GGT2.rev.compl                 agccaactccagaaaaataaaaggctgcagcgagaaacaagtggggccacagagagaaag   78458

chimp.LOC112206744.LOC107973052.GGT2-.646279-708040.rev.compl   ------------------------------------------------------------   61762
FAM230B-LOC105372935-GGT2.NCBI.GGT2.rev.compl                 gggttttgaaaaagttctaaatccccatggctagggacagtgacatcagtgtgtttacca   78518

chimp.LOC112206744.LOC107973052.GGT2-.646279-708040.rev.compl   ------------------------------------------------------------   61762
FAM230B-LOC105372935-GGT2.NCBI.GGT2.rev.compl                 agtatttgatttacttgctcaggcaaaattccagaagagacactgggacctgtgtggaga   78578

chimp.LOC112206744.LOC107973052.GGT2-.646279-708040.rev.compl   ------------------------------------------------------------   61762
FAM230B-LOC105372935-GGT2.NCBI.GGT2.rev.compl                 gtttcttcactgcagctatcctgccaagctcattgtaaagtgacgaattttgtggacact   78638

chimp.LOC112206744.LOC107973052.GGT2-.646279-708040.rev.compl   ------------------------------------------------------------   61762
FAM230B-LOC105372935-GGT2.NCBI.GGT2.rev.compl                 tacagagaaaaaatagcttcaaatgctatagaccaatgctgcacgctgcagagacacagt   78698

chimp.LOC112206744.LOC107973052.GGT2-.646279-708040.rev.compl   ------------------------------------------------------------   61762
FAM230B-LOC105372935-GGT2.NCBI.GGT2.rev.compl                 ctttgggttctgtgagctgtatttgggcattacgcagagttcagctgccatctgtccctt   78758

chimp.LOC112206744.LOC107973052.GGT2-.646279-708040.rev.compl   ------------------------------------------------------------   61762
FAM230B-LOC105372935-GGT2.NCBI.GGT2.rev.compl                 tgaaggccagctaagaagttggtcccagcctgggcatggtaccactgtcctacattccag   78818

chimp.LOC112206744.LOC107973052.GGT2-.646279-708040.rev.compl   ------------------------------------------------------------   61762
FAM230B-LOC105372935-GGT2.NCBI.GGT2.rev.compl                 ccatttgggatatcctgggttcaagcattccctcccattgctggatcaagcacttcccac   78878

chimp.LOC112206744.LOC107973052.GGT2-.646279-708040.rev.compl   ------------------------------------------------------------   61762
FAM230B-LOC105372935-GGT2.NCBI.GGT2.rev.compl                 ctcctgctctgggaaatcagatctcagtaggtcatcaaccatcccagaaagtccagcaga   78938

chimp.LOC112206744.LOC107973052.GGT2-.646279-708040.rev.compl   ------------------------------------------------------------   61762
FAM230B-LOC105372935-GGT2.NCBI.GGT2.rev.compl                 gttggttttggtttggggggtgtttttttgagttgggtggaagaagagaccttacataatct   78998

chimp.LOC112206744.LOC107973052.GGT2-.646279-708040.rev.compl   ------------------------------------------------------------   61762
FAM230B-LOC105372935-GGT2.NCBI.GGT2.rev.compl                 tttaactttttctctaaggctgaattccctataagcaaaattttaaattttttttaaag   79058

chimp.LOC112206744.LOC107973052.GGT2-.646279-708040.rev.compl   ------------------------------------------------------------   61762
FAM230B-LOC105372935-GGT2.NCBI.GGT2.rev.compl                 aagaaaaacacccagacaatccttaaaatcccctttcagctccaagactgtgtaattctg   79118

chimp.LOC112206744.LOC107973052.GGT2-.646279-708040.rev.compl   ------------------------------------------------------------   61762
FAM230B-LOC105372935-GGT2.NCBI.GGT2.rev.compl                 tgataccagtattgaaagcccctgcatttccctaactatatattcccctatgccagggaa   79178

chimp.LOC112206744.LOC107973052.GGT2-.646279-708040.rev.compl   ------------------------------------------------------------   61762
FAM230B-LOC105372935-GGT2.NCBI.GGT2.rev.compl                 aggtgcatgtaagtgagagaacaaaaaatttcacaatcgtggggacctcttctttcggct   79238

chimp.LOC112206744.LOC107973052.GGT2-.646279-708040.rev.compl   ------------------------------------------------------------   61762
FAM230B-LOC105372935-GGT2.NCBI.GGT2.rev.compl                 ttggagccccccttccctctgtctctgtacggggcagctttttccttctgtcttctccctt   79298

chimp.LOC112206744.LOC107973052.GGT2-.646279-708040.rev.compl   ------------------------------------------------------------   61762
FAM230B-LOC105372935-GGT2.NCBI.GGT2.rev.compl                 ccttcttgtctactaaactctctgcgccttagaaccaaaaataaaaattcaaaatcatga   79358

chimp.LOC112206744.LOC107973052.GGT2-.646279-708040.rev.compl   ------------------------------------------------------------   61762
FAM230B-LOC105372935-GGT2.NCBI.GGT2.rev.compl                 tcttaggggtgaaagaacactgagaaaccatagaaagcaaaaacgaaaagccaacataga   79418

chimp.LOC112206744.LOC107973052.GGT2-.646279-708040.rev.compl   ------------------------------------------------------------   61762
FAM230B-LOC105372935-GGT2.NCBI.GGT2.rev.compl                 aaaagtctgacttgaattagaagcagcacttaaaattacaggttcagaaaaagcttgaac   79478

chimp.LOC112206744.LOC107973052.GGT2-.646279-708040.rev.compl   ------------------------------------------------------------   61762
FAM230B-LOC105372935-GGT2.NCBI.GGT2.rev.compl                 agaatccagggctgaggtcagccctgagtctgcaggctctgccctctcacaaaggtgagt   79538

chimp.LOC112206744.LOC107973052.GGT2-.646279-708040.rev.compl   ------------------------------------------------------------   61762
FAM230B-LOC105372935-GGT2.NCBI.GGT2.rev.compl                 agatgggtctggagagggctgttcatgctatggaagggcctctatgccatcgaggctaaa   79598

chimp.LOC112206744.LOC107973052.GGT2-.646279-708040.rev.compl   ------------------------------------------------------------   61762
FAM230B-LOC105372935-GGT2.NCBI.GGT2.rev.compl                 accaaagcaggagcaaatcagccctgctagtgtgtacagtcggcatcaaagaggcaacaa   79658

chimp.LOC112206744.LOC107973052.GGT2-.646279-708040.rev.compl   ------------------------------------------------------------   61762
FAM230B-LOC105372935-GGT2.NCBI.GGT2.rev.compl                 cggtagagaatatttctgaaagacacacatgaaaccagtaacggcgatggtctctggaa   79718
```

```
chimp.LOC112206744.LOC107973052.GGT2-.646279-708040.rev.compl    ------------------------------------------------------------    61762
FAM230B-LOC105372935-GGT2.NCBI.GGT2.rev.compl                    gaattaggtcagaagcagacttaatttgcactgtgtacattctttggtatcctttttaat    79778

chimp.LOC112206744.LOC107973052.GGT2-.646279-708040.rev.compl    ------------------------------------------------------------    61762
FAM230B-LOC105372935-GGT2.NCBI.GGT2.rev.compl                    tgaacgcatgtgaatgcattacctattcaaaaataaatgtgtaaaaacacatgccacgac    79838

chimp.LOC112206744.LOC107973052.GGT2-.646279-708040.rev.compl    ------------------------------------------------------------    61762
FAM230B-LOC105372935-GGT2.NCBI.GGT2.rev.compl                    aaagcaaaatgaggagacagtataaaaaagacaaggataattttaaatttctcctaataa    79898

chimp.LOC112206744.LOC107973052.GGT2-.646279-708040.rev.compl    ------------------------------------------------------------    61762
FAM230B-LOC105372935-GGT2.NCBI.GGT2.rev.compl                    aacgctgtttaaaaatctaagaggaaaaaagggtaggctccagctagtggagagaacaac    79958

chimp.LOC112206744.LOC107973052.GGT2-.646279-708040.rev.compl    ------------------------------------------------------------    61762
FAM230B-LOC105372935-GGT2.NCBI.GGT2.rev.compl                    tgaaaaacaagagccacgtgttcccatggtcctggccaaccccacttctttcccatcctc    80018

chimp.LOC112206744.LOC107973052.GGT2-.646279-708040.rev.compl    ------------------------------------------------------------    61762
FAM230B-LOC105372935-GGT2.NCBI.GGT2.rev.compl                    actcgttcacatctgccttggagcacccacagcaagcctggcttcatgcccagaacatca    80078

chimp.LOC112206744.LOC107973052.GGT2-.646279-708040.rev.compl    ------------------------------------------------------------    61762
FAM230B-LOC105372935-GGT2.NCBI.GGT2.rev.compl                    cctggctttcctcctctctccctgcccacatctcctcagtcactcgctgcctcctctctt    80138

chimp.LOC112206744.LOC107973052.GGT2-.646279-708040.rev.compl    ------------------------------------------------------------    61762
FAM230B-LOC105372935-GGT2.NCBI.GGT2.rev.compl                    gcttctctcccagttctcagtttgcccatcaaatctcaggtggtactctccattctctgc    80198

chimp.LOC112206744.LOC107973052.GGT2-.646279-708040.rev.compl    ------------------------------------------------------------    61762
FAM230B-LOC105372935-GGT2.NCBI.GGT2.rev.compl                    tatgagggagttcatttgttcatgcaatttccccaccggagacaaaatgtatttcccaaa    80258

chimp.LOC112206744.LOC107973052.GGT2-.646279-708040.rev.compl    ------------------------------------------------------------    61762
FAM230B-LOC105372935-GGT2.NCBI.GGT2.rev.compl                    gttggcttgcaatgacatctcccatgcctgctctgcttacccatgactttgacactcctt    80318

chimp.LOC112206744.LOC107973052.GGT2-.646279-708040.rev.compl    ------------------------------------------------------------    61762
FAM230B-LOC105372935-GGT2.NCBI.GGT2.rev.compl                    ccatcgagagggcagaacctgtgtttcctctctctgaatctaggtgggctacagctctgg    80378

chimp.LOC112206744.LOC107973052.GGT2-.646279-708040.rev.compl    ------------------------------------------------------------    61762
FAM230B-LOC105372935-GGT2.NCBI.GGT2.rev.compl                    tagaagtgacactctgtgaccttaaggctaggtcacaaaaagggaatgcagcttctgcca    80438

chimp.LOC112206744.LOC107973052.GGT2-.646279-708040.rev.compl    ------------------------------------------------------------    61762
FAM230B-LOC105372935-GGT2.NCBI.GGT2.rev.compl                    gcaatcctgggacacacacttggagcccgagctgctgtgtaagccatctgcctgctctga    80498

chimp.LOC112206744.LOC107973052.GGT2-.646279-708040.rev.compl    ------------------------------------------------------------    61762
FAM230B-LOC105372935-GGT2.NCBI.GGT2.rev.compl                    gcccaccacactataaggaagcccaagctagcccacaggagagacgaggtgaagagaatg    80558

chimp.LOC112206744.LOC107973052.GGT2-.646279-708040.rev.compl    ------------------------------------------------------------    61762
FAM230B-LOC105372935-GGT2.NCBI.GGT2.rev.compl                    agatgccagtctccactgctccagctctctagtgtcccagccccagccactatctgactg    80618

chimp.LOC112206744.LOC107973052.GGT2-.646279-708040.rev.compl    ------------------------------------------------------------    61762
FAM230B-LOC105372935-GGT2.NCBI.GGT2.rev.compl                    caaccccatgaaagactgagccagaaccacgagcccagcccttcctgatgttgacataca    80678

chimp.LOC112206744.LOC107973052.GGT2-.646279-708040.rev.compl    ------------------------------------------------------------    61762
FAM230B-LOC105372935-GGT2.NCBI.GGT2.rev.compl                    gacttcatgaaagagaataaactaattttgctttaagccactatgatttgggatgattt    80738

chimp.LOC112206744.LOC107973052.GGT2-.646279-708040.rev.compl    ------------------------------------------------------------    61762
FAM230B-LOC105372935-GGT2.NCBI.GGT2.rev.compl                    gttatgcagcactggataactgcaacacctgcttctaaacagtagaatcccaaacgtcta    80798

chimp.LOC112206744.LOC107973052.GGT2-.646279-708040.rev.compl    ------------------------------------------------------------    61762
FAM230B-LOC105372935-GGT2.NCBI.GGT2.rev.compl                    ggccccatctctaacttacttcatgttcctgttctcactgatttttgccactgagcttg    80858

chimp.LOC112206744.LOC107973052.GGT2-.646279-708040.rev.compl    ------------------------------------------------------------    61762
FAM230B-LOC105372935-GGT2.NCBI.GGT2.rev.compl                    accaagcccttgagtcaccaattccagatatttgtttaaacaaattcactgtgactgtct    80918

chimp.LOC112206744.LOC107973052.GGT2-.646279-708040.rev.compl    ------------------------------------------------------------    61762
FAM230B-LOC105372935-GGT2.NCBI.GGT2.rev.compl                    acactatccattcctccaaaacaatccctgttatagttggccttccccagttctcttcct    80978

chimp.LOC112206744.LOC107973052.GGT2-.646279-708040.rev.compl    ------------------------------------------------------------    61762
FAM230B-LOC105372935-GGT2.NCBI.GGT2.rev.compl                    ggcatagagtagtctgtgcccttctctcatttcttaatatgaccctcacccatggagtgt    81038

chimp.LOC112206744.LOC107973052.GGT2-.646279-708040.rev.compl    ------------------------------------------------------------    61762
FAM230B-LOC105372935-GGT2.NCBI.GGT2.rev.compl                    tcctagattctgctcattcataatgcttcttcacccaccctaacctctccctcctgtagc    81098

chimp.LOC112206744.LOC107973052.GGT2-.646279-708040.rev.compl    ------------------------------------------------------------    61762
FAM230B-LOC105372935-GGT2.NCBI.GGT2.rev.compl                    cacagatcatccacttttccaggcctctgcactggaggctgcatcagccacctcccagta    81158

chimp.LOC112206744.LOC107973052.GGT2-.646279-708040.rev.compl    ------------------------------------------------------------    61762
FAM230B-LOC105372935-GGT2.NCBI.GGT2.rev.compl                    cgtcatcctttccaatgccctacaacacggggcagaaaggactctttttgctgcacatc    81218

chimp.LOC112206744.LOC107973052.GGT2-.646279-708040.rev.compl    ------------------------------------------------------------    61762
FAM230B-LOC105372935-GGT2.NCBI.GGT2.rev.compl                    caggctgtttaagaaaaggcttaaggcagcttaagcacaaagaatgtcatcttcaaacat    81278

chimp.LOC112206744.LOC107973052.GGT2-.646279-708040.rev.compl    ------------------------------------------------------------    61762
FAM230B-LOC105372935-GGT2.NCBI.GGT2.rev.compl                    tgtaacggcatcttctttttggggcactgaagtattcccatgattccaagaatcaaatga    81338

chimp.LOC112206744.LOC107973052.GGT2-.646279-708040.rev.compl    ------------------------------------------------------------    61762
FAM230B-LOC105372935-GGT2.NCBI.GGT2.rev.compl                    acagcatctggttatggggtctctggtctctgccaaggcaaactcataaagcatttcaga    81398

chimp.LOC112206744.LOC107973052.GGT2-.646279-708040.rev.compl    ------------------------------------------------------------    61762
FAM230B-LOC105372935-GGT2.NCBI.GGT2.rev.compl                    gcccttcatctggaaaggccttttctccaccccacgcccagccaatgcccactcatactt    81458

chimp.LOC112206744.LOC107973052.GGT2-.646279-708040.rev.compl    ------------------------------------------------------------    61762
FAM230B-LOC105372935-GGT2.NCBI.GGT2.rev.compl                    taaggcctagctcatccgtcatctctgctagggaaccttcctcagtgccttatgcaggca    81518

chimp.LOC112206744.LOC107973052.GGT2-.646279-708040.rev.compl    ------------------------------------------------------------    61762
FAM230B-LOC105372935-GGT2.NCBI.GGT2.rev.compl                    accattcctgccttgtgttgtgttgtgttgtgttgtgccttaagtcaaatccttaactca    81578

chimp.LOC112206744.LOC107973052.GGT2-.646279-708040.rev.compl    ------------------------------------------------------------    61762
FAM230B-LOC105372935-GGT2.NCBI.GGT2.rev.compl                    tctttgtgtgtccccttgcatttcagacagtaggtgcccaataaatgtttgtgggatgaa    81638

chimp.LOC112206744.LOC107973052.GGT2-.646279-708040.rev.compl    ------------------------------------------------------------    61762
FAM230B-LOC105372935-GGT2.NCBI.GGT2.rev.compl                    tcaactggttattctcctggaacaccctttcccagtacacctccactcgccagaacaact    81698

chimp.LOC112206744.LOC107973052.GGT2-.646279-708040.rev.compl    ------------------------------------------------------------    61762
FAM230B-LOC105372935-GGT2.NCBI.GGT2.rev.compl                    gtcttcctcacaggtgccagccatcaccccgaccacaccctctcaaaaggcctttgccag    81758

chimp.LOC112206744.LOC107973052.GGT2-.646279-708040.rev.compl    ------------------------------------------------------------    61762
FAM230B-LOC105372935-GGT2.NCBI.GGT2.rev.compl                    tcatcaaacaaagcctcttgtctcctgagagccgctccacaatcaccttcccaagaagaa    81818

chimp.LOC112206744.LOC107973052.GGT2-.646279-708040.rev.compl    ------------------------------------------------------------    61762
FAM230B-LOC105372935-GGT2.NCBI.GGT2.rev.compl                    ttccccataccgagtccaccattccttccctgatcaaagttgctttcacaggcatctttt    81878
```

```
chimp.LOC112206744.LOC107973052.GGT2-.646279-708040.rev.compl      ------------------------------------------------------------   61762
FAM230B-LOC105372935-GGT2.NCBI.GGT2.rev.compl                      cagctatccactcttcacggcttggttttcatgcactctgagttggccctatctcccctt   81938

chimp.LOC112206744.LOC107973052.GGT2-.646279-708040.rev.compl      ------------------------------------------------------------   61762
FAM230B-LOC105372935-GGT2.NCBI.GGT2.rev.compl                      tttggcctcaagcacgtcaagaggaaggctggtgacagcctttcccgagctcaggacagt   81998

chimp.LOC112206744.LOC107973052.GGT2-.646279-708040.rev.compl      ------------------------------------------------------------   61762
FAM230B-LOC105372935-GGT2.NCBI.GGT2.rev.compl                      tttcatgcaaactggttcttccaggttcttatgcctggaagaactgctgctctccaatag   82058

chimp.LOC112206744.LOC107973052.GGT2-.646279-708040.rev.compl      ------------------------------------------------------------   61762
FAM230B-LOC105372935-GGT2.NCBI.GGT2.rev.compl                      catgaagtagacaatgatgttcttagcatccagtcttaaaactagaaaaagtattggcac   82118

chimp.LOC112206744.LOC107973052.GGT2-.646279-708040.rev.compl      ------------------------------------------------------------   61762
FAM230B-LOC105372935-GGT2.NCBI.GGT2.rev.compl                      tgctggatccccagctcctagaatggtgtgtggcacctaaacagagctcaaaaaatatta   82178

chimp.LOC112206744.LOC107973052.GGT2-.646279-708040.rev.compl      ------------------------------------------------------------   61762
FAM230B-LOC105372935-GGT2.NCBI.GGT2.rev.compl                      gttgaatggaggaaagaatgaacagtcatctggcccagacttggagaagctgtgatctgt   82238

chimp.LOC112206744.LOC107973052.GGT2-.646279-708040.rev.compl      ------------------------------------------------------------   61762
FAM230B-LOC105372935-GGT2.NCBI.GGT2.rev.compl                      ctagcccttctacaccaggagcgaggcctgaaaagctgcatgactcacccaaggtcacaa   82298

chimp.LOC112206744.LOC107973052.GGT2-.646279-708040.rev.compl      ------------------------------------------------------------   61762
FAM230B-LOC105372935-GGT2.NCBI.GGT2.rev.compl                      gcaagccagtgaagaggggctggaaccctgtcttttgcctcaaagcacagggccctcctt   82358

chimp.LOC112206744.LOC107973052.GGT2-.646279-708040.rev.compl      ------------------------------------------------------------   61762
FAM230B-LOC105372935-GGT2.NCBI.GGT2.rev.compl                      ctgtcccacccagctttctcaccacccaccccaccccttccacactccaaagacagcatc   82418

chimp.LOC112206744.LOC107973052.GGT2-.646279-708040.rev.compl      ------------------------------------------------------------   61762
FAM230B-LOC105372935-GGT2.NCBI.GGT2.rev.compl                      tccgtgagaagtgttgggcagtgtccaggagtgtttttcttcctggctttaaaacaaatt   82478

chimp.LOC112206744.LOC107973052.GGT2-.646279-708040.rev.compl      ------------------------------------------------------------   61762
FAM230B-LOC105372935-GGT2.NCBI.GGT2.rev.compl                      gtgacaaatggctccacagatctaccgggccatggagctgagcagatgggactgcaattc   82538

chimp.LOC112206744.LOC107973052.GGT2-.646279-708040.rev.compl      ------------------------------------------------------------   61762
FAM230B-LOC105372935-GGT2.NCBI.GGT2.rev.compl                      acgctattaactcaaattctacttgactttctttcaaatcagggcattttatctagaaat   82598

chimp.LOC112206744.LOC107973052.GGT2-.646279-708040.rev.compl      ------------------------------------------------------------   61762
FAM230B-LOC105372935-GGT2.NCBI.GGT2.rev.compl                      tataaaaaattgttgggggagtgcctttgtaaggaagtttgccagcacattcatttacaa   82658

chimp.LOC112206744.LOC107973052.GGT2-.646279-708040.rev.compl      ------------------------------------------------------------   61762
FAM230B-LOC105372935-GGT2.NCBI.GGT2.rev.compl                      atcgacttgctcctggccaccctcactgatcacggacttttactcacgctcccttactct   82718

chimp.LOC112206744.LOC107973052.GGT2-.646279-708040.rev.compl      ------------------------------------------------------------   61762
FAM230B-LOC105372935-GGT2.NCBI.GGT2.rev.compl                      ctcccatcaaagagtccagcgcatggagagagtgatggaatcttgaggcccaccggtcta   82778

chimp.LOC112206744.LOC107973052.GGT2-.646279-708040.rev.compl      ------------------------------------------------------------   61762
FAM230B-LOC105372935-GGT2.NCBI.GGT2.rev.compl                      accacccacctaagcttgactcccctgtgcctgtggacatcctgcgtgcagatgcacatt   82838

chimp.LOC112206744.LOC107973052.GGT2-.646279-708040.rev.compl      ------------------------------------------------------------   61762
FAM230B-LOC105372935-GGT2.NCBI.GGT2.rev.compl                      cccactaatggcagccaggggaaagcactattcgaaagttcttctttactttgaactcaa   82898

chimp.LOC112206744.LOC107973052.GGT2-.646279-708040.rev.compl      ------------------------------------------------------------   61762
FAM230B-LOC105372935-GGT2.NCBI.GGT2.rev.compl                      attgggctcaggggtcctggatacaggacaaccccaatctcttatttattcaatggacaa   82958

chimp.LOC112206744.LOC107973052.GGT2-.646279-708040.rev.compl      ------------------------------------------------------------   61762
FAM230B-LOC105372935-GGT2.NCBI.GGT2.rev.compl                      caggtgagcacatactaggcctggacactgatgtaggtaggcactaaagcttcagagaga   83018

chimp.LOC112206744.LOC107973052.GGT2-.646279-708040.rev.compl      ------------------------------------------------------------   61762
FAM230B-LOC105372935-GGT2.NCBI.GGT2.rev.compl                      tgcaaatgaacaatctttaccctgaaggagctcatgttctatgggttcatttcctttccc   83078

chimp.LOC112206744.LOC107973052.GGT2-.646279-708040.rev.compl      ------------------------------------------------------------   61762
FAM230B-LOC105372935-GGT2.NCBI.GGT2.rev.compl                      tgtgtcttacaaacaggttccagtttgtttccgtccttcagagtgtactgcctagaactg   83138

chimp.LOC112206744.LOC107973052.GGT2-.646279-708040.rev.compl      ------------------------------------------------------------   61762
FAM230B-LOC105372935-GGT2.NCBI.GGT2.rev.compl                      catgctatactccacaagtggcctgaccagcttgtggacaaggatgaagcaagcagacca   83198

chimp.LOC112206744.LOC107973052.GGT2-.646279-708040.rev.compl      ------------------------------------------------------------   61762
FAM230B-LOC105372935-GGT2.NCBI.GGT2.rev.compl                      ccttctttcagtgcagacaagatggctgtttagaggccacatcacgtggcagacacctgc   83258

chimp.LOC112206744.LOC107973052.GGT2-.646279-708040.rev.compl      ------------------------------------------------------------   61762
FAM230B-LOC105372935-GGT2.NCBI.GGT2.rev.compl                      caactaaaaccctccatccttcccacagcctcccccctcctacactttgcagctggattca   83318

chimp.LOC112206744.LOC107973052.GGT2-.646279-708040.rev.compl      ------------------------------------------------------------   61762
FAM230B-LOC105372935-GGT2.NCBI.GGT2.rev.compl                      aaaccttgcattcacccttgttccatcaacagggccaggccatcagtctatctcatccgg   83378

chimp.LOC112206744.LOC107973052.GGT2-.646279-708040.rev.compl      ------------------------------------------------------------   61762
FAM230B-LOC105372935-GGT2.NCBI.GGT2.rev.compl                      actgtttttatcttacttgccaacctacacacttgctgttccttcttgccttaaagtca   83438

chimp.LOC112206744.LOC107973052.GGT2-.646279-708040.rev.compl      ------------------------------------------------------------   61762
FAM230B-LOC105372935-GGT2.NCBI.GGT2.rev.compl                      tacagatgactttaggtgattcagatgagtgtgacctctactttcaccaaacacaatagt   83498

chimp.LOC112206744.LOC107973052.GGT2-.646279-708040.rev.compl      ------------------------------------------------------------   61762
FAM230B-LOC105372935-GGT2.NCBI.GGT2.rev.compl                      caattcagaaaggtcaaggacagaaccctccagcatgctaacgcagccaaagaagtccga   83558

chimp.LOC112206744.LOC107973052.GGT2-.646279-708040.rev.compl      ------------------------------------------------------------   61762
FAM230B-LOC105372935-GGT2.NCBI.GGT2.rev.compl                      aaacaatacttggccacactgttttcctgcttacggatgtggactgctcttgtcctgcct   83618

chimp.LOC112206744.LOC107973052.GGT2-.646279-708040.rev.compl      ------------------------------------------------------------   61762
FAM230B-LOC105372935-GGT2.NCBI.GGT2.rev.compl                      gtaaatatatcggatcgctctccttcttgctggcatccacagtaactatactcaataaac   83678

chimp.LOC112206744.LOC107973052.GGT2-.646279-708040.rev.compl      ------------------------------------------------------------   61762
FAM230B-LOC105372935-GGT2.NCBI.GGT2.rev.compl                      acaccaactttgatgccaaatcctaggccatcccacacatcttaaaaataaaacaaaaat   83738

chimp.LOC112206744.LOC107973052.GGT2-.646279-708040.rev.compl      ------------------------------------------------------------   61762
FAM230B-LOC105372935-GGT2.NCBI.GGT2.rev.compl                      atgctattccagctaccctcactctttttgtggtttcctgcagtccaactagctacaaat   83798

chimp.LOC112206744.LOC107973052.GGT2-.646279-708040.rev.compl      ------------------------------------------------------------   61762
FAM230B-LOC105372935-GGT2.NCBI.GGT2.rev.compl                      ctcaaagccaaagattgtagctccctctcttttgttctttcatcaagccagggagtcctg   83858

chimp.LOC112206744.LOC107973052.GGT2-.646279-708040.rev.compl      ------------------------------------------------------------   61762
FAM230B-LOC105372935-GGT2.NCBI.GGT2.rev.compl                      ctcctctctctctagtctctcctgtgtcagtgccacttgtcaactgcctaatccagccac   83918

chimp.LOC112206744.LOC107973052.GGT2-.646279-708040.rev.compl      ------------------------------------------------------------   61762
FAM230B-LOC105372935-GGT2.NCBI.GGT2.rev.compl                      gctgctcttcaatattaaaggtcttcagcatattttaaagttagtctcctactgccactc   83978

chimp.LOC112206744.LOC107973052.GGT2-.646279-708040.rev.compl      ------------------------------------------------------------   61762
FAM230B-LOC105372935-GGT2.NCBI.GGT2.rev.compl                      ttttccctactcttgccagtccactcagtaaactactgccatacaggtctcccacagcag   84038
```

```
chimp.LOC112206744.LOC107973052.GGT2-.646279-708040.rev.compl      ------------------------------------------------------------   61762
FAM230B-LOC105372935-GGT2.NCBI.GGT2.rev.compl                      tttctcaccagataaacaatggccggagattccactttgccaatcacatcaagtccaaac   84098

chimp.LOC112206744.LOC107973052.GGT2-.646279-708040.rev.compl      ------------------------------------------------------------   61762
FAM230B-LOC105372935-GGT2.NCBI.GGT2.rev.compl                      tccgtgtcttggtatccaaggcttctaaagtctggccccactccacctactcactctgac   84158

chimp.LOC112206744.LOC107973052.GGT2-.646279-708040.rev.compl      ------------------------------------------------------------   61762
FAM230B-LOC105372935-GGT2.NCBI.GGT2.rev.compl                      acccaccactccctccaaacatccccggtacgtcttactggattttgccatttcctccaa   84218

chimp.LOC112206744.LOC107973052.GGT2-.646279-708040.rev.compl      ------------------------------------------------------------   61762
FAM230B-LOC105372935-GGT2.NCBI.GGT2.rev.compl                      gttgaccaggagtcattcacacctaaaggctatgcttacatcgttttccatctcttgtgt   84278

chimp.LOC112206744.LOC107973052.GGT2-.646279-708040.rev.compl      ------------------------------------------------------------   61762
FAM230B-LOC105372935-GGT2.NCBI.GGT2.rev.compl                      tggtgagctaaaaacaagatggaaaagaaaaaagaagaggaagagagaaaaagagaaaag   84338

chimp.LOC112206744.LOC107973052.GGT2-.646279-708040.rev.compl      ------------------------------------------------------------   61762
FAM230B-LOC105372935-GGT2.NCBI.GGT2.rev.compl                      gaagaaatggggaaaacagcaataggaccagctaactcctacagcagagtcttgccaggc   84398

chimp.LOC112206744.LOC107973052.GGT2-.646279-708040.rev.compl      ------------------------------------------------------------   61762
FAM230B-LOC105372935-GGT2.NCBI.GGT2.rev.compl                      cctcccctaattgctttatatgtgttacaatgtttaaccctcacaactacaacaggagtt   84458

chimp.LOC112206744.LOC107973052.GGT2-.646279-708040.rev.compl      ------------------------------------------------------------   61762
FAM230B-LOC105372935-GGT2.NCBI.GGT2.rev.compl                      gggtaccattccatccccacctgcagatgagtgtggcaccccaagattaagtatctgccc   84518

chimp.LOC112206744.LOC107973052.GGT2-.646279-708040.rev.compl      ------------------------------------------------------------   61762
FAM230B-LOC105372935-GGT2.NCBI.GGT2.rev.compl                      agggtcacgtgaaaagtctgcatgcatgctgaaccattccagctgcaccactgcccagga   84578

chimp.LOC112206744.LOC107973052.GGT2-.646279-708040.rev.compl      ------------------------------------------------------------   61762
FAM230B-LOC105372935-GGT2.NCBI.GGT2.rev.compl                      aaaaagtcacaggtaggaagacaagcatagaattaagaacaaagcaactaactgtaccag   84638

chimp.LOC112206744.LOC107973052.GGT2-.646279-708040.rev.compl      ------------------------------------------------------------   61762
FAM230B-LOC105372935-GGT2.NCBI.GGT2.rev.compl                      gtgctacagaagggtggtcagcctggaaaatcaggaggtgcattccttgggagattgccg   84698

chimp.LOC112206744.LOC107973052.GGT2-.646279-708040.rev.compl      ------------------------------------------------------------   61762
FAM230B-LOC105372935-GGT2.NCBI.GGT2.rev.compl                      caatcagagtctgggagtatgcaggagactggcaggcagcccagggacaggtgaggcaca   84758

chimp.LOC112206744.LOC107973052.GGT2-.646279-708040.rev.compl      ------------------------------------------------------------   61762
FAM230B-LOC105372935-GGT2.NCBI.GGT2.rev.compl                      ggcactcattcccttccatgtgagaagtcagggcacgtgcaaagccaaggccagaggaca   84818

chimp.LOC112206744.LOC107973052.GGT2-.646279-708040.rev.compl      ------------------------------------------------------------   61762
FAM230B-LOC105372935-GGT2.NCBI.GGT2.rev.compl                      aaggtgagctcgccaatagggccaatactgggtctggctggaaggggagggtggacagag   84878

chimp.LOC112206744.LOC107973052.GGT2-.646279-708040.rev.compl      ------------------------------------------------------------   61762
FAM230B-LOC105372935-GGT2.NCBI.GGT2.rev.compl                      atgggtactaagaagtgaagatggacaggcaggaccaggtcttaaagggtttgtgtcttg   84938

chimp.LOC112206744.LOC107973052.GGT2-.646279-708040.rev.compl      ------------------------------------------------------------   61762
FAM230B-LOC105372935-GGT2.NCBI.GGT2.rev.compl                      ctaaagatggacccttaattccgtgacacatctattcaaccactcctcattatctaccta   84998

chimp.LOC112206744.LOC107973052.GGT2-.646279-708040.rev.compl      ------------------------------------------------------------   61762
FAM230B-LOC105372935-GGT2.NCBI.GGT2.rev.compl                      aattctactgtgcctttgcaaagcccagctcaagtcctttttcctctcaggcctgtcctc   85058

chimp.LOC112206744.LOC107973052.GGT2-.646279-708040.rev.compl      ------------------------------------------------------------   61762
FAM230B-LOC105372935-GGT2.NCBI.GGT2.rev.compl                      cctactcctgagcaaaccacgggtgtggagacaggtcccaaccacctgcagatgacagaa   85118

chimp.LOC112206744.LOC107973052.GGT2-.646279-708040.rev.compl      ------------------------------------------------------------   61762
FAM230B-LOC105372935-GGT2.NCBI.GGT2.rev.compl                      ggaagctgcatctaggatcggtctacacagaggcaagaagggatcctcctcctctgcctt   85178

chimp.LOC112206744.LOC107973052.GGT2-.646279-708040.rev.compl      ------------------------------------------------------------   61762
FAM230B-LOC105372935-GGT2.NCBI.GGT2.rev.compl                      ctcctttcatgctgttcttactttctctcaaaaagaaagaaaatacagtgtcattgcttc   85238

chimp.LOC112206744.LOC107973052.GGT2-.646279-708040.rev.compl      ------------------------------------------------------------   61762
FAM230B-LOC105372935-GGT2.NCBI.GGT2.rev.compl                      ttccctaagcttgttaagtcaaagcaagtcaaagcatcagagcttcattacttggctttt   85298

chimp.LOC112206744.LOC107973052.GGT2-.646279-708040.rev.compl      ------------------------------------------------------------   61762
FAM230B-LOC105372935-GGT2.NCBI.GGT2.rev.compl                      tctagctataagcgtctggaaaatatcaggagtcactggcctctaggaatccattaagtt   85358

chimp.LOC112206744.LOC107973052.GGT2-.646279-708040.rev.compl      ------------------------------------------------------------   61762
FAM230B-LOC105372935-GGT2.NCBI.GGT2.rev.compl                      aaaggagctccatgatctactatcataagatgacaaagaaaatatgcactttctttggga   85418

chimp.LOC112206744.LOC107973052.GGT2-.646279-708040.rev.compl      ------------------------------------------------------------   61762
FAM230B-LOC105372935-GGT2.NCBI.GGT2.rev.compl                      ggtcaagttgggcgaatcacttgaggtcaggagttcaagaccaatctggccaacatggtg   85478

chimp.LOC112206744.LOC107973052.GGT2-.646279-708040.rev.compl      ------------------------------------------------------------   61762
FAM230B-LOC105372935-GGT2.NCBI.GGT2.rev.compl                      aaaccccgtctctactgaaaacacaaaaattagctgggcttggtggcaggcacctgtaat   85538

chimp.LOC112206744.LOC107973052.GGT2-.646279-708040.rev.compl      ------------------------------------------------------------   61762
FAM230B-LOC105372935-GGT2.NCBI.GGT2.rev.compl                      cccagctactcaggagactgaggcaggagaatcgattgaacccttacccaggagacagag   85598

chimp.LOC112206744.LOC107973052.GGT2-.646279-708040.rev.compl      ------------------------------------------------------------   61762
FAM230B-LOC105372935-GGT2.NCBI.GGT2.rev.compl                      gttgcaatgagccgagatggcaccacggcactccagcctgagcaacagcgtgactccatc   85658

chimp.LOC112206744.LOC107973052.GGT2-.646279-708040.rev.compl      ------------------------------------------------------------   61762
FAM230B-LOC105372935-GGT2.NCBI.GGT2.rev.compl                      tcaaaaaaaaaaaaaagaaagaaagaaaggaaggaaggaagaaagaagaaaatatgcatt   85718

chimp.LOC112206744.LOC107973052.GGT2-.646279-708040.rev.compl      ------------------------------------------------------------   61762
FAM230B-LOC105372935-GGT2.NCBI.GGT2.rev.compl                      ttctcaggattaaataacaaaaaatgccaagtttcttagctccagtcaagaaaatgacat   85778

chimp.LOC112206744.LOC107973052.GGT2-.646279-708040.rev.compl      ------------------------------------------------------------   61762
FAM230B-LOC105372935-GGT2.NCBI.GGT2.rev.compl                      catgtgacaacagcagccattgcaagtttatcagaagatgaacaattacagatttttcat   85838

chimp.LOC112206744.LOC107973052.GGT2-.646279-708040.rev.compl      ------------------------------------------------------------   61762
FAM230B-LOC105372935-GGT2.NCBI.GGT2.rev.compl                      ttctgaataaggaaccatttacacagacacatgtataacaaatgtctgtgcacatccagg   85898

chimp.LOC112206744.LOC107973052.GGT2-.646279-708040.rev.compl      ------------------------------------------------------------   61762
FAM230B-LOC105372935-GGT2.NCBI.GGT2.rev.compl                      acttgtcaaatcttaacattttgccacacctggtttagtttttttatttttcctttcaag   85958

chimp.LOC112206744.LOC107973052.GGT2-.646279-708040.rev.compl      ------------------------------------------------------------   61762
FAM230B-LOC105372935-GGT2.NCBI.GGT2.rev.compl                      aaagaaagcattacaggctgtgcttcaatccctatttcacccccttttcaagccccactat   86018

chimp.LOC112206744.LOC107973052.GGT2-.646279-708040.rev.compl      ------------------------------------------------------------   61762
FAM230B-LOC105372935-GGT2.NCBI.GGT2.rev.compl                      tccacccatcccagaaatgacaacatcccgaatttggtatttaccactcttgcacattta   86078

chimp.LOC112206744.LOC107973052.GGT2-.646279-708040.rev.compl      ------------------------------------------------------------   61762
FAM230B-LOC105372935-GGT2.NCBI.GGT2.rev.compl                      tttttatgcttttactacctatgtatgtagccataaacaatatacagagtttttggatgc   86138

chimp.LOC112206744.LOC107973052.GGT2-.646279-708040.rev.compl      ------------------------------------------------------------   61762
FAM230B-LOC105372935-GGT2.NCBI.GGT2.rev.compl                      tataaaactgtttaaaatggcatcacactgtatgcacccttttgcatctgttttaattga   86198

chimp.LOC112206744.LOC107973052.GGT2-.646279-708040.rev.compl      ------------------------------------------------------------   61762
```

```
FAM230B-LOC105372935-GGT2.NCBI.GGT2.rev.compl                   caaaacatttttgagggtaatccttgttgacatatgtcactccagggcattcctttcaac   86258

chimp.LOC112206744.LOC107973052.GGT2-.646279-708040.rev.compl   ------------------------------------------------------------   61762
FAM230B-LOC105372935-GGT2.NCBI.GGT2.rev.compl                   tgctttataccatggtccactgtctggataatcattatctatagcacaatgtattcattt   86318

chimp.LOC112206744.LOC107973052.GGT2-.646279-708040.rev.compl   ------------------------------------------------------------   61762
FAM230B-LOC105372935-GGT2.NCBI.GGT2.rev.compl                   tcctactgataaatttttaagtggtttccaacttttcattgcaataaatattcttgtaaa   86378

chimp.LOC112206744.LOC107973052.GGT2-.646279-708040.rev.compl   ------------------------------------------------------------   61762
FAM230B-LOC105372935-GGT2.NCBI.GGT2.rev.compl                   tcttcgtccaaggttatcttcaaccacactaggcattgcccaaattgctctgcaacatca   86438

chimp.LOC112206744.LOC107973052.GGT2-.646279-708040.rev.compl   ------------------------------------------------------------   61762
FAM230B-LOC105372935-GGT2.NCBI.GGT2.rev.compl                   gtgcaccaatttacattgccaagatcagtgtttaagtgtttctattctttgatgtctttg   86498

chimp.LOC112206744.LOC107973052.GGT2-.646279-708040.rev.compl   ------------------------------------------------------------   61762
FAM230B-LOC105372935-GGT2.NCBI.GGT2.rev.compl                   tcaaccttggtactatttgactactacacttttcaccaatcagatataaggatgaaatgg   86558

chimp.LOC112206744.LOC107973052.GGT2-.646279-708040.rev.compl   ------------------------------------------------------------   61762
FAM230B-LOC105372935-GGT2.NCBI.GGT2.rev.compl                   tgccttcttggtattctgaaaattaattttaaatgggaaaataaattactaataagtgca   86618

chimp.LOC112206744.LOC107973052.GGT2-.646279-708040.rev.compl   ------------------------------------------------------------   61762
FAM230B-LOC105372935-GGT2.NCBI.GGT2.rev.compl                   gcttaatcaacaagagttttcattaacatgtgggaatacaccaggtgttccttggatttt   86678

chimp.LOC112206744.LOC107973052.GGT2-.646279-708040.rev.compl   ------------------------------------------------------------   61762
FAM230B-LOC105372935-GGT2.NCBI.GGT2.rev.compl                   cttgaaaattctacagggagggtacgggagtgacggtacagataagattggacatgagct   86738

chimp.LOC112206744.LOC107973052.GGT2-.646279-708040.rev.compl   ------------------------------------------------------------   61762
FAM230B-LOC105372935-GGT2.NCBI.GGT2.rev.compl                   gataattgttggaactaggtgatgagtgtatgagggttaattattctgctctctccactt   86798

chimp.LOC112206744.LOC107973052.GGT2-.646279-708040.rev.compl   ------------------------------------------------------------   61762
FAM230B-LOC105372935-GGT2.NCBI.GGT2.rev.compl                   tggtacatatttgaaatttttcacaatagaaattaaaagaagaacggggcgtggtggctc   86858

chimp.LOC112206744.LOC107973052.GGT2-.646279-708040.rev.compl   ------------------------------------------------------------   61762
FAM230B-LOC105372935-GGT2.NCBI.GGT2.rev.compl                   acgcctgtaattccagcactttggaaggccatggcagaggatcacttgaggtcaggaatt   86918

chimp.LOC112206744.LOC107973052.GGT2-.646279-708040.rev.compl   ------------------------------------------------------------   61762
FAM230B-LOC105372935-GGT2.NCBI.GGT2.rev.compl                   aaagaccagctggtcaacatggggaaaccccacctctactaaaaatacaaaaattagctg   86978

chimp.LOC112206744.LOC107973052.GGT2-.646279-708040.rev.compl   ------------------------------------------------------------   61762
FAM230B-LOC105372935-GGT2.NCBI.GGT2.rev.compl                   ggtatggtggtgtgtaatcccagctattgaggaggctgaggcacgagaatcgcttggacc   87038

chimp.LOC112206744.LOC107973052.GGT2-.646279-708040.rev.compl   ------------------------------------------------------------   61762
FAM230B-LOC105372935-GGT2.NCBI.GGT2.rev.compl                   caggaggcagaggttgcagtgagccaagatcgtgccactgcactctagcctgggcaacag   87098

chimp.LOC112206744.LOC107973052.GGT2-.646279-708040.rev.compl   ------------------------------------------------------------   61762
FAM230B-LOC105372935-GGT2.NCBI.GGT2.rev.compl                   ggcaagcctctgtctcaaaaaaaaaaaaaaaaaaaaaaagaagaagaagaagaaagaaaa   87158

chimp.LOC112206744.LOC107973052.GGT2-.646279-708040.rev.compl   ------------------------------------------------------------   61762
FAM230B-LOC105372935-GGT2.NCBI.GGT2.rev.compl                   caaagaagaaagaaagaaagaaaaagaaagaaagaaagaaaattaaaagaaggaaatgcc   87218

chimp.LOC112206744.LOC107973052.GGT2-.646279-708040.rev.compl   ------------------------------------------------------------   61762
FAM230B-LOC105372935-GGT2.NCBI.GGT2.rev.compl                   ataaagcatctggagcaaagaacatccaaagaagagcactgatcactgcagcagacagcc   87278

chimp.LOC112206744.LOC107973052.GGT2-.646279-708040.rev.compl   ------------------------------------------------------------   61762
FAM230B-LOC105372935-GGT2.NCBI.GGT2.rev.compl                   attcagaatgggcagagagcagagctggcaggaagagaggacaaaaggaagaccaggaag   87338

chimp.LOC112206744.LOC107973052.GGT2-.646279-708040.rev.compl   ------------------------------------------------------------   61762
FAM230B-LOC105372935-GGT2.NCBI.GGT2.rev.compl                   ggaagccaggaaaagagattttgcattgtgttattatttcaatcagaagacttaggaaca   87398

chimp.LOC112206744.LOC107973052.GGT2-.646279-708040.rev.compl   ------------------------------------------------------------   61762
FAM230B-LOC105372935-GGT2.NCBI.GGT2.rev.compl                   ttatttaattatcaaatttgtgtgtctgatgatcctttacaacactgcagcatcaaataa   87458

chimp.LOC112206744.LOC107973052.GGT2-.646279-708040.rev.compl   ------------------------------------------------------------   61762
FAM230B-LOC105372935-GGT2.NCBI.GGT2.rev.compl                   agtgttatcagatttctttctttttttttttttgagactgggtctgattctgtagccca   87518

chimp.LOC112206744.LOC107973052.GGT2-.646279-708040.rev.compl   ------------------------------------------------------------   61762
FAM230B-LOC105372935-GGT2.NCBI.GGT2.rev.compl                   ggctggagtgcaggggtataataatagctcactgcagcttcaaactcctgagtttgtttg   87578

chimp.LOC112206744.LOC107973052.GGT2-.646279-708040.rev.compl   ------------------------------------------------------------   61762
FAM230B-LOC105372935-GGT2.NCBI.GGT2.rev.compl                   tttgtttgtctgtttgtttgtttgtttggagagacagcgtctcgaactcctgggctccag   87638

chimp.LOC112206744.LOC107973052.GGT2-.646279-708040.rev.compl   ------------------------------------------------------------   61762
FAM230B-LOC105372935-GGT2.NCBI.GGT2.rev.compl                   tgatcctcccacctcagcctcccaaagtgctgggattacaagcatgagccactgtgcccc   87698

chimp.LOC112206744.LOC107973052.GGT2-.646279-708040.rev.compl   ------------------------------------------------------------   61762
FAM230B-LOC105372935-GGT2.NCBI.GGT2.rev.compl                   agcccaacatctgatttcaaaagtactattgatattataatcaccattaaaaatcactga   87758

chimp.LOC112206744.LOC107973052.GGT2-.646279-708040.rev.compl   ------------------------------------------------------------   61762
FAM230B-LOC105372935-GGT2.NCBI.GGT2.rev.compl                   agtttttgtcacccaaagttttacgttgaaatatagaacatttcatttttttcccacaca   87818

chimp.LOC112206744.LOC107973052.GGT2-.646279-708040.rev.compl   ------------------------------------------------------------   61762
FAM230B-LOC105372935-GGT2.NCBI.GGT2.rev.compl                   cctcacatctgcttttgatttttaacatgtgcatgaattacctcttattagaaaaaaaaa   87878

chimp.LOC112206744.LOC107973052.GGT2-.646279-708040.rev.compl   ------------------------------------------------------------   61762
FAM230B-LOC105372935-GGT2.NCBI.GGT2.rev.compl                   cacaaaatttttttggtaaaagaattttccttaagtgggtatttttttccctaagtctgaaa   87938

chimp.LOC112206744.LOC107973052.GGT2-.646279-708040.rev.compl   ------------------------------------------------------------   61762
FAM230B-LOC105372935-GGT2.NCBI.GGT2.rev.compl                   tgacttagtaaggataagaggtcaaaccatatgctttgtagtgaaactgcaactatgatg   87998

chimp.LOC112206744.LOC107973052.GGT2-.646279-708040.rev.compl   ------------------------------------------------------------   61762
FAM230B-LOC105372935-GGT2.NCBI.GGT2.rev.compl                   aataacattagaattctggatgccttccagcagctggaacagggcagacctgaggcggga   88058

chimp.LOC112206744.LOC107973052.GGT2-.646279-708040.rev.compl   ------------------------------------------------------------   61762
FAM230B-LOC105372935-GGT2.NCBI.GGT2.rev.compl                   gtcctcatgcttaggggaaggggctcaggcactgggagtcggtcagggctcccaggatgt   88118

chimp.LOC112206744.LOC107973052.GGT2-.646279-708040.rev.compl   ------------------------------------------------------------   61762
FAM230B-LOC105372935-GGT2.NCBI.GGT2.rev.compl                   aaaagatacacatgaaagaaacaaacagtgggactcgggccagtgcaaaatttggcctgg   88178

chimp.LOC112206744.LOC107973052.GGT2-.646279-708040.rev.compl   ------------------------------------------------------------   61762
FAM230B-LOC105372935-GGT2.NCBI.GGT2.rev.compl                   aactgagctaaaaactgtcatactggtgggaagaagcagattggttcaacagggtaacca   88238

chimp.LOC112206744.LOC107973052.GGT2-.646279-708040.rev.compl   ------------------------------------------------------------   61762
FAM230B-LOC105372935-GGT2.NCBI.GGT2.rev.compl                   aatgtgaaaaagaaaagcctcacaagaattaaaattgatggttggaactgggcgctccga   88298

chimp.LOC112206744.LOC107973052.GGT2-.646279-708040.rev.compl   ------------------------------------------------------------   61762
FAM230B-LOC105372935-GGT2.NCBI.GGT2.rev.compl                   ggctgtgggccggtgtcctgccccgaagggtttggctgccagcaggtgcagcgctgccct   88358

chimp.LOC112206744.LOC107973052.GGT2-.646279-708040.rev.compl   ------------------------------------------------------------   61762
FAM230B-LOC105372935-GGT2.NCBI.GGT2.rev.compl                   gggataccagctcctgggatgcgctgtagtatcatcaatcaccggtgctgctcgctgggc   88418
```

```
chimp.LOC112206744.LOC107973052.GGT2-.646279-708040.rev.compl      ------------------------------------------------------------   61762
FAM230B-LOC105372935-GGT2.NCBI.GGT2.rev.compl                      cttggaagcacacagtagtagcagatgtcagcaggggccaccagggccacctgccagtg   88478

chimp.LOC112206744.LOC107973052.GGT2-.646279-708040.rev.compl      ------------------------------------------------------------   61762
FAM230B-LOC105372935-GGT2.NCBI.GGT2.rev.compl                      agcccactgcttagctgagtcatggtcaagtttctctgcagattgttgcctgaaaatatt   88538

chimp.LOC112206744.LOC107973052.GGT2-.646279-708040.rev.compl      ------------------------------------------------------------   61762
FAM230B-LOC105372935-GGT2.NCBI.GGT2.rev.compl                      tcaccacctggaacctccctgtcccacccatcttcacttgcaatactcactcccctcacc   88598

chimp.LOC112206744.LOC107973052.GGT2-.646279-708040.rev.compl      ------------------------------------------------------------   61762
FAM230B-LOC105372935-GGT2.NCBI.GGT2.rev.compl                      tctgcccctaaagcaattattcattaaccacagggatccaccatccaggaacaaaagagg   88658

chimp.LOC112206744.LOC107973052.GGT2-.646279-708040.rev.compl      ------------------------------------------------------------   61762
FAM230B-LOC105372935-GGT2.NCBI.GGT2.rev.compl                      cttgaaagctctatctgtgcctgcacttcttggtccaaaaggcgcaggggggatgggggag   88718

chimp.LOC112206744.LOC107973052.GGT2-.646279-708040.rev.compl      ------------------------------------------------------------   61762
FAM230B-LOC105372935-GGT2.NCBI.GGT2.rev.compl                      aggcaccggggtcatggtgcaggctcggctgttgagcctcattccccaaacacaatgtga   88778

chimp.LOC112206744.LOC107973052.GGT2-.646279-708040.rev.compl      ------------------------------------------------------------   61762
FAM230B-LOC105372935-GGT2.NCBI.GGT2.rev.compl                      caggtatcaatagttctggagttgatcatttcacagaaaggaaactgctgtcaacctgcc   88838

chimp.LOC112206744.LOC107973052.GGT2-.646279-708040.rev.compl      ------------------------------------------------------------   61762
FAM230B-LOC105372935-GGT2.NCBI.GGT2.rev.compl                      caaggccacacagctagtcctggataaattgacttaaagaacccaggtctctgattccca   88898

chimp.LOC112206744.LOC107973052.GGT2-.646279-708040.rev.compl      ------------------------------------------------------------   61762
FAM230B-LOC105372935-GGT2.NCBI.GGT2.rev.compl                      aatcagtgctcttacccctcccacgccacctctctcttctaaagatccacccaatggaac   88958

chimp.LOC112206744.LOC107973052.GGT2-.646279-708040.rev.compl      ------------------------------------------------------------   61762
FAM230B-LOC105372935-GGT2.NCBI.GGT2.rev.compl                      tgccccagggctggtatcccagctacttctccattcatgccgcaggtcaatgattcaggg   89018

chimp.LOC112206744.LOC107973052.GGT2-.646279-708040.rev.compl      ------------------------------------------------------------   61762
FAM230B-LOC105372935-GGT2.NCBI.GGT2.rev.compl                      cggacactcctcaagtgaccttcagaggccctgactcactcctccgacagtcctttgtgt   89078

chimp.LOC112206744.LOC107973052.GGT2-.646279-708040.rev.compl      ------------------------------------------------------------   61762
FAM230B-LOC105372935-GGT2.NCBI.GGT2.rev.compl                      ctggactcccagccgctcctcagcagctctgtcctcttttgtacatttattactgtcagc   89138

chimp.LOC112206744.LOC107973052.GGT2-.646279-708040.rev.compl      ------------------------------------------------------------   61762
FAM230B-LOC105372935-GGT2.NCBI.GGT2.rev.compl                      ctgaggctagcaggaggtctcaatgcctggtgatctgaggccagcctgtcccaggccctc   89198

chimp.LOC112206744.LOC107973052.GGT2-.646279-708040.rev.compl      ------------------------------------------------------------   61762
FAM230B-LOC105372935-GGT2.NCBI.GGT2.rev.compl                      ccagtcccgtgaggtcctcctggggatgcaggcacagcgactgacaaggataaggagcag   89258

chimp.LOC112206744.LOC107973052.GGT2-.646279-708040.rev.compl      ------------------------------------------------------------   61762
FAM230B-LOC105372935-GGT2.NCBI.GGT2.rev.compl                      aagggcaggtggctttctgggtgtgagcttggccttctgccacacccatgttcagaaagg   89318

chimp.LOC112206744.LOC107973052.GGT2-.646279-708040.rev.compl      ------------------------------------------------------------   61762
FAM230B-LOC105372935-GGT2.NCBI.GGT2.rev.compl                      tgtctacgggcacatcttgtgttgtcacaggtgtagaaagggctttgacacgcggcagat   89378

chimp.LOC112206744.LOC107973052.GGT2-.646279-708040.rev.compl      ------------------------------------------------------------   61762
FAM230B-LOC105372935-GGT2.NCBI.GGT2.rev.compl                      cctccaatccctgtgaaagagggatcattctttctattatacagtgaggacgctgaagtc   89438

chimp.LOC112206744.LOC107973052.GGT2-.646279-708040.rev.compl      ------------------------------------------------------------   61762
FAM230B-LOC105372935-GGT2.NCBI.GGT2.rev.compl                      cagaaaggtcaggtgactgatgaaaggtcataaaattagcaggaatcagggaaagggcag   89498

chimp.LOC112206744.LOC107973052.GGT2-.646279-708040.rev.compl      ------------------------------------------------------------   61762
FAM230B-LOC105372935-GGT2.NCBI.GGT2.rev.compl                      gaaagcagagcttccagttgcaaggtcaggactcttcagcccagcaggcgagtttgccag   89558

chimp.LOC112206744.LOC107973052.GGT2-.646279-708040.rev.compl      ------------------------------------------------------------   61762
FAM230B-LOC105372935-GGT2.NCBI.GGT2.rev.compl                      ccagtctgtgccctggtctctgccctccccaaacacgtgcaccaatttggaggggctgct   89618

chimp.LOC112206744.LOC107973052.GGT2-.646279-708040.rev.compl      ------------------------------------------------------------   61762
FAM230B-LOC105372935-GGT2.NCBI.GGT2.rev.compl                      cccagcacgcatgctgcaagataaacttcagtgctcagccctctcactgtgacaacaatc   89678

chimp.LOC112206744.LOC107973052.GGT2-.646279-708040.rev.compl      ------------------------------------------------------------   61762
FAM230B-LOC105372935-GGT2.NCBI.GGT2.rev.compl                      ccttagaaaggggtccacgctgtccatctgggcccctacaccaaagaacacctaattatc   89738

chimp.LOC112206744.LOC107973052.GGT2-.646279-708040.rev.compl      ------------------------------------------------------------   61762
FAM230B-LOC105372935-GGT2.NCBI.GGT2.rev.compl                      agtggtcctcactcaggaaacatgcccacaaataaagacaaaagtaaacccaactacaca   89798

chimp.LOC112206744.LOC107973052.GGT2-.646279-708040.rev.compl      ------------------------------------------------------------   61762
FAM230B-LOC105372935-GGT2.NCBI.GGT2.rev.compl                      ttcctcaaaatcctcctctccatttcacagaaacaccacaacagagcagaactgcaacct   89858

chimp.LOC112206744.LOC107973052.GGT2-.646279-708040.rev.compl      ------------------------------------------------------------   61762
FAM230B-LOC105372935-GGT2.NCBI.GGT2.rev.compl                      gatagcatccctgtctggttttggctctttcaaaacatcagggacctcacaggaaagcaa   89918

chimp.LOC112206744.LOC107973052.GGT2-.646279-708040.rev.compl      ------------------------------------------------------------   61762
FAM230B-LOC105372935-GGT2.NCBI.GGT2.rev.compl                      tgcagaaaactgcagtcatttggccgcttgttaaggagagccctgagcagcaagctccag   89978

chimp.LOC112206744.LOC107973052.GGT2-.646279-708040.rev.compl      ------------------------------------------------------------   61762
FAM230B-LOC105372935-GGT2.NCBI.GGT2.rev.compl                      accggagatgcgcatataacagcagcagttccacacagccctgcacagcagcaattcagc   90038

chimp.LOC112206744.LOC107973052.GGT2-.646279-708040.rev.compl      ------------------------------------------------------------   61762
FAM230B-LOC105372935-GGT2.NCBI.GGT2.rev.compl                      ctctggcagctgtgatgatgctgaggcagtgtccccactgccccagtggcttctgcatgc   90098

chimp.LOC112206744.LOC107973052.GGT2-.646279-708040.rev.compl      ------------------------------------------------------------   61762
FAM230B-LOC105372935-GGT2.NCBI.GGT2.rev.compl                      aggttcatttgggacaggacacctctaactccatctctggccaggagccagcatagtagg   90158

chimp.LOC112206744.LOC107973052.GGT2-.646279-708040.rev.compl      ------------------------------------------------------------   61762
FAM230B-LOC105372935-GGT2.NCBI.GGT2.rev.compl                      tgccgtgaaagccaggaagtgaactgcacagaatggactgttccagtgtccatgggctgc   90218

chimp.LOC112206744.LOC107973052.GGT2-.646279-708040.rev.compl      ------------------------------------------------------------   61762
FAM230B-LOC105372935-GGT2.NCBI.GGT2.rev.compl                      aagtgggtcccacgtcacccacaggtgaatcttaattatgaaccaaggtgacggcagaaa   90278

chimp.LOC112206744.LOC107973052.GGT2-.646279-708040.rev.compl      ------------------------------------------------------------   61762
FAM230B-LOC105372935-GGT2.NCBI.GGT2.rev.compl                      ggatgggcagcaaaggagggtgtgtgtggtgatcagccatcagaggagacggccctgtgt   90338

chimp.LOC112206744.LOC107973052.GGT2-.646279-708040.rev.compl      ------------------------------------------------------------   61762
FAM230B-LOC105372935-GGT2.NCBI.GGT2.rev.compl                      gatgaaaggacctcctcaggaaacctccccaccagcaactggctccaaatggtcagactt   90398

chimp.LOC112206744.LOC107973052.GGT2-.646279-708040.rev.compl      ------------------------------------------------------------   61762
FAM230B-LOC105372935-GGT2.NCBI.GGT2.rev.compl                      tccaagaaattcctgtgaaaaggacaggctcagggcatgtgaatgtgatagtcagcaatt   90458

chimp.LOC112206744.LOC107973052.GGT2-.646279-708040.rev.compl      ------------------------------------------------------------   61762
FAM230B-LOC105372935-GGT2.NCBI.GGT2.rev.compl                      cgcacctcagcagccagtgacactatggtatagggaaaagaacttagattaggcatcaaa   90518

chimp.LOC112206744.LOC107973052.GGT2-.646279-708040.rev.compl      ------------------------------------------------------------   61762
FAM230B-LOC105372935-GGT2.NCBI.GGT2.rev.compl                      agtacccggtccccttcatacatggctggtggggaggtaggatggtgcagccactttgga   90578
```

```
chimp.LOC112206744.LOC107973052.GGT2-.646279-708040.rev.compl      ------------------------------------------------------------   61762
FAM230B-LOC105372935-GGT2.NCBI.GGT2.rev.compl                      aaacagtctggtagtttctttcaaagttaccaaatgacccagcaatcccactcctaagta   90638

chimp.LOC112206744.LOC107973052.GGT2-.646279-708040.rev.compl      ------------------------------------------------------------   61762
FAM230B-LOC105372935-GGT2.NCBI.GGT2.rev.compl                      tttacccaagagaaataaaacctgtgtccacacaaagacttgtacacatcctaatagctg   90698

chimp.LOC112206744.LOC107973052.GGT2-.646279-708040.rev.compl      ------------------------------------------------------------   61762
FAM230B-LOC105372935-GGT2.NCBI.GGT2.rev.compl                      tattcatagtagccccaaactggaaacaacccaaatgcccatcagtttgtgaagagatgg   90758

chimp.LOC112206744.LOC107973052.GGT2-.646279-708040.rev.compl      ------------------------------------------------------------   61762
FAM230B-LOC105372935-GGT2.NCBI.GGT2.rev.compl                      acaaaatgtgctatatccataaaacggaaccctactcagcaataaaaaggtgtgaactag   90818

chimp.LOC112206744.LOC107973052.GGT2-.646279-708040.rev.compl      ------------------------------------------------------------   61762
FAM230B-LOC105372935-GGT2.NCBI.GGT2.rev.compl                      atacacacaagaacatggagggatctcaaaagcactgtgctgagcgagagaagccagatt   90878

chimp.LOC112206744.LOC107973052.GGT2-.646279-708040.rev.compl      ------------------------------------------------------------   61762
FAM230B-LOC105372935-GGT2.NCBI.GGT2.rev.compl                      caaagggctacacactgtaggatgccatgtgcatcacattctgtacagtgcaaaactgta   90938

chimp.LOC112206744.LOC107973052.GGT2-.646279-708040.rev.compl      ------------------------------------------------------------   61762
FAM230B-LOC105372935-GGT2.NCBI.GGT2.rev.compl                      gtggcaggattcaggtcccctgaggctggggtcctggaggcaggagaggggctggtagca   90998

chimp.LOC112206744.LOC107973052.GGT2-.646279-708040.rev.compl      ------------------------------------------------------------   61762
FAM230B-LOC105372935-GGT2.NCBI.GGT2.rev.compl                      aggaggcacaagggaacttttggggacagaggagtggtccatatttgataggtggtggt   91058

chimp.LOC112206744.LOC107973052.GGT2-.646279-708040.rev.compl      ------------------------------------------------------------   61762
FAM230B-LOC105372935-GGT2.NCBI.GGT2.rev.compl                      tgtgtggctgctcatgtttgtcaacaaagaattacatattttaaactggtgagttccgtt   91118

chimp.LOC112206744.LOC107973052.GGT2-.646279-708040.rev.compl      ------------------------------------------------------------   61762
FAM230B-LOC105372935-GGT2.NCBI.GGT2.rev.compl                      gtatgcaagcaaattataccacaatgaagcaaatctttaaaacactaggttccttttctt   91178

chimp.LOC112206744.LOC107973052.GGT2-.646279-708040.rev.compl      ------------------------------------------------------------   61762
FAM230B-LOC105372935-GGT2.NCBI.GGT2.rev.compl                      tatctagagctcaaagtagtggcccatgctggctatgacttggagaaatcacacaagctt   91238

chimp.LOC112206744.LOC107973052.GGT2-.646279-708040.rev.compl      ------------------------------------------------------------   61762
FAM230B-LOC105372935-GGT2.NCBI.GGT2.rev.compl                      actgggtctcaactgcttcatttgtacaggtggctgcataacactccccgggcaggggaa   91298

chimp.LOC112206744.LOC107973052.GGT2-.646279-708040.rev.compl      ------------------------------------------------------------   61762
FAM230B-LOC105372935-GGT2.NCBI.GGT2.rev.compl                      ctgtagggatgaaatcagcaaatgtgagtgaacacacagaggaagcaacacataggcctt   91358

chimp.LOC112206744.LOC107973052.GGT2-.646279-708040.rev.compl      ------------------------------------------------------------   61762
FAM230B-LOC105372935-GGT2.NCBI.GGT2.rev.compl                      ccccaaatacccacttctgttttctttcctcagtgtgagtaagtggctctaatgatccac   91418

chimp.LOC112206744.LOC107973052.GGT2-.646279-708040.rev.compl      ------------------------------------------------------------   61762
FAM230B-LOC105372935-GGT2.NCBI.GGT2.rev.compl                      aaaccttccaaccagccaagtcaaggaccagcaggaggactcaggaaagtactgaaatgc   91478

chimp.LOC112206744.LOC107973052.GGT2-.646279-708040.rev.compl      ------------------------------------------------------------   61762
FAM230B-LOC105372935-GGT2.NCBI.GGT2.rev.compl                      tggggagaaccaggacagcggctgagcacagccagacagggagggcagcctggtggggcc   91538

chimp.LOC112206744.LOC107973052.GGT2-.646279-708040.rev.compl      ------------------------------------------------------------   61762
FAM230B-LOC105372935-GGT2.NCBI.GGT2.rev.compl                      ctgggagccatggacgcgactgcgctgggagtggggatgggagatagcaggagttggtgg   91598

chimp.LOC112206744.LOC107973052.GGT2-.646279-708040.rev.compl      ------------------------------------------------------------   61762
FAM230B-LOC105372935-GGT2.NCBI.GGT2.rev.compl                      gtggggcacaaagtgggagccagggccaaattgctgctgatttgaccattttgcaaaagg   91658

chimp.LOC112206744.LOC107973052.GGT2-.646279-708040.rev.compl      ------------------------------------------------------------   61762
FAM230B-LOC105372935-GGT2.NCBI.GGT2.rev.compl                      ccagcctggctgtggcctcaggcctctcatttcctctaccataatgaggggtcacggaag   91718

chimp.LOC112206744.LOC107973052.GGT2-.646279-708040.rev.compl      ------------------------------------------------------------   61762
FAM230B-LOC105372935-GGT2.NCBI.GGT2.rev.compl                      tgccttcccacaactgcagcatctccctccacatttgggctcaaatacagagtggggccg   91778

chimp.LOC112206744.LOC107973052.GGT2-.646279-708040.rev.compl      ------------------------------------------------------------   61762
FAM230B-LOC105372935-GGT2.NCBI.GGT2.rev.compl                      gagacagaaaaagtctccgtgacacactgtgatttgtttctgcactgtgtgaacccttgg   91838

chimp.LOC112206744.LOC107973052.GGT2-.646279-708040.rev.compl      ------------------------------------------------------------   61762
FAM230B-LOC105372935-GGT2.NCBI.GGT2.rev.compl                      ctctgtgacacagaacacaggtttcagcagttcctgtcacttcccacaggcgaaacagac   91898

chimp.LOC112206744.LOC107973052.GGT2-.646279-708040.rev.compl      ------------------------------------------------------------   61762
FAM230B-LOC105372935-GGT2.NCBI.GGT2.rev.compl                      agagcattcttttgtccggcttctggtgaaacgccattatcgttattttaggaggaaga   91958

chimp.LOC112206744.LOC107973052.GGT2-.646279-708040.rev.compl      ------------------------------------------------------------   61762
FAM230B-LOC105372935-GGT2.NCBI.GGT2.rev.compl                      tggagaattggggtgggtccccacaaagaggatggattcagaagacttccaggaacctta   92018

chimp.LOC112206744.LOC107973052.GGT2-.646279-708040.rev.compl      ------------------------------------------------------------   61762
FAM230B-LOC105372935-GGT2.NCBI.GGT2.rev.compl                      aacggcttggaaactaattgtgccagttctttttgatgtctaattttattctaaaatttc   92078

chimp.LOC112206744.LOC107973052.GGT2-.646279-708040.rev.compl      ------------------------------------------------------------   61762
FAM230B-LOC105372935-GGT2.NCBI.GGT2.rev.compl                      ctctcagcacaatatattaatactgatcagccaagagaaggttaggttttggctcacaaa   92138

chimp.LOC112206744.LOC107973052.GGT2-.646279-708040.rev.compl      ------------------------------------------------------------   61762
FAM230B-LOC105372935-GGT2.NCBI.GGT2.rev.compl                      tagtcaaggagagaatctccttcattcattgagaaggcagaagatggcgttaaagagtat   92198

chimp.LOC112206744.LOC107973052.GGT2-.646279-708040.rev.compl      ------------------------------------------------------------   61762
FAM230B-LOC105372935-GGT2.NCBI.GGT2.rev.compl                      tcacaggtaactaagattgccagagggagaatcgagtgagttgtcagtggtgaaatacag   92258

chimp.LOC112206744.LOC107973052.GGT2-.646279-708040.rev.compl      ------------------------------------------------------------   61762
FAM230B-LOC105372935-GGT2.NCBI.GGT2.rev.compl                      aggaggtagagagagctacactgagacaacaaaggcagcttaagggaacaaagccagcta   92318

chimp.LOC112206744.LOC107973052.GGT2-.646279-708040.rev.compl      ------------------------------------------------------------   61762
FAM230B-LOC105372935-GGT2.NCBI.GGT2.rev.compl                      agggaacaaagccggggtccctcaggtctgtggccaccggctccagccaggacagcctgg   92378

chimp.LOC112206744.LOC107973052.GGT2-.646279-708040.rev.compl      ------------------------------------------------------------   61762
FAM230B-LOC105372935-GGT2.NCBI.GGT2.rev.compl                      tagctgaagagcacatccttgcccgttacatgcctgtgtttcaagtgatgctaacttgct   92438

chimp.LOC112206744.LOC107973052.GGT2-.646279-708040.rev.compl      ------------------------------------------------------------   61762
FAM230B-LOC105372935-GGT2.NCBI.GGT2.rev.compl                      aatgcagatatctccccaaaccctgcccctggctcaccacccagtgagcaatcactcact   92498

chimp.LOC112206744.LOC107973052.GGT2-.646279-708040.rev.compl      ------------------------------------------------------------   61762
FAM230B-LOC105372935-GGT2.NCBI.GGT2.rev.compl                      ttccccaggacaccttcagggaggtactctgccagcaccgcacacttaaacagctactgc   92558

chimp.LOC112206744.LOC107973052.GGT2-.646279-708040.rev.compl      ------------------------------------------------------------   61762
FAM230B-LOC105372935-GGT2.NCBI.GGT2.rev.compl                      aggttcttctctcagacgcagcacggttcctgagatgacgtcagcgctgctcgctgaaag   92618

chimp.LOC112206744.LOC107973052.GGT2-.646279-708040.rev.compl      ------------------------------------------------------------   61762
FAM230B-LOC105372935-GGT2.NCBI.GGT2.rev.compl                      caccaggcaaggaggcaaaacaaaggaaatcctaaagagcaggcgagagaacagtggaac   92678

chimp.LOC112206744.LOC107973052.GGT2-.646279-708040.rev.compl      ------------------------------------------------------------   61762
FAM230B-LOC105372935-GGT2.NCBI.GGT2.rev.compl                      tggcagagagaggcactggtgcggatggagagggcagcaataaaaactggggacactta   92738
```

```
chimp.LOC112206744.LOC107973052.GGT2-.646279-708040.rev.compl      ------------------------------------------------------------   61762
FAM230B-LOC105372935-GGT2.NCBI.GGT2.rev.compl                      gaggcaaaggtatggcaggtgacaaaaaagatgagctgaagctcaggagagacccaggca   92798

chimp.LOC112206744.LOC107973052.GGT2-.646279-708040.rev.compl      ------------------------------------------------------------   61762
FAM230B-LOC105372935-GGT2.NCBI.GGT2.rev.compl                      gcgatgcccaaatgaccggctgaggaatcacctcactgctgctgatgcccttgacagcga   92858

chimp.LOC112206744.LOC107973052.GGT2-.646279-708040.rev.compl      ------------------------------------------------------------   61762
FAM230B-LOC105372935-GGT2.NCBI.GGT2.rev.compl                      tgtctgaggctggcgtgagtgctccatgctcagagatacctagaacccagctacaatggc   92918

chimp.LOC112206744.LOC107973052.GGT2-.646279-708040.rev.compl      ------------------------------------------------------------   61762
FAM230B-LOC105372935-GGT2.NCBI.GGT2.rev.compl                      cttatcagactgaaacaagaacaccaagatattccaagggaggtgacaatggggtccaaa   92978

chimp.LOC112206744.LOC107973052.GGT2-.646279-708040.rev.compl      ------------------------------------------------------------   61762
FAM230B-LOC105372935-GGT2.NCBI.GGT2.rev.compl                      ggttgtgcatttatggatcacaactatgtgctgagtgtggcccaaatcccagctccatat   93038

chimp.LOC112206744.LOC107973052.GGT2-.646279-708040.rev.compl      ------------------------------------------------------------   61762
FAM230B-LOC105372935-GGT2.NCBI.GGT2.rev.compl                      ggtcacaccatgtctacagccaatcagctgaccggggtggcaacccctggctctgctcaa   93098

chimp.LOC112206744.LOC107973052.GGT2-.646279-708040.rev.compl      ------------------------------------------------------------   61762
FAM230B-LOC105372935-GGT2.NCBI.GGT2.rev.compl                      caataatgacaaccatcatttactgagcacttactatgtgccagaccgatgctaagtgct   93158

chimp.LOC112206744.LOC107973052.GGT2-.646279-708040.rev.compl      ------------------------------------------------------------   61762
FAM230B-LOC105372935-GGT2.NCBI.GGT2.rev.compl                      ttcctcagatgtttccatttaactcttagaatcaccctaatgagtatctcctattatttt   93218

chimp.LOC112206744.LOC107973052.GGT2-.646279-708040.rev.compl      ------------------------------------------------------------   61762
FAM230B-LOC105372935-GGT2.NCBI.GGT2.rev.compl                      acagatgaggaaatgagagtttagaaaagtaacttgcaaaatcatactgctggctccata   93278

chimp.LOC112206744.LOC107973052.GGT2-.646279-708040.rev.compl      ------------------------------------------------------------   61762
FAM230B-LOC105372935-GGT2.NCBI.GGT2.rev.compl                      acaaatgttgattgaatgttgtctgggtggacagaggcaaggatgaccggaacgagacaa   93338

chimp.LOC112206744.LOC107973052.GGT2-.646279-708040.rev.compl      ------------------------------------------------------------   61762
FAM230B-LOC105372935-GGT2.NCBI.GGT2.rev.compl                      acaaacaaaaggctgactgacagaccattcgggtagcacttttcctgggattaagcttag   93398

chimp.LOC112206744.LOC107973052.GGT2-.646279-708040.rev.compl      ------------------------------------------------------------   61762
FAM230B-LOC105372935-GGT2.NCBI.GGT2.rev.compl                      accttgtaggcctatcagccagaagcagtgtgtaaaataagagtggctgactgggcacgg   93458

chimp.LOC112206744.LOC107973052.GGT2-.646279-708040.rev.compl      ------------------------------------------------------------   61762
FAM230B-LOC105372935-GGT2.NCBI.GGT2.rev.compl                      tggctcaagcctgtaatccaagcactttgggaggccaaggcgggcggatcacctgaggtc   93518

chimp.LOC112206744.LOC107973052.GGT2-.646279-708040.rev.compl      ------------------------------------------------------------   61762
FAM230B-LOC105372935-GGT2.NCBI.GGT2.rev.compl                      aggagttcaagaccagcctggccaacatgatgaaaccccgtctctactacaaatacaaaa   93578

chimp.LOC112206744.LOC107973052.GGT2-.646279-708040.rev.compl      ------------------------------------------------------------   61762
FAM230B-LOC105372935-GGT2.NCBI.GGT2.rev.compl                      attagccgggtgtgatggcacacgcctgtaatcctagttactcgggaggctgaggcacga   93638

chimp.LOC112206744.LOC107973052.GGT2-.646279-708040.rev.compl      ------------------------------------------------------------   61762
FAM230B-LOC105372935-GGT2.NCBI.GGT2.rev.compl                      gaatcgcttgaacctgggaggcagaggttgcgataagctgagattgggccactgcactcc   93698

chimp.LOC112206744.LOC107973052.GGT2-.646279-708040.rev.compl      ------------------------------------------------------------   61762
FAM230B-LOC105372935-GGT2.NCBI.GGT2.rev.compl                      agcctgggcgacagagcaagactccctctgaaaaaaaaaaaaaaaaagagtggctaccca   93758

chimp.LOC112206744.LOC107973052.GGT2-.646279-708040.rev.compl      ------------------------------------------------------------   61762
FAM230B-LOC105372935-GGT2.NCBI.GGT2.rev.compl                      agaacgcaacagcagcagcaaacactgatctgatgacaccccagttttgcagaagaggaa   93818

chimp.LOC112206744.LOC107973052.GGT2-.646279-708040.rev.compl      ------------------------------------------------------------   61762
FAM230B-LOC105372935-GGT2.NCBI.GGT2.rev.compl                      ggaaaaataagaatgtcaactctccattcaacagagaaacagtgccaccactgtctttaa   93878

chimp.LOC112206744.LOC107973052.GGT2-.646279-708040.rev.compl      ------------------------------------------------------------   61762
FAM230B-LOC105372935-GGT2.NCBI.GGT2.rev.compl                      attttcaaacttatcgagaaaagctccagagaggccagtgccagcaatgccggctgtctc   93938

chimp.LOC112206744.LOC107973052.GGT2-.646279-708040.rev.compl      ------------------------------------------------------------   61762
FAM230B-LOC105372935-GGT2.NCBI.GGT2.rev.compl                      ccaagtctgaagggttttcctctgggtattaggagttgaggtgtttggttacagaaggaa   93998

chimp.LOC112206744.LOC107973052.GGT2-.646279-708040.rev.compl      ------------------------------------------------------------   61762
FAM230B-LOC105372935-GGT2.NCBI.GGT2.rev.compl                      acaggatgcacgggagatcacacacagagtaagaggtgcctctcaaatgcctttcccacg   94058

chimp.LOC112206744.LOC107973052.GGT2-.646279-708040.rev.compl      ------------------------------------------------------------   61762
FAM230B-LOC105372935-GGT2.NCBI.GGT2.rev.compl                      tttctaactctgctttttttaaagcacgatcagacccgccatctgcacttttacctgctg   94118

chimp.LOC112206744.LOC107973052.GGT2-.646279-708040.rev.compl      ------------------------------------------------------------   61762
FAM230B-LOC105372935-GGT2.NCBI.GGT2.rev.compl                      cctgacgaaaaccagatgccaaaggcctcccagcactcacggggccctttacccttagga   94178

chimp.LOC112206744.LOC107973052.GGT2-.646279-708040.rev.compl      ------------------------------------------------------------   61762
FAM230B-LOC105372935-GGT2.NCBI.GGT2.rev.compl                      aaacaccgtgattcagccaggcgagattgatgcaacattgtaacaaagcgtatgcaaagt   94238

chimp.LOC112206744.LOC107973052.GGT2-.646279-708040.rev.compl      ------------------------------------------------------------   61762
FAM230B-LOC105372935-GGT2.NCBI.GGT2.rev.compl                      gcctagctccggactgcttgcagaaaaggcacattttgactcttctgagagtgggaaaga   94298

chimp.LOC112206744.LOC107973052.GGT2-.646279-708040.rev.compl      ------------------------------------------------------------   61762
FAM230B-LOC105372935-GGT2.NCBI.GGT2.rev.compl                      tttgaaagatgggatttgcaccttattcttattcaagtctggagatctggaacttcttcc   94358

chimp.LOC112206744.LOC107973052.GGT2-.646279-708040.rev.compl      ------------------------------------------------------------   61762
FAM230B-LOC105372935-GGT2.NCBI.GGT2.rev.compl                      tgaagctccatgtttagccagataattgagacgctccacctggtcaccactggacaccag   94418

chimp.LOC112206744.LOC107973052.GGT2-.646279-708040.rev.compl      ------------------------------------------------------------   61762
FAM230B-LOC105372935-GGT2.NCBI.GGT2.rev.compl                      ccacccaaacattctcctctgcaaggcacttggcattctggtgcctgtggagagagcagc   94478

chimp.LOC112206744.LOC107973052.GGT2-.646279-708040.rev.compl      ------------------------------------------------------------   61762
FAM230B-LOC105372935-GGT2.NCBI.GGT2.rev.compl                      cttctcactagcacagctcatgaacaaggatccccttccctggacagccaagatgtaaac   94538

chimp.LOC112206744.LOC107973052.GGT2-.646279-708040.rev.compl      ------------------------------------------------------------   61762
FAM230B-LOC105372935-GGT2.NCBI.GGT2.rev.compl                      ctgcccccagaaacactgagagtcctagaggtagaatgagtcggccagcaagctccagcc   94598

chimp.LOC112206744.LOC107973052.GGT2-.646279-708040.rev.compl      ------------------------------------------------------------   61762
FAM230B-LOC105372935-GGT2.NCBI.GGT2.rev.compl                      cattgaatgcagacccctgagcaaatcaagtcaggagaaggtggacaactgtaccctgaa   94658

chimp.LOC112206744.LOC107973052.GGT2-.646279-708040.rev.compl      ------------------------------------------------------------   61762
FAM230B-LOC105372935-GGT2.NCBI.GGT2.rev.compl                      gaccacataccacccagacccaacacaaatagctgtcagcactgccacccattacagctt   94718

chimp.LOC112206744.LOC107973052.GGT2-.646279-708040.rev.compl      ------------------------------------------------------------   61762
FAM230B-LOC105372935-GGT2.NCBI.GGT2.rev.compl                      cagatctcgcttctaccagcatttcccagctgagcaacaaccgtcaaagtctggtcctgg   94778

chimp.LOC112206744.LOC107973052.GGT2-.646279-708040.rev.compl      ------------------------------------------------------------   61762
FAM230B-LOC105372935-GGT2.NCBI.GGT2.rev.compl                      aaggacagccctccccgtcttggcagccacactttctggccctagcccaaggctgctgtg   94838

chimp.LOC112206744.LOC107973052.GGT2-.646279-708040.rev.compl      ------------------------------------------------------------   61762
FAM230B-LOC105372935-GGT2.NCBI.GGT2.rev.compl                      gcatgggaggaatcaatggaggaatacatggctcttccctggttccttcctctcccactt   94898

chimp.LOC112206744.LOC107973052.GGT2-.646279-708040.rev.compl      ------------------------------------------------------------   61762
```

```
FAM230B-LOC105372935-GGT2.NCBI.GGT2.rev.compl                    cattgactaacactagtcacatggccccaactgaactgtgagggaggctggaaaatctgg   94958

chimp.LOC112206744.LOC107973052.GGT2-.646279-708040.rev.compl    ------------------------------------------------------------   61762
FAM230B-LOC105372935-GGT2.NCBI.GGT2.rev.compl                    agtatgtgcatgcctagtgagttgcctctaccacatcctcttaacccttggcctctcatt   95018

chimp.LOC112206744.LOC107973052.GGT2-.646279-708040.rev.compl    ------------------------------------------------------------   61762
FAM230B-LOC105372935-GGT2.NCBI.GGT2.rev.compl                    cccaaagtctagccctattttcaaatgaattcaacgcctcattacttagatctttcctca   95078

chimp.LOC112206744.LOC107973052.GGT2-.646279-708040.rev.compl    ------------------------------------------------------------   61762
FAM230B-LOC105372935-GGT2.NCBI.GGT2.rev.compl                    ttacctcaaagccatagtggcaaaaagcaaacagcatcttcctaaaatcttcctcttcca   95138

chimp.LOC112206744.LOC107973052.GGT2-.646279-708040.rev.compl    ------------------------------------------------------------   61762
FAM230B-LOC105372935-GGT2.NCBI.GGT2.rev.compl                    tcttcctctcctagatatccatcttacccttctttgccagttaatttattacaatcttcc   95198

chimp.LOC112206744.LOC107973052.GGT2-.646279-708040.rev.compl    ------------------------------------------------------------   61762
FAM230B-LOC105372935-GGT2.NCBI.GGT2.rev.compl                    tttgacagcttcactccctccccgcactgctatcaactcaaatgtgagttatttcagtag   95258

chimp.LOC112206744.LOC107973052.GGT2-.646279-708040.rev.compl    ------------------------------------------------------------   61762
FAM230B-LOC105372935-GGT2.NCBI.GGT2.rev.compl                    cttgagcttgtcactggccttttactttttccccttccaaagaacctggtacccagcta   95318

chimp.LOC112206744.LOC107973052.GGT2-.646279-708040.rev.compl    ------------------------------------------------------------   61762
FAM230B-LOC105372935-GGT2.NCBI.GGT2.rev.compl                    cttaatccttatccttcttcccaaggcccactttcattcctttctcccccattcatgagc   95378

chimp.LOC112206744.LOC107973052.GGT2-.646279-708040.rev.compl    ------------------------------------------------------------   61762
FAM230B-LOC105372935-GGT2.NCBI.GGT2.rev.compl                    ctgcaatacttttcccgctgcataagtcacgtgtcaccatataaactcacaatttgaggt   95438

chimp.LOC112206744.LOC107973052.GGT2-.646279-708040.rev.compl    ------------------------------------------------------------   61762
FAM230B-LOC105372935-GGT2.NCBI.GGT2.rev.compl                    gctctagaatctgtccacggaattagtcaggacataatctccctgtttttcggcagtgcc   95498

chimp.LOC112206744.LOC107973052.GGT2-.646279-708040.rev.compl    ------------------------------------------------------------   61762
FAM230B-LOC105372935-GGT2.NCBI.GGT2.rev.compl                    acatccatacctgtcttcacaggtttgctcatgccaccattcctttccccagaacgccct   95558

chimp.LOC112206744.LOC107973052.GGT2-.646279-708040.rev.compl    ------------------------------------------------------------   61762
FAM230B-LOC105372935-GGT2.NCBI.GGT2.rev.compl                    tctgctcctcccacacaaatcctaatcctccctcaggacccatttaaaatacaatctcct   95618

chimp.LOC112206744.LOC107973052.GGT2-.646279-708040.rev.compl    ------------------------------------------------------------   61762
FAM230B-LOC105372935-GGT2.NCBI.GGT2.rev.compl                    cactcaactctttcctaccaacttgctttcccctctcctttctgccttaccacaaagcta   95678

chimp.LOC112206744.LOC107973052.GGT2-.646279-708040.rev.compl    ------------------------------------------------------------   61762
FAM230B-LOC105372935-GGT2.NCBI.GGT2.rev.compl                    agctctgtctctgaataatgtgctttttgatgataatctgaatggtacaagaagaaatcc   95738

chimp.LOC112206744.LOC107973052.GGT2-.646279-708040.rev.compl    ------------------------------------------------------------   61762
FAM230B-LOC105372935-GGT2.NCBI.GGT2.rev.compl                    tatttcatgctcctttgagggagtgtctaatataggttgccttcgactataagtagtaga   95798

chimp.LOC112206744.LOC107973052.GGT2-.646279-708040.rev.compl    ------------------------------------------------------------   61762
FAM230B-LOC105372935-GGT2.NCBI.GGT2.rev.compl                    aaattcccactcaactggcttcaacagtgaggacaaaatgtatggagattggacgcttca   95858

chimp.LOC112206744.LOC107973052.GGT2-.646279-708040.rev.compl    ------------------------------------------------------------   61762
FAM230B-LOC105372935-GGT2.NCBI.GGT2.rev.compl                    gtcttcaggggcttggccctcagggtttttcccatctcttcccgtctgctggcctcaaca   95918

chimp.LOC112206744.LOC107973052.GGT2-.646279-708040.rev.compl    ------------------------------------------------------------   61762
FAM230B-LOC105372935-GGT2.NCBI.GGT2.rev.compl                    aacctgctttcctcattgcaacaagatggctgcttcacgtccgagaatctagtgaaagaa   95978

chimp.LOC112206744.LOC107973052.GGT2-.646279-708040.rev.compl    ------------------------------------------------------------   61762
FAM230B-LOC105372935-GGT2.NCBI.GGT2.rev.compl                    gctgtcccttctgcatttccttttacaaggaaaaacatctttcccagcatctcccagcag   96038

chimp.LOC112206744.LOC107973052.GGT2-.646279-708040.rev.compl    ------------------------------------------------------------   61762
FAM230B-LOC105372935-GGT2.NCBI.GGT2.rev.compl                    acttctcatacctcagtggccaaaaccatgtctaaactacgggcctccaggatgggctca   96098

chimp.LOC112206744.LOC107973052.GGT2-.646279-708040.rev.compl    ------------------------------------------------------------   61762
FAM230B-LOC105372935-GGT2.NCBI.GGT2.rev.compl                    gctcaatcagcacataccctagagctggaggcagggtcacctttccttgagtggcgtgtg   96158

chimp.LOC112206744.LOC107973052.GGT2-.646279-708040.rev.compl    ------------------------------------------------------------   61762
FAM230B-LOC105372935-GGT2.NCBI.GGT2.rev.compl                    ggagaggtgtggcatccaacccaactgggtcctgccagcaggaagagaaaagaagagcca   96218

chimp.LOC112206744.LOC107973052.GGT2-.646279-708040.rev.compl    ------------------------------------------------------------   61762
FAM230B-LOC105372935-GGT2.NCBI.GGT2.rev.compl                    ctgtaggtaaccaatcacgtctaccacaggcagggaccaataccacatatctgtgtttcc   96278

chimp.LOC112206744.LOC107973052.GGT2-.646279-708040.rev.compl    ------------------------------------------------------------   61762
FAM230B-LOC105372935-GGT2.NCBI.GGT2.rev.compl                    cctcacttgccaccccatcctgtgctggcagatgaggatgctcaacaaatagctgtgga   96338

chimp.LOC112206744.LOC107973052.GGT2-.646279-708040.rev.compl    ------------------------------------------------------------   61762
FAM230B-LOC105372935-GGT2.NCBI.GGT2.rev.compl                    actacaaattatcctgtgggtgtgagaaaagagtgtgcattataatagggtggtacaatt   96398

chimp.LOC112206744.LOC107973052.GGT2-.646279-708040.rev.compl    ------------------------------------------------------------   61762
FAM230B-LOC105372935-GGT2.NCBI.GGT2.rev.compl                    ggacatgtgtgcacggaaaatggaaaaacagccatgaaatccaaacagatgcaaaaacgt   96458

chimp.LOC112206744.LOC107973052.GGT2-.646279-708040.rev.compl    ------------------------------------------------------------   61762
FAM230B-LOC105372935-GGT2.NCBI.GGT2.rev.compl                    tcttaacacagaaggtttgggaagctctgatctaagccctagcctgggccctactcccag   96518

chimp.LOC112206744.LOC107973052.GGT2-.646279-708040.rev.compl    ------------------------------------------------------------   61762
FAM230B-LOC105372935-GGT2.NCBI.GGT2.rev.compl                    ttccacagatagggagcaggccccaggccagagtgctgggcttctcccagcaggatccag   96578

chimp.LOC112206744.LOC107973052.GGT2-.646279-708040.rev.compl    ------------------------------------------------------------   61762
FAM230B-LOC105372935-GGT2.NCBI.GGT2.rev.compl                    ctggggctttctcctcagttcaacccttgcctctgaggaccagtggtctacagtccagcc   96638

chimp.LOC112206744.LOC107973052.GGT2-.646279-708040.rev.compl    ------------------------------------------------------------   61762
FAM230B-LOC105372935-GGT2.NCBI.GGT2.rev.compl                    caccacccttcagacagaagttccaggcacaaaaaggaagctcacaataaggaacgccca   96698

chimp.LOC112206744.LOC107973052.GGT2-.646279-708040.rev.compl    ------------------------------------------------------------   61762
FAM230B-LOC105372935-GGT2.NCBI.GGT2.rev.compl                    tgcctagccataactgagcgcatcacagactcacacaaatctgtggaatgagtgcatgat   96758

chimp.LOC112206744.LOC107973052.GGT2-.646279-708040.rev.compl    ------------------------------------------------------------   61762
FAM230B-LOC105372935-GGT2.NCBI.GGT2.rev.compl                    ctcatgagctctggtctggaaaagtaagtctccctacactaggacttaaagggaaggctt   96818

chimp.LOC112206744.LOC107973052.GGT2-.646279-708040.rev.compl    ------------------------------------------------------------   61762
FAM230B-LOC105372935-GGT2.NCBI.GGT2.rev.compl                    ctttggctatgccaatatctttcctgccactttcttcttcaggccaaacagctcctatcc   96878

chimp.LOC112206744.LOC107973052.GGT2-.646279-708040.rev.compl    ------------------------------------------------------------   61762
FAM230B-LOC105372935-GGT2.NCBI.GGT2.rev.compl                    ctgccttccatcttagctttccagcatactcttcccaggttctttgggctactgacaagc   96938

chimp.LOC112206744.LOC107973052.GGT2-.646279-708040.rev.compl    ------------------------------------------------------------   61762
FAM230B-LOC105372935-GGT2.NCBI.GGT2.rev.compl                    actccagacccaaatacacatgctcccacctgaggctgggaaatgtaccaagaactgcat   96998

chimp.LOC112206744.LOC107973052.GGT2-.646279-708040.rev.compl    ------------------------------------------------------------   61762
FAM230B-LOC105372935-GGT2.NCBI.GGT2.rev.compl                    agaaaatatactggacaaaaggcactccagtctcacagatctccagtcccacagtctata   97058

chimp.LOC112206744.LOC107973052.GGT2-.646279-708040.rev.compl    ------------------------------------------------------------   61762
FAM230B-LOC105372935-GGT2.NCBI.GGT2.rev.compl                    tcaccagttttagctgtttgaagttttagaattccttttcaaaaccatagctagcatttc   97118
```

```
chimp.LOC112206744.LOC107973052.GGT2-.646279-708040.rev.compl      ------------------------------------------------------------   61762
FAM230B-LOC105372935-GGT2.NCBI.GGT2.rev.compl                      tttggctagagaaaactgtgtgaaatgcagcctaccataggatcactgattacaacatgc   97178

chimp.LOC112206744.LOC107973052.GGT2-.646279-708040.rev.compl      ------------------------------------------------------------   61762
FAM230B-LOC105372935-GGT2.NCBI.GGT2.rev.compl                      attcaacattcaatacatattgactgagcaactattaggttgaactatgtgacattattg   97238

chimp.LOC112206744.LOC107973052.GGT2-.646279-708040.rev.compl      ------------------------------------------------------------   61762
FAM230B-LOC105372935-GGT2.NCBI.GGT2.rev.compl                      acattcgactgcttctgacctatcaaaatggcaataaagctcaacttaatatttactcta   97298

chimp.LOC112206744.LOC107973052.GGT2-.646279-708040.rev.compl      ------------------------------------------------------------   61762
FAM230B-LOC105372935-GGT2.NCBI.GGT2.rev.compl                      tgccaggcagtaggctaaggaaggatttgggacaccatgacaaacaacacagaagtaacc   97358

chimp.LOC112206744.LOC107973052.GGT2-.646279-708040.rev.compl      ------------------------------------------------------------   61762
FAM230B-LOC105372935-GGT2.NCBI.GGT2.rev.compl                      cctatcttaatggaacttaaaatctagttgattaaaccattacttacaaaagtacaatct   97418

chimp.LOC112206744.LOC107973052.GGT2-.646279-708040.rev.compl      ------------------------------------------------------------   61762
FAM230B-LOC105372935-GGT2.NCBI.GGT2.rev.compl                      aacttcaggattgtaattaattataaatgcatttatttgtttactgcctgacttctttgc   97478

chimp.LOC112206744.LOC107973052.GGT2-.646279-708040.rev.compl      ------------------------------------------------------------   61762
FAM230B-LOC105372935-GGT2.NCBI.GGT2.rev.compl                      tagcctataaactacacaaagatgagggttatgtctcctccatttcacacactggcaaag   97538

chimp.LOC112206744.LOC107973052.GGT2-.646279-708040.rev.compl      ------------------------------------------------------------   61762
FAM230B-LOC105372935-GGT2.NCBI.GGT2.rev.compl                      catatagcagaatacatcatatatttgttaaatgtattgattagtgttacaacagacaat   97598

chimp.LOC112206744.LOC107973052.GGT2-.646279-708040.rev.compl      ------------------------------------------------------------   61762
FAM230B-LOC105372935-GGT2.NCBI.GGT2.rev.compl                      tgcatgtgctatggaagcgtttggcagagagatctaatctaatcttggaaggtgaggatc   97658

chimp.LOC112206744.LOC107973052.GGT2-.646279-708040.rev.compl      ------------------------------------------------------------   61762
FAM230B-LOC105372935-GGT2.NCBI.GGT2.rev.compl                      ctggccatcactaacttggaatttctgatactgatgatataaggttgagttatgtcttcg   97718

chimp.LOC112206744.LOC107973052.GGT2-.646279-708040.rev.compl      ------------------------------------------------------------   61762
FAM230B-LOC105372935-GGT2.NCBI.GGT2.rev.compl                      tgaaacaggtctctatacagcatataaacaactaccagcatgcacaggggttccgtaggg   97778

chimp.LOC112206744.LOC107973052.GGT2-.646279-708040.rev.compl      ------------------------------------------------------------   61762
FAM230B-LOC105372935-GGT2.NCBI.GGT2.rev.compl                      actatttaaaatacttaagggaacaaacggttttcaagctcagtctagtatttcagaaat   97838

chimp.LOC112206744.LOC107973052.GGT2-.646279-708040.rev.compl      ------------------------------------------------------------   61762
FAM230B-LOC105372935-GGT2.NCBI.GGT2.rev.compl                      tccatttacataaacaattggtgcactaattgcaaatcattcttatttctctcctaaagc   97898

chimp.LOC112206744.LOC107973052.GGT2-.646279-708040.rev.compl      ------------------------------------------------------------   61762
FAM230B-LOC105372935-GGT2.NCBI.GGT2.rev.compl                      aggatcccctaaggatctttattaggcattctattagcctagtttgagttacagtattta   97958

chimp.LOC112206744.LOC107973052.GGT2-.646279-708040.rev.compl      ------------------------------------------------------------   61762
FAM230B-LOC105372935-GGT2.NCBI.GGT2.rev.compl                      cttaaaagtaataacagtgaagttcttgaaaaatatttggtaattcagacattttactaa   98018

chimp.LOC112206744.LOC107973052.GGT2-.646279-708040.rev.compl      ------------------------------------------------------------   61762
FAM230B-LOC105372935-GGT2.NCBI.GGT2.rev.compl                      ttcaaactaatcttcctcattagtctacgttgttagaggtttaaccattgtattaagaga   98078

chimp.LOC112206744.LOC107973052.GGT2-.646279-708040.rev.compl      ------------------------------------------------------------   61762
FAM230B-LOC105372935-GGT2.NCBI.GGT2.rev.compl                      aagtatcaaaggcttcacttagagtaatagtcaccaaatttttttgcaagacaccttggt   98138

chimp.LOC112206744.LOC107973052.GGT2-.646279-708040.rev.compl      ------------------------------------------------------------   61762
FAM230B-LOC105372935-GGT2.NCBI.GGT2.rev.compl                      agcaggaacctgagctaacatcaggaggtaggtagaaatagaaaataaaggaatgacaat   98198

chimp.LOC112206744.LOC107973052.GGT2-.646279-708040.rev.compl      ------------------------------------------------------------   61762
FAM230B-LOC105372935-GGT2.NCBI.GGT2.rev.compl                      ttgaagagcaactgcaatccaatttagtccaacagatgacacacatatgaagtcacattc   98258

chimp.LOC112206744.LOC107973052.GGT2-.646279-708040.rev.compl      ------------------------------------------------------------   61762
FAM230B-LOC105372935-GGT2.NCBI.GGT2.rev.compl                      aaattataagcagagcacaaaaatatttagatccttatcttttgattaacctggtatca   98318

chimp.LOC112206744.LOC107973052.GGT2-.646279-708040.rev.compl      ------------------------------------------------------------   61762
FAM230B-LOC105372935-GGT2.NCBI.GGT2.rev.compl                      ttcagggaaaaatgtcagggaacaccactgaccagcagtcctctcttatctattttgcag   98378

chimp.LOC112206744.LOC107973052.GGT2-.646279-708040.rev.compl      ------------------------------------------------------------   61762
FAM230B-LOC105372935-GGT2.NCBI.GGT2.rev.compl                      atcttatctatttttctccctgagaaccagccctggacacaaactgcataaacatcaatg   98438

chimp.LOC112206744.LOC107973052.GGT2-.646279-708040.rev.compl      ------------------------------------------------------------   61762
FAM230B-LOC105372935-GGT2.NCBI.GGT2.rev.compl                      agcctagacagggtcttcatgtgtgtttggcctgccattgttctctgtaccagtacggag   98498

chimp.LOC112206744.LOC107973052.GGT2-.646279-708040.rev.compl      ------------------------------------------------------------   61762
FAM230B-LOC105372935-GGT2.NCBI.GGT2.rev.compl                      atttagtaacacactggtcacattcatggggcatcttgaactctcttgacatggtaaatg   98558

chimp.LOC112206744.LOC107973052.GGT2-.646279-708040.rev.compl      ------------------------------------------------------------   61762
FAM230B-LOC105372935-GGT2.NCBI.GGT2.rev.compl                      aacagcaggtcctgttcccatgccaaaagcaaacaaaaacctgtcaccagcaaggaagac   98618

chimp.LOC112206744.LOC107973052.GGT2-.646279-708040.rev.compl      ------------------------------------------------------------   61762
FAM230B-LOC105372935-GGT2.NCBI.GGT2.rev.compl                      acccagtctctccaaaggatgccaaagtccacatccagcctcctttgcagagcctcacaa   98678

chimp.LOC112206744.LOC107973052.GGT2-.646279-708040.rev.compl      ------------------------------------------------------------   61762
FAM230B-LOC105372935-GGT2.NCBI.GGT2.rev.compl                      atctctcggatgcaggcaggatgatcacgggctagggaagattatgctctgaatgtctta   98738

chimp.LOC112206744.LOC107973052.GGT2-.646279-708040.rev.compl      ------------------------------------------------------------   61762
FAM230B-LOC105372935-GGT2.NCBI.GGT2.rev.compl                      agagattagttacttaaaatcctccacatgacttcagtcatccccgttgcagatcacaga   98798

chimp.LOC112206744.LOC107973052.GGT2-.646279-708040.rev.compl      ------------------------------------------------------------   61762
FAM230B-LOC105372935-GGT2.NCBI.GGT2.rev.compl                      gtaaaaagcctacctgattaaagctgaatgcctatacatttgcaaaagggaaaatatgca   98858

chimp.LOC112206744.LOC107973052.GGT2-.646279-708040.rev.compl      ------------------------------------------------------------   61762
FAM230B-LOC105372935-GGT2.NCBI.GGT2.rev.compl                      gaagaaaacgagagtcagatactgatgggcatagcaccagctctaggacatcagaggtgt   98918

chimp.LOC112206744.LOC107973052.GGT2-.646279-708040.rev.compl      ------------------------------------------------------------   61762
FAM230B-LOC105372935-GGT2.NCBI.GGT2.rev.compl                      tgaataaatgatgacaaggaggcacacatagttctgatatattcaccataaataccacag   98978

chimp.LOC112206744.LOC107973052.GGT2-.646279-708040.rev.compl      ------------------------------------------------------------   61762
FAM230B-LOC105372935-GGT2.NCBI.GGT2.rev.compl                      tcctcaaggctctctcaattacttacttccccccaaatcagagactgaatctggccttat   99038

chimp.LOC112206744.LOC107973052.GGT2-.646279-708040.rev.compl      ------------------------------------------------------------   61762
FAM230B-LOC105372935-GGT2.NCBI.GGT2.rev.compl                      cttcaaatgaggcccagctctccccaaattccaaatatgggaaattacagctcctctgtg   99098

chimp.LOC112206744.LOC107973052.GGT2-.646279-708040.rev.compl      ------------------------------------------------------------   61762
FAM230B-LOC105372935-GGT2.NCBI.GGT2.rev.compl                      gtagggggaagagtgggaaggcgaatgtttgtattgcataacgttttaattaaaacaaag   99158

chimp.LOC112206744.LOC107973052.GGT2-.646279-708040.rev.compl      ------------------------------------------------------------   61762
FAM230B-LOC105372935-GGT2.NCBI.GGT2.rev.compl                      atgacgtgcaccagagtctctttctcatctcctgttaggggcgttttccagtactacttc   99218

chimp.LOC112206744.LOC107973052.GGT2-.646279-708040.rev.compl      ------------------------------------------------------------   61762
FAM230B-LOC105372935-GGT2.NCBI.GGT2.rev.compl                      caccctttacccacccaggtcaccatggtctttttcttccctctaggacaattcaccttt   99278
```

```
chimp.LOC112206744.LOC107973052.GGT2-.646279-708040.rev.compl      ------------------------------------------------------------   61762
FAM230B-LOC105372935-GGT2.NCBI.GGT2.rev.compl                      aaaataagaggattgggcaagaaatctacaaagatttttccaaaatgaactcaggcctag   99338

chimp.LOC112206744.LOC107973052.GGT2-.646279-708040.rev.compl      ------------------------------------------------------------   61762
FAM230B-LOC105372935-GGT2.NCBI.GGT2.rev.compl                      aggcatattagttctctatcttgttgctcccttctagaagtgcacagttcagagggatgc   99398

chimp.LOC112206744.LOC107973052.GGT2-.646279-708040.rev.compl      ------------------------------------------------------------   61762
FAM230B-LOC105372935-GGT2.NCBI.GGT2.rev.compl                      ccagccacagaggtgccaaaacgcagagctggaaccaatcacctgctccaacagagacaa   99458

chimp.LOC112206744.LOC107973052.GGT2-.646279-708040.rev.compl      ------------------------------------------------------------   61762
FAM230B-LOC105372935-GGT2.NCBI.GGT2.rev.compl                      ggtgaaagcactgggcagggtgctcagaatgtgctcttgcttctttgtaatgaaatatca   99518

chimp.LOC112206744.LOC107973052.GGT2-.646279-708040.rev.compl      ------------------------------------------------------------   61762
FAM230B-LOC105372935-GGT2.NCBI.GGT2.rev.compl                      gaagacagattgtaacattgttcagcaagctatttttttgtactgaaacatcagaaggcat   99578

chimp.LOC112206744.LOC107973052.GGT2-.646279-708040.rev.compl      ------------------------------------------------------------   61762
FAM230B-LOC105372935-GGT2.NCBI.GGT2.rev.compl                      tatgaaactgttcagcaaatcatctctaaacaatgtcataaagggttacctagtttgggg   99638

chimp.LOC112206744.LOC107973052.GGT2-.646279-708040.rev.compl      ------------------------------------------------------------   61762
FAM230B-LOC105372935-GGT2.NCBI.GGT2.rev.compl                      taagcccggaatgaatatctgcaccatcattcctcccgctaacaaacatgtccatctatg   99698

chimp.LOC112206744.LOC107973052.GGT2-.646279-708040.rev.compl      ------------------------------------------------------------   61762
FAM230B-LOC105372935-GGT2.NCBI.GGT2.rev.compl                      tgaagcataattcccaacacaccaccatctccaaaccaacaccaaaccactgccaaacaa   99758

chimp.LOC112206744.LOC107973052.GGT2-.646279-708040.rev.compl      ------------------------------------------------------------   61762
FAM230B-LOC105372935-GGT2.NCBI.GGT2.rev.compl                      agctgaagaggaaaaccagaaaggtctccgagaagtgctgcttgcagggagaagaatggc   99818

chimp.LOC112206744.LOC107973052.GGT2-.646279-708040.rev.compl      ------------------------------------------------------------   61762
FAM230B-LOC105372935-GGT2.NCBI.GGT2.rev.compl                      cctgcacagaggctcctgagggcagcaggtgcctgctttggagccttggcaatttccacc   99878

chimp.LOC112206744.LOC107973052.GGT2-.646279-708040.rev.compl      ------------------------------------------------------------   61762
FAM230B-LOC105372935-GGT2.NCBI.GGT2.rev.compl                      aggcagaatttacaaccgtctgggacctgcagtctcagagggctcacggagaagggggaa   99938

chimp.LOC112206744.LOC107973052.GGT2-.646279-708040.rev.compl      ------------------------------------------------------------   61762
FAM230B-LOC105372935-GGT2.NCBI.GGT2.rev.compl                      gctcttggctacccgggaggtcaccagtgactctgagcaaagtgcagatcaatgcctgct   99998

chimp.LOC112206744.LOC107973052.GGT2-.646279-708040.rev.compl      ------------------------------------------------------------   61762
FAM230B-LOC105372935-GGT2.NCBI.GGT2.rev.compl                      tagggttaccccccaggctgggagagggcatctggcaaggaacagacctggaagagcctgc   100058

chimp.LOC112206744.LOC107973052.GGT2-.646279-708040.rev.compl      ------------------------------------------------------------   61762
FAM230B-LOC105372935-GGT2.NCBI.GGT2.rev.compl                      ccctctttgagctgctacagcagcttctgggaatttgtaagagccatactgatgagggtc   100118

chimp.LOC112206744.LOC107973052.GGT2-.646279-708040.rev.compl      ------------------------------------------------------------   61762
FAM230B-LOC105372935-GGT2.NCBI.GGT2.rev.compl                      ccacaccctcccttgccagggcgccctcagggaggccacactgcaggaggcagagtatga   100178

chimp.LOC112206744.LOC107973052.GGT2-.646279-708040.rev.compl      ------------------------------------------------------------   61762
FAM230B-LOC105372935-GGT2.NCBI.GGT2.rev.compl                      ggtggggaagatggatcggccttatcctcaccaccccccatacgagtctcctctggtatg   100238

chimp.LOC112206744.LOC107973052.GGT2-.646279-708040.rev.compl      ------------------------------------------------------------   61762
FAM230B-LOC105372935-GGT2.NCBI.GGT2.rev.compl                      gatggtgtctcctctctggtgggaagtatgtgatttctctatttctatttctcttttgtt   100298

chimp.LOC112206744.LOC107973052.GGT2-.646279-708040.rev.compl      ------------------------------------------------------------   61762
FAM230B-LOC105372935-GGT2.NCBI.GGT2.rev.compl                      cgggcttcctagaaattcagagctaggtgtgaaattcccaattcccacagacaagcaaca   100358

chimp.LOC112206744.LOC107973052.GGT2-.646279-708040.rev.compl      ------------------------------------------------------------   61762
FAM230B-LOC105372935-GGT2.NCBI.GGT2.rev.compl                      tccccagcagtaataaggacgttccacacacccaggacttgggatatttgtgtattttca   100418

chimp.LOC112206744.LOC107973052.GGT2-.646279-708040.rev.compl      ------------------------------------------------------------   61762
FAM230B-LOC105372935-GGT2.NCBI.GGT2.rev.compl                      agaataaaagtttgccctttagcctcctcagagccccctcccttctcctcgctctcccct   100478

chimp.LOC112206744.LOC107973052.GGT2-.646279-708040.rev.compl      ------------------------------------------------------------   61762
FAM230B-LOC105372935-GGT2.NCBI.GGT2.rev.compl                      taccactgctattcatcatcttactcctacagatacccccctcttcgtggtggcaccgggc   100538

chimp.LOC112206744.LOC107973052.GGT2-.646279-708040.rev.compl      ------------------------------------------------------------   61762
FAM230B-LOC105372935-GGT2.NCBI.GGT2.rev.compl                      actagtaactcatggctgtaaagaagagctgagaacaaaacaggaagaaaagaggttaag   100598

chimp.LOC112206744.LOC107973052.GGT2-.646279-708040.rev.compl      ------------------------------------------------------------   61762
FAM230B-LOC105372935-GGT2.NCBI.GGT2.rev.compl                      taatggcaagaaaaaagatgtaccttaggaataaaaaagagttcccgaatgtcacagtgt   100658

chimp.LOC112206744.LOC107973052.GGT2-.646279-708040.rev.compl      ------------------------------------------------------------   61762
FAM230B-LOC105372935-GGT2.NCBI.GGT2.rev.compl                      atttaattaatgcttctatttattaaagattaaaaatacatatcaggtgacgacttaaaa   100718

chimp.LOC112206744.LOC107973052.GGT2-.646279-708040.rev.compl      ------------------------------------------------------------   61762
FAM230B-LOC105372935-GGT2.NCBI.GGT2.rev.compl                      tgttagtcatgattttaactgtgaattctgtggcaccggagataacagaacaatgagtgc   100778

chimp.LOC112206744.LOC107973052.GGT2-.646279-708040.rev.compl      ------------------------------------------------------------   61762
FAM230B-LOC105372935-GGT2.NCBI.GGT2.rev.compl                      ttgtagaaggcactcggttatttgttgaatgaacttcttgggaaagccttcttaggaaag   100838

chimp.LOC112206744.LOC107973052.GGT2-.646279-708040.rev.compl      ------------------------------------------------------------   61762
FAM230B-LOC105372935-GGT2.NCBI.GGT2.rev.compl                      tattagaagtagattctgctaaaacaggggctaccctcagaggtctcaggacaagggata   100898

chimp.LOC112206744.LOC107973052.GGT2-.646279-708040.rev.compl      ------------------------------------------------------------   61762
FAM230B-LOC105372935-GGT2.NCBI.GGT2.rev.compl                      gaaagtcaaacactgatgtttaagaagggatgcctatgacacatgaatcccacagccatc   100958

chimp.LOC112206744.LOC107973052.GGT2-.646279-708040.rev.compl      ------------------------------------------------------------   61762
FAM230B-LOC105372935-GGT2.NCBI.GGT2.rev.compl                      tctccagaagagcaactattttgactgcgcatgcgggaagatggcgggccgggcgacttg   101018

chimp.LOC112206744.LOC107973052.GGT2-.646279-708040.rev.compl      ------------------------------------------------------------   61762
FAM230B-LOC105372935-GGT2.NCBI.GGT2.rev.compl                      agatccgcgggtctccctgctccttttccgtctgcgtcgggagctcccgggcacgtgagg   101078

chimp.LOC112206744.LOC107973052.GGT2-.646279-708040.rev.compl      ------------------------------------------------------------   61762
FAM230B-LOC105372935-GGT2.NCBI.GGT2.rev.compl                      ccgtgccgcgtttactggcgggagggacggcctagccgggcgacgcctcggaggaggccg   101138

chimp.LOC112206744.LOC107973052.GGT2-.646279-708040.rev.compl      ------------------------------------------------------------   61762
FAM230B-LOC105372935-GGT2.NCBI.GGT2.rev.compl                      cggaccccttaggtgctgggcccttggaaatcggcgcgtgggggggcggtgctcgagctga   101198

chimp.LOC112206744.LOC107973052.GGT2-.646279-708040.rev.compl      ------------------------------------------------------------   61762
FAM230B-LOC105372935-GGT2.NCBI.GGT2.rev.compl                      gcgcgagagggcgggagagctcgtggggtgcgaggggagcaggacgcccggccgggcagc   101258

chimp.LOC112206744.LOC107973052.GGT2-.646279-708040.rev.compl      ------------------------------------------------------------   61762
FAM230B-LOC105372935-GGT2.NCBI.GGT2.rev.compl                      atgagtcagcagcggccggcgaggagattacccagtctcctcctgcacccgacggaggag   101318

chimp.LOC112206744.LOC107973052.GGT2-.646279-708040.rev.compl      ------------------------------------------------------------   61762
FAM230B-LOC105372935-GGT2.NCBI.GGT2.rev.compl                      accccatcaacgtggagggcctgctgccatcaaaaataaggattaatttagaagataatg   101378

chimp.LOC112206744.LOC107973052.GGT2-.646279-708040.rev.compl      ------------------------------------------------------------   61762
FAM230B-LOC105372935-GGT2.NCBI.GGT2.rev.compl                      tacaatatgtgtccatgagaaatctgctcccgggggtattcttgacttaaacaaggttgc   101438
```

```
chimp.LOC112206744.LOC107973052.GGT2-.646279-708040.rev.compl      ------------------------------------------------------------   61762
FAM230B-LOC105372935-GGT2.NCBI.GGT2.rev.compl                      aacgaaactgggagtccgaaagcggagagtgtatgacatcaccgatgtcttagatggaat   101498

chimp.LOC112206744.LOC107973052.GGT2-.646279-708040.rev.compl      ------------------------------------------------------------   61762
FAM230B-LOC105372935-GGT2.NCBI.GGT2.rev.compl                      cgacctcgttgaaaagaaatccaagaaccatattagatggataggatctgatcttagcaa   101558

chimp.LOC112206744.LOC107973052.GGT2-.646279-708040.rev.compl      ------------------------------------------------------------   61762
FAM230B-LOC105372935-GGT2.NCBI.GGT2.rev.compl                      ttttggagcagttccccaacaaaagaagctacaggaggaactttctgacttaccagcaat   101618

chimp.LOC112206744.LOC107973052.GGT2-.646279-708040.rev.compl      ------------------------------------------------------------   61762
FAM230B-LOC105372935-GGT2.NCBI.GGT2.rev.compl                      ggaagatgctttggatgagttaattaaggattgtgctcagcagctgtttgagttaacaga   101678

chimp.LOC112206744.LOC107973052.GGT2-.646279-708040.rev.compl      ------------------------------------------------------------   61762
FAM230B-LOC105372935-GGT2.NCBI.GGT2.rev.compl                      tgacaaagaaaatgaaagactagcatatgtgacctatcaagacattcatagcattcaggc   101738

chimp.LOC112206744.LOC107973052.GGT2-.646279-708040.rev.compl      ------------------------------------------------------------   61762
FAM230B-LOC105372935-GGT2.NCBI.GGT2.rev.compl                      cttccatgaacagatcgtcattgcagttaaagctccagcagaaaccagattggatgttcc   101798

chimp.LOC112206744.LOC107973052.GGT2-.646279-708040.rev.compl      ------------------------------------------------------------   61762
FAM230B-LOC105372935-GGT2.NCBI.GGT2.rev.compl                      agctcccagagaagactctatcacagtgcacataaggagcaccaacggacctatcgatgt   101858

chimp.LOC112206744.LOC107973052.GGT2-.646279-708040.rev.compl      ------------------------------------------------------------   61762
FAM230B-LOC105372935-GGT2.NCBI.GGT2.rev.compl                      ctatttgtgagaagtggagcagggtcagaccagtaacaaaaggtctgaaggtgtcaggac   101918

chimp.LOC112206744.LOC107973052.GGT2-.646279-708040.rev.compl      ------------------------------------------------------------   61762
FAM230B-LOC105372935-GGT2.NCBI.GGT2.rev.compl                      ctcttcatctgagagcactcatccagaaggccctgaggaagaagaaaatcctcagcaaag   101978

chimp.LOC112206744.LOC107973052.GGT2-.646279-708040.rev.compl      ------------------------------------------------------------   61762
FAM230B-LOC105372935-GGT2.NCBI.GGT2.rev.compl                      tgaagaattgcttgaagtaagcaactgatggcatttgagaatttatgtatcactgagttt   102038

chimp.LOC112206744.LOC107973052.GGT2-.646279-708040.rev.compl      ------------------------------------------------------------   61762
FAM230B-LOC105372935-GGT2.NCBI.GGT2.rev.compl                      tttgggaatatcttcctggagaattacgcatcaaatttgattctcagagcaataaattat   102098

chimp.LOC112206744.LOC107973052.GGT2-.646279-708040.rev.compl      ------------------------------------------------------------   61762
FAM230B-LOC105372935-GGT2.NCBI.GGT2.rev.compl                      ccatgaagtgctctcgttctcagtagcggcatcatggccagtagtgtctttgaggagttc   102158

chimp.LOC112206744.LOC107973052.GGT2-.646279-708040.rev.compl      ------------------------------------------------------------   61762
FAM230B-LOC105372935-GGT2.NCBI.GGT2.rev.compl                      accacttagattactgagtaattgtggtttccacatttgaaaacaactccttttataatt   102218

chimp.LOC112206744.LOC107973052.GGT2-.646279-708040.rev.compl      ------------------------------------------------------------   61762
FAM230B-LOC105372935-GGT2.NCBI.GGT2.rev.compl                      attcactgctttttgtcagtgaaatagacatcttgcctcctgaagtagcttcatcacaga   102278

chimp.LOC112206744.LOC107973052.GGT2-.646279-708040.rev.compl      ------------------------------------------------------------   61762
FAM230B-LOC105372935-GGT2.NCBI.GGT2.rev.compl                      gtgtcatgaagacagacagtcaggctgaaaaggacagttctttgtggactctacccttcc   102338

chimp.LOC112206744.LOC107973052.GGT2-.646279-708040.rev.compl      ------------------------------------------------------------   61762
FAM230B-LOC105372935-GGT2.NCBI.GGT2.rev.compl                      cttcaaggagtatgtcatatgtcacaaaagaaattgccttacactggttcatgtttgcag   102398

chimp.LOC112206744.LOC107973052.GGT2-.646279-708040.rev.compl      ------------------------------------------------------------   61762
FAM230B-LOC105372935-GGT2.NCBI.GGT2.rev.compl                      ttactgttgtacattgcatagatgtacacacgaatttaaatgtgatgtctttgtatatat   102458

chimp.LOC112206744.LOC107973052.GGT2-.646279-708040.rev.compl      ------------------------------------------------------------   61762
FAM230B-LOC105372935-GGT2.NCBI.GGT2.rev.compl                      ctgtataatgttgagattacttacgaaatatgtctgagtgacacttttcactctcgtaca   102518

chimp.LOC112206744.LOC107973052.GGT2-.646279-708040.rev.compl      ------------------------------------------------------------   61762
FAM230B-LOC105372935-GGT2.NCBI.GGT2.rev.compl                      gccaaaataatgtatatatggaaagtgacagacaaattctctaatctctttggtatctat   102578

chimp.LOC112206744.LOC107973052.GGT2-.646279-708040.rev.compl      ------------------------------------------------------------   61762
FAM230B-LOC105372935-GGT2.NCBI.GGT2.rev.compl                      aacttattagaatcctctggatgagggttagaagagactttttccaaacttctacatgta   102638

chimp.LOC112206744.LOC107973052.GGT2-.646279-708040.rev.compl      ------------------------------------------------------------   61762
FAM230B-LOC105372935-GGT2.NCBI.GGT2.rev.compl                      gaagtatcataaatgtgctacacatttatgtttgtggatttaattaaagtattttaatat   102698

chimp.LOC112206744.LOC107973052.GGT2-.646279-708040.rev.compl      ------------------------------------------------------------   61762
FAM230B-LOC105372935-GGT2.NCBI.GGT2.rev.compl                      ggttttcagtgctaaaattggagtcagatacttcttggttttaagctgtctacctaattg   102758

chimp.LOC112206744.LOC107973052.GGT2-.646279-708040.rev.compl      ------------------------------------------------------------   61762
FAM230B-LOC105372935-GGT2.NCBI.GGT2.rev.compl                      ctgtctcccagcagatcggtggcatgcccagtggctttgggggcaaggatagaaatgtca   102818

chimp.LOC112206744.LOC107973052.GGT2-.646279-708040.rev.compl      ------------------------------------------------------------   61762
FAM230B-LOC105372935-GGT2.NCBI.GGT2.rev.compl                      tcaggaaatagctgaattcattgtgaaacatgaattcagtcatggtgataattggaaact   102878

chimp.LOC112206744.LOC107973052.GGT2-.646279-708040.rev.compl      ------------------------------------------------------------   61762
FAM230B-LOC105372935-GGT2.NCBI.GGT2.rev.compl                      cctttcaggtttttgcaagtagattttgtaatgtttgtgtatgcagccttgctgttgagt   102938

chimp.LOC112206744.LOC107973052.GGT2-.646279-708040.rev.compl      ------------------------------------------------------------   61762
FAM230B-LOC105372935-GGT2.NCBI.GGT2.rev.compl                      cagtccaaggggttttacttaggacaagttgtaccttgccctctctccagctctgctccc   102998

chimp.LOC112206744.LOC107973052.GGT2-.646279-708040.rev.compl      ------------------------------------------------------------   61762
FAM230B-LOC105372935-GGT2.NCBI.GGT2.rev.compl                      acattttcacatacctagctatttctacctcattgggtaagtcatttaccactctgtgcc   103058

chimp.LOC112206744.LOC107973052.GGT2-.646279-708040.rev.compl      ------------------------------------------------------------   61762
FAM230B-LOC105372935-GGT2.NCBI.GGT2.rev.compl                      tcagtttactctgtagtttaccattagactgtgagctccctgagggactttgtcataatc   103118

chimp.LOC112206744.LOC107973052.GGT2-.646279-708040.rev.compl      ------------------------------------------------------------   61762
FAM230B-LOC105372935-GGT2.NCBI.GGT2.rev.compl                      actgttacatcccagtgcctcacaccatgcctggcccttaagaagtgctcaataaatgtc   103178

chimp.LOC112206744.LOC107973052.GGT2-.646279-708040.rev.compl      ------------------------------------------------------------   61762
FAM230B-LOC105372935-GGT2.NCBI.GGT2.rev.compl                      tgaacaaaaaaaaaaaaaaaaaaaaaaaaatcaggtctaaagatgagccatccaagaaa   103238

chimp.LOC112206744.LOC107973052.GGT2-.646279-708040.rev.compl      ------------------------------------------------------------   61762
FAM230B-LOC105372935-GGT2.NCBI.GGT2.rev.compl                      ctgctcttgctcccagaggttttaaagaaagtcattcccatgaacgtgagaagtaaccca   103298

chimp.LOC112206744.LOC107973052.GGT2-.646279-708040.rev.compl      ------------------------------------------------------------   61762
FAM230B-LOC105372935-GGT2.NCBI.GGT2.rev.compl                      acactttgaaatgataatctttttacagcaataactgttgctgcagggcttgagaaaaaa   103358

chimp.LOC112206744.LOC107973052.GGT2-.646279-708040.rev.compl      ------------------------------------------------------------   61762
FAM230B-LOC105372935-GGT2.NCBI.GGT2.rev.compl                      agttctcttcctttgggttaaagtcttaaatgctgagcccgtagccagcaaccttaaaga   103418

chimp.LOC112206744.LOC107973052.GGT2-.646279-708040.rev.compl      ------------------------------------------------------------   61762
FAM230B-LOC105372935-GGT2.NCBI.GGT2.rev.compl                      ctactcgggtccagttcattctatctggaggagtcacttgggaattgaaaacacaggaaa   103478

chimp.LOC112206744.LOC107973052.GGT2-.646279-708040.rev.compl      ------------------------------------------------------------   61762
FAM230B-LOC105372935-GGT2.NCBI.GGT2.rev.compl                      actttttttctggtccatgaacaaacaagaacgcttaactgaattagacacaaaactaag   103538

chimp.LOC112206744.LOC107973052.GGT2-.646279-708040.rev.compl      ------------------------------------------------------------   61762
FAM230B-LOC105372935-GGT2.NCBI.GGT2.rev.compl                      tatttatggttgtcatcttaacaggtttcaaacaggcttcattttaatttaacataattt   103598

chimp.LOC112206744.LOC107973052.GGT2-.646279-708040.rev.compl      ------------------------------------------------------------   61762
```

```
FAM230B-LOC105372935-GGT2.NCBI.GGT2.rev.compl                   caaaattatagcaaaatagtcaataaaatacgaacgtgcttaatttgatgaatttatgtg   103658

chimp.LOC112206744.LOC107973052.GGT2-.646279-708040.rev.compl   ------------------------------------------------------------   61762
FAM230B-LOC105372935-GGT2.NCBI.GGT2.rev.compl                   tttttgaagaaaatatgcagaataaacaagtcaattattctcactaaatacattaaaat   103718

chimp.LOC112206744.LOC107973052.GGT2-.646279-708040.rev.compl   ------------------------------------------------------------   61762
FAM230B-LOC105372935-GGT2.NCBI.GGT2.rev.compl                   ttaagggtaatatttttattatgacttgctctcctgatatagcaaaatttattgttatat   103778

chimp.LOC112206744.LOC107973052.GGT2-.646279-708040.rev.compl   ------------------------------------------------------------   61762
FAM230B-LOC105372935-GGT2.NCBI.GGT2.rev.compl                   agttatggaaggaggctcactactccccaatgatttcacaaattacagcttcagccagct   103838

chimp.LOC112206744.LOC107973052.GGT2-.646279-708040.rev.compl   ------------------------------------------------------------   61762
FAM230B-LOC105372935-GGT2.NCBI.GGT2.rev.compl                   ataaaataaccaaatcagttctgatatagccataaaattaagcctgaaaaaatagtgaag   103898

chimp.LOC112206744.LOC107973052.GGT2-.646279-708040.rev.compl   ------------------------------------------------------------   61762
FAM230B-LOC105372935-GGT2.NCBI.GGT2.rev.compl                   taaatcatccctttaaaatcaagttgttgaacaatatgttctgcatgattccatttatgt   103958

chimp.LOC112206744.LOC107973052.GGT2-.646279-708040.rev.compl   ------------------------------------------------------------   61762
FAM230B-LOC105372935-GGT2.NCBI.GGT2.rev.compl                   cagaaaaaatatatttgtgtaagtatatacacagaaaagggtctgggagaatatatacca   104018

chimp.LOC112206744.LOC107973052.GGT2-.646279-708040.rev.compl   ------------------------------------------------------------   61762
FAM230B-LOC105372935-GGT2.NCBI.GGT2.rev.compl                   aactataaagaatctatatcaaagtcaataaggggcatagacaaggaggggaactttcac   104078

chimp.LOC112206744.LOC107973052.GGT2-.646279-708040.rev.compl   ------------------------------------------------------------   61762
FAM230B-LOC105372935-GGT2.NCBI.GGT2.rev.compl                   attttactttccacattctttttttttttttttttgacggagtctcactctgttgcccagg   104138

chimp.LOC112206744.LOC107973052.GGT2-.646279-708040.rev.compl   ------------------------------------------------------------   61762
FAM230B-LOC105372935-GGT2.NCBI.GGT2.rev.compl                   ctggagtgcagtggcgcaatcttggctcactgcaagctccgcctcccgggttcacaccat   104198

chimp.LOC112206744.LOC107973052.GGT2-.646279-708040.rev.compl   ------------------------------------------------------------   61762
FAM230B-LOC105372935-GGT2.NCBI.GGT2.rev.compl                   tctcctgcctcagcctcccgagtagctgggactacaggcacccgccaccacgcccagcta   104258

chimp.LOC112206744.LOC107973052.GGT2-.646279-708040.rev.compl   ------------------------------------------------------------   61762
FAM230B-LOC105372935-GGT2.NCBI.GGT2.rev.compl                   atttttttatatttggggtttcactgcgttagccaggatggtctcaatctcctgaccttg   104318

chimp.LOC112206744.LOC107973052.GGT2-.646279-708040.rev.compl   ------------------------------------------------------------   61762
FAM230B-LOC105372935-GGT2.NCBI.GGT2.rev.compl                   tgatctgcccacctcacccacctcagcctcccaaagtgctgggattacaggcgtgacttt   104378

chimp.LOC112206744.LOC107973052.GGT2-.646279-708040.rev.compl   ------------------------------------------------------------   61762
FAM230B-LOC105372935-GGT2.NCBI.GGT2.rev.compl                   ctacattcttgtaacagaattatagtcaccgttacctccatcaaaaaattaaaattcaaa   104438

chimp.LOC112206744.LOC107973052.GGT2-.646279-708040.rev.compl   ------------------------------------------------------------   61762
FAM230B-LOC105372935-GGT2.NCBI.GGT2.rev.compl                   aattaggccaggcacggtggctcacgcctgtaatcccagcactttgggagtctggagcag   104498

chimp.LOC112206744.LOC107973052.GGT2-.646279-708040.rev.compl   ------------------------------------------------------------   61762
FAM230B-LOC105372935-GGT2.NCBI.GGT2.rev.compl                   cagatcacctgaggtcaggagttggagagcagcctgactggccaacatggtgaaacccca   104558

chimp.LOC112206744.LOC107973052.GGT2-.646279-708040.rev.compl   ------------------------------------------------------------   61762
FAM230B-LOC105372935-GGT2.NCBI.GGT2.rev.compl                   tctctactaaaaatacaaaaattagccgggcgtggtggagggcacctgtaatcccagcta   104618

chimp.LOC112206744.LOC107973052.GGT2-.646279-708040.rev.compl   ------------------------------------------------------------   61762
FAM230B-LOC105372935-GGT2.NCBI.GGT2.rev.compl                   cttgggaggctaaggcaggagaatcacttaaacccgggaggcggaggttgcagtgagcca   104678

chimp.LOC112206744.LOC107973052.GGT2-.646279-708040.rev.compl   ------------------------------------------------------------   61762
FAM230B-LOC105372935-GGT2.NCBI.GGT2.rev.compl                   agattgtgccactgcactccggcccgggtaacagagcgagactccatctcaaaaatgaat   104738

chimp.LOC112206744.LOC107973052.GGT2-.646279-708040.rev.compl   ------------------------------------------------------------   61762
FAM230B-LOC105372935-GGT2.NCBI.GGT2.rev.compl                   gaattaattaattaattaaagcaaacaaatataagctacctcctgaagtagaaaccagta   104798

chimp.LOC112206744.LOC107973052.GGT2-.646279-708040.rev.compl   ------------------------------------------------------------   61762
FAM230B-LOC105372935-GGT2.NCBI.GGT2.rev.compl                   tctcttcaccttacctttacataatactttaccctaatgcattttatctgctgttctaaa   104858

chimp.LOC112206744.LOC107973052.GGT2-.646279-708040.rev.compl   ------------------------------------------------------------   61762
FAM230B-LOC105372935-GGT2.NCBI.GGT2.rev.compl                   tcatgacttacttcaaatcaaatcttaagtgaaaatcaagatttccttttggggcatta   104918

chimp.LOC112206744.LOC107973052.GGT2-.646279-708040.rev.compl   ------------------------------------------------------------   61762
FAM230B-LOC105372935-GGT2.NCBI.GGT2.rev.compl                   tacaaaaaagactgtaaaatttcattctctgaaagtctttttaaaaagggatagattgta   104978

chimp.LOC112206744.LOC107973052.GGT2-.646279-708040.rev.compl   ------------------------------------------------------------   61762
FAM230B-LOC105372935-GGT2.NCBI.GGT2.rev.compl                   tctggctgggatagggtaggtgtgatcggacccggaggaaatgacggctgctcaactgtg   105038

chimp.LOC112206744.LOC107973052.GGT2-.646279-708040.rev.compl   ------------------------------------------------------------   61762
FAM230B-LOC105372935-GGT2.NCBI.GGT2.rev.compl                   catctcctttcctcaccctgctgaaaagtgccacctagcatggtcaatcaaggtcaccaa   105098

chimp.LOC112206744.LOC107973052.GGT2-.646279-708040.rev.compl   ------------------------------------------------------------   61762
FAM230B-LOC105372935-GGT2.NCBI.GGT2.rev.compl                   tgcaaagcagtagtgggcaaacaccacccaggttatccaaagcagcacatggggttggga   105158

chimp.LOC112206744.LOC107973052.GGT2-.646279-708040.rev.compl   ------------------------------------------------------------   61762
FAM230B-LOC105372935-GGT2.NCBI.GGT2.rev.compl                   aactgatgaccttggagatttcaaaatcacctaacaataaccccaccttccctaaccaca   105218

chimp.LOC112206744.LOC107973052.GGT2-.646279-708040.rev.compl   ------------------------------------------------------------   61762
FAM230B-LOC105372935-GGT2.NCBI.GGT2.rev.compl                   gggctaagcaacctaatggatggaacatatgtgtgctttggtcatccgcaacccagacca   105278

chimp.LOC112206744.LOC107973052.GGT2-.646279-708040.rev.compl   ------------------------------------------------------------   61762
FAM230B-LOC105372935-GGT2.NCBI.GGT2.rev.compl                   ctagactaattttgctttagaactggactatatcaagtgtgaacgatcatacttgttatc   105338

chimp.LOC112206744.LOC107973052.GGT2-.646279-708040.rev.compl   ------------------------------------------------------------   61762
FAM230B-LOC105372935-GGT2.NCBI.GGT2.rev.compl                   atgtgggaaaaaaggtttaattactccctaattgtctgggaaacaaaagatttcaaaata   105398

chimp.LOC112206744.LOC107973052.GGT2-.646279-708040.rev.compl   ------------------------------------------------------------   61762
FAM230B-LOC105372935-GGT2.NCBI.GGT2.rev.compl                   tggagctcctgggggaaggagagttaataaaatgggtccttgagaagtgaagtgggaagc   105458

chimp.LOC112206744.LOC107973052.GGT2-.646279-708040.rev.compl   ------------------------------------------------------------   61762
FAM230B-LOC105372935-GGT2.NCBI.GGT2.rev.compl                   ctgcctcatgccatggcaccagcgatgccacaggtgtacccagtgccctgccgagagcag   105518

chimp.LOC112206744.LOC107973052.GGT2-.646279-708040.rev.compl   ------------------------------------------------------------   61762
FAM230B-LOC105372935-GGT2.NCBI.GGT2.rev.compl                   agacggtgttctcagagttgctatcaaaaaggaggcagcagactgcattctgaagcatca   105578

chimp.LOC112206744.LOC107973052.GGT2-.646279-708040.rev.compl   ------------------------------------------------------------   61762
FAM230B-LOC105372935-GGT2.NCBI.GGT2.rev.compl                   gacaatgggtcacatgcggaaacagaattaaatacacaaaaatagcaccaacttaaggca   105638

chimp.LOC112206744.LOC107973052.GGT2-.646279-708040.rev.compl   ------------------------------------------------------------   61762
FAM230B-LOC105372935-GGT2.NCBI.GGT2.rev.compl                   agcctgggcaaactgcactacatcaaagaaagaacacacaactggtatcagattttttt   105698

chimp.LOC112206744.LOC107973052.GGT2-.646279-708040.rev.compl   ------------------------------------------------------------   61762
FAM230B-LOC105372935-GGT2.NCBI.GGT2.rev.compl                   tctgttttaataaagagaaggcctggacacctgaaagttgacgggaagcaggcatcacta   105758

chimp.LOC112206744.LOC107973052.GGT2-.646279-708040.rev.compl   ------------------------------------------------------------   61762
FAM230B-LOC105372935-GGT2.NCBI.GGT2.rev.compl                   atttctccgagttcacagttttgttagaatgtctttgctctgattatggttttatttatt   105818
```

```
chimp.LOC112206744.LOC107973052.GGT2-.646279-708040.rev.compl      ------------------------------------------------------------   61762
FAM230B-LOC105372935-GGT2.NCBI.GGT2.rev.compl                      tatttttttgagatggagtttttttttgttttgttttatttgcttttgttgccctggctgg   105878

chimp.LOC112206744.LOC107973052.GGT2-.646279-708040.rev.compl      ------------------------------------------------------------   61762
FAM230B-LOC105372935-GGT2.NCBI.GGT2.rev.compl                      agtgcagtggtgcaatctcggctcaccgcaaactccgcctcccaggttcaagagattctc   105938

chimp.LOC112206744.LOC107973052.GGT2-.646279-708040.rev.compl      ------------------------------------------------------------   61762
FAM230B-LOC105372935-GGT2.NCBI.GGT2.rev.compl                      ctgcctcagcctcccaagtagctaggattacaggcacccactaccacgtctggcttcttt   105998

chimp.LOC112206744.LOC107973052.GGT2-.646279-708040.rev.compl      ------------------------------------------------------------   61762
FAM230B-LOC105372935-GGT2.NCBI.GGT2.rev.compl                      gtgtttttagtagagatggggtttcaccatgttggtcaggctggtcttgaactcttgacc   106058

chimp.LOC112206744.LOC107973052.GGT2-.646279-708040.rev.compl      ------------------------------------------------------------   61762
FAM230B-LOC105372935-GGT2.NCBI.GGT2.rev.compl                      tcaagcgatccacccgcctcagccccccaaagtgctggaattacaggcatgagccaccac   106118

chimp.LOC112206744.LOC107973052.GGT2-.646279-708040.rev.compl      ------------------------------------------------------------   61762
FAM230B-LOC105372935-GGT2.NCBI.GGT2.rev.compl                      gcctggcctgattatgttttgatggatatacagaagctcttgctttggtacagaaacagg   106178

chimp.LOC112206744.LOC107973052.GGT2-.646279-708040.rev.compl      ------------------------------------------------------------   61762
FAM230B-LOC105372935-GGT2.NCBI.GGT2.rev.compl                      ccccgctttgcttcgacctctgggccccccaccttcagaaatgtctggtgccattcctgg   106238

chimp.LOC112206744.LOC107973052.GGT2-.646279-708040.rev.compl      ------------------------------------------------------------   61762
FAM230B-LOC105372935-GGT2.NCBI.GGT2.rev.compl                      ctcctcctctcagtggtcttccaggatatccactcacagctttctgatgtcacccacctg   106298

chimp.LOC112206744.LOC107973052.GGT2-.646279-708040.rev.compl      ------------------------------------------------------------   61762
FAM230B-LOC105372935-GGT2.NCBI.GGT2.rev.compl                      tctctagcccagatacccagtcctcatggcaagtgactgcagcaatttttacaaatctac   106358

chimp.LOC112206744.LOC107973052.GGT2-.646279-708040.rev.compl      ------------------------------------------------------------   61762
FAM230B-LOC105372935-GGT2.NCBI.GGT2.rev.compl                      gcccttagggttcccatcccaggacgttcaggtttctccacccctttagaagtgcccaga   106418

chimp.LOC112206744.LOC107973052.GGT2-.646279-708040.rev.compl      ------------------------------------------------------------   61762
FAM230B-LOC105372935-GGT2.NCBI.GGT2.rev.compl                      atatcccctaccccacgattccaacaggacgtcactctcctcttacttaagaaaaatgcc   106478

chimp.LOC112206744.LOC107973052.GGT2-.646279-708040.rev.compl      ------------------------------------------------------------   61762
FAM230B-LOC105372935-GGT2.NCBI.GGT2.rev.compl                      taaaattccagcccttccaaaaggcctttctagattgaccactcaggagttactctggcc   106538

chimp.LOC112206744.LOC107973052.GGT2-.646279-708040.rev.compl      ------------------------------------------------------------   61762
FAM230B-LOC105372935-GGT2.NCBI.GGT2.rev.compl                      tccccttcactcctgctagactgaaaacaaggacagacttgcctggcctctctcccccag   106598

chimp.LOC112206744.LOC107973052.GGT2-.646279-708040.rev.compl      ------------------------------------------------------------   61762
FAM230B-LOC105372935-GGT2.NCBI.GGT2.rev.compl                      tagacacctacccccagtgaacacatcctgtcagtgtcctcccgtccatttgcaaattga   106658

chimp.LOC112206744.LOC107973052.GGT2-.646279-708040.rev.compl      ------------------------------------------------------------   61762
FAM230B-LOC105372935-GGT2.NCBI.GGT2.rev.compl                      ttgactgtttcattgctggccttggatgacctctcctagagggcagggacggtggtgttc   106718

chimp.LOC112206744.LOC107973052.GGT2-.646279-708040.rev.compl      ------------------------------------------------------------   61762
FAM230B-LOC105372935-GGT2.NCBI.GGT2.rev.compl                      ggcaaagccttggtcgtctggataatgggaaatggggtcttctggtgaatttaattacag   106778

chimp.LOC112206744.LOC107973052.GGT2-.646279-708040.rev.compl      ------------------------------------------------------------   61762
FAM230B-LOC105372935-GGT2.NCBI.GGT2.rev.compl                      ggttaggtggaaactttgcttcaactctcattattagaaagtctattgtaaagcttttta   106838

chimp.LOC112206744.LOC107973052.GGT2-.646279-708040.rev.compl      ------------------------------------------------------------   61762
FAM230B-LOC105372935-GGT2.NCBI.GGT2.rev.compl                      aaatacaactgaccactttccagcgagggtagagttaatatgccttaatctttgatttag   106898

chimp.LOC112206744.LOC107973052.GGT2-.646279-708040.rev.compl      ------------------------------------------------------------   61762
FAM230B-LOC105372935-GGT2.NCBI.GGT2.rev.compl                      gatcttgcagtgccccaggctcaccattctcttctgctactataattatagatgcggaag   106958

chimp.LOC112206744.LOC107973052.GGT2-.646279-708040.rev.compl      ------------------------------------------------------------   61762
FAM230B-LOC105372935-GGT2.NCBI.GGT2.rev.compl                      acatacagactgggctgacaactgcttggaagctaatttctagaataaacccttgctacc   107018

chimp.LOC112206744.LOC107973052.GGT2-.646279-708040.rev.compl      ------------------------------------------------------------   61762
FAM230B-LOC105372935-GGT2.NCBI.GGT2.rev.compl                      acatcttgtgattttttgcccttctgaaggtggagttccaaaataaaaacaaaatcccgct   107078

chimp.LOC112206744.LOC107973052.GGT2-.646279-708040.rev.compl      ------------------------------------------------------------   61762
FAM230B-LOC105372935-GGT2.NCBI.GGT2.rev.compl                      taactgagggttccagttgagggaggtctgctgccatggtctttgttaagtgcctgccac   107138

chimp.LOC112206744.LOC107973052.GGT2-.646279-708040.rev.compl      ------------------------------------------------------------   61762
FAM230B-LOC105372935-GGT2.NCBI.GGT2.rev.compl                      catggcccacaattagtcagttaataaagggttcttgatgagagaatttggcaacatgtg   107198

chimp.LOC112206744.LOC107973052.GGT2-.646279-708040.rev.compl      ------------------------------------------------------------   61762
FAM230B-LOC105372935-GGT2.NCBI.GGT2.rev.compl                      attttccacaaatataagcttgattttgatgcaaccactcatttctttgaaaatcagggg   107258

chimp.LOC112206744.LOC107973052.GGT2-.646279-708040.rev.compl      ------------------------------------------------------------   61762
FAM230B-LOC105372935-GGT2.NCBI.GGT2.rev.compl                      gcattcacacacaggtgttctagatttccttacagtaaaaccaacccttttacaaggaata   107318

chimp.LOC112206744.LOC107973052.GGT2-.646279-708040.rev.compl      ------------------------------------------------------------   61762
FAM230B-LOC105372935-GGT2.NCBI.GGT2.rev.compl                      gctgtgttcactttgaagcagtaaagaagcaggtgtccactcatggcaaatgaggaactc   107378

chimp.LOC112206744.LOC107973052.GGT2-.646279-708040.rev.compl      ------------------------------------------------------------   61762
FAM230B-LOC105372935-GGT2.NCBI.GGT2.rev.compl                      ccttttttttttttttttttttttttttttttttgagacggggtctcgctctttttttgt   107438

chimp.LOC112206744.LOC107973052.GGT2-.646279-708040.rev.compl      ------------------------------------------------------------   61762
FAM230B-LOC105372935-GGT2.NCBI.GGT2.rev.compl                      gtgtgtgagatgggggccacccaggctagagtgcagtagtgcaattacagctcactgtag   107498

chimp.LOC112206744.LOC107973052.GGT2-.646279-708040.rev.compl      ------------------------------------------------------------   61762
FAM230B-LOC105372935-GGT2.NCBI.GGT2.rev.compl                      cctcaaccttctgggctcaaccagttcttcccacctcaacttctgaagcagctgagacta   107558

chimp.LOC112206744.LOC107973052.GGT2-.646279-708040.rev.compl      ------------------------------------------------------------   61762
FAM230B-LOC105372935-GGT2.NCBI.GGT2.rev.compl                      caggtgtgtaccaccatgcccaggtaatgcttttgatttttagtaagatgaggtctcgtt   107618

chimp.LOC112206744.LOC107973052.GGT2-.646279-708040.rev.compl      ------------------------------------------------------------   61762
FAM230B-LOC105372935-GGT2.NCBI.GGT2.rev.compl                      atattgcccaggctggtctcaaactcctgagctcaagtgatcctcctgcctcagcctccc   107678

chimp.LOC112206744.LOC107973052.GGT2-.646279-708040.rev.compl      ------------------------------------------------------------   61762
FAM230B-LOC105372935-GGT2.NCBI.GGT2.rev.compl                      aaagtgctaggattatcggcatgagccactatgcccagccagaagctaaaaatattaatc   107738

chimp.LOC112206744.LOC107973052.GGT2-.646279-708040.rev.compl      ------------------------------------------------------------   61762
FAM230B-LOC105372935-GGT2.NCBI.GGT2.rev.compl                      aaggagtcacaactaaggatgcactgcacacccgaagccagggaccacaagaggtcagca   107798

chimp.LOC112206744.LOC107973052.GGT2-.646279-708040.rev.compl      ------------------------------------------------------------   61762
FAM230B-LOC105372935-GGT2.NCBI.GGT2.rev.compl                      actgggactgcttcctccagttcccaggctggggctcagcagcagtggcctgggctctgg   107858

chimp.LOC112206744.LOC107973052.GGT2-.646279-708040.rev.compl      ------------------------------------------------------------   61762
FAM230B-LOC105372935-GGT2.NCBI.GGT2.rev.compl                      ggacttactttaagccgaaatcccaaactcattatcattcaaaccaattttctgtgtcca   107918

chimp.LOC112206744.LOC107973052.GGT2-.646279-708040.rev.compl      ------------------------------------------------------------   61762
FAM230B-LOC105372935-GGT2.NCBI.GGT2.rev.compl                      gcatcatcccacctaaactggtccttaaaaccactgacagtcatttctccaatctgtaaa   107978
```

```
chimp.LOC112206744.LOC107973052.GGT2-.646279-708040.rev.compl      ------------------------------------------------------------   61762
FAM230B-LOC105372935-GGT2.NCBI.GGT2.rev.compl                       aggcctgctgaggcctttgcattatcaggacttaactgggagtagaatgccctccccatc   108038

chimp.LOC112206744.LOC107973052.GGT2-.646279-708040.rev.compl      ------------------------------------------------------------   61762
FAM230B-LOC105372935-GGT2.NCBI.GGT2.rev.compl                       caaatcctgaccatttctgaacactcagcccatgtcctcttcctgagtgtcttccccaac   108098

chimp.LOC112206744.LOC107973052.GGT2-.646279-708040.rev.compl      ------------------------------------------------------------   61762
FAM230B-LOC105372935-GGT2.NCBI.GGT2.rev.compl                       aacaaccacctacactgaactttactttgatctctgcaggaaactgtagatagagatggg   108158

chimp.LOC112206744.LOC107973052.GGT2-.646279-708040.rev.compl      ------------------------------------------------------------   61762
FAM230B-LOC105372935-GGT2.NCBI.GGT2.rev.compl                       gtctcactatgttgcctagctggtcttgaactgctgagctcaagtgatctccctgcttct   108218

chimp.LOC112206744.LOC107973052.GGT2-.646279-708040.rev.compl      ------------------------------------------------------------   61762
FAM230B-LOC105372935-GGT2.NCBI.GGT2.rev.compl                       gcctcccaaagtgctaggattacagcagtgagccaccacacctggcctatgtatgctttt   108278

chimp.LOC112206744.LOC107973052.GGT2-.646279-708040.rev.compl      ------------------------------------------------------------   61762
FAM230B-LOC105372935-GGT2.NCBI.GGT2.rev.compl                       tttttcttttcttttttttttttttttgagacagagtcttgctctgtggcccagtccgc   108338

chimp.LOC112206744.LOC107973052.GGT2-.646279-708040.rev.compl      ------------------------------------------------------------   61762
FAM230B-LOC105372935-GGT2.NCBI.GGT2.rev.compl                       agtgcagtggtgcaatctcggctcatggcaacctccaggttcaaggcattttcctgcctc   108398

chimp.LOC112206744.LOC107973052.GGT2-.646279-708040.rev.compl      ------------------------------------------------------------   61762
FAM230B-LOC105372935-GGT2.NCBI.GGT2.rev.compl                       agccttccaaggagctgggattacaggcacgcaccatcacgcccagctaattggccaggc   108458

chimp.LOC112206744.LOC107973052.GGT2-.646279-708040.rev.compl      ------------------------------------------------------------   61762
FAM230B-LOC105372935-GGT2.NCBI.GGT2.rev.compl                       tggtctcaaactcctgacctcaggtgatccacccaccttggcctcccaaagtgctgggat   108518

chimp.LOC112206744.LOC107973052.GGT2-.646279-708040.rev.compl      ------------------------------------------------------------   61762
FAM230B-LOC105372935-GGT2.NCBI.GGT2.rev.compl                       tacaggtgtaagccaccacaccgggccacgtatgcattttaaaacgttatgctcagacgg   108578

chimp.LOC112206744.LOC107973052.GGT2-.646279-708040.rev.compl      ------------------------------------------------------------   61762
FAM230B-LOC105372935-GGT2.NCBI.GGT2.rev.compl                       ggtctataggtttcaccagactgccaaaaggatccatgacacaaaaaaagtactctcagt   108638

chimp.LOC112206744.LOC107973052.GGT2-.646279-708040.rev.compl      ------------------------------------------------------------   61762
FAM230B-LOC105372935-GGT2.NCBI.GGT2.rev.compl                       tgcatccttcggaacctttcaacacatacacatatacctcactcatccttttgtgaagag   108698

chimp.LOC112206744.LOC107973052.GGT2-.646279-708040.rev.compl      ------------------------------------------------------------   61762
FAM230B-LOC105372935-GGT2.NCBI.GGT2.rev.compl                       tgtctattaatcactagattaagaggatttggataaaatttataaatgtacaaactttgc   108758

chimp.LOC112206744.LOC107973052.GGT2-.646279-708040.rev.compl      ------------------------------------------------------------   61762
FAM230B-LOC105372935-GGT2.NCBI.GGT2.rev.compl                       taccaaggactgacacatttacttttctagctcatcagtttatgtcctagaaaaattcca   108818

chimp.LOC112206744.LOC107973052.GGT2-.646279-708040.rev.compl      ------------------------------------------------------------   61762
FAM230B-LOC105372935-GGT2.NCBI.GGT2.rev.compl                       gagccacattaaccagctgtttcctatttttcattttacgatggtgtttccactttcacc   108878

chimp.LOC112206744.LOC107973052.GGT2-.646279-708040.rev.compl      ------------------------------------------------------------   61762
FAM230B-LOC105372935-GGT2.NCBI.GGT2.rev.compl                       tgcaaagtgactaatagtaatacctattacagtaatttaagtaacaccatttagagtaat   108938

chimp.LOC112206744.LOC107973052.GGT2-.646279-708040.rev.compl      ------------------------------------------------------------   61762
FAM230B-LOC105372935-GGT2.NCBI.GGT2.rev.compl                       ttcatctttagtactgtatttatactactaagaaagccgccaacgattcaagctgcactc   108998

chimp.LOC112206744.LOC107973052.GGT2-.646279-708040.rev.compl      ------------------------------------------------------------   61762
FAM230B-LOC105372935-GGT2.NCBI.GGT2.rev.compl                       actggtgaaaccctatgctatatatgtaattaatttatttcctaaattcttacctccctc   109058

chimp.LOC112206744.LOC107973052.GGT2-.646279-708040.rev.compl      ------------------------------------------------------------   61762
FAM230B-LOC105372935-GGT2.NCBI.GGT2.rev.compl                       cacccctccattggaaaataaactctaagaatgcaaaagtttttgtgtgtttttcttca   109118

chimp.LOC112206744.LOC107973052.GGT2-.646279-708040.rev.compl      ------------------------------------------------------------   61762
FAM230B-LOC105372935-GGT2.NCBI.GGT2.rev.compl                       acgcttagagcagtgcctggcacctaacagtcaatatctgctaactttgatgaatgaatg   109178

chimp.LOC112206744.LOC107973052.GGT2-.646279-708040.rev.compl      ------------------------------------------------------------   61762
FAM230B-LOC105372935-GGT2.NCBI.GGT2.rev.compl                       gtatatttatattatgcaatattactatgcagccatgaaacacaatgagttagagtggta   109238

chimp.LOC112206744.LOC107973052.GGT2-.646279-708040.rev.compl      ------------------------------------------------------------   61762
FAM230B-LOC105372935-GGT2.NCBI.GGT2.rev.compl                       ccagctaatttaatggaatttccagaagtatgaaattaagaaaacaagaagaaaagtgtc   109298

chimp.LOC112206744.LOC107973052.GGT2-.646279-708040.rev.compl      ------------------------------------------------------------   61762
FAM230B-LOC105372935-GGT2.NCBI.GGT2.rev.compl                       aaaaatatcatcccactttttttttttcttccctgaaacggagtcgtgctctgacacca   109358

chimp.LOC112206744.LOC107973052.GGT2-.646279-708040.rev.compl      ------------------------------------------------------------   61762
FAM230B-LOC105372935-GGT2.NCBI.GGT2.rev.compl                       ggctacagtatagtgctgccatcatggctcactgcaacctcaaccttctgggctcaggga   109418

chimp.LOC112206744.LOC107973052.GGT2-.646279-708040.rev.compl      ------------------------------------------------------------   61762
FAM230B-LOC105372935-GGT2.NCBI.GGT2.rev.compl                       atccttccacctcagcctcccatgtagctgggactacaggtgtgtgccacaacgcccagc   109478

chimp.LOC112206744.LOC107973052.GGT2-.646279-708040.rev.compl      ------------------------------------------------------------   61762
FAM230B-LOC105372935-GGT2.NCBI.GGT2.rev.compl                       taatttattttaaattattttttgtagagacagggtctccctatgttgcccaggctggtc   109538

chimp.LOC112206744.LOC107973052.GGT2-.646279-708040.rev.compl      ------------------------------------------------------------   61762
FAM230B-LOC105372935-GGT2.NCBI.GGT2.rev.compl                       tcaaactactaggctcaagtgatcctcccacctcagcttcccaaagtgttgggattacag   109598

chimp.LOC112206744.LOC107973052.GGT2-.646279-708040.rev.compl      ------------------------------------------------------------   61762
FAM230B-LOC105372935-GGT2.NCBI.GGT2.rev.compl                       gtgtgagccactgtgcccagcccaatcccactttttaaaaagtataccccaaatccata   109658

chimp.LOC112206744.LOC107973052.GGT2-.646279-708040.rev.compl      ------------------------------------------------------------   61762
FAM230B-LOC105372935-GGT2.NCBI.GGT2.rev.compl                       cactggtgaggaagtggagaacgtggaatcctcacacacagaatgtaaaatggtacagct   109718

chimp.LOC112206744.LOC107973052.GGT2-.646279-708040.rev.compl      ------------------------------------------------------------   61762
FAM230B-LOC105372935-GGT2.NCBI.GGT2.rev.compl                       ctggagaacattttagcagtttctcaaaatgtttaacgtagctactacacgacccagcaa   109778

chimp.LOC112206744.LOC107973052.GGT2-.646279-708040.rev.compl      ------------------------------------------------------------   61762
FAM230B-LOC105372935-GGT2.NCBI.GGT2.rev.compl                       ttcaactcccagatatacacctgagagaactgaaaggatatgtccacacaaaaatttgca   109838

chimp.LOC112206744.LOC107973052.GGT2-.646279-708040.rev.compl      ------------------------------------------------------------   61762
FAM230B-LOC105372935-GGT2.NCBI.GGT2.rev.compl                       gaggaacttgtacacacttgttcacagtagcattattcataatagcccaaagtggcaaaa   109898

chimp.LOC112206744.LOC107973052.GGT2-.646279-708040.rev.compl      ------------------------------------------------------------   61762
FAM230B-LOC105372935-GGT2.NCBI.GGT2.rev.compl                       acgcaaatgttcatcaactgatggacaaataaagcatgaacaactgatggacaaataaag   109958

chimp.LOC112206744.LOC107973052.GGT2-.646279-708040.rev.compl      ------------------------------------------------------------   61762
FAM230B-LOC105372935-GGT2.NCBI.GGT2.rev.compl                       tatatcattcagcatataagaaatgaagtgctaatacatgctacaacaggaatagacctt   110018

chimp.LOC112206744.LOC107973052.GGT2-.646279-708040.rev.compl      ------------------------------------------------------------   61762
FAM230B-LOC105372935-GGT2.NCBI.GGT2.rev.compl                       aataacattttgctaaatgaaaaaagccaaacaaaaggtcacatgttgtatgattctatt   110078

chimp.LOC112206744.LOC107973052.GGT2-.646279-708040.rev.compl      ------------------------------------------------------------   61762
FAM230B-LOC105372935-GGT2.NCBI.GGT2.rev.compl                       tacataaaaatacctagactaggcaaatccatagagacacaaagatcagtgactgcccag   110138
```

```
chimp.LOC112206744.LOC107973052.GGT2-.646279-708040.rev.compl      ------------------------------------------------------------   61762
FAM230B-LOC105372935-GGT2.NCBI.GGT2.rev.compl                      agccagagggagtggtgaatgtgaaatggctgctaatgagtacagacttctttttgaggt   110198

chimp.LOC112206744.LOC107973052.GGT2-.646279-708040.rev.compl      ------------------------------------------------------------   61762
FAM230B-LOC105372935-GGT2.NCBI.GGT2.rev.compl                      attgaacatgttatggaatcagatggtggtgatcaatgcataactttgtgaatataccaa   110258

chimp.LOC112206744.LOC107973052.GGT2-.646279-708040.rev.compl      ------------------------------------------------------------   61762
FAM230B-LOC105372935-GGT2.NCBI.GGT2.rev.compl                      aaaccacatgaaaggtacactttaaaagcctgaattttaaggtatgtgaattgtatctcc   110318

chimp.LOC112206744.LOC107973052.GGT2-.646279-708040.rev.compl      ------------------------------------------------------------   61762
FAM230B-LOC105372935-GGT2.NCBI.GGT2.rev.compl                      ataaacctgtgattttgttaatatacgcttatttgcaaatgaatacatggataatgagga   110378

chimp.LOC112206744.LOC107973052.GGT2-.646279-708040.rev.compl      ------------------------------------------------------------   61762
FAM230B-LOC105372935-GGT2.NCBI.GGT2.rev.compl                      ataatatgaaaggttaatgacaaccttggtgatagggtacaggagactgggagaggaaga   110438

chimp.LOC112206744.LOC107973052.GGT2-.646279-708040.rev.compl      ------------------------------------------------------------   61762
FAM230B-LOC105372935-GGT2.NCBI.GGT2.rev.compl                      taagtgttgaggtagaagaagacaaaaaataaaaggctaaaacaataatccagggccaga   110498

chimp.LOC112206744.LOC107973052.GGT2-.646279-708040.rev.compl      ------------------------------------------------------------   61762
FAM230B-LOC105372935-GGT2.NCBI.GGT2.rev.compl                      ggccatgagttgggagatggagcaagacgaagtgtacaaagaaatgagcataaatcataa   110558

chimp.LOC112206744.LOC107973052.GGT2-.646279-708040.rev.compl      ------------------------------------------------------------   61762
FAM230B-LOC105372935-GGT2.NCBI.GGT2.rev.compl                      aaatgagggaactagggttgaaaactccagcaagtagtttgtaatgggggagatgaagca   110618

chimp.LOC112206744.LOC107973052.GGT2-.646279-708040.rev.compl      ------------------------------------------------------------   61762
FAM230B-LOC105372935-GGT2.NCBI.GGT2.rev.compl                      aacctaagtaactgcaatgattcaacggaatttgcagaaaatgcagaagggcgagaaaag   110678

chimp.LOC112206744.LOC107973052.GGT2-.646279-708040.rev.compl      ------------------------------------------------------------   61762
FAM230B-LOC105372935-GGT2.NCBI.GGT2.rev.compl                      caatgaaaaaatggatcagaagatttaaaattggctctaccccaccaaagttttaaaaag   110738

chimp.LOC112206744.LOC107973052.GGT2-.646279-708040.rev.compl      ------------------------------------------------------------   61762
FAM230B-LOC105372935-GGT2.NCBI.GGT2.rev.compl                      caagcaagaagaaaattcagttgctcctgggcaagaattcttgggcccagtggggtagaa   110798

chimp.LOC112206744.LOC107973052.GGT2-.646279-708040.rev.compl      ------------------------------------------------------------   61762
FAM230B-LOC105372935-GGT2.NCBI.GGT2.rev.compl                      ccaaaccatgataaggagacaccagtgttttgtaagcaagagaatatattaaggagagaa   110858

chimp.LOC112206744.LOC107973052.GGT2-.646279-708040.rev.compl      ------------------------------------------------------------   61762
FAM230B-LOC105372935-GGT2.NCBI.GGT2.rev.compl                      aaggggcatttattgggagaaaaggtctcactgtgattgttctggaataaagtagcccca   110918

chimp.LOC112206744.LOC107973052.GGT2-.646279-708040.rev.compl      ------------------------------------------------------------   61762
FAM230B-LOC105372935-GGT2.NCBI.GGT2.rev.compl                      tcggactgcctcaggaactcatggtggaagaggtgccactggaaaatggctaagtcggga   110978

chimp.LOC112206744.LOC107973052.GGT2-.646279-708040.rev.compl      ------------------------------------------------------------   61762
FAM230B-LOC105372935-GGT2.NCBI.GGT2.rev.compl                      tcttccccaagaacactagaagagacagaggaagaatagaagagctttaaattttagcag   111038

chimp.LOC112206744.LOC107973052.GGT2-.646279-708040.rev.compl      ------------------------------------------------------------   61762
FAM230B-LOC105372935-GGT2.NCBI.GGT2.rev.compl                      atatttctttttcttgtttaattaaaataaatcttattgacctaatattttgagactgg   111098

chimp.LOC112206744.LOC107973052.GGT2-.646279-708040.rev.compl      ------------------------------------------------------------   61762
FAM230B-LOC105372935-GGT2.NCBI.GGT2.rev.compl                      gctcttgcaatgctccccagactgaactcaaactcctaggctcaagagattcttccactt   111158

chimp.LOC112206744.LOC107973052.GGT2-.646279-708040.rev.compl      ------------------------------------------------------------   61762
FAM230B-LOC105372935-GGT2.NCBI.GGT2.rev.compl                      cagcctcctgaaaagctggcactacaggtgcacaccactgggcccagcataatggctatt   111218

chimp.LOC112206744.LOC107973052.GGT2-.646279-708040.rev.compl      ------------------------------------------------------------   61762
FAM230B-LOC105372935-GGT2.NCBI.GGT2.rev.compl                      tcttgacaaatcttgacaatgccaccagtatgtttttctttagagacagagtctcgcttt   111278

chimp.LOC112206744.LOC107973052.GGT2-.646279-708040.rev.compl      ------------------------------------------------------------   61762
FAM230B-LOC105372935-GGT2.NCBI.GGT2.rev.compl                      gtagcccaggctggagaacagtggtgtgatcctggctcactgcagcctcaaacacctgga   111338

chimp.LOC112206744.LOC107973052.GGT2-.646279-708040.rev.compl      ------------------------------------------------------------   61762
FAM230B-LOC105372935-GGT2.NCBI.GGT2.rev.compl                      ctcaggagatccccctgcctcagcctctcaggtagctgagactacaggtgcgcaccacca   111398

chimp.LOC112206744.LOC107973052.GGT2-.646279-708040.rev.compl      ------------------------------------------------------------   61762
FAM230B-LOC105372935-GGT2.NCBI.GGT2.rev.compl                      cacttggcgaatttattttatttttttttattatactttaagttctagggtacatacac   111458

chimp.LOC112206744.LOC107973052.GGT2-.646279-708040.rev.compl      ------------------------------------------------------------   61762
FAM230B-LOC105372935-GGT2.NCBI.GGT2.rev.compl                      ctggctaattataaaatgtttatctgtacagacaggttctcgttatgttgcccaggctgg   111518

chimp.LOC112206744.LOC107973052.GGT2-.646279-708040.rev.compl      ------------------------------------------------------------   61762
FAM230B-LOC105372935-GGT2.NCBI.GGT2.rev.compl                      tctcgaactctcggcctcaagcaatcctctcctcctcaccttccaaagccccgaaatatg   111578

chimp.LOC112206744.LOC107973052.GGT2-.646279-708040.rev.compl      ------------------------------------------------------------   61762
FAM230B-LOC105372935-GGT2.NCBI.GGT2.rev.compl                      cccacatccagcctgccaccagtttctttgcatacttctttcctaggcttgaccaagacc   111638

chimp.LOC112206744.LOC107973052.GGT2-.646279-708040.rev.compl      ------------------------------------------------------------   61762
FAM230B-LOC105372935-GGT2.NCBI.GGT2.rev.compl                      tttactttcttctgtgtacagctcagcattgaaaaaaaacaaaaaagtttaggctgggca   111698

chimp.LOC112206744.LOC107973052.GGT2-.646279-708040.rev.compl      ------------------------------------------------------------   61762
FAM230B-LOC105372935-GGT2.NCBI.GGT2.rev.compl                      caatgggtcacgcctgtaatcccaacacttagggtggccaaggtgggaggtttgcttgaa   111758

chimp.LOC112206744.LOC107973052.GGT2-.646279-708040.rev.compl      ------------------------------------------------------------   61762
FAM230B-LOC105372935-GGT2.NCBI.GGT2.rev.compl                      cccaggagttccacaccagcctaggcaagatggcaagaccacatctctccaaaacaattt   111818

chimp.LOC112206744.LOC107973052.GGT2-.646279-708040.rev.compl      ------------------------------------------------------------   61762
FAM230B-LOC105372935-GGT2.NCBI.GGT2.rev.compl                      ttgtaaaattagccaggtgtgctggtgcgggcctgcagtctcagctagtgagaagactaa   111878

chimp.LOC112206744.LOC107973052.GGT2-.646279-708040.rev.compl      ------------------------------------------------------------   61762
FAM230B-LOC105372935-GGT2.NCBI.GGT2.rev.compl                      ggcaagaggaacccttgaacccaggagttcaaggctacggtgagctgtgatcacctctcc   111938

chimp.LOC112206744.LOC107973052.GGT2-.646279-708040.rev.compl      ------------------------------------------------------------   61762
FAM230B-LOC105372935-GGT2.NCBI.GGT2.rev.compl                      agcctgggtgatagggtgagacccagtctctgaaaacaattaaaaataaaaaaagttgaa   111998

chimp.LOC112206744.LOC107973052.GGT2-.646279-708040.rev.compl      ------------------------------------------------------------   61762
FAM230B-LOC105372935-GGT2.NCBI.GGT2.rev.compl                      taatgagtcttctttttagaggcaatgtatttttcctctttttgttttccttaggctata   112058

chimp.LOC112206744.LOC107973052.GGT2-.646279-708040.rev.compl      ------------------------------------------------------------   61762
FAM230B-LOC105372935-GGT2.NCBI.GGT2.rev.compl                      tgcacaatagacaaagttgaaatttaatcttaatctcgtaaacccttactgtccaaacac   112118

chimp.LOC112206744.LOC107973052.GGT2-.646279-708040.rev.compl      ------------------------------------------------------------   61762
FAM230B-LOC105372935-GGT2.NCBI.GGT2.rev.compl                      atcaaactcaaaaaaaaaaaaagacaaaaacaaaaaaggtgaaaggcgtacaagagaaaa   112178

chimp.LOC112206744.LOC107973052.GGT2-.646279-708040.rev.compl      ------------------------------------------------------------   61762
FAM230B-LOC105372935-GGT2.NCBI.GGT2.rev.compl                      aagggaagtaatctgtgccaaatgtacattccttccacaatttgggacaaaaatgatgcc   112238

chimp.LOC112206744.LOC107973052.GGT2-.646279-708040.rev.compl      ------------------------------------------------------------   61762
FAM230B-LOC105372935-GGT2.NCBI.GGT2.rev.compl                      tacctggtctctgggtgccaaaataagacattagagacactcccctccttcaagaaataa   112298

chimp.LOC112206744.LOC107973052.GGT2-.646279-708040.rev.compl      ------------------------------------------------------------   61762
```

```
FAM230B-LOC105372935-GGT2.NCBI.GGT2.rev.compl                  aattggaatcagacactaagatacaattaaacagactttgaaagttggtatgcactactg   112358

chimp.LOC112206744.LOC107973052.GGT2-.646279-708040.rev.compl  ------------------------------------------------------------   61762
FAM230B-LOC105372935-GGT2.NCBI.GGT2.rev.compl                  ggaggtgtctcacagctggcagggtcaactgcaactagaagaaaagctcctaagggcaga   112418

chimp.LOC112206744.LOC107973052.GGT2-.646279-708040.rev.compl  ------------------------------------------------------------   61762
FAM230B-LOC105372935-GGT2.NCBI.GGT2.rev.compl                  aactaagtggctcacttcctctgtaacactgccgtgcccagcactgtaccagcaccccag   112478

chimp.LOC112206744.LOC107973052.GGT2-.646279-708040.rev.compl  ------------------------------------------------------------   61762
FAM230B-LOC105372935-GGT2.NCBI.GGT2.rev.compl                  cagagtcaacaactttccgtcaagccacacacttacggtctcatcgtccagtggaggccg   112538

chimp.LOC112206744.LOC107973052.GGT2-.646279-708040.rev.compl  ------------------------------------------------------------   61762
FAM230B-LOC105372935-GGT2.NCBI.GGT2.rev.compl                  cagatgagacatagtgtgccaaatcccaaatgcacagggtgccagacaagtctgcgggag   112598

chimp.LOC112206744.LOC107973052.GGT2-.646279-708040.rev.compl  ------------------------------------------------------------   61762
FAM230B-LOC105372935-GGT2.NCBI.GGT2.rev.compl                  gcaggaggactagggaagccttccaggcaaggtgatgcctgagctgagttgtttttcttt   112658

chimp.LOC112206744.LOC107973052.GGT2-.646279-708040.rev.compl  ------------------------------------------------------------   61762
FAM230B-LOC105372935-GGT2.NCBI.GGT2.rev.compl                  tgagtttggaacattaatttaatcggaggaaaacaagggacatgaaaagggcaaggaaat   112718

chimp.LOC112206744.LOC107973052.GGT2-.646279-708040.rev.compl  ------------------------------------------------------------   61762
FAM230B-LOC105372935-GGT2.NCBI.GGT2.rev.compl                  aagaaaatgagctgagttttgaaggcaatgatcagatgaagtgagagaagaatattccag   112778

chimp.LOC112206744.LOC107973052.GGT2-.646279-708040.rev.compl  ------------------------------------------------------------   61762
FAM230B-LOC105372935-GGT2.NCBI.GGT2.rev.compl                  cagagggagacaaggagactgcagagatttcaaggggccaagagctgttcggtgcagctc   112838

chimp.LOC112206744.LOC107973052.GGT2-.646279-708040.rev.compl  ------------------------------------------------------------   61762
FAM230B-LOC105372935-GGT2.NCBI.GGT2.rev.compl                  agtgaagcacctaaacagaggtgggggaaggagaacaggggtcaaagcccagtattgaag   112898

chimp.LOC112206744.LOC107973052.GGT2-.646279-708040.rev.compl  ------------------------------------------------------------   61762
FAM230B-LOC105372935-GGT2.NCBI.GGT2.rev.compl                  tgttctggataccaagctaaggagaccttattacgaaggtactggggctgctgaagggtt   112958

chimp.LOC112206744.LOC107973052.GGT2-.646279-708040.rev.compl  ------------------------------------------------------------   61762
FAM230B-LOC105372935-GGT2.NCBI.GGT2.rev.compl                  taaatgggagagtgattgtgaacagattaatagtttggaaagatacaggcagaagtaaac   113018

chimp.LOC112206744.LOC107973052.GGT2-.646279-708040.rev.compl  ------------------------------------------------------------   61762
FAM230B-LOC105372935-GGT2.NCBI.GGT2.rev.compl                  ctctggatttaaccaaaaaataaaggacagaaccacaatgaagatgttggaaaagatcag   113078

chimp.LOC112206744.LOC107973052.GGT2-.646279-708040.rev.compl  ------------------------------------------------------------   61762
FAM230B-LOC105372935-GGT2.NCBI.GGT2.rev.compl                  gaaaaacactgttaaaagatattcactctcggggccgggcgctgtggctcacgccttaaa   113138

chimp.LOC112206744.LOC107973052.GGT2-.646279-708040.rev.compl  ------------------------------------------------------------   61762
FAM230B-LOC105372935-GGT2.NCBI.GGT2.rev.compl                  tcccggcactttggtaggccgaggcgggctgatcacgaggtcaggagatcgagaccatcc   113198

chimp.LOC112206744.LOC107973052.GGT2-.646279-708040.rev.compl  ------------------------------------------------------------   61762
FAM230B-LOC105372935-GGT2.NCBI.GGT2.rev.compl                  tggctaacacggtcaaaccccgtctctactaaaaatacaaaaaattagccgggtgtggtg   113258

chimp.LOC112206744.LOC107973052.GGT2-.646279-708040.rev.compl  ------------------------------------------------------------   61762
FAM230B-LOC105372935-GGT2.NCBI.GGT2.rev.compl                  gcgggcgcctgtagtcccaactactccagaggctgaggcaggagaatggcgtgaacccgg   113318

chimp.LOC112206744.LOC107973052.GGT2-.646279-708040.rev.compl  ------------------------------------------------------------   61762
FAM230B-LOC105372935-GGT2.NCBI.GGT2.rev.compl                  taggcggagcttgcggtgagccgagatcaggccactggaatccagcctgggcgacagagg   113378

chimp.LOC112206744.LOC107973052.GGT2-.646279-708040.rev.compl  ------------------------------------------------------------   61762
FAM230B-LOC105372935-GGT2.NCBI.GGT2.rev.compl                  gagactccatctcaaaaaaaaaaaaaaaaaaaaaaagagagagagtctgactctgttgcc   113438

chimp.LOC112206744.LOC107973052.GGT2-.646279-708040.rev.compl  ------------------------------------------------------------   61762
FAM230B-LOC105372935-GGT2.NCBI.GGT2.rev.compl                  aagactggagtgcagtggcgcgatctaggcacatcacaatcccttccccgcccccgggtt   113498

chimp.LOC112206744.LOC107973052.GGT2-.646279-708040.rev.compl  ------------------------------------------------------------   61762
FAM230B-LOC105372935-GGT2.NCBI.GGT2.rev.compl                  caagtgattctcctgtctcagccgccggagtagctgggactacaggcgcgtgccaccatg   113558

chimp.LOC112206744.LOC107973052.GGT2-.646279-708040.rev.compl  ------------------------------------------------------------   61762
FAM230B-LOC105372935-GGT2.NCBI.GGT2.rev.compl                  tctgattaaatctgtatttttactagagacggggtttcactctgttggccaggctggtct   113618

chimp.LOC112206744.LOC107973052.GGT2-.646279-708040.rev.compl  ------------------------------------------------------------   61762
FAM230B-LOC105372935-GGT2.NCBI.GGT2.rev.compl                  ccaactcctgatctcgtgatccgcccgccccgacctcccaaagtgctgggatgcgtgagc   113678

chimp.LOC112206744.LOC107973052.GGT2-.646279-708040.rev.compl  ------------------------------------------------------------   61762
FAM230B-LOC105372935-GGT2.NCBI.GGT2.rev.compl                  ctccacgcctggccactaatttttctttctttctttcgtgtgtgtgtgtgtgtgtgtgtg   113738

chimp.LOC112206744.LOC107973052.GGT2-.646279-708040.rev.compl  ------------------------------------------------------------   61762
FAM230B-LOC105372935-GGT2.NCBI.GGT2.rev.compl                  tgtgtgtgtgtgtgtgtgtgtgtgtgacgaagtttcgctcttgttgcccaggttggagtg   113798

chimp.LOC112206744.LOC107973052.GGT2-.646279-708040.rev.compl  ------------------------------------------------------------   61762
FAM230B-LOC105372935-GGT2.NCBI.GGT2.rev.compl                  caatggtgtgatctcagctcactgcaatccccgcctcaacaggagagcaggaatcttcag   113858

chimp.LOC112206744.LOC107973052.GGT2-.646279-708040.rev.compl  ------------------------------------------------------------   61762
FAM230B-LOC105372935-GGT2.NCBI.GGT2.rev.compl                  tgatccactggcggatctgcagccattgtgcgcgccaggtcttcccaagtcttttgtgcg   113918

chimp.LOC112206744.LOC107973052.GGT2-.646279-708040.rev.compl  ------------------------------------------------------------   61762
FAM230B-LOC105372935-GGT2.NCBI.GGT2.rev.compl                  cgcgcctctccttccagtacctatgccgccagcgtcccctagcctcccgccattgccagc   113978

chimp.LOC112206744.LOC107973052.GGT2-.646279-708040.rev.compl  ------------------------------------------------------------   61762
FAM230B-LOC105372935-GGT2.NCBI.GGT2.rev.compl                  aggtgctgagatcgcgccattgcactccagcctgggggacaagagcgaaactccatgtca   114038

chimp.LOC112206744.LOC107973052.GGT2-.646279-708040.rev.compl  ------------------------------------------------------------   61762
FAM230B-LOC105372935-GGT2.NCBI.GGT2.rev.compl                  aaaaaaaaaaaaggatgaaaattttgggaaaatatggaagaaaccaaatggatttctagc   114098

chimp.LOC112206744.LOC107973052.GGT2-.646279-708040.rev.compl  ------------------------------------------------------------   61762
FAM230B-LOC105372935-GGT2.NCBI.GGT2.rev.compl                  tcaactaaatcgtaattattcagtgtctagttttggcaagaaaccacatatttcatgtcc   114158

chimp.LOC112206744.LOC107973052.GGT2-.646279-708040.rev.compl  ------------------------------------------------------------   61762
FAM230B-LOC105372935-GGT2.NCBI.GGT2.rev.compl                  acaatcagggacacagtcccagctctcaagtgtgggtctttcctaagcaaattgaagaac   114218

chimp.LOC112206744.LOC107973052.GGT2-.646279-708040.rev.compl  ------------------------------------------------------------   61762
FAM230B-LOC105372935-GGT2.NCBI.GGT2.rev.compl                  acaggcataaaagtacattaaatcaataaaccttttctctctgtctctctctctctttt   114278

chimp.LOC112206744.LOC107973052.GGT2-.646279-708040.rev.compl  ------------------------------------------------------------   61762
FAM230B-LOC105372935-GGT2.NCBI.GGT2.rev.compl                  tttttttttttttgagacggaggttggcactgtcacccaggctggagtgctgtggtgaga   114338

chimp.LOC112206744.LOC107973052.GGT2-.646279-708040.rev.compl  ------------------------------------------------------------   61762
FAM230B-LOC105372935-GGT2.NCBI.GGT2.rev.compl                  tctcgggtcactgcaagctccgcctcccgagttcacgccatcctcctgcctcagcctcca   114398

chimp.LOC112206744.LOC107973052.GGT2-.646279-708040.rev.compl  ------------------------------------------------------------   61762
FAM230B-LOC105372935-GGT2.NCBI.GGT2.rev.compl                  gagtagctgggactacaagcgcccgccacgacgcccggctaatttttttgtagttttagta   114458

chimp.LOC112206744.LOC107973052.GGT2-.646279-708040.rev.compl  ------------------------------------------------------------   61762
FAM230B-LOC105372935-GGT2.NCBI.GGT2.rev.compl                  gagacggggtttcgctatgttggccaggctggtctccaactcgtgacatcgtgatccgcc   114518
```

```
chimp.LOC112206744.LOC107973052.GGT2-.646279-708040.rev.compl      ------------------------------------------------------------   61762
FAM230B-LOC105372935-GGT2.NCBI.GGT2.rev.compl                      cgcctccgactcccaaagtgccacagcccaggcctttttttttaagacagagtctcggc   114578

chimp.LOC112206744.LOC107973052.GGT2-.646279-708040.rev.compl      ------------------------------------------------------------   61762
FAM230B-LOC105372935-GGT2.NCBI.GGT2.rev.compl                      ccggcgcggtggctcacgcctttaatcccagcactttgggaggccgaggcgggctgatca   114638

chimp.LOC112206744.LOC107973052.GGT2-.646279-708040.rev.compl      ------------------------------------------------------------   61762
FAM230B-LOC105372935-GGT2.NCBI.GGT2.rev.compl                      cgaggtcaggaggtcgagaccatcctggctaacacggtcaaaccccgtctctactaaaaa   114698

chimp.LOC112206744.LOC107973052.GGT2-.646279-708040.rev.compl      ------------------------------------------------------------   61762
FAM230B-LOC105372935-GGT2.NCBI.GGT2.rev.compl                      tacaaaaaattagccgggtgtggtggcgggcgcctgtagtccgagctactcccgaggctg   114758

chimp.LOC112206744.LOC107973052.GGT2-.646279-708040.rev.compl      ------------------------------------------------------------   61762
FAM230B-LOC105372935-GGT2.NCBI.GGT2.rev.compl                      aggcaggagaatggcgtgaacccggtaggcggagcttgcggtgagccgagatcaggccac   114818

chimp.LOC112206744.LOC107973052.GGT2-.646279-708040.rev.compl      ------------------------------------------------------------   61762
FAM230B-LOC105372935-GGT2.NCBI.GGT2.rev.compl                      tggaatccagcctgggcgacagagggagactccgtctcaaaaaaaaaaaaaaagagtctg   114878

chimp.LOC112206744.LOC107973052.GGT2-.646279-708040.rev.compl      ------------------------------------------------------------   61762
FAM230B-LOC105372935-GGT2.NCBI.GGT2.rev.compl                      gctctgttgcccaggctggagtgcagtggcgcgatctcggcgcatcgcaatcccttcccc   114938

chimp.LOC112206744.LOC107973052.GGT2-.646279-708040.rev.compl      ------------------------------------------------------------   61762
FAM230B-LOC105372935-GGT2.NCBI.GGT2.rev.compl                      gcccccgggttcaagtgattctccagtctcagccgccggagtagctgggactacaggcgc   114998

chimp.LOC112206744.LOC107973052.GGT2-.646279-708040.rev.compl      ------------------------------------------------------------   61762
FAM230B-LOC105372935-GGT2.NCBI.GGT2.rev.compl                      gtgccaccatgtctgactaaatttgtatttttactagagacggggtttcactatgttggc   115058

chimp.LOC112206744.LOC107973052.GGT2-.646279-708040.rev.compl      ------------------------------------------------------------   61762
FAM230B-LOC105372935-GGT2.NCBI.GGT2.rev.compl                      caggctggtctccaactcctgatctcgtgatccgtctgccccgacctcccaaagtgctag   115118

chimp.LOC112206744.LOC107973052.GGT2-.646279-708040.rev.compl      ------------------------------------------------------------   61762
FAM230B-LOC105372935-GGT2.NCBI.GGT2.rev.compl                      gattataggcataagccaccacgcccggcctctttttttctttttcttttctttatctg   115178

chimp.LOC112206744.LOC107973052.GGT2-.646279-708040.rev.compl      ------------------------------------------------------------   61762
FAM230B-LOC105372935-GGT2.NCBI.GGT2.rev.compl                      gagactgagttttgctcttgttgcccaggctggagtgcaatggtgcgatctcagctcact   115238

chimp.LOC112206744.LOC107973052.GGT2-.646279-708040.rev.compl      ------------------------------------------------------------   61762
FAM230B-LOC105372935-GGT2.NCBI.GGT2.rev.compl                      gcaatctccacctcagcaggagagcaggaatcttcagtgatccacgggcagatctgccgc   115298

chimp.LOC112206744.LOC107973052.GGT2-.646279-708040.rev.compl      ------------------------------------------------------------   61762
FAM230B-LOC105372935-GGT2.NCBI.GGT2.rev.compl                      cattgtgggcacctgtttctcccgcaacctttgtgcccgcctctctccttcccgtaccta   115358

chimp.LOC112206744.LOC107973052.GGT2-.646279-708040.rev.compl      ------------------------------------------------------------   61762
FAM230B-LOC105372935-GGT2.NCBI.GGT2.rev.compl                      ttgcatgaccccccacgtccgcctcccgccattgccagcaagtgcctcgcgcgggtacct   115418

chimp.LOC112206744.LOC107973052.GGT2-.646279-708040.rev.compl      ------------------------------------------------------------   61762
FAM230B-LOC105372935-GGT2.NCBI.GGT2.rev.compl                      ggctgcgcttattaatccgttaagctcgctctgtcacgggcgccgtgatgtgctcacgcg   115478

chimp.LOC112206744.LOC107973052.GGT2-.646279-708040.rev.compl      ------------------------------------------------------------   61762
FAM230B-LOC105372935-GGT2.NCBI.GGT2.rev.compl                      cccgctccctcaggtttaaaaggcgcgttgcccggcaacagaagaaactgctggcttagc   115538

chimp.LOC112206744.LOC107973052.GGT2-.646279-708040.rev.compl      ------------------------------------------------------------   61762
FAM230B-LOC105372935-GGT2.NCBI.GGT2.rev.compl                      cgttggccgagttggcggctggacgaggacgctcagagcccagctctcgagagttcaagc   115598

chimp.LOC112206744.LOC107973052.GGT2-.646279-708040.rev.compl      ------------------------------------------------------------   61762
FAM230B-LOC105372935-GGT2.NCBI.GGT2.rev.compl                      aaccgacggttccccactgctcccaggagcggttacctgggcactctgtgcccctccttc   115658

chimp.LOC112206744.LOC107973052.GGT2-.646279-708040.rev.compl      ------------------------------------------------------------   61762
FAM230B-LOC105372935-GGT2.NCBI.GGT2.rev.compl                      ctgttcgggcccaggccgaggacctgccagtagggctcagttgcctggagcccgttcagc   115718

chimp.LOC112206744.LOC107973052.GGT2-.646279-708040.rev.compl      ------------------------------------------------------------   61762
FAM230B-LOC105372935-GGT2.NCBI.GGT2.rev.compl                      ccatcccccagttcactttgcttgtgggatctccccgttgctcctgcccgtggactgagt   115778

chimp.LOC112206744.LOC107973052.GGT2-.646279-708040.rev.compl      ------------------------------------------------------------   61762
FAM230B-LOC105372935-GGT2.NCBI.GGT2.rev.compl                      ggcaggccatcctacaagcacccggacacttgacatcagtggtgtcaagacaactctaag   115838

chimp.LOC112206744.LOC107973052.GGT2-.646279-708040.rev.compl      ------------------------------------------------------------   61762
FAM230B-LOC105372935-GGT2.NCBI.GGT2.rev.compl                      aaggttttccgtgatcctgcaagccctgccttccttcctgggatcctgccttcaatttga   115898

chimp.LOC112206744.LOC107973052.GGT2-.646279-708040.rev.compl      ------------------------------------------------------------   61762
FAM230B-LOC105372935-GGT2.NCBI.GGT2.rev.compl                      ttgcacaggtaccacagcaagccagtgctgcgtgctccgagttccagggcgtcctccagc   115958

chimp.LOC112206744.LOC107973052.GGT2-.646279-708040.rev.compl      ------------------------------------------------------------   61762
FAM230B-LOC105372935-GGT2.NCBI.GGT2.rev.compl                      tcagccactgcactgagaacatggactctctgtggggcccaggagccgggagtcacccct   116018

chimp.LOC112206744.LOC107973052.GGT2-.646279-708040.rev.compl      ------------------------------------------------------------   61762
FAM230B-LOC105372935-GGT2.NCBI.GGT2.rev.compl                      ttggggtccacaacagccggctgtccccagacttgtgtccagggaagatagtgttgaggg   116078

chimp.LOC112206744.LOC107973052.GGT2-.646279-708040.rev.compl      -------------------------------------------     61762
FAM230B-LOC105372935-GGT2.NCBI.GGT2.rev.compl                      ccctcaaggagagcggggcagggatgcctgagcaggacaagga     116121
```
